# A Non-Canonical Role of SMAD4 in Regulating 3D Genome Architecture to Inhibit Lung Squamous Cell Carcinoma Development

**DOI:** 10.64898/2026.03.28.714793

**Authors:** Qian Tang, Chen Lian, Xinyan Han, Xuhan Zhang, Zihan Wang, Boyu Wang, Taoyu Zhu, Xinrui Lin, Xiaolei Wang, Yaping Xu, Manyu Xiao, Zijin Wang, Junmin Li, Silin Chen, Yunze Wang, Yufei Liu, Songsong Li, Zuolin Shen, Xi Lu, Xueqi Han, Yilu Zhou, Mingyang Xiao, Jiayi Ran, Xiaoran Cao, Xinyi Xu, Hugo Sámano-Sánchez, Alfredo Rodríguez, Lian Wang, Shuifang Chen, Zhanyu Xu, Shirong Zhang, Nuo Yang, Yong Tang, Jian Liu

## Abstract

Lung squamous cell carcinoma (LUSC) lacks clearly defined key drivers and effective targeted therapies, reflecting an incomplete understanding of its molecular pathogenesis. Here, we identify SMAD4 as a critical regulator of three-dimensional (3D) genome organization in LUSC and uncover a mechanistic link between tumor suppressor loss and oncogenic transcriptional activation. By integrating clinical datasets, genetically engineered mouse models, human and murine LUSC cell lines, and multi-omics analyses, we demonstrate that SMAD4 deficiency promotes LUSC progression by unleashing EP300-mediated enhancer-promoter looping at the *SOX2* locus. Mechanistically, SMAD4 does not directly bind *SOX2* regulatory elements but instead constrains chromatin looping by sequestering EP300 away from loop anchor regions. Loss of SMAD4 leads to enhanced H3K27ac deposition, aberrant *SOX2* activation, and increased LUSC tumor cell proliferation. Together, these findings reveal a non-canonical role for a transcription factor (*e.g.*, SMAD4) in regulating dysregulated 3D genome architecture to inhibit tumor development.

## 1. Introduction

Lung squamous cell carcinoma (LUSC) is a major histological subtype of lung cancer, accounting for approximately 20-30% of cases worldwide.^[1,2]^ In contrast to lung adenocarcinoma (LUAD), for which multiple key oncogenic drivers and targeted therapies have been successfully developed, LUSC lacks well-defined driver alterations and remains without effective targeted treatments.^[3,4]^ This therapeutic deficiency reflects, at least in part, an incomplete understanding of LUSC molecular pathogenesis, underscoring the urgent need to elucidate the regulatory mechanisms governing LUSC initiation and progression.

Unlike LUAD, where single genetic alterations such as *EGFR* mutations are sufficient to drive tumor development,^[5]^ LUSC is rarely induced by a single oncogenic event. To date, *Lkb1* deletion and *IKK*α^K44A^ knock-in have been reported as single genetic alterations capable of inducing LUSC in mouse models.^[6,7]^ Although *IKK*α^K44A^ effectively induces LUSC,^[7]^ this mutation has not been identified in human LUSC, probably limiting the translational relevance of this model. Notably, LKB1 expression is markedly downregulated in *IKK*α^K44A^-induced LUSC,^[7]^ highlighting the critical role of LKB1 in suppressing LUSC development. Consistently, *Lkb1* ablation using Adeno-Cre or Lenti-Cre delivery via nasal or bronchial routes failed to induce pathological phenotypes.^[8–10]^ In contrast, LUSC developed when *Lkb1* deletion was combined with *KRAS^G12D^* knock-in, *Sox2* overexpression, or *Pten* deletion using the same delivery approach.^[8–10]^ One likely explanation is that Adeno-Cre or Lenti-Cre administration through nasal or bronchial routes inefficiently infects bronchial epithelial cells, the presumed cells of origin for LUSC, due to mucociliary clearance.^[11]^ Consequently, identifying a single key LUSC driver by introducing genetic alterations into bronchial epithelial cells has proven challenging. To overcome this technical limitation, we generated a CCSP^iCre^ mouse line that enabled highly efficient Cre-LoxP-mediated recombination in bronchial epithelial cells.^[12]^ Using this model, we demonstrated that *Lkb1* ablation alone induced spontaneous LUSC development at 11-14 months of age.^[6]^ These findings established LKB1 as a critical suppressor of LUSC and further indicate that additional genetic alterations cooperate with LKB1 deficiency to promote tumorigenesis, consistent with the mutational complexity observed in human LUSC.

Recent advances in cancer genomics and epigenomics have demonstrated that tumor development is governed not only by genetic alterations but also by higher-order regulatory mechanisms that shape transcriptional programs.^[13]^ In LUAD, accumulating evidence indicated that disruption of three-dimensional (3D) genome organization, including chromatin looping, topologically associating domains (TADs), and enhancer-promoter interactions, played a central role in oncogene activation and lineage plasticity.^[14]^ However, despite the extensive chromosomal instability and epigenetic dysregulation characteristic of LUSC, the contribution of 3D genome architecture to LUSC tumorigenesis remains largely unexplored. A systematic understanding of spatial genome regulation in LUSC, therefore, represents a major unmet need in the field.

Among the molecular hallmarks of LUSC, amplification and overexpression of the lineage-defining transcription factor *SOX2* are among the most recurrent and functionally significant events. SOX2 is essential for squamous lineage specification and promotes tumor cell proliferation, survival, and malignant progression in LUSC.^[9]^ Although multiple transcriptional and epigenetic regulators of SOX2 expression have been identified, previous studies have largely focused on linear regulatory mechanisms, such as promoter activity and enhancer-associated histone modifications.^[15,16]^ Whether *SOX2* activation in LUSC is governed by aberrant 3D chromatin organization, and how such spatial regulation is established or constrained, remains largely unknown.

LUSC development is predominantly driven by multiple genetic alterations.^[17]^ Among these, SMAD4, a central mediator of transforming growth factor-β (TGF-β) signaling, functions as a frequently altered tumor suppressor in LUSC by repressing SOX2 expression in cooperation with ERK1/2 signaling.^[18]^ However, the transcriptional mechanisms underlying SMAD4-mediated regulation of *SOX2* remain unclear, including whether SMAD4 directly binds the *SOX2* locus to exert its canonical transcription factor (TF) activity.^[18]^ Moreover, accumulating evidence showed that SMAD4 loss alone was insufficient to initiate tumorigenesis,^[12]^ instead exerting strong cooperative effects with other oncogenic alterations. For instance, SMAD4 was reported to cooperate with ERK1/2 to repress *SOX2* expression, such that LUSC developed in Ad-Cre *Smad4*^f/f^ *KRAS^G12D^* mice only in the presence of ERK1/2 inhibition.^[18]^ Consequently, the role of SMAD4 in regulating spontaneous LUSC development in the absence of pharmacological intervention remains unclear. Notably, a purely linear transcriptional dysregulation model appears insufficient to explain the pronounced tumor-promoting synergy associated with SMAD4 loss, raising the possibility that SMAD4 may exert broader regulatory functions at the level of chromatin architecture.

Emerging evidence has further suggested that transcription factors can play non-canonical roles in shaping 3D genome organization beyond regulating gene expression through linear promoter/enhancer binding. For example, the ubiquitously expressed transcription factor YY1 was shown to act as a structural regulator of enhancer–promoter looping by dimerizing and facilitating DNA-DNA contacts, thereby supporting transcriptional activation programs.^[19]^ In parallel, master lineage factors have also been implicated in higher-order genome rewiring. For example, OCT4 was reported to regulate topologically associating domain (TAD) reorganization during somatic cell reprogramming through phase separation-associated chromatin looping dynamics.^[20]^ Moreover, cell-type-specific transcriptional complexes can directly instruct long-range enhancer-promoter communication; in the erythroid system, LDB1-centered complexes (working with lineage TFs such as GATA1/TAL1) were demonstrated to be required for establishing and/or maintaining enhancer-driven chromatin looping and transcriptional activation.^[21]^ Together, these precedents support the concept that TFs and their cofactors can function as active organizers of genome topology, raising the possibility that dysregulation of such TF-associated architectural mechanisms may contribute to oncogenic transcriptional programs in LUSC.

In this study, by integrating clinical datasets, genetically engineered mouse models, murine and human LUSC cell lines, and multi-omics analyses, we uncovered a previously unrecognized role of SMAD4 in regulating 3D genome organization in LUSC. We demonstrated that SMAD4 deficiency unleashed EP300-mediated chromatin looping at the *SOX2* locus, resulting in enhanced enhancer-promoter interactions, increased H3K27ac deposition, and aberrant *SOX2* activation. This chromatin architecture rewiring drove tumor cell proliferation and LUSC progression without altering squamous lineage identity. Collectively, our findings established SMAD4 as a critical guardian of chromatin loop homeostasis in LUSC and revealed an epigenetic mechanism by which tumor suppressor loss indirectly activates oncogenic transcriptional programs. These insights advanced our understanding of LUSC pathogenesis and highlighted chromatin looping regulators as potential therapeutic vulnerabilities in squamous cell lung cancer.

## 2. Results

### 2.1 SMAD4 Deficiency Promotes Mouse LUSC Development

We first queried the role of SMAD4 in regulating LUSC tumorigenesis. Clinical data from the TRACERX database indicated that multiple key genes exhibited mutations in human LUSC tumors, with the *SMAD4* mutation rate even surpassing that of the key driver genes *KRAS* and *STK11* (Figure 1A). In LUSC patients with *KRAS*/*TP53* mutations or low *STK11* expression level, SMAD4 showed lower expression than in those without *KRAS/TP53* mutations or high *STK11* expression (Figure 1B). Reduced *SMAD4* expression in LUSC correlated with these specific oncogenic backgrounds, suggesting a potential upstream regulatory mechanism that downregulates *SMAD4* during LUSC tumorigenesis. These suggested that oncogenic alterations, such as *KRAS*/*TP53* mutations or *STK11* loss, might cause selective pressures to reduce *SMAD4* expression. Low *SMAD4* expression was significantly associated with poor prognosis in LUSC patients (Figure 1C). Together, these results suggested that SMAD4 affected LUSC tumorigenesis.

**Figure 1.**
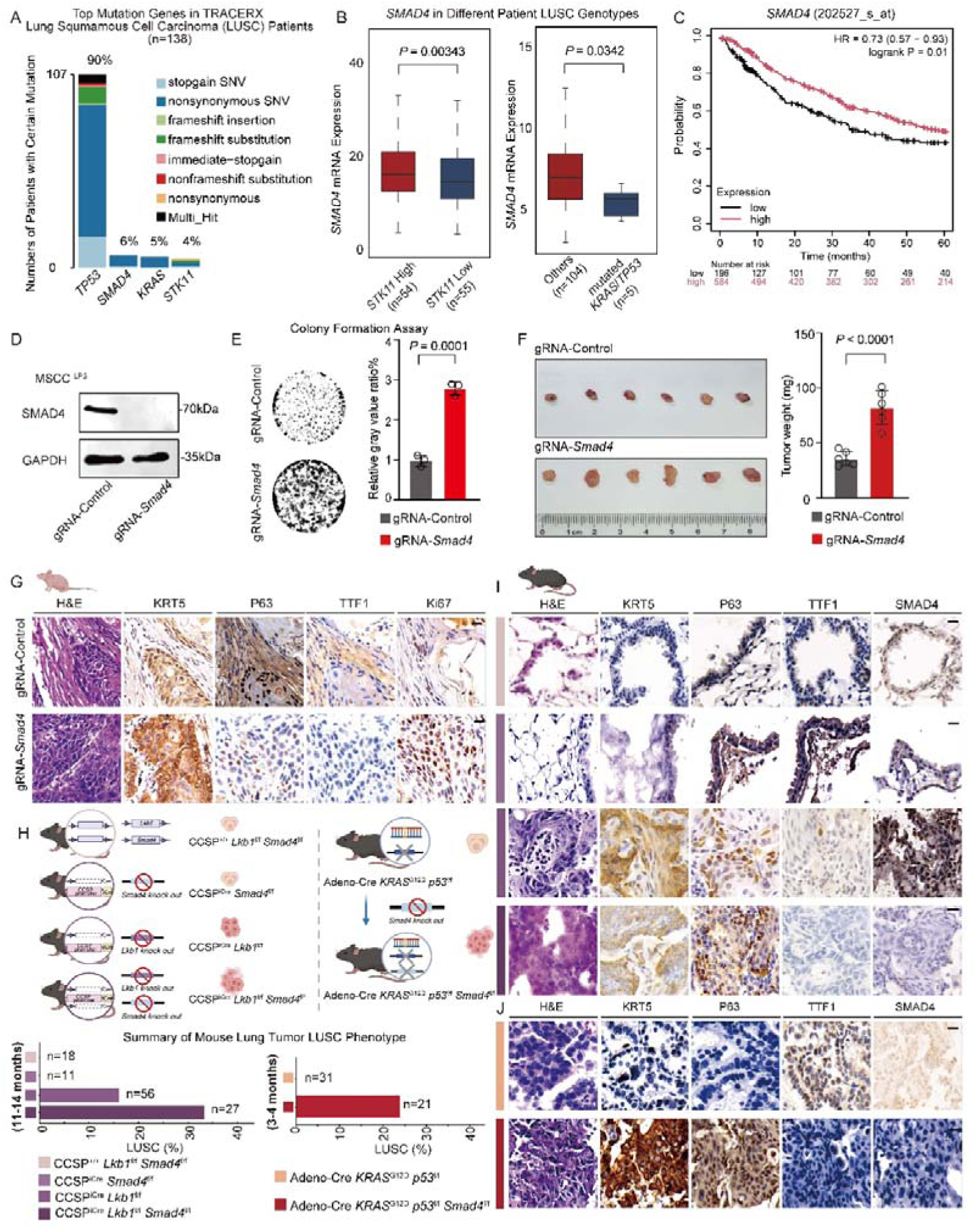
SMAD4 deficiency promotes mouse LUSC development. (A) The proportion of samples with mutations in *TP53*, *SMAD4*, *KRAS*, and *STK11* in TRACERX LUSC cohort was shown by bar plots. (B) *SMAD4* mRNA expression (log2 scale) between different genotypes of LUSC patients. (C) Survival curves showed the relationship between *SMAD4* expression levels and prognosis. (D) Western Blot analysis of SMAD4 protein levels in *Smad4* knockout (gRNA-*Smad4*) and non-knockout (gRNA-Control) in MSCC ^LP.3^ cell lines. GAPDH served as the loading control. (E) Colony formation assay under *Smad4* knockout or non-knockout conditions and the quantification of the relative gray value ratio in MSCC ^LP.3^ cell line using ImageJ. Data were presented as mean ± SD. (F) Tumor size and weight of MSCC ^LP.3^ gRNA-Control tumor groups and gRNA-*Smad4* tumor groups in vivo. (G) Phenotype analysis by H&E staining and IHC staining of LUSC markers KRT5 and TP63, LUAD marker TTF1, and cell proliferation marker Ki67 in xenograft tumors with or without *Smad4* knockout. The scale bar presented 20µm. (H) Mouse model design workflow and phenotype statistics of LUSC in different genetic backgrounds. (I) LUSC markers KRT5 and TP63, LUAD marker TTF1, and SMAD4 IHC staining along with H&E staining in CCSP*^+/+^ Lkb1*^f/f^ *Smad4*^f/f^, CCSP*^iCre^ Smad4*^f/f^, CCSP*^iCre^ Lkb1*^f/f^, and CCSP*^iCre^ Lkb1*^f/f^ *Smad4*^f/f^ mice. (J) IHC staining of SMAD4, LUSC markers KRT5 and TP63, and LUAD marker TTF1, and H&E staining in Adeno-Cre *KRAS*^G12D^ *Trp53*^f/f^ and Adeno-Cre *Smad4*^f/f^ *KRAS*^G12D^ *Trp53*^f/f^ mice.

To investigate the role of SMAD4 in regulating LUSC development, we first knocked out *Smad4* in the murine LUSC cell line MSCC^LP.3^ (Figure 1D). This cell line was derived from CCSP^iCre^*Lkb1*^f/f^*Pten*^f/f^ mice through Cre/LoxP-mediated deletion of *Lkb1* and *Pten* in lung epithelial cells.^[6]^ *Smad4* knockout significantly enhanced cell colony formation (Figure 1E). Consistently, subcutaneous xenograft assays showed that tumors derived from *Smad4*-deficient cells were larger than those from control tumors (Figure 1F). Histological analysis confirmed classical squamous features, including keratinization and keratin pearl formation (Figure 1G and Supplementary Figure 1A). At the same time, immunohistochemistry showed positive staining for LUSC markers KRT5 and TP63 and negative staining for the lung adenocarcinoma (LUAD) marker TTF1 (Figure 1G and Supplementary Figure 1A). Notably, Ki67 staining was substantially elevated in SMAD4-deficient tumors, indicating enhanced proliferative activity *in vivo* (Figure 1G and Supplementary Figure 1A). Together, these results demonstrated that SMAD4 suppressed LUSC cell proliferation without affecting squamous differentiation.

To further assess the role of SMAD4 during LUSC initiation *in vivo*, we employed genetically engineered mouse models. Given that LKB1 deficiency was shown to be the key driver of LUSC^[6]^ and reduced *SMAD4* expression was frequently observed in human tumors harboring *STK11* alterations (Figure 1B), we examined the effect of *Smad4* loss in an *Lkb1*-deficient background. CCSP^iCre^*Lkb1*^f/f^ mice developed LUSC as early as 11-14 months of age, and there was no tumor formation in CCSP^iCre^*Smad4*^f/f^ mice (Figure 1H-I and Supplementary Figure 1B), as our previously published findings.^[6]^ *Smad4^f/f^* mice were crossed with CCSP^iCre^*Lkb1*^f/f^ mice to generate *Lkb1/Smad4* double-knockout mice, which further promoted the LUSC development, doubling the incidence rate compared to LUSC in *Lkb1* knockout mice alone (Figure 1H-I and Supplementary Figure 1B). In sum, SMAD4 repressed LKB1-deficiency-driven LUSC tumorigenesis.

Reduced *SMAD4* expression was frequently observed in human tumors harboring *TP53-KRAS* co-mutations (Figure 1B). Therefore, we also examined the role of SMAD4 in a *Trp53^-/-^*/*KRAS^G12D^*-driven LUSC mouse model generated by adenoviral Cre delivery. Consistent with observations in the *Lkb1*-deficient background, *Smad4* deletion in *p53^-/-^ KRAS^G12D^* mutant mice drove the *de novo* LUSC development (Figure 1H, Figure 1J, and Supplementary Figure 1C), although we previously showed that there was only LUAD development in *p53^-/-^ KRAS^G12D^* mutant mice.^[22]^ These lung tumors in Adeno-Cre *KRAS^G12D^ p53^f/f^ Smad4^f/f^* mice exhibited molecular features closely resembling human LUSC, showing positive staining for LUSC markers (*e.g.*, TP63/KRT5) and negative staining for LUAD markers (*e.g.*, TTF1) (Figure 1J and Supplementary Figure 1C). This intriguing phenomenon suggested that SMAD4 functioned as a lineage-stabilizing factor, whose loss promoted the conversion of glandular tissue into squamous epithelium in a permissive or active manner under specific genetic and signaling contexts. Meanwhile, the *p53^-/-^*-*KRAS^G12D^* mutant mouse lung tumors showed the positive LUAD marker profile staining (Figure 1J and Supplementary Figure 1C). These findings indicated that SMAD4 exerted a tumor-suppressive role in LUSC across distinct oncogenic contexts, including both *STK11* loss (Figure 1I) and *TP53-KRAS* co-mutation backgrounds (Figure 1J).

### 2.2 SMAD4 Deficiency Drives Mouse LUSC Oncogenic Transcriptome Program (*e.g.*, *Sox2* overexpression)

To elucidate the mechanisms by which SMAD4 repressed LUSC tumorigenesis, we performed transcriptomic sequencing (RNA-Seq) on mouse LUSC cell lines (MSCC^LP.3^) with or without *Smad4* knockout (Figure 2A and Supplementary Table 1). Gene Ontology (GO) enrichment analysis of differentially expressed genes (DEGs) revealed that *Smad4*-regulated targets were significantly enriched in tumor-promoting pathways, including cell cycle progression and DNA replication (Figure 2B and Supplementary Figure 2A). Notably, *Smad4* knockout also led to marked alterations in pathways associated with chromatin remodeling, a key hallmark of tumorigenesis (Figure 2B). Consistently, Gene Set Enrichment Analysis (GSEA) further demonstrated significant enrichment of the cell cycle pathway, with differentially expressed genes including *Sox2*, a well-established LUSC marker and driver (Figure 2C). Protein–protein interaction (PPI) network analysis identified *Sox2* as a central hub gene within the cell cycle regulatory network, suggesting a pivotal role for *Sox2* in SMAD4-suppressive tumor progression (Figure 2D). Based on these observations, we selected *Sox2* as a representative target to further dissect the molecular mechanisms by which SMAD4 loss promoted LUSC progression. Immunoblotting analysis showed that SOX2 protein levels were markedly increased following *Smad4* deletion (Figure 2E). Concordantly, quantitative PCR (qPCR) analysis revealed a significant upregulation of *Sox2* mRNA expression level upon *Smad4* knockout (Figure 2F). To observe the dynamic regulation of *Sox2* by *Smad4* deficiency, we monitored the expression changes of *Smad4* and *Sox2* at regular intervals over 48-hour siRNA treatment. *Sox2* expression progressively increased as *Smad4* levels decreased, further indicating the dynamic regulatory relationship between SMAD4 and *Sox2* (Figure 2G). Consistent with this, we observed a significant downregulation of *Sox2* expression following overexpression of *Smad4* (*Smad4*_OV) (Figure 2H).

**Figure 2.**
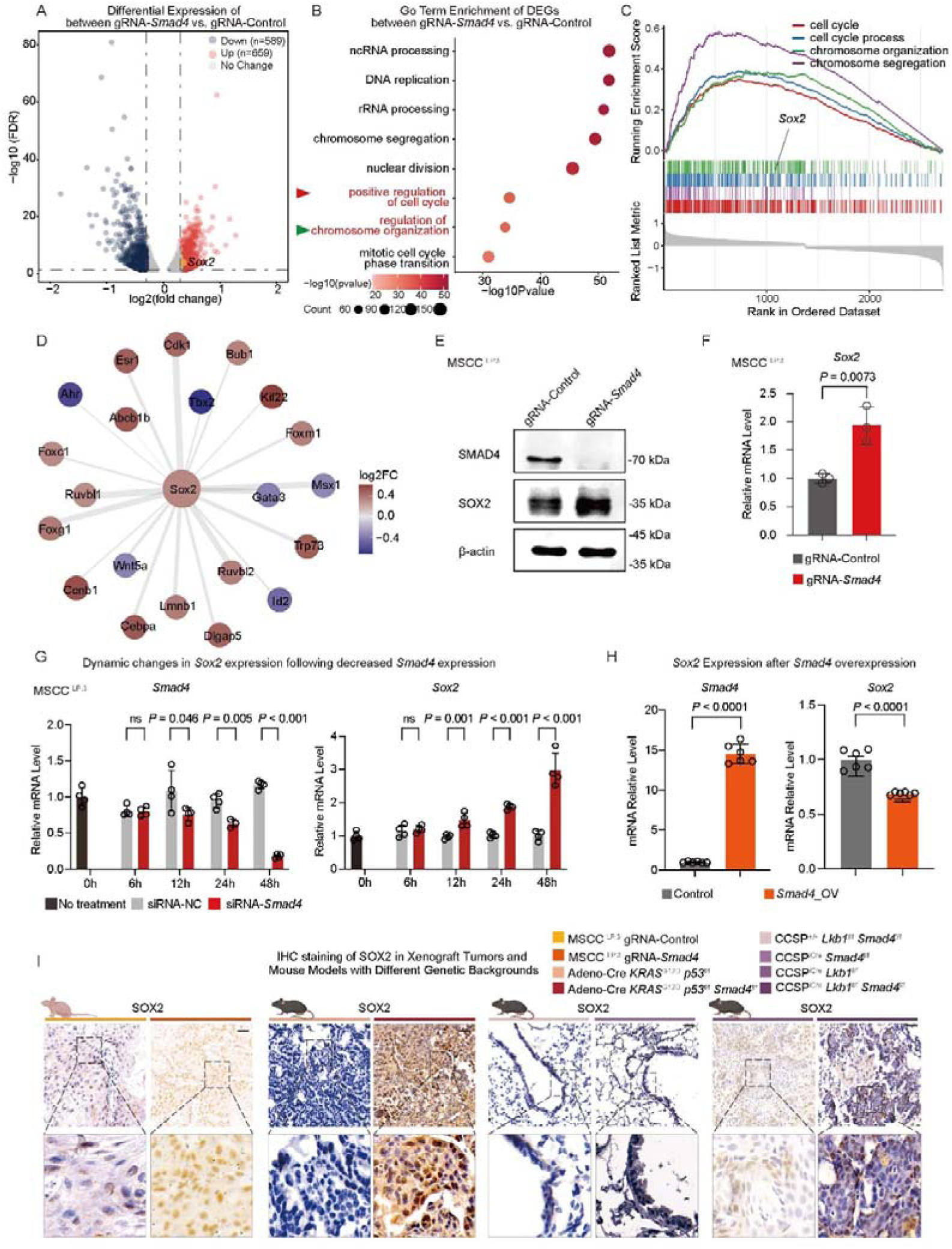
SMAD4 deficiency drives mouse LUSC oncogenic transcriptome program (*e.g.*, *Sox2* overexpression). (A) Differentially expressed genes between MSCC^LP.3^ gRNA-*Smad4* and gRNA-Control illustrated by the volcano plot. (B) The top enriched signaling pathways obtained from differentially expressed genes by GO enrichment. (C) Pathways enriched from differentially expressed genes by GSEA enrichment. (D) Gene Interaction Network in the Cell Cycle Process Extracted from the STRING Database. (E) Western Blot analysis of SOX*2* protein levels in *Smad4* knockout (gRNA-*Smad4*) and non-knockout (gRNA-Control) in MSCC^LP.3^ cell lines. β-actin served as a loading control. (F) RT-qPCR analysis of *Sox2* mRNA in *Smad4* knockout (gRNA-Smad4) and non-knockout (gRNA-Control) in MSCC^LP.3^ cell lines. 18S served as the RT-qPCR loading control. (G) *Smad4* expression levels and *Sox2* expression levels at different time points following *Smad4* siRNA knockdown in MSCC^LP.3^ cell lines. (H) *Sox2* expression level before and after overexpression of *Smad4* (*Smad4_*OV). (I) IHC staining of SOX2 in xenograft tumors with or without *Smad4* knockout and mouse models with different genetic backgrounds.

Furthermore, immunohistochemical staining demonstrated elevated SOX2 expression in both SMAD4-deficient cell lines and spontaneous tumors arising from *Smad4* knockout mice across different genetic backgrounds. In agreement with the *in vitro* findings, *Smad4* loss *in vivo* robustly enhanced *Sox2* expression (Figure 2I). Collectively, these results demonstrated that SMAD4 inhibited the expression of LUSC oncogenic drivers (*e.g.*, *Sox2*) *in vitro* and *in vivo*.

### 2.3 SMAD4 Deficiency Drives Mouse LUSC Oncogenic Transcriptome Program (*e.g.*, *Sox2* overexpression) by Remodeling 3D Genome

Since SMAD4 is a typical TF, we conducted a SMAD4 CUT&RUN to examine its genome-wide binding profile in mouse LUSC cells (MSCC^LP.3^) (Figure 3A). Annotation of SMAD4 binding sites revealed a strong enrichment in distal intergenic and intronic regions, rather than the promoter-proximal regions (Figure 3B and Supplementary Table 2). Similar profiles were observed in the CUT&RUN analyses of transcriptionally active enhancer marker (H3K27ac) (Figure 3A, Figure 3B, and Supplementary Table 3). Quantitative analysis of binding site distribution showed that SMAD4 preferentially localizes at genomic regions approximately 1Mb away from transcription start sites in MSCC^LP.3^ cell lines, suggesting a predominant role in regulating distal regulatory elements (*e.g.*, H3K27ac) (Figure 3C). Unexpectedly, despite its distal localization pattern, SMAD4 exhibited limited direct occupancy at active enhancers (Figure 3D). Specifically, there was no detectable SMAD4 binding at the *Sox2* promoter enhancers or distant enhancers (Figure 3E). Compared the published H3K27ac ChIP-Seq in mouse normal lungs with our mouse LUSC H3K27ac profile at *Sox2* and its nearby region, we found a H3K27ac peak specifically enriched in the mouse LUSC, indicated by the red arrow (Figure 3E). These results suggested the distant enhancers might affect *Sox2* expression through a nonlinear mechanism (Figure 3E). Therefore, we designed gRNAs to delete this distant enhancer and observed significantly downregulated *Sox2* expression (Figure 3F and Supplementary Figure 1D) and inhibited LUSC colony formation (Figure 3G). To further validate the specificity of this mechanism, we also designed antisense oligonucleotides (ASO) targeting the *Sox2* enhancer to suppress its function. Similarly, *Sox2* expression was significantly reduced in the anti-enhancer cell line (Figure 3H). These data suggested that SMAD4 repressed *Sox2* transcription and cell survival in LUSC partially via a distant enhancer through a non-direct pathway.

**Figure 3.**
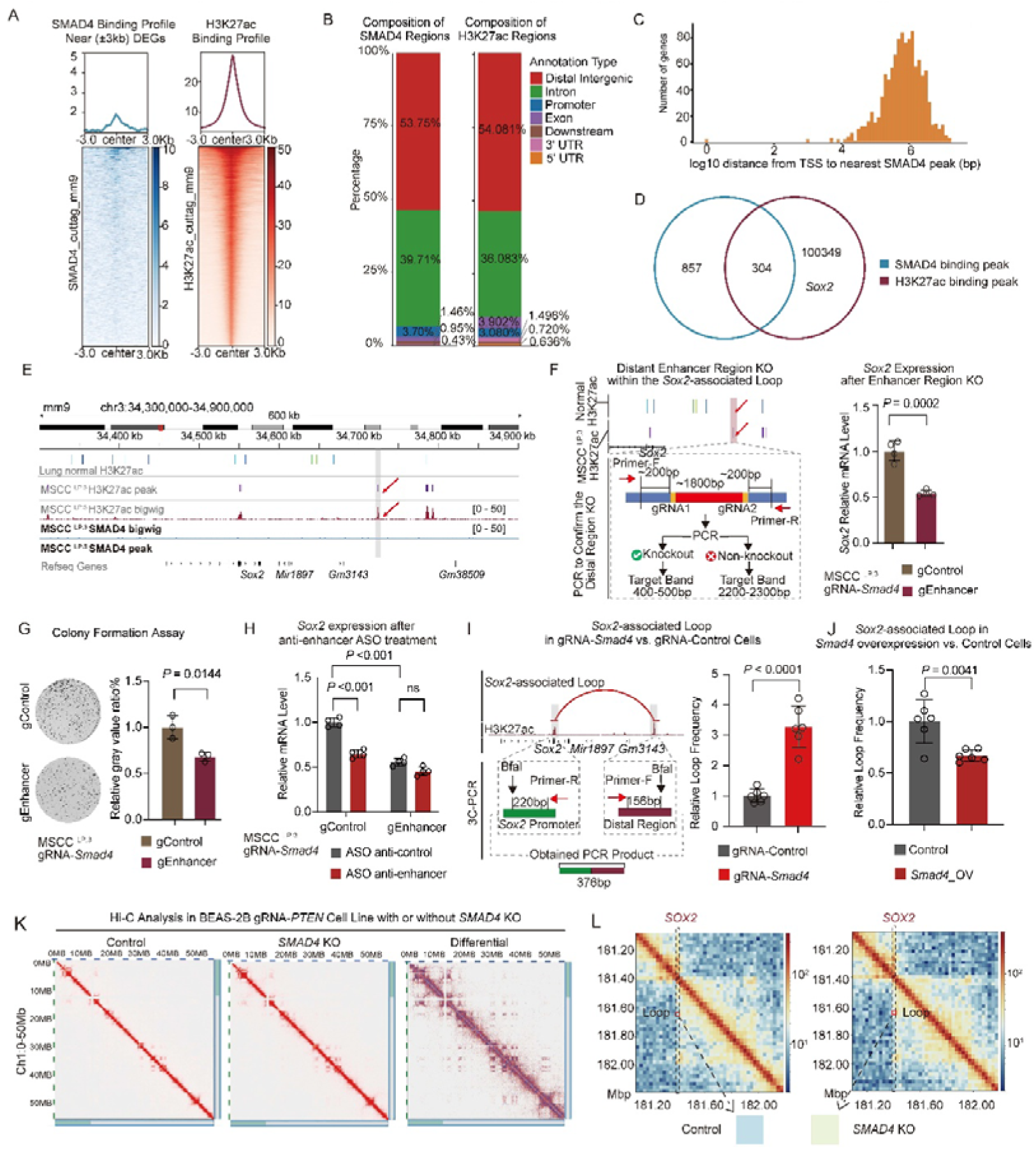
SMAD4 deficiency drives mouse LUSC oncogenic transcriptome program (*e.g.*, *Sox2* overexpression) by remodeling 3D Genome. (A) Profiling of SMAD4 and H3K27ac binding on DEGs in MSCC^LP.3^. (B) The location distribution of SMAD4 binding on different genomic regions. (C) Statistics of SMAD4 binding distribution around gene transcription start sites (TSS). (D) Venn plot of genes regulated by SMAD4 binding and H3K27ac modification. (E) Liftover H3K27ac ChIP-Seq peak regions of normal human lung tissue from the reference genome hg19 to the mouse reference genome mm9, matching with the H3K27ac CUT&RUN result in MSCC ^LP.3^ shown in IGV browser. (F) RT-qPCR of *Sox2* expression followed by the knockout of the enhancer region of *Sox2*-associated loop in MSCC^LP.3^ gRNA-*Smad4* cell line. (G) Colony formation assay after the knockout enhancer region of *Sox2*-associated loops in MSCC^LP.3^ gRNA-*Smad4* cell line and the quantification of the relative gray value ratio in MSCC^LP.3^ cell line using ImageJ. (H) *Sox2* expression before and after *Sox2*-associated enhancer region targeted by antisense oligonucleotides (ASO) in MSCC^LP.3^ with or without *Smad4* KO. Cells were incubated with 15nM ASO oligos for 48□h. (I) The 3C-qPCR analysis of the frequency of *Sox2*-associated loop by 3C-qPCR after *Smad4* knockout. (J) The 3C-qPCR analysis of the frequency of *Sox2*-associated loop by 3C-qPCR after *Smad4* overexpression (*Smad4*_OV). (K) Hi-C analysis of BEAS-2B gRNA-*PTEN* cell line with or without *SMAD4* knockout. (L) Hi-C heatmap around the *SOX2* locus exhibiting the location of the *SOX2*-associated loops in BEAS-2B gRNA-*PTEN* cells with or without *SMAD4* knockout.

Transcriptome profiling suggested that *Smad4* deficiency might remodel higher-order chromatin architecture (Figure 2C), thereby regulating key downstream oncogenic programs. We therefore sought to elucidate the molecular basis of three-dimensional (3D) genome remodeling and its contribution to SMAD4-dependent regulation of downstream targets, such as *Sox2*. Using Chromosome Conformation Capture (3C)-qPCR, we validated the existence of this *Sox2*-associated chromatin loop and demonstrated that its interaction frequency was significantly enhanced upon *Smad4* knockout (Figure 3I and Supplementary Figure 1E). Besides, *Smad4* overexpression reduced *Sox2*-associated loops in MSCC^LP.3^ (Figure 3J).

To further investigate how SMAD4 regulates 3D genomic remodeling in human LUSC tumorigenesis, we conducted High-throughput/resolution chromosome conformation capture (Hi-C) analysis of human lung bronchial epithelial cells (BEAS-2B) before and after *SMAD4* knockout in the PTEN-deficiency background, given that lung bronchial epithelial cells are the major cell of origin for LUSC development. Moreover, we previously showed that *Smad4* deletion in PTEN-deficient mouse lung epithelial cells induced adeno-squamous cell tumors with 100% incidence.^[12]^ In contrast, the tumor incidence of Adeno-Cre *KRAS^G12D^ p53^f/f^ Smad4^f/f^* or CCSP^iCre^ *Lkb1^f/f^ Smad4^f/f^* was approximately 25-35% (Figure 1H).

Genome-wide analysis revealed extensive 3D chromatin reorganization upon *SMAD4* loss, affecting numerous genomic regions (Figure 3K). Notably, inspection of the *SOX2* locus uncovered pronounced alterations in chromatin looping interactions directly associated with *SOX2* regulation following *SMAD4* deletion (Figure 3L). These data suggested that SMAD4 loss increased *Sox2* transcription and cell survival in LUSC, partially via enhancing enhancer-promoter interactions through a chromatin loop.

### 2.4 SMAD4 Deficiency Drives Mouse LUSC Oncogenic Transcriptome Program (*e.g.*, *Sox2* overexpression)

Indirectly Remodeling 3D Genome through EP300 To elucidate the molecular mechanism by which SMAD4 indirectly regulates chromatin looping, including the *Sox2*-associated chromatin loop, we first performed motif enrichment analysis at the anchoring regions of the *Sox2*-associated chromatin loop to identify potential loop-associated binding factors (Figure 4A). In parallel, we screened for transcriptional regulators known to interact with SMAD4 from the BioGRID database (Figure 4A). This integrative analysis identified SMAD3, SMAD2, and EP300 as the most significantly enriched candidate proteins (Figure 4A). To further prioritize functional mediators, we analyzed publicly available ChIP-Seq datasets from mouse lung cell lines obtained from the ChIP-Atlas database. Notably, EP300 displayed robust binding at both anchor regions of the *Sox2*-associated chromatin loop (Figure 4B). Co-immunoprecipitation (Co-IP) assays confirmed a direct physical interaction between SMAD4 and EP300 in MSCC^LP.3^ cells (Figure 4C). Moreover, ChIP-qPCR analysis demonstrated significant enrichment of EP300 at both the promoter and distal enhancer regions of *Sox2*, and *Smad4* knockout markedly enhanced EP300 occupancy at these sites (Figure 4D). Given the well-established role of EP300 as a histone acetyltransferase, we next examined histone modification status at the *Sox2*-associated chromatin loop. Consistent with increased EP300 recruitment, SMAD4 deficiency led to a pronounced elevation of H3K27ac levels at both the promoter and the distal enhancer regions of the *Sox2*-associated chromatin loop (Figure 4E). These results indicated that EP300 mediated SMAD4-dependent regulation of *Sox2*-associated chromatin architecture by modulating enhancer activity. To directly assess the functional requirement of EP300 in this process, we generated *Ep300*-knockout cells in a SMAD4-deficiency background (Figure 4F). Loss of EP300 significantly reduced the interaction frequency of *Sox2*-associated chromatin loops, as determined by 3C-qPCR (Figure 4G). Functionally, *Ep300* knockout effectively abrogated the enhanced proliferative capacity induced by SMAD4 loss (Figure 4H). Collectively, these findings supported a model in which SMAD4 restrained EP300 from participating in the formation of *Sox2*-associated chromatin loop through direct protein-protein interaction. Upon SMAD4 loss, EP300 was released and recruited to both the *Sox2* promoter and the distal enhancer, where it promoted enhancer activation, chromatin loop stabilization, and subsequent upregulation of *Sox2* transcription, ultimately driving tumor cell proliferation.

**Figure 4.**
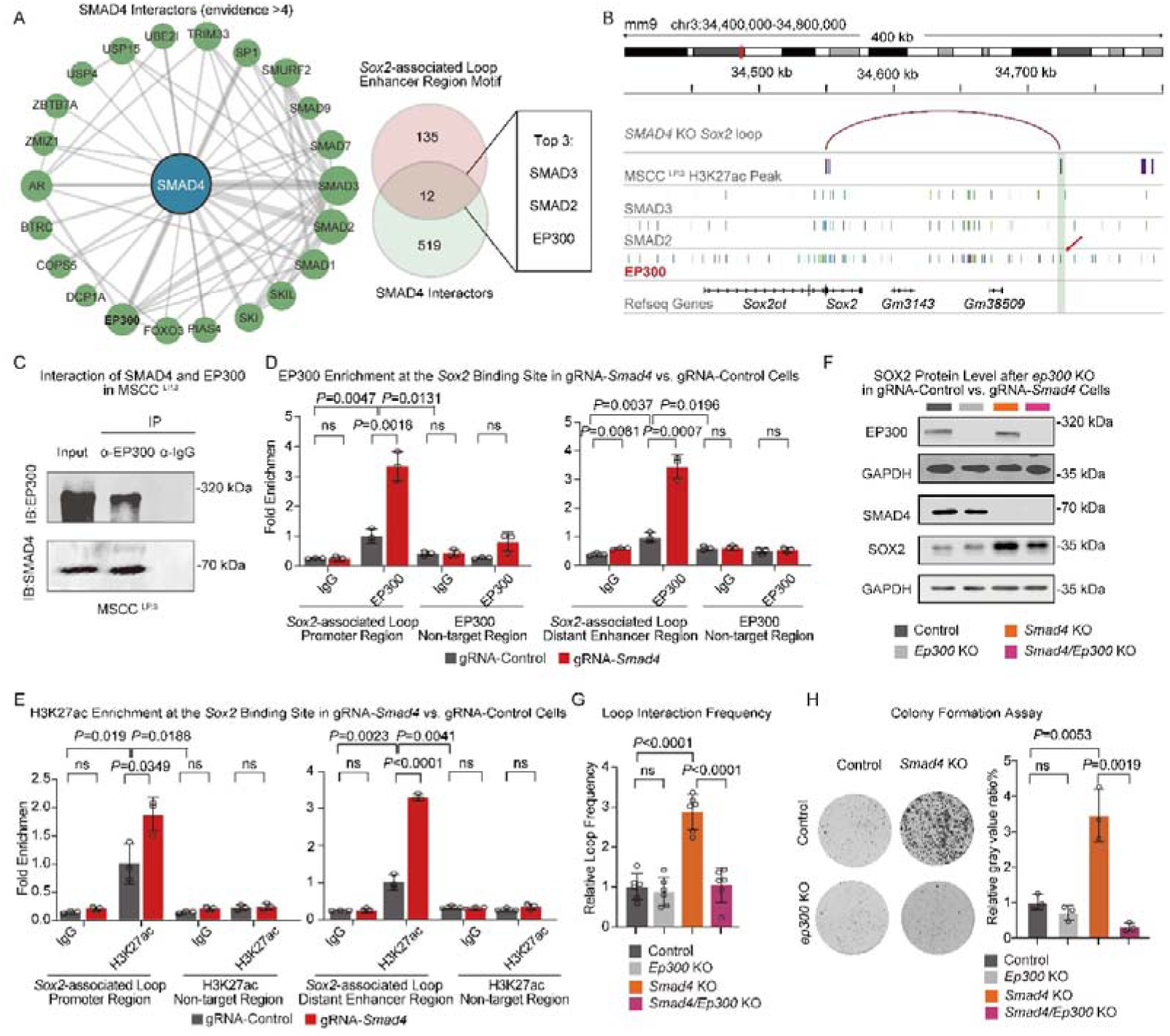
SMAD4 deficiency drives mouse LUSC oncogenic transcriptome program (*e.g.*, *Sox2* overexpression) by indirectly remodeling 3D genome through EP300. (A) Overlap analysis of proteins that interact with the SMAD4 protein and the TFs whose motifs were enriched at *Sox2*-associated loop target regions. The SMAD4 interactors were identified from the BioGRID database. The overlapped proteins were ranked by the evidence of interaction with SMAD4 according to BioGRID. (B) The ChIP-Seq peaks of the top three candidates enriched in *Sox2*-associated loop regions. Anchor regions of the *Sox2*-associated loop were shown in green background, and the EP300 binding site were indicated by red arrows. The ChIP-Seq data were from the ChIP-atlas database, and all collected cell types of the candidate proteins were selected. (C) Co-IP confirmed the interaction between EP300 and SMAD4 in the MSCC^LP.3^ cells. Input served as the positive control to validate antibody binding specificity, while IgG acted as the negative control to rule out nonspecific binding. (D) EP300 enrichment at *Sox2*-associated loop target region on MSCC^LP.3^ gRNA-Control and gRNA-*Smad4* cells detected by ChIP-qPCR. (E) H3K27ac enrichment at *Sox2*-associated loop target region on MSCC^LP.3^ gRNA-Control and gRNA-*Smad4* cell lines detected by ChIP-qPCR. Input served as the positive control to validate antibody binding specificity, while IgG acted as the negative control to rule out nonspecific binding. (F) The knockout efficiency of *Ep300* in MSCC^LP.3^ cells with or without *SMAD4* depletion by Western Blot, and the analysis of SMAD4 and SOX2 expression changes after *Ep300* knockout in these cells. GAPDH served as the loading control. (G) Analysis of the changes in the frequency of *Sox2*-associated loop by 3C-qPCR after *Ep300* knockout in the MSCC^LP.3^ cell line with or without SMAD4 depletion. (H) Colony formation assay after the knock-out of the *Sox2*-associated loop target regions in *Smad4* and/or *Ep300* knockout in the MSCC^LP.3^ cell line. Quantification of the relative gray value ratio was performed using ImageJ.

### 2.5 SMAD4 Deficiency Drives Human LUSC Oncogenic Transcriptome Program (*e.g.*, *SOX2* overexpression) by Indirectly Remodeling 3D Genome through EP300

To investigate whether SMAD4 deficiency indirectly remodels LUSC through EP300 to drive the oncogenic transcriptomic program (*e.g.*, *SOX2* overexpression), we conducted Hi-C analyses of human LUSC samples, which demonstrated a significant enhancement of *SOX2*-associated chromatin looping in tumor tissues (Figure 5A). Integration of EP300 and H3K27ac ChIP-seq data from the ChIP-Atlas database revealed vigorous enhancer activity and EP300 occupancy downstream of the *SOX2* locus, precisely overlapping with the reinforced chromatin loop anchor regions observed in tumors (Figure 5B). To investigate the functions of the *SOX2*-associated chromatin loop in human LUSC, we generated *SMAD4*-knockout cells in the human LUSC cell line H2170. Consistent with our observations in mouse models, loss of SMAD4 led to a pronounced upregulation of *SOX2* at both the mRNA and protein levels (Figures 5C-E). Functionally, *SMAD4* deletion significantly enhanced the colony formation of human LUSC cells (Figure 5F). Concomitantly, 3C-qPCR revealed a marked increase in the interaction frequency of *SOX2*-associated chromatin loop upon *SMAD4* knockout (Figure 5G and Supplementary Figure 1F). To directly assess the functional contribution of the distal enhancer involved in *SOX2*-associated loop formation, we deleted the enhancer region anchoring the *SOX2*-associated chromatin loop in SMAD4-deficient cells (Figure 5H and Supplementary Figure 1G). Disruption of this distal enhancer in SMAD4-deficient H2170 cells led to a substantial reduction in SOX2 expression (Figure 5I) and significantly impaired cell viability (Figure 5J), effectively reversing the oncogenic phenotype induced by SMAD4 loss. In parallel, the Co-IP assay further confirmed the physical interaction between SMAD4 and EP300 in human LUSC cells (Figurfe 5K). Collectively, these results demonstrated that SMAD4 regulated SOX2 expression by remodeling the three-dimensional chromatin architecture through EP300-mediated mechanisms in human LUSC, consistent with the mechanism identified in mouse models.

**Figure 5.**
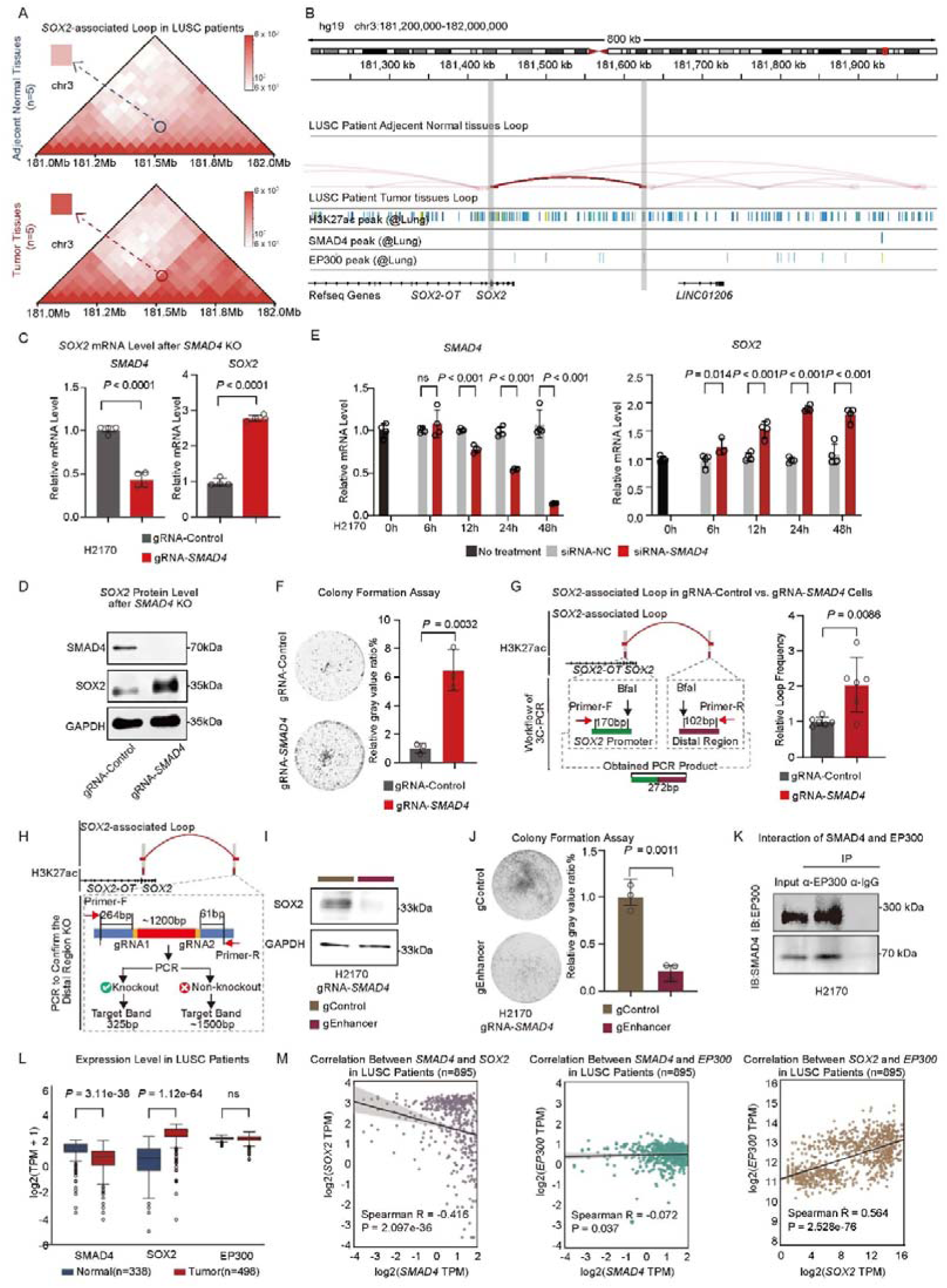
SMAD4 deficiency drives human LUSC oncogenic transcriptome program (*e.g.*, *SOX2* overexpression) by indirectly remodeling 3D genome through EP300. (A) *SOX2*-associated loops detected in Hi-C analysis of LUSC and adjacent normal samples. (B) Published ChIP-Seq data indicated the binding of EP300 and H3K27ac histone modification on *SOX2*-associated loops detected in clinical LUSC samples. (C) RT-qPCR analysis of *SMAD4* and *SOX2* expression in *SMAD4* knockout (gRNA-*SMAD4*) and non-knockout (gRNA-Control) in H2170. (D) Western Blot analysis of *SMAD4* and *SOX2* protein levels in *SMAD4* knockout (gRNA-*SMAD4*) and non-knockout (gRNA-Control) in H2170. (E) *SMAD4* expression levels and *SOX2* expression levels at different time points following *SMAD4* knockdown. (F) Colony formation assay in H2170 cell line with or without SMAD4 depletion and the quantification of the relative gray value ratio using ImageJ. (G) 3C-qPCR analysis of the relative loop frequency in H2170 cell lines with or without *SMAD4* knockout. (H) Schematic diagram of the *SOX2*-associated enhancer knockout strategy. (I) Western Blot analysis of *SOX2* expression in the H2170 cells with or without the knockout of the enhancer of the *SOX2*-associated loop. (J) Colony formation assay in the H2170 gRNA-*SMAD4* cells with or without the knockout of the enhancer of *SOX2*-assocaited loop, and the quantification of the relative gray value ratio using ImageJ. (K) Co-IP confirmed the interaction between EP300 and SMAD4 in the H2170 cells. Input served as the positive control to validate antibody binding specificity, while IgG acted as the negative control to rule out nonspecific binding. (L) Expression level of *SMAD4*, *EP300*, and *SOX2* in clinical LUSC and normal samples. (M) The expression relevance between *SMAD4*, *EP300*, and *SOX2* in LUSC samples.

To further evaluate the clinical relevance of the SMAD4-EP300-SOX2 axis, we analyzed publicly available transcriptomic datasets from LUSC patient samples. Compared with normal lung tissues, *SMAD4* expression was significantly downregulated, whereas *SOX2* expression was markedly upregulated in tumor samples (Figure 5L). In contrast, *EP300* expression showed no significant difference between tumor and normal tissues, indicating that SMAD4 modulates EP300 function primarily at the chromatin level rather than through transcriptional regulation (Figure 5L). Correlation analysis revealed a significant negative correlation between *SMAD4* and *SOX2* expression (Figure 5M), consistent with the positive correlation between *EP300* and *SOX2* expression (Figure 5M). Moreover, no significant correlation was observed between *SMAD4* and *EP300* expression, further supporting a regulatory model independent of *EP300* transcriptional abundance (Figure 5M).

### 2.6 SMAD4-dependent Chromatin Looping Regulates a Subset of Oncogenic Genes in LUSC

Although *SOX2* emerged as a prominent SMAD4-regulated target in our analyses, our data indicated that it might not represent an isolated regulatory event. Integrative analysis of Hi-C and RNA-Seq datasets identified a set of 34 genes (in addition to *SOX2*) whose transcriptional changes were accompanied by alternations of SMAD4-loss-associated chromatin loops (Figure 6A). Importantly, similar looping patterns at these loci were also observed in LUSC patient samples (Figure 6B). These results suggested that this regulatory mode might not be restricted to experimental models but reflected a disease-relevant chromatin regulatory program. At the *PRR19* locus, for example, we found that SMAD4 deficiency strengthened enhancer-promoter contacts, accompanied by increased gene expression (Figures 6A-B). Notably, the same loop configuration was detected in primary tumor samples (Figure 6B). Together, these findings substantiate the clinical relevance of the SMAD4-EP300-SOX2 axis and highlight SMAD4-regulated 3D genome reorganization as a conserved and pathogenic mechanism driving *SOX2* activation in LUSC (Figure 6C).

**Figure 6.**
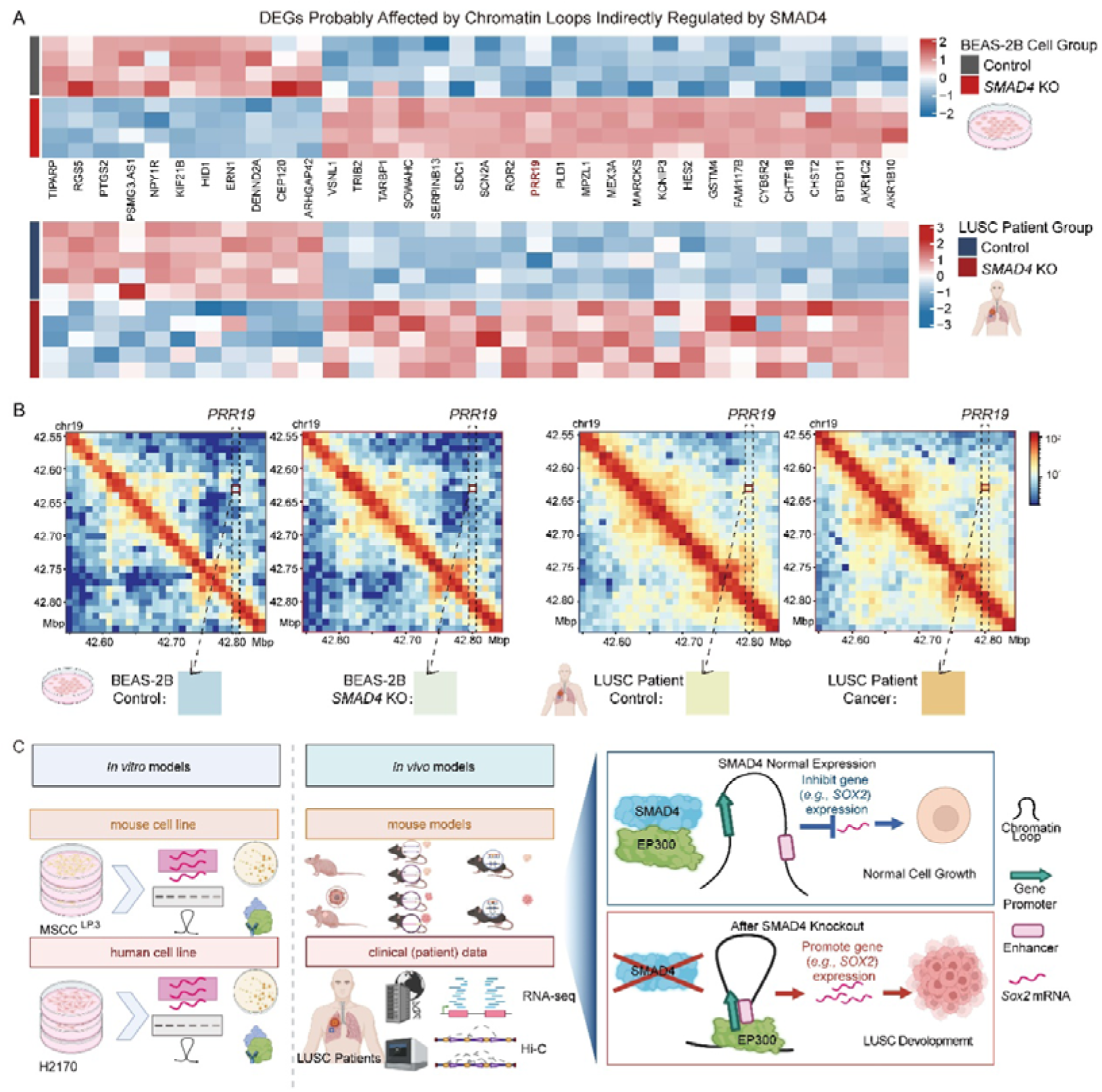
SMAD4-dependent chromatin looping regulates a subset of oncogenic genes in LUSC. (A) DEGs regulated by remodeled chromatin loops indirectly regulated by SMAD4. (B) *PRR19*-associated loops detected in BEAS-2B with or without *SMAD4* KO cells and clinical LUSC samples. (C) Working model.

## 3. Discussion

LUSC remains a clinically challenging malignancy due to largely unclear key regulators and underlying mechanisms. In this study, we identified a previously unrecognized role of SMAD4 as a guardian of three-dimensional (3D) genome architecture in LUSC, and uncovered a chromatin-based mechanism by which SMAD4 loss indirectly activated the lineage-defining oncogene *SOX2* through EP300-mediated enhancer-promoter looping. This revealed a non-canonical role for a TF in regulating the dysregulated 3D genome, rather than direct binding, to affect tumor development. Additionally, our data indicated that SMAD4 loss did not merely affect isolated enhancer-promoter contacts but also induced alterations across multiple hierarchical levels of genome architecture. Notably, approximately 8.9% of the genome exhibited compartment switching upon *SMAD4* deficiency, suggesting that SMAD4 contributes to maintaining large-scale chromatin spatial segregation (Supplementary Figures 2B-C). However, our study focused on LUSC tumor-relevant regulatory circuits (*e.g.*, *SOX2* induction) rather than providing a comprehensive catalog of architectural alterations. Consequently, we provided the brief global profiling here to stay our focus on the non-canonical role of SMAD4 in regulating the 3D genome to repress LUSC tumorigenesis.

Our work expanded the functional repertoire of SMAD4 beyond its canonical role as a TGF-β-responsive TF. While SMAD4 has long been recognized as a tumor suppressor that directly regulates gene expression through promoter-proximal binding, accumulating evidence suggests that transcriptional dysregulation alone cannot fully explain the profound tumor-promoting effects observed upon SMAD4 loss. Here, by integrating mouse models, human cell lines, and clinical datasets, we demonstrated that SMAD4 deficiency induced widespread reorganization of higher-order chromatin structure, with particularly pronounced effects at the *SOX2* locus. These findings support a paradigm in which tumor suppressor loss rewires the spatial genome to unlock oncogenic transcriptional programs, rather than acting solely through linear regulatory mechanisms.

A key insight from this study was that SMAD4 regulated chromatin looping indirectly, despite its preferential localization to distal genomic regions. Unexpectedly, SMAD4 did not directly occupy the *SOX2*-associated enhancer or promoter, nor did it broadly bind active enhancers. Instead, SMAD4 physically interacted with EP300, a master histone acetyltransferase and architectural cofactor, and restrained its recruitment to loop anchor regions. Upon SMAD4 loss, EP300 was released and preferentially accumulated at both ends of the *SOX2*-associated chromatin loop, leading to enhanced H3K27ac deposition, stabilized enhancer-promoter interactions, and robust *SOX2* activation. This mechanism revealed an additional layer of chromatin regulation in which tumor suppressors constrained the looping capacity of architectural cofactors, thereby maintaining transcriptional homeostasis.

*SOX2* amplification and overexpression are among the most recurrent and functionally important events in LUSC. Although previous studies have implicated TFs, super-enhancers, and epigenetic modifiers in *SOX2* regulation, these models have focused mainly on linear chromatin features. Our findings demonstrated that *SOX2* activation in LUSC was critically dependent on aberrant 3D genome organization and that enhancer-promoter looping constituted a decisive regulatory step downstream of the loss of tumor suppressors. The consistency of our observations across mouse and human systems, together with similar alterations detected in two additional mouse models of distinct genetic backgrounds, suggests that the SMAD4-*SOX2* axis may have broader relevance in LUSC. At the same time, given the molecular heterogeneity of LUSC, it remains possible that this regulatory relationship is modulated by context-specific factors in selected genetic backgrounds. Significantly, disruption of the *SOX2*-associated chromatin loop or deletion of its distal enhancer effectively reversed the proliferative advantage conferred by SMAD4 deficiency, highlighting chromatin looping as a functional, rather than merely correlative, driver of tumor progression. Additionally, deletion of the *Sox2*-associated distal enhancer reduced *Sox2* expression even in *Smad4*-wild-type MSCC^LP.3^ cells, indicating that this enhancer provides essential positive regulatory input for basal *Sox2* transcription (Supplementary Figure 1H). Consistently, *Smad4* overexpression further suppressed *Sox2* expression and weakened enhancer-promoter interactions (Figure 2H and Figure 3J), whereas *Smad4* loss strengthened chromatin looping and activated *Sox2* (Figure 2F and Figure 3I). These findings indicated that SMAD4 did not function as a binary repressor of *Sox2*, but rather acted as a quantitative modulator that constrained enhancer-mediated chromatin interactions to maintain *Sox2* expression within a controlled range. Our data indicate that SMAD4 loss enhances *SOX2*-associated enhancer-promoter looping and is accompanied by increased *SOX2* transcription, supporting a functional connection between chromatin architecture and transcriptional output in this context. However, the current study does not resolve whether looping reinforcement temporally precedes transcriptional activation or whether these events occur in a tightly coupled manner.^[23,24]^ As enhancer-promoter interactions are thought to establish a permissive regulatory configuration, yet are themselves highly dynamic, we favor a model in which SMAD4 deficiency promotes a chromatin state conducive to *SOX2* activation, with looping reinforcement and transcriptional induction representing mechanistically linked and dynamically coordinated processes. Given that SMAD4 is a TGF-β-responsive TF, we further investigated whether the mechanism by whether SMAD4 regulation on *Sox2* was subject to TGF-β control. Results showed that the effects of *Smad4* deletion and enhancer disruption on *SOX2* transcriptional regulation persisted in the presence or absence of TGF-β (Supplementary Figure 2D).

The clinical relevance of the SMAD4-EP300-SOX2 axis was further supported by analyses of human LUSC datasets. *SMAD4* expression was consistently reduced in tumors, whereas *SOX2* was upregulated, accompanied by the strengthened *SOX2*-associated chromatin loops in patient samples (Figure 5A and Figure 5L). Importantly, *EP300* expression itself was not altered, reinforcing the notion that SMAD4 modulated EP300 function at the chromatin level rather than through transcriptional control. This distinction was particularly relevant from a therapeutic perspective, as it suggested that targeting EP300 activity or enhancer-promoter interactions might selectively impair oncogenic transcriptional circuits without globally suppressing EP300 expression. We noted that the number of human LUSC Hi-C samples analyzed here was limited, and future studies incorporating larger patient cohorts can fully investigate inter-patient heterogeneity.

More broadly, our findings have implications beyond LUSC. The relevance of the SMAD4-*SOX2* axis may extend beyond LUSC to other squamous cell carcinomas. *SOX2* frequently acts as an oncogene through amplification or overexpression in multiple squamous malignancies,^[25–27]^ whereas loss of *SMAD4* is commonly implicated in squamous tumorigenesis through attenuation of tumor-suppressive pathways.^[28,29]^ Besides, we additionally examined published single-cell RNA-seq datasets^[30,31]^ from head and neck squamous cell carcinoma (HNSCC) and esophageal squamous cell carcinoma (ESCC) (Supplementary Figures 3A-B). Notably, in malignant tumor cell populations from both cancer types, *SMAD4* and *SOX2* expression showed a negative correlation, consistent with the trend observed in LUSC. These observations suggest that SMAD4-dependent tumor-suppressive pathways may converge on *SOX2* regulation more broadly in squamous epithelial cancers. However, whether this occurs through the same chromatin-looping mechanism across distinct squamous tumor types remains to be determined. The concept that chromatin loop homeostasis was actively maintained by tumor suppressors added a new dimension to our understanding of cancer epigenetics, suggesting that dysregulated genome topology might be an early and permissive event in tumor evolution.

Although EP300 and CREBBP are generally considered functionally transcriptional co-activators, our data suggested a locus-selective role for EP300 at the *SOX2* regulatory element. CBP/CREBBP did not show consistent enrichment at SMAD4-dependent chromatin loop anchors, whereas EP300 was robustly associated with the *SOX2* enhancer region (Supplementary Figure 2F). The loop-associated regions displayed elevated H3K27ac levels (Figure 4E), consistent with EP300 acetyltransferase activity, and perturbation of EP300 markedly attenuated *SOX2* activation and weakened enhancer-promoter interaction strength (Figure 4G). Together, these findings indicated that EP300 functioned not merely as a co-activator but as a principal chromatin effector mediating loop-associated transcriptional activation in this context. While other co-activators may contribute at additional loci, the selective occupancy and functional dependence observed here support a non-redundant role for EP300 in the SMAD4-regulated *SOX2* transcriptional program. Therefore, although a systematic dissection of co-activator usage at SMAD4-regulated chromatin domains will require broader comparative analyses, it is still worth conducting future research.

In addition to restraining EP300 redistribution to oncogenic enhancers, an alternative but not mutually exclusive possibility was that SMAD4 might contribute to the context-dependent positioning of EP300 at SMAD4 complex-occupied regulatory regions, such as *Zfp654* and *Cggpb1* gene promoter regions (Supplementary Figure 2E). In this case, SMAD4-containing complexes might help maintain EP300 activity within appropriate regulatory domains, thereby preventing promiscuous activation.

Loss of *SMAD4* could therefore lead not only to derepression of EP300 at non-canonical enhancers but also to impaired EP300 function at canonical SMAD4-associated regulatory elements, collectively contributing to transcriptional imbalance during tumorigenesis.

Although the functional importance of the SMAD4-EP300 interaction was strongly supported by both our data and previous studies, the precise molecular and structural basis of this interaction remained to be fully defined. Multiple independent studies curated in BioGRID reported a physical association between SMAD4 and EP300, including early biochemical evidence demonstrating that the C-terminal MH2 domain of SMAD4 interacts with the C-terminal region of EP300^[32]^. While our study established the functional relevance of this interaction in mediating enhancer-promoter looping and 3D genome reorganization, detailed mapping of the interaction interfaces and high-resolution structural characterization of the SMAD4-EP300 complex will be important future directions to elucidate the mechanistic basis of this non-canonical regulatory pathway.

Because EP300 lacks intrinsic DNA-binding activity, its association with specific genomic loci involves additional regulatory mechanisms. Our results indicated that SMAD4 restricted aberrant chromatin looping primarily by limiting EP300 engagement with chromatin, rather than through direct DNA binding at *SOX2* regulatory regions. In this context, sequence-specific transcription factors might provide auxiliary specificity for EP300 recruitment. Consistent with this notion, motif enrichment analysis revealed an enrichment of SP1-binding motifs at *SOX2*-associated chromatin loop anchors (Supplementary Figure 2F), suggesting SP1 occupancy at these regions. SP1 has been reported to facilitate the EP300 recruitment. Although we did not position SP1 as a central regulator in this study, these observations raised the possibility that SP1 contributed to locus-specific EP300 positioning once the SMAD4-mediated constraint on EP300 was relieved. Under this model, SMAD4 acts as the dominant upstream regulator, preventing inappropriate EP300-chromatin interactions, while SP1 may provide local sequence guidance to fine-tune EP300 engagement at selected regulatory elements. In future research, it would be warranted to conduct experimental dissection to determine the extent to which SP1 or other transcription factors participate in this process. Mechanistically, our co-IP and EP300 ChIP-qPCR data support a model in which SMAD4 suppresses enhancer activity by directly associating with EP300 and limiting its recruitment to specific chromatin loci. Given the potential role of SP1 in EP300 recruitment, SMAD4 may antagonize SP1-dependent EP300 loading at enhancers such as the Sox2-associated enhancer.

From a translational perspective, although pharmacological inhibition of EP300 acetyltransferase activity can effectively suppress enhancer-driven oncogenic transcription, the broad requirement of EP300 across diverse transcriptional programs raises concerns regarding specificity and potential toxicity. Our findings instead indicated that in SMAD4-deficient LUSC, *SOX2* activation was driven by selective strengthening of specific enhancer-promoter loops rather than global locus activation. It suggested that targeting disease-specific regulatory interactions might represent a more precise therapeutic strategy. In this context, the *SOX2*-centered enhancer hub identified in *SMAD4*-deficient cells may constitute a structural vulnerability that can be selectively targeted to attenuate oncogenic transcription while preserving basal chromatin architecture and physiological gene regulation. This translational possibility was partially evidenced by our sequence-specific oligonucleotides targeting the *Sox2* enhancer region to inhibit *Sox2* expression (Figure 3H).

In summary, this study established SMAD4 as a critical regulator of chromatin loop homeostasis in LUSC development, revealing a previously unappreciated epigenetic mechanism by which tumor suppressor loss indirectly activated oncogenic transcriptional programs. Although our study mainly examined proliferative phenotypes, transcriptomic analysis suggested broader effects of SMAD4 loss. For example, differentially expressed genes were enriched in epithelial differentiation and cell fate programs (Supplementary Figure 2A). Thus, SMAD4-mediated repression of *Sox2* likely regulates not only tumor growth but also cellular identity. By linking SMAD4 deficiency to EP300-driven enhancer-promoter looping and *SOX2* activation, our work provided new mechanistic insights into LUSC development and highlighted 3D genome regulation as a promising therapeutic vulnerability in LUSC.

## 4. Experimental Methods

### Mice

All Mice were maintained under specific pathogen-free (SPF) conditions with free access to food and water. The *Smad4* floxed mice, *KRAS^G12D^* knock-in mice, and *Trp53* floxed mice were bred and maintained at the Shanghai Key Laboratory of Regulatory Biology, East China Normal University. Experiments were operated following the ethical standards by the Animal Center at Shanghai Key Laboratory of Regulatory Biology. To generate *Trp53*^f/f^ *KRAS^G12D^* and *Smad4*^f/f^ *Trp53*^f/f^ *KRAS^G12D^* models, 6-8-week-old mice were administered 1 × 10□ pfu Adeno-Cre via intranasal instillation, as previously described.^[22]^

The CCSP*^iCre^* strain was generated by us previously. ^[12]^ *Lkb1*^f/f^ (FVB;129S6 *Stk11* tm1Rdp/Nci) mice were obtained from the NCI Frederick Mouse Repository, and *Pten*^f/f^ (C;129S4-*Pten* tm1Hwu/J) mice were purchased from The Jackson Laboratory. *Smad4*^f/f^ (*Smad4* RobCA) mice were obtained from Dr. Elizabeth J. Robertson.

CCSP*^iCre^* mice were crossed with these strains to generate conditional knockout models in the lung epithelium. All strains were maintained on a B6.129 background.^[6,12]^

For xenograft assays, nude mice (GemPharmatech, China) were kept under SPF conditions. MSCC^LP.3^ gRNA-Control and gRNA-*Smad4* cells (5 × 10□ cells in 100 µL Matrigel) suspended at 5 × 10□ cells in 100 µL of Matrigel (Corning, Cat. #356237), were subcutaneously injected into nude mice. Tumors were harvested after 3 weeks, fixed in 4% paraformaldehyde, paraffin-embedded, and processed for H&E and IHC staining following standard protocols.

### Histopathology and immunohistochemistry

Paraffin-embedded lung sections (5 μm) were deparaffinized, rehydrated through graded ethanol, and subjected to H&E staining using standard protocols for histological evaluation.

For IHC staining, deparaffinized sections underwent antigen retrieval in Antigen Unmasking Solution (Vector Laboratories, H3300) and inactivation of endogenous peroxidase with 3% H□O□. After blocking with 1% BSA, sections were incubated with primary antibodies against SMAD4 (1:1000, CST#38454), SOX2 (1:250, CST#14962), KRT5 (1:1000, ab52635), TTF1 (1:500, DAKO IR056), and TP63 (1:1000, CST#39692) overnight at 4 °C. Corresponding biotin-labeled secondary antibodies were applied, and signals were developed using a DAB substrate kit (Vector Laboratories, SK-4105). Slides were counterstained with hematoxylin, dehydrated, mounted, and scanned with a scanner (3DHISTECH, PANNORAMIC MIDI).

### Real-time quantitative PCR and western blotting

Total RNA was extracted from cells using TRIzol method followed by purification using AG RNAex Pro Reagent (Accurate Biotechnology, AG21102) and purified according to the manufacturer’s protocol. RNA quality was assessed with a NanoDrop spectrophotometer (Thermo Fisher Scientific, NanoDrop One). Extracted RNA was reverse-transcribed into cDNA using the HiScript® III All-in-one RT SuperMix Perfect for qPCR (Vazyme, R333). Quantitative real-time PCR was performed with the ChamQ Universal SYBR qPCR Master Mix (Vazyme, Q711) followed the manufacturer’s protocol. Primer sequences are listed in Supplementary Table 4.

For protein analysis, cells or tissues were lysed in RIPA buffer containing 1% PMSF (Beyotime, ST506) on ice, and protein concentrations were measured using the BCA Protein Assay Kit (Beyotime, P0012). Equal amounts of protein (20 µg) were resolved on 6%-10% SDS-PAGE gels and transferred to nitrocellulose membranes (MerckMillipore, HATF00010). After blocking with 5% non-fat milk for 1 hour at room temperature (RT), membranes were incubated overnight at 4 °C with primary antibodies against SMAD4 (1:1000, CST#38454), SOX2 (1:250, CST#14962), EP300 (1:500, Sigma-Merk 05-257), GAPDH (1:1000, CST#2118), or β-Actin (1:500, HANGZHOU FUDE BIOLOGICAL TECHNOLOGY CO., LTD., FD0060), followed by HRP-conjugated secondary antibodies (1:5000, Easy bio, BE0101 and BE0102). Signals were detected using Western Lighting Plus (Pekin Elmer, NEL105001EA) and visualized with the Odyssey FC Imaging System (LI-COR Biosciences, 2800 S/N OFC-1403).

### 3C-PCR and 3C-qPCR

2-5 × 10□ cells were crosslinked with 1% paraformaldehyde for 10 min at RT, and the reaction was quenched with 2 M glycine. Fixed nuclei were isolated using lysis buffer and digested overnight at 37 °C with BfaI restriction enzyme (New England Biolabs, R3539S). After enzyme inactivation, chromatin fragments were ligated with T4 DNA ligase (TaKaRa, SD0267) at 16 °C for 2 h and at RT for another 2h, followed by reverse crosslinking with Proteinase K (Shanghai LuoWen Biotechnology, LWK0301) at 55 °C overnight. DNA was purified by ethanol precipitation and resuspended in TE buffer.

The interaction between genomic loci was detected using PCR. For 3C-PCR, 5 ng of 3C template DNA was amplified using locus-specific primers under standard cycling conditions (94 °C 5 min; 40 cycles of 94 °C 30 s, 60 °C 30 s, 72 °C 30 s; final extension at 72 °C 5 min). PCR products were analyzed on 1.5% agarose gels. For quantitative analysis, 3C-qPCR was performed using ChamQ Universal SYBR qPCR Master Mix (Vazyme, Q711) according to the manufacturer’s instructions. The relative crosslinking frequency was calculated after normalization to the control. Primer sequences are listed in Supplementary Table 4.

### ChIP-qPCR

Approximately 1 × 10□ cells were crosslinked with 1% paraformaldehyde for 10 min at RT, quenched with 2 M glycine, and washed twice with cold PBS. Cell nuclei were isolated using hypotonic buffer and lysed in freshly prepared lysis buffer supplemented with protease inhibitors (Sigma, 539128). Chromatin was sheared to 200-500 bp fragments using a Covaris Focused-Ultrasonicator (Covaris, S220) under optimized conditions for transcription factors (EP300) or histone marks (H3K27ac).

The sheared chromatin was incubated overnight at 4°C with 4 μg of specific antibody, including anti-EP300 (Sigma-Merck, 05-257) and anti-H3K27ac (Abcam), or with normal IgG pre-bound to Protein A/G magnetic beads. Immunocomplexes were sequentially washed with low-salt, high-salt, LiCl, and TE buffers, followed by elution and reverse crosslinking at 65°C overnight. DNA was purified using phenol-chloroform extraction and ethanol precipitation.

Purified DNA was quantified using Qubit (Invitrogen, Qubit 4 Fluorometer), and enrichment of specific genomic regions was analyzed by quantitative PCR with ChamQ Universal SYBR qPCR Master Mix (Vazyme, Q711) following the manufacturer’s instructions. The relative enrichment was calculated after normalization to input DNA. Primer sequences are listed in Supplementary Table 4.

### Co-Immunoprecipitation

Cells were lysed in RIPA buffer supplemented with 1% PMSF on ice for 15 min, and the lysates were clarified by centrifugation. Protein concentrations were determined using a BCA assay. For each reaction, 400-600 µg of total protein were incubated overnight at 4 °C with 1-4 µg of the indicated primary antibody or control IgG, followed by incubation with Protein A/G magnetic beads for 1 h. The immune complexes were washed three times with RIPA buffer and once with cold PBS, then eluted in 2× loading buffer by boiling at 100 °C for 10 min. Co-precipitated proteins were analyzed by Western blot.

### Cell culture and cell colony formation

MSCC^LP.3^ cell line was isolated from CCSP*^icre^ Lkb1*^f/f^ *Pten*^f/f^ mouse by following a cancer cell isolation kit protocol, which has been published previously^[6]^. BEAS-2B and H2170 cells were purchased from ATCC. The completed medium for culturing MSCC^LP.3^ related cell lines was DMEM (Hang Zhou Ke Yi, C006) with 10% FBS (Excell Bio, FSP500) and 1% 100 × P/S (Beyotime Biotechnology, C0222).

BEAS-2B was cultured using the BEGM kit (Lonza, No. CC-3170). H2170 related cells were cultured using RPMI 1640 (Hang Zhou Ke Yi, C003) with 10% FBS and 1% 100 × P/S. Cells were seeded into 6-well plates. For MSCC^LP.3^ related cell lines, 1,000 cells were seeded per well, while for H2170 related cell lines, 2,000 cells were seeded per well and they were maintained in culture for 10-12 days. After fixation with 4% paraformaldehyde for 20 min, colonies were stained with 1% crystal violet and washed with distilled water. The quantitative analyses were performed using ImageJ Fiji software.

### Plasmids and cell lines generation

The gRNA sequences are included in Supplementary Table 5. These gRNAs were cloned into the LentiCRISPRv2 vector.^[33]^ CRISPR-Lenti nontargeting control plasmid was purchased from MilliporeSigma (Billerica, MA, CRISPR12-1EA). Plasmids Lenti-gRNA-*Smad4* and Lenti-gRNA-*SMAD4* were used to knockout *Smad4* on MSCC^LP.3^ cell line and knockout *SMAD4* on H2170, H157, BEAS-2B cell lines. Plasmid Lenti-gRNA-*PTEN* was used to knockout *PTEN* on the BEAS-2B cell line. Plasmid extraction was conducted following the protocol of the TIANprep Mini Plasmid Kit (TIANGEN, DP103).

Lentiviral particles were produced by transfecting HEK293T cells with LentiCRISPRv2 constructs, psPAX2 (Addgene, 12260), and pMD2.G (Addgene, 12259) plasmids using Polyethylenimine, Linear (PEI 40000) (YEASEN, 40816ES02). Viral supernatants were collected, filtered, and used to infect target cell lines in the presence of 10 µg/mL polybrene (YEASEN, 40804ES). Infected cells were selected with 2 µg ml^-1^ puromycin (Beyotime, ST551) until stable populations were established. The pooled gRNA-infected cells were subjected to RT-qPCR and Western blot to validate knockout efficiency, with gRNA-Control-infected cells serving as the control. For the construction of *Smad4*-overexpressing cells, 293T cells were co-transfected with pLV3-CMV-*Smad4* (mouse)-3×FLAG-CopGFP-Puro (Miaolingbio, P63084), psPAX2 and pMD2.G. Cells infected with pLV3-CMV-MCS-3×FLAG-CopGFP-Puro (Miaolingbio, DB00135) served as the corresponding control.

### Sox2-associated loop targeted region knockout cell lines generation

For the knockout of the loop distal anchor regions, the gRNA was designed upstream and downstream of the distal anchor regions within 100-200 bp, respectively, based on the precise location of the distal anchor regions identified by 3C-PCR for *Sox2*-associated loop. The gRNA sequences are included in Supplementary Table 5. These gRNAs were cloned into the Plenti-H1-hU6-hygro-dual-gRNA vector.

Plasmids Plenti-H1-hU6-hygro-dual-gRNA-Control and Plenti-H1-hU6-hygro-dual-gRNA-*Sox2* loop were used to knockout targeted region of *Sox2*-associated loop on MSCC^LP.3^ SMAD4 ablation and non-ablation cells. Following plasmid transfection and viral infection, the infected cells were selected using 0.4 mg/mL hygromycin (Solarbio, H8081) until stable populations were established. Genomic DNA was extracted using the TIANamp Genomic DNA Kit (Vazyme, DC301) following its protocol. Target regions flanking the sgRNA editing sites were amplified using Rapid Taq Master Mix (Vazyme, P222100) and gene-specific primers. PCR products were analyzed by 1.5% agarose gel electrophoresis. The primer design and cell line construction for *Ep300* knockout also followed the same principles and procedures, verified by RT-qPCR and Western blot.

### siRNA knockdown

MSCC ^LP.3^ cells were seeded into 6 wells plate. After 24 hours, cellular confluence reaches 60∼70%, wash cells twice with 1 × PBS. Prepared siRNA and RNA transfection reagents following the TSnanofect RNAi (Tsingke, DLH101). Cells were incubated with siRNA for 0□h, 6□h, 12□h, 24□h, or 48□h before harvesting for analysis. All siRNA oligos were purchased from the company Tsingke. The siRNA oligo sequences are included in Supplementary Table 6.

### TGF-β1 and ASO treatment

Cells were seeded into 6-well plates. After overnight treatment with serum-free medium, cells were incubated in medium containing 5 ng/mL TGF-β1 (MCE, HY-P7118) for 0 h, 0.5 h, or 4 h before harvesting for analysis; the concentration of TGF-β1 was determined based on the study by Jian, et al.^[12]^. Cells were also incubated with 15 nM ASOs for 48 h before harvesting, a protocol for ASO utilization adapted from previous studies.^[34,35]^ All ASOs were purchased from GenePharma, and their specific sequences are included in Supplementary Table 7.

### Experiments quantification and statistical analysis

Colony formation grayscale quantification and Western blot image analyses were performed using ImageJ Fiji software. RT-qPCR and colony formation quantified results were analyzed with GraphPad Prism 9. Values were shown as mean ± SD.

### WES analysis

Somatic variants (SNVs and indels) were called using MuTect2 (GATK v.4.2.0.0).^[36]^ Variants flagged as ‘PASS’ by the Mutect2 filter were retained and further processed with GATK’s FilterMutectCalls and SelectVariants tools. Annotation was performed using vcf2maf against the Variant Effect Predictor (VEP100) database. Additionally, variant calling was performed with VarScan2, applying a somatic P□value cutoff of ≤0.05. For both callers, a minimum sequencing depth of 30× and at least 10 supporting reads were required per genomic region.

### RNA-seq and pathway enrichment analysis

Total RNA of cell lines was extracted by the TRIzol method, and libraries were constructed according to a standard protocol (Illumina). All RNA-seq libraries were prepared in the same batch and sequenced on the same platform to minimize technical variation. Raw sequencing reads were initially subjected to adapter removal and quality filtering using Trim Galore (v0.6.10)^[37]^ with default parameters. Quality control metrics, including mapping rate, duplication level, and fragment size profile, were evaluated for each sample. The resulting high-quality reads were aligned to the human reference genome (hg19, GENCODE release) using HISAT2 (v2.2.1)^[38]^ under default settings. PCR duplicate reads were subsequently identified and removed with Picard MarkDuplicates (v2.18.29).^[39]^ To normalize sequencing depth across samples, the deduplicated BAM files were randomly subsampled to an equal number of reads using SAMtools (v1.6).^[40]^

Gene expression quantification for RNA-seq data was performed with featureCounts (v2.0.3)^[41]^ from the Subread package, using the parameters -p -t gene -f -g gene_name in conjunction with the GENCODE hg19 gene annotation. For lncRNA-seq datasets, expression levels were quantified using featureCounts with a RepeatMasker-based lncRNA annotation, applying the parameters -M --fraction -p -T 8 -t exon -f -g gene_id.

Differential expression analysis was conducted using DESeq2, which models count data based on a negative binomial distribution via the DESeq and results functions. Heatmaps illustrating differentially expressed genes or repetitive elements were generated using the ComplexHeatmap R package (v2.14.0). Functional enrichment analysis of differentially expressed genes was carried out using the enrichGO and gseGO function implemented in clusterProfiler (v4.6.2).^[42]^

### CUT&RUN analysis

Perform the CUT&RUN (nuclease-targeted cleavage and release) experiment according to the operating procedures for the Hyperactive pG-MNase CUT&RUN Kit (Vazyme, HD102). Quality control metrics, including mapping rate, duplication level, and fragment size profile, were evaluated for each sample. All CUT&RUN libraries were prepared in the same batch and sequenced on the same platform to minimize technical variation. Raw CUT&RUN sequencing reads were first processed using Trim Galore (v0.6.10)^[37]^ with default parameters to remove adapters and low-quality bases. The trimmed reads were then aligned to the human reference genome (hg19) using Bowtie2 (v2.5.1)^[43]^ with the following options: --end-to-end --very-sensitive --no-mixed --no-discordant --phred33. PCR duplicates were removed from the aligned reads using Picard MarkDuplicates (v2.18.29)^[39]^ with default settings.Signal tracks were generated by converting deduplicated BAM files into BigWig format using bamCoverage from the deepTools package (v3.3.1)^[44]^ with the parameters -of bigwig -bs 10 -e 0 --ignoreDuplicates, and negative signal values were set to zero. CUT&RUN signal profiles were computed and visualized using computeMatrix and plotProfile implemented in deepTools (v3.5.1). BigWig tracks were visualized using IGV (v2.4.13).^[45]^ Peak calling for CUT&RUN data was performed using MACS2,^[46]^ and differential peak enrichment between groups was assessed using the DiffBind R package. To improve comparability across samples, sequencing depth was normalized prior to differential peak and signal comparison.

### Hi-C analysis

Five LUSC patients who underwent surgical resection at the Affiliated Hangzhou First People’s Hospital, School of Medicine, Westlake University, between August 2015 and October 2021 were prospectively enrolled in this study. Clinicopathological information, including age, sex, tumor size, location, histological type, and TNM stage, was collected. The study was approved by the Ethics Committee of the Affiliated Hangzhou First People’s Hospital, School of Medicine, Westlake University, and written informed consent was obtained from all participants. The Hi-C library construction of patients’ samples, BEAS-2B gRNA-*PTEN* and BEAS-2B gRNA-*PTEN* gRNA-*SMAD4* cell lines was performed using the Arima-HiC Kit (Arima, A51108) protocol.

All Hi-C libraries were prepared in the same experimental batch and processed using a uniform sequencing workflow to minimize technical variation. All samples were analyzed using a consistent Hi-C processing pipeline. To improve comparability across samples, the number of valid interaction pairs was unified prior to downstream analyses, and contact matrices were normalized using the iterative correction and eigenvector decomposition (ICE) method. The raw data is aligned to the human reference genome (hg19) and processed by HiC-Pro^[47]^ standard pipeline. Interaction matrices derived from .hic files were visualized using Juicerbox (v2.20.00)^[48]^ and HiCExplorer.^[49]^ Chromatin loops were detected using the hiccups algorithm within the Juicer pipeline. The strength of identified loops was statistically assessed and visualized with Genova, and differential loop analysis was performed by extracting loop-associated interaction intensities using cooltools.

### Batch effect control and multi-omics integration

Potential batch effects were considered and controlled primarily within each omics modality rather than through a single post hoc correction step across platforms.

RNA-seq, H3K27ac CUT&RUN, and Hi-C data were processed independently using modality-specific pipelines, with matched experimental design, sample-level quality control, and appropriate normalization procedures applied within each data type.

Cross-platform integration was performed at the level of downstream biological interpretation based on concordant changes across assays, rather than by directly merging raw data matrices.

### Clinical correlation analysis

The transcriptomic data utilized in this study were obtained from a combined cohort of The Cancer Genome Atlas (TCGA), Therapeutically Applicable Research to Generate Effective Therapies (TARGET), and the Genotype-Tissue Expression (GTEx) project. We collected tumor tissue samples from the TCGA-LUSC cohort and the normal samples was constructed by combining the adjacent normal lung tissue samples from the TCGA-LUSC cohort with normal lung tissue samples from the GTEx database. Pearson correlation analysis was performed to assess the relationships between the expression levels of key genes (*SMAD4*, *SOX2*, and *EP300*). For the correlations between *SMAD4* and *SOX2* and between *SMAD4* and *EP300*, gene expression levels were quantified using Transcripts Per Million (TPM) values from transcript expression RNA-seq data. The analysis for the correlation between *EP300* and *SOX2* was conducted using gene expression RNA-seq data, also with TPM-normalized expression values.

### Ethics statement

The NSCLC tumor tissues and matched adjacent normal tissues used for Hi-C sequencing in this study were obtained from Hangzhou First People’s Hospital. All procedures involving human participants were approved by the Institutional Ethics Committee of Hangzhou First People’s Hospital (Approval No. 2026ZN017-1) and were carried out in accordance with the ethical standards of the institutional and/or national research committee. All animal studies were approved by the Biomedical Research Ethics Committee of Zhejiang University, China (Approval No. ZJU20240483 and ZJU20250257). All experimental procedures were conducted in strict accordance with institutional and national guidelines and regulations.

## Supporting information

file:///Users/tq/A%20Non-Canonical%20Role%20of%20SMAD4%20in%20Regulating%203D%20Genome%20Architecture%20to%20Inhibit%20Lung%20Squamous%20Cell%20Carcin

## Acknowledgments

This work was supported by grants to Dr. Jian Liu from the Natural Science Foundation (NSF) of China (General Grant: 82172899 and 82472637), the NSF of Zhejiang Province (Continuation Grant of Distinguished Young Scholars: LRG26H160001), Noncommunicable Chronic Diseases-National Science and Technology Major Project (2023ZD0502900/2023ZD0502902 and 2023ZD0507500/2023ZD0507501), ZJE Career Development Programme of ZJU international campus-lung and multi-organ research hub, Dr. Li Dak Sum & Yip Yio Chin Development Fund for Regenerative Medicine, Zhejiang University, the Open Fund of Zhejiang Provincial Key Laboratory of Pulmonology (KF202302), ZJE seed funding, ZJE 2024 International Campus Talent Special Funding Program, ZJE-UoE Joint Research Project, and Startup Funding of Tenure-track Assistant Professor of Zhejiang University. This work was also supported by the Zhejiang Province Pioneer Research and Development Project (No.2025C02091).

This work was also supported by Zhejiang University-University of Edinburgh Institute (ZJE) and the Department of Respiratory and Critical Care Medicine, The Second Affiliated Hospital, Zhejiang University School of Medicine, Zhejiang University.Haining People’s Hospital Tertiary-A Public Hospital Establishment Project Municipal Finance Support Fund; the Joint University-Local Cooperation Fund between ZJU-UoE Institute and Haining People’s Hospital. Supported by Sanming Project of Medicine in Shenzhen (No. SZSM202403006). This study was partially supported by the Guangxi Key Research and Development Program (GuiKe-AB25069017), the Joint Project on Regional High-Incidence Diseases Research of Guangxi Natural Science Foundation (2024GXNSFAA010101. 2024GXNSFBA010032).

We also acknowledged the help of members in Dr. Jian Liu’s lab. We thanked the ZJE core facility, especially the ZJE mouse core facility, and the Biomed-X Laboratory of ZJE Institute, School of Medicine, Zhejiang University, for continuous support. The Zhejiang University–University of Edinburgh Institute (ZJE) undergraduate authors, such as Zihan Wang, Taoyu Zhu, Xinrui Lin, Xiaolei Wang, Mingyang Xiao, Jiayi Ran, Xiaoran Cao, and Xinyi Xu gratefully acknowledged the profound impact of the ZJE HDRC3A (Human Disease: From Research to Clinic 3A) course, whose innovative pedagogical framework was instrumental in developing the critical perspective and interdisciplinary approach essential for this comprehensive work. The conceptualization and critical analysis presented in this work were directly shaped by the research-driven and AI-enhanced learning paradigm of the HDRC3A course.

## Data Availability Statement

The data that support the findings of this study are available on request from the corresponding author. The data are not publicly available due to privacy or ethical restrictions.

## Consent to publish

All authors have consented to publication.

## Author Contributions

J. Liu provided supervision, the concept, and the major resources for the project. Q. T. conducted most of the biochemical experiments. C. L. and X. H. conducted most of the bioinformatic analyses. X. Z. contributed the experiments and analyses. Z. W. helped the figure organization. N. Y. and Y. T. provided the project’s resources. S. Z. provided the Hi-C human samples. X. H. and J. Liu wrote the manuscript. Q. T., C.L., X. Z., H. S. A. R., N. Y., and Y. T. revised the manuscript. B. W., Y. X., M.X., Z. S., X.L., X.H., J.L., and Y. Z. contributed to mouse management. T. Z. and Z. W. contributed to the bioinformatic analyses. All authors contributed to the manuscript publication and revision.

## TOC Figure

**A Non-Canonical Role of SMAD4 in Regulating 3D Genome Architecture to Inhibit Lung Squamous Cell Carcinoma Development**

The mechanisms driving lung squamous cell carcinoma (LUSC) remain poorly understood, with few key regulatory factors identified. This study uncovers SMAD4 as a central regulator that reshapes the three-dimensional genome to promote LUSC development. Rather than acting through direct gene binding, SMAD4 indirectly controls genome folding, revealing a non-canonical way that transcription factors drive cancer progression.

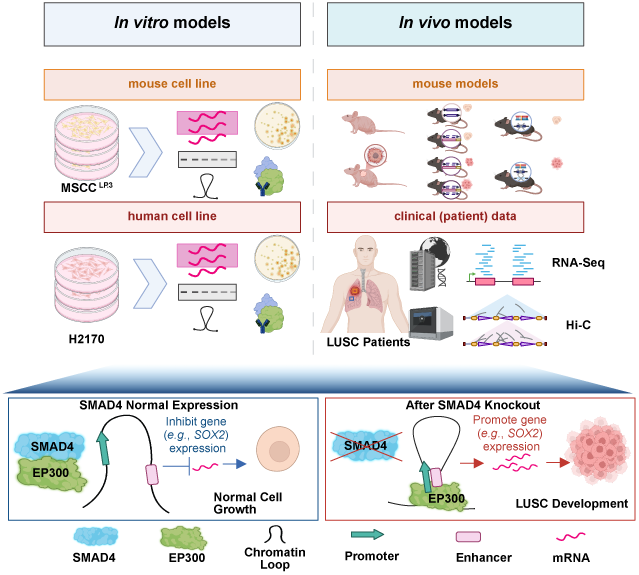

---

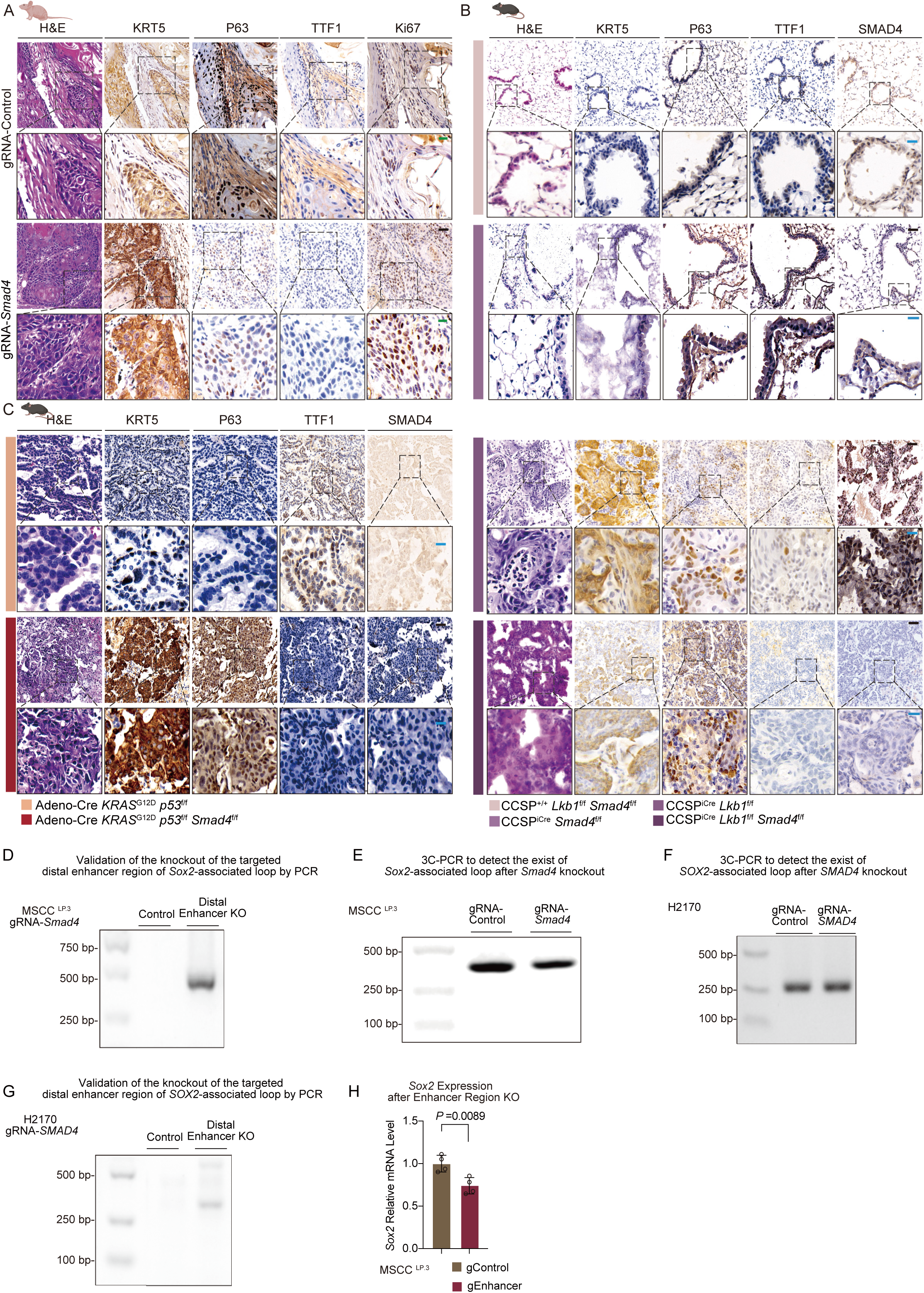

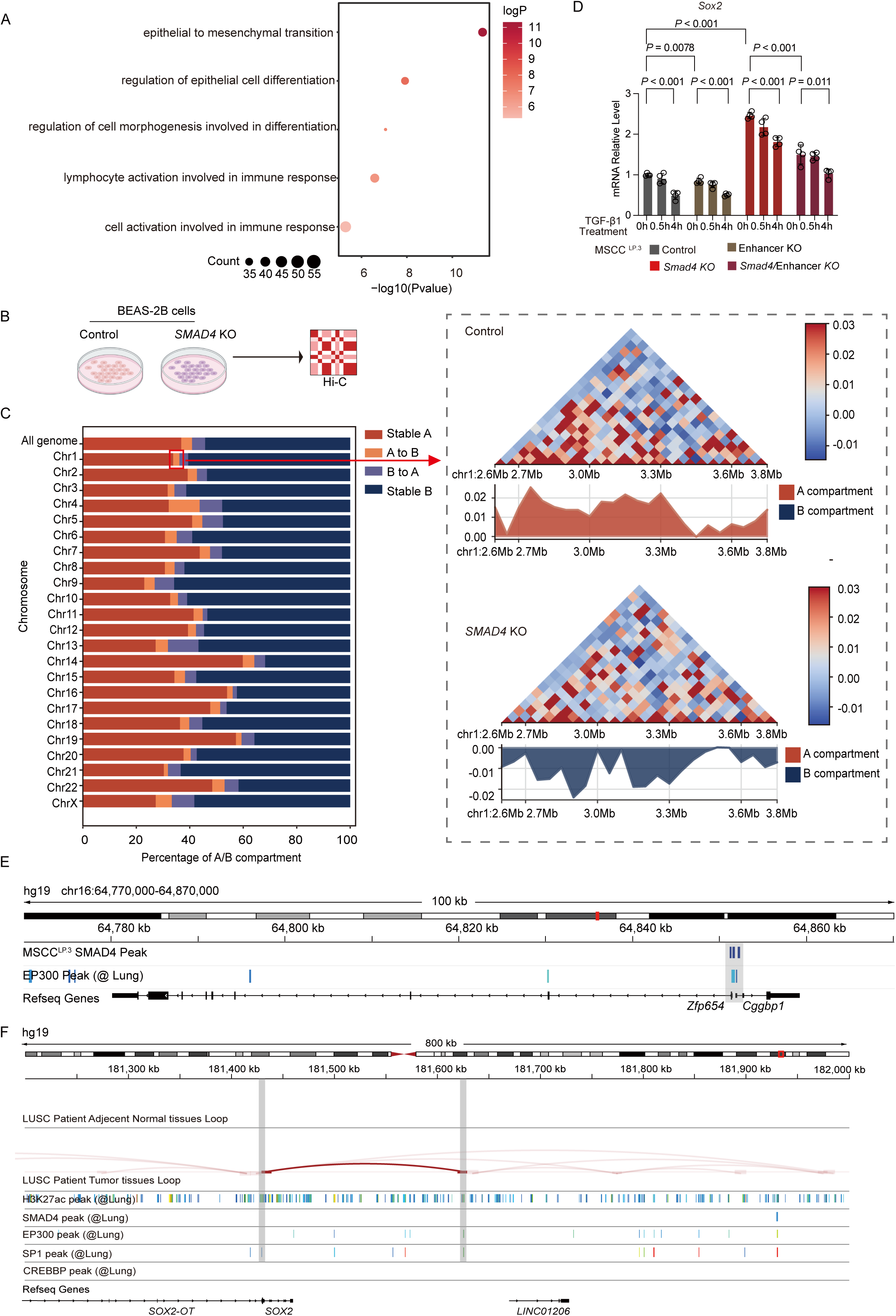

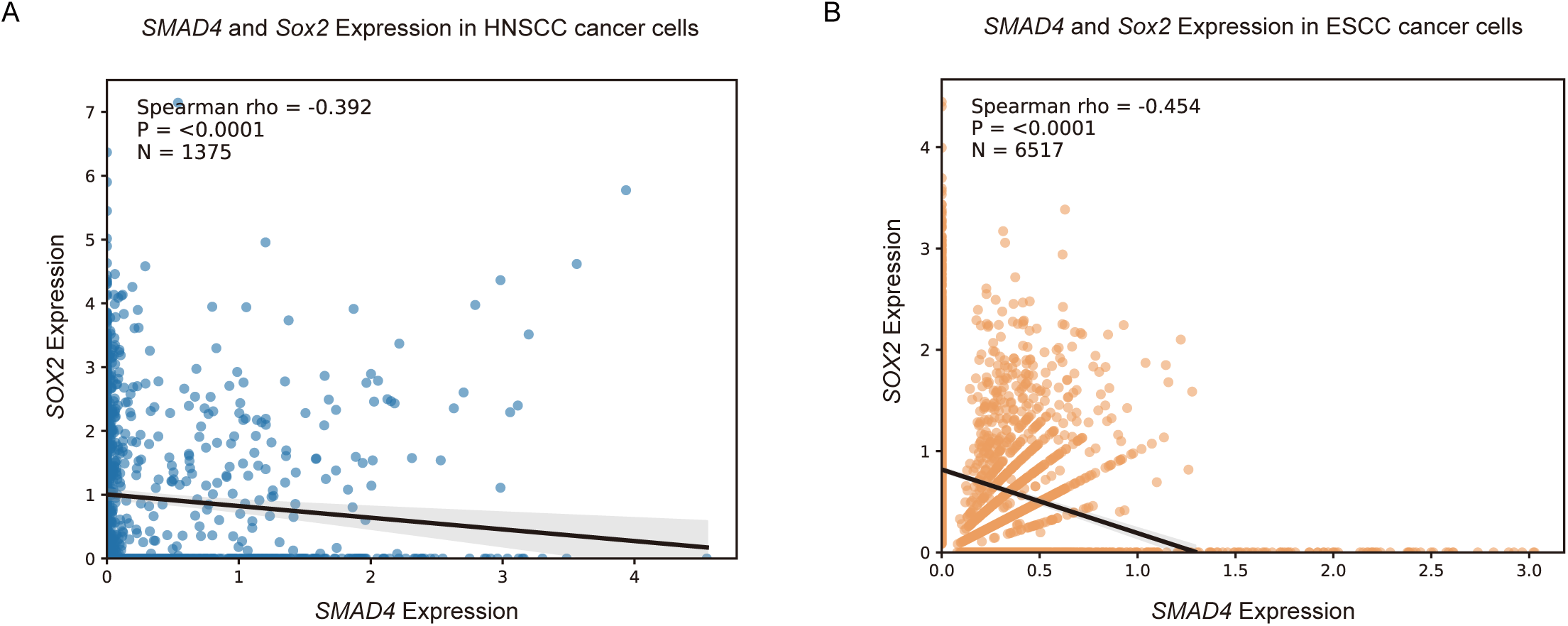

