## Supplementary material for "A Non-Canonical Role of SMAD4 in Regulating 3D Genome Architecture to Inhibit Lung Squamous Cell Carcinoma Development": file:///Users/tq/A%20Non-Canonical%20Role%20of%20SMAD4%20in%20Regulating%203D%20Genome%20Architecture%20to%20Inhibit%20Lung%20Squamous%20Cell%20Carcin

**Supplementary Materials**

**Supplementary Figure 1**
**Supplementary Figure 2**
**Supplementary Table 1. gRNA-*Smad4* vs. gRNA-Control DEGs**
**Supplementary Table 2. SMAD4 Peak Annotation**
**Supplementary Table 3. H3K27ac Peak Annotation**
**Supplementary Table 4. Summary of Primers**
**Supplementary Table 5. Summary of Guide RNA (gRNA) Sequences**
**Supplementary Table 6. Summary of siRNA Sequences**
**Supplementary Table 7. Summary of antisense oligonucleotides (ASO)**

**Supplementary Figure 1**

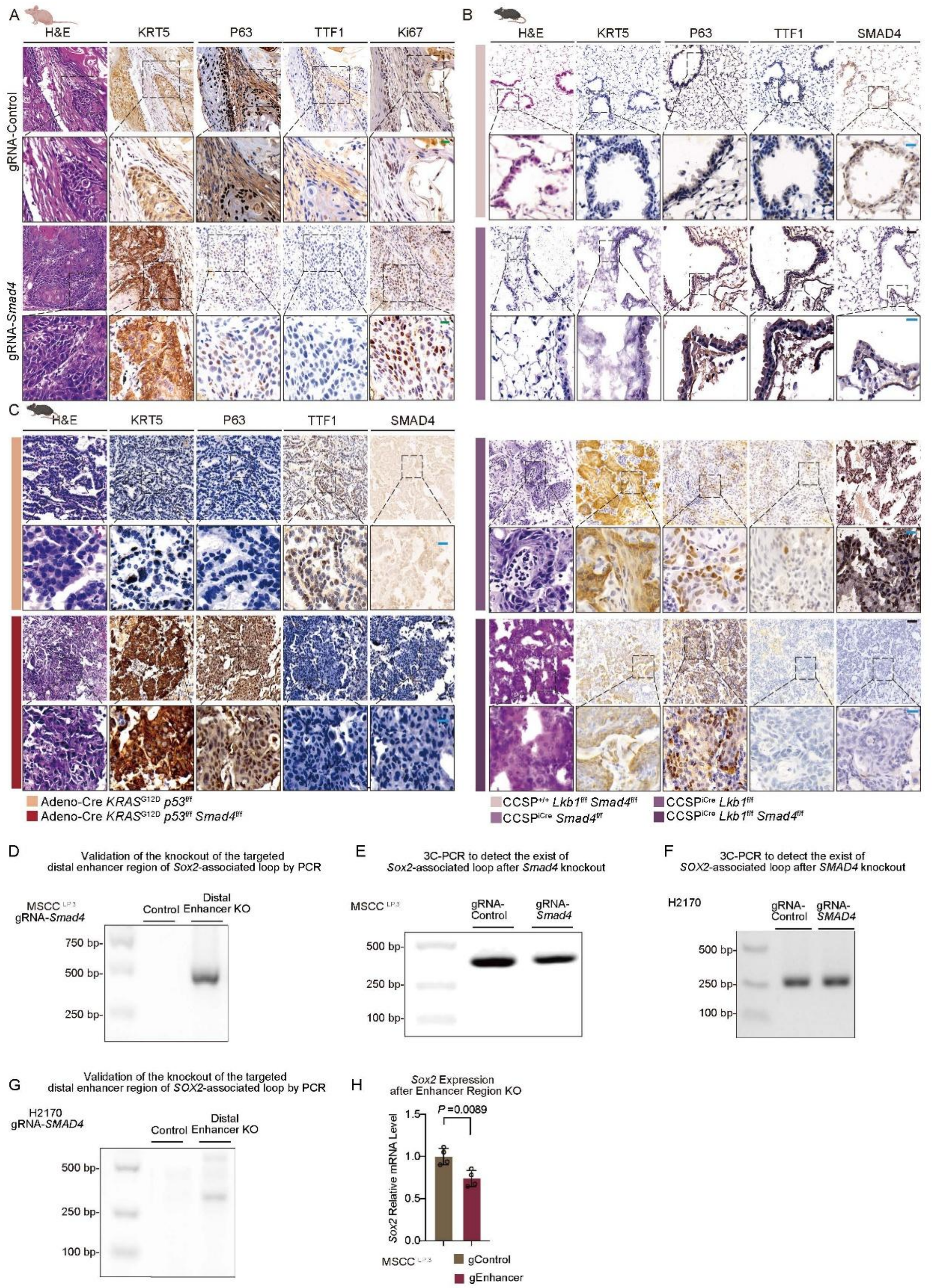


A. H&E staining and IHC staining of LUSC markers KRT5 and TP63, LUAD marker TTF1, and cell proliferation marker Ki67 in MSCC$^{LP.3}$ gRNA-Control and gRNA-*Smad4* xenograft tumors.

B. H&E staining and KRT5, TP63, TTF1 and SMAD4 IHC staining in CCSP$^{+/+}$ *Lkb1*$^{f/f}$ *Smad4*$^{f/f}$, CCSP$^{iCre}$*Smad4*$^{f/f}$, CCSP$^{iCre}$*Lkb1*$^{f/f}$, and CCSP$^{iCre}$*Lkb1*$^{f/f}$*Pten*$^{f/f}$ mice.

C. H&E staining and KRT5, TP63, TTF1 and SMAD4 IHC staining in Adeno-Cre *Kras*$^{G12D}$ *Trp53*$^{f/f}$ and Adeno-Cre *Smad4*$^{f/f}$ *Kras*$^{G12D}$ *Trp53*$^{f/f}$ mice.

D. PCR to validate the knockout of the targeted distal enhancer region of *Sox2*-associated loop in MSCC$^{LP.3}$ gRNA-*Smad4* cells.

E. 3C-PCR to detect the exist of *Sox2*-associated loop before and after *Smad4* knockout in MSCC$^{LP.3}$ cell line.

F. 3C-PCR to detect the exist of *SOX2*-associated loop before and after *SMAD4* knockout in H2170 cell line.

G. PCR to validate the knockout of the targeted distal enhancer region of *SOX2*-associated loop in H2170 gRNA-*SMAD4* cells.

H. RT-qPCR to detect *Sox2* expression followed by the knockout of the enhancer region of *Sox2*-associated loop in MSCC$^{LP.3}$ gRNA-Control cell line.

**Supplementary Figure 2**

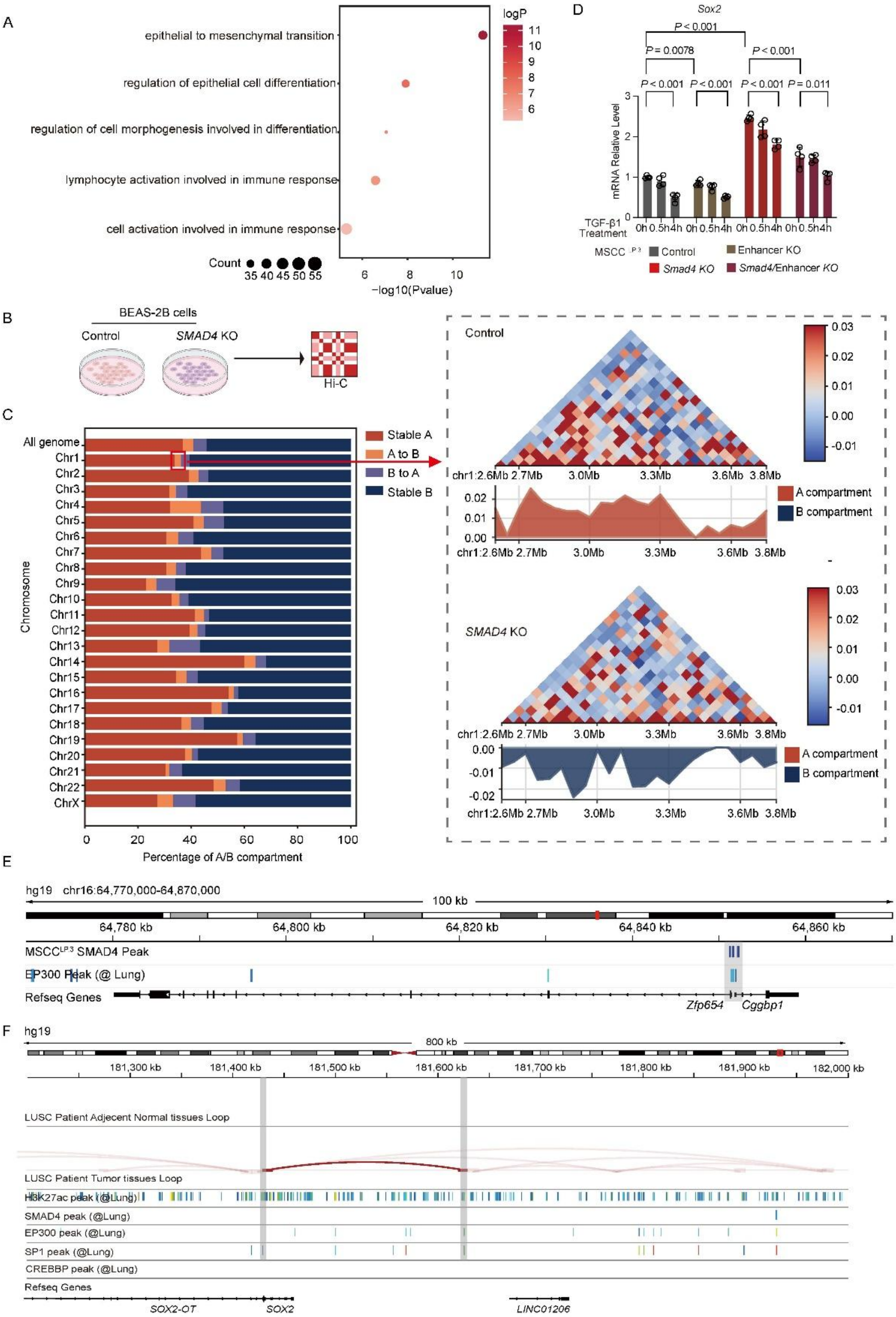


A. GO term pathway enrichment results based on differentially expressed genes between *Smad4* knockout and control groups.

B. qPCR analyses of the regulation of TGF- β on the *Sox2* transcription.

C. Hi-C analysis of *SMAD4* knockout and control BEAS-2B cell lines.

D. Changes in A/B compartments on various chromosomes before and after *SMAD4* knockout in BEAS-2B cell lines.

E. SMAD4 and EP300 binding at the *Zfp654* and *Cggpb1* promoter region in mouse lung cell line.

F. Proteins binding profiling (ChIP-Seq data of lung cell lines in public databases ChIP-Atlas) on the *SOX2*-associated loop anchor regions.

**Supplementary Figure 3**

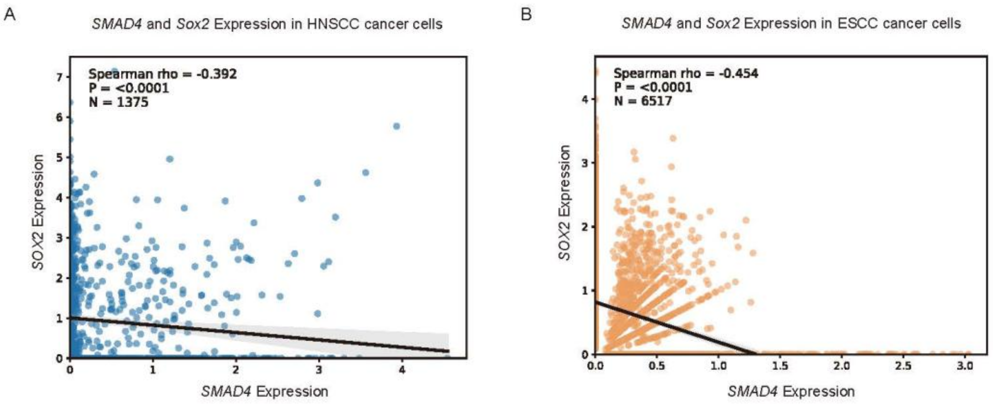


A. Correlation between *SMAD4* and *SOX2* Expression in Tumor Cells in Single-Cell RNA-Seq Data from Human Head and Neck Squamous Cell Carcinoma (HNSCC).

B. Correlation between *SMAD4* and *SOX2* expression in tumor cells in single-cell RNA-Seq Data from Esophageal Squamous Cell Carcinoma (ESCC).

**Supplementary Table 1. gRNA-*Smad4* vs. gRNA-Control DEGs**

| gene_id | log2FoldChange | FDR |
|---|---|---|
| *Ccl20* | 1.276047989 | 4.62E-14 |
| *Slc29a2* | 1.168429943 | 8.26E-20 |
| *Krt15* | 1.098585554 | 1.12E-10 |
| *Bmp2* | 0.955949521 | 3.30E-08 |
| *Scd2* | 0.94391639 | 4.61E-63 |
| *Krt7* | 0.913203583 | 3.73E-31 |
| *Aldh3a1* | 0.905833848 | 1.12E-08 |
| *Tnfaip2* | 0.870119104 | 1.88E-20 |
| *Tmprss11e* | 0.851693114 | 7.48E-17 |
| *Ces1g* | 0.844728874 | 1.12E-10 |
| *Akap12* | 0.797176595 | 1.32E-15 |
| *Prl2a1* | 0.736194317 | 5.09E-11 |
| *Ly6c1* | 0.735054685 | 5.53E-08 |

| | | |
|---|---|---|
| *Adgrl3* | 0.730877598 | 3.96E-05 |
| *Fgfbp1* | 0.719500494 | 4.95E-13 |
| *Stfa3* | 0.707331967 | 7.59E-05 |
| *Prodh* | 0.702424165 | 7.14E-07 |
| *Eps8* | 0.700150182 | 6.26E-05 |
| *Ska1* | 0.697680801 | 7.31E-05 |
| *Nrg2* | 0.697385472 | 1.50E-15 |
| *Efemp1* | 0.683304321 | 0.000174 |
| *Gm22973* | 0.681324539 | 0.000108 |
| *Hist1h3d* | 0.679971086 | 3.13E-14 |
| *Krt18* | 0.67374866 | 1.28E-11 |
| *Hist1h1b* | 0.661629691 | 2.46E-27 |
| *Ltbp1* | 0.657874252 | 1.58E-09 |
| *Lhfp* | 0.649401268 | 1.92E-18 |
| *Hist1h2ae* | 0.647896656 | 4.92E-10 |
| *Robo2* | 0.646371234 | 3.86E-11 |
| *Hist1h3h* | 0.644857447 | 2.66E-14 |
| *Hist1h2af* | 0.642038226 | 1.21E-08 |
| *Barx2* | 0.641615691 | 0.000362 |
| *Igf2r* | 0.640859194 | 5.50E-18 |
| *Depdc1b* | 0.628455467 | 1.41E-06 |
| *Cdc20* | 0.623487504 | 1.29E-22 |
| *Hist1h2bj* | 0.623103543 | 2.09E-15 |
| *Kif22* | 0.618595471 | 3.02E-16 |
| *Serpine2* | 0.616285127 | 1.44E-17 |
| *Hist2h2ac* | 0.61536397 | 1.04E-12 |
| *Nr4a1* | 0.614678251 | 0.000213 |
| *Slc16a3* | 0.614163952 | 5.59E-20 |
| *Gm24407* | 0.612603954 | 2.80E-05 |
| *Hist1h3c* | 0.610781673 | 2.30E-15 |
| *Hist1h2an* | 0.609786935 | 1.56E-06 |
| *Fam213a* | 0.609374153 | 0.000217 |
| *Gap43* | 0.605243457 | 3.68E-05 |
| *Cdk14* | 0.601528528 | 0.00052 |
| *Hist1h2bh* | 0.600623974 | 3.39E-16 |
| *Hist1h2ad* | 0.600504638 | 3.38E-08 |
| *Hs3st3b1* | 0.594626737 | 8.21E-18 |
| *Bdnf* | 0.58796371 | 0.001558 |
| *Psrc1* | 0.587381685 | 6.83E-08 |
| *Fam111a* | 0.584364407 | 2.27E-13 |
| *Hist1h4d* | 0.584262152 | 7.85E-24 |
| *Hist1h2bb* | 0.579586589 | 1.38E-22 |
| *Hist1h2bn* | 0.579184885 | 9.54E-08 |
| *Hist1h2be* | 0.578515061 | 1.39E-15 |

| | | |
|---|---|---|
| *Lcn2* | 0.578453416 | 0.002033 |
| *Tfap4* | 0.577998682 | 4.26E-07 |
| *Hist1h2bk* | 0.577459404 | 4.34E-14 |
| *Glyat* | 0.575172708 | 0.001919 |
| *Hist1h2ag* | 0.575107875 | 2.46E-06 |
| *Hist1h3b* | 0.571499825 | 0.000104 |
| *Hist1h2bg* | 0.570122518 | 1.05E-17 |
| *Lrrc8b* | 0.56905291 | 4.57E-06 |
| *B4galnt4* | 0.568551841 | 9.96E-07 |
| *Hist1h2ac* | 0.568378278 | 4.03E-11 |
| *Hist1h1e* | 0.566606489 | 6.19E-20 |
| *Pif1* | 0.56612094 | 8.61E-08 |
| *Iqgap3* | 0.564341554 | 1.27E-13 |
| *Slc1a5* | 0.564275893 | 5.51E-14 |
| *Hist1h4b* | 0.563495824 | 9.25E-14 |
| *Ak3* | 0.560830803 | 3.08E-10 |
| *Slc7a5* | 0.559384633 | 2.26E-10 |
| *Evc2* | 0.558877982 | 0.003075 |
| *Tmem229b* | 0.557473852 | 0.001666 |
| *Tgm1* | 0.556942839 | 0.001552 |
| *Ncaph* | 0.552591528 | 1.32E-15 |
| *Hist1h1d* | 0.550621563 | 6.64E-18 |
| *Ccnb1* | 0.549795724 | 3.45E-12 |
| *Hist1h4h* | 0.542773478 | 7.48E-17 |
| *Pde3b* | 0.542717509 | 0.000648 |
| *Hist1h3g* | 0.542680369 | 4.60E-11 |
| *Rfc5* | 0.542221132 | 2.90E-15 |
| *Hist1h4k* | 0.541978465 | 1.65E-15 |
| *Nek2* | 0.541853321 | 6.00E-15 |
| *Slc6a14* | 0.537116077 | 0.004796 |
| *Hist1h2bp* | 0.533897395 | 5.55E-08 |
| *Hist1h3i* | 0.532988898 | 9.81E-11 |
| *Ncapg* | 0.531957031 | 4.24E-14 |
| *Hist2h2bb* | 0.531411923 | 2.67E-05 |
| *Car12* | 0.530677271 | 6.53E-16 |
| *Hist1h1a* | 0.530027486 | 4.00E-11 |
| *Cenpu* | 0.529961898 | 2.48E-05 |
| *Ect2* | 0.528224145 | 2.28E-11 |
| *Hist1h4a* | 0.52791201 | 2.21E-13 |
| *Lyar* | 0.527691032 | 9.00E-10 |
| *Trip13* | 0.527111341 | 1.62E-08 |
| *Bub1b* | 0.525073573 | 2.25E-13 |
| *Fkbp5* | 0.524509772 | 0.00013 |
| *Hist1h2bm* | 0.522194007 | 1.01E-07 |

| | | |
|---|---|---|
| *Shmt1* | 0.522139545 | 6.32E-08 |
| *Cebpa* | 0.517530198 | 0.00697 |
| *Kif2c* | 0.516521871 | 5.52E-11 |
| *Il1f6* | 0.515962199 | 8.23E-06 |
| *Nsl1* | 0.515882623 | 4.65E-07 |
| *Abcb1b* | 0.514811539 | 3.30E-15 |
| *Hist1h2bc* | 0.513177377 | 7.84E-11 |
| *Cks2* | 0.511461976 | 1.23E-06 |
| *Plk1* | 0.51085464 | 6.75E-11 |
| *P3h1* | 0.510379491 | 0.005592 |
| *Cep128* | 0.508355215 | 3.51E-05 |
| *Hist1h4f* | 0.508231736 | 1.04E-14 |
| *Chchd4* | 0.508089074 | 4.14E-11 |
| *Ifrd2* | 0.50624756 | 1.84E-09 |
| *Slc35f2* | 0.505704368 | 0.00817 |
| *RP24-112H7.1* | 0.504180773 | 0.004862 |
| *Dhrs1* | 0.503564193 | 9.51E-13 |
| *Hirip3* | 0.502179003 | 2.82E-06 |
| *Cad* | 0.502165736 | 9.05E-18 |
| *Hist1h3e* | 0.49998519 | 7.23E-11 |
| *Zwilch* | 0.49979647 | 6.18E-10 |
| *Hist1h3f* | 0.499567542 | 3.22E-11 |
| *Cdyl2* | 0.499263162 | 0.008696 |
| *Esco2* | 0.498726872 | 1.26E-06 |
| *Hist4h4* | 0.497423254 | 1.54E-13 |
| *Tacc3* | 0.497064347 | 5.56E-17 |
| *Anln* | 0.495610369 | 5.41E-12 |
| *Trp73* | 0.494864416 | 0.001211 |
| *Apln* | 0.494523711 | 0.010365 |
| *Arrdc4* | 0.494354168 | 5.12E-08 |
| *Ccna2* | 0.494145771 | 3.57E-05 |
| *Rrp1b* | 0.492711577 | 7.89E-11 |
| *Cenpn* | 0.492479548 | 2.09E-05 |
| *Zbed5* | 0.491916128 | 0.009328 |
| *Slc25a48* | 0.490744666 | 2.84E-09 |
| *Itgb7* | 0.490059905 | 8.77E-06 |
| *Esr1* | 0.489456051 | 0.010582 |
| *Diaph3* | 0.488174014 | 1.99E-11 |
| *Krt19* | 0.487899979 | 0.000144 |
| *Rrp12* | 0.48753969 | 1.23E-14 |
| *Hist1h2ab* | 0.485683369 | 0.001293 |
| *Aurkb* | 0.485095562 | 4.05E-09 |
| *Dtwd1* | 0.485062541 | 7.78E-09 |
| *Dut* | 0.485003268 | 8.42E-12 |

| | | |
|---|---|---|
| *Apcdd1* | 0.484923845 | 0.010426 |
| *Aunip* | 0.484722796 | 0.002347 |
| *Tmem74* | 0.483781146 | 0.01034 |
| *Sh3rf2* | 0.483494153 | 1.28E-05 |
| *Uhrf1* | 0.482504952 | 2.74E-12 |
| *Dctpp1* | 0.482298837 | 1.13E-06 |
| *Hist1h4j* | 0.482162683 | 1.67E-08 |
| *Ndc1* | 0.480424592 | 1.34E-10 |
| *Krt8* | 0.480329401 | 2.66E-13 |
| *Hist1h4i* | 0.478972524 | 4.28E-10 |
| *Snord17* | 0.478275367 | 8.34E-05 |
| *Tdp1* | 0.47818147 | 9.27E-05 |
| *Muc15* | 0.478016629 | 0.014069 |
| *Recql4* | 0.477170555 | 5.41E-06 |
| *Dlgap5* | 0.476823062 | 9.21E-11 |
| *Gm22620* | 0.476379223 | 0.011796 |
| *Slc19a1* | 0.476138893 | 2.36E-06 |
| *Htr5b* | 0.475089414 | 0.01472 |
| *Srm* | 0.474770766 | 4.08E-10 |
| *Gm23130* | 0.474595159 | 0.004987 |
| *Prc1* | 0.474266766 | 2.69E-12 |
| *Dgkh* | 0.472504005 | 2.14E-05 |
| *Hist2h4* | 0.471772688 | 7.98E-08 |
| *Kif11* | 0.471037472 | 2.62E-16 |
| *Cmss1* | 0.470374597 | 0.000117 |
| *Tmem194* | 0.469931997 | 0.000109 |
| *Gm25395* | 0.469603228 | 0.002959 |
| *Aspm* | 0.469203308 | 3.01E-12 |
| *Snora78* | 0.468882704 | 0.000185 |
| *Lrig1* | 0.468506035 | 2.24E-16 |
| *Akip1* | 0.468153172 | 0.016098 |
| *Hist1h2ak* | 0.468032671 | 0.00058 |
| *Eme1* | 0.467869999 | 4.84E-05 |
| *Fam72a* | 0.467829433 | 0.005995 |
| *Cbx6* | 0.467828459 | 3.56E-08 |
| *Spc24* | 0.467288483 | 5.09E-09 |
| *Pbk* | 0.467032963 | 6.82E-08 |
| *Snord93* | 0.466456998 | 0.017147 |
| *Chtf18* | 0.465725521 | 1.37E-08 |
| *Card14* | 0.465665089 | 0.008029 |
| *Cdca2* | 0.46427333 | 5.88E-08 |
| *Espl1* | 0.464117122 | 1.79E-06 |
| *Racgap1* | 0.463285607 | 1.56E-12 |
| *Fanci* | 0.463125084 | 2.97E-07 |

| | | |
|---|---|---|
| *Rad54l* | 0.462861568 | 4.11E-05 |
| *Hist1h2bl* | 0.461702028 | 0.001092 |
| *Hist1h4c* | 0.461257678 | 2.27E-10 |
| *Pla2r1* | 0.461068692 | 0.00036 |
| *Tcof1* | 0.460987499 | 1.21E-08 |
| *Mns1* | 0.460563844 | 0.007417 |
| *Mad2l1* | 0.46009432 | 3.87E-07 |
| *Gm22422* | 0.459574477 | 0.008713 |
| *Timm8a1* | 0.459010159 | 0.00871 |
| *Lgals7* | 0.45884349 | 0.014865 |
| *RP23-461C4.3* | 0.4585796 | 0.001001 |
| *Cdk1* | 0.458322296 | 7.23E-11 |
| *Hmmr* | 0.457352558 | 4.87E-12 |
| *Wdr12* | 0.456260826 | 1.04E-13 |
| *H2afx* | 0.45604946 | 9.30E-10 |
| *Gm25835* | 0.454383796 | 0.004033 |
| *Atp13a4* | 0.454145421 | 0.018069 |
| *Ttk* | 0.453805288 | 9.81E-11 |
| *Rabggta* | 0.453387348 | 1.34E-05 |
| *Hist2h3b* | 0.453312366 | 0.000244 |
| *Tgfbr3* | 0.452891772 | 0.003241 |
| *Aif1l* | 0.45284105 | 0.000173 |
| *Mis18bp1* | 0.452739062 | 9.94E-06 |
| *Gdpd2* | 0.451283769 | 0.01845 |
| *Hist1h1c* | 0.451274402 | 8.45E-12 |
| *Snord22* | 0.450665159 | 0.00087 |
| *Ampd1* | 0.450165768 | 0.017343 |
| *Ccnf* | 0.449763558 | 5.37E-08 |
| *Gm22068* | 0.44856128 | 0.018812 |
| *Pdss1* | 0.448503171 | 0.000113 |
| *Irs1* | 0.448412027 | 7.78E-06 |
| *Itgb2* | 0.447900786 | 0.018173 |
| *Ppat* | 0.4472818 | 3.25E-11 |
| *Pus7* | 0.446190854 | 5.27E-11 |
| *Rpl7l1* | 0.445346029 | 1.98E-18 |
| *Aurka* | 0.445180464 | 2.67E-12 |
| *Ramp3* | 0.44517528 | 0.006154 |
| *Kif18b* | 0.443815192 | 5.39E-07 |
| *RP23-110C17.2* | 0.442941713 | 0.014231 |
| *Mat2a* | 0.442215544 | 2.15E-16 |
| *Gen1* | 0.441302374 | 2.32E-05 |
| *Fancb* | 0.44097926 | 0.000822 |
| *Ung* | 0.440768489 | 2.38E-05 |
| *Lsm3* | 0.43910431 | 3.37E-06 |

| | | |
|---|---|---|
| *Eri2* | 0.438121601 | 1.16E-05 |
| *Hist1h2ai* | 0.437724336 | 0.013565 |
| *Rnu3a* | 0.437647879 | 0.024782 |
| *Lin28b* | 0.437500628 | 0.004129 |
| *Hist1h2bf* | 0.436613832 | 6.40E-06 |
| *Nup37* | 0.436175577 | 8.61E-08 |
| *Kif4* | 0.436136474 | 9.72E-12 |
| *Kif20b* | 0.436018564 | 4.21E-13 |
| *Stom* | 0.435338081 | 5.38E-12 |
| *RP23-63D8.8* | 0.434994993 | 0.020208 |
| *Dhfr* | 0.434965054 | 3.90E-09 |
| *Gart* | 0.434474938 | 1.82E-16 |
| *Nup85* | 0.433270641 | 2.56E-12 |
| *Snora21* | 0.432823153 | 0.0297 |
| *Hdac5* | 0.432741199 | 0.015572 |
| *Mfsd2a* | 0.432553867 | 0.000148 |
| *Kntc1* | 0.432472482 | 1.74E-10 |
| *Gm24265* | 0.431959806 | 8.44E-06 |
| *Xrcc2* | 0.431691059 | 7.23E-06 |
| *RP24-160H16.10* | 0.431384835 | 0.009854 |
| *Exo1* | 0.431007856 | 5.17E-08 |
| *Gm25813* | 0.430228981 | 0.031104 |
| *Mphosph9* | 0.430181732 | 5.77E-05 |
| *Knstrn* | 0.429954104 | 3.66E-10 |
| *Bora* | 0.429684438 | 2.38E-05 |
| *Pim3* | 0.429053641 | 7.31E-06 |
| *Gemin6* | 0.428424161 | 0.00028 |
| *Sgol1* | 0.427973654 | 6.33E-06 |
| *Gm25360* | 0.427951614 | 0.009021 |
| *Suv39h1* | 0.427174871 | 9.55E-06 |
| *Atf3* | 0.427162241 | 0.006266 |
| *Snora73b* | 0.427130126 | 0.015845 |
| *Cenpf* | 0.426584389 | 2.93E-11 |
| *Siah1b* | 0.426580619 | 0.000139 |
| *Flot2* | 0.426495305 | 0.029437 |
| *Cenpi* | 0.426348164 | 1.10E-07 |
| *Birc5* | 0.425714809 | 3.29E-08 |
| *Rcc1* | 0.425628532 | 2.25E-07 |
| *Kifc1* | 0.425568822 | 4.82E-05 |
| *Cecr2* | 0.425327312 | 0.025923 |
| *Ankle1* | 0.423906664 | 0.00111 |
| *Spry1* | 0.423690879 | 0.034246 |
| *Pdzrn3* | 0.423676502 | 2.45E-10 |
| *Nup133* | 0.423562102 | 7.26E-12 |

| | | |
|---|---|---|
| *Htra2* | 0.423314186 | 6.20E-06 |
| *Fignl1* | 0.422603412 | 3.71E-09 |
| *Plk4* | 0.422507626 | 1.49E-08 |
| *Mthfd2* | 0.422163342 | 3.10E-07 |
| *Cdca8* | 0.42215112 | 1.41E-07 |
| *Ube2c* | 0.422086018 | 5.32E-09 |
| *Pole* | 0.421588474 | 8.08E-09 |
| *Fbxo5* | 0.421444375 | 2.15E-06 |
| *Cenph* | 0.42135587 | 7.23E-05 |
| *Nav3* | 0.421146076 | 2.10E-05 |
| *Trmt61a* | 0.420167562 | 9.97E-05 |
| *Nup107* | 0.419484818 | 3.52E-10 |
| *Slc39a4* | 0.419433315 | 0.031532 |
| *Ovol2* | 0.41905999 | 0.025053 |
| *Glt1d1* | 0.418639048 | 0.000327 |
| *Dctd* | 0.417955657 | 1.31E-05 |
| *Mcm7* | 0.417233541 | 5.05E-11 |
| *Sema3e* | 0.417053281 | 8.34E-05 |
| *Hist1h2ah* | 0.416782276 | 0.02397 |
| *Camk1d* | 0.416736528 | 0.038159 |
| *Nuf2* | 0.416650518 | 1.10E-07 |
| *Gm24830* | 0.415550721 | 0.038839 |
| *Top2a* | 0.414485365 | 4.25E-11 |
| *Nop16* | 0.414434997 | 4.15E-08 |
| *Pold1* | 0.414014524 | 3.79E-07 |
| *Rny1* | 0.413444407 | 1.13E-05 |
| *Dis3* | 0.41328851 | 1.52E-09 |
| *Kif18a* | 0.412904121 | 1.37E-05 |
| *Tma16* | 0.412479374 | 0.000134 |
| *Mir17hg* | 0.412393594 | 0.001486 |
| *RP23-480B19.17* | 0.411283776 | 0.031966 |
| *Exosc8* | 0.411134883 | 2.88E-07 |
| *Traip* | 0.411128281 | 2.73E-05 |
| *Gsg2* | 0.410275869 | 6.87E-05 |
| *Grwd1* | 0.410148271 | 2.76E-08 |
| *Spag5* | 0.409732015 | 8.43E-07 |
| *Smc4* | 0.409682542 | 1.66E-10 |
| *RP23-79J21.2* | 0.40902355 | 0.031116 |
| *Ncapg2* | 0.408632253 | 9.22E-10 |
| *Fam83d* | 0.40835068 | 7.35E-06 |
| *Stfa3* | 0.408267266 | 0.039633 |
| *Rcl1* | 0.408254308 | 1.10E-10 |
| *Nrp1* | 0.407826752 | 4.57E-15 |
| *Plekhh1* | 0.4074827 | 0.000117 |

| | | |
|---|---|---|
| *Lsm2* | 0.40646222 | 3.83E-06 |
| *RP23-300H4.1* | 0.406089104 | 0.001286 |
| *Serpina3g* | 0.406010467 | 0.040753 |
| *Cinp* | 0.40589474 | 1.01E-07 |
| *Kif14* | 0.405665243 | 2.32E-05 |
| *Man1a* | 0.40519762 | 6.51E-06 |
| *Txnip* | 0.405084844 | 3.82E-05 |
| *Cenpe* | 0.404415864 | 5.62E-08 |
| *Rbm19* | 0.404413584 | 8.46E-09 |
| *Troap* | 0.403903239 | 0.003236 |
| *Cyp3a13* | 0.403726241 | 0.000718 |
| *Cdc25c* | 0.40353349 | 3.50E-05 |
| *Mrto4* | 0.403140171 | 1.90E-07 |
| *Twist1* | 0.402962626 | 0.045992 |
| *Ptpn13* | 0.402333447 | 2.08E-20 |
| *Gsto1* | 0.401690217 | 1.44E-25 |
| *RP23-81H23.3* | 0.401607107 | 0.032796 |
| *Pros1* | 0.40079998 | 0.04778 |
| *Miip* | 0.400683403 | 2.45E-05 |
| *Serpinb10* | 0.400529667 | 8.07E-13 |
| *Snora31* | 0.400317624 | 0.048 |
| *Nop9* | 0.399736658 | 2.08E-08 |
| *Timeless* | 0.39966043 | 2.03E-07 |
| *Pi15* | 0.399492063 | 0.044859 |
| *Sorbs1* | 0.398994125 | 0.00655 |
| *Gsta1* | 0.398022808 | 0.02931 |
| *Ckap2l* | 0.397776881 | 1.14E-06 |
| *Clec2g* | 0.397351534 | 0.045426 |
| *Dph2* | 0.396643926 | 0.000358 |
| *Ptprz1* | 0.396249145 | 0.042832 |
| *Pla2g4a* | 0.395419036 | 5.40E-07 |
| *RP23-69P8.2* | 0.395240004 | 0.005209 |
| *Umps* | 0.394357086 | 3.50E-11 |
| *Asf1b* | 0.393946417 | 0.000189 |
| *Kif15* | 0.393750882 | 0.000241 |
| *Nrm* | 0.392800166 | 0.005557 |
| *Rrp15* | 0.391738787 | 3.69E-08 |
| *RP24-201C14.10* | 0.391199578 | 0.014516 |
| *Kifc5b* | 0.390917212 | 0.003046 |
| *Tnik* | 0.390315739 | 0.003601 |
| *Dnph1* | 0.390251409 | 0.001845 |
| *Fancd2* | 0.389453119 | 7.78E-06 |
| *Cenpw* | 0.388977951 | 0.000409 |
| *Incenp* | 0.387240225 | 0.004434 |

| | | |
|---|---|---|
| *Zfp59* | 0.386677869 | 0.027656 |
| *Ndc80* | 0.386392538 | 3.14E-05 |
| *Cdca3* | 0.385316194 | 1.40E-08 |
| *Brip1* | 0.385015663 | 3.36E-06 |
| *Cyb5r1* | 0.38496207 | 1.52E-06 |
| *Rpp40* | 0.384802845 | 0.001554 |
| *Mettl1* | 0.384631549 | 0.000144 |
| *Slc16a1* | 0.384351772 | 6.92E-09 |
| *Ddx11* | 0.384139739 | 1.10E-06 |
| *Lbr* | 0.384075253 | 1.61E-12 |
| *Eppk1* | 0.383942735 | 1.21E-07 |
| *Rtel1* | 0.38387706 | 4.22E-07 |
| *Arhgap11a* | 0.383705974 | 1.28E-07 |
| *Adk* | 0.383016784 | 2.60E-07 |
| *Ppid* | 0.382923708 | 1.96E-09 |
| *Shcbp1* | 0.382735349 | 4.41E-06 |
| *Gpd2* | 0.382691678 | 1.15E-14 |
| *Foxg1* | 0.382672776 | 0.017721 |
| *Tyms* | 0.382478808 | 9.70E-08 |
| *St3gal6* | 0.381838055 | 0.012095 |
| *Hist2h2ab* | 0.381570185 | 0.023407 |
| *Ube2t* | 0.381301045 | 0.007748 |
| *Nolc1* | 0.380902506 | 8.16E-11 |
| *Cit* | 0.379183312 | 9.04E-05 |
| *Cenpq* | 0.379180416 | 0.001879 |
| *Endou* | 0.378901619 | 0.015771 |
| *3110082I17Rik* | 0.378719515 | 0.003624 |
| *Cks1b* | 0.377984108 | 2.31E-05 |
| *Cdc45* | 0.377979003 | 6.51E-07 |
| *Dbf4* | 0.37758975 | 8.80E-09 |
| *Polr1b* | 0.377451282 | 3.16E-10 |
| *Lmnb1* | 0.377380906 | 2.66E-10 |
| *Slc5a6* | 0.37674036 | 0.004756 |
| *Vmn1r43* | 0.37672585 | 0.011919 |
| *Casc5* | 0.376569799 | 2.64E-06 |
| *Patz1* | 0.376240916 | 0.017992 |
| *Metrn* | 0.376169767 | 0.023277 |
| *Lig1* | 0.375494173 | 2.10E-06 |
| *Tbl3* | 0.375042055 | 1.07E-07 |
| *Ddx21* | 0.375002689 | 3.05E-15 |
| *Nudt11* | 0.374962541 | 0.043981 |
| *Wnt3a* | 0.37474663 | 7.37E-06 |
| *Cdt1* | 0.373600172 | 5.20E-07 |
| *Gm26224* | 0.373276881 | 0.011228 |

| | | |
|---|---|---|
| *Cenpm* | 0.373013315 | 0.011136 |
| *Tk1* | 0.372534349 | 5.96E-05 |
| *Nle1* | 0.37248581 | 0.00056 |
| *Mdm1* | 0.372328882 | 0.006394 |
| *RP23-313J14.6* | 0.372138096 | 0.000762 |
| *Mir1949* | 0.37187963 | 0.006532 |
| *Fen1* | 0.371855356 | 1.31E-09 |
| *Arhgap19* | 0.371309809 | 1.30E-06 |
| *Mybl2* | 0.370882904 | 0.000121 |
| *Hyls1* | 0.37010545 | 0.00142 |
| *Blm* | 0.36949868 | 0.000269 |
| *Ddx31* | 0.369375373 | 4.17E-08 |
| *Haus4* | 0.369249357 | 0.000104 |
| *Pik3ip1* | 0.369124932 | 0.042403 |
| *Ppan* | 0.368651516 | 1.34E-07 |
| *Gpsm2* | 0.368259174 | 6.14E-09 |
| *Csta1* | 0.367784324 | 0.002806 |
| *Ttc27* | 0.367502559 | 5.57E-06 |
| *Tfb2m* | 0.36688399 | 0.006872 |
| *RP23-139J18.1* | 0.366831623 | 0.000511 |
| *Wdr43* | 0.366830681 | 3.67E-10 |
| *Atic* | 0.365197116 | 3.02E-12 |
| *Gtpbp10* | 0.365088925 | 0.000227 |
| *Zfp850* | 0.36490757 | 0.007352 |
| *Plk2* | 0.364676771 | 3.28E-13 |
| *Gnl3* | 0.364335912 | 1.60E-08 |
| *Nol8* | 0.364082219 | 1.36E-07 |
| *Smyd5* | 0.36353718 | 1.12E-05 |
| *Cdc42ep3* | 0.363369684 | 5.47E-05 |
| *Grk6* | 0.362681404 | 4.95E-10 |
| *Hk2* | 0.362169477 | 4.99E-09 |
| *Ifitm3* | 0.361788461 | 1.22E-09 |
| *Ruvbl2* | 0.361765048 | 3.18E-09 |
| *Ckap2* | 0.361651943 | 2.96E-07 |
| *Lncenc1* | 0.361482138 | 1.20E-15 |
| *Prmt7* | 0.361335087 | 3.06E-07 |
| *Nol9* | 0.36125982 | 1.80E-09 |
| *Polq* | 0.361199099 | 0.000126 |
| *RP23-413D14.9* | 0.361176671 | 0.036325 |
| *Gar1* | 0.361078321 | 0.003107 |
| *Wdr46* | 0.361044567 | 2.10E-07 |
| *Rad51* | 0.360645986 | 0.000372 |
| *Ccdc85c* | 0.360287218 | 7.43E-05 |
| *Ncapd2* | 0.360161803 | 9.30E-06 |

| | | |
|---|---|---|
| *G2e3* | 0.359474068 | 1.79E-05 |
| *Dkc1* | 0.359468659 | 1.09E-06 |
| *Foxm1* | 0.359416277 | 0.000119 |
| *Wdhd1* | 0.358447842 | 8.13E-06 |
| *Enoph1* | 0.357950685 | 1.68E-06 |
| *Wdr4* | 0.357581706 | 1.31E-05 |
| *Fam64a* | 0.357433787 | 7.37E-05 |
| *Snora68* | 0.357393341 | 0.025768 |
| *RP23-376B19.2* | 0.356902391 | 1.43E-05 |
| *Gls2* | 0.355964585 | 0.013766 |
| *Otud4* | 0.355487628 | 1.18E-07 |
| *Gm24044* | 0.354862165 | 0.041218 |
| *Dscc1* | 0.354859874 | 0.028013 |
| *Mcm2* | 0.354827934 | 1.08E-09 |
| *Cdca7l* | 0.354673361 | 0.00046 |
| *Trub1* | 0.354590885 | 0.000112 |
| *Nsun5* | 0.353294928 | 0.001302 |
| *Cenpa* | 0.352251335 | 2.15E-05 |
| *Hist1h3a* | 0.351825744 | 0.0004 |
| *Cenpp* | 0.351778246 | 0.042201 |
| *Shmt2* | 0.35128836 | 9.44E-09 |
| *Bub1* | 0.350854353 | 1.90E-06 |
| *Sapcd2* | 0.349094921 | 0.001358 |
| *L3mbtl2* | 0.348706382 | 3.96E-05 |
| *Cep85* | 0.348597665 | 0.0008 |
| *Ipmk* | 0.347764283 | 9.50E-06 |
| *Ercc6l* | 0.347152076 | 3.28E-06 |
| *Gtse1* | 0.346712131 | 7.16E-06 |
| *Rrp8* | 0.346257261 | 2.75E-06 |
| *Wdr73* | 0.344992352 | 0.000272 |
| *Wdr3* | 0.344599386 | 8.99E-08 |
| *Rad18* | 0.344063585 | 0.000342 |
| *Slc2a9* | 0.343757877 | 0.042179 |
| *Cdca5* | 0.343274006 | 0.00037 |
| *Epgn* | 0.342711246 | 3.34E-08 |
| *Cep89* | 0.342186691 | 0.001637 |
| *Cep55* | 0.341637276 | 1.47E-05 |
| *Csrp2* | 0.341474859 | 0.033875 |
| *Ska3* | 0.341435594 | 0.003548 |
| *Prkg2* | 0.341318669 | 2.52E-06 |
| *Polr1a* | 0.34076979 | 1.83E-09 |
| *Zbtb10* | 0.340420457 | 0.042401 |
| *Hnrnpdl* | 0.340356614 | 9.20E-10 |
| *Hilpda* | 0.340204009 | 0.019669 |

| | | |
|---|---|---|
| *Rfc4* | 0.34009948 | 0.000103 |
| *Itpr1* | 0.339064954 | 0.002495 |
| *Melk* | 0.338902898 | 6.71E-05 |
| *Zc3h8* | 0.338705988 | 0.007187 |
| *Sox2* | 0.338648759 | 0.000124 |
| *Nusap1* | 0.338415996 | 7.61E-08 |
| *Slc30a4* | 0.338400231 | 3.94E-08 |
| *Spdl1* | 0.338340479 | 7.71E-05 |
| *Rrm1* | 0.338260335 | 8.13E-12 |
| *Npm3* | 0.338165114 | 6.83E-08 |
| *Cdc7* | 0.337084447 | 0.00085 |
| *Sacs* | 0.336869238 | 0.000922 |
| *Fpgs* | 0.33650483 | 6.65E-05 |
| *Cenpo* | 0.336228725 | 0.00853 |
| *Mboat1* | 0.33618362 | 1.74E-06 |
| *Wdr77* | 0.335930048 | 2.25E-05 |
| *Cep152* | 0.33586817 | 9.81E-05 |
| *Serpinb2* | 0.335457553 | 0.000263 |
| *Tmem109* | 0.335404725 | 3.42E-05 |
| *Pdcd2* | 0.335282737 | 0.002424 |
| *Nek7* | 0.334949504 | 5.16E-07 |
| *Rmi2* | 0.334474212 | 0.037482 |
| *Plk3* | 0.334403419 | 0.000113 |
| *Mtcl1* | 0.334356069 | 1.61E-06 |
| *RP23-9L16.5* | 0.33385995 | 7.32E-06 |
| *Nop56* | 0.333788457 | 1.23E-08 |
| *Wee1* | 0.333732603 | 0.000722 |
| *Vars* | 0.333704534 | 3.29E-08 |
| *Heatr1* | 0.333494627 | 5.32E-09 |
| *Mcm10* | 0.333361041 | 8.09E-06 |
| *Xrcc6* | 0.333292133 | 2.14E-05 |
| *Hjurp* | 0.332718614 | 1.63E-08 |
| *Parpbp* | 0.332424528 | 0.001236 |
| *Prmt1* | 0.332288835 | 3.93E-06 |
| *D030056L22Rik* | 0.332171765 | 0.00093 |
| *Pwp2* | 0.331679942 | 4.86E-08 |
| *Rpp14* | 0.33167441 | 6.11E-05 |
| *Suv39h2* | 0.331508978 | 0.003149 |
| *Npm1* | 0.331302137 | 2.02E-08 |
| *Sephs2* | 0.331192403 | 3.54E-07 |
| *Slfn9* | 0.330774858 | 0.000261 |
| *Utp15* | 0.330699805 | 1.26E-06 |
| *Timm9* | 0.329978488 | 0.004833 |
| *Fam110c* | 0.329577263 | 0.000532 |

| | | |
|---|---|---|
| *Plekho2* | 0.329186018 | 0.000309 |
| *Apobr* | 0.329098123 | 0.020931 |
| *Mcm6* | 0.327912329 | 1.70E-12 |
| *Usp1* | 0.327478379 | 6.63E-05 |
| *Hnrnpd* | 0.327443988 | 3.10E-05 |
| *Dpy19l1* | 0.32742721 | 9.56E-07 |
| *Ddias* | 0.326722379 | 9.64E-06 |
| *Anp32e* | 0.32645772 | 0.000151 |
| *Pecr* | 0.32633265 | 0.025316 |
| *Sgol2a* | 0.326190288 | 1.79E-05 |
| *Gins3* | 0.326051292 | 0.009088 |
| *Rnf220* | 0.326033393 | 0.000131 |
| *Cdkn3* | 0.325681838 | 0.00839 |
| *Pmf1* | 0.325643536 | 0.001915 |
| *Pcnt* | 0.325316575 | 2.00E-08 |
| *Mdc1* | 0.325271637 | 0.007072 |
| *Qtrt1* | 0.324923148 | 0.002529 |
| *Ttll4* | 0.324682053 | 2.86E-07 |
| *Chka* | 0.324496534 | 2.96E-07 |
| *Zfp472* | 0.32383271 | 0.029622 |
| *Hspd1* | 0.323472503 | 1.75E-10 |
| *Cbx3* | 0.322745922 | 0.001631 |
| *Snhg5* | 0.322723372 | 1.53E-06 |
| *Rrs1* | 0.3216548 | 6.74E-06 |
| *Dnmt1* | 0.320970635 | 7.27E-08 |
| *Far1* | 0.320647022 | 1.92E-05 |
| *Ddx18* | 0.320450708 | 3.43E-06 |
| *Foxc1* | 0.320371323 | 0.000995 |
| *Nrp2* | 0.319949123 | 4.80E-06 |
| *Dram1* | 0.319669434 | 0.000955 |
| *Rfc3* | 0.319626069 | 0.000225 |
| *Mdn1* | 0.319579784 | 3.31E-07 |
| *Nop14* | 0.319530008 | 8.52E-07 |
| *Arhgef39* | 0.319461988 | 0.021863 |
| *Noa1* | 0.319084338 | 0.000281 |
| *Nup43* | 0.318898201 | 9.96E-06 |
| *Fjx1* | 0.318782859 | 2.94E-05 |
| *Rbmxl1* | 0.318738871 | 0.001302 |
| *Ssscа1* | 0.318632736 | 0.000182 |
| *Dusp6* | 0.318630727 | 0.00127 |
| *Taf1d* | 0.318458626 | 4.14E-05 |
| *Dtl* | 0.318346423 | 4.75E-06 |
| *Chaf1a* | 0.318149361 | 8.07E-06 |
| *Arhgef1* | 0.317724202 | 4.57E-05 |

| | | |
|---|---|---|
| *Adgrl2* | 0.317589278 | 0.01161 |
| *Nup205* | 0.316856916 | 3.43E-10 |
| *Mpp6* | 0.316085267 | 5.31E-05 |
| *Ppih* | 0.315938411 | 0.00748 |
| *Ces2e* | 0.315930456 | 6.47E-05 |
| *RP23-95L9.5* | 0.315927661 | 0.045373 |
| *Kif23* | 0.315821114 | 3.76E-06 |
| *Cenpl* | 0.315716405 | 0.003806 |
| *Nsd2* | 0.315610069 | 6.07E-08 |
| *Kpna2* | 0.315541797 | 9.00E-07 |
| *Socs1* | 0.315368419 | 0.03102 |
| *Pms2* | 0.315278127 | 0.003138 |
| *Gas2l3* | 0.315104907 | 0.000108 |
| *Gramd4* | 0.314963784 | 0.000151 |
| *Mettl13* | 0.314608803 | 0.000334 |
| *Utp14a* | 0.31434641 | 1.86E-06 |
| *Telo2* | 0.314152756 | 0.000981 |
| *Noc4l* | 0.314068189 | 8.21E-05 |
| *Lin54* | 0.313809864 | 7.34E-05 |
| *Mcam* | 0.313800303 | 6.12E-08 |
| *Uck2* | 0.313544472 | 3.94E-08 |
| *Urb1* | 0.31330314 | 9.75E-08 |
| *Sdad1* | 0.313259439 | 8.04E-08 |
| *RP23-185B14.1* | 0.313226615 | 0.040563 |
| *Ccnb2* | 0.312258269 | 1.10E-05 |
| *Psat1* | 0.311498115 | 2.97E-11 |
| *Dnajc9* | 0.311387122 | 7.39E-05 |
| *Prim1* | 0.311352189 | 5.11E-05 |
| *Tmpo* | 0.311334471 | 4.17E-08 |
| *Rfc1* | 0.311244255 | 2.87E-09 |
| *Vars2* | 0.310656997 | 0.000555 |
| *Mcm8* | 0.3105868 | 0.007687 |
| *Slc12a2* | 0.310028119 | 1.01E-07 |
| *Dsn1* | 0.309887039 | 0.001486 |
| *Fancm* | 0.30898157 | 0.000141 |
| *Map3k14* | 0.3088933 | 0.008617 |
| *Gm25878* | 0.308492173 | 0.03186 |
| *Gfer* | 0.308363422 | 0.000329 |
| *Chek2* | 0.308308106 | 0.000244 |
| *Gpn3* | 0.308150775 | 0.000722 |
| *Fxn* | 0.30813873 | 0.022627 |
| *Dgkd* | 0.307942707 | 1.39E-05 |
| *Slbp* | 0.307356571 | 8.62E-06 |
| *Paics* | 0.307190217 | 7.25E-08 |

| | | |
|---|---|---|
| *Rhoc* | 0.30717742 | 0.040641 |
| *Sbsn* | 0.307153088 | 1.20E-06 |
| *Fam216a* | 0.307027693 | 0.000661 |
| *Ruvbl1* | 0.30699062 | 0.000811 |
| *Ddx20* | 0.30673715 | 3.90E-05 |
| *Rpp38* | 0.306608749 | 0.003726 |
| *D2Wsu81e* | 0.306357007 | 0.000553 |
| *Gnb1l* | 0.306088186 | 0.015464 |
| *Noc2l* | 0.305952203 | 1.12E-05 |
| *Polr2h* | 0.305726206 | 0.000177 |
| *Tyw1* | 0.305669148 | 0.00033 |
| *P2rx7* | 0.305527863 | 0.044166 |
| *Nob1* | 0.305248446 | 0.001509 |
| *Galnt18* | 0.304666995 | 0.000811 |
| *Acer3* | 0.304264689 | 1.23E-05 |
| *Otud1* | 0.304234413 | 0.000778 |
| *Gcsh* | 0.30410999 | 1.61E-05 |
| *Faap24* | 0.303730465 | 0.039115 |
| *Mnd1* | 0.303448274 | 0.028645 |
| *Polr3g* | 0.303445354 | 4.77E-05 |
| *Spata5* | 0.302842491 | 0.000181 |
| *Ipo5* | 0.302204595 | 1.99E-12 |
| *Haus6* | 0.302199585 | 9.31E-05 |
| *Wdr90* | 0.301943561 | 0.001779 |
| *Rps6ka2* | 0.301473614 | 0.016483 |
| *Cpsf3* | 0.301179248 | 7.61E-05 |
| *Nop58* | 0.30076325 | 3.52E-07 |
| *Sart3* | 0.300304856 | 5.48E-07 |
| *Ilf3* | 0.300287269 | 1.02E-06 |
| *Fhl1* | 0.300232914 | 0.000195 |
| *Sgsm2* | -0.300183167 | 0.003089 |
| *Samd5* | -0.300413566 | 0.0002 |
| *Sh3pxd2a* | -0.300426868 | 1.28E-13 |
| *Il24* | -0.300716506 | 7.36E-05 |
| *Aff1* | -0.300771993 | 1.70E-06 |
| *Eef2k* | -0.30095853 | 3.83E-06 |
| *Baiap2l1* | -0.301088615 | 0.017602 |
| *Slc2a1* | -0.301593128 | 2.49E-08 |
| *Slc35f6* | -0.301794983 | 1.00E-05 |
| *Pthlh* | -0.301851933 | 8.66E-09 |
| *Ocrl* | -0.302162452 | 0.001358 |
| *Sphk1* | -0.302349816 | 0.036029 |
| *Npepps* | -0.303042672 | 7.61E-07 |
| *Nhsl1* | -0.303478668 | 0.022513 |

| | | |
|---|---|---|
| *Mical3* | -0.303547553 | 3.92E-06 |
| *Slc44a1* | -0.303721106 | 3.61E-09 |
| *Nectin1* | -0.304635424 | 0.000137 |
| *Cpt2* | -0.304734232 | 0.017733 |
| *Igfbp7* | -0.305512097 | 0.000218 |
| *Lynx1* | -0.305538043 | 0.003431 |
| *Rab8b* | -0.306663892 | 5.57E-06 |
| *Plcd1* | -0.30669871 | 5.57E-06 |
| *Ube2j1* | -0.306817824 | 3.30E-07 |
| *Slc29a3* | -0.30693697 | 0.020657 |
| *Tbc1d2b* | -0.307155486 | 3.62E-09 |
| *Fmn1* | -0.307579164 | 0.000704 |
| *Creb3* | -0.308271299 | 1.65E-05 |
| *Plscr3* | -0.30869928 | 0.000137 |
| *Frmd4b* | -0.309249145 | 4.84E-10 |
| *Bcl9l* | -0.309452699 | 0.014054 |
| *Zfp185* | -0.309477071 | 0.004254 |
| *Spint1* | -0.309805354 | 1.35E-06 |
| *Fndc3b* | -0.310107348 | 1.09E-07 |
| *Dag1* | -0.310225749 | 1.22E-08 |
| *Sec11c* | -0.310334053 | 9.50E-05 |
| *Tenm4* | -0.310609814 | 0.000252 |
| *Atp2c1* | -0.31063257 | 5.17E-08 |
| *Nfkbiz* | -0.310904295 | 3.22E-06 |
| *RP23-128C4.4* | -0.311267834 | 0.026035 |
| *Tmprss13* | -0.311586389 | 0.005813 |
| *Ttll5* | -0.312349296 | 0.00012 |
| *Tmem50b* | -0.312495828 | 0.001587 |
| *Bmp2k* | -0.312893606 | 8.94E-07 |
| *Lgr4* | -0.313262134 | 2.58E-06 |
| *Rnase4* | -0.313509692 | 1.12E-05 |
| *Zfp9* | -0.313731922 | 0.00562 |
| *Vamp7* | -0.314319713 | 0.041294 |
| *mt-Rnr1* | -0.315116739 | 1.06E-06 |
| *Aplp2* | -0.315683193 | 2.47E-09 |
| *Zdhhc9* | -0.316022418 | 0.009938 |
| *Chsy1* | -0.316415978 | 2.60E-07 |
| *Cxcl5* | -0.316472265 | 2.55E-06 |
| *Tapbpl* | -0.317250244 | 0.002293 |
| *Laptm4a* | -0.317727549 | 8.93E-10 |
| *Tacc2* | -0.317956116 | 0.000118 |
| *Slc35a2* | -0.318011239 | 0.003062 |
| *Pxk* | -0.318092802 | 0.000185 |
| *Nog* | -0.318254721 | 0.007994 |

| | | |
|---|---|---|
| *Ell2* | -0.318718863 | 7.71E-05 |
| *Klhdc1* | -0.318808826 | 0.047222 |
| *Prl2c2* | -0.318991518 | 6.30E-05 |
| *Trp53bp2* | -0.319065091 | 0.004033 |
| *Reep5* | -0.319290584 | 4.57E-09 |
| *Pard6g* | -0.319697344 | 2.19E-08 |
| *Ube2h* | -0.319850725 | 8.81E-05 |
| *Ttc39c* | -0.320059482 | 7.23E-07 |
| *Fgd6* | -0.320511675 | 0.000118 |
| *Runx1* | -0.320872695 | 2.89E-06 |
| *Pcdhgb6* | -0.321362565 | 0.007366 |
| *Atp6v1e1* | -0.321799518 | 1.36E-08 |
| *Gpr68* | -0.322434437 | 0.042401 |
| *Tubb2a* | -0.322926135 | 3.22E-07 |
| *Mrc2* | -0.323120879 | 0.042278 |
| *Zhx3* | -0.32325563 | 0.010102 |
| *Fabp5* | -0.323612981 | 9.86E-09 |
| *Gpr160* | -0.323859311 | 0.044656 |
| *Elavl2* | -0.324540401 | 0.045827 |
| *Fgfrl1* | -0.32460253 | 0.033705 |
| *Fzd2* | -0.325794601 | 0.000467 |
| *Slc16a9* | -0.326084025 | 0.001632 |
| *Klhl24* | -0.326574417 | 0.000792 |
| *Krt6a* | -0.326599242 | 3.01E-12 |
| *Map7d1* | -0.327048883 | 3.45E-09 |
| *Lxn* | -0.327304404 | 0.000436 |
| *Tiparp* | -0.327320684 | 3.30E-07 |
| *Sqrdl* | -0.327428232 | 7.36E-05 |
| *Las1l* | -0.32766078 | 0.001705 |
| *Lonrf1* | -0.328569018 | 0.009055 |
| *Grip1* | -0.328859442 | 7.40E-06 |
| *Atp6v1g1* | -0.32892776 | 1.51E-07 |
| *Ppfibp2* | -0.329031413 | 0.005274 |
| *Ahnak2* | -0.329177992 | 0.000868 |
| *Ugcg* | -0.329246418 | 4.84E-06 |
| *Capn2* | -0.329506787 | 1.59E-14 |
| *Hmox1* | -0.329895305 | 3.54E-05 |
| *Cd59a* | -0.330199762 | 0.000328 |
| *Lgals9* | -0.330238822 | 0.004034 |
| *Synrg* | -0.330978048 | 0.0008 |
| *Srl* | -0.331300949 | 0.005712 |
| *Fam214a* | -0.331308087 | 0.010423 |
| *Map4* | -0.331933498 | 5.02E-09 |
| *Tmcc1* | -0.332094921 | 0.000308 |

| | | |
|---|---|---|
| *Mfsd9* | -0.332251665 | 0.013829 |
| *Map6* | -0.332418075 | 6.54E-07 |
| *Nlgn2* | -0.332717531 | 0.041645 |
| *Dsg2* | -0.332786265 | 2.18E-06 |
| *St5* | -0.332932203 | 2.37E-07 |
| *Aak1* | -0.333132024 | 0.000718 |
| *Plec* | -0.333581656 | 3.16E-09 |
| *Ak1* | -0.334002457 | 9.18E-08 |
| *Bag3* | -0.334757934 | 6.90E-06 |
| *Natd1* | -0.334891596 | 0.007423 |
| *Atxn1* | -0.335646982 | 4.96E-10 |
| *Sec23a* | -0.335977012 | 1.06E-07 |
| *Lamp2* | -0.337331463 | 1.38E-09 |
| *Shb* | -0.33885984 | 1.56E-06 |
| *Sh3bgrl2* | -0.339071397 | 2.44E-08 |
| *Prrc1* | -0.339325379 | 0.000253 |
| *Slitrk6* | -0.340333613 | 7.14E-06 |
| *Samd4b* | -0.340457679 | 0.002697 |
| *Megf9* | -0.340672632 | 3.14E-05 |
| *Ptpn23* | -0.341460058 | 0.006026 |
| *Tnfaip3* | -0.341684334 | 7.20E-11 |
| *Gramd2* | -0.341988825 | 6.25E-05 |
| *Ptgfrn* | -0.34259666 | 8.37E-16 |
| *Pex11g* | -0.34261801 | 0.027427 |
| *Jup* | -0.34346749 | 4.67E-13 |
| *Cpeb2* | -0.344089704 | 0.000221 |
| *Fam210b* | -0.344552748 | 0.000905 |
| *Lrp1* | -0.345200891 | 1.30E-11 |
| *Itpripl2* | -0.345316909 | 4.74E-16 |
| *P4ha2* | -0.345516775 | 1.31E-05 |
| *Zfp182* | -0.345744657 | 0.014656 |
| *Itgb5* | -0.345898251 | 1.88E-11 |
| *Slit3* | -0.345946675 | 0.011346 |
| *Blnk* | -0.346152644 | 1.34E-07 |
| *Itga6* | -0.34633415 | 4.65E-16 |
| *S100a9* | -0.346585835 | 0.000275 |
| *Pcdhb22* | -0.347613277 | 0.017986 |
| *Rap1gap2* | -0.348093633 | 0.001431 |
| *Samd9l* | -0.348566308 | 0.000191 |
| *Sh3rf1* | -0.348750959 | 2.69E-05 |
| *Cpeb3* | -0.349024573 | 0.038777 |
| *Duoxa1* | -0.349118366 | 1.84E-07 |
| *Rnf152* | -0.349629243 | 0.022043 |
| *Il1rn* | -0.350128221 | 0.001153 |

| | | |
|---|---|---|
| *Dnmbp* | -0.35077974 | 1.24E-06 |
| *Adrb2* | -0.35188931 | 5.53E-06 |
| *Atp10d* | -0.351914224 | 2.72E-13 |
| *Ngfrap1* | -0.352201844 | 5.66E-07 |
| *Fam101b* | -0.352384766 | 5.91E-10 |
| *Atp6ap2* | -0.352544137 | 3.08E-08 |
| *Msx1* | -0.353036872 | 0.012706 |
| *Mvb12b* | -0.353150649 | 0.001544 |
| *Sema4c* | -0.353212763 | 6.47E-05 |
| *Fgd4* | -0.353494986 | 0.00697 |
| *Pcyt1b* | -0.354693386 | 0.001081 |
| *Foxj2* | -0.354966763 | 0.004784 |
| *Plpp5* | -0.355186654 | 0.000847 |
| *Lrrc32* | -0.356689499 | 3.14E-06 |
| *Il4ra* | -0.35681881 | 4.69E-10 |
| *Klhl18* | -0.357359598 | 0.000555 |
| *Wdtc1* | -0.357438499 | 7.61E-05 |
| *Calcoco1* | -0.357728259 | 0.023903 |
| *Nucb2* | -0.358676288 | 3.28E-05 |
| *Abca2* | -0.359463367 | 0.000143 |
| *Kdm5b* | -0.359896745 | 4.45E-11 |
| *Fam43a* | -0.360321426 | 0.000282 |
| *Gpd1l* | -0.36066851 | 0.001758 |
| *Map1lc3b* | -0.360803592 | 9.64E-06 |
| *Rnf122* | -0.362445908 | 0.001277 |
| *Dnajb9* | -0.362469399 | 0.024158 |
| *Wnt5a* | -0.362553676 | 7.23E-11 |
| *8-Mar* | -0.362886741 | 9.76E-09 |
| *Gne* | -0.363094033 | 0.03195 |
| *Aph1b* | -0.363460416 | 0.000862 |
| *Pfkfb3* | -0.3641724 | 2.03E-07 |
| *Dlg5* | -0.364181545 | 1.20E-06 |
| *Garem1* | -0.365078361 | 0.004434 |
| *Lhx6* | -0.365088317 | 0.016599 |
| *Wnk1* | -0.365317444 | 1.34E-09 |
| *Klhl7* | -0.365334019 | 1.03E-05 |
| *RP23-73N8.4* | -0.366151819 | 0.042728 |
| *Pik3r3* | -0.36706068 | 0.000455 |
| *Unc13d* | -0.367345672 | 0.000765 |
| *Hk1* | -0.368066734 | 1.80E-09 |
| *Pitpnm3* | -0.368477233 | 0.000164 |
| *Cobl* | -0.369934083 | 4.63E-05 |
| *Thnsl1* | -0.370821678 | 0.007423 |
| *Ndrg1* | -0.371171344 | 2.00E-07 |

| | | |
|---|---|---|
| *Camsap3* | -0.371347401 | 0.00288 |
| *Trim29* | -0.373159034 | 2.89E-09 |
| *Ampd3* | -0.373656237 | 9.27E-10 |
| *Ispd* | -0.373869851 | 0.044874 |
| *Ptpru* | -0.374175436 | 9.88E-07 |
| *Evpl* | -0.374450577 | 8.70E-09 |
| *Sh3tc2* | -0.374626021 | 1.34E-05 |
| *Wdpcp* | -0.374710113 | 0.012429 |
| *Trnp1* | -0.374787672 | 0.010255 |
| *Sh3kbp1* | -0.375180915 | 2.31E-14 |
| *L3mbtl3* | -0.375742806 | 4.15E-08 |
| *Dsg3* | -0.376249575 | 1.93E-17 |
| *Serinc2* | -0.377137268 | 3.64E-09 |
| *Ap3m2* | -0.377689028 | 0.038551 |
| *Tuft1* | -0.378107926 | 7.89E-11 |
| *Phlda1* | -0.378350358 | 0.001616 |
| *Cpt1a* | -0.378930535 | 7.78E-08 |
| *Wipi1* | -0.37893474 | 1.85E-09 |
| *Loxl4* | -0.379495399 | 1.82E-05 |
| *Cd24a* | -0.379513509 | 1.51E-15 |
| *Ifi203* | -0.380349629 | 0.026202 |
| *Nt5dc2* | -0.381027502 | 2.57E-09 |
| *Gata3* | -0.381238886 | 0.027317 |
| *Dmxl1* | -0.381406182 | 1.70E-08 |
| *Fyco1* | -0.381922591 | 0.023268 |
| *Nipal2* | -0.382579599 | 0.00057 |
| *Rab27b* | -0.383459536 | 6.53E-06 |
| *Afap1l2* | -0.383510378 | 4.11E-09 |
| *Abr* | -0.384399083 | 5.01E-13 |
| *Smpd1* | -0.384845943 | 6.49E-05 |
| *Fam214b* | -0.384879206 | 6.47E-08 |
| *RP24-180L12.6* | -0.385161699 | 0.018752 |
| *Cited2* | -0.385567353 | 0.002496 |
| *Aes* | -0.385861537 | 7.45E-06 |
| *Metrnl* | -0.386036155 | 0.000358 |
| *Cldn4* | -0.386195031 | 2.18E-09 |
| *Gatm* | -0.386631741 | 8.42E-05 |
| *Ulk2* | -0.38698906 | 0.000138 |
| *Tspan13* | -0.387093368 | 0.020481 |
| *Apol9a* | -0.387412077 | 0.037081 |
| *Wisp1* | -0.387631614 | 3.66E-06 |
| *Hsdl2* | -0.387844497 | 8.34E-07 |
| *Erp44* | -0.388107146 | 7.15E-10 |
| *Strbp* | -0.389363197 | 4.17E-12 |

| | | |
|---|---|---|
| *Tinagl1* | -0.389364742 | 2.24E-10 |
| *Ctns* | -0.389445801 | 0.012347 |
| *Sox4* | -0.389641283 | 1.93E-18 |
| *Trim12c* | -0.390465201 | 0.001095 |
| *Kank1* | -0.390573046 | 6.70E-13 |
| *Pcdhb7* | -0.39151339 | 0.041428 |
| *Ykt6* | -0.391878618 | 2.14E-14 |
| *Gas6* | -0.39260883 | 0.002345 |
| *Zfp937* | -0.393256274 | 0.008378 |
| *Aqp3* | -0.393849751 | 2.41E-07 |
| *Ttyh3* | -0.394634186 | 6.33E-07 |
| *Dhrs9* | -0.394771466 | 0.006403 |
| *Gadd45a* | -0.395591618 | 4.98E-09 |
| *C1s1* | -0.39601358 | 0.00158 |
| *Dhrs3* | -0.396162234 | 0.030829 |
| *Dok2* | -0.396560712 | 0.035714 |
| *Alg2* | -0.397265693 | 0.007773 |
| *Rnf144b* | -0.39742531 | 5.53E-06 |
| *Shpk* | -0.399523469 | 0.048495 |
| *Tcf7l2* | -0.399794409 | 0.044967 |
| *Sparc* | -0.40016722 | 3.17E-17 |
| *Gba2* | -0.40051655 | 0.001558 |
| *Trim30a* | -0.401322645 | 0.04754 |
| *Rtn4rl1* | -0.40189747 | 0.034401 |
| *RP24-528H20.5* | -0.403424002 | 0.038454 |
| *Herc3* | -0.403508137 | 0.001632 |
| *Pde4dip* | -0.403621756 | 1.61E-08 |
| *Naaa* | -0.403759552 | 1.90E-05 |
| *Kifc3* | -0.404214741 | 1.10E-05 |
| *Mrgpre* | -0.404624309 | 0.034785 |
| *Zbp1* | -0.404810903 | 0.043504 |
| *Adarb1* | -0.405377951 | 7.02E-12 |
| *Pold4* | -0.405694742 | 0.000138 |
| *Ret* | -0.406006575 | 0.036027 |
| *Amica1* | -0.406161447 | 0.043847 |
| *Etl4* | -0.407208341 | 5.02E-09 |
| *Orai2* | -0.407718254 | 9.79E-08 |
| *Pnpla1* | -0.408025733 | 0.043326 |
| *Mcub* | -0.408807183 | 0.0033 |
| *Snrk* | -0.409103587 | 2.59E-09 |
| *Clic4* | -0.409718581 | 3.93E-20 |
| *Bmpr2* | -0.410011256 | 7.41E-10 |
| *RP24-90D21.3* | -0.410021255 | 1.74E-06 |
| *Bmp1* | -0.410860481 | 1.99E-18 |

| | | |
|---|---|---|
| *Jag1* | -0.411791592 | 1.28E-20 |
| *Cadm4* | -0.412666829 | 3.17E-07 |
| *Shank2* | -0.413149707 | 0.039778 |
| *Col4a4* | -0.413245808 | 0.039529 |
| *Adrb1* | -0.413686226 | 0.000123 |
| *Pld1* | -0.414142802 | 0.00039 |
| *Gm26888* | -0.414279966 | 0.037191 |
| *Fam46a* | -0.41491679 | 0.005001 |
| *Dnpep* | -0.415516167 | 5.09E-10 |
| *Numbl* | -0.41623545 | 0.006979 |
| *Lamc2* | -0.416580987 | 3.77E-16 |
| *Bcl3* | -0.417271579 | 0.02416 |
| *Neu3* | -0.417470248 | 0.000191 |
| *Pard3b* | -0.41757403 | 0.019017 |
| *Tlr2* | -0.417842437 | 3.50E-06 |
| *Ror2* | -0.418206937 | 0.013013 |
| *Rbms3* | -0.418861112 | 0.00725 |
| *Saa1* | -0.419696534 | 0.029485 |
| *Micall1* | -0.420703381 | 1.19E-15 |
| *Cyp2u1* | -0.420853133 | 0.030028 |
| *Ddr1* | -0.421106375 | 4.00E-08 |
| *Plpp3* | -0.422217345 | 0.001328 |
| *Kcnab2* | -0.423138416 | 0.022113 |
| *Chst2* | -0.423256558 | 4.04E-08 |
| *Zmiz2* | -0.423965313 | 0.005115 |
| *Fbxl20* | -0.424114716 | 2.15E-05 |
| *Ndfip1* | -0.424427477 | 1.61E-10 |
| *Dlc1* | -0.424750567 | 5.28E-05 |
| *Plekha2* | -0.424902543 | 3.34E-12 |
| *Clca3a2* | -0.425581222 | 0.026631 |
| *Map2k6* | -0.426368438 | 0.00206 |
| *Ugt1a7c* | -0.427117333 | 0.002614 |
| *Casq2* | -0.427133508 | 1.00E-10 |
| *Pim1* | -0.428049681 | 0.000555 |
| *Kif26a* | -0.428480128 | 9.49E-11 |
| *Sprr2h* | -0.428639547 | 0.026191 |
| *Tns4* | -0.42969601 | 3.53E-20 |
| *Rcan3* | -0.429790116 | 7.80E-06 |
| *Dstyk* | -0.430893263 | 3.58E-10 |
| *Pcdhb17* | -0.431167136 | 9.50E-05 |
| *Dclk1* | -0.431286838 | 7.54E-05 |
| *Spsb1* | -0.432505812 | 0.000447 |
| *Tceal9* | -0.433339382 | 1.91E-10 |
| *Gab3* | -0.434299259 | 0.026354 |

| | | |
|---|---|---|
| *Papss2* | -0.436182814 | 1.36E-10 |
| *Nacc2* | -0.43619259 | 4.84E-10 |
| *Krt6b* | -0.437068287 | 8.99E-20 |
| *Itgb4* | -0.437246544 | 3.80E-22 |
| *Ikbip* | -0.437564872 | 3.30E-06 |
| *Kdelr3* | -0.438805263 | 3.41E-06 |
| *Cux1* | -0.43907873 | 2.90E-18 |
| *Cbx4* | -0.439272348 | 1.08E-09 |
| *Plxnd1* | -0.440898068 | 1.17E-06 |
| *Spire1* | -0.441445275 | 1.96E-09 |
| *RP23-49E21.2* | -0.441490199 | 0.025538 |
| *Prag1* | -0.441906444 | 4.28E-08 |
| *Foxp1* | -0.441971217 | 1.10E-14 |
| *Mxi1* | -0.442164693 | 3.90E-09 |
| *Phldb2* | -0.442748356 | 1.62E-22 |
| *Ric3* | -0.445821526 | 0.023339 |
| *Ccdc68* | -0.446878716 | 0.009735 |
| *Zbtb4* | -0.446937701 | 1.30E-06 |
| *Ppbp* | -0.447288406 | 1.23E-16 |
| *Neurl1a* | -0.447907228 | 2.23E-07 |
| *Pdcd4* | -0.448582594 | 0.000664 |
| *Mgll* | -0.448906113 | 0.00132 |
| *Nhs* | -0.449813271 | 0.001112 |
| *Pof1b* | -0.451193059 | 3.92E-07 |
| *Hpgds* | -0.45185428 | 0.000955 |
| *RP23-230O9.4* | -0.452489874 | 0.012691 |
| *Ccser1* | -0.454050829 | 0.008081 |
| *Arhgef3* | -0.455841954 | 8.19E-08 |
| *Mroh3* | -0.456083535 | 0.008713 |
| *Dsc3* | -0.456888845 | 1.34E-28 |
| *Mfsd6* | -0.457778014 | 1.80E-13 |
| *Rab11fip2* | -0.45966426 | 1.77E-11 |
| *Sash1* | -0.459799745 | 4.83E-13 |
| *Hoxb9* | -0.459864984 | 0.003039 |
| *Nfat5* | -0.460140866 | 9.64E-06 |
| *Rap1gap* | -0.461406289 | 0.011511 |
| *Creb3l2* | -0.464998522 | 1.87E-07 |
| *RP24-332N10.3* | -0.465168022 | 0.008044 |
| *Ppp1r12b* | -0.465316249 | 2.50E-07 |
| *Ggt5* | -0.465327217 | 0.016846 |
| *Prl8a1* | -0.466807997 | 5.78E-12 |
| *Mgat4a* | -0.466904166 | 1.02E-07 |
| *Orm1* | -0.468255771 | 0.015146 |
| *Ttll7* | -0.469126124 | 1.32E-07 |

| | | |
|---|---|---|
| *Flrt2* | -0.470745701 | 7.43E-21 |
| *Smoc2* | -0.470767213 | 0.014799 |
| *Fam20a* | -0.47102843 | 2.97E-07 |
| *Chdh* | -0.473238655 | 0.01545 |
| *Gprc5a* | -0.473422553 | 3.63E-08 |
| *Kctd11* | -0.474374692 | 1.66E-08 |
| *Kcnh1* | -0.474873847 | 2.03E-07 |
| *Sh3tc1* | -0.476123508 | 3.22E-06 |
| *Id2* | -0.479814576 | 1.19E-10 |
| *Palmd* | -0.480922489 | 0.001419 |
| *Itpkb* | -0.483051252 | 5.91E-10 |
| *Nav1* | -0.484375062 | 7.05E-11 |
| *Dlx1* | -0.485390278 | 0.003254 |
| *Sulf1* | -0.485955918 | 3.54E-16 |
| *Nebl* | -0.486213546 | 8.08E-06 |
| *RP23-44N5.4* | -0.486755281 | 0.008687 |
| *Rora* | -0.488226852 | 0.004588 |
| *Maged2* | -0.489391679 | 9.47E-14 |
| *Lamb3* | -0.490070353 | 1.07E-26 |
| *Raph1* | -0.490168485 | 1.84E-06 |
| *Mtss1* | -0.491577333 | 5.43E-10 |
| *Grhl3* | -0.49217342 | 0.000235 |
| *Ripk3* | -0.492911843 | 8.23E-06 |
| *Shroom1* | -0.492960105 | 9.84E-06 |
| *Mmp9* | -0.49554733 | 0.009055 |
| *Fam167a* | -0.496142259 | 5.46E-24 |
| *RP24-263P17.1* | -0.496158139 | 0.009386 |
| *Wnt11* | -0.49620413 | 0.006107 |
| *Fxyd4* | -0.498254127 | 0.008884 |
| *Cst6* | -0.49827448 | 1.26E-06 |
| *RP23-442L7.4* | -0.501574029 | 0.000101 |
| *Fam114a1* | -0.502931828 | 1.22E-09 |
| *Vwa7* | -0.503080112 | 0.00033 |
| *Pkd1l3* | -0.503248087 | 0.008988 |
| *Prl2c3* | -0.504096028 | 3.76E-06 |
| *RP23-133N21.3* | -0.505037267 | 0.005221 |
| *S100a8* | -0.5052839 | 1.72E-08 |
| *Myl7* | -0.506224399 | 0.004585 |
| *Narf* | -0.511311801 | 3.76E-09 |
| *Celf2* | -0.512503494 | 0.001568 |
| *Dok4* | -0.513042134 | 6.41E-06 |
| *Dsg1b* | -0.514029902 | 1.39E-08 |
| *Wnt9a* | -0.514210595 | 1.68E-15 |
| *Fam83c* | -0.515345583 | 0.006299 |

| | | |
|---|---|---|
| *Sostdc1* | -0.516299481 | 0.00132 |
| *Ap5b1* | -0.516767461 | 3.60E-06 |
| *Spon2* | -0.518023575 | 7.57E-06 |
| *Lce1g* | -0.518931491 | 0.006452 |
| *Vdr* | -0.5191204 | 2.05E-11 |
| *Tgm2* | -0.522503528 | 9.20E-10 |
| *Mras* | -0.522971433 | 0.000458 |
| *Dusp1* | -0.524212298 | 3.16E-07 |
| *Arid3a* | -0.525057404 | 2.56E-06 |
| *Cobll1* | -0.52594396 | 2.77E-09 |
| *Col5a1* | -0.526916153 | 0.003772 |
| *Klhdc8a* | -0.526959712 | 1.81E-05 |
| *Spata13* | -0.527531275 | 3.83E-05 |
| *Serpinb1a* | -0.530979055 | 7.30E-05 |
| *Grhl1* | -0.531497656 | 7.16E-11 |
| *Itgb6* | -0.531642991 | 6.15E-28 |
| *Ccng2* | -0.532997056 | 1.42E-05 |
| *Skil* | -0.53444084 | 1.22E-27 |
| *Trps1* | -0.534736404 | 0.000534 |
| *Epb41l3* | -0.536917326 | 3.11E-07 |
| *Gse1* | -0.536924885 | 2.02E-06 |
| *Dsg1a* | -0.537438214 | 0.000136 |
| *Ifi27l2b* | -0.537933319 | 0.004735 |
| *Scube3* | -0.538154551 | 0.000372 |
| *Tgm3* | -0.540427038 | 3.90E-06 |
| *Pkp2* | -0.541077947 | 2.26E-15 |
| *Col5a2* | -0.541165573 | 0.001711 |
| *Sox9* | -0.544357405 | 4.61E-07 |
| *Ids* | -0.546243183 | 2.83E-07 |
| *Bhlhe41* | -0.547347926 | 0.000243 |
| *Adam10* | -0.548638962 | 1.01E-22 |
| *Atp2c2* | -0.552066355 | 6.18E-10 |
| *Pcdh1* | -0.552573193 | 0.0002 |
| *Paqr9* | -0.553072616 | 0.003464 |
| *Phyhipl* | -0.553527506 | 2.17E-07 |
| *Cpeb4* | -0.554465603 | 3.12E-10 |
| *Mcf2l* | -0.555129142 | 0.001197 |
| *Pcdhb19* | -0.555638026 | 0.001234 |
| *Parm1* | -0.55565045 | 9.81E-11 |
| *Gck* | -0.559681229 | 0.002185 |
| *Acta1* | -0.559745274 | 0.000981 |
| *Ctsf* | -0.55997888 | 0.002882 |
| *Atp9a* | -0.563767409 | 1.46E-15 |
| *Epb41l1* | -0.564623494 | 3.64E-11 |

| | | |
|---|---|---|
| *Lama3* | -0.565852211 | 2.23E-37 |
| *Stab1* | -0.571225711 | 7.03E-31 |
| *Ifi202b* | -0.572568924 | 1.23E-16 |
| *Soat1* | -0.576438305 | 1.93E-37 |
| *Hr* | -0.576534645 | 7.85E-24 |
| *Megf6* | -0.576761395 | 7.83E-15 |
| *Olfm2* | -0.577385917 | 0.000703 |
| *Hephl1* | -0.577405198 | 0.000783 |
| *Gpr35* | -0.577628749 | 4.72E-05 |
| *CAAA01093181.2* | -0.57914973 | 1.83E-05 |
| *Pla2g4e* | -0.57953598 | 0.001235 |
| *Cmah* | -0.579642079 | 2.17E-06 |
| *Bnip3* | -0.581035156 | 7.65E-17 |
| *Nuak1* | -0.581470766 | 8.58E-30 |
| *Vegfa* | -0.581765118 | 1.50E-12 |
| *Adgrf4* | -0.581773825 | 3.50E-09 |
| *Gadd45g* | -0.583334536 | 3.04E-08 |
| *C3ar1* | -0.583705609 | 0.001667 |
| *Ctsw* | -0.588226475 | 0.00138 |
| *Inhba* | -0.590256807 | 7.28E-08 |
| *Gnpda1* | -0.593499862 | 5.38E-15 |
| *Sorcs2* | -0.594185499 | 1.19E-14 |
| *Vgll3* | -0.594463933 | 1.52E-17 |
| *Cyth1* | -0.594627947 | 9.94E-28 |
| *Akap6* | -0.595480221 | 0.000365 |
| *Vldlr* | -0.596616925 | 1.24E-17 |
| *Ypel5* | -0.5993129 | 2.90E-15 |
| *Peg13* | -0.599956634 | 4.84E-09 |
| *Ncam1* | -0.60103124 | 9.11E-26 |
| *Mxd4* | -0.601041645 | 0.000159 |
| *Cpa4* | -0.601496642 | 4.22E-06 |
| *Dlx3* | -0.602400377 | 0.001027 |
| *Adgrg1* | -0.603445113 | 3.25E-13 |
| *Sptbn2* | -0.605414255 | 1.66E-20 |
| *Ndrg4* | -0.606791231 | 3.36E-07 |
| *Ctla2a* | -0.608429483 | 0.000651 |
| *Stmn2* | -0.609305747 | 0.000976 |
| *Adamts15* | -0.614571574 | 1.09E-09 |
| *Ccbe1* | -0.617641954 | 0.000846 |
| *Rab6b* | -0.618276567 | 0.000266 |
| *Fam19a5* | -0.618476908 | 0.000651 |
| *Id3* | -0.621224116 | 4.09E-18 |
| *Slc7a8* | -0.621918938 | 1.98E-08 |
| *Tmem140* | -0.623301125 | 4.79E-07 |

| | | |
|---|---|---|
| *Id1* | -0.624976818 | 3.80E-30 |
| *Socs2* | -0.628994325 | 7.85E-08 |
| *Hook1* | -0.634131824 | 9.01E-09 |
| *Zfp703* | -0.635225164 | 9.16E-06 |
| *Mylip* | -0.635231884 | 8.07E-06 |
| *Myh14* | -0.635680539 | 1.62E-06 |
| *Ahr* | -0.637293109 | 3.05E-17 |
| *Pvrl4* | -0.637588967 | 1.38E-22 |
| *Pmepa1* | -0.638570255 | 7.43E-41 |
| *Gsta3* | -0.640542276 | 3.95E-16 |
| *Camkk1* | -0.6436207 | 6.11E-08 |
| *Bhlhe40* | -0.644564381 | 6.75E-21 |
| *Zmiz1* | -0.647565279 | 7.67E-07 |
| *Pcdhb16* | -0.648140777 | 3.57E-13 |
| *Mapre2* | -0.649225583 | 1.49E-17 |
| *Unc13b* | -0.650399418 | 9.38E-06 |
| *Pkdcc* | -0.65153292 | 1.77E-07 |
| *Adra1b* | -0.656536554 | 0.000329 |
| *Dsc2* | -0.659395429 | 4.45E-23 |
| *Trp53inp2* | -0.663795407 | 8.38E-15 |
| *Arrb1* | -0.664517063 | 1.24E-07 |
| *Fstl1* | -0.664748233 | 1.74E-30 |
| *Pxylp1* | -0.667658063 | 1.44E-06 |
| *Optn* | -0.668891936 | 3.62E-19 |
| *Krt16* | -0.670371033 | 4.42E-30 |
| *Slc4a4* | -0.672403368 | 2.47E-05 |
| *Apol9b* | -0.676429061 | 0.000178 |
| *Trabd2b* | -0.676707038 | 0.000202 |
| *Dusp10* | -0.67713394 | 1.22E-29 |
| *Timp1* | -0.678062257 | 2.44E-20 |
| *Kcnk7* | -0.681080807 | 6.42E-06 |
| *Slc43a2* | -0.685376182 | 3.05E-12 |
| *Fbln2* | -0.690930535 | 2.12E-55 |
| *Tmcc3* | -0.692193558 | 7.55E-32 |
| *Ralgps1* | -0.69340894 | 3.46E-06 |
| *Rusc2* | -0.694657265 | 9.92E-18 |
| *Havcr2* | -0.697358832 | 3.30E-08 |
| *Vegfc* | -0.702109164 | 1.87E-11 |
| *Sesn3* | -0.704577959 | 4.83E-18 |
| *Fam63b* | -0.704959764 | 5.46E-24 |
| *Cpn1* | -0.705483625 | 9.25E-06 |
| *Thsd1* | -0.7058591 | 3.23E-09 |
| *Slc30a1* | -0.707501134 | 2.31E-11 |
| *Apol7a* | -0.708123319 | 2.08E-07 |

| | | |
|---|---|---|
| *Kif26b* | -0.712502177 | 3.93E-12 |
| *Sdc3* | -0.722871261 | 7.54E-22 |
| *Serpine1* | -0.727692039 | 1.38E-22 |
| *Tbx2* | -0.731140369 | 2.26E-06 |
| *Mill1* | -0.733707422 | 1.73E-05 |
| *Emx2* | -0.737026984 | 1.10E-09 |
| *Nat8l* | -0.740436791 | 1.59E-31 |
| *Snai3* | -0.750964565 | 1.37E-05 |
| *Atp6v0a1* | -0.75864267 | 3.80E-22 |
| *Hoxc13* | -0.758812866 | 4.31E-10 |
| *Fgf1* | -0.761850961 | 1.81E-05 |
| *Ldlrad4* | -0.762430315 | 8.16E-18 |
| *Smad6* | -0.762442356 | 1.44E-13 |
| *Gjb2* | -0.770055033 | 1.41E-14 |
| *Ifitm10* | -0.785144549 | 3.64E-07 |
| *Hs3st3a1* | -0.796115816 | 5.38E-12 |
| *Ifih1* | -0.801810623 | 7.91E-19 |
| *Smad7* | -0.802576664 | 2.05E-35 |
| *Tppp* | -0.814009151 | 3.03E-06 |
| *Itga2* | -0.825252386 | 4.57E-29 |
| *Foxq1* | -0.825718957 | 9.67E-11 |
| *Adam12* | -0.829636886 | 1.36E-25 |
| *Cd200* | -0.834432781 | 2.12E-14 |
| *Ly6d* | -0.837834385 | 4.07E-33 |
| *Serpinb8* | -0.86234898 | 5.11E-24 |
| *Adcy9* | -0.86996471 | 3.52E-28 |
| *Ovol1* | -0.872847094 | 4.38E-18 |
| *Slco2a1* | -0.875688447 | 2.53E-09 |
| *Maf* | -0.898359486 | 4.79E-30 |
| *Pdgfb* | -0.905142338 | 3.22E-12 |
| *Tgfbi* | -0.905650145 | 3.30E-51 |
| *Prl8a9* | -0.915471662 | 1.48E-81 |
| *Bpifc* | -0.95851444 | 1.88E-11 |
| *Has2* | -0.98448221 | 2.34E-12 |
| *Moxd1* | -0.986933432 | 3.63E-37 |
| *Cyp1b1* | -1.008340665 | 2.07E-18 |
| *Col5a3* | -1.011356556 | 5.20E-09 |
| *Myo7a* | -1.014096311 | 5.40E-15 |
| *RP23-121N17.2* | -1.022357923 | 2.58E-19 |
| *Slc24a3* | -1.063068649 | 2.23E-30 |
| *Smad4* | -1.094934585 | 2.65E-69 |
| *Scnn1g* | -1.142133194 | 2.50E-11 |
| *Clec2e* | -1.185569663 | 3.21E-21 |
| *Prr9* | -1.261706538 | 2.47E-17 |

| | | |
|---|---|---|
| *Ccl7* | -1.304052356 | 2.19E-21 |
| *Nckap5* | -1.351112351 | 8.80E-38 |
| *Lgr6* | -1.814441152 | 1.35E-28 |

**Supplementary Table 2. SMAD4 Peak Annotation**

| Chromsome | start | end | annotation |
|---|---|---|---|
| chr1 | 4246133 | 4246341 | Distal Intergenic |
| chr1 | 4246495 | 4249452 | Distal Intergenic |
| chr1 | 7376716 | 7376953 | Intron |
| chr1 | 7377063 | 7379112 | Intron |
| chr1 | 9627081 | 9627314 | Distal Intergenic |
| chr1 | 12738435 | 12738849 | Intron |
| chr1 | 16582814 | 16583686 | Intron |
| chr1 | 20711841 | 20712944 | Distal Intergenic |
| chr1 | 21622778 | 21623149 | Intron |
| chr1 | 23562514 | 23564486 | Distal Intergenic |
| chr1 | 25161594 | 25162301 | Intron |
| chr1 | 35713317 | 35714046 | Distal Intergenic |
| chr1 | 36080482 | 36080837 | Distal Intergenic |
| chr1 | 36255807 | 36256006 | Intron |
| chr1 | 36836819 | 36837680 | Promoter |
| chr1 | 37080804 | 37081969 | Distal Intergenic |
| chr1 | 37577567 | 37577982 | Intron |
| chr1 | 37580272 | 37581070 | Intron |
| chr1 | 38895142 | 38895393 | Distal Intergenic |
| chr1 | 39812571 | 39813000 | Distal Intergenic |
| chr1 | 39813037 | 39813493 | Distal Intergenic |
| chr1 | 41043970 | 41044536 | Distal Intergenic |
| chr1 | 42754334 | 42754639 | Promoter |
| chr1 | 42754944 | 42755159 | Promoter |
| chr1 | 42900900 | 42901299 | Distal Intergenic |
| chr1 | 43496484 | 43497171 | Distal Intergenic |
| chr1 | 46132797 | 46133000 | Intron |
| chr1 | 55245076 | 55246457 | Intron |
| chr1 | 58168824 | 58169144 | Promoter |
| chr1 | 59820305 | 59820536 | Promoter |
| chr1 | 64368128 | 64368389 | Distal Intergenic |
| chr1 | 70659010 | 70659415 | Intron |
| chr1 | 73621506 | 73621868 | Distal Intergenic |
| chr1 | 74030713 | 74030981 | Intron |
| chr1 | 75457258 | 75458178 | Intron |
| chr1 | 75458234 | 75459162 | Intron |
| chr1 | 76726575 | 76726960 | Distal Intergenic |

| | | | |
|---|---|---|---|
| chr1 | 78854856 | 78855423 | Distal Intergenic |
| chr1 | 84274080 | 84274920 | Intron |
| chr1 | 84321030 | 84321415 | Intron |
| chr1 | 92341268 | 92341589 | Distal Intergenic |
| chr1 | 92341809 | 92345349 | Distal Intergenic |
| chr1 | 98467118 | 98467317 | Distal Intergenic |
| chr1 | 102076721 | 102076928 | Intron |
| chr1 | 104524683 | 104524895 | Distal Intergenic |
| chr1 | 105554706 | 105554957 | Distal Intergenic |
| chr1 | 107212948 | 107213540 | Intron |
| chr1 | 120651971 | 120652225 | Distal Intergenic |
| chr1 | 122526154 | 122526698 | Distal Intergenic |
| chr1 | 123380209 | 123381635 | Intron |
| chr1 | 125199204 | 125199403 | Distal Intergenic |
| chr1 | 125199759 | 125200653 | Distal Intergenic |
| chr1 | 125200737 | 125200953 | Distal Intergenic |
| chr1 | 125933818 | 125934140 | Intron |
| chr1 | 126393264 | 126393662 | Distal Intergenic |
| chr1 | 127250690 | 127251086 | Distal Intergenic |
| chr1 | 131207591 | 131207838 | Intron |
| chr1 | 131207882 | 131208586 | Intron |
| chr1 | 131752989 | 131753371 | Intron |
| chr1 | 134240279 | 134240580 | Intron |
| chr1 | 135491231 | 135492493 | Distal Intergenic |
| chr1 | 137100115 | 137100314 | Distal Intergenic |
| chr1 | 137100512 | 137100738 | Distal Intergenic |
| chr1 | 138233265 | 138233557 | Intron |
| chr1 | 138234860 | 138235150 | Intron |
| chr1 | 145207298 | 145207497 | Distal Intergenic |
| chr1 | 156879427 | 156879726 | Distal Intergenic |
| chr1 | 162210056 | 162210522 | Intron |
| chr1 | 162402542 | 162403410 | Intron |
| chr1 | 166529666 | 166529920 | Distal Intergenic |
| chr1 | 168423367 | 168424247 | Intron |
| chr1 | 169270428 | 169270632 | Distal Intergenic |
| chr1 | 173769391 | 173769606 | Promoter |
| chr1 | 179371853 | 179372315 | Promoter |
| chr1 | 181816326 | 181816920 | Distal Intergenic |
| chr1 | 182280828 | 182281824 | Intron |
| chr1 | 183673106 | 183673982 | Intron |
| chr1 | 186681483 | 186681782 | Distal Intergenic |
| chr1 | 187497622 | 187498140 | Distal Intergenic |
| chr1 | 189011019 | 189011304 | Intron |
| chr1 | 189227677 | 189227888 | Distal Intergenic |

| | | | |
|---|---|---|---|
| chr1 | 189566463 | 189567055 | Intron |
| chr1 | 190391659 | 190391983 | Intron |
| chr1 | 190617002 | 190618638 | Intron |
| chr1 | 191548724 | 191549204 | Distal Intergenic |
| chr1 | 192307877 | 192308456 | Distal Intergenic |
| chr1 | 192579790 | 192580800 | Distal Intergenic |
| chr1 | 192695875 | 192696839 | Intron |
| chr1 | 192944815 | 192945031 | Distal Intergenic |
| chr1 | 194488694 | 194492753 | Intron |
| chr1 | 194493590 | 194493839 | Intron |
| chr1 | 194495838 | 194496069 | Intron |
| chr1 | 194496114 | 194496342 | Intron |
| chr1 | 194573123 | 194574080 | Intron |
| chr1 | 194574405 | 194574930 | Intron |
| chr1 | 194575287 | 194575486 | Intron |
| chr10 | 5080268 | 5080794 | Distal Intergenic |
| chr10 | 5728902 | 5729316 | Intron |
| chr10 | 8681882 | 8682137 | Distal Intergenic |
| chr10 | 11072356 | 11072867 | Intron |
| chr10 | 13094778 | 13094990 | Intron |
| chr10 | 13139988 | 13140422 | Intron |
| chr10 | 18429342 | 18429613 | Intron |
| chr10 | 19333186 | 19334249 | Distal Intergenic |
| chr10 | 20765924 | 20766285 | Intron |
| chr10 | 24660419 | 24661002 | Distal Intergenic |
| chr10 | 26582412 | 26582611 | Intron |
| chr10 | 34071017 | 34071345 | Intron |
| chr10 | 36637700 | 36638064 | Distal Intergenic |
| chr10 | 40944240 | 40946324 | Intron |
| chr10 | 60427304 | 60427551 | Distal Intergenic |
| chr10 | 60836629 | 60836837 | Intron |
| chr10 | 61051718 | 61052197 | Distal Intergenic |
| chr10 | 64088273 | 64088687 | Intron |
| chr10 | 64124592 | 64124885 | Intron |
| chr10 | 67166735 | 67167028 | Distal Intergenic |
| chr10 | 69686126 | 69686445 | Distal Intergenic |
| chr10 | 72634432 | 72634728 | Intron |
| chr10 | 77071641 | 77071931 | Intron |
| chr10 | 79819758 | 79820201 | Promoter |
| chr10 | 79823324 | 79826052 | Exon |
| chr10 | 79826842 | 79827323 | Exon |
| chr10 | 80869511 | 80869771 | Intron |
| chr10 | 82574181 | 82574520 | Intron |
| chr10 | 83662255 | 83662761 | Distal Intergenic |

| | | | |
|---|---|---|---|
| chr10 | 83860932 | 83861294 | Intron |
| chr10 | 86930534 | 86930778 | Distal Intergenic |
| chr10 | 86962070 | 86962593 | Distal Intergenic |
| chr10 | 94036250 | 94036950 | Intron |
| chr10 | 94458355 | 94458659 | Distal Intergenic |
| chr10 | 97009518 | 97010562 | Distal Intergenic |
| chr10 | 98632851 | 98634226 | Intron |
| chr10 | 100400128 | 100400605 | Distal Intergenic |
| chr10 | 104403035 | 104403457 | Distal Intergenic |
| chr10 | 106840473 | 106840938 | Intron |
| chr10 | 106874701 | 106875097 | Distal Intergenic |
| chr10 | 109625439 | 109626210 | Distal Intergenic |
| chr10 | 114931306 | 114931853 | Intron |
| chr10 | 115555961 | 115556754 | Intron |
| chr10 | 115749042 | 115749422 | Intron |
| chr10 | 116190104 | 116190978 | Distal Intergenic |
| chr10 | 116226454 | 116226814 | Intron |
| chr10 | 116314917 | 116317015 | Intron |
| chr10 | 118388695 | 118389650 | Distal Intergenic |
| chr10 | 118389686 | 118389885 | Distal Intergenic |
| chr10 | 118591740 | 118592400 | Distal Intergenic |
| chr10 | 120437370 | 120437716 | Distal Intergenic |
| chr10 | 122484243 | 122485253 | Intron |
| chr10 | 125343542 | 125343940 | Distal Intergenic |
| chr10 | 125356306 | 125356732 | Distal Intergenic |
| chr10 | 125532080 | 125532487 | Distal Intergenic |
| chr10 | 125599704 | 125600002 | Distal Intergenic |
| chr11 | 3046896 | 3047101 | Intron |
| chr11 | 3474829 | 3475066 | Distal Intergenic |
| chr11 | 4921138 | 4921458 | Intron |
| chr11 | 6565255 | 6565454 | Intron |
| chr11 | 7005969 | 7006432 | Intron |
| chr11 | 7399153 | 7399742 | Distal Intergenic |
| chr11 | 19489431 | 19490749 | Distal Intergenic |
| chr11 | 24169587 | 24170262 | Distal Intergenic |
| chr11 | 24170605 | 24170849 | Distal Intergenic |
| chr11 | 26083656 | 26083867 | Intron |
| chr11 | 31972482 | 31972811 | Distal Intergenic |
| chr11 | 32448513 | 32448891 | Intron |
| chr11 | 35817936 | 35818555 | Downstream |
| chr11 | 35842691 | 35842936 | Intron |
| chr11 | 44105603 | 44107644 | Distal Intergenic |
| chr11 | 45330842 | 45331716 | Distal Intergenic |
| chr11 | 50232832 | 50233054 | Intron |

| | | | |
|---|---|---|---|
| chr11 | 50647845 | 50648061 | Intron |
| chr11 | 52527238 | 52527870 | Distal Intergenic |
| chr11 | 53067029 | 53067622 | Distal Intergenic |
| chr11 | 57285216 | 57286531 | Distal Intergenic |
| chr11 | 58207212 | 58207499 | Distal Intergenic |
| chr11 | 68296230 | 68296651 | Intron |
| chr11 | 70423196 | 70423530 | Exon |
| chr11 | 77463247 | 77463912 | Distal Intergenic |
| chr11 | 78945117 | 78945459 | Intron |
| chr11 | 79695739 | 79696157 | Distal Intergenic |
| chr11 | 83368404 | 83369841 | Distal Intergenic |
| chr11 | 86201935 | 86202134 | Distal Intergenic |
| chr11 | 86287458 | 86287784 | Distal Intergenic |
| chr11 | 88942905 | 88943418 | Intron |
| chr11 | 91439251 | 91439677 | Promoter |
| chr11 | 97810679 | 97810938 | Intron |
| chr11 | 99122950 | 99123857 | Distal Intergenic |
| chr11 | 99268939 | 99269222 | Distal Intergenic |
| chr11 | 102875060 | 102875983 | Intron |
| chr11 | 104106408 | 104106664 | Intron |
| chr11 | 104888866 | 104889190 | Distal Intergenic |
| chr11 | 104948880 | 104949316 | Intron |
| chr11 | 107519612 | 107519818 | Intron |
| chr11 | 108123563 | 108123852 | Intron |
| chr11 | 108124061 | 108124260 | Intron |
| chr11 | 108872983 | 108873329 | Distal Intergenic |
| chr11 | 109847216 | 109847416 | Intron |
| chr11 | 109847795 | 109848355 | Intron |
| chr11 | 110314851 | 110315334 | Intron |
| chr11 | 111742343 | 111743410 | Distal Intergenic |
| chr11 | 111743451 | 111743679 | Distal Intergenic |
| chr11 | 113741149 | 113741685 | Intron |
| chr11 | 114253817 | 114254063 | Distal Intergenic |
| chr11 | 118052364 | 118052864 | Intron |
| chr11 | 118796844 | 118797263 | Distal Intergenic |
| chr11 | 119177134 | 119177862 | Intron |
| chr11 | 119489900 | 119490360 | Intron |
| chr11 | 119720472 | 119720980 | Intron |
| chr12 | 3320886 | 3321515 | Distal Intergenic |
| chr12 | 4110845 | 4112322 | Distal Intergenic |
| chr12 | 4292179 | 4293011 | Intron |
| chr12 | 4536822 | 4538116 | Distal Intergenic |
| chr12 | 12722791 | 12722990 | Distal Intergenic |
| chr12 | 12724455 | 12724660 | Distal Intergenic |

| | | | |
|---|---|---|---|
| chr12 | 14665985 | 14666879 | Distal Intergenic |
| chr12 | 18136385 | 18136690 | Distal Intergenic |
| chr12 | 18140466 | 18140984 | Distal Intergenic |
| chr12 | 19178730 | 19179638 | Distal Intergenic |
| chr12 | 25173052 | 25173699 | Distal Intergenic |
| chr12 | 35578986 | 35579400 | Intron |
| chr12 | 54027695 | 54033510 | Intron |
| chr12 | 57163120 | 57163408 | Distal Intergenic |
| chr12 | 63278293 | 63278503 | Distal Intergenic |
| chr12 | 70755068 | 70755323 | Intron |
| chr12 | 78269503 | 78270270 | Distal Intergenic |
| chr12 | 79451900 | 79452115 | Intron |
| chr12 | 84297056 | 84297595 | Distal Intergenic |
| chr12 | 89420941 | 89421195 | Promoter |
| chr12 | 92493430 | 92493629 | Intron |
| chr12 | 92740705 | 92741259 | Intron |
| chr12 | 100883366 | 100883585 | Distal Intergenic |
| chr12 | 104542378 | 104542577 | Distal Intergenic |
| chr12 | 107197512 | 107197959 | Distal Intergenic |
| chr12 | 109284211 | 109285130 | Distal Intergenic |
| chr12 | 109285173 | 109285394 | Distal Intergenic |
| chr12 | 110032024 | 110032308 | Promoter |
| chr12 | 111095284 | 111095549 | Distal Intergenic |
| chr12 | 111257645 | 111257844 | Intron |
| chr12 | 111484198 | 111484488 | Intron |
| chr12 | 114174409 | 114174753 | Intron |
| chr12 | 114175010 | 114175509 | Intron |
| chr12 | 114499767 | 114500091 | Intron |
| chr13 | 3963446 | 3964770 | Distal Intergenic |
| chr13 | 5346202 | 5346781 | Distal Intergenic |
| chr13 | 9833540 | 9833758 | Distal Intergenic |
| chr13 | 12051846 | 12052592 | Intron |
| chr13 | 14649864 | 14650185 | Distal Intergenic |
| chr13 | 27863546 | 27863818 | Downstream |
| chr13 | 35130109 | 35131008 | Distal Intergenic |
| chr13 | 37726846 | 37727418 | Distal Intergenic |
| chr13 | 38429731 | 38430960 | Distal Intergenic |
| chr13 | 40765804 | 40766012 | Distal Intergenic |
| chr13 | 42190048 | 42190670 | Intron |
| chr13 | 44964953 | 44965380 | Intron |
| chr13 | 45885192 | 45885695 | Intron |
| chr13 | 48370330 | 48371420 | Distal Intergenic |
| chr13 | 48529272 | 48529485 | Distal Intergenic |
| chr13 | 52752597 | 52752804 | Distal Intergenic |

| | | | |
|---|---|---|---|
| chr13 | 53832617 | 53833199 | Distal Intergenic |
| chr13 | 53837139 | 53838423 | Distal Intergenic |
| chr13 | 56332087 | 56332548 | Distal Intergenic |
| chr13 | 58080778 | 58081019 | Distal Intergenic |
| chr13 | 60198250 | 60198948 | Distal Intergenic |
| chr13 | 64596590 | 64597334 | Intron |
| chr13 | 69673304 | 69673586 | Promoter |
| chr13 | 70631358 | 70632479 | Distal Intergenic |
| chr13 | 72931753 | 72932320 | Distal Intergenic |
| chr13 | 74665814 | 74668431 | Distal Intergenic |
| chr13 | 75353129 | 75353845 | Distal Intergenic |
| chr13 | 83181027 | 83181390 | Distal Intergenic |
| chr13 | 90516622 | 90517613 | Distal Intergenic |
| chr13 | 93699488 | 93699691 | Distal Intergenic |
| chr13 | 97960480 | 97960700 | Intron |
| chr13 | 100560885 | 100561150 | Distal Intergenic |
| chr13 | 105012564 | 105012778 | Intron |
| chr13 | 108679396 | 108679662 | Promoter |
| chr13 | 109503753 | 109504285 | Intron |
| chr13 | 111435405 | 111436395 | Distal Intergenic |
| chr13 | 112888463 | 112889055 | Distal Intergenic |
| chr13 | 113368277 | 113368612 | Intron |
| chr13 | 114521938 | 114522311 | Distal Intergenic |
| chr14 | 19877142 | 19877346 | Distal Intergenic |
| chr14 | 19877460 | 19877821 | Distal Intergenic |
| chr14 | 19877993 | 19878290 | Distal Intergenic |
| chr14 | 19878383 | 19878981 | Distal Intergenic |
| chr14 | 19879307 | 19879925 | Distal Intergenic |
| chr14 | 19880657 | 19880887 | Distal Intergenic |
| chr14 | 19880993 | 19881580 | Distal Intergenic |
| chr14 | 19881888 | 19884248 | Distal Intergenic |
| chr14 | 19884367 | 19889384 | Distal Intergenic |
| chr14 | 19889527 | 19890402 | Distal Intergenic |
| chr14 | 19890524 | 19891693 | Distal Intergenic |
| chr14 | 23764573 | 23764784 | Intron |
| chr14 | 25836134 | 25836334 | Distal Intergenic |
| chr14 | 27700796 | 27701034 | Intron |
| chr14 | 30664481 | 30664692 | Distal Intergenic |
| chr14 | 47492729 | 47493114 | Distal Intergenic |
| chr14 | 51795014 | 51795311 | Distal Intergenic |
| chr14 | 51795441 | 51795644 | Distal Intergenic |
| chr14 | 51886686 | 51886979 | Distal Intergenic |
| chr14 | 57312720 | 57312964 | Intron |
| chr14 | 65109695 | 65110324 | Distal Intergenic |

| | | | |
|---|---|---|---|
| chr14 | 70331913 | 70332146 | Intron |
| chr14 | 72748463 | 72748759 | Distal Intergenic |
| chr14 | 75789466 | 75789725 | Distal Intergenic |
| chr14 | 75789811 | 75790010 | Distal Intergenic |
| chr14 | 86921244 | 86922607 | Distal Intergenic |
| chr14 | 88292341 | 88292602 | Distal Intergenic |
| chr14 | 99866950 | 99867627 | Distal Intergenic |
| chr14 | 103863347 | 103864010 | Intron |
| chr14 | 105114979 | 105115247 | Intron |
| chr14 | 115607250 | 115607663 | Intron |
| chr14 | 118869324 | 118869749 | Distal Intergenic |
| chr15 | 5711120 | 5711356 | Distal Intergenic |
| chr15 | 6477872 | 6478628 | Distal Intergenic |
| chr15 | 6478793 | 6479465 | Distal Intergenic |
| chr15 | 6479503 | 6479712 | Distal Intergenic |
| chr15 | 6479836 | 6480962 | Distal Intergenic |
| chr15 | 6481078 | 6481777 | Distal Intergenic |
| chr15 | 6481843 | 6482159 | Distal Intergenic |
| chr15 | 6482195 | 6482876 | Distal Intergenic |
| chr15 | 6482934 | 6483617 | Distal Intergenic |
| chr15 | 6822713 | 6823231 | Intron |
| chr15 | 7256989 | 7257208 | Intron |
| chr15 | 7693210 | 7693455 | Distal Intergenic |
| chr15 | 10509587 | 10510060 | Intron |
| chr15 | 10510301 | 10511760 | Intron |
| chr15 | 13727492 | 13727902 | Distal Intergenic |
| chr15 | 13728054 | 13728266 | Distal Intergenic |
| chr15 | 13728566 | 13728814 | Distal Intergenic |
| chr15 | 20574659 | 20575512 | Intron |
| chr15 | 31017414 | 31017684 | Distal Intergenic |
| chr15 | 38806129 | 38806374 | Distal Intergenic |
| chr15 | 38806430 | 38806787 | Distal Intergenic |
| chr15 | 39581196 | 39581556 | Intron |
| chr15 | 47371465 | 47371664 | Distal Intergenic |
| chr15 | 55633031 | 55633258 | Intron |
| chr15 | 64870510 | 64870791 | Distal Intergenic |
| chr15 | 66135297 | 66135927 | Distal Intergenic |
| chr15 | 72352984 | 72354800 | Intron |
| chr15 | 74170304 | 74170508 | Distal Intergenic |
| chr15 | 74170682 | 74171273 | Distal Intergenic |
| chr15 | 74916995 | 74917372 | Distal Intergenic |
| chr15 | 75563641 | 75564374 | Distal Intergenic |
| chr15 | 76831457 | 76833693 | Distal Intergenic |
| chr15 | 83567590 | 83569670 | Distal Intergenic |

| | | | |
|---|---|---|---|
| chr15 | 83572559 | 83573050 | Distal Intergenic |
| chr15 | 83867791 | 83871472 | Intron |
| chr15 | 86025212 | 86025681 | Distal Intergenic |
| chr15 | 86234248 | 86234463 | Intron |
| chr15 | 86253199 | 86253527 | Intron |
| chr15 | 86583395 | 86584563 | Distal Intergenic |
| chr15 | 89919550 | 89920064 | Distal Intergenic |
| chr15 | 93842848 | 93843884 | Distal Intergenic |
| chr15 | 96548526 | 96548807 | Distal Intergenic |
| chr15 | 98220757 | 98220962 | Distal Intergenic |
| chr15 | 98330941 | 98332140 | Distal Intergenic |
| chr15 | 100584942 | 100585255 | Distal Intergenic |
| chr15 | 100697216 | 100698126 | Distal Intergenic |
| chr15 | 101299687 | 101300432 | Intron |
| chr15 | 101314562 | 101315114 | Intron |
| chr16 | 5246396 | 5247075 | Intron |
| chr16 | 9464191 | 9464450 | Distal Intergenic |
| chr16 | 9464791 | 9466789 | Distal Intergenic |
| chr16 | 11144018 | 11144404 | Intron |
| chr16 | 11656258 | 11657419 | Exon |
| chr16 | 17222381 | 17222588 | Distal Intergenic |
| chr16 | 17560388 | 17560676 | Promoter |
| chr16 | 17972863 | 17973330 | Distal Intergenic |
| chr16 | 18225354 | 18225557 | Intron |
| chr16 | 18533189 | 18533514 | Intron |
| chr16 | 21487698 | 21487949 | Intron |
| chr16 | 22830876 | 22831114 | Intron |
| chr16 | 24351437 | 24352248 | Distal Intergenic |
| chr16 | 26496733 | 26499781 | Distal Intergenic |
| chr16 | 26992935 | 26994310 | Distal Intergenic |
| chr16 | 30717547 | 30718052 | Distal Intergenic |
| chr16 | 30718116 | 30718324 | Distal Intergenic |
| chr16 | 30718573 | 30718899 | Distal Intergenic |
| chr16 | 31026165 | 31027170 | Intron |
| chr16 | 31162392 | 31163867 | Intron |
| chr16 | 32211426 | 32212537 | Distal Intergenic |
| chr16 | 33018547 | 33018806 | Intron |
| chr16 | 35580733 | 35581297 | Intron |
| chr16 | 35981705 | 35981992 | Exon |
| chr16 | 45352533 | 45353413 | Distal Intergenic |
| chr16 | 45840381 | 45840580 | Intron |
| chr16 | 55852037 | 55852642 | Intron |
| chr16 | 55854447 | 55855413 | Intron |
| chr16 | 56230603 | 56230852 | Intron |

| | | | |
|---|---|---|---|
| chr16 | 57256410 | 57256811 | Distal Intergenic |
| chr16 | 57391290 | 57391809 | Intron |
| chr16 | 60947345 | 60948340 | Distal Intergenic |
| chr16 | 62833022 | 62833299 | Intron |
| chr16 | 64851188 | 64851397 | Promoter |
| chr16 | 64851493 | 64851766 | Promoter |
| chr16 | 64852130 | 64852366 | Promoter |
| chr16 | 65034288 | 65035426 | Intron |
| chr16 | 72644429 | 72645027 | Distal Intergenic |
| chr16 | 78252723 | 78253506 | Intron |
| chr16 | 78660552 | 78660761 | Distal Intergenic |
| chr16 | 78660851 | 78661218 | Distal Intergenic |
| chr16 | 80491400 | 80491793 | Distal Intergenic |
| chr16 | 86000514 | 86001505 | Distal Intergenic |
| chr16 | 86263636 | 86263835 | Distal Intergenic |
| chr16 | 86820998 | 86821470 | Distal Intergenic |
| chr16 | 87165374 | 87165730 | Distal Intergenic |
| chr16 | 87543664 | 87544692 | Distal Intergenic |
| chr16 | 88762870 | 88763201 | Distal Intergenic |
| chr16 | 90178068 | 90178268 | Distal Intergenic |
| chr16 | 90913629 | 90914109 | Distal Intergenic |
| chr16 | 91438178 | 91439404 | Distal Intergenic |
| chr16 | 91439455 | 91439685 | Distal Intergenic |
| chr16 | 92250683 | 92251792 | Distal Intergenic |
| chr16 | 92323332 | 92324604 | Distal Intergenic |
| chr16 | 92655503 | 92655928 | Intron |
| chr16 | 92807057 | 92807344 | Intron |
| chr16 | 92954834 | 92955776 | Distal Intergenic |
| chr16 | 93110075 | 93110316 | Distal Intergenic |
| chr16 | 93772185 | 93772960 | Intron |
| chr16 | 93773171 | 93773396 | Intron |
| chr16 | 93913334 | 93913699 | Distal Intergenic |
| chr16 | 94017187 | 94017425 | Distal Intergenic |
| chr16 | 94953822 | 94954079 | Distal Intergenic |
| chr16 | 94959817 | 94960038 | Distal Intergenic |
| chr16 | 94960101 | 94960473 | Distal Intergenic |
| chr16 | 95959364 | 95960010 | Distal Intergenic |
| chr16 | 96676254 | 96676633 | Intron |
| chr16 | 97124847 | 97125199 | Intron |
| chr17 | 3059596 | 3060111 | Distal Intergenic |
| chr17 | 3060166 | 3060862 | Distal Intergenic |
| chr17 | 3061122 | 3061321 | Downstream |
| chr17 | 3061650 | 3062141 | Downstream |
| chr17 | 3070003 | 3070206 | Intron |

| | | | |
|---|---|---|---|
| chr17 | 3070344 | 3073873 | Intron |
| chr17 | 3074004 | 3074594 | Intron |
| chr17 | 5241033 | 5241232 | Intron |
| chr17 | 5863896 | 5867255 | Intron |
| chr17 | 5879519 | 5884978 | Intron |
| chr17 | 10139933 | 10140132 | Distal Intergenic |
| chr17 | 10918674 | 10919179 | Intron |
| chr17 | 12482261 | 12483400 | Intron |
| chr17 | 12829073 | 12829378 | Distal Intergenic |
| chr17 | 13498714 | 13499273 | Intron |
| chr17 | 13743462 | 13746166 | Intron |
| chr17 | 16047782 | 16047981 | Distal Intergenic |
| chr17 | 16154152 | 16154415 | Distal Intergenic |
| chr17 | 24132294 | 24132493 | Intron |
| chr17 | 24479619 | 24480080 | Intron |
| chr17 | 25484556 | 25485442 | Distal Intergenic |
| chr17 | 27573430 | 27573733 | Intron |
| chr17 | 28405166 | 28405389 | Intron |
| chr17 | 28495634 | 28495962 | Promoter |
| chr17 | 30220335 | 30220726 | Intron |
| chr17 | 30822013 | 30822421 | Intron |
| chr17 | 36368215 | 36368524 | Distal Intergenic |
| chr17 | 38087232 | 38087442 | Downstream |
| chr17 | 39980003 | 39985754 | Promoter |
| chr17 | 43440990 | 43441408 | Intron |
| chr17 | 46237065 | 46237916 | Distal Intergenic |
| chr17 | 46472223 | 46472426 | Distal Intergenic |
| chr17 | 46616337 | 46616590 | Intron |
| chr17 | 50056138 | 50056567 | Distal Intergenic |
| chr17 | 50060572 | 50060857 | Distal Intergenic |
| chr17 | 50064345 | 50064913 | Distal Intergenic |
| chr17 | 50064966 | 50066171 | Distal Intergenic |
| chr17 | 50066212 | 50066713 | Distal Intergenic |
| chr17 | 50484979 | 50485424 | Distal Intergenic |
| chr17 | 50668259 | 50669375 | Intron |
| chr17 | 50669431 | 50669658 | Intron |
| chr17 | 56794503 | 56794788 | Exon |
| chr17 | 63848621 | 63849088 | Promoter |
| chr17 | 64184199 | 64184867 | Distal Intergenic |
| chr17 | 67574888 | 67575089 | Intron |
| chr17 | 68939805 | 68940812 | Intron |
| chr17 | 69905748 | 69905966 | Distal Intergenic |
| chr17 | 71643413 | 71643730 | Intron |
| chr17 | 73703619 | 73703931 | Intron |

| | | | |
|---|---|---|---|
| chr17 | 75522774 | 75523499 | Intron |
| chr17 | 81226807 | 81227226 | Distal Intergenic |
| chr17 | 85346380 | 85346579 | Intron |
| chr17 | 87296656 | 87296873 | Distal Intergenic |
| chr17 | 87411767 | 87413651 | Intron |
| chr17 | 87696449 | 87696648 | Intron |
| chr17 | 87899638 | 87899841 | Distal Intergenic |
| chr17 | 87899913 | 87900177 | Distal Intergenic |
| chr17 | 88701590 | 88701933 | Intron |
| chr17 | 89148283 | 89148898 | Intron |
| chr17_random | 323668 | 326663 | Distal Intergenic |
| chr17_random | 339011 | 339978 | Distal Intergenic |
| chr18 | 3661334 | 3661685 | Distal Intergenic |
| chr18 | 4502923 | 4504993 | Distal Intergenic |
| chr18 | 5788138 | 5790995 | Distal Intergenic |
| chr18 | 6621020 | 6621904 | Intron |
| chr18 | 6787571 | 6787963 | Intron |
| chr18 | 7608412 | 7609737 | Promoter |
| chr18 | 10548269 | 10548604 | Intron |
| chr18 | 11636868 | 11637785 | Distal Intergenic |
| chr18 | 12050742 | 12051790 | Intron |
| chr18 | 12878602 | 12879073 | Intron |
| chr18 | 14080236 | 14080892 | Intron |
| chr18 | 20747611 | 20748293 | Intron |
| chr18 | 28243445 | 28243757 | Distal Intergenic |
| chr18 | 30588483 | 30589029 | Distal Intergenic |
| chr18 | 37942419 | 37942871 | Intron |
| chr18 | 41301131 | 41301462 | Distal Intergenic |
| chr18 | 42744421 | 42745143 | Distal Intergenic |
| chr18 | 47578003 | 47578318 | Distal Intergenic |
| chr18 | 54858619 | 54858967 | Distal Intergenic |
| chr18 | 55025645 | 55025881 | Distal Intergenic |
| chr18 | 60798392 | 60798650 | Distal Intergenic |
| chr18 | 62468618 | 62469266 | Distal Intergenic |
| chr18 | 67286364 | 67286658 | Intron |
| chr18 | 68132506 | 68132732 | Intron |
| chr18 | 68851650 | 68851947 | Distal Intergenic |
| chr18 | 72066685 | 72067163 | Intron |
| chr18 | 76522184 | 76522947 | Distal Intergenic |
| chr18 | 76840394 | 76841569 | Distal Intergenic |
| chr18 | 76843440 | 76843642 | Distal Intergenic |
| chr18 | 77769574 | 77769796 | Intron |
| chr18 | 80529555 | 80530239 | Intron |
| chr18 | 81361081 | 81361435 | Distal Intergenic |

| | | | |
|---|---|---|---|
| chr18 | 81600792 | 81601341 | Distal Intergenic |
| chr18 | 81865113 | 81866078 | Distal Intergenic |
| chr18 | 82137905 | 82138104 | Distal Intergenic |
| chr18 | 82841434 | 82841940 | Intron |
| chr18 | 83710464 | 83711748 | Distal Intergenic |
| chr18 | 85867357 | 85867646 | Distal Intergenic |
| chr19 | 4306479 | 4306708 | Promoter |
| chr19 | 5241429 | 5241892 | Intron |
| chr19 | 6558628 | 6558918 | Distal Intergenic |
| chr19 | 10582454 | 10584474 | Intron |
| chr19 | 10746685 | 10747876 | Intron |
| chr19 | 11874599 | 11875330 | Intron |
| chr19 | 12091591 | 12091938 | Distal Intergenic |
| chr19 | 17119948 | 17120337 | Intron |
| chr19 | 24932108 | 24932927 | Intron |
| chr19 | 25780817 | 25781691 | Distal Intergenic |
| chr19 | 29291228 | 29291498 | Distal Intergenic |
| chr19 | 29395024 | 29395286 | Downstream |
| chr19 | 29552191 | 29552395 | Intron |
| chr19 | 35741006 | 35741205 | Distal Intergenic |
| chr19 | 35963523 | 35963740 | Intron |
| chr19 | 38784777 | 38785223 | Intron |
| chr19 | 39064787 | 39066240 | Distal Intergenic |
| chr19 | 42339832 | 42340400 | Intron |
| chr19 | 42773453 | 42773814 | Distal Intergenic |
| chr19 | 42773937 | 42774439 | Distal Intergenic |
| chr19 | 46095432 | 46095950 | Distal Intergenic |
| chr19 | 47282939 | 47283216 | Intron |
| chr19 | 47283274 | 47283854 | Intron |
| chr19 | 47708405 | 47709324 | Intron |
| chr19 | 48125981 | 48126323 | Distal Intergenic |
| chr19 | 54093203 | 54094089 | Intron |
| chr19 | 56294289 | 56294492 | Distal Intergenic |
| chr19 | 56922084 | 56922791 | Intron |
| chr19 | 58004305 | 58004665 | Intron |
| chr19 | 58358014 | 58358694 | Intron |
| chr19 | 61340160 | 61340422 | Distal Intergenic |
| chr1_random | 1033047 | 1033329 | Intron |
| chr2 | 3069508 | 3069935 | Intron |
| chr2 | 3881799 | 3885591 | Distal Intergenic |
| chr2 | 4442775 | 4443540 | Intron |
| chr2 | 5004351 | 5004761 | Distal Intergenic |
| chr2 | 5300122 | 5300422 | Intron |
| chr2 | 6117991 | 6118287 | Intron |

| | | | |
|---|---|---|---|
| chr2 | 8631081 | 8631312 | Distal Intergenic |
| chr2 | 9062161 | 9062527 | Distal Intergenic |
| chr2 | 9719135 | 9719859 | Distal Intergenic |
| chr2 | 20821481 | 20821788 | Intron |
| chr2 | 22600369 | 22600625 | Distal Intergenic |
| chr2 | 24674169 | 24674403 | Intron |
| chr2 | 25952075 | 25952600 | Intron |
| chr2 | 26919346 | 26919545 | Intron |
| chr2 | 26919585 | 26921107 | Intron |
| chr2 | 28040518 | 28041001 | Distal Intergenic |
| chr2 | 28227484 | 28227700 | Distal Intergenic |
| chr2 | 28262860 | 28263069 | Distal Intergenic |
| chr2 | 28458107 | 28458909 | Distal Intergenic |
| chr2 | 29071003 | 29072248 | Intron |
| chr2 | 29072462 | 29072853 | Intron |
| chr2 | 29427098 | 29427297 | Distal Intergenic |
| chr2 | 29449039 | 29449872 | Distal Intergenic |
| chr2 | 31221247 | 31221490 | Distal Intergenic |
| chr2 | 38510765 | 38511967 | Distal Intergenic |
| chr2 | 44245727 | 44246365 | Distal Intergenic |
| chr2 | 44246772 | 44247020 | Distal Intergenic |
| chr2 | 45275575 | 45276879 | Distal Intergenic |
| chr2 | 45785701 | 45786593 | Distal Intergenic |
| chr2 | 49463287 | 49464202 | Distal Intergenic |
| chr2 | 49651010 | 49652251 | Intron |
| chr2 | 50214498 | 50214912 | Intron |
| chr2 | 52887913 | 52888228 | Intron |
| chr2 | 53868242 | 53868700 | Distal Intergenic |
| chr2 | 54567653 | 54568051 | Intron |
| chr2 | 58278367 | 58278885 | Distal Intergenic |
| chr2 | 59200562 | 59200761 | Distal Intergenic |
| chr2 | 59200840 | 59201091 | Distal Intergenic |
| chr2 | 66250049 | 66250462 | Intron |
| chr2 | 69211671 | 69213156 | Distal Intergenic |
| chr2 | 70244333 | 70245113 | Intron |
| chr2 | 76876932 | 76877889 | Exon |
| chr2 | 79019472 | 79021361 | Distal Intergenic |
| chr2 | 93367857 | 93368553 | Distal Intergenic |
| chr2 | 94396476 | 94398141 | Distal Intergenic |
| chr2 | 98506877 | 98507304 | Distal Intergenic |
| chr2 | 102743138 | 102744257 | Distal Intergenic |
| chr2 | 102979719 | 102980534 | Distal Intergenic |
| chr2 | 105015001 | 105015279 | Distal Intergenic |
| chr2 | 105098010 | 105098870 | Intron |

| | | | |
|---|---|---|---|
| chr2 | 105098909 | 105099201 | Intron |
| chr2 | 112758052 | 112758313 | Intron |
| chr2 | 117032975 | 117033208 | Distal Intergenic |
| chr2 | 122243566 | 122243974 | Distal Intergenic |
| chr2 | 124683455 | 124683679 | Distal Intergenic |
| chr2 | 124687603 | 124687946 | Distal Intergenic |
| chr2 | 126902864 | 126903279 | Intron |
| chr2 | 127280553 | 127281057 | Downstream |
| chr2 | 127339651 | 127339850 | Intron |
| chr2 | 127489324 | 127490600 | Distal Intergenic |
| chr2 | 128737746 | 128738048 | Distal Intergenic |
| chr2 | 131567889 | 131568342 | Distal Intergenic |
| chr2 | 131990290 | 131990588 | Distal Intergenic |
| chr2 | 143629794 | 143630172 | Intron |
| chr2 | 143819795 | 143820425 | Intron |
| chr2 | 148174513 | 148175021 | Distal Intergenic |
| chr2 | 150725303 | 150725883 | Intron |
| chr2 | 152849118 | 152849441 | Distal Intergenic |
| chr2 | 152919844 | 152922149 | Distal Intergenic |
| chr2 | 152922196 | 152926160 | Distal Intergenic |
| chr2 | 153661562 | 153661761 | Distal Intergenic |
| chr2 | 155205045 | 155205285 | Distal Intergenic |
| chr2 | 155477984 | 155478455 | Intron |
| chr2 | 158045246 | 158045554 | Intron |
| chr2 | 158057563 | 158057961 | Distal Intergenic |
| chr2 | 160366766 | 160367775 | Distal Intergenic |
| chr2 | 160430996 | 160432519 | Distal Intergenic |
| chr2 | 161387561 | 161388698 | Intron |
| chr2 | 164489183 | 164489709 | Intron |
| chr2 | 164530283 | 164531113 | Distal Intergenic |
| chr2 | 167115352 | 167116465 | Distal Intergenic |
| chr2 | 167514522 | 167514772 | Promoter |
| chr2 | 167744418 | 167745014 | Distal Intergenic |
| chr2 | 167745144 | 167745546 | Distal Intergenic |
| chr2 | 167745594 | 167745856 | Distal Intergenic |
| chr2 | 168187324 | 168187773 | Distal Intergenic |
| chr2 | 168235880 | 168236283 | Distal Intergenic |
| chr2 | 168422532 | 168422959 | Intron |
| chr2 | 169016116 | 169016356 | Distal Intergenic |
| chr2 | 169248086 | 169248306 | Distal Intergenic |
| chr2 | 169392388 | 169393476 | Distal Intergenic |
| chr2 | 170584588 | 170584865 | Intron |
| chr2 | 171640211 | 171641473 | Distal Intergenic |
| chr2 | 171724689 | 171725075 | Distal Intergenic |

| | | | |
|---|---|---|---|
| chr2 | 172027442 | 172027664 | Distal Intergenic |
| chr2 | 172153263 | 172153520 | Distal Intergenic |
| chr2 | 172420452 | 172421232 | Distal Intergenic |
| chr2 | 172421470 | 172421669 | Distal Intergenic |
| chr2 | 172578925 | 172579204 | Distal Intergenic |
| chr2 | 172579277 | 172580130 | Distal Intergenic |
| chr2 | 172580172 | 172580372 | Distal Intergenic |
| chr2 | 172626719 | 172626930 | Distal Intergenic |
| chr2 | 172627799 | 172628498 | Distal Intergenic |
| chr2 | 172924080 | 172926596 | Intron |
| chr2 | 172953115 | 172953728 | Intron |
| chr2 | 172953766 | 172954113 | Intron |
| chr2 | 173083440 | 173083639 | Intron |
| chr2 | 173100634 | 173100868 | Intron |
| chr2 | 173100904 | 173101227 | Promoter |
| chr2 | 173219539 | 173219816 | Distal Intergenic |
| chr2 | 178497690 | 178498308 | Distal Intergenic |
| chr2 | 178737495 | 178738428 | Distal Intergenic |
| chr2 | 179642303 | 179642608 | Distal Intergenic |
| chr2 | 179711462 | 179711898 | Promoter |
| chr2 | 181665342 | 181665675 | Distal Intergenic |
| chr3 | 5860388 | 5860796 | Distal Intergenic |
| chr3 | 9501623 | 9502137 | Intron |
| chr3 | 14109889 | 14110307 | Intron |
| chr3 | 26623908 | 26624600 | Intron |
| chr3 | 27278494 | 27278858 | Distal Intergenic |
| chr3 | 30977928 | 30978548 | Distal Intergenic |
| chr3 | 30993239 | 30994402 | Promoter |
| chr3 | 30994474 | 30994681 | Promoter |
| chr3 | 31110553 | 31111125 | Intron |
| chr3 | 31585143 | 31585342 | Distal Intergenic |
| chr3 | 48189233 | 48189956 | Distal Intergenic |
| chr3 | 48190034 | 48190494 | Distal Intergenic |
| chr3 | 50617673 | 50618155 | Distal Intergenic |
| chr3 | 51574866 | 51575454 | Intron |
| chr3 | 53305404 | 53306294 | Intron |
| chr3 | 53324589 | 53324791 | Intron |
| chr3 | 53487107 | 53487394 | Distal Intergenic |
| chr3 | 54400245 | 54400722 | Distal Intergenic |
| chr3 | 58055630 | 58057394 | Distal Intergenic |
| chr3 | 58278156 | 58279062 | Distal Intergenic |
| chr3 | 59848977 | 59849965 | Distal Intergenic |
| chr3 | 60539979 | 60540287 | Distal Intergenic |
| chr3 | 61963071 | 61963311 | Distal Intergenic |

| | | | |
|---|---|---|---|
| chr3 | 63975878 | 63976094 | Distal Intergenic |
| chr3 | 64218195 | 64218546 | Distal Intergenic |
| chr3 | 66497642 | 66497937 | Distal Intergenic |
| chr3 | 68550260 | 68550573 | Distal Intergenic |
| chr3 | 69819042 | 69819403 | Intron |
| chr3 | 87554297 | 87554892 | Intron |
| chr3 | 93008659 | 93008932 | Intron |
| chr3 | 93017243 | 93017749 | Exon |
| chr3 | 93018250 | 93018475 | Exon |
| chr3 | 96042975 | 96043386 | Promoter |
| chr3 | 97266093 | 97266609 | Distal Intergenic |
| chr3 | 97755300 | 97756113 | Intron |
| chr3 | 98967165 | 98967442 | Intron |
| chr3 | 108349429 | 108349952 | Distal Intergenic |
| chr3 | 116762269 | 116762604 | Distal Intergenic |
| chr3 | 120074747 | 120074963 | Distal Intergenic |
| chr3 | 120885377 | 120885797 | Distal Intergenic |
| chr3 | 120952719 | 120952930 | Intron |
| chr3 | 132868307 | 132868596 | Intron |
| chr3 | 133014603 | 133015611 | Intron |
| chr3 | 133015752 | 133016446 | Intron |
| chr3 | 135239930 | 135240841 | Distal Intergenic |
| chr3 | 142128097 | 142128405 | Distal Intergenic |
| chr3 | 144261652 | 144261995 | Distal Intergenic |
| chr3 | 144712250 | 144712738 | Distal Intergenic |
| chr3 | 145015083 | 145015473 | Intron |
| chr3 | 145617222 | 145617524 | Distal Intergenic |
| chr3 | 146222946 | 146223388 | Distal Intergenic |
| chr3 | 148715473 | 148715834 | Distal Intergenic |
| chr3 | 150426958 | 150427220 | Distal Intergenic |
| chr3 | 151792063 | 151792411 | Intron |
| chr3 | 152312647 | 152312951 | Intron |
| chr3 | 152508188 | 152508502 | Intron |
| chr3 | 153287103 | 153288120 | Intron |
| chr3 | 154271686 | 154272403 | Intron |
| chr3 | 154272625 | 154273013 | Intron |
| chr4 | 3154863 | 3155990 | Distal Intergenic |
| chr4 | 3348692 | 3348928 | Distal Intergenic |
| chr4 | 3348967 | 3350758 | Distal Intergenic |
| chr4 | 10844506 | 10845353 | Distal Intergenic |
| chr4 | 12285028 | 12285846 | Distal Intergenic |
| chr4 | 16010415 | 16010696 | Distal Intergenic |
| chr4 | 16010887 | 16011776 | Distal Intergenic |
| chr4 | 20118996 | 20119199 | Intron |

| | | | |
|---|---|---|---|
| chr4 | 32366935 | 32367235 | Intron |
| chr4 | 36780589 | 36781379 | Intron |
| chr4 | 40303178 | 40304050 | Distal Intergenic |
| chr4 | 40481270 | 40481939 | Distal Intergenic |
| chr4 | 54113785 | 54114271 | Distal Intergenic |
| chr4 | 56703625 | 56704366 | Distal Intergenic |
| chr4 | 56705796 | 56706428 | Distal Intergenic |
| chr4 | 56855619 | 56855818 | Intron |
| chr4 | 58587322 | 58587970 | Distal Intergenic |
| chr4 | 58588229 | 58588474 | Distal Intergenic |
| chr4 | 60093765 | 60094475 | Intron |
| chr4 | 60163228 | 60163789 | Intron |
| chr4 | 60445480 | 60445917 | Intron |
| chr4 | 60900114 | 60900316 | Intron |
| chr4 | 61108383 | 61108822 | Intron |
| chr4 | 61264467 | 61264884 | Intron |
| chr4 | 61453657 | 61454317 | Distal Intergenic |
| chr4 | 61914333 | 61914694 | Distal Intergenic |
| chr4 | 62920347 | 62920546 | Intron |
| chr4 | 80975436 | 80976436 | Intron |
| chr4 | 104552858 | 104553688 | Distal Intergenic |
| chr4 | 106781089 | 106782256 | Distal Intergenic |
| chr4 | 118076268 | 118077278 | Intron |
| chr4 | 118228153 | 118229463 | Intron |
| chr4 | 118229634 | 118229959 | Intron |
| chr4 | 119513830 | 119514664 | Intron |
| chr4 | 119892853 | 119893199 | Distal Intergenic |
| chr4 | 122132473 | 122132672 | Distal Intergenic |
| chr4 | 124056066 | 124056342 | Distal Intergenic |
| chr4 | 128063281 | 128063484 | Intron |
| chr4 | 128430368 | 128430584 | Distal Intergenic |
| chr4 | 128542311 | 128543002 | Distal Intergenic |
| chr4 | 128721091 | 128721406 | Distal Intergenic |
| chr4 | 130551172 | 130551752 | Distal Intergenic |
| chr4 | 130683554 | 130683753 | Distal Intergenic |
| chr4 | 131844972 | 131845197 | Intron |
| chr4 | 132498083 | 132499082 | Distal Intergenic |
| chr4 | 133309163 | 133309405 | Promoter |
| chr4 | 135364615 | 135364899 | Exon |
| chr4 | 136929688 | 136929938 | Distal Intergenic |
| chr4 | 136930520 | 136931108 | Distal Intergenic |
| chr4 | 139831644 | 139831843 | Distal Intergenic |
| chr4 | 139831892 | 139832100 | Distal Intergenic |
| chr4 | 141544105 | 141544305 | Intron |

| | | | |
|---|---|---|---|
| chr4 | 142157576 | 142158279 | Distal Intergenic |
| chr4 | 142158431 | 142158649 | Distal Intergenic |
| chr4 | 142225366 | 142225694 | Intron |
| chr4 | 146777742 | 146777941 | Intron |
| chr4 | 147761622 | 147761863 | Distal Intergenic |
| chr4 | 148996952 | 148997261 | Intron |
| chr4 | 150018962 | 150019290 | Intron |
| chr4 | 150905933 | 150906176 | Intron |
| chr4 | 151662894 | 151663856 | Intron |
| chr4 | 151817716 | 151818592 | Intron |
| chr4 | 152045140 | 152045667 | Distal Intergenic |
| chr4 | 152492726 | 152492969 | Distal Intergenic |
| chr4 | 152493005 | 152493204 | Distal Intergenic |
| chr4 | 152494187 | 152494579 | Distal Intergenic |
| chr4 | 152882710 | 152882984 | Distal Intergenic |
| chr4 | 152883346 | 152883556 | Distal Intergenic |
| chr4 | 153440631 | 153440839 | Intron |
| chr4 | 154427085 | 154427298 | Distal Intergenic |
| chr4 | 155375642 | 155375887 | Intron |
| chr5 | 9381303 | 9381563 | Intron |
| chr5 | 13089854 | 13090830 | Distal Intergenic |
| chr5 | 20774226 | 20774466 | Intron |
| chr5 | 20774528 | 20774836 | Intron |
| chr5 | 21412483 | 21412920 | Intron |
| chr5 | 22001289 | 22001548 | Intron |
| chr5 | 23318070 | 23318269 | Intron |
| chr5 | 23318371 | 23318576 | Intron |
| chr5 | 23318717 | 23319058 | Intron |
| chr5 | 24161953 | 24163273 | Distal Intergenic |
| chr5 | 24221785 | 24222109 | Intron |
| chr5 | 25321803 | 25322113 | Intron |
| chr5 | 25322211 | 25322442 | Intron |
| chr5 | 25322899 | 25324256 | Intron |
| chr5 | 27079893 | 27080103 | Distal Intergenic |
| chr5 | 28051245 | 28051718 | Intron |
| chr5 | 28052372 | 28052790 | 3' UTR |
| chr5 | 28423709 | 28424008 | Distal Intergenic |
| chr5 | 28424161 | 28426693 | Distal Intergenic |
| chr5 | 30796501 | 30797088 | Intron |
| chr5 | 36301660 | 36302539 | Intron |
| chr5 | 36623676 | 36624116 | Intron |
| chr5 | 39742037 | 39742299 | Distal Intergenic |
| chr5 | 43625949 | 43626148 | Promoter |
| chr5 | 44084986 | 44085241 | Intron |

| | | | |
|---|---|---|---|
| chr5 | 50566048 | 50566399 | Distal Intergenic |
| chr5 | 63243318 | 63243613 | Intron |
| chr5 | 64735168 | 64735747 | Intron |
| chr5 | 65435679 | 65436611 | Distal Intergenic |
| chr5 | 67818485 | 67818766 | Promoter |
| chr5 | 75560157 | 75560377 | Intron |
| chr5 | 77187050 | 77187746 | Distal Intergenic |
| chr5 | 86855222 | 86856208 | Intron |
| chr5 | 91402205 | 91402488 | Intron |
| chr5 | 92223279 | 92223960 | Distal Intergenic |
| chr5 | 96830579 | 96832012 | Intron |
| chr5 | 101639006 | 101639780 | Distal Intergenic |
| chr5 | 101845056 | 101845374 | Distal Intergenic |
| chr5 | 101845810 | 101846532 | Distal Intergenic |
| chr5 | 102685482 | 102685863 | Intron |
| chr5 | 104759722 | 104761314 | Intron |
| chr5 | 107237204 | 107237640 | Distal Intergenic |
| chr5 | 107815378 | 107815917 | 3' UTR |
| chr5 | 111007090 | 111007362 | Intron |
| chr5 | 111869765 | 111870259 | Intron |
| chr5 | 111987710 | 111988522 | Distal Intergenic |
| chr5 | 112004520 | 112005510 | Distal Intergenic |
| chr5 | 112121697 | 112122224 | Distal Intergenic |
| chr5 | 113025004 | 113025526 | Distal Intergenic |
| chr5 | 113199443 | 113200349 | Intron |
| chr5 | 113451140 | 113451506 | Distal Intergenic |
| chr5 | 113601928 | 113602148 | Distal Intergenic |
| chr5 | 116555674 | 116555893 | Distal Intergenic |
| chr5 | 116612913 | 116613531 | Exon |
| chr5 | 116649550 | 116649751 | Intron |
| chr5 | 117476469 | 117476812 | Distal Intergenic |
| chr5 | 118739894 | 118740148 | Distal Intergenic |
| chr5 | 118767038 | 118767528 | Distal Intergenic |
| chr5 | 118773675 | 118774541 | Distal Intergenic |
| chr5 | 119441695 | 119444450 | Distal Intergenic |
| chr5 | 119856945 | 119857283 | Distal Intergenic |
| chr5 | 120060206 | 120060557 | Intron |
| chr5 | 120060685 | 120060918 | Intron |
| chr5 | 120221428 | 120221636 | Distal Intergenic |
| chr5 | 121086630 | 121087408 | Distal Intergenic |
| chr5 | 125258451 | 125258656 | Intron |
| chr5 | 125258724 | 125259090 | Intron |
| chr5 | 127607662 | 127608136 | Distal Intergenic |
| chr5 | 130984003 | 130984296 | Intron |

| | | | |
|---|---|---|---|
| chr5 | 131041610 | 131041809 | Intron |
| chr5 | 132960971 | 132961170 | Intron |
| chr5 | 133427438 | 133427880 | Distal Intergenic |
| chr5 | 133898677 | 133899898 | Distal Intergenic |
| chr5 | 134435117 | 134435850 | Distal Intergenic |
| chr5 | 134474527 | 134475160 | Distal Intergenic |
| chr5 | 136531541 | 136531771 | Distal Intergenic |
| chr5 | 137042480 | 137042679 | Promoter |
| chr5 | 137043052 | 137043457 | Promoter |
| chr5 | 137347665 | 137348025 | Intron |
| chr5 | 139579904 | 139580906 | Intron |
| chr5 | 140415370 | 140415637 | Intron |
| chr5 | 141433443 | 141434333 | Intron |
| chr5 | 141450854 | 141452314 | Intron |
| chr5 | 145153565 | 145153773 | Intron |
| chr5 | 147072575 | 147072923 | Intron |
| chr5 | 147179714 | 147180148 | Distal Intergenic |
| chr5 | 148771054 | 148771335 | Intron |
| chr5 | 150283611 | 150284236 | Distal Intergenic |
| chr5 | 151776440 | 151776771 | Intron |
| chr6 | 3150795 | 3151622 | Distal Intergenic |
| chr6 | 4942933 | 4943262 | Intron |
| chr6 | 5306485 | 5307227 | Distal Intergenic |
| chr6 | 7156808 | 7157057 | Distal Intergenic |
| chr6 | 7960792 | 7961363 | Intron |
| chr6 | 12664640 | 12665643 | Intron |
| chr6 | 12665981 | 12666277 | Intron |
| chr6 | 14370215 | 14370498 | Distal Intergenic |
| chr6 | 17442653 | 17443723 | Intron |
| chr6 | 21159160 | 21159359 | Distal Intergenic |
| chr6 | 22069606 | 22069950 | Intron |
| chr6 | 29697690 | 29699139 | Intron |
| chr6 | 31789435 | 31789647 | Distal Intergenic |
| chr6 | 44742951 | 44743173 | Distal Intergenic |
| chr6 | 48161607 | 48161925 | Intron |
| chr6 | 49186459 | 49186667 | Distal Intergenic |
| chr6 | 49287659 | 49288463 | Intron |
| chr6 | 51234214 | 51234413 | Distal Intergenic |
| chr6 | 52999406 | 52999618 | Intron |
| chr6 | 53304019 | 53304401 | Intron |
| chr6 | 53383290 | 53383595 | Intron |
| chr6 | 61898744 | 61899149 | Intron |
| chr6 | 65797943 | 65798142 | Intron |
| chr6 | 65798209 | 65798423 | Intron |

| | | | |
|---|---|---|---|
| chr6 | 65798540 | 65798766 | Intron |
| chr6 | 81926764 | 81927871 | Distal Intergenic |
| chr6 | 82425512 | 82425796 | Intron |
| chr6 | 83480079 | 83480471 | Intron |
| chr6 | 86590311 | 86590901 | Distal Intergenic |
| chr6 | 95158041 | 95158274 | Distal Intergenic |
| chr6 | 95158322 | 95159009 | Distal Intergenic |
| chr6 | 96960146 | 96960724 | Intron |
| chr6 | 99642414 | 99642668 | Promoter |
| chr6 | 99832973 | 99833406 | Distal Intergenic |
| chr6 | 99903691 | 99903922 | Distal Intergenic |
| chr6 | 100237022 | 100237519 | Promoter |
| chr6 | 103264452 | 103264860 | Distal Intergenic |
| chr6 | 104183580 | 104183874 | Distal Intergenic |
| chr6 | 105111414 | 105111695 | Distal Intergenic |
| chr6 | 114291028 | 114292674 | Distal Intergenic |
| chr6 | 122474590 | 122474789 | Intron |
| chr6 | 122792329 | 122792674 | Intron |
| chr6 | 124940017 | 124940257 | Distal Intergenic |
| chr6 | 125055387 | 125055899 | Exon |
| chr6 | 125254644 | 125254907 | Downstream |
| chr6 | 125255057 | 125255308 | Downstream |
| chr6 | 125256124 | 125256408 | Downstream |
| chr6 | 125967464 | 125968029 | Intron |
| chr6 | 125968066 | 125968355 | Intron |
| chr6 | 126700569 | 126701254 | Distal Intergenic |
| chr6 | 126781782 | 126782290 | Intron |
| chr6 | 127209842 | 127210053 | Distal Intergenic |
| chr6 | 128802344 | 128802645 | Intron |
| chr6 | 134344825 | 134345196 | Intron |
| chr6 | 134516796 | 134517550 | Promoter |
| chr6 | 134519942 | 134521027 | Distal Intergenic |
| chr6 | 134521537 | 134521760 | Distal Intergenic |
| chr6 | 134522570 | 134522769 | Distal Intergenic |
| chr6 | 135287689 | 135288064 | Distal Intergenic |
| chr6 | 136951520 | 136951765 | Distal Intergenic |
| chr6 | 137791957 | 137792249 | Distal Intergenic |
| chr6 | 138033281 | 138033571 | Distal Intergenic |
| chr6 | 138033898 | 138035071 | Distal Intergenic |
| chr6 | 139701393 | 139701592 | Intron |
| chr6 | 140763924 | 140764461 | Distal Intergenic |
| chr6 | 140930141 | 140930363 | Intron |
| chr6 | 142143081 | 142143280 | Distal Intergenic |
| chr6 | 142345338 | 142348628 | Intron |

| | | | |
|---|---|---|---|
| chr6 | 142719271 | 142719698 | Exon |
| chr6 | 143005397 | 143006868 | Intron |
| chr6 | 143199470 | 143199677 | Intron |
| chr6 | 143345801 | 143346606 | Distal Intergenic |
| chr6 | 145117057 | 145118138 | 3' UTR |
| chr6 | 145306879 | 145307163 | Distal Intergenic |
| chr6 | 145622437 | 145622905 | Distal Intergenic |
| chr6 | 145881298 | 145881526 | Distal Intergenic |
| chr6 | 145968083 | 145968604 | Distal Intergenic |
| chr6 | 147863748 | 147864350 | Distal Intergenic |
| chr7 | 6811030 | 6811296 | Intron |
| chr7 | 7230526 | 7231059 | Promoter |
| chr7 | 20257444 | 20258165 | Intron |
| chr7 | 25533520 | 25534170 | Intron |
| chr7 | 28205687 | 28206008 | Intron |
| chr7 | 28881001 | 28883596 | Intron |
| chr7 | 29884124 | 29884455 | Intron |
| chr7 | 29884502 | 29884905 | Intron |
| chr7 | 51876493 | 51876848 | Intron |
| chr7 | 53266441 | 53267046 | Intron |
| chr7 | 54915555 | 54915825 | Distal Intergenic |
| chr7 | 61901759 | 61902251 | Distal Intergenic |
| chr7 | 71610578 | 71641007 | Distal Intergenic |
| chr7 | 79898759 | 79899003 | Distal Intergenic |
| chr7 | 86962897 | 86963151 | Distal Intergenic |
| chr7 | 86973493 | 86975317 | Intron |
| chr7 | 89919542 | 89920545 | Intron |
| chr7 | 91378236 | 91378820 | Distal Intergenic |
| chr7 | 99971331 | 99971648 | Intron |
| chr7 | 104592068 | 104592787 | Distal Intergenic |
| chr7 | 104944696 | 104944902 | Intron |
| chr7 | 107090875 | 107091530 | Exon |
| chr7 | 108174628 | 108175006 | Distal Intergenic |
| chr7 | 112641213 | 112642223 | Distal Intergenic |
| chr7 | 114322642 | 114322946 | Intron |
| chr7 | 114323175 | 114323640 | Intron |
| chr7 | 120316415 | 120316726 | Distal Intergenic |
| chr7 | 120615428 | 120616054 | Distal Intergenic |
| chr7 | 125618003 | 125618289 | Distal Intergenic |
| chr7 | 127189437 | 127189636 | Distal Intergenic |
| chr7 | 128648593 | 128648795 | Intron |
| chr7 | 129088133 | 129089189 | Intron |
| chr7 | 129694401 | 129694600 | Intron |
| chr7 | 131388771 | 131389450 | Distal Intergenic |

| | | | |
|---|---|---|---|
| chr7 | 133302033 | 133302423 | Intron |
| chr7 | 133394404 | 133394773 | Distal Intergenic |
| chr7 | 134905111 | 134905854 | Intron |
| chr7 | 135226185 | 135226480 | Intron |
| chr7 | 135342581 | 135343046 | Intron |
| chr7 | 135343097 | 135343519 | Intron |
| chr7 | 137004594 | 137004953 | Distal Intergenic |
| chr7 | 137005089 | 137005733 | Distal Intergenic |
| chr7 | 137942505 | 137942764 | Intron |
| chr7 | 137977871 | 137978134 | Distal Intergenic |
| chr7 | 137994760 | 137995011 | Distal Intergenic |
| chr7 | 139249183 | 139249389 | 3' UTR |
| chr7 | 139249438 | 139250466 | Exon |
| chr7 | 141001843 | 141004753 | Intron |
| chr7 | 146688530 | 146688775 | Intron |
| chr7 | 148691749 | 148694982 | Intron |
| chr8 | 10692142 | 10692433 | Distal Intergenic |
| chr8 | 11594976 | 11595176 | Distal Intergenic |
| chr8 | 11674823 | 11675212 | Intron |
| chr8 | 13070151 | 13070373 | Intron |
| chr8 | 13557112 | 13557332 | Intron |
| chr8 | 13686800 | 13687052 | Distal Intergenic |
| chr8 | 13966923 | 13967904 | Intron |
| chr8 | 15519802 | 15520092 | Distal Intergenic |
| chr8 | 22044394 | 22044639 | Distal Intergenic |
| chr8 | 23946506 | 23946962 | Distal Intergenic |
| chr8 | 24363753 | 24364036 | Intron |
| chr8 | 24918448 | 24918647 | Intron |
| chr8 | 33932668 | 33933049 | Distal Intergenic |
| chr8 | 33933109 | 33933308 | Distal Intergenic |
| chr8 | 36044098 | 36044489 | Distal Intergenic |
| chr8 | 42455155 | 42455566 | Intron |
| chr8 | 46468706 | 46469266 | Distal Intergenic |
| chr8 | 55181041 | 55181242 | Intron |
| chr8 | 61933691 | 61933974 | Distal Intergenic |
| chr8 | 66077241 | 66077988 | Distal Intergenic |
| chr8 | 81040777 | 81041142 | Intron |
| chr8 | 81276659 | 81277103 | Distal Intergenic |
| chr8 | 81350719 | 81351020 | Distal Intergenic |
| chr8 | 84742829 | 84743033 | Distal Intergenic |
| chr8 | 86647068 | 86649561 | Promoter |
| chr8 | 90712548 | 90713421 | Distal Intergenic |
| chr8 | 93617558 | 93617813 | Intron |
| chr8 | 94205615 | 94205940 | Distal Intergenic |

| | | | |
|---|---|---|---|
| chr8 | 97914331 | 97914732 | Distal Intergenic |
| chr8 | 112979721 | 112980280 | Intron |
| chr8 | 116532450 | 116532649 | Distal Intergenic |
| chr8 | 118071837 | 118072196 | Distal Intergenic |
| chr8 | 119607086 | 119607285 | Promoter |
| chr8 | 119607503 | 119607769 | Distal Intergenic |
| chr8 | 122195863 | 122196120 | Intron |
| chr8 | 122216352 | 122216676 | Distal Intergenic |
| chr8 | 122523289 | 122524236 | Intron |
| chr8 | 124345965 | 124347127 | Intron |
| chr8 | 124347325 | 124348355 | Intron |
| chr8 | 124348668 | 124350085 | Intron |
| chr8 | 124350147 | 124350958 | Intron |
| chr8 | 126124567 | 126124961 | Distal Intergenic |
| chr8 | 126142447 | 126142712 | Distal Intergenic |
| chr8 | 126147428 | 126147756 | Distal Intergenic |
| chr8 | 126163841 | 126165069 | Distal Intergenic |
| chr8 | 126165480 | 126165773 | Distal Intergenic |
| chr8 | 126708796 | 126709189 | Distal Intergenic |
| chr8 | 127908003 | 127908323 | Distal Intergenic |
| chr8 | 128651523 | 128651901 | Distal Intergenic |
| chr8 | 129433850 | 129434220 | Distal Intergenic |
| chr8_random | 162830 | 163029 | Promoter |
| chr8_random | 304327 | 304751 | Distal Intergenic |
| chr9 | 3000191 | 3000849 | Distal Intergenic |
| chr9 | 3009553 | 3009768 | Distal Intergenic |
| chr9 | 3013016 | 3013219 | Distal Intergenic |
| chr9 | 3014154 | 3014434 | Distal Intergenic |
| chr9 | 3024257 | 3024464 | Distal Intergenic |
| chr9 | 3032428 | 3032627 | Distal Intergenic |
| chr9 | 3258676 | 3259089 | Distal Intergenic |
| chr9 | 13923734 | 13924096 | Distal Intergenic |
| chr9 | 14744537 | 14745378 | Distal Intergenic |
| chr9 | 15424251 | 15424662 | Exon |
| chr9 | 20759267 | 20759587 | Intron |
| chr9 | 22024582 | 22024907 | Distal Intergenic |
| chr9 | 22249205 | 22250054 | Downstream |
| chr9 | 22414865 | 22416128 | Intron |
| chr9 | 22516063 | 22517078 | Intron |
| chr9 | 26274795 | 26275151 | Distal Intergenic |
| chr9 | 29291435 | 29291889 | Intron |
| chr9 | 30399588 | 30399891 | Distal Intergenic |
| chr9 | 31539220 | 31539434 | Distal Intergenic |
| chr9 | 32094323 | 32094584 | Distal Intergenic |

| | | | |
|---|---|---|---|
| chr9 | 32115191 | 32116875 | Distal Intergenic |
| chr9 | 34404195 | 34404505 | Intron |
| chr9 | 40986505 | 40987405 | Distal Intergenic |
| chr9 | 42603332 | 42603532 | Intron |
| chr9 | 45339683 | 45340059 | Intron |
| chr9 | 45518847 | 45519570 | Intron |
| chr9 | 46823907 | 46824280 | Distal Intergenic |
| chr9 | 48015607 | 48016301 | Intron |
| chr9 | 48198704 | 48199022 | Promoter |
| chr9 | 48470004 | 48470496 | Intron |
| chr9 | 50394567 | 50395055 | Distal Intergenic |
| chr9 | 56071384 | 56071740 | Intron |
| chr9 | 56619766 | 56620052 | Distal Intergenic |
| chr9 | 56620225 | 56620424 | Distal Intergenic |
| chr9 | 58224476 | 58225448 | Distal Intergenic |
| chr9 | 61102006 | 61102663 | Distal Intergenic |
| chr9 | 61438580 | 61438809 | Distal Intergenic |
| chr9 | 62516383 | 62516725 | Distal Intergenic |
| chr9 | 67287980 | 67289077 | Intron |
| chr9 | 67289114 | 67289505 | Intron |
| chr9 | 68947359 | 68947955 | Intron |
| chr9 | 70470745 | 70471419 | Intron |
| chr9 | 70804196 | 70804923 | Distal Intergenic |
| chr9 | 70835792 | 70836271 | Distal Intergenic |
| chr9 | 73097391 | 73097653 | Distal Intergenic |
| chr9 | 73097819 | 73098034 | Distal Intergenic |
| chr9 | 77500403 | 77501043 | Intron |
| chr9 | 85064784 | 85065887 | Distal Intergenic |
| chr9 | 88147861 | 88148107 | Distal Intergenic |
| chr9 | 94263246 | 94264193 | Distal Intergenic |
| chr9 | 99889491 | 99889933 | Distal Intergenic |
| chr9 | 101312283 | 101312620 | Distal Intergenic |
| chr9 | 105473859 | 105474262 | Distal Intergenic |
| chr9 | 112378095 | 112378332 | Distal Intergenic |
| chr9 | 112936153 | 112936760 | Distal Intergenic |
| chr9 | 115853976 | 115854281 | Intron |
| chr9 | 116746515 | 116747615 | Intron |
| chr9 | 117144178 | 117144519 | Intron |
| chr9 | 117997210 | 117999177 | Intron |
| chr9 | 117999321 | 118002046 | Intron |
| chr9 | 120950732 | 120951272 | Intron |
| chr9 | 121788640 | 121788839 | Distal Intergenic |
| chr9 | 122813426 | 122815467 | Distal Intergenic |
| chr9 | 123370962 | 123371245 | 3' UTR |

| | | | |
|---|---|---|---|
| chrX | 6589946 | 6592899 | Intron |
| chrX | 35309657 | 35309914 | Distal Intergenic |
| chrX | 35309961 | 35310445 | Distal Intergenic |
| chrX | 35310512 | 35315747 | Distal Intergenic |
| chrX | 35315958 | 35316473 | Distal Intergenic |
| chrX | 35316544 | 35316756 | Distal Intergenic |
| chrX | 35316816 | 35324757 | Distal Intergenic |
| chrX | 35324980 | 35325808 | Distal Intergenic |
| chrX | 35332039 | 35332481 | Distal Intergenic |
| chrX | 35332595 | 35332853 | Distal Intergenic |
| chrX | 35333460 | 35333969 | Distal Intergenic |
| chrX | 80828573 | 80849202 | Intron |
| chrX | 108419663 | 108419874 | Distal Intergenic |
| chrX | 108472161 | 108472493 | Distal Intergenic |
| chrX | 115912685 | 115912915 | Distal Intergenic |
| chrX | 147089030 | 147089436 | Promoter |
| chrX | 153695973 | 153696183 | Intron |
| chrX | 153696734 | 153696958 | Intron |
| chrX | 153698304 | 153698503 | Intron |
| chrX | 166394537 | 166394784 | Intron |
| chrX | 166394930 | 166395152 | Intron |
| chrX | 166421281 | 166421649 | Intron |
| chrX | 166424842 | 166425092 | Exon |
| chrX | 166425194 | 166426888 | Promoter |

**Supplementary Table 3. H3K27ac Peak Annotation**

| chromosome | start | end | annotation |
|---|---|---|---|
| chr1 | 3659367 | 3660513 | Exon |
| chr1 | 4775184 | 4775563 | Intron |
| chr1 | 4847458 | 4849049 | 5' UTR |
| chr1 | 5005189 | 5005818 | Distal Intergenic |
| chr1 | 5072993 | 5073908 | Distal Intergenic |
| chr1 | 5077133 | 5077339 | Distal Intergenic |
| chr1 | 6178409 | 6178742 | Intron |
| chr1 | 6178794 | 6179363 | Intron |
| chr1 | 6204043 | 6204986 | Promoter |
| chr1 | 6269533 | 6269835 | Promoter |
| chr1 | 6270005 | 6270954 | 3' UTR |
| chr1 | 6279205 | 6281129 | 3' UTR |
| chr1 | 6301364 | 6302242 | Distal Intergenic |
| chr1 | 6302465 | 6302805 | Distal Intergenic |
| chr1 | 6477028 | 6477346 | 3' UTR |
| chr1 | 6489133 | 6489343 | Promoter |

| | | | |
|---|---|---|---|
| chr1 | 6918222 | 6918783 | Intron |
| chr1 | 6918888 | 6919326 | Intron |
| chr1 | 6929209 | 6930310 | Intron |
| chr1 | 7079333 | 7079726 | Intron |
| chr1 | 7079821 | 7080124 | Intron |
| chr1 | 7115175 | 7116832 | Intron |
| chr1 | 7189911 | 7190409 | Intron |
| chr1 | 9122221 | 9122502 | Intron |
| chr1 | 9122608 | 9124315 | Intron |
| chr1 | 9143384 | 9144326 | Intron |
| chr1 | 9535487 | 9535720 | Distal Intergenic |
| chr1 | 9536047 | 9536822 | Distal Intergenic |
| chr1 | 9621398 | 9621663 | Intron |
| chr1 | 9628433 | 9629652 | Intron |
| chr1 | 9737554 | 9737768 | Intron |
| chr1 | 9737820 | 9738552 | Intron |
| chr1 | 9744715 | 9745192 | Intron |
| chr1 | 9745233 | 9746399 | Intron |
| chr1 | 9788511 | 9790619 | 3' UTR |
| chr1 | 9931302 | 9932054 | Exon |
| chr1 | 9932414 | 9934450 | Intron |
| chr1 | 9973002 | 9973703 | Distal Intergenic |
| chr1 | 10027325 | 10029226 | Intron |
| chr1 | 10029399 | 10029615 | Intron |
| chr1 | 10221493 | 10223262 | Intron |
| chr1 | 10243560 | 10244887 | Distal Intergenic |
| chr1 | 10534743 | 10535457 | Promoter |
| chr1 | 10560360 | 10561915 | Intron |
| chr1 | 10582633 | 10583839 | Intron |
| chr1 | 10626595 | 10628238 | Intron |
| chr1 | 10862575 | 10863000 | Distal Intergenic |
| chr1 | 10870162 | 10870491 | Distal Intergenic |
| chr1 | 12610119 | 12614376 | Distal Intergenic |
| chr1 | 12766559 | 12767760 | Distal Intergenic |
| chr1 | 13206347 | 13206983 | Distal Intergenic |
| chr1 | 13294446 | 13294645 | Distal Intergenic |
| chr1 | 13295090 | 13295794 | Distal Intergenic |
| chr1 | 13362106 | 13362535 | Intron |
| chr1 | 13362667 | 13363077 | Intron |
| chr1 | 13363454 | 13363662 | Intron |
| chr1 | 13568926 | 13570376 | Distal Intergenic |
| chr1 | 13605552 | 13606695 | Downstream |
| chr1 | 13629408 | 13629734 | Distal Intergenic |
| chr1 | 13629802 | 13630254 | Exon |

| | | | |
|---|---|---|---|
| chr1 | 13633282 | 13633507 | Exon |
| chr1 | 13649192 | 13651816 | Distal Intergenic |
| chr1 | 13863984 | 13864205 | Distal Intergenic |
| chr1 | 13991673 | 13991945 | Distal Intergenic |
| chr1 | 14263801 | 14264113 | Distal Intergenic |
| chr1 | 14474066 | 14475019 | Distal Intergenic |
| chr1 | 15709446 | 15710698 | 3' UTR |
| chr1 | 15774287 | 15775657 | Distal Intergenic |
| chr1 | 15781180 | 15781565 | Distal Intergenic |
| chr1 | 15781613 | 15782179 | Distal Intergenic |
| chr1 | 15795202 | 15795600 | Intron |
| chr1 | 15795740 | 15796746 | Intron |
| chr1 | 15988053 | 15988832 | Exon |
| chr1 | 16093615 | 16095130 | Exon |
| chr1 | 16119515 | 16120994 | Distal Intergenic |
| chr1 | 16162003 | 16162289 | Exon |
| chr1 | 16162403 | 16162684 | Exon |
| chr1 | 16218173 | 16218833 | Intron |
| chr1 | 16409666 | 16409874 | Distal Intergenic |
| chr1 | 16608069 | 16608268 | Intron |
| chr1 | 16608310 | 16608774 | Intron |
| chr1 | 16609002 | 16609207 | Intron |
| chr1 | 16646084 | 16647491 | Intron |
| chr1 | 16651064 | 16652216 | Intron |
| chr1 | 16655025 | 16655480 | Intron |
| chr1 | 16655814 | 16656375 | Intron |
| chr1 | 16665736 | 16665940 | Intron |
| chr1 | 16678046 | 16678717 | Promoter |
| chr1 | 16678936 | 16679253 | Promoter |
| chr1 | 16686547 | 16686747 | Distal Intergenic |
| chr1 | 16879911 | 16880325 | Distal Intergenic |
| chr1 | 16997917 | 16998228 | Distal Intergenic |
| chr1 | 16998360 | 16999141 | Distal Intergenic |
| chr1 | 17478299 | 17481534 | Distal Intergenic |
| chr1 | 17594485 | 17594740 | Intron |
| chr1 | 17594941 | 17596302 | Intron |
| chr1 | 18120233 | 18120482 | Intron |
| chr1 | 19764054 | 19765253 | Intron |
| chr1 | 19784499 | 19784793 | Intron |
| chr1 | 19941881 | 19942091 | Intron |
| chr1 | 19946790 | 19947492 | Intron |
| chr1 | 20861225 | 20861439 | Distal Intergenic |
| chr1 | 20992336 | 20992881 | Exon |
| chr1 | 20998390 | 20999449 | Exon |

| | | | |
|---|---|---|---|
| chr1 | 21216230 | 21217988 | Intron |
| chr1 | 21224898 | 21226213 | Exon |
| chr1 | 21229975 | 21231368 | Intron |
| chr1 | 23196709 | 23197508 | Intron |
| chr1 | 23262609 | 23262808 | Distal Intergenic |
| chr1 | 23274676 | 23274891 | Distal Intergenic |
| chr1 | 23276657 | 23276898 | Distal Intergenic |
| chr1 | 23277401 | 23279446 | Distal Intergenic |
| chr1 | 23919452 | 23920612 | Intron |
| chr1 | 23920667 | 23920944 | Intron |
| chr1 | 23924079 | 23925296 | Intron |
| chr1 | 23925576 | 23926683 | Intron |
| chr1 | 23927688 | 23928100 | Intron |
| chr1 | 24109331 | 24109923 | Intron |
| chr1 | 24113896 | 24115697 | 3' UTR |
| chr1 | 24685568 | 24685856 | Intron |
| chr1 | 24686060 | 24686294 | Intron |
| chr1 | 26340634 | 26340833 | Distal Intergenic |
| chr1 | 26355245 | 26355461 | Intron |
| chr1 | 26446026 | 26446419 | Intron |
| chr1 | 26451374 | 26451659 | 3' UTR |
| chr1 | 26451868 | 26453870 | 3' UTR |
| chr1 | 28898267 | 28898552 | Exon |
| chr1 | 30239990 | 30242119 | Distal Intergenic |
| chr1 | 30271784 | 30272092 | Distal Intergenic |
| chr1 | 30272159 | 30272602 | Distal Intergenic |
| chr1 | 30930586 | 30933222 | Distal Intergenic |
| chr1 | 30960346 | 30960765 | Distal Intergenic |
| chr1 | 31005299 | 31005990 | Distal Intergenic |
| chr1 | 31006341 | 31006543 | Distal Intergenic |
| chr1 | 31006704 | 31007697 | Distal Intergenic |
| chr1 | 31070754 | 31071235 | Distal Intergenic |
| chr1 | 31152753 | 31153682 | Distal Intergenic |
| chr1 | 31834992 | 31835823 | Intron |
| chr1 | 33582563 | 33583046 | Intron |
| chr1 | 33586695 | 33587147 | Distal Intergenic |
| chr1 | 33607853 | 33608227 | Exon |
| chr1 | 33813506 | 33814062 | Intron |
| chr1 | 33814276 | 33814536 | Promoter |
| chr1 | 33814775 | 33815040 | Promoter |
| chr1 | 33870700 | 33871401 | Intron |
| chr1 | 34130367 | 34130566 | Intron |
| chr1 | 34130952 | 34131266 | Intron |
| chr1 | 34131343 | 34131770 | Intron |

| | | | |
|---|---|---|---|
| chr1 | 34166219 | 34167281 | Exon |
| chr1 | 34169040 | 34169258 | Intron |
| chr1 | 34177463 | 34178803 | Intron |
| chr1 | 34178883 | 34179098 | Intron |
| chr1 | 34179239 | 34179440 | Intron |
| chr1 | 34182584 | 34183005 | Intron |
| chr1 | 34184149 | 34185240 | Intron |
| chr1 | 34196703 | 34197023 | Intron |
| chr1 | 34197145 | 34198488 | Intron |
| chr1 | 34198563 | 34200669 | Intron |
| chr1 | 34204413 | 34205899 | Exon |
| chr1 | 34206817 | 34208317 | Intron |
| chr1 | 34215362 | 34219733 | Intron |
| chr1 | 34220163 | 34220372 | Intron |
| chr1 | 34220574 | 34221180 | Intron |
| chr1 | 34221224 | 34221670 | Intron |
| chr1 | 34240781 | 34241475 | Intron |
| chr1 | 34241633 | 34243486 | Intron |
| chr1 | 34699650 | 34699991 | Distal Intergenic |
| chr1 | 34734533 | 34735626 | Distal Intergenic |
| chr1 | 34757674 | 34758460 | Distal Intergenic |
| chr1 | 34758524 | 34758759 | Distal Intergenic |
| chr1 | 34768711 | 34769184 | Distal Intergenic |
| chr1 | 34769235 | 34769968 | Distal Intergenic |
| chr1 | 34771068 | 34771283 | Distal Intergenic |
| chr1 | 34772890 | 34774443 | Distal Intergenic |
| chr1 | 34776285 | 34780423 | Distal Intergenic |
| chr1 | 34855952 | 34856425 | Distal Intergenic |
| chr1 | 34898605 | 34898916 | Distal Intergenic |
| chr1 | 34899062 | 34899338 | Distal Intergenic |
| chr1 | 34900131 | 34900331 | Distal Intergenic |
| chr1 | 34906784 | 34906984 | Distal Intergenic |
| chr1 | 34907043 | 34909679 | Distal Intergenic |
| chr1 | 35719600 | 35720185 | Distal Intergenic |
| chr1 | 35970263 | 35971217 | Intron |
| chr1 | 36073778 | 36074004 | Intron |
| chr1 | 36116257 | 36118427 | Distal Intergenic |
| chr1 | 36126639 | 36126922 | Distal Intergenic |
| chr1 | 36145051 | 36145370 | Distal Intergenic |
| chr1 | 36146469 | 36147503 | Distal Intergenic |
| chr1 | 36152334 | 36153097 | Distal Intergenic |
| chr1 | 36173359 | 36174290 | Distal Intergenic |
| chr1 | 36180534 | 36181012 | 3' UTR |
| chr1 | 36216995 | 36217692 | Exon |

| | | | |
|---|---|---|---|
| chr1 | 36300843 | 36301500 | Exon |
| chr1 | 36301765 | 36302060 | Intron |
| chr1 | 36302297 | 36302542 | Intron |
| chr1 | 36366089 | 36366288 | Intron |
| chr1 | 36382458 | 36383027 | Intron |
| chr1 | 36383108 | 36383427 | Exon |
| chr1 | 36425363 | 36425625 | Intron |
| chr1 | 36490109 | 36490420 | Intron |
| chr1 | 36626600 | 36627160 | Intron |
| chr1 | 36630431 | 36631158 | Intron |
| chr1 | 36637671 | 36638230 | Promoter |
| chr1 | 36640455 | 36641358 | 5' UTR |
| chr1 | 36704064 | 36705002 | Promoter |
| chr1 | 36748338 | 36748767 | Intron |
| chr1 | 36887679 | 36889193 | Exon |
| chr1 | 36889282 | 36889558 | Intron |
| chr1 | 36915614 | 36916922 | Promoter |
| chr1 | 36960799 | 36961951 | Distal Intergenic |
| chr1 | 36971954 | 36972806 | Distal Intergenic |
| chr1 | 36976559 | 36977938 | Distal Intergenic |
| chr1 | 36979846 | 36981020 | Distal Intergenic |
| chr1 | 36994093 | 36994340 | Distal Intergenic |
| chr1 | 36994409 | 36996020 | Distal Intergenic |
| chr1 | 36996268 | 36997061 | Distal Intergenic |
| chr1 | 37005614 | 37007660 | Distal Intergenic |
| chr1 | 37022188 | 37022867 | Distal Intergenic |
| chr1 | 37145757 | 37146526 | Distal Intergenic |
| chr1 | 37272251 | 37273697 | Intron |
| chr1 | 37486311 | 37486900 | Intron |
| chr1 | 37486987 | 37487625 | Intron |
| chr1 | 37643140 | 37643594 | Distal Intergenic |
| chr1 | 37651999 | 37652943 | Distal Intergenic |
| chr1 | 37928605 | 37929045 | Intron |
| chr1 | 37929321 | 37929843 | Intron |
| chr1 | 37946639 | 37947248 | Promoter |
| chr1 | 38028473 | 38029086 | Intron |
| chr1 | 38053476 | 38055936 | Intron |
| chr1 | 38086521 | 38088309 | Intron |
| chr1 | 38095178 | 38096336 | 5' UTR |
| chr1 | 38159737 | 38161006 | Intron |
| chr1 | 38361196 | 38361754 | Intron |
| chr1 | 38375297 | 38376099 | Intron |
| chr1 | 38376143 | 38378761 | Intron |
| chr1 | 38438324 | 38439190 | Intron |

| | | | |
|---|---|---|---|
| chr1 | 38450001 | 38452942 | Exon |
| chr1 | 38481313 | 38481574 | Intron |
| chr1 | 38494333 | 38494729 | Distal Intergenic |
| chr1 | 38604354 | 38606256 | Distal Intergenic |
| chr1 | 38752807 | 38753122 | Distal Intergenic |
| chr1 | 39028684 | 39030843 | Distal Intergenic |
| chr1 | 39044145 | 39044442 | Distal Intergenic |
| chr1 | 39044946 | 39045449 | Distal Intergenic |
| chr1 | 39312846 | 39314109 | Intron |
| chr1 | 39424197 | 39424396 | Distal Intergenic |
| chr1 | 39424772 | 39424986 | Distal Intergenic |
| chr1 | 39425026 | 39425294 | Distal Intergenic |
| chr1 | 39425447 | 39426097 | Distal Intergenic |
| chr1 | 39691700 | 39692052 | Intron |
| chr1 | 39713215 | 39714930 | Intron |
| chr1 | 39815953 | 39816529 | Exon |
| chr1 | 39816708 | 39817152 | Intron |
| chr1 | 39817820 | 39819219 | Exon |
| chr1 | 39945870 | 39946913 | Exon |
| chr1 | 39952021 | 39952316 | 3' UTR |
| chr1 | 39957121 | 39957602 | Promoter |
| chr1 | 39957765 | 39958172 | Promoter |
| chr1 | 39958215 | 39959992 | Promoter |
| chr1 | 39963369 | 39964136 | Intron |
| chr1 | 39988387 | 39989905 | Exon |
| chr1 | 39997813 | 39998992 | Intron |
| chr1 | 40024396 | 40026511 | Promoter |
| chr1 | 40228668 | 40228890 | Exon |
| chr1 | 40230996 | 40231496 | Intron |
| chr1 | 40328281 | 40329820 | Intron |
| chr1 | 40335457 | 40337079 | Intron |
| chr1 | 40411966 | 40412734 | Distal Intergenic |
| chr1 | 40415197 | 40415400 | Distal Intergenic |
| chr1 | 40437969 | 40438189 | Distal Intergenic |
| chr1 | 40441813 | 40442685 | Distal Intergenic |
| chr1 | 40450318 | 40450888 | Distal Intergenic |
| chr1 | 40501152 | 40502292 | Distal Intergenic |
| chr1 | 40502340 | 40502588 | Distal Intergenic |
| chr1 | 40599091 | 40600040 | Distal Intergenic |
| chr1 | 40643567 | 40643766 | Intron |
| chr1 | 42908282 | 42908481 | Intron |
| chr1 | 43155060 | 43155761 | Intron |
| chr1 | 43155797 | 43156091 | Intron |
| chr1 | 43198369 | 43199635 | 3' UTR |

| | | | |
|---|---|---|---|
| chr1 | 43200578 | 43201018 | 3' UTR |
| chr1 | 43201219 | 43205763 | Promoter |
| chr1 | 43205968 | 43206430 | Promoter |
| chr1 | 43206637 | 43207000 | Promoter |
| chr1 | 43208007 | 43208896 | Distal Intergenic |
| chr1 | 43208945 | 43209215 | Downstream |
| chr1 | 43214949 | 43215158 | Intron |
| chr1 | 43215363 | 43215592 | Intron |
| chr1 | 43216720 | 43217215 | Intron |
| chr1 | 43217985 | 43218201 | Exon |
| chr1 | 43219969 | 43220523 | Intron |
| chr1 | 43220726 | 43221200 | Exon |
| chr1 | 43223768 | 43226228 | 3' UTR |
| chr1 | 43226483 | 43226762 | 3' UTR |
| chr1 | 43259512 | 43260620 | Exon |
| chr1 | 43280838 | 43281503 | Intron |
| chr1 | 43359857 | 43360093 | Distal Intergenic |
| chr1 | 43468926 | 43469137 | Distal Intergenic |
| chr1 | 43469634 | 43469896 | Distal Intergenic |
| chr1 | 43469977 | 43470749 | Distal Intergenic |
| chr1 | 43470789 | 43471079 | Distal Intergenic |
| chr1 | 43528112 | 43529272 | Distal Intergenic |
| chr1 | 43529332 | 43529606 | Distal Intergenic |
| chr1 | 43553283 | 43554040 | Distal Intergenic |
| chr1 | 43560527 | 43561167 | Distal Intergenic |
| chr1 | 43841236 | 43842637 | Distal Intergenic |
| chr1 | 43876084 | 43876474 | Intron |
| chr1 | 43990243 | 43990733 | Distal Intergenic |
| chr1 | 43990779 | 43991885 | Distal Intergenic |
| chr1 | 44025844 | 44026574 | Intron |
| chr1 | 44053680 | 44054356 | Intron |
| chr1 | 44175505 | 44176470 | Intron |
| chr1 | 44204589 | 44205806 | Intron |
| chr1 | 45480652 | 45480938 | Exon |
| chr1 | 45786486 | 45787688 | Distal Intergenic |
| chr1 | 45852004 | 45852232 | Intron |
| chr1 | 45852269 | 45852469 | Intron |
| chr1 | 46906795 | 46907714 | Exon |
| chr1 | 46908275 | 46909184 | Exon |
| chr1 | 46910142 | 46910387 | Intron |
| chr1 | 46910430 | 46911165 | Exon |
| chr1 | 46911428 | 46911733 | Distal Intergenic |
| chr1 | 47019882 | 47020189 | Intron |
| chr1 | 47020276 | 47020976 | Intron |

| | | | |
|---|---|---|---|
| chr1 | 51122652 | 51122881 | Intron |
| chr1 | 51458849 | 51459268 | Distal Intergenic |
| chr1 | 51532821 | 51536039 | Distal Intergenic |
| chr1 | 51559434 | 51561631 | Distal Intergenic |
| chr1 | 51561710 | 51564739 | Distal Intergenic |
| chr1 | 51565144 | 51566951 | Promoter |
| chr1 | 51582677 | 51583299 | Intron |
| chr1 | 51586499 | 51587922 | Intron |
| chr1 | 51700732 | 51701389 | Promoter |
| chr1 | 51757720 | 51758914 | Intron |
| chr1 | 51804156 | 51805245 | Intron |
| chr1 | 51872479 | 51873465 | Intron |
| chr1 | 51885992 | 51886682 | Intron |
| chr1 | 51905482 | 51907057 | Exon |
| chr1 | 51907121 | 51907364 | Intron |
| chr1 | 51908793 | 51914222 | Exon |
| chr1 | 52169985 | 52170670 | Intron |
| chr1 | 52288654 | 52288890 | Intron |
| chr1 | 52311262 | 52311962 | Intron |
| chr1 | 52339305 | 52339896 | Intron |
| chr1 | 52556462 | 52556879 | Distal Intergenic |
| chr1 | 52557611 | 52559013 | Distal Intergenic |
| chr1 | 52635266 | 52635672 | Intron |
| chr1 | 52654734 | 52655412 | Intron |
| chr1 | 52779212 | 52780067 | Intron |
| chr1 | 52780694 | 52780929 | Intron |
| chr1 | 52781024 | 52781477 | Intron |
| chr1 | 52855919 | 52857075 | Intron |
| chr1 | 52859320 | 52859574 | Exon |
| chr1 | 53841241 | 53841493 | Distal Intergenic |
| chr1 | 53894751 | 53896031 | Distal Intergenic |
| chr1 | 53901758 | 53902565 | Downstream |
| chr1 | 53902608 | 53903689 | Downstream |
| chr1 | 54770303 | 54770775 | Intron |
| chr1 | 55083227 | 55084477 | Intron |
| chr1 | 55108241 | 55109447 | Promoter |
| chr1 | 55109679 | 55109910 | Intron |
| chr1 | 55143136 | 55143433 | Intron |
| chr1 | 55143948 | 55144696 | Exon |
| chr1 | 55146267 | 55146552 | Intron |
| chr1 | 55187751 | 55187975 | Intron |
| chr1 | 55188123 | 55188808 | Exon |
| chr1 | 55189219 | 55189466 | Intron |
| chr1 | 55250154 | 55250381 | 3' UTR |

| | | | |
|---|---|---|---|
| chr1 | 55252499 | 55254180 | Exon |
| chr1 | 55257981 | 55259431 | Intron |
| chr1 | 56433235 | 56433492 | Distal Intergenic |
| chr1 | 56738029 | 56738540 | Distal Intergenic |
| chr1 | 56914532 | 56915747 | Distal Intergenic |
| chr1 | 57028937 | 57030125 | Intron |
| chr1 | 57462791 | 57464079 | 3' UTR |
| chr1 | 58045660 | 58046633 | Intron |
| chr1 | 58052930 | 58053297 | Intron |
| chr1 | 58086094 | 58088189 | Intron |
| chr1 | 58360737 | 58361679 | Intron |
| chr1 | 58416034 | 58416958 | Intron |
| chr1 | 58423208 | 58424635 | Intron |
| chr1 | 58448710 | 58448945 | Intron |
| chr1 | 58449983 | 58450200 | Intron |
| chr1 | 58450413 | 58451050 | Intron |
| chr1 | 58479843 | 58480506 | Intron |
| chr1 | 58480652 | 58480888 | Intron |
| chr1 | 58480943 | 58481205 | Intron |
| chr1 | 58502038 | 58503131 | Intron |
| chr1 | 58643223 | 58643639 | Intron |
| chr1 | 58708137 | 58710594 | Intron |
| chr1 | 58715001 | 58715583 | Promoter |
| chr1 | 58719554 | 58719902 | Intron |
| chr1 | 58720120 | 58720962 | Intron |
| chr1 | 58767316 | 58769618 | Intron |
| chr1 | 58769813 | 58771104 | Intron |
| chr1 | 58798742 | 58800226 | Intron |
| chr1 | 58817688 | 58817903 | Intron |
| chr1 | 58818101 | 58819090 | Intron |
| chr1 | 58851835 | 58853307 | Intron |
| chr1 | 58853671 | 58853993 | Intron |
| chr1 | 58854039 | 58854244 | Intron |
| chr1 | 58912836 | 58913068 | Intron |
| chr1 | 58913244 | 58914952 | Intron |
| chr1 | 58916443 | 58917197 | Intron |
| chr1 | 59219286 | 59220127 | Distal Intergenic |
| chr1 | 59248008 | 59248326 | Exon |
| chr1 | 59294385 | 59294614 | Intron |
| chr1 | 59485145 | 59485362 | Distal Intergenic |
| chr1 | 59485430 | 59486926 | Distal Intergenic |
| chr1 | 59487533 | 59487916 | Distal Intergenic |
| chr1 | 59623307 | 59624669 | Distal Intergenic |
| chr1 | 59636681 | 59637527 | Distal Intergenic |

| | | | |
|---|---|---|---|
| chr1 | 59726682 | 59727965 | Distal Intergenic |
| chr1 | 59741778 | 59742675 | Distal Intergenic |
| chr1 | 59796025 | 59797594 | Intron |
| chr1 | 59819669 | 59820198 | Intron |
| chr1 | 59820315 | 59821084 | Intron |
| chr1 | 59821138 | 59821740 | Intron |
| chr1 | 59821858 | 59822376 | Intron |
| chr1 | 59852110 | 59852419 | Intron |
| chr1 | 59936988 | 59938093 | Intron |
| chr1 | 59939044 | 59940154 | Intron |
| chr1 | 60057351 | 60057682 | Intron |
| chr1 | 60154326 | 60155010 | Intron |
| chr1 | 60237354 | 60237817 | Distal Intergenic |
| chr1 | 60601241 | 60602820 | Distal Intergenic |
| chr1 | 60622564 | 60622829 | Distal Intergenic |
| chr1 | 60665770 | 60667275 | Distal Intergenic |
| chr1 | 60753227 | 60753688 | Distal Intergenic |
| chr1 | 61024531 | 61025520 | Distal Intergenic |
| chr1 | 61036244 | 61036942 | Distal Intergenic |
| chr1 | 61151586 | 61152986 | Intron |
| chr1 | 61541835 | 61542280 | Promoter |
| chr1 | 61542322 | 61542932 | Promoter |
| chr1 | 62131085 | 62133171 | Distal Intergenic |
| chr1 | 62444899 | 62445101 | Intron |
| chr1 | 62445209 | 62445468 | Intron |
| chr1 | 62767418 | 62767645 | Intron |
| chr1 | 62767900 | 62768974 | Intron |
| chr1 | 62809938 | 62811205 | Distal Intergenic |
| chr1 | 62811949 | 62812417 | Distal Intergenic |
| chr1 | 62812575 | 62812909 | Distal Intergenic |
| chr1 | 62813501 | 62815442 | Distal Intergenic |
| chr1 | 62818442 | 62818727 | Distal Intergenic |
| chr1 | 62840784 | 62841781 | Distal Intergenic |
| chr1 | 62841911 | 62842156 | Distal Intergenic |
| chr1 | 62869528 | 62870478 | Distal Intergenic |
| chr1 | 62870539 | 62870877 | Distal Intergenic |
| chr1 | 62954634 | 62955214 | Exon |
| chr1 | 62955335 | 62955549 | Intron |
| chr1 | 62968370 | 62968689 | Intron |
| chr1 | 63159336 | 63161865 | Distal Intergenic |
| chr1 | 63181569 | 63181785 | Distal Intergenic |
| chr1 | 63222791 | 63224541 | Distal Intergenic |
| chr1 | 63742912 | 63743257 | Intron |
| chr1 | 63743293 | 63743592 | Intron |

| | | | |
|---|---|---|---|
| chr1 | 64023616 | 64024088 | Intron |
| chr1 | 64027424 | 64027715 | Exon |
| chr1 | 64027986 | 64029115 | Intron |
| chr1 | 64051323 | 64051796 | Distal Intergenic |
| chr1 | 64051989 | 64052618 | Distal Intergenic |
| chr1 | 64128486 | 64129004 | Distal Intergenic |
| chr1 | 64165364 | 64166398 | Distal Intergenic |
| chr1 | 64166543 | 64168007 | Distal Intergenic |
| chr1 | 64168408 | 64169336 | Distal Intergenic |
| chr1 | 64255354 | 64258633 | Intron |
| chr1 | 64272250 | 64273645 | Intron |
| chr1 | 64298125 | 64299803 | Intron |
| chr1 | 64351769 | 64352368 | Intron |
| chr1 | 64480125 | 64481054 | Intron |
| chr1 | 64481194 | 64481486 | Intron |
| chr1 | 64579512 | 64579802 | Intron |
| chr1 | 64580192 | 64580860 | Intron |
| chr1 | 64585716 | 64585954 | Intron |
| chr1 | 64593236 | 64593533 | Intron |
| chr1 | 64647578 | 64648773 | Distal Intergenic |
| chr1 | 64661603 | 64663673 | Distal Intergenic |
| chr1 | 64663787 | 64664074 | Distal Intergenic |
| chr1 | 64802027 | 64802240 | Distal Intergenic |
| chr1 | 64802339 | 64803760 | Distal Intergenic |
| chr1 | 64841960 | 64844200 | Distal Intergenic |
| chr1 | 64855046 | 64855323 | Distal Intergenic |
| chr1 | 64855473 | 64855805 | Distal Intergenic |
| chr1 | 64860397 | 64861633 | Distal Intergenic |
| chr1 | 64987800 | 64989463 | Intron |
| chr1 | 64991480 | 64992220 | Intron |
| chr1 | 65164548 | 65166032 | Distal Intergenic |
| chr1 | 65225313 | 65225684 | Intron |
| chr1 | 65233279 | 65233599 | Intron |
| chr1 | 65436458 | 65436942 | Distal Intergenic |
| chr1 | 65878900 | 65880011 | 3' UTR |
| chr1 | 65978279 | 65980293 | Intron |
| chr1 | 65984429 | 65984924 | Intron |
| chr1 | 66108679 | 66109725 | Distal Intergenic |
| chr1 | 66186861 | 66187318 | Distal Intergenic |
| chr1 | 66191296 | 66191994 | Distal Intergenic |
| chr1 | 66327113 | 66327387 | Intron |
| chr1 | 66444273 | 66445599 | Intron |
| chr1 | 66747470 | 66747720 | Intron |
| chr1 | 66747911 | 66748360 | Intron |

| | | | |
|---|---|---|---|
| chr1 | 66834680 | 66836076 | Intron |
| chr1 | 66861193 | 66861467 | Distal Intergenic |
| chr1 | 66862145 | 66863478 | Distal Intergenic |
| chr1 | 66892038 | 66893958 | Distal Intergenic |
| chr1 | 67083365 | 67085162 | Intron |
| chr1 | 67085202 | 67085428 | Intron |
| chr1 | 67088537 | 67089216 | Intron |
| chr1 | 67090630 | 67092555 | Exon |
| chr1 | 67095340 | 67095826 | Intron |
| chr1 | 67496954 | 67498547 | Intron |
| chr1 | 68826464 | 68827065 | Distal Intergenic |
| chr1 | 69331959 | 69333643 | Distal Intergenic |
| chr1 | 69413884 | 69414250 | Distal Intergenic |
| chr1 | 69414501 | 69414922 | Distal Intergenic |
| chr1 | 69426329 | 69426718 | Distal Intergenic |
| chr1 | 69443601 | 69448965 | Distal Intergenic |
| chr1 | 69479481 | 69481522 | Distal Intergenic |
| chr1 | 69487064 | 69487707 | Distal Intergenic |
| chr1 | 69499879 | 69500159 | Distal Intergenic |
| chr1 | 69504048 | 69504406 | Distal Intergenic |
| chr1 | 69506758 | 69507045 | Distal Intergenic |
| chr1 | 69507143 | 69508437 | Distal Intergenic |
| chr1 | 69513595 | 69514108 | Distal Intergenic |
| chr1 | 69514273 | 69516431 | Distal Intergenic |
| chr1 | 69554188 | 69555682 | Distal Intergenic |
| chr1 | 69701505 | 69702953 | Distal Intergenic |
| chr1 | 69703904 | 69704189 | Distal Intergenic |
| chr1 | 69704373 | 69705506 | Distal Intergenic |
| chr1 | 69732905 | 69733505 | Distal Intergenic |
| chr1 | 69734407 | 69734711 | Distal Intergenic |
| chr1 | 70124991 | 70125591 | Intron |
| chr1 | 70181590 | 70181833 | Intron |
| chr1 | 70182142 | 70182759 | Intron |
| chr1 | 70965114 | 70965328 | Distal Intergenic |
| chr1 | 71046523 | 71047317 | Distal Intergenic |
| chr1 | 71049940 | 71050217 | Distal Intergenic |
| chr1 | 71050433 | 71051898 | Distal Intergenic |
| chr1 | 71142075 | 71142479 | Distal Intergenic |
| chr1 | 71143149 | 71143366 | Distal Intergenic |
| chr1 | 71148730 | 71149017 | Distal Intergenic |
| chr1 | 71149109 | 71149530 | Distal Intergenic |
| chr1 | 71149663 | 71150130 | Distal Intergenic |
| chr1 | 71151722 | 71153281 | Distal Intergenic |
| chr1 | 71381810 | 71382344 | Intron |

| | | | |
|---|---|---|---|
| chr1 | 71385197 | 71386492 | Intron |
| chr1 | 71416524 | 71416723 | Intron |
| chr1 | 71416821 | 71417048 | Intron |
| chr1 | 71442390 | 71445252 | Intron |
| chr1 | 71459353 | 71460939 | Intron |
| chr1 | 71461107 | 71461472 | Intron |
| chr1 | 71461517 | 71461980 | Intron |
| chr1 | 71504207 | 71508590 | Intron |
| chr1 | 71524905 | 71526049 | Intron |
| chr1 | 71603628 | 71603867 | Intron |
| chr1 | 71604158 | 71604390 | Intron |
| chr1 | 71713333 | 71713584 | Distal Intergenic |
| chr1 | 71713790 | 71714933 | Distal Intergenic |
| chr1 | 71737890 | 71738281 | Distal Intergenic |
| chr1 | 71738398 | 71738718 | Distal Intergenic |
| chr1 | 71801598 | 71802674 | Distal Intergenic |
| chr1 | 71927010 | 71927263 | Intron |
| chr1 | 71927311 | 71927638 | Intron |
| chr1 | 71940419 | 71940775 | Intron |
| chr1 | 72116902 | 72117403 | Intron |
| chr1 | 72232722 | 72233033 | Intron |
| chr1 | 72239593 | 72239851 | Intron |
| chr1 | 72240068 | 72241223 | Intron |
| chr1 | 72248548 | 72250555 | Intron |
| chr1 | 72251692 | 72252000 | Intron |
| chr1 | 72257426 | 72258607 | Intron |
| chr1 | 72273090 | 72273852 | Intron |
| chr1 | 72284020 | 72285274 | Intron |
| chr1 | 72289478 | 72290434 | Intron |
| chr1 | 72290541 | 72291691 | Intron |
| chr1 | 72301806 | 72302415 | Promoter |
| chr1 | 72353626 | 72353898 | Intron |
| chr1 | 72630118 | 72630357 | Intron |
| chr1 | 72758157 | 72758872 | Distal Intergenic |
| chr1 | 72779153 | 72781180 | Distal Intergenic |
| chr1 | 72782009 | 72783637 | Distal Intergenic |
| chr1 | 72785420 | 72786209 | Distal Intergenic |
| chr1 | 72794071 | 72794910 | Distal Intergenic |
| chr1 | 72795015 | 72795840 | Distal Intergenic |
| chr1 | 72799639 | 72800660 | Distal Intergenic |
| chr1 | 72801630 | 72802879 | Distal Intergenic |
| chr1 | 72824403 | 72828455 | Distal Intergenic |
| chr1 | 72828715 | 72829683 | Distal Intergenic |
| chr1 | 72829856 | 72830982 | Distal Intergenic |

| | | | |
|---|---|---|---|
| chr1 | 72840181 | 72840380 | Distal Intergenic |
| chr1 | 72867102 | 72868354 | Distal Intergenic |
| chr1 | 72871921 | 72872863 | Distal Intergenic |
| chr1 | 72874724 | 72874923 | Distal Intergenic |
| chr1 | 72894567 | 72894778 | Distal Intergenic |
| chr1 | 73517922 | 73518831 | Distal Intergenic |
| chr1 | 73550240 | 73551809 | Distal Intergenic |
| chr1 | 73553314 | 73553990 | Distal Intergenic |
| chr1 | 73554161 | 73554905 | Distal Intergenic |
| chr1 | 74002082 | 74002517 | Distal Intergenic |
| chr1 | 74055458 | 74055739 | Distal Intergenic |
| chr1 | 74195535 | 74195957 | Distal Intergenic |
| chr1 | 74196090 | 74196302 | Distal Intergenic |
| chr1 | 74283137 | 74286558 | Distal Intergenic |
| chr1 | 74286652 | 74287208 | Distal Intergenic |
| chr1 | 74294746 | 74294987 | Distal Intergenic |
| chr1 | 74295228 | 74295795 | Distal Intergenic |
| chr1 | 74330983 | 74331336 | Distal Intergenic |
| chr1 | 74341559 | 74346313 | Distal Intergenic |
| chr1 | 74346812 | 74347397 | Distal Intergenic |
| chr1 | 74347538 | 74347766 | Distal Intergenic |
| chr1 | 74348990 | 74349667 | Distal Intergenic |
| chr1 | 74349862 | 74350713 | Distal Intergenic |
| chr1 | 74350751 | 74351174 | Distal Intergenic |
| chr1 | 74433266 | 74433889 | Distal Intergenic |
| chr1 | 74443219 | 74443685 | Distal Intergenic |
| chr1 | 74443848 | 74444389 | Distal Intergenic |
| chr1 | 74552607 | 74554050 | Intron |
| chr1 | 74634383 | 74634658 | Intron |
| chr1 | 74647422 | 74647954 | Intron |
| chr1 | 74772682 | 74773092 | Intron |
| chr1 | 74773182 | 74773764 | Intron |
| chr1 | 74838585 | 74841982 | Intron |
| chr1 | 74842077 | 74842405 | Intron |
| chr1 | 74850306 | 74853637 | Intron |
| chr1 | 75121200 | 75121753 | Intron |
| chr1 | 75138413 | 75138690 | Promoter |
| chr1 | 75163867 | 75165165 | Distal Intergenic |
| chr1 | 75189055 | 75189493 | Intron |
| chr1 | 75207179 | 75207588 | Intron |
| chr1 | 75210807 | 75211345 | Intron |
| chr1 | 75211399 | 75212202 | Intron |
| chr1 | 75214843 | 75215839 | Intron |
| chr1 | 75226961 | 75229230 | Intron |

| | | | |
|---|---|---|---|
| chr1 | 75231412 | 75231846 | 3' UTR |
| chr1 | 75232480 | 75232894 | Distal Intergenic |
| chr1 | 75233006 | 75233348 | Distal Intergenic |
| chr1 | 75233430 | 75233750 | Distal Intergenic |
| chr1 | 75244877 | 75245098 | Distal Intergenic |
| chr1 | 75245229 | 75245449 | Distal Intergenic |
| chr1 | 75289307 | 75290343 | Distal Intergenic |
| chr1 | 75313448 | 75313849 | Distal Intergenic |
| chr1 | 75432532 | 75432733 | Distal Intergenic |
| chr1 | 75517716 | 75518297 | Distal Intergenic |
| chr1 | 75584489 | 75585407 | Distal Intergenic |
| chr1 | 75585508 | 75585872 | Distal Intergenic |
| chr1 | 76020896 | 76022101 | Intron |
| chr1 | 76163774 | 76164221 | Intron |
| chr1 | 76293111 | 76293647 | Intron |
| chr1 | 76638137 | 76638394 | Intron |
| chr1 | 76641631 | 76641851 | Intron |
| chr1 | 76641917 | 76643467 | Intron |
| chr1 | 76648259 | 76648554 | Intron |
| chr1 | 76660593 | 76660796 | Intron |
| chr1 | 76714197 | 76714647 | Intron |
| chr1 | 76714752 | 76715016 | Intron |
| chr1 | 76715057 | 76715986 | Intron |
| chr1 | 76831643 | 76831867 | Intron |
| chr1 | 76869525 | 76871398 | Intron |
| chr1 | 76876712 | 76877559 | Intron |
| chr1 | 76923554 | 76924990 | Intron |
| chr1 | 77004015 | 77005120 | Intron |
| chr1 | 77077469 | 77078046 | Intron |
| chr1 | 77457081 | 77460094 | Intron |
| chr1 | 77461705 | 77461914 | Intron |
| chr1 | 77462036 | 77462281 | Intron |
| chr1 | 77462377 | 77462634 | Intron |
| chr1 | 77463988 | 77464467 | Intron |
| chr1 | 77467206 | 77468369 | Intron |
| chr1 | 77691640 | 77691878 | Distal Intergenic |
| chr1 | 77692088 | 77692545 | Distal Intergenic |
| chr1 | 77735556 | 77736129 | Distal Intergenic |
| chr1 | 77736168 | 77736446 | Distal Intergenic |
| chr1 | 77753923 | 77755285 | Intron |
| chr1 | 77836900 | 77837884 | Intron |
| chr1 | 77842060 | 77843490 | Intron |
| chr1 | 78485835 | 78486143 | Distal Intergenic |
| chr1 | 78529313 | 78529665 | Intron |

| | | | |
|---|---|---|---|
| chr1 | 78533957 | 78534306 | Intron |
| chr1 | 78534396 | 78535277 | Intron |
| chr1 | 78587905 | 78588619 | Intron |
| chr1 | 78588842 | 78589181 | Intron |
| chr1 | 78589617 | 78589821 | Intron |
| chr1 | 78640551 | 78641534 | Distal Intergenic |
| chr1 | 78652724 | 78653884 | Distal Intergenic |
| chr1 | 78654415 | 78654828 | Distal Intergenic |
| chr1 | 78654904 | 78656247 | Distal Intergenic |
| chr1 | 78677138 | 78677529 | Distal Intergenic |
| chr1 | 79140902 | 79141375 | Distal Intergenic |
| chr1 | 79173512 | 79173997 | Distal Intergenic |
| chr1 | 79577749 | 79578080 | Distal Intergenic |
| chr1 | 79687827 | 79688548 | Distal Intergenic |
| chr1 | 79742343 | 79743252 | Distal Intergenic |
| chr1 | 79757245 | 79757444 | Distal Intergenic |
| chr1 | 79826688 | 79827120 | Distal Intergenic |
| chr1 | 79827158 | 79827363 | Distal Intergenic |
| chr1 | 80022015 | 80022499 | Distal Intergenic |
| chr1 | 80217081 | 80218321 | Distal Intergenic |
| chr1 | 80335899 | 80336686 | Distal Intergenic |
| chr1 | 81363027 | 81364252 | Distal Intergenic |
| chr1 | 81365654 | 81366124 | Distal Intergenic |
| chr1 | 81620621 | 81620845 | Distal Intergenic |
| chr1 | 81620898 | 81621158 | Distal Intergenic |
| chr1 | 81626879 | 81627316 | Distal Intergenic |
| chr1 | 81667335 | 81668532 | Distal Intergenic |
| chr1 | 81673586 | 81675146 | Distal Intergenic |
| chr1 | 81755024 | 81757157 | Distal Intergenic |
| chr1 | 81910207 | 81910421 | Intron |
| chr1 | 82028787 | 82029131 | Intron |
| chr1 | 82152594 | 82153308 | Intron |
| chr1 | 82313199 | 82313416 | Intron |
| chr1 | 82368544 | 82368971 | Intron |
| chr1 | 82425881 | 82426131 | Intron |
| chr1 | 82593633 | 82596198 | Distal Intergenic |
| chr1 | 82596252 | 82596556 | Distal Intergenic |
| chr1 | 82721497 | 82721746 | Distal Intergenic |
| chr1 | 82789503 | 82790356 | Distal Intergenic |
| chr1 | 82791543 | 82793069 | Distal Intergenic |
| chr1 | 82828187 | 82828579 | Distal Intergenic |
| chr1 | 82836060 | 82836300 | Distal Intergenic |
| chr1 | 82875709 | 82875932 | Distal Intergenic |
| chr1 | 82905092 | 82905907 | Distal Intergenic |

| | | | |
|---|---|---|---|
| chr1 | 82943440 | 82944791 | Distal Intergenic |
| chr1 | 82951418 | 82951799 | Distal Intergenic |
| chr1 | 84403198 | 84403935 | Exon |
| chr1 | 84769234 | 84769775 | Promoter |
| chr1 | 84834611 | 84835441 | Intron |
| chr1 | 84836249 | 84837359 | Intron |
| chr1 | 84877297 | 84879013 | 5' UTR |
| chr1 | 84935584 | 84937314 | Distal Intergenic |
| chr1 | 84945382 | 84946913 | Promoter |
| chr1 | 87690524 | 87691542 | Distal Intergenic |
| chr1 | 87791765 | 87794049 | Promoter |
| chr1 | 87794839 | 87795218 | Promoter |
| chr1 | 87795330 | 87795669 | Intron |
| chr1 | 87796558 | 87796829 | Promoter |
| chr1 | 87796998 | 87797352 | Promoter |
| chr1 | 87961000 | 87962153 | Distal Intergenic |
| chr1 | 88042708 | 88043526 | Distal Intergenic |
| chr1 | 88044121 | 88044324 | Distal Intergenic |
| chr1 | 88046799 | 88048941 | Distal Intergenic |
| chr1 | 88049010 | 88049226 | Distal Intergenic |
| chr1 | 88050451 | 88051309 | Distal Intergenic |
| chr1 | 88140257 | 88140805 | Distal Intergenic |
| chr1 | 88255598 | 88256067 | Distal Intergenic |
| chr1 | 88256163 | 88256781 | Distal Intergenic |
| chr1 | 88272372 | 88273608 | Distal Intergenic |
| chr1 | 88331157 | 88331685 | Distal Intergenic |
| chr1 | 88334093 | 88337218 | Distal Intergenic |
| chr1 | 88366902 | 88369704 | Distal Intergenic |
| chr1 | 88412317 | 88412893 | Distal Intergenic |
| chr1 | 88418849 | 88419735 | Distal Intergenic |
| chr1 | 88424303 | 88424557 | Distal Intergenic |
| chr1 | 88424761 | 88425047 | Distal Intergenic |
| chr1 | 88425233 | 88425546 | Distal Intergenic |
| chr1 | 88427658 | 88427880 | Distal Intergenic |
| chr1 | 88428214 | 88429083 | Distal Intergenic |
| chr1 | 88518826 | 88520763 | Distal Intergenic |
| chr1 | 88576398 | 88576650 | Distal Intergenic |
| chr1 | 88576745 | 88577154 | Distal Intergenic |
| chr1 | 88920877 | 88921181 | Distal Intergenic |
| chr1 | 88921218 | 88921570 | Distal Intergenic |
| chr1 | 88925528 | 88925933 | Distal Intergenic |
| chr1 | 88932252 | 88932675 | Distal Intergenic |
| chr1 | 88933715 | 88935766 | Distal Intergenic |
| chr1 | 88936522 | 88937839 | Distal Intergenic |

| | | | |
|---|---|---|---|
| chr1 | 89077227 | 89078070 | Intron |
| chr1 | 89110234 | 89110466 | Intron |
| chr1 | 89111057 | 89111470 | Intron |
| chr1 | 89149364 | 89149651 | Promoter |
| chr1 | 89151892 | 89152095 | Intron |
| chr1 | 89224026 | 89224448 | Intron |
| chr1 | 89306671 | 89307922 | Distal Intergenic |
| chr1 | 89308746 | 89309559 | Distal Intergenic |
| chr1 | 89391271 | 89391482 | Distal Intergenic |
| chr1 | 89445547 | 89445770 | 3' UTR |
| chr1 | 89446925 | 89448251 | 3' UTR |
| chr1 | 89455846 | 89456148 | Intron |
| chr1 | 89456198 | 89457195 | Intron |
| chr1 | 89778230 | 89778768 | Distal Intergenic |
| chr1 | 89867084 | 89867513 | Distal Intergenic |
| chr1 | 89990879 | 89991931 | Promoter |
| chr1 | 89993521 | 89995130 | Intron |
| chr1 | 89998919 | 90000654 | Intron |
| chr1 | 90005547 | 90005950 | Intron |
| chr1 | 90025701 | 90025970 | Intron |
| chr1 | 90026238 | 90026563 | Intron |
| chr1 | 90029946 | 90032618 | Intron |
| chr1 | 90036903 | 90037780 | Intron |
| chr1 | 90049478 | 90050898 | Exon |
| chr1 | 90134307 | 90134646 | Intron |
| chr1 | 90158818 | 90159506 | Intron |
| chr1 | 90173645 | 90174254 | Intron |
| chr1 | 90257236 | 90259351 | Distal Intergenic |
| chr1 | 90276785 | 90278403 | Distal Intergenic |
| chr1 | 90318882 | 90320164 | Intron |
| chr1 | 90329675 | 90330240 | Intron |
| chr1 | 90333400 | 90334995 | Intron |
| chr1 | 90339722 | 90340207 | Intron |
| chr1 | 90370412 | 90372845 | Intron |
| chr1 | 90402494 | 90402806 | Distal Intergenic |
| chr1 | 90404299 | 90405839 | Distal Intergenic |
| chr1 | 90421780 | 90422404 | Distal Intergenic |
| chr1 | 90422546 | 90423779 | Distal Intergenic |
| chr1 | 90430117 | 90431800 | Distal Intergenic |
| chr1 | 90432902 | 90434466 | Distal Intergenic |
| chr1 | 90439211 | 90440693 | Distal Intergenic |
| chr1 | 90453066 | 90453521 | Exon |
| chr1 | 90483474 | 90484497 | Exon |
| chr1 | 90535809 | 90536626 | Distal Intergenic |

| | | | |
|---|---|---|---|
| chr1 | 90576819 | 90577595 | Distal Intergenic |
| chr1 | 90595636 | 90597164 | Distal Intergenic |
| chr1 | 90597231 | 90598204 | Distal Intergenic |
| chr1 | 90598309 | 90598601 | Distal Intergenic |
| chr1 | 90598953 | 90599161 | Distal Intergenic |
| chr1 | 90765611 | 90766813 | Distal Intergenic |
| chr1 | 90813666 | 90814020 | Distal Intergenic |
| chr1 | 90814394 | 90814600 | Distal Intergenic |
| chr1 | 90911722 | 90912911 | Distal Intergenic |
| chr1 | 90915980 | 90917158 | Distal Intergenic |
| chr1 | 90932892 | 90933291 | Distal Intergenic |
| chr1 | 90965262 | 90966712 | Distal Intergenic |
| chr1 | 90967003 | 90967202 | Distal Intergenic |
| chr1 | 90997826 | 90998182 | Distal Intergenic |
| chr1 | 90998264 | 90999228 | Distal Intergenic |
| chr1 | 91017868 | 91021134 | Distal Intergenic |
| chr1 | 91027814 | 91028617 | Distal Intergenic |
| chr1 | 91029874 | 91030114 | Distal Intergenic |
| chr1 | 91370707 | 91372007 | Distal Intergenic |
| chr1 | 91409859 | 91411356 | Intron |
| chr1 | 91471335 | 91471682 | Intron |
| chr1 | 91473434 | 91473797 | Intron |
| chr1 | 91499454 | 91499690 | Distal Intergenic |
| chr1 | 91544301 | 91545281 | Distal Intergenic |
| chr1 | 91552498 | 91554338 | Distal Intergenic |
| chr1 | 91582741 | 91584192 | Distal Intergenic |
| chr1 | 91593011 | 91593359 | Distal Intergenic |
| chr1 | 91593418 | 91594453 | Distal Intergenic |
| chr1 | 91597003 | 91597202 | Distal Intergenic |
| chr1 | 91597482 | 91598087 | Distal Intergenic |
| chr1 | 91626566 | 91627508 | Distal Intergenic |
| chr1 | 91701393 | 91702229 | Distal Intergenic |
| chr1 | 91947547 | 91948089 | Distal Intergenic |
| chr1 | 91993857 | 91994618 | Distal Intergenic |
| chr1 | 91996500 | 91997304 | Distal Intergenic |
| chr1 | 92060347 | 92061698 | Distal Intergenic |
| chr1 | 92079949 | 92081063 | Distal Intergenic |
| chr1 | 92081323 | 92081699 | Distal Intergenic |
| chr1 | 92082366 | 92083227 | Distal Intergenic |
| chr1 | 92100581 | 92102667 | Distal Intergenic |
| chr1 | 92103155 | 92103370 | Distal Intergenic |
| chr1 | 92118011 | 92118847 | Distal Intergenic |
| chr1 | 92120609 | 92124407 | Distal Intergenic |
| chr1 | 92126995 | 92127580 | Distal Intergenic |

| | | | |
|---|---|---|---|
| chr1 | 92130567 | 92131303 | Distal Intergenic |
| chr1 | 92131369 | 92131653 | Distal Intergenic |
| chr1 | 92131699 | 92131957 | Distal Intergenic |
| chr1 | 92155078 | 92156196 | Intron |
| chr1 | 92312593 | 92314044 | Intron |
| chr1 | 92336328 | 92338114 | Intron |
| chr1 | 92574077 | 92574655 | Intron |
| chr1 | 92919889 | 92920616 | Distal Intergenic |
| chr1 | 92920747 | 92921090 | Distal Intergenic |
| chr1 | 92950017 | 92951076 | Promoter |
| chr1 | 92978051 | 92978865 | 3' UTR |
| chr1 | 92991410 | 92991925 | Intron |
| chr1 | 92991988 | 92994198 | Intron |
| chr1 | 93023427 | 93023827 | Intron |
| chr1 | 93144384 | 93145762 | Intron |
| chr1 | 93146401 | 93146874 | Intron |
| chr1 | 93147267 | 93147885 | Intron |
| chr1 | 93159552 | 93159992 | Promoter |
| chr1 | 93160080 | 93160285 | Promoter |
| chr1 | 93168729 | 93169449 | Exon |
| chr1 | 93174837 | 93176159 | Intron |
| chr1 | 93294958 | 93295495 | Distal Intergenic |
| chr1 | 93296856 | 93298473 | Promoter |
| chr1 | 93386725 | 93387097 | Intron |
| chr1 | 93537138 | 93537738 | Distal Intergenic |
| chr1 | 93695848 | 93697569 | Intron |
| chr1 | 93956052 | 93956713 | Intron |
| chr1 | 93982329 | 93985322 | Intron |
| chr1 | 93995806 | 93996679 | Exon |
| chr1 | 94002755 | 94003033 | Intron |
| chr1 | 94003204 | 94003690 | Intron |
| chr1 | 94008046 | 94008493 | Intron |
| chr1 | 94008671 | 94008871 | Promoter |
| chr1 | 94009232 | 94009458 | Promoter |
| chr1 | 94015951 | 94016682 | Exon |
| chr1 | 94026019 | 94027600 | 3' UTR |
| chr1 | 94027695 | 94028452 | 3' UTR |
| chr1 | 94028505 | 94028704 | Intron |
| chr1 | 94040535 | 94042416 | Exon |
| chr1 | 94058548 | 94059709 | Intron |
| chr1 | 94062794 | 94063556 | Intron |
| chr1 | 94063776 | 94064740 | Exon |
| chr1 | 94075919 | 94076199 | Intron |
| chr1 | 94077225 | 94077758 | Intron |

| | | | |
|---|---|---|---|
| chr1 | 94077850 | 94078162 | Intron |
| chr1 | 94078226 | 94078675 | Intron |
| chr1 | 94078895 | 94081087 | Promoter |
| chr1 | 94369808 | 94370354 | Exon |
| chr1 | 94579573 | 94580538 | Intron |
| chr1 | 94641834 | 94642036 | Intron |
| chr1 | 94695920 | 94696148 | Intron |
| chr1 | 94696485 | 94697241 | 5' UTR |
| chr1 | 94712200 | 94712643 | Intron |
| chr1 | 94799920 | 94801441 | Distal Intergenic |
| chr1 | 94808177 | 94808387 | Distal Intergenic |
| chr1 | 94827088 | 94827599 | Distal Intergenic |
| chr1 | 94830417 | 94831383 | Distal Intergenic |
| chr1 | 94868650 | 94869451 | Distal Intergenic |
| chr1 | 94875216 | 94875635 | Distal Intergenic |
| chr1 | 95026950 | 95027381 | Distal Intergenic |
| chr1 | 95027491 | 95028081 | Distal Intergenic |
| chr1 | 95056819 | 95057563 | Distal Intergenic |
| chr1 | 95084890 | 95085180 | Distal Intergenic |
| chr1 | 95093490 | 95095062 | Distal Intergenic |
| chr1 | 95196078 | 95197330 | Intron |
| chr1 | 95341129 | 95342717 | Intron |
| chr1 | 95374162 | 95374416 | Intron |
| chr1 | 95374454 | 95374976 | Intron |
| chr1 | 95375133 | 95375434 | Intron |
| chr1 | 95375512 | 95376812 | Intron |
| chr1 | 95425377 | 95427275 | Exon |
| chr1 | 95576830 | 95577176 | Intron |
| chr1 | 95601604 | 95603061 | Intron |
| chr1 | 95645142 | 95645556 | Intron |
| chr1 | 95650343 | 95650579 | Intron |
| chr1 | 95721839 | 95722092 | Distal Intergenic |
| chr1 | 96045894 | 96046939 | Distal Intergenic |
| chr1 | 97364776 | 97365080 | Distal Intergenic |
| chr1 | 99666899 | 99667154 | Distal Intergenic |
| chr1 | 99807225 | 99808488 | Distal Intergenic |
| chr1 | 99860604 | 99861016 | Distal Intergenic |
| chr1 | 102045693 | 102047687 | Distal Intergenic |
| chr1 | 102067596 | 102069409 | Distal Intergenic |
| chr1 | 102069640 | 102070311 | Distal Intergenic |
| chr1 | 103029753 | 103031568 | Distal Intergenic |
| chr1 | 103689359 | 103690374 | Distal Intergenic |
| chr1 | 107186441 | 107188808 | Distal Intergenic |
| chr1 | 107249753 | 107250378 | Distal Intergenic |

| | | | |
|---|---|---|---|
| chr1 | 107250448 | 107250782 | Distal Intergenic |
| chr1 | 107327261 | 107327734 | Distal Intergenic |
| chr1 | 107328357 | 107328945 | Distal Intergenic |
| chr1 | 107339075 | 107339323 | Distal Intergenic |
| chr1 | 107383390 | 107384548 | Distal Intergenic |
| chr1 | 107559432 | 107560285 | Distal Intergenic |
| chr1 | 107762286 | 107762558 | Intron |
| chr1 | 107762596 | 107764008 | Intron |
| chr1 | 107888775 | 107889887 | Intron |
| chr1 | 107889976 | 107890202 | Intron |
| chr1 | 107954876 | 107955472 | Intron |
| chr1 | 108059138 | 108059376 | Distal Intergenic |
| chr1 | 108107501 | 108108327 | Distal Intergenic |
| chr1 | 108137500 | 108139110 | Exon |
| chr1 | 108164397 | 108165282 | Intron |
| chr1 | 108303568 | 108304633 | Intron |
| chr1 | 108304684 | 108305107 | Intron |
| chr1 | 108357420 | 108358629 | Intron |
| chr1 | 108498967 | 108499290 | Intron |
| chr1 | 108656402 | 108657003 | Distal Intergenic |
| chr1 | 108661794 | 108663718 | Distal Intergenic |
| chr1 | 108692168 | 108693891 | Intron |
| chr1 | 108718510 | 108719446 | Intron |
| chr1 | 108719506 | 108719875 | Intron |
| chr1 | 108753608 | 108754040 | Distal Intergenic |
| chr1 | 108754104 | 108755330 | Distal Intergenic |
| chr1 | 108756642 | 108760396 | Distal Intergenic |
| chr1 | 108761520 | 108762824 | Distal Intergenic |
| chr1 | 109399310 | 109399509 | Promoter |
| chr1 | 109401810 | 109403701 | Promoter |
| chr1 | 109424724 | 109428211 | 5' UTR |
| chr1 | 109434814 | 109436153 | Intron |
| chr1 | 109486589 | 109487882 | Exon |
| chr1 | 109487921 | 109488173 | Intron |
| chr1 | 109942971 | 109943180 | 3' UTR |
| chr1 | 113761047 | 113761275 | Distal Intergenic |
| chr1 | 116337958 | 116339501 | Distal Intergenic |
| chr1 | 116339541 | 116339825 | Distal Intergenic |
| chr1 | 120207376 | 120208017 | Distal Intergenic |
| chr1 | 120265011 | 120266089 | 5' UTR |
| chr1 | 120276896 | 120277236 | Intron |
| chr1 | 120283697 | 120285900 | Exon |
| chr1 | 120286524 | 120286835 | 3' UTR |
| chr1 | 120321040 | 120321769 | Distal Intergenic |

| | | | |
|---|---|---|---|
| chr1 | 120321853 | 120322080 | Distal Intergenic |
| chr1 | 120355054 | 120355263 | Promoter |
| chr1 | 120355393 | 120356050 | Distal Intergenic |
| chr1 | 120435374 | 120435640 | Downstream |
| chr1 | 120704114 | 120704814 | Distal Intergenic |
| chr1 | 120721428 | 120722968 | Distal Intergenic |
| chr1 | 120748554 | 120749095 | Distal Intergenic |
| chr1 | 120769886 | 120771066 | Distal Intergenic |
| chr1 | 120784156 | 120784364 | Distal Intergenic |
| chr1 | 120849592 | 120850022 | Intron |
| chr1 | 120872866 | 120873532 | Distal Intergenic |
| chr1 | 120961582 | 120961952 | Distal Intergenic |
| chr1 | 121021336 | 121021756 | Distal Intergenic |
| chr1 | 121021794 | 121022583 | Distal Intergenic |
| chr1 | 121063713 | 121064049 | Distal Intergenic |
| chr1 | 121064105 | 121064500 | Distal Intergenic |
| chr1 | 121121426 | 121121868 | Intron |
| chr1 | 121196892 | 121197282 | Distal Intergenic |
| chr1 | 121400348 | 121400560 | Distal Intergenic |
| chr1 | 121400600 | 121400861 | Distal Intergenic |
| chr1 | 121433155 | 121435298 | Distal Intergenic |
| chr1 | 121979570 | 121980062 | Distal Intergenic |
| chr1 | 121980166 | 121980807 | Distal Intergenic |
| chr1 | 122016667 | 122017372 | Distal Intergenic |
| chr1 | 122032626 | 122032962 | Distal Intergenic |
| chr1 | 122033015 | 122033345 | Distal Intergenic |
| chr1 | 122636444 | 122636824 | Distal Intergenic |
| chr1 | 122640546 | 122640759 | Distal Intergenic |
| chr1 | 122653252 | 122655957 | Distal Intergenic |
| chr1 | 122880350 | 122880796 | Distal Intergenic |
| chr1 | 122931600 | 122931915 | Distal Intergenic |
| chr1 | 123257317 | 123257709 | Distal Intergenic |
| chr1 | 123273723 | 123274490 | Distal Intergenic |
| chr1 | 123274535 | 123274880 | Distal Intergenic |
| chr1 | 123454876 | 123455212 | Distal Intergenic |
| chr1 | 123550428 | 123550777 | Distal Intergenic |
| chr1 | 125743573 | 125743960 | Distal Intergenic |
| chr1 | 125886712 | 125886993 | Distal Intergenic |
| chr1 | 126383804 | 126385919 | Distal Intergenic |
| chr1 | 127330431 | 127332094 | Distal Intergenic |
| chr1 | 127346058 | 127348028 | Distal Intergenic |
| chr1 | 127378799 | 127379282 | Distal Intergenic |
| chr1 | 127379588 | 127380277 | Distal Intergenic |
| chr1 | 127422316 | 127422528 | Distal Intergenic |

| | | | |
|---|---|---|---|
| chr1 | 127422564 | 127423146 | Distal Intergenic |
| chr1 | 127446350 | 127447274 | Distal Intergenic |
| chr1 | 127456586 | 127456916 | Distal Intergenic |
| chr1 | 127464386 | 127465273 | Distal Intergenic |
| chr1 | 127597727 | 127598254 | Distal Intergenic |
| chr1 | 127598418 | 127599060 | Distal Intergenic |
| chr1 | 128489428 | 128490462 | Distal Intergenic |
| chr1 | 128498730 | 128499261 | Distal Intergenic |
| chr1 | 128505292 | 128506137 | Distal Intergenic |
| chr1 | 128514990 | 128516668 | Distal Intergenic |
| chr1 | 128555031 | 128555947 | Distal Intergenic |
| chr1 | 128567114 | 128567655 | Distal Intergenic |
| chr1 | 128575882 | 128577030 | Distal Intergenic |
| chr1 | 128580117 | 128581624 | Distal Intergenic |
| chr1 | 128634014 | 128634224 | Distal Intergenic |
| chr1 | 128713776 | 128714929 | Distal Intergenic |
| chr1 | 128897701 | 128898934 | Distal Intergenic |
| chr1 | 128899002 | 128899759 | Distal Intergenic |
| chr1 | 129115751 | 129116035 | Distal Intergenic |
| chr1 | 129116082 | 129116688 | Distal Intergenic |
| chr1 | 129129739 | 129130182 | Distal Intergenic |
| chr1 | 129153856 | 129154238 | Distal Intergenic |
| chr1 | 129670806 | 129671576 | Distal Intergenic |
| chr1 | 129757212 | 129757428 | Distal Intergenic |
| chr1 | 129879626 | 129879842 | Distal Intergenic |
| chr1 | 129880024 | 129880708 | Distal Intergenic |
| chr1 | 129887932 | 129888572 | Distal Intergenic |
| chr1 | 130140925 | 130141841 | Distal Intergenic |
| chr1 | 130253966 | 130254637 | Distal Intergenic |
| chr1 | 131547992 | 131548432 | Distal Intergenic |
| chr1 | 132587389 | 132587613 | Distal Intergenic |
| chr1 | 132612715 | 132613776 | Distal Intergenic |
| chr1 | 132614134 | 132614451 | Distal Intergenic |
| chr1 | 132628270 | 132628538 | Distal Intergenic |
| chr1 | 132629225 | 132629494 | Distal Intergenic |
| chr1 | 132629548 | 132632104 | Distal Intergenic |
| chr1 | 132637644 | 132637880 | Distal Intergenic |
| chr1 | 132643526 | 132644124 | Distal Intergenic |
| chr1 | 132666292 | 132668654 | Distal Intergenic |
| chr1 | 132699468 | 132700065 | Distal Intergenic |
| chr1 | 132700178 | 132700692 | Distal Intergenic |
| chr1 | 132776287 | 132777868 | Distal Intergenic |
| chr1 | 132777965 | 132778323 | Distal Intergenic |
| chr1 | 132782807 | 132784056 | Distal Intergenic |

| | | | |
|---|---|---|---|
| chr1 | 132788670 | 132790036 | Distal Intergenic |
| chr1 | 132814791 | 132816052 | Distal Intergenic |
| chr1 | 132838981 | 132841376 | Distal Intergenic |
| chr1 | 132843591 | 132844879 | Distal Intergenic |
| chr1 | 132906836 | 132907690 | Distal Intergenic |
| chr1 | 132960682 | 132962037 | Distal Intergenic |
| chr1 | 132993530 | 132993782 | Distal Intergenic |
| chr1 | 133020171 | 133020542 | Distal Intergenic |
| chr1 | 133020711 | 133021204 | Distal Intergenic |
| chr1 | 133022677 | 133023363 | Distal Intergenic |
| chr1 | 133031969 | 133032241 | Distal Intergenic |
| chr1 | 133035003 | 133036417 | Distal Intergenic |
| chr1 | 133077144 | 133077441 | Distal Intergenic |
| chr1 | 133081454 | 133081752 | Distal Intergenic |
| chr1 | 133117700 | 133118976 | Distal Intergenic |
| chr1 | 133134712 | 133136991 | Distal Intergenic |
| chr1 | 133137042 | 133137333 | Distal Intergenic |
| chr1 | 133141981 | 133142277 | Distal Intergenic |
| chr1 | 133175614 | 133177147 | Distal Intergenic |
| chr1 | 133190169 | 133191122 | Distal Intergenic |
| chr1 | 133229796 | 133230225 | Distal Intergenic |
| chr1 | 133397754 | 133399984 | Distal Intergenic |
| chr1 | 133401807 | 133402364 | Distal Intergenic |
| chr1 | 133402502 | 133404257 | Distal Intergenic |
| chr1 | 133405393 | 133405771 | Distal Intergenic |
| chr1 | 133410211 | 133410628 | Distal Intergenic |
| chr1 | 133422950 | 133423400 | Distal Intergenic |
| chr1 | 133586101 | 133588245 | Distal Intergenic |
| chr1 | 133807081 | 133807697 | Distal Intergenic |
| chr1 | 133890265 | 133890561 | Distal Intergenic |
| chr1 | 133898580 | 133899093 | Distal Intergenic |
| chr1 | 133901209 | 133901408 | Distal Intergenic |
| chr1 | 133903824 | 133904110 | Distal Intergenic |
| chr1 | 133904219 | 133905233 | Distal Intergenic |
| chr1 | 133907058 | 133907419 | Distal Intergenic |
| chr1 | 134035352 | 134035553 | Distal Intergenic |
| chr1 | 134035820 | 134036030 | Distal Intergenic |
| chr1 | 134046923 | 134048858 | Distal Intergenic |
| chr1 | 134092035 | 134092483 | Distal Intergenic |
| chr1 | 134092962 | 134093186 | Distal Intergenic |
| chr1 | 134151593 | 134151844 | Distal Intergenic |
| chr1 | 134151914 | 134152221 | Distal Intergenic |
| chr1 | 134161001 | 134162310 | Distal Intergenic |
| chr1 | 134177057 | 134177878 | Distal Intergenic |

| | | | |
|---|---|---|---|
| chr1 | 134183436 | 134184171 | Distal Intergenic |
| chr1 | 134187769 | 134189583 | Distal Intergenic |
| chr1 | 134191466 | 134192862 | Distal Intergenic |
| chr1 | 134212663 | 134212887 | Distal Intergenic |
| chr1 | 134213463 | 134216211 | Distal Intergenic |
| chr1 | 134231400 | 134232732 | Distal Intergenic |
| chr1 | 134243038 | 134243721 | Distal Intergenic |
| chr1 | 134249607 | 134251376 | Distal Intergenic |
| chr1 | 134314187 | 134314471 | Distal Intergenic |
| chr1 | 134314525 | 134314811 | Distal Intergenic |
| chr1 | 134327864 | 134330438 | Distal Intergenic |
| chr1 | 134462363 | 134465906 | Distal Intergenic |
| chr1 | 134467428 | 134467802 | Distal Intergenic |
| chr1 | 134467913 | 134468112 | Distal Intergenic |
| chr1 | 134915384 | 134915592 | Distal Intergenic |
| chr1 | 134915649 | 134918027 | Distal Intergenic |
| chr1 | 134920836 | 134921207 | Distal Intergenic |
| chr1 | 134921483 | 134922352 | Distal Intergenic |
| chr1 | 135027246 | 135029120 | Distal Intergenic |
| chr1 | 135029267 | 135029492 | Distal Intergenic |
| chr1 | 135029868 | 135030067 | Distal Intergenic |
| chr1 | 135128169 | 135128522 | Distal Intergenic |
| chr1 | 135128620 | 135129090 | Distal Intergenic |
| chr1 | 135173405 | 135173742 | Distal Intergenic |
| chr1 | 135173896 | 135174149 | Distal Intergenic |
| chr1 | 135373497 | 135373927 | Distal Intergenic |
| chr1 | 135506228 | 135506455 | Distal Intergenic |
| chr1 | 135506677 | 135506899 | Distal Intergenic |
| chr1 | 135557049 | 135558165 | Distal Intergenic |
| chr1 | 135590482 | 135593283 | Distal Intergenic |
| chr1 | 135673990 | 135674465 | Distal Intergenic |
| chr1 | 135674554 | 135674793 | Distal Intergenic |
| chr1 | 135674844 | 135675965 | Distal Intergenic |
| chr1 | 135751966 | 135752729 | Distal Intergenic |
| chr1 | 135861344 | 135862627 | Distal Intergenic |
| chr1 | 135863983 | 135865208 | Distal Intergenic |
| chr1 | 135962354 | 135962753 | Distal Intergenic |
| chr1 | 135975980 | 135976263 | Distal Intergenic |
| chr1 | 135995444 | 135998583 | Distal Intergenic |
| chr1 | 136010010 | 136011837 | Distal Intergenic |
| chr1 | 136302494 | 136302781 | Distal Intergenic |
| chr1 | 136311244 | 136311519 | Distal Intergenic |
| chr1 | 136311623 | 136311884 | Distal Intergenic |
| chr1 | 136312604 | 136313004 | Distal Intergenic |

| | | | |
|---|---|---|---|
| chr1 | 136314396 | 136314672 | Distal Intergenic |
| chr1 | 136314709 | 136315211 | Distal Intergenic |
| chr1 | 136315255 | 136317027 | Distal Intergenic |
| chr1 | 136317178 | 136317583 | Distal Intergenic |
| chr1 | 136477228 | 136477726 | Distal Intergenic |
| chr1 | 136480014 | 136480882 | Distal Intergenic |
| chr1 | 136699919 | 136701465 | Distal Intergenic |
| chr1 | 137043616 | 137044073 | Distal Intergenic |
| chr1 | 137079772 | 137079971 | Distal Intergenic |
| chr1 | 137080017 | 137080225 | Distal Intergenic |
| chr1 | 137080319 | 137081137 | Distal Intergenic |
| chr1 | 137155164 | 137155401 | Distal Intergenic |
| chr1 | 137155537 | 137155825 | Distal Intergenic |
| chr1 | 137156913 | 137158131 | Distal Intergenic |
| chr1 | 137164485 | 137165520 | Distal Intergenic |
| chr1 | 137179636 | 137179852 | Distal Intergenic |
| chr1 | 137179902 | 137180249 | Distal Intergenic |
| chr1 | 137209770 | 137210141 | Distal Intergenic |
| chr1 | 137255268 | 137255806 | Distal Intergenic |
| chr1 | 137307452 | 137308198 | Distal Intergenic |
| chr1 | 137363225 | 137364215 | Distal Intergenic |
| chr1 | 137373709 | 137374967 | Distal Intergenic |
| chr1 | 137375012 | 137377850 | Distal Intergenic |
| chr1 | 137386470 | 137389813 | Distal Intergenic |
| chr1 | 137390770 | 137391246 | Distal Intergenic |
| chr1 | 137391367 | 137393053 | Distal Intergenic |
| chr1 | 137405902 | 137406737 | Distal Intergenic |
| chr1 | 137406778 | 137407492 | Distal Intergenic |
| chr1 | 137431525 | 137433315 | Distal Intergenic |
| chr1 | 137565303 | 137566025 | Distal Intergenic |
| chr1 | 137611048 | 137611584 | Distal Intergenic |
| chr1 | 137611625 | 137612350 | Distal Intergenic |
| chr1 | 137627009 | 137627475 | Distal Intergenic |
| chr1 | 137627620 | 137628975 | Distal Intergenic |
| chr1 | 137651135 | 137651356 | Distal Intergenic |
| chr1 | 137651479 | 137653269 | Distal Intergenic |
| chr1 | 137655728 | 137657791 | Distal Intergenic |
| chr1 | 137660347 | 137661011 | Distal Intergenic |
| chr1 | 137661691 | 137662624 | Distal Intergenic |
| chr1 | 137662686 | 137668974 | Distal Intergenic |
| chr1 | 137688673 | 137689644 | Distal Intergenic |
| chr1 | 137689902 | 137691341 | Distal Intergenic |
| chr1 | 137709221 | 137709476 | Distal Intergenic |
| chr1 | 137715534 | 137717060 | Distal Intergenic |

| | | | |
|---|---|---|---|
| chr1 | 137717260 | 137718669 | Distal Intergenic |
| chr1 | 137743240 | 137743536 | Distal Intergenic |
| chr1 | 137748702 | 137749894 | Distal Intergenic |
| chr1 | 137753191 | 137754846 | Distal Intergenic |
| chr1 | 137758179 | 137761000 | Distal Intergenic |
| chr1 | 137764480 | 137767103 | Distal Intergenic |
| chr1 | 137771119 | 137771356 | Distal Intergenic |
| chr1 | 137775198 | 137775427 | Distal Intergenic |
| chr1 | 137779484 | 137781407 | Distal Intergenic |
| chr1 | 137786966 | 137787178 | Distal Intergenic |
| chr1 | 137787375 | 137788320 | Distal Intergenic |
| chr1 | 137788464 | 137791599 | Distal Intergenic |
| chr1 | 137791741 | 137793475 | Distal Intergenic |
| chr1 | 137794542 | 137794760 | Distal Intergenic |
| chr1 | 137794808 | 137801056 | Distal Intergenic |
| chr1 | 137801321 | 137801643 | Distal Intergenic |
| chr1 | 137801940 | 137802198 | Distal Intergenic |
| chr1 | 137802288 | 137804552 | Distal Intergenic |
| chr1 | 137804736 | 137804976 | Distal Intergenic |
| chr1 | 137805952 | 137806163 | Distal Intergenic |
| chr1 | 137806379 | 137806736 | Distal Intergenic |
| chr1 | 137806858 | 137807386 | Distal Intergenic |
| chr1 | 137809496 | 137810064 | Distal Intergenic |
| chr1 | 137810289 | 137811330 | Distal Intergenic |
| chr1 | 137811489 | 137814734 | Distal Intergenic |
| chr1 | 137814888 | 137815934 | Distal Intergenic |
| chr1 | 137821754 | 137823053 | Distal Intergenic |
| chr1 | 137826326 | 137827256 | Distal Intergenic |
| chr1 | 137829631 | 137830991 | Distal Intergenic |
| chr1 | 137834587 | 137835440 | Distal Intergenic |
| chr1 | 137837418 | 137838534 | Distal Intergenic |
| chr1 | 137838693 | 137839769 | Distal Intergenic |
| chr1 | 137840554 | 137840823 | Distal Intergenic |
| chr1 | 138022668 | 138024635 | Distal Intergenic |
| chr1 | 138026186 | 138026674 | Distal Intergenic |
| chr1 | 138031721 | 138032153 | Distal Intergenic |
| chr1 | 138033440 | 138034299 | Distal Intergenic |
| chr1 | 138036246 | 138036445 | Distal Intergenic |
| chr1 | 138036731 | 138037823 | Distal Intergenic |
| chr1 | 138056572 | 138058110 | Distal Intergenic |
| chr1 | 138061303 | 138061509 | Distal Intergenic |
| chr1 | 138120320 | 138121453 | Distal Intergenic |
| chr1 | 138124974 | 138127697 | Distal Intergenic |
| chr1 | 138224385 | 138224606 | Distal Intergenic |

| | | | |
|---|---|---|---|
| chr1 | 138224740 | 138227435 | Distal Intergenic |
| chr1 | 138240311 | 138240759 | Distal Intergenic |
| chr1 | 138241235 | 138241434 | Distal Intergenic |
| chr1 | 138241589 | 138242702 | Distal Intergenic |
| chr1 | 138252041 | 138252832 | Distal Intergenic |
| chr1 | 138296411 | 138296761 | Distal Intergenic |
| chr1 | 138297803 | 138298433 | Distal Intergenic |
| chr1 | 138304471 | 138307795 | Distal Intergenic |
| chr1 | 138311490 | 138311727 | Distal Intergenic |
| chr1 | 138324052 | 138324411 | Distal Intergenic |
| chr1 | 138492285 | 138493318 | Distal Intergenic |
| chr1 | 138582287 | 138582518 | Distal Intergenic |
| chr1 | 138592014 | 138592405 | Distal Intergenic |
| chr1 | 138592556 | 138592880 | Distal Intergenic |
| chr1 | 138593040 | 138593275 | Distal Intergenic |
| chr1 | 138976047 | 138976266 | Distal Intergenic |
| chr1 | 140225657 | 140225946 | Distal Intergenic |
| chr1 | 140244384 | 140246220 | Distal Intergenic |
| chr1 | 140246346 | 140246545 | Distal Intergenic |
| chr1 | 140249228 | 140249748 | Distal Intergenic |
| chr1 | 140250132 | 140250366 | Distal Intergenic |
| chr1 | 140252436 | 140253806 | Distal Intergenic |
| chr1 | 140391570 | 140392467 | Distal Intergenic |
| chr1 | 140403441 | 140403815 | Distal Intergenic |
| chr1 | 140407385 | 140408818 | Distal Intergenic |
| chr1 | 140411957 | 140412744 | Distal Intergenic |
| chr1 | 140432614 | 140434215 | Distal Intergenic |
| chr1 | 140467677 | 140468477 | Distal Intergenic |
| chr1 | 140514454 | 140514670 | Distal Intergenic |
| chr1 | 140515266 | 140515575 | Distal Intergenic |
| chr1 | 140553089 | 140553298 | Distal Intergenic |
| chr1 | 141350644 | 141351072 | Distal Intergenic |
| chr1 | 143141332 | 143141594 | Intron |
| chr1 | 145383919 | 145384620 | Intron |
| chr1 | 145389268 | 145389839 | Intron |
| chr1 | 145549429 | 145549851 | Promoter |
| chr1 | 145788805 | 145789062 | Promoter |
| chr1 | 147804841 | 147805158 | Intron |
| chr1 | 148204113 | 148204849 | Intron |
| chr1 | 149649113 | 149650555 | Exon |
| chr1 | 149898756 | 149899757 | Promoter |
| chr1 | 149932381 | 149932623 | Intron |
| chr1 | 150640840 | 150641121 | Intron |
| chr1 | 151908215 | 151908896 | Distal Intergenic |

| | | | |
|---|---|---|---|
| chr1 | 151911989 | 151912191 | Distal Intergenic |
| chr1 | 151944886 | 151945205 | Distal Intergenic |
| chr1 | 151945378 | 151947221 | Distal Intergenic |
| chr1 | 151962208 | 151963158 | Intron |
| chr1 | 151977411 | 151979685 | Intron |
| chr1 | 151997949 | 152000583 | Intron |
| chr1 | 152011139 | 152012736 | Intron |
| chr1 | 152016599 | 152018136 | Distal Intergenic |
| chr1 | 152044570 | 152044856 | Distal Intergenic |
| chr1 | 152065652 | 152066064 | Distal Intergenic |
| chr1 | 152098450 | 152099327 | Intron |
| chr1 | 152240404 | 152240779 | Distal Intergenic |
| chr1 | 152240852 | 152241143 | Distal Intergenic |
| chr1 | 153019068 | 153019338 | Distal Intergenic |
| chr1 | 153027894 | 153028227 | Downstream |
| chr1 | 153028302 | 153028654 | 3' UTR |
| chr1 | 153073400 | 153076467 | Intron |
| chr1 | 153136257 | 153137044 | Distal Intergenic |
| chr1 | 153192188 | 153192421 | Distal Intergenic |
| chr1 | 153192675 | 153192894 | Distal Intergenic |
| chr1 | 153192967 | 153193368 | Distal Intergenic |
| chr1 | 153194285 | 153194535 | Distal Intergenic |
| chr1 | 153275664 | 153276960 | Exon |
| chr1 | 153800275 | 153800794 | Exon |
| chr1 | 153800885 | 153802076 | Intron |
| chr1 | 153809813 | 153810019 | Intron |
| chr1 | 153891695 | 153892206 | Intron |
| chr1 | 153900231 | 153900918 | Downstream |
| chr1 | 153908120 | 153908370 | Intron |
| chr1 | 153908531 | 153909913 | Exon |
| chr1 | 153909997 | 153911375 | Exon |
| chr1 | 154973330 | 154974778 | Promoter |
| chr1 | 154975143 | 154975378 | Promoter |
| chr1 | 154975566 | 154978783 | Promoter |
| chr1 | 154979485 | 154979697 | Exon |
| chr1 | 154981452 | 154981916 | Intron |
| chr1 | 155021801 | 155023513 | Promoter |
| chr1 | 155024539 | 155026137 | Promoter |
| chr1 | 155027265 | 155028374 | 5' UTR |
| chr1 | 155028485 | 155028964 | 5' UTR |
| chr1 | 155029651 | 155030325 | Exon |
| chr1 | 155032014 | 155033216 | Exon |
| chr1 | 155033304 | 155033614 | Exon |
| chr1 | 155034015 | 155034454 | 3' UTR |

| | | | |
|---|---|---|---|
| chr1 | 155045340 | 155045733 | Intron |
| chr1 | 155046308 | 155046724 | Intron |
| chr1 | 155047411 | 155051997 | Promoter |
| chr1 | 155052062 | 155055120 | Promoter |
| chr1 | 155147277 | 155147476 | Promoter |
| chr1 | 155180287 | 155181021 | Exon |
| chr1 | 155389606 | 155390946 | Intron |
| chr1 | 155412664 | 155413044 | Intron |
| chr1 | 155413146 | 155413433 | Intron |
| chr1 | 155419470 | 155420440 | Intron |
| chr1 | 155432490 | 155432698 | Intron |
| chr1 | 155432837 | 155433329 | Intron |
| chr1 | 155912124 | 155913040 | Promoter |
| chr1 | 156897821 | 156898294 | Intron |
| chr1 | 156899370 | 156900538 | Exon |
| chr1 | 156903507 | 156903786 | Downstream |
| chr1 | 156919043 | 156919301 | Promoter |
| chr1 | 156919753 | 156921934 | Promoter |
| chr1 | 156921987 | 156922348 | Intron |
| chr1 | 156944679 | 156945686 | Intron |
| chr1 | 156985936 | 156986542 | Intron |
| chr1 | 157062115 | 157065135 | 3' UTR |
| chr1 | 157787956 | 157789475 | Promoter |
| chr1 | 157882567 | 157882843 | Distal Intergenic |
| chr1 | 157882936 | 157883293 | Distal Intergenic |
| chr1 | 157883341 | 157885135 | Distal Intergenic |
| chr1 | 157887774 | 157888941 | Distal Intergenic |
| chr1 | 158256401 | 158257386 | Distal Intergenic |
| chr1 | 158407472 | 158407744 | Distal Intergenic |
| chr1 | 158437420 | 158437660 | Distal Intergenic |
| chr1 | 158520381 | 158521021 | Distal Intergenic |
| chr1 | 158523198 | 158524123 | Distal Intergenic |
| chr1 | 158528322 | 158529530 | Downstream |
| chr1 | 158544782 | 158548486 | Downstream |
| chr1 | 158548557 | 158548876 | Promoter |
| chr1 | 158549911 | 158550129 | Promoter |
| chr1 | 158590166 | 158590577 | Exon |
| chr1 | 158597629 | 158598164 | Intron |
| chr1 | 158604358 | 158605370 | Exon |
| chr1 | 158607089 | 158609046 | Exon |
| chr1 | 158609236 | 158610924 | Exon |
| chr1 | 158844317 | 158844529 | Distal Intergenic |
| chr1 | 158844976 | 158845720 | Distal Intergenic |
| chr1 | 159162461 | 159163366 | Exon |

| | | | |
|---|---|---|---|
| chr1 | 159178460 | 159178744 | Distal Intergenic |
| chr1 | 159185939 | 159186149 | Distal Intergenic |
| chr1 | 159186560 | 159187245 | Distal Intergenic |
| chr1 | 159206080 | 159207033 | Distal Intergenic |
| chr1 | 159250248 | 159251723 | Distal Intergenic |
| chr1 | 159273040 | 159274697 | Exon |
| chr1 | 159309450 | 159311764 | Distal Intergenic |
| chr1 | 159335787 | 159336001 | Intron |
| chr1 | 159370310 | 159370859 | Intron |
| chr1 | 159389661 | 159390214 | Intron |
| chr1 | 161182947 | 161183424 | Exon |
| chr1 | 161185382 | 161185667 | Promoter |
| chr1 | 161186257 | 161186494 | Intron |
| chr1 | 161186530 | 161186763 | Intron |
| chr1 | 161186806 | 161187099 | Intron |
| chr1 | 161970953 | 161972134 | Intron |
| chr1 | 161974231 | 161974641 | Intron |
| chr1 | 161989998 | 161990210 | Intron |
| chr1 | 161990310 | 161990827 | Intron |
| chr1 | 162104864 | 162105420 | Intron |
| chr1 | 162134341 | 162134940 | Intron |
| chr1 | 162141192 | 162141391 | Intron |
| chr1 | 162141468 | 162142304 | Intron |
| chr1 | 162143269 | 162143708 | Intron |
| chr1 | 162193092 | 162194923 | Intron |
| chr1 | 162205170 | 162205923 | Intron |
| chr1 | 162299904 | 162301591 | Intron |
| chr1 | 162450914 | 162451150 | Distal Intergenic |
| chr1 | 162451206 | 162451517 | Distal Intergenic |
| chr1 | 162451566 | 162451867 | Distal Intergenic |
| chr1 | 162525995 | 162528014 | Distal Intergenic |
| chr1 | 162837037 | 162837315 | Intron |
| chr1 | 162964314 | 162964513 | Distal Intergenic |
| chr1 | 162965344 | 162968943 | Distal Intergenic |
| chr1 | 162969074 | 162970840 | Distal Intergenic |
| chr1 | 162971167 | 162971403 | Distal Intergenic |
| chr1 | 163001123 | 163001969 | Distal Intergenic |
| chr1 | 163061790 | 163062012 | Distal Intergenic |
| chr1 | 163174367 | 163175181 | Intron |
| chr1 | 163178537 | 163178922 | Intron |
| chr1 | 163179070 | 163179575 | Intron |
| chr1 | 163179752 | 163180200 | Intron |
| chr1 | 163180334 | 163180824 | Intron |
| chr1 | 163180910 | 163181137 | Intron |

| | | | |
|---|---|---|---|
| chr1 | 163198255 | 163198507 | Intron |
| chr1 | 163412159 | 163412726 | Distal Intergenic |
| chr1 | 163537302 | 163537996 | Distal Intergenic |
| chr1 | 163805420 | 163806497 | Distal Intergenic |
| chr1 | 163825701 | 163827203 | Distal Intergenic |
| chr1 | 163836988 | 163838851 | Distal Intergenic |
| chr1 | 163843249 | 163844446 | Distal Intergenic |
| chr1 | 163844662 | 163844861 | Distal Intergenic |
| chr1 | 163845909 | 163846131 | Distal Intergenic |
| chr1 | 164056216 | 164056453 | Distal Intergenic |
| chr1 | 164094808 | 164095039 | Distal Intergenic |
| chr1 | 164095161 | 164095457 | Distal Intergenic |
| chr1 | 164172936 | 164174504 | Distal Intergenic |
| chr1 | 164225037 | 164226859 | Distal Intergenic |
| chr1 | 164227137 | 164227389 | Distal Intergenic |
| chr1 | 164489772 | 164491056 | Distal Intergenic |
| chr1 | 164669648 | 164670040 | Intron |
| chr1 | 164670095 | 164670371 | Intron |
| chr1 | 164670414 | 164670674 | Intron |
| chr1 | 165043727 | 165044075 | Distal Intergenic |
| chr1 | 165044281 | 165045252 | Distal Intergenic |
| chr1 | 165182313 | 165182983 | Promoter |
| chr1 | 165183019 | 165183843 | Promoter |
| chr1 | 165203061 | 165203940 | Intron |
| chr1 | 165204724 | 165205082 | Intron |
| chr1 | 165215179 | 165215794 | Intron |
| chr1 | 165310148 | 165310905 | Intron |
| chr1 | 165311033 | 165311368 | Intron |
| chr1 | 165312252 | 165313214 | Intron |
| chr1 | 165313306 | 165313976 | Intron |
| chr1 | 165333872 | 165334106 | Distal Intergenic |
| chr1 | 165491109 | 165492986 | Exon |
| chr1 | 166179577 | 166181047 | Distal Intergenic |
| chr1 | 166224397 | 166224813 | Distal Intergenic |
| chr1 | 166224865 | 166225087 | Distal Intergenic |
| chr1 | 166237179 | 166237613 | Distal Intergenic |
| chr1 | 166817870 | 166819336 | Exon |
| chr1 | 166923553 | 166923823 | Intron |
| chr1 | 167010351 | 167010668 | Distal Intergenic |
| chr1 | 167220970 | 167222497 | Intron |
| chr1 | 167231913 | 167234190 | Intron |
| chr1 | 167516075 | 167516824 | Intron |
| chr1 | 167572776 | 167573144 | Distal Intergenic |
| chr1 | 167915793 | 167916723 | Intron |

| | | | |
|---|---|---|---|
| chr1 | 167916775 | 167917021 | Intron |
| chr1 | 168898581 | 168898780 | Distal Intergenic |
| chr1 | 168926879 | 168927341 | Distal Intergenic |
| chr1 | 168949914 | 168950531 | Distal Intergenic |
| chr1 | 169113675 | 169114496 | Intron |
| chr1 | 169172777 | 169173136 | Intron |
| chr1 | 169173182 | 169173530 | Intron |
| chr1 | 169198662 | 169200265 | Exon |
| chr1 | 169200944 | 169201310 | Intron |
| chr1 | 169219012 | 169219248 | Intron |
| chr1 | 170208553 | 170209553 | Distal Intergenic |
| chr1 | 170253325 | 170255024 | Promoter |
| chr1 | 170258176 | 170259520 | Distal Intergenic |
| chr1 | 170264579 | 170266423 | Distal Intergenic |
| chr1 | 170266624 | 170267215 | Distal Intergenic |
| chr1 | 170267265 | 170268762 | Distal Intergenic |
| chr1 | 170268811 | 170269246 | Distal Intergenic |
| chr1 | 170292147 | 170293468 | Distal Intergenic |
| chr1 | 170321009 | 170321272 | Distal Intergenic |
| chr1 | 170321391 | 170322969 | Distal Intergenic |
| chr1 | 170356140 | 170358494 | Distal Intergenic |
| chr1 | 170358557 | 170361971 | Distal Intergenic |
| chr1 | 170362643 | 170362867 | Distal Intergenic |
| chr1 | 170363868 | 170364067 | Distal Intergenic |
| chr1 | 170364236 | 170365903 | Distal Intergenic |
| chr1 | 170431947 | 170432999 | Intron |
| chr1 | 170989683 | 170989955 | Intron |
| chr1 | 171488499 | 171488759 | Intron |
| chr1 | 171899596 | 171900334 | Intron |
| chr1 | 172118857 | 172119309 | Intron |
| chr1 | 172145191 | 172145508 | Intron |
| chr1 | 172700838 | 172701052 | Distal Intergenic |
| chr1 | 172974297 | 172974597 | Distal Intergenic |
| chr1 | 172975293 | 172975759 | Distal Intergenic |
| chr1 | 172976221 | 172977037 | Distal Intergenic |
| chr1 | 173177006 | 173177219 | Promoter |
| chr1 | 173200810 | 173201340 | Downstream |
| chr1 | 173224850 | 173225158 | Intron |
| chr1 | 173258563 | 173259490 | Intron |
| chr1 | 173305481 | 173309085 | Intron |
| chr1 | 173314011 | 173314685 | Intron |
| chr1 | 173314739 | 173315033 | Intron |
| chr1 | 173315076 | 173315343 | Intron |
| chr1 | 173315400 | 173315750 | Intron |

| | | | |
|---|---|---|---|
| chr1 | 173367660 | 173370337 | Intron |
| chr1 | 173370648 | 173370849 | Intron |
| chr1 | 173371102 | 173371675 | Intron |
| chr1 | 173372032 | 173372361 | Intron |
| chr1 | 173375111 | 173375520 | Intron |
| chr1 | 173399239 | 173401776 | Intron |
| chr1 | 173403049 | 173404632 | Intron |
| chr1 | 173408057 | 173408843 | Intron |
| chr1 | 173408941 | 173409238 | Intron |
| chr1 | 173433313 | 173433538 | Intron |
| chr1 | 173433611 | 173433877 | Intron |
| chr1 | 173948911 | 173949342 | Intron |
| chr1 | 173949565 | 173950083 | Exon |
| chr1 | 174012290 | 174012868 | Distal Intergenic |
| chr1 | 174012919 | 174013340 | Distal Intergenic |
| chr1 | 174057242 | 174057441 | Distal Intergenic |
| chr1 | 174078036 | 174079222 | Distal Intergenic |
| chr1 | 174121278 | 174121851 | Distal Intergenic |
| chr1 | 174172580 | 174172883 | Intron |
| chr1 | 174219892 | 174220209 | Promoter |
| chr1 | 174240487 | 174241752 | 5' UTR |
| chr1 | 174242605 | 174242805 | Intron |
| chr1 | 174341354 | 174342231 | Intron |
| chr1 | 174353301 | 174354020 | Intron |
| chr1 | 174371893 | 174372103 | Intron |
| chr1 | 174372281 | 174372758 | Intron |
| chr1 | 174429832 | 174432427 | Intron |
| chr1 | 174432524 | 174433434 | Intron |
| chr1 | 174433471 | 174433923 | Intron |
| chr1 | 174487697 | 174487910 | Intron |
| chr1 | 174488054 | 174488513 | Intron |
| chr1 | 174488577 | 174488961 | Intron |
| chr1 | 174495660 | 174495907 | Intron |
| chr1 | 174495991 | 174496229 | Intron |
| chr1 | 174496496 | 174496736 | Intron |
| chr1 | 174609574 | 174611614 | Intron |
| chr1 | 175696510 | 175697949 | Intron |
| chr1 | 175776425 | 175777135 | Distal Intergenic |
| chr1 | 175809904 | 175810337 | Distal Intergenic |
| chr1 | 175842274 | 175842571 | Distal Intergenic |
| chr1 | 175842631 | 175844076 | Distal Intergenic |
| chr1 | 175872411 | 175872646 | Distal Intergenic |
| chr1 | 175912662 | 175913707 | Downstream |
| chr1 | 178757984 | 178758427 | Intron |

| | | | |
|---|---|---|---|
| chr1 | 178762784 | 178762983 | Intron |
| chr1 | 179316137 | 179316467 | Intron |
| chr1 | 179648832 | 179649996 | Intron |
| chr1 | 179678257 | 179678521 | Distal Intergenic |
| chr1 | 179724933 | 179725308 | Intron |
| chr1 | 179725399 | 179725614 | Intron |
| chr1 | 179725806 | 179726132 | Intron |
| chr1 | 180227733 | 180229178 | Intron |
| chr1 | 180248185 | 180248405 | Distal Intergenic |
| chr1 | 180248445 | 180250644 | Distal Intergenic |
| chr1 | 180251060 | 180251301 | Distal Intergenic |
| chr1 | 180265762 | 180266978 | Intron |
| chr1 | 180267442 | 180268494 | Intron |
| chr1 | 180335857 | 180336950 | Intron |
| chr1 | 180971546 | 180973618 | Exon |
| chr1 | 181733400 | 181733608 | Intron |
| chr1 | 181778185 | 181778706 | Distal Intergenic |
| chr1 | 181900204 | 181900797 | Distal Intergenic |
| chr1 | 181944670 | 181944869 | Distal Intergenic |
| chr1 | 181944920 | 181946589 | Distal Intergenic |
| chr1 | 182242681 | 182243014 | Distal Intergenic |
| chr1 | 182258805 | 182259927 | Distal Intergenic |
| chr1 | 182843953 | 182844181 | Exon |
| chr1 | 182871712 | 182873859 | Exon |
| chr1 | 182873971 | 182875408 | Exon |
| chr1 | 182885674 | 182886557 | Intron |
| chr1 | 182896523 | 182896760 | Intron |
| chr1 | 182896827 | 182898011 | Intron |
| chr1 | 182945295 | 182947264 | Distal Intergenic |
| chr1 | 183056827 | 183057065 | Intron |
| chr1 | 183074480 | 183074717 | Intron |
| chr1 | 183161271 | 183161470 | Intron |
| chr1 | 183890067 | 183890302 | Intron |
| chr1 | 183890419 | 183890709 | Intron |
| chr1 | 183915914 | 183916135 | Promoter |
| chr1 | 183916195 | 183916563 | Promoter |
| chr1 | 183927590 | 183927876 | Intron |
| chr1 | 183943415 | 183943683 | Intron |
| chr1 | 183943753 | 183944270 | Exon |
| chr1 | 183944381 | 183949724 | 5' UTR |
| chr1 | 183950782 | 183952830 | Intron |
| chr1 | 183982686 | 183984124 | Intron |
| chr1 | 183987442 | 183988872 | Intron |
| chr1 | 183988933 | 183989630 | Intron |

| | | | |
|---|---|---|---|
| chr1 | 183989727 | 183990137 | Intron |
| chr1 | 183996442 | 183999530 | Intron |
| chr1 | 184004120 | 184005496 | Intron |
| chr1 | 184017722 | 184023556 | Promoter |
| chr1 | 184024125 | 184024918 | Intron |
| chr1 | 184024965 | 184025486 | Intron |
| chr1 | 184025629 | 184025858 | Intron |
| chr1 | 184119568 | 184120519 | Distal Intergenic |
| chr1 | 184445310 | 184445664 | Intron |
| chr1 | 184445713 | 184447367 | 5' UTR |
| chr1 | 184458304 | 184458840 | Intron |
| chr1 | 184472162 | 184472428 | Intron |
| chr1 | 184472475 | 184472748 | Intron |
| chr1 | 184472789 | 184473149 | Intron |
| chr1 | 184473306 | 184473513 | Intron |
| chr1 | 184522347 | 184522596 | Intron |
| chr1 | 184874398 | 184875633 | Intron |
| chr1 | 184886167 | 184886471 | Intron |
| chr1 | 185493597 | 185493930 | Distal Intergenic |
| chr1 | 185602667 | 185602866 | Distal Intergenic |
| chr1 | 185688710 | 185689822 | Distal Intergenic |
| chr1 | 185728973 | 185729389 | Intron |
| chr1 | 185729447 | 185730075 | Intron |
| chr1 | 185737664 | 185738630 | Intron |
| chr1 | 185740504 | 185740703 | Intron |
| chr1 | 185741305 | 185741856 | Intron |
| chr1 | 185744866 | 185745073 | Intron |
| chr1 | 185745278 | 185745999 | Intron |
| chr1 | 185754211 | 185754877 | Intron |
| chr1 | 185766415 | 185766799 | Intron |
| chr1 | 185768587 | 185769630 | Intron |
| chr1 | 185787474 | 185788055 | Intron |
| chr1 | 185858264 | 185858463 | Intron |
| chr1 | 185875412 | 185875823 | Intron |
| chr1 | 185876091 | 185876381 | Intron |
| chr1 | 185881047 | 185881456 | Intron |
| chr1 | 185900255 | 185900518 | Intron |
| chr1 | 185900819 | 185901424 | Intron |
| chr1 | 185907736 | 185909196 | Intron |
| chr1 | 185990761 | 185990992 | Intron |
| chr1 | 186631933 | 186633327 | Distal Intergenic |
| chr1 | 186646970 | 186647279 | Intron |
| chr1 | 186976503 | 186977155 | Distal Intergenic |
| chr1 | 188285434 | 188285745 | Distal Intergenic |

| | | | |
|---|---|---|---|
| chr1 | 189039131 | 189039340 | Distal Intergenic |
| chr1 | 190743493 | 190743916 | Intron |
| chr1 | 190743955 | 190744650 | Intron |
| chr1 | 190744791 | 190745004 | Intron |
| chr1 | 191025485 | 191025956 | Distal Intergenic |
| chr1 | 191027784 | 191029589 | Distal Intergenic |
| chr1 | 191038225 | 191039959 | Distal Intergenic |
| chr1 | 191183223 | 191184402 | Distal Intergenic |
| chr1 | 191551544 | 191551781 | Distal Intergenic |
| chr1 | 191578690 | 191579741 | Distal Intergenic |
| chr1 | 191638228 | 191638919 | Distal Intergenic |
| chr1 | 193925090 | 193925355 | Distal Intergenic |
| chr1 | 194237541 | 194238144 | Distal Intergenic |
| chr1 | 194268410 | 194269513 | Distal Intergenic |
| chr1 | 194271300 | 194274431 | Distal Intergenic |
| chr1 | 194274660 | 194274859 | Distal Intergenic |
| chr1 | 194276064 | 194276310 | Distal Intergenic |
| chr1 | 194571583 | 194571782 | Distal Intergenic |
| chr1 | 194605037 | 194605860 | Distal Intergenic |
| chr1 | 194612143 | 194612541 | Distal Intergenic |
| chr1 | 194633106 | 194633737 | Distal Intergenic |
| chr1 | 194654950 | 194655756 | Distal Intergenic |
| chr1 | 194657679 | 194657891 | Distal Intergenic |
| chr1 | 194956597 | 194958137 | Distal Intergenic |
| chr1 | 194965480 | 194968965 | Distal Intergenic |
| chr1 | 194975952 | 194976311 | Distal Intergenic |
| chr1 | 194976759 | 194978013 | Distal Intergenic |
| chr1 | 194978214 | 194978484 | Distal Intergenic |
| chr1 | 194978846 | 194979058 | Distal Intergenic |
| chr1 | 194979419 | 194980601 | Distal Intergenic |
| chr1 | 194981088 | 194981651 | Distal Intergenic |
| chr1 | 195124154 | 195124455 | Distal Intergenic |
| chr1 | 195125992 | 195127138 | Distal Intergenic |
| chr1 | 195127271 | 195128481 | Distal Intergenic |
| chr1 | 195224503 | 195227194 | Distal Intergenic |
| chr1 | 195330909 | 195331303 | Distal Intergenic |
| chr1 | 195331937 | 195333443 | Distal Intergenic |
| chr1 | 195333498 | 195340310 | Distal Intergenic |
| chr1 | 195340379 | 195341176 | Distal Intergenic |
| chr1 | 195341256 | 195341456 | Distal Intergenic |
| chr1 | 195342158 | 195349726 | Distal Intergenic |
| chr1 | 195352477 | 195353480 | Distal Intergenic |
| chr1 | 195371747 | 195371946 | Distal Intergenic |
| chr1 | 195372240 | 195374242 | Distal Intergenic |

| | | | |
|---|---|---|---|
| chr1 | 195374506 | 195377832 | Distal Intergenic |
| chr1 | 195378098 | 195378328 | Distal Intergenic |
| chr1 | 195379966 | 195381297 | Distal Intergenic |
| chr1 | 195381396 | 195381618 | Distal Intergenic |
| chr1 | 195387658 | 195389450 | Distal Intergenic |
| chr1 | 195389643 | 195391713 | Distal Intergenic |
| chr1 | 195392540 | 195396112 | Distal Intergenic |
| chr1 | 195400515 | 195401859 | Distal Intergenic |
| chr1 | 195414361 | 195414899 | Distal Intergenic |
| chr1 | 195415104 | 195415324 | Distal Intergenic |
| chr1 | 196478399 | 196478600 | Intron |
| chr1 | 196501758 | 196502008 | Intron |
| chr1 | 196502171 | 196503057 | Intron |
| chr1 | 196503284 | 196503580 | Intron |
| chr1 | 196591035 | 196591301 | Distal Intergenic |
| chr1 | 196955962 | 196956374 | Intron |
| chr1 | 196956486 | 196957733 | Intron |
| chr10 | 3086226 | 3086462 | Distal Intergenic |
| chr10 | 3087084 | 3088500 | Distal Intergenic |
| chr10 | 3133814 | 3134201 | Intron |
| chr10 | 3156566 | 3157629 | Intron |
| chr10 | 3159794 | 3160726 | 5' UTR |
| chr10 | 3165652 | 3166104 | Intron |
| chr10 | 3501202 | 3502355 | Intron |
| chr10 | 3502436 | 3502644 | Intron |
| chr10 | 3503019 | 3504714 | Intron |
| chr10 | 3521369 | 3522069 | Intron |
| chr10 | 4522274 | 4522840 | Distal Intergenic |
| chr10 | 4767718 | 4768254 | Distal Intergenic |
| chr10 | 5057317 | 5057560 | Intron |
| chr10 | 5137767 | 5138390 | Intron |
| chr10 | 5235430 | 5235880 | Distal Intergenic |
| chr10 | 5259996 | 5261138 | 3' UTR |
| chr10 | 5437503 | 5438173 | Intron |
| chr10 | 5724745 | 5726142 | Promoter |
| chr10 | 5733763 | 5735140 | Exon |
| chr10 | 5735360 | 5735599 | Intron |
| chr10 | 5913141 | 5913412 | Intron |
| chr10 | 5913532 | 5913912 | Intron |
| chr10 | 5951033 | 5951273 | Exon |
| chr10 | 6224862 | 6225835 | Intron |
| chr10 | 6238151 | 6239515 | Intron |
| chr10 | 6370776 | 6371133 | Exon |
| chr10 | 6371181 | 6371474 | Exon |

| | | | |
|---|---|---|---|
| chr10 | 6372169 | 6372682 | Intron |
| chr10 | 6372780 | 6373394 | Exon |
| chr10 | 6403970 | 6405544 | Distal Intergenic |
| chr10 | 6411264 | 6412474 | Distal Intergenic |
| chr10 | 6481955 | 6483500 | Intron |
| chr10 | 6483680 | 6484427 | Exon |
| chr10 | 6509903 | 6512293 | Intron |
| chr10 | 6879650 | 6880642 | Exon |
| chr10 | 6961113 | 6963744 | Distal Intergenic |
| chr10 | 6985714 | 6985951 | Distal Intergenic |
| chr10 | 7108807 | 7109678 | Distal Intergenic |
| chr10 | 7383173 | 7383741 | Intron |
| chr10 | 7400181 | 7401995 | Intron |
| chr10 | 7445936 | 7446767 | Intron |
| chr10 | 7496253 | 7497645 | Distal Intergenic |
| chr10 | 7507292 | 7508001 | Distal Intergenic |
| chr10 | 7512054 | 7513266 | Distal Intergenic |
| chr10 | 7581388 | 7581588 | Distal Intergenic |
| chr10 | 7632149 | 7632362 | Intron |
| chr10 | 7674487 | 7674693 | Intron |
| chr10 | 7675014 | 7675299 | Intron |
| chr10 | 7675392 | 7676052 | Intron |
| chr10 | 7676143 | 7676940 | Intron |
| chr10 | 7996164 | 7997903 | Intron |
| chr10 | 8060868 | 8061398 | Distal Intergenic |
| chr10 | 8125278 | 8125495 | Distal Intergenic |
| chr10 | 8211614 | 8212022 | Distal Intergenic |
| chr10 | 8225585 | 8226378 | Distal Intergenic |
| chr10 | 8467367 | 8467633 | Distal Intergenic |
| chr10 | 8481784 | 8482443 | Distal Intergenic |
| chr10 | 8487679 | 8489583 | Distal Intergenic |
| chr10 | 8491721 | 8491920 | Distal Intergenic |
| chr10 | 8531093 | 8532222 | Distal Intergenic |
| chr10 | 8532422 | 8532704 | Distal Intergenic |
| chr10 | 8593150 | 8593999 | Distal Intergenic |
| chr10 | 8605902 | 8606115 | Distal Intergenic |
| chr10 | 8606186 | 8606435 | Distal Intergenic |
| chr10 | 8609681 | 8609942 | Distal Intergenic |
| chr10 | 8609981 | 8610929 | Distal Intergenic |
| chr10 | 8698108 | 8701254 | Exon |
| chr10 | 8991937 | 8992409 | Distal Intergenic |
| chr10 | 9002367 | 9002599 | Distal Intergenic |
| chr10 | 9215978 | 9216584 | Distal Intergenic |
| chr10 | 9216746 | 9217225 | Distal Intergenic |

| | | | |
|---|---|---|---|
| chr10 | 9248609 | 9250000 | Distal Intergenic |
| chr10 | 9250062 | 9250295 | Distal Intergenic |
| chr10 | 9264379 | 9265197 | Distal Intergenic |
| chr10 | 9318701 | 9318984 | Distal Intergenic |
| chr10 | 9319163 | 9320350 | Distal Intergenic |
| chr10 | 9350097 | 9351467 | Distal Intergenic |
| chr10 | 9360205 | 9360413 | Distal Intergenic |
| chr10 | 9360473 | 9360721 | Distal Intergenic |
| chr10 | 9601553 | 9601977 | Distal Intergenic |
| chr10 | 9621076 | 9622170 | Distal Intergenic |
| chr10 | 9779289 | 9779605 | Distal Intergenic |
| chr10 | 9971652 | 9972474 | Distal Intergenic |
| chr10 | 9972592 | 9973029 | Distal Intergenic |
| chr10 | 10117658 | 10118103 | Distal Intergenic |
| chr10 | 10263988 | 10264454 | Distal Intergenic |
| chr10 | 10264835 | 10265927 | Distal Intergenic |
| chr10 | 10274443 | 10276207 | Distal Intergenic |
| chr10 | 10276258 | 10277117 | Distal Intergenic |
| chr10 | 10277214 | 10277812 | Distal Intergenic |
| chr10 | 10917452 | 10917851 | Distal Intergenic |
| chr10 | 10917888 | 10918216 | Distal Intergenic |
| chr10 | 11000233 | 11000956 | Distal Intergenic |
| chr10 | 11001567 | 11002500 | Distal Intergenic |
| chr10 | 12324700 | 12325206 | Distal Intergenic |
| chr10 | 12414541 | 12415054 | Intron |
| chr10 | 12445915 | 12446145 | Intron |
| chr10 | 12446191 | 12446598 | Intron |
| chr10 | 12507484 | 12508518 | Intron |
| chr10 | 12530346 | 12530629 | Intron |
| chr10 | 12530834 | 12531249 | Intron |
| chr10 | 12547124 | 12547580 | Intron |
| chr10 | 12586773 | 12588081 | Intron |
| chr10 | 12588140 | 12588968 | Intron |
| chr10 | 12594625 | 12595200 | Intron |
| chr10 | 12596021 | 12596352 | Intron |
| chr10 | 12598169 | 12598573 | Intron |
| chr10 | 12598642 | 12599165 | Intron |
| chr10 | 12599508 | 12599720 | Intron |
| chr10 | 12599788 | 12600479 | Intron |
| chr10 | 12612804 | 12613743 | Intron |
| chr10 | 12631941 | 12632651 | Intron |
| chr10 | 12642623 | 12642822 | Intron |
| chr10 | 12683043 | 12683625 | Intron |
| chr10 | 12727843 | 12728124 | Intron |

| | | | |
|---|---|---|---|
| chr10 | 12860069 | 12860275 | Intron |
| chr10 | 12860443 | 12860676 | Intron |
| chr10 | 12912360 | 12912994 | Distal Intergenic |
| chr10 | 13089352 | 13090377 | Intron |
| chr10 | 13186487 | 13187045 | Distal Intergenic |
| chr10 | 13272571 | 13272901 | Intron |
| chr10 | 13273029 | 13273634 | Intron |
| chr10 | 13296328 | 13297979 | Distal Intergenic |
| chr10 | 13315042 | 13315242 | Distal Intergenic |
| chr10 | 13315445 | 13316580 | Distal Intergenic |
| chr10 | 13379562 | 13380093 | Promoter |
| chr10 | 13383783 | 13384115 | Intron |
| chr10 | 13403441 | 13404654 | Distal Intergenic |
| chr10 | 13408014 | 13409053 | Distal Intergenic |
| chr10 | 13495248 | 13495480 | Intron |
| chr10 | 13512176 | 13512576 | Intron |
| chr10 | 13512780 | 13513068 | Intron |
| chr10 | 13552195 | 13553089 | Distal Intergenic |
| chr10 | 13589076 | 13589614 | Distal Intergenic |
| chr10 | 13685682 | 13685908 | 3' UTR |
| chr10 | 13686490 | 13686823 | 3' UTR |
| chr10 | 13737486 | 13737924 | Intron |
| chr10 | 14027426 | 14027742 | Intron |
| chr10 | 14031805 | 14032304 | Intron |
| chr10 | 14032722 | 14034677 | Intron |
| chr10 | 14104437 | 14107612 | Intron |
| chr10 | 14247380 | 14248278 | Intron |
| chr10 | 14382065 | 14382323 | Intron |
| chr10 | 14384422 | 14385415 | Intron |
| chr10 | 14424516 | 14425320 | Intron |
| chr10 | 16227694 | 16228007 | Distal Intergenic |
| chr10 | 16264640 | 16265909 | Distal Intergenic |
| chr10 | 16272704 | 16273286 | Distal Intergenic |
| chr10 | 16598032 | 16598598 | Distal Intergenic |
| chr10 | 16731342 | 16733923 | Intron |
| chr10 | 16734009 | 16734263 | Intron |
| chr10 | 16734439 | 16734871 | Intron |
| chr10 | 16754938 | 16755188 | Intron |
| chr10 | 16971998 | 16973294 | Intron |
| chr10 | 17018308 | 17018931 | Intron |
| chr10 | 17067150 | 17068216 | Intron |
| chr10 | 17068417 | 17068683 | Intron |
| chr10 | 17268994 | 17269918 | Promoter |
| chr10 | 17304111 | 17306233 | Distal Intergenic |

| | | | |
|---|---|---|---|
| chr10 | 17443097 | 17443416 | Intron |
| chr10 | 17471570 | 17473568 | Intron |
| chr10 | 17478885 | 17481327 | Intron |
| chr10 | 17491909 | 17492463 | Intron |
| chr10 | 17496195 | 17497179 | Promoter |
| chr10 | 17588802 | 17589882 | Distal Intergenic |
| chr10 | 17649915 | 17651476 | Intron |
| chr10 | 17651523 | 17651761 | Intron |
| chr10 | 17725135 | 17726036 | Intron |
| chr10 | 17728272 | 17728504 | Intron |
| chr10 | 17741664 | 17742361 | 5' UTR |
| chr10 | 17742594 | 17743247 | Intron |
| chr10 | 17828604 | 17831673 | Intron |
| chr10 | 18025001 | 18025297 | Intron |
| chr10 | 18043089 | 18044328 | Intron |
| chr10 | 18049089 | 18049994 | Intron |
| chr10 | 18070198 | 18071418 | Intron |
| chr10 | 18158546 | 18158840 | Intron |
| chr10 | 18183037 | 18183318 | Exon |
| chr10 | 18190949 | 18192360 | Exon |
| chr10 | 18200373 | 18201180 | Distal Intergenic |
| chr10 | 18564933 | 18567905 | Intron |
| chr10 | 18567982 | 18571126 | Intron |
| chr10 | 18571712 | 18572254 | Intron |
| chr10 | 18578748 | 18581665 | Intron |
| chr10 | 18582890 | 18583899 | Intron |
| chr10 | 18595393 | 18596851 | Intron |
| chr10 | 18604150 | 18604380 | Intron |
| chr10 | 18707217 | 18707494 | Intron |
| chr10 | 18707554 | 18708069 | Intron |
| chr10 | 18728098 | 18730826 | Intron |
| chr10 | 18731530 | 18731729 | Intron |
| chr10 | 18731770 | 18732938 | Intron |
| chr10 | 18733178 | 18733494 | Intron |
| chr10 | 18746414 | 18746733 | Intron |
| chr10 | 18746815 | 18747014 | Intron |
| chr10 | 18747617 | 18748954 | Intron |
| chr10 | 18753993 | 18755827 | Intron |
| chr10 | 18763588 | 18763805 | Intron |
| chr10 | 18765353 | 18767393 | Intron |
| chr10 | 18784808 | 18785906 | Intron |
| chr10 | 18801041 | 18803630 | Exon |
| chr10 | 18811102 | 18812288 | Intron |
| chr10 | 18838592 | 18838953 | Intron |

| | | | |
|---|---|---|---|
| chr10 | 18887702 | 18888067 | Intron |
| chr10 | 18896292 | 18896665 | Intron |
| chr10 | 18896742 | 18897860 | Intron |
| chr10 | 18902766 | 18903086 | Intron |
| chr10 | 19183556 | 19185574 | Distal Intergenic |
| chr10 | 19217802 | 19218882 | Distal Intergenic |
| chr10 | 19219251 | 19219506 | Distal Intergenic |
| chr10 | 19797084 | 19797606 | Intron |
| chr10 | 19810665 | 19811606 | Intron |
| chr10 | 19823992 | 19824211 | Intron |
| chr10 | 19824872 | 19825145 | Intron |
| chr10 | 19869362 | 19869629 | Intron |
| chr10 | 19869785 | 19870098 | Intron |
| chr10 | 19870154 | 19870399 | Intron |
| chr10 | 19878896 | 19879733 | Intron |
| chr10 | 19889073 | 19890511 | Intron |
| chr10 | 19893674 | 19894414 | Intron |
| chr10 | 19898340 | 19899427 | Distal Intergenic |
| chr10 | 20031374 | 20032538 | Distal Intergenic |
| chr10 | 20032823 | 20033428 | Distal Intergenic |
| chr10 | 20287835 | 20290135 | Intron |
| chr10 | 20290318 | 20290568 | Intron |
| chr10 | 20290611 | 20292645 | Exon |
| chr10 | 20377244 | 20378334 | Intron |
| chr10 | 20839029 | 20839230 | Downstream |
| chr10 | 20892813 | 20893793 | Distal Intergenic |
| chr10 | 21015791 | 21016595 | Distal Intergenic |
| chr10 | 21018891 | 21019116 | Distal Intergenic |
| chr10 | 21019516 | 21022120 | Distal Intergenic |
| chr10 | 21024124 | 21024416 | Distal Intergenic |
| chr10 | 21024463 | 21024757 | Distal Intergenic |
| chr10 | 21053730 | 21056364 | Distal Intergenic |
| chr10 | 21062354 | 21063076 | Distal Intergenic |
| chr10 | 21118817 | 21119188 | Intron |
| chr10 | 21493369 | 21493595 | Distal Intergenic |
| chr10 | 21509775 | 21511065 | Distal Intergenic |
| chr10 | 21548259 | 21550077 | Distal Intergenic |
| chr10 | 21552930 | 21554547 | Distal Intergenic |
| chr10 | 21558636 | 21560130 | Distal Intergenic |
| chr10 | 21565941 | 21568226 | Distal Intergenic |
| chr10 | 21714154 | 21714353 | Distal Intergenic |
| chr10 | 21714516 | 21715063 | Distal Intergenic |
| chr10 | 22415377 | 22415663 | Distal Intergenic |
| chr10 | 22416853 | 22417053 | Distal Intergenic |

| | | | |
|---|---|---|---|
| chr10 | 22418054 | 22418770 | Distal Intergenic |
| chr10 | 22451064 | 22451863 | Distal Intergenic |
| chr10 | 22876810 | 22877067 | Intron |
| chr10 | 22896429 | 22898079 | Exon |
| chr10 | 22904782 | 22905994 | Intron |
| chr10 | 23502957 | 23505773 | Distal Intergenic |
| chr10 | 23506125 | 23506808 | Distal Intergenic |
| chr10 | 23506863 | 23508582 | Distal Intergenic |
| chr10 | 23929386 | 23931466 | Distal Intergenic |
| chr10 | 24002107 | 24002406 | Intron |
| chr10 | 24002447 | 24002659 | Intron |
| chr10 | 24165521 | 24166765 | Intron |
| chr10 | 24506976 | 24507175 | Intron |
| chr10 | 24507260 | 24507877 | Intron |
| chr10 | 24602353 | 24604458 | Intron |
| chr10 | 24604621 | 24604857 | Intron |
| chr10 | 24692570 | 24693806 | Intron |
| chr10 | 24704353 | 24705120 | Intron |
| chr10 | 25145894 | 25147117 | Exon |
| chr10 | 25152947 | 25153659 | Intron |
| chr10 | 25508495 | 25508748 | Intron |
| chr10 | 25680938 | 25681401 | Intron |
| chr10 | 25968998 | 25970047 | Intron |
| chr10 | 25970208 | 25970431 | Intron |
| chr10 | 26077951 | 26079900 | Distal Intergenic |
| chr10 | 26093536 | 26093745 | Distal Intergenic |
| chr10 | 26284091 | 26284970 | Intron |
| chr10 | 26387770 | 26388215 | Intron |
| chr10 | 26388315 | 26388833 | Intron |
| chr10 | 26398682 | 26398989 | Intron |
| chr10 | 26399190 | 26399547 | Intron |
| chr10 | 26482196 | 26482635 | Exon |
| chr10 | 26492306 | 26492512 | Intron |
| chr10 | 26615440 | 26615987 | Distal Intergenic |
| chr10 | 26715884 | 26716176 | Distal Intergenic |
| chr10 | 27373887 | 27374182 | Intron |
| chr10 | 27374343 | 27374581 | Intron |
| chr10 | 27702846 | 27706131 | Promoter |
| chr10 | 27706193 | 27709570 | Distal Intergenic |
| chr10 | 27795024 | 27796166 | Intron |
| chr10 | 27806890 | 27807918 | Intron |
| chr10 | 28023587 | 28024787 | Exon |
| chr10 | 28863252 | 28863932 | Intron |
| chr10 | 28864093 | 28864292 | Intron |

| | | | |
|---|---|---|---|
| chr10 | 29427497 | 29427726 | Distal Intergenic |
| chr10 | 29427766 | 29429014 | Distal Intergenic |
| chr10 | 29429234 | 29429654 | Distal Intergenic |
| chr10 | 29917917 | 29918313 | Intron |
| chr10 | 29919943 | 29920294 | Intron |
| chr10 | 30320211 | 30320733 | Intron |
| chr10 | 30327960 | 30328372 | Intron |
| chr10 | 30337797 | 30338145 | Intron |
| chr10 | 30338325 | 30338951 | Intron |
| chr10 | 31252115 | 31255507 | Intron |
| chr10 | 31257345 | 31259197 | Intron |
| chr10 | 31260127 | 31261086 | Intron |
| chr10 | 31272674 | 31274072 | 5' UTR |
| chr10 | 31277208 | 31277431 | Intron |
| chr10 | 31327141 | 31327973 | Distal Intergenic |
| chr10 | 31328125 | 31328378 | Distal Intergenic |
| chr10 | 31328433 | 31328714 | Distal Intergenic |
| chr10 | 31328894 | 31329097 | Distal Intergenic |
| chr10 | 31587475 | 31588323 | Distal Intergenic |
| chr10 | 32703967 | 32706018 | Distal Intergenic |
| chr10 | 33738608 | 33739225 | Distal Intergenic |
| chr10 | 33906178 | 33907411 | Distal Intergenic |
| chr10 | 33914354 | 33914559 | Distal Intergenic |
| chr10 | 33916162 | 33918208 | Distal Intergenic |
| chr10 | 33918903 | 33919102 | Distal Intergenic |
| chr10 | 33926135 | 33927661 | Distal Intergenic |
| chr10 | 33927997 | 33928274 | Distal Intergenic |
| chr10 | 33928399 | 33928663 | Distal Intergenic |
| chr10 | 33948998 | 33949881 | Distal Intergenic |
| chr10 | 33949965 | 33950202 | Distal Intergenic |
| chr10 | 34202987 | 34204625 | Distal Intergenic |
| chr10 | 34226784 | 34228800 | Distal Intergenic |
| chr10 | 34252134 | 34253165 | Distal Intergenic |
| chr10 | 34316017 | 34316375 | Distal Intergenic |
| chr10 | 34316450 | 34317006 | Distal Intergenic |
| chr10 | 34338238 | 34338793 | Distal Intergenic |
| chr10 | 36421764 | 36423683 | Distal Intergenic |
| chr10 | 36694119 | 36694745 | Distal Intergenic |
| chr10 | 36856995 | 36858797 | Distal Intergenic |
| chr10 | 36858836 | 36860092 | Distal Intergenic |
| chr10 | 36860174 | 36860793 | Distal Intergenic |
| chr10 | 36861304 | 36861659 | Distal Intergenic |
| chr10 | 37050288 | 37050965 | Distal Intergenic |
| chr10 | 37054225 | 37057891 | Distal Intergenic |

| | | | |
|---|---|---|---|
| chr10 | 37080328 | 37082005 | Distal Intergenic |
| chr10 | 37087068 | 37088325 | Distal Intergenic |
| chr10 | 37176784 | 37177167 | Distal Intergenic |
| chr10 | 37177220 | 37177628 | Distal Intergenic |
| chr10 | 37320137 | 37320835 | Distal Intergenic |
| chr10 | 37322816 | 37323299 | Distal Intergenic |
| chr10 | 37333767 | 37333979 | Distal Intergenic |
| chr10 | 37523160 | 37523370 | Distal Intergenic |
| chr10 | 37966644 | 37966856 | Distal Intergenic |
| chr10 | 37967019 | 37967225 | Distal Intergenic |
| chr10 | 38714178 | 38714919 | Distal Intergenic |
| chr10 | 39135407 | 39135749 | Distal Intergenic |
| chr10 | 39290313 | 39290724 | Distal Intergenic |
| chr10 | 39290761 | 39291198 | Distal Intergenic |
| chr10 | 39292435 | 39293446 | Distal Intergenic |
| chr10 | 39332042 | 39334274 | Distal Intergenic |
| chr10 | 39966400 | 39966626 | Distal Intergenic |
| chr10 | 39976686 | 39979476 | Distal Intergenic |
| chr10 | 39981662 | 39984452 | Distal Intergenic |
| chr10 | 39987757 | 39989158 | Distal Intergenic |
| chr10 | 40018187 | 40018766 | Distal Intergenic |
| chr10 | 40018975 | 40019190 | Distal Intergenic |
| chr10 | 40020071 | 40020372 | Distal Intergenic |
| chr10 | 40020423 | 40020920 | Distal Intergenic |
| chr10 | 40021077 | 40021688 | Distal Intergenic |
| chr10 | 40021743 | 40022711 | Distal Intergenic |
| chr10 | 40036692 | 40037288 | Distal Intergenic |
| chr10 | 40057589 | 40059476 | Distal Intergenic |
| chr10 | 40067566 | 40068566 | Distal Intergenic |
| chr10 | 40090248 | 40092855 | Distal Intergenic |
| chr10 | 40099244 | 40099483 | Distal Intergenic |
| chr10 | 40136647 | 40137741 | Distal Intergenic |
| chr10 | 40162005 | 40162508 | Distal Intergenic |
| chr10 | 40162907 | 40164029 | Distal Intergenic |
| chr10 | 40230177 | 40232087 | Distal Intergenic |
| chr10 | 40241370 | 40241704 | Distal Intergenic |
| chr10 | 40602136 | 40603147 | Distal Intergenic |
| chr10 | 40622892 | 40623176 | Distal Intergenic |
| chr10 | 41238322 | 41238880 | Distal Intergenic |
| chr10 | 41239288 | 41240302 | Distal Intergenic |
| chr10 | 41314740 | 41314952 | Distal Intergenic |
| chr10 | 41450369 | 41451797 | Distal Intergenic |
| chr10 | 41600780 | 41602090 | Distal Intergenic |
| chr10 | 41612980 | 41613779 | Distal Intergenic |

| | | | |
|---|---|---|---|
| chr10 | 41753384 | 41753846 | Distal Intergenic |
| chr10 | 41802606 | 41803065 | Distal Intergenic |
| chr10 | 41806312 | 41806772 | Distal Intergenic |
| chr10 | 41821196 | 41821412 | Distal Intergenic |
| chr10 | 41823176 | 41824672 | Distal Intergenic |
| chr10 | 41847668 | 41848096 | Distal Intergenic |
| chr10 | 41848137 | 41848686 | Distal Intergenic |
| chr10 | 41850165 | 41850713 | Distal Intergenic |
| chr10 | 41939339 | 41939635 | Distal Intergenic |
| chr10 | 41941844 | 41942129 | Distal Intergenic |
| chr10 | 41942173 | 41942509 | Distal Intergenic |
| chr10 | 41967647 | 41967864 | Distal Intergenic |
| chr10 | 41993987 | 41994653 | Distal Intergenic |
| chr10 | 41995180 | 41995468 | Distal Intergenic |
| chr10 | 41996170 | 41996514 | Distal Intergenic |
| chr10 | 42194344 | 42195183 | Distal Intergenic |
| chr10 | 42195275 | 42195532 | Distal Intergenic |
| chr10 | 42197745 | 42198429 | Distal Intergenic |
| chr10 | 42221954 | 42223190 | Distal Intergenic |
| chr10 | 42478605 | 42479201 | Distal Intergenic |
| chr10 | 42479333 | 42480061 | Distal Intergenic |
| chr10 | 42480780 | 42481152 | Distal Intergenic |
| chr10 | 42481372 | 42481911 | Distal Intergenic |
| chr10 | 42482112 | 42482904 | Distal Intergenic |
| chr10 | 42500941 | 42501606 | Distal Intergenic |
| chr10 | 42584619 | 42584945 | Distal Intergenic |
| chr10 | 42936842 | 42938218 | Intron |
| chr10 | 42940325 | 42942009 | Intron |
| chr10 | 43298951 | 43301137 | Intron |
| chr10 | 43372970 | 43373566 | Distal Intergenic |
| chr10 | 43377363 | 43378126 | Distal Intergenic |
| chr10 | 43404275 | 43404498 | Distal Intergenic |
| chr10 | 43409179 | 43410104 | Distal Intergenic |
| chr10 | 43639940 | 43641625 | Intron |
| chr10 | 43715741 | 43716429 | Intron |
| chr10 | 43718216 | 43718442 | Intron |
| chr10 | 43722580 | 43723545 | Intron |
| chr10 | 43723610 | 43726840 | Promoter |
| chr10 | 43739748 | 43739966 | Intron |
| chr10 | 43740006 | 43740545 | Intron |
| chr10 | 43763351 | 43764240 | Promoter |
| chr10 | 43798609 | 43799292 | Distal Intergenic |
| chr10 | 43831600 | 43831954 | Distal Intergenic |
| chr10 | 43857681 | 43858235 | Distal Intergenic |

| | | | |
|---|---|---|---|
| chr10 | 43858286 | 43858485 | Distal Intergenic |
| chr10 | 43987693 | 43987892 | Distal Intergenic |
| chr10 | 44139651 | 44140443 | Promoter |
| chr10 | 44284631 | 44285059 | Promoter |
| chr10 | 44285151 | 44285611 | Promoter |
| chr10 | 44435211 | 44437476 | Promoter |
| chr10 | 44443191 | 44443486 | Intron |
| chr10 | 44475037 | 44475249 | Distal Intergenic |
| chr10 | 44652426 | 44652697 | Distal Intergenic |
| chr10 | 44723000 | 44723369 | Distal Intergenic |
| chr10 | 44786365 | 44786585 | Distal Intergenic |
| chr10 | 44787197 | 44790025 | Promoter |
| chr10 | 44989381 | 44989893 | Distal Intergenic |
| chr10 | 45117390 | 45118645 | Distal Intergenic |
| chr10 | 45301534 | 45302976 | Distal Intergenic |
| chr10 | 45355408 | 45355638 | Intron |
| chr10 | 45355744 | 45357042 | Intron |
| chr10 | 45357542 | 45361490 | Intron |
| chr10 | 45371116 | 45371315 | Intron |
| chr10 | 46504809 | 46506418 | Distal Intergenic |
| chr10 | 47786071 | 47787734 | Distal Intergenic |
| chr10 | 50311604 | 50311840 | 3' UTR |
| chr10 | 50311994 | 50312419 | Intron |
| chr10 | 50312702 | 50313189 | Intron |
| chr10 | 51290956 | 51291796 | Exon |
| chr10 | 51997329 | 51997888 | Intron |
| chr10 | 51997943 | 51998211 | Intron |
| chr10 | 52102144 | 52102423 | Intron |
| chr10 | 52137978 | 52139495 | Intron |
| chr10 | 53114599 | 53115516 | Intron |
| chr10 | 53316785 | 53317051 | Intron |
| chr10 | 53317513 | 53317719 | Intron |
| chr10 | 53317885 | 53318139 | Intron |
| chr10 | 53318235 | 53318807 | Intron |
| chr10 | 54070482 | 54070870 | Intron |
| chr10 | 55882043 | 55883106 | Intron |
| chr10 | 56096562 | 56096958 | Intron |
| chr10 | 56097075 | 56100040 | Intron |
| chr10 | 56100193 | 56100480 | Intron |
| chr10 | 56100546 | 56100932 | Intron |
| chr10 | 56145161 | 56145399 | Intron |
| chr10 | 56188910 | 56189109 | Intron |
| chr10 | 56205849 | 56207628 | Intron |
| chr10 | 56259722 | 56259972 | Intron |

| | | | |
|---|---|---|---|
| chr10 | 56260142 | 56260399 | Intron |
| chr10 | 56260435 | 56260660 | Intron |
| chr10 | 56279579 | 56281010 | Intron |
| chr10 | 56302081 | 56302280 | Intron |
| chr10 | 56329474 | 56332420 | Intron |
| chr10 | 56501760 | 56504371 | Intron |
| chr10 | 56664389 | 56664588 | Intron |
| chr10 | 56664701 | 56665319 | Intron |
| chr10 | 56713989 | 56715748 | Intron |
| chr10 | 57251337 | 57251714 | Intron |
| chr10 | 57251893 | 57252542 | Intron |
| chr10 | 57784003 | 57784476 | Distal Intergenic |
| chr10 | 57784785 | 57786005 | Distal Intergenic |
| chr10 | 57786343 | 57787281 | Distal Intergenic |
| chr10 | 57793231 | 57795569 | Distal Intergenic |
| chr10 | 57795674 | 57797008 | Distal Intergenic |
| chr10 | 57808005 | 57809644 | Distal Intergenic |
| chr10 | 57822779 | 57823055 | Distal Intergenic |
| chr10 | 57823234 | 57823464 | Distal Intergenic |
| chr10 | 57823506 | 57823754 | Distal Intergenic |
| chr10 | 57827371 | 57828065 | Distal Intergenic |
| chr10 | 57833022 | 57833442 | Distal Intergenic |
| chr10 | 57833804 | 57834615 | Distal Intergenic |
| chr10 | 57909618 | 57910206 | Distal Intergenic |
| chr10 | 57959374 | 57960868 | Distal Intergenic |
| chr10 | 58071638 | 58072661 | Distal Intergenic |
| chr10 | 58076523 | 58077992 | Distal Intergenic |
| chr10 | 58078947 | 58079365 | Distal Intergenic |
| chr10 | 58079403 | 58080418 | Distal Intergenic |
| chr10 | 58080617 | 58082104 | Distal Intergenic |
| chr10 | 58113907 | 58115603 | Downstream |
| chr10 | 58684882 | 58685081 | Distal Intergenic |
| chr10 | 58685905 | 58686700 | Distal Intergenic |
| chr10 | 58703696 | 58704730 | Distal Intergenic |
| chr10 | 58706427 | 58707705 | Distal Intergenic |
| chr10 | 58786554 | 58786980 | Distal Intergenic |
| chr10 | 59040497 | 59042193 | Distal Intergenic |
| chr10 | 59342514 | 59342883 | Distal Intergenic |
| chr10 | 59361790 | 59364868 | Distal Intergenic |
| chr10 | 59370112 | 59371639 | Distal Intergenic |
| chr10 | 59371748 | 59372532 | Distal Intergenic |
| chr10 | 59405977 | 59406646 | Distal Intergenic |
| chr10 | 59410406 | 59413240 | Distal Intergenic |
| chr10 | 59413287 | 59414050 | Distal Intergenic |

| | | | |
|---|---|---|---|
| chr10 | 59414145 | 59414575 | Distal Intergenic |
| chr10 | 59466103 | 59466361 | Distal Intergenic |
| chr10 | 59471619 | 59471829 | Distal Intergenic |
| chr10 | 59472046 | 59472706 | Distal Intergenic |
| chr10 | 59541560 | 59543236 | Distal Intergenic |
| chr10 | 59672291 | 59673528 | Distal Intergenic |
| chr10 | 59692959 | 59694294 | Distal Intergenic |
| chr10 | 59817062 | 59818417 | Distal Intergenic |
| chr10 | 59818606 | 59818805 | Distal Intergenic |
| chr10 | 60267866 | 60268095 | Distal Intergenic |
| chr10 | 60268182 | 60268740 | Distal Intergenic |
| chr10 | 60593476 | 60593882 | Distal Intergenic |
| chr10 | 60596084 | 60597027 | Distal Intergenic |
| chr10 | 60608609 | 60608827 | Distal Intergenic |
| chr10 | 60609548 | 60610294 | Distal Intergenic |
| chr10 | 60613070 | 60613628 | Distal Intergenic |
| chr10 | 60780291 | 60780945 | Distal Intergenic |
| chr10 | 60914308 | 60914634 | Distal Intergenic |
| chr10 | 61032669 | 61033005 | Intron |
| chr10 | 61044846 | 61046083 | Intron |
| chr10 | 61051586 | 61052250 | Intron |
| chr10 | 61090091 | 61090293 | Intron |
| chr10 | 61111334 | 61112148 | Promoter |
| chr10 | 61124586 | 61124939 | Distal Intergenic |
| chr10 | 61132264 | 61133114 | Distal Intergenic |
| chr10 | 61133152 | 61133399 | Distal Intergenic |
| chr10 | 61143081 | 61143319 | Distal Intergenic |
| chr10 | 61762477 | 61763643 | Distal Intergenic |
| chr10 | 61803096 | 61803339 | Intron |
| chr10 | 61911927 | 61913341 | Intron |
| chr10 | 61933390 | 61934283 | Intron |
| chr10 | 61948608 | 61949308 | Intron |
| chr10 | 62063923 | 62064125 | Intron |
| chr10 | 62064288 | 62065240 | Intron |
| chr10 | 62113226 | 62113457 | Intron |
| chr10 | 62113575 | 62113907 | Intron |
| chr10 | 62254369 | 62254606 | Intron |
| chr10 | 62255238 | 62255542 | Intron |
| chr10 | 62485158 | 62486579 | Intron |
| chr10 | 62487069 | 62487770 | Intron |
| chr10 | 62640225 | 62640792 | Intron |
| chr10 | 62706224 | 62706425 | Intron |
| chr10 | 62706852 | 62707246 | Intron |
| chr10 | 63304162 | 63304474 | Distal Intergenic |

| | | | |
|---|---|---|---|
| chr10 | 63306642 | 63308586 | Distal Intergenic |
| chr10 | 63486336 | 63486654 | Intron |
| chr10 | 63538271 | 63540160 | Distal Intergenic |
| chr10 | 64298694 | 64300237 | Intron |
| chr10 | 65802808 | 65803653 | Distal Intergenic |
| chr10 | 65816100 | 65818815 | Distal Intergenic |
| chr10 | 66312457 | 66313324 | Distal Intergenic |
| chr10 | 66326893 | 66327728 | Distal Intergenic |
| chr10 | 66383316 | 66384482 | Distal Intergenic |
| chr10 | 66438412 | 66438650 | Distal Intergenic |
| chr10 | 66465520 | 66466735 | Distal Intergenic |
| chr10 | 66472937 | 66474104 | Distal Intergenic |
| chr10 | 66559881 | 66562259 | Distal Intergenic |
| chr10 | 66569276 | 66570252 | Distal Intergenic |
| chr10 | 66617907 | 66620818 | Distal Intergenic |
| chr10 | 66647811 | 66650601 | Distal Intergenic |
| chr10 | 66681137 | 66682518 | Distal Intergenic |
| chr10 | 67308417 | 67309614 | Distal Intergenic |
| chr10 | 67334696 | 67335857 | Distal Intergenic |
| chr10 | 67361677 | 67362404 | Distal Intergenic |
| chr10 | 67446990 | 67447200 | Distal Intergenic |
| chr10 | 67470834 | 67471590 | Distal Intergenic |
| chr10 | 67502327 | 67502550 | Distal Intergenic |
| chr10 | 67502713 | 67503259 | Distal Intergenic |
| chr10 | 67565689 | 67568208 | Distal Intergenic |
| chr10 | 67599499 | 67599970 | Distal Intergenic |
| chr10 | 67618473 | 67621297 | Distal Intergenic |
| chr10 | 67628635 | 67629738 | Distal Intergenic |
| chr10 | 67709334 | 67710200 | Intron |
| chr10 | 67710366 | 67713093 | Intron |
| chr10 | 67735493 | 67737115 | Intron |
| chr10 | 67738016 | 67740383 | Intron |
| chr10 | 67740485 | 67741096 | Intron |
| chr10 | 67741686 | 67742135 | Intron |
| chr10 | 67743969 | 67744178 | Intron |
| chr10 | 67817932 | 67819271 | Intron |
| chr10 | 67909297 | 67909879 | Intron |
| chr10 | 68163773 | 68164828 | Intron |
| chr10 | 68168431 | 68169405 | Intron |
| chr10 | 68354897 | 68356562 | Intron |
| chr10 | 68360248 | 68360515 | Intron |
| chr10 | 68369849 | 68370601 | Intron |
| chr10 | 68452962 | 68453507 | Intron |
| chr10 | 68453639 | 68454047 | Intron |

| | | | |
|---|---|---|---|
| chr10 | 68648054 | 68649432 | Intron |
| chr10 | 68814950 | 68815692 | Intron |
| chr10 | 68815802 | 68816022 | Intron |
| chr10 | 68970402 | 68970637 | Intron |
| chr10 | 69239785 | 69240289 | Intron |
| chr10 | 69334771 | 69336126 | Intron |
| chr10 | 69425008 | 69425518 | Promoter |
| chr10 | 69557289 | 69559300 | Intron |
| chr10 | 69559347 | 69559721 | Intron |
| chr10 | 69725049 | 69728605 | 3' UTR |
| chr10 | 69729117 | 69729594 | Intron |
| chr10 | 70057193 | 70057524 | Promoter |
| chr10 | 70057675 | 70058916 | Promoter |
| chr10 | 70059027 | 70059328 | Intron |
| chr10 | 70160744 | 70161682 | 5' UTR |
| chr10 | 70261278 | 70263017 | Intron |
| chr10 | 70729273 | 70729503 | Intron |
| chr10 | 71330319 | 71330522 | Downstream |
| chr10 | 72096397 | 72096660 | Intron |
| chr10 | 72096820 | 72097868 | Intron |
| chr10 | 72116528 | 72118361 | Intron |
| chr10 | 72126472 | 72126784 | Intron |
| chr10 | 72701206 | 72701406 | Distal Intergenic |
| chr10 | 74000871 | 74002041 | Distal Intergenic |
| chr10 | 74125826 | 74126060 | Intron |
| chr10 | 74143052 | 74143277 | Intron |
| chr10 | 74533172 | 74533371 | Intron |
| chr10 | 74547999 | 74548815 | Intron |
| chr10 | 74555896 | 74556634 | Intron |
| chr10 | 74566289 | 74567493 | Intron |
| chr10 | 74567627 | 74568966 | Intron |
| chr10 | 74575372 | 74575586 | Intron |
| chr10 | 74592320 | 74592684 | Intron |
| chr10 | 74684789 | 74686237 | Intron |
| chr10 | 74980291 | 74980692 | Intron |
| chr10 | 74980746 | 74981083 | Intron |
| chr10 | 75011022 | 75011251 | Intron |
| chr10 | 75321700 | 75322032 | Intron |
| chr10 | 75322578 | 75322863 | Intron |
| chr10 | 75323097 | 75323451 | Intron |
| chr10 | 75335191 | 75336666 | Promoter |
| chr10 | 75341502 | 75342313 | Intron |
| chr10 | 75343590 | 75344595 | Intron |
| chr10 | 75361406 | 75361848 | Distal Intergenic |

| | | | |
|---|---|---|---|
| chr10 | 75384005 | 75384314 | Distal Intergenic |
| chr10 | 75384606 | 75384819 | Distal Intergenic |
| chr10 | 75494800 | 75495170 | Exon |
| chr10 | 75495483 | 75496170 | Distal Intergenic |
| chr10 | 75725594 | 75726139 | Distal Intergenic |
| chr10 | 75751160 | 75751368 | Distal Intergenic |
| chr10 | 75799478 | 75799990 | Intron |
| chr10 | 75808337 | 75808748 | Intron |
| chr10 | 75820028 | 75821957 | Intron |
| chr10 | 75999474 | 76000890 | Intron |
| chr10 | 76004731 | 76006158 | Intron |
| chr10 | 76336033 | 76337377 | Intron |
| chr10 | 76337416 | 76337769 | Intron |
| chr10 | 76365395 | 76365987 | Intron |
| chr10 | 76390094 | 76390294 | Intron |
| chr10 | 76393819 | 76394050 | Intron |
| chr10 | 76394097 | 76394456 | Intron |
| chr10 | 76546875 | 76547726 | Distal Intergenic |
| chr10 | 76552274 | 76553413 | Distal Intergenic |
| chr10 | 76574314 | 76574758 | Distal Intergenic |
| chr10 | 76584044 | 76584711 | Distal Intergenic |
| chr10 | 76588636 | 76589100 | Intron |
| chr10 | 76604889 | 76605088 | Intron |
| chr10 | 76614276 | 76615292 | Intron |
| chr10 | 76615904 | 76617362 | Intron |
| chr10 | 76624538 | 76625504 | Intron |
| chr10 | 76626844 | 76627058 | Intron |
| chr10 | 76628000 | 76628897 | Intron |
| chr10 | 76629996 | 76630939 | Intron |
| chr10 | 76663930 | 76665530 | Intron |
| chr10 | 76669218 | 76669811 | Intron |
| chr10 | 76673064 | 76674904 | Intron |
| chr10 | 76700276 | 76700644 | Intron |
| chr10 | 76700793 | 76701120 | Intron |
| chr10 | 76714759 | 76715293 | Intron |
| chr10 | 76741484 | 76742260 | Exon |
| chr10 | 76841886 | 76842417 | Distal Intergenic |
| chr10 | 76842486 | 76845757 | Distal Intergenic |
| chr10 | 76849306 | 76851663 | Downstream |
| chr10 | 76860664 | 76861561 | Intron |
| chr10 | 76869218 | 76869804 | Promoter |
| chr10 | 76873850 | 76874902 | Intron |
| chr10 | 76893438 | 76893654 | Intron |
| chr10 | 76928721 | 76929370 | Intron |

| | | | |
|---|---|---|---|
| chr10 | 76972154 | 76972505 | Intron |
| chr10 | 76972557 | 76973038 | Intron |
| chr10 | 77032413 | 77033852 | Distal Intergenic |
| chr10 | 77044808 | 77045007 | Distal Intergenic |
| chr10 | 77045053 | 77045291 | Distal Intergenic |
| chr10 | 77046265 | 77046522 | Distal Intergenic |
| chr10 | 77085707 | 77085995 | Intron |
| chr10 | 77100311 | 77100632 | Intron |
| chr10 | 77100835 | 77101426 | Intron |
| chr10 | 77171652 | 77172546 | Distal Intergenic |
| chr10 | 77436818 | 77437857 | Distal Intergenic |
| chr10 | 77438605 | 77439186 | Distal Intergenic |
| chr10 | 77439358 | 77439729 | Distal Intergenic |
| chr10 | 77471719 | 77472122 | Distal Intergenic |
| chr10 | 77472158 | 77472483 | Distal Intergenic |
| chr10 | 77624998 | 77626152 | Intron |
| chr10 | 77633787 | 77634144 | Intron |
| chr10 | 77646004 | 77646238 | Intron |
| chr10 | 77646824 | 77647023 | Intron |
| chr10 | 77647274 | 77647772 | Intron |
| chr10 | 77680728 | 77681200 | Intron |
| chr10 | 77681364 | 77681640 | Intron |
| chr10 | 77883509 | 77884591 | Intron |
| chr10 | 77888486 | 77889218 | Intron |
| chr10 | 77891072 | 77891483 | Intron |
| chr10 | 77891557 | 77892239 | Intron |
| chr10 | 78027950 | 78028197 | Intron |
| chr10 | 78037455 | 78037684 | Intron |
| chr10 | 78037745 | 78038416 | Intron |
| chr10 | 79145328 | 79145731 | Intron |
| chr10 | 79145771 | 79146427 | Intron |
| chr10 | 79146589 | 79146856 | Intron |
| chr10 | 79147051 | 79147250 | Intron |
| chr10 | 79167238 | 79167824 | Intron |
| chr10 | 79169576 | 79169974 | Intron |
| chr10 | 79170281 | 79171077 | Intron |
| chr10 | 79315634 | 79317286 | Intron |
| chr10 | 79317375 | 79317584 | Exon |
| chr10 | 79318375 | 79318574 | Exon |
| chr10 | 79319403 | 79319845 | Exon |
| chr10 | 79320914 | 79321201 | Intron |
| chr10 | 79371443 | 79371709 | Intron |
| chr10 | 79447704 | 79448050 | Distal Intergenic |
| chr10 | 79448304 | 79448714 | Distal Intergenic |

| | | | |
|---|---|---|---|
| chr10 | 79487092 | 79487315 | Distal Intergenic |
| chr10 | 79516690 | 79516925 | Distal Intergenic |
| chr10 | 79533997 | 79534261 | Distal Intergenic |
| chr10 | 79535420 | 79536049 | Distal Intergenic |
| chr10 | 79538166 | 79538640 | Distal Intergenic |
| chr10 | 79563631 | 79563842 | Intron |
| chr10 | 79564100 | 79564608 | Intron |
| chr10 | 79565222 | 79568141 | Exon |
| chr10 | 79610564 | 79611017 | Intron |
| chr10 | 79611684 | 79612077 | Intron |
| chr10 | 79612457 | 79612667 | Promoter |
| chr10 | 79613209 | 79614407 | Promoter |
| chr10 | 79614800 | 79615384 | Intron |
| chr10 | 79620958 | 79622543 | Intron |
| chr10 | 79622816 | 79623542 | Intron |
| chr10 | 79623587 | 79624763 | Intron |
| chr10 | 79624862 | 79625122 | Intron |
| chr10 | 79625919 | 79627150 | 5' UTR |
| chr10 | 79627285 | 79628153 | 3' UTR |
| chr10 | 79628329 | 79628528 | 3' UTR |
| chr10 | 79635629 | 79635896 | Intron |
| chr10 | 79712030 | 79712444 | Distal Intergenic |
| chr10 | 79713972 | 79715433 | Distal Intergenic |
| chr10 | 79723982 | 79724625 | Distal Intergenic |
| chr10 | 79799296 | 79799496 | Intron |
| chr10 | 79807163 | 79808548 | Intron |
| chr10 | 79848625 | 79849335 | Distal Intergenic |
| chr10 | 79898272 | 79898665 | Distal Intergenic |
| chr10 | 80024055 | 80024527 | Intron |
| chr10 | 80065991 | 80066221 | Intron |
| chr10 | 80086161 | 80086523 | Intron |
| chr10 | 80126215 | 80126562 | Intron |
| chr10 | 80126604 | 80127772 | Exon |
| chr10 | 80138052 | 80138293 | Intron |
| chr10 | 80139455 | 80141444 | Exon |
| chr10 | 80563795 | 80564482 | Distal Intergenic |
| chr10 | 80568023 | 80568596 | Distal Intergenic |
| chr10 | 80599435 | 80599678 | Distal Intergenic |
| chr10 | 80600631 | 80601515 | Distal Intergenic |
| chr10 | 80604153 | 80605167 | Distal Intergenic |
| chr10 | 80605224 | 80606147 | Distal Intergenic |
| chr10 | 80608127 | 80610261 | Distal Intergenic |
| chr10 | 80698041 | 80699047 | Distal Intergenic |
| chr10 | 80783664 | 80783910 | Intron |

| | | | |
|---|---|---|---|
| chr10 | 80847169 | 80847409 | Intron |
| chr10 | 80985792 | 80988012 | Intron |
| chr10 | 80988130 | 80988615 | Intron |
| chr10 | 81089898 | 81090756 | Distal Intergenic |
| chr10 | 81640861 | 81641086 | Distal Intergenic |
| chr10 | 81816956 | 81817449 | Intron |
| chr10 | 81967640 | 81968668 | Promoter |
| chr10 | 82092208 | 82092448 | Distal Intergenic |
| chr10 | 82092961 | 82093468 | Downstream |
| chr10 | 82225712 | 82225936 | Intron |
| chr10 | 82258097 | 82258432 | Intron |
| chr10 | 82261002 | 82262091 | Intron |
| chr10 | 82262298 | 82263237 | Intron |
| chr10 | 82306785 | 82307228 | Intron |
| chr10 | 82307306 | 82307606 | Intron |
| chr10 | 82321057 | 82324247 | Intron |
| chr10 | 82438770 | 82438992 | Distal Intergenic |
| chr10 | 82451934 | 82454176 | Distal Intergenic |
| chr10 | 82455383 | 82455610 | Distal Intergenic |
| chr10 | 82455744 | 82456080 | Distal Intergenic |
| chr10 | 82516906 | 82517537 | Distal Intergenic |
| chr10 | 82518666 | 82519750 | Distal Intergenic |
| chr10 | 82614923 | 82615860 | Distal Intergenic |
| chr10 | 82640868 | 82641157 | Distal Intergenic |
| chr10 | 83358351 | 83359033 | Distal Intergenic |
| chr10 | 83360181 | 83360726 | Distal Intergenic |
| chr10 | 83361006 | 83361350 | Distal Intergenic |
| chr10 | 83371294 | 83371779 | Distal Intergenic |
| chr10 | 83371894 | 83372206 | Distal Intergenic |
| chr10 | 83431319 | 83432140 | Distal Intergenic |
| chr10 | 83432183 | 83434317 | Distal Intergenic |
| chr10 | 83447532 | 83447911 | Distal Intergenic |
| chr10 | 83448117 | 83448474 | Distal Intergenic |
| chr10 | 83466113 | 83466583 | Distal Intergenic |
| chr10 | 83486134 | 83487445 | Distal Intergenic |
| chr10 | 83519230 | 83519439 | Distal Intergenic |
| chr10 | 83528846 | 83529780 | Distal Intergenic |
| chr10 | 83542338 | 83544234 | Distal Intergenic |
| chr10 | 83563288 | 83564804 | Distal Intergenic |
| chr10 | 83577609 | 83578226 | Distal Intergenic |
| chr10 | 83584466 | 83585251 | Distal Intergenic |
| chr10 | 83656664 | 83657703 | Intron |
| chr10 | 83697255 | 83697661 | Intron |
| chr10 | 83697733 | 83697932 | Intron |

| | | | |
|---|---|---|---|
| chr10 | 83701512 | 83701899 | Intron |
| chr10 | 83855495 | 83855912 | Intron |
| chr10 | 83903751 | 83904418 | Intron |
| chr10 | 83994856 | 83996297 | Intron |
| chr10 | 84010448 | 84010686 | Intron |
| chr10 | 84010931 | 84011404 | Intron |
| chr10 | 84039379 | 84039591 | Intron |
| chr10 | 84084658 | 84085532 | Intron |
| chr10 | 84152442 | 84152783 | Intron |
| chr10 | 84817696 | 84818421 | Distal Intergenic |
| chr10 | 84829211 | 84830379 | Distal Intergenic |
| chr10 | 84834269 | 84835703 | Distal Intergenic |
| chr10 | 84851876 | 84852618 | Distal Intergenic |
| chr10 | 84852718 | 84853765 | Distal Intergenic |
| chr10 | 84883647 | 84883930 | Distal Intergenic |
| chr10 | 84914160 | 84914905 | Distal Intergenic |
| chr10 | 84931961 | 84932164 | Distal Intergenic |
| chr10 | 84944660 | 84945369 | Distal Intergenic |
| chr10 | 84948020 | 84949834 | Distal Intergenic |
| chr10 | 84957378 | 84958411 | Distal Intergenic |
| chr10 | 84978238 | 84978964 | Distal Intergenic |
| chr10 | 84985101 | 84985462 | Distal Intergenic |
| chr10 | 84989352 | 84989631 | Distal Intergenic |
| chr10 | 84990395 | 84990654 | Distal Intergenic |
| chr10 | 85040001 | 85041681 | Distal Intergenic |
| chr10 | 85188122 | 85188781 | Distal Intergenic |
| chr10 | 85435655 | 85435987 | Distal Intergenic |
| chr10 | 85451594 | 85452855 | Distal Intergenic |
| chr10 | 85473289 | 85475048 | Distal Intergenic |
| chr10 | 85485010 | 85485473 | Distal Intergenic |
| chr10 | 85558485 | 85559369 | Distal Intergenic |
| chr10 | 85559445 | 85559644 | Distal Intergenic |
| chr10 | 85563246 | 85563481 | Distal Intergenic |
| chr10 | 85633378 | 85634400 | Distal Intergenic |
| chr10 | 85635841 | 85636899 | Distal Intergenic |
| chr10 | 85660972 | 85661896 | Distal Intergenic |
| chr10 | 85748796 | 85749559 | Distal Intergenic |
| chr10 | 85753191 | 85753416 | Distal Intergenic |
| chr10 | 85801361 | 85804132 | Distal Intergenic |
| chr10 | 85913664 | 85914564 | Distal Intergenic |
| chr10 | 86074814 | 86076091 | Distal Intergenic |
| chr10 | 86167931 | 86168130 | Intron |
| chr10 | 86168575 | 86169585 | Intron |
| chr10 | 87268466 | 87268747 | Distal Intergenic |

| | | | |
|---|---|---|---|
| chr10 | 87268930 | 87269168 | Distal Intergenic |
| chr10 | 87343011 | 87343428 | Intron |
| chr10 | 87609696 | 87610090 | Intron |
| chr10 | 87612818 | 87613550 | Intron |
| chr10 | 87663951 | 87664268 | Intron |
| chr10 | 87808797 | 87809728 | Intron |
| chr10 | 87868386 | 87868611 | Intron |
| chr10 | 87868722 | 87868930 | Intron |
| chr10 | 87977065 | 87977404 | Intron |
| chr10 | 88190796 | 88191032 | Distal Intergenic |
| chr10 | 88288745 | 88289204 | Distal Intergenic |
| chr10 | 88289327 | 88289526 | Distal Intergenic |
| chr10 | 88433861 | 88434300 | Intron |
| chr10 | 88905397 | 88905838 | Intron |
| chr10 | 88906166 | 88906389 | Intron |
| chr10 | 88989577 | 88990204 | Intron |
| chr10 | 89013736 | 89015365 | Intron |
| chr10 | 89062044 | 89062634 | Intron |
| chr10 | 89067065 | 89068056 | Promoter |
| chr10 | 89208291 | 89208582 | Distal Intergenic |
| chr10 | 89208655 | 89209251 | Distal Intergenic |
| chr10 | 89531618 | 89531842 | Intron |
| chr10 | 89655878 | 89656741 | Intron |
| chr10 | 89843054 | 89843566 | Distal Intergenic |
| chr10 | 90555145 | 90555691 | Distal Intergenic |
| chr10 | 90585348 | 90586192 | Exon |
| chr10 | 90586454 | 90586822 | Intron |
| chr10 | 90586899 | 90587718 | Intron |
| chr10 | 90619263 | 90619764 | Distal Intergenic |
| chr10 | 90619806 | 90620017 | Distal Intergenic |
| chr10 | 90620616 | 90622263 | Distal Intergenic |
| chr10 | 90629760 | 90630715 | Distal Intergenic |
| chr10 | 90630872 | 90631489 | Distal Intergenic |
| chr10 | 90631572 | 90631809 | Distal Intergenic |
| chr10 | 90632014 | 90632601 | Distal Intergenic |
| chr10 | 90633021 | 90633271 | Distal Intergenic |
| chr10 | 90634763 | 90635051 | Distal Intergenic |
| chr10 | 90643201 | 90644503 | Promoter |
| chr10 | 90644962 | 90646731 | Promoter |
| chr10 | 90696623 | 90698550 | Exon |
| chr10 | 90703727 | 90704195 | Intron |
| chr10 | 90706660 | 90707165 | 5' UTR |
| chr10 | 92394875 | 92395784 | Distal Intergenic |
| chr10 | 92405580 | 92406012 | Distal Intergenic |

| | | | |
|---|---|---|---|
| chr10 | 92532057 | 92532404 | Intron |
| chr10 | 92557708 | 92558324 | Intron |
| chr10 | 92570525 | 92570748 | Intron |
| chr10 | 92600951 | 92601644 | Intron |
| chr10 | 92623634 | 92624598 | Distal Intergenic |
| chr10 | 92624635 | 92625403 | Distal Intergenic |
| chr10 | 92648723 | 92649445 | Intron |
| chr10 | 92650265 | 92652255 | Intron |
| chr10 | 92723895 | 92725174 | Distal Intergenic |
| chr10 | 92725229 | 92725581 | Distal Intergenic |
| chr10 | 92730673 | 92732291 | Distal Intergenic |
| chr10 | 92733342 | 92736659 | Distal Intergenic |
| chr10 | 92750310 | 92751482 | Distal Intergenic |
| chr10 | 92768126 | 92772075 | Distal Intergenic |
| chr10 | 92772128 | 92773388 | Distal Intergenic |
| chr10 | 92779430 | 92780021 | Distal Intergenic |
| chr10 | 92805291 | 92807249 | Promoter |
| chr10 | 92819503 | 92821054 | Intron |
| chr10 | 92884618 | 92886716 | Distal Intergenic |
| chr10 | 92887999 | 92888594 | Distal Intergenic |
| chr10 | 92971452 | 92973663 | Intron |
| chr10 | 93104321 | 93106247 | Intron |
| chr10 | 93110544 | 93111819 | Exon |
| chr10 | 93123526 | 93124255 | Intron |
| chr10 | 93124295 | 93124766 | Intron |
| chr10 | 93139578 | 93139812 | Intron |
| chr10 | 93139929 | 93140128 | Intron |
| chr10 | 93245998 | 93246229 | Intron |
| chr10 | 93351627 | 93352801 | Intron |
| chr10 | 93395985 | 93396872 | Distal Intergenic |
| chr10 | 93510710 | 93511334 | Distal Intergenic |
| chr10 | 93521990 | 93522847 | Distal Intergenic |
| chr10 | 93544576 | 93545313 | Distal Intergenic |
| chr10 | 93545357 | 93545640 | Distal Intergenic |
| chr10 | 93579324 | 93581610 | Exon |
| chr10 | 93611049 | 93611410 | Exon |
| chr10 | 93637199 | 93638233 | Distal Intergenic |
| chr10 | 93661305 | 93661727 | Distal Intergenic |
| chr10 | 93843331 | 93844183 | Intron |
| chr10 | 93979650 | 93980115 | Intron |
| chr10 | 93987400 | 93988302 | Intron |
| chr10 | 94007573 | 94009478 | Intron |
| chr10 | 94013473 | 94014454 | Intron |
| chr10 | 94014571 | 94014845 | Intron |

| | | | |
|---|---|---|---|
| chr10 | 94015180 | 94015408 | Intron |
| chr10 | 94016579 | 94019455 | Intron |
| chr10 | 94020848 | 94021133 | Intron |
| chr10 | 94023620 | 94024580 | Intron |
| chr10 | 94032485 | 94032885 | Intron |
| chr10 | 94033044 | 94033317 | Intron |
| chr10 | 94033546 | 94033839 | Intron |
| chr10 | 94151925 | 94152203 | Distal Intergenic |
| chr10 | 94675948 | 94676469 | Intron |
| chr10 | 94744629 | 94744925 | Intron |
| chr10 | 94902663 | 94902901 | Distal Intergenic |
| chr10 | 94920239 | 94921517 | Distal Intergenic |
| chr10 | 94921714 | 94921938 | Distal Intergenic |
| chr10 | 94970533 | 94972332 | Distal Intergenic |
| chr10 | 94977699 | 94979089 | Distal Intergenic |
| chr10 | 94986116 | 94986338 | Distal Intergenic |
| chr10 | 94986681 | 94988674 | Distal Intergenic |
| chr10 | 94988799 | 94989047 | Distal Intergenic |
| chr10 | 94989172 | 94990247 | Distal Intergenic |
| chr10 | 95093627 | 95093866 | Exon |
| chr10 | 95094471 | 95094787 | Intron |
| chr10 | 95324724 | 95326112 | Promoter |
| chr10 | 95326674 | 95327462 | Promoter |
| chr10 | 95346345 | 95347212 | Exon |
| chr10 | 95583559 | 95585311 | Distal Intergenic |
| chr10 | 95693989 | 95694290 | Intron |
| chr10 | 95878183 | 95879681 | Intron |
| chr10 | 96079692 | 96079957 | Intron |
| chr10 | 96080216 | 96080792 | Intron |
| chr10 | 96080828 | 96081058 | Intron |
| chr10 | 96090667 | 96091774 | Downstream |
| chr10 | 96095351 | 96095888 | Intron |
| chr10 | 96160308 | 96160517 | Distal Intergenic |
| chr10 | 96160556 | 96160902 | Distal Intergenic |
| chr10 | 96184710 | 96184909 | Intron |
| chr10 | 96184966 | 96185191 | Intron |
| chr10 | 96776013 | 96776256 | Distal Intergenic |
| chr10 | 97795048 | 97795404 | Intron |
| chr10 | 97795547 | 97795784 | Intron |
| chr10 | 97971428 | 97971953 | Intron |
| chr10 | 98406688 | 98406969 | Intron |
| chr10 | 98413905 | 98414359 | Intron |
| chr10 | 98416918 | 98417295 | Intron |
| chr10 | 98417342 | 98417628 | Intron |

| | | | |
|---|---|---|---|
| chr10 | 98436459 | 98437606 | Intron |
| chr10 | 98525163 | 98526063 | Distal Intergenic |
| chr10 | 98544277 | 98544788 | Distal Intergenic |
| chr10 | 98635967 | 98636166 | Intron |
| chr10 | 98699722 | 98700473 | Intron |
| chr10 | 98704101 | 98704866 | Intron |
| chr10 | 98704914 | 98705398 | Intron |
| chr10 | 98705577 | 98705808 | Intron |
| chr10 | 98705964 | 98706563 | Intron |
| chr10 | 98724177 | 98725641 | 3' UTR |
| chr10 | 98725889 | 98726207 | Intron |
| chr10 | 98771426 | 98771762 | Intron |
| chr10 | 98772013 | 98772449 | Intron |
| chr10 | 98853720 | 98853953 | Intron |
| chr10 | 98854048 | 98854921 | Intron |
| chr10 | 98855444 | 98855653 | Intron |
| chr10 | 98858528 | 98860150 | Intron |
| chr10 | 99372616 | 99372815 | Intron |
| chr10 | 99372861 | 99373257 | Intron |
| chr10 | 99495138 | 99495570 | Distal Intergenic |
| chr10 | 99923311 | 99924264 | Intron |
| chr10 | 99946760 | 99949957 | Intron |
| chr10 | 100029971 | 100030363 | Distal Intergenic |
| chr10 | 102122103 | 102122341 | 3' UTR |
| chr10 | 102122470 | 102122969 | 3' UTR |
| chr10 | 104960448 | 104961066 | Intron |
| chr10 | 106212125 | 106212399 | Intron |
| chr10 | 106212567 | 106213523 | Intron |
| chr10 | 107095182 | 107095537 | Distal Intergenic |
| chr10 | 107095610 | 107096631 | Distal Intergenic |
| chr10 | 107266108 | 107266369 | Distal Intergenic |
| chr10 | 107429495 | 107429801 | Distal Intergenic |
| chr10 | 107599404 | 107599645 | Distal Intergenic |
| chr10 | 107607569 | 107607998 | Distal Intergenic |
| chr10 | 107609698 | 107610623 | Distal Intergenic |
| chr10 | 107621692 | 107623892 | Distal Intergenic |
| chr10 | 107638419 | 107638778 | Distal Intergenic |
| chr10 | 107730761 | 107731824 | Distal Intergenic |
| chr10 | 107734518 | 107734979 | Distal Intergenic |
| chr10 | 107735244 | 107735902 | Distal Intergenic |
| chr10 | 107744761 | 107745852 | Distal Intergenic |
| chr10 | 107766998 | 107768931 | Distal Intergenic |
| chr10 | 107769443 | 107769670 | Distal Intergenic |
| chr10 | 107774899 | 107775263 | Distal Intergenic |

| | | | |
|---|---|---|---|
| chr10 | 107812782 | 107813202 | Distal Intergenic |
| chr10 | 107813518 | 107813717 | Distal Intergenic |
| chr10 | 107813912 | 107814111 | Distal Intergenic |
| chr10 | 107814155 | 107816250 | Distal Intergenic |
| chr10 | 109255019 | 109255524 | Distal Intergenic |
| chr10 | 109340463 | 109341420 | Distal Intergenic |
| chr10 | 109760607 | 109761333 | Distal Intergenic |
| chr10 | 109761548 | 109761851 | Distal Intergenic |
| chr10 | 110671038 | 110671468 | Distal Intergenic |
| chr10 | 110943706 | 110944289 | Distal Intergenic |
| chr10 | 110944761 | 110945244 | Distal Intergenic |
| chr10 | 110945423 | 110945900 | Distal Intergenic |
| chr10 | 110946146 | 110946780 | Distal Intergenic |
| chr10 | 110952255 | 110954513 | Distal Intergenic |
| chr10 | 110976780 | 110977034 | Distal Intergenic |
| chr10 | 110980385 | 110981258 | Distal Intergenic |
| chr10 | 110981346 | 110981547 | Distal Intergenic |
| chr10 | 110981656 | 110981869 | Distal Intergenic |
| chr10 | 111006311 | 111006703 | Distal Intergenic |
| chr10 | 111007225 | 111008599 | Distal Intergenic |
| chr10 | 111012678 | 111014239 | Distal Intergenic |
| chr10 | 111019976 | 111020261 | Distal Intergenic |
| chr10 | 111020371 | 111020748 | Distal Intergenic |
| chr10 | 111020876 | 111021759 | Distal Intergenic |
| chr10 | 111021802 | 111022140 | Distal Intergenic |
| chr10 | 111055076 | 111057663 | Distal Intergenic |
| chr10 | 111057949 | 111061967 | Distal Intergenic |
| chr10 | 111072608 | 111074996 | Distal Intergenic |
| chr10 | 111090982 | 111094193 | Distal Intergenic |
| chr10 | 111109432 | 111110476 | Distal Intergenic |
| chr10 | 111121819 | 111122603 | Distal Intergenic |
| chr10 | 111128622 | 111129146 | Distal Intergenic |
| chr10 | 111150534 | 111150767 | Distal Intergenic |
| chr10 | 111151682 | 111152705 | Distal Intergenic |
| chr10 | 111279809 | 111280033 | Distal Intergenic |
| chr10 | 111280113 | 111280724 | Distal Intergenic |
| chr10 | 111432699 | 111433898 | Distal Intergenic |
| chr10 | 111918771 | 111919140 | Distal Intergenic |
| chr10 | 112365513 | 112366015 | Distal Intergenic |
| chr10 | 114752891 | 114753144 | Intron |
| chr10 | 114768672 | 114770076 | Intron |
| chr10 | 114785973 | 114787013 | Intron |
| chr10 | 114791603 | 114792058 | Intron |
| chr10 | 114799604 | 114800167 | Promoter |

| | | | |
|---|---|---|---|
| chr10 | 114821370 | 114821609 | Intron |
| chr10 | 114821990 | 114822238 | Intron |
| chr10 | 114822448 | 114822726 | Intron |
| chr10 | 114822826 | 114823263 | Intron |
| chr10 | 115675790 | 115676217 | Exon |
| chr10 | 115703355 | 115704920 | Distal Intergenic |
| chr10 | 115704974 | 115705600 | Distal Intergenic |
| chr10 | 115705726 | 115705955 | Distal Intergenic |
| chr10 | 115965606 | 115966748 | Exon |
| chr10 | 115973372 | 115974683 | Exon |
| chr10 | 116017041 | 116018087 | Intron |
| chr10 | 116018457 | 116018829 | Intron |
| chr10 | 116130455 | 116131535 | Intron |
| chr10 | 116143503 | 116144545 | Intron |
| chr10 | 116144587 | 116144821 | Intron |
| chr10 | 116162384 | 116162622 | Intron |
| chr10 | 116170090 | 116171598 | Distal Intergenic |
| chr10 | 116185572 | 116187508 | Distal Intergenic |
| chr10 | 116203800 | 116204100 | Exon |
| chr10 | 116205787 | 116206041 | Intron |
| chr10 | 116206165 | 116206959 | Intron |
| chr10 | 116208010 | 116209383 | Intron |
| chr10 | 116433052 | 116433270 | Intron |
| chr10 | 116471804 | 116472841 | Intron |
| chr10 | 116500381 | 116500847 | Intron |
| chr10 | 116501029 | 116501239 | Intron |
| chr10 | 116812751 | 116812991 | Distal Intergenic |
| chr10 | 117146014 | 117147386 | Intron |
| chr10 | 117147543 | 117147743 | Intron |
| chr10 | 117147971 | 117148190 | Intron |
| chr10 | 117148255 | 117149103 | Intron |
| chr10 | 117154569 | 117155229 | Intron |
| chr10 | 117158445 | 117158697 | Intron |
| chr10 | 117158733 | 117159258 | Intron |
| chr10 | 117281724 | 117282192 | Intron |
| chr10 | 118246546 | 118247022 | Distal Intergenic |
| chr10 | 118425001 | 118425764 | 3' UTR |
| chr10 | 118425802 | 118426050 | Intron |
| chr10 | 118428650 | 118428926 | Promoter |
| chr10 | 118437985 | 118438946 | Intron |
| chr10 | 118441128 | 118441344 | Exon |
| chr10 | 118676933 | 118677139 | Intron |
| chr10 | 118796173 | 118796780 | Intron |
| chr10 | 118805169 | 118806409 | Intron |

| | | | |
|---|---|---|---|
| chr10 | 118881648 | 118882444 | Intron |
| chr10 | 118883676 | 118884437 | Intron |
| chr10 | 118891309 | 118891521 | 3' UTR |
| chr10 | 119247296 | 119248684 | Exon |
| chr10 | 119313957 | 119314384 | Distal Intergenic |
| chr10 | 119641676 | 119642201 | Distal Intergenic |
| chr10 | 119646377 | 119646636 | Distal Intergenic |
| chr10 | 119663952 | 119664262 | Distal Intergenic |
| chr10 | 119664619 | 119664997 | Distal Intergenic |
| chr10 | 119745746 | 119746072 | Distal Intergenic |
| chr10 | 119746340 | 119746777 | Distal Intergenic |
| chr10 | 119746829 | 119747143 | Distal Intergenic |
| chr10 | 119911720 | 119912018 | Intron |
| chr10 | 120153968 | 120154663 | Distal Intergenic |
| chr10 | 121024136 | 121024353 | Intron |
| chr10 | 121062798 | 121062997 | Intron |
| chr10 | 121063156 | 121063355 | Intron |
| chr10 | 121340825 | 121341446 | 5' UTR |
| chr10 | 122244388 | 122244638 | Intron |
| chr10 | 122247841 | 122248739 | Intron |
| chr10 | 122299062 | 122299438 | Intron |
| chr10 | 122299493 | 122299805 | Intron |
| chr10 | 122306858 | 122307236 | Intron |
| chr10 | 122307287 | 122308157 | Intron |
| chr10 | 122308205 | 122308873 | Intron |
| chr10 | 122328665 | 122328955 | Intron |
| chr10 | 122330024 | 122330480 | Intron |
| chr10 | 122330633 | 122331260 | Intron |
| chr10 | 122331537 | 122331832 | Intron |
| chr10 | 122421256 | 122421522 | Distal Intergenic |
| chr10 | 122421646 | 122422494 | Distal Intergenic |
| chr10 | 122423422 | 122424017 | Distal Intergenic |
| chr10 | 122431814 | 122432159 | Distal Intergenic |
| chr10 | 122442035 | 122442425 | Distal Intergenic |
| chr10 | 126540176 | 126540665 | Distal Intergenic |
| chr10 | 126563633 | 126563884 | Distal Intergenic |
| chr10 | 126581178 | 126581965 | Distal Intergenic |
| chr10 | 126726470 | 126727273 | Intron |
| chr10 | 126867880 | 126868250 | Distal Intergenic |
| chr10 | 127000106 | 127000557 | Distal Intergenic |
| chr10 | 127027659 | 127028934 | Distal Intergenic |
| chr10 | 127029426 | 127029947 | Distal Intergenic |
| chr10 | 127030064 | 127030347 | Distal Intergenic |
| chr10 | 127033693 | 127034133 | Distal Intergenic |

| | | | |
|---|---|---|---|
| chr10 | 127034423 | 127034707 | Distal Intergenic |
| chr10 | 127058379 | 127058735 | Distal Intergenic |
| chr10 | 127072493 | 127073661 | Distal Intergenic |
| chr10 | 127079224 | 127081279 | Distal Intergenic |
| chr10 | 127104011 | 127104254 | Distal Intergenic |
| chr10 | 127104435 | 127105113 | Distal Intergenic |
| chr10 | 127105311 | 127105918 | Distal Intergenic |
| chr10 | 127341518 | 127342349 | Intron |
| chr10 | 127359150 | 127360968 | Intron |
| chr10 | 127361086 | 127363081 | Intron |
| chr10 | 127364672 | 127364977 | Intron |
| chr10 | 127472420 | 127472821 | Distal Intergenic |
| chr10 | 127473123 | 127473372 | Distal Intergenic |
| chr10 | 127473488 | 127474341 | Downstream |
| chr10 | 127496199 | 127497091 | Promoter |
| chr10 | 127520331 | 127520565 | Intron |
| chr10 | 127527821 | 127529807 | Exon |
| chr10 | 127529870 | 127530387 | 3' UTR |
| chr10 | 127531298 | 127531725 | 3' UTR |
| chr10 | 127541185 | 127542012 | 5' UTR |
| chr10 | 127563348 | 127563598 | Intron |
| chr10 | 127613067 | 127613515 | Intron |
| chr10 | 127613596 | 127613852 | Intron |
| chr10 | 127622538 | 127622825 | Intron |
| chr10 | 127734933 | 127735423 | Intron |
| chr10 | 127740425 | 127740645 | Intron |
| chr10 | 127759513 | 127759885 | Intron |
| chr10 | 127775359 | 127775852 | Intron |
| chr10 | 127781727 | 127782658 | Exon |
| chr10 | 127847614 | 127847850 | Intron |
| chr10 | 127983617 | 127984384 | Intron |
| chr10 | 127985789 | 127986192 | Intron |
| chr10 | 128063356 | 128063825 | Intron |
| chr10 | 128178951 | 128180460 | Intron |
| chr10 | 128184594 | 128184794 | Intron |
| chr10 | 128184868 | 128185235 | Intron |
| chr10 | 128259064 | 128259268 | Intron |
| chr10 | 128358608 | 128359490 | Promoter |
| chr10 | 128359642 | 128360080 | Promoter |
| chr11 | 3023678 | 3024358 | Exon |
| chr11 | 3051313 | 3051530 | Exon |
| chr11 | 3052313 | 3053140 | Exon |
| chr11 | 3053859 | 3054274 | Intron |
| chr11 | 3056938 | 3058767 | Intron |

| | | | |
|---|---|---|---|
| chr11 | 3166276 | 3166621 | Intron |
| chr11 | 3232608 | 3232807 | Distal Intergenic |
| chr11 | 3234662 | 3235849 | Distal Intergenic |
| chr11 | 3236827 | 3237026 | Downstream |
| chr11 | 3237312 | 3238077 | Downstream |
| chr11 | 3238165 | 3238607 | Promoter |
| chr11 | 3238682 | 3239384 | Promoter |
| chr11 | 3248708 | 3249463 | 3' UTR |
| chr11 | 3249510 | 3249714 | Exon |
| chr11 | 3265487 | 3265820 | Distal Intergenic |
| chr11 | 3265864 | 3268237 | Distal Intergenic |
| chr11 | 3268687 | 3272279 | Distal Intergenic |
| chr11 | 3272365 | 3272666 | Distal Intergenic |
| chr11 | 3273648 | 3275451 | Distal Intergenic |
| chr11 | 3308388 | 3309022 | Distal Intergenic |
| chr11 | 3337294 | 3338647 | Distal Intergenic |
| chr11 | 3342959 | 3343644 | Distal Intergenic |
| chr11 | 3351244 | 3351483 | Distal Intergenic |
| chr11 | 3351604 | 3352975 | Distal Intergenic |
| chr11 | 3438088 | 3439398 | Distal Intergenic |
| chr11 | 3459594 | 3459987 | Distal Intergenic |
| chr11 | 3460145 | 3460582 | Distal Intergenic |
| chr11 | 3467286 | 3467643 | Distal Intergenic |
| chr11 | 3467898 | 3468473 | Distal Intergenic |
| chr11 | 3515523 | 3516060 | Distal Intergenic |
| chr11 | 3516172 | 3516457 | Distal Intergenic |
| chr11 | 3547887 | 3548086 | Distal Intergenic |
| chr11 | 3548435 | 3549156 | Distal Intergenic |
| chr11 | 3568298 | 3569815 | Distal Intergenic |
| chr11 | 3571037 | 3571713 | Distal Intergenic |
| chr11 | 3574629 | 3575059 | Distal Intergenic |
| chr11 | 3732311 | 3732773 | Intron |
| chr11 | 3793793 | 3795662 | Exon |
| chr11 | 3804381 | 3806403 | Intron |
| chr11 | 3808843 | 3809937 | Intron |
| chr11 | 3811132 | 3811603 | Intron |
| chr11 | 3811978 | 3812634 | Intron |
| chr11 | 3813021 | 3814856 | Intron |
| chr11 | 3863542 | 3864744 | Distal Intergenic |
| chr11 | 3907581 | 3907982 | Intron |
| chr11 | 4002119 | 4002468 | Intron |
| chr11 | 4060115 | 4061096 | Intron |
| chr11 | 4087035 | 4087730 | Intron |
| chr11 | 4087938 | 4089895 | Intron |

| | | | |
|---|---|---|---|
| chr11 | 4089941 | 4091992 | Exon |
| chr11 | 4094852 | 4096530 | Exon |
| chr11 | 4097554 | 4097897 | Intron |
| chr11 | 4097941 | 4100163 | Intron |
| chr11 | 4115779 | 4118389 | Promoter |
| chr11 | 4118552 | 4118833 | Intron |
| chr11 | 4160682 | 4160972 | Intron |
| chr11 | 4161222 | 4163256 | Intron |
| chr11 | 4163484 | 4167652 | Intron |
| chr11 | 4167711 | 4168000 | Intron |
| chr11 | 4198837 | 4199709 | Intron |
| chr11 | 4199748 | 4199997 | Intron |
| chr11 | 4202109 | 4204135 | Intron |
| chr11 | 4207326 | 4209991 | Promoter |
| chr11 | 4244909 | 4245127 | Distal Intergenic |
| chr11 | 4301235 | 4302069 | Distal Intergenic |
| chr11 | 4389204 | 4390899 | Promoter |
| chr11 | 4478128 | 4479397 | Distal Intergenic |
| chr11 | 4493852 | 4494478 | Distal Intergenic |
| chr11 | 4494565 | 4494806 | Distal Intergenic |
| chr11 | 4505622 | 4507797 | Distal Intergenic |
| chr11 | 4537069 | 4537550 | Distal Intergenic |
| chr11 | 4537699 | 4538724 | Distal Intergenic |
| chr11 | 4561201 | 4561447 | Distal Intergenic |
| chr11 | 4561570 | 4561788 | Distal Intergenic |
| chr11 | 4603988 | 4604618 | Distal Intergenic |
| chr11 | 4629433 | 4630082 | Promoter |
| chr11 | 4738288 | 4738487 | Distal Intergenic |
| chr11 | 4743659 | 4743980 | Distal Intergenic |
| chr11 | 4748844 | 4749567 | Distal Intergenic |
| chr11 | 4749615 | 4750234 | Distal Intergenic |
| chr11 | 4760009 | 4762134 | Distal Intergenic |
| chr11 | 4762354 | 4762653 | Distal Intergenic |
| chr11 | 4795070 | 4795796 | Distal Intergenic |
| chr11 | 4799344 | 4800471 | Distal Intergenic |
| chr11 | 4909548 | 4910377 | Distal Intergenic |
| chr11 | 4922611 | 4922923 | Distal Intergenic |
| chr11 | 4937526 | 4938373 | Promoter |
| chr11 | 4942849 | 4943513 | Downstream |
| chr11 | 4964420 | 4964817 | Downstream |
| chr11 | 4965049 | 4965745 | Downstream |
| chr11 | 4997065 | 4997338 | Distal Intergenic |
| chr11 | 4997617 | 5001987 | Distal Intergenic |
| chr11 | 5140212 | 5140411 | Distal Intergenic |

| | | | |
|---|---|---|---|
| chr11 | 5144460 | 5145353 | Exon |
| chr11 | 5145400 | 5145602 | Intron |
| chr11 | 5161794 | 5163739 | Distal Intergenic |
| chr11 | 5171305 | 5172503 | Downstream |
| chr11 | 5195142 | 5195564 | Distal Intergenic |
| chr11 | 5255080 | 5255884 | Promoter |
| chr11 | 5298525 | 5298788 | Intron |
| chr11 | 5327281 | 5327499 | Intron |
| chr11 | 5332315 | 5332735 | Intron |
| chr11 | 5332839 | 5333360 | Intron |
| chr11 | 5334078 | 5334477 | Intron |
| chr11 | 5334538 | 5334745 | Intron |
| chr11 | 5420936 | 5422203 | Intron |
| chr11 | 5441431 | 5441685 | Intron |
| chr11 | 5441947 | 5442216 | Intron |
| chr11 | 5637144 | 5637767 | Intron |
| chr11 | 5641352 | 5641594 | Promoter |
| chr11 | 5661937 | 5662136 | Intron |
| chr11 | 5662180 | 5662379 | Intron |
| chr11 | 5687978 | 5688485 | Intron |
| chr11 | 5688530 | 5689141 | Exon |
| chr11 | 5697896 | 5698348 | Intron |
| chr11 | 5734946 | 5735781 | Intron |
| chr11 | 5736549 | 5738017 | Intron |
| chr11 | 5796110 | 5796395 | Intron |
| chr11 | 5855783 | 5856701 | Intron |
| chr11 | 6068591 | 6070300 | Distal Intergenic |
| chr11 | 6099982 | 6100662 | Distal Intergenic |
| chr11 | 6167510 | 6168236 | Distal Intergenic |
| chr11 | 6191919 | 6192291 | Promoter |
| chr11 | 6192493 | 6192768 | Promoter |
| chr11 | 6246655 | 6247412 | Intron |
| chr11 | 6277304 | 6277593 | Distal Intergenic |
| chr11 | 6314289 | 6316049 | Distal Intergenic |
| chr11 | 6316452 | 6316859 | Distal Intergenic |
| chr11 | 6316925 | 6317648 | Distal Intergenic |
| chr11 | 6374257 | 6375693 | Distal Intergenic |
| chr11 | 6376148 | 6376587 | Distal Intergenic |
| chr11 | 6426049 | 6428756 | Promoter |
| chr11 | 6446054 | 6446724 | Distal Intergenic |
| chr11 | 6446929 | 6447145 | Distal Intergenic |
| chr11 | 6447274 | 6447658 | Distal Intergenic |
| chr11 | 6486788 | 6487131 | 5' UTR |
| chr11 | 6487340 | 6487776 | Intron |

| | | | |
|---|---|---|---|
| chr11 | 6525075 | 6525373 | Intron |
| chr11 | 6525856 | 6526263 | Intron |
| chr11 | 6594865 | 6598472 | Distal Intergenic |
| chr11 | 7111087 | 7111381 | Promoter |
| chr11 | 7111675 | 7112519 | 3' UTR |
| chr11 | 7138030 | 7138601 | Distal Intergenic |
| chr11 | 7304484 | 7305523 | Intron |
| chr11 | 7322780 | 7325423 | Exon |
| chr11 | 7351979 | 7352380 | Intron |
| chr11 | 7353937 | 7354156 | Intron |
| chr11 | 7356414 | 7356764 | Intron |
| chr11 | 7357503 | 7357708 | Intron |
| chr11 | 7426376 | 7426768 | Intron |
| chr11 | 7461458 | 7462771 | Exon |
| chr11 | 7502191 | 7504676 | Intron |
| chr11 | 7504716 | 7505210 | Intron |
| chr11 | 7515349 | 7515840 | Intron |
| chr11 | 7583301 | 7583571 | Intron |
| chr11 | 7583736 | 7583994 | Intron |
| chr11 | 7605181 | 7606102 | Intron |
| chr11 | 7679787 | 7681928 | Distal Intergenic |
| chr11 | 7693348 | 7696239 | Promoter |
| chr11 | 7797570 | 7798459 | Distal Intergenic |
| chr11 | 7798728 | 7799273 | Distal Intergenic |
| chr11 | 7799784 | 7801270 | Distal Intergenic |
| chr11 | 7895494 | 7896320 | Intron |
| chr11 | 8343967 | 8344754 | Distal Intergenic |
| chr11 | 8344985 | 8345200 | Distal Intergenic |
| chr11 | 8351886 | 8352489 | Distal Intergenic |
| chr11 | 8363478 | 8363823 | Distal Intergenic |
| chr11 | 8365777 | 8365982 | Distal Intergenic |
| chr11 | 8366116 | 8369212 | Distal Intergenic |
| chr11 | 8423395 | 8423651 | Intron |
| chr11 | 8423852 | 8424193 | Intron |
| chr11 | 8443940 | 8445452 | Intron |
| chr11 | 8469414 | 8469948 | Intron |
| chr11 | 8470015 | 8470334 | Intron |
| chr11 | 8470424 | 8470927 | Intron |
| chr11 | 8479629 | 8480058 | Intron |
| chr11 | 8480126 | 8480531 | Intron |
| chr11 | 8482750 | 8485173 | 5' UTR |
| chr11 | 8489065 | 8490338 | Intron |
| chr11 | 8542128 | 8542374 | Intron |
| chr11 | 8563958 | 8564638 | Intron |

| | | | |
|---|---|---|---|
| chr11 | 8567969 | 8568595 | Intron |
| chr11 | 8574359 | 8576112 | Intron |
| chr11 | 8578051 | 8578291 | Intron |
| chr11 | 8578348 | 8579073 | Intron |
| chr11 | 8584856 | 8586636 | Intron |
| chr11 | 8720852 | 8722254 | Exon |
| chr11 | 8730443 | 8730741 | Intron |
| chr11 | 8910921 | 8911407 | Intron |
| chr11 | 8939321 | 8939592 | 3' UTR |
| chr11 | 9456389 | 9457824 | Exon |
| chr11 | 9458013 | 9458235 | Intron |
| chr11 | 12270922 | 12271140 | Intron |
| chr11 | 12284693 | 12285074 | 3' UTR |
| chr11 | 12310842 | 12311324 | Intron |
| chr11 | 12311391 | 12311683 | Intron |
| chr11 | 12320633 | 12322536 | Intron |
| chr11 | 12335006 | 12335429 | Intron |
| chr11 | 12341891 | 12343246 | Intron |
| chr11 | 12370229 | 12371416 | Exon |
| chr11 | 12768116 | 12768862 | Intron |
| chr11 | 12770844 | 12771080 | Intron |
| chr11 | 12771884 | 12772375 | Intron |
| chr11 | 12772535 | 12773167 | Intron |
| chr11 | 16407319 | 16408104 | Intron |
| chr11 | 16491090 | 16492352 | Intron |
| chr11 | 16492389 | 16492625 | Intron |
| chr11 | 16583244 | 16584190 | Intron |
| chr11 | 16635004 | 16635696 | Intron |
| chr11 | 16635796 | 16636184 | Intron |
| chr11 | 16637319 | 16637546 | Intron |
| chr11 | 16652242 | 16653798 | Intron |
| chr11 | 16689704 | 16691752 | Intron |
| chr11 | 16695731 | 16695950 | Intron |
| chr11 | 16700497 | 16701210 | Intron |
| chr11 | 16704088 | 16704679 | Intron |
| chr11 | 16709670 | 16710372 | Intron |
| chr11 | 16710491 | 16710868 | Intron |
| chr11 | 16729059 | 16729258 | Intron |
| chr11 | 16729301 | 16731380 | Intron |
| chr11 | 16731594 | 16731797 | Intron |
| chr11 | 16732599 | 16733453 | Intron |
| chr11 | 16744027 | 16744960 | Intron |
| chr11 | 16749138 | 16749877 | Intron |
| chr11 | 16750605 | 16751893 | Intron |

| | | | |
|---|---|---|---|
| chr11 | 16757185 | 16757824 | Intron |
| chr11 | 16850768 | 16850980 | Intron |
| chr11 | 17059503 | 17059791 | Distal Intergenic |
| chr11 | 17059837 | 17060685 | Distal Intergenic |
| chr11 | 17072021 | 17073116 | Promoter |
| chr11 | 17076520 | 17077138 | Distal Intergenic |
| chr11 | 17111613 | 17111891 | Intron |
| chr11 | 17149394 | 17149958 | Intron |
| chr11 | 17157610 | 17158141 | Exon |
| chr11 | 17229620 | 17229876 | Promoter |
| chr11 | 17286510 | 17287392 | Intron |
| chr11 | 18798099 | 18798863 | Intron |
| chr11 | 18799013 | 18799437 | Intron |
| chr11 | 18804791 | 18806050 | Intron |
| chr11 | 18818752 | 18821150 | Distal Intergenic |
| chr11 | 18853148 | 18854392 | Distal Intergenic |
| chr11 | 18854600 | 18854846 | Distal Intergenic |
| chr11 | 18857024 | 18857331 | Distal Intergenic |
| chr11 | 18876124 | 18876573 | Distal Intergenic |
| chr11 | 18918549 | 18918748 | Distal Intergenic |
| chr11 | 18919064 | 18919869 | Distal Intergenic |
| chr11 | 18919908 | 18920451 | Distal Intergenic |
| chr11 | 19566483 | 19567420 | Intron |
| chr11 | 19637161 | 19638027 | Intron |
| chr11 | 19639969 | 19640327 | Intron |
| chr11 | 19640368 | 19641356 | Intron |
| chr11 | 19729323 | 19730763 | Intron |
| chr11 | 19734199 | 19734934 | Promoter |
| chr11 | 19735038 | 19737175 | Promoter |
| chr11 | 19738781 | 19739434 | Intron |
| chr11 | 19742718 | 19743970 | Intron |
| chr11 | 19747423 | 19750262 | Intron |
| chr11 | 19750383 | 19751146 | Intron |
| chr11 | 19761457 | 19761656 | Intron |
| chr11 | 19769127 | 19770935 | Intron |
| chr11 | 19773393 | 19774610 | Intron |
| chr11 | 19776883 | 19778126 | Intron |
| chr11 | 19819694 | 19820722 | Intron |
| chr11 | 19824310 | 19825003 | Intron |
| chr11 | 19825111 | 19825354 | Intron |
| chr11 | 19825504 | 19826013 | Intron |
| chr11 | 19832563 | 19833445 | Intron |
| chr11 | 19854891 | 19855329 | Intron |
| chr11 | 19855556 | 19855901 | Intron |

| | | | |
|---|---|---|---|
| chr11 | 19857993 | 19861602 | Intron |
| chr11 | 19869160 | 19870612 | Intron |
| chr11 | 19878707 | 19879942 | Intron |
| chr11 | 19892315 | 19892549 | Intron |
| chr11 | 19892766 | 19893489 | Intron |
| chr11 | 20007147 | 20009168 | Intron |
| chr11 | 20011747 | 20012525 | Intron |
| chr11 | 20012604 | 20013806 | Intron |
| chr11 | 20101247 | 20101548 | Intron |
| chr11 | 20101587 | 20102879 | Exon |
| chr11 | 20136900 | 20138214 | Intron |
| chr11 | 20232908 | 20234383 | Distal Intergenic |
| chr11 | 20247349 | 20249007 | Distal Intergenic |
| chr11 | 20257438 | 20258370 | Distal Intergenic |
| chr11 | 20269831 | 20271613 | Distal Intergenic |
| chr11 | 20272446 | 20272734 | Distal Intergenic |
| chr11 | 20272837 | 20273226 | Distal Intergenic |
| chr11 | 20377329 | 20378136 | Distal Intergenic |
| chr11 | 20442067 | 20442436 | Intron |
| chr11 | 20443260 | 20443460 | Intron |
| chr11 | 20443537 | 20443878 | Intron |
| chr11 | 20470774 | 20472582 | Intron |
| chr11 | 20486359 | 20486953 | Intron |
| chr11 | 20494242 | 20495990 | Intron |
| chr11 | 20530812 | 20531463 | 3' UTR |
| chr11 | 20531506 | 20531805 | Distal Intergenic |
| chr11 | 20532116 | 20533398 | Distal Intergenic |
| chr11 | 20547534 | 20547938 | Distal Intergenic |
| chr11 | 20640626 | 20640925 | Intron |
| chr11 | 20641415 | 20641634 | Intron |
| chr11 | 20641689 | 20642609 | Intron |
| chr11 | 20786360 | 20787236 | Intron |
| chr11 | 20796396 | 20796710 | Intron |
| chr11 | 20982874 | 20983351 | Intron |
| chr11 | 20991268 | 20991563 | Intron |
| chr11 | 20991805 | 20992006 | Intron |
| chr11 | 21138289 | 21139180 | Intron |
| chr11 | 21139254 | 21140526 | Intron |
| chr11 | 21141361 | 21141634 | Intron |
| chr11 | 21270038 | 21270468 | Intron |
| chr11 | 21270716 | 21270973 | Intron |
| chr11 | 21298262 | 21300366 | Intron |
| chr11 | 21481129 | 21482227 | Intron |
| chr11 | 21482715 | 21484439 | Intron |

| | | | |
|---|---|---|---|
| chr11 | 21588238 | 21589030 | Intron |
| chr11 | 21851077 | 21852603 | Distal Intergenic |
| chr11 | 21901320 | 21901714 | Distal Intergenic |
| chr11 | 21901818 | 21902378 | Distal Intergenic |
| chr11 | 21902431 | 21904123 | Distal Intergenic |
| chr11 | 21904226 | 21905923 | Distal Intergenic |
| chr11 | 21908291 | 21908951 | Distal Intergenic |
| chr11 | 22147284 | 22147911 | Distal Intergenic |
| chr11 | 22360095 | 22361730 | Promoter |
| chr11 | 22424393 | 22425055 | Distal Intergenic |
| chr11 | 22430270 | 22430568 | Distal Intergenic |
| chr11 | 22511673 | 22511946 | Distal Intergenic |
| chr11 | 22521706 | 22523024 | Distal Intergenic |
| chr11 | 22528657 | 22530370 | Distal Intergenic |
| chr11 | 22619701 | 22619905 | Distal Intergenic |
| chr11 | 22633048 | 22633699 | Distal Intergenic |
| chr11 | 22711562 | 22711831 | Intron |
| chr11 | 22743128 | 22744809 | Intron |
| chr11 | 22751042 | 22751320 | Intron |
| chr11 | 22751362 | 22752160 | Intron |
| chr11 | 22754943 | 22755149 | Intron |
| chr11 | 22756306 | 22757256 | Intron |
| chr11 | 22757311 | 22758782 | Intron |
| chr11 | 22759275 | 22759691 | Exon |
| chr11 | 22865142 | 22865851 | Downstream |
| chr11 | 22882056 | 22882284 | Promoter |
| chr11 | 22890280 | 22890541 | Distal Intergenic |
| chr11 | 22890728 | 22891593 | Distal Intergenic |
| chr11 | 22968574 | 22968805 | Distal Intergenic |
| chr11 | 23155031 | 23155504 | Distal Intergenic |
| chr11 | 23156161 | 23156568 | Distal Intergenic |
| chr11 | 23156746 | 23157055 | Distal Intergenic |
| chr11 | 23157119 | 23157791 | Distal Intergenic |
| chr11 | 23205500 | 23205801 | Distal Intergenic |
| chr11 | 23205902 | 23206270 | Distal Intergenic |
| chr11 | 23206830 | 23207865 | Distal Intergenic |
| chr11 | 23208046 | 23208975 | Distal Intergenic |
| chr11 | 23564623 | 23564899 | Distal Intergenic |
| chr11 | 23565036 | 23567009 | Distal Intergenic |
| chr11 | 23612270 | 23613187 | Distal Intergenic |
| chr11 | 23623287 | 23623658 | Distal Intergenic |
| chr11 | 23623708 | 23627499 | Distal Intergenic |
| chr11 | 23646223 | 23647443 | Distal Intergenic |
| chr11 | 23669174 | 23670972 | Distal Intergenic |

| | | | |
|---|---|---|---|
| chr11 | 24090050 | 24090422 | Distal Intergenic |
| chr11 | 24242998 | 24243352 | Distal Intergenic |
| chr11 | 24271997 | 24272356 | Distal Intergenic |
| chr11 | 24272402 | 24272713 | Distal Intergenic |
| chr11 | 24272988 | 24273236 | Distal Intergenic |
| chr11 | 26492813 | 26494395 | Intron |
| chr11 | 26499367 | 26500651 | Intron |
| chr11 | 27229982 | 27231631 | Distal Intergenic |
| chr11 | 27243733 | 27244658 | Distal Intergenic |
| chr11 | 27571660 | 27572269 | Intron |
| chr11 | 28712067 | 28712335 | Distal Intergenic |
| chr11 | 28712395 | 28713521 | Distal Intergenic |
| chr11 | 28733158 | 28734545 | Distal Intergenic |
| chr11 | 28780911 | 28782239 | Distal Intergenic |
| chr11 | 28790224 | 28790500 | Distal Intergenic |
| chr11 | 28806066 | 28806272 | Distal Intergenic |
| chr11 | 28806312 | 28806716 | Distal Intergenic |
| chr11 | 29030759 | 29030962 | Distal Intergenic |
| chr11 | 29072538 | 29073671 | Distal Intergenic |
| chr11 | 29073734 | 29073933 | Distal Intergenic |
| chr11 | 29414650 | 29414896 | Distal Intergenic |
| chr11 | 29446441 | 29447555 | Distal Intergenic |
| chr11 | 29447619 | 29448796 | Distal Intergenic |
| chr11 | 29579578 | 29579781 | Distal Intergenic |
| chr11 | 29579869 | 29580401 | Distal Intergenic |
| chr11 | 29592292 | 29592698 | Distal Intergenic |
| chr11 | 29592955 | 29593188 | Distal Intergenic |
| chr11 | 29593582 | 29593952 | Distal Intergenic |
| chr11 | 29594032 | 29594690 | Distal Intergenic |
| chr11 | 29739955 | 29740226 | Distal Intergenic |
| chr11 | 30017931 | 30019072 | Distal Intergenic |
| chr11 | 30058211 | 30059241 | Distal Intergenic |
| chr11 | 30082880 | 30083249 | Distal Intergenic |
| chr11 | 30083317 | 30083523 | Distal Intergenic |
| chr11 | 30096026 | 30097323 | Distal Intergenic |
| chr11 | 30097554 | 30099160 | Distal Intergenic |
| chr11 | 30109492 | 30109925 | Distal Intergenic |
| chr11 | 30111221 | 30111721 | Distal Intergenic |
| chr11 | 30510591 | 30512136 | Intron |
| chr11 | 30672246 | 30672593 | Distal Intergenic |
| chr11 | 30673097 | 30673410 | Distal Intergenic |
| chr11 | 30853461 | 30853882 | Intron |
| chr11 | 30854364 | 30854582 | Intron |
| chr11 | 30885890 | 30886944 | 3' UTR |

| | | | |
|---|---|---|---|
| chr11 | 31423952 | 31424886 | Intron |
| chr11 | 31570405 | 31570903 | Intron |
| chr11 | 31571296 | 31571517 | Intron |
| chr11 | 32118485 | 32119307 | Promoter |
| chr11 | 32120830 | 32122228 | Exon |
| chr11 | 32150903 | 32151543 | Distal Intergenic |
| chr11 | 32162891 | 32163269 | Distal Intergenic |
| chr11 | 32167097 | 32167947 | Distal Intergenic |
| chr11 | 32212483 | 32215431 | Distal Intergenic |
| chr11 | 32236358 | 32236947 | Distal Intergenic |
| chr11 | 32245446 | 32247100 | Distal Intergenic |
| chr11 | 32247409 | 32247809 | Distal Intergenic |
| chr11 | 32247912 | 32250152 | Distal Intergenic |
| chr11 | 32335371 | 32336153 | Distal Intergenic |
| chr11 | 32487153 | 32487797 | Distal Intergenic |
| chr11 | 32505105 | 32505898 | Distal Intergenic |
| chr11 | 32534957 | 32535406 | Distal Intergenic |
| chr11 | 32542512 | 32543349 | Distal Intergenic |
| chr11 | 32543390 | 32544170 | Distal Intergenic |
| chr11 | 32561625 | 32562377 | Distal Intergenic |
| chr11 | 32566834 | 32568702 | Distal Intergenic |
| chr11 | 32605010 | 32605209 | Promoter |
| chr11 | 32665904 | 32666412 | Intron |
| chr11 | 33048296 | 33048623 | Intron |
| chr11 | 33061431 | 33061940 | Promoter |
| chr11 | 33062070 | 33062792 | Promoter |
| chr11 | 34715470 | 34715680 | Distal Intergenic |
| chr11 | 34716020 | 34716258 | Distal Intergenic |
| chr11 | 34879835 | 34880144 | Distal Intergenic |
| chr11 | 34969069 | 34971207 | Exon |
| chr11 | 35075467 | 35075715 | Distal Intergenic |
| chr11 | 35076060 | 35076343 | Distal Intergenic |
| chr11 | 35076490 | 35076818 | Distal Intergenic |
| chr11 | 35130481 | 35132296 | Distal Intergenic |
| chr11 | 35144409 | 35145916 | Distal Intergenic |
| chr11 | 35228856 | 35230117 | 5' UTR |
| chr11 | 35286845 | 35288097 | 3' UTR |
| chr11 | 35481486 | 35481949 | Intron |
| chr11 | 35520486 | 35521156 | Intron |
| chr11 | 35582275 | 35583213 | Distal Intergenic |
| chr11 | 35583293 | 35584355 | Distal Intergenic |
| chr11 | 35584427 | 35584698 | Distal Intergenic |
| chr11 | 35647251 | 35647750 | Distal Intergenic |
| chr11 | 35651017 | 35652094 | Distal Intergenic |

| | | | |
|---|---|---|---|
| chr11 | 35652147 | 35652996 | Distal Intergenic |
| chr11 | 35658033 | 35658250 | Distal Intergenic |
| chr11 | 35691549 | 35691947 | Intron |
| chr11 | 35692164 | 35692363 | Intron |
| chr11 | 35739006 | 35739709 | Intron |
| chr11 | 35754248 | 35755201 | Intron |
| chr11 | 35756377 | 35756623 | Intron |
| chr11 | 35764444 | 35764696 | Intron |
| chr11 | 35768639 | 35770094 | Intron |
| chr11 | 35783962 | 35785681 | Intron |
| chr11 | 35792661 | 35793534 | Intron |
| chr11 | 35796079 | 35796682 | Intron |
| chr11 | 37909481 | 37909685 | Distal Intergenic |
| chr11 | 37931447 | 37931830 | Distal Intergenic |
| chr11 | 40505617 | 40507376 | Intron |
| chr11 | 40507539 | 40508185 | Intron |
| chr11 | 40546492 | 40546914 | Intron |
| chr11 | 40547453 | 40548263 | Intron |
| chr11 | 40554790 | 40555791 | Intron |
| chr11 | 40567402 | 40569569 | Intron |
| chr11 | 40603331 | 40603543 | Intron |
| chr11 | 40603776 | 40604197 | Intron |
| chr11 | 42199786 | 42200797 | Distal Intergenic |
| chr11 | 42752907 | 42753192 | Distal Intergenic |
| chr11 | 42836375 | 42836980 | Distal Intergenic |
| chr11 | 42845717 | 42845916 | Distal Intergenic |
| chr11 | 43239585 | 43240479 | Distal Intergenic |
| chr11 | 43253187 | 43253449 | Distal Intergenic |
| chr11 | 43706035 | 43707205 | Intron |
| chr11 | 43732922 | 43733121 | Intron |
| chr11 | 43733184 | 43734142 | Intron |
| chr11 | 43740292 | 43740753 | Intron |
| chr11 | 43746887 | 43747092 | Intron |
| chr11 | 43952421 | 43953599 | Distal Intergenic |
| chr11 | 44136286 | 44137452 | Intron |
| chr11 | 44333037 | 44335166 | Distal Intergenic |
| chr11 | 44347990 | 44349396 | Distal Intergenic |
| chr11 | 44988499 | 44988714 | Distal Intergenic |
| chr11 | 45618675 | 45619340 | Distal Intergenic |
| chr11 | 45665939 | 45666449 | Downstream |
| chr11 | 45689985 | 45691365 | Distal Intergenic |
| chr11 | 45973672 | 45973871 | Intron |
| chr11 | 45973973 | 45975092 | Exon |
| chr11 | 45978222 | 45978612 | Intron |

| | | | |
|---|---|---|---|
| chr11 | 45979213 | 45979503 | Intron |
| chr11 | 45979613 | 45980090 | Intron |
| chr11 | 45990375 | 45991326 | Intron |
| chr11 | 46038399 | 46039700 | Intron |
| chr11 | 46134623 | 46135161 | Intron |
| chr11 | 46250499 | 46250702 | Distal Intergenic |
| chr11 | 46250893 | 46251094 | Distal Intergenic |
| chr11 | 46275322 | 46275604 | Distal Intergenic |
| chr11 | 47910553 | 47912238 | Distal Intergenic |
| chr11 | 48612944 | 48615400 | Distal Intergenic |
| chr11 | 48631172 | 48631745 | Distal Intergenic |
| chr11 | 48632823 | 48634617 | Distal Intergenic |
| chr11 | 48651344 | 48653290 | Distal Intergenic |
| chr11 | 48668670 | 48668889 | Distal Intergenic |
| chr11 | 48787596 | 48788891 | Distal Intergenic |
| chr11 | 48859596 | 48860279 | Distal Intergenic |
| chr11 | 48860334 | 48860596 | Distal Intergenic |
| chr11 | 49016237 | 49018174 | Distal Intergenic |
| chr11 | 49059590 | 49062968 | Distal Intergenic |
| chr11 | 49063092 | 49064450 | Distal Intergenic |
| chr11 | 49280772 | 49281315 | Distal Intergenic |
| chr11 | 49345179 | 49345531 | Distal Intergenic |
| chr11 | 49346243 | 49346745 | Distal Intergenic |
| chr11 | 49526082 | 49526313 | Distal Intergenic |
| chr11 | 49526491 | 49526874 | Distal Intergenic |
| chr11 | 49580153 | 49580715 | Promoter |
| chr11 | 50023275 | 50027395 | Distal Intergenic |
| chr11 | 50038074 | 50038772 | Distal Intergenic |
| chr11 | 50065805 | 50066227 | Distal Intergenic |
| chr11 | 50088593 | 50088798 | Distal Intergenic |
| chr11 | 50088931 | 50089661 | Distal Intergenic |
| chr11 | 50104722 | 50105418 | Distal Intergenic |
| chr11 | 50105455 | 50105808 | Distal Intergenic |
| chr11 | 50105999 | 50106820 | Distal Intergenic |
| chr11 | 50126316 | 50126634 | Distal Intergenic |
| chr11 | 50126724 | 50126968 | Distal Intergenic |
| chr11 | 50127565 | 50127868 | Distal Intergenic |
| chr11 | 50138029 | 50139221 | Distal Intergenic |
| chr11 | 50139359 | 50139806 | Distal Intergenic |
| chr11 | 50139916 | 50140169 | Distal Intergenic |
| chr11 | 50191206 | 50191883 | Distal Intergenic |
| chr11 | 50254978 | 50255735 | Intron |
| chr11 | 50487595 | 50488008 | Distal Intergenic |
| chr11 | 51076656 | 51076873 | Distal Intergenic |

| | | | |
|---|---|---|---|
| chr11 | 51076966 | 51077326 | Distal Intergenic |
| chr11 | 51077521 | 51078444 | Distal Intergenic |
| chr11 | 51242973 | 51243200 | Distal Intergenic |
| chr11 | 51243241 | 51243559 | Distal Intergenic |
| chr11 | 51243616 | 51244392 | Distal Intergenic |
| chr11 | 51244432 | 51245411 | Distal Intergenic |
| chr11 | 51397288 | 51398001 | Distal Intergenic |
| chr11 | 51398224 | 51398693 | Distal Intergenic |
| chr11 | 51419307 | 51419553 | Distal Intergenic |
| chr11 | 51419808 | 51420039 | Distal Intergenic |
| chr11 | 51420440 | 51420712 | Distal Intergenic |
| chr11 | 51428665 | 51429600 | Distal Intergenic |
| chr11 | 51433436 | 51433746 | Distal Intergenic |
| chr11 | 51433808 | 51434041 | Distal Intergenic |
| chr11 | 51438491 | 51438754 | Distal Intergenic |
| chr11 | 51501770 | 51502019 | Distal Intergenic |
| chr11 | 51502378 | 51502795 | Distal Intergenic |
| chr11 | 51502857 | 51503065 | Distal Intergenic |
| chr11 | 51614286 | 51614559 | Distal Intergenic |
| chr11 | 51614821 | 51615241 | Distal Intergenic |
| chr11 | 51668650 | 51668912 | Distal Intergenic |
| chr11 | 51669318 | 51669530 | Distal Intergenic |
| chr11 | 51679458 | 51680054 | Distal Intergenic |
| chr11 | 51687556 | 51688010 | Distal Intergenic |
| chr11 | 51688046 | 51688876 | Distal Intergenic |
| chr11 | 51728084 | 51728355 | Distal Intergenic |
| chr11 | 51729158 | 51730549 | Distal Intergenic |
| chr11 | 51735733 | 51736713 | Distal Intergenic |
| chr11 | 51781610 | 51781867 | Distal Intergenic |
| chr11 | 51813446 | 51813851 | Distal Intergenic |
| chr11 | 51816912 | 51817625 | Distal Intergenic |
| chr11 | 51817778 | 51818510 | Distal Intergenic |
| chr11 | 51848436 | 51849728 | Distal Intergenic |
| chr11 | 51859922 | 51860443 | Distal Intergenic |
| chr11 | 51885054 | 51885416 | Distal Intergenic |
| chr11 | 51891172 | 51892164 | Distal Intergenic |
| chr11 | 51893175 | 51894553 | Distal Intergenic |
| chr11 | 51912402 | 51913565 | Distal Intergenic |
| chr11 | 51937860 | 51939251 | Distal Intergenic |
| chr11 | 51962125 | 51962556 | Distal Intergenic |
| chr11 | 52045592 | 52045827 | Distal Intergenic |
| chr11 | 52045899 | 52046102 | Distal Intergenic |
| chr11 | 52146459 | 52147822 | Distal Intergenic |
| chr11 | 52149368 | 52149650 | Distal Intergenic |

| | | | |
|---|---|---|---|
| chr11 | 52150904 | 52151997 | Distal Intergenic |
| chr11 | 52174396 | 52175372 | Distal Intergenic |
| chr11 | 52209815 | 52210120 | Distal Intergenic |
| chr11 | 52210283 | 52210546 | Distal Intergenic |
| chr11 | 52261701 | 52263153 | Distal Intergenic |
| chr11 | 53112622 | 53113844 | Distal Intergenic |
| chr11 | 53114022 | 53114658 | Distal Intergenic |
| chr11 | 53144354 | 53145402 | Distal Intergenic |
| chr11 | 53145578 | 53145803 | Distal Intergenic |
| chr11 | 53164525 | 53165796 | Distal Intergenic |
| chr11 | 53243811 | 53244348 | Distal Intergenic |
| chr11 | 53273054 | 53274484 | Distal Intergenic |
| chr11 | 53296832 | 53297039 | Distal Intergenic |
| chr11 | 53297136 | 53297460 | Distal Intergenic |
| chr11 | 53333646 | 53334318 | Distal Intergenic |
| chr11 | 53520227 | 53520746 | Distal Intergenic |
| chr11 | 53572244 | 53573492 | Distal Intergenic |
| chr11 | 53574930 | 53575174 | Distal Intergenic |
| chr11 | 53575210 | 53576546 | Distal Intergenic |
| chr11 | 53576584 | 53576800 | Distal Intergenic |
| chr11 | 53582157 | 53583154 | Distal Intergenic |
| chr11 | 53593147 | 53593412 | Distal Intergenic |
| chr11 | 53608366 | 53609159 | Distal Intergenic |
| chr11 | 53609326 | 53610957 | Distal Intergenic |
| chr11 | 53616244 | 53618293 | Distal Intergenic |
| chr11 | 53627077 | 53627279 | Distal Intergenic |
| chr11 | 53831465 | 53831851 | Distal Intergenic |
| chr11 | 53831964 | 53833276 | Distal Intergenic |
| chr11 | 54252248 | 54252485 | Distal Intergenic |
| chr11 | 54265977 | 54266188 | Distal Intergenic |
| chr11 | 54305353 | 54306560 | Distal Intergenic |
| chr11 | 54336400 | 54336961 | Distal Intergenic |
| chr11 | 54542686 | 54543426 | Distal Intergenic |
| chr11 | 54561995 | 54562788 | Distal Intergenic |
| chr11 | 54600179 | 54600503 | Distal Intergenic |
| chr11 | 54673182 | 54673722 | Distal Intergenic |
| chr11 | 54674126 | 54674694 | Distal Intergenic |
| chr11 | 54769192 | 54769391 | Distal Intergenic |
| chr11 | 54941650 | 54942245 | Distal Intergenic |
| chr11 | 55093530 | 55094415 | Distal Intergenic |
| chr11 | 55102977 | 55104344 | Distal Intergenic |
| chr11 | 55126877 | 55127093 | Distal Intergenic |
| chr11 | 55127250 | 55128748 | Distal Intergenic |
| chr11 | 55128965 | 55130403 | Distal Intergenic |

| | | | |
|---|---|---|---|
| chr11 | 55140339 | 55142624 | Distal Intergenic |
| chr11 | 55178543 | 55178785 | Distal Intergenic |
| chr11 | 55214566 | 55214779 | Distal Intergenic |
| chr11 | 55217978 | 55220550 | Distal Intergenic |
| chr11 | 55229476 | 55229850 | Distal Intergenic |
| chr11 | 55230270 | 55230592 | Distal Intergenic |
| chr11 | 55230757 | 55233870 | Distal Intergenic |
| chr11 | 55237501 | 55237774 | Distal Intergenic |
| chr11 | 55241309 | 55242550 | Distal Intergenic |
| chr11 | 55259910 | 55260234 | Distal Intergenic |
| chr11 | 55266625 | 55267968 | Distal Intergenic |
| chr11 | 55268763 | 55270157 | Distal Intergenic |
| chr11 | 55270213 | 55271166 | Distal Intergenic |
| chr11 | 55273949 | 55274222 | Distal Intergenic |
| chr11 | 55281845 | 55282708 | Distal Intergenic |
| chr11 | 55314438 | 55314840 | Distal Intergenic |
| chr11 | 55314930 | 55315259 | Distal Intergenic |
| chr11 | 55316034 | 55318565 | Distal Intergenic |
| chr11 | 55318622 | 55319624 | Distal Intergenic |
| chr11 | 55405637 | 55405843 | Promoter |
| chr11 | 55408293 | 55409712 | Distal Intergenic |
| chr11 | 55447079 | 55447823 | Distal Intergenic |
| chr11 | 55557628 | 55558711 | Distal Intergenic |
| chr11 | 55560337 | 55562042 | Promoter |
| chr11 | 55780165 | 55780548 | Distal Intergenic |
| chr11 | 57331430 | 57331878 | Intron |
| chr11 | 57532962 | 57533280 | Intron |
| chr11 | 57712790 | 57713046 | Distal Intergenic |
| chr11 | 57724954 | 57725234 | Distal Intergenic |
| chr11 | 57726752 | 57728520 | Distal Intergenic |
| chr11 | 57775673 | 57776252 | Distal Intergenic |
| chr11 | 57821852 | 57822056 | Intron |
| chr11 | 57822961 | 57824255 | 5' UTR |
| chr11 | 57828676 | 57830501 | Intron |
| chr11 | 57834275 | 57835571 | Intron |
| chr11 | 57837342 | 57838930 | Intron |
| chr11 | 57854451 | 57855565 | Intron |
| chr11 | 57856499 | 57856754 | Intron |
| chr11 | 57881501 | 57882989 | Intron |
| chr11 | 57917593 | 57917931 | Intron |
| chr11 | 58050210 | 58051374 | Distal Intergenic |
| chr11 | 58057087 | 58057915 | Distal Intergenic |
| chr11 | 58062810 | 58064251 | Distal Intergenic |
| chr11 | 58143860 | 58144347 | Distal Intergenic |

| | | | |
|---|---|---|---|
| chr11 | 58147646 | 58148206 | Distal Intergenic |
| chr11 | 58792189 | 58794543 | Intron |
| chr11 | 59031689 | 59033162 | Distal Intergenic |
| chr11 | 59039371 | 59041746 | Distal Intergenic |
| chr11 | 59091903 | 59092189 | Distal Intergenic |
| chr11 | 59092274 | 59092910 | Distal Intergenic |
| chr11 | 59093076 | 59093314 | Distal Intergenic |
| chr11 | 59095258 | 59097155 | Distal Intergenic |
| chr11 | 59097199 | 59098445 | Distal Intergenic |
| chr11 | 59137149 | 59138101 | Distal Intergenic |
| chr11 | 59198316 | 59198582 | Distal Intergenic |
| chr11 | 59215795 | 59217466 | Distal Intergenic |
| chr11 | 59222877 | 59223076 | Downstream |
| chr11 | 59223128 | 59228853 | Promoter |
| chr11 | 59233030 | 59235878 | Distal Intergenic |
| chr11 | 59245022 | 59245260 | Promoter |
| chr11 | 59263399 | 59264138 | Distal Intergenic |
| chr11 | 59269989 | 59270335 | Promoter |
| chr11 | 59270457 | 59270664 | Promoter |
| chr11 | 59271247 | 59274193 | Promoter |
| chr11 | 59285205 | 59285980 | Distal Intergenic |
| chr11 | 59319519 | 59319739 | Distal Intergenic |
| chr11 | 59413826 | 59415701 | Exon |
| chr11 | 59447423 | 59448355 | Distal Intergenic |
| chr11 | 59448471 | 59449915 | Distal Intergenic |
| chr11 | 59450207 | 59450882 | Distal Intergenic |
| chr11 | 59468157 | 59470001 | Distal Intergenic |
| chr11 | 59476580 | 59476945 | Distal Intergenic |
| chr11 | 59476997 | 59477236 | Distal Intergenic |
| chr11 | 59477286 | 59477549 | Downstream |
| chr11 | 59496034 | 59496787 | Distal Intergenic |
| chr11 | 59531200 | 59531426 | Intron |
| chr11 | 59544407 | 59544867 | Intron |
| chr11 | 59545024 | 59546696 | Intron |
| chr11 | 59556566 | 59557793 | Intron |
| chr11 | 59623344 | 59624121 | Exon |
| chr11 | 59903894 | 59904254 | Distal Intergenic |
| chr11 | 59919078 | 59920468 | Distal Intergenic |
| chr11 | 59921853 | 59922655 | Distal Intergenic |
| chr11 | 59926978 | 59927368 | Distal Intergenic |
| chr11 | 59927420 | 59927704 | Distal Intergenic |
| chr11 | 59927820 | 59928021 | Distal Intergenic |
| chr11 | 59928081 | 59928823 | Distal Intergenic |
| chr11 | 59928866 | 59929287 | Distal Intergenic |

| | | | |
|---|---|---|---|
| chr11 | 60033180 | 60034082 | Distal Intergenic |
| chr11 | 60034377 | 60034602 | Distal Intergenic |
| chr11 | 60170575 | 60170794 | Intron |
| chr11 | 60179941 | 60180140 | Intron |
| chr11 | 60350313 | 60350703 | Distal Intergenic |
| chr11 | 60350746 | 60350976 | Distal Intergenic |
| chr11 | 60351516 | 60351715 | Distal Intergenic |
| chr11 | 60521040 | 60521679 | Distal Intergenic |
| chr11 | 60540164 | 60540383 | Intron |
| chr11 | 60540610 | 60541332 | Exon |
| chr11 | 60541460 | 60541663 | Intron |
| chr11 | 60589999 | 60591006 | Distal Intergenic |
| chr11 | 60591104 | 60591379 | Distal Intergenic |
| chr11 | 60644202 | 60644409 | Distal Intergenic |
| chr11 | 60677195 | 60678277 | Distal Intergenic |
| chr11 | 60691714 | 60691929 | Promoter |
| chr11 | 60691986 | 60692325 | Promoter |
| chr11 | 60692433 | 60692801 | Promoter |
| chr11 | 60744423 | 60745396 | Intron |
| chr11 | 60745699 | 60745968 | Intron |
| chr11 | 60746018 | 60746312 | Intron |
| chr11 | 60747069 | 60747268 | Intron |
| chr11 | 60748768 | 60749035 | Intron |
| chr11 | 60749073 | 60749433 | Intron |
| chr11 | 60754442 | 60755106 | Intron |
| chr11 | 61013299 | 61013787 | Exon |
| chr11 | 61080750 | 61081043 | Exon |
| chr11 | 61087059 | 61088087 | Intron |
| chr11 | 61092205 | 61092598 | Intron |
| chr11 | 61451030 | 61452045 | Intron |
| chr11 | 61455253 | 61455923 | Intron |
| chr11 | 61457008 | 61457287 | Intron |
| chr11 | 61457572 | 61458761 | Intron |
| chr11 | 61460670 | 61462129 | Intron |
| chr11 | 61488789 | 61490737 | Exon |
| chr11 | 61497646 | 61497845 | Intron |
| chr11 | 61497894 | 61502206 | Exon |
| chr11 | 61502263 | 61502860 | Exon |
| chr11 | 61503009 | 61503349 | Exon |
| chr11 | 61504095 | 61504338 | Intron |
| chr11 | 61506059 | 61506819 | Intron |
| chr11 | 61506876 | 61508267 | Exon |
| chr11 | 61510436 | 61510792 | Intron |
| chr11 | 61512088 | 61512502 | 3' UTR |

| | | | |
|---|---|---|---|
| chr11 | 61512630 | 61513632 | 3' UTR |
| chr11 | 61519518 | 61520246 | Promoter |
| chr11 | 61521699 | 61522619 | Promoter |
| chr11 | 61836175 | 61836429 | Distal Intergenic |
| chr11 | 61836465 | 61837336 | Distal Intergenic |
| chr11 | 61888583 | 61889334 | Distal Intergenic |
| chr11 | 61890501 | 61890764 | Promoter |
| chr11 | 61891246 | 61891834 | Promoter |
| chr11 | 61903295 | 61903636 | Intron |
| chr11 | 61954299 | 61954820 | Distal Intergenic |
| chr11 | 61957530 | 61957945 | Promoter |
| chr11 | 61958094 | 61958350 | Promoter |
| chr11 | 61958718 | 61959159 | Intron |
| chr11 | 61959215 | 61959422 | Intron |
| chr11 | 61961294 | 61961553 | Distal Intergenic |
| chr11 | 61998433 | 61999138 | Distal Intergenic |
| chr11 | 62057559 | 62057802 | Distal Intergenic |
| chr11 | 62063117 | 62063483 | Downstream |
| chr11 | 62063536 | 62065002 | 3' UTR |
| chr11 | 62081300 | 62081618 | Distal Intergenic |
| chr11 | 62081713 | 62083276 | Distal Intergenic |
| chr11 | 62122833 | 62123279 | Intron |
| chr11 | 62270186 | 62270419 | Intron |
| chr11 | 62270580 | 62270879 | Intron |
| chr11 | 62270920 | 62271823 | Intron |
| chr11 | 62362922 | 62364565 | Exon |
| chr11 | 62364727 | 62365677 | 5' UTR |
| chr11 | 62416308 | 62418344 | Exon |
| chr11 | 62418762 | 62419016 | Intron |
| chr11 | 62627438 | 62628340 | Intron |
| chr11 | 62633431 | 62635638 | Intron |
| chr11 | 62638626 | 62639811 | Exon |
| chr11 | 62912622 | 62913502 | Promoter |
| chr11 | 62928282 | 62928910 | Intron |
| chr11 | 62949644 | 62950237 | Intron |
| chr11 | 63575864 | 63576065 | Distal Intergenic |
| chr11 | 63576335 | 63577807 | Distal Intergenic |
| chr11 | 63660533 | 63660904 | Intron |
| chr11 | 63671849 | 63673513 | Exon |
| chr11 | 63676248 | 63676647 | Exon |
| chr11 | 63704985 | 63705502 | Promoter |
| chr11 | 63723113 | 63724318 | 3' UTR |
| chr11 | 63892994 | 63893443 | Intron |
| chr11 | 64049793 | 64050172 | Intron |

| | | | |
|---|---|---|---|
| chr11 | 64195673 | 64195872 | Distal Intergenic |
| chr11 | 64196031 | 64196695 | Distal Intergenic |
| chr11 | 64282233 | 64282472 | Distal Intergenic |
| chr11 | 64282575 | 64282877 | Distal Intergenic |
| chr11 | 64285513 | 64287240 | Distal Intergenic |
| chr11 | 64314069 | 64314329 | Distal Intergenic |
| chr11 | 64315773 | 64317176 | Distal Intergenic |
| chr11 | 64457277 | 64458189 | Promoter |
| chr11 | 64750380 | 64751001 | Distal Intergenic |
| chr11 | 64759956 | 64760322 | Intron |
| chr11 | 65601543 | 65601744 | Promoter |
| chr11 | 65619952 | 65620386 | Exon |
| chr11 | 65631506 | 65631717 | Promoter |
| chr11 | 66865692 | 66866100 | Distal Intergenic |
| chr11 | 67037802 | 67038203 | Intron |
| chr11 | 67376762 | 67377002 | Promoter |
| chr11 | 67903591 | 67904009 | Distal Intergenic |
| chr11 | 67905638 | 67906331 | Distal Intergenic |
| chr11 | 68665741 | 68666533 | Intron |
| chr11 | 68666735 | 68668033 | 5' UTR |
| chr11 | 68673169 | 68674818 | Exon |
| chr11 | 68685184 | 68687181 | Exon |
| chr11 | 68715097 | 68715756 | Distal Intergenic |
| chr11 | 68715988 | 68716422 | Distal Intergenic |
| chr11 | 68759821 | 68760618 | Distal Intergenic |
| chr11 | 68780587 | 68780867 | Promoter |
| chr11 | 68780931 | 68781418 | Promoter |
| chr11 | 68828948 | 68830700 | Exon |
| chr11 | 68852940 | 68853743 | Exon |
| chr11 | 68858739 | 68859006 | Distal Intergenic |
| chr11 | 68859134 | 68859781 | Distal Intergenic |
| chr11 | 68873245 | 68873510 | Distal Intergenic |
| chr11 | 68886689 | 68887107 | Distal Intergenic |
| chr11 | 68887143 | 68887587 | Distal Intergenic |
| chr11 | 68908918 | 68909393 | Distal Intergenic |
| chr11 | 68910234 | 68910493 | Distal Intergenic |
| chr11 | 68910596 | 68910828 | Distal Intergenic |
| chr11 | 68910882 | 68911130 | Distal Intergenic |
| chr11 | 68913118 | 68913488 | Distal Intergenic |
| chr11 | 68938824 | 68939294 | Distal Intergenic |
| chr11 | 69178990 | 69181180 | Distal Intergenic |
| chr11 | 69196143 | 69196633 | Distal Intergenic |
| chr11 | 69208731 | 69209212 | Distal Intergenic |
| chr11 | 69222136 | 69224960 | Distal Intergenic |

| | | | |
|---|---|---|---|
| chr11 | 69225014 | 69225567 | Distal Intergenic |
| chr11 | 69227200 | 69228224 | Distal Intergenic |
| chr11 | 69228389 | 69229580 | Exon |
| chr11 | 69229942 | 69230169 | Exon |
| chr11 | 69231260 | 69231638 | Exon |
| chr11 | 69231738 | 69232102 | Distal Intergenic |
| chr11 | 69392459 | 69392907 | Distal Intergenic |
| chr11 | 69393029 | 69394621 | Distal Intergenic |
| chr11 | 69394753 | 69395253 | Distal Intergenic |
| chr11 | 69468143 | 69468500 | 3' UTR |
| chr11 | 69468574 | 69469007 | 3' UTR |
| chr11 | 69469105 | 69470897 | 3' UTR |
| chr11 | 69475899 | 69476105 | Intron |
| chr11 | 69479458 | 69479708 | Intron |
| chr11 | 69479839 | 69480042 | Intron |
| chr11 | 69484956 | 69485235 | Intron |
| chr11 | 69485289 | 69485527 | Intron |
| chr11 | 69485618 | 69486494 | 3' UTR |
| chr11 | 69495036 | 69495957 | Distal Intergenic |
| chr11 | 69571871 | 69572249 | Distal Intergenic |
| chr11 | 69572286 | 69572529 | Distal Intergenic |
| chr11 | 69572615 | 69573095 | Distal Intergenic |
| chr11 | 69573139 | 69573421 | Distal Intergenic |
| chr11 | 69618021 | 69618397 | Distal Intergenic |
| chr11 | 69659207 | 69661018 | Distal Intergenic |
| chr11 | 69691483 | 69695694 | Distal Intergenic |
| chr11 | 69696507 | 69697925 | Distal Intergenic |
| chr11 | 69727234 | 69727471 | Distal Intergenic |
| chr11 | 69732896 | 69734461 | Distal Intergenic |
| chr11 | 69734612 | 69734830 | Distal Intergenic |
| chr11 | 69735326 | 69735921 | Distal Intergenic |
| chr11 | 69774479 | 69775751 | Distal Intergenic |
| chr11 | 69804118 | 69804486 | Distal Intergenic |
| chr11 | 69804738 | 69805711 | Distal Intergenic |
| chr11 | 69809403 | 69810251 | Distal Intergenic |
| chr11 | 69810306 | 69810680 | Distal Intergenic |
| chr11 | 69810885 | 69811114 | Distal Intergenic |
| chr11 | 70002474 | 70003801 | Exon |
| chr11 | 70005662 | 70006149 | Intron |
| chr11 | 70006193 | 70007969 | Exon |
| chr11 | 70008037 | 70008322 | Intron |
| chr11 | 70010947 | 70012487 | Exon |
| chr11 | 70023774 | 70025054 | Intron |
| chr11 | 70033341 | 70034023 | Exon |

| | | | |
|---|---|---|---|
| chr11 | 70034092 | 70034617 | 3' UTR |
| chr11 | 70051714 | 70052097 | Intron |
| chr11 | 70052429 | 70052654 | 3' UTR |
| chr11 | 70072808 | 70073185 | Distal Intergenic |
| chr11 | 70101598 | 70102214 | Distal Intergenic |
| chr11 | 70104713 | 70105022 | Distal Intergenic |
| chr11 | 70106872 | 70107236 | Distal Intergenic |
| chr11 | 70265452 | 70265684 | Promoter |
| chr11 | 70265740 | 70266856 | Promoter |
| chr11 | 70271786 | 70273870 | Intron |
| chr11 | 70280875 | 70281688 | 3' UTR |
| chr11 | 70281957 | 70282199 | 3' UTR |
| chr11 | 70338844 | 70339282 | Intron |
| chr11 | 70375211 | 70376129 | Intron |
| chr11 | 70386225 | 70386607 | Intron |
| chr11 | 70387073 | 70387311 | Intron |
| chr11 | 70387493 | 70387775 | Intron |
| chr11 | 70389975 | 70394200 | Intron |
| chr11 | 70396564 | 70398298 | Intron |
| chr11 | 70398595 | 70400915 | Intron |
| chr11 | 70460785 | 70461232 | Intron |
| chr11 | 70464814 | 70465452 | Intron |
| chr11 | 70465627 | 70466185 | Intron |
| chr11 | 70467188 | 70468150 | Intron |
| chr11 | 70468776 | 70469243 | Intron |
| chr11 | 70481743 | 70482234 | Intron |
| chr11 | 70482503 | 70482790 | Intron |
| chr11 | 70514342 | 70514632 | Intron |
| chr11 | 70515110 | 70515316 | Intron |
| chr11 | 70528867 | 70529148 | Intron |
| chr11 | 70529290 | 70529542 | Intron |
| chr11 | 70648913 | 70649399 | Intron |
| chr11 | 70833150 | 70833572 | Intron |
| chr11 | 71855459 | 71855730 | Distal Intergenic |
| chr11 | 71895727 | 71897959 | Distal Intergenic |
| chr11 | 71898088 | 71898287 | Distal Intergenic |
| chr11 | 71903789 | 71904450 | Promoter |
| chr11 | 71923684 | 71925057 | Distal Intergenic |
| chr11 | 71925257 | 71925625 | Distal Intergenic |
| chr11 | 71925693 | 71926732 | Distal Intergenic |
| chr11 | 71945543 | 71946523 | Exon |
| chr11 | 71956481 | 71958201 | Distal Intergenic |
| chr11 | 71966646 | 71967541 | Distal Intergenic |
| chr11 | 71976735 | 71977171 | Distal Intergenic |

| | | | |
|---|---|---|---|
| chr11 | 71977458 | 71977805 | Distal Intergenic |
| chr11 | 72009730 | 72011051 | Intron |
| chr11 | 72020085 | 72023317 | Promoter |
| chr11 | 72248754 | 72249222 | Distal Intergenic |
| chr11 | 72254624 | 72255393 | Distal Intergenic |
| chr11 | 72309642 | 72309850 | Intron |
| chr11 | 72314670 | 72314869 | Intron |
| chr11 | 72314907 | 72315121 | Intron |
| chr11 | 72315241 | 72315707 | Intron |
| chr11 | 72420749 | 72420972 | Exon |
| chr11 | 72421692 | 72422166 | Exon |
| chr11 | 72503495 | 72504024 | Promoter |
| chr11 | 72509771 | 72510857 | Distal Intergenic |
| chr11 | 72565290 | 72565633 | Intron |
| chr11 | 72604991 | 72607963 | Intron |
| chr11 | 72608114 | 72609571 | Intron |
| chr11 | 72655908 | 72656469 | Intron |
| chr11 | 72656630 | 72656929 | Intron |
| chr11 | 72657019 | 72657299 | Intron |
| chr11 | 72659996 | 72662384 | Intron |
| chr11 | 72835232 | 72835431 | Intron |
| chr11 | 72861367 | 72861851 | Distal Intergenic |
| chr11 | 72989737 | 72990154 | Intron |
| chr11 | 72990289 | 72990543 | Intron |
| chr11 | 72990713 | 72992528 | Intron |
| chr11 | 73079659 | 73080156 | 3' UTR |
| chr11 | 73080279 | 73080917 | 3' UTR |
| chr11 | 73139953 | 73140370 | Intron |
| chr11 | 74295039 | 74295340 | Distal Intergenic |
| chr11 | 74296845 | 74297513 | Distal Intergenic |
| chr11 | 74369375 | 74369775 | Distal Intergenic |
| chr11 | 74370274 | 74370478 | Distal Intergenic |
| chr11 | 74370634 | 74371531 | Distal Intergenic |
| chr11 | 74455719 | 74455970 | Distal Intergenic |
| chr11 | 74456069 | 74456455 | Distal Intergenic |
| chr11 | 74484425 | 74484827 | Intron |
| chr11 | 74485012 | 74485243 | Intron |
| chr11 | 74486245 | 74487616 | Intron |
| chr11 | 74517365 | 74517965 | Intron |
| chr11 | 74536730 | 74537531 | Intron |
| chr11 | 74537652 | 74538299 | Intron |
| chr11 | 74538348 | 74538569 | Intron |
| chr11 | 74576764 | 74577916 | Intron |
| chr11 | 74643409 | 74643695 | Intron |

| | | | |
|---|---|---|---|
| chr11 | 74645475 | 74645695 | Intron |
| chr11 | 74645971 | 74646283 | Intron |
| chr11 | 74684743 | 74685216 | Intron |
| chr11 | 74687549 | 74688325 | 3' UTR |
| chr11 | 74711618 | 74711974 | Intron |
| chr11 | 74724291 | 74724542 | Distal Intergenic |
| chr11 | 74859134 | 74859488 | Intron |
| chr11 | 74866421 | 74867069 | Intron |
| chr11 | 74870061 | 74871022 | Promoter |
| chr11 | 74873987 | 74874553 | Intron |
| chr11 | 75201376 | 75201849 | Promoter |
| chr11 | 75202909 | 75203129 | Intron |
| chr11 | 75215906 | 75216105 | Intron |
| chr11 | 75216226 | 75217897 | Exon |
| chr11 | 75273163 | 75274984 | Promoter |
| chr11 | 75275123 | 75275384 | Intron |
| chr11 | 75275508 | 75277051 | Intron |
| chr11 | 75300364 | 75300891 | Intron |
| chr11 | 75364885 | 75365785 | Intron |
| chr11 | 75377269 | 75378410 | Intron |
| chr11 | 75406019 | 75406311 | Distal Intergenic |
| chr11 | 75433946 | 75434266 | Intron |
| chr11 | 75434619 | 75435092 | Intron |
| chr11 | 75444041 | 75444340 | Distal Intergenic |
| chr11 | 75465058 | 75465885 | Distal Intergenic |
| chr11 | 75466897 | 75467449 | Downstream |
| chr11 | 75468561 | 75469775 | Exon |
| chr11 | 75491292 | 75492211 | Intron |
| chr11 | 75492311 | 75493465 | Intron |
| chr11 | 75493503 | 75493900 | Intron |
| chr11 | 75546486 | 75547477 | Intron |
| chr11 | 75549403 | 75550754 | Promoter |
| chr11 | 75673898 | 75675620 | Intron |
| chr11 | 75793473 | 75795053 | Intron |
| chr11 | 75873327 | 75874172 | Distal Intergenic |
| chr11 | 75934223 | 75934457 | Distal Intergenic |
| chr11 | 75934560 | 75934769 | Distal Intergenic |
| chr11 | 75944052 | 75944698 | Distal Intergenic |
| chr11 | 75948707 | 75949886 | Distal Intergenic |
| chr11 | 76022101 | 76023824 | Distal Intergenic |
| chr11 | 76029962 | 76031042 | Distal Intergenic |
| chr11 | 76049198 | 76050093 | Distal Intergenic |
| chr11 | 76078702 | 76079021 | Intron |
| chr11 | 76079203 | 76079492 | Intron |

| | | | |
|---|---|---|---|
| chr11 | 76097166 | 76098282 | Intron |
| chr11 | 76098388 | 76100398 | Intron |
| chr11 | 76100453 | 76101090 | Intron |
| chr11 | 76113713 | 76115320 | Exon |
| chr11 | 76115594 | 76116594 | Intron |
| chr11 | 76125436 | 76127218 | Exon |
| chr11 | 76128432 | 76128689 | Distal Intergenic |
| chr11 | 76132804 | 76133417 | Distal Intergenic |
| chr11 | 76133463 | 76134041 | Distal Intergenic |
| chr11 | 76144597 | 76146945 | Distal Intergenic |
| chr11 | 76187733 | 76189922 | Intron |
| chr11 | 76211060 | 76211318 | Intron |
| chr11 | 76211378 | 76211974 | Intron |
| chr11 | 76220428 | 76220869 | Intron |
| chr11 | 76278579 | 76278792 | Distal Intergenic |
| chr11 | 76282872 | 76284173 | Distal Intergenic |
| chr11 | 76286005 | 76286623 | Distal Intergenic |
| chr11 | 76289777 | 76291657 | Distal Intergenic |
| chr11 | 76291871 | 76294561 | Distal Intergenic |
| chr11 | 76306871 | 76307112 | Distal Intergenic |
| chr11 | 76307237 | 76307742 | Distal Intergenic |
| chr11 | 76307789 | 76308786 | Distal Intergenic |
| chr11 | 76385212 | 76385411 | Distal Intergenic |
| chr11 | 76715027 | 76716000 | Intron |
| chr11 | 76716171 | 76716990 | Intron |
| chr11 | 76815965 | 76816362 | Intron |
| chr11 | 76817979 | 76819450 | Intron |
| chr11 | 76891200 | 76892594 | Promoter |
| chr11 | 76973785 | 76974464 | Intron |
| chr11 | 77041981 | 77043442 | Intron |
| chr11 | 77047977 | 77049057 | Exon |
| chr11 | 77278152 | 77279212 | Distal Intergenic |
| chr11 | 77279283 | 77280519 | Distal Intergenic |
| chr11 | 77307513 | 77307727 | Intron |
| chr11 | 77420146 | 77420789 | Intron |
| chr11 | 77421538 | 77421760 | Intron |
| chr11 | 77499300 | 77499703 | Intron |
| chr11 | 77499956 | 77501593 | Intron |
| chr11 | 77501661 | 77501997 | Intron |
| chr11 | 77577095 | 77577894 | Intron |
| chr11 | 77598862 | 77600120 | Intron |
| chr11 | 77605535 | 77605743 | Intron |
| chr11 | 77606949 | 77607330 | Intron |
| chr11 | 77649990 | 77651801 | Intron |

| | | | |
|---|---|---|---|
| chr11 | 77796971 | 77797605 | Distal Intergenic |
| chr11 | 77977870 | 77978365 | Intron |
| chr11 | 77979106 | 77980045 | Intron |
| chr11 | 77990027 | 77990816 | Intron |
| chr11 | 77991167 | 77992024 | 5' UTR |
| chr11 | 77992570 | 77993646 | Intron |
| chr11 | 77995323 | 77995522 | Intron |
| chr11 | 77995714 | 77996000 | Intron |
| chr11 | 77996179 | 77997396 | Intron |
| chr11 | 78059002 | 78060168 | Intron |
| chr11 | 78060255 | 78060702 | Intron |
| chr11 | 78196326 | 78196629 | Intron |
| chr11 | 78260659 | 78260943 | Intron |
| chr11 | 78279184 | 78279935 | 5' UTR |
| chr11 | 78325203 | 78325440 | Distal Intergenic |
| chr11 | 78325730 | 78325938 | Distal Intergenic |
| chr11 | 78348485 | 78349302 | Distal Intergenic |
| chr11 | 78804738 | 78804971 | Intron |
| chr11 | 78815874 | 78816253 | Intron |
| chr11 | 79027748 | 79028364 | Intron |
| chr11 | 79066275 | 79067649 | Intron |
| chr11 | 79067927 | 79068184 | Intron |
| chr11 | 79226944 | 79227550 | Distal Intergenic |
| chr11 | 79296998 | 79298872 | Distal Intergenic |
| chr11 | 79308689 | 79308907 | Distal Intergenic |
| chr11 | 79517599 | 79517808 | Distal Intergenic |
| chr11 | 79552815 | 79553059 | Distal Intergenic |
| chr11 | 79685232 | 79685698 | Distal Intergenic |
| chr11 | 79721776 | 79722910 | Distal Intergenic |
| chr11 | 79775522 | 79775877 | Distal Intergenic |
| chr11 | 79807105 | 79807502 | Distal Intergenic |
| chr11 | 79807726 | 79807925 | Distal Intergenic |
| chr11 | 80196470 | 80196695 | Distal Intergenic |
| chr11 | 80196755 | 80197839 | Distal Intergenic |
| chr11 | 80198217 | 80198682 | Distal Intergenic |
| chr11 | 80226753 | 80226985 | Distal Intergenic |
| chr11 | 80242129 | 80242806 | Distal Intergenic |
| chr11 | 80253424 | 80253658 | Distal Intergenic |
| chr11 | 80328918 | 80329235 | Distal Intergenic |
| chr11 | 80346214 | 80346467 | Distal Intergenic |
| chr11 | 80367238 | 80368204 | Distal Intergenic |
| chr11 | 80588760 | 80589000 | Distal Intergenic |
| chr11 | 80592726 | 80592973 | Distal Intergenic |
| chr11 | 80597803 | 80598269 | Distal Intergenic |

| | | | |
|---|---|---|---|
| chr11 | 81722082 | 81722385 | Intron |
| chr11 | 81722541 | 81722929 | Intron |
| chr11 | 81820507 | 81820840 | Intron |
| chr11 | 81825174 | 81825614 | Intron |
| chr11 | 81828429 | 81829486 | Intron |
| chr11 | 81843581 | 81844483 | Intron |
| chr11 | 81846197 | 81847604 | Intron |
| chr11 | 81887207 | 81887486 | Intron |
| chr11 | 81929487 | 81929776 | Intron |
| chr11 | 81952729 | 81953047 | Intron |
| chr11 | 81953084 | 81953557 | Intron |
| chr11 | 81982533 | 81983794 | Intron |
| chr11 | 81984908 | 81985816 | Intron |
| chr11 | 82577943 | 82578142 | Intron |
| chr11 | 82653379 | 82654820 | Intron |
| chr11 | 82659542 | 82660029 | Intron |
| chr11 | 82852946 | 82853153 | Distal Intergenic |
| chr11 | 82853311 | 82853518 | Distal Intergenic |
| chr11 | 82853564 | 82854044 | Distal Intergenic |
| chr11 | 82857210 | 82857921 | Distal Intergenic |
| chr11 | 83104915 | 83105137 | Intron |
| chr11 | 83112044 | 83112447 | Intron |
| chr11 | 83112529 | 83112761 | Intron |
| chr11 | 83115337 | 83115559 | Intron |
| chr11 | 83115690 | 83117316 | Intron |
| chr11 | 83268241 | 83269955 | Intron |
| chr11 | 83270349 | 83270548 | Intron |
| chr11 | 83270875 | 83272457 | Intron |
| chr11 | 83282020 | 83282908 | Intron |
| chr11 | 83285668 | 83286236 | Intron |
| chr11 | 83286924 | 83287234 | Intron |
| chr11 | 83377393 | 83378020 | Intron |
| chr11 | 83411055 | 83412900 | Intron |
| chr11 | 83419461 | 83420814 | Intron |
| chr11 | 83429916 | 83430501 | Intron |
| chr11 | 83705189 | 83705479 | Intron |
| chr11 | 83733899 | 83734268 | Intron |
| chr11 | 83751985 | 83752403 | Intron |
| chr11 | 83755687 | 83755905 | Intron |
| chr11 | 83856550 | 83856906 | Intron |
| chr11 | 83860844 | 83861064 | Intron |
| chr11 | 83876355 | 83877040 | Intron |
| chr11 | 83880988 | 83881507 | Intron |
| chr11 | 83882109 | 83883120 | Intron |

| | | | |
|---|---|---|---|
| chr11 | 83893786 | 83894170 | Intron |
| chr11 | 83914420 | 83916498 | Intron |
| chr11 | 83992567 | 83992949 | Intron |
| chr11 | 83993799 | 83994109 | Intron |
| chr11 | 83994241 | 83994502 | Intron |
| chr11 | 84627314 | 84628032 | Intron |
| chr11 | 84632077 | 84632320 | Intron |
| chr11 | 84682912 | 84683416 | Intron |
| chr11 | 84684220 | 84684579 | Intron |
| chr11 | 84684640 | 84685002 | Intron |
| chr11 | 84693132 | 84693400 | Intron |
| chr11 | 84693520 | 84695635 | Intron |
| chr11 | 84695770 | 84695974 | Intron |
| chr11 | 84729994 | 84730920 | Intron |
| chr11 | 84953779 | 84954253 | Intron |
| chr11 | 84970718 | 84971279 | Intron |
| chr11 | 85047838 | 85048088 | Intron |
| chr11 | 85087057 | 85089337 | Intron |
| chr11 | 85090652 | 85091514 | Intron |
| chr11 | 85125384 | 85125784 | Intron |
| chr11 | 85180751 | 85182223 | Intron |
| chr11 | 85192315 | 85192815 | Intron |
| chr11 | 85205251 | 85205500 | Intron |
| chr11 | 85205701 | 85206014 | Intron |
| chr11 | 85206159 | 85206358 | Intron |
| chr11 | 85603176 | 85603925 | Intron |
| chr11 | 85605919 | 85606452 | Exon |
| chr11 | 85625986 | 85626411 | Intron |
| chr11 | 85631654 | 85632699 | Distal Intergenic |
| chr11 | 85944482 | 85945551 | Distal Intergenic |
| chr11 | 86014970 | 86015265 | Promoter |
| chr11 | 86137586 | 86138323 | Distal Intergenic |
| chr11 | 86138369 | 86138568 | Distal Intergenic |
| chr11 | 86169782 | 86171012 | Intron |
| chr11 | 86171416 | 86171778 | Intron |
| chr11 | 86205425 | 86206326 | Intron |
| chr11 | 86206365 | 86206664 | Intron |
| chr11 | 86320170 | 86322096 | Intron |
| chr11 | 86358067 | 86358282 | Intron |
| chr11 | 86390725 | 86393094 | Distal Intergenic |
| chr11 | 86393130 | 86402330 | Distal Intergenic |
| chr11 | 86402636 | 86402854 | Distal Intergenic |
| chr11 | 86402965 | 86403297 | Distal Intergenic |
| chr11 | 86403450 | 86405705 | Distal Intergenic |

| | | | |
|---|---|---|---|
| chr11 | 86405760 | 86408369 | Distal Intergenic |
| chr11 | 86413155 | 86416039 | Distal Intergenic |
| chr11 | 86437120 | 86439282 | Distal Intergenic |
| chr11 | 86441558 | 86442047 | Distal Intergenic |
| chr11 | 86447789 | 86449680 | Distal Intergenic |
| chr11 | 86459334 | 86466777 | Distal Intergenic |
| chr11 | 86497538 | 86497766 | Distal Intergenic |
| chr11 | 86570505 | 86571019 | Intron |
| chr11 | 86571412 | 86572450 | Intron |
| chr11 | 86573241 | 86573489 | Intron |
| chr11 | 86573530 | 86574034 | Intron |
| chr11 | 86923107 | 86923458 | Intron |
| chr11 | 87123496 | 87124005 | Distal Intergenic |
| chr11 | 87235817 | 87236328 | Distal Intergenic |
| chr11 | 87236979 | 87237222 | Distal Intergenic |
| chr11 | 87262315 | 87262778 | Distal Intergenic |
| chr11 | 87275853 | 87276158 | Distal Intergenic |
| chr11 | 87284974 | 87285176 | Distal Intergenic |
| chr11 | 87404939 | 87405143 | Distal Intergenic |
| chr11 | 87405917 | 87407010 | Distal Intergenic |
| chr11 | 87492200 | 87492430 | Distal Intergenic |
| chr11 | 87539009 | 87539337 | Distal Intergenic |
| chr11 | 87539541 | 87539823 | Distal Intergenic |
| chr11 | 87860900 | 87861102 | Intron |
| chr11 | 87861207 | 87861839 | Intron |
| chr11 | 87917886 | 87918943 | Distal Intergenic |
| chr11 | 87919485 | 87919684 | Distal Intergenic |
| chr11 | 87932043 | 87932263 | Distal Intergenic |
| chr11 | 87932360 | 87932816 | Distal Intergenic |
| chr11 | 87936332 | 87937879 | Distal Intergenic |
| chr11 | 87964806 | 87965297 | Distal Intergenic |
| chr11 | 87967134 | 87967457 | Distal Intergenic |
| chr11 | 87967701 | 87968352 | Distal Intergenic |
| chr11 | 87968794 | 87969143 | Distal Intergenic |
| chr11 | 87974647 | 87974846 | Distal Intergenic |
| chr11 | 87978055 | 87979367 | Distal Intergenic |
| chr11 | 87985069 | 87986101 | Distal Intergenic |
| chr11 | 87986160 | 87986827 | Distal Intergenic |
| chr11 | 87987114 | 87987368 | Distal Intergenic |
| chr11 | 88281269 | 88281468 | Intron |
| chr11 | 88308994 | 88309965 | Intron |
| chr11 | 88383956 | 88384752 | Intron |
| chr11 | 88799194 | 88799631 | Promoter |
| chr11 | 88857519 | 88858469 | Distal Intergenic |

| | | | |
|---|---|---|---|
| chr11 | 88861458 | 88861809 | Distal Intergenic |
| chr11 | 88884919 | 88885278 | Distal Intergenic |
| chr11 | 88885589 | 88885925 | Distal Intergenic |
| chr11 | 89048396 | 89049644 | Distal Intergenic |
| chr11 | 89067575 | 89068432 | Intron |
| chr11 | 89100889 | 89101182 | Intron |
| chr11 | 89293080 | 89294032 | Intron |
| chr11 | 89294185 | 89294765 | Intron |
| chr11 | 89400953 | 89401485 | Intron |
| chr11 | 90201495 | 90202927 | Distal Intergenic |
| chr11 | 90211098 | 90211997 | Distal Intergenic |
| chr11 | 90212750 | 90215285 | Distal Intergenic |
| chr11 | 90215949 | 90217725 | Distal Intergenic |
| chr11 | 90502739 | 90502963 | Intron |
| chr11 | 90526199 | 90526959 | Intron |
| chr11 | 90540056 | 90540419 | Intron |
| chr11 | 92356589 | 92356799 | Intron |
| chr11 | 92867347 | 92867918 | Distal Intergenic |
| chr11 | 93529294 | 93530066 | Exon |
| chr11 | 93801693 | 93802683 | Intron |
| chr11 | 93816088 | 93816360 | Intron |
| chr11 | 93816465 | 93816787 | Intron |
| chr11 | 93816910 | 93817215 | Intron |
| chr11 | 93817405 | 93819001 | Intron |
| chr11 | 93828465 | 93829598 | Intron |
| chr11 | 93848253 | 93850804 | Intron |
| chr11 | 93855396 | 93857169 | Intron |
| chr11 | 94002958 | 94003607 | Distal Intergenic |
| chr11 | 94088789 | 94089027 | Distal Intergenic |
| chr11 | 94089085 | 94089298 | Distal Intergenic |
| chr11 | 94135849 | 94137555 | Distal Intergenic |
| chr11 | 94183077 | 94183279 | Intron |
| chr11 | 94231342 | 94232447 | Promoter |
| chr11 | 94277685 | 94277952 | Promoter |
| chr11 | 94316552 | 94316802 | Exon |
| chr11 | 94317098 | 94318508 | Intron |
| chr11 | 94318640 | 94318856 | Exon |
| chr11 | 94359074 | 94360920 | Distal Intergenic |
| chr11 | 94491106 | 94492161 | Distal Intergenic |
| chr11 | 94819129 | 94819763 | Distal Intergenic |
| chr11 | 94849069 | 94849400 | Intron |
| chr11 | 94849442 | 94852165 | Intron |
| chr11 | 94852207 | 94852493 | Intron |
| chr11 | 94852749 | 94853000 | Intron |

| | | | |
|---|---|---|---|
| chr11 | 94853916 | 94854124 | Intron |
| chr11 | 94930530 | 94935997 | Intron |
| chr11 | 94936060 | 94938011 | Intron |
| chr11 | 94938111 | 94938613 | Intron |
| chr11 | 94938803 | 94940277 | Intron |
| chr11 | 94946162 | 94948664 | Intron |
| chr11 | 94954267 | 94955683 | Intron |
| chr11 | 94957414 | 94960146 | Intron |
| chr11 | 94960227 | 94960435 | Intron |
| chr11 | 94970376 | 94970670 | Distal Intergenic |
| chr11 | 94998098 | 94998700 | Distal Intergenic |
| chr11 | 95014709 | 95015226 | Distal Intergenic |
| chr11 | 95134811 | 95136389 | Distal Intergenic |
| chr11 | 95161115 | 95161795 | Distal Intergenic |
| chr11 | 95171159 | 95171544 | Distal Intergenic |
| chr11 | 95171651 | 95171925 | Distal Intergenic |
| chr11 | 95245996 | 95246418 | Distal Intergenic |
| chr11 | 95254354 | 95254574 | Distal Intergenic |
| chr11 | 95254621 | 95255027 | Distal Intergenic |
| chr11 | 95263751 | 95265265 | Distal Intergenic |
| chr11 | 95276128 | 95276561 | Distal Intergenic |
| chr11 | 95277091 | 95277648 | Distal Intergenic |
| chr11 | 95294118 | 95294872 | Distal Intergenic |
| chr11 | 95411005 | 95411236 | Distal Intergenic |
| chr11 | 95411295 | 95412581 | Distal Intergenic |
| chr11 | 95437784 | 95437983 | Distal Intergenic |
| chr11 | 95638320 | 95639808 | Intron |
| chr11 | 95661554 | 95661896 | Distal Intergenic |
| chr11 | 95662049 | 95662424 | Distal Intergenic |
| chr11 | 95679913 | 95681335 | Distal Intergenic |
| chr11 | 95843212 | 95843475 | Intron |
| chr11 | 95843566 | 95844886 | Intron |
| chr11 | 95849564 | 95850430 | Intron |
| chr11 | 95865740 | 95866287 | Intron |
| chr11 | 95866771 | 95867242 | Intron |
| chr11 | 95879357 | 95879607 | Intron |
| chr11 | 95925864 | 95926190 | Intron |
| chr11 | 96112869 | 96113085 | Intron |
| chr11 | 96113497 | 96114528 | Intron |
| chr11 | 96120907 | 96121106 | Promoter |
| chr11 | 96121303 | 96121744 | Promoter |
| chr11 | 96320100 | 96320482 | Distal Intergenic |
| chr11 | 96608684 | 96609509 | Distal Intergenic |
| chr11 | 96610131 | 96611214 | Distal Intergenic |

| | | | |
|---|---|---|---|
| chr11 | 96651382 | 96651632 | Distal Intergenic |
| chr11 | 96683849 | 96685265 | Distal Intergenic |
| chr11 | 96685483 | 96686956 | Distal Intergenic |
| chr11 | 96690215 | 96690574 | Distal Intergenic |
| chr11 | 96691532 | 96691808 | Distal Intergenic |
| chr11 | 96831034 | 96831900 | Distal Intergenic |
| chr11 | 96833710 | 96835105 | Distal Intergenic |
| chr11 | 96835678 | 96835914 | Distal Intergenic |
| chr11 | 96836106 | 96837042 | Distal Intergenic |
| chr11 | 96837168 | 96837459 | Distal Intergenic |
| chr11 | 96837511 | 96838451 | Distal Intergenic |
| chr11 | 96838768 | 96839003 | Distal Intergenic |
| chr11 | 96841218 | 96841554 | Distal Intergenic |
| chr11 | 96880459 | 96880724 | Distal Intergenic |
| chr11 | 96880770 | 96881144 | Distal Intergenic |
| chr11 | 97049863 | 97050098 | Distal Intergenic |
| chr11 | 97050147 | 97050411 | Distal Intergenic |
| chr11 | 97053257 | 97053668 | Distal Intergenic |
| chr11 | 97068478 | 97068973 | Distal Intergenic |
| chr11 | 97140726 | 97141175 | Distal Intergenic |
| chr11 | 97163888 | 97164097 | Distal Intergenic |
| chr11 | 97170091 | 97171630 | Distal Intergenic |
| chr11 | 97177252 | 97177576 | Distal Intergenic |
| chr11 | 97183241 | 97183579 | Distal Intergenic |
| chr11 | 97233252 | 97234205 | Distal Intergenic |
| chr11 | 97280716 | 97280935 | Distal Intergenic |
| chr11 | 97281132 | 97281349 | Distal Intergenic |
| chr11 | 97288937 | 97289616 | Distal Intergenic |
| chr11 | 97289893 | 97290185 | Distal Intergenic |
| chr11 | 97295904 | 97297958 | Distal Intergenic |
| chr11 | 97299533 | 97299984 | Distal Intergenic |
| chr11 | 97301150 | 97301349 | Distal Intergenic |
| chr11 | 97301906 | 97303782 | Distal Intergenic |
| chr11 | 97310464 | 97311884 | Distal Intergenic |
| chr11 | 97332164 | 97333335 | Distal Intergenic |
| chr11 | 97561135 | 97561496 | Distal Intergenic |
| chr11 | 97561666 | 97561918 | Distal Intergenic |
| chr11 | 97564885 | 97565084 | Distal Intergenic |
| chr11 | 97610117 | 97610319 | Distal Intergenic |
| chr11 | 97627593 | 97628182 | Distal Intergenic |
| chr11 | 97641443 | 97641760 | Distal Intergenic |
| chr11 | 97642046 | 97642357 | Distal Intergenic |
| chr11 | 97642812 | 97643011 | Distal Intergenic |
| chr11 | 97643357 | 97643640 | Distal Intergenic |

| | | | |
|---|---|---|---|
| chr11 | 97661311 | 97661918 | Distal Intergenic |
| chr11 | 97661961 | 97663164 | Distal Intergenic |
| chr11 | 97665188 | 97665392 | Distal Intergenic |
| chr11 | 97669158 | 97669519 | Distal Intergenic |
| chr11 | 97670529 | 97670738 | Distal Intergenic |
| chr11 | 97670832 | 97672454 | Distal Intergenic |
| chr11 | 97672566 | 97673231 | Distal Intergenic |
| chr11 | 97689007 | 97689360 | Distal Intergenic |
| chr11 | 97810930 | 97811167 | Distal Intergenic |
| chr11 | 97811283 | 97812121 | Distal Intergenic |
| chr11 | 97823601 | 97823909 | Distal Intergenic |
| chr11 | 97824041 | 97824382 | Distal Intergenic |
| chr11 | 97849756 | 97850117 | Distal Intergenic |
| chr11 | 97887751 | 97888502 | Distal Intergenic |
| chr11 | 97888574 | 97888859 | Distal Intergenic |
| chr11 | 97918715 | 97919484 | Distal Intergenic |
| chr11 | 97923158 | 97923357 | Distal Intergenic |
| chr11 | 97923568 | 97923878 | Distal Intergenic |
| chr11 | 98064082 | 98064507 | Distal Intergenic |
| chr11 | 98065434 | 98065668 | Distal Intergenic |
| chr11 | 98280110 | 98280309 | Distal Intergenic |
| chr11 | 98280541 | 98280921 | Distal Intergenic |
| chr11 | 98449068 | 98449743 | Distal Intergenic |
| chr11 | 98570061 | 98570304 | Distal Intergenic |
| chr11 | 98570706 | 98570920 | Distal Intergenic |
| chr11 | 98633790 | 98634316 | Distal Intergenic |
| chr11 | 98634450 | 98635122 | Distal Intergenic |
| chr11 | 98635458 | 98635663 | Distal Intergenic |
| chr11 | 98635773 | 98636247 | Distal Intergenic |
| chr11 | 98637065 | 98637544 | Distal Intergenic |
| chr11 | 98655599 | 98656255 | Distal Intergenic |
| chr11 | 98656365 | 98656584 | Distal Intergenic |
| chr11 | 98656745 | 98657447 | Distal Intergenic |
| chr11 | 98657552 | 98657797 | Distal Intergenic |
| chr11 | 98697328 | 98698013 | Distal Intergenic |
| chr11 | 98699607 | 98700660 | Distal Intergenic |
| chr11 | 98793976 | 98794191 | Distal Intergenic |
| chr11 | 98794440 | 98794709 | Distal Intergenic |
| chr11 | 98797200 | 98797597 | Distal Intergenic |
| chr11 | 98797770 | 98798815 | Distal Intergenic |
| chr11 | 98885020 | 98885512 | Distal Intergenic |
| chr11 | 98937810 | 98938876 | Intron |
| chr11 | 98939045 | 98939268 | Intron |
| chr11 | 98941934 | 98950740 | Intron |

| | | | |
|---|---|---|---|
| chr11 | 98950909 | 98951185 | Intron |
| chr11 | 98956222 | 98956894 | Intron |
| chr11 | 98959676 | 98960679 | Intron |
| chr11 | 98965743 | 98966931 | Intron |
| chr11 | 98966986 | 98971584 | Intron |
| chr11 | 98971653 | 98972007 | Intron |
| chr11 | 99060956 | 99061538 | Intron |
| chr11 | 99090812 | 99091439 | Intron |
| chr11 | 99091615 | 99092462 | Intron |
| chr11 | 99130138 | 99131140 | Intron |
| chr11 | 99333586 | 99333858 | Intron |
| chr11 | 99361380 | 99361866 | Intron |
| chr11 | 99377788 | 99379246 | Intron |
| chr11 | 99419427 | 99420025 | Intron |
| chr11 | 99420097 | 99420481 | Intron |
| chr11 | 99442535 | 99442994 | Intron |
| chr11 | 99917194 | 99917963 | Intron |
| chr11 | 99927347 | 99927572 | Intron |
| chr11 | 99927609 | 99929022 | Intron |
| chr11 | 99929073 | 99929483 | Intron |
| chr11 | 99974657 | 99975530 | Intron |
| chr11 | 99975586 | 99976006 | Intron |
| chr11 | 100062808 | 100064708 | Promoter |
| chr11 | 100065387 | 100065714 | Intron |
| chr11 | 100066062 | 100071605 | Intron |
| chr11 | 100109667 | 100109879 | Intron |
| chr11 | 100109925 | 100111354 | Intron |
| chr11 | 100113915 | 100114229 | Intron |
| chr11 | 100114323 | 100115957 | Intron |
| chr11 | 100116129 | 100117375 | Intron |
| chr11 | 100118621 | 100119002 | Intron |
| chr11 | 100119208 | 100122422 | Intron |
| chr11 | 100122674 | 100123729 | Intron |
| chr11 | 100139004 | 100140880 | Intron |
| chr11 | 100142961 | 100143419 | Intron |
| chr11 | 100144091 | 100144546 | Intron |
| chr11 | 100144738 | 100153008 | Intron |
| chr11 | 100153061 | 100153391 | Intron |
| chr11 | 100159805 | 100160614 | Intron |
| chr11 | 100165989 | 100167434 | Intron |
| chr11 | 100181274 | 100181679 | Intron |
| chr11 | 100247823 | 100251133 | Distal Intergenic |
| chr11 | 100251862 | 100252084 | Distal Intergenic |
| chr11 | 100252663 | 100252862 | Distal Intergenic |

| | | | |
|---|---|---|---|
| chr11 | 100254306 | 100255617 | Distal Intergenic |
| chr11 | 100255825 | 100258807 | Distal Intergenic |
| chr11 | 100259522 | 100260030 | Distal Intergenic |
| chr11 | 100267295 | 100269106 | Distal Intergenic |
| chr11 | 100269212 | 100270062 | Distal Intergenic |
| chr11 | 100272083 | 100275035 | Distal Intergenic |
| chr11 | 100450339 | 100451361 | Distal Intergenic |
| chr11 | 100452139 | 100452579 | Distal Intergenic |
| chr11 | 100452770 | 100453058 | Distal Intergenic |
| chr11 | 100480688 | 100480918 | Distal Intergenic |
| chr11 | 100484227 | 100484525 | Distal Intergenic |
| chr11 | 100579587 | 100580068 | Intron |
| chr11 | 100580136 | 100581203 | Intron |
| chr11 | 100581413 | 100584709 | Intron |
| chr11 | 100598240 | 100599032 | Intron |
| chr11 | 100599134 | 100599507 | Intron |
| chr11 | 100792940 | 100793772 | Intron |
| chr11 | 100823209 | 100825415 | Intron |
| chr11 | 100826006 | 100827275 | Intron |
| chr11 | 100827325 | 100827652 | Intron |
| chr11 | 100830921 | 100831410 | Promoter |
| chr11 | 100831498 | 100831714 | Promoter |
| chr11 | 100870438 | 100870651 | Distal Intergenic |
| chr11 | 100943938 | 100944303 | Intron |
| chr11 | 100963498 | 100965944 | Intron |
| chr11 | 100972663 | 100972909 | Intron |
| chr11 | 100999923 | 101000332 | Promoter |
| chr11 | 101033496 | 101033899 | Intron |
| chr11 | 101034169 | 101034404 | Intron |
| chr11 | 101114130 | 101114943 | Distal Intergenic |
| chr11 | 101114999 | 101116216 | Distal Intergenic |
| chr11 | 101118925 | 101119240 | Distal Intergenic |
| chr11 | 101139048 | 101139281 | Distal Intergenic |
| chr11 | 101139382 | 101140130 | Distal Intergenic |
| chr11 | 101163223 | 101163542 | Distal Intergenic |
| chr11 | 101309170 | 101309449 | Distal Intergenic |
| chr11 | 101309524 | 101310369 | Distal Intergenic |
| chr11 | 101488502 | 101489038 | Distal Intergenic |
| chr11 | 101489079 | 101489827 | Distal Intergenic |
| chr11 | 101548608 | 101549984 | Distal Intergenic |
| chr11 | 101648073 | 101648272 | Distal Intergenic |
| chr11 | 102046157 | 102046356 | Intron |
| chr11 | 102106786 | 102106985 | Distal Intergenic |
| chr11 | 102146216 | 102146461 | Distal Intergenic |

| | | | |
|---|---|---|---|
| chr11 | 102156516 | 102157678 | Distal Intergenic |
| chr11 | 102158564 | 102159046 | Distal Intergenic |
| chr11 | 102178668 | 102178949 | Distal Intergenic |
| chr11 | 102179402 | 102180477 | Distal Intergenic |
| chr11 | 102244409 | 102244922 | Intron |
| chr11 | 102249089 | 102249314 | 3' UTR |
| chr11 | 102270053 | 102270318 | Intron |
| chr11 | 102286323 | 102286664 | Intron |
| chr11 | 102327615 | 102328871 | Distal Intergenic |
| chr11 | 102397414 | 102398632 | 3' UTR |
| chr11 | 102416811 | 102417117 | Distal Intergenic |
| chr11 | 102505152 | 102506163 | Distal Intergenic |
| chr11 | 102517407 | 102518062 | Distal Intergenic |
| chr11 | 102742119 | 102742472 | Exon |
| chr11 | 102785797 | 102786408 | Distal Intergenic |
| chr11 | 102786554 | 102786979 | Distal Intergenic |
| chr11 | 102820008 | 102820441 | Intron |
| chr11 | 102857591 | 102858251 | Distal Intergenic |
| chr11 | 102864537 | 102865233 | Distal Intergenic |
| chr11 | 102889364 | 102889575 | Distal Intergenic |
| chr11 | 102912317 | 102914125 | Distal Intergenic |
| chr11 | 102955298 | 102955653 | Intron |
| chr11 | 102955719 | 102956285 | Intron |
| chr11 | 102956346 | 102956741 | Intron |
| chr11 | 102957930 | 102959662 | Intron |
| chr11 | 102961581 | 102961991 | Promoter |
| chr11 | 102962373 | 102962665 | Promoter |
| chr11 | 102963030 | 102963500 | Promoter |
| chr11 | 102976376 | 102977344 | Distal Intergenic |
| chr11 | 102993611 | 102994046 | Exon |
| chr11 | 103098468 | 103098667 | Intron |
| chr11 | 103107394 | 103108651 | Intron |
| chr11 | 103123783 | 103124693 | Exon |
| chr11 | 103131266 | 103131465 | Intron |
| chr11 | 103131648 | 103131878 | Intron |
| chr11 | 103160204 | 103160890 | Intron |
| chr11 | 103173129 | 103173982 | Exon |
| chr11 | 103195792 | 103197319 | Intron |
| chr11 | 103201576 | 103202732 | Intron |
| chr11 | 103222987 | 103223228 | Intron |
| chr11 | 103223345 | 103223578 | Intron |
| chr11 | 103223629 | 103225453 | Intron |
| chr11 | 103239469 | 103240137 | Intron |
| chr11 | 103242954 | 103244408 | Intron |

| | | | |
|---|---|---|---|
| chr11 | 103257835 | 103258721 | Intron |
| chr11 | 103545597 | 103545847 | Distal Intergenic |
| chr11 | 103546019 | 103547263 | Distal Intergenic |
| chr11 | 103550254 | 103550586 | Distal Intergenic |
| chr11 | 103551052 | 103551284 | Distal Intergenic |
| chr11 | 103558451 | 103559825 | Distal Intergenic |
| chr11 | 104149832 | 104150802 | Distal Intergenic |
| chr11 | 104150842 | 104151543 | Distal Intergenic |
| chr11 | 104164808 | 104165044 | Distal Intergenic |
| chr11 | 104239189 | 104239452 | Distal Intergenic |
| chr11 | 104302037 | 104302704 | Distal Intergenic |
| chr11 | 104302776 | 104303015 | Distal Intergenic |
| chr11 | 104303827 | 104305146 | Distal Intergenic |
| chr11 | 104411384 | 104411621 | Distal Intergenic |
| chr11 | 104534779 | 104535602 | Distal Intergenic |
| chr11 | 104553298 | 104554003 | Distal Intergenic |
| chr11 | 105021163 | 105021362 | Distal Intergenic |
| chr11 | 105042941 | 105043268 | Distal Intergenic |
| chr11 | 105043320 | 105044865 | Distal Intergenic |
| chr11 | 105191359 | 105192788 | Distal Intergenic |
| chr11 | 105301117 | 105301485 | Distal Intergenic |
| chr11 | 105328647 | 105329130 | Distal Intergenic |
| chr11 | 105329180 | 105329382 | Distal Intergenic |
| chr11 | 105654048 | 105654263 | Intron |
| chr11 | 105684331 | 105685094 | Intron |
| chr11 | 105744899 | 105745124 | Intron |
| chr11 | 105962770 | 105963138 | Intron |
| chr11 | 106054467 | 106054836 | Distal Intergenic |
| chr11 | 106055147 | 106055408 | Distal Intergenic |
| chr11 | 106078252 | 106078934 | Distal Intergenic |
| chr11 | 106116765 | 106117960 | Distal Intergenic |
| chr11 | 106133279 | 106133538 | Exon |
| chr11 | 106277282 | 106278485 | Distal Intergenic |
| chr11 | 106331349 | 106332531 | Distal Intergenic |
| chr11 | 106345017 | 106345812 | Distal Intergenic |
| chr11 | 106362128 | 106363917 | Distal Intergenic |
| chr11 | 106510315 | 106510572 | Distal Intergenic |
| chr11 | 106519252 | 106519995 | Distal Intergenic |
| chr11 | 106526532 | 106528490 | Distal Intergenic |
| chr11 | 106532272 | 106532699 | Distal Intergenic |
| chr11 | 106535766 | 106536770 | Distal Intergenic |
| chr11 | 106574205 | 106576237 | Intron |
| chr11 | 106583786 | 106584408 | Intron |
| chr11 | 106641045 | 106641502 | Intron |

| | | | |
|---|---|---|---|
| chr11 | 106648537 | 106648827 | Intron |
| chr11 | 106649577 | 106649786 | Intron |
| chr11 | 106650292 | 106651129 | Intron |
| chr11 | 106758084 | 106758334 | Intron |
| chr11 | 106860028 | 106860312 | Intron |
| chr11 | 106860520 | 106860825 | Intron |
| chr11 | 106873454 | 106873963 | Intron |
| chr11 | 106993775 | 106994012 | Distal Intergenic |
| chr11 | 106999170 | 106999690 | Distal Intergenic |
| chr11 | 107009842 | 107010051 | Distal Intergenic |
| chr11 | 107010163 | 107011917 | Distal Intergenic |
| chr11 | 107018101 | 107018678 | Distal Intergenic |
| chr11 | 107018928 | 107019212 | Distal Intergenic |
| chr11 | 107050234 | 107050464 | Distal Intergenic |
| chr11 | 107220206 | 107220567 | Intron |
| chr11 | 107278199 | 107279019 | Intron |
| chr11 | 107289258 | 107290138 | Intron |
| chr11 | 107309965 | 107311357 | Intron |
| chr11 | 107319528 | 107320085 | Intron |
| chr11 | 107331511 | 107331947 | Distal Intergenic |
| chr11 | 107340836 | 107341387 | Distal Intergenic |
| chr11 | 107880769 | 107882214 | Intron |
| chr11 | 107982431 | 107982643 | Distal Intergenic |
| chr11 | 108076907 | 108077159 | Intron |
| chr11 | 108078117 | 108078867 | Intron |
| chr11 | 108129236 | 108129476 | Intron |
| chr11 | 108144057 | 108144802 | Intron |
| chr11 | 108209467 | 108209829 | Intron |
| chr11 | 108226726 | 108227204 | Exon |
| chr11 | 108613782 | 108614045 | Intron |
| chr11 | 108614340 | 108615079 | Intron |
| chr11 | 108618981 | 108619252 | Intron |
| chr11 | 108619294 | 108619821 | Intron |
| chr11 | 108758949 | 108760244 | Intron |
| chr11 | 109216922 | 109217916 | Distal Intergenic |
| chr11 | 109221557 | 109221791 | Distal Intergenic |
| chr11 | 109221885 | 109223850 | Distal Intergenic |
| chr11 | 109470275 | 109470576 | Distal Intergenic |
| chr11 | 109471110 | 109471593 | Distal Intergenic |
| chr11 | 109471699 | 109472034 | Distal Intergenic |
| chr11 | 109522815 | 109523014 | Distal Intergenic |
| chr11 | 109523505 | 109523817 | Distal Intergenic |
| chr11 | 109530789 | 109531342 | Distal Intergenic |
| chr11 | 109531413 | 109533181 | Distal Intergenic |

| | | | |
|---|---|---|---|
| chr11 | 109533219 | 109533741 | Distal Intergenic |
| chr11 | 109559238 | 109560526 | Distal Intergenic |
| chr11 | 109678290 | 109678491 | Distal Intergenic |
| chr11 | 109721475 | 109721940 | Distal Intergenic |
| chr11 | 110240559 | 110241660 | Intron |
| chr11 | 110300212 | 110300411 | Promoter |
| chr11 | 110309983 | 110312125 | Intron |
| chr11 | 110333206 | 110333484 | 3' UTR |
| chr11 | 110333565 | 110333971 | 3' UTR |
| chr11 | 110640357 | 110641272 | Distal Intergenic |
| chr11 | 110643178 | 110643719 | Distal Intergenic |
| chr11 | 110643823 | 110644034 | Distal Intergenic |
| chr11 | 112027126 | 112027692 | Intron |
| chr11 | 112060510 | 112061715 | Intron |
| chr11 | 112112884 | 112113128 | Intron |
| chr11 | 112113282 | 112115212 | Intron |
| chr11 | 112163072 | 112164544 | Distal Intergenic |
| chr11 | 112164597 | 112164821 | Distal Intergenic |
| chr11 | 112331959 | 112332272 | Distal Intergenic |
| chr11 | 112481815 | 112482903 | Distal Intergenic |
| chr11 | 112492650 | 112493371 | Distal Intergenic |
| chr11 | 112522287 | 112522583 | Distal Intergenic |
| chr11 | 112522678 | 112523022 | Distal Intergenic |
| chr11 | 112523126 | 112523686 | Distal Intergenic |
| chr11 | 112849262 | 112849609 | Intron |
| chr11 | 112918029 | 112918528 | Intron |
| chr11 | 112921797 | 112923023 | Intron |
| chr11 | 112924171 | 112925152 | Intron |
| chr11 | 112935489 | 112936677 | Intron |
| chr11 | 112961729 | 112965483 | Intron |
| chr11 | 112965806 | 112967841 | Intron |
| chr11 | 112994203 | 112994440 | Intron |
| chr11 | 113057642 | 113057854 | Intron |
| chr11 | 113057954 | 113058337 | Intron |
| chr11 | 113059582 | 113060353 | Intron |
| chr11 | 113060414 | 113061447 | Intron |
| chr11 | 113070597 | 113070946 | Intron |
| chr11 | 113398142 | 113398988 | Distal Intergenic |
| chr11 | 113612058 | 113612390 | Intron |
| chr11 | 113612516 | 113613119 | Intron |
| chr11 | 113616273 | 113617843 | Intron |
| chr11 | 113628192 | 113628494 | Exon |
| chr11 | 114529927 | 114530170 | Distal Intergenic |
| chr11 | 114530838 | 114531073 | Distal Intergenic |

| | | | |
|---|---|---|---|
| chr11 | 115033921 | 115035678 | Distal Intergenic |
| chr11 | 115375319 | 115375718 | Promoter |
| chr11 | 115425560 | 115425763 | Distal Intergenic |
| chr11 | 115608260 | 115608544 | Distal Intergenic |
| chr11 | 115637202 | 115637457 | Distal Intergenic |
| chr11 | 115637720 | 115638050 | Distal Intergenic |
| chr11 | 115638132 | 115639084 | Distal Intergenic |
| chr11 | 115639153 | 115639548 | Distal Intergenic |
| chr11 | 115688508 | 115688724 | Distal Intergenic |
| chr11 | 115730459 | 115730889 | Distal Intergenic |
| chr11 | 115794938 | 115795148 | Distal Intergenic |
| chr11 | 115805432 | 115808068 | Distal Intergenic |
| chr11 | 115815572 | 115817192 | Distal Intergenic |
| chr11 | 115817285 | 115819440 | Distal Intergenic |
| chr11 | 115830444 | 115833158 | Distal Intergenic |
| chr11 | 115833239 | 115833519 | Distal Intergenic |
| chr11 | 115835096 | 115836308 | Distal Intergenic |
| chr11 | 115836714 | 115838324 | Distal Intergenic |
| chr11 | 115885200 | 115885467 | Distal Intergenic |
| chr11 | 115892079 | 115892440 | Distal Intergenic |
| chr11 | 116028584 | 116028848 | Distal Intergenic |
| chr11 | 116028924 | 116029200 | Distal Intergenic |
| chr11 | 116029242 | 116029489 | Distal Intergenic |
| chr11 | 116116208 | 116116609 | Distal Intergenic |
| chr11 | 116117226 | 116118341 | Distal Intergenic |
| chr11 | 116267502 | 116267936 | Distal Intergenic |
| chr11 | 116383122 | 116383718 | Distal Intergenic |
| chr11 | 116482318 | 116482686 | Distal Intergenic |
| chr11 | 116482735 | 116483013 | Distal Intergenic |
| chr11 | 116483317 | 116483606 | Distal Intergenic |
| chr11 | 116703906 | 116704168 | Downstream |
| chr11 | 116704311 | 116704612 | Downstream |
| chr11 | 116714245 | 116714560 | 3' UTR |
| chr11 | 116714625 | 116714839 | 3' UTR |
| chr11 | 116942577 | 116943048 | Intron |
| chr11 | 116943880 | 116944079 | Intron |
| chr11 | 117061158 | 117061515 | Exon |
| chr11 | 117063112 | 117064105 | Exon |
| chr11 | 117064166 | 117065089 | Exon |
| chr11 | 117065630 | 117067571 | 3' UTR |
| chr11 | 117068841 | 117069545 | Promoter |
| chr11 | 117117217 | 117117521 | Intron |
| chr11 | 117120617 | 117120949 | Intron |
| chr11 | 117120994 | 117122296 | Intron |

| | | | |
|---|---|---|---|
| chr11 | 117124856 | 117125120 | Intron |
| chr11 | 117125171 | 117126021 | Intron |
| chr11 | 117126161 | 117126378 | Intron |
| chr11 | 117126419 | 117127102 | Intron |
| chr11 | 117185082 | 117185438 | Intron |
| chr11 | 117185571 | 117185938 | Intron |
| chr11 | 117195277 | 117195476 | Intron |
| chr11 | 117195517 | 117196161 | Intron |
| chr11 | 117429514 | 117430863 | Intron |
| chr11 | 117568390 | 117568921 | Intron |
| chr11 | 117642144 | 117643383 | Intron |
| chr11 | 117649749 | 117649948 | Intron |
| chr11 | 117650232 | 117650481 | Intron |
| chr11 | 117651748 | 117652002 | Intron |
| chr11 | 117652111 | 117652633 | Intron |
| chr11 | 117652750 | 117652963 | Intron |
| chr11 | 117672101 | 117672338 | Intron |
| chr11 | 117685594 | 117686109 | Intron |
| chr11 | 117686531 | 117687954 | Promoter |
| chr11 | 117704645 | 117705367 | Intron |
| chr11 | 117706323 | 117706564 | Intron |
| chr11 | 117709509 | 117709836 | Exon |
| chr11 | 117710045 | 117710356 | Exon |
| chr11 | 117710640 | 117711051 | Exon |
| chr11 | 117803053 | 117803508 | Distal Intergenic |
| chr11 | 117819073 | 117819517 | Distal Intergenic |
| chr11 | 117819587 | 117820022 | Distal Intergenic |
| chr11 | 117848743 | 117849013 | Distal Intergenic |
| chr11 | 117851372 | 117851602 | Distal Intergenic |
| chr11 | 117855243 | 117855886 | Distal Intergenic |
| chr11 | 117868354 | 117868947 | Intron |
| chr11 | 117869222 | 117869755 | Exon |
| chr11 | 117871057 | 117871583 | 3' UTR |
| chr11 | 117873378 | 117873662 | Distal Intergenic |
| chr11 | 117873774 | 117874776 | Distal Intergenic |
| chr11 | 117890608 | 117890861 | Intron |
| chr11 | 118014270 | 118015481 | Promoter |
| chr11 | 118017647 | 118019358 | Intron |
| chr11 | 118019454 | 118019714 | Intron |
| chr11 | 118021565 | 118022576 | Intron |
| chr11 | 118035932 | 118037166 | 3' UTR |
| chr11 | 118048777 | 118049021 | Distal Intergenic |
| chr11 | 118049107 | 118049307 | Distal Intergenic |
| chr11 | 118073953 | 118074233 | Promoter |

| | | | |
|---|---|---|---|
| chr11 | 118074272 | 118074484 | Promoter |
| chr11 | 118076839 | 118078628 | Intron |
| chr11 | 118208016 | 118209030 | Downstream |
| chr11 | 118712874 | 118713631 | Distal Intergenic |
| chr11 | 118966572 | 118967709 | Promoter |
| chr11 | 118968713 | 118969404 | 3' UTR |
| chr11 | 118989313 | 118989687 | Distal Intergenic |
| chr11 | 119011625 | 119012328 | Distal Intergenic |
| chr11 | 119012765 | 119013754 | Distal Intergenic |
| chr11 | 119046110 | 119046454 | Intron |
| chr11 | 119254688 | 119254900 | Intron |
| chr11 | 119263382 | 119264830 | Intron |
| chr11 | 119264896 | 119265128 | Intron |
| chr11 | 119546984 | 119547736 | Intron |
| chr11 | 119556953 | 119557938 | Intron |
| chr11 | 119679141 | 119679397 | Distal Intergenic |
| chr11 | 119718021 | 119718511 | Distal Intergenic |
| chr11 | 119794678 | 119795501 | Distal Intergenic |
| chr11 | 119798205 | 119799302 | Distal Intergenic |
| chr11 | 119829646 | 119830913 | Distal Intergenic |
| chr11 | 119836716 | 119838113 | Distal Intergenic |
| chr11 | 119850059 | 119850556 | Distal Intergenic |
| chr11 | 120114380 | 120114579 | Intron |
| chr11 | 120210166 | 120211276 | Intron |
| chr11 | 120211469 | 120211725 | Intron |
| chr11 | 120368496 | 120370407 | Distal Intergenic |
| chr11 | 120371276 | 120371603 | Distal Intergenic |
| chr11 | 120414325 | 120414534 | Intron |
| chr11 | 120416105 | 120416400 | Intron |
| chr11 | 120416445 | 120416692 | Intron |
| chr11 | 120435191 | 120435560 | Promoter |
| chr11 | 120435702 | 120435901 | Promoter |
| chr11 | 120491306 | 120492270 | Intron |
| chr11 | 120492419 | 120492634 | Intron |
| chr11 | 120493387 | 120493647 | Intron |
| chr11 | 120493754 | 120494171 | Intron |
| chr11 | 120534689 | 120534895 | Intron |
| chr11 | 120559036 | 120559459 | Intron |
| chr11 | 120559517 | 120559798 | Intron |
| chr11 | 120565414 | 120565681 | Intron |
| chr11 | 120569197 | 120569896 | Intron |
| chr11 | 120593375 | 120593871 | Intron |
| chr11 | 120594184 | 120594726 | Intron |
| chr11 | 120684614 | 120684975 | Intron |

| | | | |
|---|---|---|---|
| chr11 | 120807525 | 120807747 | Intron |
| chr11 | 120807796 | 120808007 | Intron |
| chr11 | 120845460 | 120846712 | Intron |
| chr11 | 120846921 | 120847233 | Intron |
| chr11 | 120852285 | 120852613 | Intron |
| chr11 | 121098029 | 121098248 | Distal Intergenic |
| chr11 | 121376858 | 121378533 | Intron |
| chr11 | 121380797 | 121381190 | Intron |
| chr11 | 121381398 | 121381936 | Intron |
| chr11 | 121405019 | 121405419 | Intron |
| chr11 | 121405516 | 121405920 | Intron |
| chr11 | 121405961 | 121407145 | Intron |
| chr11 | 121458523 | 121458952 | Exon |
| chr11 | 121462660 | 121463329 | Intron |
| chr11 | 121507529 | 121507853 | Distal Intergenic |
| chr11 | 121507910 | 121508761 | Distal Intergenic |
| chr12 | 3082039 | 3082857 | Intron |
| chr12 | 3174126 | 3174331 | Distal Intergenic |
| chr12 | 3174382 | 3174608 | Distal Intergenic |
| chr12 | 3284745 | 3285300 | Intron |
| chr12 | 3308618 | 3309014 | Intron |
| chr12 | 3310153 | 3310963 | 5' UTR |
| chr12 | 3319376 | 3319811 | Intron |
| chr12 | 3329553 | 3330404 | Intron |
| chr12 | 3348946 | 3351143 | Intron |
| chr12 | 3351991 | 3353029 | Intron |
| chr12 | 3426088 | 3426692 | Distal Intergenic |
| chr12 | 3426775 | 3427106 | Distal Intergenic |
| chr12 | 3427176 | 3428031 | Exon |
| chr12 | 3537313 | 3538768 | Intron |
| chr12 | 3582648 | 3582854 | Intron |
| chr12 | 3719408 | 3719891 | Intron |
| chr12 | 3746137 | 3746412 | Intron |
| chr12 | 4233442 | 4234718 | Distal Intergenic |
| chr12 | 4292533 | 4292755 | Distal Intergenic |
| chr12 | 4357022 | 4357539 | Distal Intergenic |
| chr12 | 4595556 | 4595859 | Downstream |
| chr12 | 4596968 | 4598278 | 3' UTR |
| chr12 | 4598827 | 4599505 | 3' UTR |
| chr12 | 4599615 | 4600006 | Exon |
| chr12 | 4600170 | 4600740 | Exon |
| chr12 | 4606129 | 4606635 | Intron |
| chr12 | 4613368 | 4613941 | Intron |
| chr12 | 4824054 | 4824253 | Distal Intergenic |

| | | | |
|---|---|---|---|
| chr12 | 4824318 | 4825884 | Distal Intergenic |
| chr12 | 4825925 | 4826854 | Distal Intergenic |
| chr12 | 4924174 | 4925547 | Intron |
| chr12 | 4981351 | 4981600 | Distal Intergenic |
| chr12 | 5352909 | 5353128 | Distal Intergenic |
| chr12 | 5353250 | 5354066 | Distal Intergenic |
| chr12 | 5356284 | 5357896 | Distal Intergenic |
| chr12 | 5357936 | 5358177 | Distal Intergenic |
| chr12 | 8077481 | 8078525 | Exon |
| chr12 | 8082410 | 8082687 | Exon |
| chr12 | 8082746 | 8083021 | Intron |
| chr12 | 8083104 | 8083434 | Exon |
| chr12 | 8214964 | 8215346 | Intron |
| chr12 | 8369274 | 8369591 | Distal Intergenic |
| chr12 | 8371151 | 8371755 | Downstream |
| chr12 | 8378611 | 8380004 | Promoter |
| chr12 | 8385401 | 8385890 | Intron |
| chr12 | 8385947 | 8386399 | Intron |
| chr12 | 8386519 | 8387429 | Exon |
| chr12 | 8393070 | 8394708 | Promoter |
| chr12 | 8394905 | 8396890 | Promoter |
| chr12 | 8494912 | 8495131 | Distal Intergenic |
| chr12 | 8506555 | 8506832 | Downstream |
| chr12 | 8516434 | 8518777 | Intron |
| chr12 | 8526226 | 8526593 | Intron |
| chr12 | 8526750 | 8527479 | Intron |
| chr12 | 8527521 | 8528557 | Intron |
| chr12 | 8528609 | 8528975 | Intron |
| chr12 | 8559023 | 8559278 | Distal Intergenic |
| chr12 | 8680638 | 8680837 | Distal Intergenic |
| chr12 | 8681444 | 8682748 | Distal Intergenic |
| chr12 | 8767336 | 8768720 | Distal Intergenic |
| chr12 | 8778810 | 8791841 | Distal Intergenic |
| chr12 | 8791962 | 8792642 | Distal Intergenic |
| chr12 | 8793445 | 8794013 | Distal Intergenic |
| chr12 | 8803815 | 8805346 | Exon |
| chr12 | 8805916 | 8807188 | Exon |
| chr12 | 8812179 | 8813156 | Intron |
| chr12 | 8813750 | 8814192 | Intron |
| chr12 | 8814405 | 8814618 | Promoter |
| chr12 | 8815834 | 8820092 | Promoter |
| chr12 | 8824592 | 8824946 | Distal Intergenic |
| chr12 | 8825106 | 8825376 | Distal Intergenic |
| chr12 | 8828744 | 8829022 | Distal Intergenic |

| | | | |
|---|---|---|---|
| chr12 | 8831859 | 8832242 | Distal Intergenic |
| chr12 | 8832511 | 8832723 | Distal Intergenic |
| chr12 | 8833416 | 8834660 | Promoter |
| chr12 | 8861096 | 8861733 | Intron |
| chr12 | 8864957 | 8867386 | 5' UTR |
| chr12 | 8867547 | 8868492 | Intron |
| chr12 | 8871268 | 8871553 | Intron |
| chr12 | 8871593 | 8871842 | Intron |
| chr12 | 8871884 | 8872267 | Intron |
| chr12 | 8880425 | 8883191 | Intron |
| chr12 | 8891396 | 8892744 | Intron |
| chr12 | 8903656 | 8904066 | Intron |
| chr12 | 8922839 | 8923711 | Intron |
| chr12 | 8927720 | 8930486 | 3' UTR |
| chr12 | 8930548 | 8930824 | Intron |
| chr12 | 8934630 | 8935069 | Intron |
| chr12 | 8935493 | 8935797 | Exon |
| chr12 | 8976571 | 8976962 | Intron |
| chr12 | 8981444 | 8981817 | Intron |
| chr12 | 9072836 | 9073367 | Intron |
| chr12 | 9118533 | 9120113 | Distal Intergenic |
| chr12 | 9236587 | 9236993 | Intron |
| chr12 | 9238815 | 9239566 | Intron |
| chr12 | 9303274 | 9303517 | Exon |
| chr12 | 9303685 | 9304079 | Intron |
| chr12 | 9304116 | 9304381 | Exon |
| chr12 | 9305273 | 9305502 | Exon |
| chr12 | 9305665 | 9306495 | Exon |
| chr12 | 9310383 | 9311784 | Exon |
| chr12 | 9694658 | 9696932 | Distal Intergenic |
| chr12 | 9862539 | 9862874 | Distal Intergenic |
| chr12 | 10397595 | 10398306 | Distal Intergenic |
| chr12 | 11271902 | 11272766 | Intron |
| chr12 | 11273438 | 11274193 | Intron |
| chr12 | 11394215 | 11399749 | Intron |
| chr12 | 12549431 | 12549786 | Intron |
| chr12 | 12895264 | 12895652 | Intron |
| chr12 | 12988276 | 12988475 | Distal Intergenic |
| chr12 | 12988644 | 12989327 | Distal Intergenic |
| chr12 | 12989474 | 12990430 | Distal Intergenic |
| chr12 | 12990648 | 12991229 | Distal Intergenic |
| chr12 | 13083209 | 13084414 | Distal Intergenic |
| chr12 | 13099373 | 13100074 | Intron |
| chr12 | 13182633 | 13184083 | Intron |

| | | | |
|---|---|---|---|
| chr12 | 14075632 | 14075999 | Intron |
| chr12 | 14076208 | 14076542 | Intron |
| chr12 | 14216390 | 14216620 | Distal Intergenic |
| chr12 | 14216678 | 14217320 | Distal Intergenic |
| chr12 | 14219516 | 14220089 | Distal Intergenic |
| chr12 | 14293136 | 14293602 | Distal Intergenic |
| chr12 | 14293654 | 14293853 | Distal Intergenic |
| chr12 | 14307970 | 14309911 | Distal Intergenic |
| chr12 | 15839409 | 15839623 | Intron |
| chr12 | 15839739 | 15839968 | Intron |
| chr12 | 16673891 | 16674149 | Distal Intergenic |
| chr12 | 16674209 | 16674531 | Distal Intergenic |
| chr12 | 16840986 | 16841689 | Distal Intergenic |
| chr12 | 16841749 | 16842216 | Distal Intergenic |
| chr12 | 16847648 | 16848446 | Distal Intergenic |
| chr12 | 16852183 | 16852499 | Distal Intergenic |
| chr12 | 16860199 | 16860580 | Distal Intergenic |
| chr12 | 16901758 | 16903386 | Distal Intergenic |
| chr12 | 16926831 | 16927756 | Distal Intergenic |
| chr12 | 17006024 | 17006290 | Distal Intergenic |
| chr12 | 17006979 | 17007224 | Distal Intergenic |
| chr12 | 17046215 | 17046622 | Distal Intergenic |
| chr12 | 17046888 | 17047559 | Distal Intergenic |
| chr12 | 17227237 | 17228311 | Distal Intergenic |
| chr12 | 17273767 | 17273970 | Distal Intergenic |
| chr12 | 17306367 | 17307424 | Distal Intergenic |
| chr12 | 17309951 | 17310722 | Distal Intergenic |
| chr12 | 17310901 | 17311155 | Distal Intergenic |
| chr12 | 17354624 | 17354893 | Distal Intergenic |
| chr12 | 17354983 | 17355714 | Distal Intergenic |
| chr12 | 17394121 | 17394906 | Distal Intergenic |
| chr12 | 17397069 | 17397462 | Distal Intergenic |
| chr12 | 17426470 | 17428110 | Distal Intergenic |
| chr12 | 17516899 | 17518364 | Distal Intergenic |
| chr12 | 17547390 | 17547658 | Distal Intergenic |
| chr12 | 17549133 | 17550215 | Distal Intergenic |
| chr12 | 17559115 | 17559771 | Distal Intergenic |
| chr12 | 17700619 | 17701433 | Distal Intergenic |
| chr12 | 17701472 | 17701798 | Distal Intergenic |
| chr12 | 17727907 | 17728521 | Distal Intergenic |
| chr12 | 17735867 | 17736790 | Distal Intergenic |
| chr12 | 17739541 | 17739970 | Distal Intergenic |
| chr12 | 17768647 | 17768846 | Distal Intergenic |
| chr12 | 17768885 | 17770231 | Distal Intergenic |

| | | | |
|---|---|---|---|
| chr12 | 21172283 | 21172701 | Exon |
| chr12 | 21182279 | 21182637 | Intron |
| chr12 | 21232824 | 21233605 | Intron |
| chr12 | 21378779 | 21380144 | Intron |
| chr12 | 21422491 | 21422897 | Exon |
| chr12 | 21422943 | 21423451 | Intron |
| chr12 | 21430626 | 21431574 | Intron |
| chr12 | 22077763 | 22077963 | 3' UTR |
| chr12 | 25183409 | 25183643 | Distal Intergenic |
| chr12 | 25202626 | 25203896 | Distal Intergenic |
| chr12 | 25210179 | 25210585 | Intron |
| chr12 | 25220278 | 25223608 | 5' UTR |
| chr12 | 25235389 | 25236081 | Intron |
| chr12 | 25237420 | 25238092 | Intron |
| chr12 | 25277648 | 25277894 | Intron |
| chr12 | 25278018 | 25278232 | Intron |
| chr12 | 25278578 | 25278777 | Intron |
| chr12 | 25284491 | 25285582 | Intron |
| chr12 | 25391898 | 25393203 | Intron |
| chr12 | 25393300 | 25394993 | Intron |
| chr12 | 25408407 | 25408820 | Distal Intergenic |
| chr12 | 25408866 | 25409351 | Distal Intergenic |
| chr12 | 25561605 | 25562826 | Distal Intergenic |
| chr12 | 25647950 | 25649579 | 3' UTR |
| chr12 | 25651549 | 25652003 | Intron |
| chr12 | 25690507 | 25690717 | Intron |
| chr12 | 25697462 | 25698010 | Intron |
| chr12 | 25700295 | 25700592 | Intron |
| chr12 | 25779737 | 25779936 | Intron |
| chr12 | 25780151 | 25780482 | Intron |
| chr12 | 25781080 | 25781364 | Intron |
| chr12 | 25795138 | 25796237 | Intron |
| chr12 | 26112534 | 26114479 | Promoter |
| chr12 | 26859803 | 26860043 | Intron |
| chr12 | 28174769 | 28175708 | Distal Intergenic |
| chr12 | 28176913 | 28177737 | Distal Intergenic |
| chr12 | 28683412 | 28685257 | Intron |
| chr12 | 28685321 | 28685612 | Intron |
| chr12 | 28685649 | 28685932 | Intron |
| chr12 | 28686933 | 28687223 | Intron |
| chr12 | 29320166 | 29321095 | Intron |
| chr12 | 29412083 | 29412451 | Intron |
| chr12 | 29482406 | 29483143 | Intron |
| chr12 | 29489288 | 29490644 | Downstream |

| | | | |
|---|---|---|---|
| chr12 | 30434319 | 30434685 | Distal Intergenic |
| chr12 | 30677149 | 30677482 | Distal Intergenic |
| chr12 | 30677703 | 30677978 | Distal Intergenic |
| chr12 | 30688437 | 30688754 | Distal Intergenic |
| chr12 | 31596206 | 31596405 | Intron |
| chr12 | 31596572 | 31596771 | Intron |
| chr12 | 31743941 | 31745101 | Promoter |
| chr12 | 31745204 | 31745810 | Intron |
| chr12 | 31745914 | 31746331 | Intron |
| chr12 | 31746761 | 31747404 | Exon |
| chr12 | 31753979 | 31754818 | Exon |
| chr12 | 31758955 | 31759381 | Intron |
| chr12 | 31760044 | 31760368 | Intron |
| chr12 | 31760906 | 31761107 | Intron |
| chr12 | 31762986 | 31763499 | Intron |
| chr12 | 31768045 | 31769011 | Exon |
| chr12 | 31870132 | 31870489 | Intron |
| chr12 | 31871471 | 31872098 | Intron |
| chr12 | 31955830 | 31956478 | Distal Intergenic |
| chr12 | 31973571 | 31974380 | Distal Intergenic |
| chr12 | 31976929 | 31977196 | Distal Intergenic |
| chr12 | 31977249 | 31977778 | Distal Intergenic |
| chr12 | 32036058 | 32036257 | Distal Intergenic |
| chr12 | 32100924 | 32101216 | Distal Intergenic |
| chr12 | 32101319 | 32101546 | Distal Intergenic |
| chr12 | 32198481 | 32198937 | Distal Intergenic |
| chr12 | 32248653 | 32249481 | Distal Intergenic |
| chr12 | 32339226 | 32339645 | Intron |
| chr12 | 32532985 | 32534368 | Distal Intergenic |
| chr12 | 32634089 | 32634308 | Distal Intergenic |
| chr12 | 32658583 | 32658976 | Intron |
| chr12 | 32676954 | 32677292 | Intron |
| chr12 | 32681561 | 32681839 | Intron |
| chr12 | 32742088 | 32742504 | Intron |
| chr12 | 33119478 | 33120523 | Distal Intergenic |
| chr12 | 33306187 | 33306463 | Distal Intergenic |
| chr12 | 33306580 | 33307764 | Distal Intergenic |
| chr12 | 33307852 | 33308264 | Distal Intergenic |
| chr12 | 33308652 | 33309115 | Distal Intergenic |
| chr12 | 33504425 | 33504881 | Distal Intergenic |
| chr12 | 33639397 | 33639807 | Distal Intergenic |
| chr12 | 33679504 | 33679771 | Distal Intergenic |
| chr12 | 33680143 | 33680409 | Distal Intergenic |
| chr12 | 33704779 | 33706276 | Distal Intergenic |

| | | | |
|---|---|---|---|
| chr12 | 33857430 | 33858155 | Distal Intergenic |
| chr12 | 33858558 | 33859011 | Distal Intergenic |
| chr12 | 33866625 | 33866837 | Distal Intergenic |
| chr12 | 33970490 | 33970689 | Distal Intergenic |
| chr12 | 34114631 | 34114865 | Distal Intergenic |
| chr12 | 34724782 | 34725953 | Distal Intergenic |
| chr12 | 34760766 | 34760967 | Distal Intergenic |
| chr12 | 35731878 | 35732298 | Distal Intergenic |
| chr12 | 35893889 | 35894629 | Distal Intergenic |
| chr12 | 35894714 | 35895521 | Distal Intergenic |
| chr12 | 35902097 | 35902484 | Distal Intergenic |
| chr12 | 36218889 | 36219088 | Distal Intergenic |
| chr12 | 36247460 | 36248310 | Distal Intergenic |
| chr12 | 36337186 | 36337385 | Distal Intergenic |
| chr12 | 36340971 | 36341237 | Distal Intergenic |
| chr12 | 36341396 | 36341603 | Distal Intergenic |
| chr12 | 36355578 | 36357865 | Distal Intergenic |
| chr12 | 36387105 | 36388179 | Distal Intergenic |
| chr12 | 36388358 | 36388633 | Distal Intergenic |
| chr12 | 36397337 | 36397833 | Distal Intergenic |
| chr12 | 36418920 | 36419223 | Distal Intergenic |
| chr12 | 36860741 | 36861070 | Distal Intergenic |
| chr12 | 36883762 | 36884132 | Distal Intergenic |
| chr12 | 36884171 | 36884562 | Distal Intergenic |
| chr12 | 36942086 | 36942320 | Distal Intergenic |
| chr12 | 36942398 | 36944206 | Distal Intergenic |
| chr12 | 36989009 | 36989223 | Distal Intergenic |
| chr12 | 36993720 | 36995591 | Distal Intergenic |
| chr12 | 37016954 | 37017218 | Distal Intergenic |
| chr12 | 37107959 | 37108337 | Distal Intergenic |
| chr12 | 37108487 | 37108768 | Distal Intergenic |
| chr12 | 37186685 | 37187391 | Distal Intergenic |
| chr12 | 40556829 | 40558927 | Distal Intergenic |
| chr12 | 40558981 | 40559259 | Distal Intergenic |
| chr12 | 40608816 | 40609577 | Distal Intergenic |
| chr12 | 40646839 | 40647270 | Intron |
| chr12 | 40657042 | 40657863 | Exon |
| chr12 | 40948560 | 40949116 | Intron |
| chr12 | 40960555 | 40960773 | Intron |
| chr12 | 41310833 | 41312210 | Intron |
| chr12 | 41312299 | 41312502 | Exon |
| chr12 | 41614383 | 41614879 | Intron |
| chr12 | 41615193 | 41616177 | Intron |
| chr12 | 41625128 | 41625900 | Intron |

| | | | |
|---|---|---|---|
| chr12 | 41634437 | 41636566 | Intron |
| chr12 | 41806288 | 41808006 | Intron |
| chr12 | 41808384 | 41808592 | Intron |
| chr12 | 41810686 | 41812745 | Intron |
| chr12 | 41812786 | 41813353 | Intron |
| chr12 | 41816143 | 41816414 | Intron |
| chr12 | 41816875 | 41817273 | Intron |
| chr12 | 43050221 | 43051347 | Distal Intergenic |
| chr12 | 46148322 | 46148638 | Intron |
| chr12 | 46160905 | 46161286 | Intron |
| chr12 | 46161349 | 46161947 | Intron |
| chr12 | 46162226 | 46162614 | Intron |
| chr12 | 46641486 | 46641800 | Intron |
| chr12 | 49055415 | 49055776 | Intron |
| chr12 | 49120877 | 49121339 | Promoter |
| chr12 | 50146988 | 50147333 | Intron |
| chr12 | 50577924 | 50578225 | Intron |
| chr12 | 50613402 | 50614446 | Intron |
| chr12 | 51070377 | 51070588 | Intron |
| chr12 | 51070809 | 51071008 | Intron |
| chr12 | 52384722 | 52385208 | Intron |
| chr12 | 52448820 | 52449877 | Promoter |
| chr12 | 52478555 | 52479092 | Intron |
| chr12 | 52791419 | 52792689 | Intron |
| chr12 | 52792982 | 52793227 | Intron |
| chr12 | 52793308 | 52793572 | Intron |
| chr12 | 52793615 | 52793833 | Exon |
| chr12 | 52793913 | 52794148 | Exon |
| chr12 | 52794289 | 52796577 | Exon |
| chr12 | 52804777 | 52805228 | Distal Intergenic |
| chr12 | 52822574 | 52823838 | Intron |
| chr12 | 52834948 | 52835156 | Distal Intergenic |
| chr12 | 52835210 | 52837382 | Distal Intergenic |
| chr12 | 52929197 | 52930338 | Distal Intergenic |
| chr12 | 53604149 | 53604554 | 3' UTR |
| chr12 | 53605605 | 53606253 | Exon |
| chr12 | 53630290 | 53631268 | Distal Intergenic |
| chr12 | 53705036 | 53705235 | Intron |
| chr12 | 53705275 | 53705509 | Intron |
| chr12 | 53705836 | 53706044 | Intron |
| chr12 | 53726388 | 53726996 | Intron |
| chr12 | 53736768 | 53737004 | Distal Intergenic |
| chr12 | 53737172 | 53737775 | Distal Intergenic |
| chr12 | 53793198 | 53793748 | Intron |

| | | | |
|---|---|---|---|
| chr12 | 53801458 | 53801657 | Intron |
| chr12 | 53801714 | 53802564 | Intron |
| chr12 | 53802692 | 53803895 | Exon |
| chr12 | 53882423 | 53882626 | Intron |
| chr12 | 53883116 | 53884984 | Intron |
| chr12 | 53935323 | 53937735 | 3' UTR |
| chr12 | 53937789 | 53938026 | Intron |
| chr12 | 54051650 | 54051962 | Distal Intergenic |
| chr12 | 54247946 | 54248242 | Distal Intergenic |
| chr12 | 54989490 | 54990002 | Distal Intergenic |
| chr12 | 55049649 | 55050582 | Distal Intergenic |
| chr12 | 55288675 | 55289709 | Distal Intergenic |
| chr12 | 55292855 | 55293101 | Distal Intergenic |
| chr12 | 55293147 | 55293736 | Distal Intergenic |
| chr12 | 55301728 | 55302222 | Distal Intergenic |
| chr12 | 55302452 | 55302876 | Distal Intergenic |
| chr12 | 55342840 | 55343130 | 3' UTR |
| chr12 | 55387601 | 55388980 | Distal Intergenic |
| chr12 | 55699113 | 55699406 | Distal Intergenic |
| chr12 | 55720243 | 55722592 | Distal Intergenic |
| chr12 | 55741116 | 55742101 | Distal Intergenic |
| chr12 | 55796281 | 55797794 | Distal Intergenic |
| chr12 | 55811974 | 55812188 | Distal Intergenic |
| chr12 | 55896401 | 55897387 | Distal Intergenic |
| chr12 | 55963066 | 55963418 | Distal Intergenic |
| chr12 | 56041730 | 56042483 | Distal Intergenic |
| chr12 | 56086666 | 56087545 | Exon |
| chr12 | 56087637 | 56087860 | Exon |
| chr12 | 56096285 | 56096552 | Promoter |
| chr12 | 56096934 | 56097258 | Promoter |
| chr12 | 56108809 | 56109033 | Promoter |
| chr12 | 56181503 | 56181711 | Intron |
| chr12 | 56256350 | 56256734 | Distal Intergenic |
| chr12 | 56331727 | 56332028 | 5' UTR |
| chr12 | 56403785 | 56404162 | Distal Intergenic |
| chr12 | 56499251 | 56500217 | Intron |
| chr12 | 56500758 | 56501045 | Exon |
| chr12 | 56563523 | 56564343 | Exon |
| chr12 | 56564968 | 56565745 | Exon |
| chr12 | 56565976 | 56568630 | Exon |
| chr12 | 56572660 | 56572948 | Exon |
| chr12 | 56579678 | 56579916 | Intron |
| chr12 | 56590022 | 56590309 | Distal Intergenic |
| chr12 | 56591650 | 56591877 | Distal Intergenic |

| | | | |
|---|---|---|---|
| chr12 | 56592045 | 56592364 | Distal Intergenic |
| chr12 | 57419502 | 57420557 | Downstream |
| chr12 | 57426613 | 57427756 | Intron |
| chr12 | 57458657 | 57459302 | Intron |
| chr12 | 57880205 | 57881569 | Promoter |
| chr12 | 57895569 | 57896294 | Intron |
| chr12 | 57904979 | 57905894 | Promoter |
| chr12 | 60112272 | 60112563 | Intron |
| chr12 | 60112599 | 60112814 | Intron |
| chr12 | 60113178 | 60114347 | Intron |
| chr12 | 60161856 | 60162428 | 5' UTR |
| chr12 | 60162479 | 60162678 | Intron |
| chr12 | 60168077 | 60168307 | Intron |
| chr12 | 60177422 | 60179576 | 3' UTR |
| chr12 | 66066294 | 66066924 | Distal Intergenic |
| chr12 | 66067177 | 66067552 | Distal Intergenic |
| chr12 | 66071609 | 66071981 | Distal Intergenic |
| chr12 | 66076915 | 66077611 | Distal Intergenic |
| chr12 | 66137502 | 66137758 | Distal Intergenic |
| chr12 | 66138038 | 66138516 | Distal Intergenic |
| chr12 | 66174735 | 66174972 | Intron |
| chr12 | 66273039 | 66273440 | Intron |
| chr12 | 70259489 | 70260155 | Distal Intergenic |
| chr12 | 70336565 | 70336940 | Intron |
| chr12 | 70340699 | 70341063 | Intron |
| chr12 | 70462581 | 70463418 | Distal Intergenic |
| chr12 | 70463496 | 70463790 | Distal Intergenic |
| chr12 | 70464954 | 70465231 | Distal Intergenic |
| chr12 | 70472485 | 70472874 | Distal Intergenic |
| chr12 | 70473107 | 70473404 | Distal Intergenic |
| chr12 | 70473879 | 70474925 | Distal Intergenic |
| chr12 | 70523584 | 70524488 | Distal Intergenic |
| chr12 | 70537695 | 70538714 | Distal Intergenic |
| chr12 | 70547494 | 70547892 | Distal Intergenic |
| chr12 | 70548186 | 70548748 | Distal Intergenic |
| chr12 | 70548863 | 70549095 | Distal Intergenic |
| chr12 | 70603162 | 70603514 | Distal Intergenic |
| chr12 | 70603558 | 70603757 | Distal Intergenic |
| chr12 | 70638096 | 70640493 | Promoter |
| chr12 | 70844764 | 70845833 | Distal Intergenic |
| chr12 | 70925656 | 70926772 | Exon |
| chr12 | 70935176 | 70935788 | Intron |
| chr12 | 70979794 | 70981547 | Exon |
| chr12 | 70981619 | 70982132 | Intron |

| | | | |
|---|---|---|---|
| chr12 | 70982197 | 70982481 | Intron |
| chr12 | 70982580 | 70983261 | Intron |
| chr12 | 70985912 | 70986712 | 3' UTR |
| chr12 | 70989733 | 70989977 | Exon |
| chr12 | 70992868 | 70993815 | Intron |
| chr12 | 71087459 | 71087902 | Intron |
| chr12 | 71098457 | 71100849 | Intron |
| chr12 | 71645485 | 71646637 | Intron |
| chr12 | 71655116 | 71657631 | Intron |
| chr12 | 71657669 | 71657941 | Intron |
| chr12 | 71678673 | 71679534 | Intron |
| chr12 | 71690477 | 71690704 | Intron |
| chr12 | 71691174 | 71692176 | Intron |
| chr12 | 71701783 | 71702136 | Intron |
| chr12 | 71717388 | 71717780 | Intron |
| chr12 | 71724814 | 71727665 | Intron |
| chr12 | 71727763 | 71730907 | Intron |
| chr12 | 71748080 | 71752217 | Intron |
| chr12 | 71752287 | 71753254 | Intron |
| chr12 | 71754171 | 71754414 | Intron |
| chr12 | 71760396 | 71762105 | Intron |
| chr12 | 71771319 | 71775506 | Intron |
| chr12 | 71795775 | 71797265 | Intron |
| chr12 | 71806791 | 71809356 | Intron |
| chr12 | 71812692 | 71814299 | Intron |
| chr12 | 71821787 | 71822381 | Intron |
| chr12 | 71840415 | 71841806 | Intron |
| chr12 | 71925738 | 71926012 | Intron |
| chr12 | 71926054 | 71926253 | Intron |
| chr12 | 71926913 | 71927636 | Intron |
| chr12 | 71931173 | 71931402 | Intron |
| chr12 | 71931578 | 71932244 | Intron |
| chr12 | 71936032 | 71936391 | Intron |
| chr12 | 71991795 | 71992833 | Distal Intergenic |
| chr12 | 72001878 | 72002115 | Downstream |
| chr12 | 72008121 | 72009101 | Exon |
| chr12 | 72009143 | 72009579 | Intron |
| chr12 | 72009891 | 72010091 | Intron |
| chr12 | 72010925 | 72011228 | Intron |
| chr12 | 72038557 | 72038869 | Exon |
| chr12 | 72068959 | 72069449 | Intron |
| chr12 | 72069520 | 72069906 | Intron |
| chr12 | 72075959 | 72076461 | Distal Intergenic |
| chr12 | 72971727 | 72971996 | Intron |

| | | | |
|---|---|---|---|
| chr12 | 72985981 | 72986306 | Intron |
| chr12 | 72988567 | 72991210 | Intron |
| chr12 | 73084025 | 73084360 | Distal Intergenic |
| chr12 | 73084648 | 73084998 | Distal Intergenic |
| chr12 | 73103778 | 73104524 | Distal Intergenic |
| chr12 | 73639517 | 73639921 | Distal Intergenic |
| chr12 | 73845844 | 73846051 | Distal Intergenic |
| chr12 | 73861449 | 73862013 | Distal Intergenic |
| chr12 | 74248002 | 74248478 | Distal Intergenic |
| chr12 | 74248559 | 74248788 | Distal Intergenic |
| chr12 | 74329980 | 74331000 | Distal Intergenic |
| chr12 | 74331074 | 74331291 | Distal Intergenic |
| chr12 | 74331365 | 74331644 | Distal Intergenic |
| chr12 | 74395533 | 74397081 | Distal Intergenic |
| chr12 | 74717117 | 74718996 | Distal Intergenic |
| chr12 | 74819171 | 74820007 | Distal Intergenic |
| chr12 | 74826400 | 74827193 | Distal Intergenic |
| chr12 | 74827353 | 74827642 | Distal Intergenic |
| chr12 | 74840280 | 74840500 | Distal Intergenic |
| chr12 | 74840554 | 74841210 | Distal Intergenic |
| chr12 | 74847261 | 74847460 | Distal Intergenic |
| chr12 | 74855066 | 74856733 | Distal Intergenic |
| chr12 | 74856847 | 74857920 | Distal Intergenic |
| chr12 | 74865778 | 74866052 | Distal Intergenic |
| chr12 | 74879345 | 74880990 | Distal Intergenic |
| chr12 | 74891290 | 74892395 | Distal Intergenic |
| chr12 | 74900234 | 74901169 | Distal Intergenic |
| chr12 | 74907812 | 74908011 | Distal Intergenic |
| chr12 | 74908239 | 74909735 | Distal Intergenic |
| chr12 | 74911802 | 74912004 | Distal Intergenic |
| chr12 | 74915591 | 74916635 | Distal Intergenic |
| chr12 | 74939254 | 74939597 | Distal Intergenic |
| chr12 | 74943422 | 74943621 | Distal Intergenic |
| chr12 | 74981326 | 74981791 | Distal Intergenic |
| chr12 | 74995890 | 74997832 | Distal Intergenic |
| chr12 | 74997893 | 74998191 | Distal Intergenic |
| chr12 | 75008311 | 75008663 | Distal Intergenic |
| chr12 | 75009007 | 75009276 | Distal Intergenic |
| chr12 | 75009706 | 75011287 | Distal Intergenic |
| chr12 | 75011476 | 75012158 | Distal Intergenic |
| chr12 | 75194016 | 75194698 | Distal Intergenic |
| chr12 | 75194846 | 75196201 | Distal Intergenic |
| chr12 | 75625083 | 75625697 | Distal Intergenic |
| chr12 | 75925522 | 75926420 | Distal Intergenic |

| | | | |
|---|---|---|---|
| chr12 | 75926604 | 75926817 | Distal Intergenic |
| chr12 | 76629758 | 76630387 | Distal Intergenic |
| chr12 | 76644876 | 76647867 | Distal Intergenic |
| chr12 | 76655952 | 76656631 | Distal Intergenic |
| chr12 | 76656669 | 76657025 | Distal Intergenic |
| chr12 | 76658791 | 76659655 | Distal Intergenic |
| chr12 | 76664236 | 76665435 | Distal Intergenic |
| chr12 | 76665531 | 76665809 | Distal Intergenic |
| chr12 | 76697475 | 76697770 | Distal Intergenic |
| chr12 | 76770748 | 76773201 | Exon |
| chr12 | 76781259 | 76782778 | Intron |
| chr12 | 76803131 | 76804598 | Exon |
| chr12 | 76927490 | 76929491 | 5' UTR |
| chr12 | 76967839 | 76968426 | Distal Intergenic |
| chr12 | 76980812 | 76981067 | Distal Intergenic |
| chr12 | 77115619 | 77115818 | Distal Intergenic |
| chr12 | 77116551 | 77116906 | Distal Intergenic |
| chr12 | 77143623 | 77144303 | Distal Intergenic |
| chr12 | 77187108 | 77188193 | Intron |
| chr12 | 77356860 | 77357173 | Distal Intergenic |
| chr12 | 77471185 | 77471653 | Distal Intergenic |
| chr12 | 77559834 | 77560205 | Distal Intergenic |
| chr12 | 77560247 | 77560631 | Distal Intergenic |
| chr12 | 77580862 | 77581414 | Distal Intergenic |
| chr12 | 77604171 | 77604838 | Distal Intergenic |
| chr12 | 77631934 | 77632326 | Distal Intergenic |
| chr12 | 77632711 | 77632984 | Distal Intergenic |
| chr12 | 77633227 | 77637208 | Distal Intergenic |
| chr12 | 77637280 | 77638869 | Distal Intergenic |
| chr12 | 77641421 | 77642501 | Distal Intergenic |
| chr12 | 77648631 | 77648844 | Distal Intergenic |
| chr12 | 77648946 | 77649178 | Distal Intergenic |
| chr12 | 77652984 | 77653951 | Distal Intergenic |
| chr12 | 77656001 | 77656403 | Distal Intergenic |
| chr12 | 77657168 | 77657382 | Distal Intergenic |
| chr12 | 77658398 | 77659955 | Distal Intergenic |
| chr12 | 77694744 | 77695177 | Distal Intergenic |
| chr12 | 77695229 | 77695467 | Distal Intergenic |
| chr12 | 77895310 | 77895771 | Distal Intergenic |
| chr12 | 77895831 | 77896980 | Distal Intergenic |
| chr12 | 77910605 | 77910865 | Distal Intergenic |
| chr12 | 77919103 | 77919460 | Distal Intergenic |
| chr12 | 78120889 | 78121145 | Distal Intergenic |
| chr12 | 78186213 | 78186424 | Distal Intergenic |

| | | | |
|---|---|---|---|
| chr12 | 79071265 | 79071503 | Distal Intergenic |
| chr12 | 79741375 | 79741848 | Intron |
| chr12 | 79742054 | 79742297 | Intron |
| chr12 | 79868959 | 79869296 | Distal Intergenic |
| chr12 | 79869346 | 79869904 | Distal Intergenic |
| chr12 | 79870136 | 79870563 | Distal Intergenic |
| chr12 | 79942532 | 79942749 | Distal Intergenic |
| chr12 | 79943207 | 79944194 | Distal Intergenic |
| chr12 | 79961927 | 79962470 | Distal Intergenic |
| chr12 | 79962736 | 79963169 | Distal Intergenic |
| chr12 | 80008886 | 80009128 | Intron |
| chr12 | 80011391 | 80013015 | Intron |
| chr12 | 80053003 | 80053884 | Intron |
| chr12 | 80148588 | 80148800 | Distal Intergenic |
| chr12 | 80269007 | 80270098 | Intron |
| chr12 | 80272907 | 80273114 | Intron |
| chr12 | 80291702 | 80291910 | Intron |
| chr12 | 80292476 | 80292853 | Intron |
| chr12 | 80330761 | 80331170 | Distal Intergenic |
| chr12 | 80348695 | 80350231 | Distal Intergenic |
| chr12 | 80355191 | 80356292 | Distal Intergenic |
| chr12 | 80431499 | 80431783 | Distal Intergenic |
| chr12 | 80448865 | 80450307 | Distal Intergenic |
| chr12 | 80452489 | 80453800 | Distal Intergenic |
| chr12 | 80453840 | 80454272 | Distal Intergenic |
| chr12 | 80483973 | 80485307 | Distal Intergenic |
| chr12 | 80521464 | 80522507 | Distal Intergenic |
| chr12 | 80534028 | 80536744 | Distal Intergenic |
| chr12 | 80566692 | 80567938 | Distal Intergenic |
| chr12 | 80572152 | 80573673 | Distal Intergenic |
| chr12 | 80589820 | 80590308 | Distal Intergenic |
| chr12 | 80596204 | 80596508 | Distal Intergenic |
| chr12 | 80599007 | 80600038 | Distal Intergenic |
| chr12 | 80602499 | 80603694 | Promoter |
| chr12 | 80603796 | 80606101 | Promoter |
| chr12 | 80606672 | 80606929 | Intron |
| chr12 | 80607035 | 80609292 | Intron |
| chr12 | 80615288 | 80616329 | Exon |
| chr12 | 80624565 | 80624978 | Intron |
| chr12 | 80625262 | 80627823 | Exon |
| chr12 | 80628167 | 80628604 | Intron |
| chr12 | 80646517 | 80647409 | Exon |
| chr12 | 80652252 | 80654499 | Intron |
| chr12 | 80654566 | 80654850 | Intron |

| | | | |
|---|---|---|---|
| chr12 | 80656041 | 80656688 | Intron |
| chr12 | 80660330 | 80660730 | Exon |
| chr12 | 80660819 | 80661031 | Intron |
| chr12 | 80684218 | 80685102 | Intron |
| chr12 | 80698763 | 80699942 | Exon |
| chr12 | 80699980 | 80700856 | Intron |
| chr12 | 80701021 | 80702999 | Intron |
| chr12 | 80703340 | 80703623 | Intron |
| chr12 | 80711717 | 80713140 | Exon |
| chr12 | 80749812 | 80751014 | Promoter |
| chr12 | 80965397 | 80966461 | Intron |
| chr12 | 81091304 | 81092373 | Distal Intergenic |
| chr12 | 81135874 | 81136639 | Distal Intergenic |
| chr12 | 81198781 | 81199258 | Intron |
| chr12 | 81199400 | 81199600 | Intron |
| chr12 | 81205026 | 81205307 | Exon |
| chr12 | 81212963 | 81213524 | Intron |
| chr12 | 81214630 | 81214851 | Intron |
| chr12 | 81215169 | 81215796 | Intron |
| chr12 | 81216069 | 81216784 | Intron |
| chr12 | 81216828 | 81217213 | Intron |
| chr12 | 81217331 | 81217730 | Intron |
| chr12 | 81321003 | 81321922 | Intron |
| chr12 | 81324134 | 81325225 | Intron |
| chr12 | 81325322 | 81325589 | Intron |
| chr12 | 81325652 | 81325989 | Intron |
| chr12 | 81329066 | 81337689 | Promoter |
| chr12 | 81341303 | 81341786 | Distal Intergenic |
| chr12 | 81344817 | 81346489 | Distal Intergenic |
| chr12 | 81352149 | 81352856 | Distal Intergenic |
| chr12 | 81356736 | 81357115 | Distal Intergenic |
| chr12 | 81357459 | 81359342 | Distal Intergenic |
| chr12 | 81360109 | 81360833 | Distal Intergenic |
| chr12 | 81361533 | 81362188 | Distal Intergenic |
| chr12 | 81444971 | 81448162 | Distal Intergenic |
| chr12 | 81459523 | 81459922 | Distal Intergenic |
| chr12 | 81466731 | 81467701 | Distal Intergenic |
| chr12 | 81468960 | 81469612 | Distal Intergenic |
| chr12 | 81475006 | 81476464 | Intron |
| chr12 | 81535875 | 81536668 | Intron |
| chr12 | 81744667 | 81745289 | Intron |
| chr12 | 81891599 | 81891827 | Intron |
| chr12 | 81892009 | 81892645 | Intron |
| chr12 | 81892718 | 81893539 | Intron |

| | | | |
|---|---|---|---|
| chr12 | 81944532 | 81944849 | Intron |
| chr12 | 81945000 | 81946031 | Intron |
| chr12 | 81954251 | 81955603 | Intron |
| chr12 | 82037970 | 82039228 | Intron |
| chr12 | 82046509 | 82046715 | Intron |
| chr12 | 82257294 | 82257596 | Distal Intergenic |
| chr12 | 82257683 | 82258117 | Distal Intergenic |
| chr12 | 82302338 | 82302537 | Distal Intergenic |
| chr12 | 82336643 | 82336884 | Distal Intergenic |
| chr12 | 82585699 | 82586107 | Distal Intergenic |
| chr12 | 82633679 | 82633887 | Distal Intergenic |
| chr12 | 82798607 | 82800367 | Intron |
| chr12 | 82824152 | 82824918 | Exon |
| chr12 | 82824960 | 82825160 | Intron |
| chr12 | 82848744 | 82851747 | Exon |
| chr12 | 82863436 | 82863984 | Intron |
| chr12 | 82880619 | 82881084 | Distal Intergenic |
| chr12 | 82881246 | 82881519 | Distal Intergenic |
| chr12 | 82882334 | 82882868 | Distal Intergenic |
| chr12 | 82941844 | 82944084 | Distal Intergenic |
| chr12 | 82944163 | 82944983 | Distal Intergenic |
| chr12 | 83072652 | 83073164 | Distal Intergenic |
| chr12 | 83255257 | 83256335 | Intron |
| chr12 | 84866621 | 84867943 | Distal Intergenic |
| chr12 | 84972829 | 84973558 | Distal Intergenic |
| chr12 | 84973663 | 84974575 | Distal Intergenic |
| chr12 | 85028791 | 85029367 | Distal Intergenic |
| chr12 | 85029434 | 85030292 | Distal Intergenic |
| chr12 | 85062443 | 85062691 | Distal Intergenic |
| chr12 | 85175989 | 85176483 | Distal Intergenic |
| chr12 | 85215453 | 85215810 | Distal Intergenic |
| chr12 | 85515757 | 85516012 | Intron |
| chr12 | 85516507 | 85517237 | Intron |
| chr12 | 85517641 | 85518247 | Exon |
| chr12 | 85518787 | 85519103 | Intron |
| chr12 | 85519274 | 85519591 | Intron |
| chr12 | 85521580 | 85521878 | Exon |
| chr12 | 85526473 | 85527158 | Intron |
| chr12 | 85533520 | 85533722 | Intron |
| chr12 | 85535456 | 85535868 | Intron |
| chr12 | 85542679 | 85545232 | Intron |
| chr12 | 85557901 | 85559619 | Intron |
| chr12 | 85559854 | 85560465 | Intron |
| chr12 | 85563281 | 85563795 | Intron |

| | | | |
|---|---|---|---|
| chr12 | 85564086 | 85564323 | Intron |
| chr12 | 85573657 | 85575270 | Intron |
| chr12 | 85576528 | 85577927 | Intron |
| chr12 | 85725523 | 85726469 | Distal Intergenic |
| chr12 | 85736054 | 85736361 | Distal Intergenic |
| chr12 | 85736529 | 85736738 | Distal Intergenic |
| chr12 | 85955533 | 85956266 | Distal Intergenic |
| chr12 | 85972853 | 85973215 | Distal Intergenic |
| chr12 | 86108478 | 86108752 | Distal Intergenic |
| chr12 | 86187660 | 86187913 | Distal Intergenic |
| chr12 | 86208348 | 86208572 | Intron |
| chr12 | 86391939 | 86392250 | Intron |
| chr12 | 86574284 | 86574494 | Intron |
| chr12 | 86679278 | 86679839 | Intron |
| chr12 | 86702387 | 86702643 | Intron |
| chr12 | 86703310 | 86704068 | Intron |
| chr12 | 86714600 | 86716374 | Intron |
| chr12 | 86758761 | 86759477 | Intron |
| chr12 | 86759842 | 86760055 | Intron |
| chr12 | 86761705 | 86761930 | Intron |
| chr12 | 86792529 | 86793504 | Intron |
| chr12 | 86910538 | 86910737 | Intron |
| chr12 | 86910842 | 86911411 | Intron |
| chr12 | 87165122 | 87165325 | Intron |
| chr12 | 87165692 | 87165909 | Intron |
| chr12 | 87210509 | 87210742 | Intron |
| chr12 | 87210816 | 87211188 | Intron |
| chr12 | 87300109 | 87301866 | Distal Intergenic |
| chr12 | 87302270 | 87302480 | Distal Intergenic |
| chr12 | 87302530 | 87302762 | Distal Intergenic |
| chr12 | 87310360 | 87311578 | Distal Intergenic |
| chr12 | 87312556 | 87313806 | Distal Intergenic |
| chr12 | 87314209 | 87314421 | Distal Intergenic |
| chr12 | 87341165 | 87341455 | Distal Intergenic |
| chr12 | 87341574 | 87341831 | Distal Intergenic |
| chr12 | 87352201 | 87353518 | Distal Intergenic |
| chr12 | 87423759 | 87424517 | Distal Intergenic |
| chr12 | 87424585 | 87427254 | Distal Intergenic |
| chr12 | 87483016 | 87483908 | Distal Intergenic |
| chr12 | 87582302 | 87582779 | Distal Intergenic |
| chr12 | 88116548 | 88117294 | Distal Intergenic |
| chr12 | 88128055 | 88128636 | Distal Intergenic |
| chr12 | 88136375 | 88136936 | Distal Intergenic |
| chr12 | 88137307 | 88137562 | Distal Intergenic |

| | | | |
|---|---|---|---|
| chr12 | 88139848 | 88140917 | Distal Intergenic |
| chr12 | 88140986 | 88143251 | Distal Intergenic |
| chr12 | 88143331 | 88143627 | Distal Intergenic |
| chr12 | 88164780 | 88166040 | Distal Intergenic |
| chr12 | 88166218 | 88166685 | Distal Intergenic |
| chr12 | 88172811 | 88176041 | Downstream |
| chr12 | 88177411 | 88177799 | Promoter |
| chr12 | 88208273 | 88209261 | Distal Intergenic |
| chr12 | 88213297 | 88213566 | Distal Intergenic |
| chr12 | 88224941 | 88225222 | Distal Intergenic |
| chr12 | 88230053 | 88230336 | Distal Intergenic |
| chr12 | 88488710 | 88488984 | Intron |
| chr12 | 88607368 | 88607595 | Distal Intergenic |
| chr12 | 88607928 | 88608199 | Distal Intergenic |
| chr12 | 88775353 | 88776928 | Distal Intergenic |
| chr12 | 88783889 | 88784089 | Distal Intergenic |
| chr12 | 88784445 | 88784936 | Distal Intergenic |
| chr12 | 88812456 | 88812662 | Distal Intergenic |
| chr12 | 88812808 | 88813134 | Distal Intergenic |
| chr12 | 89108904 | 89109561 | Distal Intergenic |
| chr12 | 89453124 | 89453535 | Distal Intergenic |
| chr12 | 89453595 | 89455215 | Distal Intergenic |
| chr12 | 89515962 | 89516578 | Distal Intergenic |
| chr12 | 92020330 | 92020548 | Distal Intergenic |
| chr12 | 92143322 | 92143561 | Distal Intergenic |
| chr12 | 92159460 | 92159896 | Distal Intergenic |
| chr12 | 92159977 | 92160176 | Distal Intergenic |
| chr12 | 92826703 | 92827414 | Intron |
| chr12 | 92828092 | 92828413 | Intron |
| chr12 | 92924260 | 92924554 | Distal Intergenic |
| chr12 | 92924686 | 92925191 | Distal Intergenic |
| chr12 | 92931529 | 92932474 | Distal Intergenic |
| chr12 | 92984036 | 92984765 | Distal Intergenic |
| chr12 | 92986329 | 92986748 | Distal Intergenic |
| chr12 | 93013161 | 93013974 | Distal Intergenic |
| chr12 | 96614516 | 96615889 | Intron |
| chr12 | 96685547 | 96686209 | Intron |
| chr12 | 96707965 | 96708298 | Intron |
| chr12 | 96729251 | 96731066 | Intron |
| chr12 | 96776791 | 96777118 | Intron |
| chr12 | 96798745 | 96800201 | Distal Intergenic |
| chr12 | 96801762 | 96802123 | Distal Intergenic |
| chr12 | 96837244 | 96839504 | Distal Intergenic |
| chr12 | 96930810 | 96931015 | Intron |

| | | | |
|---|---|---|---|
| chr12 | 96931231 | 96933241 | Exon |
| chr12 | 96933535 | 96933789 | Intron |
| chr12 | 99520668 | 99522279 | Intron |
| chr12 | 99951791 | 99952126 | Intron |
| chr12 | 99975150 | 99975433 | Intron |
| chr12 | 100232657 | 100233193 | Intron |
| chr12 | 100233229 | 100233465 | Intron |
| chr12 | 100233638 | 100233967 | Intron |
| chr12 | 100234069 | 100235088 | Intron |
| chr12 | 100262789 | 100263005 | Intron |
| chr12 | 100302321 | 100303011 | Intron |
| chr12 | 100568806 | 100569005 | Distal Intergenic |
| chr12 | 100573885 | 100576200 | Distal Intergenic |
| chr12 | 100594554 | 100595963 | Promoter |
| chr12 | 100599235 | 100599536 | 5' UTR |
| chr12 | 100599665 | 100599936 | Intron |
| chr12 | 100615059 | 100616309 | Exon |
| chr12 | 101217777 | 101218725 | Intron |
| chr12 | 101448988 | 101449330 | Intron |
| chr12 | 101518357 | 101518649 | Intron |
| chr12 | 101759784 | 101760096 | Intron |
| chr12 | 101943117 | 101943387 | Distal Intergenic |
| chr12 | 101994898 | 101995097 | Intron |
| chr12 | 102016551 | 102016911 | Intron |
| chr12 | 102017322 | 102017803 | Intron |
| chr12 | 102219191 | 102220400 | Intron |
| chr12 | 102240837 | 102241739 | Distal Intergenic |
| chr12 | 102321268 | 102321653 | Distal Intergenic |
| chr12 | 103151156 | 103151594 | Distal Intergenic |
| chr12 | 103161416 | 103162419 | Distal Intergenic |
| chr12 | 103163441 | 103163789 | Distal Intergenic |
| chr12 | 103213515 | 103214995 | Exon |
| chr12 | 103963621 | 103964009 | Distal Intergenic |
| chr12 | 103964214 | 103964645 | Distal Intergenic |
| chr12 | 103982004 | 103982635 | Promoter |
| chr12 | 104115447 | 104115860 | Intron |
| chr12 | 104519418 | 104519674 | Intron |
| chr12 | 104519832 | 104520366 | Exon |
| chr12 | 104618382 | 104618710 | Exon |
| chr12 | 104618787 | 104619244 | Intron |
| chr12 | 104803131 | 104803508 | Distal Intergenic |
| chr12 | 105241032 | 105241547 | Intron |
| chr12 | 105302771 | 105303659 | Exon |
| chr12 | 105361483 | 105361829 | Distal Intergenic |

| | | | |
|---|---|---|---|
| chr12 | 105362126 | 105362401 | Distal Intergenic |
| chr12 | 105362531 | 105363213 | Distal Intergenic |
| chr12 | 105433437 | 105433636 | Exon |
| chr12 | 105450215 | 105450416 | Intron |
| chr12 | 105485170 | 105485893 | Distal Intergenic |
| chr12 | 105485993 | 105486353 | Distal Intergenic |
| chr12 | 105500390 | 105501601 | Promoter |
| chr12 | 105509285 | 105509870 | 5' UTR |
| chr12 | 105654258 | 105654621 | Distal Intergenic |
| chr12 | 106237108 | 106237560 | Distal Intergenic |
| chr12 | 106811245 | 106813149 | Intron |
| chr12 | 107022963 | 107023176 | Intron |
| chr12 | 107081353 | 107082527 | Intron |
| chr12 | 107806588 | 107806908 | Intron |
| chr12 | 107806998 | 107807458 | Intron |
| chr12 | 108001604 | 108001817 | Intron |
| chr12 | 108001879 | 108002787 | Intron |
| chr12 | 108003140 | 108003392 | Intron |
| chr12 | 108117312 | 108117767 | Distal Intergenic |
| chr12 | 108118250 | 108118469 | Distal Intergenic |
| chr12 | 108855020 | 108855607 | Distal Intergenic |
| chr12 | 108883827 | 108884093 | Distal Intergenic |
| chr12 | 108990717 | 108990994 | Promoter |
| chr12 | 108992893 | 108994843 | Promoter |
| chr12 | 109045454 | 109046616 | Exon |
| chr12 | 109094278 | 109094792 | Intron |
| chr12 | 109095095 | 109095632 | 5' UTR |
| chr12 | 109189447 | 109189921 | Intron |
| chr12 | 109691954 | 109692594 | Exon |
| chr12 | 109917576 | 109917775 | Intron |
| chr12 | 109987053 | 109989409 | Downstream |
| chr12 | 110152166 | 110152691 | Promoter |
| chr12 | 110152745 | 110153114 | Promoter |
| chr12 | 110864902 | 110865245 | Distal Intergenic |
| chr12 | 111724020 | 111724312 | Intron |
| chr12 | 111749547 | 111749934 | Exon |
| chr12 | 111750203 | 111751371 | Intron |
| chr12 | 111751531 | 111751779 | Intron |
| chr12 | 111770852 | 111772356 | Exon |
| chr12 | 111790460 | 111792016 | Distal Intergenic |
| chr12 | 111793634 | 111793925 | Distal Intergenic |
| chr12 | 111794014 | 111794214 | Distal Intergenic |
| chr12 | 111794271 | 111794919 | Distal Intergenic |
| chr12 | 112036155 | 112036354 | Promoter |

| | | | |
|---|---|---|---|
| chr12 | 112424990 | 112425375 | Intron |
| chr12 | 112476539 | 112477088 | Exon |
| chr12 | 112587659 | 112587954 | Exon |
| chr12 | 112588065 | 112588303 | Intron |
| chr12 | 112591203 | 112591696 | 3' UTR |
| chr12 | 112709589 | 112710271 | Intron |
| chr12 | 112775748 | 112776109 | Intron |
| chr12 | 112790912 | 112792142 | Intron |
| chr12 | 112794811 | 112795236 | Intron |
| chr12 | 112798092 | 112798404 | Intron |
| chr12 | 112803686 | 112803889 | Intron |
| chr12 | 112806749 | 112807708 | Intron |
| chr12 | 112813409 | 112813608 | Intron |
| chr12 | 112936191 | 112937044 | Intron |
| chr12 | 112942820 | 112943888 | 3' UTR |
| chr12 | 112991277 | 112991916 | Distal Intergenic |
| chr12 | 112996088 | 112996813 | Distal Intergenic |
| chr12 | 112998308 | 112998936 | Distal Intergenic |
| chr12 | 113001149 | 113001441 | Distal Intergenic |
| chr12 | 113388168 | 113388455 | Intron |
| chr12 | 113420135 | 113420415 | Intron |
| chr12 | 113420754 | 113421030 | Intron |
| chr12 | 113421186 | 113421440 | Intron |
| chr12 | 113421612 | 113421861 | Intron |
| chr12 | 113422513 | 113422802 | Intron |
| chr12 | 113532741 | 113533250 | Exon |
| chr12 | 113533482 | 113533796 | Intron |
| chr12 | 113533962 | 113534266 | Intron |
| chr12 | 113578033 | 113578656 | Distal Intergenic |
| chr12 | 113816791 | 113817000 | Intron |
| chr12 | 113817317 | 113817724 | Intron |
| chr12 | 113817935 | 113818215 | Intron |
| chr12 | 113832725 | 113834516 | Intron |
| chr12 | 113854985 | 113855221 | Distal Intergenic |
| chr12 | 113912616 | 113913017 | Distal Intergenic |
| chr12 | 113913287 | 113913519 | Distal Intergenic |
| chr12 | 114032212 | 114032800 | Distal Intergenic |
| chr12 | 114032856 | 114034425 | Distal Intergenic |
| chr12 | 114035399 | 114035788 | Distal Intergenic |
| chr12 | 114036423 | 114036647 | Distal Intergenic |
| chr12 | 114060556 | 114062980 | Distal Intergenic |
| chr12 | 114066859 | 114067573 | Distal Intergenic |
| chr12 | 114089347 | 114090730 | Distal Intergenic |
| chr12 | 114114845 | 114115375 | Distal Intergenic |

| | | | |
|---|---|---|---|
| chr12 | 114116100 | 114116512 | Distal Intergenic |
| chr12 | 114116555 | 114116934 | Distal Intergenic |
| chr12 | 114118829 | 114119181 | Distal Intergenic |
| chr12 | 114119412 | 114119770 | Distal Intergenic |
| chr12 | 114165789 | 114166070 | Distal Intergenic |
| chr12 | 114178524 | 114180084 | Distal Intergenic |
| chr12 | 114261191 | 114261390 | Intron |
| chr12 | 114415330 | 114415559 | Distal Intergenic |
| chr12 | 114416034 | 114416267 | Distal Intergenic |
| chr12 | 117518933 | 117519300 | Intron |
| chr12 | 117531904 | 117532833 | Intron |
| chr12 | 117712615 | 117713747 | Intron |
| chr12 | 118669007 | 118669990 | Intron |
| chr12 | 118680614 | 118681211 | Intron |
| chr12 | 118681870 | 118682100 | Intron |
| chr12 | 118687098 | 118687618 | Intron |
| chr12 | 118771627 | 118771851 | Intron |
| chr12 | 120052834 | 120053245 | Intron |
| chr12 | 120144933 | 120145188 | Intron |
| chr12 | 120145229 | 120145603 | Intron |
| chr12 | 120145702 | 120145959 | Intron |
| chr12 | 120664455 | 120664731 | Promoter |
| chr12 | 120679547 | 120679753 | Intron |
| chr13 | 3094316 | 3094515 | Distal Intergenic |
| chr13 | 3537412 | 3538395 | Distal Intergenic |
| chr13 | 3609200 | 3609768 | Distal Intergenic |
| chr13 | 3745611 | 3746237 | Distal Intergenic |
| chr13 | 3778294 | 3778570 | Distal Intergenic |
| chr13 | 3817602 | 3818371 | Distal Intergenic |
| chr13 | 3820876 | 3821766 | Distal Intergenic |
| chr13 | 3830953 | 3831232 | Distal Intergenic |
| chr13 | 3831443 | 3831692 | Distal Intergenic |
| chr13 | 3861734 | 3862586 | Distal Intergenic |
| chr13 | 3864726 | 3865330 | Distal Intergenic |
| chr13 | 3866216 | 3866714 | Distal Intergenic |
| chr13 | 3876038 | 3877611 | Distal Intergenic |
| chr13 | 3886101 | 3886747 | Distal Intergenic |
| chr13 | 3891033 | 3891258 | Distal Intergenic |
| chr13 | 3891529 | 3891762 | Distal Intergenic |
| chr13 | 3891808 | 3892765 | Distal Intergenic |
| chr13 | 5301638 | 5302116 | Distal Intergenic |
| chr13 | 5314188 | 5314655 | Distal Intergenic |
| chr13 | 5423275 | 5423659 | Distal Intergenic |
| chr13 | 5443417 | 5445148 | Distal Intergenic |

| | | | |
|---|---|---|---|
| chr13 | 5723129 | 5723410 | Distal Intergenic |
| chr13 | 5723453 | 5723697 | Distal Intergenic |
| chr13 | 5802476 | 5804008 | Distal Intergenic |
| chr13 | 5824733 | 5826141 | Distal Intergenic |
| chr13 | 5836288 | 5837082 | Distal Intergenic |
| chr13 | 5839755 | 5841460 | Distal Intergenic |
| chr13 | 5843145 | 5845231 | Distal Intergenic |
| chr13 | 5858171 | 5859332 | Distal Intergenic |
| chr13 | 5859418 | 5860474 | Distal Intergenic |
| chr13 | 5860724 | 5863254 | Distal Intergenic |
| chr13 | 5863339 | 5864454 | Distal Intergenic |
| chr13 | 5864528 | 5866319 | Distal Intergenic |
| chr13 | 5868541 | 5870201 | Distal Intergenic |
| chr13 | 5870402 | 5871177 | Distal Intergenic |
| chr13 | 5885665 | 5886793 | Distal Intergenic |
| chr13 | 5891293 | 5893489 | Distal Intergenic |
| chr13 | 5912233 | 5912659 | Distal Intergenic |
| chr13 | 6164650 | 6165253 | Distal Intergenic |
| chr13 | 6190935 | 6191189 | Distal Intergenic |
| chr13 | 6191302 | 6193335 | Distal Intergenic |
| chr13 | 6203922 | 6204121 | Distal Intergenic |
| chr13 | 6612983 | 6614274 | Distal Intergenic |
| chr13 | 6623601 | 6624036 | Distal Intergenic |
| chr13 | 6628185 | 6628529 | Distal Intergenic |
| chr13 | 6630788 | 6631137 | Distal Intergenic |
| chr13 | 6646544 | 6646875 | Distal Intergenic |
| chr13 | 6658419 | 6658623 | Distal Intergenic |
| chr13 | 8828582 | 8828984 | Distal Intergenic |
| chr13 | 8870244 | 8870726 | Distal Intergenic |
| chr13 | 8883993 | 8885451 | Distal Intergenic |
| chr13 | 8885517 | 8885841 | Distal Intergenic |
| chr13 | 8885935 | 8886154 | Distal Intergenic |
| chr13 | 8886253 | 8886472 | Distal Intergenic |
| chr13 | 8994522 | 8995555 | Distal Intergenic |
| chr13 | 9093775 | 9094187 | Distal Intergenic |
| chr13 | 9700051 | 9700826 | Distal Intergenic |
| chr13 | 9748828 | 9749470 | Distal Intergenic |
| chr13 | 9762400 | 9763680 | Distal Intergenic |
| chr13 | 9772285 | 9772990 | Distal Intergenic |
| chr13 | 11641365 | 11642023 | Distal Intergenic |
| chr13 | 11642392 | 11642616 | Distal Intergenic |
| chr13 | 12384574 | 12385114 | Distal Intergenic |
| chr13 | 12385261 | 12385830 | Distal Intergenic |
| chr13 | 12385877 | 12386352 | Distal Intergenic |

| | | | |
|---|---|---|---|
| chr13 | 12552672 | 12553546 | Distal Intergenic |
| chr13 | 12577864 | 12578739 | Distal Intergenic |
| chr13 | 12599299 | 12599667 | Distal Intergenic |
| chr13 | 12612135 | 12612373 | Distal Intergenic |
| chr13 | 12612605 | 12612824 | Distal Intergenic |
| chr13 | 12795904 | 12796344 | Distal Intergenic |
| chr13 | 12798563 | 12798823 | Distal Intergenic |
| chr13 | 12892325 | 12892865 | Distal Intergenic |
| chr13 | 12892932 | 12893546 | Distal Intergenic |
| chr13 | 12894031 | 12895476 | Distal Intergenic |
| chr13 | 12900507 | 12902760 | Distal Intergenic |
| chr13 | 12904775 | 12907116 | Distal Intergenic |
| chr13 | 12910119 | 12911410 | Distal Intergenic |
| chr13 | 13097318 | 13098793 | Distal Intergenic |
| chr13 | 13099602 | 13100548 | Distal Intergenic |
| chr13 | 13106105 | 13106478 | Distal Intergenic |
| chr13 | 13106639 | 13107095 | Distal Intergenic |
| chr13 | 13116921 | 13118957 | Distal Intergenic |
| chr13 | 13122466 | 13123217 | Distal Intergenic |
| chr13 | 13166698 | 13167345 | Distal Intergenic |
| chr13 | 13168425 | 13168974 | Distal Intergenic |
| chr13 | 13255470 | 13256510 | Distal Intergenic |
| chr13 | 13260018 | 13262619 | Distal Intergenic |
| chr13 | 13265629 | 13266176 | Distal Intergenic |
| chr13 | 13272340 | 13272890 | Distal Intergenic |
| chr13 | 13273900 | 13274628 | Distal Intergenic |
| chr13 | 13533514 | 13533713 | Distal Intergenic |
| chr13 | 13665097 | 13665383 | Distal Intergenic |
| chr13 | 13840540 | 13841006 | Distal Intergenic |
| chr13 | 14060443 | 14061848 | Distal Intergenic |
| chr13 | 14126577 | 14126832 | Distal Intergenic |
| chr13 | 14131236 | 14131530 | Distal Intergenic |
| chr13 | 14131624 | 14132911 | Distal Intergenic |
| chr13 | 14152158 | 14152674 | Distal Intergenic |
| chr13 | 14154670 | 14155415 | Distal Intergenic |
| chr13 | 14155980 | 14156439 | Distal Intergenic |
| chr13 | 14705491 | 14706253 | Distal Intergenic |
| chr13 | 14722223 | 14722436 | Distal Intergenic |
| chr13 | 14722621 | 14722954 | Distal Intergenic |
| chr13 | 15263607 | 15263876 | Distal Intergenic |
| chr13 | 15275812 | 15276864 | Distal Intergenic |
| chr13 | 15513201 | 15514114 | Distal Intergenic |
| chr13 | 15518587 | 15519299 | Distal Intergenic |
| chr13 | 15602654 | 15602893 | Distal Intergenic |

| | | | |
|---|---|---|---|
| chr13 | 15603391 | 15604364 | Distal Intergenic |
| chr13 | 15604821 | 15606447 | Distal Intergenic |
| chr13 | 15629514 | 15630098 | Distal Intergenic |
| chr13 | 15752253 | 15752758 | Distal Intergenic |
| chr13 | 15895759 | 15895958 | Distal Intergenic |
| chr13 | 15896076 | 15896448 | Distal Intergenic |
| chr13 | 15898214 | 15899671 | Distal Intergenic |
| chr13 | 16557356 | 16558456 | Distal Intergenic |
| chr13 | 16578534 | 16579238 | Distal Intergenic |
| chr13 | 16589885 | 16590439 | Distal Intergenic |
| chr13 | 16641846 | 16642581 | Distal Intergenic |
| chr13 | 16733407 | 16734604 | Distal Intergenic |
| chr13 | 16749305 | 16749736 | Distal Intergenic |
| chr13 | 16750896 | 16752338 | Distal Intergenic |
| chr13 | 16761224 | 16762780 | Distal Intergenic |
| chr13 | 16827592 | 16828606 | Distal Intergenic |
| chr13 | 16829151 | 16832251 | Distal Intergenic |
| chr13 | 16852905 | 16853908 | Distal Intergenic |
| chr13 | 17549298 | 17550497 | Distal Intergenic |
| chr13 | 17584169 | 17587244 | Distal Intergenic |
| chr13 | 17603118 | 17604007 | Distal Intergenic |
| chr13 | 17695250 | 17697196 | Distal Intergenic |
| chr13 | 17698310 | 17698663 | Distal Intergenic |
| chr13 | 17728447 | 17728978 | Distal Intergenic |
| chr13 | 17781602 | 17782046 | Distal Intergenic |
| chr13 | 17785723 | 17786474 | Distal Intergenic |
| chr13 | 17786915 | 17787192 | Distal Intergenic |
| chr13 | 17787231 | 17787509 | Distal Intergenic |
| chr13 | 17787610 | 17788079 | Distal Intergenic |
| chr13 | 17797066 | 17797492 | Distal Intergenic |
| chr13 | 17895032 | 17895697 | Distal Intergenic |
| chr13 | 17896161 | 17896382 | Distal Intergenic |
| chr13 | 17992265 | 17993899 | Distal Intergenic |
| chr13 | 18035040 | 18035246 | Distal Intergenic |
| chr13 | 18035294 | 18035768 | Distal Intergenic |
| chr13 | 19486705 | 19487140 | Intron |
| chr13 | 21271253 | 21272035 | Distal Intergenic |
| chr13 | 21494437 | 21494648 | Distal Intergenic |
| chr13 | 21507601 | 21507930 | Distal Intergenic |
| chr13 | 21507993 | 21508207 | Distal Intergenic |
| chr13 | 21508698 | 21509933 | Distal Intergenic |
| chr13 | 21544840 | 21545292 | Downstream |
| chr13 | 21598235 | 21599037 | Intron |
| chr13 | 21599176 | 21600658 | Intron |

| | | | |
|---|---|---|---|
| chr13 | 21600856 | 21601056 | Intron |
| chr13 | 21807710 | 21808119 | Distal Intergenic |
| chr13 | 21808276 | 21808680 | Distal Intergenic |
| chr13 | 21808933 | 21809416 | Distal Intergenic |
| chr13 | 21809537 | 21811050 | Distal Intergenic |
| chr13 | 21812967 | 21814593 | Distal Intergenic |
| chr13 | 21826246 | 21826821 | Distal Intergenic |
| chr13 | 21827041 | 21827642 | Distal Intergenic |
| chr13 | 21841008 | 21841599 | Distal Intergenic |
| chr13 | 21841688 | 21842909 | Distal Intergenic |
| chr13 | 21843965 | 21844822 | Distal Intergenic |
| chr13 | 21845715 | 21847234 | Distal Intergenic |
| chr13 | 21870303 | 21872758 | Promoter |
| chr13 | 21873784 | 21875450 | Intron |
| chr13 | 21877523 | 21879266 | Intron |
| chr13 | 21879338 | 21880136 | Intron |
| chr13 | 21900513 | 21902191 | Intron |
| chr13 | 21902501 | 21904126 | Intron |
| chr13 | 21921077 | 21921276 | Intron |
| chr13 | 21922719 | 21923120 | Intron |
| chr13 | 21923330 | 21925395 | Exon |
| chr13 | 21925533 | 21927086 | Intron |
| chr13 | 22125564 | 22126048 | Intron |
| chr13 | 22126677 | 22126926 | Intron |
| chr13 | 22127027 | 22127383 | Intron |
| chr13 | 22127940 | 22128749 | Intron |
| chr13 | 22131580 | 22134682 | Intron |
| chr13 | 22135505 | 22135756 | Intron |
| chr13 | 22135794 | 22136161 | Intron |
| chr13 | 23460515 | 23461076 | Distal Intergenic |
| chr13 | 23536694 | 23537395 | Distal Intergenic |
| chr13 | 23537453 | 23537751 | Distal Intergenic |
| chr13 | 23597929 | 23598141 | Distal Intergenic |
| chr13 | 23622324 | 23622818 | Distal Intergenic |
| chr13 | 23623016 | 23625438 | Distal Intergenic |
| chr13 | 23625895 | 23628523 | Distal Intergenic |
| chr13 | 23633921 | 23634121 | Distal Intergenic |
| chr13 | 23634202 | 23636051 | Distal Intergenic |
| chr13 | 23636307 | 23637835 | Distal Intergenic |
| chr13 | 23641587 | 23641822 | Distal Intergenic |
| chr13 | 23642359 | 23644173 | Distal Intergenic |
| chr13 | 23646602 | 23649121 | Distal Intergenic |
| chr13 | 23652773 | 23654221 | Distal Intergenic |
| chr13 | 23654893 | 23655211 | Distal Intergenic |

| | | | |
|---|---|---|---|
| chr13 | 23655390 | 23655648 | Distal Intergenic |
| chr13 | 23661472 | 23661687 | Distal Intergenic |
| chr13 | 23661992 | 23663040 | Distal Intergenic |
| chr13 | 23663148 | 23666333 | Distal Intergenic |
| chr13 | 23666386 | 23667022 | Distal Intergenic |
| chr13 | 23667134 | 23668628 | Distal Intergenic |
| chr13 | 23668676 | 23669185 | Distal Intergenic |
| chr13 | 23672960 | 23675383 | Distal Intergenic |
| chr13 | 23675566 | 23678082 | Distal Intergenic |
| chr13 | 23678300 | 23678660 | Distal Intergenic |
| chr13 | 23712622 | 23713976 | Distal Intergenic |
| chr13 | 23751240 | 23751505 | Distal Intergenic |
| chr13 | 23751757 | 23752375 | Distal Intergenic |
| chr13 | 23774145 | 23774723 | Intron |
| chr13 | 23774815 | 23775507 | Intron |
| chr13 | 23775568 | 23775884 | Intron |
| chr13 | 23775927 | 23777720 | Intron |
| chr13 | 23789483 | 23790227 | Intron |
| chr13 | 23830191 | 23832706 | Intron |
| chr13 | 23835952 | 23838494 | Intron |
| chr13 | 23838647 | 23843037 | Intron |
| chr13 | 23843173 | 23845736 | Intron |
| chr13 | 23845778 | 23845991 | Intron |
| chr13 | 23848348 | 23848734 | Intron |
| chr13 | 23848803 | 23850101 | Intron |
| chr13 | 23852076 | 23853029 | Intron |
| chr13 | 24188251 | 24188558 | Intron |
| chr13 | 24195569 | 24196955 | Intron |
| chr13 | 24197060 | 24198013 | Intron |
| chr13 | 24466650 | 24468286 | Exon |
| chr13 | 24489016 | 24489617 | Intron |
| chr13 | 24612871 | 24613083 | Intron |
| chr13 | 24613171 | 24613801 | Intron |
| chr13 | 24613841 | 24614421 | Intron |
| chr13 | 24628913 | 24629303 | Intron |
| chr13 | 24656622 | 24658692 | Intron |
| chr13 | 24766007 | 24766501 | Intron |
| chr13 | 24770753 | 24771856 | Intron |
| chr13 | 24892741 | 24893702 | Exon |
| chr13 | 24893756 | 24894002 | Intron |
| chr13 | 24894046 | 24894367 | Intron |
| chr13 | 25163028 | 25163232 | Intron |
| chr13 | 25840662 | 25840932 | Intron |
| chr13 | 27032094 | 27032448 | Distal Intergenic |

| | | | |
|---|---|---|---|
| chr13 | 27095812 | 27096534 | Distal Intergenic |
| chr13 | 27119075 | 27119422 | Distal Intergenic |
| chr13 | 27204654 | 27206261 | Intron |
| chr13 | 27240429 | 27241141 | Intron |
| chr13 | 27241376 | 27241604 | Intron |
| chr13 | 27674210 | 27674523 | Intron |
| chr13 | 27677911 | 27678330 | Intron |
| chr13 | 27678456 | 27678726 | Intron |
| chr13 | 27752133 | 27752774 | Distal Intergenic |
| chr13 | 27810804 | 27811527 | Distal Intergenic |
| chr13 | 27816617 | 27817906 | Distal Intergenic |
| chr13 | 27820867 | 27823937 | Distal Intergenic |
| chr13 | 27825434 | 27827470 | Promoter |
| chr13 | 27890644 | 27891605 | Distal Intergenic |
| chr13 | 27891643 | 27892020 | Distal Intergenic |
| chr13 | 28548787 | 28550366 | Downstream |
| chr13 | 28721183 | 28721941 | Intron |
| chr13 | 28810887 | 28811307 | Intron |
| chr13 | 28829546 | 28830105 | Intron |
| chr13 | 28976067 | 28976385 | Intron |
| chr13 | 29008877 | 29010284 | Intron |
| chr13 | 29042129 | 29042707 | Intron |
| chr13 | 29042757 | 29043488 | Intron |
| chr13 | 29043772 | 29045318 | Intron |
| chr13 | 29593172 | 29596226 | Distal Intergenic |
| chr13 | 29596416 | 29597126 | Distal Intergenic |
| chr13 | 29719287 | 29720672 | Intron |
| chr13 | 30075373 | 30075741 | Intron |
| chr13 | 30076650 | 30076849 | Intron |
| chr13 | 30076892 | 30077541 | 3' UTR |
| chr13 | 30200362 | 30200572 | Distal Intergenic |
| chr13 | 30200979 | 30201596 | Distal Intergenic |
| chr13 | 30241007 | 30241496 | Distal Intergenic |
| chr13 | 30253554 | 30253939 | Distal Intergenic |
| chr13 | 30285744 | 30286093 | Distal Intergenic |
| chr13 | 30286203 | 30287167 | Distal Intergenic |
| chr13 | 31023690 | 31024183 | Distal Intergenic |
| chr13 | 31064574 | 31064773 | Intron |
| chr13 | 31064944 | 31065781 | Intron |
| chr13 | 31343400 | 31344413 | Distal Intergenic |
| chr13 | 31419498 | 31419915 | Distal Intergenic |
| chr13 | 31422668 | 31423777 | Distal Intergenic |
| chr13 | 31423971 | 31424603 | Distal Intergenic |
| chr13 | 31426526 | 31427162 | Distal Intergenic |

| | | | |
|---|---|---|---|
| chr13 | 31433332 | 31433556 | Distal Intergenic |
| chr13 | 31497357 | 31497636 | Intron |
| chr13 | 31499951 | 31500546 | Intron |
| chr13 | 31500747 | 31501024 | Intron |
| chr13 | 31659147 | 31660043 | Distal Intergenic |
| chr13 | 31669309 | 31669709 | Distal Intergenic |
| chr13 | 31669926 | 31670487 | Distal Intergenic |
| chr13 | 31899181 | 31899567 | Intron |
| chr13 | 31900661 | 31901215 | Intron |
| chr13 | 31901316 | 31901984 | Intron |
| chr13 | 31902196 | 31902395 | Intron |
| chr13 | 31955195 | 31956420 | Distal Intergenic |
| chr13 | 31968707 | 31968948 | Distal Intergenic |
| chr13 | 32226335 | 32227369 | Distal Intergenic |
| chr13 | 32570602 | 32571597 | Distal Intergenic |
| chr13 | 32683526 | 32684311 | Intron |
| chr13 | 32752127 | 32752396 | 5' UTR |
| chr13 | 32752498 | 32752697 | 5' UTR |
| chr13 | 32924067 | 32925210 | Intron |
| chr13 | 32937895 | 32938379 | Intron |
| chr13 | 32976462 | 32976802 | 3' UTR |
| chr13 | 32995361 | 32995711 | Intron |
| chr13 | 33004498 | 33005465 | Downstream |
| chr13 | 33005507 | 33005709 | Downstream |
| chr13 | 33015898 | 33016133 | Intron |
| chr13 | 33016284 | 33017683 | Exon |
| chr13 | 33018030 | 33020823 | 5' UTR |
| chr13 | 33030872 | 33031883 | Intron |
| chr13 | 33055586 | 33057074 | Promoter |
| chr13 | 33057112 | 33057546 | Intron |
| chr13 | 33095250 | 33097290 | 5' UTR |
| chr13 | 33118476 | 33118944 | Distal Intergenic |
| chr13 | 33119174 | 33119446 | Distal Intergenic |
| chr13 | 33119515 | 33119757 | Distal Intergenic |
| chr13 | 33119824 | 33121624 | Distal Intergenic |
| chr13 | 33142488 | 33143220 | Distal Intergenic |
| chr13 | 33143757 | 33144083 | Distal Intergenic |
| chr13 | 33158620 | 33158936 | Distal Intergenic |
| chr13 | 33158994 | 33160190 | Promoter |
| chr13 | 33169767 | 33170004 | Intron |
| chr13 | 33170055 | 33171677 | Intron |
| chr13 | 33181354 | 33181666 | Intron |
| chr13 | 33181813 | 33182872 | Intron |
| chr13 | 33182932 | 33183441 | Intron |

| | | | |
|---|---|---|---|
| chr13 | 34026725 | 34027409 | Intron |
| chr13 | 34027474 | 34027824 | Intron |
| chr13 | 34027959 | 34028365 | Intron |
| chr13 | 34094206 | 34094459 | Intron |
| chr13 | 34171946 | 34172325 | Intron |
| chr13 | 34187942 | 34189032 | Intron |
| chr13 | 34201378 | 34202206 | Intron |
| chr13 | 34203590 | 34203801 | Intron |
| chr13 | 34203890 | 34205813 | Intron |
| chr13 | 34221750 | 34222354 | Intron |
| chr13 | 34254345 | 34254844 | Distal Intergenic |
| chr13 | 34281309 | 34284829 | Distal Intergenic |
| chr13 | 34284896 | 34285126 | Distal Intergenic |
| chr13 | 34285203 | 34285408 | Distal Intergenic |
| chr13 | 34301557 | 34302585 | Distal Intergenic |
| chr13 | 34304042 | 34304888 | Distal Intergenic |
| chr13 | 34304994 | 34305454 | Distal Intergenic |
| chr13 | 34418606 | 34419031 | Intron |
| chr13 | 34424344 | 34424774 | Intron |
| chr13 | 34424832 | 34425088 | Intron |
| chr13 | 34435068 | 34435336 | Intron |
| chr13 | 34435387 | 34435726 | Intron |
| chr13 | 34461648 | 34462365 | Intron |
| chr13 | 34462538 | 34464423 | Intron |
| chr13 | 34489935 | 34492481 | Intron |
| chr13 | 34493690 | 34495105 | Intron |
| chr13 | 34632236 | 34632488 | Distal Intergenic |
| chr13 | 34742699 | 34742909 | Distal Intergenic |
| chr13 | 34743059 | 34743935 | Distal Intergenic |
| chr13 | 34744339 | 34744639 | Distal Intergenic |
| chr13 | 34744860 | 34745333 | Distal Intergenic |
| chr13 | 34757784 | 34758320 | Distal Intergenic |
| chr13 | 34767871 | 34769695 | Distal Intergenic |
| chr13 | 34774830 | 34775033 | Distal Intergenic |
| chr13 | 34775342 | 34776636 | Distal Intergenic |
| chr13 | 34781591 | 34782543 | Distal Intergenic |
| chr13 | 34788653 | 34789552 | Distal Intergenic |
| chr13 | 34807033 | 34807550 | Distal Intergenic |
| chr13 | 34807618 | 34807979 | Distal Intergenic |
| chr13 | 34818109 | 34818367 | Distal Intergenic |
| chr13 | 34966842 | 34967614 | Distal Intergenic |
| chr13 | 34967697 | 34968042 | Distal Intergenic |
| chr13 | 35375677 | 35376056 | Distal Intergenic |
| chr13 | 35376101 | 35376563 | Distal Intergenic |

| | | | |
|---|---|---|---|
| chr13 | 35604223 | 35604689 | Intron |
| chr13 | 35925869 | 35926279 | Intron |
| chr13 | 36625943 | 36626900 | Intron |
| chr13 | 36702421 | 36702885 | Intron |
| chr13 | 37382419 | 37383177 | Distal Intergenic |
| chr13 | 37545070 | 37546817 | Exon |
| chr13 | 37875275 | 37877026 | Distal Intergenic |
| chr13 | 37919355 | 37919681 | Distal Intergenic |
| chr13 | 37920093 | 37920297 | Distal Intergenic |
| chr13 | 37922577 | 37922933 | Distal Intergenic |
| chr13 | 37938597 | 37939897 | Distal Intergenic |
| chr13 | 37957979 | 37959033 | Distal Intergenic |
| chr13 | 37959091 | 37959665 | Distal Intergenic |
| chr13 | 38000104 | 38001563 | Distal Intergenic |
| chr13 | 38001675 | 38002005 | Distal Intergenic |
| chr13 | 38002092 | 38003304 | Distal Intergenic |
| chr13 | 38015907 | 38016226 | Distal Intergenic |
| chr13 | 38162283 | 38162920 | Intron |
| chr13 | 38163066 | 38163361 | Intron |
| chr13 | 38171582 | 38174292 | Promoter |
| chr13 | 38174367 | 38175700 | Intron |
| chr13 | 38187790 | 38189446 | Distal Intergenic |
| chr13 | 38190294 | 38191156 | Distal Intergenic |
| chr13 | 38210319 | 38210912 | 3' UTR |
| chr13 | 38214955 | 38215183 | Intron |
| chr13 | 38215268 | 38216460 | Intron |
| chr13 | 38218879 | 38219899 | Intron |
| chr13 | 38236600 | 38237873 | Exon |
| chr13 | 38242251 | 38245744 | Intron |
| chr13 | 38245785 | 38255851 | 5' UTR |
| chr13 | 38296827 | 38297384 | Intron |
| chr13 | 38302229 | 38303451 | Intron |
| chr13 | 38303488 | 38303703 | Intron |
| chr13 | 38441202 | 38442897 | Intron |
| chr13 | 38806743 | 38806972 | Distal Intergenic |
| chr13 | 38807008 | 38807392 | Distal Intergenic |
| chr13 | 38868141 | 38868779 | Distal Intergenic |
| chr13 | 38885065 | 38885313 | Distal Intergenic |
| chr13 | 38885351 | 38885801 | Distal Intergenic |
| chr13 | 38907215 | 38907860 | Distal Intergenic |
| chr13 | 38907898 | 38908128 | Distal Intergenic |
| chr13 | 38908216 | 38908422 | Distal Intergenic |
| chr13 | 39703314 | 39703834 | Distal Intergenic |
| chr13 | 39999331 | 39999574 | Intron |

| | | | |
|---|---|---|---|
| chr13 | 40018454 | 40019083 | Intron |
| chr13 | 40019497 | 40020810 | Intron |
| chr13 | 40023974 | 40024792 | Intron |
| chr13 | 40779367 | 40779660 | Intron |
| chr13 | 40813810 | 40815737 | Distal Intergenic |
| chr13 | 40818822 | 40820115 | Distal Intergenic |
| chr13 | 40820169 | 40820370 | Distal Intergenic |
| chr13 | 40820465 | 40821118 | Distal Intergenic |
| chr13 | 40821430 | 40822015 | Distal Intergenic |
| chr13 | 40822149 | 40826247 | Distal Intergenic |
| chr13 | 40827023 | 40828957 | Distal Intergenic |
| chr13 | 40944295 | 40944757 | Intron |
| chr13 | 41011979 | 41012966 | Intron |
| chr13 | 41078347 | 41078589 | Intron |
| chr13 | 41081212 | 41081946 | Intron |
| chr13 | 41096136 | 41096827 | Intron |
| chr13 | 41111671 | 41112035 | Intron |
| chr13 | 41419367 | 41419797 | Intron |
| chr13 | 41419852 | 41420456 | Intron |
| chr13 | 41426253 | 41429606 | Exon |
| chr13 | 41430163 | 41430711 | Intron |
| chr13 | 41434317 | 41434807 | Intron |
| chr13 | 41443674 | 41443899 | Intron |
| chr13 | 41444784 | 41445078 | Intron |
| chr13 | 41445445 | 41445817 | Intron |
| chr13 | 41571802 | 41572469 | Intron |
| chr13 | 42328677 | 42329857 | Intron |
| chr13 | 42423947 | 42424316 | Intron |
| chr13 | 42573152 | 42574220 | Distal Intergenic |
| chr13 | 42650102 | 42650316 | Intron |
| chr13 | 42650525 | 42650773 | Intron |
| chr13 | 42650833 | 42651114 | Intron |
| chr13 | 42653278 | 42654661 | Intron |
| chr13 | 43243867 | 43245190 | Distal Intergenic |
| chr13 | 43249946 | 43250152 | Distal Intergenic |
| chr13 | 43268159 | 43268495 | Distal Intergenic |
| chr13 | 43288774 | 43289231 | Distal Intergenic |
| chr13 | 43330556 | 43330777 | Distal Intergenic |
| chr13 | 43388263 | 43388674 | Distal Intergenic |
| chr13 | 43574453 | 43574749 | Distal Intergenic |
| chr13 | 44182684 | 44182897 | Intron |
| chr13 | 44628373 | 44628607 | Distal Intergenic |
| chr13 | 44726607 | 44727494 | Intron |
| chr13 | 44755134 | 44755821 | Distal Intergenic |

| | | | |
|---|---|---|---|
| chr13 | 44755976 | 44756232 | Distal Intergenic |
| chr13 | 44798659 | 44799690 | Distal Intergenic |
| chr13 | 44826174 | 44826556 | Distal Intergenic |
| chr13 | 44827585 | 44827875 | Distal Intergenic |
| chr13 | 44828117 | 44828571 | Distal Intergenic |
| chr13 | 44829066 | 44829704 | Distal Intergenic |
| chr13 | 44914667 | 44915145 | Distal Intergenic |
| chr13 | 44941237 | 44941473 | Distal Intergenic |
| chr13 | 45034324 | 45036467 | Intron |
| chr13 | 45064446 | 45064657 | Intron |
| chr13 | 45370777 | 45371169 | Downstream |
| chr13 | 45371636 | 45371893 | Downstream |
| chr13 | 45373153 | 45373768 | Exon |
| chr13 | 45401425 | 45401681 | Distal Intergenic |
| chr13 | 45401789 | 45402224 | Distal Intergenic |
| chr13 | 45497200 | 45498865 | Distal Intergenic |
| chr13 | 45665084 | 45666052 | Distal Intergenic |
| chr13 | 45752861 | 45754310 | Intron |
| chr13 | 45779680 | 45781294 | Intron |
| chr13 | 45781387 | 45781774 | Exon |
| chr13 | 45790269 | 45790972 | Intron |
| chr13 | 45814511 | 45814710 | Intron |
| chr13 | 45814779 | 45815088 | Intron |
| chr13 | 45846568 | 45847046 | Intron |
| chr13 | 45897549 | 45899285 | Distal Intergenic |
| chr13 | 45923128 | 45923407 | Intron |
| chr13 | 45992564 | 45994432 | Promoter |
| chr13 | 46001833 | 46002236 | Distal Intergenic |
| chr13 | 46002358 | 46003652 | Distal Intergenic |
| chr13 | 46006922 | 46007583 | Distal Intergenic |
| chr13 | 46017576 | 46019126 | Distal Intergenic |
| chr13 | 46019170 | 46020992 | Distal Intergenic |
| chr13 | 46026624 | 46027035 | Distal Intergenic |
| chr13 | 46060832 | 46061727 | Intron |
| chr13 | 46062055 | 46062330 | Intron |
| chr13 | 46137903 | 46138179 | Exon |
| chr13 | 46138248 | 46141049 | Intron |
| chr13 | 46142354 | 46142759 | Exon |
| chr13 | 46203179 | 46204595 | Distal Intergenic |
| chr13 | 46204727 | 46205288 | Distal Intergenic |
| chr13 | 46322628 | 46323343 | Distal Intergenic |
| chr13 | 46377123 | 46378693 | Intron |
| chr13 | 46379140 | 46379438 | Intron |
| chr13 | 46379537 | 46381080 | Intron |

| | | | |
|---|---|---|---|
| chr13 | 46811669 | 46812211 | Intron |
| chr13 | 46820732 | 46821199 | Exon |
| chr13 | 46821248 | 46823359 | Intron |
| chr13 | 46935015 | 46938821 | Exon |
| chr13 | 47023466 | 47024661 | Distal Intergenic |
| chr13 | 47025341 | 47026478 | Distal Intergenic |
| chr13 | 47050116 | 47052111 | Distal Intergenic |
| chr13 | 47054127 | 47055471 | Distal Intergenic |
| chr13 | 47059658 | 47064310 | Distal Intergenic |
| chr13 | 47263411 | 47267179 | Exon |
| chr13 | 47275463 | 47275903 | Intron |
| chr13 | 47289883 | 47290523 | Intron |
| chr13 | 47311685 | 47312228 | Intron |
| chr13 | 47312379 | 47312994 | Intron |
| chr13 | 47340038 | 47340400 | Distal Intergenic |
| chr13 | 47340470 | 47340778 | Distal Intergenic |
| chr13 | 47368275 | 47368474 | Intron |
| chr13 | 47376487 | 47376962 | Distal Intergenic |
| chr13 | 47489729 | 47490485 | Distal Intergenic |
| chr13 | 47581910 | 47582109 | Distal Intergenic |
| chr13 | 48112281 | 48112527 | Distal Intergenic |
| chr13 | 48205192 | 48205532 | Distal Intergenic |
| chr13 | 48205651 | 48206492 | Distal Intergenic |
| chr13 | 48611045 | 48611366 | Promoter |
| chr13 | 48661123 | 48663826 | Intron |
| chr13 | 48666628 | 48666980 | Intron |
| chr13 | 48710141 | 48710374 | Distal Intergenic |
| chr13 | 48965007 | 48965450 | Intron |
| chr13 | 49028234 | 49028468 | Intron |
| chr13 | 49028529 | 49028885 | Intron |
| chr13 | 49054953 | 49055326 | 3' UTR |
| chr13 | 49055578 | 49058883 | 3' UTR |
| chr13 | 49061097 | 49062987 | Downstream |
| chr13 | 49063830 | 49064206 | 3' UTR |
| chr13 | 49279146 | 49279756 | Intron |
| chr13 | 49279794 | 49280003 | Promoter |
| chr13 | 49282214 | 49282657 | 3' UTR |
| chr13 | 49389323 | 49389604 | Distal Intergenic |
| chr13 | 49389654 | 49390031 | Distal Intergenic |
| chr13 | 49393665 | 49395257 | Distal Intergenic |
| chr13 | 49395312 | 49395521 | Distal Intergenic |
| chr13 | 49395867 | 49396469 | Distal Intergenic |
| chr13 | 49402736 | 49403077 | Distal Intergenic |
| chr13 | 49403322 | 49403537 | Distal Intergenic |

| | | | |
|---|---|---|---|
| chr13 | 49434513 | 49435013 | Distal Intergenic |
| chr13 | 49437752 | 49438492 | Distal Intergenic |
| chr13 | 49441219 | 49441515 | Distal Intergenic |
| chr13 | 49457354 | 49458579 | Distal Intergenic |
| chr13 | 49459398 | 49460975 | Distal Intergenic |
| chr13 | 49461037 | 49461495 | Distal Intergenic |
| chr13 | 49461875 | 49464212 | Distal Intergenic |
| chr13 | 49466756 | 49467034 | Distal Intergenic |
| chr13 | 49467078 | 49467668 | Distal Intergenic |
| chr13 | 49467869 | 49468449 | Distal Intergenic |
| chr13 | 49491760 | 49492058 | Distal Intergenic |
| chr13 | 49492175 | 49492544 | Distal Intergenic |
| chr13 | 49497743 | 49498142 | Distal Intergenic |
| chr13 | 49645210 | 49645499 | Intron |
| chr13 | 49679130 | 49679869 | Intron |
| chr13 | 49748481 | 49748869 | Exon |
| chr13 | 49777905 | 49778105 | Intron |
| chr13 | 50538142 | 50538468 | Distal Intergenic |
| chr13 | 51052687 | 51054919 | Intron |
| chr13 | 51055787 | 51056756 | Intron |
| chr13 | 51106109 | 51106590 | Intron |
| chr13 | 51106658 | 51106964 | Intron |
| chr13 | 51107044 | 51107270 | Intron |
| chr13 | 51109969 | 51110230 | Intron |
| chr13 | 51134368 | 51134587 | Intron |
| chr13 | 51134625 | 51135340 | Intron |
| chr13 | 51135896 | 51138002 | Intron |
| chr13 | 51157632 | 51159026 | Intron |
| chr13 | 51185167 | 51185652 | Intron |
| chr13 | 51185734 | 51186084 | Intron |
| chr13 | 51196835 | 51197681 | Intron |
| chr13 | 51208077 | 51212094 | Distal Intergenic |
| chr13 | 51223310 | 51223560 | Distal Intergenic |
| chr13 | 51396531 | 51397627 | 3' UTR |
| chr13 | 51423902 | 51424917 | Exon |
| chr13 | 51427973 | 51429593 | Distal Intergenic |
| chr13 | 51495115 | 51495594 | Intron |
| chr13 | 51495670 | 51496345 | Intron |
| chr13 | 51740667 | 51741091 | Intron |
| chr13 | 51741229 | 51741653 | Intron |
| chr13 | 51746616 | 51746826 | Promoter |
| chr13 | 51770653 | 51771256 | Distal Intergenic |
| chr13 | 51779988 | 51781417 | Distal Intergenic |
| chr13 | 51834065 | 51834275 | Intron |

| | | | |
|---|---|---|---|
| chr13 | 51851645 | 51851892 | Intron |
| chr13 | 51857771 | 51858097 | 3' UTR |
| chr13 | 51858135 | 51859245 | 3' UTR |
| chr13 | 51943364 | 51943590 | Exon |
| chr13 | 52060070 | 52060276 | Distal Intergenic |
| chr13 | 52064476 | 52064839 | Distal Intergenic |
| chr13 | 52070963 | 52071527 | Distal Intergenic |
| chr13 | 52284926 | 52285749 | Intron |
| chr13 | 52331198 | 52331456 | Intron |
| chr13 | 52337149 | 52337902 | Distal Intergenic |
| chr13 | 52401834 | 52403047 | Intron |
| chr13 | 52550140 | 52551134 | Intron |
| chr13 | 52573411 | 52574053 | Intron |
| chr13 | 52577114 | 52577323 | Intron |
| chr13 | 52577381 | 52577729 | Intron |
| chr13 | 52621646 | 52622024 | Distal Intergenic |
| chr13 | 52636747 | 52638393 | Downstream |
| chr13 | 52683256 | 52683590 | Intron |
| chr13 | 52754072 | 52754274 | Intron |
| chr13 | 52754397 | 52755746 | Exon |
| chr13 | 52770229 | 52772376 | Intron |
| chr13 | 52775321 | 52775573 | Intron |
| chr13 | 52975221 | 52975434 | Intron |
| chr13 | 53104068 | 53105029 | Intron |
| chr13 | 53113067 | 53113438 | Intron |
| chr13 | 53113513 | 53113838 | Intron |
| chr13 | 53113875 | 53114566 | Intron |
| chr13 | 53114635 | 53115011 | Intron |
| chr13 | 53325921 | 53326720 | Distal Intergenic |
| chr13 | 53434288 | 53434948 | Distal Intergenic |
| chr13 | 53435125 | 53435437 | Distal Intergenic |
| chr13 | 53813073 | 53813829 | Distal Intergenic |
| chr13 | 54196457 | 54197237 | Distal Intergenic |
| chr13 | 54635276 | 54635475 | Distal Intergenic |
| chr13 | 54691800 | 54692022 | Distal Intergenic |
| chr13 | 54706564 | 54707586 | Exon |
| chr13 | 54707821 | 54709139 | Distal Intergenic |
| chr13 | 54709497 | 54711033 | Distal Intergenic |
| chr13 | 54729995 | 54731238 | Distal Intergenic |
| chr13 | 54787823 | 54788898 | Distal Intergenic |
| chr13 | 55290702 | 55291654 | Distal Intergenic |
| chr13 | 55420228 | 55421482 | Distal Intergenic |
| chr13 | 55431425 | 55431624 | Distal Intergenic |
| chr13 | 55463609 | 55464202 | Distal Intergenic |

| | | | |
|---|---|---|---|
| chr13 | 55501319 | 55501660 | Distal Intergenic |
| chr13 | 55505293 | 55506400 | Distal Intergenic |
| chr13 | 55611408 | 55614478 | Distal Intergenic |
| chr13 | 55705944 | 55706143 | Distal Intergenic |
| chr13 | 55724399 | 55724598 | Distal Intergenic |
| chr13 | 55793701 | 55794579 | Distal Intergenic |
| chr13 | 55828924 | 55829193 | Distal Intergenic |
| chr13 | 55927492 | 55928256 | Distal Intergenic |
| chr13 | 55928360 | 55931705 | Distal Intergenic |
| chr13 | 55932141 | 55932581 | Distal Intergenic |
| chr13 | 55933364 | 55933563 | Distal Intergenic |
| chr13 | 55933737 | 55934042 | Distal Intergenic |
| chr13 | 55994231 | 55994850 | Distal Intergenic |
| chr13 | 55996580 | 55997415 | Distal Intergenic |
| chr13 | 56038135 | 56038334 | Distal Intergenic |
| chr13 | 56038509 | 56038798 | Distal Intergenic |
| chr13 | 56039190 | 56039443 | Distal Intergenic |
| chr13 | 56061426 | 56062144 | Distal Intergenic |
| chr13 | 56062342 | 56062621 | Distal Intergenic |
| chr13 | 56062709 | 56063351 | Distal Intergenic |
| chr13 | 56063535 | 56064057 | Distal Intergenic |
| chr13 | 56077123 | 56079068 | Distal Intergenic |
| chr13 | 56093914 | 56094476 | Distal Intergenic |
| chr13 | 56105405 | 56106419 | Distal Intergenic |
| chr13 | 56106803 | 56108455 | Distal Intergenic |
| chr13 | 56129180 | 56129379 | Distal Intergenic |
| chr13 | 56129528 | 56129780 | Distal Intergenic |
| chr13 | 56425482 | 56425725 | Distal Intergenic |
| chr13 | 56505812 | 56507644 | Distal Intergenic |
| chr13 | 56531735 | 56532109 | Distal Intergenic |
| chr13 | 56532158 | 56532557 | Distal Intergenic |
| chr13 | 56539013 | 56539320 | Distal Intergenic |
| chr13 | 56539374 | 56539602 | Distal Intergenic |
| chr13 | 56542185 | 56543009 | Distal Intergenic |
| chr13 | 56544017 | 56545700 | Distal Intergenic |
| chr13 | 56683477 | 56683779 | Distal Intergenic |
| chr13 | 56684248 | 56684552 | Distal Intergenic |
| chr13 | 56687011 | 56688054 | Distal Intergenic |
| chr13 | 56690001 | 56690900 | Distal Intergenic |
| chr13 | 56695355 | 56695573 | Distal Intergenic |
| chr13 | 56699881 | 56700667 | Distal Intergenic |
| chr13 | 56701101 | 56701327 | Distal Intergenic |
| chr13 | 56701463 | 56701777 | Distal Intergenic |
| chr13 | 56701830 | 56704049 | Distal Intergenic |

| | | | |
|---|---|---|---|
| chr13 | 56709049 | 56709269 | Distal Intergenic |
| chr13 | 56709373 | 56709790 | Distal Intergenic |
| chr13 | 56710155 | 56711008 | Distal Intergenic |
| chr13 | 56711225 | 56711468 | Distal Intergenic |
| chr13 | 56714387 | 56715877 | Distal Intergenic |
| chr13 | 56804409 | 56804647 | Distal Intergenic |
| chr13 | 56813351 | 56813636 | Distal Intergenic |
| chr13 | 57959946 | 57960154 | Distal Intergenic |
| chr13 | 58228036 | 58228292 | Intron |
| chr13 | 58229238 | 58229870 | Intron |
| chr13 | 58230361 | 58231603 | Intron |
| chr13 | 58315987 | 58316336 | Distal Intergenic |
| chr13 | 58502030 | 58502799 | Distal Intergenic |
| chr13 | 58503029 | 58503341 | Distal Intergenic |
| chr13 | 58504032 | 58504246 | Distal Intergenic |
| chr13 | 58504397 | 58504618 | Distal Intergenic |
| chr13 | 58621887 | 58622139 | Distal Intergenic |
| chr13 | 58635719 | 58636785 | Distal Intergenic |
| chr13 | 59725362 | 59726330 | Distal Intergenic |
| chr13 | 59747931 | 59748877 | Distal Intergenic |
| chr13 | 59871641 | 59871857 | Distal Intergenic |
| chr13 | 59925878 | 59926091 | Distal Intergenic |
| chr13 | 59958080 | 59958378 | Distal Intergenic |
| chr13 | 59977149 | 59977722 | Distal Intergenic |
| chr13 | 60120505 | 60121768 | Distal Intergenic |
| chr13 | 60145727 | 60146744 | Distal Intergenic |
| chr13 | 60146829 | 60147046 | Distal Intergenic |
| chr13 | 60148450 | 60150387 | Distal Intergenic |
| chr13 | 60152408 | 60152752 | Distal Intergenic |
| chr13 | 60411192 | 60411392 | Intron |
| chr13 | 60528758 | 60529036 | Intron |
| chr13 | 60529277 | 60529695 | Intron |
| chr13 | 60719630 | 60720231 | Promoter |
| chr13 | 60759490 | 60760794 | Distal Intergenic |
| chr13 | 60761371 | 60762849 | Distal Intergenic |
| chr13 | 60763425 | 60764077 | Distal Intergenic |
| chr13 | 60998926 | 60999447 | Intron |
| chr13 | 63147409 | 63147823 | Distal Intergenic |
| chr13 | 63147926 | 63148417 | Distal Intergenic |
| chr13 | 63166784 | 63168905 | Distal Intergenic |
| chr13 | 63171884 | 63172225 | Distal Intergenic |
| chr13 | 63172593 | 63176418 | Distal Intergenic |
| chr13 | 63188893 | 63191939 | Distal Intergenic |
| chr13 | 63193521 | 63195524 | Distal Intergenic |

| | | | |
|---|---|---|---|
| chr13 | 63218883 | 63219748 | Distal Intergenic |
| chr13 | 63220946 | 63222944 | Distal Intergenic |
| chr13 | 63257438 | 63257797 | Distal Intergenic |
| chr13 | 63287866 | 63289359 | Distal Intergenic |
| chr13 | 63294523 | 63294722 | Distal Intergenic |
| chr13 | 63318086 | 63318299 | Distal Intergenic |
| chr13 | 63318410 | 63318612 | Distal Intergenic |
| chr13 | 63341511 | 63343105 | Distal Intergenic |
| chr13 | 63343143 | 63344052 | Distal Intergenic |
| chr13 | 63344169 | 63344411 | Distal Intergenic |
| chr13 | 63344466 | 63344843 | Distal Intergenic |
| chr13 | 63353460 | 63355316 | Distal Intergenic |
| chr13 | 63356056 | 63356481 | Distal Intergenic |
| chr13 | 63367316 | 63367898 | Distal Intergenic |
| chr13 | 63375428 | 63376302 | Distal Intergenic |
| chr13 | 63379379 | 63380198 | Distal Intergenic |
| chr13 | 63380270 | 63380575 | Distal Intergenic |
| chr13 | 63380784 | 63382124 | Distal Intergenic |
| chr13 | 63382314 | 63382513 | Distal Intergenic |
| chr13 | 63405605 | 63406921 | Distal Intergenic |
| chr13 | 63407104 | 63407679 | Distal Intergenic |
| chr13 | 63665151 | 63666217 | Distal Intergenic |
| chr13 | 63675909 | 63676111 | Distal Intergenic |
| chr13 | 63677130 | 63677393 | Distal Intergenic |
| chr13 | 63677593 | 63678299 | Distal Intergenic |
| chr13 | 63773775 | 63774148 | Distal Intergenic |
| chr13 | 63783811 | 63784382 | Distal Intergenic |
| chr13 | 64255470 | 64255822 | Distal Intergenic |
| chr13 | 64256061 | 64256279 | Distal Intergenic |
| chr13 | 64357869 | 64360093 | Distal Intergenic |
| chr13 | 64408988 | 64409507 | Intron |
| chr13 | 64409699 | 64409921 | Intron |
| chr13 | 64432850 | 64434445 | Distal Intergenic |
| chr13 | 64438572 | 64440204 | Distal Intergenic |
| chr13 | 64471277 | 64471580 | Distal Intergenic |
| chr13 | 64471781 | 64472128 | Distal Intergenic |
| chr13 | 64478825 | 64479068 | Distal Intergenic |
| chr13 | 64489752 | 64490585 | Distal Intergenic |
| chr13 | 64501920 | 64503702 | Distal Intergenic |
| chr13 | 64522747 | 64523134 | Distal Intergenic |
| chr13 | 64523229 | 64524132 | Distal Intergenic |
| chr13 | 65213860 | 65214517 | Distal Intergenic |
| chr13 | 67005644 | 67006019 | Intron |
| chr13 | 67033582 | 67033868 | Intron |

| | | | |
|---|---|---|---|
| chr13 | 67034022 | 67034404 | Intron |
| chr13 | 67209315 | 67209566 | Intron |
| chr13 | 67461146 | 67461360 | Intron |
| chr13 | 67653534 | 67654039 | Intron |
| chr13 | 67885058 | 67885266 | Distal Intergenic |
| chr13 | 67885318 | 67885727 | Distal Intergenic |
| chr13 | 70861933 | 70863911 | Distal Intergenic |
| chr13 | 70883505 | 70883769 | Distal Intergenic |
| chr13 | 70932106 | 70932305 | Distal Intergenic |
| chr13 | 70932598 | 70932910 | Distal Intergenic |
| chr13 | 70932998 | 70933210 | Distal Intergenic |
| chr13 | 70933315 | 70933834 | Distal Intergenic |
| chr13 | 71785501 | 71787897 | Distal Intergenic |
| chr13 | 71789238 | 71789597 | Distal Intergenic |
| chr13 | 71796328 | 71796527 | Distal Intergenic |
| chr13 | 71948782 | 71949430 | Distal Intergenic |
| chr13 | 71949637 | 71949869 | Distal Intergenic |
| chr13 | 72100069 | 72100367 | Intron |
| chr13 | 72335088 | 72337053 | Intron |
| chr13 | 73511213 | 73512060 | Intron |
| chr13 | 73512100 | 73512657 | Intron |
| chr13 | 73567371 | 73567953 | Intron |
| chr13 | 73568338 | 73568555 | Intron |
| chr13 | 73568686 | 73570562 | Intron |
| chr13 | 73643255 | 73643504 | Intron |
| chr13 | 74163007 | 74163972 | Distal Intergenic |
| chr13 | 74307415 | 74307634 | Intron |
| chr13 | 74438657 | 74440042 | Intron |
| chr13 | 74454291 | 74454772 | Intron |
| chr13 | 74776694 | 74777693 | Distal Intergenic |
| chr13 | 74902802 | 74903011 | Distal Intergenic |
| chr13 | 74904823 | 74905120 | Distal Intergenic |
| chr13 | 74905185 | 74905468 | Distal Intergenic |
| chr13 | 74906263 | 74907548 | Distal Intergenic |
| chr13 | 74907618 | 74910808 | Distal Intergenic |
| chr13 | 74944707 | 74946469 | Distal Intergenic |
| chr13 | 74959139 | 74959893 | Distal Intergenic |
| chr13 | 75597971 | 75598765 | Distal Intergenic |
| chr13 | 75680826 | 75681083 | Distal Intergenic |
| chr13 | 75846083 | 75846314 | Distal Intergenic |
| chr13 | 75846353 | 75846570 | Distal Intergenic |
| chr13 | 75889670 | 75890248 | Promoter |
| chr13 | 75891015 | 75891240 | Promoter |
| chr13 | 75891292 | 75892481 | Promoter |

| | | | |
|---|---|---|---|
| chr13 | 75926340 | 75927535 | Intron |
| chr13 | 75935000 | 75935349 | Intron |
| chr13 | 75935557 | 75936593 | Exon |
| chr13 | 75962553 | 75962958 | Intron |
| chr13 | 76155472 | 76155943 | Intron |
| chr13 | 76239020 | 76239988 | Intron |
| chr13 | 76260062 | 76261494 | Intron |
| chr13 | 81024574 | 81024792 | Distal Intergenic |
| chr13 | 81025069 | 81025338 | Distal Intergenic |
| chr13 | 81025491 | 81025690 | Distal Intergenic |
| chr13 | 81235387 | 81235616 | Distal Intergenic |
| chr13 | 81315280 | 81315623 | Distal Intergenic |
| chr13 | 81315815 | 81316136 | Distal Intergenic |
| chr13 | 81316562 | 81317324 | Distal Intergenic |
| chr13 | 81818459 | 81818772 | Distal Intergenic |
| chr13 | 81822419 | 81822680 | Distal Intergenic |
| chr13 | 81822993 | 81824318 | Distal Intergenic |
| chr13 | 82121203 | 82121473 | Distal Intergenic |
| chr13 | 82121601 | 82121800 | Distal Intergenic |
| chr13 | 85427658 | 85428069 | Distal Intergenic |
| chr13 | 85428555 | 85428786 | Distal Intergenic |
| chr13 | 86185686 | 86186279 | Distal Intergenic |
| chr13 | 90356311 | 90356806 | Distal Intergenic |
| chr13 | 90714935 | 90715135 | Intron |
| chr13 | 90936265 | 90937628 | Distal Intergenic |
| chr13 | 91062546 | 91063001 | Distal Intergenic |
| chr13 | 91116727 | 91117307 | Distal Intergenic |
| chr13 | 91120205 | 91122445 | Distal Intergenic |
| chr13 | 91177597 | 91181632 | Exon |
| chr13 | 91303751 | 91304900 | Distal Intergenic |
| chr13 | 91304990 | 91306323 | Distal Intergenic |
| chr13 | 91317313 | 91319284 | Distal Intergenic |
| chr13 | 91345746 | 91346279 | Distal Intergenic |
| chr13 | 91356575 | 91357094 | Distal Intergenic |
| chr13 | 91359242 | 91360087 | Distal Intergenic |
| chr13 | 91363360 | 91363585 | Distal Intergenic |
| chr13 | 91363767 | 91364687 | Distal Intergenic |
| chr13 | 91445315 | 91446758 | Distal Intergenic |
| chr13 | 91447478 | 91448326 | Distal Intergenic |
| chr13 | 91465919 | 91467376 | Distal Intergenic |
| chr13 | 91489066 | 91489586 | Distal Intergenic |
| chr13 | 91506345 | 91506695 | Distal Intergenic |
| chr13 | 91789037 | 91789512 | Intron |
| chr13 | 91789713 | 91789937 | Intron |

| | | | |
|---|---|---|---|
| chr13 | 91941029 | 91941952 | Distal Intergenic |
| chr13 | 91947030 | 91947229 | Distal Intergenic |
| chr13 | 91947289 | 91947517 | Distal Intergenic |
| chr13 | 92842377 | 92842727 | Intron |
| chr13 | 93445259 | 93445886 | Intron |
| chr13 | 93446073 | 93446948 | Intron |
| chr13 | 93961006 | 93961408 | Intron |
| chr13 | 93961563 | 93962116 | Intron |
| chr13 | 94192970 | 94193615 | Intron |
| chr13 | 94254359 | 94254712 | Intron |
| chr13 | 94262406 | 94262792 | Intron |
| chr13 | 94263371 | 94263780 | Intron |
| chr13 | 94267003 | 94267300 | Intron |
| chr13 | 94267611 | 94267898 | Intron |
| chr13 | 94762092 | 94763161 | Intron |
| chr13 | 94767690 | 94767929 | Intron |
| chr13 | 94768198 | 94768804 | Intron |
| chr13 | 95199547 | 95199777 | Distal Intergenic |
| chr13 | 95587623 | 95588402 | Exon |
| chr13 | 95597757 | 95598281 | Distal Intergenic |
| chr13 | 95598367 | 95600830 | Distal Intergenic |
| chr13 | 96254666 | 96255009 | Intron |
| chr13 | 96258783 | 96259595 | Intron |
| chr13 | 96284762 | 96285124 | Intron |
| chr13 | 96285177 | 96285410 | Intron |
| chr13 | 96410514 | 96411794 | Intron |
| chr13 | 96413730 | 96414172 | Intron |
| chr13 | 96560685 | 96562424 | Intron |
| chr13 | 96564052 | 96564766 | Intron |
| chr13 | 96780852 | 96781552 | Intron |
| chr13 | 96785211 | 96785707 | Intron |
| chr13 | 96785911 | 96786132 | Intron |
| chr13 | 97257406 | 97257788 | Intron |
| chr13 | 97280567 | 97281579 | Intron |
| chr13 | 97312259 | 97312557 | Intron |
| chr13 | 97344064 | 97344272 | Intron |
| chr13 | 97344367 | 97344597 | Intron |
| chr13 | 97439484 | 97439747 | Intron |
| chr13 | 97439892 | 97440838 | Intron |
| chr13 | 97441262 | 97441710 | Intron |
| chr13 | 97450534 | 97451180 | Intron |
| chr13 | 97451247 | 97451471 | Intron |
| chr13 | 97907963 | 97908162 | Intron |
| chr13 | 97971130 | 97971467 | Intron |

| | | | |
|---|---|---|---|
| chr13 | 98018277 | 98018509 | Intron |
| chr13 | 98018827 | 98020165 | Intron |
| chr13 | 98252599 | 98252848 | Distal Intergenic |
| chr13 | 98253302 | 98253755 | Distal Intergenic |
| chr13 | 98254320 | 98256187 | Distal Intergenic |
| chr13 | 98257925 | 98258129 | Distal Intergenic |
| chr13 | 98267055 | 98267285 | Distal Intergenic |
| chr13 | 98267841 | 98268078 | Distal Intergenic |
| chr13 | 98612342 | 98612555 | Intron |
| chr13 | 98697946 | 98698359 | Distal Intergenic |
| chr13 | 98698488 | 98698687 | Distal Intergenic |
| chr13 | 98783545 | 98784130 | Distal Intergenic |
| chr13 | 98784251 | 98785441 | Distal Intergenic |
| chr13 | 98785509 | 98786051 | Distal Intergenic |
| chr13 | 98786100 | 98786393 | Distal Intergenic |
| chr13 | 98797134 | 98798267 | Intron |
| chr13 | 98808593 | 98808921 | Intron |
| chr13 | 98808963 | 98809360 | Intron |
| chr13 | 98891566 | 98892863 | Intron |
| chr13 | 98902835 | 98903108 | Intron |
| chr13 | 99086360 | 99086623 | Intron |
| chr13 | 99086670 | 99086945 | Intron |
| chr13 | 99204420 | 99204678 | Intron |
| chr13 | 99204757 | 99205315 | Intron |
| chr13 | 99393004 | 99393272 | Intron |
| chr13 | 99402275 | 99402746 | Intron |
| chr13 | 99404932 | 99405217 | Promoter |
| chr13 | 99449570 | 99450287 | Exon |
| chr13 | 99638329 | 99638668 | Intron |
| chr13 | 99638763 | 99639013 | Intron |
| chr13 | 99663915 | 99664493 | Intron |
| chr13 | 99695243 | 99695457 | Intron |
| chr13 | 99940487 | 99940748 | Intron |
| chr13 | 99945235 | 99945506 | Intron |
| chr13 | 100008662 | 100008873 | Promoter |
| chr13 | 100077788 | 100078019 | Distal Intergenic |
| chr13 | 100078651 | 100078874 | Distal Intergenic |
| chr13 | 100101509 | 100101955 | Distal Intergenic |
| chr13 | 100148478 | 100148920 | Intron |
| chr13 | 100154030 | 100154386 | Promoter |
| chr13 | 100154472 | 100155472 | Promoter |
| chr13 | 100161340 | 100161832 | Intron |
| chr13 | 100161881 | 100162125 | Intron |
| chr13 | 100212720 | 100213595 | Intron |

| | | | |
|---|---|---|---|
| chr13 | 100217280 | 100218061 | Distal Intergenic |
| chr13 | 100219142 | 100220532 | Distal Intergenic |
| chr13 | 100229147 | 100229694 | Distal Intergenic |
| chr13 | 100245561 | 100245793 | Distal Intergenic |
| chr13 | 100362677 | 100362928 | Intron |
| chr13 | 100531320 | 100531909 | Intron |
| chr13 | 100708828 | 100709389 | Distal Intergenic |
| chr13 | 100717611 | 100717972 | Distal Intergenic |
| chr13 | 100733666 | 100733901 | Distal Intergenic |
| chr13 | 100873401 | 100873736 | Intron |
| chr13 | 101286508 | 101286794 | Intron |
| chr13 | 101421099 | 101421416 | Intron |
| chr13 | 101500260 | 101500872 | Intron |
| chr13 | 103342768 | 103343879 | Exon |
| chr13 | 103352556 | 103352767 | Intron |
| chr13 | 103354119 | 103354648 | Intron |
| chr13 | 103364370 | 103364677 | Distal Intergenic |
| chr13 | 103638781 | 103639338 | Distal Intergenic |
| chr13 | 103670215 | 103670446 | Distal Intergenic |
| chr13 | 103670558 | 103671071 | Distal Intergenic |
| chr13 | 103678913 | 103679639 | Distal Intergenic |
| chr13 | 103685975 | 103686498 | Distal Intergenic |
| chr13 | 103687764 | 103688346 | Distal Intergenic |
| chr13 | 103688409 | 103689416 | Distal Intergenic |
| chr13 | 103689479 | 103690185 | Distal Intergenic |
| chr13 | 103690744 | 103690948 | Distal Intergenic |
| chr13 | 103695392 | 103696571 | 3' UTR |
| chr13 | 103748618 | 103749029 | Distal Intergenic |
| chr13 | 104176866 | 104177163 | Distal Intergenic |
| chr13 | 104177361 | 104177659 | Distal Intergenic |
| chr13 | 104564057 | 104564286 | Distal Intergenic |
| chr13 | 104709631 | 104710087 | Distal Intergenic |
| chr13 | 104710303 | 104710713 | Distal Intergenic |
| chr13 | 104768098 | 104769022 | Distal Intergenic |
| chr13 | 104831166 | 104831840 | Distal Intergenic |
| chr13 | 104864885 | 104865238 | Distal Intergenic |
| chr13 | 105310413 | 105311762 | Distal Intergenic |
| chr13 | 105606574 | 105607209 | Distal Intergenic |
| chr13 | 108601454 | 108603098 | Distal Intergenic |
| chr13 | 108603289 | 108605284 | Distal Intergenic |
| chr13 | 108617184 | 108617454 | Distal Intergenic |
| chr13 | 108617789 | 108619140 | Distal Intergenic |
| chr13 | 108620025 | 108621438 | Distal Intergenic |
| chr13 | 108623843 | 108624042 | Distal Intergenic |

| | | | |
|---|---|---|---|
| chr13 | 108656623 | 108656888 | Distal Intergenic |
| chr13 | 108833339 | 108833776 | Distal Intergenic |
| chr13 | 108833860 | 108834566 | Distal Intergenic |
| chr13 | 108834840 | 108835104 | Distal Intergenic |
| chr13 | 108835207 | 108835532 | Distal Intergenic |
| chr13 | 108835779 | 108836335 | Distal Intergenic |
| chr13 | 109127759 | 109128080 | Distal Intergenic |
| chr13 | 110033125 | 110033327 | Distal Intergenic |
| chr13 | 110050194 | 110051576 | Distal Intergenic |
| chr13 | 110058805 | 110059635 | Distal Intergenic |
| chr13 | 110068589 | 110069020 | Distal Intergenic |
| chr13 | 110069089 | 110069804 | Distal Intergenic |
| chr13 | 110609388 | 110610974 | Distal Intergenic |
| chr13 | 110611543 | 110612921 | Distal Intergenic |
| chr13 | 110613146 | 110614388 | Distal Intergenic |
| chr13 | 110688143 | 110688797 | Distal Intergenic |
| chr13 | 111008687 | 111008939 | Intron |
| chr13 | 111009032 | 111009281 | Intron |
| chr13 | 111185260 | 111185744 | Intron |
| chr13 | 111331797 | 111331996 | Intron |
| chr13 | 111514427 | 111516403 | Exon |
| chr13 | 111698061 | 111698417 | Distal Intergenic |
| chr13 | 111704004 | 111704328 | Distal Intergenic |
| chr13 | 111704455 | 111704673 | Distal Intergenic |
| chr13 | 111715436 | 111716059 | Distal Intergenic |
| chr13 | 111753610 | 111754081 | Distal Intergenic |
| chr13 | 111756736 | 111758390 | Distal Intergenic |
| chr13 | 112279841 | 112280091 | Distal Intergenic |
| chr13 | 112817799 | 112818842 | Distal Intergenic |
| chr13 | 112845159 | 112845374 | Distal Intergenic |
| chr13 | 112861195 | 112862356 | Distal Intergenic |
| chr13 | 113172991 | 113173509 | Exon |
| chr13 | 113548300 | 113548851 | Distal Intergenic |
| chr13 | 113560693 | 113560924 | Intron |
| chr13 | 113561003 | 113561241 | Intron |
| chr13 | 113954391 | 113955025 | Intron |
| chr13 | 114048899 | 114049168 | Distal Intergenic |
| chr13 | 114134424 | 114135231 | Exon |
| chr13 | 114141147 | 114141670 | Intron |
| chr13 | 114285324 | 114286687 | Exon |
| chr13 | 114391534 | 114391735 | Intron |
| chr13 | 114635008 | 114635216 | Distal Intergenic |
| chr13 | 115419755 | 115420208 | Distal Intergenic |
| chr13 | 115420371 | 115420599 | Distal Intergenic |

| | | | |
|---|---|---|---|
| chr13 | 115425488 | 115426548 | Distal Intergenic |
| chr13 | 115428750 | 115430822 | Distal Intergenic |
| chr13 | 115433443 | 115433645 | Distal Intergenic |
| chr13 | 115433837 | 115434631 | Distal Intergenic |
| chr13 | 115469505 | 115469720 | Distal Intergenic |
| chr13 | 115525044 | 115525379 | Distal Intergenic |
| chr13 | 115525613 | 115526804 | Distal Intergenic |
| chr13 | 115720733 | 115720941 | Distal Intergenic |
| chr13 | 115735302 | 115736961 | Distal Intergenic |
| chr13 | 117617750 | 117617964 | Distal Intergenic |
| chr13 | 117739045 | 117740529 | Distal Intergenic |
| chr14 | 3271936 | 3272306 | Distal Intergenic |
| chr14 | 3279989 | 3280385 | Distal Intergenic |
| chr14 | 3627463 | 3627812 | Distal Intergenic |
| chr14 | 3858840 | 3859146 | Distal Intergenic |
| chr14 | 3866766 | 3867066 | Distal Intergenic |
| chr14 | 4706341 | 4706651 | Distal Intergenic |
| chr14 | 5051181 | 5051512 | Distal Intergenic |
| chr14 | 7166690 | 7166931 | Distal Intergenic |
| chr14 | 7490510 | 7490995 | Distal Intergenic |
| chr14 | 8640784 | 8641587 | Distal Intergenic |
| chr14 | 8641711 | 8642664 | Distal Intergenic |
| chr14 | 8642858 | 8643070 | Distal Intergenic |
| chr14 | 8645119 | 8645340 | Distal Intergenic |
| chr14 | 8650575 | 8650821 | Distal Intergenic |
| chr14 | 8651067 | 8651606 | Distal Intergenic |
| chr14 | 8651643 | 8652349 | Distal Intergenic |
| chr14 | 8658763 | 8659529 | Distal Intergenic |
| chr14 | 8678558 | 8679560 | Distal Intergenic |
| chr14 | 8679648 | 8685294 | Distal Intergenic |
| chr14 | 8686033 | 8686757 | Distal Intergenic |
| chr14 | 8693306 | 8693662 | Distal Intergenic |
| chr14 | 8693848 | 8694688 | Distal Intergenic |
| chr14 | 8756931 | 8757377 | Distal Intergenic |
| chr14 | 8784726 | 8787861 | Distal Intergenic |
| chr14 | 8788059 | 8788571 | Distal Intergenic |
| chr14 | 8788952 | 8789480 | Distal Intergenic |
| chr14 | 8793780 | 8794438 | Distal Intergenic |
| chr14 | 8794482 | 8795048 | Distal Intergenic |
| chr14 | 8795117 | 8795339 | Distal Intergenic |
| chr14 | 8831036 | 8832088 | Distal Intergenic |
| chr14 | 8913090 | 8913289 | Distal Intergenic |
| chr14 | 8931055 | 8932810 | Distal Intergenic |
| chr14 | 8934298 | 8934532 | Distal Intergenic |

| | | | |
|---|---|---|---|
| chr14 | 8941525 | 8942063 | Distal Intergenic |
| chr14 | 8943464 | 8944757 | Distal Intergenic |
| chr14 | 8957278 | 8960014 | Distal Intergenic |
| chr14 | 8960293 | 8960798 | Distal Intergenic |
| chr14 | 9126631 | 9127324 | Distal Intergenic |
| chr14 | 9189610 | 9189809 | Distal Intergenic |
| chr14 | 9274953 | 9275687 | Distal Intergenic |
| chr14 | 9442678 | 9443070 | Distal Intergenic |
| chr14 | 10055759 | 10056738 | Distal Intergenic |
| chr14 | 10057411 | 10057628 | Distal Intergenic |
| chr14 | 10111020 | 10111235 | Distal Intergenic |
| chr14 | 10113774 | 10114736 | Distal Intergenic |
| chr14 | 10117063 | 10117302 | Distal Intergenic |
| chr14 | 10117688 | 10118135 | Distal Intergenic |
| chr14 | 10152073 | 10152811 | Distal Intergenic |
| chr14 | 10264645 | 10265711 | Distal Intergenic |
| chr14 | 10378754 | 10380648 | Distal Intergenic |
| chr14 | 10380701 | 10381094 | Distal Intergenic |
| chr14 | 10960934 | 10961391 | Distal Intergenic |
| chr14 | 10961497 | 10961918 | Distal Intergenic |
| chr14 | 12386307 | 12386665 | Distal Intergenic |
| chr14 | 12448311 | 12449055 | Distal Intergenic |
| chr14 | 12516090 | 12516404 | Distal Intergenic |
| chr14 | 12643452 | 12643899 | Distal Intergenic |
| chr14 | 12898197 | 12898773 | Distal Intergenic |
| chr14 | 12898989 | 12899720 | Distal Intergenic |
| chr14 | 13116745 | 13117510 | Distal Intergenic |
| chr14 | 13488888 | 13489311 | Distal Intergenic |
| chr14 | 14249025 | 14249927 | Distal Intergenic |
| chr14 | 14249986 | 14250185 | Distal Intergenic |
| chr14 | 14793167 | 14793386 | Distal Intergenic |
| chr14 | 15323303 | 15323582 | Distal Intergenic |
| chr14 | 15350545 | 15352737 | Distal Intergenic |
| chr14 | 15465541 | 15465740 | Distal Intergenic |
| chr14 | 16264665 | 16265565 | Distal Intergenic |
| chr14 | 17075900 | 17076648 | Distal Intergenic |
| chr14 | 17139966 | 17140529 | Distal Intergenic |
| chr14 | 17140668 | 17140957 | Distal Intergenic |
| chr14 | 17141006 | 17141406 | Distal Intergenic |
| chr14 | 17141451 | 17141763 | Distal Intergenic |
| chr14 | 17141941 | 17142569 | Distal Intergenic |
| chr14 | 17197671 | 17198072 | Distal Intergenic |
| chr14 | 17198269 | 17198763 | Distal Intergenic |
| chr14 | 17198825 | 17199099 | Distal Intergenic |

| | | | |
|---|---|---|---|
| chr14 | 17201295 | 17202211 | Distal Intergenic |
| chr14 | 17316426 | 17316695 | Distal Intergenic |
| chr14 | 17316743 | 17317112 | Distal Intergenic |
| chr14 | 17345559 | 17346066 | Distal Intergenic |
| chr14 | 17427125 | 17427356 | Distal Intergenic |
| chr14 | 17430522 | 17430721 | Distal Intergenic |
| chr14 | 17896015 | 17896467 | Distal Intergenic |
| chr14 | 19057814 | 19058024 | Distal Intergenic |
| chr14 | 19066075 | 19067299 | Distal Intergenic |
| chr14 | 19071413 | 19071612 | Distal Intergenic |
| chr14 | 19103020 | 19103257 | Distal Intergenic |
| chr14 | 19103311 | 19103569 | Distal Intergenic |
| chr14 | 19104091 | 19104330 | Distal Intergenic |
| chr14 | 19397092 | 19398345 | Distal Intergenic |
| chr14 | 19606306 | 19607473 | Exon |
| chr14 | 19680594 | 19681094 | Promoter |
| chr14 | 20305862 | 20306555 | Distal Intergenic |
| chr14 | 20787408 | 20788246 | Intron |
| chr14 | 20902145 | 20902344 | Intron |
| chr14 | 20902522 | 20902739 | Intron |
| chr14 | 21167034 | 21167329 | Intron |
| chr14 | 21167417 | 21167980 | 5' UTR |
| chr14 | 21296786 | 21299488 | Distal Intergenic |
| chr14 | 21322115 | 21322684 | Distal Intergenic |
| chr14 | 21364700 | 21365543 | Distal Intergenic |
| chr14 | 21365581 | 21365811 | Distal Intergenic |
| chr14 | 21428067 | 21429974 | Distal Intergenic |
| chr14 | 21450058 | 21450830 | Distal Intergenic |
| chr14 | 21460231 | 21460506 | 3' UTR |
| chr14 | 21492877 | 21493176 | Promoter |
| chr14 | 21493502 | 21494013 | Promoter |
| chr14 | 21494062 | 21494301 | Promoter |
| chr14 | 21521330 | 21521671 | Intron |
| chr14 | 21522042 | 21522750 | Intron |
| chr14 | 21623746 | 21624066 | Promoter |
| chr14 | 21624240 | 21626358 | Promoter |
| chr14 | 21646707 | 21647111 | Distal Intergenic |
| chr14 | 21647186 | 21647989 | Distal Intergenic |
| chr14 | 21648211 | 21648463 | Distal Intergenic |
| chr14 | 21651051 | 21651501 | Distal Intergenic |
| chr14 | 21652880 | 21653320 | Distal Intergenic |
| chr14 | 21655957 | 21656297 | Distal Intergenic |
| chr14 | 21663803 | 21667657 | Downstream |
| chr14 | 21669431 | 21670361 | Exon |

| | | | |
|---|---|---|---|
| chr14 | 21722326 | 21722536 | Intron |
| chr14 | 21749199 | 21749616 | Distal Intergenic |
| chr14 | 21757374 | 21758057 | Intron |
| chr14 | 21788543 | 21788849 | Promoter |
| chr14 | 21788901 | 21789818 | Promoter |
| chr14 | 21791813 | 21793458 | Promoter |
| chr14 | 21884859 | 21885573 | Intron |
| chr14 | 21905252 | 21907967 | Promoter |
| chr14 | 21954422 | 21954858 | Promoter |
| chr14 | 22170562 | 22170877 | Distal Intergenic |
| chr14 | 22366427 | 22367553 | Intron |
| chr14 | 22567184 | 22567393 | Intron |
| chr14 | 22567506 | 22567914 | Intron |
| chr14 | 22570237 | 22570637 | Intron |
| chr14 | 22649996 | 22650414 | Intron |
| chr14 | 22650465 | 22651338 | Intron |
| chr14 | 22884958 | 22885396 | Intron |
| chr14 | 23468661 | 23469995 | Promoter |
| chr14 | 23470323 | 23470584 | Promoter |
| chr14 | 24576084 | 24577649 | Intron |
| chr14 | 24577753 | 24577953 | Intron |
| chr14 | 25001638 | 25002037 | Distal Intergenic |
| chr14 | 25002073 | 25002798 | Distal Intergenic |
| chr14 | 25012121 | 25014460 | Distal Intergenic |
| chr14 | 25025252 | 25025805 | Distal Intergenic |
| chr14 | 25025971 | 25027251 | Distal Intergenic |
| chr14 | 25032111 | 25032414 | Distal Intergenic |
| chr14 | 25032505 | 25033342 | Distal Intergenic |
| chr14 | 25042209 | 25042434 | Downstream |
| chr14 | 25047067 | 25048487 | Distal Intergenic |
| chr14 | 25063332 | 25063553 | Distal Intergenic |
| chr14 | 25063624 | 25063990 | Distal Intergenic |
| chr14 | 25064055 | 25064718 | Distal Intergenic |
| chr14 | 25091544 | 25092290 | Distal Intergenic |
| chr14 | 25092480 | 25092735 | Distal Intergenic |
| chr14 | 25130613 | 25131037 | Distal Intergenic |
| chr14 | 25288022 | 25289015 | Promoter |
| chr14 | 25309895 | 25310103 | Intron |
| chr14 | 25851060 | 25851692 | Distal Intergenic |
| chr14 | 25957321 | 25957553 | Distal Intergenic |
| chr14 | 26006350 | 26006831 | Distal Intergenic |
| chr14 | 26182273 | 26182974 | Distal Intergenic |
| chr14 | 26184767 | 26185007 | Distal Intergenic |
| chr14 | 26185043 | 26185416 | Distal Intergenic |

| | | | |
|---|---|---|---|
| chr14 | 26185494 | 26185779 | Distal Intergenic |
| chr14 | 26197432 | 26198947 | Distal Intergenic |
| chr14 | 26200226 | 26200656 | Distal Intergenic |
| chr14 | 26233895 | 26235173 | Distal Intergenic |
| chr14 | 26278740 | 26279826 | Distal Intergenic |
| chr14 | 26280024 | 26280375 | Distal Intergenic |
| chr14 | 26280899 | 26281238 | Distal Intergenic |
| chr14 | 26283866 | 26284226 | Distal Intergenic |
| chr14 | 26284300 | 26284536 | Distal Intergenic |
| chr14 | 26293502 | 26297179 | Distal Intergenic |
| chr14 | 26306246 | 26306818 | Distal Intergenic |
| chr14 | 26346290 | 26346712 | Distal Intergenic |
| chr14 | 26346963 | 26347380 | Distal Intergenic |
| chr14 | 26353590 | 26353852 | Distal Intergenic |
| chr14 | 26405723 | 26409397 | Distal Intergenic |
| chr14 | 26418699 | 26420741 | Distal Intergenic |
| chr14 | 26425849 | 26426166 | Distal Intergenic |
| chr14 | 26446658 | 26447812 | Distal Intergenic |
| chr14 | 26449380 | 26449590 | Distal Intergenic |
| chr14 | 26449759 | 26450107 | Distal Intergenic |
| chr14 | 26455502 | 26456768 | Distal Intergenic |
| chr14 | 26520199 | 26520752 | Distal Intergenic |
| chr14 | 26520869 | 26521127 | Distal Intergenic |
| chr14 | 26662051 | 26662458 | Distal Intergenic |
| chr14 | 26681210 | 26681886 | Distal Intergenic |
| chr14 | 26746798 | 26747112 | Distal Intergenic |
| chr14 | 26886492 | 26886745 | Distal Intergenic |
| chr14 | 27326806 | 27327349 | Distal Intergenic |
| chr14 | 27344340 | 27344574 | Distal Intergenic |
| chr14 | 27352730 | 27352984 | Distal Intergenic |
| chr14 | 27353050 | 27353490 | Distal Intergenic |
| chr14 | 27354126 | 27354645 | Distal Intergenic |
| chr14 | 27360134 | 27361013 | Distal Intergenic |
| chr14 | 27398747 | 27398946 | Distal Intergenic |
| chr14 | 27399440 | 27399668 | Distal Intergenic |
| chr14 | 27457144 | 27457544 | Distal Intergenic |
| chr14 | 27457704 | 27459008 | Distal Intergenic |
| chr14 | 27488641 | 27489631 | Distal Intergenic |
| chr14 | 27882732 | 27883544 | Intron |
| chr14 | 28062463 | 28062821 | Intron |
| chr14 | 28062858 | 28063124 | Intron |
| chr14 | 28112989 | 28113331 | Intron |
| chr14 | 28114291 | 28114531 | Intron |
| chr14 | 28115024 | 28115390 | Intron |

| | | | |
|---|---|---|---|
| chr14 | 28154311 | 28155005 | Distal Intergenic |
| chr14 | 28160228 | 28160853 | Distal Intergenic |
| chr14 | 28163880 | 28165311 | Distal Intergenic |
| chr14 | 28188976 | 28190025 | Distal Intergenic |
| chr14 | 28242030 | 28242229 | Distal Intergenic |
| chr14 | 28242368 | 28243027 | Distal Intergenic |
| chr14 | 28747161 | 28747382 | Distal Intergenic |
| chr14 | 29182231 | 29184843 | Distal Intergenic |
| chr14 | 29243778 | 29243992 | Intron |
| chr14 | 29319019 | 29319474 | Distal Intergenic |
| chr14 | 29319604 | 29319950 | Distal Intergenic |
| chr14 | 29320056 | 29320349 | Distal Intergenic |
| chr14 | 29320648 | 29321122 | Distal Intergenic |
| chr14 | 29338365 | 29338893 | Distal Intergenic |
| chr14 | 29355465 | 29356021 | Distal Intergenic |
| chr14 | 29357585 | 29358041 | Distal Intergenic |
| chr14 | 29358299 | 29358603 | Distal Intergenic |
| chr14 | 29358853 | 29359327 | Distal Intergenic |
| chr14 | 29375658 | 29377249 | Distal Intergenic |
| chr14 | 29377363 | 29377580 | Distal Intergenic |
| chr14 | 29621444 | 29621645 | Distal Intergenic |
| chr14 | 29885712 | 29886401 | Distal Intergenic |
| chr14 | 29893153 | 29895183 | Downstream |
| chr14 | 29895411 | 29896060 | Downstream |
| chr14 | 30082050 | 30082291 | Intron |
| chr14 | 30095697 | 30097446 | Exon |
| chr14 | 30309996 | 30310303 | Intron |
| chr14 | 30310418 | 30310664 | Intron |
| chr14 | 30781509 | 30782191 | Distal Intergenic |
| chr14 | 31292827 | 31293182 | Distal Intergenic |
| chr14 | 31316345 | 31317276 | Distal Intergenic |
| chr14 | 31344055 | 31344254 | Promoter |
| chr14 | 31344657 | 31346426 | Promoter |
| chr14 | 31346504 | 31346752 | Intron |
| chr14 | 31362038 | 31362589 | Intron |
| chr14 | 31374463 | 31375422 | Exon |
| chr14 | 31406429 | 31406953 | Intron |
| chr14 | 31407032 | 31407296 | Intron |
| chr14 | 31409240 | 31409748 | Intron |
| chr14 | 31434084 | 31436178 | Intron |
| chr14 | 31436241 | 31436458 | Intron |
| chr14 | 31436602 | 31437585 | Intron |
| chr14 | 31437786 | 31438744 | Intron |
| chr14 | 31438828 | 31439036 | Intron |

| | | | |
|---|---|---|---|
| chr14 | 31439226 | 31439454 | Intron |
| chr14 | 31787190 | 31787716 | Exon |
| chr14 | 31813970 | 31814529 | Exon |
| chr14 | 31831786 | 31832243 | Intron |
| chr14 | 31952871 | 31953321 | 5' UTR |
| chr14 | 31967054 | 31967383 | Distal Intergenic |
| chr14 | 31973005 | 31974382 | Distal Intergenic |
| chr14 | 31978104 | 31980242 | Distal Intergenic |
| chr14 | 32011016 | 32011659 | Distal Intergenic |
| chr14 | 32011695 | 32012109 | Distal Intergenic |
| chr14 | 32018324 | 32019488 | Distal Intergenic |
| chr14 | 32217162 | 32218189 | Intron |
| chr14 | 32261111 | 32261581 | Intron |
| chr14 | 32261969 | 32262180 | Intron |
| chr14 | 32308044 | 32309213 | Intron |
| chr14 | 32475023 | 32475689 | Distal Intergenic |
| chr14 | 32475788 | 32476159 | Distal Intergenic |
| chr14 | 32482220 | 32482508 | Distal Intergenic |
| chr14 | 32482709 | 32483849 | Distal Intergenic |
| chr14 | 32594664 | 32595193 | Intron |
| chr14 | 32757904 | 32758122 | Distal Intergenic |
| chr14 | 32758351 | 32758550 | Distal Intergenic |
| chr14 | 32758690 | 32759391 | Distal Intergenic |
| chr14 | 32759429 | 32759749 | Distal Intergenic |
| chr14 | 32898201 | 32898436 | Intron |
| chr14 | 32899686 | 32899885 | Intron |
| chr14 | 32976952 | 32978972 | Intron |
| chr14 | 33014313 | 33015285 | Exon |
| chr14 | 33306266 | 33306488 | Distal Intergenic |
| chr14 | 34163951 | 34164167 | Intron |
| chr14 | 34164242 | 34166060 | Intron |
| chr14 | 34537113 | 34538560 | Distal Intergenic |
| chr14 | 34637192 | 34638824 | Distal Intergenic |
| chr14 | 34643020 | 34643535 | Distal Intergenic |
| chr14 | 34898297 | 34901407 | Downstream |
| chr14 | 35110295 | 35110532 | Distal Intergenic |
| chr14 | 35124633 | 35124851 | Distal Intergenic |
| chr14 | 35175838 | 35177333 | Downstream |
| chr14 | 35186889 | 35188320 | Distal Intergenic |
| chr14 | 35188640 | 35190519 | Distal Intergenic |
| chr14 | 35191754 | 35192566 | Distal Intergenic |
| chr14 | 35280785 | 35280984 | Intron |
| chr14 | 35314820 | 35315139 | Intron |
| chr14 | 35316003 | 35318139 | Intron |

| | | | |
|---|---|---|---|
| chr14 | 35341040 | 35341246 | Intron |
| chr14 | 35361512 | 35362664 | Distal Intergenic |
| chr14 | 35481641 | 35481990 | Intron |
| chr14 | 35487226 | 35488315 | Exon |
| chr14 | 37512079 | 37512730 | Intron |
| chr14 | 37512782 | 37513180 | Intron |
| chr14 | 37698173 | 37698627 | Intron |
| chr14 | 37698828 | 37699344 | Intron |
| chr14 | 37745958 | 37746199 | Intron |
| chr14 | 37948279 | 37948506 | Intron |
| chr14 | 41716100 | 41716494 | Distal Intergenic |
| chr14 | 41760713 | 41764278 | Distal Intergenic |
| chr14 | 41768981 | 41770371 | Distal Intergenic |
| chr14 | 41808765 | 41810151 | Distal Intergenic |
| chr14 | 41844501 | 41844918 | Distal Intergenic |
| chr14 | 44448973 | 44449474 | Distal Intergenic |
| chr14 | 45660444 | 45660757 | Intron |
| chr14 | 45810343 | 45810881 | Distal Intergenic |
| chr14 | 45810919 | 45811342 | Distal Intergenic |
| chr14 | 45949522 | 45949999 | Distal Intergenic |
| chr14 | 46062401 | 46062871 | Distal Intergenic |
| chr14 | 46098758 | 46099097 | Distal Intergenic |
| chr14 | 46111971 | 46112492 | Distal Intergenic |
| chr14 | 46791860 | 46792691 | Intron |
| chr14 | 46792750 | 46793081 | Intron |
| chr14 | 47136432 | 47136631 | Distal Intergenic |
| chr14 | 47137242 | 47137462 | Distal Intergenic |
| chr14 | 47256266 | 47256465 | Distal Intergenic |
| chr14 | 47407372 | 47407587 | Intron |
| chr14 | 47410115 | 47410810 | Intron |
| chr14 | 47420204 | 47421474 | Intron |
| chr14 | 47585565 | 47586062 | Intron |
| chr14 | 47586166 | 47586418 | Intron |
| chr14 | 47641704 | 47642653 | Intron |
| chr14 | 47896648 | 47897269 | Intron |
| chr14 | 47917862 | 47918264 | Intron |
| chr14 | 47918436 | 47918858 | Intron |
| chr14 | 47970664 | 47971239 | Intron |
| chr14 | 47973525 | 47974060 | Intron |
| chr14 | 47994016 | 47994215 | Intron |
| chr14 | 48138526 | 48138739 | Intron |
| chr14 | 48152217 | 48153438 | Distal Intergenic |
| chr14 | 48235402 | 48236193 | Exon |
| chr14 | 48244195 | 48245959 | Intron |

| | | | |
|---|---|---|---|
| chr14 | 48253655 | 48254647 | Intron |
| chr14 | 48771979 | 48772231 | Distal Intergenic |
| chr14 | 49685068 | 49685715 | Distal Intergenic |
| chr14 | 49686143 | 49687056 | Distal Intergenic |
| chr14 | 49692433 | 49694445 | Distal Intergenic |
| chr14 | 49694505 | 49694704 | Distal Intergenic |
| chr14 | 51543625 | 51543993 | Intron |
| chr14 | 51544077 | 51544365 | Intron |
| chr14 | 51550844 | 51552050 | Intron |
| chr14 | 51552550 | 51553011 | Intron |
| chr14 | 51563625 | 51563919 | Distal Intergenic |
| chr14 | 51564172 | 51564381 | Distal Intergenic |
| chr14 | 51564465 | 51566896 | Distal Intergenic |
| chr14 | 51576894 | 51578160 | Distal Intergenic |
| chr14 | 51710450 | 51710649 | 5' UTR |
| chr14 | 51710866 | 51712762 | 5' UTR |
| chr14 | 51713108 | 51714394 | Exon |
| chr14 | 51720554 | 51721069 | Exon |
| chr14 | 51818747 | 51819084 | Intron |
| chr14 | 51819271 | 51820286 | Intron |
| chr14 | 52477766 | 52477966 | Intron |
| chr14 | 52485018 | 52487139 | Exon |
| chr14 | 52490558 | 52490821 | Intron |
| chr14 | 52552104 | 52552893 | Distal Intergenic |
| chr14 | 52579942 | 52580169 | Distal Intergenic |
| chr14 | 52633359 | 52633772 | Distal Intergenic |
| chr14 | 52634496 | 52636104 | Distal Intergenic |
| chr14 | 52639079 | 52639370 | Distal Intergenic |
| chr14 | 52639541 | 52639785 | Distal Intergenic |
| chr14 | 52639842 | 52640481 | Distal Intergenic |
| chr14 | 52640525 | 52640880 | Distal Intergenic |
| chr14 | 52642834 | 52643206 | Distal Intergenic |
| chr14 | 52723365 | 52723688 | Distal Intergenic |
| chr14 | 52816238 | 52816689 | Distal Intergenic |
| chr14 | 52816730 | 52817557 | Distal Intergenic |
| chr14 | 52899187 | 52899438 | Exon |
| chr14 | 52924228 | 52925007 | Intron |
| chr14 | 54841854 | 54843010 | Distal Intergenic |
| chr14 | 55049949 | 55050557 | Intron |
| chr14 | 55050967 | 55054573 | Intron |
| chr14 | 55082572 | 55082919 | Intron |
| chr14 | 55111136 | 55112041 | Intron |
| chr14 | 55191023 | 55191491 | Intron |
| chr14 | 55191543 | 55191849 | Intron |

| | | | |
|---|---|---|---|
| chr14 | 55192067 | 55194217 | Intron |
| chr14 | 55194261 | 55194606 | Intron |
| chr14 | 55194958 | 55195205 | Intron |
| chr14 | 55195297 | 55196081 | Intron |
| chr14 | 55196310 | 55196650 | Intron |
| chr14 | 55236321 | 55237075 | Exon |
| chr14 | 55304915 | 55305696 | Distal Intergenic |
| chr14 | 55349052 | 55349531 | Intron |
| chr14 | 55349605 | 55350013 | Intron |
| chr14 | 55385661 | 55385986 | Distal Intergenic |
| chr14 | 55386137 | 55388464 | Distal Intergenic |
| chr14 | 55501822 | 55502166 | Intron |
| chr14 | 55502359 | 55502734 | Intron |
| chr14 | 56209441 | 56209794 | Distal Intergenic |
| chr14 | 56209841 | 56210449 | Distal Intergenic |
| chr14 | 56254573 | 56254773 | Intron |
| chr14 | 56261721 | 56262665 | Promoter |
| chr14 | 56290227 | 56290489 | Distal Intergenic |
| chr14 | 56290529 | 56291172 | Distal Intergenic |
| chr14 | 56291234 | 56291509 | Distal Intergenic |
| chr14 | 56333624 | 56334868 | Distal Intergenic |
| chr14 | 56342807 | 56343023 | Distal Intergenic |
| chr14 | 56343063 | 56343296 | Distal Intergenic |
| chr14 | 56343606 | 56343824 | Distal Intergenic |
| chr14 | 56348910 | 56349119 | Distal Intergenic |
| chr14 | 56381967 | 56382166 | Distal Intergenic |
| chr14 | 56383810 | 56385023 | Distal Intergenic |
| chr14 | 56386342 | 56387794 | Distal Intergenic |
| chr14 | 57431141 | 57431671 | Distal Intergenic |
| chr14 | 57494034 | 57494815 | Distal Intergenic |
| chr14 | 57504543 | 57504777 | Distal Intergenic |
| chr14 | 57506016 | 57506483 | Distal Intergenic |
| chr14 | 57506606 | 57506892 | Distal Intergenic |
| chr14 | 57724709 | 57724997 | Intron |
| chr14 | 57757001 | 57757556 | Distal Intergenic |
| chr14 | 57757806 | 57758298 | Distal Intergenic |
| chr14 | 57779775 | 57779983 | Distal Intergenic |
| chr14 | 57910117 | 57910876 | Distal Intergenic |
| chr14 | 57914276 | 57914758 | Distal Intergenic |
| chr14 | 57915217 | 57915490 | Distal Intergenic |
| chr14 | 57916148 | 57916735 | Distal Intergenic |
| chr14 | 57916995 | 57917214 | Distal Intergenic |
| chr14 | 58185263 | 58185479 | Intron |
| chr14 | 58283063 | 58283614 | Intron |

| | | | |
|---|---|---|---|
| chr14 | 58284075 | 58284311 | Intron |
| chr14 | 58324188 | 58324690 | Intron |
| chr14 | 58325954 | 58327235 | Intron |
| chr14 | 58327350 | 58327631 | Intron |
| chr14 | 58364540 | 58364924 | Distal Intergenic |
| chr14 | 58469047 | 58470183 | Downstream |
| chr14 | 58501839 | 58502094 | Intron |
| chr14 | 58513137 | 58513443 | Intron |
| chr14 | 59406062 | 59406345 | Distal Intergenic |
| chr14 | 59406383 | 59406975 | Distal Intergenic |
| chr14 | 59407022 | 59407542 | Distal Intergenic |
| chr14 | 60870444 | 60870692 | Distal Intergenic |
| chr14 | 61344915 | 61345979 | Intron |
| chr14 | 61943474 | 61943815 | Promoter |
| chr14 | 61973360 | 61973608 | Intron |
| chr14 | 61973829 | 61974292 | Intron |
| chr14 | 62040019 | 62040246 | Intron |
| chr14 | 62128341 | 62129490 | Distal Intergenic |
| chr14 | 62129526 | 62129726 | Distal Intergenic |
| chr14 | 62214851 | 62218184 | Promoter |
| chr14 | 62218670 | 62218924 | Distal Intergenic |
| chr14 | 62220976 | 62221200 | Distal Intergenic |
| chr14 | 62221355 | 62221767 | Distal Intergenic |
| chr14 | 62221854 | 62222112 | Distal Intergenic |
| chr14 | 62264303 | 62264559 | Distal Intergenic |
| chr14 | 62265672 | 62266873 | Distal Intergenic |
| chr14 | 62267375 | 62267685 | Distal Intergenic |
| chr14 | 63335823 | 63336448 | Intron |
| chr14 | 63379019 | 63380010 | Intron |
| chr14 | 63443995 | 63444278 | Intron |
| chr14 | 63444516 | 63445454 | Intron |
| chr14 | 63559349 | 63559781 | Intron |
| chr14 | 63741558 | 63741784 | Intron |
| chr14 | 63742014 | 63742628 | Intron |
| chr14 | 63750225 | 63750463 | Intron |
| chr14 | 63750616 | 63751668 | Intron |
| chr14 | 63767320 | 63768154 | Distal Intergenic |
| chr14 | 64022389 | 64022649 | Distal Intergenic |
| chr14 | 64046130 | 64046385 | Distal Intergenic |
| chr14 | 64046433 | 64046834 | Distal Intergenic |
| chr14 | 64065678 | 64067537 | 5' UTR |
| chr14 | 64069555 | 64073893 | Intron |
| chr14 | 64535703 | 64536686 | Intron |
| chr14 | 64596906 | 64597307 | Exon |

| | | | |
|---|---|---|---|
| chr14 | 64597345 | 64598246 | Intron |
| chr14 | 64598314 | 64599377 | Exon |
| chr14 | 64601544 | 64601992 | Intron |
| chr14 | 64616684 | 64617011 | Intron |
| chr14 | 64880056 | 64880998 | Intron |
| chr14 | 64882610 | 64883552 | Intron |
| chr14 | 64968569 | 64969242 | Intron |
| chr14 | 65059008 | 65059411 | Distal Intergenic |
| chr14 | 65059465 | 65059771 | Distal Intergenic |
| chr14 | 65059983 | 65060397 | Distal Intergenic |
| chr14 | 65578337 | 65578865 | Distal Intergenic |
| chr14 | 65601282 | 65601682 | Distal Intergenic |
| chr14 | 65716178 | 65716548 | Distal Intergenic |
| chr14 | 65719237 | 65720580 | Distal Intergenic |
| chr14 | 65731628 | 65732086 | Distal Intergenic |
| chr14 | 65732247 | 65732855 | Distal Intergenic |
| chr14 | 66157395 | 66157603 | Intron |
| chr14 | 66158386 | 66158641 | Intron |
| chr14 | 66486072 | 66487167 | Distal Intergenic |
| chr14 | 66646921 | 66647194 | Distal Intergenic |
| chr14 | 66723871 | 66725341 | Distal Intergenic |
| chr14 | 66849500 | 66849699 | Distal Intergenic |
| chr14 | 66875975 | 66878539 | Distal Intergenic |
| chr14 | 66885224 | 66885585 | Distal Intergenic |
| chr14 | 67595453 | 67596089 | Intron |
| chr14 | 67596161 | 67596454 | Intron |
| chr14 | 67665658 | 67666444 | Exon |
| chr14 | 67666492 | 67666753 | Exon |
| chr14 | 67669500 | 67669727 | Intron |
| chr14 | 67716004 | 67716384 | Intron |
| chr14 | 67716538 | 67717226 | Intron |
| chr14 | 68334675 | 68335188 | Intron |
| chr14 | 68335258 | 68335469 | Intron |
| chr14 | 68495503 | 68495883 | Intron |
| chr14 | 68496119 | 68496449 | Intron |
| chr14 | 68541626 | 68542107 | Intron |
| chr14 | 68638195 | 68638579 | Intron |
| chr14 | 68653069 | 68653300 | Intron |
| chr14 | 68653533 | 68653907 | Intron |
| chr14 | 69264690 | 69264927 | Distal Intergenic |
| chr14 | 69885090 | 69886453 | Intron |
| chr14 | 70212295 | 70212564 | Intron |
| chr14 | 70212662 | 70212940 | Intron |
| chr14 | 70213024 | 70213461 | Intron |

| | | | |
|---|---|---|---|
| chr14 | 70215547 | 70215985 | Intron |
| chr14 | 70216051 | 70216267 | Intron |
| chr14 | 70586958 | 70587159 | Intron |
| chr14 | 70853775 | 70854366 | Intron |
| chr14 | 70897479 | 70898747 | Distal Intergenic |
| chr14 | 70900393 | 70901506 | Distal Intergenic |
| chr14 | 70903861 | 70904060 | Distal Intergenic |
| chr14 | 70915931 | 70918468 | Promoter |
| chr14 | 70954848 | 70955094 | Distal Intergenic |
| chr14 | 70974454 | 70975419 | Distal Intergenic |
| chr14 | 70975470 | 70975683 | Distal Intergenic |
| chr14 | 70976460 | 70979002 | Distal Intergenic |
| chr14 | 70979898 | 70980397 | Distal Intergenic |
| chr14 | 70983701 | 70986137 | Downstream |
| chr14 | 70999160 | 70999421 | Intron |
| chr14 | 71166837 | 71167185 | Distal Intergenic |
| chr14 | 71172349 | 71173374 | Distal Intergenic |
| chr14 | 71364860 | 71366421 | Distal Intergenic |
| chr14 | 71482444 | 71482686 | Intron |
| chr14 | 71630176 | 71630376 | Distal Intergenic |
| chr14 | 72195609 | 72196829 | Exon |
| chr14 | 73076051 | 73076665 | Exon |
| chr14 | 73125143 | 73125673 | Intron |
| chr14 | 73183991 | 73184235 | Intron |
| chr14 | 73184291 | 73184835 | Intron |
| chr14 | 73234716 | 73235112 | Intron |
| chr14 | 73235253 | 73235557 | Intron |
| chr14 | 73286622 | 73286979 | Intron |
| chr14 | 73323643 | 73324123 | Intron |
| chr14 | 73324756 | 73325142 | Intron |
| chr14 | 73510042 | 73511244 | Distal Intergenic |
| chr14 | 73569474 | 73570993 | Exon |
| chr14 | 73580731 | 73580964 | Intron |
| chr14 | 73633790 | 73636012 | Intron |
| chr14 | 73637075 | 73639271 | Exon |
| chr14 | 73639307 | 73639685 | Intron |
| chr14 | 73653466 | 73653852 | Exon |
| chr14 | 73768611 | 73772370 | Intron |
| chr14 | 73782892 | 73783100 | Exon |
| chr14 | 73783531 | 73785089 | Intron |
| chr14 | 73785224 | 73786110 | Intron |
| chr14 | 73787866 | 73789039 | Intron |
| chr14 | 73794350 | 73794658 | Intron |
| chr14 | 73796854 | 73797245 | Intron |

| | | | |
|---|---|---|---|
| chr14 | 73802098 | 73802741 | Intron |
| chr14 | 73809546 | 73810057 | Intron |
| chr14 | 73810139 | 73810438 | Intron |
| chr14 | 73845097 | 73846300 | Intron |
| chr14 | 73848208 | 73850538 | Intron |
| chr14 | 73850796 | 73851422 | Intron |
| chr14 | 73860292 | 73860678 | Intron |
| chr14 | 73883518 | 73884146 | Intron |
| chr14 | 73885854 | 73886535 | Intron |
| chr14 | 73896200 | 73896451 | Intron |
| chr14 | 73907510 | 73908016 | Intron |
| chr14 | 74107465 | 74108146 | Distal Intergenic |
| chr14 | 74288132 | 74288509 | Distal Intergenic |
| chr14 | 74302332 | 74302531 | Distal Intergenic |
| chr14 | 74418029 | 74418237 | Promoter |
| chr14 | 74498963 | 74499209 | Promoter |
| chr14 | 74663798 | 74664512 | Intron |
| chr14 | 74727468 | 74727814 | 3' UTR |
| chr14 | 75132043 | 75133862 | Intron |
| chr14 | 75136417 | 75137144 | Exon |
| chr14 | 75184245 | 75184605 | Intron |
| chr14 | 75187325 | 75187578 | Intron |
| chr14 | 75187701 | 75188010 | Intron |
| chr14 | 75222325 | 75223897 | Distal Intergenic |
| chr14 | 75280474 | 75281570 | Intron |
| chr14 | 75282737 | 75283344 | Exon |
| chr14 | 75283501 | 75285054 | Exon |
| chr14 | 75310073 | 75310886 | Distal Intergenic |
| chr14 | 75345253 | 75346949 | Distal Intergenic |
| chr14 | 75396127 | 75396919 | Distal Intergenic |
| chr14 | 75445162 | 75445503 | Distal Intergenic |
| chr14 | 75653688 | 75653942 | Distal Intergenic |
| chr14 | 76233815 | 76235013 | Intron |
| chr14 | 76244055 | 76244456 | Intron |
| chr14 | 76337318 | 76338404 | Intron |
| chr14 | 76338558 | 76339489 | Intron |
| chr14 | 76349895 | 76350784 | Intron |
| chr14 | 76354108 | 76355123 | Intron |
| chr14 | 76410100 | 76410675 | Intron |
| chr14 | 76483768 | 76484429 | Intron |
| chr14 | 76499497 | 76500880 | Intron |
| chr14 | 76510895 | 76511158 | Intron |
| chr14 | 76678246 | 76678913 | Distal Intergenic |
| chr14 | 76679117 | 76679507 | Distal Intergenic |

| | | | |
|---|---|---|---|
| chr14 | 76878425 | 76879527 | Intron |
| chr14 | 76904667 | 76905395 | Intron |
| chr14 | 76905440 | 76905683 | Intron |
| chr14 | 76934734 | 76935335 | Intron |
| chr14 | 77052528 | 77052727 | Distal Intergenic |
| chr14 | 77064460 | 77065323 | Distal Intergenic |
| chr14 | 77065521 | 77065957 | Distal Intergenic |
| chr14 | 77141781 | 77142783 | Distal Intergenic |
| chr14 | 77155160 | 77156085 | Distal Intergenic |
| chr14 | 77161017 | 77162211 | Distal Intergenic |
| chr14 | 77187048 | 77187288 | Distal Intergenic |
| chr14 | 77346635 | 77347476 | Distal Intergenic |
| chr14 | 77351147 | 77351750 | Distal Intergenic |
| chr14 | 77436837 | 77438174 | Distal Intergenic |
| chr14 | 78419993 | 78420550 | Distal Intergenic |
| chr14 | 79335184 | 79335785 | Intron |
| chr14 | 79734015 | 79734745 | Intron |
| chr14 | 79735032 | 79736588 | Intron |
| chr14 | 79789424 | 79790038 | Intron |
| chr14 | 79881576 | 79882009 | Intron |
| chr14 | 79903192 | 79903832 | Intron |
| chr14 | 79903918 | 79904163 | Intron |
| chr14 | 79915085 | 79916774 | Intron |
| chr14 | 79916855 | 79917304 | Intron |
| chr14 | 79926259 | 79928727 | Intron |
| chr14 | 79933353 | 79933634 | Exon |
| chr14 | 79934108 | 79934484 | Intron |
| chr14 | 79934548 | 79935341 | Intron |
| chr14 | 79949069 | 79950060 | Intron |
| chr14 | 79951674 | 79952094 | Intron |
| chr14 | 79986839 | 79987038 | Intron |
| chr14 | 80883874 | 80884247 | Intron |
| chr14 | 80884337 | 80884539 | Intron |
| chr14 | 82275560 | 82276155 | Distal Intergenic |
| chr14 | 87451321 | 87451533 | Distal Intergenic |
| chr14 | 87540056 | 87540902 | Distal Intergenic |
| chr14 | 90622426 | 90623002 | Intron |
| chr14 | 92248528 | 92249078 | 3' UTR |
| chr14 | 98870738 | 98871985 | Distal Intergenic |
| chr14 | 98881886 | 98882717 | Distal Intergenic |
| chr14 | 98882835 | 98883087 | Distal Intergenic |
| chr14 | 99436909 | 99437294 | Distal Intergenic |
| chr14 | 99573599 | 99574198 | Distal Intergenic |
| chr14 | 99698834 | 99699275 | Intron |

| | | | |
|---|---|---|---|
| chr14 | 99699403 | 99700109 | Intron |
| chr14 | 99700211 | 99700763 | Intron |
| chr14 | 99723924 | 99724524 | Exon |
| chr14 | 99772877 | 99773076 | Distal Intergenic |
| chr14 | 99848480 | 99849324 | Distal Intergenic |
| chr14 | 99865036 | 99865858 | Exon |
| chr14 | 99947641 | 99947943 | Promoter |
| chr14 | 99947980 | 99948217 | Promoter |
| chr14 | 99977738 | 99978547 | 3' UTR |
| chr14 | 99978713 | 99978999 | 3' UTR |
| chr14 | 99979168 | 99979674 | 3' UTR |
| chr14 | 100023630 | 100024969 | Intron |
| chr14 | 100030171 | 100032594 | Intron |
| chr14 | 101274713 | 101274962 | Distal Intergenic |
| chr14 | 102269828 | 102270032 | Intron |
| chr14 | 105666813 | 105668151 | Distal Intergenic |
| chr14 | 105727371 | 105728688 | Intron |
| chr14 | 105740730 | 105740988 | Intron |
| chr14 | 105768720 | 105769739 | Intron |
| chr14 | 105776672 | 105777057 | Intron |
| chr14 | 106561625 | 106561948 | Exon |
| chr14 | 110925971 | 110927438 | Distal Intergenic |
| chr14 | 111103084 | 111103354 | Distal Intergenic |
| chr14 | 111103430 | 111103823 | Distal Intergenic |
| chr14 | 111569764 | 111570171 | Distal Intergenic |
| chr14 | 111570211 | 111570575 | Distal Intergenic |
| chr14 | 116700646 | 116700881 | Distal Intergenic |
| chr14 | 117328065 | 117328285 | Distal Intergenic |
| chr14 | 118785196 | 118785576 | Distal Intergenic |
| chr14 | 118792579 | 118793628 | Distal Intergenic |
| chr14 | 118901579 | 118902351 | Distal Intergenic |
| chr14 | 118965013 | 118965741 | Distal Intergenic |
| chr14 | 118965862 | 118966175 | Distal Intergenic |
| chr14 | 120675461 | 120677239 | Distal Intergenic |
| chr14 | 120677709 | 120677929 | Distal Intergenic |
| chr14 | 120691051 | 120692433 | Distal Intergenic |
| chr14 | 120705512 | 120705871 | Distal Intergenic |
| chr14 | 120705977 | 120706213 | Distal Intergenic |
| chr14 | 120706394 | 120707696 | Distal Intergenic |
| chr14 | 120717420 | 120718041 | Distal Intergenic |
| chr14 | 120720114 | 120721490 | Distal Intergenic |
| chr14 | 120721563 | 120723091 | Distal Intergenic |
| chr14 | 120723324 | 120723536 | Distal Intergenic |
| chr14 | 120740413 | 120742248 | Distal Intergenic |

| | | | |
|---|---|---|---|
| chr14 | 120771865 | 120773026 | Distal Intergenic |
| chr14 | 120773113 | 120773346 | Distal Intergenic |
| chr14 | 121245121 | 121245852 | Distal Intergenic |
| chr14 | 121324874 | 121325085 | Distal Intergenic |
| chr14 | 121755664 | 121756148 | Distal Intergenic |
| chr14 | 122010600 | 122011536 | Distal Intergenic |
| chr14 | 122108996 | 122109394 | Distal Intergenic |
| chr14 | 122109528 | 122109730 | Distal Intergenic |
| chr14 | 122109914 | 122110911 | Distal Intergenic |
| chr14 | 122111373 | 122112121 | Distal Intergenic |
| chr14 | 122112337 | 122113393 | Distal Intergenic |
| chr14 | 122197369 | 122197603 | Distal Intergenic |
| chr14 | 122221251 | 122221897 | Distal Intergenic |
| chr14 | 122221952 | 122222567 | Distal Intergenic |
| chr14 | 122223885 | 122224094 | Distal Intergenic |
| chr14 | 122224367 | 122225557 | Distal Intergenic |
| chr14 | 122243608 | 122243920 | Distal Intergenic |
| chr14 | 122504393 | 122504605 | Distal Intergenic |
| chr14 | 122505726 | 122505974 | Distal Intergenic |
| chr14 | 122582902 | 122583378 | Distal Intergenic |
| chr14 | 124061419 | 124061827 | Distal Intergenic |
| chr14 | 124061896 | 124062148 | Distal Intergenic |
| chr15 | 3087090 | 3087417 | Distal Intergenic |
| chr15 | 3929011 | 3929216 | Distal Intergenic |
| chr15 | 3929522 | 3929945 | Distal Intergenic |
| chr15 | 3950196 | 3950436 | Distal Intergenic |
| chr15 | 3972403 | 3972745 | Distal Intergenic |
| chr15 | 3973379 | 3974715 | Distal Intergenic |
| chr15 | 4117894 | 4118160 | Distal Intergenic |
| chr15 | 4118227 | 4118473 | Distal Intergenic |
| chr15 | 4118620 | 4119850 | Distal Intergenic |
| chr15 | 5065587 | 5066318 | Distal Intergenic |
| chr15 | 5066619 | 5067480 | Distal Intergenic |
| chr15 | 5067672 | 5067925 | Distal Intergenic |
| chr15 | 5135488 | 5135692 | Distal Intergenic |
| chr15 | 5144510 | 5144949 | Distal Intergenic |
| chr15 | 5145874 | 5146706 | Distal Intergenic |
| chr15 | 5148742 | 5148947 | Distal Intergenic |
| chr15 | 5408838 | 5409850 | Distal Intergenic |
| chr15 | 5409897 | 5410366 | Distal Intergenic |
| chr15 | 5411183 | 5412700 | Distal Intergenic |
| chr15 | 5487213 | 5487528 | Distal Intergenic |
| chr15 | 5489167 | 5489779 | Distal Intergenic |
| chr15 | 6009242 | 6009441 | Distal Intergenic |

| | | | |
|---|---|---|---|
| chr15 | 6183507 | 6184174 | Distal Intergenic |
| chr15 | 6263356 | 6264394 | Distal Intergenic |
| chr15 | 6334023 | 6334869 | Distal Intergenic |
| chr15 | 6442636 | 6442872 | Distal Intergenic |
| chr15 | 6514990 | 6515470 | Distal Intergenic |
| chr15 | 6524164 | 6525193 | Distal Intergenic |
| chr15 | 6675568 | 6675857 | Distal Intergenic |
| chr15 | 6675997 | 6676411 | Distal Intergenic |
| chr15 | 6823729 | 6824003 | Distal Intergenic |
| chr15 | 6824146 | 6825069 | Distal Intergenic |
| chr15 | 6876925 | 6878613 | Distal Intergenic |
| chr15 | 6901022 | 6901223 | Distal Intergenic |
| chr15 | 6945574 | 6945784 | Distal Intergenic |
| chr15 | 7547043 | 7547253 | Distal Intergenic |
| chr15 | 7547445 | 7548196 | Distal Intergenic |
| chr15 | 7548248 | 7548601 | Distal Intergenic |
| chr15 | 7628687 | 7629504 | Distal Intergenic |
| chr15 | 7630648 | 7632008 | Distal Intergenic |
| chr15 | 7641263 | 7641477 | Distal Intergenic |
| chr15 | 7642738 | 7644116 | Distal Intergenic |
| chr15 | 7644179 | 7644450 | Distal Intergenic |
| chr15 | 7666006 | 7666355 | Distal Intergenic |
| chr15 | 7700232 | 7700563 | Distal Intergenic |
| chr15 | 7761124 | 7761769 | Distal Intergenic |
| chr15 | 7788419 | 7789496 | Distal Intergenic |
| chr15 | 7812413 | 7813505 | Distal Intergenic |
| chr15 | 7827649 | 7828038 | Distal Intergenic |
| chr15 | 7881641 | 7882453 | Distal Intergenic |
| chr15 | 7912063 | 7912299 | Distal Intergenic |
| chr15 | 7912408 | 7913904 | Distal Intergenic |
| chr15 | 8059287 | 8059858 | Distal Intergenic |
| chr15 | 8139651 | 8140665 | Distal Intergenic |
| chr15 | 8140746 | 8140945 | Distal Intergenic |
| chr15 | 8391916 | 8392182 | Distal Intergenic |
| chr15 | 8392375 | 8394455 | Distal Intergenic |
| chr15 | 8395516 | 8396619 | Distal Intergenic |
| chr15 | 8396666 | 8397063 | Distal Intergenic |
| chr15 | 8617802 | 8618048 | Distal Intergenic |
| chr15 | 8618161 | 8618479 | Distal Intergenic |
| chr15 | 8735526 | 8735725 | Distal Intergenic |
| chr15 | 8944677 | 8945315 | Distal Intergenic |
| chr15 | 8945415 | 8945958 | Distal Intergenic |
| chr15 | 9165080 | 9166078 | Distal Intergenic |
| chr15 | 10414572 | 10415692 | Distal Intergenic |

| | | | |
|---|---|---|---|
| chr15 | 10415743 | 10416025 | Distal Intergenic |
| chr15 | 10598641 | 10598867 | Distal Intergenic |
| chr15 | 10600107 | 10600719 | Distal Intergenic |
| chr15 | 10615165 | 10615786 | Distal Intergenic |
| chr15 | 10620623 | 10620904 | Distal Intergenic |
| chr15 | 10620941 | 10621229 | Distal Intergenic |
| chr15 | 10747733 | 10748996 | Distal Intergenic |
| chr15 | 10752646 | 10753482 | Distal Intergenic |
| chr15 | 10794056 | 10795410 | Distal Intergenic |
| chr15 | 10940914 | 10941660 | Distal Intergenic |
| chr15 | 10947660 | 10947913 | Distal Intergenic |
| chr15 | 10948945 | 10949181 | Distal Intergenic |
| chr15 | 11013880 | 11015305 | Distal Intergenic |
| chr15 | 11088467 | 11089460 | Distal Intergenic |
| chr15 | 11383561 | 11385531 | Distal Intergenic |
| chr15 | 11386010 | 11386397 | Distal Intergenic |
| chr15 | 11386906 | 11387842 | Distal Intergenic |
| chr15 | 11626142 | 11626870 | Distal Intergenic |
| chr15 | 11629558 | 11630106 | Distal Intergenic |
| chr15 | 11925879 | 11926099 | Distal Intergenic |
| chr15 | 12047105 | 12047505 | Distal Intergenic |
| chr15 | 12047653 | 12048028 | Distal Intergenic |
| chr15 | 12048400 | 12048619 | Distal Intergenic |
| chr15 | 12063523 | 12063799 | Distal Intergenic |
| chr15 | 12064060 | 12064893 | Distal Intergenic |
| chr15 | 12065118 | 12065352 | Distal Intergenic |
| chr15 | 12134112 | 12134629 | Distal Intergenic |
| chr15 | 12151322 | 12152349 | Distal Intergenic |
| chr15 | 12251846 | 12252162 | Distal Intergenic |
| chr15 | 12252230 | 12252490 | Distal Intergenic |
| chr15 | 12503214 | 12503771 | Distal Intergenic |
| chr15 | 13296138 | 13296768 | Distal Intergenic |
| chr15 | 13297063 | 13298763 | Distal Intergenic |
| chr15 | 19551897 | 19552683 | Distal Intergenic |
| chr15 | 20365957 | 20366231 | Distal Intergenic |
| chr15 | 22662430 | 22663467 | Intron |
| chr15 | 25278370 | 25278746 | 3' UTR |
| chr15 | 25298586 | 25299797 | Promoter |
| chr15 | 25303210 | 25303873 | Promoter |
| chr15 | 25325863 | 25327500 | Promoter |
| chr15 | 25342086 | 25342303 | Promoter |
| chr15 | 25342389 | 25342647 | Promoter |
| chr15 | 25343123 | 25343352 | Promoter |
| chr15 | 25420641 | 25421175 | Promoter |

| | | | |
|---|---|---|---|
| chr15 | 25424246 | 25424878 | Promoter |
| chr15 | 25552294 | 25555604 | Intron |
| chr15 | 25555953 | 25556195 | Intron |
| chr15 | 25567378 | 25567612 | Intron |
| chr15 | 25567676 | 25568334 | Intron |
| chr15 | 25581935 | 25583574 | 3' UTR |
| chr15 | 25792588 | 25792823 | Distal Intergenic |
| chr15 | 25914061 | 25914281 | Distal Intergenic |
| chr15 | 26283178 | 26284006 | Intron |
| chr15 | 26304032 | 26304878 | Distal Intergenic |
| chr15 | 27347969 | 27348216 | Intron |
| chr15 | 27348326 | 27349217 | Intron |
| chr15 | 27563636 | 27564468 | Intron |
| chr15 | 27731668 | 27733099 | Intron |
| chr15 | 27853121 | 27853525 | Distal Intergenic |
| chr15 | 27853833 | 27854234 | Distal Intergenic |
| chr15 | 27855192 | 27857523 | Distal Intergenic |
| chr15 | 27897666 | 27898761 | Distal Intergenic |
| chr15 | 27913540 | 27913833 | Distal Intergenic |
| chr15 | 27914034 | 27914509 | Distal Intergenic |
| chr15 | 27921647 | 27923989 | Distal Intergenic |
| chr15 | 27924071 | 27924432 | Distal Intergenic |
| chr15 | 27926761 | 27928220 | Distal Intergenic |
| chr15 | 27933908 | 27935153 | Distal Intergenic |
| chr15 | 27954821 | 27955115 | Distal Intergenic |
| chr15 | 27989399 | 27990146 | Distal Intergenic |
| chr15 | 28010575 | 28010863 | Intron |
| chr15 | 28011176 | 28011769 | Intron |
| chr15 | 28061919 | 28063861 | Intron |
| chr15 | 28088681 | 28089062 | Intron |
| chr15 | 28109503 | 28110017 | Intron |
| chr15 | 28110135 | 28110398 | Intron |
| chr15 | 29329929 | 29330182 | Intron |
| chr15 | 30275054 | 30276678 | Distal Intergenic |
| chr15 | 30312870 | 30313328 | Distal Intergenic |
| chr15 | 30329452 | 30331221 | Distal Intergenic |
| chr15 | 31021871 | 31022305 | Intron |
| chr15 | 31114499 | 31114988 | 5' UTR |
| chr15 | 31116446 | 31117861 | Exon |
| chr15 | 31117907 | 31118779 | Intron |
| chr15 | 31152239 | 31152495 | Distal Intergenic |
| chr15 | 31153725 | 31153971 | Distal Intergenic |
| chr15 | 31154901 | 31155167 | Distal Intergenic |
| chr15 | 31227433 | 31227895 | Intron |

| | | | |
|---|---|---|---|
| chr15 | 31241387 | 31241779 | Promoter |
| chr15 | 31241834 | 31242172 | Promoter |
| chr15 | 31261697 | 31262130 | Intron |
| chr15 | 31530883 | 31531242 | Distal Intergenic |
| chr15 | 31531425 | 31532049 | Distal Intergenic |
| chr15 | 32171099 | 32171423 | Distal Intergenic |
| chr15 | 32249931 | 32251526 | Distal Intergenic |
| chr15 | 32251610 | 32253225 | Distal Intergenic |
| chr15 | 32310299 | 32310708 | Distal Intergenic |
| chr15 | 32331436 | 32332020 | Intron |
| chr15 | 32332096 | 32332327 | Intron |
| chr15 | 32846327 | 32846844 | Intron |
| chr15 | 32850068 | 32850460 | Intron |
| chr15 | 32850592 | 32851146 | Intron |
| chr15 | 32929749 | 32930162 | 3' UTR |
| chr15 | 33573368 | 33573876 | Distal Intergenic |
| chr15 | 33574074 | 33574385 | Distal Intergenic |
| chr15 | 33836334 | 33837645 | Intron |
| chr15 | 33845333 | 33845567 | Intron |
| chr15 | 33845667 | 33846247 | Intron |
| chr15 | 33881184 | 33882154 | Intron |
| chr15 | 34012453 | 34013835 | Intron |
| chr15 | 34039552 | 34040598 | Exon |
| chr15 | 34076253 | 34077805 | Intron |
| chr15 | 34168497 | 34168705 | Intron |
| chr15 | 34182495 | 34182857 | Intron |
| chr15 | 34183883 | 34184664 | Intron |
| chr15 | 34262201 | 34262737 | Intron |
| chr15 | 34342091 | 34344109 | Intron |
| chr15 | 34371533 | 34373755 | Downstream |
| chr15 | 34425067 | 34425473 | Distal Intergenic |
| chr15 | 34608068 | 34608269 | Intron |
| chr15 | 34608638 | 34609176 | Intron |
| chr15 | 34950187 | 34950395 | Distal Intergenic |
| chr15 | 34950482 | 34951138 | Distal Intergenic |
| chr15 | 35239424 | 35239705 | Intron |
| chr15 | 35248977 | 35249194 | Intron |
| chr15 | 35301019 | 35301594 | Distal Intergenic |
| chr15 | 35994392 | 35995896 | Intron |
| chr15 | 36211849 | 36212881 | Distal Intergenic |
| chr15 | 36251992 | 36253727 | Distal Intergenic |
| chr15 | 36254110 | 36254866 | Distal Intergenic |
| chr15 | 36303754 | 36304350 | Distal Intergenic |
| chr15 | 36304469 | 36304971 | Distal Intergenic |

| | | | |
|---|---|---|---|
| chr15 | 36535946 | 36537865 | Distal Intergenic |
| chr15 | 36538021 | 36538539 | Distal Intergenic |
| chr15 | 36549258 | 36549487 | Distal Intergenic |
| chr15 | 36550946 | 36551161 | Distal Intergenic |
| chr15 | 36551382 | 36551581 | Distal Intergenic |
| chr15 | 36586016 | 36587895 | Distal Intergenic |
| chr15 | 36672438 | 36673188 | Distal Intergenic |
| chr15 | 36689182 | 36690106 | Distal Intergenic |
| chr15 | 36691304 | 36693180 | Distal Intergenic |
| chr15 | 36697279 | 36697520 | Distal Intergenic |
| chr15 | 36716367 | 36716640 | Distal Intergenic |
| chr15 | 36716925 | 36719284 | Distal Intergenic |
| chr15 | 36721278 | 36722391 | Distal Intergenic |
| chr15 | 36722513 | 36723043 | Distal Intergenic |
| chr15 | 36723118 | 36723357 | Distal Intergenic |
| chr15 | 36723886 | 36724517 | Distal Intergenic |
| chr15 | 36771709 | 36774268 | Distal Intergenic |
| chr15 | 36781784 | 36782200 | Distal Intergenic |
| chr15 | 36784442 | 36786215 | Distal Intergenic |
| chr15 | 36857027 | 36858015 | Distal Intergenic |
| chr15 | 36868095 | 36868361 | Distal Intergenic |
| chr15 | 36935995 | 36936673 | 5' UTR |
| chr15 | 36937250 | 36937477 | 5' UTR |
| chr15 | 36960812 | 36961673 | Intron |
| chr15 | 37089385 | 37089676 | Intron |
| chr15 | 37122243 | 37123519 | Distal Intergenic |
| chr15 | 37126841 | 37127491 | Distal Intergenic |
| chr15 | 37127595 | 37131760 | Distal Intergenic |
| chr15 | 37162759 | 37164975 | Intron |
| chr15 | 37193515 | 37193733 | Intron |
| chr15 | 37193779 | 37194012 | Intron |
| chr15 | 37194188 | 37194884 | Intron |
| chr15 | 37196145 | 37196490 | Intron |
| chr15 | 37196527 | 37198718 | Intron |
| chr15 | 37204966 | 37205365 | Intron |
| chr15 | 37210164 | 37212142 | Intron |
| chr15 | 37235445 | 37235699 | Intron |
| chr15 | 37242886 | 37243092 | Intron |
| chr15 | 37243180 | 37243471 | Intron |
| chr15 | 37406860 | 37407090 | Distal Intergenic |
| chr15 | 37425514 | 37426176 | Distal Intergenic |
| chr15 | 37536802 | 37537113 | Distal Intergenic |
| chr15 | 37537183 | 37537402 | Distal Intergenic |
| chr15 | 37890643 | 37890851 | Distal Intergenic |

| | | | |
|---|---|---|---|
| chr15 | 38106028 | 38106334 | Distal Intergenic |
| chr15 | 38253902 | 38254110 | Intron |
| chr15 | 38256206 | 38256517 | Intron |
| chr15 | 38314858 | 38315216 | Intron |
| chr15 | 38320678 | 38322454 | Intron |
| chr15 | 38381977 | 38382353 | Distal Intergenic |
| chr15 | 38382393 | 38385288 | Distal Intergenic |
| chr15 | 38385902 | 38387998 | Distal Intergenic |
| chr15 | 38388118 | 38388389 | Distal Intergenic |
| chr15 | 38399968 | 38400406 | Distal Intergenic |
| chr15 | 38447633 | 38447947 | Intron |
| chr15 | 38502086 | 38502727 | Intron |
| chr15 | 38515213 | 38515632 | Intron |
| chr15 | 38620630 | 38621916 | Intron |
| chr15 | 38639027 | 38640948 | Intron |
| chr15 | 38651427 | 38652152 | Distal Intergenic |
| chr15 | 38655229 | 38656185 | Distal Intergenic |
| chr15 | 38658082 | 38658291 | Distal Intergenic |
| chr15 | 38754604 | 38754809 | Intron |
| chr15 | 38763459 | 38764921 | Intron |
| chr15 | 38781044 | 38783621 | 3' UTR |
| chr15 | 38787752 | 38788829 | Intron |
| chr15 | 38804515 | 38806039 | Exon |
| chr15 | 38809724 | 38811565 | Promoter |
| chr15 | 38838194 | 38838881 | Intron |
| chr15 | 38944444 | 38944989 | Distal Intergenic |
| chr15 | 40195266 | 40197526 | Intron |
| chr15 | 40197694 | 40197917 | Intron |
| chr15 | 40487645 | 40487875 | Intron |
| chr15 | 40785370 | 40788195 | Distal Intergenic |
| chr15 | 40788320 | 40790376 | Distal Intergenic |
| chr15 | 40800911 | 40802144 | Distal Intergenic |
| chr15 | 40859403 | 40861327 | Promoter |
| chr15 | 41558996 | 41559269 | Intron |
| chr15 | 41566694 | 41568164 | Intron |
| chr15 | 41573248 | 41573740 | 3' UTR |
| chr15 | 43113655 | 43114097 | Intron |
| chr15 | 43638824 | 43639127 | Intron |
| chr15 | 43696894 | 43697632 | 3' UTR |
| chr15 | 44258426 | 44258686 | Intron |
| chr15 | 44258818 | 44259164 | Intron |
| chr15 | 44259272 | 44259478 | Intron |
| chr15 | 44259969 | 44260182 | Intron |
| chr15 | 44530523 | 44530765 | Distal Intergenic |

| | | | |
|---|---|---|---|
| chr15 | 49237942 | 49238192 | Intron |
| chr15 | 50003907 | 50004106 | Distal Intergenic |
| chr15 | 50535518 | 50535970 | Intron |
| chr15 | 50773979 | 50774240 | Exon |
| chr15 | 50774291 | 50774658 | Intron |
| chr15 | 50804090 | 50804764 | Intron |
| chr15 | 50806315 | 50806537 | Intron |
| chr15 | 50806707 | 50807053 | Intron |
| chr15 | 51297147 | 51299415 | 3' UTR |
| chr15 | 51452762 | 51453140 | Distal Intergenic |
| chr15 | 51453181 | 51453559 | Distal Intergenic |
| chr15 | 51511115 | 51511836 | Promoter |
| chr15 | 51513640 | 51514340 | 3' UTR |
| chr15 | 51562319 | 51563118 | Intron |
| chr15 | 51635992 | 51636257 | Intron |
| chr15 | 51636486 | 51636865 | Intron |
| chr15 | 51638024 | 51638231 | Intron |
| chr15 | 51638272 | 51638602 | Intron |
| chr15 | 51684091 | 51684357 | Intron |
| chr15 | 51684406 | 51686502 | Intron |
| chr15 | 51686542 | 51686763 | Intron |
| chr15 | 51696354 | 51697072 | 3' UTR |
| chr15 | 51708924 | 51709176 | Distal Intergenic |
| chr15 | 51709390 | 51709609 | Distal Intergenic |
| chr15 | 51817206 | 51818033 | Intron |
| chr15 | 51821796 | 51822863 | Intron |
| chr15 | 51823069 | 51823268 | Intron |
| chr15 | 52935054 | 52936548 | Intron |
| chr15 | 52944269 | 52944884 | Promoter |
| chr15 | 53001693 | 53003865 | Distal Intergenic |
| chr15 | 53027823 | 53029017 | Distal Intergenic |
| chr15 | 53053981 | 53055013 | Intron |
| chr15 | 53089787 | 53090034 | Distal Intergenic |
| chr15 | 53090323 | 53091204 | Distal Intergenic |
| chr15 | 53218083 | 53219271 | Distal Intergenic |
| chr15 | 53222263 | 53223808 | Distal Intergenic |
| chr15 | 53298819 | 53299164 | Distal Intergenic |
| chr15 | 53507620 | 53507822 | Distal Intergenic |
| chr15 | 53520476 | 53521242 | Distal Intergenic |
| chr15 | 53521422 | 53521700 | Distal Intergenic |
| chr15 | 53728673 | 53733864 | Distal Intergenic |
| chr15 | 53764592 | 53765064 | Distal Intergenic |
| chr15 | 53837408 | 53837641 | Intron |
| chr15 | 53838115 | 53838325 | Intron |

| | | | |
|---|---|---|---|
| chr15 | 54966093 | 54967306 | Distal Intergenic |
| chr15 | 55017850 | 55018912 | Distal Intergenic |
| chr15 | 55147063 | 55147491 | Distal Intergenic |
| chr15 | 55165085 | 55165373 | Distal Intergenic |
| chr15 | 55388537 | 55389353 | Distal Intergenic |
| chr15 | 55413101 | 55413852 | Distal Intergenic |
| chr15 | 55658147 | 55658364 | Intron |
| chr15 | 55849040 | 55849244 | Intron |
| chr15 | 56385534 | 56385733 | 3' UTR |
| chr15 | 56517030 | 56517469 | Intron |
| chr15 | 56552831 | 56553327 | Distal Intergenic |
| chr15 | 56971848 | 56972047 | Intron |
| chr15 | 57399312 | 57399723 | Intron |
| chr15 | 57399795 | 57400220 | Intron |
| chr15 | 57443753 | 57444028 | Intron |
| chr15 | 57444069 | 57444412 | Intron |
| chr15 | 57447674 | 57449829 | Intron |
| chr15 | 57477519 | 57478452 | Intron |
| chr15 | 57479792 | 57480200 | Intron |
| chr15 | 57480299 | 57480535 | Intron |
| chr15 | 57481262 | 57483138 | Intron |
| chr15 | 57485842 | 57486461 | Intron |
| chr15 | 57491194 | 57492304 | Intron |
| chr15 | 57493798 | 57494031 | Intron |
| chr15 | 57592998 | 57593197 | Promoter |
| chr15 | 57593294 | 57593769 | Promoter |
| chr15 | 57615801 | 57616903 | Distal Intergenic |
| chr15 | 57722986 | 57723658 | Intron |
| chr15 | 57738214 | 57739172 | Intron |
| chr15 | 57807945 | 57808190 | Intron |
| chr15 | 57849282 | 57849883 | Distal Intergenic |
| chr15 | 57995948 | 57997115 | Intron |
| chr15 | 58025947 | 58026146 | Distal Intergenic |
| chr15 | 58087602 | 58088169 | Distal Intergenic |
| chr15 | 58088390 | 58089563 | Distal Intergenic |
| chr15 | 58090051 | 58091342 | Distal Intergenic |
| chr15 | 58092053 | 58092836 | Distal Intergenic |
| chr15 | 58132484 | 58132762 | Distal Intergenic |
| chr15 | 58133192 | 58133830 | Distal Intergenic |
| chr15 | 58156716 | 58157785 | Distal Intergenic |
| chr15 | 58163034 | 58163503 | Distal Intergenic |
| chr15 | 58189451 | 58190293 | Distal Intergenic |
| chr15 | 58194936 | 58195711 | Distal Intergenic |
| chr15 | 58198849 | 58199481 | Distal Intergenic |

| | | | |
|---|---|---|---|
| chr15 | 58209810 | 58213552 | Distal Intergenic |
| chr15 | 58213645 | 58214510 | Distal Intergenic |
| chr15 | 58214724 | 58214925 | Distal Intergenic |
| chr15 | 58244301 | 58244963 | Downstream |
| chr15 | 58248282 | 58248506 | Intron |
| chr15 | 58648681 | 58648880 | Distal Intergenic |
| chr15 | 58649732 | 58650882 | Distal Intergenic |
| chr15 | 58745643 | 58745889 | Intron |
| chr15 | 58758993 | 58760085 | Intron |
| chr15 | 58763872 | 58765809 | Intron |
| chr15 | 58829300 | 58829709 | Intron |
| chr15 | 58912127 | 58912551 | Intron |
| chr15 | 58922117 | 58922639 | Intron |
| chr15 | 58982932 | 58983445 | 3' UTR |
| chr15 | 58990015 | 58990445 | Intron |
| chr15 | 58990583 | 58991333 | Intron |
| chr15 | 58999759 | 59002090 | Intron |
| chr15 | 59015794 | 59015998 | Intron |
| chr15 | 59016155 | 59017046 | Intron |
| chr15 | 59017140 | 59017408 | Intron |
| chr15 | 59017463 | 59017662 | Intron |
| chr15 | 59017792 | 59018011 | Intron |
| chr15 | 59018063 | 59018304 | Intron |
| chr15 | 59018352 | 59018553 | Intron |
| chr15 | 59029177 | 59029669 | Intron |
| chr15 | 59033365 | 59034508 | Intron |
| chr15 | 59043618 | 59044821 | Distal Intergenic |
| chr15 | 59053934 | 59054156 | Distal Intergenic |
| chr15 | 59054237 | 59056108 | Distal Intergenic |
| chr15 | 59145561 | 59146298 | Intron |
| chr15 | 59146353 | 59148541 | 3' UTR |
| chr15 | 59217464 | 59219861 | Intron |
| chr15 | 59240761 | 59241350 | Distal Intergenic |
| chr15 | 59243733 | 59244018 | Distal Intergenic |
| chr15 | 59279663 | 59280715 | Promoter |
| chr15 | 59309006 | 59309388 | Intron |
| chr15 | 59309568 | 59309833 | Intron |
| chr15 | 59312268 | 59313397 | Intron |
| chr15 | 59381343 | 59381778 | Intron |
| chr15 | 59386616 | 59387859 | 3' UTR |
| chr15 | 59428087 | 59428798 | 3' UTR |
| chr15 | 59429121 | 59429862 | 3' UTR |
| chr15 | 59429981 | 59431673 | Exon |
| chr15 | 59431727 | 59432067 | Intron |

| | | | |
|---|---|---|---|
| chr15 | 59557666 | 59558064 | Intron |
| chr15 | 59590882 | 59591361 | Intron |
| chr15 | 59591526 | 59592128 | Intron |
| chr15 | 59654267 | 59654914 | Intron |
| chr15 | 59932956 | 59933325 | Intron |
| chr15 | 59933559 | 59935094 | Exon |
| chr15 | 59935294 | 59936133 | Intron |
| chr15 | 59936271 | 59936630 | Intron |
| chr15 | 59941889 | 59944655 | 5' UTR |
| chr15 | 59957837 | 59958052 | Intron |
| chr15 | 59958259 | 59958464 | Intron |
| chr15 | 60936751 | 60938016 | Intron |
| chr15 | 60940470 | 60942038 | Intron |
| chr15 | 60946387 | 60946790 | Intron |
| chr15 | 60955802 | 60956093 | Intron |
| chr15 | 60956374 | 60957841 | Intron |
| chr15 | 60958186 | 60960015 | Intron |
| chr15 | 60960179 | 60960777 | Intron |
| chr15 | 60969810 | 60971600 | Exon |
| chr15 | 60989374 | 60990557 | Intron |
| chr15 | 60996863 | 60997567 | Intron |
| chr15 | 61038422 | 61040302 | Intron |
| chr15 | 61072365 | 61074265 | Intron |
| chr15 | 61079028 | 61079227 | Intron |
| chr15 | 61125383 | 61127696 | Intron |
| chr15 | 61128180 | 61132438 | Intron |
| chr15 | 61283298 | 61283534 | Intron |
| chr15 | 61283690 | 61287095 | Intron |
| chr15 | 61338201 | 61338778 | Intron |
| chr15 | 61339988 | 61340466 | Intron |
| chr15 | 61363242 | 61366323 | Intron |
| chr15 | 61366451 | 61368015 | Intron |
| chr15 | 61439530 | 61442234 | Intron |
| chr15 | 61449929 | 61451308 | Intron |
| chr15 | 61479070 | 61479363 | Intron |
| chr15 | 61479415 | 61479632 | Intron |
| chr15 | 61516806 | 61517063 | Intron |
| chr15 | 61517117 | 61517361 | Intron |
| chr15 | 61517436 | 61517791 | Intron |
| chr15 | 61743552 | 61744591 | Distal Intergenic |
| chr15 | 61744998 | 61745799 | Distal Intergenic |
| chr15 | 61816465 | 61816770 | Distal Intergenic |
| chr15 | 61816811 | 61817057 | Distal Intergenic |
| chr15 | 61817336 | 61818391 | Distal Intergenic |

| | | | |
|---|---|---|---|
| chr15 | 61823158 | 61823646 | Distal Intergenic |
| chr15 | 61830607 | 61830984 | Distal Intergenic |
| chr15 | 61869918 | 61870525 | Distal Intergenic |
| chr15 | 61870808 | 61871191 | Distal Intergenic |
| chr15 | 61875089 | 61875401 | Distal Intergenic |
| chr15 | 61910314 | 61911043 | Distal Intergenic |
| chr15 | 61911138 | 61912131 | Distal Intergenic |
| chr15 | 61914747 | 61915527 | Distal Intergenic |
| chr15 | 61923450 | 61924913 | Distal Intergenic |
| chr15 | 61950408 | 61950976 | Distal Intergenic |
| chr15 | 61970357 | 61971082 | Distal Intergenic |
| chr15 | 61993215 | 61994387 | Distal Intergenic |
| chr15 | 62446414 | 62446687 | Distal Intergenic |
| chr15 | 63172171 | 63172395 | Distal Intergenic |
| chr15 | 63229413 | 63229631 | Distal Intergenic |
| chr15 | 63819909 | 63821355 | 5' UTR |
| chr15 | 63821425 | 63821688 | Intron |
| chr15 | 63828635 | 63828894 | Promoter |
| chr15 | 63829050 | 63829273 | Promoter |
| chr15 | 63836704 | 63838982 | Intron |
| chr15 | 63844903 | 63845941 | 5' UTR |
| chr15 | 63850178 | 63851029 | 5' UTR |
| chr15 | 63863434 | 63863644 | Intron |
| chr15 | 63864254 | 63865388 | Intron |
| chr15 | 63890639 | 63891077 | Intron |
| chr15 | 63983834 | 63984903 | Exon |
| chr15 | 64016392 | 64016964 | Intron |
| chr15 | 64162392 | 64162591 | Distal Intergenic |
| chr15 | 64162836 | 64163035 | Distal Intergenic |
| chr15 | 64213004 | 64214031 | Intron |
| chr15 | 64214533 | 64215702 | Intron |
| chr15 | 64308960 | 64309312 | Intron |
| chr15 | 65768094 | 65768346 | Intron |
| chr15 | 66371933 | 66375585 | Intron |
| chr15 | 66675837 | 66676040 | Distal Intergenic |
| chr15 | 66720519 | 66721415 | Intron |
| chr15 | 66722368 | 66723059 | Intron |
| chr15 | 66763148 | 66764036 | Intron |
| chr15 | 66978677 | 66979270 | Distal Intergenic |
| chr15 | 67126594 | 67126793 | Distal Intergenic |
| chr15 | 67612414 | 67612625 | Intron |
| chr15 | 67991970 | 67992176 | Intron |
| chr15 | 67992278 | 67992501 | Intron |
| chr15 | 67992559 | 67992886 | Intron |

| | | | |
|---|---|---|---|
| chr15 | 67997206 | 67997800 | Intron |
| chr15 | 72485475 | 72486401 | Intron |
| chr15 | 72489849 | 72491719 | Promoter |
| chr15 | 72491806 | 72492359 | 5' UTR |
| chr15 | 72492508 | 72493136 | 5' UTR |
| chr15 | 72493330 | 72493782 | Intron |
| chr15 | 72498635 | 72498907 | Intron |
| chr15 | 72921339 | 72921694 | Distal Intergenic |
| chr15 | 72977947 | 72978823 | Promoter |
| chr15 | 72980243 | 72980653 | Intron |
| chr15 | 72980909 | 72982096 | Intron |
| chr15 | 72993182 | 72993887 | Intron |
| chr15 | 73014475 | 73015013 | Promoter |
| chr15 | 73065039 | 73066281 | Intron |
| chr15 | 73066553 | 73066861 | Promoter |
| chr15 | 73072904 | 73074456 | Promoter |
| chr15 | 73074633 | 73074916 | Promoter |
| chr15 | 73075030 | 73075229 | Promoter |
| chr15 | 73075578 | 73075845 | Promoter |
| chr15 | 73076079 | 73078140 | Promoter |
| chr15 | 73098595 | 73099086 | Distal Intergenic |
| chr15 | 73253754 | 73254146 | Distal Intergenic |
| chr15 | 73579081 | 73580335 | Intron |
| chr15 | 74131728 | 74131948 | Distal Intergenic |
| chr15 | 74132005 | 74132217 | Distal Intergenic |
| chr15 | 74419518 | 74420317 | 3' UTR |
| chr15 | 74452111 | 74452310 | Distal Intergenic |
| chr15 | 74452472 | 74454404 | Distal Intergenic |
| chr15 | 74591501 | 74592381 | Intron |
| chr15 | 74594584 | 74597072 | Intron |
| chr15 | 74614090 | 74616115 | Intron |
| chr15 | 74721939 | 74722767 | Intron |
| chr15 | 74751356 | 74752445 | 5' UTR |
| chr15 | 74760776 | 74764313 | Intron |
| chr15 | 74764442 | 74766045 | Intron |
| chr15 | 74784786 | 74785977 | Distal Intergenic |
| chr15 | 74786051 | 74786370 | Distal Intergenic |
| chr15 | 74822578 | 74822829 | Distal Intergenic |
| chr15 | 74822891 | 74823093 | Distal Intergenic |
| chr15 | 74825796 | 74826910 | Distal Intergenic |
| chr15 | 74827002 | 74830195 | Distal Intergenic |
| chr15 | 74830246 | 74830894 | Distal Intergenic |
| chr15 | 74831014 | 74831256 | Distal Intergenic |
| chr15 | 74879076 | 74879532 | Intron |

| | | | |
|---|---|---|---|
| chr15 | 74879589 | 74881414 | Intron |
| chr15 | 74893904 | 74894249 | Exon |
| chr15 | 75100233 | 75101719 | Distal Intergenic |
| chr15 | 75156453 | 75157626 | Intron |
| chr15 | 75159217 | 75161769 | Intron |
| chr15 | 75200034 | 75200495 | Promoter |
| chr15 | 75502351 | 75502960 | Intron |
| chr15 | 75627925 | 75628714 | Promoter |
| chr15 | 75681341 | 75681708 | Intron |
| chr15 | 75753896 | 75754195 | Distal Intergenic |
| chr15 | 75759825 | 75760024 | 3' UTR |
| chr15 | 75819421 | 75819701 | Exon |
| chr15 | 75829894 | 75830991 | Intron |
| chr15 | 75831039 | 75831818 | Intron |
| chr15 | 75837745 | 75838001 | Intron |
| chr15 | 75838484 | 75839923 | Intron |
| chr15 | 75840065 | 75840332 | Intron |
| chr15 | 75843679 | 75844749 | Intron |
| chr15 | 75845050 | 75845272 | Intron |
| chr15 | 75911699 | 75912052 | Intron |
| chr15 | 75987408 | 75987642 | Intron |
| chr15 | 75987816 | 75988017 | Intron |
| chr15 | 75988130 | 75989743 | Intron |
| chr15 | 75989963 | 75990264 | Intron |
| chr15 | 76022597 | 76023300 | Exon |
| chr15 | 76023422 | 76024161 | Intron |
| chr15 | 76024828 | 76026718 | Intron |
| chr15 | 76027033 | 76027241 | Intron |
| chr15 | 76027999 | 76028247 | Intron |
| chr15 | 76028383 | 76028629 | Intron |
| chr15 | 76028799 | 76030316 | Promoter |
| chr15 | 76034371 | 76035798 | Distal Intergenic |
| chr15 | 76037187 | 76038369 | Distal Intergenic |
| chr15 | 76149812 | 76150012 | Intron |
| chr15 | 76157405 | 76157649 | Intron |
| chr15 | 76157862 | 76158074 | Intron |
| chr15 | 76291524 | 76292098 | Intron |
| chr15 | 76307868 | 76308286 | Distal Intergenic |
| chr15 | 76524952 | 76525866 | Intron |
| chr15 | 76554648 | 76554940 | Intron |
| chr15 | 76707276 | 76708162 | Intron |
| chr15 | 76709788 | 76710115 | Intron |
| chr15 | 76734570 | 76734849 | Intron |
| chr15 | 76779078 | 76779324 | Intron |

| | | | |
|---|---|---|---|
| chr15 | 76865870 | 76868433 | Exon |
| chr15 | 76886190 | 76886389 | Intron |
| chr15 | 76889255 | 76889666 | Intron |
| chr15 | 76907301 | 76907701 | Intron |
| chr15 | 76980170 | 76985319 | Intron |
| chr15 | 77022816 | 77023527 | Intron |
| chr15 | 77036109 | 77037108 | Intron |
| chr15 | 77049969 | 77050995 | Intron |
| chr15 | 77227971 | 77229630 | Exon |
| chr15 | 77571115 | 77572173 | Intron |
| chr15 | 77572228 | 77572627 | Intron |
| chr15 | 77629470 | 77632365 | Intron |
| chr15 | 77632413 | 77632641 | Intron |
| chr15 | 77633010 | 77633319 | Intron |
| chr15 | 77633443 | 77634715 | Intron |
| chr15 | 77634950 | 77635187 | Intron |
| chr15 | 77637121 | 77637962 | Intron |
| chr15 | 77645633 | 77648379 | Intron |
| chr15 | 77648522 | 77650109 | Intron |
| chr15 | 77650701 | 77654749 | Intron |
| chr15 | 77654828 | 77655778 | Intron |
| chr15 | 77655843 | 77656120 | Intron |
| chr15 | 77656214 | 77656555 | Intron |
| chr15 | 77658437 | 77658645 | Intron |
| chr15 | 77659879 | 77662175 | Intron |
| chr15 | 77662304 | 77665755 | Intron |
| chr15 | 77665810 | 77672010 | Intron |
| chr15 | 77707126 | 77707599 | Intron |
| chr15 | 77707635 | 77707978 | Intron |
| chr15 | 77718965 | 77719305 | Intron |
| chr15 | 77720147 | 77720500 | Intron |
| chr15 | 77720614 | 77720813 | Intron |
| chr15 | 77743562 | 77744182 | Intron |
| chr15 | 77752247 | 77752851 | Intron |
| chr15 | 77800870 | 77801224 | Distal Intergenic |
| chr15 | 77801316 | 77801892 | Distal Intergenic |
| chr15 | 78280722 | 78280965 | Distal Intergenic |
| chr15 | 78637845 | 78638233 | Intron |
| chr15 | 78638300 | 78639106 | Intron |
| chr15 | 78658300 | 78658527 | Distal Intergenic |
| chr15 | 78658593 | 78659548 | Distal Intergenic |
| chr15 | 78673356 | 78677006 | Distal Intergenic |
| chr15 | 78705979 | 78706185 | Distal Intergenic |
| chr15 | 78706386 | 78706637 | Distal Intergenic |

| | | | |
|---|---|---|---|
| chr15 | 78706758 | 78707589 | Distal Intergenic |
| chr15 | 78730833 | 78731877 | Promoter |
| chr15 | 78927573 | 78928996 | 5' UTR |
| chr15 | 78936549 | 78936756 | Intron |
| chr15 | 78937115 | 78937367 | Intron |
| chr15 | 78937425 | 78939403 | Intron |
| chr15 | 78939770 | 78941160 | Intron |
| chr15 | 79122823 | 79123805 | Distal Intergenic |
| chr15 | 79128732 | 79130992 | Distal Intergenic |
| chr15 | 79153977 | 79154316 | Distal Intergenic |
| chr15 | 79157505 | 79158634 | Distal Intergenic |
| chr15 | 79158745 | 79159044 | Distal Intergenic |
| chr15 | 79160328 | 79160783 | Distal Intergenic |
| chr15 | 79160908 | 79161139 | Distal Intergenic |
| chr15 | 79489290 | 79489553 | Intron |
| chr15 | 79595572 | 79595841 | Distal Intergenic |
| chr15 | 79722760 | 79723789 | Distal Intergenic |
| chr15 | 79730020 | 79730951 | Intron |
| chr15 | 79884201 | 79885046 | Distal Intergenic |
| chr15 | 79891778 | 79892137 | Distal Intergenic |
| chr15 | 79892558 | 79892867 | Distal Intergenic |
| chr15 | 79893565 | 79893830 | Distal Intergenic |
| chr15 | 79906024 | 79907371 | Distal Intergenic |
| chr15 | 79909987 | 79910242 | Distal Intergenic |
| chr15 | 79910407 | 79910627 | Distal Intergenic |
| chr15 | 79911179 | 79911397 | Distal Intergenic |
| chr15 | 79911557 | 79914351 | Distal Intergenic |
| chr15 | 79963672 | 79964029 | Distal Intergenic |
| chr15 | 80084731 | 80085496 | Distal Intergenic |
| chr15 | 80085571 | 80085836 | Distal Intergenic |
| chr15 | 80085956 | 80087004 | Distal Intergenic |
| chr15 | 80087566 | 80087887 | Distal Intergenic |
| chr15 | 80088130 | 80088995 | Distal Intergenic |
| chr15 | 80180610 | 80181616 | Exon |
| chr15 | 80479453 | 80480289 | Distal Intergenic |
| chr15 | 80483235 | 80484725 | Distal Intergenic |
| chr15 | 80502350 | 80502624 | Distal Intergenic |
| chr15 | 80502870 | 80504361 | Distal Intergenic |
| chr15 | 80504417 | 80505095 | Distal Intergenic |
| chr15 | 80505188 | 80511288 | Distal Intergenic |
| chr15 | 80511489 | 80511778 | Distal Intergenic |
| chr15 | 80511963 | 80512170 | Distal Intergenic |
| chr15 | 80532601 | 80532800 | Distal Intergenic |
| chr15 | 80532940 | 80534103 | Distal Intergenic |

| | | | |
|---|---|---|---|
| chr15 | 80628496 | 80628958 | Intron |
| chr15 | 80629028 | 80629294 | Intron |
| chr15 | 80629662 | 80629972 | Intron |
| chr15 | 80630062 | 80630261 | Intron |
| chr15 | 80643051 | 80643492 | Distal Intergenic |
| chr15 | 80679647 | 80680933 | Distal Intergenic |
| chr15 | 80685764 | 80686353 | Distal Intergenic |
| chr15 | 80691044 | 80691301 | Distal Intergenic |
| chr15 | 80691583 | 80691793 | Distal Intergenic |
| chr15 | 80691897 | 80692338 | Distal Intergenic |
| chr15 | 80728994 | 80729258 | Intron |
| chr15 | 80872115 | 80872854 | Exon |
| chr15 | 80898119 | 80902561 | Distal Intergenic |
| chr15 | 80904283 | 80905198 | Distal Intergenic |
| chr15 | 80909768 | 80909967 | Distal Intergenic |
| chr15 | 80916136 | 80916564 | Distal Intergenic |
| chr15 | 80916645 | 80918522 | Distal Intergenic |
| chr15 | 80967824 | 80968239 | Distal Intergenic |
| chr15 | 80988740 | 80990831 | Intron |
| chr15 | 80993931 | 80999691 | Intron |
| chr15 | 81021537 | 81023997 | Intron |
| chr15 | 81077847 | 81078550 | Intron |
| chr15 | 81094897 | 81095254 | Intron |
| chr15 | 81183805 | 81184317 | Intron |
| chr15 | 81186946 | 81187878 | Exon |
| chr15 | 81190629 | 81191093 | Intron |
| chr15 | 81230562 | 81230761 | Intron |
| chr15 | 81348349 | 81349603 | Distal Intergenic |
| chr15 | 81352858 | 81353146 | Distal Intergenic |
| chr15 | 81415891 | 81416541 | Intron |
| chr15 | 81416649 | 81417193 | Intron |
| chr15 | 81559911 | 81560306 | Intron |
| chr15 | 81640756 | 81641290 | Intron |
| chr15 | 81688791 | 81690414 | Distal Intergenic |
| chr15 | 81790281 | 81790927 | Distal Intergenic |
| chr15 | 81976398 | 81977371 | Distal Intergenic |
| chr15 | 81977926 | 81979506 | Distal Intergenic |
| chr15 | 81981919 | 81982221 | Distal Intergenic |
| chr15 | 81982588 | 81983117 | Distal Intergenic |
| chr15 | 81983158 | 81983357 | Distal Intergenic |
| chr15 | 81989655 | 81989979 | Distal Intergenic |
| chr15 | 81990193 | 81990681 | Distal Intergenic |
| chr15 | 81990881 | 81991367 | Distal Intergenic |
| chr15 | 81991403 | 81991997 | Distal Intergenic |

| | | | |
|---|---|---|---|
| chr15 | 81994693 | 81995425 | Distal Intergenic |
| chr15 | 81995668 | 81997211 | Distal Intergenic |
| chr15 | 82551838 | 82552079 | Intron |
| chr15 | 82552457 | 82552656 | Intron |
| chr15 | 82622051 | 82622415 | Exon |
| chr15 | 82731708 | 82732052 | Distal Intergenic |
| chr15 | 82732225 | 82732441 | Distal Intergenic |
| chr15 | 82770647 | 82771597 | Intron |
| chr15 | 82972932 | 82974162 | Exon |
| chr15 | 82979149 | 82979396 | Intron |
| chr15 | 82980041 | 82980511 | Exon |
| chr15 | 82987522 | 82988510 | Intron |
| chr15 | 82995617 | 82995844 | Intron |
| chr15 | 82997432 | 82997703 | Intron |
| chr15 | 83122586 | 83122905 | Intron |
| chr15 | 83122955 | 83124221 | Intron |
| chr15 | 83168598 | 83168797 | Intron |
| chr15 | 83168838 | 83169327 | Intron |
| chr15 | 83239009 | 83239289 | Intron |
| chr15 | 83271231 | 83272177 | Intron |
| chr15 | 83297797 | 83298224 | Promoter |
| chr15 | 83393098 | 83393604 | Downstream |
| chr15 | 83394311 | 83394705 | 3' UTR |
| chr15 | 83394801 | 83396630 | 5' UTR |
| chr15 | 83404718 | 83406337 | Intron |
| chr15 | 83424389 | 83424793 | Promoter |
| chr15 | 84022341 | 84022540 | Distal Intergenic |
| chr15 | 84330639 | 84331233 | Intron |
| chr15 | 84754636 | 84755400 | Intron |
| chr15 | 85062309 | 85062900 | Distal Intergenic |
| chr15 | 85063269 | 85064314 | Distal Intergenic |
| chr15 | 85064367 | 85064643 | Distal Intergenic |
| chr15 | 85112505 | 85112914 | Promoter |
| chr15 | 85127572 | 85129904 | Distal Intergenic |
| chr15 | 85177357 | 85177606 | Exon |
| chr15 | 85177659 | 85178324 | Intron |
| chr15 | 85254429 | 85255243 | Intron |
| chr15 | 85309262 | 85309511 | Intron |
| chr15 | 85384031 | 85384271 | Exon |
| chr15 | 85386440 | 85387056 | Intron |
| chr15 | 85465246 | 85465445 | Intron |
| chr15 | 85475601 | 85476787 | Exon |
| chr15 | 85477313 | 85477582 | Intron |
| chr15 | 85494839 | 85501003 | Intron |

| | | | |
|---|---|---|---|
| chr15 | 85502012 | 85504982 | 3' UTR |
| chr15 | 85505033 | 85505617 | Intron |
| chr15 | 85505803 | 85506228 | Intron |
| chr15 | 85506506 | 85507127 | Intron |
| chr15 | 85507222 | 85508142 | Intron |
| chr15 | 85508298 | 85509793 | Intron |
| chr15 | 85509850 | 85510716 | Intron |
| chr15 | 85510955 | 85511254 | Intron |
| chr15 | 85514206 | 85515725 | Intron |
| chr15 | 85515802 | 85516131 | Intron |
| chr15 | 85516172 | 85516417 | Intron |
| chr15 | 85516697 | 85517621 | Intron |
| chr15 | 85724623 | 85724862 | Distal Intergenic |
| chr15 | 85725076 | 85725296 | Distal Intergenic |
| chr15 | 85766317 | 85766568 | Intron |
| chr15 | 85796670 | 85797248 | Distal Intergenic |
| chr15 | 85800128 | 85800614 | Distal Intergenic |
| chr15 | 85800714 | 85801191 | Distal Intergenic |
| chr15 | 85821856 | 85822117 | Distal Intergenic |
| chr15 | 85822346 | 85822819 | Distal Intergenic |
| chr15 | 85834489 | 85835951 | Distal Intergenic |
| chr15 | 85838064 | 85838700 | Distal Intergenic |
| chr15 | 85839078 | 85839747 | Distal Intergenic |
| chr15 | 85846174 | 85847656 | Distal Intergenic |
| chr15 | 85865017 | 85865369 | Distal Intergenic |
| chr15 | 85924718 | 85925281 | Promoter |
| chr15 | 85925328 | 85925618 | Intron |
| chr15 | 85942855 | 85943312 | Intron |
| chr15 | 85943358 | 85944152 | Intron |
| chr15 | 85944229 | 85944453 | Intron |
| chr15 | 85997228 | 85998087 | Intron |
| chr15 | 86016724 | 86017048 | Intron |
| chr15 | 86017286 | 86017790 | Intron |
| chr15 | 87677791 | 87678827 | Distal Intergenic |
| chr15 | 88563511 | 88563951 | Intron |
| chr15 | 89000205 | 89000472 | Downstream |
| chr15 | 89005249 | 89005467 | 3' UTR |
| chr15 | 89005547 | 89006401 | 3' UTR |
| chr15 | 89007257 | 89011192 | Promoter |
| chr15 | 89011278 | 89011539 | Promoter |
| chr15 | 89019532 | 89020409 | Exon |
| chr15 | 89186604 | 89187568 | Intron |
| chr15 | 89198124 | 89198361 | Intron |
| chr15 | 89203596 | 89204134 | Distal Intergenic |

| | | | |
|---|---|---|---|
| chr15 | 89260123 | 89260339 | Distal Intergenic |
| chr15 | 90232075 | 90232965 | Intron |
| chr15 | 90261944 | 90263585 | Promoter |
| chr15 | 90342656 | 90343778 | Exon |
| chr15 | 90437428 | 90437737 | Promoter |
| chr15 | 90437871 | 90440619 | Promoter |
| chr15 | 90446609 | 90447163 | Exon |
| chr15 | 90486718 | 90487664 | Distal Intergenic |
| chr15 | 90548839 | 90549741 | Intron |
| chr15 | 90549802 | 90550056 | Intron |
| chr15 | 90746228 | 90747490 | Intron |
| chr15 | 90858956 | 90859642 | Distal Intergenic |
| chr15 | 90868883 | 90869159 | Distal Intergenic |
| chr15 | 90869250 | 90869450 | Distal Intergenic |
| chr15 | 91506699 | 91507304 | Downstream |
| chr15 | 91533448 | 91533959 | Intron |
| chr15 | 92755647 | 92756817 | Distal Intergenic |
| chr15 | 92836110 | 92836315 | Distal Intergenic |
| chr15 | 92836585 | 92837000 | Distal Intergenic |
| chr15 | 93017251 | 93018763 | Intron |
| chr15 | 93019012 | 93019319 | Intron |
| chr15 | 93020033 | 93020334 | Intron |
| chr15 | 93020690 | 93020921 | Intron |
| chr15 | 93106109 | 93106323 | Distal Intergenic |
| chr15 | 93228370 | 93228584 | Intron |
| chr15 | 93564208 | 93565484 | 3' UTR |
| chr15 | 94403826 | 94406211 | Intron |
| chr15 | 94419309 | 94419545 | Intron |
| chr15 | 95201303 | 95201819 | Distal Intergenic |
| chr15 | 95633477 | 95636222 | Distal Intergenic |
| chr15 | 95653500 | 95654869 | Distal Intergenic |
| chr15 | 95689556 | 95689919 | Distal Intergenic |
| chr15 | 96117981 | 96118404 | Distal Intergenic |
| chr15 | 96118811 | 96119423 | Distal Intergenic |
| chr15 | 96138855 | 96139159 | Distal Intergenic |
| chr15 | 96290169 | 96290796 | Distal Intergenic |
| chr15 | 96452091 | 96452456 | Distal Intergenic |
| chr15 | 96528310 | 96528516 | Distal Intergenic |
| chr15 | 96535038 | 96535299 | Distal Intergenic |
| chr15 | 96535398 | 96535619 | Distal Intergenic |
| chr15 | 96586457 | 96586891 | Distal Intergenic |
| chr15 | 96586950 | 96587552 | Distal Intergenic |
| chr15 | 96587597 | 96587911 | Distal Intergenic |
| chr15 | 96600211 | 96601542 | Distal Intergenic |

| | | | |
|---|---|---|---|
| chr15 | 96601622 | 96602046 | Distal Intergenic |
| chr15 | 96652464 | 96653232 | Distal Intergenic |
| chr15 | 96660073 | 96661919 | Distal Intergenic |
| chr15 | 96662357 | 96662757 | Distal Intergenic |
| chr15 | 96662804 | 96663471 | Distal Intergenic |
| chr15 | 96673154 | 96673877 | Distal Intergenic |
| chr15 | 96673925 | 96676204 | Distal Intergenic |
| chr15 | 96707735 | 96708959 | Distal Intergenic |
| chr15 | 96709009 | 96709698 | Distal Intergenic |
| chr15 | 96872568 | 96872767 | Intron |
| chr15 | 97574361 | 97574738 | Distal Intergenic |
| chr15 | 97574790 | 97575076 | Distal Intergenic |
| chr15 | 97586355 | 97586625 | Distal Intergenic |
| chr15 | 97603480 | 97604142 | Distal Intergenic |
| chr15 | 97604213 | 97605058 | Distal Intergenic |
| chr15 | 97605300 | 97606509 | Distal Intergenic |
| chr15 | 97654962 | 97655182 | Distal Intergenic |
| chr15 | 97662146 | 97662345 | Distal Intergenic |
| chr15 | 97667325 | 97668574 | Distal Intergenic |
| chr15 | 97701408 | 97703158 | Distal Intergenic |
| chr15 | 97705334 | 97706667 | Distal Intergenic |
| chr15 | 97811881 | 97812088 | Distal Intergenic |
| chr15 | 97812137 | 97812971 | Distal Intergenic |
| chr15 | 97922965 | 97923764 | Distal Intergenic |
| chr15 | 97923807 | 97924021 | Distal Intergenic |
| chr15 | 98049512 | 98049717 | Distal Intergenic |
| chr15 | 98049941 | 98050340 | Distal Intergenic |
| chr15 | 98356356 | 98356595 | Intron |
| chr15 | 98356637 | 98357720 | Intron |
| chr15 | 98362895 | 98364294 | Intron |
| chr15 | 98397597 | 98398161 | Intron |
| chr15 | 98398221 | 98398725 | Intron |
| chr15 | 98492442 | 98492707 | Intron |
| chr15 | 98492880 | 98493340 | Intron |
| chr15 | 98495224 | 98495429 | Intron |
| chr15 | 98501333 | 98501563 | Intron |
| chr15 | 98503029 | 98503801 | Promoter |
| chr15 | 98574710 | 98575446 | Distal Intergenic |
| chr15 | 98765656 | 98765902 | Distal Intergenic |
| chr15 | 98815282 | 98816718 | Distal Intergenic |
| chr15 | 98823944 | 98824372 | Distal Intergenic |
| chr15 | 98859753 | 98860534 | Distal Intergenic |
| chr15 | 98860790 | 98861468 | Distal Intergenic |
| chr15 | 98861704 | 98864043 | Distal Intergenic |

| | | | |
|---|---|---|---|
| chr15 | 98877245 | 98877541 | Distal Intergenic |
| chr15 | 99082463 | 99082766 | Distal Intergenic |
| chr15 | 99223396 | 99224881 | Intron |
| chr15 | 99226822 | 99227073 | Intron |
| chr15 | 99228029 | 99228568 | Intron |
| chr15 | 99628238 | 99628437 | Distal Intergenic |
| chr15 | 99628601 | 99628832 | Distal Intergenic |
| chr15 | 99628908 | 99629470 | Distal Intergenic |
| chr15 | 99629608 | 99630129 | Distal Intergenic |
| chr15 | 99647728 | 99648404 | Intron |
| chr15 | 99648522 | 99651122 | Intron |
| chr15 | 99651230 | 99651849 | Intron |
| chr15 | 99651953 | 99652263 | Intron |
| chr15 | 99803429 | 99803689 | Intron |
| chr15 | 99869644 | 99870072 | Intron |
| chr15 | 99871561 | 99871823 | Intron |
| chr15 | 99872014 | 99873652 | Intron |
| chr15 | 99881169 | 99882126 | Intron |
| chr15 | 99890325 | 99891324 | Intron |
| chr15 | 99912306 | 99912631 | Intron |
| chr15 | 99912689 | 99912954 | Intron |
| chr15 | 99913339 | 99914008 | Intron |
| chr15 | 99917890 | 99919768 | Intron |
| chr15 | 99919818 | 99920623 | Intron |
| chr15 | 99926986 | 99927970 | 3' UTR |
| chr15 | 99940210 | 99940748 | Distal Intergenic |
| chr15 | 99949411 | 99950338 | Distal Intergenic |
| chr15 | 100050771 | 100051179 | Distal Intergenic |
| chr15 | 100051273 | 100052275 | Distal Intergenic |
| chr15 | 100052322 | 100052565 | Distal Intergenic |
| chr15 | 100057754 | 100057993 | Distal Intergenic |
| chr15 | 100058819 | 100059229 | Distal Intergenic |
| chr15 | 100081494 | 100082282 | Distal Intergenic |
| chr15 | 100100069 | 100101402 | Distal Intergenic |
| chr15 | 100129646 | 100129940 | Intron |
| chr15 | 100155422 | 100155860 | Intron |
| chr15 | 100156035 | 100156548 | Intron |
| chr15 | 100251833 | 100252037 | Intron |
| chr15 | 100550127 | 100550687 | Intron |
| chr15 | 100900364 | 100903895 | Distal Intergenic |
| chr15 | 101034776 | 101034982 | Intron |
| chr15 | 101051159 | 101051469 | Intron |
| chr15 | 101159534 | 101164285 | Intron |
| chr15 | 101165484 | 101166117 | Intron |

| | | | |
|---|---|---|---|
| chr15 | 101166362 | 101169322 | Intron |
| chr15 | 101169679 | 101169963 | Exon |
| chr15 | 101202753 | 101204070 | Distal Intergenic |
| chr15 | 101204257 | 101204481 | Distal Intergenic |
| chr15 | 101284888 | 101285094 | Distal Intergenic |
| chr15 | 101519552 | 101519893 | Intron |
| chr15 | 101523088 | 101524410 | Exon |
| chr15 | 101524509 | 101524750 | Intron |
| chr15 | 101524872 | 101526927 | Intron |
| chr15 | 101527341 | 101528558 | Intron |
| chr15 | 101535808 | 101536511 | Intron |
| chr15 | 101536556 | 101536811 | Intron |
| chr15 | 101538228 | 101538533 | Intron |
| chr15 | 101541120 | 101542134 | Intron |
| chr15 | 101542186 | 101545119 | Intron |
| chr15 | 101545293 | 101546658 | Intron |
| chr15 | 101546762 | 101549333 | Exon |
| chr15 | 101600129 | 101603161 | Exon |
| chr15 | 101804383 | 101807855 | Distal Intergenic |
| chr15 | 101808061 | 101808329 | Downstream |
| chr15 | 101810268 | 101811234 | 3' UTR |
| chr15 | 101811472 | 101812276 | 3' UTR |
| chr15 | 101846121 | 101846461 | Intron |
| chr15 | 101846505 | 101847213 | Intron |
| chr15 | 101876176 | 101876605 | Intron |
| chr15 | 101905016 | 101905254 | 3' UTR |
| chr15 | 102065614 | 102066240 | Distal Intergenic |
| chr15 | 102074417 | 102074806 | Distal Intergenic |
| chr15 | 102076252 | 102076767 | Distal Intergenic |
| chr15 | 102092872 | 102093563 | Distal Intergenic |
| chr15 | 102156599 | 102156820 | Distal Intergenic |
| chr15 | 102180170 | 102181194 | Intron |
| chr15 | 102232600 | 102233029 | Intron |
| chr15 | 102235322 | 102236255 | Intron |
| chr15 | 102236746 | 102236957 | Intron |
| chr15 | 102237387 | 102237899 | Intron |
| chr15 | 102348743 | 102349340 | Distal Intergenic |
| chr15 | 102632556 | 102632843 | Distal Intergenic |
| chr15 | 102666777 | 102668852 | Distal Intergenic |
| chr15 | 102668892 | 102669134 | Distal Intergenic |
| chr15 | 102669439 | 102670301 | Distal Intergenic |
| chr15 | 102958456 | 102958677 | Distal Intergenic |
| chr15 | 102961467 | 102961703 | Distal Intergenic |
| chr15 | 103045006 | 103046253 | Distal Intergenic |

| | | | |
|---|---|---|---|
| chr15 | 103051047 | 103051984 | Distal Intergenic |
| chr15 | 103070885 | 103071247 | Distal Intergenic |
| chr15 | 103176819 | 103177018 | Distal Intergenic |
| chr16 | 3743451 | 3743924 | Intron |
| chr16 | 3744095 | 3745092 | Intron |
| chr16 | 3819577 | 3820680 | Exon |
| chr16 | 3884338 | 3884577 | Intron |
| chr16 | 4077718 | 4078188 | Intron |
| chr16 | 4447705 | 4448921 | Intron |
| chr16 | 4464739 | 4466768 | Promoter |
| chr16 | 4469205 | 4470555 | Distal Intergenic |
| chr16 | 4473017 | 4473236 | Distal Intergenic |
| chr16 | 4473297 | 4473545 | Distal Intergenic |
| chr16 | 4477039 | 4479154 | Intron |
| chr16 | 4558045 | 4558671 | Exon |
| chr16 | 4586858 | 4588121 | Promoter |
| chr16 | 4624257 | 4624962 | Exon |
| chr16 | 4683440 | 4684025 | Intron |
| chr16 | 4776982 | 4778197 | Promoter |
| chr16 | 4878170 | 4879814 | Intron |
| chr16 | 4879889 | 4880100 | Intron |
| chr16 | 4880254 | 4880463 | Intron |
| chr16 | 4880581 | 4881454 | Intron |
| chr16 | 4913110 | 4913538 | Intron |
| chr16 | 4959271 | 4959730 | Intron |
| chr16 | 4959801 | 4960103 | Intron |
| chr16 | 5049343 | 5050377 | Intron |
| chr16 | 5135108 | 5137040 | 3' UTR |
| chr16 | 6640886 | 6641088 | Intron |
| chr16 | 7148030 | 7148357 | Intron |
| chr16 | 7951032 | 7951268 | Distal Intergenic |
| chr16 | 7951354 | 7951858 | Distal Intergenic |
| chr16 | 8636810 | 8638616 | Distal Intergenic |
| chr16 | 8665971 | 8668442 | Distal Intergenic |
| chr16 | 8671158 | 8671357 | Distal Intergenic |
| chr16 | 8671538 | 8671900 | Distal Intergenic |
| chr16 | 8790596 | 8791497 | Intron |
| chr16 | 8811102 | 8811434 | Intron |
| chr16 | 8829188 | 8829692 | 5' UTR |
| chr16 | 8830220 | 8831703 | Intron |
| chr16 | 8848561 | 8849799 | Intron |
| chr16 | 8849930 | 8851268 | Intron |
| chr16 | 8853775 | 8853977 | Intron |
| chr16 | 8861986 | 8863182 | Exon |

| | | | |
|---|---|---|---|
| chr16 | 8864059 | 8864535 | Intron |
| chr16 | 9038368 | 9039446 | Intron |
| chr16 | 9259238 | 9259641 | Distal Intergenic |
| chr16 | 10286399 | 10287466 | Distal Intergenic |
| chr16 | 10330716 | 10331400 | Distal Intergenic |
| chr16 | 10347328 | 10347597 | Distal Intergenic |
| chr16 | 10347740 | 10348171 | Distal Intergenic |
| chr16 | 10365519 | 10367761 | Distal Intergenic |
| chr16 | 10373317 | 10375217 | Distal Intergenic |
| chr16 | 10558465 | 10558887 | Intron |
| chr16 | 10640680 | 10641027 | Intron |
| chr16 | 10754367 | 10754708 | Intron |
| chr16 | 10754904 | 10755154 | Intron |
| chr16 | 10782093 | 10782934 | Intron |
| chr16 | 10785695 | 10786829 | Intron |
| chr16 | 10836861 | 10837131 | Promoter |
| chr16 | 10837274 | 10839607 | Promoter |
| chr16 | 10857166 | 10858111 | Exon |
| chr16 | 10872858 | 10873188 | Intron |
| chr16 | 10975579 | 10977694 | Intron |
| chr16 | 10991339 | 10993232 | Exon |
| chr16 | 11007657 | 11009664 | Exon |
| chr16 | 11015813 | 11017040 | Exon |
| chr16 | 11017464 | 11022865 | 3' UTR |
| chr16 | 11049837 | 11050895 | Intron |
| chr16 | 11057353 | 11058324 | Intron |
| chr16 | 11058395 | 11059733 | Intron |
| chr16 | 11062773 | 11063859 | Exon |
| chr16 | 11085772 | 11086071 | Intron |
| chr16 | 11133831 | 11134428 | Intron |
| chr16 | 11134846 | 11135482 | Intron |
| chr16 | 11172895 | 11173262 | Intron |
| chr16 | 11173391 | 11173590 | Intron |
| chr16 | 11173716 | 11176049 | Intron |
| chr16 | 11176540 | 11177018 | Intron |
| chr16 | 11193011 | 11194333 | Intron |
| chr16 | 11202456 | 11203400 | Intron |
| chr16 | 11203475 | 11204406 | Intron |
| chr16 | 11252836 | 11254445 | Intron |
| chr16 | 11254519 | 11255196 | Intron |
| chr16 | 11263530 | 11263762 | Intron |
| chr16 | 11286169 | 11286481 | Distal Intergenic |
| chr16 | 11501093 | 11502244 | Distal Intergenic |
| chr16 | 11505884 | 11506181 | Distal Intergenic |

| | | | |
|---|---|---|---|
| chr16 | 11511395 | 11511754 | Distal Intergenic |
| chr16 | 11511895 | 11512432 | Distal Intergenic |
| chr16 | 11776332 | 11776795 | Intron |
| chr16 | 11778270 | 11779177 | Intron |
| chr16 | 11867890 | 11869767 | Exon |
| chr16 | 11908596 | 11909805 | Distal Intergenic |
| chr16 | 12742072 | 12743948 | Distal Intergenic |
| chr16 | 13119107 | 13119565 | Intron |
| chr16 | 13159846 | 13160350 | Intron |
| chr16 | 13160481 | 13161716 | Intron |
| chr16 | 13240160 | 13241077 | Intron |
| chr16 | 13323313 | 13324441 | Intron |
| chr16 | 13350824 | 13352141 | Distal Intergenic |
| chr16 | 13366263 | 13366481 | Distal Intergenic |
| chr16 | 13366555 | 13367218 | Distal Intergenic |
| chr16 | 13367304 | 13367792 | Distal Intergenic |
| chr16 | 13370936 | 13372654 | Distal Intergenic |
| chr16 | 13381894 | 13382125 | Distal Intergenic |
| chr16 | 13667858 | 13668229 | Distal Intergenic |
| chr16 | 13672028 | 13672234 | Distal Intergenic |
| chr16 | 13780157 | 13781331 | Distal Intergenic |
| chr16 | 13819714 | 13819982 | Distal Intergenic |
| chr16 | 13825832 | 13827397 | Distal Intergenic |
| chr16 | 13872369 | 13872568 | Distal Intergenic |
| chr16 | 13903302 | 13903613 | Distal Intergenic |
| chr16 | 13968919 | 13969175 | Distal Intergenic |
| chr16 | 14159006 | 14159296 | Distal Intergenic |
| chr16 | 14159637 | 14159924 | Distal Intergenic |
| chr16 | 14162696 | 14163322 | Distal Intergenic |
| chr16 | 14163392 | 14163705 | Distal Intergenic |
| chr16 | 14163775 | 14164028 | Distal Intergenic |
| chr16 | 14177618 | 14177897 | Intron |
| chr16 | 14178153 | 14179331 | Intron |
| chr16 | 14316819 | 14317049 | Intron |
| chr16 | 14317115 | 14317332 | Intron |
| chr16 | 14317581 | 14318275 | Intron |
| chr16 | 14319513 | 14320423 | Intron |
| chr16 | 14350368 | 14353871 | Exon |
| chr16 | 14361196 | 14362776 | Distal Intergenic |
| chr16 | 14363177 | 14363376 | Distal Intergenic |
| chr16 | 14369465 | 14370924 | Distal Intergenic |
| chr16 | 14371532 | 14373976 | Distal Intergenic |
| chr16 | 14375755 | 14378649 | Distal Intergenic |
| chr16 | 14378786 | 14381643 | Exon |

| | | | |
|---|---|---|---|
| chr16 | 14416630 | 14417972 | Distal Intergenic |
| chr16 | 14704952 | 14705831 | Intron |
| chr16 | 14705937 | 14706408 | Intron |
| chr16 | 14706603 | 14707698 | Intron |
| chr16 | 14709261 | 14709460 | Intron |
| chr16 | 14799911 | 14800821 | Distal Intergenic |
| chr16 | 14883978 | 14886574 | Distal Intergenic |
| chr16 | 14926894 | 14929215 | Promoter |
| chr16 | 14929848 | 14930116 | Intron |
| chr16 | 14930155 | 14932269 | Exon |
| chr16 | 14932315 | 14934027 | Exon |
| chr16 | 14935736 | 14938875 | Exon |
| chr16 | 14939093 | 14941651 | Exon |
| chr16 | 14941786 | 14942172 | Intron |
| chr16 | 14942972 | 14944225 | Intron |
| chr16 | 14983805 | 14984678 | Intron |
| chr16 | 15112344 | 15116220 | Exon |
| chr16 | 15157684 | 15158062 | Intron |
| chr16 | 15250823 | 15251222 | Distal Intergenic |
| chr16 | 15251349 | 15252883 | Distal Intergenic |
| chr16 | 15354665 | 15355425 | Distal Intergenic |
| chr16 | 15355578 | 15358351 | Distal Intergenic |
| chr16 | 15358633 | 15358937 | Distal Intergenic |
| chr16 | 15360606 | 15360981 | Distal Intergenic |
| chr16 | 15493259 | 15494213 | Intron |
| chr16 | 15541759 | 15542150 | Intron |
| chr16 | 15546631 | 15547622 | Intron |
| chr16 | 15549942 | 15551170 | Intron |
| chr16 | 15551617 | 15551989 | Intron |
| chr16 | 15594006 | 15594223 | Intron |
| chr16 | 15594371 | 15594864 | Intron |
| chr16 | 15595296 | 15595675 | Promoter |
| chr16 | 15637253 | 15637452 | Intron |
| chr16 | 15641057 | 15641270 | Intron |
| chr16 | 15684854 | 15685808 | Downstream |
| chr16 | 15709936 | 15710174 | Intron |
| chr16 | 15984499 | 15984956 | Distal Intergenic |
| chr16 | 15991333 | 15991756 | Distal Intergenic |
| chr16 | 15991964 | 15992942 | Distal Intergenic |
| chr16 | 16060444 | 16061214 | Intron |
| chr16 | 16082558 | 16083612 | Intron |
| chr16 | 16215932 | 16217811 | Exon |
| chr16 | 16221806 | 16222289 | Intron |
| chr16 | 16223852 | 16225297 | Intron |

| | | | |
|---|---|---|---|
| chr16 | 16302667 | 16304145 | Exon |
| chr16 | 16359109 | 16359460 | Intron |
| chr16 | 16865950 | 16867032 | Distal Intergenic |
| chr16 | 16870385 | 16870777 | Distal Intergenic |
| chr16 | 16870966 | 16871439 | Distal Intergenic |
| chr16 | 16911891 | 16913178 | Distal Intergenic |
| chr16 | 16983505 | 16983901 | Distal Intergenic |
| chr16 | 16984225 | 16984653 | Distal Intergenic |
| chr16 | 16987804 | 16989604 | Distal Intergenic |
| chr16 | 17024298 | 17024506 | Distal Intergenic |
| chr16 | 17111073 | 17111316 | Distal Intergenic |
| chr16 | 17167575 | 17167807 | Distal Intergenic |
| chr16 | 17170039 | 17170293 | Distal Intergenic |
| chr16 | 17190753 | 17191313 | Distal Intergenic |
| chr16 | 17200473 | 17202171 | 3' UTR |
| chr16 | 17249133 | 17249729 | Intron |
| chr16 | 17451606 | 17451806 | Intron |
| chr16 | 17452074 | 17453218 | Intron |
| chr16 | 17453342 | 17453734 | Intron |
| chr16 | 17456568 | 17457808 | Intron |
| chr16 | 17508611 | 17509670 | Intron |
| chr16 | 17722023 | 17723040 | Distal Intergenic |
| chr16 | 17733076 | 17733757 | Distal Intergenic |
| chr16 | 17865563 | 17866023 | Distal Intergenic |
| chr16 | 17866110 | 17866724 | Distal Intergenic |
| chr16 | 17876498 | 17876868 | Distal Intergenic |
| chr16 | 17891163 | 17891362 | Distal Intergenic |
| chr16 | 17922067 | 17922958 | Distal Intergenic |
| chr16 | 17923434 | 17923635 | Distal Intergenic |
| chr16 | 17928313 | 17929093 | Distal Intergenic |
| chr16 | 17938306 | 17939323 | Distal Intergenic |
| chr16 | 17949122 | 17950494 | Distal Intergenic |
| chr16 | 18182848 | 18183517 | Distal Intergenic |
| chr16 | 18194603 | 18195472 | Distal Intergenic |
| chr16 | 18247811 | 18248790 | Distal Intergenic |
| chr16 | 18249091 | 18249722 | Distal Intergenic |
| chr16 | 18249766 | 18250022 | Distal Intergenic |
| chr16 | 18338536 | 18338904 | Distal Intergenic |
| chr16 | 18364026 | 18364231 | Distal Intergenic |
| chr16 | 18376386 | 18377488 | Distal Intergenic |
| chr16 | 18394855 | 18395411 | Distal Intergenic |
| chr16 | 18416138 | 18416337 | Intron |
| chr16 | 18416427 | 18416735 | Intron |
| chr16 | 18425764 | 18426064 | Intron |

| | | | |
|---|---|---|---|
| chr16 | 18426213 | 18426894 | Intron |
| chr16 | 18482521 | 18483351 | Distal Intergenic |
| chr16 | 18484279 | 18485760 | Exon |
| chr16 | 18498457 | 18498801 | Distal Intergenic |
| chr16 | 18547810 | 18548194 | Intron |
| chr16 | 18687076 | 18687547 | Distal Intergenic |
| chr16 | 18811536 | 18812663 | Promoter |
| chr16 | 18812711 | 18812987 | Promoter |
| chr16 | 18836400 | 18836888 | Intron |
| chr16 | 18876885 | 18877288 | Intron |
| chr16 | 18877489 | 18877702 | Intron |
| chr16 | 18877797 | 18878368 | Exon |
| chr16 | 18949847 | 18950900 | Distal Intergenic |
| chr16 | 19997566 | 19997934 | Distal Intergenic |
| chr16 | 20301491 | 20301715 | Distal Intergenic |
| chr16 | 20301826 | 20302385 | Distal Intergenic |
| chr16 | 20353789 | 20354051 | Intron |
| chr16 | 20354255 | 20354454 | Intron |
| chr16 | 20428663 | 20429531 | Exon |
| chr16 | 20429580 | 20430024 | Exon |
| chr16 | 20438878 | 20439359 | Exon |
| chr16 | 20445604 | 20447183 | Intron |
| chr16 | 20449738 | 20450095 | Intron |
| chr16 | 20476899 | 20477361 | Exon |
| chr16 | 20496504 | 20496797 | Intron |
| chr16 | 20499263 | 20499574 | Exon |
| chr16 | 20510988 | 20512052 | Distal Intergenic |
| chr16 | 20517137 | 20517710 | Distal Intergenic |
| chr16 | 20535032 | 20535441 | Distal Intergenic |
| chr16 | 20535656 | 20538422 | Distal Intergenic |
| chr16 | 20547241 | 20547576 | Downstream |
| chr16 | 20547665 | 20548517 | 3' UTR |
| chr16 | 20610833 | 20611184 | Distal Intergenic |
| chr16 | 20651740 | 20652165 | Exon |
| chr16 | 20673245 | 20673505 | Intron |
| chr16 | 20677748 | 20678275 | Intron |
| chr16 | 20716880 | 20717569 | Distal Intergenic |
| chr16 | 20718053 | 20718526 | Distal Intergenic |
| chr16 | 20910961 | 20911621 | Promoter |
| chr16 | 20931191 | 20931398 | Intron |
| chr16 | 20938187 | 20940359 | Distal Intergenic |
| chr16 | 20975219 | 20976518 | Exon |
| chr16 | 21138455 | 21139728 | Exon |
| chr16 | 21139833 | 21140667 | Intron |

| | | | |
|---|---|---|---|
| chr16 | 21151247 | 21152185 | Exon |
| chr16 | 21163305 | 21163521 | Intron |
| chr16 | 21207023 | 21209189 | 3' UTR |
| chr16 | 21209244 | 21210061 | Intron |
| chr16 | 21210142 | 21210912 | Exon |
| chr16 | 21211113 | 21211947 | Exon |
| chr16 | 21225712 | 21227311 | Distal Intergenic |
| chr16 | 21255959 | 21256221 | Intron |
| chr16 | 21332814 | 21333672 | Distal Intergenic |
| chr16 | 21339301 | 21339579 | Distal Intergenic |
| chr16 | 21361243 | 21361537 | Promoter |
| chr16 | 21361655 | 21362049 | Promoter |
| chr16 | 21381264 | 21383712 | Intron |
| chr16 | 21401719 | 21402766 | Distal Intergenic |
| chr16 | 21402948 | 21403719 | Distal Intergenic |
| chr16 | 21818466 | 21818888 | Intron |
| chr16 | 21998120 | 21998407 | Intron |
| chr16 | 21998490 | 21998689 | Intron |
| chr16 | 22009572 | 22010019 | Intron |
| chr16 | 22027758 | 22028057 | Intron |
| chr16 | 22077297 | 22077538 | Intron |
| chr16 | 22125413 | 22125710 | Intron |
| chr16 | 22151561 | 22151833 | Exon |
| chr16 | 22152044 | 22152387 | Intron |
| chr16 | 22153506 | 22153705 | Intron |
| chr16 | 22160888 | 22161260 | Exon |
| chr16 | 22161405 | 22161826 | Intron |
| chr16 | 22162197 | 22162627 | Intron |
| chr16 | 22162728 | 22162944 | Intron |
| chr16 | 22163605 | 22164093 | Exon |
| chr16 | 22201220 | 22202121 | Distal Intergenic |
| chr16 | 22263098 | 22263389 | Intron |
| chr16 | 22263660 | 22263859 | Intron |
| chr16 | 22264065 | 22266047 | Intron |
| chr16 | 22266138 | 22266971 | Intron |
| chr16 | 22365655 | 22366733 | Intron |
| chr16 | 22920077 | 22920277 | Intron |
| chr16 | 22920362 | 22920625 | Intron |
| chr16 | 23107218 | 23107421 | Intron |
| chr16 | 23107540 | 23109573 | Intron |
| chr16 | 23109612 | 23109916 | Intron |
| chr16 | 23109985 | 23110402 | Intron |
| chr16 | 23110472 | 23111066 | Intron |
| chr16 | 23127295 | 23127586 | Intron |

| | | | |
|---|---|---|---|
| chr16 | 23127709 | 23128068 | Intron |
| chr16 | 23128175 | 23128449 | Intron |
| chr16 | 23266611 | 23267049 | Distal Intergenic |
| chr16 | 23272364 | 23273006 | Distal Intergenic |
| chr16 | 23988521 | 23988738 | Intron |
| chr16 | 23998219 | 23998479 | Intron |
| chr16 | 24012359 | 24013476 | Intron |
| chr16 | 24037898 | 24038146 | Intron |
| chr16 | 24053239 | 24054716 | Intron |
| chr16 | 24055364 | 24056222 | Intron |
| chr16 | 24066429 | 24067655 | Intron |
| chr16 | 24092781 | 24093307 | Intron |
| chr16 | 24236306 | 24237647 | Distal Intergenic |
| chr16 | 24364462 | 24365094 | Intron |
| chr16 | 24392502 | 24393735 | Distal Intergenic |
| chr16 | 24393868 | 24395863 | Distal Intergenic |
| chr16 | 24401079 | 24401393 | Distal Intergenic |
| chr16 | 24427339 | 24428545 | Distal Intergenic |
| chr16 | 24430790 | 24431731 | Distal Intergenic |
| chr16 | 24433858 | 24435706 | Distal Intergenic |
| chr16 | 24443280 | 24444026 | Distal Intergenic |
| chr16 | 24474443 | 24474702 | Distal Intergenic |
| chr16 | 24474750 | 24476437 | Distal Intergenic |
| chr16 | 24495352 | 24496011 | Distal Intergenic |
| chr16 | 24496069 | 24496372 | Distal Intergenic |
| chr16 | 24497989 | 24498191 | Distal Intergenic |
| chr16 | 24498392 | 24498951 | Distal Intergenic |
| chr16 | 24566033 | 24567316 | Promoter |
| chr16 | 24678527 | 24679470 | Intron |
| chr16 | 24695227 | 24695712 | Distal Intergenic |
| chr16 | 24696523 | 24697561 | Distal Intergenic |
| chr16 | 24732866 | 24733925 | Distal Intergenic |
| chr16 | 24737288 | 24737987 | Distal Intergenic |
| chr16 | 24910714 | 24911287 | Intron |
| chr16 | 24913430 | 24914367 | Intron |
| chr16 | 24948561 | 24949481 | Promoter |
| chr16 | 24949524 | 24949817 | Intron |
| chr16 | 25011336 | 25011664 | Intron |
| chr16 | 25147326 | 25147942 | Intron |
| chr16 | 25281993 | 25283576 | Distal Intergenic |
| chr16 | 25384801 | 25385902 | Distal Intergenic |
| chr16 | 25407445 | 25407797 | Distal Intergenic |
| chr16 | 25407908 | 25408142 | Distal Intergenic |
| chr16 | 25465600 | 25466636 | Distal Intergenic |

| | | | |
|---|---|---|---|
| chr16 | 25483399 | 25484037 | Distal Intergenic |
| chr16 | 25484077 | 25486150 | Distal Intergenic |
| chr16 | 25531693 | 25532486 | Distal Intergenic |
| chr16 | 25584864 | 25585159 | Distal Intergenic |
| chr16 | 25615514 | 25616136 | Distal Intergenic |
| chr16 | 25623727 | 25626157 | Distal Intergenic |
| chr16 | 25724917 | 25726361 | Intron |
| chr16 | 25777757 | 25778324 | Intron |
| chr16 | 25778741 | 25778963 | Intron |
| chr16 | 25779032 | 25779260 | Intron |
| chr16 | 25779388 | 25780893 | Intron |
| chr16 | 25801619 | 25801852 | Intron |
| chr16 | 25801900 | 25806283 | Intron |
| chr16 | 25806748 | 25807018 | Intron |
| chr16 | 25807369 | 25807568 | Intron |
| chr16 | 25811863 | 25812687 | Intron |
| chr16 | 25832522 | 25832766 | Intron |
| chr16 | 25833348 | 25835832 | Intron |
| chr16 | 25835927 | 25837051 | Intron |
| chr16 | 25837855 | 25839046 | Intron |
| chr16 | 25839090 | 25839305 | Intron |
| chr16 | 25839523 | 25839851 | Intron |
| chr16 | 25841867 | 25845027 | Intron |
| chr16 | 25845165 | 25846231 | Intron |
| chr16 | 25849422 | 25849664 | Intron |
| chr16 | 25851629 | 25852150 | Intron |
| chr16 | 25853089 | 25853386 | Intron |
| chr16 | 25853436 | 25854203 | Intron |
| chr16 | 25859160 | 25859609 | Intron |
| chr16 | 25859662 | 25859943 | Intron |
| chr16 | 25860430 | 25861761 | Intron |
| chr16 | 25898627 | 25901432 | Intron |
| chr16 | 25901817 | 25903319 | Intron |
| chr16 | 25904361 | 25906796 | Intron |
| chr16 | 25912227 | 25913355 | Intron |
| chr16 | 25933883 | 25934763 | Intron |
| chr16 | 25941762 | 25943474 | Intron |
| chr16 | 25946785 | 25947211 | Intron |
| chr16 | 25947298 | 25949210 | Intron |
| chr16 | 25951427 | 25952227 | Intron |
| chr16 | 25953048 | 25954293 | Intron |
| chr16 | 26041175 | 26041988 | Intron |
| chr16 | 26053771 | 26054062 | Intron |
| chr16 | 26088567 | 26088802 | Intron |

| | | | |
|---|---|---|---|
| chr16 | 26146238 | 26148150 | 3' UTR |
| chr16 | 26211298 | 26211746 | Distal Intergenic |
| chr16 | 26248064 | 26248315 | Distal Intergenic |
| chr16 | 26248351 | 26249031 | Distal Intergenic |
| chr16 | 26430199 | 26430989 | Distal Intergenic |
| chr16 | 26444819 | 26445064 | Distal Intergenic |
| chr16 | 27365979 | 27367069 | Intron |
| chr16 | 27384929 | 27385132 | Distal Intergenic |
| chr16 | 27385267 | 27386165 | Distal Intergenic |
| chr16 | 27388751 | 27389017 | Distal Intergenic |
| chr16 | 27389097 | 27390090 | Distal Intergenic |
| chr16 | 28870234 | 28871491 | Intron |
| chr16 | 28871877 | 28872262 | Intron |
| chr16 | 28878398 | 28882665 | 5' UTR |
| chr16 | 28882707 | 28883635 | Exon |
| chr16 | 28883848 | 28886040 | 3' UTR |
| chr16 | 28929325 | 28929757 | Intron |
| chr16 | 28930086 | 28930486 | Intron |
| chr16 | 29169840 | 29170041 | Intron |
| chr16 | 29775431 | 29776720 | Intron |
| chr16 | 29814490 | 29815076 | Exon |
| chr16 | 29815285 | 29816616 | Promoter |
| chr16 | 29822593 | 29824319 | Promoter |
| chr16 | 29903142 | 29903624 | Intron |
| chr16 | 29903675 | 29904108 | Intron |
| chr16 | 29909898 | 29912280 | Promoter |
| chr16 | 29915218 | 29915459 | Intron |
| chr16 | 29915591 | 29916301 | Exon |
| chr16 | 29944049 | 29945388 | Intron |
| chr16 | 29945570 | 29946931 | Intron |
| chr16 | 29977753 | 29978376 | Intron |
| chr16 | 30024428 | 30025746 | Promoter |
| chr16 | 30033834 | 30034787 | Promoter |
| chr16 | 30043161 | 30043391 | Promoter |
| chr16 | 30056815 | 30058759 | Intron |
| chr16 | 30058970 | 30060442 | Intron |
| chr16 | 30060480 | 30060820 | Intron |
| chr16 | 30062458 | 30063425 | Promoter |
| chr16 | 30064520 | 30065148 | Promoter |
| chr16 | 30065363 | 30067037 | Promoter |
| chr16 | 30067198 | 30067584 | Intron |
| chr16 | 30068264 | 30068523 | Intron |
| chr16 | 30068722 | 30069311 | Intron |
| chr16 | 30069373 | 30070284 | Intron |

| | | | |
|---|---|---|---|
| chr16 | 30070783 | 30071077 | Intron |
| chr16 | 30133390 | 30133694 | Promoter |
| chr16 | 30187672 | 30188258 | Intron |
| chr16 | 30332312 | 30333328 | Intron |
| chr16 | 30356295 | 30358017 | Distal Intergenic |
| chr16 | 30375135 | 30376075 | Intron |
| chr16 | 30387257 | 30387525 | Exon |
| chr16 | 30387752 | 30388379 | Exon |
| chr16 | 30388968 | 30389233 | Promoter |
| chr16 | 30439481 | 30440583 | Promoter |
| chr16 | 30442058 | 30443196 | Promoter |
| chr16 | 30465146 | 30465345 | Distal Intergenic |
| chr16 | 30465478 | 30466564 | Distal Intergenic |
| chr16 | 30499733 | 30500001 | Intron |
| chr16 | 30500258 | 30500679 | Exon |
| chr16 | 30502301 | 30504566 | Intron |
| chr16 | 30505305 | 30505926 | Exon |
| chr16 | 30510816 | 30514696 | Promoter |
| chr16 | 30517660 | 30519511 | Exon |
| chr16 | 30524477 | 30526826 | Exon |
| chr16 | 30527385 | 30527589 | Intron |
| chr16 | 30527636 | 30529260 | Exon |
| chr16 | 30572359 | 30572715 | Distal Intergenic |
| chr16 | 30587136 | 30588308 | Intron |
| chr16 | 30597315 | 30597916 | Promoter |
| chr16 | 30771094 | 30773000 | Promoter |
| chr16 | 30780591 | 30780995 | Exon |
| chr16 | 30835522 | 30836592 | Distal Intergenic |
| chr16 | 30955999 | 30957144 | Intron |
| chr16 | 31066994 | 31067233 | Distal Intergenic |
| chr16 | 31185845 | 31186079 | Distal Intergenic |
| chr16 | 31189358 | 31189672 | Distal Intergenic |
| chr16 | 31191259 | 31191599 | Promoter |
| chr16 | 31191637 | 31193220 | Promoter |
| chr16 | 31200896 | 31201154 | Exon |
| chr16 | 31233953 | 31234505 | Exon |
| chr16 | 31255897 | 31257079 | Distal Intergenic |
| chr16 | 31274454 | 31275837 | Intron |
| chr16 | 31290381 | 31291291 | Intron |
| chr16 | 31291378 | 31291778 | Intron |
| chr16 | 31407562 | 31408697 | Exon |
| chr16 | 31506450 | 31507641 | Intron |
| chr16 | 31526803 | 31527092 | Distal Intergenic |
| chr16 | 31592100 | 31593949 | Distal Intergenic |

| | | | |
|---|---|---|---|
| chr16 | 31598145 | 31599581 | Distal Intergenic |
| chr16 | 31948464 | 31949233 | Distal Intergenic |
| chr16 | 31979363 | 31980166 | Distal Intergenic |
| chr16 | 31994137 | 31994949 | Distal Intergenic |
| chr16 | 32002804 | 32004103 | Distal Intergenic |
| chr16 | 32005547 | 32006353 | Distal Intergenic |
| chr16 | 32008663 | 32008889 | Distal Intergenic |
| chr16 | 32010100 | 32010597 | Distal Intergenic |
| chr16 | 32045981 | 32046957 | Distal Intergenic |
| chr16 | 32058907 | 32059632 | Intron |
| chr16 | 32068269 | 32069812 | Intron |
| chr16 | 32077393 | 32079412 | Exon |
| chr16 | 32231611 | 32232633 | Distal Intergenic |
| chr16 | 32260901 | 32261624 | Distal Intergenic |
| chr16 | 32277030 | 32278550 | Distal Intergenic |
| chr16 | 32332474 | 32332692 | Distal Intergenic |
| chr16 | 32332735 | 32333551 | Distal Intergenic |
| chr16 | 32640285 | 32640690 | Distal Intergenic |
| chr16 | 32641656 | 32642185 | Distal Intergenic |
| chr16 | 32644861 | 32646658 | Distal Intergenic |
| chr16 | 32657013 | 32657281 | Distal Intergenic |
| chr16 | 32671885 | 32673549 | Distal Intergenic |
| chr16 | 32675969 | 32677688 | Distal Intergenic |
| chr16 | 32821967 | 32823547 | Distal Intergenic |
| chr16 | 32858148 | 32860485 | Distal Intergenic |
| chr16 | 32863471 | 32863755 | Distal Intergenic |
| chr16 | 32863821 | 32864257 | Distal Intergenic |
| chr16 | 32867538 | 32868479 | Distal Intergenic |
| chr16 | 32914468 | 32915611 | Exon |
| chr16 | 32915723 | 32916165 | Intron |
| chr16 | 33056253 | 33058291 | Intron |
| chr16 | 33061612 | 33062456 | Intron |
| chr16 | 33129713 | 33131760 | Intron |
| chr16 | 33185655 | 33186046 | Intron |
| chr16 | 33379882 | 33381122 | Distal Intergenic |
| chr16 | 33381524 | 33382184 | Distal Intergenic |
| chr16 | 33382278 | 33382514 | Distal Intergenic |
| chr16 | 33394440 | 33394801 | Distal Intergenic |
| chr16 | 33400009 | 33400959 | Distal Intergenic |
| chr16 | 33410972 | 33411182 | Distal Intergenic |
| chr16 | 33411276 | 33411783 | Distal Intergenic |
| chr16 | 33667477 | 33667890 | Distal Intergenic |
| chr16 | 33830127 | 33831308 | Distal Intergenic |
| chr16 | 33832429 | 33832682 | Distal Intergenic |

| | | | |
|---|---|---|---|
| chr16 | 33832735 | 33834414 | Distal Intergenic |
| chr16 | 33836103 | 33837953 | Distal Intergenic |
| chr16 | 33852962 | 33853814 | Distal Intergenic |
| chr16 | 33863138 | 33863671 | Distal Intergenic |
| chr16 | 33868999 | 33874248 | Distal Intergenic |
| chr16 | 33877504 | 33879443 | Distal Intergenic |
| chr16 | 33883634 | 33884137 | Distal Intergenic |
| chr16 | 33932937 | 33935147 | Distal Intergenic |
| chr16 | 33940900 | 33941184 | Distal Intergenic |
| chr16 | 33941232 | 33941722 | Distal Intergenic |
| chr16 | 34602930 | 34603451 | Intron |
| chr16 | 34603746 | 34604087 | Intron |
| chr16 | 35022999 | 35023760 | Distal Intergenic |
| chr16 | 35033804 | 35034293 | Distal Intergenic |
| chr16 | 35046679 | 35048443 | Distal Intergenic |
| chr16 | 35363606 | 35363906 | Distal Intergenic |
| chr16 | 35938995 | 35939596 | Distal Intergenic |
| chr16 | 36068834 | 36072946 | Distal Intergenic |
| chr16 | 36089737 | 36091885 | Distal Intergenic |
| chr16 | 36095388 | 36096211 | Distal Intergenic |
| chr16 | 36130520 | 36131767 | Distal Intergenic |
| chr16 | 36132829 | 36136359 | Distal Intergenic |
| chr16 | 36241301 | 36242057 | Distal Intergenic |
| chr16 | 36335286 | 36335841 | Distal Intergenic |
| chr16 | 36455046 | 36455447 | Distal Intergenic |
| chr16 | 36455513 | 36455730 | Distal Intergenic |
| chr16 | 36874624 | 36875081 | Distal Intergenic |
| chr16 | 36875212 | 36875416 | Distal Intergenic |
| chr16 | 37057168 | 37058738 | Distal Intergenic |
| chr16 | 37440478 | 37441053 | Distal Intergenic |
| chr16 | 37539302 | 37539521 | Distal Intergenic |
| chr16 | 37539566 | 37540585 | Distal Intergenic |
| chr16 | 37654043 | 37654427 | Distal Intergenic |
| chr16 | 37675579 | 37676667 | Distal Intergenic |
| chr16 | 37704708 | 37705583 | Distal Intergenic |
| chr16 | 37725588 | 37726252 | Distal Intergenic |
| chr16 | 37728497 | 37729139 | Distal Intergenic |
| chr16 | 37787794 | 37791731 | Distal Intergenic |
| chr16 | 37800667 | 37802182 | Distal Intergenic |
| chr16 | 37868704 | 37869381 | Distal Intergenic |
| chr16 | 37940588 | 37940812 | Distal Intergenic |
| chr16 | 38012478 | 38012933 | Distal Intergenic |
| chr16 | 38017103 | 38018764 | Distal Intergenic |
| chr16 | 38088003 | 38088406 | Distal Intergenic |

| | | | |
|---|---|---|---|
| chr16 | 38089149 | 38089548 | Distal Intergenic |
| chr16 | 38089690 | 38091106 | Distal Intergenic |
| chr16 | 38092097 | 38092347 | Distal Intergenic |
| chr16 | 38119290 | 38119650 | Distal Intergenic |
| chr16 | 38138571 | 38138774 | Distal Intergenic |
| chr16 | 38160092 | 38161027 | Distal Intergenic |
| chr16 | 38162001 | 38162206 | Distal Intergenic |
| chr16 | 38204783 | 38205948 | Distal Intergenic |
| chr16 | 38346885 | 38347984 | Distal Intergenic |
| chr16 | 38348093 | 38348487 | Distal Intergenic |
| chr16 | 38368383 | 38368656 | Distal Intergenic |
| chr16 | 38370341 | 38372428 | Distal Intergenic |
| chr16 | 38393797 | 38394946 | Distal Intergenic |
| chr16 | 38395011 | 38395666 | Distal Intergenic |
| chr16 | 38395791 | 38397033 | Distal Intergenic |
| chr16 | 38402755 | 38404859 | Distal Intergenic |
| chr16 | 38406282 | 38407329 | Distal Intergenic |
| chr16 | 38452467 | 38453436 | Distal Intergenic |
| chr16 | 38458093 | 38460614 | Distal Intergenic |
| chr16 | 38461110 | 38461406 | Distal Intergenic |
| chr16 | 38521959 | 38522789 | Distal Intergenic |
| chr16 | 38522848 | 38523443 | Distal Intergenic |
| chr16 | 38549872 | 38550108 | Distal Intergenic |
| chr16 | 38550204 | 38550408 | Distal Intergenic |
| chr16 | 38562642 | 38562869 | Distal Intergenic |
| chr16 | 38562909 | 38563181 | Distal Intergenic |
| chr16 | 38563342 | 38563569 | Distal Intergenic |
| chr16 | 42297722 | 42298410 | Distal Intergenic |
| chr16 | 42385575 | 42386093 | Distal Intergenic |
| chr16 | 42440181 | 42441655 | Distal Intergenic |
| chr16 | 42442662 | 42443502 | Distal Intergenic |
| chr16 | 42443770 | 42443985 | Distal Intergenic |
| chr16 | 42452467 | 42452791 | Distal Intergenic |
| chr16 | 42452858 | 42456025 | Distal Intergenic |
| chr16 | 42478876 | 42481241 | Distal Intergenic |
| chr16 | 42498898 | 42500330 | Distal Intergenic |
| chr16 | 42585565 | 42587532 | Distal Intergenic |
| chr16 | 42638622 | 42639012 | Distal Intergenic |
| chr16 | 42648356 | 42653298 | Distal Intergenic |
| chr16 | 42661570 | 42663251 | Distal Intergenic |
| chr16 | 42663293 | 42663816 | Distal Intergenic |
| chr16 | 42674951 | 42675211 | Distal Intergenic |
| chr16 | 42675663 | 42675915 | Distal Intergenic |
| chr16 | 42675967 | 42678632 | Distal Intergenic |

| | | | |
|---|---|---|---|
| chr16 | 42745872 | 42746800 | Distal Intergenic |
| chr16 | 42781433 | 42782969 | Distal Intergenic |
| chr16 | 42874900 | 42875271 | Distal Intergenic |
| chr16 | 43252708 | 43253696 | Distal Intergenic |
| chr16 | 43256971 | 43257387 | Distal Intergenic |
| chr16 | 43261171 | 43261685 | Distal Intergenic |
| chr16 | 43458405 | 43460090 | Distal Intergenic |
| chr16 | 43463801 | 43466216 | Distal Intergenic |
| chr16 | 43655011 | 43655211 | Distal Intergenic |
| chr16 | 43663510 | 43665504 | Distal Intergenic |
| chr16 | 43697532 | 43700340 | Distal Intergenic |
| chr16 | 43701361 | 43701560 | Distal Intergenic |
| chr16 | 43701697 | 43702411 | Distal Intergenic |
| chr16 | 43889046 | 43890482 | Distal Intergenic |
| chr16 | 44100910 | 44103018 | Distal Intergenic |
| chr16 | 44131489 | 44133643 | Distal Intergenic |
| chr16 | 44136987 | 44137186 | Distal Intergenic |
| chr16 | 44138455 | 44138655 | Distal Intergenic |
| chr16 | 44138836 | 44140538 | Distal Intergenic |
| chr16 | 44140800 | 44141149 | Distal Intergenic |
| chr16 | 44724408 | 44725663 | Distal Intergenic |
| chr16 | 44726017 | 44726326 | Distal Intergenic |
| chr16 | 44745409 | 44745765 | Distal Intergenic |
| chr16 | 44746210 | 44746422 | Distal Intergenic |
| chr16 | 45110567 | 45110952 | Distal Intergenic |
| chr16 | 45110994 | 45111327 | Distal Intergenic |
| chr16 | 45124168 | 45125056 | Distal Intergenic |
| chr16 | 45158851 | 45159722 | Distal Intergenic |
| chr16 | 45336023 | 45336732 | Distal Intergenic |
| chr16 | 45355970 | 45357129 | Distal Intergenic |
| chr16 | 45360686 | 45361005 | Distal Intergenic |
| chr16 | 45363072 | 45363344 | Distal Intergenic |
| chr16 | 45363386 | 45363729 | Distal Intergenic |
| chr16 | 45376949 | 45380315 | Distal Intergenic |
| chr16 | 45383862 | 45384365 | Distal Intergenic |
| chr16 | 45384683 | 45384882 | Distal Intergenic |
| chr16 | 45742604 | 45743081 | Distal Intergenic |
| chr16 | 45837555 | 45839226 | Distal Intergenic |
| chr16 | 45839579 | 45839940 | Distal Intergenic |
| chr16 | 45843150 | 45845541 | Distal Intergenic |
| chr16 | 45848759 | 45849837 | Distal Intergenic |
| chr16 | 45877593 | 45877824 | Distal Intergenic |
| chr16 | 45886546 | 45887829 | Distal Intergenic |
| chr16 | 45892531 | 45892931 | Distal Intergenic |

| | | | |
|---|---|---|---|
| chr16 | 45893006 | 45893711 | Distal Intergenic |
| chr16 | 45902990 | 45904408 | Distal Intergenic |
| chr16 | 45930337 | 45931272 | Distal Intergenic |
| chr16 | 45944761 | 45947204 | Distal Intergenic |
| chr16 | 45952514 | 45952857 | Distal Intergenic |
| chr16 | 48992857 | 48995340 | Distal Intergenic |
| chr16 | 49276340 | 49276574 | Distal Intergenic |
| chr16 | 49276629 | 49277033 | Distal Intergenic |
| chr16 | 49277085 | 49278339 | Distal Intergenic |
| chr16 | 49288562 | 49290327 | Distal Intergenic |
| chr16 | 49507498 | 49509735 | Distal Intergenic |
| chr16 | 49854910 | 49855651 | Promoter |
| chr16 | 49856063 | 49856473 | Promoter |
| chr16 | 49856513 | 49859720 | Promoter |
| chr16 | 49864151 | 49864437 | Intron |
| chr16 | 49864473 | 49864695 | Intron |
| chr16 | 49904525 | 49904758 | Distal Intergenic |
| chr16 | 49905041 | 49906481 | Distal Intergenic |
| chr16 | 49914788 | 49915033 | Distal Intergenic |
| chr16 | 49915086 | 49917092 | Distal Intergenic |
| chr16 | 49923665 | 49923924 | Distal Intergenic |
| chr16 | 49935931 | 49939865 | Distal Intergenic |
| chr16 | 49951170 | 49951837 | Distal Intergenic |
| chr16 | 49951954 | 49953073 | Distal Intergenic |
| chr16 | 49969648 | 49971866 | Distal Intergenic |
| chr16 | 50009519 | 50009797 | Distal Intergenic |
| chr16 | 50188230 | 50188429 | Promoter |
| chr16 | 50188659 | 50188872 | Intron |
| chr16 | 50288875 | 50289346 | Distal Intergenic |
| chr16 | 50289552 | 50290320 | Distal Intergenic |
| chr16 | 50302878 | 50303135 | Intron |
| chr16 | 50364529 | 50365131 | Intron |
| chr16 | 50432272 | 50432486 | Distal Intergenic |
| chr16 | 50462183 | 50462470 | Distal Intergenic |
| chr16 | 50528356 | 50529753 | Distal Intergenic |
| chr16 | 50529797 | 50530211 | Distal Intergenic |
| chr16 | 50731995 | 50732431 | Promoter |
| chr16 | 50737172 | 50737839 | Intron |
| chr16 | 51811060 | 51811847 | Distal Intergenic |
| chr16 | 51918246 | 51918793 | Distal Intergenic |
| chr16 | 51926746 | 51927003 | Distal Intergenic |
| chr16 | 51999737 | 52000862 | Distal Intergenic |
| chr16 | 52000906 | 52001177 | Distal Intergenic |
| chr16 | 52030935 | 52031201 | Distal Intergenic |

| | | | |
|---|---|---|---|
| chr16 | 52031343 | 52031605 | Distal Intergenic |
| chr16 | 52031671 | 52032263 | Distal Intergenic |
| chr16 | 52079080 | 52080377 | Intron |
| chr16 | 52134844 | 52136183 | Distal Intergenic |
| chr16 | 52179708 | 52180838 | Distal Intergenic |
| chr16 | 52196314 | 52197622 | Distal Intergenic |
| chr16 | 54041729 | 54042198 | Intron |
| chr16 | 54263650 | 54263902 | Distal Intergenic |
| chr16 | 55298139 | 55299594 | Distal Intergenic |
| chr16 | 55533046 | 55533396 | Intron |
| chr16 | 55540218 | 55541142 | 3' UTR |
| chr16 | 55555121 | 55556524 | Intron |
| chr16 | 55556578 | 55557091 | Intron |
| chr16 | 55608195 | 55608559 | Exon |
| chr16 | 55608616 | 55609103 | Exon |
| chr16 | 55636112 | 55637184 | Distal Intergenic |
| chr16 | 55770199 | 55770412 | Exon |
| chr16 | 55783175 | 55784896 | Exon |
| chr16 | 55785079 | 55785425 | Distal Intergenic |
| chr16 | 55787198 | 55789348 | Distal Intergenic |
| chr16 | 55789593 | 55790096 | Distal Intergenic |
| chr16 | 55819808 | 55822302 | Distal Intergenic |
| chr16 | 55950916 | 55951216 | Intron |
| chr16 | 55966187 | 55967738 | Intron |
| chr16 | 55970451 | 55970783 | Intron |
| chr16 | 55972448 | 55972651 | Intron |
| chr16 | 55973233 | 55975148 | Intron |
| chr16 | 56027921 | 56028131 | Distal Intergenic |
| chr16 | 56028366 | 56028595 | Distal Intergenic |
| chr16 | 56028669 | 56030192 | Distal Intergenic |
| chr16 | 56037063 | 56037716 | Distal Intergenic |
| chr16 | 56716639 | 56716972 | Promoter |
| chr16 | 56717756 | 56718341 | 3' UTR |
| chr16 | 56852670 | 56853567 | Intron |
| chr16 | 56869859 | 56874438 | Exon |
| chr16 | 57121806 | 57122076 | Distal Intergenic |
| chr16 | 57122130 | 57122701 | Distal Intergenic |
| chr16 | 57122934 | 57123328 | Distal Intergenic |
| chr16 | 57167535 | 57167957 | Intron |
| chr16 | 57226379 | 57227386 | Intron |
| chr16 | 57383497 | 57384332 | Distal Intergenic |
| chr16 | 57418811 | 57422885 | 3' UTR |
| chr16 | 57424762 | 57424987 | Distal Intergenic |
| chr16 | 57425331 | 57425533 | Distal Intergenic |

| | | | |
|---|---|---|---|
| chr16 | 57427128 | 57428849 | Distal Intergenic |
| chr16 | 57452583 | 57452886 | Distal Intergenic |
| chr16 | 57452974 | 57453469 | Distal Intergenic |
| chr16 | 57453666 | 57454975 | Distal Intergenic |
| chr16 | 57467202 | 57468211 | 3' UTR |
| chr16 | 57525196 | 57526480 | Distal Intergenic |
| chr16 | 57526749 | 57527766 | Distal Intergenic |
| chr16 | 57528227 | 57528612 | Distal Intergenic |
| chr16 | 57531991 | 57532926 | Distal Intergenic |
| chr16 | 57536005 | 57536273 | Distal Intergenic |
| chr16 | 57536402 | 57537398 | Distal Intergenic |
| chr16 | 57543378 | 57543664 | Downstream |
| chr16 | 57545552 | 57547644 | 3' UTR |
| chr16 | 57548909 | 57552168 | Exon |
| chr16 | 57552252 | 57553680 | Exon |
| chr16 | 57553787 | 57554559 | Intron |
| chr16 | 57554734 | 57555626 | Exon |
| chr16 | 57555790 | 57558699 | Intron |
| chr16 | 57558972 | 57560025 | Exon |
| chr16 | 57560219 | 57561018 | Intron |
| chr16 | 57561082 | 57561457 | Intron |
| chr16 | 57561877 | 57567211 | 5' UTR |
| chr16 | 57671156 | 57671514 | Intron |
| chr16 | 57671724 | 57672924 | Promoter |
| chr16 | 57745862 | 57746112 | Intron |
| chr16 | 57746862 | 57753955 | Exon |
| chr16 | 57754061 | 57754330 | Intron |
| chr16 | 57757417 | 57757776 | Intron |
| chr16 | 58352048 | 58352751 | Distal Intergenic |
| chr16 | 58409183 | 58409425 | Distal Intergenic |
| chr16 | 58553237 | 58553977 | 3' UTR |
| chr16 | 58586604 | 58587357 | Promoter |
| chr16 | 58590898 | 58592180 | Intron |
| chr16 | 58669151 | 58670541 | Distal Intergenic |
| chr16 | 58670769 | 58671423 | Distal Intergenic |
| chr16 | 58671519 | 58671866 | Distal Intergenic |
| chr16 | 58701687 | 58702900 | Intron |
| chr16 | 58703040 | 58703526 | Intron |
| chr16 | 58703662 | 58704323 | Exon |
| chr16 | 58728071 | 58729101 | Distal Intergenic |
| chr16 | 58745337 | 58747662 | Intron |
| chr16 | 59472523 | 59472722 | Distal Intergenic |
| chr16 | 59474187 | 59474592 | Distal Intergenic |
| chr16 | 59474819 | 59475623 | Distal Intergenic |

| | | | |
|---|---|---|---|
| chr16 | 59487070 | 59488343 | Distal Intergenic |
| chr16 | 59547430 | 59548776 | Distal Intergenic |
| chr16 | 59565957 | 59566997 | Distal Intergenic |
| chr16 | 59583737 | 59584047 | Distal Intergenic |
| chr16 | 59600152 | 59600560 | Distal Intergenic |
| chr16 | 62729432 | 62729705 | Distal Intergenic |
| chr16 | 62729906 | 62730747 | Distal Intergenic |
| chr16 | 62786198 | 62786507 | Distal Intergenic |
| chr16 | 62798249 | 62798617 | Distal Intergenic |
| chr16 | 62798701 | 62800237 | Distal Intergenic |
| chr16 | 62846234 | 62846784 | Distal Intergenic |
| chr16 | 64758846 | 64759250 | Distal Intergenic |
| chr16 | 64774995 | 64776252 | Distal Intergenic |
| chr16 | 64779117 | 64779763 | Distal Intergenic |
| chr16 | 64796453 | 64796739 | Distal Intergenic |
| chr16 | 64797073 | 64797409 | Distal Intergenic |
| chr16 | 64850512 | 64850926 | Distal Intergenic |
| chr16 | 64851213 | 64851616 | Distal Intergenic |
| chr16 | 64851789 | 64852041 | Distal Intergenic |
| chr16 | 64873179 | 64874534 | Distal Intergenic |
| chr16 | 65562102 | 65562471 | Intron |
| chr16 | 65562707 | 65562907 | Intron |
| chr16 | 65814753 | 65815035 | Distal Intergenic |
| chr16 | 65815487 | 65815827 | Distal Intergenic |
| chr16 | 65816035 | 65817738 | Distal Intergenic |
| chr16 | 65817875 | 65818240 | Distal Intergenic |
| chr16 | 65818505 | 65819898 | Distal Intergenic |
| chr16 | 65822304 | 65822575 | Distal Intergenic |
| chr16 | 65868694 | 65870210 | Distal Intergenic |
| chr16 | 65870281 | 65870504 | Distal Intergenic |
| chr16 | 66921404 | 66921656 | 3' UTR |
| chr16 | 66921884 | 66922721 | 3' UTR |
| chr16 | 66922813 | 66923086 | 3' UTR |
| chr16 | 68205287 | 68205730 | Intron |
| chr16 | 69382054 | 69382281 | Intron |
| chr16 | 69577016 | 69578202 | Distal Intergenic |
| chr16 | 69943788 | 69944508 | 3' UTR |
| chr16 | 70007071 | 70007854 | Intron |
| chr16 | 70307025 | 70307259 | Intron |
| chr16 | 70310160 | 70311752 | Exon |
| chr16 | 70330722 | 70331266 | Intron |
| chr16 | 70492480 | 70493015 | Intron |
| chr16 | 70495867 | 70496649 | Intron |
| chr16 | 70523813 | 70525179 | Exon |

| | | | |
|---|---|---|---|
| chr16 | 70525216 | 70525458 | Intron |
| chr16 | 71511989 | 71512236 | 5' UTR |
| chr16 | 72688611 | 72689272 | Distal Intergenic |
| chr16 | 73770376 | 73771582 | Distal Intergenic |
| chr16 | 73856500 | 73856761 | Distal Intergenic |
| chr16 | 74095589 | 74096270 | Distal Intergenic |
| chr16 | 74135116 | 74135433 | Distal Intergenic |
| chr16 | 74159783 | 74159987 | Distal Intergenic |
| chr16 | 74160647 | 74161208 | Distal Intergenic |
| chr16 | 74388946 | 74389330 | Intron |
| chr16 | 74389451 | 74390230 | Intron |
| chr16 | 74400865 | 74401825 | Promoter |
| chr16 | 74519284 | 74519495 | Intron |
| chr16 | 74526472 | 74527635 | Exon |
| chr16 | 74530779 | 74531042 | Intron |
| chr16 | 74531923 | 74532129 | Intron |
| chr16 | 74609361 | 74610438 | Intron |
| chr16 | 74610523 | 74612836 | Intron |
| chr16 | 74937293 | 74937615 | Promoter |
| chr16 | 75753118 | 75753326 | Distal Intergenic |
| chr16 | 75836319 | 75838109 | Distal Intergenic |
| chr16 | 75838189 | 75838388 | Distal Intergenic |
| chr16 | 76201314 | 76202356 | Distal Intergenic |
| chr16 | 76296041 | 76296556 | Distal Intergenic |
| chr16 | 76296699 | 76296913 | Distal Intergenic |
| chr16 | 76371184 | 76371695 | Intron |
| chr16 | 76371745 | 76372686 | Intron |
| chr16 | 76373442 | 76373957 | Intron |
| chr16 | 76471623 | 76472194 | Intron |
| chr16 | 76472251 | 76472890 | Intron |
| chr16 | 76499762 | 76500033 | Intron |
| chr16 | 76500082 | 76500323 | Intron |
| chr16 | 76525698 | 76526895 | Intron |
| chr16 | 76539686 | 76542329 | Intron |
| chr16 | 76621814 | 76622174 | Distal Intergenic |
| chr16 | 76958804 | 76959052 | Distal Intergenic |
| chr16 | 76959102 | 76959337 | Distal Intergenic |
| chr16 | 76959433 | 76959735 | Distal Intergenic |
| chr16 | 77005322 | 77006073 | Distal Intergenic |
| chr16 | 77014883 | 77015112 | Distal Intergenic |
| chr16 | 77015150 | 77015423 | Distal Intergenic |
| chr16 | 77015475 | 77015689 | Distal Intergenic |
| chr16 | 78302515 | 78303199 | Intron |
| chr16 | 78318143 | 78318364 | Intron |

| | | | |
|---|---|---|---|
| chr16 | 78423017 | 78423733 | Intron |
| chr16 | 78423788 | 78424102 | Intron |
| chr16 | 78456271 | 78457257 | Intron |
| chr16 | 78473375 | 78474619 | Intron |
| chr16 | 78504121 | 78504469 | Intron |
| chr16 | 78505005 | 78505286 | Intron |
| chr16 | 78531896 | 78533108 | Intron |
| chr16 | 78563285 | 78565141 | Intron |
| chr16 | 78575069 | 78576392 | Intron |
| chr16 | 78576695 | 78576929 | Intron |
| chr16 | 78577251 | 78577739 | Intron |
| chr16 | 78619010 | 78619209 | Intron |
| chr16 | 78619328 | 78620127 | Intron |
| chr16 | 78631250 | 78633227 | Intron |
| chr16 | 79570329 | 79571240 | Distal Intergenic |
| chr16 | 79571309 | 79571723 | Distal Intergenic |
| chr16 | 79591026 | 79591378 | Distal Intergenic |
| chr16 | 79839528 | 79840398 | Distal Intergenic |
| chr16 | 84807329 | 84807555 | Intron |
| chr16 | 84834614 | 84835806 | Distal Intergenic |
| chr16 | 84835959 | 84836585 | Distal Intergenic |
| chr16 | 84838294 | 84839165 | Distal Intergenic |
| chr16 | 85134108 | 85134549 | Intron |
| chr16 | 85137164 | 85138546 | Intron |
| chr16 | 85138607 | 85138831 | Intron |
| chr16 | 85141164 | 85144310 | Exon |
| chr16 | 85144841 | 85145363 | Promoter |
| chr16 | 85147879 | 85150469 | Distal Intergenic |
| chr16 | 85152395 | 85153652 | Distal Intergenic |
| chr16 | 85158652 | 85159680 | Distal Intergenic |
| chr16 | 85171697 | 85172008 | Promoter |
| chr16 | 85172105 | 85172786 | Intron |
| chr16 | 85172910 | 85173777 | Intron |
| chr16 | 85195730 | 85197799 | Distal Intergenic |
| chr16 | 85297709 | 85297921 | Distal Intergenic |
| chr16 | 85422875 | 85423308 | Distal Intergenic |
| chr16 | 85445560 | 85446864 | Distal Intergenic |
| chr16 | 85801167 | 85801523 | Distal Intergenic |
| chr16 | 85803571 | 85803797 | Distal Intergenic |
| chr16 | 85804147 | 85804350 | Distal Intergenic |
| chr16 | 86138272 | 86138539 | Distal Intergenic |
| chr16 | 86268709 | 86269277 | Distal Intergenic |
| chr16 | 86269391 | 86269645 | Distal Intergenic |
| chr16 | 86346658 | 86346987 | Distal Intergenic |

| | | | |
|---|---|---|---|
| chr16 | 86347033 | 86348932 | Distal Intergenic |
| chr16 | 86853258 | 86854154 | Distal Intergenic |
| chr16 | 87043956 | 87044398 | Distal Intergenic |
| chr16 | 87085673 | 87086423 | Distal Intergenic |
| chr16 | 87103560 | 87104141 | Distal Intergenic |
| chr16 | 87214786 | 87215360 | Intron |
| chr16 | 87354369 | 87354643 | Distal Intergenic |
| chr16 | 87440634 | 87441126 | 3' UTR |
| chr16 | 87495545 | 87495925 | Intron |
| chr16 | 87599846 | 87600105 | Distal Intergenic |
| chr16 | 87625201 | 87626335 | Distal Intergenic |
| chr16 | 87656924 | 87657624 | Intron |
| chr16 | 87658182 | 87658484 | Intron |
| chr16 | 87658539 | 87659495 | Intron |
| chr16 | 87659546 | 87659790 | Intron |
| chr16 | 87661174 | 87663940 | Intron |
| chr16 | 87699246 | 87700719 | Intron |
| chr16 | 87712863 | 87713847 | Intron |
| chr16 | 87784096 | 87785275 | Promoter |
| chr16 | 88478650 | 88478879 | Distal Intergenic |
| chr16 | 88505172 | 88505513 | Exon |
| chr16 | 89835751 | 89838114 | Exon |
| chr16 | 89952677 | 89953980 | Intron |
| chr16 | 89954343 | 89955527 | Intron |
| chr16 | 89973119 | 89975827 | Exon |
| chr16 | 90095004 | 90096402 | Promoter |
| chr16 | 90096711 | 90097048 | Promoter |
| chr16 | 90097672 | 90099303 | 5' UTR |
| chr16 | 90126826 | 90127743 | Promoter |
| chr16 | 90144867 | 90145187 | Distal Intergenic |
| chr16 | 90150842 | 90151145 | Distal Intergenic |
| chr16 | 90181650 | 90181851 | Intron |
| chr16 | 90210878 | 90211085 | Distal Intergenic |
| chr16 | 90284355 | 90284657 | Intron |
| chr16 | 90393826 | 90394045 | Distal Intergenic |
| chr16 | 90468266 | 90469551 | Distal Intergenic |
| chr16 | 90479096 | 90479321 | Distal Intergenic |
| chr16 | 90621711 | 90622947 | Distal Intergenic |
| chr16 | 90830246 | 90830520 | Distal Intergenic |
| chr16 | 90830559 | 90830917 | Distal Intergenic |
| chr16 | 90831115 | 90831438 | Distal Intergenic |
| chr16 | 90831558 | 90831916 | Distal Intergenic |
| chr16 | 90844722 | 90845864 | Distal Intergenic |
| chr16 | 90856645 | 90857785 | Distal Intergenic |

| | | | |
|---|---|---|---|
| chr16 | 90865889 | 90866226 | Distal Intergenic |
| chr16 | 90873577 | 90873778 | Distal Intergenic |
| chr16 | 90885142 | 90887076 | Distal Intergenic |
| chr16 | 91010985 | 91011354 | Distal Intergenic |
| chr16 | 91011851 | 91012074 | Distal Intergenic |
| chr16 | 91015829 | 91016320 | Distal Intergenic |
| chr16 | 91044066 | 91044287 | Distal Intergenic |
| chr16 | 91398044 | 91398946 | Distal Intergenic |
| chr16 | 91399041 | 91399246 | Distal Intergenic |
| chr16 | 91405732 | 91406070 | Distal Intergenic |
| chr16 | 91406150 | 91406639 | Distal Intergenic |
| chr16 | 91485573 | 91485772 | Distal Intergenic |
| chr16 | 91646424 | 91646739 | Distal Intergenic |
| chr16 | 91647348 | 91647582 | Distal Intergenic |
| chr16 | 91648055 | 91648368 | Distal Intergenic |
| chr16 | 91648562 | 91648888 | Distal Intergenic |
| chr16 | 92399104 | 92400399 | Distal Intergenic |
| chr16 | 92400533 | 92401431 | Distal Intergenic |
| chr16 | 92402411 | 92402838 | Distal Intergenic |
| chr16 | 92418998 | 92420374 | Distal Intergenic |
| chr16 | 92421070 | 92421269 | Distal Intergenic |
| chr16 | 92425783 | 92426325 | Distal Intergenic |
| chr16 | 92550053 | 92553814 | Distal Intergenic |
| chr16 | 92608260 | 92608861 | Distal Intergenic |
| chr16 | 92608898 | 92611561 | Distal Intergenic |
| chr16 | 92615173 | 92615372 | Distal Intergenic |
| chr16 | 92615411 | 92616680 | Distal Intergenic |
| chr16 | 92616733 | 92618187 | Distal Intergenic |
| chr16 | 92620207 | 92622757 | Distal Intergenic |
| chr16 | 92622858 | 92623213 | Distal Intergenic |
| chr16 | 92623603 | 92625285 | Distal Intergenic |
| chr16 | 92626527 | 92627756 | Distal Intergenic |
| chr16 | 92684202 | 92685933 | Distal Intergenic |
| chr16 | 92693107 | 92693321 | Distal Intergenic |
| chr16 | 92693648 | 92695020 | Distal Intergenic |
| chr16 | 92695382 | 92696958 | Distal Intergenic |
| chr16 | 92697648 | 92697953 | Distal Intergenic |
| chr16 | 92698402 | 92699017 | Distal Intergenic |
| chr16 | 92824280 | 92825100 | Distal Intergenic |
| chr16 | 92827813 | 92828054 | Distal Intergenic |
| chr16 | 92828132 | 92828700 | Distal Intergenic |
| chr16 | 92828839 | 92829247 | Distal Intergenic |
| chr16 | 92903932 | 92904287 | Distal Intergenic |
| chr16 | 92904488 | 92904796 | Distal Intergenic |

| | | | |
|---|---|---|---|
| chr16 | 92912184 | 92912730 | Distal Intergenic |
| chr16 | 93150603 | 93150812 | Distal Intergenic |
| chr16 | 93151019 | 93151218 | Distal Intergenic |
| chr16 | 93381997 | 93382196 | Distal Intergenic |
| chr16 | 93446036 | 93447300 | Distal Intergenic |
| chr16 | 93542341 | 93542677 | Distal Intergenic |
| chr16 | 93542880 | 93543215 | Distal Intergenic |
| chr16 | 94023967 | 94024396 | Distal Intergenic |
| chr16 | 94733623 | 94733861 | Distal Intergenic |
| chr16 | 94740983 | 94741262 | Distal Intergenic |
| chr16 | 94741321 | 94741596 | Distal Intergenic |
| chr16 | 94791246 | 94791668 | Distal Intergenic |
| chr16 | 94792214 | 94793344 | Distal Intergenic |
| chr16 | 94794001 | 94794200 | Distal Intergenic |
| chr16 | 94807103 | 94808800 | Distal Intergenic |
| chr16 | 94808845 | 94809126 | Distal Intergenic |
| chr16 | 94823715 | 94824860 | Distal Intergenic |
| chr16 | 94824930 | 94825168 | Distal Intergenic |
| chr16 | 94830801 | 94831278 | Distal Intergenic |
| chr16 | 95259592 | 95259846 | Distal Intergenic |
| chr16 | 95259923 | 95260787 | Distal Intergenic |
| chr16 | 95500564 | 95501150 | Distal Intergenic |
| chr16 | 95505657 | 95507750 | Distal Intergenic |
| chr16 | 95661993 | 95662256 | Distal Intergenic |
| chr16 | 95880154 | 95881589 | Distal Intergenic |
| chr16 | 95890266 | 95891045 | Distal Intergenic |
| chr16 | 95895769 | 95896744 | Distal Intergenic |
| chr16 | 95896987 | 95897585 | Distal Intergenic |
| chr16 | 95903656 | 95904055 | Distal Intergenic |
| chr16 | 95919753 | 95920587 | Distal Intergenic |
| chr16 | 95924055 | 95925627 | Distal Intergenic |
| chr16 | 95937792 | 95938141 | Distal Intergenic |
| chr16 | 95950321 | 95950976 | Distal Intergenic |
| chr16 | 95962929 | 95963696 | Distal Intergenic |
| chr16 | 95981534 | 95982195 | Distal Intergenic |
| chr16 | 95988823 | 95989140 | Distal Intergenic |
| chr16 | 96048981 | 96049638 | Distal Intergenic |
| chr16 | 96119445 | 96120633 | Distal Intergenic |
| chr16 | 96312571 | 96313920 | Distal Intergenic |
| chr16 | 96348038 | 96348447 | Distal Intergenic |
| chr16 | 97755314 | 97755551 | Distal Intergenic |
| chr16 | 97781923 | 97782942 | Distal Intergenic |
| chr16 | 97783016 | 97783231 | Distal Intergenic |
| chr16 | 97815480 | 97815713 | Distal Intergenic |

| | | | |
|---|---|---|---|
| chr16 | 97892683 | 97892928 | Distal Intergenic |
| chr16 | 97893135 | 97893404 | Distal Intergenic |
| chr16 | 97899190 | 97900056 | Distal Intergenic |
| chr16 | 97901849 | 97902300 | Distal Intergenic |
| chr16 | 97902379 | 97903409 | Distal Intergenic |
| chr16 | 97907319 | 97907518 | Distal Intergenic |
| chr16 | 97929737 | 97930572 | Distal Intergenic |
| chr16 | 97930748 | 97930966 | Distal Intergenic |
| chr16 | 97936117 | 97936331 | Distal Intergenic |
| chr16 | 97936381 | 97936783 | Distal Intergenic |
| chr16 | 97937897 | 97938133 | Distal Intergenic |
| chr16 | 97939146 | 97939660 | Distal Intergenic |
| chr16 | 97958186 | 97958507 | Distal Intergenic |
| chr16 | 97958631 | 97958910 | Distal Intergenic |
| chr16 | 97972463 | 97972667 | Distal Intergenic |
| chr16 | 97973494 | 97974415 | Distal Intergenic |
| chr16 | 97974600 | 97974926 | Distal Intergenic |
| chr16 | 97978604 | 97978849 | Distal Intergenic |
| chr16 | 97979055 | 97979704 | Distal Intergenic |
| chr16 | 97979814 | 97980063 | Distal Intergenic |
| chr16 | 97980455 | 97980717 | Distal Intergenic |
| chr16 | 97980818 | 97981865 | Distal Intergenic |
| chr16 | 97982048 | 97982415 | Distal Intergenic |
| chr16 | 97982489 | 97985008 | Distal Intergenic |
| chr16 | 97985257 | 97985512 | Distal Intergenic |
| chr16 | 97990266 | 97990632 | Distal Intergenic |
| chr16 | 98093472 | 98093734 | Distal Intergenic |
| chr16 | 98093842 | 98094632 | Distal Intergenic |
| chr16 | 98118937 | 98119370 | Distal Intergenic |
| chr16 | 98149230 | 98149556 | Distal Intergenic |
| chr16 | 98218201 | 98218661 | Distal Intergenic |
| chr16 | 98303200 | 98303600 | Distal Intergenic |
| chr17 | 3083772 | 3084140 | Distal Intergenic |
| chr17 | 3114976 | 3115175 | Distal Intergenic |
| chr17 | 3115391 | 3115823 | Distal Intergenic |
| chr17 | 3162084 | 3163563 | Distal Intergenic |
| chr17 | 3165348 | 3166265 | Distal Intergenic |
| chr17 | 3331835 | 3332254 | Distal Intergenic |
| chr17 | 3349435 | 3349859 | Exon |
| chr17 | 3349981 | 3350712 | Intron |
| chr17 | 4993726 | 4994536 | Intron |
| chr17 | 4996187 | 4996552 | 3' UTR |
| chr17 | 4997097 | 4997325 | 3' UTR |
| chr17 | 5125907 | 5126227 | Intron |

| | | | |
|---|---|---|---|
| chr17 | 5131649 | 5132331 | Intron |
| chr17 | 5132625 | 5133340 | Intron |
| chr17 | 5189056 | 5189955 | Intron |
| chr17 | 5237774 | 5238571 | Exon |
| chr17 | 5238608 | 5238823 | Exon |
| chr17 | 5822493 | 5822695 | Intron |
| chr17 | 5846367 | 5846720 | Distal Intergenic |
| chr17 | 5846810 | 5847140 | Distal Intergenic |
| chr17 | 5974712 | 5976113 | Promoter |
| chr17 | 5989907 | 5990134 | Intron |
| chr17 | 5991162 | 5991630 | Exon |
| chr17 | 5991673 | 5991887 | Intron |
| chr17 | 5994440 | 5995188 | Intron |
| chr17 | 6000862 | 6001268 | Intron |
| chr17 | 6001516 | 6001728 | Intron |
| chr17 | 6001769 | 6001984 | Intron |
| chr17 | 6005879 | 6006269 | Intron |
| chr17 | 6008148 | 6008360 | Intron |
| chr17 | 6016544 | 6017175 | Intron |
| chr17 | 6017687 | 6018271 | Intron |
| chr17 | 6107242 | 6108431 | Distal Intergenic |
| chr17 | 6117169 | 6117933 | Distal Intergenic |
| chr17 | 6127416 | 6128824 | Distal Intergenic |
| chr17 | 6140633 | 6141084 | Distal Intergenic |
| chr17 | 6141145 | 6141344 | Distal Intergenic |
| chr17 | 6153201 | 6154802 | Distal Intergenic |
| chr17 | 6172222 | 6173490 | Distal Intergenic |
| chr17 | 6174017 | 6174559 | Distal Intergenic |
| chr17 | 6201968 | 6203943 | Distal Intergenic |
| chr17 | 6955507 | 6956639 | Distal Intergenic |
| chr17 | 6959681 | 6960805 | Distal Intergenic |
| chr17 | 6961210 | 6963311 | Distal Intergenic |
| chr17 | 6982793 | 6982999 | Promoter |
| chr17 | 6983443 | 6984072 | Promoter |
| chr17 | 6984268 | 6984547 | Promoter |
| chr17 | 7012366 | 7014207 | Intron |
| chr17 | 7031275 | 7033320 | Distal Intergenic |
| chr17 | 7164852 | 7165072 | Intron |
| chr17 | 7165589 | 7165794 | Promoter |
| chr17 | 8358728 | 8359090 | Intron |
| chr17 | 8701421 | 8702288 | Promoter |
| chr17 | 8745119 | 8746016 | Intron |
| chr17 | 10012992 | 10015507 | Intron |
| chr17 | 10247527 | 10247740 | Intron |

| | | | |
|---|---|---|---|
| chr17 | 10375049 | 10375889 | Intron |
| chr17 | 10442293 | 10442518 | Intron |
| chr17 | 10510353 | 10511334 | Intron |
| chr17 | 11676313 | 11676827 | Intron |
| chr17 | 12468433 | 12468650 | Intron |
| chr17 | 12468698 | 12468915 | Intron |
| chr17 | 12933393 | 12934148 | Distal Intergenic |
| chr17 | 12943294 | 12943900 | Distal Intergenic |
| chr17 | 12946988 | 12948520 | Distal Intergenic |
| chr17 | 12983692 | 12985071 | Distal Intergenic |
| chr17 | 13108959 | 13109786 | Distal Intergenic |
| chr17 | 13183947 | 13184162 | Distal Intergenic |
| chr17 | 13184229 | 13184494 | Distal Intergenic |
| chr17 | 13735556 | 13735884 | Distal Intergenic |
| chr17 | 13819356 | 13821274 | Distal Intergenic |
| chr17 | 13836269 | 13836525 | Distal Intergenic |
| chr17 | 13836640 | 13838003 | Distal Intergenic |
| chr17 | 13846614 | 13846885 | Distal Intergenic |
| chr17 | 13851843 | 13853323 | Distal Intergenic |
| chr17 | 13939746 | 13940089 | Intron |
| chr17 | 13992388 | 13992976 | Intron |
| chr17 | 13993195 | 13993481 | Intron |
| chr17 | 13993531 | 13993926 | Intron |
| chr17 | 14003807 | 14004970 | Intron |
| chr17 | 14057540 | 14058732 | Intron |
| chr17 | 14061182 | 14061891 | Intron |
| chr17 | 14061986 | 14062430 | Intron |
| chr17 | 14062691 | 14063330 | 5' UTR |
| chr17 | 14309692 | 14310180 | Distal Intergenic |
| chr17 | 14324650 | 14325240 | Distal Intergenic |
| chr17 | 14330886 | 14331718 | Distal Intergenic |
| chr17 | 14473630 | 14476111 | Distal Intergenic |
| chr17 | 14476151 | 14476359 | Distal Intergenic |
| chr17 | 14476547 | 14477556 | Distal Intergenic |
| chr17 | 14561620 | 14561942 | Distal Intergenic |
| chr17 | 14623251 | 14624094 | Distal Intergenic |
| chr17 | 14650936 | 14651206 | Distal Intergenic |
| chr17 | 14651522 | 14651721 | Distal Intergenic |
| chr17 | 14890312 | 14890536 | Distal Intergenic |
| chr17 | 15110233 | 15110690 | Distal Intergenic |
| chr17 | 15110792 | 15111047 | Distal Intergenic |
| chr17 | 15114870 | 15115759 | Distal Intergenic |
| chr17 | 15116059 | 15116652 | Distal Intergenic |
| chr17 | 15126735 | 15127736 | Distal Intergenic |

| | | | |
|---|---|---|---|
| chr17 | 15127879 | 15129701 | Distal Intergenic |
| chr17 | 15141411 | 15142544 | Intron |
| chr17 | 15146333 | 15147083 | Intron |
| chr17 | 15147276 | 15147875 | Intron |
| chr17 | 15157873 | 15160660 | Intron |
| chr17 | 15172763 | 15173842 | Distal Intergenic |
| chr17 | 15177450 | 15178506 | Distal Intergenic |
| chr17 | 15178554 | 15179248 | Distal Intergenic |
| chr17 | 15189186 | 15189468 | Distal Intergenic |
| chr17 | 15189900 | 15190340 | Distal Intergenic |
| chr17 | 15190560 | 15190872 | Distal Intergenic |
| chr17 | 15191147 | 15191824 | Distal Intergenic |
| chr17 | 15277476 | 15277877 | Distal Intergenic |
| chr17 | 15278185 | 15278386 | Distal Intergenic |
| chr17 | 15285907 | 15287200 | Distal Intergenic |
| chr17 | 15360206 | 15362028 | Intron |
| chr17 | 15450206 | 15450885 | 5' UTR |
| chr17 | 15462562 | 15465011 | Intron |
| chr17 | 15493357 | 15494188 | Intron |
| chr17 | 15502439 | 15503705 | Promoter |
| chr17 | 15512328 | 15512622 | Intron |
| chr17 | 15647335 | 15647864 | 3' UTR |
| chr17 | 15647993 | 15648274 | 3' UTR |
| chr17 | 15841284 | 15841526 | Distal Intergenic |
| chr17 | 15842598 | 15842917 | Distal Intergenic |
| chr17 | 15958557 | 15958805 | Intron |
| chr17 | 15960701 | 15961476 | Promoter |
| chr17 | 15961843 | 15962227 | Promoter |
| chr17 | 16205147 | 16205699 | Intron |
| chr17 | 16871239 | 16871438 | Intron |
| chr17 | 17201248 | 17201448 | Distal Intergenic |
| chr17 | 17510870 | 17511105 | Distal Intergenic |
| chr17 | 17511267 | 17511936 | Distal Intergenic |
| chr17 | 17511984 | 17512183 | Distal Intergenic |
| chr17 | 17755403 | 17756258 | Intron |
| chr17 | 17760424 | 17761025 | Intron |
| chr17 | 17761156 | 17761659 | Exon |
| chr17 | 17761722 | 17762489 | Intron |
| chr17 | 17928924 | 17929282 | Intron |
| chr17 | 17929483 | 17930225 | Exon |
| chr17 | 17960446 | 17963941 | Exon |
| chr17 | 17963981 | 17964431 | Intron |
| chr17 | 17964570 | 17965617 | Exon |
| chr17 | 17971259 | 17972198 | 3' UTR |

| | | | |
|---|---|---|---|
| chr17 | 17973396 | 17973817 | Distal Intergenic |
| chr17 | 17973856 | 17976214 | Distal Intergenic |
| chr17 | 17981979 | 17983263 | Distal Intergenic |
| chr17 | 21081623 | 21083117 | Exon |
| chr17 | 21145945 | 21146289 | 3' UTR |
| chr17 | 21146435 | 21146684 | 3' UTR |
| chr17 | 21586878 | 21587583 | Distal Intergenic |
| chr17 | 21609888 | 21611296 | Distal Intergenic |
| chr17 | 21612761 | 21612996 | Distal Intergenic |
| chr17 | 21615010 | 21615660 | Distal Intergenic |
| chr17 | 21652913 | 21653976 | Distal Intergenic |
| chr17 | 21654056 | 21654255 | Distal Intergenic |
| chr17 | 21668892 | 21672046 | Distal Intergenic |
| chr17 | 21672460 | 21674926 | Distal Intergenic |
| chr17 | 21694204 | 21694680 | Distal Intergenic |
| chr17 | 21882604 | 21883395 | Distal Intergenic |
| chr17 | 22099251 | 22099450 | Distal Intergenic |
| chr17 | 22099703 | 22099945 | Distal Intergenic |
| chr17 | 22362423 | 22362622 | Distal Intergenic |
| chr17 | 22451409 | 22451834 | Distal Intergenic |
| chr17 | 22498423 | 22498865 | Distal Intergenic |
| chr17 | 22876461 | 22876830 | Distal Intergenic |
| chr17 | 23680515 | 23681164 | Distal Intergenic |
| chr17 | 23683818 | 23684248 | Distal Intergenic |
| chr17 | 23687527 | 23688093 | Distal Intergenic |
| chr17 | 23794573 | 23794937 | Distal Intergenic |
| chr17 | 23794980 | 23795312 | Distal Intergenic |
| chr17 | 23810969 | 23813433 | Distal Intergenic |
| chr17 | 23813617 | 23814269 | Distal Intergenic |
| chr17 | 23817116 | 23817647 | Distal Intergenic |
| chr17 | 23831109 | 23831768 | Distal Intergenic |
| chr17 | 23831811 | 23832310 | Distal Intergenic |
| chr17 | 23863920 | 23864152 | Distal Intergenic |
| chr17 | 23940249 | 23940448 | Distal Intergenic |
| chr17 | 23940613 | 23941744 | Distal Intergenic |
| chr17 | 23965148 | 23966051 | Distal Intergenic |
| chr17 | 24061489 | 24061804 | Distal Intergenic |
| chr17 | 24061997 | 24062282 | Distal Intergenic |
| chr17 | 24116319 | 24117732 | Distal Intergenic |
| chr17 | 24134561 | 24137102 | Distal Intergenic |
| chr17 | 24137225 | 24137424 | Distal Intergenic |
| chr17 | 24152209 | 24152529 | Distal Intergenic |
| chr17 | 24152615 | 24152893 | Distal Intergenic |
| chr17 | 24171683 | 24172425 | Distal Intergenic |

| | | | |
|---|---|---|---|
| chr17 | 24210397 | 24210601 | Distal Intergenic |
| chr17 | 24283659 | 24284166 | Distal Intergenic |
| chr17 | 24284571 | 24284810 | Distal Intergenic |
| chr17 | 24289008 | 24289727 | Distal Intergenic |
| chr17 | 24305575 | 24305815 | Distal Intergenic |
| chr17 | 24306442 | 24306643 | Distal Intergenic |
| chr17 | 24357854 | 24358092 | Distal Intergenic |
| chr17 | 24380772 | 24382409 | Distal Intergenic |
| chr17 | 24387425 | 24388384 | Distal Intergenic |
| chr17 | 24388583 | 24389220 | Distal Intergenic |
| chr17 | 24551744 | 24552690 | Distal Intergenic |
| chr17 | 24615787 | 24615986 | Distal Intergenic |
| chr17 | 24664211 | 24667669 | Distal Intergenic |
| chr17 | 24667846 | 24668054 | Distal Intergenic |
| chr17 | 24804740 | 24804939 | Distal Intergenic |
| chr17 | 24805519 | 24806554 | Distal Intergenic |
| chr17 | 24807070 | 24807421 | Distal Intergenic |
| chr17 | 24808995 | 24809226 | Distal Intergenic |
| chr17 | 24832485 | 24832892 | Distal Intergenic |
| chr17 | 24856468 | 24856890 | Distal Intergenic |
| chr17 | 24857060 | 24857305 | Distal Intergenic |
| chr17 | 24888250 | 24888867 | Distal Intergenic |
| chr17 | 24954654 | 24956105 | Distal Intergenic |
| chr17 | 24976066 | 24978625 | Distal Intergenic |
| chr17 | 24985981 | 24986215 | Distal Intergenic |
| chr17 | 24986429 | 24986712 | Distal Intergenic |
| chr17 | 24986848 | 24987291 | Distal Intergenic |
| chr17 | 24992217 | 24993314 | Distal Intergenic |
| chr17 | 24993364 | 24993652 | Distal Intergenic |
| chr17 | 24996647 | 24997368 | Distal Intergenic |
| chr17 | 25031697 | 25032448 | Distal Intergenic |
| chr17 | 25032562 | 25032898 | Distal Intergenic |
| chr17 | 25033046 | 25033401 | Distal Intergenic |
| chr17 | 25066301 | 25066983 | Distal Intergenic |
| chr17 | 25085126 | 25085531 | Distal Intergenic |
| chr17 | 25087431 | 25088456 | Distal Intergenic |
| chr17 | 25359664 | 25360182 | Distal Intergenic |
| chr17 | 25408092 | 25409043 | Distal Intergenic |
| chr17 | 25409191 | 25410891 | Distal Intergenic |
| chr17 | 25893331 | 25894611 | Intron |
| chr17 | 26055746 | 26055979 | Distal Intergenic |
| chr17 | 26078862 | 26079190 | Distal Intergenic |
| chr17 | 26122019 | 26122342 | Intron |
| chr17 | 26315376 | 26315660 | Distal Intergenic |

| | | | |
|---|---|---|---|
| chr17 | 26389619 | 26390319 | Intron |
| chr17 | 26390591 | 26390973 | Intron |
| chr17 | 26497865 | 26498431 | Intron |
| chr17 | 26639413 | 26640118 | Distal Intergenic |
| chr17 | 26640337 | 26641082 | Distal Intergenic |
| chr17 | 26670638 | 26671530 | Exon |
| chr17 | 26674772 | 26675398 | 3' UTR |
| chr17 | 26686011 | 26686496 | Exon |
| chr17 | 26697941 | 26698394 | Promoter |
| chr17 | 26741805 | 26742754 | Distal Intergenic |
| chr17 | 26742799 | 26743056 | Distal Intergenic |
| chr17 | 26757870 | 26758871 | Distal Intergenic |
| chr17 | 26767454 | 26767826 | Distal Intergenic |
| chr17 | 26768142 | 26769540 | Distal Intergenic |
| chr17 | 26813502 | 26814073 | Intron |
| chr17 | 27164491 | 27164698 | Intron |
| chr17 | 27164780 | 27165559 | Intron |
| chr17 | 27175148 | 27175370 | Distal Intergenic |
| chr17 | 27175423 | 27175681 | Distal Intergenic |
| chr17 | 27175744 | 27176117 | Distal Intergenic |
| chr17 | 27180345 | 27180573 | Distal Intergenic |
| chr17 | 27185024 | 27185299 | Promoter |
| chr17 | 27187353 | 27188668 | Promoter |
| chr17 | 27193140 | 27194105 | Distal Intergenic |
| chr17 | 27203826 | 27204160 | Downstream |
| chr17 | 27206802 | 27207312 | 3' UTR |
| chr17 | 27210155 | 27210976 | Exon |
| chr17 | 27211014 | 27213424 | Exon |
| chr17 | 27219239 | 27220095 | Intron |
| chr17 | 27220266 | 27221384 | Intron |
| chr17 | 27340208 | 27341034 | Distal Intergenic |
| chr17 | 27399891 | 27400227 | Downstream |
| chr17 | 27724473 | 27724721 | Intron |
| chr17 | 27759329 | 27759584 | Intron |
| chr17 | 27759788 | 27759987 | Intron |
| chr17 | 27771521 | 27771734 | Intron |
| chr17 | 27771968 | 27772210 | Intron |
| chr17 | 27934143 | 27938192 | 5' UTR |
| chr17 | 27956256 | 27956854 | 3' UTR |
| chr17 | 27957104 | 27957336 | 3' UTR |
| chr17 | 27962684 | 27963187 | Exon |
| chr17 | 27963246 | 27963568 | Exon |
| chr17 | 28113411 | 28114002 | Intron |
| chr17 | 28114167 | 28114584 | Intron |

| | | | |
|---|---|---|---|
| chr17 | 28159412 | 28160630 | Intron |
| chr17 | 28162377 | 28163390 | Intron |
| chr17 | 28344398 | 28345966 | Intron |
| chr17 | 28360382 | 28360797 | Intron |
| chr17 | 28361269 | 28362362 | Exon |
| chr17 | 28367601 | 28367918 | Intron |
| chr17 | 28368057 | 28368460 | Intron |
| chr17 | 28368551 | 28368780 | Intron |
| chr17 | 28369667 | 28374252 | Intron |
| chr17 | 28374295 | 28374720 | Intron |
| chr17 | 28393235 | 28394520 | Intron |
| chr17 | 28395478 | 28395944 | Intron |
| chr17 | 28395998 | 28397590 | Intron |
| chr17 | 28405181 | 28405464 | Exon |
| chr17 | 28406011 | 28407756 | Exon |
| chr17 | 28450665 | 28450913 | Intron |
| chr17 | 28466303 | 28467410 | Intron |
| chr17 | 28467526 | 28467872 | Intron |
| chr17 | 28468324 | 28468596 | Intron |
| chr17 | 28477307 | 28477873 | Intron |
| chr17 | 28487459 | 28487750 | Intron |
| chr17 | 28487793 | 28488162 | Intron |
| chr17 | 28834052 | 28835573 | Intron |
| chr17 | 28929981 | 28930374 | Intron |
| chr17 | 29062877 | 29063134 | Intron |
| chr17 | 29089570 | 29090012 | Intron |
| chr17 | 29144464 | 29144723 | Intron |
| chr17 | 29144862 | 29145416 | Intron |
| chr17 | 29168669 | 29169034 | Intron |
| chr17 | 29169315 | 29170091 | Intron |
| chr17 | 29170206 | 29171177 | Exon |
| chr17 | 29171222 | 29171492 | Intron |
| chr17 | 29215432 | 29216373 | Intron |
| chr17 | 29218959 | 29219371 | Intron |
| chr17 | 29227498 | 29229562 | Exon |
| chr17 | 29230032 | 29232219 | 3' UTR |
| chr17 | 29232758 | 29234171 | Promoter |
| chr17 | 29292706 | 29292973 | Distal Intergenic |
| chr17 | 29293283 | 29293482 | Distal Intergenic |
| chr17 | 29296435 | 29297826 | Promoter |
| chr17 | 29317570 | 29317872 | Intron |
| chr17 | 29400617 | 29401146 | Distal Intergenic |
| chr17 | 29468080 | 29469158 | Intron |
| chr17 | 29512515 | 29513230 | Intron |

| | | | |
|---|---|---|---|
| chr17 | 29513359 | 29513597 | Intron |
| chr17 | 29538171 | 29539385 | Intron |
| chr17 | 29797761 | 29798353 | Intron |
| chr17 | 29861242 | 29862333 | Promoter |
| chr17 | 30140983 | 30141319 | Distal Intergenic |
| chr17 | 30209561 | 30209790 | Intron |
| chr17 | 30209990 | 30210206 | Intron |
| chr17 | 30748239 | 30749283 | Distal Intergenic |
| chr17 | 31204203 | 31204402 | Promoter |
| chr17 | 31210939 | 31211138 | Distal Intergenic |
| chr17 | 31412390 | 31412848 | Intron |
| chr17 | 31442091 | 31442532 | Intron |
| chr17 | 31648628 | 31648849 | Intron |
| chr17 | 31794280 | 31795067 | Intron |
| chr17 | 31795107 | 31795496 | Intron |
| chr17 | 31880832 | 31881488 | Intron |
| chr17 | 31883027 | 31886024 | Intron |
| chr17 | 31890573 | 31891147 | Intron |
| chr17 | 31964911 | 31965209 | Intron |
| chr17 | 31967306 | 31968755 | Intron |
| chr17 | 31970425 | 31971650 | Intron |
| chr17 | 31971699 | 31972162 | Intron |
| chr17 | 31978575 | 31980053 | Intron |
| chr17 | 32033615 | 32033882 | Intron |
| chr17 | 32033979 | 32034475 | Intron |
| chr17 | 32140177 | 32140429 | Intron |
| chr17 | 32140507 | 32140723 | Intron |
| chr17 | 32167492 | 32168176 | Intron |
| chr17 | 32228008 | 32228542 | Intron |
| chr17 | 32228633 | 32229125 | Intron |
| chr17 | 32339600 | 32340396 | Intron |
| chr17 | 32420414 | 32420750 | Intron |
| chr17 | 32420849 | 32421092 | Intron |
| chr17 | 32457275 | 32457551 | Intron |
| chr17 | 32457755 | 32458101 | Intron |
| chr17 | 32591243 | 32591503 | Distal Intergenic |
| chr17 | 32594796 | 32595289 | Distal Intergenic |
| chr17 | 32595436 | 32595635 | Distal Intergenic |
| chr17 | 32611855 | 32612106 | Promoter |
| chr17 | 32637477 | 32637676 | Distal Intergenic |
| chr17 | 32923555 | 32923755 | Intron |
| chr17 | 32924069 | 32924609 | Intron |
| chr17 | 32924840 | 32925246 | Intron |
| chr17 | 33226249 | 33227263 | Distal Intergenic |

| | | | |
|---|---|---|---|
| chr17 | 33766333 | 33766667 | 3' UTR |
| chr17 | 33821597 | 33822343 | Intron |
| chr17 | 33897013 | 33897304 | Distal Intergenic |
| chr17 | 33920676 | 33920875 | Intron |
| chr17 | 33961480 | 33961896 | Intron |
| chr17 | 33961994 | 33962193 | Intron |
| chr17 | 34046394 | 34047336 | Intron |
| chr17 | 34052863 | 34053083 | 3' UTR |
| chr17 | 34055804 | 34057135 | Distal Intergenic |
| chr17 | 34076989 | 34077767 | Exon |
| chr17 | 34088552 | 34088921 | Promoter |
| chr17 | 34089632 | 34089862 | Intron |
| chr17 | 34090618 | 34090855 | Intron |
| chr17 | 34092702 | 34092977 | 3' UTR |
| chr17 | 34167876 | 34168140 | Intron |
| chr17 | 34168393 | 34168592 | Intron |
| chr17 | 34256622 | 34259216 | Promoter |
| chr17 | 34260049 | 34260873 | Downstream |
| chr17 | 34394818 | 34395284 | Intron |
| chr17 | 34397334 | 34398158 | Exon |
| chr17 | 34405893 | 34406687 | Distal Intergenic |
| chr17 | 34741230 | 34742197 | Intron |
| chr17 | 34987773 | 34988006 | Distal Intergenic |
| chr17 | 35034269 | 35035317 | Distal Intergenic |
| chr17 | 35036204 | 35037153 | Distal Intergenic |
| chr17 | 35081338 | 35081563 | Distal Intergenic |
| chr17 | 35086394 | 35086634 | Distal Intergenic |
| chr17 | 35086780 | 35090019 | Distal Intergenic |
| chr17 | 35096355 | 35098865 | Distal Intergenic |
| chr17 | 35101577 | 35102216 | Distal Intergenic |
| chr17 | 35102276 | 35103220 | Distal Intergenic |
| chr17 | 35117762 | 35119053 | Distal Intergenic |
| chr17 | 35119824 | 35120023 | Distal Intergenic |
| chr17 | 35137830 | 35138389 | Distal Intergenic |
| chr17 | 35153665 | 35153970 | Distal Intergenic |
| chr17 | 35179343 | 35179583 | Distal Intergenic |
| chr17 | 35180145 | 35183186 | Distal Intergenic |
| chr17 | 35183252 | 35183558 | Distal Intergenic |
| chr17 | 35184627 | 35184863 | Distal Intergenic |
| chr17 | 35186669 | 35186920 | Distal Intergenic |
| chr17 | 35186957 | 35189241 | Distal Intergenic |
| chr17 | 35258388 | 35259132 | Distal Intergenic |
| chr17 | 35271722 | 35272359 | Distal Intergenic |
| chr17 | 35272404 | 35272781 | Distal Intergenic |

| | | | |
|---|---|---|---|
| chr17 | 35300338 | 35300681 | 3' UTR |
| chr17 | 35300786 | 35301766 | 3' UTR |
| chr17 | 35323090 | 35323312 | Intron |
| chr17 | 35323682 | 35323983 | Intron |
| chr17 | 35335061 | 35335660 | Intron |
| chr17 | 35335702 | 35336603 | Intron |
| chr17 | 35336640 | 35337102 | Intron |
| chr17 | 35337168 | 35338195 | Intron |
| chr17 | 35371634 | 35371958 | Intron |
| chr17 | 35372237 | 35372874 | Intron |
| chr17 | 35378741 | 35380256 | Intron |
| chr17 | 35694073 | 35694447 | Intron |
| chr17 | 35725551 | 35725890 | Intron |
| chr17 | 35725960 | 35726326 | 5' UTR |
| chr17 | 35727361 | 35727920 | Intron |
| chr17 | 35729041 | 35729412 | Intron |
| chr17 | 35732549 | 35732794 | Promoter |
| chr17 | 35817989 | 35818188 | Intron |
| chr17 | 35834418 | 35835042 | Exon |
| chr17 | 35835121 | 35836164 | Intron |
| chr17 | 35836339 | 35837389 | 3' UTR |
| chr17 | 35837999 | 35838296 | Distal Intergenic |
| chr17 | 35838600 | 35838819 | Distal Intergenic |
| chr17 | 35880851 | 35881460 | Intron |
| chr17 | 35887020 | 35887241 | Intron |
| chr17 | 35887409 | 35887635 | Intron |
| chr17 | 35932498 | 35933185 | Intron |
| chr17 | 35933883 | 35934144 | Intron |
| chr17 | 35944956 | 35945832 | Exon |
| chr17 | 35948983 | 35951081 | Intron |
| chr17 | 35958068 | 35958319 | Intron |
| chr17 | 35959113 | 35960061 | Promoter |
| chr17 | 35973941 | 35974153 | 3' UTR |
| chr17 | 35974209 | 35974665 | 3' UTR |
| chr17 | 35974839 | 35976182 | Intron |
| chr17 | 36002707 | 36008232 | Promoter |
| chr17 | 36011399 | 36011783 | Distal Intergenic |
| chr17 | 36047140 | 36047339 | 3' UTR |
| chr17 | 36052504 | 36052968 | Intron |
| chr17 | 36053092 | 36053357 | Intron |
| chr17 | 36054068 | 36054489 | Intron |
| chr17 | 36054574 | 36056546 | Intron |
| chr17 | 36106408 | 36106705 | Distal Intergenic |
| chr17 | 36131429 | 36131938 | Distal Intergenic |

| | | | |
|---|---|---|---|
| chr17 | 36137739 | 36139687 | Distal Intergenic |
| chr17 | 36160600 | 36160972 | Distal Intergenic |
| chr17 | 36179843 | 36180124 | Distal Intergenic |
| chr17 | 36408259 | 36408479 | Intron |
| chr17 | 36408546 | 36409201 | Intron |
| chr17 | 37068907 | 37069263 | Intron |
| chr17 | 37088024 | 37088267 | Distal Intergenic |
| chr17 | 37088344 | 37088668 | Distal Intergenic |
| chr17 | 40939599 | 40940822 | 5' UTR |
| chr17 | 40943406 | 40943984 | Intron |
| chr17 | 41014187 | 41015920 | Distal Intergenic |
| chr17 | 41015961 | 41016198 | Distal Intergenic |
| chr17 | 41016330 | 41016613 | Distal Intergenic |
| chr17 | 41016732 | 41017943 | Distal Intergenic |
| chr17 | 41021055 | 41021434 | 3' UTR |
| chr17 | 41021624 | 41023059 | Distal Intergenic |
| chr17 | 42432699 | 42433196 | Exon |
| chr17 | 42827471 | 42829677 | 3' UTR |
| chr17 | 43275246 | 43275817 | Distal Intergenic |
| chr17 | 43276015 | 43276259 | Distal Intergenic |
| chr17 | 43356492 | 43356817 | Intron |
| chr17 | 43356988 | 43357371 | Intron |
| chr17 | 43544269 | 43544484 | Intron |
| chr17 | 43694878 | 43695373 | Distal Intergenic |
| chr17 | 44636723 | 44638421 | Intron |
| chr17 | 44638491 | 44638826 | Intron |
| chr17 | 44725105 | 44725330 | Intron |
| chr17 | 44791829 | 44792040 | Intron |
| chr17 | 44826846 | 44827930 | Exon |
| chr17 | 44846273 | 44847639 | Exon |
| chr17 | 44847792 | 44848248 | Intron |
| chr17 | 44904385 | 44904895 | Distal Intergenic |
| chr17 | 45202281 | 45202480 | Intron |
| chr17 | 45202579 | 45202874 | Intron |
| chr17 | 45496570 | 45496974 | Intron |
| chr17 | 45709005 | 45709946 | Distal Intergenic |
| chr17 | 45709987 | 45710334 | Distal Intergenic |
| chr17 | 45821879 | 45822934 | 3' UTR |
| chr17 | 45831541 | 45832732 | Distal Intergenic |
| chr17 | 45883595 | 45883822 | Downstream |
| chr17 | 45883873 | 45884427 | Downstream |
| chr17 | 45935905 | 45936108 | Distal Intergenic |
| chr17 | 46042486 | 46042729 | Distal Intergenic |
| chr17 | 46296756 | 46297101 | Intron |

| | | | |
|---|---|---|---|
| chr17 | 46338856 | 46339055 | Intron |
| chr17 | 46384505 | 46384737 | Intron |
| chr17 | 46403329 | 46405508 | Intron |
| chr17 | 46405866 | 46406314 | Intron |
| chr17 | 46406380 | 46407465 | Intron |
| chr17 | 46410414 | 46411872 | Intron |
| chr17 | 46413978 | 46414946 | Intron |
| chr17 | 46414982 | 46415342 | Intron |
| chr17 | 46464783 | 46465188 | Intron |
| chr17 | 46465279 | 46465507 | Intron |
| chr17 | 46690052 | 46690317 | 3' UTR |
| chr17 | 46692860 | 46693080 | Promoter |
| chr17 | 46696639 | 46696980 | Downstream |
| chr17 | 46711553 | 46712077 | Distal Intergenic |
| chr17 | 46746630 | 46747458 | Distal Intergenic |
| chr17 | 46760584 | 46761998 | Distal Intergenic |
| chr17 | 46765435 | 46765825 | Distal Intergenic |
| chr17 | 46810515 | 46811230 | Distal Intergenic |
| chr17 | 46818024 | 46818409 | Distal Intergenic |
| chr17 | 46910125 | 46910480 | Intron |
| chr17 | 46919395 | 46920076 | Intron |
| chr17 | 46920113 | 46920321 | Intron |
| chr17 | 46993208 | 46995409 | Exon |
| chr17 | 47003646 | 47004766 | 3' UTR |
| chr17 | 47011111 | 47011898 | Intron |
| chr17 | 47012770 | 47014721 | Exon |
| chr17 | 47016944 | 47017370 | Intron |
| chr17 | 47066054 | 47066253 | Distal Intergenic |
| chr17 | 47066570 | 47066861 | Distal Intergenic |
| chr17 | 47068796 | 47070892 | Distal Intergenic |
| chr17 | 47084639 | 47085811 | Intron |
| chr17 | 47146496 | 47146985 | Distal Intergenic |
| chr17 | 47202440 | 47202656 | Distal Intergenic |
| chr17 | 47278872 | 47280152 | Distal Intergenic |
| chr17 | 47280277 | 47280723 | Downstream |
| chr17 | 47281328 | 47282386 | Downstream |
| chr17 | 47403408 | 47404937 | Intron |
| chr17 | 47430697 | 47432182 | Intron |
| chr17 | 47433388 | 47434678 | Intron |
| chr17 | 47505619 | 47505970 | Distal Intergenic |
| chr17 | 47506182 | 47506402 | Distal Intergenic |
| chr17 | 47573789 | 47574554 | Intron |
| chr17 | 47574793 | 47575444 | Intron |
| chr17 | 47575482 | 47575686 | Intron |

| | | | |
|---|---|---|---|
| chr17 | 47575801 | 47576217 | Intron |
| chr17 | 47587418 | 47588440 | Exon |
| chr17 | 47621603 | 47621802 | Distal Intergenic |
| chr17 | 47748230 | 47748879 | Intron |
| chr17 | 47765724 | 47766130 | Distal Intergenic |
| chr17 | 47970671 | 47972065 | Distal Intergenic |
| chr17 | 47978393 | 47979752 | Distal Intergenic |
| chr17 | 48024605 | 48025806 | Distal Intergenic |
| chr17 | 48495389 | 48495772 | Distal Intergenic |
| chr17 | 48547990 | 48548189 | Intron |
| chr17 | 48549450 | 48550121 | Exon |
| chr17 | 51317716 | 51318181 | Distal Intergenic |
| chr17 | 51318328 | 51318579 | Distal Intergenic |
| chr17 | 51792701 | 51793142 | Distal Intergenic |
| chr17 | 52895421 | 52895862 | Distal Intergenic |
| chr17 | 53617881 | 53618484 | Distal Intergenic |
| chr17 | 53618632 | 53619511 | Distal Intergenic |
| chr17 | 53622071 | 53622822 | Distal Intergenic |
| chr17 | 53828786 | 53829027 | Promoter |
| chr17 | 56017520 | 56017733 | Intron |
| chr17 | 56018443 | 56018645 | Intron |
| chr17 | 56098375 | 56098765 | Distal Intergenic |
| chr17 | 56166500 | 56166775 | 3' UTR |
| chr17 | 56175034 | 56175233 | Distal Intergenic |
| chr17 | 56207233 | 56207662 | Distal Intergenic |
| chr17 | 56232126 | 56232331 | Promoter |
| chr17 | 56405470 | 56405891 | Promoter |
| chr17 | 56415962 | 56416231 | Intron |
| chr17 | 56416423 | 56416724 | Intron |
| chr17 | 56430047 | 56430637 | Promoter |
| chr17 | 56499539 | 56500712 | 3' UTR |
| chr17 | 56604244 | 56604473 | Promoter |
| chr17 | 56617513 | 56617713 | Promoter |
| chr17 | 56748463 | 56749385 | Intron |
| chr17 | 56812200 | 56812468 | Distal Intergenic |
| chr17 | 56812721 | 56813381 | Distal Intergenic |
| chr17 | 56958399 | 56958610 | Intron |
| chr17 | 56958656 | 56960136 | Intron |
| chr17 | 56961606 | 56961928 | Intron |
| chr17 | 56961996 | 56962307 | Intron |
| chr17 | 57016749 | 57017010 | Intron |
| chr17 | 57046004 | 57046838 | Intron |
| chr17 | 57047015 | 57047216 | Exon |
| chr17 | 57150409 | 57150733 | Intron |

| | | | |
|---|---|---|---|
| chr17 | 57366568 | 57368259 | Distal Intergenic |
| chr17 | 57387976 | 57388201 | Distal Intergenic |
| chr17 | 57389530 | 57389807 | Distal Intergenic |
| chr17 | 59215526 | 59215728 | Intron |
| chr17 | 59215805 | 59216382 | Intron |
| chr17 | 59216576 | 59216831 | Intron |
| chr17 | 59408805 | 59410020 | Intron |
| chr17 | 61360777 | 61360976 | Intron |
| chr17 | 61420399 | 61420855 | Intron |
| chr17 | 61440753 | 61441215 | Intron |
| chr17 | 61441408 | 61441856 | Intron |
| chr17 | 62635336 | 62636354 | Intron |
| chr17 | 62636503 | 62638578 | Intron |
| chr17 | 62697898 | 62698178 | Distal Intergenic |
| chr17 | 62727962 | 62728585 | Distal Intergenic |
| chr17 | 62728669 | 62728965 | Distal Intergenic |
| chr17 | 62764038 | 62764459 | Intron |
| chr17 | 62854218 | 62855024 | Exon |
| chr17 | 62983738 | 62983960 | Distal Intergenic |
| chr17 | 62984500 | 62984759 | Distal Intergenic |
| chr17 | 62984835 | 62985060 | Distal Intergenic |
| chr17 | 63025466 | 63026629 | Intron |
| chr17 | 63036447 | 63036970 | Intron |
| chr17 | 63037066 | 63037569 | Intron |
| chr17 | 63106540 | 63106739 | Distal Intergenic |
| chr17 | 63132891 | 63133258 | Promoter |
| chr17 | 63133481 | 63134003 | Promoter |
| chr17 | 63229264 | 63229758 | Distal Intergenic |
| chr17 | 63510982 | 63511332 | Distal Intergenic |
| chr17 | 64146243 | 64146645 | Intron |
| chr17 | 64147238 | 64149191 | Intron |
| chr17 | 64150396 | 64150640 | Intron |
| chr17 | 64150738 | 64151598 | Intron |
| chr17 | 64298186 | 64298729 | Promoter |
| chr17 | 64298780 | 64299024 | Promoter |
| chr17 | 64352545 | 64352979 | Intron |
| chr17 | 64353015 | 64353577 | Intron |
| chr17 | 64949259 | 64949458 | Distal Intergenic |
| chr17 | 64994184 | 64994590 | Intron |
| chr17 | 65115170 | 65115566 | Intron |
| chr17 | 65961801 | 65962132 | Intron |
| chr17 | 65963381 | 65964039 | Intron |
| chr17 | 66052814 | 66054104 | Distal Intergenic |
| chr17 | 66068465 | 66070601 | Distal Intergenic |

| | | | |
|---|---|---|---|
| chr17 | 66075273 | 66075541 | Distal Intergenic |
| chr17 | 66075724 | 66076667 | Distal Intergenic |
| chr17 | 66086146 | 66087990 | Distal Intergenic |
| chr17 | 66088997 | 66090918 | Distal Intergenic |
| chr17 | 66121177 | 66121408 | Intron |
| chr17 | 66131965 | 66132261 | Exon |
| chr17 | 66133169 | 66134142 | Distal Intergenic |
| chr17 | 66163277 | 66163545 | Distal Intergenic |
| chr17 | 66182701 | 66183508 | Distal Intergenic |
| chr17 | 66254482 | 66254835 | Promoter |
| chr17 | 66255003 | 66256220 | Promoter |
| chr17 | 66262256 | 66264067 | 3' UTR |
| chr17 | 66265020 | 66265282 | 3' UTR |
| chr17 | 66265347 | 66265991 | Exon |
| chr17 | 66451030 | 66451242 | Intron |
| chr17 | 66451284 | 66451484 | Intron |
| chr17 | 66586470 | 66586829 | 5' UTR |
| chr17 | 66586903 | 66587263 | Intron |
| chr17 | 66618514 | 66618743 | Distal Intergenic |
| chr17 | 66671426 | 66671745 | Distal Intergenic |
| chr17 | 66671819 | 66673677 | Distal Intergenic |
| chr17 | 66674029 | 66674932 | Distal Intergenic |
| chr17 | 66714554 | 66714753 | Distal Intergenic |
| chr17 | 66715301 | 66715526 | Distal Intergenic |
| chr17 | 66769200 | 66769411 | Distal Intergenic |
| chr17 | 67473303 | 67473695 | Intron |
| chr17 | 67524946 | 67525295 | Intron |
| chr17 | 67525342 | 67525557 | Intron |
| chr17 | 67974541 | 67974850 | Distal Intergenic |
| chr17 | 69576803 | 69577633 | Distal Intergenic |
| chr17 | 69578521 | 69579434 | Distal Intergenic |
| chr17 | 69766254 | 69766654 | Distal Intergenic |
| chr17 | 69766762 | 69767307 | Distal Intergenic |
| chr17 | 69767590 | 69767789 | Distal Intergenic |
| chr17 | 69771162 | 69771604 | Distal Intergenic |
| chr17 | 71069471 | 71069896 | Intron |
| chr17 | 71075553 | 71078248 | Intron |
| chr17 | 71078381 | 71079677 | Intron |
| chr17 | 71083434 | 71084592 | Intron |
| chr17 | 71085164 | 71085505 | Intron |
| chr17 | 71085973 | 71089006 | Promoter |
| chr17 | 71090652 | 71091799 | Distal Intergenic |
| chr17 | 71095594 | 71095902 | Distal Intergenic |
| chr17 | 71095951 | 71098325 | Distal Intergenic |

| | | | |
|---|---|---|---|
| chr17 | 71102572 | 71103539 | Distal Intergenic |
| chr17 | 71104544 | 71104770 | Distal Intergenic |
| chr17 | 71104967 | 71105318 | Distal Intergenic |
| chr17 | 71185969 | 71186260 | Distal Intergenic |
| chr17 | 71186385 | 71186635 | Distal Intergenic |
| chr17 | 71196535 | 71200549 | Exon |
| chr17 | 71200851 | 71201095 | Intron |
| chr17 | 71201509 | 71202177 | Exon |
| chr17 | 71202399 | 71202886 | Exon |
| chr17 | 71208117 | 71208439 | Intron |
| chr17 | 71208481 | 71208707 | Intron |
| chr17 | 71237669 | 71238157 | Intron |
| chr17 | 71238246 | 71238448 | Exon |
| chr17 | 71248565 | 71249613 | Exon |
| chr17 | 71257781 | 71258325 | Promoter |
| chr17 | 71258865 | 71259064 | Promoter |
| chr17 | 71349495 | 71350000 | Intron |
| chr17 | 71350038 | 71351229 | Intron |
| chr17 | 71351628 | 71352543 | Intron |
| chr17 | 71508524 | 71509164 | Intron |
| chr17 | 71542215 | 71542431 | Intron |
| chr17 | 71542647 | 71543272 | Intron |
| chr17 | 71544093 | 71545151 | Intron |
| chr17 | 71545235 | 71545467 | Intron |
| chr17 | 71876323 | 71876671 | Distal Intergenic |
| chr17 | 71965373 | 71965645 | Distal Intergenic |
| chr17 | 71966119 | 71966519 | Distal Intergenic |
| chr17 | 72001178 | 72003076 | Distal Intergenic |
| chr17 | 72091431 | 72092040 | Distal Intergenic |
| chr17 | 72092308 | 72092556 | Distal Intergenic |
| chr17 | 72129911 | 72130269 | Distal Intergenic |
| chr17 | 72347105 | 72347678 | 3' UTR |
| chr17 | 72491662 | 72491981 | Distal Intergenic |
| chr17 | 72648890 | 72649122 | Distal Intergenic |
| chr17 | 72688328 | 72689574 | Intron |
| chr17 | 72705442 | 72707102 | Intron |
| chr17 | 73177033 | 73178007 | Exon |
| chr17 | 73306654 | 73306853 | Distal Intergenic |
| chr17 | 73306910 | 73307235 | Distal Intergenic |
| chr17 | 73307499 | 73307785 | Distal Intergenic |
| chr17 | 73307841 | 73309610 | Distal Intergenic |
| chr17 | 73315503 | 73315874 | 3' UTR |
| chr17 | 74167467 | 74167747 | Intron |
| chr17 | 74242640 | 74243875 | Distal Intergenic |

| | | | |
|---|---|---|---|
| chr17 | 74245810 | 74247454 | Distal Intergenic |
| chr17 | 74282716 | 74284877 | 5' UTR |
| chr17 | 74285089 | 74285290 | Intron |
| chr17 | 74285726 | 74287681 | 5' UTR |
| chr17 | 74287718 | 74287949 | Exon |
| chr17 | 74290755 | 74292434 | Intron |
| chr17 | 74297318 | 74299966 | 5' UTR |
| chr17 | 74332402 | 74332650 | Intron |
| chr17 | 74333070 | 74333272 | Intron |
| chr17 | 74338729 | 74339626 | Intron |
| chr17 | 74366545 | 74367393 | Distal Intergenic |
| chr17 | 74678516 | 74678755 | Intron |
| chr17 | 74679245 | 74679494 | Intron |
| chr17 | 74682108 | 74682595 | Intron |
| chr17 | 74685284 | 74685670 | Promoter |
| chr17 | 74687719 | 74688807 | Intron |
| chr17 | 74693345 | 74693712 | Intron |
| chr17 | 74694027 | 74694234 | Intron |
| chr17 | 74695090 | 74695911 | Intron |
| chr17 | 74809063 | 74810027 | Distal Intergenic |
| chr17 | 74818071 | 74818976 | Distal Intergenic |
| chr17 | 74925711 | 74926026 | Intron |
| chr17 | 74926083 | 74927397 | Intron |
| chr17 | 74928446 | 74928812 | Exon |
| chr17 | 74945674 | 74946078 | 3' UTR |
| chr17 | 75098522 | 75098949 | Intron |
| chr17 | 75446392 | 75446991 | Promoter |
| chr17 | 75509612 | 75510665 | Distal Intergenic |
| chr17 | 75510715 | 75511182 | Distal Intergenic |
| chr17 | 75598911 | 75599327 | Distal Intergenic |
| chr17 | 75655144 | 75655359 | Distal Intergenic |
| chr17 | 75657566 | 75658530 | Distal Intergenic |
| chr17 | 75744393 | 75744683 | Distal Intergenic |
| chr17 | 75806393 | 75808017 | Distal Intergenic |
| chr17 | 76154984 | 76156658 | Promoter |
| chr17 | 76336188 | 76336391 | Distal Intergenic |
| chr17 | 77993246 | 77993685 | Intron |
| chr17 | 77993767 | 77994021 | Intron |
| chr17 | 78564567 | 78564836 | Intron |
| chr17 | 78564953 | 78565265 | Intron |
| chr17 | 78566156 | 78566383 | Intron |
| chr17 | 78606031 | 78606394 | Intron |
| chr17 | 78606472 | 78606911 | Intron |
| chr17 | 78611494 | 78613332 | Intron |

| | | | |
|---|---|---|---|
| chr17 | 78720252 | 78720773 | Intron |
| chr17 | 78720900 | 78721219 | Intron |
| chr17 | 78721533 | 78721991 | Intron |
| chr17 | 78722161 | 78722414 | Intron |
| chr17 | 78787013 | 78788370 | Intron |
| chr17 | 78791709 | 78791915 | Intron |
| chr17 | 78791974 | 78792326 | Intron |
| chr17 | 78801164 | 78801627 | Intron |
| chr17 | 78801830 | 78802061 | Intron |
| chr17 | 78802254 | 78802690 | Intron |
| chr17 | 78802748 | 78803296 | Intron |
| chr17 | 79123820 | 79124231 | Intron |
| chr17 | 79125966 | 79126347 | Intron |
| chr17 | 79126411 | 79126724 | Intron |
| chr17 | 79684411 | 79685247 | Exon |
| chr17 | 79771524 | 79772813 | 3' UTR |
| chr17 | 79793911 | 79794397 | Promoter |
| chr17 | 79794436 | 79794791 | Distal Intergenic |
| chr17 | 79989219 | 79989617 | Promoter |
| chr17 | 79998839 | 79999047 | Distal Intergenic |
| chr17 | 80145485 | 80145956 | Intron |
| chr17 | 80461173 | 80461421 | Distal Intergenic |
| chr17 | 81128105 | 81128310 | Distal Intergenic |
| chr17 | 81164434 | 81164772 | Distal Intergenic |
| chr17 | 81200305 | 81200523 | Distal Intergenic |
| chr17 | 81285372 | 81285814 | Distal Intergenic |
| chr17 | 81896855 | 81897713 | Distal Intergenic |
| chr17 | 83479797 | 83481035 | Distal Intergenic |
| chr17 | 83484964 | 83485830 | Distal Intergenic |
| chr17 | 83505384 | 83505874 | Distal Intergenic |
| chr17 | 83506001 | 83506495 | Distal Intergenic |
| chr17 | 83509008 | 83509550 | Distal Intergenic |
| chr17 | 83580798 | 83581642 | Distal Intergenic |
| chr17 | 84294841 | 84295260 | Distal Intergenic |
| chr17 | 84296253 | 84297730 | Distal Intergenic |
| chr17 | 84298536 | 84299547 | Distal Intergenic |
| chr17 | 84301238 | 84302447 | Distal Intergenic |
| chr17 | 84321624 | 84321823 | Distal Intergenic |
| chr17 | 84402452 | 84403745 | Distal Intergenic |
| chr17 | 84424680 | 84425360 | Distal Intergenic |
| chr17 | 84470129 | 84470574 | Distal Intergenic |
| chr17 | 84474222 | 84475033 | Distal Intergenic |
| chr17 | 84483293 | 84484028 | Distal Intergenic |
| chr17 | 84484353 | 84485822 | Distal Intergenic |

| | | | |
|---|---|---|---|
| chr17 | 84548392 | 84548892 | Distal Intergenic |
| chr17 | 84584182 | 84584388 | Distal Intergenic |
| chr17 | 84584428 | 84584922 | Distal Intergenic |
| chr17 | 84724874 | 84725937 | Distal Intergenic |
| chr17 | 84727729 | 84728592 | Distal Intergenic |
| chr17 | 85189951 | 85190150 | Distal Intergenic |
| chr17 | 85355983 | 85356215 | Distal Intergenic |
| chr17 | 85562062 | 85562409 | Distal Intergenic |
| chr17 | 85562525 | 85562868 | Distal Intergenic |
| chr17 | 86844510 | 86845124 | Distal Intergenic |
| chr17 | 86953534 | 86953757 | Distal Intergenic |
| chr17 | 86972654 | 86973313 | Distal Intergenic |
| chr17 | 86990370 | 86990719 | Distal Intergenic |
| chr17 | 86990845 | 86991413 | Distal Intergenic |
| chr17 | 87249888 | 87250173 | Distal Intergenic |
| chr17 | 87452138 | 87452358 | Distal Intergenic |
| chr17 | 87569299 | 87570760 | Distal Intergenic |
| chr17 | 87571011 | 87571210 | Distal Intergenic |
| chr17 | 87606296 | 87607068 | Distal Intergenic |
| chr17 | 87653121 | 87653426 | Distal Intergenic |
| chr17 | 87653471 | 87653918 | Distal Intergenic |
| chr17 | 87681544 | 87682140 | Distal Intergenic |
| chr17 | 87970645 | 87971033 | Distal Intergenic |
| chr17 | 88012710 | 88013543 | Distal Intergenic |
| chr17 | 88248115 | 88251704 | Distal Intergenic |
| chr17 | 88396962 | 88397555 | Distal Intergenic |
| chr17 | 88463344 | 88463623 | Distal Intergenic |
| chr17 | 88464780 | 88465101 | Distal Intergenic |
| chr17 | 88594950 | 88595196 | Distal Intergenic |
| chr17 | 88655150 | 88657032 | Distal Intergenic |
| chr17 | 88657387 | 88657993 | Distal Intergenic |
| chr17 | 88908018 | 88909249 | Distal Intergenic |
| chr17 | 88909746 | 88910728 | Distal Intergenic |
| chr17 | 88941854 | 88942904 | Distal Intergenic |
| chr17 | 89010565 | 89010907 | Distal Intergenic |
| chr17 | 89027923 | 89028649 | Distal Intergenic |
| chr17 | 89028737 | 89030283 | Distal Intergenic |
| chr17 | 89673415 | 89674735 | Distal Intergenic |
| chr17 | 89680500 | 89681087 | Distal Intergenic |
| chr17 | 89685974 | 89686385 | Distal Intergenic |
| chr17 | 91865924 | 91866599 | Distal Intergenic |
| chr17 | 95148984 | 95149252 | Distal Intergenic |
| chr17 | 95263990 | 95264221 | Distal Intergenic |
| chr18 | 3280529 | 3281275 | Distal Intergenic |

| | | | |
|---|---|---|---|
| chr18 | 3336553 | 3337346 | Distal Intergenic |
| chr18 | 3382900 | 3383171 | Distal Intergenic |
| chr18 | 3383618 | 3383868 | Distal Intergenic |
| chr18 | 3767355 | 3767897 | Intron |
| chr18 | 3963550 | 3964218 | Intron |
| chr18 | 3964361 | 3964560 | Intron |
| chr18 | 4156080 | 4156666 | Intron |
| chr18 | 4157785 | 4158004 | Intron |
| chr18 | 4310881 | 4311225 | Intron |
| chr18 | 4375348 | 4375780 | Intron |
| chr18 | 4553190 | 4553454 | Distal Intergenic |
| chr18 | 4737639 | 4738214 | Distal Intergenic |
| chr18 | 4785245 | 4786111 | Distal Intergenic |
| chr18 | 4845472 | 4846001 | Distal Intergenic |
| chr18 | 4920938 | 4921566 | Distal Intergenic |
| chr18 | 4921613 | 4922999 | Distal Intergenic |
| chr18 | 4959548 | 4960091 | Distal Intergenic |
| chr18 | 5001977 | 5002205 | Distal Intergenic |
| chr18 | 5002269 | 5002849 | Distal Intergenic |
| chr18 | 5012816 | 5013666 | Distal Intergenic |
| chr18 | 5016223 | 5017283 | Distal Intergenic |
| chr18 | 5045821 | 5047627 | Distal Intergenic |
| chr18 | 5077531 | 5079179 | Distal Intergenic |
| chr18 | 5334601 | 5334806 | Distal Intergenic |
| chr18 | 5395835 | 5396227 | Exon |
| chr18 | 5396335 | 5396995 | Intron |
| chr18 | 5402544 | 5403340 | Intron |
| chr18 | 5414377 | 5415038 | Intron |
| chr18 | 5945852 | 5946572 | Distal Intergenic |
| chr18 | 6119723 | 6120296 | Intron |
| chr18 | 6135260 | 6135571 | Intron |
| chr18 | 6135718 | 6135946 | Intron |
| chr18 | 6136527 | 6136753 | Intron |
| chr18 | 6192953 | 6193185 | Intron |
| chr18 | 6231100 | 6232465 | Intron |
| chr18 | 6235543 | 6235829 | Intron |
| chr18 | 6237835 | 6238855 | Exon |
| chr18 | 6239440 | 6239639 | Intron |
| chr18 | 6240283 | 6242235 | Exon |
| chr18 | 6275368 | 6275964 | Intron |
| chr18 | 6280816 | 6282938 | Intron |
| chr18 | 6285491 | 6286132 | Intron |
| chr18 | 6412409 | 6412651 | Intron |
| chr18 | 6419580 | 6419868 | Distal Intergenic |

| | | | |
|---|---|---|---|
| chr18 | 6489567 | 6489790 | Distal Intergenic |
| chr18 | 6489981 | 6490853 | Distal Intergenic |
| chr18 | 6516269 | 6516506 | Intron |
| chr18 | 6726112 | 6727323 | Distal Intergenic |
| chr18 | 6727441 | 6728870 | Distal Intergenic |
| chr18 | 6760268 | 6761422 | Distal Intergenic |
| chr18 | 6765224 | 6765776 | Distal Intergenic |
| chr18 | 6959736 | 6960264 | Intron |
| chr18 | 7613816 | 7614394 | Intron |
| chr18 | 7626068 | 7626858 | Intron |
| chr18 | 7649480 | 7650479 | Intron |
| chr18 | 7651242 | 7653054 | Intron |
| chr18 | 7868177 | 7868505 | Intron |
| chr18 | 7868866 | 7870428 | Intron |
| chr18 | 8161143 | 8161470 | Intron |
| chr18 | 8161507 | 8161706 | Intron |
| chr18 | 8693500 | 8693701 | Distal Intergenic |
| chr18 | 8694018 | 8694491 | Distal Intergenic |
| chr18 | 8719805 | 8720019 | Intron |
| chr18 | 8720094 | 8720586 | Exon |
| chr18 | 9356764 | 9356998 | Intron |
| chr18 | 9357043 | 9358282 | Intron |
| chr18 | 9363935 | 9364194 | Intron |
| chr18 | 9373971 | 9375691 | Intron |
| chr18 | 9375850 | 9376555 | Intron |
| chr18 | 9386427 | 9389931 | 5' UTR |
| chr18 | 9390307 | 9390517 | Intron |
| chr18 | 9390652 | 9391082 | Intron |
| chr18 | 9394017 | 9394608 | Intron |
| chr18 | 9428248 | 9429804 | Distal Intergenic |
| chr18 | 9450242 | 9450487 | Distal Intergenic |
| chr18 | 9552317 | 9552757 | Intron |
| chr18 | 9552864 | 9553492 | Exon |
| chr18 | 9734972 | 9735563 | Intron |
| chr18 | 9814585 | 9814823 | Intron |
| chr18 | 9957767 | 9958749 | 3' UTR |
| chr18 | 10012995 | 10013335 | Distal Intergenic |
| chr18 | 10013479 | 10014382 | Distal Intergenic |
| chr18 | 10028934 | 10029729 | Distal Intergenic |
| chr18 | 10181953 | 10182168 | Distal Intergenic |
| chr18 | 10290062 | 10290615 | Distal Intergenic |
| chr18 | 10533087 | 10533570 | 5' UTR |
| chr18 | 10617897 | 10618619 | Distal Intergenic |
| chr18 | 10725588 | 10725982 | Intron |

| | | | |
|---|---|---|---|
| chr18 | 10726334 | 10726599 | Intron |
| chr18 | 10726636 | 10726867 | Intron |
| chr18 | 11200321 | 11200849 | Distal Intergenic |
| chr18 | 11356174 | 11356401 | Distal Intergenic |
| chr18 | 11357063 | 11357526 | Distal Intergenic |
| chr18 | 11719601 | 11721662 | Intron |
| chr18 | 11742454 | 11744329 | Intron |
| chr18 | 11752472 | 11752734 | Promoter |
| chr18 | 11754876 | 11757294 | Intron |
| chr18 | 11791209 | 11791716 | Intron |
| chr18 | 11791774 | 11793616 | Intron |
| chr18 | 11815059 | 11815300 | Intron |
| chr18 | 11815551 | 11815807 | Intron |
| chr18 | 12035306 | 12035956 | Distal Intergenic |
| chr18 | 12041588 | 12042865 | Distal Intergenic |
| chr18 | 12124687 | 12125950 | Exon |
| chr18 | 12136239 | 12136831 | Distal Intergenic |
| chr18 | 12136893 | 12137092 | Distal Intergenic |
| chr18 | 12225372 | 12225688 | Distal Intergenic |
| chr18 | 12286923 | 12287126 | Distal Intergenic |
| chr18 | 12287281 | 12287508 | Distal Intergenic |
| chr18 | 12287544 | 12288070 | Distal Intergenic |
| chr18 | 12288115 | 12288314 | Distal Intergenic |
| chr18 | 12288350 | 12290187 | Distal Intergenic |
| chr18 | 12413414 | 12414548 | Intron |
| chr18 | 12414591 | 12414802 | Intron |
| chr18 | 12420051 | 12421360 | Promoter |
| chr18 | 12476596 | 12477308 | Intron |
| chr18 | 12478579 | 12479890 | Exon |
| chr18 | 12549956 | 12550515 | Intron |
| chr18 | 12614943 | 12615628 | Intron |
| chr18 | 12620704 | 12621001 | Intron |
| chr18 | 12621141 | 12622565 | Intron |
| chr18 | 12642202 | 12643775 | Intron |
| chr18 | 12656228 | 12656439 | Promoter |
| chr18 | 12656548 | 12656949 | Promoter |
| chr18 | 12660957 | 12665978 | 3' UTR |
| chr18 | 12669709 | 12669916 | Intron |
| chr18 | 12670027 | 12670818 | Intron |
| chr18 | 12671030 | 12671807 | Intron |
| chr18 | 12769009 | 12772294 | Distal Intergenic |
| chr18 | 12785827 | 12788188 | Exon |
| chr18 | 12801630 | 12802163 | Exon |
| chr18 | 12802507 | 12804055 | Intron |

| | | | |
|---|---|---|---|
| chr18 | 12806204 | 12806420 | Intron |
| chr18 | 12806700 | 12807590 | Intron |
| chr18 | 12809468 | 12810539 | Intron |
| chr18 | 12811248 | 12814552 | Exon |
| chr18 | 12844959 | 12845690 | Intron |
| chr18 | 12884057 | 12884874 | Promoter |
| chr18 | 12898581 | 12899711 | Distal Intergenic |
| chr18 | 13099759 | 13100012 | Intron |
| chr18 | 13113894 | 13114224 | Exon |
| chr18 | 13114386 | 13114652 | Exon |
| chr18 | 14840742 | 14840986 | Intron |
| chr18 | 14841039 | 14841530 | Intron |
| chr18 | 14931285 | 14931874 | Distal Intergenic |
| chr18 | 17449397 | 17449634 | Distal Intergenic |
| chr18 | 20117209 | 20120722 | Distal Intergenic |
| chr18 | 20130277 | 20130769 | Distal Intergenic |
| chr18 | 20151514 | 20151719 | Distal Intergenic |
| chr18 | 20151782 | 20152016 | Distal Intergenic |
| chr18 | 20152411 | 20152650 | Distal Intergenic |
| chr18 | 20156967 | 20157562 | Distal Intergenic |
| chr18 | 20157621 | 20157934 | Distal Intergenic |
| chr18 | 20158307 | 20158532 | Distal Intergenic |
| chr18 | 20158806 | 20161044 | Distal Intergenic |
| chr18 | 20164148 | 20165748 | Distal Intergenic |
| chr18 | 20185321 | 20185915 | Distal Intergenic |
| chr18 | 20192855 | 20193324 | Distal Intergenic |
| chr18 | 20193362 | 20193718 | Distal Intergenic |
| chr18 | 20264729 | 20265044 | Distal Intergenic |
| chr18 | 20265199 | 20265519 | Distal Intergenic |
| chr18 | 20300269 | 20302958 | Distal Intergenic |
| chr18 | 20304689 | 20304893 | Distal Intergenic |
| chr18 | 20307179 | 20307494 | Distal Intergenic |
| chr18 | 20307602 | 20307947 | Distal Intergenic |
| chr18 | 20315724 | 20317219 | Distal Intergenic |
| chr18 | 20345782 | 20350213 | Distal Intergenic |
| chr18 | 20386329 | 20387358 | Distal Intergenic |
| chr18 | 20533489 | 20535112 | Intron |
| chr18 | 20647658 | 20649118 | Distal Intergenic |
| chr18 | 20667868 | 20671124 | Distal Intergenic |
| chr18 | 20671201 | 20672111 | Distal Intergenic |
| chr18 | 20672279 | 20672494 | Distal Intergenic |
| chr18 | 20672588 | 20673053 | Distal Intergenic |
| chr18 | 20673278 | 20673725 | Distal Intergenic |
| chr18 | 20673844 | 20674955 | Distal Intergenic |

| | | | |
|---|---|---|---|
| chr18 | 20675189 | 20677095 | Distal Intergenic |
| chr18 | 20677191 | 20677507 | Distal Intergenic |
| chr18 | 20711142 | 20711910 | Distal Intergenic |
| chr18 | 20712128 | 20713676 | Promoter |
| chr18 | 20717042 | 20717320 | Intron |
| chr18 | 20941149 | 20941404 | Intron |
| chr18 | 21160435 | 21161036 | Intron |
| chr18 | 21310319 | 21311798 | Intron |
| chr18 | 23265144 | 23265343 | Distal Intergenic |
| chr18 | 23265392 | 23265594 | Distal Intergenic |
| chr18 | 23265715 | 23266016 | Distal Intergenic |
| chr18 | 23807537 | 23807747 | Intron |
| chr18 | 23846712 | 23847304 | Intron |
| chr18 | 23941461 | 23942646 | Intron |
| chr18 | 24025113 | 24026201 | Distal Intergenic |
| chr18 | 24034239 | 24035700 | 3' UTR |
| chr18 | 24035768 | 24037428 | 3' UTR |
| chr18 | 24113250 | 24113616 | Intron |
| chr18 | 24143488 | 24143933 | Intron |
| chr18 | 24178632 | 24178890 | Intron |
| chr18 | 24208952 | 24210879 | 5' UTR |
| chr18 | 24253539 | 24253777 | Distal Intergenic |
| chr18 | 24253946 | 24254369 | Distal Intergenic |
| chr18 | 24269591 | 24269790 | Exon |
| chr18 | 24269878 | 24271590 | Exon |
| chr18 | 24274833 | 24276489 | Intron |
| chr18 | 24334951 | 24335296 | Distal Intergenic |
| chr18 | 24361054 | 24361635 | Distal Intergenic |
| chr18 | 24688286 | 24688609 | Intron |
| chr18 | 24757309 | 24759460 | Intron |
| chr18 | 24760748 | 24761634 | Intron |
| chr18 | 24761690 | 24762317 | Intron |
| chr18 | 24762424 | 24762831 | Intron |
| chr18 | 24878506 | 24878737 | Distal Intergenic |
| chr18 | 24878805 | 24879012 | Distal Intergenic |
| chr18 | 24880557 | 24882265 | Distal Intergenic |
| chr18 | 24900623 | 24900986 | Distal Intergenic |
| chr18 | 25019194 | 25021025 | Intron |
| chr18 | 25129939 | 25130369 | Intron |
| chr18 | 25135442 | 25135641 | Intron |
| chr18 | 25139846 | 25140237 | Intron |
| chr18 | 25140290 | 25140589 | Intron |
| chr18 | 25210669 | 25210957 | Distal Intergenic |
| chr18 | 25250394 | 25250634 | Distal Intergenic |

| | | | |
|---|---|---|---|
| chr18 | 25443286 | 25443933 | Distal Intergenic |
| chr18 | 25525493 | 25526673 | Distal Intergenic |
| chr18 | 25643769 | 25645333 | Intron |
| chr18 | 25669279 | 25669512 | Intron |
| chr18 | 25788267 | 25789630 | Distal Intergenic |
| chr18 | 25796584 | 25797116 | Distal Intergenic |
| chr18 | 25797321 | 25797956 | Distal Intergenic |
| chr18 | 25797998 | 25798369 | Distal Intergenic |
| chr18 | 30109636 | 30110537 | Distal Intergenic |
| chr18 | 30494614 | 30495176 | Distal Intergenic |
| chr18 | 31499661 | 31500059 | Intron |
| chr18 | 31597187 | 31599251 | Intron |
| chr18 | 31919493 | 31920418 | Distal Intergenic |
| chr18 | 31948289 | 31948508 | Distal Intergenic |
| chr18 | 31948563 | 31949171 | Distal Intergenic |
| chr18 | 31963733 | 31963967 | Distal Intergenic |
| chr18 | 32211726 | 32212828 | Intron |
| chr18 | 32214528 | 32215305 | Intron |
| chr18 | 32227684 | 32228228 | Intron |
| chr18 | 32312618 | 32312924 | Intron |
| chr18 | 32399504 | 32400181 | Intron |
| chr18 | 32439846 | 32440121 | Intron |
| chr18 | 32495229 | 32495886 | Distal Intergenic |
| chr18 | 32502477 | 32502863 | Distal Intergenic |
| chr18 | 32502927 | 32503307 | Distal Intergenic |
| chr18 | 32506591 | 32507873 | Distal Intergenic |
| chr18 | 32537212 | 32537580 | Distal Intergenic |
| chr18 | 32779068 | 32779352 | Distal Intergenic |
| chr18 | 32793758 | 32793957 | Distal Intergenic |
| chr18 | 32794001 | 32795067 | Distal Intergenic |
| chr18 | 32798427 | 32799390 | Distal Intergenic |
| chr18 | 32800551 | 32802513 | Distal Intergenic |
| chr18 | 32803792 | 32804125 | Distal Intergenic |
| chr18 | 32804238 | 32806340 | Distal Intergenic |
| chr18 | 32806606 | 32808411 | Distal Intergenic |
| chr18 | 32808754 | 32809943 | Distal Intergenic |
| chr18 | 32996956 | 32997462 | Distal Intergenic |
| chr18 | 33372512 | 33373529 | Distal Intergenic |
| chr18 | 33373621 | 33374810 | Distal Intergenic |
| chr18 | 33954597 | 33956302 | Intron |
| chr18 | 33957127 | 33957384 | Intron |
| chr18 | 34008954 | 34010053 | Intron |
| chr18 | 34028799 | 34029050 | Intron |
| chr18 | 34047976 | 34051718 | Intron |

| | | | |
|---|---|---|---|
| chr18 | 34054658 | 34055697 | Intron |
| chr18 | 34067633 | 34068071 | Intron |
| chr18 | 34110211 | 34110584 | Intron |
| chr18 | 34146954 | 34147993 | Intron |
| chr18 | 34380644 | 34381758 | Intron |
| chr18 | 34428317 | 34428526 | Intron |
| chr18 | 34490241 | 34490710 | Intron |
| chr18 | 34531071 | 34532663 | Intron |
| chr18 | 34538149 | 34538469 | Promoter |
| chr18 | 34538509 | 34538841 | Promoter |
| chr18 | 34666623 | 34668348 | Intron |
| chr18 | 34783559 | 34785173 | Intron |
| chr18 | 34918504 | 34919351 | Intron |
| chr18 | 34936860 | 34937722 | Intron |
| chr18 | 35057124 | 35057515 | Intron |
| chr18 | 35060287 | 35061872 | Intron |
| chr18 | 35081496 | 35081935 | Intron |
| chr18 | 35089727 | 35090023 | Intron |
| chr18 | 35090136 | 35092443 | Intron |
| chr18 | 35092665 | 35092901 | Intron |
| chr18 | 35107914 | 35108722 | Intron |
| chr18 | 35111295 | 35111623 | Intron |
| chr18 | 35111801 | 35112144 | Intron |
| chr18 | 35112223 | 35112431 | Intron |
| chr18 | 35112571 | 35112786 | Intron |
| chr18 | 35112884 | 35113179 | Intron |
| chr18 | 35113416 | 35114258 | Intron |
| chr18 | 35114497 | 35115242 | Intron |
| chr18 | 35117676 | 35118346 | Intron |
| chr18 | 35118459 | 35118795 | Intron |
| chr18 | 35181872 | 35182742 | Distal Intergenic |
| chr18 | 35225248 | 35225706 | Distal Intergenic |
| chr18 | 35225795 | 35226060 | Distal Intergenic |
| chr18 | 35277965 | 35279863 | Distal Intergenic |
| chr18 | 35299233 | 35299493 | Distal Intergenic |
| chr18 | 35349815 | 35350160 | Distal Intergenic |
| chr18 | 35350389 | 35350911 | Distal Intergenic |
| chr18 | 35351200 | 35352402 | Distal Intergenic |
| chr18 | 35458701 | 35459719 | Distal Intergenic |
| chr18 | 35713004 | 35716387 | Distal Intergenic |
| chr18 | 35719795 | 35720287 | Distal Intergenic |
| chr18 | 35720818 | 35721066 | Distal Intergenic |
| chr18 | 35721119 | 35721489 | Distal Intergenic |
| chr18 | 35721858 | 35723918 | Distal Intergenic |

| | | | |
|---|---|---|---|
| chr18 | 35758308 | 35758684 | Distal Intergenic |
| chr18 | 35931225 | 35932365 | Distal Intergenic |
| chr18 | 36192829 | 36193554 | Distal Intergenic |
| chr18 | 36200837 | 36202701 | Distal Intergenic |
| chr18 | 36202974 | 36203471 | Distal Intergenic |
| chr18 | 36203518 | 36203727 | Distal Intergenic |
| chr18 | 36208225 | 36208482 | Distal Intergenic |
| chr18 | 36231166 | 36231409 | Distal Intergenic |
| chr18 | 36231750 | 36232050 | Distal Intergenic |
| chr18 | 36233990 | 36234431 | Distal Intergenic |
| chr18 | 36258817 | 36260157 | Distal Intergenic |
| chr18 | 36261936 | 36263279 | Distal Intergenic |
| chr18 | 36277293 | 36278078 | Distal Intergenic |
| chr18 | 36297916 | 36298989 | Distal Intergenic |
| chr18 | 36473840 | 36474060 | Distal Intergenic |
| chr18 | 36564146 | 36566151 | Distal Intergenic |
| chr18 | 36574379 | 36576807 | Distal Intergenic |
| chr18 | 36613604 | 36613838 | Distal Intergenic |
| chr18 | 36613904 | 36614127 | Exon |
| chr18 | 36652134 | 36653181 | Distal Intergenic |
| chr18 | 36673765 | 36674155 | Distal Intergenic |
| chr18 | 36674389 | 36674614 | Distal Intergenic |
| chr18 | 36675741 | 36676141 | Distal Intergenic |
| chr18 | 36676860 | 36677373 | Distal Intergenic |
| chr18 | 36681452 | 36681948 | Distal Intergenic |
| chr18 | 36693817 | 36694872 | Distal Intergenic |
| chr18 | 36718926 | 36719527 | Distal Intergenic |
| chr18 | 36752295 | 36752651 | Distal Intergenic |
| chr18 | 36762233 | 36763486 | Distal Intergenic |
| chr18 | 36816504 | 36816728 | Intron |
| chr18 | 36828936 | 36829175 | Intron |
| chr18 | 36829289 | 36830406 | Intron |
| chr18 | 36839260 | 36839569 | Intron |
| chr18 | 36868517 | 36868779 | Intron |
| chr18 | 36885854 | 36886494 | Intron |
| chr18 | 36894047 | 36895349 | Intron |
| chr18 | 36910137 | 36910336 | Intron |
| chr18 | 36942631 | 36943294 | Intron |
| chr18 | 36953518 | 36953833 | Intron |
| chr18 | 37803439 | 37803840 | Distal Intergenic |
| chr18 | 37803989 | 37804203 | Distal Intergenic |
| chr18 | 37966094 | 37966330 | Distal Intergenic |
| chr18 | 37985979 | 37986706 | Distal Intergenic |
| chr18 | 38007462 | 38007661 | Distal Intergenic |

| | | | |
|---|---|---|---|
| chr18 | 38008277 | 38008559 | Distal Intergenic |
| chr18 | 38008652 | 38010370 | Distal Intergenic |
| chr18 | 38035844 | 38039981 | Distal Intergenic |
| chr18 | 38040035 | 38040620 | Distal Intergenic |
| chr18 | 38043307 | 38044264 | Distal Intergenic |
| chr18 | 38083424 | 38084958 | Distal Intergenic |
| chr18 | 38085327 | 38086051 | Distal Intergenic |
| chr18 | 38087217 | 38088007 | Distal Intergenic |
| chr18 | 38093946 | 38094854 | Distal Intergenic |
| chr18 | 38123854 | 38124184 | Distal Intergenic |
| chr18 | 38239955 | 38241082 | Distal Intergenic |
| chr18 | 38304089 | 38304288 | Distal Intergenic |
| chr18 | 38365147 | 38365777 | Distal Intergenic |
| chr18 | 38409287 | 38410097 | Distal Intergenic |
| chr18 | 38410145 | 38410736 | Distal Intergenic |
| chr18 | 38411193 | 38413146 | Distal Intergenic |
| chr18 | 38414758 | 38415439 | Distal Intergenic |
| chr18 | 38415496 | 38415711 | Distal Intergenic |
| chr18 | 38418874 | 38420389 | Distal Intergenic |
| chr18 | 38433774 | 38434104 | Distal Intergenic |
| chr18 | 38434187 | 38434447 | Distal Intergenic |
| chr18 | 38456708 | 38457637 | Distal Intergenic |
| chr18 | 38507661 | 38509775 | Distal Intergenic |
| chr18 | 38543440 | 38545330 | Distal Intergenic |
| chr18 | 38566544 | 38566977 | Distal Intergenic |
| chr18 | 38578932 | 38579350 | Distal Intergenic |
| chr18 | 39040983 | 39041185 | Distal Intergenic |
| chr18 | 39041600 | 39042153 | Distal Intergenic |
| chr18 | 39471585 | 39471956 | Distal Intergenic |
| chr18 | 39478677 | 39479058 | Distal Intergenic |
| chr18 | 39479104 | 39479373 | Distal Intergenic |
| chr18 | 39533891 | 39535318 | Promoter |
| chr18 | 39538158 | 39538501 | Intron |
| chr18 | 39540422 | 39541468 | Intron |
| chr18 | 39736858 | 39737132 | Distal Intergenic |
| chr18 | 39737262 | 39737542 | Distal Intergenic |
| chr18 | 39896727 | 39897827 | Intron |
| chr18 | 39900947 | 39901178 | Intron |
| chr18 | 39959737 | 39960085 | Intron |
| chr18 | 40359831 | 40360242 | Intron |
| chr18 | 40378813 | 40379083 | Intron |
| chr18 | 40642477 | 40642730 | Intron |
| chr18 | 42207617 | 42207965 | Distal Intergenic |
| chr18 | 42208012 | 42208247 | Distal Intergenic |

| | | | |
|---|---|---|---|
| chr18 | 42212080 | 42212447 | Distal Intergenic |
| chr18 | 42212654 | 42213232 | Distal Intergenic |
| chr18 | 42221300 | 42221770 | Distal Intergenic |
| chr18 | 42282395 | 42282997 | Intron |
| chr18 | 42283189 | 42284796 | Intron |
| chr18 | 42421147 | 42421598 | Intron |
| chr18 | 42434594 | 42434861 | Intron |
| chr18 | 42435012 | 42435436 | Intron |
| chr18 | 42435953 | 42436357 | Intron |
| chr18 | 42467188 | 42468533 | Intron |
| chr18 | 42471571 | 42472447 | Intron |
| chr18 | 42493906 | 42494332 | Intron |
| chr18 | 42671024 | 42671938 | Distal Intergenic |
| chr18 | 42672106 | 42672561 | Distal Intergenic |
| chr18 | 42688444 | 42689904 | Distal Intergenic |
| chr18 | 42723024 | 42723265 | Distal Intergenic |
| chr18 | 42926703 | 42927498 | Intron |
| chr18 | 42929769 | 42931639 | Intron |
| chr18 | 42952393 | 42953157 | Intron |
| chr18 | 42988504 | 42989138 | Intron |
| chr18 | 43592549 | 43594201 | Intron |
| chr18 | 43595788 | 43597347 | Exon |
| chr18 | 43627626 | 43629866 | Intron |
| chr18 | 43968424 | 43968664 | Intron |
| chr18 | 43976276 | 43977545 | Intron |
| chr18 | 43977884 | 43978700 | Intron |
| chr18 | 43986232 | 43986444 | Intron |
| chr18 | 44006525 | 44007827 | Intron |
| chr18 | 44022831 | 44024136 | Intron |
| chr18 | 44090515 | 44091146 | Intron |
| chr18 | 44121004 | 44121910 | Exon |
| chr18 | 44122813 | 44123022 | 5' UTR |
| chr18 | 44126871 | 44128391 | 5' UTR |
| chr18 | 44200277 | 44200555 | Intron |
| chr18 | 44236007 | 44236279 | Promoter |
| chr18 | 44540182 | 44540424 | Intron |
| chr18 | 44540528 | 44541078 | Intron |
| chr18 | 44554358 | 44554600 | Exon |
| chr18 | 44554700 | 44554907 | Exon |
| chr18 | 44631697 | 44633244 | Downstream |
| chr18 | 44699778 | 44701266 | Intron |
| chr18 | 44701817 | 44703346 | Promoter |
| chr18 | 44756430 | 44757837 | Exon |
| chr18 | 44794821 | 44795140 | Distal Intergenic |

| | | | |
|---|---|---|---|
| chr18 | 44803320 | 44803527 | Distal Intergenic |
| chr18 | 44814260 | 44815134 | Intron |
| chr18 | 44815617 | 44815909 | Intron |
| chr18 | 44820913 | 44822740 | Intron |
| chr18 | 44827507 | 44829021 | Intron |
| chr18 | 45420232 | 45420598 | Intron |
| chr18 | 45420754 | 45421048 | Intron |
| chr18 | 45438131 | 45439736 | Intron |
| chr18 | 46439785 | 46440075 | Distal Intergenic |
| chr18 | 46440483 | 46440688 | Distal Intergenic |
| chr18 | 46440804 | 46441003 | Distal Intergenic |
| chr18 | 46484639 | 46484844 | Distal Intergenic |
| chr18 | 46540859 | 46541276 | Distal Intergenic |
| chr18 | 46549712 | 46550050 | Distal Intergenic |
| chr18 | 46683090 | 46683333 | Intron |
| chr18 | 46683918 | 46685269 | Intron |
| chr18 | 46685955 | 46686168 | Intron |
| chr18 | 46701036 | 46701915 | Intron |
| chr18 | 46756451 | 46756964 | Intron |
| chr18 | 46757409 | 46759053 | Intron |
| chr18 | 46901248 | 46901458 | Intron |
| chr18 | 47122705 | 47122964 | Distal Intergenic |
| chr18 | 47603240 | 47603584 | Intron |
| chr18 | 50145958 | 50146246 | Intron |
| chr18 | 50146293 | 50147114 | Intron |
| chr18 | 50149922 | 50150149 | Intron |
| chr18 | 50177454 | 50178330 | Intron |
| chr18 | 50182591 | 50184506 | Intron |
| chr18 | 50189871 | 50191710 | Intron |
| chr18 | 50191748 | 50192227 | Intron |
| chr18 | 50192293 | 50192632 | Intron |
| chr18 | 50220730 | 50220935 | Intron |
| chr18 | 50221099 | 50223501 | Intron |
| chr18 | 50223576 | 50223833 | Intron |
| chr18 | 50223875 | 50226172 | Intron |
| chr18 | 50255558 | 50256169 | Intron |
| chr18 | 51733811 | 51734908 | Intron |
| chr18 | 53062317 | 53063825 | Intron |
| chr18 | 53099895 | 53100459 | Intron |
| chr18 | 53357632 | 53358077 | Distal Intergenic |
| chr18 | 53397958 | 53398252 | Distal Intergenic |
| chr18 | 53409036 | 53409371 | Distal Intergenic |
| chr18 | 53409649 | 53410309 | Distal Intergenic |
| chr18 | 53505219 | 53507094 | Distal Intergenic |

| | | | |
|---|---|---|---|
| chr18 | 53576920 | 53577125 | Distal Intergenic |
| chr18 | 53850156 | 53852012 | Distal Intergenic |
| chr18 | 53904483 | 53904733 | Distal Intergenic |
| chr18 | 53961962 | 53962171 | Distal Intergenic |
| chr18 | 54021413 | 54021619 | Distal Intergenic |
| chr18 | 54021835 | 54022149 | Distal Intergenic |
| chr18 | 54022498 | 54022793 | Distal Intergenic |
| chr18 | 54063021 | 54063640 | Distal Intergenic |
| chr18 | 55019925 | 55020927 | Promoter |
| chr18 | 55089222 | 55089519 | Distal Intergenic |
| chr18 | 55340072 | 55340480 | Intron |
| chr18 | 55355727 | 55356503 | Exon |
| chr18 | 55425733 | 55425981 | Intron |
| chr18 | 55783608 | 55784056 | Intron |
| chr18 | 55806875 | 55808264 | Intron |
| chr18 | 56579253 | 56579547 | Intron |
| chr18 | 56603114 | 56603330 | Intron |
| chr18 | 56637020 | 56637781 | Intron |
| chr18 | 56650526 | 56651085 | Intron |
| chr18 | 56773594 | 56775109 | Distal Intergenic |
| chr18 | 56846717 | 56849111 | Distal Intergenic |
| chr18 | 56866777 | 56867166 | Distal Intergenic |
| chr18 | 56867488 | 56867729 | Distal Intergenic |
| chr18 | 56868681 | 56869028 | Distal Intergenic |
| chr18 | 56881925 | 56882277 | Distal Intergenic |
| chr18 | 56882337 | 56884299 | Distal Intergenic |
| chr18 | 56885370 | 56887047 | Promoter |
| chr18 | 56896035 | 56897467 | Intron |
| chr18 | 57241683 | 57242574 | Intron |
| chr18 | 57244134 | 57244819 | Intron |
| chr18 | 57945935 | 57946509 | Distal Intergenic |
| chr18 | 57947697 | 57947972 | Distal Intergenic |
| chr18 | 57948017 | 57948539 | Distal Intergenic |
| chr18 | 57964260 | 57965738 | Distal Intergenic |
| chr18 | 57970729 | 57970994 | Distal Intergenic |
| chr18 | 57996296 | 57997932 | Distal Intergenic |
| chr18 | 58015872 | 58016242 | Distal Intergenic |
| chr18 | 58017467 | 58018281 | Distal Intergenic |
| chr18 | 58727634 | 58728138 | Distal Intergenic |
| chr18 | 60640390 | 60641577 | Intron |
| chr18 | 60660058 | 60663173 | Distal Intergenic |
| chr18 | 60683146 | 60683468 | Distal Intergenic |
| chr18 | 60684951 | 60685826 | Distal Intergenic |
| chr18 | 60686301 | 60687530 | Distal Intergenic |

| | | | |
|---|---|---|---|
| chr18 | 60808238 | 60809539 | Intron |
| chr18 | 60828667 | 60829264 | Intron |
| chr18 | 60895935 | 60897242 | Intron |
| chr18 | 60899813 | 60901492 | Intron |
| chr18 | 60905394 | 60907859 | Intron |
| chr18 | 60908274 | 60909126 | Intron |
| chr18 | 60909794 | 60910761 | Intron |
| chr18 | 60910945 | 60912372 | Intron |
| chr18 | 60915830 | 60916664 | Intron |
| chr18 | 60934015 | 60934454 | Intron |
| chr18 | 60934496 | 60935390 | Intron |
| chr18 | 60935573 | 60935854 | Intron |
| chr18 | 61336005 | 61336417 | Distal Intergenic |
| chr18 | 61336546 | 61336805 | Distal Intergenic |
| chr18 | 61374563 | 61374814 | Intron |
| chr18 | 61375169 | 61375472 | Intron |
| chr18 | 61383339 | 61384650 | Exon |
| chr18 | 61676610 | 61676919 | Distal Intergenic |
| chr18 | 61710264 | 61710580 | Distal Intergenic |
| chr18 | 61714272 | 61714648 | Distal Intergenic |
| chr18 | 61714904 | 61717074 | Distal Intergenic |
| chr18 | 61778826 | 61779040 | Intron |
| chr18 | 61785471 | 61787216 | Intron |
| chr18 | 61791220 | 61794372 | Intron |
| chr18 | 61794629 | 61794829 | Exon |
| chr18 | 61795651 | 61800558 | Intron |
| chr18 | 61850762 | 61851022 | Intron |
| chr18 | 61904107 | 61906240 | Intron |
| chr18 | 61934120 | 61934535 | Intron |
| chr18 | 61934842 | 61935189 | Intron |
| chr18 | 61997109 | 61997712 | Intron |
| chr18 | 62101037 | 62101236 | Distal Intergenic |
| chr18 | 62111432 | 62113033 | Distal Intergenic |
| chr18 | 62175877 | 62176589 | Distal Intergenic |
| chr18 | 62176941 | 62177356 | Distal Intergenic |
| chr18 | 62191913 | 62194436 | Distal Intergenic |
| chr18 | 62260600 | 62263769 | Distal Intergenic |
| chr18 | 62281059 | 62282911 | Distal Intergenic |
| chr18 | 62328359 | 62329825 | Distal Intergenic |
| chr18 | 62338047 | 62338598 | Distal Intergenic |
| chr18 | 62340066 | 62340433 | Distal Intergenic |
| chr18 | 62362423 | 62364257 | Distal Intergenic |
| chr18 | 62708106 | 62708323 | Distal Intergenic |
| chr18 | 62708593 | 62708804 | Distal Intergenic |

| | | | |
|---|---|---|---|
| chr18 | 62744868 | 62745090 | Distal Intergenic |
| chr18 | 63065731 | 63066783 | Distal Intergenic |
| chr18 | 63153376 | 63153983 | Distal Intergenic |
| chr18 | 63200230 | 63200959 | Distal Intergenic |
| chr18 | 63202300 | 63202507 | Distal Intergenic |
| chr18 | 63560375 | 63560805 | Distal Intergenic |
| chr18 | 63622724 | 63623451 | Distal Intergenic |
| chr18 | 63850337 | 63850558 | Distal Intergenic |
| chr18 | 63850819 | 63851991 | Distal Intergenic |
| chr18 | 63852100 | 63852407 | Distal Intergenic |
| chr18 | 63990531 | 63990880 | Distal Intergenic |
| chr18 | 64041395 | 64041605 | Distal Intergenic |
| chr18 | 64641891 | 64643014 | Distal Intergenic |
| chr18 | 64672125 | 64672446 | Distal Intergenic |
| chr18 | 64675457 | 64676025 | Distal Intergenic |
| chr18 | 65098831 | 65099964 | Distal Intergenic |
| chr18 | 65104575 | 65105990 | Distal Intergenic |
| chr18 | 65210081 | 65211609 | Intron |
| chr18 | 65269174 | 65269817 | Intron |
| chr18 | 65540746 | 65542016 | Exon |
| chr18 | 65725132 | 65725430 | Distal Intergenic |
| chr18 | 65725523 | 65725824 | Distal Intergenic |
| chr18 | 65739591 | 65740367 | Distal Intergenic |
| chr18 | 65740488 | 65741048 | Distal Intergenic |
| chr18 | 65757573 | 65758479 | Distal Intergenic |
| chr18 | 65762992 | 65763831 | Distal Intergenic |
| chr18 | 65765222 | 65765429 | Distal Intergenic |
| chr18 | 65766004 | 65766308 | Distal Intergenic |
| chr18 | 65766354 | 65767335 | Distal Intergenic |
| chr18 | 65775701 | 65775900 | Distal Intergenic |
| chr18 | 65785871 | 65788498 | Distal Intergenic |
| chr18 | 65829306 | 65830357 | Distal Intergenic |
| chr18 | 65857777 | 65859517 | Distal Intergenic |
| chr18 | 65864325 | 65866253 | Distal Intergenic |
| chr18 | 65897908 | 65898252 | Distal Intergenic |
| chr18 | 65898971 | 65899762 | Distal Intergenic |
| chr18 | 65954727 | 65955184 | Distal Intergenic |
| chr18 | 66618418 | 66618626 | Intron |
| chr18 | 66631501 | 66631881 | Intron |
| chr18 | 66632258 | 66632737 | Intron |
| chr18 | 66637606 | 66637950 | Intron |
| chr18 | 67126462 | 67126712 | Intron |
| chr18 | 67354012 | 67355232 | Intron |
| chr18 | 67458941 | 67459572 | Intron |

| | | | |
|---|---|---|---|
| chr18 | 67461198 | 67461404 | Intron |
| chr18 | 67461669 | 67461888 | Intron |
| chr18 | 67469595 | 67470529 | Intron |
| chr18 | 67470578 | 67470786 | Intron |
| chr18 | 67471773 | 67472024 | Intron |
| chr18 | 67477346 | 67478645 | Intron |
| chr18 | 67550828 | 67551033 | Intron |
| chr18 | 67551257 | 67551901 | Intron |
| chr18 | 67709360 | 67709810 | Intron |
| chr18 | 67737156 | 67738268 | Intron |
| chr18 | 67801293 | 67801604 | Intron |
| chr18 | 67865930 | 67866185 | Promoter |
| chr18 | 67870600 | 67871465 | 5' UTR |
| chr18 | 67882916 | 67883438 | Distal Intergenic |
| chr18 | 67934697 | 67935099 | Distal Intergenic |
| chr18 | 67974947 | 67975317 | Intron |
| chr18 | 68105453 | 68105697 | Distal Intergenic |
| chr18 | 68142964 | 68145571 | Distal Intergenic |
| chr18 | 68145624 | 68145916 | Distal Intergenic |
| chr18 | 68216994 | 68217813 | Distal Intergenic |
| chr18 | 68217888 | 68218719 | Distal Intergenic |
| chr18 | 68243747 | 68244491 | Distal Intergenic |
| chr18 | 68278929 | 68279539 | Distal Intergenic |
| chr18 | 68314972 | 68315175 | Distal Intergenic |
| chr18 | 68316069 | 68318380 | Distal Intergenic |
| chr18 | 68319769 | 68320006 | Distal Intergenic |
| chr18 | 68320121 | 68320475 | Distal Intergenic |
| chr18 | 68349730 | 68350289 | Distal Intergenic |
| chr18 | 68351192 | 68351433 | Distal Intergenic |
| chr18 | 68460030 | 68460302 | Distal Intergenic |
| chr18 | 68947620 | 68948325 | Distal Intergenic |
| chr18 | 68948577 | 68948979 | Distal Intergenic |
| chr18 | 68988867 | 68989238 | Distal Intergenic |
| chr18 | 69504945 | 69505211 | Distal Intergenic |
| chr18 | 69505444 | 69505729 | Distal Intergenic |
| chr18 | 69814833 | 69815196 | Distal Intergenic |
| chr18 | 69973753 | 69974567 | Distal Intergenic |
| chr18 | 70085423 | 70085673 | Distal Intergenic |
| chr18 | 70152054 | 70152454 | Distal Intergenic |
| chr18 | 70182178 | 70182443 | Distal Intergenic |
| chr18 | 70182593 | 70182945 | Distal Intergenic |
| chr18 | 70403773 | 70404717 | Distal Intergenic |
| chr18 | 70412844 | 70414011 | 3' UTR |
| chr18 | 70690093 | 70690424 | Distal Intergenic |

| | | | |
|---|---|---|---|
| chr18 | 70690636 | 70690891 | Distal Intergenic |
| chr18 | 70728788 | 70729017 | Distal Intergenic |
| chr18 | 70760613 | 70761703 | Distal Intergenic |
| chr18 | 70803858 | 70804480 | Distal Intergenic |
| chr18 | 71293236 | 71293501 | Distal Intergenic |
| chr18 | 73462849 | 73463094 | Distal Intergenic |
| chr18 | 73463214 | 73464420 | Distal Intergenic |
| chr18 | 73731142 | 73731772 | Distal Intergenic |
| chr18 | 73732424 | 73732819 | Distal Intergenic |
| chr18 | 73732973 | 73733207 | Distal Intergenic |
| chr18 | 73733329 | 73734597 | Distal Intergenic |
| chr18 | 73734730 | 73734931 | Distal Intergenic |
| chr18 | 73876589 | 73877335 | Distal Intergenic |
| chr18 | 73877729 | 73878210 | Distal Intergenic |
| chr18 | 74507372 | 74507937 | Exon |
| chr18 | 75106607 | 75107052 | Distal Intergenic |
| chr18 | 75164597 | 75165400 | Distal Intergenic |
| chr18 | 75330400 | 75331843 | Distal Intergenic |
| chr18 | 75352813 | 75353810 | Distal Intergenic |
| chr18 | 75501493 | 75502381 | Distal Intergenic |
| chr18 | 75502594 | 75502847 | Distal Intergenic |
| chr18 | 75504394 | 75505161 | Distal Intergenic |
| chr18 | 75522746 | 75522976 | Distal Intergenic |
| chr18 | 75523267 | 75524593 | Distal Intergenic |
| chr18 | 75526434 | 75526800 | Distal Intergenic |
| chr18 | 75527698 | 75528645 | Distal Intergenic |
| chr18 | 75528757 | 75529066 | Distal Intergenic |
| chr18 | 75529228 | 75529735 | Distal Intergenic |
| chr18 | 75529900 | 75530254 | Distal Intergenic |
| chr18 | 75530491 | 75531881 | Distal Intergenic |
| chr18 | 75532428 | 75532747 | Distal Intergenic |
| chr18 | 75532819 | 75533933 | Distal Intergenic |
| chr18 | 75534290 | 75535382 | Distal Intergenic |
| chr18 | 75539742 | 75539986 | Distal Intergenic |
| chr18 | 75546483 | 75546785 | Distal Intergenic |
| chr18 | 75546943 | 75547168 | Distal Intergenic |
| chr18 | 75614997 | 75616186 | Distal Intergenic |
| chr18 | 75616255 | 75616482 | Distal Intergenic |
| chr18 | 75654220 | 75654891 | Distal Intergenic |
| chr18 | 75715454 | 75716162 | Distal Intergenic |
| chr18 | 75716341 | 75716585 | Distal Intergenic |
| chr18 | 76018003 | 76018448 | Distal Intergenic |
| chr18 | 76020595 | 76021645 | Distal Intergenic |
| chr18 | 76030873 | 76031103 | Distal Intergenic |

| | | | |
|---|---|---|---|
| chr18 | 76212797 | 76213656 | Distal Intergenic |
| chr18 | 76256477 | 76257479 | Distal Intergenic |
| chr18 | 76269988 | 76271409 | Distal Intergenic |
| chr18 | 76443035 | 76443259 | Distal Intergenic |
| chr18 | 76495076 | 76495861 | Distal Intergenic |
| chr18 | 77293263 | 77293935 | Distal Intergenic |
| chr18 | 77489738 | 77489981 | Intron |
| chr18 | 77902454 | 77903299 | Distal Intergenic |
| chr18 | 78038315 | 78038904 | Distal Intergenic |
| chr18 | 78118606 | 78119227 | Distal Intergenic |
| chr18 | 78141439 | 78141948 | Distal Intergenic |
| chr18 | 78145557 | 78145787 | Distal Intergenic |
| chr18 | 78677129 | 78677924 | Distal Intergenic |
| chr18 | 78690448 | 78690746 | Distal Intergenic |
| chr18 | 78694701 | 78694910 | Distal Intergenic |
| chr18 | 78774226 | 78774537 | Distal Intergenic |
| chr18 | 79205202 | 79205979 | Distal Intergenic |
| chr18 | 79393246 | 79393528 | Distal Intergenic |
| chr18 | 79663635 | 79664482 | Distal Intergenic |
| chr18 | 80201983 | 80203710 | Distal Intergenic |
| chr18 | 80243829 | 80245229 | Distal Intergenic |
| chr18 | 80246770 | 80247031 | Distal Intergenic |
| chr18 | 80247208 | 80249265 | Distal Intergenic |
| chr18 | 80253448 | 80254697 | Distal Intergenic |
| chr18 | 80273177 | 80274650 | Distal Intergenic |
| chr18 | 80296996 | 80297300 | Distal Intergenic |
| chr18 | 80300125 | 80300332 | Distal Intergenic |
| chr18 | 80300623 | 80300840 | Distal Intergenic |
| chr18 | 80306721 | 80309944 | Distal Intergenic |
| chr18 | 80312301 | 80313439 | Distal Intergenic |
| chr18 | 80346953 | 80347648 | Distal Intergenic |
| chr18 | 80348038 | 80348474 | Distal Intergenic |
| chr18 | 80366125 | 80367739 | Distal Intergenic |
| chr18 | 80369586 | 80369871 | Distal Intergenic |
| chr18 | 80449152 | 80450543 | Distal Intergenic |
| chr18 | 80452008 | 80452896 | Distal Intergenic |
| chr18 | 80453063 | 80453353 | Distal Intergenic |
| chr18 | 80453934 | 80454239 | Distal Intergenic |
| chr18 | 80457615 | 80460848 | Distal Intergenic |
| chr18 | 80511795 | 80512015 | Distal Intergenic |
| chr18 | 80666963 | 80667209 | Distal Intergenic |
| chr18 | 80751372 | 80751651 | Distal Intergenic |
| chr18 | 80751800 | 80752186 | Distal Intergenic |
| chr18 | 80780475 | 80780742 | Distal Intergenic |

| | | | |
|---|---|---|---|
| chr18 | 80852615 | 80852814 | Distal Intergenic |
| chr18 | 80853329 | 80853532 | Distal Intergenic |
| chr18 | 81129530 | 81129821 | Distal Intergenic |
| chr18 | 81130056 | 81130863 | Distal Intergenic |
| chr18 | 81248089 | 81248448 | Distal Intergenic |
| chr18 | 81248646 | 81248866 | Distal Intergenic |
| chr18 | 82104303 | 82104516 | Distal Intergenic |
| chr18 | 82657817 | 82659997 | Distal Intergenic |
| chr18 | 82661810 | 82662602 | Distal Intergenic |
| chr18 | 82662800 | 82664320 | Distal Intergenic |
| chr18 | 82667717 | 82667916 | Distal Intergenic |
| chr18 | 82668144 | 82668735 | Distal Intergenic |
| chr18 | 82687294 | 82687596 | Distal Intergenic |
| chr18 | 82691292 | 82692911 | Distal Intergenic |
| chr18 | 82694040 | 82694246 | Distal Intergenic |
| chr18 | 82861368 | 82861930 | Distal Intergenic |
| chr18 | 82862364 | 82862802 | Distal Intergenic |
| chr18 | 82907185 | 82907487 | Distal Intergenic |
| chr18 | 83082812 | 83083105 | Distal Intergenic |
| chr18 | 83084609 | 83084810 | Distal Intergenic |
| chr18 | 83085702 | 83086505 | Distal Intergenic |
| chr18 | 83503303 | 83503706 | Distal Intergenic |
| chr18 | 84064368 | 84064656 | Distal Intergenic |
| chr18 | 84478746 | 84479430 | Distal Intergenic |
| chr18 | 84758904 | 84759862 | Distal Intergenic |
| chr18 | 85049222 | 85050269 | Distal Intergenic |
| chr18 | 85050378 | 85050953 | Distal Intergenic |
| chr18 | 85063395 | 85063701 | Distal Intergenic |
| chr18 | 85063738 | 85063962 | Distal Intergenic |
| chr18 | 85593934 | 85594924 | Distal Intergenic |
| chr18 | 87639611 | 87639814 | Distal Intergenic |
| chr18 | 89004551 | 89004908 | Distal Intergenic |
| chr18 | 89019932 | 89020255 | Distal Intergenic |
| chr18 | 89062557 | 89063445 | Distal Intergenic |
| chr18 | 89068642 | 89069808 | Distal Intergenic |
| chr18 | 89077287 | 89078016 | Distal Intergenic |
| chr18 | 89096863 | 89097452 | Distal Intergenic |
| chr18 | 89097624 | 89098691 | Distal Intergenic |
| chr18 | 89267206 | 89267691 | Distal Intergenic |
| chr18 | 89728959 | 89729357 | Distal Intergenic |
| chr19 | 3282612 | 3283277 | Intron |
| chr19 | 3324298 | 3325167 | Distal Intergenic |
| chr19 | 3333768 | 3334167 | Distal Intergenic |
| chr19 | 3353666 | 3353930 | Distal Intergenic |

| | | | |
|---|---|---|---|
| chr19 | 3519231 | 3520047 | Intron |
| chr19 | 3525965 | 3526817 | Exon |
| chr19 | 3554985 | 3555392 | Intron |
| chr19 | 3574776 | 3575152 | Intron |
| chr19 | 3575477 | 3575702 | Exon |
| chr19 | 3624452 | 3626617 | Promoter |
| chr19 | 3654801 | 3655069 | Intron |
| chr19 | 3663627 | 3663867 | Intron |
| chr19 | 3664098 | 3664382 | Intron |
| chr19 | 3668830 | 3670112 | Intron |
| chr19 | 3767947 | 3768704 | Downstream |
| chr19 | 3832986 | 3833185 | Intron |
| chr19 | 3838856 | 3840693 | Intron |
| chr19 | 3852491 | 3852690 | Exon |
| chr19 | 3903864 | 3904965 | Intron |
| chr19 | 3905229 | 3906868 | Exon |
| chr19 | 3907144 | 3907514 | Intron |
| chr19 | 3913087 | 3913366 | Intron |
| chr19 | 4000315 | 4001190 | Distal Intergenic |
| chr19 | 4045479 | 4047471 | 3' UTR |
| chr19 | 4093628 | 4094210 | Intron |
| chr19 | 4099206 | 4099944 | Exon |
| chr19 | 4112984 | 4113605 | Intron |
| chr19 | 4113984 | 4114245 | Intron |
| chr19 | 4125790 | 4127367 | Distal Intergenic |
| chr19 | 4134842 | 4136353 | Distal Intergenic |
| chr19 | 4148560 | 4148855 | Distal Intergenic |
| chr19 | 4192196 | 4192916 | Intron |
| chr19 | 4201237 | 4201611 | Intron |
| chr19 | 4213594 | 4214810 | Intron |
| chr19 | 4214861 | 4215232 | Intron |
| chr19 | 4231886 | 4232110 | Intron |
| chr19 | 4243983 | 4245835 | Distal Intergenic |
| chr19 | 4268528 | 4269379 | 3' UTR |
| chr19 | 4298828 | 4299175 | Intron |
| chr19 | 4299561 | 4300822 | Intron |
| chr19 | 4305419 | 4306064 | Promoter |
| chr19 | 4326051 | 4326818 | Intron |
| chr19 | 4332491 | 4334653 | Exon |
| chr19 | 4395038 | 4396721 | Intron |
| chr19 | 4396870 | 4397119 | Intron |
| chr19 | 4419043 | 4421747 | Intron |
| chr19 | 4429884 | 4431398 | Exon |
| chr19 | 4437845 | 4441362 | Intron |

| | | | |
|---|---|---|---|
| chr19 | 4478046 | 4478263 | Intron |
| chr19 | 4481727 | 4482120 | Intron |
| chr19 | 4614750 | 4614959 | Distal Intergenic |
| chr19 | 4615563 | 4615804 | Distal Intergenic |
| chr19 | 4625367 | 4625643 | Distal Intergenic |
| chr19 | 4625817 | 4626087 | Distal Intergenic |
| chr19 | 4665051 | 4665953 | Intron |
| chr19 | 4670650 | 4672890 | Promoter |
| chr19 | 4686447 | 4687422 | Intron |
| chr19 | 4720619 | 4721052 | Intron |
| chr19 | 4721223 | 4721475 | Intron |
| chr19 | 4729514 | 4730001 | Distal Intergenic |
| chr19 | 4741384 | 4741697 | Distal Intergenic |
| chr19 | 4741774 | 4742072 | Distal Intergenic |
| chr19 | 4755814 | 4756105 | Distal Intergenic |
| chr19 | 4793265 | 4794773 | 3' UTR |
| chr19 | 4811177 | 4811584 | Distal Intergenic |
| chr19 | 4811841 | 4812280 | Distal Intergenic |
| chr19 | 4858849 | 4859661 | Exon |
| chr19 | 4928406 | 4928623 | Intron |
| chr19 | 4962053 | 4962584 | 3' UTR |
| chr19 | 5012829 | 5013851 | Intron |
| chr19 | 5015918 | 5017012 | Promoter |
| chr19 | 5017117 | 5019442 | Promoter |
| chr19 | 5041116 | 5041760 | Exon |
| chr19 | 5049658 | 5052012 | Intron |
| chr19 | 5057526 | 5059528 | Intron |
| chr19 | 5060302 | 5060649 | Intron |
| chr19 | 5060696 | 5061336 | Intron |
| chr19 | 5061409 | 5061763 | Intron |
| chr19 | 5088109 | 5088308 | Intron |
| chr19 | 5256890 | 5257521 | Intron |
| chr19 | 5294856 | 5295440 | Intron |
| chr19 | 5300391 | 5301036 | Intron |
| chr19 | 5345292 | 5346475 | Distal Intergenic |
| chr19 | 5366706 | 5367693 | Distal Intergenic |
| chr19 | 5388533 | 5389385 | Distal Intergenic |
| chr19 | 5424333 | 5424808 | Distal Intergenic |
| chr19 | 5424998 | 5425708 | Distal Intergenic |
| chr19 | 5428075 | 5431029 | Distal Intergenic |
| chr19 | 5431688 | 5433252 | Distal Intergenic |
| chr19 | 5433464 | 5434132 | Distal Intergenic |
| chr19 | 5445157 | 5445376 | Distal Intergenic |
| chr19 | 5447677 | 5449202 | Distal Intergenic |

| | | | |
|---|---|---|---|
| chr19 | 5449373 | 5450222 | Distal Intergenic |
| chr19 | 5488747 | 5489053 | Distal Intergenic |
| chr19 | 5489218 | 5489461 | Distal Intergenic |
| chr19 | 5490577 | 5491047 | Distal Intergenic |
| chr19 | 5491215 | 5491643 | Distal Intergenic |
| chr19 | 5529575 | 5530104 | Distal Intergenic |
| chr19 | 5566836 | 5567596 | Promoter |
| chr19 | 5572278 | 5572946 | Distal Intergenic |
| chr19 | 5609327 | 5610535 | Exon |
| chr19 | 5637495 | 5637694 | Intron |
| chr19 | 5637792 | 5638058 | Intron |
| chr19 | 5688850 | 5690216 | Promoter |
| chr19 | 5703587 | 5705539 | Intron |
| chr19 | 5713640 | 5714311 | 5' UTR |
| chr19 | 5724393 | 5726300 | Exon |
| chr19 | 5729111 | 5729327 | Intron |
| chr19 | 5729364 | 5729706 | Intron |
| chr19 | 5749609 | 5749822 | Intron |
| chr19 | 5793205 | 5793767 | Distal Intergenic |
| chr19 | 5793908 | 5794338 | Distal Intergenic |
| chr19 | 5794399 | 5795795 | Distal Intergenic |
| chr19 | 5795949 | 5796185 | Distal Intergenic |
| chr19 | 5799159 | 5805004 | Distal Intergenic |
| chr19 | 5805055 | 5805295 | Distal Intergenic |
| chr19 | 5808813 | 5809285 | Distal Intergenic |
| chr19 | 5809456 | 5811535 | Distal Intergenic |
| chr19 | 5811581 | 5812851 | Distal Intergenic |
| chr19 | 5814443 | 5816725 | Distal Intergenic |
| chr19 | 5817291 | 5818095 | Distal Intergenic |
| chr19 | 5818168 | 5818382 | Distal Intergenic |
| chr19 | 5831357 | 5832605 | 5' UTR |
| chr19 | 5836447 | 5836886 | Intron |
| chr19 | 5837147 | 5845880 | Promoter |
| chr19 | 5846110 | 5846476 | 5' UTR |
| chr19 | 5846621 | 5850851 | Promoter |
| chr19 | 5851076 | 5852470 | Promoter |
| chr19 | 5877538 | 5878069 | Intron |
| chr19 | 5911942 | 5912807 | Distal Intergenic |
| chr19 | 5912950 | 5913349 | Promoter |
| chr19 | 5944854 | 5945965 | Intron |
| chr19 | 5953766 | 5954417 | Intron |
| chr19 | 5963927 | 5964151 | Intron |
| chr19 | 5964772 | 5965213 | Intron |
| chr19 | 5965451 | 5965832 | Intron |

| | | | |
|---|---|---|---|
| chr19 | 5965997 | 5967365 | Intron |
| chr19 | 5991666 | 5992171 | Intron |
| chr19 | 6014057 | 6015970 | Intron |
| chr19 | 6046112 | 6046460 | Intron |
| chr19 | 6057959 | 6058288 | Intron |
| chr19 | 6058336 | 6058976 | Intron |
| chr19 | 6059751 | 6059950 | Intron |
| chr19 | 6060276 | 6061417 | Intron |
| chr19 | 6061710 | 6062409 | Intron |
| chr19 | 6076647 | 6077250 | Intron |
| chr19 | 6084906 | 6085919 | Intron |
| chr19 | 6117389 | 6119653 | Distal Intergenic |
| chr19 | 6125006 | 6125709 | Distal Intergenic |
| chr19 | 6127564 | 6128278 | Distal Intergenic |
| chr19 | 6136026 | 6137228 | Promoter |
| chr19 | 6235951 | 6236328 | Intron |
| chr19 | 6277427 | 6279059 | Promoter |
| chr19 | 6295748 | 6298213 | Distal Intergenic |
| chr19 | 6305806 | 6306042 | Downstream |
| chr19 | 6306170 | 6306426 | Downstream |
| chr19 | 6306617 | 6308566 | 3' UTR |
| chr19 | 6334512 | 6335202 | Promoter |
| chr19 | 6363702 | 6364480 | Exon |
| chr19 | 6893951 | 6894609 | Intron |
| chr19 | 6894876 | 6895524 | Intron |
| chr19 | 6895572 | 6895771 | Intron |
| chr19 | 6914506 | 6916605 | Exon |
| chr19 | 6934858 | 6936032 | Exon |
| chr19 | 6951622 | 6951874 | Downstream |
| chr19 | 6984081 | 6985105 | 5' UTR |
| chr19 | 6995352 | 6995808 | Distal Intergenic |
| chr19 | 7016376 | 7017284 | Distal Intergenic |
| chr19 | 7048959 | 7049616 | Promoter |
| chr19 | 7049733 | 7051096 | Promoter |
| chr19 | 7054162 | 7054361 | Downstream |
| chr19 | 7055119 | 7055481 | Downstream |
| chr19 | 7066779 | 7066994 | Distal Intergenic |
| chr19 | 7067153 | 7067356 | Distal Intergenic |
| chr19 | 7092234 | 7093113 | Distal Intergenic |
| chr19 | 7097701 | 7098584 | Distal Intergenic |
| chr19 | 7113565 | 7114501 | 3' UTR |
| chr19 | 7115352 | 7115691 | 3' UTR |
| chr19 | 7128959 | 7130114 | Exon |
| chr19 | 7130427 | 7130950 | Intron |

| | | | |
|---|---|---|---|
| chr19 | 7165816 | 7166905 | Exon |
| chr19 | 7222199 | 7223881 | Intron |
| chr19 | 7233220 | 7235119 | Intron |
| chr19 | 7235205 | 7236273 | Intron |
| chr19 | 7279036 | 7279272 | Intron |
| chr19 | 7279355 | 7281234 | Intron |
| chr19 | 7291517 | 7292615 | Intron |
| chr19 | 7332148 | 7332443 | Distal Intergenic |
| chr19 | 7369494 | 7370384 | Distal Intergenic |
| chr19 | 7372063 | 7372655 | Distal Intergenic |
| chr19 | 7375859 | 7377679 | Distal Intergenic |
| chr19 | 7377810 | 7378036 | Distal Intergenic |
| chr19 | 7381905 | 7383664 | Distal Intergenic |
| chr19 | 7384087 | 7386949 | Distal Intergenic |
| chr19 | 7410126 | 7410766 | Distal Intergenic |
| chr19 | 7413958 | 7415066 | Distal Intergenic |
| chr19 | 7415298 | 7416356 | Distal Intergenic |
| chr19 | 7416418 | 7416949 | Distal Intergenic |
| chr19 | 7456561 | 7457504 | Intron |
| chr19 | 7533327 | 7534615 | Exon |
| chr19 | 7557939 | 7558245 | Distal Intergenic |
| chr19 | 7566393 | 7567620 | Intron |
| chr19 | 7567958 | 7568206 | Intron |
| chr19 | 7568550 | 7569153 | Exon |
| chr19 | 7569333 | 7569716 | Promoter |
| chr19 | 7573223 | 7573815 | 3' UTR |
| chr19 | 8773477 | 8774085 | Distal Intergenic |
| chr19 | 8776520 | 8777457 | Distal Intergenic |
| chr19 | 8777561 | 8777898 | Distal Intergenic |
| chr19 | 8779446 | 8782381 | Distal Intergenic |
| chr19 | 8783878 | 8788597 | Distal Intergenic |
| chr19 | 8796744 | 8801769 | Distal Intergenic |
| chr19 | 8801805 | 8802795 | Distal Intergenic |
| chr19 | 8815645 | 8815861 | Distal Intergenic |
| chr19 | 8815979 | 8816532 | Distal Intergenic |
| chr19 | 8830827 | 8831269 | Distal Intergenic |
| chr19 | 8831554 | 8832158 | Distal Intergenic |
| chr19 | 8848128 | 8849171 | Distal Intergenic |
| chr19 | 8872414 | 8872643 | Distal Intergenic |
| chr19 | 8876493 | 8876796 | Distal Intergenic |
| chr19 | 8877008 | 8878302 | Distal Intergenic |
| chr19 | 8893027 | 8893902 | Distal Intergenic |
| chr19 | 8894188 | 8894683 | Distal Intergenic |
| chr19 | 8945729 | 8945965 | Distal Intergenic |

| | | | |
|---|---|---|---|
| chr19 | 8946036 | 8947078 | Distal Intergenic |
| chr19 | 8961178 | 8961448 | Intron |
| chr19 | 8961484 | 8962434 | Exon |
| chr19 | 8962495 | 8964123 | Intron |
| chr19 | 8967514 | 8967871 | Intron |
| chr19 | 8972262 | 8972470 | Intron |
| chr19 | 8972543 | 8972822 | Intron |
| chr19 | 8973092 | 8973452 | Intron |
| chr19 | 8994451 | 8995214 | Exon |
| chr19 | 8995311 | 8995607 | Intron |
| chr19 | 9016151 | 9019328 | Exon |
| chr19 | 9027519 | 9029064 | Exon |
| chr19 | 9040626 | 9042963 | Intron |
| chr19 | 9043122 | 9043358 | Intron |
| chr19 | 9051055 | 9051762 | Intron |
| chr19 | 9053934 | 9054458 | Exon |
| chr19 | 9054507 | 9059144 | Exon |
| chr19 | 9062130 | 9062637 | Exon |
| chr19 | 9062972 | 9064451 | Exon |
| chr19 | 9064499 | 9065297 | Exon |
| chr19 | 9065398 | 9067874 | Exon |
| chr19 | 9068549 | 9071154 | Exon |
| chr19 | 9071224 | 9073000 | Exon |
| chr19 | 9107248 | 9108526 | Distal Intergenic |
| chr19 | 9133810 | 9135301 | Distal Intergenic |
| chr19 | 9135491 | 9135883 | Distal Intergenic |
| chr19 | 9140158 | 9141061 | Distal Intergenic |
| chr19 | 9147804 | 9149413 | Distal Intergenic |
| chr19 | 9180678 | 9182604 | Distal Intergenic |
| chr19 | 9972070 | 9974055 | Intron |
| chr19 | 9974108 | 9974937 | Intron |
| chr19 | 10012789 | 10013009 | Intron |
| chr19 | 10013151 | 10014034 | Intron |
| chr19 | 10028514 | 10029600 | Intron |
| chr19 | 10057208 | 10057538 | Distal Intergenic |
| chr19 | 10114555 | 10115231 | Exon |
| chr19 | 10115964 | 10118540 | Exon |
| chr19 | 10174866 | 10175105 | Intron |
| chr19 | 10175235 | 10176149 | Intron |
| chr19 | 10176652 | 10177112 | Intron |
| chr19 | 10187092 | 10187474 | Distal Intergenic |
| chr19 | 10187527 | 10188060 | Distal Intergenic |
| chr19 | 10211257 | 10213198 | Promoter |
| chr19 | 10232212 | 10232463 | Distal Intergenic |

| | | | |
|---|---|---|---|
| chr19 | 10237375 | 10238135 | Distal Intergenic |
| chr19 | 10269510 | 10269717 | Intron |
| chr19 | 10269797 | 10270101 | Intron |
| chr19 | 10270168 | 10272124 | Exon |
| chr19 | 10277786 | 10278885 | Promoter |
| chr19 | 10364050 | 10365004 | Intron |
| chr19 | 10599106 | 10599713 | Intron |
| chr19 | 10599766 | 10600163 | Exon |
| chr19 | 10635472 | 10637241 | Distal Intergenic |
| chr19 | 10676544 | 10680674 | Promoter |
| chr19 | 10731223 | 10732313 | Intron |
| chr19 | 10762945 | 10763196 | Exon |
| chr19 | 10763514 | 10764300 | Promoter |
| chr19 | 10764407 | 10764663 | Promoter |
| chr19 | 10908175 | 10908400 | Exon |
| chr19 | 10943762 | 10945220 | 3' UTR |
| chr19 | 10954586 | 10954834 | Distal Intergenic |
| chr19 | 10954884 | 10955240 | Distal Intergenic |
| chr19 | 10956333 | 10956661 | Distal Intergenic |
| chr19 | 11022948 | 11023854 | Promoter |
| chr19 | 11026246 | 11026931 | Intron |
| chr19 | 11395569 | 11395894 | Distal Intergenic |
| chr19 | 11844909 | 11845470 | Intron |
| chr19 | 11884476 | 11884675 | Intron |
| chr19 | 11884841 | 11885676 | Intron |
| chr19 | 11887170 | 11888247 | Intron |
| chr19 | 11892144 | 11892593 | Exon |
| chr19 | 11892970 | 11893499 | 3' UTR |
| chr19 | 12079396 | 12079626 | Intron |
| chr19 | 12080059 | 12080429 | Intron |
| chr19 | 12084651 | 12085006 | Intron |
| chr19 | 12547786 | 12548289 | Intron |
| chr19 | 12548552 | 12548758 | Intron |
| chr19 | 12549837 | 12550138 | Intron |
| chr19 | 12550186 | 12550505 | Intron |
| chr19 | 12565500 | 12566542 | Distal Intergenic |
| chr19 | 12649322 | 12649760 | Intron |
| chr19 | 12707587 | 12707840 | Intron |
| chr19 | 12832738 | 12832952 | Promoter |
| chr19 | 12833047 | 12833956 | Promoter |
| chr19 | 12836919 | 12838045 | Distal Intergenic |
| chr19 | 12840684 | 12840933 | Downstream |
| chr19 | 12868582 | 12869178 | 3' UTR |
| chr19 | 12869388 | 12869632 | Distal Intergenic |

| | | | |
|---|---|---|---|
| chr19 | 12870427 | 12870655 | Distal Intergenic |
| chr19 | 12870717 | 12871412 | Downstream |
| chr19 | 12871512 | 12871753 | Downstream |
| chr19 | 14272114 | 14272921 | Exon |
| chr19 | 15998158 | 15998677 | Intron |
| chr19 | 16015362 | 16016954 | Distal Intergenic |
| chr19 | 16020715 | 16020914 | Downstream |
| chr19 | 16021043 | 16021467 | Downstream |
| chr19 | 16090235 | 16091627 | Distal Intergenic |
| chr19 | 16150026 | 16150437 | Exon |
| chr19 | 16160909 | 16161262 | Distal Intergenic |
| chr19 | 16161439 | 16163123 | Distal Intergenic |
| chr19 | 16241070 | 16242884 | Intron |
| chr19 | 16244978 | 16245411 | Promoter |
| chr19 | 16256517 | 16256749 | Intron |
| chr19 | 16277034 | 16277887 | Intron |
| chr19 | 16278810 | 16280355 | Exon |
| chr19 | 16284528 | 16284769 | Promoter |
| chr19 | 16325177 | 16325861 | Intron |
| chr19 | 16344095 | 16346093 | 3' UTR |
| chr19 | 16346178 | 16346377 | Distal Intergenic |
| chr19 | 16346952 | 16348388 | Distal Intergenic |
| chr19 | 16360596 | 16361871 | Distal Intergenic |
| chr19 | 16400128 | 16400817 | Distal Intergenic |
| chr19 | 16446173 | 16446841 | Distal Intergenic |
| chr19 | 16460547 | 16461550 | Distal Intergenic |
| chr19 | 16483494 | 16485415 | Intron |
| chr19 | 16485573 | 16485779 | Intron |
| chr19 | 16485822 | 16486098 | Intron |
| chr19 | 16492879 | 16493118 | Intron |
| chr19 | 16587840 | 16589312 | Downstream |
| chr19 | 16718942 | 16720254 | Intron |
| chr19 | 17085867 | 17086081 | Exon |
| chr19 | 17237908 | 17238169 | Intron |
| chr19 | 17330651 | 17331924 | Distal Intergenic |
| chr19 | 17335047 | 17335461 | Distal Intergenic |
| chr19 | 17337731 | 17338340 | Promoter |
| chr19 | 17338395 | 17338596 | Intron |
| chr19 | 17435108 | 17435456 | Intron |
| chr19 | 17875077 | 17875483 | 5' UTR |
| chr19 | 18687906 | 18688189 | 3' UTR |
| chr19 | 18696936 | 18698812 | Promoter |
| chr19 | 18704481 | 18705831 | Exon |
| chr19 | 18706101 | 18706302 | Intron |

| | | | |
|---|---|---|---|
| chr19 | 18732750 | 18733953 | Distal Intergenic |
| chr19 | 18734292 | 18734546 | Distal Intergenic |
| chr19 | 18734760 | 18736760 | Distal Intergenic |
| chr19 | 19320765 | 19322303 | Promoter |
| chr19 | 20460527 | 20461413 | Intron |
| chr19 | 20462367 | 20465466 | Intron |
| chr19 | 20465653 | 20466022 | Intron |
| chr19 | 20467647 | 20469264 | Intron |
| chr19 | 20476964 | 20477879 | Intron |
| chr19 | 20480816 | 20481585 | Intron |
| chr19 | 21125773 | 21126753 | 3' UTR |
| chr19 | 21151998 | 21153163 | Distal Intergenic |
| chr19 | 21346768 | 21349624 | 5' UTR |
| chr19 | 21678764 | 21680700 | Exon |
| chr19 | 21718098 | 21718297 | Intron |
| chr19 | 21718527 | 21719140 | Exon |
| chr19 | 21720117 | 21721068 | 3' UTR |
| chr19 | 21726371 | 21727548 | Intron |
| chr19 | 21847945 | 21848268 | Distal Intergenic |
| chr19 | 21848310 | 21848720 | Distal Intergenic |
| chr19 | 21852868 | 21853474 | Distal Intergenic |
| chr19 | 21888568 | 21888832 | Distal Intergenic |
| chr19 | 21894328 | 21895051 | Distal Intergenic |
| chr19 | 21936897 | 21938660 | Intron |
| chr19 | 22109411 | 22110148 | Distal Intergenic |
| chr19 | 22110195 | 22111066 | Distal Intergenic |
| chr19 | 22990874 | 22991483 | Distal Intergenic |
| chr19 | 23254657 | 23255517 | Distal Intergenic |
| chr19 | 23403732 | 23404011 | Downstream |
| chr19 | 23404049 | 23404636 | Downstream |
| chr19 | 23429078 | 23429608 | Intron |
| chr19 | 23594801 | 23597141 | Intron |
| chr19 | 24150272 | 24151638 | Distal Intergenic |
| chr19 | 24162828 | 24163165 | Distal Intergenic |
| chr19 | 24164280 | 24165155 | Distal Intergenic |
| chr19 | 24240695 | 24241700 | Intron |
| chr19 | 24247798 | 24248443 | Exon |
| chr19 | 24248890 | 24250036 | Exon |
| chr19 | 24267097 | 24268122 | Intron |
| chr19 | 24271325 | 24272197 | Intron |
| chr19 | 24298672 | 24298972 | Intron |
| chr19 | 24299055 | 24299264 | Intron |
| chr19 | 24299749 | 24300324 | Intron |
| chr19 | 24336129 | 24336870 | Distal Intergenic |

| | | | |
|---|---|---|---|
| chr19 | 24551834 | 24552382 | Distal Intergenic |
| chr19 | 24806606 | 24807439 | Distal Intergenic |
| chr19 | 25069131 | 25069337 | Distal Intergenic |
| chr19 | 25134217 | 25135605 | Distal Intergenic |
| chr19 | 25241216 | 25242170 | Distal Intergenic |
| chr19 | 25242331 | 25242553 | Distal Intergenic |
| chr19 | 25245011 | 25248368 | Distal Intergenic |
| chr19 | 25249030 | 25249573 | Distal Intergenic |
| chr19 | 25259907 | 25266160 | Distal Intergenic |
| chr19 | 25338137 | 25339793 | Distal Intergenic |
| chr19 | 25340927 | 25343296 | Distal Intergenic |
| chr19 | 25364819 | 25365267 | Distal Intergenic |
| chr19 | 25415630 | 25417628 | Distal Intergenic |
| chr19 | 25418952 | 25419279 | Distal Intergenic |
| chr19 | 25426829 | 25427072 | Distal Intergenic |
| chr19 | 25427194 | 25427397 | Distal Intergenic |
| chr19 | 25431257 | 25432449 | Distal Intergenic |
| chr19 | 25446886 | 25449898 | Distal Intergenic |
| chr19 | 25473960 | 25475641 | Distal Intergenic |
| chr19 | 25500490 | 25502242 | Distal Intergenic |
| chr19 | 25502437 | 25503593 | Distal Intergenic |
| chr19 | 25535938 | 25536921 | Distal Intergenic |
| chr19 | 25654851 | 25655132 | Distal Intergenic |
| chr19 | 26165012 | 26165250 | Distal Intergenic |
| chr19 | 26165306 | 26166720 | Distal Intergenic |
| chr19 | 26214843 | 26215708 | Distal Intergenic |
| chr19 | 26218123 | 26218358 | Distal Intergenic |
| chr19 | 26287586 | 26288614 | Distal Intergenic |
| chr19 | 26512962 | 26513181 | Distal Intergenic |
| chr19 | 26717341 | 26718233 | Distal Intergenic |
| chr19 | 26807644 | 26808637 | Distal Intergenic |
| chr19 | 26820648 | 26823159 | Distal Intergenic |
| chr19 | 26920021 | 26920253 | Distal Intergenic |
| chr19 | 26965593 | 26966694 | Distal Intergenic |
| chr19 | 26967569 | 26967904 | Distal Intergenic |
| chr19 | 27012956 | 27013301 | Distal Intergenic |
| chr19 | 27013499 | 27014138 | Distal Intergenic |
| chr19 | 27089921 | 27091171 | Distal Intergenic |
| chr19 | 27114556 | 27114820 | Distal Intergenic |
| chr19 | 27136587 | 27136816 | Distal Intergenic |
| chr19 | 27136853 | 27137244 | Distal Intergenic |
| chr19 | 27139941 | 27140249 | Distal Intergenic |
| chr19 | 27200567 | 27201517 | Distal Intergenic |
| chr19 | 27208831 | 27209526 | Distal Intergenic |

| | | | |
|---|---|---|---|
| chr19 | 27251107 | 27252196 | Distal Intergenic |
| chr19 | 27253386 | 27256920 | Distal Intergenic |
| chr19 | 27257665 | 27258896 | Distal Intergenic |
| chr19 | 27300963 | 27302413 | Distal Intergenic |
| chr19 | 27372223 | 27373047 | Distal Intergenic |
| chr19 | 27425926 | 27426476 | Distal Intergenic |
| chr19 | 27503705 | 27503904 | Distal Intergenic |
| chr19 | 27504140 | 27504374 | Distal Intergenic |
| chr19 | 27504472 | 27504852 | Distal Intergenic |
| chr19 | 27978692 | 27978942 | Distal Intergenic |
| chr19 | 27978997 | 27979968 | Distal Intergenic |
| chr19 | 28085887 | 28086119 | Distal Intergenic |
| chr19 | 28086380 | 28086796 | Distal Intergenic |
| chr19 | 28273030 | 28274717 | Distal Intergenic |
| chr19 | 28277899 | 28278338 | Distal Intergenic |
| chr19 | 28278402 | 28278721 | Downstream |
| chr19 | 28320660 | 28321499 | Distal Intergenic |
| chr19 | 28348719 | 28349138 | Distal Intergenic |
| chr19 | 28370682 | 28370992 | Distal Intergenic |
| chr19 | 28371116 | 28371364 | Distal Intergenic |
| chr19 | 28373467 | 28375461 | Distal Intergenic |
| chr19 | 28463007 | 28464616 | Distal Intergenic |
| chr19 | 28549953 | 28550335 | Distal Intergenic |
| chr19 | 28592623 | 28593004 | Distal Intergenic |
| chr19 | 28593062 | 28593438 | Distal Intergenic |
| chr19 | 28593616 | 28593954 | Distal Intergenic |
| chr19 | 28632184 | 28633785 | Distal Intergenic |
| chr19 | 28690466 | 28690733 | Distal Intergenic |
| chr19 | 28697809 | 28699632 | Distal Intergenic |
| chr19 | 28877671 | 28878369 | Distal Intergenic |
| chr19 | 29064921 | 29065333 | Intron |
| chr19 | 29115904 | 29116267 | Intron |
| chr19 | 29116526 | 29117464 | Intron |
| chr19 | 29325367 | 29325827 | Distal Intergenic |
| chr19 | 29345324 | 29346599 | Distal Intergenic |
| chr19 | 29374753 | 29375583 | Distal Intergenic |
| chr19 | 29458585 | 29460198 | Exon |
| chr19 | 29596964 | 29597599 | Distal Intergenic |
| chr19 | 29598232 | 29598435 | Distal Intergenic |
| chr19 | 29678453 | 29679031 | Distal Intergenic |
| chr19 | 29715830 | 29716040 | Distal Intergenic |
| chr19 | 29967005 | 29967658 | Intron |
| chr19 | 29968087 | 29969537 | Intron |
| chr19 | 30105850 | 30106437 | 3' UTR |

| | | | |
|---|---|---|---|
| chr19 | 30647210 | 30648141 | Distal Intergenic |
| chr19 | 30925308 | 30925676 | Intron |
| chr19 | 30927635 | 30929043 | Intron |
| chr19 | 31193631 | 31193874 | Distal Intergenic |
| chr19 | 31924318 | 31926559 | Distal Intergenic |
| chr19 | 32140840 | 32141904 | Distal Intergenic |
| chr19 | 32142254 | 32142520 | Distal Intergenic |
| chr19 | 32142777 | 32143025 | Distal Intergenic |
| chr19 | 32143083 | 32145239 | Distal Intergenic |
| chr19 | 32145440 | 32145884 | Distal Intergenic |
| chr19 | 32162464 | 32163939 | Distal Intergenic |
| chr19 | 32181845 | 32182493 | Distal Intergenic |
| chr19 | 32182677 | 32182915 | Distal Intergenic |
| chr19 | 32247066 | 32247404 | Distal Intergenic |
| chr19 | 32254183 | 32254704 | Distal Intergenic |
| chr19 | 32256187 | 32256449 | Distal Intergenic |
| chr19 | 32256571 | 32256852 | Distal Intergenic |
| chr19 | 32264383 | 32265383 | Distal Intergenic |
| chr19 | 32271122 | 32273804 | Distal Intergenic |
| chr19 | 32284547 | 32286553 | Distal Intergenic |
| chr19 | 32338205 | 32339007 | Distal Intergenic |
| chr19 | 32341098 | 32341376 | Distal Intergenic |
| chr19 | 32341441 | 32342142 | Distal Intergenic |
| chr19 | 32346874 | 32349106 | Distal Intergenic |
| chr19 | 32349151 | 32349357 | Distal Intergenic |
| chr19 | 32359969 | 32360187 | Distal Intergenic |
| chr19 | 32365510 | 32367105 | Distal Intergenic |
| chr19 | 32378196 | 32381016 | Distal Intergenic |
| chr19 | 32390662 | 32391207 | Distal Intergenic |
| chr19 | 32391779 | 32391991 | Distal Intergenic |
| chr19 | 32420526 | 32421299 | Distal Intergenic |
| chr19 | 32463216 | 32463542 | Distal Intergenic |
| chr19 | 32468304 | 32468733 | Distal Intergenic |
| chr19 | 32591680 | 32592055 | Distal Intergenic |
| chr19 | 32592174 | 32593272 | Distal Intergenic |
| chr19 | 32598136 | 32598503 | Distal Intergenic |
| chr19 | 32598591 | 32598851 | Distal Intergenic |
| chr19 | 32747592 | 32747816 | Distal Intergenic |
| chr19 | 32747987 | 32748470 | Distal Intergenic |
| chr19 | 32748653 | 32748852 | Distal Intergenic |
| chr19 | 32750084 | 32750961 | Distal Intergenic |
| chr19 | 32786051 | 32786797 | Distal Intergenic |
| chr19 | 32786907 | 32787288 | Distal Intergenic |
| chr19 | 32830300 | 32830935 | Distal Intergenic |

| | | | |
|---|---|---|---|
| chr19 | 32831519 | 32831834 | Distal Intergenic |
| chr19 | 32832227 | 32832731 | Distal Intergenic |
| chr19 | 32833258 | 32835268 | Distal Intergenic |
| chr19 | 33048809 | 33049179 | Distal Intergenic |
| chr19 | 34096013 | 34096236 | Distal Intergenic |
| chr19 | 34096309 | 34097013 | Distal Intergenic |
| chr19 | 34116911 | 34117646 | Intron |
| chr19 | 34117764 | 34118102 | Intron |
| chr19 | 34125774 | 34127512 | Intron |
| chr19 | 34132401 | 34133093 | Intron |
| chr19 | 34173238 | 34173461 | Intron |
| chr19 | 34193001 | 34195806 | Intron |
| chr19 | 34238449 | 34241171 | Intron |
| chr19 | 34253861 | 34254080 | Intron |
| chr19 | 34437486 | 34438082 | Distal Intergenic |
| chr19 | 34483351 | 34483875 | Distal Intergenic |
| chr19 | 34484097 | 34484339 | Distal Intergenic |
| chr19 | 34484710 | 34485217 | Distal Intergenic |
| chr19 | 34485499 | 34485730 | Distal Intergenic |
| chr19 | 34498647 | 34498906 | Distal Intergenic |
| chr19 | 34516008 | 34517355 | Distal Intergenic |
| chr19 | 34519660 | 34520909 | Distal Intergenic |
| chr19 | 34522458 | 34522698 | Distal Intergenic |
| chr19 | 34522899 | 34524191 | Distal Intergenic |
| chr19 | 34524862 | 34525380 | Distal Intergenic |
| chr19 | 34525427 | 34525797 | Distal Intergenic |
| chr19 | 34529010 | 34530287 | Distal Intergenic |
| chr19 | 34760540 | 34760739 | Intron |
| chr19 | 34934454 | 34934929 | Exon |
| chr19 | 34952021 | 34952955 | Intron |
| chr19 | 34953382 | 34953604 | Intron |
| chr19 | 34953698 | 34954053 | Intron |
| chr19 | 34996543 | 34997472 | Distal Intergenic |
| chr19 | 35201331 | 35202711 | Distal Intergenic |
| chr19 | 36157622 | 36158375 | Promoter |
| chr19 | 36203842 | 36204089 | Promoter |
| chr19 | 36436830 | 36437087 | Distal Intergenic |
| chr19 | 36525317 | 36526118 | 3' UTR |
| chr19 | 36832237 | 36832772 | Exon |
| chr19 | 36907207 | 36908726 | Promoter |
| chr19 | 36908794 | 36910527 | Promoter |
| chr19 | 36990748 | 36993343 | Distal Intergenic |
| chr19 | 36993420 | 36993619 | Distal Intergenic |
| chr19 | 36999852 | 37000502 | Downstream |

| | | | |
|---|---|---|---|
| chr19 | 37000618 | 37001931 | 3' UTR |
| chr19 | 37002003 | 37002240 | 3' UTR |
| chr19 | 37027057 | 37027545 | Distal Intergenic |
| chr19 | 37174744 | 37178317 | Promoter |
| chr19 | 37179503 | 37180781 | Promoter |
| chr19 | 37281991 | 37283645 | Distal Intergenic |
| chr19 | 37338601 | 37339329 | Intron |
| chr19 | 37339452 | 37339675 | Intron |
| chr19 | 37407202 | 37408447 | Promoter |
| chr19 | 37449870 | 37451262 | Intron |
| chr19 | 38030044 | 38031892 | 3' UTR |
| chr19 | 38033182 | 38035707 | 3' UTR |
| chr19 | 38035786 | 38035986 | Distal Intergenic |
| chr19 | 38040191 | 38041114 | Promoter |
| chr19 | 38044496 | 38045155 | Intron |
| chr19 | 38059595 | 38061033 | Exon |
| chr19 | 38062720 | 38063172 | Intron |
| chr19 | 38071306 | 38071722 | Intron |
| chr19 | 38089043 | 38091574 | 5' UTR |
| chr19 | 38091929 | 38092227 | Exon |
| chr19 | 38097269 | 38098135 | Intron |
| chr19 | 38098801 | 38099909 | Intron |
| chr19 | 38099959 | 38101727 | Intron |
| chr19 | 38106799 | 38108301 | Distal Intergenic |
| chr19 | 38112145 | 38113735 | Distal Intergenic |
| chr19 | 38116638 | 38117881 | Distal Intergenic |
| chr19 | 38129470 | 38130063 | Intron |
| chr19 | 38130387 | 38130907 | Intron |
| chr19 | 39005584 | 39005934 | Exon |
| chr19 | 39005981 | 39006278 | Intron |
| chr19 | 40305689 | 40306597 | Distal Intergenic |
| chr19 | 40316251 | 40316454 | 3' UTR |
| chr19 | 40316619 | 40317361 | Exon |
| chr19 | 40321374 | 40321778 | Exon |
| chr19 | 40321883 | 40322720 | 5' UTR |
| chr19 | 40324265 | 40324479 | Promoter |
| chr19 | 40324691 | 40324890 | Promoter |
| chr19 | 40327651 | 40333892 | Promoter |
| chr19 | 40338335 | 40339340 | Distal Intergenic |
| chr19 | 40341236 | 40345823 | Distal Intergenic |
| chr19 | 40570003 | 40571046 | Distal Intergenic |
| chr19 | 40626534 | 40627352 | Distal Intergenic |
| chr19 | 40629128 | 40630296 | Distal Intergenic |
| chr19 | 40700694 | 40700980 | Intron |

| | | | |
|---|---|---|---|
| chr19 | 40956152 | 40956544 | Intron |
| chr19 | 40968969 | 40969836 | Intron |
| chr19 | 41050787 | 41051045 | Intron |
| chr19 | 41060823 | 41061681 | 3' UTR |
| chr19 | 41063974 | 41065324 | Intron |
| chr19 | 41067938 | 41071026 | Intron |
| chr19 | 41072740 | 41073142 | Intron |
| chr19 | 41337790 | 41338071 | Distal Intergenic |
| chr19 | 41338167 | 41338403 | Distal Intergenic |
| chr19 | 41338545 | 41338766 | Distal Intergenic |
| chr19 | 41569703 | 41569902 | Distal Intergenic |
| chr19 | 41569938 | 41570165 | Distal Intergenic |
| chr19 | 41821489 | 41822989 | Exon |
| chr19 | 41840451 | 41842017 | Intron |
| chr19 | 41847095 | 41847718 | Intron |
| chr19 | 41931849 | 41932054 | Exon |
| chr19 | 41970083 | 41970487 | Intron |
| chr19 | 41970563 | 41970891 | Intron |
| chr19 | 41980391 | 41982242 | Intron |
| chr19 | 41985840 | 41986291 | Intron |
| chr19 | 41986525 | 41988908 | Intron |
| chr19 | 42008048 | 42008575 | Distal Intergenic |
| chr19 | 42067492 | 42067832 | Intron |
| chr19 | 42068414 | 42068711 | Intron |
| chr19 | 42090532 | 42092770 | Exon |
| chr19 | 42092876 | 42093282 | 3' UTR |
| chr19 | 42096129 | 42097612 | Distal Intergenic |
| chr19 | 42154198 | 42154648 | Distal Intergenic |
| chr19 | 42173526 | 42177339 | 3' UTR |
| chr19 | 42194676 | 42195347 | Distal Intergenic |
| chr19 | 42199609 | 42206403 | Distal Intergenic |
| chr19 | 42209772 | 42210953 | Distal Intergenic |
| chr19 | 42222327 | 42223557 | Intron |
| chr19 | 42226058 | 42226331 | Intron |
| chr19 | 42241004 | 42242395 | Distal Intergenic |
| chr19 | 42324492 | 42325290 | Distal Intergenic |
| chr19 | 42661881 | 42662182 | Distal Intergenic |
| chr19 | 42662235 | 42663059 | Distal Intergenic |
| chr19 | 42663120 | 42663343 | Distal Intergenic |
| chr19 | 42672663 | 42673262 | Distal Intergenic |
| chr19 | 42687733 | 42687968 | Distal Intergenic |
| chr19 | 42706021 | 42707127 | 5' UTR |
| chr19 | 42734225 | 42735069 | 3' UTR |
| chr19 | 42838791 | 42839060 | Intron |

| | | | |
|---|---|---|---|
| chr19 | 42839101 | 42839654 | Exon |
| chr19 | 43573057 | 43574257 | Intron |
| chr19 | 43605221 | 43605550 | Intron |
| chr19 | 43625390 | 43626351 | Intron |
| chr19 | 43743395 | 43744036 | Intron |
| chr19 | 43750127 | 43750528 | Intron |
| chr19 | 43785459 | 43786237 | Distal Intergenic |
| chr19 | 43826857 | 43827469 | Distal Intergenic |
| chr19 | 43987408 | 43987641 | Intron |
| chr19 | 43987808 | 43988511 | Intron |
| chr19 | 43990775 | 43992051 | Exon |
| chr19 | 43992091 | 43992388 | Intron |
| chr19 | 43995758 | 43996350 | Intron |
| chr19 | 43996560 | 43996959 | Intron |
| chr19 | 44012616 | 44014187 | Exon |
| chr19 | 44043571 | 44043799 | Distal Intergenic |
| chr19 | 44044000 | 44044261 | Distal Intergenic |
| chr19 | 44126312 | 44127512 | 3' UTR |
| chr19 | 44170187 | 44170608 | Intron |
| chr19 | 44181918 | 44182276 | Distal Intergenic |
| chr19 | 44210118 | 44210317 | Distal Intergenic |
| chr19 | 44220121 | 44220639 | Promoter |
| chr19 | 44322458 | 44322677 | Intron |
| chr19 | 44322867 | 44323402 | Intron |
| chr19 | 44355339 | 44356068 | Distal Intergenic |
| chr19 | 44482444 | 44482849 | Distal Intergenic |
| chr19 | 44752370 | 44753073 | Intron |
| chr19 | 44753254 | 44755635 | Intron |
| chr19 | 44755855 | 44756143 | Intron |
| chr19 | 44783238 | 44783609 | Intron |
| chr19 | 44783667 | 44784413 | Intron |
| chr19 | 44829523 | 44829813 | Downstream |
| chr19 | 44856274 | 44857404 | Intron |
| chr19 | 44973588 | 44974256 | Distal Intergenic |
| chr19 | 45000779 | 45001737 | 5' UTR |
| chr19 | 45005684 | 45005888 | Distal Intergenic |
| chr19 | 45080962 | 45081566 | Distal Intergenic |
| chr19 | 45122065 | 45123145 | Intron |
| chr19 | 45129958 | 45130159 | Intron |
| chr19 | 45132214 | 45133987 | Intron |
| chr19 | 45148999 | 45149369 | Intron |
| chr19 | 45250779 | 45251228 | Promoter |
| chr19 | 45283775 | 45284949 | Exon |
| chr19 | 45726509 | 45726726 | Intron |

| | | | |
|---|---|---|---|
| chr19 | 45729783 | 45731181 | Exon |
| chr19 | 45733375 | 45735001 | Intron |
| chr19 | 45856762 | 45857531 | Intron |
| chr19 | 45857787 | 45858088 | Exon |
| chr19 | 45864899 | 45865316 | Promoter |
| chr19 | 45911100 | 45912372 | 3' UTR |
| chr19 | 45933759 | 45936467 | Intron |
| chr19 | 46072667 | 46073049 | Intron |
| chr19 | 46077068 | 46077712 | Intron |
| chr19 | 46099419 | 46099919 | Intron |
| chr19 | 46112060 | 46114062 | 3' UTR |
| chr19 | 46114215 | 46114414 | Intron |
| chr19 | 46131039 | 46131998 | Intron |
| chr19 | 46150658 | 46150904 | Distal Intergenic |
| chr19 | 46378836 | 46379042 | Distal Intergenic |
| chr19 | 46379241 | 46379749 | Distal Intergenic |
| chr19 | 46379872 | 46380114 | Distal Intergenic |
| chr19 | 46426366 | 46427308 | Distal Intergenic |
| chr19 | 46464751 | 46465726 | Intron |
| chr19 | 46465825 | 46466069 | Intron |
| chr19 | 46468629 | 46468882 | Intron |
| chr19 | 46469782 | 46470231 | Intron |
| chr19 | 46522918 | 46523204 | Promoter |
| chr19 | 46553487 | 46555156 | Distal Intergenic |
| chr19 | 46557657 | 46560219 | Distal Intergenic |
| chr19 | 46561806 | 46562022 | Exon |
| chr19 | 46562199 | 46563276 | Exon |
| chr19 | 46563421 | 46565046 | Exon |
| chr19 | 46574684 | 46575157 | Intron |
| chr19 | 46575508 | 46575721 | Intron |
| chr19 | 46577385 | 46577641 | Intron |
| chr19 | 46577901 | 46579011 | Intron |
| chr19 | 46579273 | 46579535 | Intron |
| chr19 | 46595895 | 46596526 | Distal Intergenic |
| chr19 | 46596674 | 46596941 | Distal Intergenic |
| chr19 | 46603663 | 46604455 | Distal Intergenic |
| chr19 | 46673586 | 46674722 | Distal Intergenic |
| chr19 | 46674802 | 46675047 | Distal Intergenic |
| chr19 | 46694708 | 46695022 | Intron |
| chr19 | 46695231 | 46695572 | Intron |
| chr19 | 46881264 | 46881619 | Intron |
| chr19 | 46967139 | 46967860 | Downstream |
| chr19 | 47035333 | 47035583 | Intron |
| chr19 | 47035970 | 47036453 | Intron |

| | | | |
|---|---|---|---|
| chr19 | 47059528 | 47059982 | Intron |
| chr19 | 47062362 | 47062586 | Intron |
| chr19 | 47141735 | 47142107 | Distal Intergenic |
| chr19 | 47164153 | 47164393 | Promoter |
| chr19 | 47164557 | 47165743 | Promoter |
| chr19 | 47211555 | 47213349 | Promoter |
| chr19 | 47251430 | 47251748 | Intron |
| chr19 | 47251818 | 47252486 | 5' UTR |
| chr19 | 47252635 | 47253332 | Intron |
| chr19 | 47293487 | 47295796 | Distal Intergenic |
| chr19 | 47295853 | 47296122 | Distal Intergenic |
| chr19 | 47306295 | 47308080 | Distal Intergenic |
| chr19 | 47323868 | 47324119 | Distal Intergenic |
| chr19 | 47324377 | 47324841 | Distal Intergenic |
| chr19 | 47355510 | 47357533 | Distal Intergenic |
| chr19 | 47357600 | 47357799 | Distal Intergenic |
| chr19 | 47385181 | 47385389 | Distal Intergenic |
| chr19 | 47385529 | 47391199 | Distal Intergenic |
| chr19 | 47402652 | 47406660 | Distal Intergenic |
| chr19 | 47420388 | 47420782 | Distal Intergenic |
| chr19 | 47517712 | 47518254 | Distal Intergenic |
| chr19 | 47565049 | 47566170 | Downstream |
| chr19 | 47566253 | 47569870 | 3' UTR |
| chr19 | 47574390 | 47576485 | Exon |
| chr19 | 47581827 | 47582111 | Intron |
| chr19 | 47582561 | 47585813 | Exon |
| chr19 | 47589726 | 47590172 | Exon |
| chr19 | 47745347 | 47746551 | Distal Intergenic |
| chr19 | 47762064 | 47762524 | Intron |
| chr19 | 47762750 | 47767115 | Exon |
| chr19 | 47771818 | 47775192 | 3' UTR |
| chr19 | 47781837 | 47782036 | Distal Intergenic |
| chr19 | 47782135 | 47783404 | Distal Intergenic |
| chr19 | 47784121 | 47785980 | Distal Intergenic |
| chr19 | 47800456 | 47800861 | Distal Intergenic |
| chr19 | 47803205 | 47805782 | Distal Intergenic |
| chr19 | 47909146 | 47910033 | 3' UTR |
| chr19 | 47918037 | 47919291 | Promoter |
| chr19 | 47926612 | 47927097 | Distal Intergenic |
| chr19 | 47928220 | 47931181 | Downstream |
| chr19 | 47931220 | 47933287 | 3' UTR |
| chr19 | 47943974 | 47945007 | Exon |
| chr19 | 47974607 | 47976372 | Promoter |
| chr19 | 47977034 | 47977901 | Downstream |

| | | | |
|---|---|---|---|
| chr19 | 47983096 | 47983295 | Exon |
| chr19 | 47990714 | 47991711 | 3' UTR |
| chr19 | 47991966 | 47992279 | Intron |
| chr19 | 47992420 | 47994530 | Exon |
| chr19 | 48002524 | 48004485 | Exon |
| chr19 | 48005615 | 48006865 | Exon |
| chr19 | 50996014 | 50996466 | Distal Intergenic |
| chr19 | 51019476 | 51019719 | Downstream |
| chr19 | 51041827 | 51042352 | Intron |
| chr19 | 52191837 | 52192354 | Distal Intergenic |
| chr19 | 52192648 | 52192847 | Distal Intergenic |
| chr19 | 52314095 | 52314328 | Intron |
| chr19 | 52707163 | 52707712 | Intron |
| chr19 | 53096085 | 53096461 | Distal Intergenic |
| chr19 | 53112646 | 53113085 | Downstream |
| chr19 | 53113208 | 53113707 | Downstream |
| chr19 | 53114613 | 53116121 | 3' UTR |
| chr19 | 53353972 | 53354312 | Intron |
| chr19 | 53354507 | 53354706 | Intron |
| chr19 | 53358068 | 53359306 | Intron |
| chr19 | 53464129 | 53465058 | Intron |
| chr19 | 53465146 | 53465816 | Promoter |
| chr19 | 53486396 | 53487295 | Intron |
| chr19 | 53495034 | 53495517 | Intron |
| chr19 | 53495610 | 53496493 | Promoter |
| chr19 | 53523941 | 53526125 | Distal Intergenic |
| chr19 | 53605970 | 53606395 | Promoter |
| chr19 | 53674643 | 53674842 | Intron |
| chr19 | 53806431 | 53807022 | Distal Intergenic |
| chr19 | 53969312 | 53970090 | Promoter |
| chr19 | 54017968 | 54018727 | Distal Intergenic |
| chr19 | 54019384 | 54019586 | Distal Intergenic |
| chr19 | 55057084 | 55057302 | Distal Intergenic |
| chr19 | 55057441 | 55057691 | Distal Intergenic |
| chr19 | 55295726 | 55296147 | 3' UTR |
| chr19 | 55306416 | 55307070 | Intron |
| chr19 | 55327807 | 55329993 | Promoter |
| chr19 | 55331685 | 55333098 | Exon |
| chr19 | 55341362 | 55341663 | Exon |
| chr19 | 55497376 | 55498879 | Exon |
| chr19 | 55537795 | 55538342 | Intron |
| chr19 | 55616365 | 55617138 | Intron |
| chr19 | 55617176 | 55617788 | Intron |
| chr19 | 55644632 | 55646391 | Exon |

| | | | |
|---|---|---|---|
| chr19 | 55786666 | 55787912 | Intron |
| chr19 | 55788110 | 55788534 | Intron |
| chr19 | 55807192 | 55807598 | Intron |
| chr19 | 55869375 | 55869645 | 3' UTR |
| chr19 | 55869851 | 55870500 | 3' UTR |
| chr19 | 56070083 | 56070982 | Distal Intergenic |
| chr19 | 56112113 | 56112353 | Promoter |
| chr19 | 56471795 | 56472407 | Intron |
| chr19 | 56525360 | 56526749 | Intron |
| chr19 | 56622242 | 56622618 | Intron |
| chr19 | 56622670 | 56623214 | Intron |
| chr19 | 56623321 | 56624207 | Intron |
| chr19 | 56645766 | 56646420 | Distal Intergenic |
| chr19 | 56653711 | 56654738 | Intron |
| chr19 | 56662493 | 56663522 | Intron |
| chr19 | 56674350 | 56677916 | Distal Intergenic |
| chr19 | 56778235 | 56778521 | Intron |
| chr19 | 56798328 | 56798543 | Intron |
| chr19 | 56885462 | 56885742 | Intron |
| chr19 | 56960352 | 56960610 | Intron |
| chr19 | 56980098 | 56980638 | Intron |
| chr19 | 56989182 | 56989943 | Promoter |
| chr19 | 57009002 | 57011854 | Distal Intergenic |
| chr19 | 57033060 | 57034546 | Intron |
| chr19 | 57051584 | 57053110 | Intron |
| chr19 | 57055072 | 57055437 | Intron |
| chr19 | 57064210 | 57066043 | 3' UTR |
| chr19 | 57067517 | 57070506 | 3' UTR |
| chr19 | 57076346 | 57077600 | Intron |
| chr19 | 57081137 | 57082651 | 5' UTR |
| chr19 | 57089096 | 57090492 | 3' UTR |
| chr19 | 57130046 | 57130699 | Intron |
| chr19 | 57181726 | 57181986 | Intron |
| chr19 | 57191656 | 57192736 | Distal Intergenic |
| chr19 | 57192790 | 57193532 | Distal Intergenic |
| chr19 | 57255546 | 57258308 | Intron |
| chr19 | 57272457 | 57273077 | Intron |
| chr19 | 57312675 | 57314226 | Intron |
| chr19 | 57315827 | 57316519 | Intron |
| chr19 | 57316605 | 57316935 | Intron |
| chr19 | 57347610 | 57347888 | Promoter |
| chr19 | 57347949 | 57348368 | Promoter |
| chr19 | 57426182 | 57427476 | Distal Intergenic |
| chr19 | 57436098 | 57436773 | Distal Intergenic |

| | | | |
|---|---|---|---|
| chr19 | 57438541 | 57439526 | Distal Intergenic |
| chr19 | 57680738 | 57681075 | Distal Intergenic |
| chr19 | 57801502 | 57801895 | Intron |
| chr19 | 58178959 | 58179726 | Promoter |
| chr19 | 58295113 | 58295887 | Intron |
| chr19 | 58596217 | 58598813 | 3' UTR |
| chr19 | 59145837 | 59146270 | Distal Intergenic |
| chr19 | 59420013 | 59420283 | Distal Intergenic |
| chr19 | 59474766 | 59475361 | Distal Intergenic |
| chr19 | 59532986 | 59533318 | Distal Intergenic |
| chr19 | 59661767 | 59662384 | Distal Intergenic |
| chr19 | 59919528 | 59923299 | Distal Intergenic |
| chr19 | 59933260 | 59933658 | Distal Intergenic |
| chr19 | 59937436 | 59938977 | Distal Intergenic |
| chr19 | 59954580 | 59956215 | Distal Intergenic |
| chr19 | 59956314 | 59956608 | Distal Intergenic |
| chr19 | 59962424 | 59963451 | Distal Intergenic |
| chr19 | 59963751 | 59964090 | Distal Intergenic |
| chr19 | 59964521 | 59965280 | Distal Intergenic |
| chr19 | 59973085 | 59974838 | Distal Intergenic |
| chr19 | 60018835 | 60019174 | Distal Intergenic |
| chr19 | 60116321 | 60118271 | Distal Intergenic |
| chr19 | 60655790 | 60656139 | Distal Intergenic |
| chr19 | 60657109 | 60657870 | Distal Intergenic |
| chr19 | 60692294 | 60692799 | Distal Intergenic |
| chr19 | 60692960 | 60693589 | Distal Intergenic |
| chr19 | 60862265 | 60862920 | Distal Intergenic |
| chr19 | 60864734 | 60865335 | Distal Intergenic |
| chr19 | 60865520 | 60866032 | Distal Intergenic |
| chr19 | 60866172 | 60866933 | Distal Intergenic |
| chr19 | 60950051 | 60950420 | Distal Intergenic |
| chr19 | 61031794 | 61032187 | Distal Intergenic |
| chr19 | 61283138 | 61283426 | Distal Intergenic |
| chr2 | 3058856 | 3059101 | Intron |
| chr2 | 3134944 | 3135421 | Distal Intergenic |
| chr2 | 3154728 | 3155112 | Distal Intergenic |
| chr2 | 3156014 | 3157050 | Distal Intergenic |
| chr2 | 3248204 | 3248536 | Intron |
| chr2 | 3248738 | 3249371 | Intron |
| chr2 | 3249522 | 3249879 | Intron |
| chr2 | 3370167 | 3370376 | Intron |
| chr2 | 3629473 | 3630634 | Distal Intergenic |
| chr2 | 3630689 | 3630938 | Distal Intergenic |
| chr2 | 3643806 | 3644444 | Intron |

| | | | |
|---|---|---|---|
| chr2 | 3665410 | 3667172 | Intron |
| chr2 | 3670909 | 3671294 | Intron |
| chr2 | 3677197 | 3677412 | Intron |
| chr2 | 3677460 | 3678324 | Intron |
| chr2 | 3702966 | 3703271 | Distal Intergenic |
| chr2 | 4018786 | 4019039 | Intron |
| chr2 | 4183402 | 4184071 | Distal Intergenic |
| chr2 | 4342084 | 4342550 | Distal Intergenic |
| chr2 | 4366451 | 4367343 | Distal Intergenic |
| chr2 | 4427670 | 4428059 | Distal Intergenic |
| chr2 | 4480090 | 4481002 | Distal Intergenic |
| chr2 | 4496276 | 4496587 | Distal Intergenic |
| chr2 | 4572777 | 4573147 | Distal Intergenic |
| chr2 | 4803281 | 4803634 | Distal Intergenic |
| chr2 | 4826310 | 4826552 | Distal Intergenic |
| chr2 | 5207119 | 5207638 | Distal Intergenic |
| chr2 | 5534965 | 5536075 | Distal Intergenic |
| chr2 | 5765801 | 5766752 | Distal Intergenic |
| chr2 | 5766888 | 5767087 | Distal Intergenic |
| chr2 | 5809744 | 5811267 | Distal Intergenic |
| chr2 | 5871272 | 5872129 | Distal Intergenic |
| chr2 | 5872442 | 5872777 | Distal Intergenic |
| chr2 | 5879322 | 5879533 | Distal Intergenic |
| chr2 | 5879620 | 5879831 | Distal Intergenic |
| chr2 | 5975685 | 5975897 | Distal Intergenic |
| chr2 | 6000689 | 6003397 | Distal Intergenic |
| chr2 | 6038558 | 6040244 | Distal Intergenic |
| chr2 | 6049955 | 6050678 | Distal Intergenic |
| chr2 | 6050873 | 6051113 | Distal Intergenic |
| chr2 | 6083315 | 6085571 | Intron |
| chr2 | 6089122 | 6089444 | Intron |
| chr2 | 6089498 | 6090114 | Intron |
| chr2 | 6134087 | 6134371 | Distal Intergenic |
| chr2 | 6269970 | 6271511 | Distal Intergenic |
| chr2 | 6273705 | 6274073 | Distal Intergenic |
| chr2 | 6279307 | 6280585 | Distal Intergenic |
| chr2 | 6280896 | 6281195 | Distal Intergenic |
| chr2 | 6287510 | 6287831 | Distal Intergenic |
| chr2 | 6299803 | 6300233 | Distal Intergenic |
| chr2 | 6300459 | 6301495 | Distal Intergenic |
| chr2 | 6468898 | 6471281 | Distal Intergenic |
| chr2 | 6490405 | 6491685 | Distal Intergenic |
| chr2 | 6503638 | 6504140 | Distal Intergenic |
| chr2 | 6514615 | 6516467 | Distal Intergenic |

| | | | |
|---|---|---|---|
| chr2 | 7456372 | 7456636 | Distal Intergenic |
| chr2 | 8197240 | 8197480 | Intron |
| chr2 | 8197521 | 8198761 | Intron |
| chr2 | 8702570 | 8703993 | Distal Intergenic |
| chr2 | 9062989 | 9063615 | Intron |
| chr2 | 9403315 | 9403760 | Intron |
| chr2 | 9406956 | 9407550 | Intron |
| chr2 | 9408739 | 9409273 | Intron |
| chr2 | 9654834 | 9655454 | Intron |
| chr2 | 9712902 | 9714595 | Distal Intergenic |
| chr2 | 9716985 | 9717297 | Distal Intergenic |
| chr2 | 9724743 | 9724992 | 3' UTR |
| chr2 | 9725059 | 9725269 | 3' UTR |
| chr2 | 9757028 | 9757308 | Intron |
| chr2 | 9788618 | 9789063 | Distal Intergenic |
| chr2 | 9800351 | 9801214 | Distal Intergenic |
| chr2 | 9877476 | 9878066 | Distal Intergenic |
| chr2 | 10001532 | 10001735 | Intron |
| chr2 | 10001838 | 10002056 | Intron |
| chr2 | 10008665 | 10009524 | Intron |
| chr2 | 10502669 | 10502930 | Intron |
| chr2 | 11260330 | 11261200 | Exon |
| chr2 | 11298965 | 11299196 | Intron |
| chr2 | 11316745 | 11316958 | Intron |
| chr2 | 11339425 | 11339637 | Intron |
| chr2 | 11351003 | 11351408 | Intron |
| chr2 | 11379510 | 11380822 | Intron |
| chr2 | 11385965 | 11388544 | Intron |
| chr2 | 11524109 | 11524786 | Distal Intergenic |
| chr2 | 11545332 | 11545695 | Distal Intergenic |
| chr2 | 11545747 | 11545978 | Distal Intergenic |
| chr2 | 11597790 | 11598253 | Intron |
| chr2 | 11626689 | 11629528 | Distal Intergenic |
| chr2 | 11629588 | 11631583 | Distal Intergenic |
| chr2 | 11975056 | 11975502 | Downstream |
| chr2 | 11998981 | 11999180 | Distal Intergenic |
| chr2 | 12333821 | 12334271 | Intron |
| chr2 | 12340144 | 12340720 | Intron |
| chr2 | 12340854 | 12341064 | Intron |
| chr2 | 12686317 | 12686546 | Intron |
| chr2 | 12687816 | 12688087 | Intron |
| chr2 | 12845838 | 12846220 | Distal Intergenic |
| chr2 | 13061337 | 13063283 | Distal Intergenic |
| chr2 | 13292999 | 13293746 | Distal Intergenic |

| | | | |
|---|---|---|---|
| chr2 | 13294017 | 13294569 | Distal Intergenic |
| chr2 | 13326261 | 13327748 | Distal Intergenic |
| chr2 | 13328839 | 13330019 | Distal Intergenic |
| chr2 | 13330056 | 13330261 | Distal Intergenic |
| chr2 | 13330928 | 13331976 | Distal Intergenic |
| chr2 | 13332038 | 13332499 | Distal Intergenic |
| chr2 | 13372404 | 13372914 | Distal Intergenic |
| chr2 | 13478932 | 13479421 | Distal Intergenic |
| chr2 | 13994760 | 13997261 | Distal Intergenic |
| chr2 | 14970485 | 14971237 | Distal Intergenic |
| chr2 | 14976215 | 14976486 | Distal Intergenic |
| chr2 | 14976593 | 14976799 | Distal Intergenic |
| chr2 | 14976916 | 14977726 | Distal Intergenic |
| chr2 | 14977808 | 14978063 | Distal Intergenic |
| chr2 | 14981910 | 14982748 | Distal Intergenic |
| chr2 | 15827810 | 15828030 | Distal Intergenic |
| chr2 | 16090071 | 16090832 | Distal Intergenic |
| chr2 | 16135729 | 16136093 | Distal Intergenic |
| chr2 | 16327528 | 16329638 | Distal Intergenic |
| chr2 | 16367280 | 16367673 | Distal Intergenic |
| chr2 | 17570188 | 17570395 | Distal Intergenic |
| chr2 | 17663075 | 17664643 | Distal Intergenic |
| chr2 | 17766931 | 17767437 | Intron |
| chr2 | 17949289 | 17950553 | Exon |
| chr2 | 17977825 | 17978194 | Intron |
| chr2 | 17978567 | 17979554 | Intron |
| chr2 | 17985335 | 17986082 | Distal Intergenic |
| chr2 | 17986202 | 17986861 | Distal Intergenic |
| chr2 | 17987144 | 17987648 | Distal Intergenic |
| chr2 | 18038132 | 18038363 | Distal Intergenic |
| chr2 | 18056225 | 18056996 | Distal Intergenic |
| chr2 | 18135663 | 18136000 | Distal Intergenic |
| chr2 | 18313720 | 18313992 | Distal Intergenic |
| chr2 | 18314034 | 18314432 | Distal Intergenic |
| chr2 | 18332978 | 18334235 | Distal Intergenic |
| chr2 | 18593563 | 18593826 | Distal Intergenic |
| chr2 | 18594096 | 18595108 | Distal Intergenic |
| chr2 | 18615689 | 18616132 | Distal Intergenic |
| chr2 | 18687380 | 18689278 | Distal Intergenic |
| chr2 | 18849718 | 18850377 | Distal Intergenic |
| chr2 | 19328406 | 19329332 | Distal Intergenic |
| chr2 | 19462326 | 19462860 | Distal Intergenic |
| chr2 | 19500263 | 19502755 | Distal Intergenic |
| chr2 | 19527274 | 19527794 | Distal Intergenic |

| | | | |
|---|---|---|---|
| chr2 | 19527830 | 19528419 | Distal Intergenic |
| chr2 | 19579584 | 19579816 | Distal Intergenic |
| chr2 | 19580398 | 19580689 | Distal Intergenic |
| chr2 | 19580740 | 19581093 | Distal Intergenic |
| chr2 | 19604797 | 19605155 | Distal Intergenic |
| chr2 | 19606987 | 19607321 | Distal Intergenic |
| chr2 | 19624655 | 19624985 | Distal Intergenic |
| chr2 | 20440979 | 20443628 | Distal Intergenic |
| chr2 | 20468206 | 20468427 | Intron |
| chr2 | 20468476 | 20472340 | Intron |
| chr2 | 20476788 | 20478042 | Promoter |
| chr2 | 20480929 | 20482538 | Intron |
| chr2 | 20482798 | 20484489 | Exon |
| chr2 | 20484657 | 20485072 | Intron |
| chr2 | 20518349 | 20519843 | 5' UTR |
| chr2 | 20519991 | 20521549 | Intron |
| chr2 | 20521662 | 20522765 | Intron |
| chr2 | 20522870 | 20523128 | Intron |
| chr2 | 20538314 | 20538528 | Intron |
| chr2 | 20540135 | 20542466 | Intron |
| chr2 | 20563195 | 20563436 | Distal Intergenic |
| chr2 | 20563599 | 20565402 | Distal Intergenic |
| chr2 | 20573621 | 20577209 | Distal Intergenic |
| chr2 | 20577549 | 20577917 | Distal Intergenic |
| chr2 | 20578087 | 20578350 | Distal Intergenic |
| chr2 | 20579856 | 20580083 | Distal Intergenic |
| chr2 | 20581748 | 20585520 | Distal Intergenic |
| chr2 | 20585601 | 20585961 | Distal Intergenic |
| chr2 | 20586113 | 20586816 | Distal Intergenic |
| chr2 | 20586987 | 20588605 | Distal Intergenic |
| chr2 | 20612229 | 20613230 | Distal Intergenic |
| chr2 | 20648615 | 20650153 | 3' UTR |
| chr2 | 20657315 | 20659316 | Distal Intergenic |
| chr2 | 20673742 | 20676140 | Distal Intergenic |
| chr2 | 20794720 | 20795968 | Distal Intergenic |
| chr2 | 20865073 | 20865537 | Promoter |
| chr2 | 20865573 | 20866104 | Promoter |
| chr2 | 20870332 | 20870546 | Exon |
| chr2 | 20870830 | 20871173 | Exon |
| chr2 | 20888517 | 20891327 | Intron |
| chr2 | 20891364 | 20891587 | Intron |
| chr2 | 20951026 | 20953195 | Intron |
| chr2 | 20958902 | 20959221 | Intron |
| chr2 | 20975223 | 20976533 | Intron |

| | | | |
|---|---|---|---|
| chr2 | 20986251 | 20987259 | Intron |
| chr2 | 21103958 | 21105048 | Distal Intergenic |
| chr2 | 21155841 | 21156275 | Distal Intergenic |
| chr2 | 21156331 | 21159317 | Distal Intergenic |
| chr2 | 21161496 | 21161808 | Distal Intergenic |
| chr2 | 21220966 | 21221487 | Downstream |
| chr2 | 21444463 | 21444712 | Distal Intergenic |
| chr2 | 21864273 | 21865758 | Distal Intergenic |
| chr2 | 22579337 | 22580110 | Distal Intergenic |
| chr2 | 22861477 | 22862448 | Distal Intergenic |
| chr2 | 22862607 | 22862870 | Distal Intergenic |
| chr2 | 22863495 | 22863743 | Distal Intergenic |
| chr2 | 22870367 | 22870889 | Distal Intergenic |
| chr2 | 22876991 | 22878134 | Distal Intergenic |
| chr2 | 22878239 | 22878463 | Distal Intergenic |
| chr2 | 22889914 | 22892469 | Distal Intergenic |
| chr2 | 22894222 | 22895818 | Distal Intergenic |
| chr2 | 22923001 | 22924761 | Distal Intergenic |
| chr2 | 22924931 | 22925191 | Distal Intergenic |
| chr2 | 23010636 | 23011489 | Distal Intergenic |
| chr2 | 23011583 | 23012433 | Distal Intergenic |
| chr2 | 23426950 | 23427179 | Distal Intergenic |
| chr2 | 23427369 | 23427568 | Distal Intergenic |
| chr2 | 23929440 | 23930513 | 3' UTR |
| chr2 | 24041541 | 24042219 | Intron |
| chr2 | 24070280 | 24070479 | Intron |
| chr2 | 24123015 | 24123214 | Intron |
| chr2 | 24146532 | 24146732 | Intron |
| chr2 | 24149626 | 24151416 | Promoter |
| chr2 | 24172772 | 24172971 | Intron |
| chr2 | 24191856 | 24192491 | Exon |
| chr2 | 24197160 | 24197365 | Intron |
| chr2 | 24223261 | 24223941 | 3' UTR |
| chr2 | 24240186 | 24243074 | Exon |
| chr2 | 24262908 | 24264354 | Intron |
| chr2 | 24294194 | 24294432 | Intron |
| chr2 | 24294482 | 24296277 | Intron |
| chr2 | 24308383 | 24309470 | Promoter |
| chr2 | 24418099 | 24419406 | Distal Intergenic |
| chr2 | 24761546 | 24762013 | Distal Intergenic |
| chr2 | 24762169 | 24762694 | Distal Intergenic |
| chr2 | 24774181 | 24774386 | Distal Intergenic |
| chr2 | 24783260 | 24784520 | Distal Intergenic |
| chr2 | 24784573 | 24785176 | Distal Intergenic |

| | | | |
|---|---|---|---|
| chr2 | 24830702 | 24830938 | Intron |
| chr2 | 25028108 | 25028422 | Intron |
| chr2 | 25031807 | 25032740 | Intron |
| chr2 | 25035090 | 25036008 | Intron |
| chr2 | 25051214 | 25053120 | Intron |
| chr2 | 25053243 | 25053449 | Intron |
| chr2 | 25066450 | 25066906 | Intron |
| chr2 | 25079189 | 25080083 | Intron |
| chr2 | 25080916 | 25081444 | Intron |
| chr2 | 25082972 | 25083962 | Intron |
| chr2 | 25092883 | 25094309 | Intron |
| chr2 | 25095411 | 25095660 | Exon |
| chr2 | 25110360 | 25111185 | Intron |
| chr2 | 25111636 | 25111854 | Intron |
| chr2 | 25117764 | 25117963 | Intron |
| chr2 | 25121132 | 25121910 | Intron |
| chr2 | 25141278 | 25143511 | Promoter |
| chr2 | 25187782 | 25188074 | Intron |
| chr2 | 25316556 | 25316838 | Intron |
| chr2 | 25356321 | 25356897 | Intron |
| chr2 | 25357938 | 25358172 | Intron |
| chr2 | 25358316 | 25358520 | Exon |
| chr2 | 25389298 | 25390744 | Promoter |
| chr2 | 25401861 | 25402077 | Distal Intergenic |
| chr2 | 25430022 | 25430404 | Distal Intergenic |
| chr2 | 25430499 | 25430746 | Distal Intergenic |
| chr2 | 25463363 | 25463670 | Exon |
| chr2 | 25463814 | 25464074 | Intron |
| chr2 | 25512716 | 25513325 | Intron |
| chr2 | 25516941 | 25517495 | Intron |
| chr2 | 25831083 | 25831334 | Intron |
| chr2 | 25841757 | 25845080 | Intron |
| chr2 | 25974454 | 25974776 | Intron |
| chr2 | 25988641 | 25989470 | Intron |
| chr2 | 25999661 | 26000380 | Intron |
| chr2 | 26085665 | 26085864 | Intron |
| chr2 | 26086423 | 26087883 | Intron |
| chr2 | 26090949 | 26091224 | Intron |
| chr2 | 26107987 | 26108282 | Distal Intergenic |
| chr2 | 26108531 | 26108861 | Distal Intergenic |
| chr2 | 26167517 | 26167975 | Intron |
| chr2 | 26206419 | 26206885 | Promoter |
| chr2 | 26206939 | 26207575 | Distal Intergenic |
| chr2 | 26244964 | 26245366 | Distal Intergenic |

| | | | |
|---|---|---|---|
| chr2 | 26300278 | 26300769 | Intron |
| chr2 | 26301467 | 26301872 | Intron |
| chr2 | 26302119 | 26302323 | Intron |
| chr2 | 26356208 | 26359619 | 3' UTR |
| chr2 | 26361466 | 26364050 | Distal Intergenic |
| chr2 | 26387108 | 26387510 | Distal Intergenic |
| chr2 | 26387557 | 26387757 | Distal Intergenic |
| chr2 | 26403569 | 26404068 | Promoter |
| chr2 | 26406211 | 26412987 | 3' UTR |
| chr2 | 26494244 | 26496520 | Exon |
| chr2 | 26498944 | 26499388 | Intron |
| chr2 | 26757702 | 26759347 | Intron |
| chr2 | 26765293 | 26765803 | Intron |
| chr2 | 26765965 | 26766971 | Intron |
| chr2 | 26769861 | 26770446 | Intron |
| chr2 | 26770723 | 26770929 | Intron |
| chr2 | 26771042 | 26772538 | Intron |
| chr2 | 26788285 | 26788927 | Intron |
| chr2 | 26819383 | 26820497 | Intron |
| chr2 | 26865816 | 26866038 | Distal Intergenic |
| chr2 | 27173715 | 27174285 | Distal Intergenic |
| chr2 | 27182467 | 27182666 | Distal Intergenic |
| chr2 | 27220311 | 27220610 | Intron |
| chr2 | 27272007 | 27272369 | Distal Intergenic |
| chr2 | 27319253 | 27319504 | Intron |
| chr2 | 27319663 | 27319987 | Exon |
| chr2 | 27360259 | 27362136 | Promoter |
| chr2 | 27371109 | 27371324 | Promoter |
| chr2 | 27408417 | 27409389 | Distal Intergenic |
| chr2 | 27409601 | 27409914 | Distal Intergenic |
| chr2 | 27410012 | 27410231 | Distal Intergenic |
| chr2 | 27486204 | 27487245 | Promoter |
| chr2 | 27527541 | 27527777 | Intron |
| chr2 | 27531868 | 27532195 | Promoter |
| chr2 | 27533355 | 27533640 | Intron |
| chr2 | 27534550 | 27534796 | Exon |
| chr2 | 27535183 | 27536614 | Exon |
| chr2 | 27557262 | 27560482 | Promoter |
| chr2 | 27569098 | 27571960 | Intron |
| chr2 | 27572121 | 27572384 | Intron |
| chr2 | 27572827 | 27575453 | 5' UTR |
| chr2 | 27575498 | 27578117 | Intron |
| chr2 | 27590085 | 27590486 | 3' UTR |
| chr2 | 27595456 | 27595856 | 5' UTR |

| | | | |
|---|---|---|---|
| chr2 | 27598792 | 27598991 | Exon |
| chr2 | 27599083 | 27600795 | 3' UTR |
| chr2 | 27600903 | 27602081 | Exon |
| chr2 | 27613601 | 27615155 | Intron |
| chr2 | 28372974 | 28373461 | Intron |
| chr2 | 28379178 | 28379414 | Intron |
| chr2 | 28379453 | 28380078 | Intron |
| chr2 | 28380496 | 28380695 | Intron |
| chr2 | 28380757 | 28381884 | Intron |
| chr2 | 28384837 | 28385746 | Intron |
| chr2 | 28438329 | 28438762 | Intron |
| chr2 | 28496517 | 28496727 | Intron |
| chr2 | 28695674 | 28696060 | Distal Intergenic |
| chr2 | 28696176 | 28696408 | Distal Intergenic |
| chr2 | 28696533 | 28697064 | Distal Intergenic |
| chr2 | 28915685 | 28915979 | Distal Intergenic |
| chr2 | 28916042 | 28916257 | Distal Intergenic |
| chr2 | 28920536 | 28920997 | Distal Intergenic |
| chr2 | 28921173 | 28922129 | Distal Intergenic |
| chr2 | 28944087 | 28945379 | Distal Intergenic |
| chr2 | 28956165 | 28956908 | Distal Intergenic |
| chr2 | 29090809 | 29091310 | Intron |
| chr2 | 29091414 | 29091644 | Intron |
| chr2 | 29278011 | 29278843 | Distal Intergenic |
| chr2 | 29381582 | 29382308 | Intron |
| chr2 | 29386475 | 29386677 | Intron |
| chr2 | 29386962 | 29387235 | Intron |
| chr2 | 29468966 | 29469222 | Intron |
| chr2 | 29474228 | 29475128 | Intron |
| chr2 | 29475529 | 29475875 | Intron |
| chr2 | 29476040 | 29476617 | Intron |
| chr2 | 29480019 | 29480369 | Intron |
| chr2 | 29480447 | 29480686 | Intron |
| chr2 | 29483533 | 29483980 | Intron |
| chr2 | 29484516 | 29485097 | Intron |
| chr2 | 29485160 | 29486863 | Intron |
| chr2 | 29487644 | 29489043 | Intron |
| chr2 | 29526169 | 29526729 | Intron |
| chr2 | 29546515 | 29548054 | Intron |
| chr2 | 29550397 | 29552630 | Exon |
| chr2 | 29561249 | 29561628 | Intron |
| chr2 | 29566299 | 29569348 | Intron |
| chr2 | 29571681 | 29573083 | Intron |
| chr2 | 29584712 | 29585725 | Intron |

| | | | |
|---|---|---|---|
| chr2 | 29627107 | 29627899 | Intron |
| chr2 | 29682136 | 29682525 | Intron |
| chr2 | 29682636 | 29683170 | Intron |
| chr2 | 29745719 | 29746137 | Intron |
| chr2 | 29816136 | 29817685 | Intron |
| chr2 | 29819702 | 29820821 | Intron |
| chr2 | 29822077 | 29824868 | Intron |
| chr2 | 29827462 | 29828831 | Intron |
| chr2 | 29917625 | 29919707 | Exon |
| chr2 | 29921318 | 29921537 | Intron |
| chr2 | 29922109 | 29922502 | Intron |
| chr2 | 29922544 | 29923450 | Intron |
| chr2 | 29988860 | 29989068 | Intron |
| chr2 | 30092280 | 30093135 | Intron |
| chr2 | 30093743 | 30094015 | Intron |
| chr2 | 30105926 | 30108019 | Intron |
| chr2 | 30113094 | 30115466 | Intron |
| chr2 | 30121223 | 30122312 | Intron |
| chr2 | 30142101 | 30142360 | Intron |
| chr2 | 30142636 | 30142941 | Exon |
| chr2 | 30178307 | 30179938 | Distal Intergenic |
| chr2 | 30180463 | 30181738 | Distal Intergenic |
| chr2 | 30206189 | 30207151 | Distal Intergenic |
| chr2 | 30240975 | 30241694 | Distal Intergenic |
| chr2 | 30295536 | 30298799 | Distal Intergenic |
| chr2 | 30298867 | 30299109 | Distal Intergenic |
| chr2 | 30299369 | 30299573 | Distal Intergenic |
| chr2 | 30300112 | 30301840 | Distal Intergenic |
| chr2 | 30318859 | 30320015 | Distal Intergenic |
| chr2 | 30325579 | 30326758 | Distal Intergenic |
| chr2 | 30326891 | 30327564 | Distal Intergenic |
| chr2 | 30327627 | 30328095 | Distal Intergenic |
| chr2 | 30403511 | 30404095 | Distal Intergenic |
| chr2 | 30555523 | 30555907 | Distal Intergenic |
| chr2 | 30562816 | 30563126 | Distal Intergenic |
| chr2 | 30563162 | 30563632 | Distal Intergenic |
| chr2 | 30563755 | 30565529 | Distal Intergenic |
| chr2 | 30567036 | 30567272 | Distal Intergenic |
| chr2 | 30567836 | 30568956 | Distal Intergenic |
| chr2 | 30574009 | 30574970 | Exon |
| chr2 | 30575103 | 30575398 | Exon |
| chr2 | 30575520 | 30575936 | Distal Intergenic |
| chr2 | 30663508 | 30663844 | Distal Intergenic |
| chr2 | 30753033 | 30753294 | Intron |

| | | | |
|---|---|---|---|
| chr2 | 30753369 | 30753576 | Intron |
| chr2 | 30754445 | 30755506 | Intron |
| chr2 | 30757397 | 30758504 | Intron |
| chr2 | 30773055 | 30774575 | Intron |
| chr2 | 30774669 | 30776322 | 5' UTR |
| chr2 | 30783909 | 30786223 | 5' UTR |
| chr2 | 30788336 | 30789696 | Intron |
| chr2 | 30898030 | 30899190 | Distal Intergenic |
| chr2 | 30978870 | 30979547 | Intron |
| chr2 | 31007455 | 31007695 | Intron |
| chr2 | 31007812 | 31009104 | Intron |
| chr2 | 31017931 | 31021558 | Intron |
| chr2 | 31023115 | 31024753 | Intron |
| chr2 | 31027455 | 31027870 | Intron |
| chr2 | 31118970 | 31120595 | Distal Intergenic |
| chr2 | 31297789 | 31298899 | Intron |
| chr2 | 31309059 | 31310810 | Intron |
| chr2 | 31325894 | 31326699 | Intron |
| chr2 | 31379784 | 31380007 | Distal Intergenic |
| chr2 | 31397456 | 31397719 | 3' UTR |
| chr2 | 31427980 | 31428241 | 5' UTR |
| chr2 | 31526286 | 31526512 | Distal Intergenic |
| chr2 | 31543818 | 31544483 | Distal Intergenic |
| chr2 | 31615217 | 31616781 | Intron |
| chr2 | 31625007 | 31627163 | Exon |
| chr2 | 31790258 | 31790672 | Intron |
| chr2 | 31830070 | 31830377 | Distal Intergenic |
| chr2 | 31862310 | 31863996 | Distal Intergenic |
| chr2 | 31993220 | 31993432 | Distal Intergenic |
| chr2 | 32007149 | 32008367 | Distal Intergenic |
| chr2 | 32008414 | 32008861 | Distal Intergenic |
| chr2 | 32016055 | 32016651 | Distal Intergenic |
| chr2 | 32020200 | 32021016 | Distal Intergenic |
| chr2 | 32022519 | 32023550 | Distal Intergenic |
| chr2 | 32029137 | 32031494 | Distal Intergenic |
| chr2 | 32037139 | 32037470 | Distal Intergenic |
| chr2 | 32037515 | 32039503 | Distal Intergenic |
| chr2 | 32091483 | 32091996 | Downstream |
| chr2 | 32092116 | 32092433 | Downstream |
| chr2 | 32180759 | 32180958 | Intron |
| chr2 | 32197653 | 32198389 | Intron |
| chr2 | 32198519 | 32198899 | Intron |
| chr2 | 32218730 | 32219267 | Intron |
| chr2 | 32278591 | 32280734 | Distal Intergenic |

| | | | |
|---|---|---|---|
| chr2 | 32283309 | 32284374 | Distal Intergenic |
| chr2 | 32292498 | 32292761 | Intron |
| chr2 | 32326259 | 32326835 | Intron |
| chr2 | 32426403 | 32426960 | Intron |
| chr2 | 32483863 | 32485321 | Intron |
| chr2 | 32535330 | 32538031 | Distal Intergenic |
| chr2 | 32541187 | 32542170 | Distal Intergenic |
| chr2 | 32567431 | 32568121 | Distal Intergenic |
| chr2 | 32569758 | 32569966 | Distal Intergenic |
| chr2 | 32570436 | 32571192 | Distal Intergenic |
| chr2 | 32576085 | 32576294 | Distal Intergenic |
| chr2 | 32630839 | 32631097 | Intron |
| chr2 | 32631369 | 32631731 | Exon |
| chr2 | 32690160 | 32690555 | Exon |
| chr2 | 32722590 | 32722981 | Intron |
| chr2 | 32723026 | 32723688 | Intron |
| chr2 | 32725167 | 32726744 | Exon |
| chr2 | 32730519 | 32736016 | Exon |
| chr2 | 32736167 | 32738472 | Exon |
| chr2 | 32738557 | 32738955 | Intron |
| chr2 | 32750025 | 32750583 | Exon |
| chr2 | 32751390 | 32753297 | Intron |
| chr2 | 32753512 | 32753832 | Intron |
| chr2 | 32754402 | 32754654 | Intron |
| chr2 | 32757638 | 32758564 | Promoter |
| chr2 | 32772944 | 32773168 | Exon |
| chr2 | 32817259 | 32818615 | Intron |
| chr2 | 32818692 | 32820598 | Exon |
| chr2 | 32821140 | 32821348 | Intron |
| chr2 | 32821387 | 32823088 | Exon |
| chr2 | 32883235 | 32883521 | Intron |
| chr2 | 33052161 | 33052608 | Intron |
| chr2 | 33060177 | 33062703 | Intron |
| chr2 | 33071582 | 33072877 | Intron |
| chr2 | 33074787 | 33077728 | Intron |
| chr2 | 33148236 | 33149121 | Intron |
| chr2 | 33158757 | 33159131 | Intron |
| chr2 | 33159328 | 33159547 | Intron |
| chr2 | 33169688 | 33171209 | Exon |
| chr2 | 33173378 | 33173589 | Intron |
| chr2 | 33174451 | 33176980 | Intron |
| chr2 | 33195367 | 33196488 | Intron |
| chr2 | 33197596 | 33198701 | Intron |
| chr2 | 33203480 | 33204084 | Intron |

| | | | |
|---|---|---|---|
| chr2 | 33204154 | 33204716 | Intron |
| chr2 | 33212136 | 33213533 | Intron |
| chr2 | 33353648 | 33354040 | Intron |
| chr2 | 33354158 | 33354450 | Intron |
| chr2 | 34609825 | 34610177 | Distal Intergenic |
| chr2 | 34610305 | 34612136 | Distal Intergenic |
| chr2 | 34625508 | 34625754 | Distal Intergenic |
| chr2 | 34625963 | 34626224 | Distal Intergenic |
| chr2 | 34626314 | 34628018 | Distal Intergenic |
| chr2 | 34655041 | 34655240 | Distal Intergenic |
| chr2 | 34655302 | 34655779 | Distal Intergenic |
| chr2 | 34655847 | 34656100 | Distal Intergenic |
| chr2 | 34657487 | 34658600 | Distal Intergenic |
| chr2 | 34681922 | 34682189 | Distal Intergenic |
| chr2 | 34688401 | 34688863 | Distal Intergenic |
| chr2 | 34987196 | 34988303 | Distal Intergenic |
| chr2 | 35049567 | 35049969 | Distal Intergenic |
| chr2 | 35055286 | 35055869 | Distal Intergenic |
| chr2 | 35056793 | 35056992 | Distal Intergenic |
| chr2 | 35059298 | 35059925 | Distal Intergenic |
| chr2 | 35085725 | 35086001 | Distal Intergenic |
| chr2 | 35121066 | 35122802 | Distal Intergenic |
| chr2 | 35122891 | 35126550 | Distal Intergenic |
| chr2 | 35127266 | 35127778 | Distal Intergenic |
| chr2 | 35165335 | 35166098 | Distal Intergenic |
| chr2 | 35170949 | 35171292 | Distal Intergenic |
| chr2 | 35189451 | 35189650 | Distal Intergenic |
| chr2 | 35197877 | 35198745 | Distal Intergenic |
| chr2 | 35201418 | 35202829 | Distal Intergenic |
| chr2 | 35253766 | 35254070 | Distal Intergenic |
| chr2 | 35279348 | 35280446 | Distal Intergenic |
| chr2 | 35289348 | 35289633 | Distal Intergenic |
| chr2 | 35300234 | 35303588 | Distal Intergenic |
| chr2 | 35316252 | 35318562 | Distal Intergenic |
| chr2 | 35333736 | 35334150 | Distal Intergenic |
| chr2 | 35342673 | 35342925 | Distal Intergenic |
| chr2 | 35345403 | 35346411 | Distal Intergenic |
| chr2 | 35357115 | 35358032 | Distal Intergenic |
| chr2 | 35363287 | 35363530 | Distal Intergenic |
| chr2 | 35517141 | 35517904 | Distal Intergenic |
| chr2 | 35518073 | 35518272 | Distal Intergenic |
| chr2 | 35546890 | 35547823 | Distal Intergenic |
| chr2 | 35551942 | 35552420 | Distal Intergenic |
| chr2 | 35558394 | 35559302 | Distal Intergenic |

| | | | |
|---|---|---|---|
| chr2 | 35570018 | 35570374 | Distal Intergenic |
| chr2 | 35894393 | 35895970 | Distal Intergenic |
| chr2 | 35991702 | 35992135 | Distal Intergenic |
| chr2 | 35992185 | 35992432 | Distal Intergenic |
| chr2 | 36073044 | 36073392 | Distal Intergenic |
| chr2 | 36077357 | 36078974 | Distal Intergenic |
| chr2 | 36081475 | 36083352 | Distal Intergenic |
| chr2 | 36089624 | 36092159 | Distal Intergenic |
| chr2 | 36094114 | 36094321 | Distal Intergenic |
| chr2 | 36094412 | 36094938 | Distal Intergenic |
| chr2 | 36095671 | 36096211 | Distal Intergenic |
| chr2 | 36130580 | 36132145 | Distal Intergenic |
| chr2 | 37214593 | 37215089 | Intron |
| chr2 | 37277484 | 37278511 | Intron |
| chr2 | 37278619 | 37278897 | Intron |
| chr2 | 37285983 | 37287330 | Exon |
| chr2 | 37298006 | 37298625 | Intron |
| chr2 | 37298800 | 37299617 | Exon |
| chr2 | 37307249 | 37307599 | Intron |
| chr2 | 37307803 | 37308095 | Intron |
| chr2 | 37514129 | 37514474 | Intron |
| chr2 | 37564632 | 37565042 | Distal Intergenic |
| chr2 | 37565088 | 37567147 | Distal Intergenic |
| chr2 | 37567328 | 37567565 | Distal Intergenic |
| chr2 | 37858544 | 37858772 | Distal Intergenic |
| chr2 | 38367988 | 38368225 | Intron |
| chr2 | 38476747 | 38477701 | Distal Intergenic |
| chr2 | 38498657 | 38499439 | Distal Intergenic |
| chr2 | 38595262 | 38595576 | Intron |
| chr2 | 38598295 | 38598494 | Intron |
| chr2 | 38640690 | 38640927 | Distal Intergenic |
| chr2 | 38721637 | 38723262 | Distal Intergenic |
| chr2 | 38730318 | 38730517 | Distal Intergenic |
| chr2 | 38781381 | 38781592 | Distal Intergenic |
| chr2 | 38781669 | 38781913 | Distal Intergenic |
| chr2 | 38782031 | 38782285 | Distal Intergenic |
| chr2 | 38805855 | 38806445 | Intron |
| chr2 | 38829124 | 38830361 | Promoter |
| chr2 | 38842020 | 38842601 | Distal Intergenic |
| chr2 | 38842732 | 38844330 | Distal Intergenic |
| chr2 | 38859564 | 38861809 | Distal Intergenic |
| chr2 | 38862829 | 38863545 | Distal Intergenic |
| chr2 | 38863839 | 38864290 | Distal Intergenic |
| chr2 | 38920694 | 38920910 | Intron |

| | | | |
|---|---|---|---|
| chr2 | 38921173 | 38921531 | Intron |
| chr2 | 39081116 | 39082023 | Promoter |
| chr2 | 42828458 | 42828799 | Intron |
| chr2 | 44674258 | 44674727 | Intron |
| chr2 | 44776086 | 44776412 | Promoter |
| chr2 | 44782043 | 44782407 | Intron |
| chr2 | 44800965 | 44801473 | Intron |
| chr2 | 44808110 | 44808967 | Intron |
| chr2 | 44811683 | 44813243 | Intron |
| chr2 | 45309603 | 45311189 | Distal Intergenic |
| chr2 | 45311333 | 45311664 | Distal Intergenic |
| chr2 | 45632434 | 45633164 | Intron |
| chr2 | 46086141 | 46087070 | Intron |
| chr2 | 46110324 | 46111413 | Intron |
| chr2 | 46180719 | 46181629 | Intron |
| chr2 | 46298599 | 46299603 | Intron |
| chr2 | 46346607 | 46347061 | Intron |
| chr2 | 46368540 | 46368739 | Intron |
| chr2 | 47733502 | 47734017 | Intron |
| chr2 | 47932715 | 47933753 | Distal Intergenic |
| chr2 | 48295051 | 48295271 | Distal Intergenic |
| chr2 | 48296617 | 48296824 | Distal Intergenic |
| chr2 | 48804372 | 48804716 | Intron |
| chr2 | 48804971 | 48805304 | Intron |
| chr2 | 49664846 | 49665341 | Distal Intergenic |
| chr2 | 49666613 | 49668202 | Distal Intergenic |
| chr2 | 49669075 | 49669795 | Distal Intergenic |
| chr2 | 49672894 | 49673095 | Distal Intergenic |
| chr2 | 49764322 | 49767558 | Distal Intergenic |
| chr2 | 49767611 | 49768091 | Distal Intergenic |
| chr2 | 49986154 | 49986386 | Distal Intergenic |
| chr2 | 50151243 | 50151575 | Intron |
| chr2 | 50151611 | 50152131 | Intron |
| chr2 | 50221063 | 50221361 | Intron |
| chr2 | 50223451 | 50224556 | Intron |
| chr2 | 50232016 | 50232804 | Intron |
| chr2 | 50317470 | 50317736 | Intron |
| chr2 | 50317782 | 50318819 | Exon |
| chr2 | 50318932 | 50319369 | Intron |
| chr2 | 50409066 | 50410392 | Intron |
| chr2 | 50519632 | 50520085 | Intron |
| chr2 | 50523636 | 50523989 | Intron |
| chr2 | 50542267 | 50545509 | Intron |
| chr2 | 50545559 | 50547133 | Intron |

| | | | |
|---|---|---|---|
| chr2 | 50647577 | 50649911 | Intron |
| chr2 | 50680331 | 50681911 | Intron |
| chr2 | 50741294 | 50741758 | Intron |
| chr2 | 50742023 | 50742287 | Intron |
| chr2 | 50818837 | 50820292 | Intron |
| chr2 | 50870313 | 50871464 | Intron |
| chr2 | 50917354 | 50917582 | Intron |
| chr2 | 50978808 | 50979134 | Intron |
| chr2 | 50979187 | 50979420 | Intron |
| chr2 | 50993049 | 50993459 | Intron |
| chr2 | 50994252 | 50994511 | Intron |
| chr2 | 50994672 | 50995438 | Intron |
| chr2 | 50995490 | 51000798 | Intron |
| chr2 | 51001066 | 51004659 | Intron |
| chr2 | 51026435 | 51026697 | Intron |
| chr2 | 51026836 | 51027567 | Intron |
| chr2 | 51029444 | 51031133 | Intron |
| chr2 | 51032507 | 51032989 | Intron |
| chr2 | 51033099 | 51033584 | Intron |
| chr2 | 51034524 | 51035518 | Intron |
| chr2 | 51082721 | 51084067 | Intron |
| chr2 | 51134126 | 51135466 | Intron |
| chr2 | 51160255 | 51160845 | Intron |
| chr2 | 51160922 | 51161136 | Intron |
| chr2 | 51274235 | 51274605 | Distal Intergenic |
| chr2 | 51294564 | 51294786 | Distal Intergenic |
| chr2 | 51294974 | 51295282 | Distal Intergenic |
| chr2 | 51790174 | 51790409 | Distal Intergenic |
| chr2 | 52600741 | 52601431 | Distal Intergenic |
| chr2 | 52728139 | 52728558 | Distal Intergenic |
| chr2 | 52732627 | 52733155 | Distal Intergenic |
| chr2 | 52742636 | 52742965 | Distal Intergenic |
| chr2 | 52743628 | 52745160 | Distal Intergenic |
| chr2 | 52745677 | 52747868 | Distal Intergenic |
| chr2 | 52749750 | 52750935 | Distal Intergenic |
| chr2 | 52797262 | 52797463 | Distal Intergenic |
| chr2 | 52797565 | 52797922 | Distal Intergenic |
| chr2 | 52801868 | 52802067 | Distal Intergenic |
| chr2 | 52802245 | 52803294 | Distal Intergenic |
| chr2 | 52810878 | 52811872 | Distal Intergenic |
| chr2 | 52813828 | 52815378 | Distal Intergenic |
| chr2 | 52887837 | 52888068 | Distal Intergenic |
| chr2 | 52888194 | 52888393 | Distal Intergenic |
| chr2 | 53049334 | 53050270 | Distal Intergenic |

| | | | |
|---|---|---|---|
| chr2 | 53050797 | 53051059 | Distal Intergenic |
| chr2 | 53051486 | 53051888 | Distal Intergenic |
| chr2 | 55001848 | 55002047 | Intron |
| chr2 | 55002342 | 55002719 | Intron |
| chr2 | 55057095 | 55059803 | Intron |
| chr2 | 55059947 | 55060188 | Intron |
| chr2 | 55069143 | 55070206 | Intron |
| chr2 | 55070252 | 55070660 | Intron |
| chr2 | 55105573 | 55105936 | Intron |
| chr2 | 55106072 | 55106522 | Intron |
| chr2 | 56969291 | 56969891 | Distal Intergenic |
| chr2 | 57082229 | 57082917 | Distal Intergenic |
| chr2 | 57082961 | 57083369 | Distal Intergenic |
| chr2 | 57083626 | 57084558 | Distal Intergenic |
| chr2 | 57090068 | 57091525 | Distal Intergenic |
| chr2 | 57091684 | 57092119 | Distal Intergenic |
| chr2 | 57112994 | 57113731 | Distal Intergenic |
| chr2 | 57113932 | 57114247 | Distal Intergenic |
| chr2 | 57115680 | 57116070 | Distal Intergenic |
| chr2 | 57116398 | 57116980 | Distal Intergenic |
| chr2 | 57206120 | 57206632 | Distal Intergenic |
| chr2 | 58333050 | 58333636 | Intron |
| chr2 | 58333824 | 58334094 | Intron |
| chr2 | 58334266 | 58334631 | Intron |
| chr2 | 58335555 | 58335881 | Intron |
| chr2 | 58335990 | 58336208 | Intron |
| chr2 | 58336323 | 58338816 | Intron |
| chr2 | 58338856 | 58339781 | Intron |
| chr2 | 58349446 | 58349789 | Intron |
| chr2 | 58357868 | 58359084 | Exon |
| chr2 | 58382341 | 58382540 | Intron |
| chr2 | 58394585 | 58395248 | Intron |
| chr2 | 58398737 | 58400136 | Intron |
| chr2 | 58400338 | 58400605 | Intron |
| chr2 | 58400839 | 58402525 | Intron |
| chr2 | 58414289 | 58414579 | Intron |
| chr2 | 58524061 | 58524694 | Distal Intergenic |
| chr2 | 58524864 | 58525653 | Distal Intergenic |
| chr2 | 58682424 | 58683130 | Distal Intergenic |
| chr2 | 58898031 | 58898230 | Intron |
| chr2 | 59009213 | 59009736 | Intron |
| chr2 | 59040979 | 59041645 | Intron |
| chr2 | 59041721 | 59041920 | Intron |
| chr2 | 59177707 | 59178659 | Intron |

| | | | |
|---|---|---|---|
| chr2 | 59189469 | 59189732 | Intron |
| chr2 | 59276394 | 59276618 | Intron |
| chr2 | 59285410 | 59286826 | Exon |
| chr2 | 59322660 | 59323219 | Distal Intergenic |
| chr2 | 59323268 | 59323660 | Distal Intergenic |
| chr2 | 59425461 | 59426222 | Distal Intergenic |
| chr2 | 59450596 | 59450829 | Distal Intergenic |
| chr2 | 59455538 | 59456034 | Distal Intergenic |
| chr2 | 59507897 | 59509032 | Distal Intergenic |
| chr2 | 59514124 | 59515578 | Distal Intergenic |
| chr2 | 59517479 | 59518274 | Distal Intergenic |
| chr2 | 59607051 | 59607568 | Distal Intergenic |
| chr2 | 59607682 | 59608050 | Distal Intergenic |
| chr2 | 59613480 | 59614752 | Distal Intergenic |
| chr2 | 59616727 | 59618588 | Distal Intergenic |
| chr2 | 59638625 | 59638872 | Distal Intergenic |
| chr2 | 59658632 | 59658854 | Distal Intergenic |
| chr2 | 59661817 | 59662460 | Distal Intergenic |
| chr2 | 59734429 | 59734865 | Distal Intergenic |
| chr2 | 59836070 | 59836333 | Distal Intergenic |
| chr2 | 59932087 | 59932483 | Distal Intergenic |
| chr2 | 59962129 | 59963028 | Distal Intergenic |
| chr2 | 59963203 | 59963990 | Distal Intergenic |
| chr2 | 60009673 | 60010018 | Distal Intergenic |
| chr2 | 60010056 | 60010382 | Distal Intergenic |
| chr2 | 60048264 | 60049392 | Distal Intergenic |
| chr2 | 60076775 | 60077954 | Distal Intergenic |
| chr2 | 60376855 | 60378066 | Distal Intergenic |
| chr2 | 60381955 | 60383598 | Distal Intergenic |
| chr2 | 60387843 | 60388455 | Distal Intergenic |
| chr2 | 60471019 | 60472436 | Distal Intergenic |
| chr2 | 60472878 | 60473655 | Distal Intergenic |
| chr2 | 60475365 | 60479725 | Distal Intergenic |
| chr2 | 60509917 | 60510116 | Distal Intergenic |
| chr2 | 60510159 | 60512239 | Distal Intergenic |
| chr2 | 60512325 | 60512587 | Distal Intergenic |
| chr2 | 60527000 | 60528338 | Distal Intergenic |
| chr2 | 60529609 | 60534642 | Distal Intergenic |
| chr2 | 60535870 | 60536359 | Distal Intergenic |
| chr2 | 60537942 | 60539703 | Distal Intergenic |
| chr2 | 60549035 | 60550267 | Distal Intergenic |
| chr2 | 60645349 | 60646913 | Distal Intergenic |
| chr2 | 60648251 | 60649051 | Distal Intergenic |
| chr2 | 60649471 | 60649681 | Distal Intergenic |

| | | | |
|---|---|---|---|
| chr2 | 60663848 | 60666016 | Distal Intergenic |
| chr2 | 60666754 | 60668183 | Distal Intergenic |
| chr2 | 60668831 | 60669200 | Distal Intergenic |
| chr2 | 60692541 | 60694915 | Intron |
| chr2 | 60694951 | 60695451 | Intron |
| chr2 | 60695510 | 60695828 | Intron |
| chr2 | 60702383 | 60704429 | Intron |
| chr2 | 60707990 | 60708302 | Intron |
| chr2 | 60708399 | 60709884 | Intron |
| chr2 | 60710272 | 60710575 | Intron |
| chr2 | 60715331 | 60719105 | Intron |
| chr2 | 60719790 | 60720327 | Intron |
| chr2 | 60739281 | 60739510 | Intron |
| chr2 | 60739617 | 60740816 | Intron |
| chr2 | 60800543 | 60800945 | Distal Intergenic |
| chr2 | 60902443 | 60902794 | Distal Intergenic |
| chr2 | 61105648 | 61106167 | Intron |
| chr2 | 61106511 | 61108028 | Promoter |
| chr2 | 61146907 | 61148512 | Exon |
| chr2 | 61431207 | 61432654 | Promoter |
| chr2 | 61549811 | 61551047 | Intron |
| chr2 | 61579380 | 61579579 | Intron |
| chr2 | 64877832 | 64878508 | Intron |
| chr2 | 64979227 | 64979798 | Distal Intergenic |
| chr2 | 64985721 | 64985949 | Distal Intergenic |
| chr2 | 65041835 | 65042074 | Distal Intergenic |
| chr2 | 65045865 | 65046281 | Distal Intergenic |
| chr2 | 65059613 | 65061805 | Distal Intergenic |
| chr2 | 65065122 | 65066202 | Distal Intergenic |
| chr2 | 65074685 | 65075878 | Exon |
| chr2 | 65076213 | 65076436 | Intron |
| chr2 | 65077206 | 65077513 | Intron |
| chr2 | 65103158 | 65103387 | Distal Intergenic |
| chr2 | 65104972 | 65107569 | Distal Intergenic |
| chr2 | 65248033 | 65248410 | 3' UTR |
| chr2 | 65961974 | 65962181 | Intron |
| chr2 | 66018583 | 66018871 | Intron |
| chr2 | 68060219 | 68060418 | Distal Intergenic |
| chr2 | 68060482 | 68061075 | Distal Intergenic |
| chr2 | 68098289 | 68099031 | Distal Intergenic |
| chr2 | 68141961 | 68143989 | Distal Intergenic |
| chr2 | 68700127 | 68700935 | Intron |
| chr2 | 68716823 | 68717777 | Exon |
| chr2 | 68748669 | 68749565 | Intron |

| | | | |
|---|---|---|---|
| chr2 | 68753693 | 68754188 | Intron |
| chr2 | 68754637 | 68755716 | Intron |
| chr2 | 68762667 | 68765472 | 5' UTR |
| chr2 | 68772880 | 68774699 | Intron |
| chr2 | 68818051 | 68818598 | Intron |
| chr2 | 68819095 | 68819294 | Intron |
| chr2 | 68820499 | 68823122 | Intron |
| chr2 | 68823486 | 68824019 | Intron |
| chr2 | 68824168 | 68827296 | Intron |
| chr2 | 68832304 | 68834012 | Intron |
| chr2 | 68894683 | 68895558 | Distal Intergenic |
| chr2 | 68911254 | 68913173 | Intron |
| chr2 | 68914113 | 68916196 | Intron |
| chr2 | 69160792 | 69161377 | Distal Intergenic |
| chr2 | 69238841 | 69239854 | Promoter |
| chr2 | 69484832 | 69485132 | Distal Intergenic |
| chr2 | 69560888 | 69561264 | Intron |
| chr2 | 69561372 | 69561831 | Exon |
| chr2 | 69627925 | 69628140 | Intron |
| chr2 | 69628196 | 69628483 | Intron |
| chr2 | 69698973 | 69699208 | 3' UTR |
| chr2 | 69699565 | 69700676 | 3' UTR |
| chr2 | 70499762 | 70500243 | Intron |
| chr2 | 70649724 | 70649935 | Distal Intergenic |
| chr2 | 70893798 | 70893999 | Intron |
| chr2 | 71049778 | 71050448 | Distal Intergenic |
| chr2 | 71069583 | 71070366 | Distal Intergenic |
| chr2 | 71380300 | 71380761 | Distal Intergenic |
| chr2 | 71427319 | 71427941 | Intron |
| chr2 | 71431839 | 71433614 | Intron |
| chr2 | 71435972 | 71437066 | Intron |
| chr2 | 71460583 | 71460792 | Distal Intergenic |
| chr2 | 71460866 | 71461122 | Distal Intergenic |
| chr2 | 71461208 | 71461621 | Distal Intergenic |
| chr2 | 71464403 | 71464961 | Distal Intergenic |
| chr2 | 71477865 | 71479483 | Distal Intergenic |
| chr2 | 71481798 | 71482157 | Distal Intergenic |
| chr2 | 71482386 | 71483015 | Distal Intergenic |
| chr2 | 71489239 | 71491831 | Distal Intergenic |
| chr2 | 71527975 | 71529250 | Intron |
| chr2 | 71576261 | 71578420 | Promoter |
| chr2 | 71609815 | 71610040 | Intron |
| chr2 | 71625723 | 71628698 | 5' UTR |
| chr2 | 71628770 | 71629142 | 5' UTR |

| | | | |
|---|---|---|---|
| chr2 | 71638236 | 71638506 | Intron |
| chr2 | 71638596 | 71641109 | Intron |
| chr2 | 71641161 | 71641402 | Intron |
| chr2 | 71642041 | 71644736 | Promoter |
| chr2 | 71652149 | 71652387 | Intron |
| chr2 | 72079560 | 72079954 | Distal Intergenic |
| chr2 | 72079994 | 72080225 | Distal Intergenic |
| chr2 | 72091111 | 72091350 | Distal Intergenic |
| chr2 | 72118704 | 72119360 | Distal Intergenic |
| chr2 | 72123206 | 72127911 | Distal Intergenic |
| chr2 | 72127958 | 72128297 | Distal Intergenic |
| chr2 | 72134972 | 72136643 | Distal Intergenic |
| chr2 | 72146200 | 72146465 | Distal Intergenic |
| chr2 | 72146629 | 72148657 | Distal Intergenic |
| chr2 | 72168479 | 72169464 | Distal Intergenic |
| chr2 | 72188492 | 72188733 | Distal Intergenic |
| chr2 | 72188988 | 72190649 | Distal Intergenic |
| chr2 | 72282162 | 72282495 | Distal Intergenic |
| chr2 | 72282554 | 72282783 | Distal Intergenic |
| chr2 | 72282941 | 72285491 | Distal Intergenic |
| chr2 | 72313808 | 72314154 | Distal Intergenic |
| chr2 | 72314352 | 72314611 | Distal Intergenic |
| chr2 | 72314827 | 72315511 | Distal Intergenic |
| chr2 | 72570859 | 72574068 | Intron |
| chr2 | 72574360 | 72575418 | Intron |
| chr2 | 72719823 | 72720469 | Intron |
| chr2 | 72746651 | 72747791 | Intron |
| chr2 | 72818368 | 72820710 | Intron |
| chr2 | 72833220 | 72835729 | Intron |
| chr2 | 72843674 | 72844145 | Intron |
| chr2 | 72861861 | 72862071 | Intron |
| chr2 | 72873597 | 72874502 | Intron |
| chr2 | 72879064 | 72880269 | Intron |
| chr2 | 73051856 | 73052055 | Intron |
| chr2 | 73150335 | 73150686 | Intron |
| chr2 | 73423278 | 73423818 | Distal Intergenic |
| chr2 | 73749055 | 73749289 | Intron |
| chr2 | 73749472 | 73749916 | Intron |
| chr2 | 74688775 | 74689456 | Exon |
| chr2 | 75447041 | 75449114 | Distal Intergenic |
| chr2 | 75450672 | 75451212 | Distal Intergenic |
| chr2 | 75451275 | 75451985 | Distal Intergenic |
| chr2 | 75452149 | 75453069 | Distal Intergenic |
| chr2 | 75469428 | 75469790 | Distal Intergenic |

| | | | |
|---|---|---|---|
| chr2 | 75478546 | 75478828 | Distal Intergenic |
| chr2 | 75479002 | 75482344 | Distal Intergenic |
| chr2 | 75495863 | 75497218 | Distal Intergenic |
| chr2 | 75497344 | 75497598 | Distal Intergenic |
| chr2 | 75497698 | 75499630 | Distal Intergenic |
| chr2 | 75499806 | 75500453 | Distal Intergenic |
| chr2 | 75501716 | 75501930 | Distal Intergenic |
| chr2 | 75507171 | 75507960 | Distal Intergenic |
| chr2 | 75520762 | 75524360 | Distal Intergenic |
| chr2 | 75527137 | 75527422 | Distal Intergenic |
| chr2 | 75533230 | 75533432 | Distal Intergenic |
| chr2 | 75533622 | 75534588 | Distal Intergenic |
| chr2 | 75534751 | 75534972 | Distal Intergenic |
| chr2 | 75539006 | 75542963 | Distal Intergenic |
| chr2 | 75603242 | 75603834 | Distal Intergenic |
| chr2 | 75605349 | 75607097 | Distal Intergenic |
| chr2 | 75607184 | 75607668 | Distal Intergenic |
| chr2 | 75610265 | 75611187 | Distal Intergenic |
| chr2 | 75669761 | 75669975 | Distal Intergenic |
| chr2 | 75670337 | 75671139 | Distal Intergenic |
| chr2 | 75707424 | 75707768 | Distal Intergenic |
| chr2 | 75895851 | 75896637 | Intron |
| chr2 | 75898137 | 75898537 | Intron |
| chr2 | 76895174 | 76895373 | Distal Intergenic |
| chr2 | 77106314 | 77107356 | Intron |
| chr2 | 77783543 | 77784569 | Distal Intergenic |
| chr2 | 78962818 | 78963228 | Distal Intergenic |
| chr2 | 78998933 | 79000194 | Distal Intergenic |
| chr2 | 79004433 | 79006049 | Distal Intergenic |
| chr2 | 79007465 | 79009041 | Distal Intergenic |
| chr2 | 79238704 | 79239055 | Distal Intergenic |
| chr2 | 79372664 | 79373310 | Distal Intergenic |
| chr2 | 79396002 | 79396201 | Distal Intergenic |
| chr2 | 79406392 | 79406611 | Distal Intergenic |
| chr2 | 79436955 | 79438069 | Intron |
| chr2 | 79443055 | 79444729 | Intron |
| chr2 | 79445081 | 79445328 | Intron |
| chr2 | 79450723 | 79451278 | Intron |
| chr2 | 79458309 | 79460245 | Intron |
| chr2 | 79462152 | 79462448 | Intron |
| chr2 | 79462495 | 79463191 | Intron |
| chr2 | 79475907 | 79476644 | Intron |
| chr2 | 79496888 | 79497282 | Intron |
| chr2 | 79538228 | 79538493 | Intron |

| | | | |
|---|---|---|---|
| chr2 | 79538622 | 79538904 | Intron |
| chr2 | 80019730 | 80020332 | Intron |
| chr2 | 80419817 | 80421107 | Intron |
| chr2 | 80478656 | 80478918 | Intron |
| chr2 | 80478988 | 80479345 | Intron |
| chr2 | 80528067 | 80528766 | Intron |
| chr2 | 81733113 | 81733331 | Distal Intergenic |
| chr2 | 82018444 | 82018784 | Distal Intergenic |
| chr2 | 82018949 | 82019378 | Distal Intergenic |
| chr2 | 83485029 | 83485371 | Distal Intergenic |
| chr2 | 83511975 | 83512309 | Distal Intergenic |
| chr2 | 83515034 | 83515280 | Distal Intergenic |
| chr2 | 83563902 | 83564524 | Distal Intergenic |
| chr2 | 83564805 | 83566236 | Distal Intergenic |
| chr2 | 83568016 | 83569505 | Distal Intergenic |
| chr2 | 83584436 | 83585076 | Distal Intergenic |
| chr2 | 84333617 | 84334260 | Distal Intergenic |
| chr2 | 84351870 | 84352108 | Distal Intergenic |
| chr2 | 84369099 | 84369358 | Distal Intergenic |
| chr2 | 84379222 | 84380214 | Distal Intergenic |
| chr2 | 84380318 | 84380533 | Distal Intergenic |
| chr2 | 84380663 | 84381583 | Distal Intergenic |
| chr2 | 84384107 | 84386359 | Distal Intergenic |
| chr2 | 84388454 | 84392502 | Distal Intergenic |
| chr2 | 84458197 | 84460201 | Distal Intergenic |
| chr2 | 84465898 | 84469354 | Distal Intergenic |
| chr2 | 84470434 | 84470723 | Distal Intergenic |
| chr2 | 84472349 | 84477298 | Distal Intergenic |
| chr2 | 84478361 | 84478587 | Distal Intergenic |
| chr2 | 84479917 | 84480495 | Distal Intergenic |
| chr2 | 84487948 | 84488203 | Distal Intergenic |
| chr2 | 84488325 | 84491650 | Distal Intergenic |
| chr2 | 84510524 | 84510751 | Distal Intergenic |
| chr2 | 84510916 | 84511472 | Distal Intergenic |
| chr2 | 84544034 | 84546506 | Distal Intergenic |
| chr2 | 84546741 | 84547114 | Distal Intergenic |
| chr2 | 84553957 | 84555389 | Distal Intergenic |
| chr2 | 84555875 | 84556713 | Distal Intergenic |
| chr2 | 84583592 | 84583945 | Distal Intergenic |
| chr2 | 84584422 | 84584646 | Distal Intergenic |
| chr2 | 84667174 | 84667376 | Intron |
| chr2 | 84877111 | 84877437 | Intron |
| chr2 | 84877677 | 84877967 | Intron |
| chr2 | 84887677 | 84888128 | Intron |

| | | | |
|---|---|---|---|
| chr2 | 84888195 | 84888468 | Intron |
| chr2 | 84888697 | 84888956 | Intron |
| chr2 | 84889938 | 84891547 | Intron |
| chr2 | 84900158 | 84901904 | Intron |
| chr2 | 84906437 | 84909163 | Exon |
| chr2 | 85349984 | 85350354 | Distal Intergenic |
| chr2 | 85350398 | 85351381 | Distal Intergenic |
| chr2 | 85351525 | 85352089 | Distal Intergenic |
| chr2 | 85394590 | 85395642 | Intron |
| chr2 | 90336188 | 90336994 | Intron |
| chr2 | 90340754 | 90341442 | Intron |
| chr2 | 90382631 | 90382974 | Intron |
| chr2 | 90403275 | 90403487 | Intron |
| chr2 | 90405624 | 90406516 | Intron |
| chr2 | 90451245 | 90453059 | Intron |
| chr2 | 90453137 | 90453498 | Intron |
| chr2 | 90585153 | 90585467 | Distal Intergenic |
| chr2 | 90586053 | 90586530 | Distal Intergenic |
| chr2 | 90744305 | 90744826 | Distal Intergenic |
| chr2 | 90744932 | 90745362 | Distal Intergenic |
| chr2 | 90757062 | 90757814 | Distal Intergenic |
| chr2 | 90757852 | 90758707 | Distal Intergenic |
| chr2 | 90894193 | 90894580 | Distal Intergenic |
| chr2 | 90901722 | 90902261 | Distal Intergenic |
| chr2 | 90909258 | 90910420 | Distal Intergenic |
| chr2 | 90910769 | 90911281 | Distal Intergenic |
| chr2 | 91077324 | 91077742 | Distal Intergenic |
| chr2 | 91077787 | 91078040 | Distal Intergenic |
| chr2 | 91082714 | 91084044 | Distal Intergenic |
| chr2 | 91097180 | 91097386 | Distal Intergenic |
| chr2 | 91097422 | 91098251 | Distal Intergenic |
| chr2 | 91104970 | 91105297 | Distal Intergenic |
| chr2 | 91105372 | 91105950 | Distal Intergenic |
| chr2 | 91139668 | 91139982 | Distal Intergenic |
| chr2 | 91355599 | 91355933 | Distal Intergenic |
| chr2 | 91366948 | 91367166 | Distal Intergenic |
| chr2 | 91377402 | 91377708 | Distal Intergenic |
| chr2 | 91489333 | 91490447 | Distal Intergenic |
| chr2 | 91536170 | 91536521 | Distal Intergenic |
| chr2 | 91536599 | 91536844 | Distal Intergenic |
| chr2 | 91550580 | 91550979 | Distal Intergenic |
| chr2 | 91551040 | 91551816 | Distal Intergenic |
| chr2 | 91570657 | 91571057 | Distal Intergenic |
| chr2 | 91787660 | 91789134 | Distal Intergenic |

| | | | |
|---|---|---|---|
| chr2 | 91804304 | 91806422 | Exon |
| chr2 | 91852313 | 91853186 | Distal Intergenic |
| chr2 | 92018853 | 92019956 | Distal Intergenic |
| chr2 | 92020205 | 92020749 | Distal Intergenic |
| chr2 | 92024257 | 92024713 | Distal Intergenic |
| chr2 | 92044496 | 92044911 | Distal Intergenic |
| chr2 | 92097704 | 92098773 | Distal Intergenic |
| chr2 | 92211748 | 92212017 | Distal Intergenic |
| chr2 | 92212238 | 92214257 | Distal Intergenic |
| chr2 | 92214890 | 92215860 | Distal Intergenic |
| chr2 | 92267607 | 92268606 | Distal Intergenic |
| chr2 | 92518679 | 92518887 | Distal Intergenic |
| chr2 | 92776891 | 92778759 | Distal Intergenic |
| chr2 | 92779097 | 92779751 | Distal Intergenic |
| chr2 | 92785340 | 92785807 | Distal Intergenic |
| chr2 | 92788773 | 92789100 | Distal Intergenic |
| chr2 | 92835512 | 92835798 | Distal Intergenic |
| chr2 | 92973230 | 92974533 | Distal Intergenic |
| chr2 | 93023261 | 93028090 | Distal Intergenic |
| chr2 | 93616649 | 93617230 | Distal Intergenic |
| chr2 | 93742954 | 93743190 | Distal Intergenic |
| chr2 | 93747580 | 93747854 | Distal Intergenic |
| chr2 | 93997137 | 93998064 | Distal Intergenic |
| chr2 | 94246565 | 94246768 | Distal Intergenic |
| chr2 | 94246886 | 94247611 | Distal Intergenic |
| chr2 | 95230549 | 95231023 | Distal Intergenic |
| chr2 | 97278163 | 97278488 | Exon |
| chr2 | 97474239 | 97475490 | 3' UTR |
| chr2 | 97550471 | 97551385 | Intron |
| chr2 | 99973321 | 99973573 | Intron |
| chr2 | 101518599 | 101518802 | Intron |
| chr2 | 101632283 | 101633436 | Intron |
| chr2 | 101826900 | 101827099 | Distal Intergenic |
| chr2 | 101945292 | 101946038 | Distal Intergenic |
| chr2 | 102143096 | 102143582 | Distal Intergenic |
| chr2 | 102143644 | 102144358 | Distal Intergenic |
| chr2 | 102273970 | 102274245 | Distal Intergenic |
| chr2 | 102287949 | 102289249 | Distal Intergenic |
| chr2 | 102289332 | 102290035 | Distal Intergenic |
| chr2 | 102290228 | 102290584 | Distal Intergenic |
| chr2 | 102347541 | 102348795 | Intron |
| chr2 | 102713009 | 102713246 | Intron |
| chr2 | 102713402 | 102714963 | Intron |
| chr2 | 102721436 | 102723597 | Intron |

| | | | |
|---|---|---|---|
| chr2 | 102723812 | 102726135 | Intron |
| chr2 | 102728199 | 102728398 | Intron |
| chr2 | 102729198 | 102729421 | Intron |
| chr2 | 102729683 | 102730086 | Intron |
| chr2 | 102730128 | 102730663 | Intron |
| chr2 | 102733646 | 102738352 | Intron |
| chr2 | 102738484 | 102740198 | Intron |
| chr2 | 102740284 | 102741932 | Intron |
| chr2 | 102784236 | 102785131 | Exon |
| chr2 | 102785238 | 102785709 | Intron |
| chr2 | 102787885 | 102789677 | Exon |
| chr2 | 102796461 | 102797701 | Distal Intergenic |
| chr2 | 102797756 | 102799845 | Distal Intergenic |
| chr2 | 102820806 | 102822383 | Intron |
| chr2 | 102913891 | 102914104 | Distal Intergenic |
| chr2 | 102957324 | 102957666 | Intron |
| chr2 | 103078708 | 103079298 | Distal Intergenic |
| chr2 | 103111454 | 103111920 | Intron |
| chr2 | 103350581 | 103350953 | Intron |
| chr2 | 103351013 | 103351255 | Intron |
| chr2 | 103471069 | 103471370 | Distal Intergenic |
| chr2 | 103472525 | 103473947 | Distal Intergenic |
| chr2 | 103495852 | 103496280 | Distal Intergenic |
| chr2 | 103497807 | 103498038 | Distal Intergenic |
| chr2 | 103510238 | 103510464 | Distal Intergenic |
| chr2 | 103527041 | 103528060 | Distal Intergenic |
| chr2 | 103530091 | 103531818 | Distal Intergenic |
| chr2 | 103536323 | 103536777 | Distal Intergenic |
| chr2 | 103601497 | 103601787 | Distal Intergenic |
| chr2 | 103635796 | 103637106 | Distal Intergenic |
| chr2 | 103936147 | 103936454 | Distal Intergenic |
| chr2 | 103936536 | 103936946 | Distal Intergenic |
| chr2 | 103938776 | 103939654 | Distal Intergenic |
| chr2 | 103942632 | 103943381 | Distal Intergenic |
| chr2 | 104306566 | 104307684 | Distal Intergenic |
| chr2 | 104333322 | 104334318 | Distal Intergenic |
| chr2 | 104430576 | 104431259 | Distal Intergenic |
| chr2 | 104431310 | 104431618 | Distal Intergenic |
| chr2 | 104470099 | 104470298 | Distal Intergenic |
| chr2 | 104470347 | 104470617 | Distal Intergenic |
| chr2 | 104539776 | 104540332 | Distal Intergenic |
| chr2 | 104544557 | 104547112 | Distal Intergenic |
| chr2 | 104572063 | 104573459 | Distal Intergenic |
| chr2 | 104581778 | 104582101 | Distal Intergenic |

| | | | |
|---|---|---|---|
| chr2 | 104585363 | 104585688 | Distal Intergenic |
| chr2 | 104603142 | 104603405 | Distal Intergenic |
| chr2 | 104603463 | 104603722 | Distal Intergenic |
| chr2 | 104655669 | 104656359 | Distal Intergenic |
| chr2 | 104657179 | 104657392 | Distal Intergenic |
| chr2 | 104657444 | 104657682 | Distal Intergenic |
| chr2 | 104688550 | 104689765 | Distal Intergenic |
| chr2 | 104695833 | 104696371 | Distal Intergenic |
| chr2 | 104856654 | 104857102 | Distal Intergenic |
| chr2 | 104857233 | 104857750 | Distal Intergenic |
| chr2 | 105212744 | 105214706 | Distal Intergenic |
| chr2 | 105214815 | 105215698 | Distal Intergenic |
| chr2 | 105215830 | 105216570 | Distal Intergenic |
| chr2 | 105238672 | 105239318 | Distal Intergenic |
| chr2 | 105239859 | 105240210 | Distal Intergenic |
| chr2 | 105256643 | 105258312 | Distal Intergenic |
| chr2 | 105636577 | 105636901 | Distal Intergenic |
| chr2 | 105637376 | 105637575 | Distal Intergenic |
| chr2 | 105637793 | 105637992 | Distal Intergenic |
| chr2 | 105744779 | 105745012 | Distal Intergenic |
| chr2 | 105756333 | 105757064 | Distal Intergenic |
| chr2 | 106814319 | 106814522 | Distal Intergenic |
| chr2 | 106830866 | 106832525 | Distal Intergenic |
| chr2 | 106845015 | 106845240 | Distal Intergenic |
| chr2 | 106845353 | 106847337 | Distal Intergenic |
| chr2 | 106916521 | 106916753 | Distal Intergenic |
| chr2 | 106916952 | 106917419 | Distal Intergenic |
| chr2 | 107066909 | 107068886 | Exon |
| chr2 | 107068988 | 107069380 | Exon |
| chr2 | 107072376 | 107073810 | Exon |
| chr2 | 107181392 | 107182082 | Distal Intergenic |
| chr2 | 107232911 | 107233172 | Distal Intergenic |
| chr2 | 108238940 | 108240831 | Distal Intergenic |
| chr2 | 108244572 | 108244931 | Distal Intergenic |
| chr2 | 108462621 | 108464165 | Intron |
| chr2 | 108501471 | 108501736 | Intron |
| chr2 | 108501901 | 108502253 | Intron |
| chr2 | 109121019 | 109121397 | Intron |
| chr2 | 109121467 | 109121687 | Intron |
| chr2 | 109683201 | 109683436 | Distal Intergenic |
| chr2 | 109700675 | 109701621 | Distal Intergenic |
| chr2 | 109730718 | 109730968 | Distal Intergenic |
| chr2 | 109760759 | 109762419 | Intron |
| chr2 | 109763069 | 109763705 | Intron |

| | | | |
|---|---|---|---|
| chr2 | 109799244 | 109799514 | Intron |
| chr2 | 109809614 | 109809844 | Intron |
| chr2 | 109810018 | 109810351 | Intron |
| chr2 | 109824845 | 109825891 | Intron |
| chr2 | 109892570 | 109894022 | Intron |
| chr2 | 109995954 | 109996175 | Intron |
| chr2 | 109996253 | 109997202 | Intron |
| chr2 | 110012488 | 110014121 | Intron |
| chr2 | 110016183 | 110017876 | Intron |
| chr2 | 110025734 | 110027718 | Intron |
| chr2 | 110032032 | 110033400 | Intron |
| chr2 | 110360212 | 110361793 | Intron |
| chr2 | 110549928 | 110550859 | Promoter |
| chr2 | 110558571 | 110559502 | Exon |
| chr2 | 112043044 | 112044027 | Intron |
| chr2 | 112077958 | 112078703 | Promoter |
| chr2 | 112124932 | 112125144 | Exon |
| chr2 | 112147773 | 112148253 | Intron |
| chr2 | 112148306 | 112149817 | Intron |
| chr2 | 112213192 | 112214176 | Intron |
| chr2 | 112217939 | 112218431 | Intron |
| chr2 | 112219717 | 112220562 | Intron |
| chr2 | 112238401 | 112239744 | Intron |
| chr2 | 112294808 | 112295115 | Distal Intergenic |
| chr2 | 112295600 | 112295829 | Distal Intergenic |
| chr2 | 113214646 | 113214996 | Distal Intergenic |
| chr2 | 113216242 | 113216778 | Distal Intergenic |
| chr2 | 113216816 | 113217139 | Distal Intergenic |
| chr2 | 113236538 | 113237037 | Distal Intergenic |
| chr2 | 113259753 | 113261922 | Exon |
| chr2 | 113280364 | 113281434 | Intron |
| chr2 | 113657998 | 113658898 | Distal Intergenic |
| chr2 | 113661185 | 113661797 | Distal Intergenic |
| chr2 | 113687948 | 113689336 | Distal Intergenic |
| chr2 | 114071796 | 114072279 | Distal Intergenic |
| chr2 | 114480274 | 114480853 | Exon |
| chr2 | 114622871 | 114623975 | Distal Intergenic |
| chr2 | 115406823 | 115407841 | Intron |
| chr2 | 115434306 | 115436493 | Intron |
| chr2 | 115605815 | 115606999 | Intron |
| chr2 | 115796171 | 115796417 | Intron |
| chr2 | 115891868 | 115892178 | Intron |
| chr2 | 116552952 | 116553687 | Intron |
| chr2 | 116791808 | 116792111 | Distal Intergenic |

| | | | |
|---|---|---|---|
| chr2 | 116797812 | 116798343 | Distal Intergenic |
| chr2 | 117075490 | 117075933 | Distal Intergenic |
| chr2 | 117186195 | 117187267 | Distal Intergenic |
| chr2 | 117468769 | 117469189 | Distal Intergenic |
| chr2 | 117469364 | 117470316 | Distal Intergenic |
| chr2 | 117493578 | 117494056 | Distal Intergenic |
| chr2 | 117556347 | 117557607 | Distal Intergenic |
| chr2 | 117612856 | 117613243 | Distal Intergenic |
| chr2 | 117613410 | 117615198 | Distal Intergenic |
| chr2 | 117628161 | 117628630 | Distal Intergenic |
| chr2 | 117628853 | 117629662 | Distal Intergenic |
| chr2 | 117629773 | 117630185 | Distal Intergenic |
| chr2 | 117630608 | 117630912 | Distal Intergenic |
| chr2 | 117708180 | 117708444 | Distal Intergenic |
| chr2 | 117728796 | 117729176 | Distal Intergenic |
| chr2 | 117814240 | 117814552 | Distal Intergenic |
| chr2 | 117825336 | 117825573 | Distal Intergenic |
| chr2 | 117919865 | 117921333 | Distal Intergenic |
| chr2 | 117935194 | 117936853 | Distal Intergenic |
| chr2 | 118445582 | 118445929 | Distal Intergenic |
| chr2 | 118446088 | 118446426 | Distal Intergenic |
| chr2 | 118484258 | 118486280 | Distal Intergenic |
| chr2 | 118489222 | 118491178 | Distal Intergenic |
| chr2 | 118494447 | 118495532 | Distal Intergenic |
| chr2 | 118499428 | 118499882 | Distal Intergenic |
| chr2 | 118502294 | 118502667 | Distal Intergenic |
| chr2 | 118502957 | 118503750 | Distal Intergenic |
| chr2 | 118586673 | 118589442 | 3' UTR |
| chr2 | 118589602 | 118589830 | 3' UTR |
| chr2 | 118590000 | 118590326 | Distal Intergenic |
| chr2 | 118590441 | 118592232 | Distal Intergenic |
| chr2 | 118631142 | 118631342 | Distal Intergenic |
| chr2 | 118631399 | 118632564 | Distal Intergenic |
| chr2 | 118640083 | 118640335 | Distal Intergenic |
| chr2 | 118787607 | 118787855 | Distal Intergenic |
| chr2 | 118938870 | 118939296 | Distal Intergenic |
| chr2 | 118998886 | 118999329 | Distal Intergenic |
| chr2 | 118999542 | 119000818 | Distal Intergenic |
| chr2 | 119034081 | 119034301 | Distal Intergenic |
| chr2 | 119089070 | 119089277 | Distal Intergenic |
| chr2 | 119170032 | 119170934 | Distal Intergenic |
| chr2 | 119240889 | 119241536 | Distal Intergenic |
| chr2 | 119241842 | 119242041 | Distal Intergenic |
| chr2 | 119302481 | 119303827 | Distal Intergenic |

| | | | |
|---|---|---|---|
| chr2 | 119372927 | 119373139 | Distal Intergenic |
| chr2 | 119373796 | 119373998 | Distal Intergenic |
| chr2 | 119420401 | 119420680 | Distal Intergenic |
| chr2 | 119500868 | 119501067 | Distal Intergenic |
| chr2 | 119612893 | 119613226 | Distal Intergenic |
| chr2 | 119798736 | 119798935 | Distal Intergenic |
| chr2 | 119799234 | 119799545 | Distal Intergenic |
| chr2 | 119820926 | 119821290 | Distal Intergenic |
| chr2 | 119832945 | 119834339 | Distal Intergenic |
| chr2 | 119930476 | 119931740 | Distal Intergenic |
| chr2 | 119954111 | 119955064 | Distal Intergenic |
| chr2 | 119957225 | 119957710 | Distal Intergenic |
| chr2 | 119957843 | 119958210 | Distal Intergenic |
| chr2 | 119958441 | 119958646 | Distal Intergenic |
| chr2 | 119959366 | 119961348 | Distal Intergenic |
| chr2 | 119962184 | 119962622 | Distal Intergenic |
| chr2 | 119965625 | 119967881 | Distal Intergenic |
| chr2 | 119972617 | 119973143 | Distal Intergenic |
| chr2 | 120212943 | 120213429 | Intron |
| chr2 | 120229455 | 120229871 | Intron |
| chr2 | 120339480 | 120339796 | Intron |
| chr2 | 120434740 | 120435322 | Distal Intergenic |
| chr2 | 120557287 | 120557548 | Intron |
| chr2 | 121048709 | 121049438 | Intron |
| chr2 | 121212314 | 121212713 | Distal Intergenic |
| chr2 | 121238764 | 121239524 | Distal Intergenic |
| chr2 | 121239983 | 121240227 | Distal Intergenic |
| chr2 | 121282425 | 121283355 | Distal Intergenic |
| chr2 | 121285724 | 121286364 | Distal Intergenic |
| chr2 | 121446823 | 121447022 | Distal Intergenic |
| chr2 | 121454324 | 121454570 | Distal Intergenic |
| chr2 | 121782258 | 121782951 | Distal Intergenic |
| chr2 | 121990327 | 121990622 | Intron |
| chr2 | 122055395 | 122058139 | Distal Intergenic |
| chr2 | 122059844 | 122060071 | Distal Intergenic |
| chr2 | 122060284 | 122060542 | Distal Intergenic |
| chr2 | 122130873 | 122131091 | Intron |
| chr2 | 122131180 | 122132242 | Intron |
| chr2 | 122132283 | 122132685 | Intron |
| chr2 | 122151795 | 122152022 | Intron |
| chr2 | 122152068 | 122153349 | Intron |
| chr2 | 122153477 | 122153971 | Intron |
| chr2 | 122170143 | 122172825 | Intron |
| chr2 | 122174891 | 122175117 | Intron |

| | | | |
|---|---|---|---|
| chr2 | 122477818 | 122478625 | Intron |
| chr2 | 122523440 | 122523715 | 3' UTR |
| chr2 | 124684040 | 124684471 | Distal Intergenic |
| chr2 | 124684523 | 124684756 | Distal Intergenic |
| chr2 | 125564235 | 125564442 | Intron |
| chr2 | 125564661 | 125565702 | Intron |
| chr2 | 125617301 | 125617571 | Intron |
| chr2 | 125618095 | 125618366 | Intron |
| chr2 | 125618486 | 125618722 | Intron |
| chr2 | 125683966 | 125685220 | Distal Intergenic |
| chr2 | 126017820 | 126018266 | Distal Intergenic |
| chr2 | 126128448 | 126129476 | Distal Intergenic |
| chr2 | 126135085 | 126137483 | Distal Intergenic |
| chr2 | 126189077 | 126189345 | Distal Intergenic |
| chr2 | 126189894 | 126190331 | Distal Intergenic |
| chr2 | 126191372 | 126191661 | Distal Intergenic |
| chr2 | 126499663 | 126502520 | Distal Intergenic |
| chr2 | 126502787 | 126503017 | Distal Intergenic |
| chr2 | 126700536 | 126701441 | Distal Intergenic |
| chr2 | 126701534 | 126701870 | Distal Intergenic |
| chr2 | 126758092 | 126758458 | Distal Intergenic |
| chr2 | 126758621 | 126759597 | Distal Intergenic |
| chr2 | 126763654 | 126765201 | Distal Intergenic |
| chr2 | 126771504 | 126771789 | Distal Intergenic |
| chr2 | 126774002 | 126774235 | Distal Intergenic |
| chr2 | 126774280 | 126774747 | Distal Intergenic |
| chr2 | 126960037 | 126960301 | Distal Intergenic |
| chr2 | 126962865 | 126963244 | Distal Intergenic |
| chr2 | 126963387 | 126963630 | Distal Intergenic |
| chr2 | 126963793 | 126964062 | Distal Intergenic |
| chr2 | 126964229 | 126964892 | Distal Intergenic |
| chr2 | 126965086 | 126965303 | Distal Intergenic |
| chr2 | 127095620 | 127095876 | Distal Intergenic |
| chr2 | 127412938 | 127413196 | Promoter |
| chr2 | 127475109 | 127475786 | Distal Intergenic |
| chr2 | 127475897 | 127476276 | Distal Intergenic |
| chr2 | 127543901 | 127544322 | Distal Intergenic |
| chr2 | 127544428 | 127544979 | Distal Intergenic |
| chr2 | 127547704 | 127548607 | Distal Intergenic |
| chr2 | 127959775 | 127960389 | Intron |
| chr2 | 128253720 | 128255610 | Exon |
| chr2 | 128644263 | 128644529 | Promoter |
| chr2 | 128793064 | 128793537 | Distal Intergenic |
| chr2 | 128955503 | 128955909 | Distal Intergenic |

| | | | |
|---|---|---|---|
| chr2 | 128991877 | 128993921 | Distal Intergenic |
| chr2 | 129001322 | 129002042 | Distal Intergenic |
| chr2 | 129002186 | 129002436 | Distal Intergenic |
| chr2 | 129002480 | 129003139 | Distal Intergenic |
| chr2 | 129032414 | 129032766 | Intron |
| chr2 | 129068546 | 129068971 | Intron |
| chr2 | 129071618 | 129071904 | Intron |
| chr2 | 129072040 | 129072610 | Intron |
| chr2 | 129083895 | 129084114 | Distal Intergenic |
| chr2 | 129088785 | 129090379 | Distal Intergenic |
| chr2 | 129122635 | 129122910 | Distal Intergenic |
| chr2 | 129123050 | 129123810 | Distal Intergenic |
| chr2 | 129133671 | 129133939 | Distal Intergenic |
| chr2 | 129134513 | 129134958 | Distal Intergenic |
| chr2 | 129135529 | 129135817 | Distal Intergenic |
| chr2 | 129149748 | 129150744 | Distal Intergenic |
| chr2 | 129156214 | 129157061 | Distal Intergenic |
| chr2 | 129160241 | 129160734 | Distal Intergenic |
| chr2 | 129163764 | 129164449 | Distal Intergenic |
| chr2 | 129232311 | 129234831 | Distal Intergenic |
| chr2 | 129237322 | 129238256 | Distal Intergenic |
| chr2 | 129246527 | 129247142 | Distal Intergenic |
| chr2 | 129418180 | 129418693 | Distal Intergenic |
| chr2 | 129419081 | 129419342 | Distal Intergenic |
| chr2 | 129419395 | 129420064 | Distal Intergenic |
| chr2 | 129425201 | 129425630 | Distal Intergenic |
| chr2 | 129457915 | 129460705 | Distal Intergenic |
| chr2 | 129626392 | 129627874 | Distal Intergenic |
| chr2 | 129643033 | 129643245 | Distal Intergenic |
| chr2 | 129645363 | 129646354 | Distal Intergenic |
| chr2 | 129751430 | 129751701 | Distal Intergenic |
| chr2 | 129754229 | 129755734 | Distal Intergenic |
| chr2 | 129779934 | 129781722 | Distal Intergenic |
| chr2 | 129788910 | 129789190 | Distal Intergenic |
| chr2 | 129789294 | 129789796 | Distal Intergenic |
| chr2 | 129825035 | 129825351 | Distal Intergenic |
| chr2 | 129825409 | 129825993 | Distal Intergenic |
| chr2 | 129859893 | 129860541 | Distal Intergenic |
| chr2 | 130004413 | 130004703 | Intron |
| chr2 | 130099736 | 130100054 | Distal Intergenic |
| chr2 | 130250000 | 130250215 | Distal Intergenic |
| chr2 | 130455628 | 130455997 | Distal Intergenic |
| chr2 | 131025210 | 131026395 | Distal Intergenic |
| chr2 | 131028834 | 131031362 | Exon |

| | | | |
|---|---|---|---|
| chr2 | 131033882 | 131034711 | Distal Intergenic |
| chr2 | 131034901 | 131035198 | Distal Intergenic |
| chr2 | 131035399 | 131036349 | Exon |
| chr2 | 131088475 | 131089284 | Distal Intergenic |
| chr2 | 131218475 | 131218749 | Downstream |
| chr2 | 131229322 | 131229691 | Intron |
| chr2 | 131229846 | 131230504 | Intron |
| chr2 | 131335591 | 131336444 | Exon |
| chr2 | 131372862 | 131373074 | Intron |
| chr2 | 131586810 | 131587093 | Distal Intergenic |
| chr2 | 131587177 | 131587881 | Distal Intergenic |
| chr2 | 131727629 | 131727835 | Intron |
| chr2 | 131727899 | 131728372 | Intron |
| chr2 | 131728577 | 131729285 | Intron |
| chr2 | 131757518 | 131759220 | Intron |
| chr2 | 131770494 | 131771378 | Intron |
| chr2 | 131774738 | 131776823 | Intron |
| chr2 | 131776991 | 131777258 | Intron |
| chr2 | 131829941 | 131832974 | Intron |
| chr2 | 131908057 | 131911336 | Intron |
| chr2 | 131923102 | 131923356 | Intron |
| chr2 | 131923407 | 131923857 | Intron |
| chr2 | 131924059 | 131924261 | Intron |
| chr2 | 131924462 | 131925364 | Intron |
| chr2 | 131936431 | 131937208 | Intron |
| chr2 | 131978427 | 131979211 | Intron |
| chr2 | 131994565 | 131995485 | Intron |
| chr2 | 132073242 | 132073511 | Intron |
| chr2 | 132073919 | 132074208 | Intron |
| chr2 | 132165728 | 132166225 | Exon |
| chr2 | 132370475 | 132370710 | Distal Intergenic |
| chr2 | 132404170 | 132404511 | Distal Intergenic |
| chr2 | 132433627 | 132433954 | Distal Intergenic |
| chr2 | 132438961 | 132439768 | Distal Intergenic |
| chr2 | 132769377 | 132769654 | Distal Intergenic |
| chr2 | 132769751 | 132770314 | Distal Intergenic |
| chr2 | 132770363 | 132771256 | Distal Intergenic |
| chr2 | 132771349 | 132772579 | Distal Intergenic |
| chr2 | 132777441 | 132778103 | Distal Intergenic |
| chr2 | 133542165 | 133542369 | Exon |
| chr2 | 133916878 | 133917368 | Intron |
| chr2 | 135350984 | 135351634 | Intron |
| chr2 | 135441448 | 135442297 | Intron |
| chr2 | 135563207 | 135563471 | Distal Intergenic |

| | | | |
|---|---|---|---|
| chr2 | 135827346 | 135828065 | Intron |
| chr2 | 136717821 | 136718312 | Intron |
| chr2 | 136743721 | 136746844 | Promoter |
| chr2 | 136890745 | 136890980 | Distal Intergenic |
| chr2 | 136893290 | 136893601 | Distal Intergenic |
| chr2 | 136933386 | 136935666 | Distal Intergenic |
| chr2 | 136935703 | 136937947 | Distal Intergenic |
| chr2 | 136938092 | 136938508 | Distal Intergenic |
| chr2 | 136938552 | 136938802 | Distal Intergenic |
| chr2 | 136939407 | 136940346 | Distal Intergenic |
| chr2 | 136940628 | 136940926 | Distal Intergenic |
| chr2 | 136941011 | 136942523 | Distal Intergenic |
| chr2 | 136942604 | 136943193 | Distal Intergenic |
| chr2 | 136974338 | 136975495 | Distal Intergenic |
| chr2 | 136975605 | 136975874 | Distal Intergenic |
| chr2 | 136976015 | 136977275 | Distal Intergenic |
| chr2 | 137050096 | 137051809 | Distal Intergenic |
| chr2 | 137088066 | 137089206 | Distal Intergenic |
| chr2 | 137199272 | 137199495 | Distal Intergenic |
| chr2 | 137234730 | 137236248 | Distal Intergenic |
| chr2 | 137238534 | 137238855 | Distal Intergenic |
| chr2 | 137238976 | 137239831 | Distal Intergenic |
| chr2 | 137255896 | 137257181 | Distal Intergenic |
| chr2 | 137280721 | 137283111 | Distal Intergenic |
| chr2 | 137284008 | 137284208 | Distal Intergenic |
| chr2 | 137284316 | 137284709 | Distal Intergenic |
| chr2 | 137319877 | 137321221 | Distal Intergenic |
| chr2 | 137391877 | 137394561 | Distal Intergenic |
| chr2 | 137398852 | 137401237 | Distal Intergenic |
| chr2 | 137419720 | 137424749 | Distal Intergenic |
| chr2 | 137430524 | 137432979 | Distal Intergenic |
| chr2 | 137479024 | 137481321 | Distal Intergenic |
| chr2 | 137536414 | 137536680 | Intron |
| chr2 | 137541586 | 137542175 | Intron |
| chr2 | 137545840 | 137546698 | Intron |
| chr2 | 137546909 | 137547141 | Intron |
| chr2 | 137547346 | 137548391 | Intron |
| chr2 | 137548537 | 137549126 | Intron |
| chr2 | 137550594 | 137550856 | Intron |
| chr2 | 137551022 | 137551275 | Intron |
| chr2 | 137551457 | 137551890 | Intron |
| chr2 | 137633242 | 137633603 | Intron |
| chr2 | 137702463 | 137703479 | Intron |
| chr2 | 138049312 | 138049757 | Intron |

| | | | |
|---|---|---|---|
| chr2 | 138145873 | 138147921 | Intron |
| chr2 | 138257547 | 138259159 | Intron |
| chr2 | 138303109 | 138303394 | Intron |
| chr2 | 138303548 | 138304053 | Intron |
| chr2 | 138329851 | 138330050 | Exon |
| chr2 | 139307965 | 139308716 | Exon |
| chr2 | 139311186 | 139313033 | Intron |
| chr2 | 140494745 | 140494971 | Distal Intergenic |
| chr2 | 140495023 | 140495289 | Distal Intergenic |
| chr2 | 140496091 | 140497086 | Distal Intergenic |
| chr2 | 140616005 | 140616960 | Distal Intergenic |
| chr2 | 140662448 | 140664362 | Distal Intergenic |
| chr2 | 140664534 | 140664785 | Distal Intergenic |
| chr2 | 140673155 | 140674145 | Distal Intergenic |
| chr2 | 140730479 | 140730685 | Distal Intergenic |
| chr2 | 140730840 | 140731070 | Distal Intergenic |
| chr2 | 140731336 | 140731537 | Distal Intergenic |
| chr2 | 141091128 | 141093466 | Exon |
| chr2 | 141096713 | 141098385 | Intron |
| chr2 | 141099557 | 141102736 | Intron |
| chr2 | 141103441 | 141103664 | Intron |
| chr2 | 141103969 | 141104255 | Intron |
| chr2 | 141104327 | 141106362 | Intron |
| chr2 | 141273823 | 141275473 | Exon |
| chr2 | 141289471 | 141290733 | Intron |
| chr2 | 142464814 | 142465077 | Intron |
| chr2 | 142521539 | 142522222 | Intron |
| chr2 | 142889132 | 142889333 | Promoter |
| chr2 | 143678461 | 143678818 | Intron |
| chr2 | 143678965 | 143679322 | Intron |
| chr2 | 143729107 | 143731244 | Intron |
| chr2 | 143734697 | 143735439 | Intron |
| chr2 | 143739900 | 143740887 | Intron |
| chr2 | 143741069 | 143741410 | Intron |
| chr2 | 143741582 | 143741870 | Intron |
| chr2 | 143755293 | 143755761 | Intron |
| chr2 | 144094357 | 144094595 | Intron |
| chr2 | 144095774 | 144096262 | Intron |
| chr2 | 144381675 | 144381875 | Exon |
| chr2 | 144565704 | 144566014 | Distal Intergenic |
| chr2 | 144566058 | 144567221 | Distal Intergenic |
| chr2 | 144772052 | 144773165 | Intron |
| chr2 | 145044091 | 145045644 | Intron |
| chr2 | 145099095 | 145101286 | Distal Intergenic |

| | | | |
|---|---|---|---|
| chr2 | 145109625 | 145109875 | Distal Intergenic |
| chr2 | 145147246 | 145147449 | Exon |
| chr2 | 145163587 | 145165142 | Intron |
| chr2 | 145187374 | 145188952 | Exon |
| chr2 | 145190387 | 145190626 | Intron |
| chr2 | 145195932 | 145196831 | Intron |
| chr2 | 145656035 | 145656352 | Intron |
| chr2 | 145668984 | 145669782 | Intron |
| chr2 | 145671928 | 145673022 | Intron |
| chr2 | 145674320 | 145675491 | Intron |
| chr2 | 145686125 | 145686324 | Intron |
| chr2 | 145686420 | 145687175 | Intron |
| chr2 | 145687221 | 145687460 | Intron |
| chr2 | 145687968 | 145688508 | Intron |
| chr2 | 145691274 | 145692899 | Intron |
| chr2 | 145729414 | 145729622 | Intron |
| chr2 | 146114279 | 146114629 | Distal Intergenic |
| chr2 | 146312843 | 146313941 | Distal Intergenic |
| chr2 | 146359033 | 146359311 | Distal Intergenic |
| chr2 | 146379252 | 146379576 | Distal Intergenic |
| chr2 | 146379769 | 146380038 | Distal Intergenic |
| chr2 | 146396876 | 146397937 | Distal Intergenic |
| chr2 | 146400132 | 146401002 | Distal Intergenic |
| chr2 | 146802503 | 146803166 | Distal Intergenic |
| chr2 | 146811708 | 146811907 | Distal Intergenic |
| chr2 | 146838278 | 146840051 | Distal Intergenic |
| chr2 | 148006269 | 148006518 | Distal Intergenic |
| chr2 | 148010032 | 148011515 | Distal Intergenic |
| chr2 | 148265356 | 148266119 | Distal Intergenic |
| chr2 | 148281400 | 148281788 | Distal Intergenic |
| chr2 | 148294324 | 148295287 | Distal Intergenic |
| chr2 | 148295383 | 148296112 | Distal Intergenic |
| chr2 | 148332688 | 148333613 | Distal Intergenic |
| chr2 | 148347077 | 148347344 | Distal Intergenic |
| chr2 | 148347680 | 148348540 | Distal Intergenic |
| chr2 | 148350525 | 148351709 | Distal Intergenic |
| chr2 | 148351826 | 148352216 | Distal Intergenic |
| chr2 | 148456711 | 148457418 | Distal Intergenic |
| chr2 | 148504826 | 148505161 | Distal Intergenic |
| chr2 | 148505453 | 148505665 | Distal Intergenic |
| chr2 | 148505978 | 148506644 | Distal Intergenic |
| chr2 | 148569644 | 148569888 | Distal Intergenic |
| chr2 | 148570123 | 148570372 | Distal Intergenic |
| chr2 | 148698658 | 148698938 | Intron |

| | | | |
|---|---|---|---|
| chr2 | 149363857 | 149364159 | Distal Intergenic |
| chr2 | 149399379 | 149399969 | Distal Intergenic |
| chr2 | 149689652 | 149689865 | Intron |
| chr2 | 149712467 | 149712735 | Intron |
| chr2 | 149823984 | 149824248 | Intron |
| chr2 | 149824402 | 149824608 | Intron |
| chr2 | 150665341 | 150665714 | Distal Intergenic |
| chr2 | 150665754 | 150666162 | Distal Intergenic |
| chr2 | 150666208 | 150666671 | Distal Intergenic |
| chr2 | 150667065 | 150668024 | Distal Intergenic |
| chr2 | 150730660 | 150730861 | Distal Intergenic |
| chr2 | 151798691 | 151799408 | Distal Intergenic |
| chr2 | 151799848 | 151800051 | Distal Intergenic |
| chr2 | 151803307 | 151804497 | Distal Intergenic |
| chr2 | 151804881 | 151805097 | Distal Intergenic |
| chr2 | 151805176 | 151805558 | Distal Intergenic |
| chr2 | 151931242 | 151931495 | Distal Intergenic |
| chr2 | 152052288 | 152052512 | Distal Intergenic |
| chr2 | 152059691 | 152060053 | Distal Intergenic |
| chr2 | 152065934 | 152066250 | Distal Intergenic |
| chr2 | 152085029 | 152085577 | Distal Intergenic |
| chr2 | 152085690 | 152085889 | Distal Intergenic |
| chr2 | 152109250 | 152109468 | Exon |
| chr2 | 152157552 | 152157800 | Distal Intergenic |
| chr2 | 152522413 | 152522819 | Exon |
| chr2 | 152525435 | 152526038 | Exon |
| chr2 | 152526235 | 152526544 | Intron |
| chr2 | 152526692 | 152526953 | Intron |
| chr2 | 152552690 | 152553065 | Intron |
| chr2 | 152560166 | 152560683 | Intron |
| chr2 | 152561055 | 152562585 | Promoter |
| chr2 | 152562637 | 152565091 | Promoter |
| chr2 | 152565167 | 152566120 | Intron |
| chr2 | 152566842 | 152567936 | Exon |
| chr2 | 152570136 | 152570581 | Intron |
| chr2 | 152612692 | 152614546 | Distal Intergenic |
| chr2 | 152618153 | 152618424 | Distal Intergenic |
| chr2 | 152618598 | 152619103 | Distal Intergenic |
| chr2 | 152635685 | 152636099 | Distal Intergenic |
| chr2 | 152636142 | 152637205 | Distal Intergenic |
| chr2 | 152646402 | 152648009 | Intron |
| chr2 | 152655851 | 152656050 | Intron |
| chr2 | 152656289 | 152657011 | Intron |
| chr2 | 152673895 | 152674147 | Intron |

| | | | |
|---|---|---|---|
| chr2 | 152714062 | 152714534 | Intron |
| chr2 | 152714694 | 152714976 | Intron |
| chr2 | 152715012 | 152716740 | Intron |
| chr2 | 152987046 | 152987563 | Intron |
| chr2 | 153168730 | 153168929 | Distal Intergenic |
| chr2 | 153172521 | 153172760 | Distal Intergenic |
| chr2 | 153567482 | 153567706 | Intron |
| chr2 | 153575978 | 153576279 | Promoter |
| chr2 | 154007729 | 154007954 | Distal Intergenic |
| chr2 | 154008596 | 154008947 | Distal Intergenic |
| chr2 | 154163879 | 154164239 | Distal Intergenic |
| chr2 | 154164475 | 154164896 | Distal Intergenic |
| chr2 | 154262364 | 154262630 | Distal Intergenic |
| chr2 | 154289364 | 154289738 | Distal Intergenic |
| chr2 | 154482791 | 154483773 | Distal Intergenic |
| chr2 | 154483957 | 154484162 | Distal Intergenic |
| chr2 | 154494816 | 154495274 | Distal Intergenic |
| chr2 | 154604800 | 154605051 | Distal Intergenic |
| chr2 | 154616856 | 154617055 | Distal Intergenic |
| chr2 | 154717361 | 154717668 | Distal Intergenic |
| chr2 | 154717817 | 154718635 | Distal Intergenic |
| chr2 | 154899650 | 154899926 | Intron |
| chr2 | 154899993 | 154900193 | Intron |
| chr2 | 154900393 | 154900660 | Intron |
| chr2 | 155062362 | 155062669 | Intron |
| chr2 | 155417824 | 155418126 | Distal Intergenic |
| chr2 | 155576441 | 155576703 | Intron |
| chr2 | 155629694 | 155630038 | Intron |
| chr2 | 155630306 | 155633179 | Intron |
| chr2 | 155634891 | 155635742 | Intron |
| chr2 | 155643933 | 155644356 | Intron |
| chr2 | 155644451 | 155644928 | Intron |
| chr2 | 155644976 | 155647597 | Intron |
| chr2 | 155650771 | 155652506 | Intron |
| chr2 | 155699801 | 155700227 | Intron |
| chr2 | 155704415 | 155705119 | Intron |
| chr2 | 155755864 | 155756183 | Distal Intergenic |
| chr2 | 155804608 | 155805506 | Distal Intergenic |
| chr2 | 155936710 | 155937322 | Distal Intergenic |
| chr2 | 155937441 | 155937643 | Distal Intergenic |
| chr2 | 155970017 | 155970673 | Distal Intergenic |
| chr2 | 156004932 | 156005357 | Distal Intergenic |
| chr2 | 156005507 | 156005960 | Distal Intergenic |
| chr2 | 156058858 | 156059522 | Distal Intergenic |

| | | | |
|---|---|---|---|
| chr2 | 156138088 | 156138561 | Distal Intergenic |
| chr2 | 156217823 | 156218053 | Distal Intergenic |
| chr2 | 156218090 | 156218656 | Distal Intergenic |
| chr2 | 156272808 | 156273102 | Distal Intergenic |
| chr2 | 156282314 | 156282531 | Distal Intergenic |
| chr2 | 156282707 | 156283127 | Distal Intergenic |
| chr2 | 156364743 | 156365054 | Distal Intergenic |
| chr2 | 156365207 | 156365457 | Distal Intergenic |
| chr2 | 156547164 | 156547615 | Distal Intergenic |
| chr2 | 156547838 | 156548152 | Distal Intergenic |
| chr2 | 156667838 | 156668197 | Distal Intergenic |
| chr2 | 156670329 | 156670570 | Distal Intergenic |
| chr2 | 156670620 | 156670851 | Distal Intergenic |
| chr2 | 156688150 | 156688399 | Distal Intergenic |
| chr2 | 156689263 | 156689732 | Distal Intergenic |
| chr2 | 156871582 | 156871944 | Distal Intergenic |
| chr2 | 156897424 | 156898974 | Intron |
| chr2 | 156899141 | 156899362 | Intron |
| chr2 | 156899718 | 156900790 | Intron |
| chr2 | 157028009 | 157030545 | Intron |
| chr2 | 157193679 | 157194071 | Exon |
| chr2 | 157201894 | 157202375 | Distal Intergenic |
| chr2 | 157218038 | 157218274 | Distal Intergenic |
| chr2 | 157218347 | 157218809 | Distal Intergenic |
| chr2 | 157236355 | 157239112 | Distal Intergenic |
| chr2 | 157248612 | 157250222 | Distal Intergenic |
| chr2 | 157250300 | 157252843 | Distal Intergenic |
| chr2 | 157253244 | 157253581 | Distal Intergenic |
| chr2 | 157253662 | 157253873 | Distal Intergenic |
| chr2 | 157254044 | 157254539 | Distal Intergenic |
| chr2 | 157255131 | 157256526 | Distal Intergenic |
| chr2 | 157259002 | 157259211 | Distal Intergenic |
| chr2 | 157259361 | 157260363 | Distal Intergenic |
| chr2 | 157260415 | 157260789 | Distal Intergenic |
| chr2 | 157271399 | 157273233 | Distal Intergenic |
| chr2 | 157273409 | 157274570 | Distal Intergenic |
| chr2 | 157275650 | 157275892 | Distal Intergenic |
| chr2 | 157276518 | 157278178 | Distal Intergenic |
| chr2 | 157289764 | 157290008 | Distal Intergenic |
| chr2 | 157297185 | 157297569 | Intron |
| chr2 | 157298599 | 157299243 | Intron |
| chr2 | 157299282 | 157299544 | Intron |
| chr2 | 157391735 | 157393738 | Intron |
| chr2 | 157633348 | 157634409 | Distal Intergenic |

| | | | |
|---|---|---|---|
| chr2 | 157636225 | 157636648 | Distal Intergenic |
| chr2 | 157639987 | 157640720 | Distal Intergenic |
| chr2 | 157671266 | 157672003 | Distal Intergenic |
| chr2 | 157853809 | 157854133 | Distal Intergenic |
| chr2 | 157854175 | 157854394 | Distal Intergenic |
| chr2 | 157854778 | 157855204 | Distal Intergenic |
| chr2 | 157958541 | 157959075 | Distal Intergenic |
| chr2 | 157980498 | 157980842 | Distal Intergenic |
| chr2 | 157985569 | 157985987 | Distal Intergenic |
| chr2 | 157993356 | 157993766 | Distal Intergenic |
| chr2 | 157993825 | 157994676 | Distal Intergenic |
| chr2 | 157994720 | 157995460 | Distal Intergenic |
| chr2 | 158186992 | 158187252 | Distal Intergenic |
| chr2 | 158187380 | 158187602 | Distal Intergenic |
| chr2 | 158241753 | 158242073 | Distal Intergenic |
| chr2 | 158336237 | 158336901 | Intron |
| chr2 | 158658979 | 158660532 | Intron |
| chr2 | 158665619 | 158666196 | Intron |
| chr2 | 158666356 | 158666839 | Intron |
| chr2 | 158919151 | 158919679 | Intron |
| chr2 | 159973514 | 159975266 | Intron |
| chr2 | 159975462 | 159976062 | Intron |
| chr2 | 160120907 | 160121106 | Intron |
| chr2 | 160139019 | 160139373 | Exon |
| chr2 | 160171580 | 160172253 | Distal Intergenic |
| chr2 | 160445694 | 160446761 | Intron |
| chr2 | 160446990 | 160447513 | Intron |
| chr2 | 160456416 | 160456631 | Intron |
| chr2 | 160456719 | 160457575 | Intron |
| chr2 | 160472119 | 160472416 | Promoter |
| chr2 | 160472468 | 160473081 | Promoter |
| chr2 | 160571169 | 160571397 | Intron |
| chr2 | 160571444 | 160573612 | 5' UTR |
| chr2 | 160713486 | 160713722 | Intron |
| chr2 | 160788876 | 160789119 | Distal Intergenic |
| chr2 | 160937479 | 160937728 | Distal Intergenic |
| chr2 | 162098164 | 162099262 | Distal Intergenic |
| chr2 | 162099310 | 162099710 | Distal Intergenic |
| chr2 | 162108243 | 162108707 | Distal Intergenic |
| chr2 | 162108800 | 162109037 | Distal Intergenic |
| chr2 | 162766600 | 162766961 | Intron |
| chr2 | 162900749 | 162901555 | Intron |
| chr2 | 163239090 | 163239594 | Intron |
| chr2 | 163561887 | 163562269 | Intron |

| | | | |
|---|---|---|---|
| chr2 | 163821288 | 163821627 | Distal Intergenic |
| chr2 | 163842583 | 163843812 | Distal Intergenic |
| chr2 | 164182139 | 164182386 | Distal Intergenic |
| chr2 | 164247027 | 164248015 | Distal Intergenic |
| chr2 | 164256907 | 164259439 | Distal Intergenic |
| chr2 | 164260529 | 164261270 | Distal Intergenic |
| chr2 | 164264359 | 164264591 | Distal Intergenic |
| chr2 | 164264903 | 164265190 | Distal Intergenic |
| chr2 | 164344308 | 164344645 | Distal Intergenic |
| chr2 | 164344849 | 164345352 | Distal Intergenic |
| chr2 | 164347166 | 164348167 | Distal Intergenic |
| chr2 | 164384388 | 164385157 | Distal Intergenic |
| chr2 | 164499379 | 164500305 | Intron |
| chr2 | 164500412 | 164500636 | Intron |
| chr2 | 164561243 | 164561766 | Intron |
| chr2 | 164562350 | 164564136 | Intron |
| chr2 | 164610769 | 164611087 | Distal Intergenic |
| chr2 | 164682261 | 164682761 | Distal Intergenic |
| chr2 | 164683217 | 164683620 | Distal Intergenic |
| chr2 | 164736679 | 164737813 | Distal Intergenic |
| chr2 | 165147290 | 165151625 | Distal Intergenic |
| chr2 | 165645745 | 165646328 | Intron |
| chr2 | 165646477 | 165646738 | Intron |
| chr2 | 165656270 | 165658697 | Exon |
| chr2 | 165681104 | 165683916 | Intron |
| chr2 | 165684033 | 165684873 | Intron |
| chr2 | 165685053 | 165685252 | Intron |
| chr2 | 165709145 | 165710209 | Distal Intergenic |
| chr2 | 165710285 | 165710589 | Distal Intergenic |
| chr2 | 165719960 | 165720270 | Distal Intergenic |
| chr2 | 165979947 | 165980161 | Intron |
| chr2 | 165995379 | 165995848 | Intron |
| chr2 | 166255032 | 166255293 | Distal Intergenic |
| chr2 | 166261059 | 166261753 | Distal Intergenic |
| chr2 | 166731954 | 166732387 | Intron |
| chr2 | 166889107 | 166889590 | Intron |
| chr2 | 166890014 | 166890674 | Intron |
| chr2 | 166891374 | 166891681 | Intron |
| chr2 | 166892046 | 166892297 | Intron |
| chr2 | 167246468 | 167247106 | Distal Intergenic |
| chr2 | 167250331 | 167251339 | Distal Intergenic |
| chr2 | 167311221 | 167312129 | Intron |
| chr2 | 167395571 | 167396283 | Distal Intergenic |
| chr2 | 167600049 | 167601147 | Distal Intergenic |

| | | | |
|---|---|---|---|
| chr2 | 167602381 | 167604013 | Distal Intergenic |
| chr2 | 167614475 | 167615980 | Distal Intergenic |
| chr2 | 167623718 | 167623925 | Distal Intergenic |
| chr2 | 167624581 | 167624789 | Distal Intergenic |
| chr2 | 167625038 | 167625415 | Distal Intergenic |
| chr2 | 167626118 | 167626729 | Distal Intergenic |
| chr2 | 167642815 | 167643194 | Distal Intergenic |
| chr2 | 167656555 | 167656867 | Distal Intergenic |
| chr2 | 167659220 | 167659670 | Distal Intergenic |
| chr2 | 167659983 | 167660557 | Distal Intergenic |
| chr2 | 167665002 | 167665279 | Distal Intergenic |
| chr2 | 167744260 | 167745046 | Promoter |
| chr2 | 167745177 | 167746153 | Promoter |
| chr2 | 167774492 | 167775010 | Intron |
| chr2 | 167775064 | 167777175 | Intron |
| chr2 | 167786623 | 167786922 | Intron |
| chr2 | 168017054 | 168018773 | Intron |
| chr2 | 168031197 | 168031910 | Intron |
| chr2 | 168032374 | 168032981 | Intron |
| chr2 | 168055877 | 168056115 | Intron |
| chr2 | 168283199 | 168283468 | Distal Intergenic |
| chr2 | 168283722 | 168284274 | Distal Intergenic |
| chr2 | 168371296 | 168372070 | Distal Intergenic |
| chr2 | 168373563 | 168373971 | Distal Intergenic |
| chr2 | 168374335 | 168375012 | Distal Intergenic |
| chr2 | 168745245 | 168745485 | Distal Intergenic |
| chr2 | 168745701 | 168745983 | Distal Intergenic |
| chr2 | 168780366 | 168782130 | Distal Intergenic |
| chr2 | 168783770 | 168785258 | Distal Intergenic |
| chr2 | 169414822 | 169416283 | Intron |
| chr2 | 169979433 | 169979654 | Distal Intergenic |
| chr2 | 169979713 | 169980223 | Distal Intergenic |
| chr2 | 169980361 | 169981364 | Downstream |
| chr2 | 170071258 | 170071877 | Intron |
| chr2 | 170071930 | 170072324 | Intron |
| chr2 | 170073232 | 170073816 | Intron |
| chr2 | 170133196 | 170135217 | Exon |
| chr2 | 170135509 | 170136362 | Exon |
| chr2 | 172469298 | 172470121 | Distal Intergenic |
| chr2 | 172686801 | 172687570 | Intron |
| chr2 | 172688918 | 172689527 | Intron |
| chr2 | 172762833 | 172763093 | Distal Intergenic |
| chr2 | 172763137 | 172763371 | Distal Intergenic |
| chr2 | 172763595 | 172763807 | Distal Intergenic |

| | | | |
|---|---|---|---|
| chr2 | 173057302 | 173059459 | Distal Intergenic |
| chr2 | 173071222 | 173071638 | Distal Intergenic |
| chr2 | 173076320 | 173076665 | Distal Intergenic |
| chr2 | 173082668 | 173084890 | Distal Intergenic |
| chr2 | 173086193 | 173094579 | Distal Intergenic |
| chr2 | 173099082 | 173099303 | Distal Intergenic |
| chr2 | 173099815 | 173101214 | Distal Intergenic |
| chr2 | 173106380 | 173107156 | Distal Intergenic |
| chr2 | 173485817 | 173486161 | Exon |
| chr2 | 173487798 | 173488549 | Intron |
| chr2 | 173505717 | 173506098 | Distal Intergenic |
| chr2 | 174155921 | 174156340 | Distal Intergenic |
| chr2 | 179905062 | 179905743 | Intron |
| chr2 | 179970097 | 179972457 | 3' UTR |
| chr2 | 180192662 | 180192911 | Distal Intergenic |
| chr2 | 180315890 | 180316211 | Intron |
| chr2 | 180316980 | 180317189 | Intron |
| chr2 | 180436143 | 180437534 | Intron |
| chr2 | 180444298 | 180444714 | Intron |
| chr2 | 180922842 | 180923463 | Distal Intergenic |
| chr2 | 180923627 | 180923884 | Distal Intergenic |
| chr2 | 181021849 | 181022390 | Distal Intergenic |
| chr2 | 181232239 | 181232448 | Distal Intergenic |
| chr2 | 181232577 | 181232793 | Distal Intergenic |
| chr2 | 181254476 | 181255121 | Distal Intergenic |
| chr2 | 181255391 | 181255922 | Distal Intergenic |
| chr2 | 181257351 | 181257949 | Distal Intergenic |
| chr2 | 181259436 | 181260869 | Distal Intergenic |
| chr2 | 181316010 | 181316209 | Distal Intergenic |
| chr3 | 5575337 | 5575908 | Distal Intergenic |
| chr3 | 7503080 | 7503338 | 5' UTR |
| chr3 | 7503460 | 7503814 | Intron |
| chr3 | 7512556 | 7514272 | Intron |
| chr3 | 8053246 | 8054390 | Intron |
| chr3 | 8084132 | 8084447 | Distal Intergenic |
| chr3 | 8084517 | 8085108 | Distal Intergenic |
| chr3 | 8085145 | 8085372 | Distal Intergenic |
| chr3 | 8307021 | 8307529 | Intron |
| chr3 | 8440870 | 8441583 | Intron |
| chr3 | 8919227 | 8920311 | 3' UTR |
| chr3 | 8988147 | 8989338 | Exon |
| chr3 | 8989647 | 8989961 | Intron |
| chr3 | 8993444 | 8994055 | Intron |
| chr3 | 9001157 | 9002288 | Intron |

| | | | |
|---|---|---|---|
| chr3 | 9003713 | 9004021 | Intron |
| chr3 | 9014766 | 9014999 | Distal Intergenic |
| chr3 | 9015122 | 9015422 | Distal Intergenic |
| chr3 | 10258177 | 10259909 | Exon |
| chr3 | 10263042 | 10265910 | Exon |
| chr3 | 10273006 | 10275160 | Intron |
| chr3 | 10276390 | 10280434 | Exon |
| chr3 | 10292223 | 10293637 | Intron |
| chr3 | 10294007 | 10294879 | Intron |
| chr3 | 14610841 | 14611261 | Distal Intergenic |
| chr3 | 14641887 | 14642442 | Distal Intergenic |
| chr3 | 14647764 | 14649892 | Distal Intergenic |
| chr3 | 14659313 | 14660616 | Distal Intergenic |
| chr3 | 14947635 | 14948830 | Intron |
| chr3 | 15007595 | 15008431 | Intron |
| chr3 | 15452378 | 15452715 | 3' UTR |
| chr3 | 16083224 | 16084841 | Intron |
| chr3 | 18113621 | 18114138 | Intron |
| chr3 | 18115014 | 18115245 | Intron |
| chr3 | 18115449 | 18116347 | Intron |
| chr3 | 18239546 | 18239746 | Intron |
| chr3 | 18275804 | 18277625 | Intron |
| chr3 | 19081494 | 19083031 | Distal Intergenic |
| chr3 | 19087390 | 19088171 | Distal Intergenic |
| chr3 | 19088223 | 19088594 | Distal Intergenic |
| chr3 | 19116060 | 19116396 | Distal Intergenic |
| chr3 | 19145409 | 19145669 | Distal Intergenic |
| chr3 | 19145802 | 19146166 | Distal Intergenic |
| chr3 | 19146451 | 19146657 | Distal Intergenic |
| chr3 | 19146708 | 19147955 | Distal Intergenic |
| chr3 | 19161990 | 19165649 | Distal Intergenic |
| chr3 | 19165746 | 19166095 | Distal Intergenic |
| chr3 | 19166305 | 19166663 | Distal Intergenic |
| chr3 | 19176550 | 19177576 | Distal Intergenic |
| chr3 | 19208324 | 19208549 | Intron |
| chr3 | 19208684 | 19211279 | Intron |
| chr3 | 19269275 | 19269774 | Intron |
| chr3 | 19305689 | 19308994 | Intron |
| chr3 | 19310924 | 19311532 | Intron |
| chr3 | 19324792 | 19326457 | Intron |
| chr3 | 19326502 | 19327054 | Intron |
| chr3 | 19327984 | 19330758 | Intron |
| chr3 | 19351336 | 19351648 | Intron |
| chr3 | 19356051 | 19357333 | Promoter |

| | | | |
|---|---|---|---|
| chr3 | 19409073 | 19409339 | Intron |
| chr3 | 19418999 | 19420747 | Intron |
| chr3 | 19957305 | 19957539 | Intron |
| chr3 | 21299644 | 21301054 | Distal Intergenic |
| chr3 | 21302159 | 21302994 | Distal Intergenic |
| chr3 | 21303055 | 21303959 | Distal Intergenic |
| chr3 | 21304135 | 21304515 | Distal Intergenic |
| chr3 | 21536758 | 21536957 | Intron |
| chr3 | 21566483 | 21567339 | Intron |
| chr3 | 21570031 | 21571285 | Intron |
| chr3 | 21571330 | 21572185 | Intron |
| chr3 | 21578281 | 21578554 | Intron |
| chr3 | 21598207 | 21598555 | Intron |
| chr3 | 21628143 | 21628711 | Intron |
| chr3 | 21659793 | 21662026 | Intron |
| chr3 | 21739787 | 21740216 | Intron |
| chr3 | 21804554 | 21805127 | Intron |
| chr3 | 21836883 | 21837777 | Intron |
| chr3 | 21976235 | 21976449 | Intron |
| chr3 | 21976781 | 21977001 | Intron |
| chr3 | 23769248 | 23769699 | Distal Intergenic |
| chr3 | 24503596 | 24503844 | Intron |
| chr3 | 24593494 | 24593801 | Distal Intergenic |
| chr3 | 27001986 | 27002185 | Distal Intergenic |
| chr3 | 27002419 | 27003105 | Distal Intergenic |
| chr3 | 27051805 | 27052219 | Distal Intergenic |
| chr3 | 27052530 | 27052788 | Distal Intergenic |
| chr3 | 27052903 | 27054450 | Distal Intergenic |
| chr3 | 27108179 | 27109944 | Distal Intergenic |
| chr3 | 27487182 | 27487896 | Intron |
| chr3 | 27489124 | 27490001 | Intron |
| chr3 | 27490054 | 27490378 | Exon |
| chr3 | 27515346 | 27515941 | Intron |
| chr3 | 27550374 | 27553181 | Distal Intergenic |
| chr3 | 27555585 | 27556040 | Distal Intergenic |
| chr3 | 27556304 | 27556572 | Distal Intergenic |
| chr3 | 27608380 | 27608883 | Distal Intergenic |
| chr3 | 27609446 | 27609846 | Distal Intergenic |
| chr3 | 27762198 | 27762397 | Intron |
| chr3 | 27881280 | 27882256 | Distal Intergenic |
| chr3 | 27882541 | 27883180 | Distal Intergenic |
| chr3 | 27887464 | 27888084 | Distal Intergenic |
| chr3 | 27907008 | 27908347 | Distal Intergenic |
| chr3 | 27920755 | 27921225 | Distal Intergenic |

| | | | |
|---|---|---|---|
| chr3 | 28088487 | 28088743 | Distal Intergenic |
| chr3 | 28366562 | 28367025 | Intron |
| chr3 | 28633413 | 28634244 | Intron |
| chr3 | 28704379 | 28704779 | Intron |
| chr3 | 28705151 | 28705564 | Intron |
| chr3 | 28705661 | 28706073 | Intron |
| chr3 | 29778379 | 29778812 | Intron |
| chr3 | 29778891 | 29779404 | Intron |
| chr3 | 29788558 | 29790752 | Intron |
| chr3 | 29802189 | 29803513 | Intron |
| chr3 | 29852384 | 29852631 | Intron |
| chr3 | 30501529 | 30501956 | Distal Intergenic |
| chr3 | 30691794 | 30692012 | 5' UTR |
| chr3 | 30692061 | 30692293 | Intron |
| chr3 | 30692550 | 30692887 | Intron |
| chr3 | 30828820 | 30829304 | Intron |
| chr3 | 30867372 | 30867728 | Intron |
| chr3 | 30895088 | 30896132 | Intron |
| chr3 | 30896455 | 30896683 | Intron |
| chr3 | 30896863 | 30899547 | Exon |
| chr3 | 30917062 | 30918742 | Intron |
| chr3 | 30920859 | 30922114 | Intron |
| chr3 | 30932535 | 30933338 | Intron |
| chr3 | 30937077 | 30938094 | Promoter |
| chr3 | 30960128 | 30960900 | Distal Intergenic |
| chr3 | 30992688 | 30993122 | Distal Intergenic |
| chr3 | 30994254 | 30995429 | Distal Intergenic |
| chr3 | 30995492 | 30995890 | Distal Intergenic |
| chr3 | 31032677 | 31034782 | Distal Intergenic |
| chr3 | 31095899 | 31096594 | Distal Intergenic |
| chr3 | 31096638 | 31096902 | Distal Intergenic |
| chr3 | 31159057 | 31160047 | Distal Intergenic |
| chr3 | 31225680 | 31226906 | Distal Intergenic |
| chr3 | 31336640 | 31338173 | Distal Intergenic |
| chr3 | 31390479 | 31390945 | Distal Intergenic |
| chr3 | 31394462 | 31394826 | Distal Intergenic |
| chr3 | 32239386 | 32239639 | Distal Intergenic |
| chr3 | 32241188 | 32241518 | Distal Intergenic |
| chr3 | 32241596 | 32242140 | Distal Intergenic |
| chr3 | 32262904 | 32263870 | Distal Intergenic |
| chr3 | 32263937 | 32264233 | Distal Intergenic |
| chr3 | 32295243 | 32295442 | Intron |
| chr3 | 32314617 | 32315020 | Intron |
| chr3 | 32316606 | 32316806 | Intron |

| | | | |
|---|---|---|---|
| chr3 | 32409924 | 32410168 | Intron |
| chr3 | 33464266 | 33464488 | Intron |
| chr3 | 33698409 | 33698953 | Intron |
| chr3 | 34310214 | 34311015 | Distal Intergenic |
| chr3 | 34549610 | 34549839 | Distal Intergenic |
| chr3 | 34550427 | 34550902 | Distal Intergenic |
| chr3 | 34551303 | 34551568 | Distal Intergenic |
| chr3 | 34551640 | 34551875 | Distal Intergenic |
| chr3 | 34552460 | 34553000 | Distal Intergenic |
| chr3 | 34723549 | 34724669 | Distal Intergenic |
| chr3 | 34784339 | 34787456 | Distal Intergenic |
| chr3 | 34791951 | 34793082 | Distal Intergenic |
| chr3 | 35129621 | 35130394 | Distal Intergenic |
| chr3 | 35130679 | 35132474 | Distal Intergenic |
| chr3 | 35133952 | 35134206 | Distal Intergenic |
| chr3 | 35144453 | 35146028 | Distal Intergenic |
| chr3 | 35619339 | 35620433 | Distal Intergenic |
| chr3 | 35620828 | 35621168 | Distal Intergenic |
| chr3 | 35797023 | 35797418 | Intron |
| chr3 | 35830945 | 35831442 | Intron |
| chr3 | 35831528 | 35831727 | Intron |
| chr3 | 35832127 | 35832618 | Intron |
| chr3 | 36285964 | 36286744 | Distal Intergenic |
| chr3 | 36471208 | 36471515 | Intron |
| chr3 | 37211613 | 37212077 | Intron |
| chr3 | 37212113 | 37212926 | Intron |
| chr3 | 37319760 | 37320159 | Intron |
| chr3 | 37372028 | 37372258 | Intron |
| chr3 | 37394959 | 37395907 | Intron |
| chr3 | 37400091 | 37400360 | Intron |
| chr3 | 37400930 | 37403792 | 3' UTR |
| chr3 | 37805803 | 37806150 | Intron |
| chr3 | 38175482 | 38175681 | Exon |
| chr3 | 38175809 | 38176829 | Intron |
| chr3 | 38181807 | 38183919 | 3' UTR |
| chr3 | 38271947 | 38273351 | 3' UTR |
| chr3 | 38280654 | 38282687 | Intron |
| chr3 | 38341413 | 38342812 | Intron |
| chr3 | 38380997 | 38381296 | Distal Intergenic |
| chr3 | 38382483 | 38383437 | Distal Intergenic |
| chr3 | 38412905 | 38413134 | Intron |
| chr3 | 38460286 | 38460903 | Distal Intergenic |
| chr3 | 38515484 | 38516734 | Intron |
| chr3 | 39008138 | 39009042 | Distal Intergenic |

| | | | |
|---|---|---|---|
| chr3 | 39168537 | 39168937 | Intron |
| chr3 | 40485807 | 40487121 | Exon |
| chr3 | 40491841 | 40492118 | Intron |
| chr3 | 40498009 | 40498716 | Promoter |
| chr3 | 40549770 | 40550172 | Intron |
| chr3 | 40550208 | 40550458 | Intron |
| chr3 | 40564841 | 40565211 | Distal Intergenic |
| chr3 | 40603255 | 40603881 | Distal Intergenic |
| chr3 | 40887417 | 40888029 | Distal Intergenic |
| chr3 | 40997983 | 40999518 | Distal Intergenic |
| chr3 | 41004052 | 41004594 | Distal Intergenic |
| chr3 | 41024281 | 41025092 | Distal Intergenic |
| chr3 | 41088413 | 41090104 | Distal Intergenic |
| chr3 | 41092013 | 41093060 | Distal Intergenic |
| chr3 | 41116168 | 41116786 | Distal Intergenic |
| chr3 | 41121655 | 41123691 | Distal Intergenic |
| chr3 | 41178573 | 41180083 | Distal Intergenic |
| chr3 | 41180771 | 41181007 | Distal Intergenic |
| chr3 | 41240491 | 41240759 | Promoter |
| chr3 | 41313133 | 41313796 | Intron |
| chr3 | 41325449 | 41327118 | Intron |
| chr3 | 41351072 | 41351295 | Intron |
| chr3 | 41351388 | 41351662 | Intron |
| chr3 | 41367909 | 41368196 | Intron |
| chr3 | 41409719 | 41410100 | Intron |
| chr3 | 47466143 | 47467074 | Exon |
| chr3 | 49635992 | 49636191 | Intron |
| chr3 | 49805512 | 49806521 | Intron |
| chr3 | 50241728 | 50242711 | Promoter |
| chr3 | 50260137 | 50261690 | Distal Intergenic |
| chr3 | 50352664 | 50353056 | Downstream |
| chr3 | 51027520 | 51027731 | Intron |
| chr3 | 51027845 | 51028051 | Intron |
| chr3 | 51080981 | 51082045 | Intron |
| chr3 | 51131485 | 51131684 | Intron |
| chr3 | 51142317 | 51142654 | Intron |
| chr3 | 51147091 | 51147370 | Intron |
| chr3 | 51212503 | 51212702 | Intron |
| chr3 | 51215852 | 51218106 | Intron |
| chr3 | 51218657 | 51221017 | Intron |
| chr3 | 51287337 | 51287857 | Intron |
| chr3 | 51364084 | 51364810 | Intron |
| chr3 | 51377902 | 51378479 | Intron |
| chr3 | 51464702 | 51465587 | Exon |

| | | | |
|---|---|---|---|
| chr3 | 51465788 | 51466264 | Intron |
| chr3 | 51469246 | 51469923 | Intron |
| chr3 | 51525206 | 51525676 | Intron |
| chr3 | 51646040 | 51646979 | Intron |
| chr3 | 51709189 | 51710459 | Intron |
| chr3 | 51710976 | 51712070 | Intron |
| chr3 | 51712111 | 51712947 | Intron |
| chr3 | 51738634 | 51739498 | Distal Intergenic |
| chr3 | 51747184 | 51747406 | Exon |
| chr3 | 51747595 | 51748062 | Exon |
| chr3 | 51821673 | 51822031 | Distal Intergenic |
| chr3 | 51849859 | 51851641 | Promoter |
| chr3 | 51851841 | 51852532 | Promoter |
| chr3 | 51873531 | 51874995 | Intron |
| chr3 | 51892348 | 51892733 | Intron |
| chr3 | 51892805 | 51893013 | Intron |
| chr3 | 51906731 | 51906930 | Intron |
| chr3 | 51944355 | 51944554 | Distal Intergenic |
| chr3 | 51954745 | 51955645 | Distal Intergenic |
| chr3 | 51961994 | 51963212 | Distal Intergenic |
| chr3 | 52000521 | 52000888 | Promoter |
| chr3 | 52001530 | 52002057 | Promoter |
| chr3 | 52002220 | 52003317 | Promoter |
| chr3 | 52101683 | 52102048 | Distal Intergenic |
| chr3 | 52106861 | 52107122 | Downstream |
| chr3 | 52146747 | 52147561 | Intron |
| chr3 | 52407307 | 52407529 | Intron |
| chr3 | 52411912 | 52412174 | Intron |
| chr3 | 52434096 | 52434602 | 3' UTR |
| chr3 | 52434643 | 52434993 | Downstream |
| chr3 | 52867101 | 52867494 | Promoter |
| chr3 | 52867600 | 52870117 | Promoter |
| chr3 | 52912339 | 52912603 | Intron |
| chr3 | 52912827 | 52913070 | Intron |
| chr3 | 53004868 | 53006777 | Intron |
| chr3 | 53007049 | 53007405 | Intron |
| chr3 | 53007742 | 53009111 | Intron |
| chr3 | 53009171 | 53009374 | Intron |
| chr3 | 53266886 | 53267085 | Intron |
| chr3 | 53267131 | 53267450 | Exon |
| chr3 | 53667484 | 53667943 | Intron |
| chr3 | 54131740 | 54131948 | Distal Intergenic |
| chr3 | 54235711 | 54235937 | Intron |
| chr3 | 54270492 | 54271458 | Intron |

| | | | |
|---|---|---|---|
| chr3 | 54283761 | 54285398 | Intron |
| chr3 | 54497270 | 54497520 | Intron |
| chr3 | 54539828 | 54540070 | Intron |
| chr3 | 54916106 | 54917238 | Intron |
| chr3 | 54917761 | 54918072 | Intron |
| chr3 | 54918184 | 54918405 | Intron |
| chr3 | 54926169 | 54927559 | Intron |
| chr3 | 55204389 | 55205652 | Distal Intergenic |
| chr3 | 55392031 | 55392246 | Distal Intergenic |
| chr3 | 55394247 | 55394648 | Distal Intergenic |
| chr3 | 55396704 | 55397882 | Distal Intergenic |
| chr3 | 55399669 | 55401007 | Distal Intergenic |
| chr3 | 55403718 | 55404362 | Distal Intergenic |
| chr3 | 55420640 | 55420875 | Distal Intergenic |
| chr3 | 55421071 | 55421322 | Distal Intergenic |
| chr3 | 55691686 | 55691936 | Exon |
| chr3 | 55989567 | 55990091 | Intron |
| chr3 | 55990476 | 55990841 | Intron |
| chr3 | 55991062 | 55991826 | Intron |
| chr3 | 56765937 | 56766635 | 3' UTR |
| chr3 | 56766782 | 56768041 | Intron |
| chr3 | 56897119 | 56898024 | Intron |
| chr3 | 57098604 | 57098910 | Intron |
| chr3 | 57098949 | 57099184 | Intron |
| chr3 | 57244794 | 57245390 | Distal Intergenic |
| chr3 | 57304727 | 57305992 | 3' UTR |
| chr3 | 57358069 | 57359185 | Intron |
| chr3 | 57359771 | 57360080 | Intron |
| chr3 | 57539247 | 57539516 | Distal Intergenic |
| chr3 | 57540084 | 57540350 | Distal Intergenic |
| chr3 | 58007950 | 58008976 | Intron |
| chr3 | 58014530 | 58015822 | Intron |
| chr3 | 58174488 | 58177580 | Downstream |
| chr3 | 58218058 | 58218522 | Distal Intergenic |
| chr3 | 58219659 | 58220525 | Distal Intergenic |
| chr3 | 58220593 | 58220860 | Distal Intergenic |
| chr3 | 58220922 | 58221444 | Distal Intergenic |
| chr3 | 58221518 | 58223929 | Promoter |
| chr3 | 58223974 | 58224791 | Promoter |
| chr3 | 58224961 | 58225224 | Intron |
| chr3 | 58225314 | 58226210 | Intron |
| chr3 | 58227071 | 58227485 | Intron |
| chr3 | 58227661 | 58227860 | Intron |
| chr3 | 58272827 | 58275234 | Intron |

| | | | |
|---|---|---|---|
| chr3 | 58380899 | 58381288 | Intron |
| chr3 | 58917771 | 58919000 | Intron |
| chr3 | 58924676 | 58925991 | Intron |
| chr3 | 58972480 | 58973469 | Intron |
| chr3 | 58973540 | 58974677 | Intron |
| chr3 | 58991282 | 58991899 | Intron |
| chr3 | 58991975 | 58992292 | Intron |
| chr3 | 58992374 | 58992690 | Intron |
| chr3 | 58992982 | 58993456 | Intron |
| chr3 | 58994154 | 58994477 | Intron |
| chr3 | 58994628 | 58999457 | Intron |
| chr3 | 58999625 | 58999829 | Intron |
| chr3 | 59000311 | 59002696 | Intron |
| chr3 | 59007557 | 59008558 | Intron |
| chr3 | 59082440 | 59083174 | Distal Intergenic |
| chr3 | 59092126 | 59095079 | Distal Intergenic |
| chr3 | 59099444 | 59101068 | Distal Intergenic |
| chr3 | 59116455 | 59116726 | Distal Intergenic |
| chr3 | 59519599 | 59521118 | Distal Intergenic |
| chr3 | 59856903 | 59858888 | Intron |
| chr3 | 60150003 | 60151479 | Intron |
| chr3 | 60163689 | 60163891 | Intron |
| chr3 | 60163935 | 60164167 | Intron |
| chr3 | 60167236 | 60168726 | Intron |
| chr3 | 60192104 | 60193640 | Intron |
| chr3 | 60212307 | 60213137 | Intron |
| chr3 | 60227719 | 60228725 | Intron |
| chr3 | 60290331 | 60290602 | Intron |
| chr3 | 60302740 | 60304863 | Intron |
| chr3 | 60305455 | 60309129 | Intron |
| chr3 | 60321917 | 60322854 | Intron |
| chr3 | 60374135 | 60374495 | Intron |
| chr3 | 60406981 | 60407333 | Intron |
| chr3 | 60419671 | 60420070 | Intron |
| chr3 | 60420584 | 60421675 | Intron |
| chr3 | 60421720 | 60422320 | Intron |
| chr3 | 60436851 | 60437050 | Intron |
| chr3 | 60602151 | 60603557 | Intron |
| chr3 | 60690926 | 60691791 | Intron |
| chr3 | 60839459 | 60840619 | Intron |
| chr3 | 61020527 | 61020763 | Intron |
| chr3 | 61143966 | 61144754 | Intron |
| chr3 | 61147620 | 61147965 | Intron |
| chr3 | 61148272 | 61148887 | Intron |

| | | | |
|---|---|---|---|
| chr3 | 61166354 | 61167321 | Intron |
| chr3 | 61167365 | 61167646 | Intron |
| chr3 | 61167733 | 61168243 | Intron |
| chr3 | 61168524 | 61168810 | Intron |
| chr3 | 61168951 | 61169482 | Intron |
| chr3 | 61169683 | 61170128 | Intron |
| chr3 | 61179532 | 61180126 | Intron |
| chr3 | 61270190 | 61270457 | Distal Intergenic |
| chr3 | 61270550 | 61271009 | Distal Intergenic |
| chr3 | 63767891 | 63768408 | Distal Intergenic |
| chr3 | 63780085 | 63780516 | Distal Intergenic |
| chr3 | 64142646 | 64142854 | Exon |
| chr3 | 64142896 | 64143295 | Exon |
| chr3 | 65195323 | 65195569 | Distal Intergenic |
| chr3 | 65195740 | 65195953 | Distal Intergenic |
| chr3 | 65569696 | 65571110 | Intron |
| chr3 | 65645073 | 65645480 | Intron |
| chr3 | 65646883 | 65650770 | Intron |
| chr3 | 65689239 | 65689695 | Intron |
| chr3 | 65726417 | 65727741 | Intron |
| chr3 | 65728771 | 65729942 | Intron |
| chr3 | 65730298 | 65730582 | Intron |
| chr3 | 65730689 | 65732403 | Intron |
| chr3 | 65732847 | 65733495 | Intron |
| chr3 | 65760683 | 65761562 | Intron |
| chr3 | 65778090 | 65778688 | Intron |
| chr3 | 67267005 | 67267542 | Distal Intergenic |
| chr3 | 68296511 | 68297714 | Intron |
| chr3 | 68298151 | 68298861 | Intron |
| chr3 | 68299057 | 68299766 | Intron |
| chr3 | 68308630 | 68308829 | Intron |
| chr3 | 68311452 | 68313084 | Intron |
| chr3 | 68315420 | 68316172 | Intron |
| chr3 | 68338203 | 68338402 | Intron |
| chr3 | 68338720 | 68339000 | Intron |
| chr3 | 68808874 | 68809912 | Intron |
| chr3 | 68846712 | 68847039 | Intron |
| chr3 | 68847177 | 68847431 | Intron |
| chr3 | 68847492 | 68852104 | Intron |
| chr3 | 68929729 | 68930381 | Exon |
| chr3 | 68985313 | 68985538 | Distal Intergenic |
| chr3 | 68985797 | 68986225 | Distal Intergenic |
| chr3 | 69515659 | 69515902 | Intron |
| chr3 | 69516285 | 69516484 | Intron |

| | | | |
|---|---|---|---|
| chr3 | 69525959 | 69526182 | Intron |
| chr3 | 69526480 | 69526755 | Intron |
| chr3 | 70607138 | 70608205 | Distal Intergenic |
| chr3 | 70608351 | 70608589 | Distal Intergenic |
| chr3 | 75303502 | 75303701 | Distal Intergenic |
| chr3 | 75733794 | 75735259 | Distal Intergenic |
| chr3 | 75735979 | 75736311 | Distal Intergenic |
| chr3 | 75739893 | 75740690 | Distal Intergenic |
| chr3 | 75740804 | 75744683 | Distal Intergenic |
| chr3 | 75745752 | 75747513 | Distal Intergenic |
| chr3 | 75752361 | 75752656 | Distal Intergenic |
| chr3 | 75756703 | 75757099 | Downstream |
| chr3 | 75758226 | 75758449 | Downstream |
| chr3 | 75759199 | 75759785 | Exon |
| chr3 | 75759844 | 75760252 | Exon |
| chr3 | 75760391 | 75760711 | Exon |
| chr3 | 75789582 | 75791055 | Exon |
| chr3 | 78933372 | 78933579 | Intron |
| chr3 | 78956874 | 78957708 | Intron |
| chr3 | 78984994 | 78986601 | Intron |
| chr3 | 79051715 | 79053097 | Intron |
| chr3 | 79198436 | 79198835 | Intron |
| chr3 | 79198943 | 79199173 | Intron |
| chr3 | 79294119 | 79294463 | Intron |
| chr3 | 79362944 | 79363162 | Intron |
| chr3 | 79363378 | 79363832 | Intron |
| chr3 | 79395456 | 79395657 | Intron |
| chr3 | 79395808 | 79396112 | Intron |
| chr3 | 80687273 | 80688392 | Distal Intergenic |
| chr3 | 80861775 | 80863156 | Distal Intergenic |
| chr3 | 80910457 | 80910711 | Distal Intergenic |
| chr3 | 80910779 | 80911090 | Distal Intergenic |
| chr3 | 80947661 | 80947911 | Distal Intergenic |
| chr3 | 80967765 | 80968192 | Distal Intergenic |
| chr3 | 80988407 | 80989334 | Distal Intergenic |
| chr3 | 81004412 | 81005354 | Distal Intergenic |
| chr3 | 81009083 | 81009808 | Distal Intergenic |
| chr3 | 81721623 | 81721945 | Intron |
| chr3 | 81722006 | 81722295 | Intron |
| chr3 | 83579026 | 83579236 | Distal Intergenic |
| chr3 | 83666800 | 83667425 | Distal Intergenic |
| chr3 | 83669271 | 83669615 | Distal Intergenic |
| chr3 | 83747463 | 83748042 | Distal Intergenic |
| chr3 | 83791430 | 83792310 | Distal Intergenic |

| | | | |
|---|---|---|---|
| chr3 | 83797177 | 83797497 | Distal Intergenic |
| chr3 | 83797812 | 83798097 | Distal Intergenic |
| chr3 | 83825316 | 83827852 | Distal Intergenic |
| chr3 | 83911922 | 83913211 | Distal Intergenic |
| chr3 | 84097347 | 84097549 | Distal Intergenic |
| chr3 | 84119060 | 84119325 | Distal Intergenic |
| chr3 | 84122314 | 84123787 | Distal Intergenic |
| chr3 | 84127398 | 84129297 | Distal Intergenic |
| chr3 | 84284477 | 84284911 | Distal Intergenic |
| chr3 | 84285075 | 84285310 | Distal Intergenic |
| chr3 | 84307265 | 84307490 | Distal Intergenic |
| chr3 | 84405950 | 84407802 | Distal Intergenic |
| chr3 | 84449888 | 84450233 | Distal Intergenic |
| chr3 | 84499190 | 84500519 | Distal Intergenic |
| chr3 | 84500577 | 84501000 | Distal Intergenic |
| chr3 | 84644242 | 84644441 | Distal Intergenic |
| chr3 | 84644562 | 84644875 | Distal Intergenic |
| chr3 | 84755640 | 84756483 | Intron |
| chr3 | 84774692 | 84776633 | Intron |
| chr3 | 84776731 | 84778780 | Intron |
| chr3 | 84826838 | 84827187 | Intron |
| chr3 | 84828414 | 84828805 | Intron |
| chr3 | 85087525 | 85089021 | Intron |
| chr3 | 85172037 | 85173947 | Intron |
| chr3 | 85282482 | 85282850 | Intron |
| chr3 | 85282991 | 85283265 | Intron |
| chr3 | 85377983 | 85378190 | Intron |
| chr3 | 85595958 | 85597497 | Intron |
| chr3 | 85684561 | 85685299 | Intron |
| chr3 | 85735433 | 85736149 | Intron |
| chr3 | 85769426 | 85769630 | Intron |
| chr3 | 85797960 | 85798457 | Intron |
| chr3 | 85844273 | 85845175 | Intron |
| chr3 | 86057178 | 86057511 | Intron |
| chr3 | 86654595 | 86655334 | Distal Intergenic |
| chr3 | 86876083 | 86878174 | Distal Intergenic |
| chr3 | 86883380 | 86884299 | Distal Intergenic |
| chr3 | 86903905 | 86904595 | Distal Intergenic |
| chr3 | 86904634 | 86905584 | Distal Intergenic |
| chr3 | 86916748 | 86917028 | Distal Intergenic |
| chr3 | 86944670 | 86946175 | Distal Intergenic |
| chr3 | 86953798 | 86955930 | Distal Intergenic |
| chr3 | 86961859 | 86962909 | Distal Intergenic |
| chr3 | 86977249 | 86977587 | Distal Intergenic |

| | | | |
|---|---|---|---|
| chr3 | 86977633 | 86977879 | Distal Intergenic |
| chr3 | 87322859 | 87323337 | Intron |
| chr3 | 87369500 | 87370326 | Distal Intergenic |
| chr3 | 87480179 | 87480480 | Distal Intergenic |
| chr3 | 87689134 | 87689768 | Distal Intergenic |
| chr3 | 87710775 | 87714544 | Distal Intergenic |
| chr3 | 87714664 | 87715090 | Distal Intergenic |
| chr3 | 87723156 | 87723359 | Distal Intergenic |
| chr3 | 87723472 | 87723922 | Distal Intergenic |
| chr3 | 87746470 | 87746747 | Distal Intergenic |
| chr3 | 87748779 | 87749109 | Distal Intergenic |
| chr3 | 87839352 | 87840176 | Distal Intergenic |
| chr3 | 87942164 | 87942363 | Distal Intergenic |
| chr3 | 87942583 | 87945475 | Distal Intergenic |
| chr3 | 87945546 | 87945901 | Distal Intergenic |
| chr3 | 87945968 | 87946218 | Distal Intergenic |
| chr3 | 87947176 | 87947565 | Distal Intergenic |
| chr3 | 87981091 | 87981404 | Distal Intergenic |
| chr3 | 87981771 | 87981994 | Distal Intergenic |
| chr3 | 88100108 | 88100699 | Downstream |
| chr3 | 88101144 | 88101371 | 3' UTR |
| chr3 | 88137568 | 88139491 | Intron |
| chr3 | 88139675 | 88140108 | Intron |
| chr3 | 88140675 | 88141135 | Intron |
| chr3 | 88141272 | 88141585 | Intron |
| chr3 | 88220238 | 88221056 | Distal Intergenic |
| chr3 | 88225919 | 88226447 | Distal Intergenic |
| chr3 | 88292363 | 88293451 | Distal Intergenic |
| chr3 | 88294643 | 88297057 | Distal Intergenic |
| chr3 | 88297114 | 88300959 | Distal Intergenic |
| chr3 | 88303923 | 88305165 | Distal Intergenic |
| chr3 | 88305303 | 88306908 | Distal Intergenic |
| chr3 | 88307336 | 88307596 | Distal Intergenic |
| chr3 | 88313642 | 88314008 | Distal Intergenic |
| chr3 | 88314084 | 88314905 | Distal Intergenic |
| chr3 | 88315538 | 88316230 | Distal Intergenic |
| chr3 | 88316283 | 88316629 | Distal Intergenic |
| chr3 | 88316686 | 88317990 | Distal Intergenic |
| chr3 | 88357112 | 88357314 | Distal Intergenic |
| chr3 | 88395893 | 88398746 | Distal Intergenic |
| chr3 | 88425503 | 88426815 | Distal Intergenic |
| chr3 | 88427755 | 88428450 | Distal Intergenic |
| chr3 | 88428487 | 88429331 | Distal Intergenic |
| chr3 | 88457845 | 88458615 | Distal Intergenic |

| | | | |
|---|---|---|---|
| chr3 | 88489306 | 88489622 | Distal Intergenic |
| chr3 | 88489704 | 88490503 | Distal Intergenic |
| chr3 | 88717935 | 88718134 | Distal Intergenic |
| chr3 | 88754846 | 88755088 | Distal Intergenic |
| chr3 | 88906040 | 88906411 | Distal Intergenic |
| chr3 | 88906447 | 88906779 | Distal Intergenic |
| chr3 | 89011790 | 89012545 | Distal Intergenic |
| chr3 | 89017603 | 89017802 | Distal Intergenic |
| chr3 | 89062942 | 89063404 | Distal Intergenic |
| chr3 | 89081301 | 89083487 | Distal Intergenic |
| chr3 | 89086038 | 89086295 | Distal Intergenic |
| chr3 | 89132197 | 89133212 | Distal Intergenic |
| chr3 | 89182993 | 89183262 | Intron |
| chr3 | 89195447 | 89196497 | Intron |
| chr3 | 89196554 | 89197168 | Intron |
| chr3 | 89197397 | 89197628 | Intron |
| chr3 | 89197726 | 89198331 | Intron |
| chr3 | 89198387 | 89198632 | Intron |
| chr3 | 89199124 | 89199354 | Intron |
| chr3 | 89215554 | 89215801 | Intron |
| chr3 | 89222110 | 89222376 | Intron |
| chr3 | 89222428 | 89224950 | Intron |
| chr3 | 89225173 | 89225395 | Intron |
| chr3 | 89262433 | 89262982 | Intron |
| chr3 | 89263457 | 89263682 | Intron |
| chr3 | 89489300 | 89490303 | Intron |
| chr3 | 89599598 | 89599921 | Distal Intergenic |
| chr3 | 89611261 | 89612863 | Distal Intergenic |
| chr3 | 89616837 | 89617236 | Distal Intergenic |
| chr3 | 89665096 | 89665812 | Distal Intergenic |
| chr3 | 89684016 | 89684728 | Distal Intergenic |
| chr3 | 89699332 | 89703050 | Distal Intergenic |
| chr3 | 89705218 | 89706251 | Distal Intergenic |
| chr3 | 89716260 | 89717679 | Distal Intergenic |
| chr3 | 89766244 | 89766908 | Distal Intergenic |
| chr3 | 89767263 | 89768311 | Distal Intergenic |
| chr3 | 89802321 | 89802581 | Distal Intergenic |
| chr3 | 89802643 | 89802894 | Distal Intergenic |
| chr3 | 89855435 | 89856017 | Distal Intergenic |
| chr3 | 89883588 | 89884681 | Distal Intergenic |
| chr3 | 89885214 | 89886141 | Distal Intergenic |
| chr3 | 89990862 | 89991114 | Distal Intergenic |
| chr3 | 89991167 | 89992539 | Distal Intergenic |
| chr3 | 90012240 | 90013698 | Distal Intergenic |

| | | | |
|---|---|---|---|
| chr3 | 90013789 | 90014319 | Distal Intergenic |
| chr3 | 90014670 | 90015238 | Distal Intergenic |
| chr3 | 90015727 | 90016608 | Distal Intergenic |
| chr3 | 90017051 | 90017622 | Distal Intergenic |
| chr3 | 90033057 | 90033368 | Distal Intergenic |
| chr3 | 90049526 | 90050355 | Distal Intergenic |
| chr3 | 90051607 | 90051985 | Distal Intergenic |
| chr3 | 90052413 | 90052714 | Distal Intergenic |
| chr3 | 90069342 | 90069687 | Distal Intergenic |
| chr3 | 90096377 | 90096885 | Distal Intergenic |
| chr3 | 90237147 | 90237480 | Distal Intergenic |
| chr3 | 90312676 | 90312897 | Distal Intergenic |
| chr3 | 90313196 | 90313423 | Distal Intergenic |
| chr3 | 90313578 | 90313782 | Distal Intergenic |
| chr3 | 90329832 | 90330224 | Distal Intergenic |
| chr3 | 90330264 | 90331687 | Distal Intergenic |
| chr3 | 90341082 | 90342146 | Distal Intergenic |
| chr3 | 90342182 | 90342381 | Distal Intergenic |
| chr3 | 90378814 | 90379407 | Distal Intergenic |
| chr3 | 90379538 | 90379783 | Distal Intergenic |
| chr3 | 90389324 | 90389616 | Distal Intergenic |
| chr3 | 90389797 | 90394292 | Distal Intergenic |
| chr3 | 90401945 | 90402167 | Distal Intergenic |
| chr3 | 90404135 | 90405629 | Distal Intergenic |
| chr3 | 90408074 | 90409160 | Distal Intergenic |
| chr3 | 90416372 | 90416925 | Distal Intergenic |
| chr3 | 90417018 | 90419420 | Distal Intergenic |
| chr3 | 91816656 | 91817297 | Distal Intergenic |
| chr3 | 91817358 | 91818027 | Distal Intergenic |
| chr3 | 91838056 | 91838409 | Distal Intergenic |
| chr3 | 91973403 | 91973950 | Distal Intergenic |
| chr3 | 91974457 | 91974683 | Distal Intergenic |
| chr3 | 92014785 | 92017021 | Distal Intergenic |
| chr3 | 92017311 | 92018032 | Distal Intergenic |
| chr3 | 92018138 | 92018526 | Distal Intergenic |
| chr3 | 92018709 | 92019892 | Distal Intergenic |
| chr3 | 92048302 | 92048715 | Distal Intergenic |
| chr3 | 92048763 | 92051842 | Distal Intergenic |
| chr3 | 92052114 | 92052859 | Distal Intergenic |
| chr3 | 92052983 | 92053500 | Distal Intergenic |
| chr3 | 92053553 | 92054918 | Distal Intergenic |
| chr3 | 92082794 | 92086536 | Distal Intergenic |
| chr3 | 92086816 | 92087500 | Distal Intergenic |
| chr3 | 92088172 | 92089543 | Distal Intergenic |

| | | | |
|---|---|---|---|
| chr3 | 92213919 | 92215309 | Distal Intergenic |
| chr3 | 92219062 | 92219429 | Distal Intergenic |
| chr3 | 92221318 | 92221667 | Distal Intergenic |
| chr3 | 92244437 | 92245316 | Distal Intergenic |
| chr3 | 92259095 | 92259564 | Distal Intergenic |
| chr3 | 92317404 | 92317793 | Distal Intergenic |
| chr3 | 92358179 | 92358456 | Distal Intergenic |
| chr3 | 92358636 | 92358843 | Distal Intergenic |
| chr3 | 92377916 | 92378115 | Distal Intergenic |
| chr3 | 92383508 | 92383775 | Distal Intergenic |
| chr3 | 92389904 | 92391022 | Distal Intergenic |
| chr3 | 92391089 | 92391696 | Distal Intergenic |
| chr3 | 92391880 | 92393454 | Distal Intergenic |
| chr3 | 92413093 | 92413331 | Distal Intergenic |
| chr3 | 92413387 | 92413924 | Distal Intergenic |
| chr3 | 92413961 | 92414267 | Distal Intergenic |
| chr3 | 92457341 | 92457740 | Distal Intergenic |
| chr3 | 92626974 | 92627716 | Distal Intergenic |
| chr3 | 92697980 | 92699581 | Distal Intergenic |
| chr3 | 92827532 | 92827889 | Distal Intergenic |
| chr3 | 92828033 | 92828280 | Distal Intergenic |
| chr3 | 92829179 | 92829601 | Distal Intergenic |
| chr3 | 92830083 | 92830282 | Distal Intergenic |
| chr3 | 92948274 | 92949147 | Distal Intergenic |
| chr3 | 92980509 | 92980949 | Distal Intergenic |
| chr3 | 93233187 | 93234081 | Distal Intergenic |
| chr3 | 93235818 | 93236152 | Distal Intergenic |
| chr3 | 93310879 | 93311110 | Distal Intergenic |
| chr3 | 93311146 | 93311741 | Distal Intergenic |
| chr3 | 93323778 | 93326219 | Distal Intergenic |
| chr3 | 93326469 | 93326874 | Distal Intergenic |
| chr3 | 93340759 | 93341153 | Distal Intergenic |
| chr3 | 93341377 | 93343230 | Distal Intergenic |
| chr3 | 93359079 | 93360034 | Distal Intergenic |
| chr3 | 93360451 | 93360657 | Distal Intergenic |
| chr3 | 93375847 | 93376075 | Distal Intergenic |
| chr3 | 93383719 | 93383924 | Distal Intergenic |
| chr3 | 93384220 | 93385441 | Distal Intergenic |
| chr3 | 94286465 | 94286853 | Distal Intergenic |
| chr3 | 94368561 | 94369750 | Distal Intergenic |
| chr3 | 94415019 | 94415825 | Distal Intergenic |
| chr3 | 94421808 | 94422017 | Distal Intergenic |
| chr3 | 94422089 | 94422589 | Distal Intergenic |
| chr3 | 94425535 | 94426136 | Distal Intergenic |

| | | | |
|---|---|---|---|
| chr3 | 94428259 | 94428613 | Distal Intergenic |
| chr3 | 94432749 | 94433322 | Distal Intergenic |
| chr3 | 94433519 | 94434590 | Distal Intergenic |
| chr3 | 94462325 | 94462767 | Distal Intergenic |
| chr3 | 94462877 | 94463735 | Distal Intergenic |
| chr3 | 94561502 | 94561874 | Distal Intergenic |
| chr3 | 94562226 | 94562528 | Distal Intergenic |
| chr3 | 94584325 | 94585869 | Distal Intergenic |
| chr3 | 94586104 | 94586341 | Distal Intergenic |
| chr3 | 94586452 | 94588151 | Distal Intergenic |
| chr3 | 94757206 | 94757789 | Intron |
| chr3 | 94819554 | 94819844 | Intron |
| chr3 | 94819887 | 94820119 | Intron |
| chr3 | 94846084 | 94846426 | Intron |
| chr3 | 94857877 | 94858093 | Intron |
| chr3 | 94858242 | 94859830 | Intron |
| chr3 | 94869860 | 94870635 | Intron |
| chr3 | 94870711 | 94871131 | Intron |
| chr3 | 94871211 | 94871565 | Intron |
| chr3 | 94909067 | 94910828 | Distal Intergenic |
| chr3 | 94912575 | 94913941 | Distal Intergenic |
| chr3 | 94914813 | 94915350 | Distal Intergenic |
| chr3 | 94937558 | 94937958 | Distal Intergenic |
| chr3 | 95021372 | 95022695 | Distal Intergenic |
| chr3 | 95031170 | 95031473 | Distal Intergenic |
| chr3 | 95031597 | 95033810 | Distal Intergenic |
| chr3 | 95033975 | 95034621 | Distal Intergenic |
| chr3 | 95034718 | 95035697 | Distal Intergenic |
| chr3 | 95036127 | 95036852 | Distal Intergenic |
| chr3 | 95036935 | 95037228 | Distal Intergenic |
| chr3 | 95065611 | 95066796 | Distal Intergenic |
| chr3 | 95086385 | 95086818 | Distal Intergenic |
| chr3 | 95238030 | 95238251 | Distal Intergenic |
| chr3 | 95454497 | 95455407 | Distal Intergenic |
| chr3 | 95463000 | 95463375 | Distal Intergenic |
| chr3 | 95463431 | 95463972 | Distal Intergenic |
| chr3 | 95465058 | 95465323 | Distal Intergenic |
| chr3 | 95467061 | 95467338 | Distal Intergenic |
| chr3 | 95467633 | 95469725 | Distal Intergenic |
| chr3 | 95469970 | 95472103 | Distal Intergenic |
| chr3 | 95477866 | 95478173 | Distal Intergenic |
| chr3 | 95478210 | 95479885 | Distal Intergenic |
| chr3 | 95479932 | 95480222 | Distal Intergenic |
| chr3 | 95500238 | 95501135 | Distal Intergenic |

| | | | |
|---|---|---|---|
| chr3 | 95505760 | 95506720 | Distal Intergenic |
| chr3 | 95529201 | 95531060 | Distal Intergenic |
| chr3 | 95537563 | 95538515 | Distal Intergenic |
| chr3 | 95542903 | 95543511 | Distal Intergenic |
| chr3 | 95562091 | 95563526 | Distal Intergenic |
| chr3 | 95564857 | 95565350 | Distal Intergenic |
| chr3 | 95565387 | 95566904 | Distal Intergenic |
| chr3 | 95621885 | 95622883 | Distal Intergenic |
| chr3 | 95623813 | 95624127 | Distal Intergenic |
| chr3 | 95659541 | 95659740 | Distal Intergenic |
| chr3 | 95891178 | 95892042 | Distal Intergenic |
| chr3 | 95897881 | 95898105 | Distal Intergenic |
| chr3 | 95963534 | 95965451 | Distal Intergenic |
| chr3 | 95965504 | 95965736 | Distal Intergenic |
| chr3 | 95975931 | 95976705 | Distal Intergenic |
| chr3 | 96001228 | 96001706 | Distal Intergenic |
| chr3 | 96023985 | 96024824 | Distal Intergenic |
| chr3 | 96030602 | 96031277 | Distal Intergenic |
| chr3 | 96031371 | 96031571 | Distal Intergenic |
| chr3 | 96041303 | 96042852 | Distal Intergenic |
| chr3 | 96043176 | 96043816 | Distal Intergenic |
| chr3 | 96048462 | 96049074 | Distal Intergenic |
| chr3 | 96049678 | 96050412 | Distal Intergenic |
| chr3 | 96051377 | 96051622 | Distal Intergenic |
| chr3 | 96051701 | 96052025 | Distal Intergenic |
| chr3 | 96065680 | 96068550 | Distal Intergenic |
| chr3 | 96071908 | 96073254 | Distal Intergenic |
| chr3 | 96077782 | 96078094 | Distal Intergenic |
| chr3 | 96218317 | 96218733 | Distal Intergenic |
| chr3 | 96433879 | 96434078 | Distal Intergenic |
| chr3 | 96477261 | 96480213 | Distal Intergenic |
| chr3 | 96482701 | 96483446 | Distal Intergenic |
| chr3 | 96483870 | 96484089 | Distal Intergenic |
| chr3 | 96617642 | 96618024 | Intron |
| chr3 | 97438526 | 97438836 | Intron |
| chr3 | 97631812 | 97632354 | Intron |
| chr3 | 97736727 | 97739079 | Intron |
| chr3 | 97739258 | 97739483 | Intron |
| chr3 | 97741105 | 97742590 | Intron |
| chr3 | 97818245 | 97818485 | Distal Intergenic |
| chr3 | 97824447 | 97824651 | Distal Intergenic |
| chr3 | 97824821 | 97825061 | Distal Intergenic |
| chr3 | 97831730 | 97831949 | Distal Intergenic |
| chr3 | 97837389 | 97837722 | Distal Intergenic |

| | | | |
|---|---|---|---|
| chr3 | 97860594 | 97861462 | Distal Intergenic |
| chr3 | 97883816 | 97884981 | Distal Intergenic |
| chr3 | 98080390 | 98080775 | Distal Intergenic |
| chr3 | 98178024 | 98179886 | Distal Intergenic |
| chr3 | 100488518 | 100488776 | Intron |
| chr3 | 100507569 | 100507813 | Intron |
| chr3 | 100600801 | 100602193 | Intron |
| chr3 | 100652369 | 100652568 | Intron |
| chr3 | 100743825 | 100745942 | Distal Intergenic |
| chr3 | 100814575 | 100814801 | Distal Intergenic |
| chr3 | 100826406 | 100827595 | Distal Intergenic |
| chr3 | 100835564 | 100836276 | Distal Intergenic |
| chr3 | 100837502 | 100838088 | Distal Intergenic |
| chr3 | 100851145 | 100851950 | Distal Intergenic |
| chr3 | 100852092 | 100852998 | Distal Intergenic |
| chr3 | 100853090 | 100853534 | Distal Intergenic |
| chr3 | 100855459 | 100855719 | Distal Intergenic |
| chr3 | 100874488 | 100874761 | Distal Intergenic |
| chr3 | 100877272 | 100878729 | Distal Intergenic |
| chr3 | 100900259 | 100900563 | Distal Intergenic |
| chr3 | 100919745 | 100920063 | Distal Intergenic |
| chr3 | 100920241 | 100920551 | Distal Intergenic |
| chr3 | 101116700 | 101117421 | Intron |
| chr3 | 101117573 | 101118080 | Exon |
| chr3 | 101122254 | 101122591 | Intron |
| chr3 | 101122867 | 101123418 | Intron |
| chr3 | 101125706 | 101126508 | Intron |
| chr3 | 101150180 | 101151507 | Intron |
| chr3 | 101246517 | 101246752 | Distal Intergenic |
| chr3 | 101247127 | 101247623 | Distal Intergenic |
| chr3 | 101251717 | 101251916 | Distal Intergenic |
| chr3 | 101252100 | 101252940 | Distal Intergenic |
| chr3 | 101254424 | 101254845 | Distal Intergenic |
| chr3 | 101254930 | 101255180 | Distal Intergenic |
| chr3 | 101265228 | 101266672 | Distal Intergenic |
| chr3 | 101374947 | 101375499 | Exon |
| chr3 | 101375645 | 101376406 | Intron |
| chr3 | 101376459 | 101376803 | Intron |
| chr3 | 101376904 | 101377107 | Intron |
| chr3 | 101382426 | 101382689 | Intron |
| chr3 | 101404365 | 101404673 | Promoter |
| chr3 | 101405652 | 101405864 | Promoter |
| chr3 | 101405980 | 101406270 | Promoter |
| chr3 | 101407253 | 101407490 | Distal Intergenic |

| | | | |
|---|---|---|---|
| chr3 | 101407801 | 101408075 | Distal Intergenic |
| chr3 | 101439783 | 101441391 | Distal Intergenic |
| chr3 | 101494885 | 101496630 | Distal Intergenic |
| chr3 | 101503782 | 101504140 | Intron |
| chr3 | 101504300 | 101505415 | 5' UTR |
| chr3 | 101505516 | 101505809 | Intron |
| chr3 | 101632237 | 101635313 | Distal Intergenic |
| chr3 | 101635549 | 101636303 | Distal Intergenic |
| chr3 | 101639170 | 101639594 | Distal Intergenic |
| chr3 | 101639732 | 101640518 | Distal Intergenic |
| chr3 | 101842121 | 101844188 | Intron |
| chr3 | 101907549 | 101908903 | Intron |
| chr3 | 101912205 | 101912409 | Intron |
| chr3 | 101915506 | 101917489 | Intron |
| chr3 | 101939282 | 101939937 | Intron |
| chr3 | 101940000 | 101940482 | Intron |
| chr3 | 101945399 | 101947329 | 5' UTR |
| chr3 | 101955729 | 101956999 | Intron |
| chr3 | 101963002 | 101964771 | Intron |
| chr3 | 101973000 | 101973249 | Intron |
| chr3 | 101973351 | 101974348 | Intron |
| chr3 | 101974501 | 101975803 | Intron |
| chr3 | 101976318 | 101976666 | Intron |
| chr3 | 101976702 | 101977451 | Intron |
| chr3 | 101977871 | 101978986 | Intron |
| chr3 | 101981774 | 101983090 | Intron |
| chr3 | 101991674 | 101992481 | Intron |
| chr3 | 101992520 | 101992933 | Intron |
| chr3 | 101996828 | 101997287 | Intron |
| chr3 | 101997384 | 101998655 | Intron |
| chr3 | 101999277 | 101999519 | Intron |
| chr3 | 102007507 | 102007859 | Intron |
| chr3 | 102136915 | 102137282 | 5' UTR |
| chr3 | 102140597 | 102141870 | Intron |
| chr3 | 102142012 | 102142368 | Intron |
| chr3 | 102168148 | 102168758 | Intron |
| chr3 | 102176005 | 102176222 | Intron |
| chr3 | 102176259 | 102177198 | Exon |
| chr3 | 102178405 | 102178683 | Intron |
| chr3 | 102178752 | 102179044 | Intron |
| chr3 | 102278678 | 102278969 | Distal Intergenic |
| chr3 | 102281353 | 102282203 | Distal Intergenic |
| chr3 | 102471782 | 102472238 | Distal Intergenic |
| chr3 | 102474691 | 102475105 | Distal Intergenic |

| | | | |
|---|---|---|---|
| chr3 | 102492250 | 102492460 | Distal Intergenic |
| chr3 | 102823935 | 102824220 | Distal Intergenic |
| chr3 | 102824505 | 102824879 | Distal Intergenic |
| chr3 | 102825007 | 102825488 | Distal Intergenic |
| chr3 | 102859756 | 102859990 | Distal Intergenic |
| chr3 | 102862243 | 102863098 | Distal Intergenic |
| chr3 | 102904626 | 102905767 | Distal Intergenic |
| chr3 | 102906022 | 102906507 | Distal Intergenic |
| chr3 | 102906545 | 102907240 | Distal Intergenic |
| chr3 | 102933982 | 102934770 | Distal Intergenic |
| chr3 | 102943221 | 102944627 | Distal Intergenic |
| chr3 | 102944845 | 102945961 | Distal Intergenic |
| chr3 | 102949348 | 102951690 | Distal Intergenic |
| chr3 | 103045744 | 103046052 | Distal Intergenic |
| chr3 | 103059966 | 103060242 | Distal Intergenic |
| chr3 | 103060389 | 103060642 | Distal Intergenic |
| chr3 | 103066695 | 103067636 | Distal Intergenic |
| chr3 | 103215880 | 103216850 | Distal Intergenic |
| chr3 | 103634675 | 103634876 | Distal Intergenic |
| chr3 | 103634931 | 103635243 | Distal Intergenic |
| chr3 | 103711425 | 103711648 | Intron |
| chr3 | 103718926 | 103719869 | Intron |
| chr3 | 103720572 | 103720824 | Intron |
| chr3 | 103771871 | 103772275 | Distal Intergenic |
| chr3 | 104163880 | 104164103 | Distal Intergenic |
| chr3 | 104544664 | 104545639 | Distal Intergenic |
| chr3 | 104545840 | 104546471 | Distal Intergenic |
| chr3 | 104546646 | 104546860 | Distal Intergenic |
| chr3 | 104666991 | 104668197 | Distal Intergenic |
| chr3 | 104856864 | 104857094 | Distal Intergenic |
| chr3 | 104857674 | 104858089 | Distal Intergenic |
| chr3 | 105187348 | 105187763 | Intron |
| chr3 | 105198661 | 105198997 | Intron |
| chr3 | 105199109 | 105199314 | Intron |
| chr3 | 105199431 | 105200494 | Intron |
| chr3 | 105507922 | 105508194 | Intron |
| chr3 | 106803021 | 106803532 | Intron |
| chr3 | 107136680 | 107137246 | Distal Intergenic |
| chr3 | 107471565 | 107471943 | Intron |
| chr3 | 107472024 | 107472702 | Intron |
| chr3 | 107578047 | 107578926 | Intron |
| chr3 | 107579074 | 107579958 | Intron |
| chr3 | 107596105 | 107596527 | Promoter |
| chr3 | 107761201 | 107761406 | Downstream |

| | | | |
|---|---|---|---|
| chr3 | 107816303 | 107816584 | Distal Intergenic |
| chr3 | 107888651 | 107889211 | Intron |
| chr3 | 107889331 | 107889839 | Intron |
| chr3 | 107929199 | 107929981 | Intron |
| chr3 | 107931745 | 107933818 | Exon |
| chr3 | 107934090 | 107935513 | Intron |
| chr3 | 107948233 | 107948670 | Distal Intergenic |
| chr3 | 107948800 | 107949378 | Distal Intergenic |
| chr3 | 107949658 | 107949886 | Distal Intergenic |
| chr3 | 108060120 | 108060365 | Intron |
| chr3 | 108204949 | 108205196 | Intron |
| chr3 | 108227094 | 108227576 | Intron |
| chr3 | 108227648 | 108228018 | Intron |
| chr3 | 108246837 | 108247135 | Intron |
| chr3 | 108365473 | 108366048 | Exon |
| chr3 | 108374712 | 108375026 | 3' UTR |
| chr3 | 108388598 | 108390397 | Intron |
| chr3 | 108565462 | 108565765 | Intron |
| chr3 | 108565958 | 108566696 | Exon |
| chr3 | 108573821 | 108574351 | Distal Intergenic |
| chr3 | 108574585 | 108575674 | Distal Intergenic |
| chr3 | 108575844 | 108576183 | Distal Intergenic |
| chr3 | 108576297 | 108576575 | Distal Intergenic |
| chr3 | 108576938 | 108578228 | Distal Intergenic |
| chr3 | 108581528 | 108582637 | Distal Intergenic |
| chr3 | 108584074 | 108584794 | Distal Intergenic |
| chr3 | 108623281 | 108623657 | Downstream |
| chr3 | 108658575 | 108659469 | Intron |
| chr3 | 108659670 | 108659944 | Intron |
| chr3 | 108659988 | 108662004 | Intron |
| chr3 | 108665674 | 108666434 | Intron |
| chr3 | 108682931 | 108683372 | Intron |
| chr3 | 108683516 | 108685424 | Intron |
| chr3 | 108713659 | 108714334 | Intron |
| chr3 | 108714451 | 108714834 | Intron |
| chr3 | 108761121 | 108761465 | Intron |
| chr3 | 108762040 | 108762422 | Intron |
| chr3 | 108762597 | 108762991 | Intron |
| chr3 | 108854783 | 108855727 | Promoter |
| chr3 | 108926257 | 108926618 | Distal Intergenic |
| chr3 | 108926806 | 108927148 | Distal Intergenic |
| chr3 | 109111842 | 109113297 | Distal Intergenic |
| chr3 | 109114346 | 109114548 | Distal Intergenic |
| chr3 | 109171410 | 109171881 | Intron |

| | | | |
|---|---|---|---|
| chr3 | 109271785 | 109272147 | Distal Intergenic |
| chr3 | 109272625 | 109273166 | Distal Intergenic |
| chr3 | 109424884 | 109425094 | Distal Intergenic |
| chr3 | 109425166 | 109426681 | Distal Intergenic |
| chr3 | 115590300 | 115590713 | Intron |
| chr3 | 115590833 | 115591292 | Intron |
| chr3 | 115817779 | 115818209 | Intron |
| chr3 | 115821847 | 115822092 | Intron |
| chr3 | 115892445 | 115892644 | Intron |
| chr3 | 116365304 | 116365532 | Distal Intergenic |
| chr3 | 116608798 | 116608999 | Distal Intergenic |
| chr3 | 116747188 | 116747509 | Distal Intergenic |
| chr3 | 116747616 | 116748676 | Distal Intergenic |
| chr3 | 116770221 | 116770790 | Distal Intergenic |
| chr3 | 116770915 | 116771478 | Distal Intergenic |
| chr3 | 116772956 | 116773546 | Distal Intergenic |
| chr3 | 116773617 | 116773930 | Distal Intergenic |
| chr3 | 116773981 | 116774358 | Distal Intergenic |
| chr3 | 116779370 | 116779726 | Distal Intergenic |
| chr3 | 116779807 | 116780895 | Distal Intergenic |
| chr3 | 116968419 | 116968703 | Distal Intergenic |
| chr3 | 116968787 | 116968986 | Distal Intergenic |
| chr3 | 116969093 | 116969414 | Distal Intergenic |
| chr3 | 119598299 | 119598551 | Intron |
| chr3 | 119658851 | 119659277 | Intron |
| chr3 | 119718557 | 119719421 | Intron |
| chr3 | 121081954 | 121082431 | Intron |
| chr3 | 121129958 | 121130291 | Intron |
| chr3 | 121304841 | 121306032 | 3' UTR |
| chr3 | 121319787 | 121321616 | Intron |
| chr3 | 121328844 | 121329478 | Intron |
| chr3 | 121352999 | 121353599 | Exon |
| chr3 | 121354953 | 121355524 | Exon |
| chr3 | 121364650 | 121365237 | Intron |
| chr3 | 121377911 | 121379743 | Promoter |
| chr3 | 121713356 | 121713792 | Intron |
| chr3 | 121765866 | 121766094 | Distal Intergenic |
| chr3 | 121785165 | 121785421 | Intron |
| chr3 | 121790531 | 121792168 | Intron |
| chr3 | 122024698 | 122025852 | Distal Intergenic |
| chr3 | 122051849 | 122052048 | Intron |
| chr3 | 122066563 | 122066773 | Distal Intergenic |
| chr3 | 122093952 | 122095629 | Intron |
| chr3 | 122096610 | 122097217 | Intron |

| | | | |
|---|---|---|---|
| chr3 | 122111372 | 122111714 | Intron |
| chr3 | 122111903 | 122112721 | Intron |
| chr3 | 122117400 | 122117618 | Intron |
| chr3 | 122122288 | 122122839 | Intron |
| chr3 | 122123527 | 122123842 | Intron |
| chr3 | 122124203 | 122124678 | Intron |
| chr3 | 122129264 | 122129463 | Downstream |
| chr3 | 122150697 | 122150928 | Intron |
| chr3 | 122151227 | 122152098 | Intron |
| chr3 | 122160664 | 122162677 | Exon |
| chr3 | 122165168 | 122166413 | Intron |
| chr3 | 122166519 | 122166718 | Intron |
| chr3 | 122174088 | 122174798 | Intron |
| chr3 | 122177821 | 122179439 | Promoter |
| chr3 | 122184012 | 122184211 | Intron |
| chr3 | 122184409 | 122185638 | Intron |
| chr3 | 122186070 | 122186422 | 5' UTR |
| chr3 | 122189358 | 122189575 | Intron |
| chr3 | 122190554 | 122191617 | Intron |
| chr3 | 122192154 | 122192393 | Intron |
| chr3 | 122200464 | 122200836 | Intron |
| chr3 | 122200931 | 122201911 | Intron |
| chr3 | 122211998 | 122213467 | Intron |
| chr3 | 122214698 | 122214897 | Promoter |
| chr3 | 122224046 | 122224608 | Intron |
| chr3 | 122227466 | 122227831 | Intron |
| chr3 | 122227870 | 122228212 | Intron |
| chr3 | 122228402 | 122228972 | Intron |
| chr3 | 122229184 | 122229430 | Intron |
| chr3 | 122229656 | 122230407 | Intron |
| chr3 | 122289416 | 122290368 | 3' UTR |
| chr3 | 122322170 | 122322372 | Intron |
| chr3 | 122623799 | 122624780 | Distal Intergenic |
| chr3 | 122756393 | 122756806 | Distal Intergenic |
| chr3 | 122757014 | 122757309 | Distal Intergenic |
| chr3 | 123041770 | 123041986 | Intron |
| chr3 | 123134654 | 123135223 | Promoter |
| chr3 | 123143976 | 123144340 | Intron |
| chr3 | 123144628 | 123144836 | Intron |
| chr3 | 123208464 | 123208797 | Distal Intergenic |
| chr3 | 123208908 | 123210373 | Downstream |
| chr3 | 123210433 | 123211304 | Downstream |
| chr3 | 123211554 | 123211792 | Downstream |
| chr3 | 123211843 | 123212209 | Downstream |

| | | | |
|---|---|---|---|
| chr3 | 126087645 | 126088378 | Distal Intergenic |
| chr3 | 126363508 | 126364127 | Intron |
| chr3 | 126447524 | 126449034 | Intron |
| chr3 | 126530736 | 126530956 | Intron |
| chr3 | 126532499 | 126533981 | Intron |
| chr3 | 126534021 | 126535083 | Intron |
| chr3 | 126535230 | 126535429 | Intron |
| chr3 | 126571882 | 126572487 | Intron |
| chr3 | 126572575 | 126573536 | Intron |
| chr3 | 127482750 | 127482979 | Intron |
| chr3 | 127483069 | 127483472 | Intron |
| chr3 | 127499965 | 127500789 | Exon |
| chr3 | 127803346 | 127803953 | Intron |
| chr3 | 127807254 | 127807653 | Intron |
| chr3 | 127807708 | 127808318 | Intron |
| chr3 | 127808423 | 127809135 | Intron |
| chr3 | 127809559 | 127809950 | Intron |
| chr3 | 127854320 | 127854538 | Intron |
| chr3 | 127854639 | 127855016 | Intron |
| chr3 | 127855203 | 127855653 | Intron |
| chr3 | 129236246 | 129236511 | Exon |
| chr3 | 129236550 | 129236767 | Intron |
| chr3 | 129236841 | 129237670 | Intron |
| chr3 | 129238003 | 129239336 | 3' UTR |
| chr3 | 129239449 | 129239761 | Distal Intergenic |
| chr3 | 130953373 | 130953995 | Intron |
| chr3 | 130955462 | 130956444 | Intron |
| chr3 | 131530543 | 131530761 | Intron |
| chr3 | 131890310 | 131890780 | Intron |
| chr3 | 132550259 | 132550525 | Intron |
| chr3 | 133263163 | 133264197 | Distal Intergenic |
| chr3 | 133287729 | 133288749 | Distal Intergenic |
| chr3 | 133289048 | 133289634 | Distal Intergenic |
| chr3 | 133289804 | 133290559 | Distal Intergenic |
| chr3 | 135102662 | 135103369 | Distal Intergenic |
| chr3 | 135255719 | 135256999 | Distal Intergenic |
| chr3 | 135257061 | 135257461 | Distal Intergenic |
| chr3 | 135257682 | 135258217 | Distal Intergenic |
| chr3 | 135304166 | 135304420 | Distal Intergenic |
| chr3 | 135314207 | 135315023 | Distal Intergenic |
| chr3 | 135333499 | 135334252 | Distal Intergenic |
| chr3 | 135334429 | 135334659 | Distal Intergenic |
| chr3 | 135335311 | 135338039 | Distal Intergenic |
| chr3 | 135352367 | 135353143 | Distal Intergenic |

| | | | |
|---|---|---|---|
| chr3 | 135353305 | 135353506 | Distal Intergenic |
| chr3 | 135353636 | 135354083 | Distal Intergenic |
| chr3 | 135375021 | 135375687 | Distal Intergenic |
| chr3 | 136094616 | 136095282 | Intron |
| chr3 | 136102972 | 136103271 | Intron |
| chr3 | 136346715 | 136347523 | Intron |
| chr3 | 136351597 | 136352162 | Intron |
| chr3 | 136354602 | 136354853 | Intron |
| chr3 | 137806924 | 137807179 | Intron |
| chr3 | 137807255 | 137807771 | Exon |
| chr3 | 141981347 | 141981621 | Distal Intergenic |
| chr3 | 142035353 | 142035636 | Intron |
| chr3 | 142035908 | 142036225 | Intron |
| chr3 | 142036338 | 142036590 | Intron |
| chr3 | 142051342 | 142051572 | Exon |
| chr3 | 142054780 | 142055178 | Intron |
| chr3 | 143790715 | 143791039 | Distal Intergenic |
| chr3 | 143866790 | 143867168 | Distal Intergenic |
| chr3 | 143973640 | 143973907 | Distal Intergenic |
| chr3 | 144094746 | 144095975 | Distal Intergenic |
| chr3 | 144325879 | 144326146 | Distal Intergenic |
| chr3 | 144326346 | 144326762 | Distal Intergenic |
| chr3 | 144656508 | 144656734 | Distal Intergenic |
| chr3 | 145314588 | 145314787 | Distal Intergenic |
| chr3 | 145579725 | 145579954 | Distal Intergenic |
| chr3 | 145580003 | 145580202 | Distal Intergenic |
| chr3 | 145587911 | 145588627 | Distal Intergenic |
| chr3 | 146163567 | 146163842 | Intron |
| chr3 | 151785383 | 151785723 | Distal Intergenic |
| chr3 | 151785885 | 151786128 | Distal Intergenic |
| chr3 | 157561455 | 157561974 | Distal Intergenic |
| chr3 | 157562028 | 157562311 | Distal Intergenic |
| chr3 | 159503429 | 159503839 | Intron |
| chr3 | 159509836 | 159511205 | Intron |
| chr3 | 159511388 | 159511705 | Intron |
| chr4 | 3424199 | 3424951 | Exon |
| chr4 | 3501762 | 3502087 | Exon |
| chr4 | 3684564 | 3685398 | Distal Intergenic |
| chr4 | 3760126 | 3760339 | Distal Intergenic |
| chr4 | 3761711 | 3762496 | Distal Intergenic |
| chr4 | 3793453 | 3793915 | Distal Intergenic |
| chr4 | 4561963 | 4562229 | Intron |
| chr4 | 4562286 | 4562698 | Intron |
| chr4 | 4562764 | 4563137 | Intron |

| | | | |
|---|---|---|---|
| chr4 | 4720087 | 4720505 | Distal Intergenic |
| chr4 | 5451594 | 5452627 | Intron |
| chr4 | 5649883 | 5650103 | Intron |
| chr4 | 5652226 | 5652769 | Intron |
| chr4 | 5695298 | 5695876 | Intron |
| chr4 | 5695932 | 5696547 | 5' UTR |
| chr4 | 5699684 | 5700139 | Intron |
| chr4 | 5719516 | 5720848 | Intron |
| chr4 | 5763238 | 5763583 | Intron |
| chr4 | 6293180 | 6293866 | Exon |
| chr4 | 6363820 | 6364355 | Intron |
| chr4 | 6364492 | 6364691 | Intron |
| chr4 | 6379815 | 6380878 | Exon |
| chr4 | 6380934 | 6381133 | Intron |
| chr4 | 8426141 | 8426779 | Intron |
| chr4 | 8437699 | 8437909 | Intron |
| chr4 | 8443522 | 8444064 | Promoter |
| chr4 | 8444163 | 8444625 | Intron |
| chr4 | 8463434 | 8463806 | Intron |
| chr4 | 8840234 | 8840483 | Distal Intergenic |
| chr4 | 8931766 | 8931985 | Distal Intergenic |
| chr4 | 8994949 | 8995544 | Distal Intergenic |
| chr4 | 9031093 | 9031298 | Distal Intergenic |
| chr4 | 10192740 | 10193041 | Distal Intergenic |
| chr4 | 10193106 | 10193598 | Distal Intergenic |
| chr4 | 10870046 | 10870842 | Distal Intergenic |
| chr4 | 11021508 | 11022062 | Distal Intergenic |
| chr4 | 11054200 | 11054477 | Distal Intergenic |
| chr4 | 11054517 | 11054723 | Distal Intergenic |
| chr4 | 11082255 | 11082474 | Distal Intergenic |
| chr4 | 11082522 | 11083428 | Distal Intergenic |
| chr4 | 11249428 | 11252068 | Distal Intergenic |
| chr4 | 11312822 | 11313967 | Distal Intergenic |
| chr4 | 11314053 | 11315342 | Distal Intergenic |
| chr4 | 11346226 | 11346980 | Distal Intergenic |
| chr4 | 11350800 | 11351017 | Distal Intergenic |
| chr4 | 11354959 | 11355201 | Distal Intergenic |
| chr4 | 11381980 | 11382267 | Distal Intergenic |
| chr4 | 11463325 | 11463609 | Distal Intergenic |
| chr4 | 11463726 | 11464156 | Distal Intergenic |
| chr4 | 11618840 | 11619252 | Distal Intergenic |
| chr4 | 11917200 | 11917980 | Distal Intergenic |
| chr4 | 14783669 | 14784809 | Intron |
| chr4 | 14790472 | 14791273 | Intron |

| | | | |
|---|---|---|---|
| chr4 | 14791396 | 14792521 | Intron |
| chr4 | 14825668 | 14826082 | Intron |
| chr4 | 14937485 | 14938116 | Intron |
| chr4 | 15126313 | 15127884 | Distal Intergenic |
| chr4 | 15132120 | 15133471 | Distal Intergenic |
| chr4 | 15998082 | 15998926 | Intron |
| chr4 | 16095318 | 16096242 | Distal Intergenic |
| chr4 | 16126809 | 16127240 | Distal Intergenic |
| chr4 | 16311342 | 16311586 | Distal Intergenic |
| chr4 | 17491835 | 17493069 | Exon |
| chr4 | 17494085 | 17494338 | Intron |
| chr4 | 17494436 | 17494641 | Intron |
| chr4 | 17495735 | 17497339 | Intron |
| chr4 | 19452132 | 19453032 | Distal Intergenic |
| chr4 | 19491044 | 19491913 | Distal Intergenic |
| chr4 | 20089393 | 20090523 | Distal Intergenic |
| chr4 | 20121435 | 20123304 | Distal Intergenic |
| chr4 | 20127899 | 20128122 | Distal Intergenic |
| chr4 | 20326038 | 20326804 | Intron |
| chr4 | 20329959 | 20330258 | Intron |
| chr4 | 20567103 | 20567358 | Intron |
| chr4 | 20603314 | 20604406 | Intron |
| chr4 | 21122733 | 21122942 | Intron |
| chr4 | 21420832 | 21421262 | Intron |
| chr4 | 21421577 | 21421830 | Intron |
| chr4 | 21654871 | 21655139 | Intron |
| chr4 | 21774980 | 21775663 | Intron |
| chr4 | 21806979 | 21807214 | Intron |
| chr4 | 23793454 | 23793769 | 3' UTR |
| chr4 | 23881339 | 23881549 | Intron |
| chr4 | 25742864 | 25743087 | Distal Intergenic |
| chr4 | 25743469 | 25745564 | Distal Intergenic |
| chr4 | 25810519 | 25811126 | Intron |
| chr4 | 25811207 | 25811568 | Intron |
| chr4 | 25811763 | 25811962 | Intron |
| chr4 | 26142894 | 26144457 | Distal Intergenic |
| chr4 | 26185987 | 26187198 | Distal Intergenic |
| chr4 | 26187244 | 26187702 | Distal Intergenic |
| chr4 | 29010177 | 29010526 | Distal Intergenic |
| chr4 | 29508979 | 29509178 | Distal Intergenic |
| chr4 | 29509406 | 29509651 | Distal Intergenic |
| chr4 | 31855243 | 31855453 | Distal Intergenic |
| chr4 | 32050371 | 32050899 | Distal Intergenic |
| chr4 | 32051264 | 32051469 | Distal Intergenic |

| | | | |
|---|---|---|---|
| chr4 | 32266994 | 32267293 | Distal Intergenic |
| chr4 | 32337468 | 32338391 | Distal Intergenic |
| chr4 | 32391106 | 32391568 | Distal Intergenic |
| chr4 | 32392199 | 32392427 | Distal Intergenic |
| chr4 | 32406788 | 32407041 | Distal Intergenic |
| chr4 | 32492355 | 32493153 | Distal Intergenic |
| chr4 | 32496205 | 32497200 | Distal Intergenic |
| chr4 | 32744058 | 32744750 | Distal Intergenic |
| chr4 | 32887040 | 32887760 | Distal Intergenic |
| chr4 | 32918703 | 32919494 | Distal Intergenic |
| chr4 | 33117617 | 33117855 | Distal Intergenic |
| chr4 | 33335653 | 33336037 | Distal Intergenic |
| chr4 | 33348008 | 33349777 | Distal Intergenic |
| chr4 | 33360420 | 33360761 | Distal Intergenic |
| chr4 | 33368613 | 33369802 | Distal Intergenic |
| chr4 | 33424552 | 33425606 | Distal Intergenic |
| chr4 | 33461597 | 33462594 | Distal Intergenic |
| chr4 | 33506160 | 33507126 | Distal Intergenic |
| chr4 | 33533057 | 33533313 | Distal Intergenic |
| chr4 | 33533439 | 33533639 | Distal Intergenic |
| chr4 | 33925400 | 33925684 | Intron |
| chr4 | 34156233 | 34156526 | Distal Intergenic |
| chr4 | 34182087 | 34182605 | Distal Intergenic |
| chr4 | 34469705 | 34470234 | Distal Intergenic |
| chr4 | 34470436 | 34470958 | Distal Intergenic |
| chr4 | 34497740 | 34498068 | Distal Intergenic |
| chr4 | 34498253 | 34499443 | Distal Intergenic |
| chr4 | 34561761 | 34562154 | Distal Intergenic |
| chr4 | 34562328 | 34562744 | Distal Intergenic |
| chr4 | 34829619 | 34829872 | Distal Intergenic |
| chr4 | 34937962 | 34938880 | Distal Intergenic |
| chr4 | 34984885 | 34986480 | Distal Intergenic |
| chr4 | 35006324 | 35006607 | Distal Intergenic |
| chr4 | 35008141 | 35008736 | Distal Intergenic |
| chr4 | 35037136 | 35037587 | Distal Intergenic |
| chr4 | 35962621 | 35963058 | Intron |
| chr4 | 35963147 | 35963752 | Intron |
| chr4 | 35982087 | 35982381 | Intron |
| chr4 | 36506512 | 36507146 | Distal Intergenic |
| chr4 | 38690204 | 38690599 | Exon |
| chr4 | 38695135 | 38695612 | Intron |
| chr4 | 39035150 | 39035582 | Distal Intergenic |
| chr4 | 40089707 | 40090269 | Intron |
| chr4 | 40215667 | 40216154 | Intron |

| | | | |
|---|---|---|---|
| chr4 | 40216202 | 40216516 | Intron |
| chr4 | 40226594 | 40226884 | Intron |
| chr4 | 40643221 | 40644688 | Distal Intergenic |
| chr4 | 40647640 | 40647890 | Distal Intergenic |
| chr4 | 40649183 | 40650869 | Distal Intergenic |
| chr4 | 40668641 | 40669385 | Distal Intergenic |
| chr4 | 40720276 | 40720652 | Distal Intergenic |
| chr4 | 40723763 | 40725160 | Distal Intergenic |
| chr4 | 40728792 | 40729666 | Distal Intergenic |
| chr4 | 40745469 | 40746734 | Distal Intergenic |
| chr4 | 40747140 | 40747360 | Distal Intergenic |
| chr4 | 40747813 | 40750586 | Distal Intergenic |
| chr4 | 40764703 | 40766009 | Intron |
| chr4 | 40767565 | 40769227 | Intron |
| chr4 | 40791823 | 40793114 | Exon |
| chr4 | 40793172 | 40795087 | Intron |
| chr4 | 40799006 | 40799532 | Intron |
| chr4 | 40799569 | 40799889 | Intron |
| chr4 | 40800091 | 40800577 | Intron |
| chr4 | 40917667 | 40917866 | Intron |
| chr4 | 40971759 | 40973881 | Intron |
| chr4 | 40997912 | 40999373 | Intron |
| chr4 | 41004189 | 41005345 | Intron |
| chr4 | 41010188 | 41010423 | Intron |
| chr4 | 41010653 | 41013689 | Intron |
| chr4 | 41032951 | 41033266 | Intron |
| chr4 | 41036921 | 41037320 | Intron |
| chr4 | 41040532 | 41040743 | Intron |
| chr4 | 41040838 | 41041996 | Intron |
| chr4 | 41048903 | 41050791 | Intron |
| chr4 | 41055407 | 41056018 | Intron |
| chr4 | 41081815 | 41082385 | Intron |
| chr4 | 41082810 | 41083973 | Intron |
| chr4 | 41099797 | 41099996 | Intron |
| chr4 | 41100152 | 41100460 | Intron |
| chr4 | 41143778 | 41145559 | 5' UTR |
| chr4 | 41213116 | 41213805 | Intron |
| chr4 | 41221407 | 41222080 | Downstream |
| chr4 | 41231391 | 41231621 | Intron |
| chr4 | 41240747 | 41241568 | Intron |
| chr4 | 41242541 | 41242881 | Intron |
| chr4 | 41260962 | 41262758 | 5' UTR |
| chr4 | 41296374 | 41296895 | Distal Intergenic |
| chr4 | 41296940 | 41297241 | Distal Intergenic |

| | | | |
|---|---|---|---|
| chr4 | 41299338 | 41300918 | Distal Intergenic |
| chr4 | 41684764 | 41685617 | Intron |
| chr4 | 41687328 | 41687585 | Intron |
| chr4 | 41687810 | 41688153 | Exon |
| chr4 | 41717171 | 41717613 | Distal Intergenic |
| chr4 | 43012193 | 43013309 | Intron |
| chr4 | 43013468 | 43013906 | Intron |
| chr4 | 43044102 | 43044325 | Distal Intergenic |
| chr4 | 43051716 | 43053164 | Distal Intergenic |
| chr4 | 43053317 | 43053684 | Distal Intergenic |
| chr4 | 43053808 | 43054266 | Distal Intergenic |
| chr4 | 43261741 | 43262057 | Distal Intergenic |
| chr4 | 43273067 | 43273305 | Distal Intergenic |
| chr4 | 43410799 | 43412639 | Distal Intergenic |
| chr4 | 43415077 | 43415823 | Distal Intergenic |
| chr4 | 43432529 | 43433385 | Distal Intergenic |
| chr4 | 43535115 | 43536225 | Distal Intergenic |
| chr4 | 43573899 | 43575359 | Distal Intergenic |
| chr4 | 43616940 | 43618500 | Distal Intergenic |
| chr4 | 43741108 | 43741833 | Distal Intergenic |
| chr4 | 44030774 | 44031019 | Distal Intergenic |
| chr4 | 44107098 | 44107297 | Distal Intergenic |
| chr4 | 44179882 | 44180518 | Intron |
| chr4 | 44181302 | 44181505 | Intron |
| chr4 | 44217829 | 44218784 | Intron |
| chr4 | 44293989 | 44295719 | Intron |
| chr4 | 44768812 | 44769011 | Distal Intergenic |
| chr4 | 44769494 | 44770236 | Distal Intergenic |
| chr4 | 44852199 | 44852441 | Distal Intergenic |
| chr4 | 45023977 | 45024642 | Distal Intergenic |
| chr4 | 45230610 | 45231159 | Distal Intergenic |
| chr4 | 45354853 | 45355105 | Distal Intergenic |
| chr4 | 45467714 | 45468588 | Distal Intergenic |
| chr4 | 45469847 | 45471178 | Distal Intergenic |
| chr4 | 45481837 | 45482411 | Distal Intergenic |
| chr4 | 45485900 | 45486867 | Distal Intergenic |
| chr4 | 45488052 | 45488753 | Distal Intergenic |
| chr4 | 45496862 | 45497224 | Distal Intergenic |
| chr4 | 45501518 | 45503818 | Distal Intergenic |
| chr4 | 45504327 | 45504541 | Distal Intergenic |
| chr4 | 45504756 | 45506992 | Distal Intergenic |
| chr4 | 45507185 | 45507466 | Distal Intergenic |
| chr4 | 45507716 | 45509155 | Distal Intergenic |
| chr4 | 45516209 | 45518074 | Distal Intergenic |

| | | | |
|---|---|---|---|
| chr4 | 45518127 | 45518820 | Distal Intergenic |
| chr4 | 45519229 | 45519488 | Distal Intergenic |
| chr4 | 45523240 | 45524544 | Distal Intergenic |
| chr4 | 45526364 | 45527926 | Distal Intergenic |
| chr4 | 45541705 | 45542459 | Distal Intergenic |
| chr4 | 46082722 | 46083589 | Intron |
| chr4 | 46150722 | 46151595 | Distal Intergenic |
| chr4 | 46151782 | 46152196 | Distal Intergenic |
| chr4 | 46366289 | 46366560 | Intron |
| chr4 | 46384164 | 46384511 | Intron |
| chr4 | 46402112 | 46402311 | Distal Intergenic |
| chr4 | 46402392 | 46402740 | Distal Intergenic |
| chr4 | 46402889 | 46403118 | Distal Intergenic |
| chr4 | 46403209 | 46403485 | Distal Intergenic |
| chr4 | 46463979 | 46465931 | Distal Intergenic |
| chr4 | 46465994 | 46466219 | Distal Intergenic |
| chr4 | 46498449 | 46499124 | Distal Intergenic |
| chr4 | 46536504 | 46536851 | Distal Intergenic |
| chr4 | 46646347 | 46647065 | Distal Intergenic |
| chr4 | 46652774 | 46653888 | Distal Intergenic |
| chr4 | 46655999 | 46656310 | Distal Intergenic |
| chr4 | 46656511 | 46657038 | Distal Intergenic |
| chr4 | 46661295 | 46661585 | Distal Intergenic |
| chr4 | 46661683 | 46662874 | Distal Intergenic |
| chr4 | 47263997 | 47264996 | Intron |
| chr4 | 47323465 | 47324123 | Intron |
| chr4 | 47341457 | 47342702 | Intron |
| chr4 | 47343125 | 47343331 | Intron |
| chr4 | 47419014 | 47419366 | Intron |
| chr4 | 47419569 | 47420845 | Intron |
| chr4 | 47487286 | 47487533 | Promoter |
| chr4 | 48238661 | 48242552 | Intron |
| chr4 | 48243182 | 48243415 | Intron |
| chr4 | 48244398 | 48246440 | Intron |
| chr4 | 48254648 | 48258438 | Intron |
| chr4 | 48266773 | 48267292 | Intron |
| chr4 | 48289783 | 48289995 | Distal Intergenic |
| chr4 | 48290087 | 48290742 | Distal Intergenic |
| chr4 | 48291272 | 48291790 | Distal Intergenic |
| chr4 | 48292681 | 48292883 | Distal Intergenic |
| chr4 | 48485758 | 48485957 | Promoter |
| chr4 | 48552850 | 48553147 | Intron |
| chr4 | 48553446 | 48554293 | Exon |
| chr4 | 48554388 | 48555744 | Exon |

| | | | |
|---|---|---|---|
| chr4 | 48556024 | 48556223 | Intron |
| chr4 | 48598745 | 48599183 | Intron |
| chr4 | 48917338 | 48919231 | Distal Intergenic |
| chr4 | 49533521 | 49533942 | Distal Intergenic |
| chr4 | 52451842 | 52452407 | Distal Intergenic |
| chr4 | 52452473 | 52452735 | Distal Intergenic |
| chr4 | 53319900 | 53320171 | Distal Intergenic |
| chr4 | 53320233 | 53320601 | Distal Intergenic |
| chr4 | 53422280 | 53422991 | Distal Intergenic |
| chr4 | 53452604 | 53453027 | Distal Intergenic |
| chr4 | 53458064 | 53459863 | 3' UTR |
| chr4 | 53464565 | 53465250 | 3' UTR |
| chr4 | 54034191 | 54034724 | Intron |
| chr4 | 54034935 | 54035259 | Intron |
| chr4 | 54353505 | 54353705 | Intron |
| chr4 | 54353794 | 54354365 | Intron |
| chr4 | 54368375 | 54368850 | Intron |
| chr4 | 54368896 | 54369113 | Intron |
| chr4 | 54369194 | 54369437 | Intron |
| chr4 | 54369485 | 54369693 | Intron |
| chr4 | 54960907 | 54961274 | Intron |
| chr4 | 54961610 | 54961813 | Intron |
| chr4 | 54962121 | 54962403 | Intron |
| chr4 | 54977840 | 54979481 | Intron |
| chr4 | 54980249 | 54981238 | Intron |
| chr4 | 54984381 | 54985417 | Intron |
| chr4 | 54985564 | 54986306 | Intron |
| chr4 | 54992253 | 54992787 | Intron |
| chr4 | 54992835 | 54993265 | Intron |
| chr4 | 55013578 | 55013793 | Intron |
| chr4 | 55013904 | 55015108 | Intron |
| chr4 | 55218784 | 55219579 | Distal Intergenic |
| chr4 | 55264393 | 55264945 | Distal Intergenic |
| chr4 | 55266280 | 55267326 | Distal Intergenic |
| chr4 | 55362257 | 55362583 | Distal Intergenic |
| chr4 | 55362967 | 55364171 | Distal Intergenic |
| chr4 | 55364334 | 55364536 | Distal Intergenic |
| chr4 | 55365214 | 55365576 | Distal Intergenic |
| chr4 | 55366495 | 55366717 | Distal Intergenic |
| chr4 | 55366771 | 55366996 | Distal Intergenic |
| chr4 | 55368555 | 55370815 | Distal Intergenic |
| chr4 | 55544313 | 55544958 | Intron |
| chr4 | 55545715 | 55546720 | Intron |
| chr4 | 55546792 | 55547850 | Intron |

| | | | |
|---|---|---|---|
| chr4 | 55547916 | 55549363 | Intron |
| chr4 | 55734362 | 55735420 | Distal Intergenic |
| chr4 | 55744922 | 55745268 | Distal Intergenic |
| chr4 | 55849899 | 55850135 | Distal Intergenic |
| chr4 | 56031344 | 56031649 | Distal Intergenic |
| chr4 | 56227578 | 56228662 | Intron |
| chr4 | 56318382 | 56319105 | Intron |
| chr4 | 56386220 | 56388212 | Intron |
| chr4 | 56399860 | 56400494 | Intron |
| chr4 | 56492225 | 56492509 | Intron |
| chr4 | 56493223 | 56498256 | Promoter |
| chr4 | 56500483 | 56500979 | Intron |
| chr4 | 56501629 | 56503310 | Promoter |
| chr4 | 56504651 | 56505470 | Distal Intergenic |
| chr4 | 56505534 | 56505805 | Distal Intergenic |
| chr4 | 56545526 | 56547018 | Distal Intergenic |
| chr4 | 56554664 | 56554912 | Distal Intergenic |
| chr4 | 56594054 | 56597341 | Distal Intergenic |
| chr4 | 56611496 | 56611804 | Distal Intergenic |
| chr4 | 56618625 | 56619559 | Distal Intergenic |
| chr4 | 56620812 | 56626011 | Distal Intergenic |
| chr4 | 56629910 | 56631360 | Distal Intergenic |
| chr4 | 56631579 | 56631824 | Distal Intergenic |
| chr4 | 56632219 | 56635934 | Distal Intergenic |
| chr4 | 56643846 | 56644263 | Distal Intergenic |
| chr4 | 56650386 | 56650600 | Distal Intergenic |
| chr4 | 56651201 | 56653834 | Distal Intergenic |
| chr4 | 56656074 | 56657922 | Distal Intergenic |
| chr4 | 56674039 | 56674274 | Distal Intergenic |
| chr4 | 56674476 | 56675117 | Distal Intergenic |
| chr4 | 56675155 | 56675411 | Distal Intergenic |
| chr4 | 56680777 | 56682595 | Distal Intergenic |
| chr4 | 56685543 | 56687440 | Promoter |
| chr4 | 56693800 | 56694550 | Intron |
| chr4 | 56876587 | 56877505 | Intron |
| chr4 | 56949560 | 56951211 | Distal Intergenic |
| chr4 | 56958865 | 56959298 | Distal Intergenic |
| chr4 | 56959341 | 56959640 | Distal Intergenic |
| chr4 | 57165879 | 57167292 | Intron |
| chr4 | 57313692 | 57314174 | Intron |
| chr4 | 57350368 | 57351655 | Exon |
| chr4 | 57365662 | 57366933 | Promoter |
| chr4 | 57572751 | 57573225 | Distal Intergenic |
| chr4 | 57623855 | 57624323 | Distal Intergenic |

| | | | |
|---|---|---|---|
| chr4 | 57810515 | 57812324 | Intron |
| chr4 | 57814496 | 57814877 | Intron |
| chr4 | 57814939 | 57815692 | Intron |
| chr4 | 57817315 | 57818487 | Intron |
| chr4 | 57967496 | 57968895 | Intron |
| chr4 | 57969339 | 57970323 | Intron |
| chr4 | 58024003 | 58024224 | Intron |
| chr4 | 58024426 | 58024651 | Intron |
| chr4 | 58044889 | 58045359 | Intron |
| chr4 | 58045481 | 58046383 | Intron |
| chr4 | 58052072 | 58053130 | Intron |
| chr4 | 58414698 | 58415205 | Distal Intergenic |
| chr4 | 58837045 | 58837334 | Distal Intergenic |
| chr4 | 58924341 | 58924768 | Distal Intergenic |
| chr4 | 58925250 | 58925978 | Distal Intergenic |
| chr4 | 59251848 | 59253557 | Distal Intergenic |
| chr4 | 59412085 | 59412483 | Distal Intergenic |
| chr4 | 59446869 | 59448291 | Distal Intergenic |
| chr4 | 59462838 | 59464481 | Distal Intergenic |
| chr4 | 59466290 | 59468438 | Distal Intergenic |
| chr4 | 59470219 | 59471408 | Distal Intergenic |
| chr4 | 59537733 | 59538055 | Distal Intergenic |
| chr4 | 59538123 | 59538421 | Distal Intergenic |
| chr4 | 59539568 | 59540030 | Distal Intergenic |
| chr4 | 59559952 | 59560409 | Distal Intergenic |
| chr4 | 59560577 | 59560966 | Distal Intergenic |
| chr4 | 59561525 | 59562108 | Distal Intergenic |
| chr4 | 59585387 | 59586034 | Distal Intergenic |
| chr4 | 59596650 | 59597172 | Distal Intergenic |
| chr4 | 59664970 | 59665959 | Distal Intergenic |
| chr4 | 59666007 | 59666276 | Distal Intergenic |
| chr4 | 59768982 | 59770996 | Distal Intergenic |
| chr4 | 59796376 | 59796641 | Distal Intergenic |
| chr4 | 59828266 | 59828673 | Distal Intergenic |
| chr4 | 59834825 | 59835146 | Distal Intergenic |
| chr4 | 61861498 | 61861974 | Distal Intergenic |
| chr4 | 61866665 | 61866883 | Distal Intergenic |
| chr4 | 61868993 | 61869268 | Distal Intergenic |
| chr4 | 61948003 | 61949996 | Distal Intergenic |
| chr4 | 61950220 | 61950419 | Distal Intergenic |
| chr4 | 62020833 | 62021111 | Distal Intergenic |
| chr4 | 62021656 | 62021925 | Distal Intergenic |
| chr4 | 62022060 | 62022504 | Distal Intergenic |
| chr4 | 62040019 | 62040227 | Distal Intergenic |

| | | | |
|---|---|---|---|
| chr4 | 62059761 | 62060705 | Distal Intergenic |
| chr4 | 62068412 | 62070622 | Intron |
| chr4 | 62133199 | 62134377 | Intron |
| chr4 | 62181073 | 62181413 | Intron |
| chr4 | 62186149 | 62186348 | Intron |
| chr4 | 62324546 | 62325136 | Intron |
| chr4 | 62391902 | 62394519 | Intron |
| chr4 | 62399234 | 62399912 | Intron |
| chr4 | 62400448 | 62400942 | Intron |
| chr4 | 62984228 | 62985029 | Distal Intergenic |
| chr4 | 63004140 | 63004376 | Distal Intergenic |
| chr4 | 63004426 | 63005651 | Distal Intergenic |
| chr4 | 63205875 | 63206384 | Distal Intergenic |
| chr4 | 63413884 | 63414806 | Distal Intergenic |
| chr4 | 63457386 | 63458461 | Distal Intergenic |
| chr4 | 63619300 | 63620825 | Distal Intergenic |
| chr4 | 63638104 | 63639667 | Distal Intergenic |
| chr4 | 63670688 | 63671998 | Distal Intergenic |
| chr4 | 63672114 | 63673358 | Distal Intergenic |
| chr4 | 63673438 | 63673757 | Distal Intergenic |
| chr4 | 63674032 | 63674233 | Distal Intergenic |
| chr4 | 63674694 | 63676272 | Distal Intergenic |
| chr4 | 63737305 | 63738069 | Distal Intergenic |
| chr4 | 63738894 | 63740151 | Distal Intergenic |
| chr4 | 63744117 | 63745584 | Distal Intergenic |
| chr4 | 63748674 | 63748944 | Distal Intergenic |
| chr4 | 63755601 | 63755931 | Distal Intergenic |
| chr4 | 63758787 | 63760121 | Distal Intergenic |
| chr4 | 63785707 | 63786000 | Distal Intergenic |
| chr4 | 63793222 | 63793791 | Distal Intergenic |
| chr4 | 63836259 | 63838050 | Distal Intergenic |
| chr4 | 63915408 | 63916644 | Distal Intergenic |
| chr4 | 64464960 | 64465635 | Distal Intergenic |
| chr4 | 64558603 | 64559827 | Distal Intergenic |
| chr4 | 64559863 | 64560118 | Distal Intergenic |
| chr4 | 64560679 | 64563320 | Distal Intergenic |
| chr4 | 65356868 | 65357424 | Distal Intergenic |
| chr4 | 65360516 | 65361649 | Distal Intergenic |
| chr4 | 65778868 | 65779614 | Downstream |
| chr4 | 65779706 | 65780005 | Exon |
| chr4 | 65787454 | 65789472 | Intron |
| chr4 | 65849043 | 65849297 | Intron |
| chr4 | 65849486 | 65850578 | Intron |
| chr4 | 65998272 | 65998753 | Distal Intergenic |

| | | | |
|---|---|---|---|
| chr4 | 66488429 | 66490193 | Intron |
| chr4 | 70191064 | 70191478 | Distal Intergenic |
| chr4 | 70191613 | 70192239 | Distal Intergenic |
| chr4 | 70196311 | 70197990 | Distal Intergenic |
| chr4 | 71801540 | 71801773 | Intron |
| chr4 | 71836848 | 71838450 | Intron |
| chr4 | 72317605 | 72317831 | Intron |
| chr4 | 72325637 | 72326105 | Intron |
| chr4 | 72331108 | 72331656 | Intron |
| chr4 | 72331719 | 72331941 | Intron |
| chr4 | 72331986 | 72333293 | Exon |
| chr4 | 76070439 | 76070682 | Distal Intergenic |
| chr4 | 77881283 | 77881639 | Intron |
| chr4 | 80562650 | 80563760 | Distal Intergenic |
| chr4 | 80563859 | 80565272 | Distal Intergenic |
| chr4 | 80596131 | 80596414 | Distal Intergenic |
| chr4 | 80617362 | 80618319 | Distal Intergenic |
| chr4 | 80817058 | 80817677 | Distal Intergenic |
| chr4 | 80817724 | 80818157 | Distal Intergenic |
| chr4 | 80898716 | 80898917 | 3' UTR |
| chr4 | 80954920 | 80955152 | Intron |
| chr4 | 80955213 | 80956203 | Intron |
| chr4 | 81189828 | 81190027 | Intron |
| chr4 | 81572059 | 81573313 | Intron |
| chr4 | 81694572 | 81694884 | Intron |
| chr4 | 81811809 | 81812098 | Intron |
| chr4 | 81812297 | 81812954 | Intron |
| chr4 | 82111826 | 82112146 | Intron |
| chr4 | 82112280 | 82113363 | Intron |
| chr4 | 82116156 | 82118086 | Intron |
| chr4 | 82149200 | 82149399 | Distal Intergenic |
| chr4 | 82149568 | 82149874 | Distal Intergenic |
| chr4 | 82150089 | 82150301 | Distal Intergenic |
| chr4 | 82937888 | 82938120 | Distal Intergenic |
| chr4 | 82947181 | 82949461 | Distal Intergenic |
| chr4 | 82949502 | 82949990 | Distal Intergenic |
| chr4 | 82961073 | 82962096 | Distal Intergenic |
| chr4 | 82962246 | 82962445 | Distal Intergenic |
| chr4 | 82969054 | 82969363 | Distal Intergenic |
| chr4 | 82969401 | 82969724 | Distal Intergenic |
| chr4 | 82978137 | 82979760 | Distal Intergenic |
| chr4 | 83019185 | 83019424 | Distal Intergenic |
| chr4 | 83019555 | 83020919 | Distal Intergenic |
| chr4 | 83021269 | 83021591 | Distal Intergenic |

| | | | |
|---|---|---|---|
| chr4 | 83024992 | 83026907 | Distal Intergenic |
| chr4 | 83062895 | 83063510 | Distal Intergenic |
| chr4 | 83064100 | 83064631 | Distal Intergenic |
| chr4 | 84030844 | 84031835 | Promoter |
| chr4 | 86280545 | 86281059 | Distal Intergenic |
| chr4 | 86314968 | 86315353 | Distal Intergenic |
| chr4 | 86315458 | 86315916 | Distal Intergenic |
| chr4 | 86316162 | 86316576 | Distal Intergenic |
| chr4 | 86317650 | 86319520 | Distal Intergenic |
| chr4 | 86319728 | 86321909 | Distal Intergenic |
| chr4 | 86322018 | 86322428 | Distal Intergenic |
| chr4 | 86322688 | 86323109 | Distal Intergenic |
| chr4 | 86325241 | 86325723 | Distal Intergenic |
| chr4 | 86356657 | 86357824 | Distal Intergenic |
| chr4 | 86357893 | 86360062 | Distal Intergenic |
| chr4 | 86502603 | 86503464 | Intron |
| chr4 | 87466546 | 87467758 | Intron |
| chr4 | 87484232 | 87484899 | Intron |
| chr4 | 87485216 | 87485551 | Intron |
| chr4 | 87495428 | 87497032 | Intron |
| chr4 | 87630298 | 87630689 | Intron |
| chr4 | 87630730 | 87631394 | Intron |
| chr4 | 87725249 | 87725454 | Intron |
| chr4 | 88161089 | 88161999 | Distal Intergenic |
| chr4 | 88176208 | 88176475 | Distal Intergenic |
| chr4 | 88176616 | 88177046 | Distal Intergenic |
| chr4 | 88215664 | 88216002 | Distal Intergenic |
| chr4 | 88287671 | 88288163 | Intron |
| chr4 | 88368529 | 88368771 | Intron |
| chr4 | 88552647 | 88552899 | Distal Intergenic |
| chr4 | 88574196 | 88574862 | Intron |
| chr4 | 88575421 | 88584117 | 5' UTR |
| chr4 | 88584668 | 88590973 | 3' UTR |
| chr4 | 88593670 | 88594084 | Distal Intergenic |
| chr4 | 88594236 | 88594960 | Distal Intergenic |
| chr4 | 88595150 | 88597827 | Distal Intergenic |
| chr4 | 88597928 | 88598176 | Distal Intergenic |
| chr4 | 88601453 | 88603436 | Distal Intergenic |
| chr4 | 88604517 | 88604745 | Distal Intergenic |
| chr4 | 88605156 | 88606411 | Distal Intergenic |
| chr4 | 88606462 | 88606980 | Distal Intergenic |
| chr4 | 88623632 | 88624713 | Distal Intergenic |
| chr4 | 88644411 | 88645303 | Distal Intergenic |
| chr4 | 88646518 | 88648318 | Distal Intergenic |

| | | | |
|---|---|---|---|
| chr4 | 88684092 | 88686158 | Distal Intergenic |
| chr4 | 88686200 | 88686411 | Distal Intergenic |
| chr4 | 88686543 | 88686895 | Distal Intergenic |
| chr4 | 88689939 | 88690406 | Distal Intergenic |
| chr4 | 88694222 | 88694427 | Distal Intergenic |
| chr4 | 88694821 | 88696470 | Distal Intergenic |
| chr4 | 88696574 | 88705508 | Distal Intergenic |
| chr4 | 88705620 | 88705819 | Distal Intergenic |
| chr4 | 88706527 | 88708548 | Distal Intergenic |
| chr4 | 88717702 | 88717990 | Distal Intergenic |
| chr4 | 88718234 | 88718589 | Distal Intergenic |
| chr4 | 88769361 | 88774868 | Distal Intergenic |
| chr4 | 88775049 | 88775349 | Distal Intergenic |
| chr4 | 88880435 | 88881363 | Distal Intergenic |
| chr4 | 88903429 | 88903804 | Exon |
| chr4 | 88941545 | 88941913 | Intron |
| chr4 | 88953948 | 88954334 | Intron |
| chr4 | 88955652 | 88955959 | Intron |
| chr4 | 88956002 | 88956251 | Intron |
| chr4 | 89094882 | 89095641 | Intron |
| chr4 | 89136536 | 89137298 | Intron |
| chr4 | 89137609 | 89137913 | Intron |
| chr4 | 89138345 | 89138860 | Intron |
| chr4 | 89145426 | 89146229 | Intron |
| chr4 | 90119006 | 90119500 | Distal Intergenic |
| chr4 | 93684256 | 93684488 | Intron |
| chr4 | 93761350 | 93761616 | Intron |
| chr4 | 93761765 | 93761994 | Intron |
| chr4 | 93914831 | 93915389 | Intron |
| chr4 | 94269129 | 94270130 | Intron |
| chr4 | 94718453 | 94718778 | Distal Intergenic |
| chr4 | 94719528 | 94720527 | Distal Intergenic |
| chr4 | 94824980 | 94825987 | Distal Intergenic |
| chr4 | 94827656 | 94828113 | Distal Intergenic |
| chr4 | 94828158 | 94829128 | Distal Intergenic |
| chr4 | 94854488 | 94855767 | Distal Intergenic |
| chr4 | 94855809 | 94856198 | Distal Intergenic |
| chr4 | 95050542 | 95050875 | Distal Intergenic |
| chr4 | 95107215 | 95107414 | Distal Intergenic |
| chr4 | 95107609 | 95107835 | Distal Intergenic |
| chr4 | 95327843 | 95328208 | Distal Intergenic |
| chr4 | 95330382 | 95330732 | Distal Intergenic |
| chr4 | 95593052 | 95593565 | Distal Intergenic |
| chr4 | 95593803 | 95594332 | Distal Intergenic |

| | | | |
|---|---|---|---|
| chr4 | 95597855 | 95599084 | Distal Intergenic |
| chr4 | 95689898 | 95690254 | Intron |
| chr4 | 95690299 | 95690526 | Intron |
| chr4 | 95697470 | 95698577 | Intron |
| chr4 | 96233514 | 96234328 | Intron |
| chr4 | 96234624 | 96234836 | Intron |
| chr4 | 96234878 | 96235115 | Intron |
| chr4 | 96457886 | 96458085 | Intron |
| chr4 | 97497901 | 97498728 | Distal Intergenic |
| chr4 | 97498767 | 97499024 | Distal Intergenic |
| chr4 | 97499267 | 97500186 | Distal Intergenic |
| chr4 | 98062887 | 98063337 | Distal Intergenic |
| chr4 | 98074071 | 98074566 | Distal Intergenic |
| chr4 | 98102533 | 98102924 | Distal Intergenic |
| chr4 | 98121743 | 98122631 | Distal Intergenic |
| chr4 | 99354760 | 99355328 | Exon |
| chr4 | 99360790 | 99361490 | Intron |
| chr4 | 99374209 | 99374446 | Distal Intergenic |
| chr4 | 99717822 | 99718674 | Distal Intergenic |
| chr4 | 99946019 | 99947326 | Exon |
| chr4 | 99958069 | 99959265 | Intron |
| chr4 | 99959332 | 99959531 | Intron |
| chr4 | 100059434 | 100059676 | Intron |
| chr4 | 100297397 | 100297612 | Distal Intergenic |
| chr4 | 101204136 | 101205456 | Distal Intergenic |
| chr4 | 101434995 | 101435353 | Intron |
| chr4 | 102183707 | 102185097 | Intron |
| chr4 | 102185144 | 102186008 | Intron |
| chr4 | 102186054 | 102186263 | Intron |
| chr4 | 102205762 | 102205979 | Intron |
| chr4 | 102229433 | 102229632 | Intron |
| chr4 | 102242699 | 102243923 | Intron |
| chr4 | 102244367 | 102245114 | Intron |
| chr4 | 102245300 | 102245811 | Intron |
| chr4 | 102245852 | 102246295 | Intron |
| chr4 | 102249752 | 102250662 | Intron |
| chr4 | 102254518 | 102254907 | Intron |
| chr4 | 102559340 | 102559641 | Intron |
| chr4 | 102792345 | 102792759 | Intron |
| chr4 | 103252559 | 103252784 | Intron |
| chr4 | 104590691 | 104591274 | Intron |
| chr4 | 104591702 | 104592367 | Intron |
| chr4 | 104608742 | 104608941 | Intron |
| chr4 | 105035752 | 105036057 | Distal Intergenic |

| | | | |
|---|---|---|---|
| chr4 | 105076231 | 105076918 | Distal Intergenic |
| chr4 | 105091822 | 105092103 | Distal Intergenic |
| chr4 | 105092281 | 105092493 | Distal Intergenic |
| chr4 | 105524267 | 105524807 | Intron |
| chr4 | 105609881 | 105610239 | Distal Intergenic |
| chr4 | 105720001 | 105720200 | Distal Intergenic |
| chr4 | 105721502 | 105722280 | Distal Intergenic |
| chr4 | 105722512 | 105722946 | Distal Intergenic |
| chr4 | 105973210 | 105974899 | Distal Intergenic |
| chr4 | 105988558 | 105988757 | Distal Intergenic |
| chr4 | 105988877 | 105990709 | Distal Intergenic |
| chr4 | 106223249 | 106223524 | Distal Intergenic |
| chr4 | 106255020 | 106255726 | Distal Intergenic |
| chr4 | 106262478 | 106263326 | Distal Intergenic |
| chr4 | 106511825 | 106512188 | Intron |
| chr4 | 106514181 | 106514498 | Intron |
| chr4 | 106515468 | 106515716 | Intron |
| chr4 | 106520541 | 106520989 | Intron |
| chr4 | 106580988 | 106582383 | Intron |
| chr4 | 106582909 | 106583283 | Intron |
| chr4 | 106619337 | 106621493 | 5' UTR |
| chr4 | 106652881 | 106653946 | Intron |
| chr4 | 106660268 | 106660769 | Intron |
| chr4 | 106850571 | 106850997 | Intron |
| chr4 | 107160929 | 107161725 | Intron |
| chr4 | 107226256 | 107226455 | Intron |
| chr4 | 107537734 | 107537950 | Distal Intergenic |
| chr4 | 107552355 | 107552591 | Distal Intergenic |
| chr4 | 108252090 | 108252456 | Distal Intergenic |
| chr4 | 108452932 | 108453191 | Distal Intergenic |
| chr4 | 108453865 | 108454064 | Distal Intergenic |
| chr4 | 108454186 | 108454541 | Distal Intergenic |
| chr4 | 108455295 | 108455897 | Distal Intergenic |
| chr4 | 108505611 | 108506173 | Distal Intergenic |
| chr4 | 108507011 | 108507619 | Distal Intergenic |
| chr4 | 108520070 | 108520384 | Distal Intergenic |
| chr4 | 108790156 | 108790560 | Intron |
| chr4 | 108885791 | 108886146 | Distal Intergenic |
| chr4 | 108886266 | 108887087 | Distal Intergenic |
| chr4 | 108893317 | 108893616 | Distal Intergenic |
| chr4 | 109132833 | 109133049 | Distal Intergenic |
| chr4 | 109339434 | 109339994 | Distal Intergenic |
| chr4 | 109348553 | 109348802 | Distal Intergenic |
| chr4 | 109349405 | 109349775 | Distal Intergenic |

| | | | |
|---|---|---|---|
| chr4 | 109492909 | 109494005 | Intron |
| chr4 | 111097439 | 111098133 | Intron |
| chr4 | 111106734 | 111107175 | Intron |
| chr4 | 111249911 | 111250177 | Distal Intergenic |
| chr4 | 111250251 | 111250518 | Distal Intergenic |
| chr4 | 114032570 | 114032810 | Intron |
| chr4 | 114033008 | 114033287 | Intron |
| chr4 | 114033826 | 114034073 | Intron |
| chr4 | 114034115 | 114034372 | Intron |
| chr4 | 114218301 | 114218510 | Intron |
| chr4 | 114250579 | 114251123 | Intron |
| chr4 | 114251170 | 114251436 | 5' UTR |
| chr4 | 114262344 | 114262721 | Promoter |
| chr4 | 114273312 | 114275114 | Exon |
| chr4 | 114630865 | 114631280 | Intron |
| chr4 | 114672846 | 114673362 | Intron |
| chr4 | 115747735 | 115747953 | Downstream |
| chr4 | 115748029 | 115748691 | Downstream |
| chr4 | 115748790 | 115749632 | 3' UTR |
| chr4 | 116228828 | 116229436 | Distal Intergenic |
| chr4 | 116229473 | 116229685 | Distal Intergenic |
| chr4 | 116230839 | 116231039 | Distal Intergenic |
| chr4 | 116267581 | 116268530 | Distal Intergenic |
| chr4 | 116268755 | 116269087 | Distal Intergenic |
| chr4 | 116269158 | 116269502 | Distal Intergenic |
| chr4 | 116323356 | 116323968 | Distal Intergenic |
| chr4 | 116324048 | 116324285 | Distal Intergenic |
| chr4 | 116352325 | 116352536 | Distal Intergenic |
| chr4 | 116358238 | 116358445 | Distal Intergenic |
| chr4 | 116422569 | 116422802 | Distal Intergenic |
| chr4 | 116790369 | 116791583 | Distal Intergenic |
| chr4 | 116793448 | 116794162 | Distal Intergenic |
| chr4 | 116795157 | 116795412 | Distal Intergenic |
| chr4 | 116797247 | 116799068 | Distal Intergenic |
| chr4 | 116799126 | 116799705 | Distal Intergenic |
| chr4 | 116799758 | 116800026 | Distal Intergenic |
| chr4 | 116862342 | 116862729 | Distal Intergenic |
| chr4 | 117504718 | 117504917 | Distal Intergenic |
| chr4 | 117619859 | 117620098 | Distal Intergenic |
| chr4 | 117852732 | 117852931 | Distal Intergenic |
| chr4 | 117901768 | 117902194 | Distal Intergenic |
| chr4 | 117902276 | 117902594 | Distal Intergenic |
| chr4 | 117902743 | 117902942 | Distal Intergenic |
| chr4 | 117912914 | 117913299 | Distal Intergenic |

| | | | |
|---|---|---|---|
| chr4 | 117918663 | 117919377 | Distal Intergenic |
| chr4 | 117944322 | 117945560 | Distal Intergenic |
| chr4 | 117946524 | 117946830 | Distal Intergenic |
| chr4 | 117946873 | 117947444 | Distal Intergenic |
| chr4 | 117947652 | 117947971 | Distal Intergenic |
| chr4 | 117948672 | 117949343 | Distal Intergenic |
| chr4 | 117949572 | 117951204 | Distal Intergenic |
| chr4 | 118100214 | 118100605 | Distal Intergenic |
| chr4 | 118100678 | 118101557 | Distal Intergenic |
| chr4 | 118101788 | 118102006 | Distal Intergenic |
| chr4 | 118109668 | 118110275 | Distal Intergenic |
| chr4 | 118594869 | 118596210 | Distal Intergenic |
| chr4 | 118685402 | 118686158 | Distal Intergenic |
| chr4 | 118691180 | 118691824 | Distal Intergenic |
| chr4 | 118692171 | 118692654 | Distal Intergenic |
| chr4 | 118729134 | 118730943 | Distal Intergenic |
| chr4 | 118732186 | 118732433 | Distal Intergenic |
| chr4 | 118739369 | 118740289 | Distal Intergenic |
| chr4 | 118774720 | 118775725 | Distal Intergenic |
| chr4 | 118776837 | 118777113 | Distal Intergenic |
| chr4 | 118777158 | 118777556 | Distal Intergenic |
| chr4 | 118778100 | 118779513 | Distal Intergenic |
| chr4 | 118780843 | 118781169 | Distal Intergenic |
| chr4 | 118781471 | 118781719 | Distal Intergenic |
| chr4 | 118782704 | 118782903 | Distal Intergenic |
| chr4 | 118783627 | 118783826 | Distal Intergenic |
| chr4 | 118791190 | 118792086 | Distal Intergenic |
| chr4 | 118792151 | 118794679 | Distal Intergenic |
| chr4 | 118797171 | 118799242 | Distal Intergenic |
| chr4 | 118799409 | 118799713 | Distal Intergenic |
| chr4 | 118813465 | 118814131 | Distal Intergenic |
| chr4 | 118814171 | 118814401 | Distal Intergenic |
| chr4 | 118830638 | 118831244 | Distal Intergenic |
| chr4 | 119604883 | 119606448 | Promoter |
| chr4 | 119608644 | 119608885 | Intron |
| chr4 | 119627989 | 119628188 | Intron |
| chr4 | 119695161 | 119696057 | Intron |
| chr4 | 119820916 | 119821340 | Intron |
| chr4 | 119842974 | 119843560 | Intron |
| chr4 | 119902372 | 119902589 | Intron |
| chr4 | 119928241 | 119929611 | Intron |
| chr4 | 119942927 | 119945502 | Promoter |
| chr4 | 120145148 | 120145722 | Intron |
| chr4 | 120145850 | 120146082 | Intron |

| | | | |
|---|---|---|---|
| chr4 | 120160113 | 120161294 | Promoter |
| chr4 | 120188757 | 120191605 | Promoter |
| chr4 | 120200534 | 120200787 | Intron |
| chr4 | 120215917 | 120217925 | 3' UTR |
| chr4 | 120240885 | 120241465 | Intron |
| chr4 | 120241665 | 120241876 | Exon |
| chr4 | 120242156 | 120242592 | Promoter |
| chr4 | 120243522 | 120245269 | Promoter |
| chr4 | 120245768 | 120246906 | Distal Intergenic |
| chr4 | 120488501 | 120488711 | Intron |
| chr4 | 122545398 | 122546096 | Distal Intergenic |
| chr4 | 122563452 | 122563721 | Distal Intergenic |
| chr4 | 122632408 | 122633117 | Distal Intergenic |
| chr4 | 122723103 | 122723997 | Promoter |
| chr4 | 123093330 | 123093576 | Intron |
| chr4 | 123099344 | 123099695 | Intron |
| chr4 | 123099811 | 123101558 | Intron |
| chr4 | 123206530 | 123206760 | Intron |
| chr4 | 123206922 | 123207861 | Exon |
| chr4 | 123207932 | 123208228 | Exon |
| chr4 | 123212351 | 123212613 | Intron |
| chr4 | 123229214 | 123229470 | Exon |
| chr4 | 123232579 | 123232843 | Intron |
| chr4 | 123233044 | 123233419 | Intron |
| chr4 | 123233497 | 123233698 | Intron |
| chr4 | 123241206 | 123244008 | Intron |
| chr4 | 123248351 | 123249396 | Exon |
| chr4 | 123249768 | 123251268 | Intron |
| chr4 | 123261860 | 123262434 | Intron |
| chr4 | 123264687 | 123266232 | 3' UTR |
| chr4 | 123266377 | 123267425 | Intron |
| chr4 | 123288226 | 123288431 | Distal Intergenic |
| chr4 | 123288553 | 123289619 | Distal Intergenic |
| chr4 | 123298714 | 123299187 | Promoter |
| chr4 | 123299340 | 123300281 | Promoter |
| chr4 | 123300517 | 123302118 | Promoter |
| chr4 | 123302308 | 123303220 | Exon |
| chr4 | 123312331 | 123313660 | Intron |
| chr4 | 123314029 | 123314998 | Exon |
| chr4 | 123342267 | 123342887 | Exon |
| chr4 | 123343957 | 123344388 | Intron |
| chr4 | 123344574 | 123344785 | Intron |
| chr4 | 123429127 | 123429374 | Distal Intergenic |
| chr4 | 123979974 | 123980308 | Intron |

| | | | |
|---|---|---|---|
| chr4 | 123980467 | 123980997 | Intron |
| chr4 | 124078414 | 124078932 | Intron |
| chr4 | 124106344 | 124106548 | Intron |
| chr4 | 124248076 | 124248427 | Distal Intergenic |
| chr4 | 124352056 | 124352255 | Distal Intergenic |
| chr4 | 124353886 | 124354607 | Distal Intergenic |
| chr4 | 124378720 | 124378927 | Distal Intergenic |
| chr4 | 124379642 | 124379841 | Distal Intergenic |
| chr4 | 124380673 | 124382398 | Distal Intergenic |
| chr4 | 124385621 | 124386744 | Distal Intergenic |
| chr4 | 124475233 | 124475501 | Distal Intergenic |
| chr4 | 124792183 | 124792443 | Intron |
| chr4 | 124793055 | 124795451 | Intron |
| chr4 | 124799272 | 124799993 | Intron |
| chr4 | 124800045 | 124803878 | Intron |
| chr4 | 125779812 | 125780501 | Distal Intergenic |
| chr4 | 125788013 | 125788492 | Distal Intergenic |
| chr4 | 125878741 | 125879378 | Distal Intergenic |
| chr4 | 125894876 | 125895144 | Distal Intergenic |
| chr4 | 125917504 | 125917864 | Distal Intergenic |
| chr4 | 125923482 | 125924597 | Distal Intergenic |
| chr4 | 125927562 | 125928598 | Distal Intergenic |
| chr4 | 125928640 | 125929125 | Distal Intergenic |
| chr4 | 125932328 | 125932810 | Distal Intergenic |
| chr4 | 125933525 | 125934049 | Distal Intergenic |
| chr4 | 125936376 | 125938700 | Distal Intergenic |
| chr4 | 126048817 | 126049432 | Distal Intergenic |
| chr4 | 127005681 | 127006569 | Distal Intergenic |
| chr4 | 127006708 | 127007687 | Distal Intergenic |
| chr4 | 127026689 | 127026973 | Distal Intergenic |
| chr4 | 127027048 | 127027553 | Distal Intergenic |
| chr4 | 127030775 | 127032628 | Distal Intergenic |
| chr4 | 128389719 | 128389918 | Distal Intergenic |
| chr4 | 128405380 | 128406093 | Distal Intergenic |
| chr4 | 128406166 | 128406600 | Distal Intergenic |
| chr4 | 128406767 | 128407977 | Distal Intergenic |
| chr4 | 128408586 | 128408861 | Distal Intergenic |
| chr4 | 128410737 | 128411292 | Distal Intergenic |
| chr4 | 128411578 | 128413143 | Distal Intergenic |
| chr4 | 128413244 | 128416149 | Distal Intergenic |
| chr4 | 128426258 | 128426457 | Distal Intergenic |
| chr4 | 128515155 | 128515883 | Distal Intergenic |
| chr4 | 128515944 | 128516151 | Distal Intergenic |
| chr4 | 128516383 | 128517653 | Distal Intergenic |

| | | | |
|---|---|---|---|
| chr4 | 128529531 | 128529991 | Distal Intergenic |
| chr4 | 128531926 | 128532661 | Distal Intergenic |
| chr4 | 128532853 | 128533810 | Distal Intergenic |
| chr4 | 128543644 | 128543862 | Distal Intergenic |
| chr4 | 128634727 | 128634949 | Intron |
| chr4 | 128635022 | 128635221 | Exon |
| chr4 | 128710655 | 128710929 | Intron |
| chr4 | 128711055 | 128711410 | Intron |
| chr4 | 128711885 | 128713717 | Intron |
| chr4 | 128713753 | 128713962 | Intron |
| chr4 | 128732094 | 128733285 | Exon |
| chr4 | 128744881 | 128745653 | Intron |
| chr4 | 128758676 | 128760113 | Exon |
| chr4 | 128881843 | 128882042 | Intron |
| chr4 | 128902967 | 128903348 | Intron |
| chr4 | 128903469 | 128904194 | 5' UTR |
| chr4 | 128908317 | 128911108 | Exon |
| chr4 | 128911559 | 128911959 | Intron |
| chr4 | 128922307 | 128922593 | Intron |
| chr4 | 128924856 | 128925241 | Intron |
| chr4 | 128936024 | 128937191 | Intron |
| chr4 | 128937507 | 128938029 | Intron |
| chr4 | 128938604 | 128939127 | Exon |
| chr4 | 128940262 | 128940471 | Intron |
| chr4 | 128940845 | 128941295 | Exon |
| chr4 | 129093300 | 129095526 | Intron |
| chr4 | 129103304 | 129104211 | Intron |
| chr4 | 129104258 | 129104479 | Intron |
| chr4 | 129317418 | 129317617 | Distal Intergenic |
| chr4 | 129419832 | 129421011 | Intron |
| chr4 | 129488364 | 129489150 | Distal Intergenic |
| chr4 | 129498298 | 129498505 | Distal Intergenic |
| chr4 | 129498564 | 129498966 | Distal Intergenic |
| chr4 | 129499439 | 129499728 | Distal Intergenic |
| chr4 | 129500248 | 129500627 | Distal Intergenic |
| chr4 | 129554733 | 129554932 | Distal Intergenic |
| chr4 | 129727576 | 129727809 | Distal Intergenic |
| chr4 | 129728028 | 129728557 | Distal Intergenic |
| chr4 | 129784885 | 129785084 | 3' UTR |
| chr4 | 129850120 | 129851766 | Intron |
| chr4 | 129851814 | 129852509 | Intron |
| chr4 | 129852669 | 129853120 | Intron |
| chr4 | 129854199 | 129855155 | Intron |
| chr4 | 129946008 | 129946643 | Intron |

| | | | |
|---|---|---|---|
| chr4 | 129950793 | 129951625 | Intron |
| chr4 | 129951974 | 129952391 | Intron |
| chr4 | 130186334 | 130186663 | Distal Intergenic |
| chr4 | 130186700 | 130187116 | Distal Intergenic |
| chr4 | 130722240 | 130722537 | Intron |
| chr4 | 130751396 | 130752184 | Intron |
| chr4 | 131380939 | 131382340 | Distal Intergenic |
| chr4 | 131429883 | 131430609 | Distal Intergenic |
| chr4 | 131767615 | 131767973 | Distal Intergenic |
| chr4 | 131864681 | 131865279 | Distal Intergenic |
| chr4 | 131865339 | 131865560 | Distal Intergenic |
| chr4 | 131865691 | 131866343 | Distal Intergenic |
| chr4 | 131897126 | 131897367 | Distal Intergenic |
| chr4 | 131900460 | 131901136 | Distal Intergenic |
| chr4 | 131909196 | 131909864 | Distal Intergenic |
| chr4 | 131958989 | 131959622 | Distal Intergenic |
| chr4 | 131975742 | 131975967 | Distal Intergenic |
| chr4 | 132080160 | 132080983 | Distal Intergenic |
| chr4 | 132611714 | 132612031 | Distal Intergenic |
| chr4 | 132625715 | 132627067 | Distal Intergenic |
| chr4 | 132668357 | 132668746 | Distal Intergenic |
| chr4 | 132673834 | 132674126 | Distal Intergenic |
| chr4 | 132686674 | 132688344 | Distal Intergenic |
| chr4 | 132740916 | 132741220 | Distal Intergenic |
| chr4 | 132818128 | 132818935 | Distal Intergenic |
| chr4 | 132903610 | 132904647 | Distal Intergenic |
| chr4 | 132909256 | 132909863 | Distal Intergenic |
| chr4 | 132934056 | 132934387 | Distal Intergenic |
| chr4 | 132934428 | 132935089 | Distal Intergenic |
| chr4 | 132961815 | 132963950 | Distal Intergenic |
| chr4 | 132964243 | 132964681 | Distal Intergenic |
| chr4 | 133045453 | 133045814 | Distal Intergenic |
| chr4 | 133047193 | 133047454 | Distal Intergenic |
| chr4 | 133048635 | 133049821 | Distal Intergenic |
| chr4 | 133049914 | 133050490 | Distal Intergenic |
| chr4 | 133140009 | 133140298 | Distal Intergenic |
| chr4 | 133150961 | 133152244 | Distal Intergenic |
| chr4 | 133154261 | 133155575 | Distal Intergenic |
| chr4 | 133155814 | 133156595 | Distal Intergenic |
| chr4 | 133156693 | 133163819 | Distal Intergenic |
| chr4 | 133165579 | 133166061 | Distal Intergenic |
| chr4 | 133169295 | 133170468 | Distal Intergenic |
| chr4 | 133188133 | 133188783 | Distal Intergenic |
| chr4 | 133215531 | 133216244 | Distal Intergenic |

| | | | |
|---|---|---|---|
| chr4 | 133227941 | 133228243 | Distal Intergenic |
| chr4 | 133302235 | 133302973 | Distal Intergenic |
| chr4 | 133307677 | 133308133 | Distal Intergenic |
| chr4 | 133308369 | 133308578 | Distal Intergenic |
| chr4 | 133308687 | 133309466 | Distal Intergenic |
| chr4 | 133371218 | 133373173 | Distal Intergenic |
| chr4 | 133396888 | 133397500 | Distal Intergenic |
| chr4 | 133398807 | 133399006 | Distal Intergenic |
| chr4 | 133399044 | 133399738 | Distal Intergenic |
| chr4 | 133442218 | 133442417 | Distal Intergenic |
| chr4 | 133465889 | 133466639 | Distal Intergenic |
| chr4 | 133496210 | 133497659 | Distal Intergenic |
| chr4 | 133793768 | 133794027 | Distal Intergenic |
| chr4 | 134290407 | 134290861 | Distal Intergenic |
| chr4 | 134305993 | 134306192 | Distal Intergenic |
| chr4 | 134306553 | 134306959 | Distal Intergenic |
| chr4 | 134313038 | 134313474 | Distal Intergenic |
| chr4 | 134313517 | 134313728 | Distal Intergenic |
| chr4 | 134314631 | 134315043 | Distal Intergenic |
| chr4 | 134318060 | 134318751 | Distal Intergenic |
| chr4 | 134318891 | 134319346 | Distal Intergenic |
| chr4 | 134321857 | 134322971 | Distal Intergenic |
| chr4 | 134323013 | 134323424 | Distal Intergenic |
| chr4 | 134531452 | 134531719 | Distal Intergenic |
| chr4 | 134635750 | 134636195 | Distal Intergenic |
| chr4 | 134636561 | 134636780 | Distal Intergenic |
| chr4 | 134827918 | 134828292 | Distal Intergenic |
| chr4 | 134853056 | 134853306 | Distal Intergenic |
| chr4 | 134853516 | 134855493 | Distal Intergenic |
| chr4 | 134908511 | 134908958 | Distal Intergenic |
| chr4 | 134909689 | 134910130 | Distal Intergenic |
| chr4 | 135128877 | 135129097 | Distal Intergenic |
| chr4 | 135131024 | 135131349 | Distal Intergenic |
| chr4 | 135153791 | 135154376 | Distal Intergenic |
| chr4 | 135336102 | 135336950 | Distal Intergenic |
| chr4 | 135337775 | 135338085 | Distal Intergenic |
| chr4 | 135428524 | 135429765 | Distal Intergenic |
| chr4 | 135432404 | 135432725 | Distal Intergenic |
| chr4 | 135612123 | 135612612 | Distal Intergenic |
| chr4 | 135645964 | 135646275 | Distal Intergenic |
| chr4 | 135678652 | 135679647 | Distal Intergenic |
| chr4 | 135696465 | 135697314 | Distal Intergenic |
| chr4 | 135698869 | 135699286 | Distal Intergenic |
| chr4 | 135699375 | 135703219 | Distal Intergenic |

| | | | |
|---|---|---|---|
| chr4 | 135835010 | 135835602 | Distal Intergenic |
| chr4 | 135842558 | 135843463 | Distal Intergenic |
| chr4 | 135843651 | 135843897 | Distal Intergenic |
| chr4 | 135989130 | 135990440 | Distal Intergenic |
| chr4 | 135997050 | 135998186 | Distal Intergenic |
| chr4 | 136026513 | 136026723 | Distal Intergenic |
| chr4 | 136061155 | 136062523 | Distal Intergenic |
| chr4 | 136066971 | 136067858 | Distal Intergenic |
| chr4 | 136073392 | 136073591 | Distal Intergenic |
| chr4 | 136073683 | 136074102 | Distal Intergenic |
| chr4 | 136074370 | 136074615 | Distal Intergenic |
| chr4 | 136154031 | 136154230 | Distal Intergenic |
| chr4 | 136154481 | 136154746 | Distal Intergenic |
| chr4 | 136157647 | 136157894 | Distal Intergenic |
| chr4 | 136357713 | 136358099 | Distal Intergenic |
| chr4 | 136358240 | 136358462 | Distal Intergenic |
| chr4 | 136757370 | 136757783 | Distal Intergenic |
| chr4 | 136758225 | 136758452 | Distal Intergenic |
| chr4 | 136763187 | 136763386 | Distal Intergenic |
| chr4 | 136763618 | 136764091 | Distal Intergenic |
| chr4 | 136782230 | 136782435 | Distal Intergenic |
| chr4 | 136837261 | 136838184 | Distal Intergenic |
| chr4 | 136838406 | 136838659 | Distal Intergenic |
| chr4 | 136839262 | 136840191 | Distal Intergenic |
| chr4 | 136842601 | 136843290 | Distal Intergenic |
| chr4 | 136847487 | 136847757 | Distal Intergenic |
| chr4 | 136847843 | 136848446 | Distal Intergenic |
| chr4 | 136901193 | 136901764 | Distal Intergenic |
| chr4 | 136923258 | 136923582 | Distal Intergenic |
| chr4 | 136943951 | 136944568 | Distal Intergenic |
| chr4 | 136944691 | 136944907 | Distal Intergenic |
| chr4 | 137004507 | 137005265 | Distal Intergenic |
| chr4 | 137025494 | 137027642 | Distal Intergenic |
| chr4 | 137409440 | 137409816 | Distal Intergenic |
| chr4 | 137419534 | 137419790 | Distal Intergenic |
| chr4 | 137420077 | 137420302 | Distal Intergenic |
| chr4 | 137420402 | 137421202 | Distal Intergenic |
| chr4 | 137424951 | 137425763 | Distal Intergenic |
| chr4 | 137452936 | 137453139 | Distal Intergenic |
| chr4 | 137463929 | 137464665 | Distal Intergenic |
| chr4 | 137484755 | 137485124 | Distal Intergenic |
| chr4 | 137485164 | 137486372 | Distal Intergenic |
| chr4 | 137486775 | 137487752 | Distal Intergenic |
| chr4 | 137510694 | 137510936 | Distal Intergenic |

| | | | |
|---|---|---|---|
| chr4 | 137511229 | 137511457 | Distal Intergenic |
| chr4 | 137770103 | 137770305 | Distal Intergenic |
| chr4 | 137773158 | 137773661 | Distal Intergenic |
| chr4 | 137802571 | 137803013 | Distal Intergenic |
| chr4 | 137803522 | 137803765 | Distal Intergenic |
| chr4 | 137805332 | 137805607 | Distal Intergenic |
| chr4 | 137912781 | 137913008 | Distal Intergenic |
| chr4 | 138257507 | 138257712 | Distal Intergenic |
| chr4 | 138257820 | 138258166 | Distal Intergenic |
| chr4 | 138258454 | 138259211 | Distal Intergenic |
| chr4 | 138348184 | 138349068 | Distal Intergenic |
| chr4 | 138351786 | 138352011 | Distal Intergenic |
| chr4 | 138352056 | 138352767 | Distal Intergenic |
| chr4 | 138528526 | 138528788 | Distal Intergenic |
| chr4 | 138772658 | 138772959 | Distal Intergenic |
| chr4 | 138773021 | 138773258 | Distal Intergenic |
| chr4 | 138942638 | 138943265 | Distal Intergenic |
| chr4 | 139037062 | 139038252 | Intron |
| chr4 | 139047977 | 139048286 | Intron |
| chr4 | 139056978 | 139057331 | Intron |
| chr4 | 139077582 | 139077917 | Intron |
| chr4 | 139091510 | 139092720 | 3' UTR |
| chr4 | 139099898 | 139100748 | Exon |
| chr4 | 139132684 | 139133077 | Intron |
| chr4 | 139136168 | 139136402 | Intron |
| chr4 | 139136595 | 139136794 | Intron |
| chr4 | 139137857 | 139138135 | Intron |
| chr4 | 139150582 | 139155422 | Exon |
| chr4 | 139166055 | 139166707 | Distal Intergenic |
| chr4 | 139352910 | 139353812 | Distal Intergenic |
| chr4 | 140222150 | 140222444 | Promoter |
| chr4 | 140255145 | 140255426 | Exon |
| chr4 | 140255594 | 140257111 | Intron |
| chr4 | 140364961 | 140365975 | Distal Intergenic |
| chr4 | 140418190 | 140418395 | Distal Intergenic |
| chr4 | 140694897 | 140695327 | Intron |
| chr4 | 140695436 | 140695635 | Intron |
| chr4 | 140798391 | 140798712 | Intron |
| chr4 | 140798796 | 140799134 | Intron |
| chr4 | 140800368 | 140800655 | Intron |
| chr4 | 140800830 | 140801359 | Intron |
| chr4 | 140815194 | 140815550 | Intron |
| chr4 | 140815616 | 140815826 | Intron |
| chr4 | 140816049 | 140816768 | Intron |

| | | | |
|---|---|---|---|
| chr4 | 140829071 | 140829274 | Intron |
| chr4 | 140829583 | 140830027 | Intron |
| chr4 | 140830139 | 140832055 | Intron |
| chr4 | 140832989 | 140834864 | Intron |
| chr4 | 140834908 | 140835132 | Intron |
| chr4 | 140843303 | 140843548 | Intron |
| chr4 | 140858046 | 140858283 | Intron |
| chr4 | 140859617 | 140860107 | Intron |
| chr4 | 140865596 | 140866129 | Intron |
| chr4 | 141010358 | 141011183 | Intron |
| chr4 | 141013862 | 141014389 | Intron |
| chr4 | 141089255 | 141089485 | Distal Intergenic |
| chr4 | 141102080 | 141102337 | Distal Intergenic |
| chr4 | 141114879 | 141115721 | Distal Intergenic |
| chr4 | 141121134 | 141121649 | Distal Intergenic |
| chr4 | 141148381 | 141148738 | Distal Intergenic |
| chr4 | 141148800 | 141149414 | Distal Intergenic |
| chr4 | 141160130 | 141160670 | Distal Intergenic |
| chr4 | 141297167 | 141298744 | Intron |
| chr4 | 141549209 | 141549563 | Exon |
| chr4 | 141613524 | 141613812 | Intron |
| chr4 | 141621495 | 141622256 | Intron |
| chr4 | 141626497 | 141628377 | Intron |
| chr4 | 141784973 | 141785426 | Downstream |
| chr4 | 142710168 | 142712158 | Distal Intergenic |
| chr4 | 142868271 | 142868737 | Distal Intergenic |
| chr4 | 142878268 | 142878672 | Distal Intergenic |
| chr4 | 142878721 | 142879279 | Distal Intergenic |
| chr4 | 142884197 | 142885671 | Distal Intergenic |
| chr4 | 142885722 | 142886106 | Distal Intergenic |
| chr4 | 142886182 | 142886518 | Distal Intergenic |
| chr4 | 142886555 | 142887043 | Distal Intergenic |
| chr4 | 142895131 | 142895433 | Distal Intergenic |
| chr4 | 143994775 | 143995295 | Distal Intergenic |
| chr4 | 147279969 | 147281490 | Intron |
| chr4 | 147861614 | 147862351 | Intron |
| chr4 | 148000288 | 148000586 | Distal Intergenic |
| chr4 | 148000669 | 148001124 | Distal Intergenic |
| chr4 | 148172004 | 148172208 | Distal Intergenic |
| chr4 | 148423545 | 148424043 | Intron |
| chr4 | 148544042 | 148545839 | Exon |
| chr4 | 148775377 | 148775593 | Intron |
| chr4 | 148779850 | 148780284 | Intron |
| chr4 | 148780394 | 148780647 | Intron |

| | | | |
|---|---|---|---|
| chr4 | 148800495 | 148800694 | Intron |
| chr4 | 148820294 | 148820500 | Intron |
| chr4 | 148859767 | 148860112 | Intron |
| chr4 | 148878678 | 148879088 | Intron |
| chr4 | 148932500 | 148932900 | Intron |
| chr4 | 148945338 | 148947418 | Promoter |
| chr4 | 148966744 | 148967423 | Intron |
| chr4 | 148967610 | 148967857 | Intron |
| chr4 | 148967934 | 148968273 | Exon |
| chr4 | 148981636 | 148982180 | Intron |
| chr4 | 149270516 | 149270715 | Intron |
| chr4 | 149272550 | 149272782 | Intron |
| chr4 | 149272896 | 149273133 | Intron |
| chr4 | 149434986 | 149435192 | Distal Intergenic |
| chr4 | 149611388 | 149611745 | Distal Intergenic |
| chr4 | 149612305 | 149612712 | Distal Intergenic |
| chr4 | 149612951 | 149614108 | Distal Intergenic |
| chr4 | 149656256 | 149656569 | Distal Intergenic |
| chr4 | 149833056 | 149833256 | Distal Intergenic |
| chr4 | 149918759 | 149919129 | Distal Intergenic |
| chr4 | 150234622 | 150235043 | Distal Intergenic |
| chr4 | 150244849 | 150245524 | Distal Intergenic |
| chr4 | 150245660 | 150246067 | Distal Intergenic |
| chr4 | 150246282 | 150246502 | Distal Intergenic |
| chr4 | 150347065 | 150347565 | Distal Intergenic |
| chr4 | 150347601 | 150347917 | Distal Intergenic |
| chr4 | 151393123 | 151393359 | Intron |
| chr4 | 151466508 | 151467388 | Intron |
| chr4 | 151539422 | 151539644 | Intron |
| chr4 | 151559440 | 151561249 | Intron |
| chr4 | 151561342 | 151561584 | Intron |
| chr4 | 151634268 | 151634501 | Intron |
| chr4 | 153464942 | 153465260 | Distal Intergenic |
| chr4 | 153592005 | 153592447 | Intron |
| chr4 | 153610334 | 153610623 | Distal Intergenic |
| chr4 | 154430523 | 154430736 | Intron |
| chr4 | 154430821 | 154432500 | Intron |
| chr4 | 154432607 | 154432905 | Intron |
| chr4 | 154500996 | 154501381 | Intron |
| chr4 | 154503573 | 154503970 | Intron |
| chr4 | 154504075 | 154504301 | Intron |
| chr4 | 154504364 | 154504954 | Exon |
| chr4 | 154507344 | 154507886 | Exon |
| chr4 | 154508401 | 154508642 | Intron |

| | | | |
|---|---|---|---|
| chr4 | 154523307 | 154523708 | Exon |
| chr4 | 154547400 | 154547704 | Intron |
| chr4 | 154547817 | 154548448 | Intron |
| chr4 | 154548664 | 154549233 | Exon |
| chr4 | 154626517 | 154626906 | 3' UTR |
| chr4 | 154649443 | 154649825 | Exon |
| chr4 | 154650018 | 154650395 | Intron |
| chr4 | 154855958 | 154856427 | Distal Intergenic |
| chr4 | 154856583 | 154857018 | Distal Intergenic |
| chr4 | 154857069 | 154857290 | Distal Intergenic |
| chr4 | 154857377 | 154857577 | Distal Intergenic |
| chr4 | 154862389 | 154863029 | Distal Intergenic |
| chr4 | 155071249 | 155071545 | Distal Intergenic |
| chr4 | 155149114 | 155150294 | Distal Intergenic |
| chr4 | 155150637 | 155151666 | Distal Intergenic |
| chr4 | 155151718 | 155152028 | Distal Intergenic |
| chr4 | 155153147 | 155153383 | Downstream |
| chr4 | 155153720 | 155154552 | Downstream |
| chr4 | 155457347 | 155457566 | 3' UTR |
| chr4 | 155566540 | 155567406 | Intron |
| chr4 | 155567795 | 155568216 | Intron |
| chr5 | 3123319 | 3124113 | Distal Intergenic |
| chr5 | 3151969 | 3152695 | Distal Intergenic |
| chr5 | 3300838 | 3302321 | Distal Intergenic |
| chr5 | 3309292 | 3309727 | Distal Intergenic |
| chr5 | 3321361 | 3322855 | Distal Intergenic |
| chr5 | 3341549 | 3341748 | Distal Intergenic |
| chr5 | 3341788 | 3344138 | Distal Intergenic |
| chr5 | 3444368 | 3444699 | Intron |
| chr5 | 3444879 | 3445706 | Intron |
| chr5 | 3494234 | 3494810 | Intron |
| chr5 | 3501273 | 3502080 | Intron |
| chr5 | 3647037 | 3647269 | Distal Intergenic |
| chr5 | 3802978 | 3804345 | Distal Intergenic |
| chr5 | 4103151 | 4103935 | Distal Intergenic |
| chr5 | 4104030 | 4105373 | Distal Intergenic |
| chr5 | 4495865 | 4497263 | Distal Intergenic |
| chr5 | 4530424 | 4531168 | Distal Intergenic |
| chr5 | 5419550 | 5419753 | Distal Intergenic |
| chr5 | 5510714 | 5511344 | Distal Intergenic |
| chr5 | 5621295 | 5621539 | Distal Intergenic |
| chr5 | 5621580 | 5623476 | Distal Intergenic |
| chr5 | 5624336 | 5626284 | Distal Intergenic |
| chr5 | 5648656 | 5649645 | Distal Intergenic |

| | | | |
|---|---|---|---|
| chr5 | 5684273 | 5687687 | Distal Intergenic |
| chr5 | 5687992 | 5688844 | Distal Intergenic |
| chr5 | 5688903 | 5689301 | Distal Intergenic |
| chr5 | 5707658 | 5708561 | Distal Intergenic |
| chr5 | 8421706 | 8422722 | Intron |
| chr5 | 8797932 | 8798960 | Distal Intergenic |
| chr5 | 9068873 | 9069084 | Intron |
| chr5 | 9069175 | 9069768 | Intron |
| chr5 | 9141961 | 9142194 | Intron |
| chr5 | 9160844 | 9161651 | Intron |
| chr5 | 9161842 | 9162355 | Intron |
| chr5 | 10645825 | 10646054 | Intron |
| chr5 | 12107042 | 12107323 | Distal Intergenic |
| chr5 | 12107401 | 12107642 | Distal Intergenic |
| chr5 | 12242120 | 12242836 | Distal Intergenic |
| chr5 | 12243119 | 12243688 | Distal Intergenic |
| chr5 | 12404416 | 12404810 | Distal Intergenic |
| chr5 | 12910758 | 12911222 | Distal Intergenic |
| chr5 | 13415963 | 13416288 | Distal Intergenic |
| chr5 | 13745441 | 13745859 | Intron |
| chr5 | 13745903 | 13746209 | Intron |
| chr5 | 13974328 | 13975192 | Distal Intergenic |
| chr5 | 13983726 | 13984665 | Distal Intergenic |
| chr5 | 13987036 | 13987590 | Distal Intergenic |
| chr5 | 15838421 | 15838634 | Intron |
| chr5 | 16864697 | 16866333 | Intron |
| chr5 | 17010350 | 17012173 | Distal Intergenic |
| chr5 | 17050190 | 17051156 | Distal Intergenic |
| chr5 | 17061484 | 17063724 | Distal Intergenic |
| chr5 | 17077281 | 17077954 | Distal Intergenic |
| chr5 | 17079803 | 17080071 | Distal Intergenic |
| chr5 | 17080699 | 17081083 | Distal Intergenic |
| chr5 | 17143950 | 17145486 | Intron |
| chr5 | 17179753 | 17179953 | Intron |
| chr5 | 17194960 | 17195218 | Intron |
| chr5 | 17195367 | 17195841 | Intron |
| chr5 | 17505037 | 17506659 | Distal Intergenic |
| chr5 | 17509010 | 17509540 | Distal Intergenic |
| chr5 | 17509604 | 17510086 | Distal Intergenic |
| chr5 | 17541080 | 17542000 | Distal Intergenic |
| chr5 | 17542137 | 17542558 | Distal Intergenic |
| chr5 | 17641747 | 17644048 | Distal Intergenic |
| chr5 | 17663984 | 17664488 | Distal Intergenic |
| chr5 | 17664535 | 17665110 | Distal Intergenic |

| | | | |
|---|---|---|---|
| chr5 | 17665412 | 17665679 | Distal Intergenic |
| chr5 | 17803399 | 17803706 | Distal Intergenic |
| chr5 | 17880795 | 17881056 | Intron |
| chr5 | 17881390 | 17881589 | Intron |
| chr5 | 20215988 | 20217419 | Intron |
| chr5 | 20254326 | 20256095 | 5' UTR |
| chr5 | 20256170 | 20256379 | Intron |
| chr5 | 20405236 | 20405810 | Intron |
| chr5 | 20559219 | 20561751 | Intron |
| chr5 | 20562226 | 20562449 | Intron |
| chr5 | 20642012 | 20642284 | Intron |
| chr5 | 20692312 | 20692674 | Intron |
| chr5 | 20694110 | 20695680 | Intron |
| chr5 | 20723384 | 20723646 | Intron |
| chr5 | 20723853 | 20724237 | Intron |
| chr5 | 20946598 | 20947203 | Distal Intergenic |
| chr5 | 20992938 | 20993302 | Distal Intergenic |
| chr5 | 20993541 | 20993740 | Distal Intergenic |
| chr5 | 20997849 | 20999919 | Distal Intergenic |
| chr5 | 21004084 | 21004545 | Distal Intergenic |
| chr5 | 21004591 | 21004865 | Distal Intergenic |
| chr5 | 21044766 | 21045129 | Distal Intergenic |
| chr5 | 21059428 | 21060096 | Distal Intergenic |
| chr5 | 21065549 | 21066077 | Distal Intergenic |
| chr5 | 21290449 | 21290673 | Distal Intergenic |
| chr5 | 21291098 | 21291546 | Distal Intergenic |
| chr5 | 21292010 | 21293194 | Distal Intergenic |
| chr5 | 21617336 | 21617848 | Intron |
| chr5 | 21618390 | 21621523 | Intron |
| chr5 | 21624294 | 21624502 | Intron |
| chr5 | 21624555 | 21624935 | Intron |
| chr5 | 21624979 | 21625179 | Intron |
| chr5 | 21625791 | 21627293 | Intron |
| chr5 | 21647072 | 21648618 | Intron |
| chr5 | 21660569 | 21663419 | Intron |
| chr5 | 21663529 | 21664377 | Intron |
| chr5 | 21664435 | 21664634 | Intron |
| chr5 | 21664954 | 21665203 | Intron |
| chr5 | 21665512 | 21665813 | Intron |
| chr5 | 21729620 | 21731539 | Intron |
| chr5 | 21833828 | 21834881 | Intron |
| chr5 | 22048476 | 22048690 | Intron |
| chr5 | 22055605 | 22056193 | Intron |
| chr5 | 22892316 | 22892873 | Distal Intergenic |

| | | | |
|---|---|---|---|
| chr5 | 22892948 | 22893260 | Distal Intergenic |
| chr5 | 22923922 | 22924801 | Distal Intergenic |
| chr5 | 22929100 | 22929302 | Distal Intergenic |
| chr5 | 22937656 | 22938800 | Distal Intergenic |
| chr5 | 22938849 | 22939164 | Distal Intergenic |
| chr5 | 22939319 | 22941902 | Distal Intergenic |
| chr5 | 23011257 | 23011992 | Distal Intergenic |
| chr5 | 23087201 | 23087490 | Distal Intergenic |
| chr5 | 23121673 | 23122643 | Distal Intergenic |
| chr5 | 23157542 | 23157791 | Distal Intergenic |
| chr5 | 23157882 | 23158952 | Distal Intergenic |
| chr5 | 23217693 | 23218147 | Distal Intergenic |
| chr5 | 23349990 | 23350217 | Distal Intergenic |
| chr5 | 23357024 | 23357223 | Distal Intergenic |
| chr5 | 23383209 | 23383868 | Distal Intergenic |
| chr5 | 23383968 | 23385030 | Distal Intergenic |
| chr5 | 23393110 | 23393491 | Distal Intergenic |
| chr5 | 23399773 | 23400586 | Distal Intergenic |
| chr5 | 23404326 | 23405141 | Distal Intergenic |
| chr5 | 23408060 | 23409597 | Distal Intergenic |
| chr5 | 23419024 | 23419855 | Distal Intergenic |
| chr5 | 23419895 | 23420243 | Distal Intergenic |
| chr5 | 23498427 | 23500834 | Distal Intergenic |
| chr5 | 23535311 | 23535625 | Distal Intergenic |
| chr5 | 23535695 | 23536452 | Distal Intergenic |
| chr5 | 23542191 | 23543754 | Distal Intergenic |
| chr5 | 23543986 | 23544185 | Distal Intergenic |
| chr5 | 23544705 | 23545617 | Distal Intergenic |
| chr5 | 23545707 | 23546054 | Distal Intergenic |
| chr5 | 23552511 | 23554804 | Distal Intergenic |
| chr5 | 23569032 | 23569439 | Distal Intergenic |
| chr5 | 23569488 | 23571103 | Distal Intergenic |
| chr5 | 23617586 | 23619299 | Distal Intergenic |
| chr5 | 23729753 | 23730274 | Distal Intergenic |
| chr5 | 23767328 | 23768861 | Distal Intergenic |
| chr5 | 23772024 | 23772607 | Distal Intergenic |
| chr5 | 23775305 | 23775817 | Distal Intergenic |
| chr5 | 23895037 | 23895503 | Distal Intergenic |
| chr5 | 23895804 | 23896037 | Distal Intergenic |
| chr5 | 23896912 | 23899791 | Distal Intergenic |
| chr5 | 23910515 | 23911501 | Distal Intergenic |
| chr5 | 23929499 | 23930563 | Distal Intergenic |
| chr5 | 23950663 | 23951106 | Distal Intergenic |
| chr5 | 23971330 | 23971532 | Intron |

| | | | |
|---|---|---|---|
| chr5 | 23977822 | 23978396 | Exon |
| chr5 | 23978863 | 23982352 | Exon |
| chr5 | 24190862 | 24191555 | Distal Intergenic |
| chr5 | 24191608 | 24193023 | Distal Intergenic |
| chr5 | 24195056 | 24196296 | Distal Intergenic |
| chr5 | 24257008 | 24258233 | Distal Intergenic |
| chr5 | 24282432 | 24283671 | Distal Intergenic |
| chr5 | 24326361 | 24326595 | Distal Intergenic |
| chr5 | 24326658 | 24328307 | Distal Intergenic |
| chr5 | 24331605 | 24332018 | Distal Intergenic |
| chr5 | 24343035 | 24344439 | Distal Intergenic |
| chr5 | 24346042 | 24346377 | Distal Intergenic |
| chr5 | 24346997 | 24348396 | Distal Intergenic |
| chr5 | 25218889 | 25219160 | Distal Intergenic |
| chr5 | 25253915 | 25254124 | Distal Intergenic |
| chr5 | 25254243 | 25254488 | Distal Intergenic |
| chr5 | 25254904 | 25255294 | Distal Intergenic |
| chr5 | 25255350 | 25256428 | Distal Intergenic |
| chr5 | 25256482 | 25258213 | Distal Intergenic |
| chr5 | 26492318 | 26492627 | Distal Intergenic |
| chr5 | 26495876 | 26498252 | Distal Intergenic |
| chr5 | 28057211 | 28059404 | Distal Intergenic |
| chr5 | 28413299 | 28413577 | Distal Intergenic |
| chr5 | 28643224 | 28643662 | Distal Intergenic |
| chr5 | 28643958 | 28644921 | Distal Intergenic |
| chr5 | 28678294 | 28678763 | Distal Intergenic |
| chr5 | 28716725 | 28717005 | Distal Intergenic |
| chr5 | 28723847 | 28724121 | Distal Intergenic |
| chr5 | 28955879 | 28958509 | Distal Intergenic |
| chr5 | 28959416 | 28959706 | Distal Intergenic |
| chr5 | 29705016 | 29705285 | Distal Intergenic |
| chr5 | 29761281 | 29761580 | Distal Intergenic |
| chr5 | 30024652 | 30027316 | Distal Intergenic |
| chr5 | 30029097 | 30030639 | Distal Intergenic |
| chr5 | 30032807 | 30033618 | Distal Intergenic |
| chr5 | 30033698 | 30036791 | Distal Intergenic |
| chr5 | 30036855 | 30038006 | Distal Intergenic |
| chr5 | 30061810 | 30063894 | Distal Intergenic |
| chr5 | 30070469 | 30070675 | Distal Intergenic |
| chr5 | 30137926 | 30138130 | Distal Intergenic |
| chr5 | 30169273 | 30169893 | Distal Intergenic |
| chr5 | 30171364 | 30172372 | Distal Intergenic |
| chr5 | 30399542 | 30399876 | Distal Intergenic |
| chr5 | 30481001 | 30481788 | Distal Intergenic |

| | | | |
|---|---|---|---|
| chr5 | 30559339 | 30559978 | Distal Intergenic |
| chr5 | 30566683 | 30569635 | Distal Intergenic |
| chr5 | 30572819 | 30573029 | Distal Intergenic |
| chr5 | 30605233 | 30606092 | Distal Intergenic |
| chr5 | 30615433 | 30615824 | Distal Intergenic |
| chr5 | 30950828 | 30951285 | Distal Intergenic |
| chr5 | 30951458 | 30952100 | Distal Intergenic |
| chr5 | 30952147 | 30952346 | Distal Intergenic |
| chr5 | 30953161 | 30954896 | Distal Intergenic |
| chr5 | 30955055 | 30955928 | Distal Intergenic |
| chr5 | 30957541 | 30958913 | Distal Intergenic |
| chr5 | 30968966 | 30969511 | Distal Intergenic |
| chr5 | 31152011 | 31153803 | Distal Intergenic |
| chr5 | 31166116 | 31166708 | Distal Intergenic |
| chr5 | 31191732 | 31191933 | Distal Intergenic |
| chr5 | 31209592 | 31210117 | Intron |
| chr5 | 31357229 | 31357440 | Distal Intergenic |
| chr5 | 31456324 | 31456621 | Intron |
| chr5 | 31456989 | 31457216 | Intron |
| chr5 | 31481537 | 31483049 | Intron |
| chr5 | 31495953 | 31496201 | Intron |
| chr5 | 31498986 | 31499263 | Intron |
| chr5 | 31503620 | 31505180 | Exon |
| chr5 | 31521999 | 31522617 | Intron |
| chr5 | 31522765 | 31523001 | Intron |
| chr5 | 31523075 | 31523696 | Intron |
| chr5 | 31554255 | 31554978 | 3' UTR |
| chr5 | 31555104 | 31555332 | 3' UTR |
| chr5 | 31556200 | 31557614 | Distal Intergenic |
| chr5 | 31828928 | 31829371 | Intron |
| chr5 | 31866162 | 31867976 | Intron |
| chr5 | 31870437 | 31871010 | Intron |
| chr5 | 31878298 | 31879066 | Intron |
| chr5 | 31879222 | 31879787 | Intron |
| chr5 | 31898674 | 31902002 | Intron |
| chr5 | 31902245 | 31905438 | Intron |
| chr5 | 31907335 | 31909219 | Intron |
| chr5 | 31916029 | 31916941 | Intron |
| chr5 | 31917193 | 31918687 | Intron |
| chr5 | 31922330 | 31923262 | Intron |
| chr5 | 31927614 | 31928113 | Intron |
| chr5 | 31931134 | 31931555 | Intron |
| chr5 | 31931723 | 31932225 | Intron |
| chr5 | 31932328 | 31933966 | Intron |

| | | | |
|---|---|---|---|
| chr5 | 31935810 | 31940087 | Promoter |
| chr5 | 31995298 | 31995531 | Intron |
| chr5 | 31995628 | 31995917 | Exon |
| chr5 | 31995997 | 31996196 | Intron |
| chr5 | 32000032 | 32000252 | Exon |
| chr5 | 32000429 | 32001880 | Intron |
| chr5 | 32002104 | 32002333 | Intron |
| chr5 | 32002553 | 32002802 | Intron |
| chr5 | 32052811 | 32053276 | Exon |
| chr5 | 32053352 | 32054012 | Exon |
| chr5 | 32054057 | 32054324 | Intron |
| chr5 | 32057258 | 32058062 | Exon |
| chr5 | 32060461 | 32061728 | Exon |
| chr5 | 32184764 | 32185154 | Distal Intergenic |
| chr5 | 32207619 | 32207888 | Distal Intergenic |
| chr5 | 32238085 | 32238418 | Intron |
| chr5 | 32243834 | 32245562 | Intron |
| chr5 | 32285720 | 32287792 | Intron |
| chr5 | 32337569 | 32338277 | Distal Intergenic |
| chr5 | 32338345 | 32338605 | Distal Intergenic |
| chr5 | 32423034 | 32424913 | Intron |
| chr5 | 32436173 | 32437212 | Intron |
| chr5 | 32438241 | 32438819 | Intron |
| chr5 | 32439057 | 32441644 | Intron |
| chr5 | 32441706 | 32441905 | Intron |
| chr5 | 32448692 | 32448962 | Distal Intergenic |
| chr5 | 32479492 | 32480055 | Distal Intergenic |
| chr5 | 32494768 | 32495174 | Distal Intergenic |
| chr5 | 32499858 | 32500550 | Distal Intergenic |
| chr5 | 32501066 | 32502133 | Distal Intergenic |
| chr5 | 32511727 | 32512043 | Distal Intergenic |
| chr5 | 32616892 | 32617125 | Distal Intergenic |
| chr5 | 32617285 | 32618008 | Distal Intergenic |
| chr5 | 32637835 | 32641752 | Distal Intergenic |
| chr5 | 32645025 | 32645478 | Distal Intergenic |
| chr5 | 32716195 | 32716488 | Intron |
| chr5 | 32716857 | 32717106 | Intron |
| chr5 | 32727807 | 32728404 | Intron |
| chr5 | 32728536 | 32728735 | Intron |
| chr5 | 32760932 | 32761198 | Intron |
| chr5 | 32761495 | 32763324 | Intron |
| chr5 | 32765678 | 32765882 | Intron |
| chr5 | 32774549 | 32775659 | Exon |
| chr5 | 32802593 | 32803209 | Distal Intergenic |

| | | | |
|---|---|---|---|
| chr5 | 32912870 | 32913074 | Distal Intergenic |
| chr5 | 32913767 | 32913969 | Distal Intergenic |
| chr5 | 32914024 | 32914241 | Distal Intergenic |
| chr5 | 33196709 | 33197148 | Distal Intergenic |
| chr5 | 33318954 | 33320039 | Distal Intergenic |
| chr5 | 33345367 | 33346483 | Distal Intergenic |
| chr5 | 33362489 | 33362707 | Distal Intergenic |
| chr5 | 33558730 | 33558997 | Intron |
| chr5 | 33562358 | 33563298 | Intron |
| chr5 | 33616769 | 33617025 | Intron |
| chr5 | 33678649 | 33680274 | Intron |
| chr5 | 33684054 | 33684440 | Exon |
| chr5 | 33684498 | 33686966 | Intron |
| chr5 | 33693926 | 33694747 | Intron |
| chr5 | 33694791 | 33696054 | Intron |
| chr5 | 33721331 | 33721624 | Intron |
| chr5 | 33971451 | 33971862 | Intron |
| chr5 | 33999943 | 34000287 | Intron |
| chr5 | 34000504 | 34001077 | Intron |
| chr5 | 34125021 | 34125354 | Distal Intergenic |
| chr5 | 34161892 | 34162393 | Distal Intergenic |
| chr5 | 34261998 | 34262813 | Distal Intergenic |
| chr5 | 34278684 | 34278916 | Distal Intergenic |
| chr5 | 34279128 | 34279327 | Distal Intergenic |
| chr5 | 34298259 | 34298975 | Distal Intergenic |
| chr5 | 34352812 | 34353036 | Distal Intergenic |
| chr5 | 34353073 | 34353576 | Distal Intergenic |
| chr5 | 34419835 | 34420545 | Distal Intergenic |
| chr5 | 34423930 | 34426854 | Distal Intergenic |
| chr5 | 34430164 | 34431064 | Distal Intergenic |
| chr5 | 34511829 | 34512100 | Distal Intergenic |
| chr5 | 34678109 | 34680120 | Intron |
| chr5 | 34702532 | 34702858 | Intron |
| chr5 | 34702913 | 34703132 | Intron |
| chr5 | 34703234 | 34703433 | Intron |
| chr5 | 34772965 | 34773183 | Intron |
| chr5 | 34924787 | 34925074 | Exon |
| chr5 | 34951346 | 34954434 | Exon |
| chr5 | 34965950 | 34967016 | Distal Intergenic |
| chr5 | 34979057 | 34980148 | Distal Intergenic |
| chr5 | 35005972 | 35006256 | Intron |
| chr5 | 35331184 | 35333372 | Distal Intergenic |
| chr5 | 35340062 | 35343645 | Distal Intergenic |
| chr5 | 35343828 | 35344155 | Distal Intergenic |

| | | | |
|---|---|---|---|
| chr5 | 35346961 | 35347173 | Distal Intergenic |
| chr5 | 35363403 | 35363793 | Distal Intergenic |
| chr5 | 35375090 | 35377717 | Distal Intergenic |
| chr5 | 35377910 | 35381285 | Distal Intergenic |
| chr5 | 35917469 | 35918304 | Intron |
| chr5 | 35936473 | 35937131 | Intron |
| chr5 | 35957993 | 35958542 | Intron |
| chr5 | 36004555 | 36005329 | Distal Intergenic |
| chr5 | 36045140 | 36046788 | Intron |
| chr5 | 36059973 | 36060799 | Intron |
| chr5 | 36082443 | 36082751 | Distal Intergenic |
| chr5 | 36097966 | 36098275 | Distal Intergenic |
| chr5 | 36236415 | 36237009 | Intron |
| chr5 | 36250021 | 36252046 | Exon |
| chr5 | 36266108 | 36266737 | Intron |
| chr5 | 36266832 | 36267094 | Intron |
| chr5 | 36282698 | 36287264 | Intron |
| chr5 | 36289692 | 36291565 | Intron |
| chr5 | 36297133 | 36297892 | Intron |
| chr5 | 36315967 | 36316842 | Distal Intergenic |
| chr5 | 36545181 | 36546728 | Distal Intergenic |
| chr5 | 36709492 | 36709732 | Distal Intergenic |
| chr5 | 36709939 | 36710277 | Distal Intergenic |
| chr5 | 36722315 | 36723876 | Distal Intergenic |
| chr5 | 36724664 | 36726584 | Distal Intergenic |
| chr5 | 36732467 | 36733138 | Distal Intergenic |
| chr5 | 36757230 | 36757976 | Distal Intergenic |
| chr5 | 36764514 | 36766177 | Distal Intergenic |
| chr5 | 36803004 | 36804706 | Distal Intergenic |
| chr5 | 36961582 | 36961785 | Promoter |
| chr5 | 37065608 | 37067391 | 3' UTR |
| chr5 | 37067824 | 37068135 | Distal Intergenic |
| chr5 | 37103982 | 37104885 | Downstream |
| chr5 | 37105646 | 37106737 | 3' UTR |
| chr5 | 37113677 | 37114130 | Intron |
| chr5 | 37114252 | 37114529 | Intron |
| chr5 | 37187211 | 37187464 | Intron |
| chr5 | 37230166 | 37232711 | Exon |
| chr5 | 37236581 | 37238099 | Intron |
| chr5 | 37239487 | 37241943 | Exon |
| chr5 | 37243186 | 37244635 | Exon |
| chr5 | 37296290 | 37296750 | Intron |
| chr5 | 37311200 | 37313368 | Intron |
| chr5 | 38871295 | 38871704 | Intron |

| | | | |
|---|---|---|---|
| chr5 | 38944934 | 38945900 | Exon |
| chr5 | 38951752 | 38952106 | Intron |
| chr5 | 38953114 | 38953313 | Exon |
| chr5 | 39824353 | 39825245 | Distal Intergenic |
| chr5 | 40153179 | 40153775 | Distal Intergenic |
| chr5 | 42542073 | 42542680 | Intron |
| chr5 | 42547116 | 42547339 | Promoter |
| chr5 | 43371966 | 43373246 | Distal Intergenic |
| chr5 | 43378641 | 43379701 | Intron |
| chr5 | 43414297 | 43414519 | Distal Intergenic |
| chr5 | 43414562 | 43415410 | Distal Intergenic |
| chr5 | 43415461 | 43416007 | Distal Intergenic |
| chr5 | 43436824 | 43437793 | Distal Intergenic |
| chr5 | 43439695 | 43440245 | Distal Intergenic |
| chr5 | 43463243 | 43464766 | Intron |
| chr5 | 43472041 | 43473790 | Intron |
| chr5 | 43623668 | 43624011 | Intron |
| chr5 | 43624459 | 43624999 | Intron |
| chr5 | 43746496 | 43746961 | Distal Intergenic |
| chr5 | 43749477 | 43749705 | Distal Intergenic |
| chr5 | 43766770 | 43767015 | Distal Intergenic |
| chr5 | 44083338 | 44084995 | Distal Intergenic |
| chr5 | 44157502 | 44157783 | Distal Intergenic |
| chr5 | 44157903 | 44158505 | Distal Intergenic |
| chr5 | 44171540 | 44172769 | Distal Intergenic |
| chr5 | 44173032 | 44173301 | Distal Intergenic |
| chr5 | 44178817 | 44179717 | Distal Intergenic |
| chr5 | 44179773 | 44180071 | Distal Intergenic |
| chr5 | 44188505 | 44189303 | Distal Intergenic |
| chr5 | 44190558 | 44191572 | Distal Intergenic |
| chr5 | 44191701 | 44192002 | Distal Intergenic |
| chr5 | 44192602 | 44193634 | Distal Intergenic |
| chr5 | 44206580 | 44208148 | Distal Intergenic |
| chr5 | 44208422 | 44208731 | Distal Intergenic |
| chr5 | 44211567 | 44211852 | Distal Intergenic |
| chr5 | 44211972 | 44212504 | Distal Intergenic |
| chr5 | 44336625 | 44336951 | Intron |
| chr5 | 44338564 | 44341201 | Intron |
| chr5 | 44365291 | 44366820 | Intron |
| chr5 | 44376130 | 44377929 | Intron |
| chr5 | 44549262 | 44549476 | Distal Intergenic |
| chr5 | 44642863 | 44643137 | Distal Intergenic |
| chr5 | 44643176 | 44643390 | Distal Intergenic |
| chr5 | 45686643 | 45686978 | Intron |

| | | | |
|---|---|---|---|
| chr5 | 45868863 | 45869619 | Distal Intergenic |
| chr5 | 45911183 | 45911444 | Distal Intergenic |
| chr5 | 46248509 | 46248708 | Distal Intergenic |
| chr5 | 47787301 | 47789258 | Distal Intergenic |
| chr5 | 47893966 | 47894409 | Distal Intergenic |
| chr5 | 48080197 | 48080455 | Distal Intergenic |
| chr5 | 48504968 | 48505218 | Distal Intergenic |
| chr5 | 48505259 | 48505549 | Distal Intergenic |
| chr5 | 48955113 | 48955455 | Distal Intergenic |
| chr5 | 49728042 | 49728260 | Intron |
| chr5 | 49728555 | 49728763 | Intron |
| chr5 | 51432664 | 51433513 | Distal Intergenic |
| chr5 | 51437224 | 51437476 | Distal Intergenic |
| chr5 | 52529202 | 52530251 | Distal Intergenic |
| chr5 | 52530310 | 52530637 | Distal Intergenic |
| chr5 | 52580579 | 52581287 | Distal Intergenic |
| chr5 | 52582327 | 52582767 | Distal Intergenic |
| chr5 | 52623099 | 52623332 | Distal Intergenic |
| chr5 | 52623373 | 52623572 | Distal Intergenic |
| chr5 | 53052520 | 53053302 | Distal Intergenic |
| chr5 | 53174399 | 53174605 | Exon |
| chr5 | 53177729 | 53178001 | Downstream |
| chr5 | 53178075 | 53178383 | Downstream |
| chr5 | 53329559 | 53330071 | Intron |
| chr5 | 53331495 | 53331760 | Intron |
| chr5 | 53432959 | 53434561 | Intron |
| chr5 | 53493213 | 53494354 | Intron |
| chr5 | 53494706 | 53494924 | Intron |
| chr5 | 53494977 | 53495945 | Intron |
| chr5 | 53528610 | 53530625 | Intron |
| chr5 | 53562687 | 53563613 | Intron |
| chr5 | 53587792 | 53588127 | Intron |
| chr5 | 53625970 | 53627071 | Distal Intergenic |
| chr5 | 53627351 | 53627762 | Distal Intergenic |
| chr5 | 53629779 | 53631867 | Distal Intergenic |
| chr5 | 53658140 | 53658799 | Distal Intergenic |
| chr5 | 53658915 | 53659181 | Distal Intergenic |
| chr5 | 53659382 | 53659693 | Distal Intergenic |
| chr5 | 54312253 | 54312663 | Distal Intergenic |
| chr5 | 54340079 | 54340602 | Distal Intergenic |
| chr5 | 54344294 | 54345825 | Distal Intergenic |
| chr5 | 54353055 | 54353355 | Distal Intergenic |
| chr5 | 54362178 | 54363170 | Distal Intergenic |
| chr5 | 58107925 | 58108140 | Intron |

| | | | |
|---|---|---|---|
| chr5 | 58109565 | 58109909 | Intron |
| chr5 | 58113722 | 58114033 | Intron |
| chr5 | 58146893 | 58147457 | 3' UTR |
| chr5 | 58206911 | 58208234 | Distal Intergenic |
| chr5 | 58310170 | 58310399 | Intron |
| chr5 | 58310486 | 58310753 | Intron |
| chr5 | 58311064 | 58311263 | Intron |
| chr5 | 58331730 | 58332139 | Intron |
| chr5 | 58333808 | 58335747 | Promoter |
| chr5 | 58365214 | 58365964 | Intron |
| chr5 | 58366932 | 58368681 | Intron |
| chr5 | 58430741 | 58432008 | Intron |
| chr5 | 58497558 | 58497922 | Intron |
| chr5 | 58498256 | 58498710 | Intron |
| chr5 | 58991032 | 58991335 | Intron |
| chr5 | 63156434 | 63156727 | Distal Intergenic |
| chr5 | 63156807 | 63157060 | Distal Intergenic |
| chr5 | 63157600 | 63158078 | Distal Intergenic |
| chr5 | 63193666 | 63194582 | Distal Intergenic |
| chr5 | 63203926 | 63204127 | Distal Intergenic |
| chr5 | 64345133 | 64345354 | Distal Intergenic |
| chr5 | 64349086 | 64349715 | Distal Intergenic |
| chr5 | 64355753 | 64356625 | Distal Intergenic |
| chr5 | 64358601 | 64359511 | Distal Intergenic |
| chr5 | 64397832 | 64398613 | Distal Intergenic |
| chr5 | 64436094 | 64436905 | Distal Intergenic |
| chr5 | 64436985 | 64437243 | Distal Intergenic |
| chr5 | 64439090 | 64439289 | Distal Intergenic |
| chr5 | 64503245 | 64503660 | Intron |
| chr5 | 64504013 | 64504215 | Intron |
| chr5 | 64537610 | 64538409 | Exon |
| chr5 | 64552093 | 64552416 | Intron |
| chr5 | 64565393 | 64566309 | Intron |
| chr5 | 64625779 | 64627157 | Intron |
| chr5 | 64630419 | 64631530 | Intron |
| chr5 | 64897157 | 64897377 | Intron |
| chr5 | 64912743 | 64913077 | Intron |
| chr5 | 64970673 | 64971666 | Intron |
| chr5 | 65094956 | 65095202 | Intron |
| chr5 | 65095549 | 65096147 | Intron |
| chr5 | 65195480 | 65198563 | Distal Intergenic |
| chr5 | 65200773 | 65200977 | Distal Intergenic |
| chr5 | 65201660 | 65204502 | Distal Intergenic |
| chr5 | 65384030 | 65384256 | Distal Intergenic |

| | | | |
|---|---|---|---|
| chr5 | 65781054 | 65781260 | Distal Intergenic |
| chr5 | 65781321 | 65781776 | Distal Intergenic |
| chr5 | 65782112 | 65782672 | Distal Intergenic |
| chr5 | 65819838 | 65820341 | Distal Intergenic |
| chr5 | 65845568 | 65845767 | Distal Intergenic |
| chr5 | 65845872 | 65846727 | Distal Intergenic |
| chr5 | 65848119 | 65850082 | Distal Intergenic |
| chr5 | 65863613 | 65863857 | Distal Intergenic |
| chr5 | 65927646 | 65928182 | Intron |
| chr5 | 65928559 | 65929818 | Intron |
| chr5 | 66037836 | 66038193 | Intron |
| chr5 | 66088645 | 66088965 | Intron |
| chr5 | 66154271 | 66154673 | Intron |
| chr5 | 66156010 | 66156236 | Intron |
| chr5 | 66425590 | 66425843 | Intron |
| chr5 | 66425981 | 66426340 | Exon |
| chr5 | 66436988 | 66437920 | Intron |
| chr5 | 66541090 | 66543488 | Distal Intergenic |
| chr5 | 66556871 | 66557304 | Distal Intergenic |
| chr5 | 66666258 | 66666457 | Distal Intergenic |
| chr5 | 66888529 | 66888992 | Distal Intergenic |
| chr5 | 66889193 | 66889423 | Distal Intergenic |
| chr5 | 66909675 | 66909885 | Distal Intergenic |
| chr5 | 66933567 | 66935871 | Distal Intergenic |
| chr5 | 66941460 | 66942575 | Distal Intergenic |
| chr5 | 66948905 | 66952012 | Distal Intergenic |
| chr5 | 66970287 | 66970894 | Distal Intergenic |
| chr5 | 66972754 | 66972953 | Distal Intergenic |
| chr5 | 66973106 | 66973359 | Distal Intergenic |
| chr5 | 66977039 | 66978032 | Distal Intergenic |
| chr5 | 66979402 | 66981566 | Distal Intergenic |
| chr5 | 66998229 | 67000317 | Distal Intergenic |
| chr5 | 67652015 | 67652542 | Distal Intergenic |
| chr5 | 67799195 | 67799575 | Distal Intergenic |
| chr5 | 67799618 | 67800394 | Distal Intergenic |
| chr5 | 67801733 | 67802171 | Distal Intergenic |
| chr5 | 68222543 | 68222990 | Distal Intergenic |
| chr5 | 68235337 | 68236034 | Distal Intergenic |
| chr5 | 68238804 | 68239037 | Distal Intergenic |
| chr5 | 68257479 | 68258080 | Distal Intergenic |
| chr5 | 68258123 | 68258742 | Distal Intergenic |
| chr5 | 72558788 | 72559160 | Distal Intergenic |
| chr5 | 72591498 | 72594471 | Distal Intergenic |
| chr5 | 72594554 | 72595256 | Distal Intergenic |

| | | | |
|---|---|---|---|
| chr5 | 72595530 | 72595812 | Distal Intergenic |
| chr5 | 72602467 | 72604029 | Distal Intergenic |
| chr5 | 72640771 | 72642094 | Distal Intergenic |
| chr5 | 72645928 | 72646342 | Distal Intergenic |
| chr5 | 73246076 | 73247362 | Distal Intergenic |
| chr5 | 73471271 | 73471484 | Exon |
| chr5 | 73495030 | 73495233 | Distal Intergenic |
| chr5 | 73624362 | 73625014 | Distal Intergenic |
| chr5 | 73625204 | 73625469 | Distal Intergenic |
| chr5 | 73733392 | 73736174 | Distal Intergenic |
| chr5 | 73760129 | 73760331 | Distal Intergenic |
| chr5 | 73764002 | 73764461 | Distal Intergenic |
| chr5 | 73894022 | 73894332 | Distal Intergenic |
| chr5 | 73955022 | 73955639 | Intron |
| chr5 | 73956792 | 73958828 | Intron |
| chr5 | 73969496 | 73970866 | Intron |
| chr5 | 74488782 | 74488984 | Intron |
| chr5 | 74489127 | 74490646 | Exon |
| chr5 | 74490927 | 74491488 | Exon |
| chr5 | 74493136 | 74493565 | Intron |
| chr5 | 74493611 | 74495132 | Intron |
| chr5 | 74497673 | 74498226 | Intron |
| chr5 | 74501561 | 74501844 | Intron |
| chr5 | 74568198 | 74568677 | Distal Intergenic |
| chr5 | 74681113 | 74681478 | Intron |
| chr5 | 74681601 | 74681800 | Exon |
| chr5 | 74888496 | 74888991 | Intron |
| chr5 | 74931583 | 74932298 | Exon |
| chr5 | 75148797 | 75149790 | Distal Intergenic |
| chr5 | 75292750 | 75294262 | Distal Intergenic |
| chr5 | 75331219 | 75331423 | Distal Intergenic |
| chr5 | 75331516 | 75332989 | Distal Intergenic |
| chr5 | 75334254 | 75334515 | Distal Intergenic |
| chr5 | 75429259 | 75429556 | Intron |
| chr5 | 75439425 | 75439757 | Intron |
| chr5 | 75812879 | 75813131 | Intron |
| chr5 | 75817526 | 75817860 | Intron |
| chr5 | 76613480 | 76614071 | Intron |
| chr5 | 76733901 | 76734532 | Exon |
| chr5 | 76774501 | 76775045 | Intron |
| chr5 | 76775091 | 76776939 | Intron |
| chr5 | 77334227 | 77334430 | Intron |
| chr5 | 77335458 | 77335807 | Intron |
| chr5 | 77391164 | 77391389 | Intron |

| | | | |
|---|---|---|---|
| chr5 | 77391492 | 77391769 | Intron |
| chr5 | 77403178 | 77403688 | Intron |
| chr5 | 77403750 | 77404027 | Intron |
| chr5 | 77404391 | 77404618 | Intron |
| chr5 | 77501519 | 77501785 | Intron |
| chr5 | 77583769 | 77583984 | Intron |
| chr5 | 77584174 | 77584704 | Intron |
| chr5 | 77629401 | 77629613 | Distal Intergenic |
| chr5 | 77629758 | 77630342 | Distal Intergenic |
| chr5 | 77677174 | 77677571 | Intron |
| chr5 | 77681454 | 77681685 | Intron |
| chr5 | 77695549 | 77696479 | Intron |
| chr5 | 77793581 | 77793825 | Intron |
| chr5 | 77794043 | 77794347 | Intron |
| chr5 | 77794669 | 77797751 | Intron |
| chr5 | 86493659 | 86493949 | Intron |
| chr5 | 86494147 | 86494615 | Intron |
| chr5 | 86669931 | 86670255 | Exon |
| chr5 | 86674032 | 86674281 | Exon |
| chr5 | 86676478 | 86676772 | Intron |
| chr5 | 86677190 | 86677841 | Intron |
| chr5 | 86969811 | 86970301 | Distal Intergenic |
| chr5 | 87059712 | 87060110 | Distal Intergenic |
| chr5 | 87105347 | 87105841 | Distal Intergenic |
| chr5 | 87124114 | 87124647 | Distal Intergenic |
| chr5 | 87151661 | 87152759 | Distal Intergenic |
| chr5 | 87155742 | 87157921 | Distal Intergenic |
| chr5 | 87158572 | 87160682 | Distal Intergenic |
| chr5 | 87170191 | 87170631 | Distal Intergenic |
| chr5 | 87171118 | 87171470 | Distal Intergenic |
| chr5 | 87171698 | 87172488 | Distal Intergenic |
| chr5 | 87233562 | 87234049 | Distal Intergenic |
| chr5 | 87234212 | 87234464 | Distal Intergenic |
| chr5 | 87285437 | 87286162 | Distal Intergenic |
| chr5 | 88982821 | 88983179 | Distal Intergenic |
| chr5 | 89116952 | 89118258 | Distal Intergenic |
| chr5 | 89149644 | 89150062 | Distal Intergenic |
| chr5 | 89412231 | 89412686 | Distal Intergenic |
| chr5 | 89899084 | 89899986 | Intron |
| chr5 | 89902088 | 89902296 | Intron |
| chr5 | 89902338 | 89902576 | Intron |
| chr5 | 90177453 | 90178054 | Intron |
| chr5 | 90653118 | 90653530 | Distal Intergenic |
| chr5 | 90653651 | 90653947 | Distal Intergenic |

| | | | |
|---|---|---|---|
| chr5 | 90793710 | 90794751 | Distal Intergenic |
| chr5 | 90794837 | 90796311 | Distal Intergenic |
| chr5 | 91023954 | 91024468 | Distal Intergenic |
| chr5 | 91040219 | 91042117 | Distal Intergenic |
| chr5 | 91063733 | 91063932 | Distal Intergenic |
| chr5 | 91088681 | 91090563 | Distal Intergenic |
| chr5 | 91116709 | 91119561 | Distal Intergenic |
| chr5 | 91120165 | 91120364 | Distal Intergenic |
| chr5 | 91120428 | 91120769 | Distal Intergenic |
| chr5 | 91120990 | 91121195 | Distal Intergenic |
| chr5 | 91122680 | 91124225 | Distal Intergenic |
| chr5 | 91127693 | 91129417 | Distal Intergenic |
| chr5 | 91134992 | 91135357 | Distal Intergenic |
| chr5 | 91135399 | 91135780 | Distal Intergenic |
| chr5 | 91187828 | 91188122 | Distal Intergenic |
| chr5 | 91188267 | 91188479 | Distal Intergenic |
| chr5 | 91188601 | 91189327 | Distal Intergenic |
| chr5 | 91189515 | 91191808 | Distal Intergenic |
| chr5 | 91191967 | 91193844 | Distal Intergenic |
| chr5 | 91206396 | 91206783 | Distal Intergenic |
| chr5 | 91206837 | 91207829 | Distal Intergenic |
| chr5 | 91208084 | 91208524 | Distal Intergenic |
| chr5 | 91304442 | 91306227 | Distal Intergenic |
| chr5 | 91308253 | 91310056 | Distal Intergenic |
| chr5 | 91310687 | 91311163 | Distal Intergenic |
| chr5 | 91320241 | 91320445 | Distal Intergenic |
| chr5 | 91320540 | 91321126 | Distal Intergenic |
| chr5 | 91328310 | 91330271 | Distal Intergenic |
| chr5 | 91360425 | 91361201 | Distal Intergenic |
| chr5 | 91400937 | 91401977 | Distal Intergenic |
| chr5 | 91406610 | 91407478 | Distal Intergenic |
| chr5 | 91416339 | 91417024 | Distal Intergenic |
| chr5 | 91444002 | 91444260 | Distal Intergenic |
| chr5 | 91448029 | 91448420 | Distal Intergenic |
| chr5 | 91456312 | 91456631 | Distal Intergenic |
| chr5 | 91457077 | 91457276 | Distal Intergenic |
| chr5 | 91463929 | 91467395 | Distal Intergenic |
| chr5 | 91502139 | 91502503 | Intron |
| chr5 | 91502618 | 91504095 | Intron |
| chr5 | 91505144 | 91505367 | Intron |
| chr5 | 91510280 | 91510625 | Intron |
| chr5 | 91530652 | 91531012 | Intron |
| chr5 | 91531074 | 91531480 | Intron |
| chr5 | 91533152 | 91533352 | Intron |

| | | | |
|---|---|---|---|
| chr5 | 91533440 | 91533929 | Intron |
| chr5 | 91534037 | 91534651 | Intron |
| chr5 | 91534855 | 91535183 | Intron |
| chr5 | 91538361 | 91540951 | Intron |
| chr5 | 91543336 | 91543642 | Intron |
| chr5 | 91670607 | 91671490 | Intron |
| chr5 | 91726077 | 91726305 | Intron |
| chr5 | 92038074 | 92038306 | Distal Intergenic |
| chr5 | 92038379 | 92039698 | Distal Intergenic |
| chr5 | 92041771 | 92042543 | Distal Intergenic |
| chr5 | 92046262 | 92046919 | Distal Intergenic |
| chr5 | 92100299 | 92100554 | Distal Intergenic |
| chr5 | 92137270 | 92138004 | Distal Intergenic |
| chr5 | 92269712 | 92270579 | Distal Intergenic |
| chr5 | 92389801 | 92392081 | Distal Intergenic |
| chr5 | 92415766 | 92417023 | Distal Intergenic |
| chr5 | 92472245 | 92472444 | Distal Intergenic |
| chr5 | 92647704 | 92650899 | Distal Intergenic |
| chr5 | 92658438 | 92659971 | Distal Intergenic |
| chr5 | 92711913 | 92712892 | Distal Intergenic |
| chr5 | 92897709 | 92899662 | Promoter |
| chr5 | 92899746 | 92899994 | Promoter |
| chr5 | 92900102 | 92901541 | Promoter |
| chr5 | 92918263 | 92919638 | Promoter |
| chr5 | 92933839 | 92934158 | Distal Intergenic |
| chr5 | 93253579 | 93253938 | Intron |
| chr5 | 93278776 | 93279169 | Intron |
| chr5 | 93390587 | 93391445 | Intron |
| chr5 | 93395888 | 93397190 | Intron |
| chr5 | 93398283 | 93398600 | Intron |
| chr5 | 93398825 | 93399025 | Intron |
| chr5 | 93399173 | 93399720 | Intron |
| chr5 | 93522236 | 93522602 | Intron |
| chr5 | 93532421 | 93534230 | Intron |
| chr5 | 93534555 | 93534851 | Intron |
| chr5 | 93555100 | 93555390 | Intron |
| chr5 | 93556994 | 93557292 | Intron |
| chr5 | 93558794 | 93559200 | Intron |
| chr5 | 93564303 | 93564502 | Intron |
| chr5 | 96241488 | 96242964 | Intron |
| chr5 | 96589456 | 96590109 | Distal Intergenic |
| chr5 | 96590733 | 96591015 | Distal Intergenic |
| chr5 | 96591117 | 96591516 | Distal Intergenic |
| chr5 | 96650296 | 96650620 | Distal Intergenic |

| | | | |
|---|---|---|---|
| chr5 | 96650686 | 96651073 | Distal Intergenic |
| chr5 | 96651128 | 96651355 | Distal Intergenic |
| chr5 | 96700253 | 96700480 | Distal Intergenic |
| chr5 | 97007056 | 97008246 | Distal Intergenic |
| chr5 | 97048681 | 97050593 | Distal Intergenic |
| chr5 | 97051362 | 97051885 | Distal Intergenic |
| chr5 | 97109757 | 97110050 | Distal Intergenic |
| chr5 | 97222287 | 97224491 | Distal Intergenic |
| chr5 | 97229547 | 97230004 | Distal Intergenic |
| chr5 | 97230141 | 97230757 | Distal Intergenic |
| chr5 | 97230935 | 97231977 | Distal Intergenic |
| chr5 | 97232056 | 97233130 | Distal Intergenic |
| chr5 | 97233305 | 97234609 | Distal Intergenic |
| chr5 | 97326651 | 97328868 | Distal Intergenic |
| chr5 | 97328969 | 97329347 | Distal Intergenic |
| chr5 | 97393484 | 97393741 | Distal Intergenic |
| chr5 | 98712866 | 98713147 | Distal Intergenic |
| chr5 | 98713322 | 98713902 | Distal Intergenic |
| chr5 | 98719529 | 98720735 | Distal Intergenic |
| chr5 | 98721731 | 98723071 | Distal Intergenic |
| chr5 | 98855739 | 98855964 | Distal Intergenic |
| chr5 | 98858446 | 98859830 | Exon |
| chr5 | 99273413 | 99274090 | Distal Intergenic |
| chr5 | 99311908 | 99313443 | Distal Intergenic |
| chr5 | 99319253 | 99320014 | Distal Intergenic |
| chr5 | 99326954 | 99327647 | Distal Intergenic |
| chr5 | 99372640 | 99372915 | Distal Intergenic |
| chr5 | 99372983 | 99373483 | Distal Intergenic |
| chr5 | 99500988 | 99502549 | Distal Intergenic |
| chr5 | 99502598 | 99502801 | Distal Intergenic |
| chr5 | 99502871 | 99503154 | Distal Intergenic |
| chr5 | 99533787 | 99534916 | Distal Intergenic |
| chr5 | 100396481 | 100397985 | Distal Intergenic |
| chr5 | 100398181 | 100398834 | Distal Intergenic |
| chr5 | 100398875 | 100399323 | Distal Intergenic |
| chr5 | 100404826 | 100405097 | Distal Intergenic |
| chr5 | 100405963 | 100407610 | Distal Intergenic |
| chr5 | 100488377 | 100488699 | Distal Intergenic |
| chr5 | 100844469 | 100844830 | Distal Intergenic |
| chr5 | 100845518 | 100845882 | Distal Intergenic |
| chr5 | 100845977 | 100846183 | Distal Intergenic |
| chr5 | 100857910 | 100858539 | Distal Intergenic |
| chr5 | 100858785 | 100859091 | Distal Intergenic |
| chr5 | 100928875 | 100929074 | Distal Intergenic |

| | | | |
|---|---|---|---|
| chr5 | 100947427 | 100947784 | Distal Intergenic |
| chr5 | 101051059 | 101052372 | Distal Intergenic |
| chr5 | 101061858 | 101062125 | Distal Intergenic |
| chr5 | 101529856 | 101531691 | Distal Intergenic |
| chr5 | 101591704 | 101593151 | Exon |
| chr5 | 101645879 | 101646370 | Distal Intergenic |
| chr5 | 102171068 | 102171747 | Distal Intergenic |
| chr5 | 102276611 | 102277258 | Intron |
| chr5 | 102835666 | 102835932 | Distal Intergenic |
| chr5 | 103793892 | 103795768 | Distal Intergenic |
| chr5 | 103795890 | 103797089 | Distal Intergenic |
| chr5 | 103805397 | 103806097 | Distal Intergenic |
| chr5 | 103806156 | 103806401 | Distal Intergenic |
| chr5 | 103824689 | 103824925 | Distal Intergenic |
| chr5 | 103848658 | 103850218 | Distal Intergenic |
| chr5 | 103855009 | 103855461 | Distal Intergenic |
| chr5 | 103879292 | 103880565 | Distal Intergenic |
| chr5 | 103887897 | 103888489 | Distal Intergenic |
| chr5 | 103888559 | 103888762 | Distal Intergenic |
| chr5 | 103947355 | 103948549 | Distal Intergenic |
| chr5 | 104024662 | 104026298 | Distal Intergenic |
| chr5 | 104120782 | 104121153 | Distal Intergenic |
| chr5 | 104161238 | 104161975 | Distal Intergenic |
| chr5 | 104165464 | 104167635 | Distal Intergenic |
| chr5 | 104181632 | 104182484 | Distal Intergenic |
| chr5 | 104182620 | 104182871 | Distal Intergenic |
| chr5 | 104183149 | 104183995 | Distal Intergenic |
| chr5 | 104187063 | 104187818 | Distal Intergenic |
| chr5 | 104201824 | 104202067 | Distal Intergenic |
| chr5 | 104202191 | 104202682 | Distal Intergenic |
| chr5 | 104203261 | 104204930 | Distal Intergenic |
| chr5 | 104207024 | 104208085 | Distal Intergenic |
| chr5 | 104208626 | 104209184 | Distal Intergenic |
| chr5 | 104248991 | 104249277 | Distal Intergenic |
| chr5 | 104250711 | 104251060 | Distal Intergenic |
| chr5 | 104251206 | 104253704 | Distal Intergenic |
| chr5 | 104257942 | 104258143 | Distal Intergenic |
| chr5 | 104258214 | 104258671 | Distal Intergenic |
| chr5 | 104475339 | 104475691 | Distal Intergenic |
| chr5 | 104836300 | 104837412 | Distal Intergenic |
| chr5 | 105583515 | 105583947 | Distal Intergenic |
| chr5 | 105949118 | 105949411 | Distal Intergenic |
| chr5 | 105949519 | 105949813 | Distal Intergenic |
| chr5 | 105950064 | 105950383 | Distal Intergenic |

| | | | |
|---|---|---|---|
| chr5 | 105950894 | 105951103 | Distal Intergenic |
| chr5 | 105951285 | 105953244 | Distal Intergenic |
| chr5 | 105993569 | 105994469 | Distal Intergenic |
| chr5 | 106065663 | 106066138 | Distal Intergenic |
| chr5 | 106066174 | 106066586 | Distal Intergenic |
| chr5 | 106206092 | 106206449 | Distal Intergenic |
| chr5 | 106207306 | 106209120 | Distal Intergenic |
| chr5 | 106221568 | 106222098 | Distal Intergenic |
| chr5 | 106222249 | 106222469 | Distal Intergenic |
| chr5 | 106227517 | 106228134 | Distal Intergenic |
| chr5 | 106229501 | 106229775 | Distal Intergenic |
| chr5 | 106235018 | 106235258 | Distal Intergenic |
| chr5 | 106250907 | 106252612 | Distal Intergenic |
| chr5 | 106304441 | 106304667 | Distal Intergenic |
| chr5 | 106304821 | 106305183 | Distal Intergenic |
| chr5 | 106306126 | 106306355 | Distal Intergenic |
| chr5 | 106353209 | 106353432 | Distal Intergenic |
| chr5 | 106353495 | 106353705 | Distal Intergenic |
| chr5 | 107430829 | 107431169 | Intron |
| chr5 | 107553758 | 107553997 | Intron |
| chr5 | 107697383 | 107697837 | Intron |
| chr5 | 107697886 | 107698243 | Intron |
| chr5 | 107920705 | 107922031 | Distal Intergenic |
| chr5 | 108222617 | 108222903 | Intron |
| chr5 | 108231898 | 108232100 | Intron |
| chr5 | 108274418 | 108274635 | Intron |
| chr5 | 108276201 | 108277153 | Intron |
| chr5 | 108330721 | 108330960 | Intron |
| chr5 | 108366834 | 108367915 | Intron |
| chr5 | 108400807 | 108401543 | Intron |
| chr5 | 108401644 | 108401914 | Intron |
| chr5 | 108416938 | 108417144 | Intron |
| chr5 | 108424737 | 108425924 | Intron |
| chr5 | 108494749 | 108495121 | Intron |
| chr5 | 108528978 | 108529338 | Exon |
| chr5 | 108560776 | 108561619 | Distal Intergenic |
| chr5 | 108697501 | 108697863 | Intron |
| chr5 | 108741231 | 108741531 | Intron |
| chr5 | 108890678 | 108891006 | Distal Intergenic |
| chr5 | 108891194 | 108891601 | Distal Intergenic |
| chr5 | 109057797 | 109058614 | Intron |
| chr5 | 109098462 | 109098692 | Intron |
| chr5 | 109143079 | 109143278 | Intron |
| chr5 | 110528811 | 110529132 | Distal Intergenic |

| | | | |
|---|---|---|---|
| chr5 | 110755268 | 110755893 | Intron |
| chr5 | 110856956 | 110858198 | Intron |
| chr5 | 111088590 | 111088886 | Intron |
| chr5 | 111089087 | 111089292 | Intron |
| chr5 | 111133155 | 111133363 | Intron |
| chr5 | 111229845 | 111232321 | Intron |
| chr5 | 111232882 | 111233385 | Intron |
| chr5 | 111233423 | 111234480 | Intron |
| chr5 | 111237266 | 111237480 | Intron |
| chr5 | 111237754 | 111237957 | Intron |
| chr5 | 111238008 | 111238260 | Intron |
| chr5 | 111739188 | 111739688 | Intron |
| chr5 | 111759936 | 111760265 | Distal Intergenic |
| chr5 | 111970689 | 111971005 | Distal Intergenic |
| chr5 | 112023052 | 112023452 | Distal Intergenic |
| chr5 | 112219284 | 112220145 | Intron |
| chr5 | 112309411 | 112309913 | Distal Intergenic |
| chr5 | 112310082 | 112310737 | Distal Intergenic |
| chr5 | 112707446 | 112707780 | Intron |
| chr5 | 112707939 | 112710992 | Intron |
| chr5 | 112724098 | 112724542 | Intron |
| chr5 | 112726381 | 112726825 | Intron |
| chr5 | 112752744 | 112753082 | Intron |
| chr5 | 112753193 | 112755191 | Intron |
| chr5 | 113534123 | 113535145 | Distal Intergenic |
| chr5 | 113535581 | 113536402 | Distal Intergenic |
| chr5 | 113581818 | 113583613 | Distal Intergenic |
| chr5 | 113583708 | 113584127 | Distal Intergenic |
| chr5 | 113644874 | 113645929 | Distal Intergenic |
| chr5 | 113808784 | 113808999 | Exon |
| chr5 | 113990247 | 113991606 | Intron |
| chr5 | 113994369 | 113994568 | Intron |
| chr5 | 114135335 | 114136051 | Distal Intergenic |
| chr5 | 114221707 | 114222199 | Distal Intergenic |
| chr5 | 114291031 | 114291260 | Distal Intergenic |
| chr5 | 114292082 | 114295339 | Distal Intergenic |
| chr5 | 114323381 | 114323668 | Distal Intergenic |
| chr5 | 114323896 | 114324162 | Distal Intergenic |
| chr5 | 114324610 | 114324944 | Distal Intergenic |
| chr5 | 114328587 | 114330116 | Distal Intergenic |
| chr5 | 114357136 | 114357914 | Distal Intergenic |
| chr5 | 114380003 | 114380287 | Distal Intergenic |
| chr5 | 114425496 | 114425695 | Distal Intergenic |
| chr5 | 114821510 | 114821950 | Distal Intergenic |

| | | | |
|---|---|---|---|
| chr5 | 114822086 | 114823889 | Distal Intergenic |
| chr5 | 114828107 | 114828344 | Distal Intergenic |
| chr5 | 114829255 | 114829958 | Distal Intergenic |
| chr5 | 114830616 | 114831215 | Distal Intergenic |
| chr5 | 114843129 | 114843341 | Distal Intergenic |
| chr5 | 114847793 | 114848614 | Distal Intergenic |
| chr5 | 114856232 | 114858524 | 3' UTR |
| chr5 | 114858573 | 114859211 | 3' UTR |
| chr5 | 115097295 | 115098280 | Distal Intergenic |
| chr5 | 115098322 | 115099663 | Distal Intergenic |
| chr5 | 115100982 | 115101339 | Distal Intergenic |
| chr5 | 115101678 | 115101945 | Distal Intergenic |
| chr5 | 115183677 | 115184162 | Intron |
| chr5 | 115201054 | 115201962 | Intron |
| chr5 | 115222089 | 115222374 | Intron |
| chr5 | 115297081 | 115297683 | Promoter |
| chr5 | 115447505 | 115447907 | Intron |
| chr5 | 115449866 | 115453143 | Intron |
| chr5 | 115457772 | 115458010 | Intron |
| chr5 | 115458127 | 115458656 | Intron |
| chr5 | 115512402 | 115513547 | Intron |
| chr5 | 115518699 | 115520023 | Intron |
| chr5 | 115521121 | 115522683 | Intron |
| chr5 | 115602374 | 115602604 | Intron |
| chr5 | 115762554 | 115764414 | Distal Intergenic |
| chr5 | 115941177 | 115941493 | Distal Intergenic |
| chr5 | 115941549 | 115942506 | Distal Intergenic |
| chr5 | 115953126 | 115953465 | Distal Intergenic |
| chr5 | 115964652 | 115965302 | Distal Intergenic |
| chr5 | 115971772 | 115972109 | Distal Intergenic |
| chr5 | 115975369 | 115977055 | Distal Intergenic |
| chr5 | 115983933 | 115985655 | Distal Intergenic |
| chr5 | 115993479 | 115993943 | Distal Intergenic |
| chr5 | 115994429 | 115995320 | Distal Intergenic |
| chr5 | 115995459 | 115996044 | Distal Intergenic |
| chr5 | 115997573 | 115997795 | Distal Intergenic |
| chr5 | 116010051 | 116010627 | Distal Intergenic |
| chr5 | 116013775 | 116014417 | Distal Intergenic |
| chr5 | 116022494 | 116022761 | Distal Intergenic |
| chr5 | 116022965 | 116023601 | Distal Intergenic |
| chr5 | 116459105 | 116459322 | Distal Intergenic |
| chr5 | 116459561 | 116460165 | Distal Intergenic |
| chr5 | 116870520 | 116871218 | Intron |
| chr5 | 116872811 | 116873538 | Intron |

| | | | |
|---|---|---|---|
| chr5 | 116877421 | 116878798 | Intron |
| chr5 | 116895996 | 116897088 | Intron |
| chr5 | 117558097 | 117558296 | Distal Intergenic |
| chr5 | 117558467 | 117559022 | Distal Intergenic |
| chr5 | 117565025 | 117565626 | Distal Intergenic |
| chr5 | 117573262 | 117573604 | Distal Intergenic |
| chr5 | 117584273 | 117585019 | Distal Intergenic |
| chr5 | 117608646 | 117609362 | Distal Intergenic |
| chr5 | 117632067 | 117632935 | Distal Intergenic |
| chr5 | 117729242 | 117729442 | Distal Intergenic |
| chr5 | 117810492 | 117811488 | Distal Intergenic |
| chr5 | 117812017 | 117812528 | Distal Intergenic |
| chr5 | 117812566 | 117813470 | Distal Intergenic |
| chr5 | 118339110 | 118340432 | Exon |
| chr5 | 118534528 | 118534808 | Intron |
| chr5 | 118534845 | 118535131 | Intron |
| chr5 | 118898668 | 118899123 | Distal Intergenic |
| chr5 | 118899330 | 118899828 | Distal Intergenic |
| chr5 | 118899953 | 118900196 | Distal Intergenic |
| chr5 | 118901877 | 118902104 | Distal Intergenic |
| chr5 | 118902192 | 118902391 | Distal Intergenic |
| chr5 | 118902482 | 118902762 | Distal Intergenic |
| chr5 | 118909335 | 118909932 | Distal Intergenic |
| chr5 | 119062828 | 119063843 | Distal Intergenic |
| chr5 | 119077386 | 119078004 | Distal Intergenic |
| chr5 | 119099129 | 119100226 | Distal Intergenic |
| chr5 | 119579029 | 119579500 | Distal Intergenic |
| chr5 | 119579992 | 119580282 | Distal Intergenic |
| chr5 | 119590132 | 119590654 | Distal Intergenic |
| chr5 | 119602490 | 119604124 | Distal Intergenic |
| chr5 | 119606388 | 119607108 | Distal Intergenic |
| chr5 | 119617562 | 119617942 | Distal Intergenic |
| chr5 | 119663507 | 119663719 | Distal Intergenic |
| chr5 | 119663892 | 119664112 | Distal Intergenic |
| chr5 | 119749998 | 119750403 | Distal Intergenic |
| chr5 | 120067873 | 120068541 | Distal Intergenic |
| chr5 | 120120187 | 120120620 | Distal Intergenic |
| chr5 | 120965995 | 120966962 | Distal Intergenic |
| chr5 | 120967045 | 120968192 | Distal Intergenic |
| chr5 | 121294954 | 121295316 | Distal Intergenic |
| chr5 | 121654476 | 121654815 | Intron |
| chr5 | 121737428 | 121737935 | Intron |
| chr5 | 121744504 | 121746208 | Intron |
| chr5 | 121843502 | 121844133 | Distal Intergenic |

| | | | |
|---|---|---|---|
| chr5 | 121847210 | 121847745 | Distal Intergenic |
| chr5 | 121888252 | 121889642 | Distal Intergenic |
| chr5 | 121902736 | 121906483 | Distal Intergenic |
| chr5 | 121987298 | 121988077 | Intron |
| chr5 | 122255992 | 122257480 | Intron |
| chr5 | 122269444 | 122269831 | Intron |
| chr5 | 122594920 | 122595935 | Distal Intergenic |
| chr5 | 122616245 | 122617524 | Distal Intergenic |
| chr5 | 122749468 | 122750604 | Intron |
| chr5 | 122754270 | 122754469 | Intron |
| chr5 | 122968584 | 122970225 | Distal Intergenic |
| chr5 | 122974257 | 122975709 | Distal Intergenic |
| chr5 | 122975776 | 122977049 | Distal Intergenic |
| chr5 | 122997348 | 122997596 | Distal Intergenic |
| chr5 | 123094319 | 123094551 | Distal Intergenic |
| chr5 | 123332605 | 123333238 | Distal Intergenic |
| chr5 | 123585780 | 123592021 | Distal Intergenic |
| chr5 | 123592222 | 123592433 | Distal Intergenic |
| chr5 | 123775629 | 123776812 | Distal Intergenic |
| chr5 | 123777092 | 123777911 | Distal Intergenic |
| chr5 | 123814227 | 123815067 | Distal Intergenic |
| chr5 | 124133128 | 124133845 | Distal Intergenic |
| chr5 | 124146048 | 124146270 | Distal Intergenic |
| chr5 | 124146350 | 124146602 | Distal Intergenic |
| chr5 | 124198754 | 124199440 | Distal Intergenic |
| chr5 | 124312694 | 124313679 | Distal Intergenic |
| chr5 | 124313761 | 124314073 | Distal Intergenic |
| chr5 | 124314284 | 124314531 | Distal Intergenic |
| chr5 | 124314674 | 124314881 | Distal Intergenic |
| chr5 | 124315217 | 124315621 | Distal Intergenic |
| chr5 | 124318920 | 124320612 | Distal Intergenic |
| chr5 | 124320649 | 124320911 | Distal Intergenic |
| chr5 | 124428121 | 124428673 | Distal Intergenic |
| chr5 | 124430384 | 124430876 | Distal Intergenic |
| chr5 | 124480541 | 124480742 | Distal Intergenic |
| chr5 | 124480930 | 124481907 | Distal Intergenic |
| chr5 | 124485493 | 124485693 | Distal Intergenic |
| chr5 | 124486015 | 124486240 | Distal Intergenic |
| chr5 | 124496922 | 124498042 | Distal Intergenic |
| chr5 | 124501886 | 124502593 | Distal Intergenic |
| chr5 | 124616139 | 124616388 | Distal Intergenic |
| chr5 | 124616429 | 124616735 | Distal Intergenic |
| chr5 | 124618399 | 124619119 | Distal Intergenic |
| chr5 | 124788745 | 124790515 | Distal Intergenic |

| | | | |
|---|---|---|---|
| chr5 | 124801706 | 124802362 | Distal Intergenic |
| chr5 | 124802524 | 124803789 | Distal Intergenic |
| chr5 | 124874793 | 124875179 | Distal Intergenic |
| chr5 | 124876450 | 124876752 | Distal Intergenic |
| chr5 | 124889931 | 124890178 | Distal Intergenic |
| chr5 | 124892157 | 124892373 | Distal Intergenic |
| chr5 | 124892571 | 124893846 | Distal Intergenic |
| chr5 | 124894196 | 124894449 | Distal Intergenic |
| chr5 | 124894889 | 124895246 | Distal Intergenic |
| chr5 | 124895916 | 124896617 | Distal Intergenic |
| chr5 | 124896757 | 124897031 | Distal Intergenic |
| chr5 | 124932540 | 124933897 | Distal Intergenic |
| chr5 | 124963180 | 124963421 | Distal Intergenic |
| chr5 | 125552735 | 125552934 | Distal Intergenic |
| chr5 | 125561120 | 125562968 | Distal Intergenic |
| chr5 | 125563045 | 125563273 | Distal Intergenic |
| chr5 | 125606091 | 125606492 | Distal Intergenic |
| chr5 | 125614667 | 125614920 | Distal Intergenic |
| chr5 | 125615126 | 125615891 | Distal Intergenic |
| chr5 | 125616125 | 125616478 | Distal Intergenic |
| chr5 | 125620305 | 125622555 | Distal Intergenic |
| chr5 | 125627118 | 125628318 | Distal Intergenic |
| chr5 | 125634955 | 125636577 | Distal Intergenic |
| chr5 | 125644598 | 125646699 | Distal Intergenic |
| chr5 | 125647232 | 125647498 | Distal Intergenic |
| chr5 | 125658773 | 125659039 | Distal Intergenic |
| chr5 | 125669736 | 125670210 | Distal Intergenic |
| chr5 | 125725588 | 125726326 | Intron |
| chr5 | 125726411 | 125727031 | Intron |
| chr5 | 125732536 | 125734186 | Intron |
| chr5 | 125734520 | 125734920 | Intron |
| chr5 | 125869590 | 125870088 | Distal Intergenic |
| chr5 | 125870187 | 125875167 | Downstream |
| chr5 | 125886398 | 125887181 | Intron |
| chr5 | 128090980 | 128091337 | Distal Intergenic |
| chr5 | 128091464 | 128091777 | Distal Intergenic |
| chr5 | 128100422 | 128100955 | Distal Intergenic |
| chr5 | 128101026 | 128101426 | Distal Intergenic |
| chr5 | 129527083 | 129527328 | Distal Intergenic |
| chr5 | 130024339 | 130024636 | Distal Intergenic |
| chr5 | 130029468 | 130029829 | Distal Intergenic |
| chr5 | 130029866 | 130031136 | Distal Intergenic |
| chr5 | 130032598 | 130033149 | Distal Intergenic |
| chr5 | 130033374 | 130033576 | Distal Intergenic |

| | | | |
|---|---|---|---|
| chr5 | 130292428 | 130292733 | Distal Intergenic |
| chr5 | 130292816 | 130293105 | Distal Intergenic |
| chr5 | 130293276 | 130293504 | Distal Intergenic |
| chr5 | 130505338 | 130505614 | Distal Intergenic |
| chr5 | 130539569 | 130539909 | 3' UTR |
| chr5 | 130564992 | 130566397 | Distal Intergenic |
| chr5 | 130584265 | 130586347 | Distal Intergenic |
| chr5 | 130647933 | 130648712 | Intron |
| chr5 | 130653600 | 130654023 | Intron |
| chr5 | 130660539 | 130660743 | Intron |
| chr5 | 130660824 | 130662027 | Intron |
| chr5 | 130662206 | 130662684 | Intron |
| chr5 | 130662861 | 130663939 | Intron |
| chr5 | 130664008 | 130664230 | Intron |
| chr5 | 130730754 | 130731207 | Distal Intergenic |
| chr5 | 131318044 | 131318963 | Intron |
| chr5 | 131991775 | 131992161 | Exon |
| chr5 | 132013120 | 132013522 | Intron |
| chr5 | 132033538 | 132034662 | Intron |
| chr5 | 132042040 | 132042277 | Exon |
| chr5 | 132088870 | 132090168 | 3' UTR |
| chr5 | 132373642 | 132374655 | Distal Intergenic |
| chr5 | 134578629 | 134578873 | Intron |
| chr5 | 134722035 | 134722483 | Intron |
| chr5 | 134912626 | 134913414 | Intron |
| chr5 | 135015994 | 135016370 | Distal Intergenic |
| chr5 | 135027815 | 135028516 | Distal Intergenic |
| chr5 | 135052515 | 135054112 | Distal Intergenic |
| chr5 | 135057023 | 135057904 | Distal Intergenic |
| chr5 | 135087937 | 135088725 | Distal Intergenic |
| chr5 | 135114238 | 135114437 | Distal Intergenic |
| chr5 | 135114610 | 135114837 | Distal Intergenic |
| chr5 | 135114927 | 135115211 | Distal Intergenic |
| chr5 | 135155904 | 135156687 | Distal Intergenic |
| chr5 | 135312304 | 135312581 | Distal Intergenic |
| chr5 | 135312639 | 135313125 | Distal Intergenic |
| chr5 | 135411525 | 135411760 | Distal Intergenic |
| chr5 | 135412061 | 135412384 | Distal Intergenic |
| chr5 | 135426669 | 135427331 | Distal Intergenic |
| chr5 | 135434923 | 135435455 | Distal Intergenic |
| chr5 | 135508740 | 135509728 | Intron |
| chr5 | 135663488 | 135663732 | Intron |
| chr5 | 135869806 | 135870373 | Distal Intergenic |
| chr5 | 136009470 | 136009741 | Distal Intergenic |

| | | | |
|---|---|---|---|
| chr5 | 136096979 | 136098036 | Distal Intergenic |
| chr5 | 136102177 | 136102796 | Distal Intergenic |
| chr5 | 136115850 | 136116385 | Distal Intergenic |
| chr5 | 136119696 | 136119899 | Distal Intergenic |
| chr5 | 136146370 | 136146683 | Distal Intergenic |
| chr5 | 136146735 | 136146942 | Distal Intergenic |
| chr5 | 136178512 | 136179122 | Distal Intergenic |
| chr5 | 136183654 | 136183936 | Distal Intergenic |
| chr5 | 136353619 | 136354182 | Intron |
| chr5 | 136354277 | 136354478 | Intron |
| chr5 | 136355028 | 136356319 | Intron |
| chr5 | 136357962 | 136359870 | Intron |
| chr5 | 136359962 | 136361275 | Intron |
| chr5 | 136361464 | 136361949 | Intron |
| chr5 | 136363275 | 136364100 | Intron |
| chr5 | 136364290 | 136364911 | Intron |
| chr5 | 136368601 | 136369679 | Intron |
| chr5 | 136405689 | 136405948 | Intron |
| chr5 | 136408315 | 136408674 | Intron |
| chr5 | 136409659 | 136410481 | Intron |
| chr5 | 136410884 | 136411093 | Intron |
| chr5 | 136421160 | 136422403 | Intron |
| chr5 | 136478538 | 136478863 | Intron |
| chr5 | 136482003 | 136482898 | Intron |
| chr5 | 136483170 | 136483418 | Intron |
| chr5 | 136490447 | 136490716 | Intron |
| chr5 | 136490753 | 136493374 | Intron |
| chr5 | 136527576 | 136528225 | Intron |
| chr5 | 136569825 | 136570286 | Intron |
| chr5 | 136612049 | 136612259 | Intron |
| chr5 | 136645918 | 136646196 | Intron |
| chr5 | 136646353 | 136648869 | Intron |
| chr5 | 136896882 | 136898197 | Distal Intergenic |
| chr5 | 136941640 | 136942569 | Distal Intergenic |
| chr5 | 136942703 | 136942910 | Distal Intergenic |
| chr5 | 136943326 | 136943727 | Distal Intergenic |
| chr5 | 136954988 | 136956298 | 3' UTR |
| chr5 | 136988936 | 136990908 | Intron |
| chr5 | 137007632 | 137009071 | Promoter |
| chr5 | 137010028 | 137010286 | Promoter |
| chr5 | 137028763 | 137029070 | Promoter |
| chr5 | 137030002 | 137030321 | Intron |
| chr5 | 137034783 | 137035383 | Intron |
| chr5 | 137035488 | 137036324 | Intron |

| | | | |
|---|---|---|---|
| chr5 | 137039848 | 137040497 | Intron |
| chr5 | 137040632 | 137041873 | Promoter |
| chr5 | 137041953 | 137043529 | Promoter |
| chr5 | 137043829 | 137044037 | Intron |
| chr5 | 137044208 | 137044502 | Intron |
| chr5 | 137059882 | 137060114 | Intron |
| chr5 | 137060279 | 137061400 | Intron |
| chr5 | 137067486 | 137068490 | Intron |
| chr5 | 137084736 | 137085051 | Intron |
| chr5 | 137106001 | 137107566 | Intron |
| chr5 | 137107720 | 137110260 | Intron |
| chr5 | 137117428 | 137119088 | Intron |
| chr5 | 137122975 | 137124184 | Intron |
| chr5 | 137128381 | 137128605 | Intron |
| chr5 | 137129194 | 137129558 | Intron |
| chr5 | 137130300 | 137130728 | Intron |
| chr5 | 137132831 | 137133954 | Intron |
| chr5 | 137398861 | 137399402 | Distal Intergenic |
| chr5 | 137429315 | 137429514 | Distal Intergenic |
| chr5 | 137438001 | 137438238 | Distal Intergenic |
| chr5 | 137438371 | 137438580 | Distal Intergenic |
| chr5 | 137548308 | 137549198 | Promoter |
| chr5 | 137555707 | 137556340 | Distal Intergenic |
| chr5 | 137578169 | 137579494 | Distal Intergenic |
| chr5 | 137735128 | 137735344 | Intron |
| chr5 | 137735455 | 137735709 | Exon |
| chr5 | 137815075 | 137815544 | Distal Intergenic |
| chr5 | 137942598 | 137943079 | Distal Intergenic |
| chr5 | 137943118 | 137943362 | Distal Intergenic |
| chr5 | 137974236 | 137974460 | Distal Intergenic |
| chr5 | 138187222 | 138187712 | Intron |
| chr5 | 138388299 | 138389362 | Intron |
| chr5 | 138558260 | 138559178 | Distal Intergenic |
| chr5 | 138628070 | 138628270 | Promoter |
| chr5 | 139183511 | 139183760 | Intron |
| chr5 | 139189324 | 139189798 | Exon |
| chr5 | 139350946 | 139351204 | Intron |
| chr5 | 139374208 | 139374612 | Intron |
| chr5 | 139374655 | 139375682 | Intron |
| chr5 | 139464108 | 139465527 | Distal Intergenic |
| chr5 | 139465720 | 139466175 | Distal Intergenic |
| chr5 | 139469033 | 139470043 | Distal Intergenic |
| chr5 | 139672695 | 139673292 | Intron |
| chr5 | 139705628 | 139706145 | Distal Intergenic |

| | | | |
|---|---|---|---|
| chr5 | 140208209 | 140209291 | Promoter |
| chr5 | 140209429 | 140209691 | Exon |
| chr5 | 140210158 | 140210357 | Intron |
| chr5 | 140212442 | 140212953 | Intron |
| chr5 | 140213495 | 140214364 | Promoter |
| chr5 | 140268682 | 140268942 | Intron |
| chr5 | 140269118 | 140269351 | Intron |
| chr5 | 140273126 | 140273359 | Intron |
| chr5 | 140556733 | 140557181 | Promoter |
| chr5 | 140797671 | 140797993 | Promoter |
| chr5 | 140808922 | 140809371 | Promoter |
| chr5 | 140894361 | 140895224 | 3' UTR |
| chr5 | 140920232 | 140920503 | Intron |
| chr5 | 140990386 | 140990732 | Intron |
| chr5 | 140991244 | 140991674 | Intron |
| chr5 | 141083857 | 141084117 | Distal Intergenic |
| chr5 | 141084613 | 141085218 | Distal Intergenic |
| chr5 | 141085270 | 141085776 | Distal Intergenic |
| chr5 | 141086756 | 141086974 | Distal Intergenic |
| chr5 | 141087014 | 141087247 | Distal Intergenic |
| chr5 | 141156779 | 141158621 | Distal Intergenic |
| chr5 | 142880389 | 142880824 | Distal Intergenic |
| chr5 | 143530915 | 143531147 | Distal Intergenic |
| chr5 | 143555845 | 143557594 | Intron |
| chr5 | 143557943 | 143558586 | Intron |
| chr5 | 143558625 | 143559711 | Intron |
| chr5 | 143575720 | 143576128 | Intron |
| chr5 | 143576312 | 143579320 | Intron |
| chr5 | 143579887 | 143585074 | Promoter |
| chr5 | 143585208 | 143585787 | Promoter |
| chr5 | 143668723 | 143668922 | Intron |
| chr5 | 143682486 | 143682930 | Intron |
| chr5 | 143723012 | 143723679 | Intron |
| chr5 | 143723745 | 143724373 | Intron |
| chr5 | 143724680 | 143725043 | Intron |
| chr5 | 143725544 | 143726102 | Intron |
| chr5 | 143810282 | 143810490 | Intron |
| chr5 | 143810564 | 143810815 | Intron |
| chr5 | 143818030 | 143818882 | Intron |
| chr5 | 143819466 | 143819694 | Intron |
| chr5 | 144026627 | 144026826 | Distal Intergenic |
| chr5 | 144033960 | 144034695 | Distal Intergenic |
| chr5 | 144141228 | 144141514 | Distal Intergenic |
| chr5 | 144175627 | 144176007 | Distal Intergenic |

| | | | |
|---|---|---|---|
| chr5 | 144272994 | 144273194 | Distal Intergenic |
| chr5 | 144278257 | 144279954 | Distal Intergenic |
| chr5 | 144286092 | 144288381 | Distal Intergenic |
| chr5 | 144289004 | 144289551 | Distal Intergenic |
| chr5 | 144379162 | 144379694 | Distal Intergenic |
| chr5 | 144396838 | 144399702 | Distal Intergenic |
| chr5 | 144402673 | 144403707 | Distal Intergenic |
| chr5 | 145049785 | 145050197 | Distal Intergenic |
| chr5 | 145050495 | 145050817 | Distal Intergenic |
| chr5 | 145079025 | 145079224 | Distal Intergenic |
| chr5 | 145595685 | 145596927 | Intron |
| chr5 | 145659120 | 145659319 | Intron |
| chr5 | 145659580 | 145660263 | Intron |
| chr5 | 145665456 | 145665856 | 3' UTR |
| chr5 | 145680098 | 145680399 | Distal Intergenic |
| chr5 | 145691106 | 145692229 | Distal Intergenic |
| chr5 | 145692406 | 145692659 | Distal Intergenic |
| chr5 | 145710846 | 145711714 | Distal Intergenic |
| chr5 | 145713798 | 145714072 | Distal Intergenic |
| chr5 | 145715028 | 145716193 | Distal Intergenic |
| chr5 | 145727972 | 145728373 | Distal Intergenic |
| chr5 | 145728732 | 145729581 | Distal Intergenic |
| chr5 | 145739427 | 145740226 | Distal Intergenic |
| chr5 | 145863260 | 145863906 | Intron |
| chr5 | 145875648 | 145875856 | Intron |
| chr5 | 145876670 | 145877525 | Intron |
| chr5 | 145952277 | 145952555 | Distal Intergenic |
| chr5 | 145965480 | 145965935 | Distal Intergenic |
| chr5 | 147084624 | 147084856 | Intron |
| chr5 | 147633630 | 147633901 | Distal Intergenic |
| chr5 | 147644493 | 147644729 | Distal Intergenic |
| chr5 | 147644774 | 147645070 | Distal Intergenic |
| chr5 | 147801963 | 147804400 | Exon |
| chr5 | 147804513 | 147805455 | Exon |
| chr5 | 147810915 | 147811204 | Intron |
| chr5 | 147887746 | 147887945 | Intron |
| chr5 | 147895094 | 147896615 | Intron |
| chr5 | 147901598 | 147902074 | Intron |
| chr5 | 147902254 | 147902931 | Exon |
| chr5 | 147970420 | 147970901 | Intron |
| chr5 | 148672608 | 148672935 | Intron |
| chr5 | 149179385 | 149179653 | Intron |
| chr5 | 149184873 | 149186732 | Intron |
| chr5 | 149231062 | 149231489 | 3' UTR |

| | | | |
|---|---|---|---|
| chr5 | 149235876 | 149238301 | 3' UTR |
| chr5 | 149246766 | 149247632 | Exon |
| chr5 | 149255335 | 149255583 | Intron |
| chr5 | 149255774 | 149255983 | Intron |
| chr5 | 149301146 | 149301378 | Exon |
| chr5 | 149444553 | 149444843 | Intron |
| chr5 | 149444885 | 149445268 | Intron |
| chr5 | 149445342 | 149445775 | Intron |
| chr5 | 149445813 | 149446125 | Intron |
| chr5 | 149459289 | 149460271 | Promoter |
| chr5 | 149588409 | 149588776 | Intron |
| chr5 | 149589006 | 149589278 | 3' UTR |
| chr5 | 149614289 | 149614927 | Intron |
| chr5 | 149777006 | 149777342 | Intron |
| chr5 | 149777482 | 149777859 | Intron |
| chr5 | 149806387 | 149806605 | Distal Intergenic |
| chr5 | 149807106 | 149807333 | Distal Intergenic |
| chr5 | 149995204 | 149995623 | Intron |
| chr5 | 150191688 | 150193385 | Distal Intergenic |
| chr5 | 151260466 | 151261520 | Intron |
| chr5 | 151402706 | 151403411 | Intron |
| chr5 | 151403489 | 151403745 | Intron |
| chr5 | 151414431 | 151414693 | Intron |
| chr5 | 151424984 | 151426511 | Intron |
| chr5 | 151428079 | 151428755 | Intron |
| chr5 | 151467269 | 151468170 | Intron |
| chr5 | 151509188 | 151509486 | Intron |
| chr5 | 151967103 | 151967776 | Distal Intergenic |
| chr5 | 151969096 | 151969352 | Distal Intergenic |
| chr5 | 151976031 | 151976865 | Distal Intergenic |
| chr5 | 152005831 | 152006359 | Intron |
| chr5 | 152054494 | 152054719 | Intron |
| chr5 | 152209890 | 152210151 | Intron |
| chr6 | 3238285 | 3238742 | Intron |
| chr6 | 4140958 | 4141178 | Intron |
| chr6 | 5079186 | 5079528 | Intron |
| chr6 | 5079929 | 5080295 | Intron |
| chr6 | 5088798 | 5090051 | Intron |
| chr6 | 5247170 | 5248332 | Intron |
| chr6 | 5250137 | 5250344 | Intron |
| chr6 | 5250385 | 5250729 | Intron |
| chr6 | 5250774 | 5251241 | Intron |
| chr6 | 6527618 | 6528857 | Intron |
| chr6 | 6536473 | 6536703 | Intron |

| | | | |
|---|---|---|---|
| chr6 | 6536877 | 6537102 | Intron |
| chr6 | 6538229 | 6539134 | Intron |
| chr6 | 6541198 | 6543412 | Intron |
| chr6 | 6607307 | 6607702 | Intron |
| chr6 | 7165701 | 7166030 | Intron |
| chr6 | 7732888 | 7734289 | Intron |
| chr6 | 7795517 | 7795789 | Intron |
| chr6 | 7795888 | 7796619 | Intron |
| chr6 | 7797016 | 7798854 | Intron |
| chr6 | 7806525 | 7807922 | Intron |
| chr6 | 7807968 | 7808261 | Intron |
| chr6 | 7891814 | 7892207 | Exon |
| chr6 | 7893405 | 7893696 | Intron |
| chr6 | 7893739 | 7894149 | Intron |
| chr6 | 7969310 | 7969649 | Intron |
| chr6 | 7998332 | 7998545 | Intron |
| chr6 | 8302338 | 8302743 | Distal Intergenic |
| chr6 | 11798396 | 11800243 | Distal Intergenic |
| chr6 | 11800349 | 11800617 | Distal Intergenic |
| chr6 | 11824514 | 11826755 | Distal Intergenic |
| chr6 | 11857122 | 11857692 | Distal Intergenic |
| chr6 | 11875420 | 11875740 | Distal Intergenic |
| chr6 | 12762872 | 12763715 | Intron |
| chr6 | 12763788 | 12764467 | Intron |
| chr6 | 13332995 | 13333241 | Distal Intergenic |
| chr6 | 13336558 | 13336803 | Distal Intergenic |
| chr6 | 13350537 | 13352091 | Distal Intergenic |
| chr6 | 13526391 | 13527200 | Distal Intergenic |
| chr6 | 13531877 | 13532094 | Distal Intergenic |
| chr6 | 13627444 | 13627656 | Intron |
| chr6 | 13821028 | 13822693 | Distal Intergenic |
| chr6 | 13832410 | 13833219 | Distal Intergenic |
| chr6 | 17015214 | 17016806 | Distal Intergenic |
| chr6 | 17022804 | 17023120 | Distal Intergenic |
| chr6 | 17023204 | 17023551 | Distal Intergenic |
| chr6 | 17058853 | 17060108 | Distal Intergenic |
| chr6 | 17092569 | 17093722 | Distal Intergenic |
| chr6 | 17120821 | 17121058 | Exon |
| chr6 | 17121189 | 17121772 | Intron |
| chr6 | 17121881 | 17122302 | Intron |
| chr6 | 17143464 | 17144941 | Distal Intergenic |
| chr6 | 17161477 | 17161684 | Distal Intergenic |
| chr6 | 17196686 | 17198267 | Distal Intergenic |
| chr6 | 17209053 | 17209360 | Distal Intergenic |

| | | | |
|---|---|---|---|
| chr6 | 17214937 | 17215485 | Distal Intergenic |
| chr6 | 17215766 | 17216540 | Distal Intergenic |
| chr6 | 17231197 | 17232944 | Distal Intergenic |
| chr6 | 17256019 | 17256750 | Distal Intergenic |
| chr6 | 17256964 | 17257440 | Distal Intergenic |
| chr6 | 17257525 | 17258081 | Distal Intergenic |
| chr6 | 17258549 | 17258888 | Distal Intergenic |
| chr6 | 17259068 | 17259483 | Distal Intergenic |
| chr6 | 17264587 | 17267902 | Distal Intergenic |
| chr6 | 17359021 | 17361257 | Distal Intergenic |
| chr6 | 17413942 | 17417005 | Intron |
| chr6 | 17427934 | 17429995 | Intron |
| chr6 | 17443444 | 17446228 | Intron |
| chr6 | 17446293 | 17450689 | Intron |
| chr6 | 17450808 | 17451260 | Intron |
| chr6 | 17451353 | 17452008 | Intron |
| chr6 | 17453190 | 17457808 | Intron |
| chr6 | 17522525 | 17524274 | Intron |
| chr6 | 17562194 | 17563229 | Distal Intergenic |
| chr6 | 17587546 | 17588327 | Distal Intergenic |
| chr6 | 17668747 | 17670112 | Exon |
| chr6 | 17760883 | 17761639 | 3' UTR |
| chr6 | 18192506 | 18192713 | Intron |
| chr6 | 18220964 | 18221671 | Intron |
| chr6 | 18798628 | 18798967 | Distal Intergenic |
| chr6 | 18811243 | 18811537 | Distal Intergenic |
| chr6 | 19611022 | 19611357 | Distal Intergenic |
| chr6 | 19623041 | 19624444 | Distal Intergenic |
| chr6 | 21580398 | 21580598 | Distal Intergenic |
| chr6 | 22328717 | 22329062 | Distal Intergenic |
| chr6 | 22367169 | 22367561 | Distal Intergenic |
| chr6 | 22393548 | 22393758 | Distal Intergenic |
| chr6 | 22393903 | 22394221 | Distal Intergenic |
| chr6 | 22397113 | 22399438 | Distal Intergenic |
| chr6 | 22741980 | 22742189 | Distal Intergenic |
| chr6 | 22752630 | 22752835 | Distal Intergenic |
| chr6 | 22817478 | 22817677 | Distal Intergenic |
| chr6 | 22894762 | 22895018 | Distal Intergenic |
| chr6 | 22895228 | 22895474 | Distal Intergenic |
| chr6 | 23118777 | 23120523 | Distal Intergenic |
| chr6 | 23132583 | 23133809 | Distal Intergenic |
| chr6 | 24613733 | 24614580 | Intron |
| chr6 | 24656174 | 24657724 | Intron |
| chr6 | 28178333 | 28179423 | Distal Intergenic |

| | | | |
|---|---|---|---|
| chr6 | 28179759 | 28180002 | Distal Intergenic |
| chr6 | 28272411 | 28272787 | Distal Intergenic |
| chr6 | 28272858 | 28273059 | Distal Intergenic |
| chr6 | 28371860 | 28372150 | Distal Intergenic |
| chr6 | 28487603 | 28487834 | Distal Intergenic |
| chr6 | 28573909 | 28574127 | Distal Intergenic |
| chr6 | 28766417 | 28766675 | Distal Intergenic |
| chr6 | 28766779 | 28767185 | Distal Intergenic |
| chr6 | 28771951 | 28774331 | Distal Intergenic |
| chr6 | 28811220 | 28811536 | Distal Intergenic |
| chr6 | 28811887 | 28812989 | Distal Intergenic |
| chr6 | 28814151 | 28815675 | Distal Intergenic |
| chr6 | 28816207 | 28818050 | Distal Intergenic |
| chr6 | 28978256 | 28978698 | Distal Intergenic |
| chr6 | 28980416 | 28980631 | Distal Intergenic |
| chr6 | 28981014 | 28983395 | Distal Intergenic |
| chr6 | 28986042 | 28987466 | Distal Intergenic |
| chr6 | 29023462 | 29024670 | Intron |
| chr6 | 29032358 | 29034566 | Exon |
| chr6 | 29039280 | 29040150 | Intron |
| chr6 | 29114259 | 29114488 | Distal Intergenic |
| chr6 | 29222701 | 29223288 | Distal Intergenic |
| chr6 | 29248851 | 29249176 | Distal Intergenic |
| chr6 | 29258324 | 29258854 | Distal Intergenic |
| chr6 | 29265181 | 29266414 | Distal Intergenic |
| chr6 | 29266726 | 29267807 | Distal Intergenic |
| chr6 | 29298476 | 29298737 | Distal Intergenic |
| chr6 | 29322457 | 29322814 | Downstream |
| chr6 | 29497126 | 29497884 | Promoter |
| chr6 | 29497962 | 29499338 | Promoter |
| chr6 | 29536208 | 29537009 | Intron |
| chr6 | 29559010 | 29560171 | Intron |
| chr6 | 29679604 | 29680706 | Distal Intergenic |
| chr6 | 29685787 | 29686926 | Distal Intergenic |
| chr6 | 29717727 | 29718087 | Promoter |
| chr6 | 29718506 | 29718803 | Promoter |
| chr6 | 29718902 | 29719787 | Promoter |
| chr6 | 29719831 | 29721173 | Promoter |
| chr6 | 29722087 | 29723008 | Distal Intergenic |
| chr6 | 29997311 | 29997675 | Intron |
| chr6 | 29998381 | 29999518 | Intron |
| chr6 | 30006979 | 30007745 | Intron |
| chr6 | 30123245 | 30123473 | Intron |
| chr6 | 30123573 | 30124758 | Exon |

| | | | |
|---|---|---|---|
| chr6 | 30202182 | 30203468 | Exon |
| chr6 | 30250731 | 30251575 | Intron |
| chr6 | 30251628 | 30251911 | Intron |
| chr6 | 30251991 | 30252539 | Intron |
| chr6 | 30252771 | 30253006 | Intron |
| chr6 | 30253260 | 30254007 | Intron |
| chr6 | 30257285 | 30258775 | Exon |
| chr6 | 30296558 | 30297364 | Promoter |
| chr6 | 30302624 | 30303316 | Intron |
| chr6 | 30340738 | 30341001 | Distal Intergenic |
| chr6 | 30458958 | 30460381 | 3' UTR |
| chr6 | 30511966 | 30512586 | 3' UTR |
| chr6 | 30512860 | 30513094 | 3' UTR |
| chr6 | 30999217 | 30999719 | Intron |
| chr6 | 31010829 | 31012560 | Distal Intergenic |
| chr6 | 31015137 | 31016929 | Distal Intergenic |
| chr6 | 31024874 | 31025218 | Intron |
| chr6 | 31025401 | 31025987 | Intron |
| chr6 | 31030496 | 31031260 | Distal Intergenic |
| chr6 | 31031455 | 31032697 | Distal Intergenic |
| chr6 | 31037714 | 31039448 | Distal Intergenic |
| chr6 | 31050579 | 31050778 | Distal Intergenic |
| chr6 | 31057015 | 31057241 | Distal Intergenic |
| chr6 | 31062237 | 31062489 | Distal Intergenic |
| chr6 | 31062570 | 31063104 | Distal Intergenic |
| chr6 | 31063181 | 31065199 | Distal Intergenic |
| chr6 | 31065236 | 31065544 | Distal Intergenic |
| chr6 | 31065877 | 31066086 | Distal Intergenic |
| chr6 | 31074958 | 31077050 | Downstream |
| chr6 | 31170371 | 31170717 | 3' UTR |
| chr6 | 32812516 | 32813658 | Promoter |
| chr6 | 32967952 | 32968944 | Distal Intergenic |
| chr6 | 33198730 | 33199550 | Distal Intergenic |
| chr6 | 33240033 | 33240769 | Promoter |
| chr6 | 33670917 | 33671502 | Intron |
| chr6 | 34297661 | 34298257 | Intron |
| chr6 | 34304123 | 34304329 | Intron |
| chr6 | 34304495 | 34304797 | Intron |
| chr6 | 34334032 | 34334375 | Intron |
| chr6 | 34448272 | 34448504 | Intron |
| chr6 | 34547600 | 34549343 | Distal Intergenic |
| chr6 | 34549474 | 34549938 | Distal Intergenic |
| chr6 | 34550305 | 34550504 | Distal Intergenic |
| chr6 | 34551872 | 34552089 | Downstream |

| | | | |
|---|---|---|---|
| chr6 | 34558333 | 34558999 | 3' UTR |
| chr6 | 34559156 | 34560134 | Intron |
| chr6 | 34560187 | 34561145 | Intron |
| chr6 | 34569431 | 34569727 | Intron |
| chr6 | 34569860 | 34570539 | Intron |
| chr6 | 34572317 | 34575204 | Exon |
| chr6 | 34657004 | 34658957 | Intron |
| chr6 | 34659012 | 34660505 | Intron |
| chr6 | 34660547 | 34661343 | Intron |
| chr6 | 34663411 | 34663753 | Promoter |
| chr6 | 34666779 | 34667814 | Distal Intergenic |
| chr6 | 34668057 | 34668390 | Distal Intergenic |
| chr6 | 34671432 | 34672959 | Distal Intergenic |
| chr6 | 34682255 | 34682564 | Distal Intergenic |
| chr6 | 34686719 | 34687303 | Distal Intergenic |
| chr6 | 34697086 | 34697538 | Distal Intergenic |
| chr6 | 34697783 | 34698220 | Distal Intergenic |
| chr6 | 34699064 | 34700729 | Distal Intergenic |
| chr6 | 34700784 | 34701123 | Distal Intergenic |
| chr6 | 34730522 | 34730862 | Intron |
| chr6 | 34731244 | 34731469 | Intron |
| chr6 | 34746183 | 34748309 | Distal Intergenic |
| chr6 | 34812754 | 34813035 | Intron |
| chr6 | 34813168 | 34813546 | Intron |
| chr6 | 34815696 | 34818038 | Intron |
| chr6 | 34818317 | 34820062 | Intron |
| chr6 | 34902350 | 34903282 | Intron |
| chr6 | 34916260 | 34916489 | Intron |
| chr6 | 34921161 | 34922076 | Intron |
| chr6 | 34970551 | 34970967 | Intron |
| chr6 | 34977376 | 34978234 | Intron |
| chr6 | 35083452 | 35083695 | Intron |
| chr6 | 35374732 | 35375465 | Intron |
| chr6 | 35376086 | 35376321 | Intron |
| chr6 | 35469243 | 35469498 | Intron |
| chr6 | 35488956 | 35489854 | Distal Intergenic |
| chr6 | 35492778 | 35493230 | Distal Intergenic |
| chr6 | 37153162 | 37154197 | Distal Intergenic |
| chr6 | 37306276 | 37307006 | Distal Intergenic |
| chr6 | 37320684 | 37321147 | Promoter |
| chr6 | 37321207 | 37321508 | Promoter |
| chr6 | 37327263 | 37328176 | Intron |
| chr6 | 37335534 | 37335811 | Intron |
| chr6 | 37338070 | 37339704 | Exon |

| | | | |
|---|---|---|---|
| chr6 | 37349258 | 37349624 | Intron |
| chr6 | 37378131 | 37378370 | Distal Intergenic |
| chr6 | 37381219 | 37382260 | Distal Intergenic |
| chr6 | 37405720 | 37406786 | Intron |
| chr6 | 37407389 | 37408893 | Intron |
| chr6 | 37415173 | 37416871 | Intron |
| chr6 | 37471502 | 37471701 | Distal Intergenic |
| chr6 | 37567758 | 37567976 | Distal Intergenic |
| chr6 | 37685622 | 37686624 | Distal Intergenic |
| chr6 | 38188443 | 38188647 | Intron |
| chr6 | 38188977 | 38189230 | Intron |
| chr6 | 38204782 | 38205332 | Intron |
| chr6 | 38303764 | 38304909 | Intron |
| chr6 | 38324389 | 38324708 | Intron |
| chr6 | 38369503 | 38370111 | Intron |
| chr6 | 38383449 | 38383760 | Intron |
| chr6 | 38431096 | 38432985 | Intron |
| chr6 | 38483001 | 38483682 | Intron |
| chr6 | 38500535 | 38500897 | Intron |
| chr6 | 38501003 | 38501202 | Intron |
| chr6 | 38501949 | 38502536 | Intron |
| chr6 | 38575493 | 38576264 | Intron |
| chr6 | 38582673 | 38584021 | Intron |
| chr6 | 38592394 | 38593646 | Intron |
| chr6 | 38594017 | 38595373 | Intron |
| chr6 | 38779628 | 38779980 | Intron |
| chr6 | 38824852 | 38825184 | Intron |
| chr6 | 39193160 | 39194447 | Intron |
| chr6 | 39284066 | 39284376 | Exon |
| chr6 | 39313821 | 39314439 | Exon |
| chr6 | 39460579 | 39460964 | Intron |
| chr6 | 39464748 | 39466308 | Intron |
| chr6 | 39542156 | 39542583 | Intron |
| chr6 | 39542637 | 39542855 | Intron |
| chr6 | 39731250 | 39732808 | Distal Intergenic |
| chr6 | 40053872 | 40054652 | Distal Intergenic |
| chr6 | 40421462 | 40421737 | Intron |
| chr6 | 40689782 | 40690140 | Distal Intergenic |
| chr6 | 41124021 | 41124269 | Downstream |
| chr6 | 41490032 | 41490460 | Distal Intergenic |
| chr6 | 41498230 | 41499122 | Distal Intergenic |
| chr6 | 41506969 | 41507372 | Distal Intergenic |
| chr6 | 41507422 | 41509893 | Distal Intergenic |
| chr6 | 41510843 | 41511256 | Distal Intergenic |

| | | | |
|---|---|---|---|
| chr6 | 42056537 | 42057132 | Distal Intergenic |
| chr6 | 42297924 | 42298276 | Intron |
| chr6 | 42298327 | 42299591 | Intron |
| chr6 | 42300603 | 42300805 | Intron |
| chr6 | 42302401 | 42302668 | Intron |
| chr6 | 42328699 | 42328898 | Intron |
| chr6 | 42329019 | 42329888 | Intron |
| chr6 | 42331526 | 42332426 | Intron |
| chr6 | 42340776 | 42341164 | Intron |
| chr6 | 42345726 | 42347538 | Intron |
| chr6 | 42369291 | 42370112 | Intron |
| chr6 | 42439558 | 42440522 | Distal Intergenic |
| chr6 | 42440701 | 42441061 | Distal Intergenic |
| chr6 | 42491990 | 42492432 | Distal Intergenic |
| chr6 | 42642342 | 42642649 | Intron |
| chr6 | 42998096 | 42999287 | Distal Intergenic |
| chr6 | 43005007 | 43005791 | 3' UTR |
| chr6 | 43009951 | 43010599 | Exon |
| chr6 | 43215619 | 43216873 | Intron |
| chr6 | 43217162 | 43217624 | Intron |
| chr6 | 43241848 | 43242606 | Intron |
| chr6 | 43242672 | 43243081 | Intron |
| chr6 | 43332085 | 43332670 | Intron |
| chr6 | 45791734 | 45792468 | Distal Intergenic |
| chr6 | 47189210 | 47189692 | Distal Intergenic |
| chr6 | 47189760 | 47190300 | Distal Intergenic |
| chr6 | 47219973 | 47220319 | Intron |
| chr6 | 47278319 | 47278580 | Promoter |
| chr6 | 47404413 | 47405498 | Distal Intergenic |
| chr6 | 47405632 | 47405941 | Distal Intergenic |
| chr6 | 47477606 | 47478037 | Intron |
| chr6 | 47510046 | 47510409 | Intron |
| chr6 | 47544448 | 47544650 | Intron |
| chr6 | 47545534 | 47545756 | Intron |
| chr6 | 47738419 | 47738627 | Distal Intergenic |
| chr6 | 47762194 | 47762727 | Intron |
| chr6 | 47763761 | 47764155 | Intron |
| chr6 | 47827870 | 47830053 | Distal Intergenic |
| chr6 | 47871016 | 47871217 | Intron |
| chr6 | 48035957 | 48036174 | Promoter |
| chr6 | 48374627 | 48374910 | Distal Intergenic |
| chr6 | 48382666 | 48382886 | Distal Intergenic |
| chr6 | 48383073 | 48383493 | Distal Intergenic |
| chr6 | 48779048 | 48780637 | Distal Intergenic |

| | | | |
|---|---|---|---|
| chr6 | 49023370 | 49023754 | Distal Intergenic |
| chr6 | 49209934 | 49210485 | Distal Intergenic |
| chr6 | 49212825 | 49214057 | Distal Intergenic |
| chr6 | 49214095 | 49214341 | Distal Intergenic |
| chr6 | 50313541 | 50313784 | Distal Intergenic |
| chr6 | 50327839 | 50330532 | Distal Intergenic |
| chr6 | 50331222 | 50333412 | Distal Intergenic |
| chr6 | 50333547 | 50333805 | Distal Intergenic |
| chr6 | 50333904 | 50334488 | Distal Intergenic |
| chr6 | 50337466 | 50337679 | Distal Intergenic |
| chr6 | 50337715 | 50338519 | Distal Intergenic |
| chr6 | 50339055 | 50341593 | Distal Intergenic |
| chr6 | 50341655 | 50341927 | Distal Intergenic |
| chr6 | 50344864 | 50345109 | Distal Intergenic |
| chr6 | 50345705 | 50346090 | Distal Intergenic |
| chr6 | 50353937 | 50355554 | Distal Intergenic |
| chr6 | 50357539 | 50359551 | Distal Intergenic |
| chr6 | 50359614 | 50359813 | Distal Intergenic |
| chr6 | 50391603 | 50392206 | Distal Intergenic |
| chr6 | 50392340 | 50392539 | Distal Intergenic |
| chr6 | 50404672 | 50405088 | Distal Intergenic |
| chr6 | 50405531 | 50405730 | Distal Intergenic |
| chr6 | 50405867 | 50406155 | Distal Intergenic |
| chr6 | 50406235 | 50406780 | Distal Intergenic |
| chr6 | 50406981 | 50408481 | Distal Intergenic |
| chr6 | 50409764 | 50409987 | Distal Intergenic |
| chr6 | 50410212 | 50410812 | Distal Intergenic |
| chr6 | 50637627 | 50637891 | Distal Intergenic |
| chr6 | 50974513 | 50975391 | Distal Intergenic |
| chr6 | 51009937 | 51011737 | Distal Intergenic |
| chr6 | 51026468 | 51027476 | Distal Intergenic |
| chr6 | 51068836 | 51069355 | Distal Intergenic |
| chr6 | 51069655 | 51070251 | Distal Intergenic |
| chr6 | 51418060 | 51419852 | Distal Intergenic |
| chr6 | 51494257 | 51495124 | Intron |
| chr6 | 51495253 | 51495730 | Intron |
| chr6 | 51503835 | 51504299 | Intron |
| chr6 | 52108747 | 52110269 | Promoter |
| chr6 | 52663490 | 52663689 | Intron |
| chr6 | 52663752 | 52664614 | 5' UTR |
| chr6 | 53384784 | 53385488 | Intron |
| chr6 | 53411897 | 53413615 | Distal Intergenic |
| chr6 | 53447627 | 53447953 | Intron |
| chr6 | 53448109 | 53449044 | Intron |

| | | | |
|---|---|---|---|
| chr6 | 53552520 | 53553070 | Distal Intergenic |
| chr6 | 53573910 | 53574230 | Distal Intergenic |
| chr6 | 53592400 | 53593106 | Distal Intergenic |
| chr6 | 53593251 | 53593870 | Distal Intergenic |
| chr6 | 54040086 | 54040618 | Intron |
| chr6 | 54040854 | 54041088 | Intron |
| chr6 | 54042537 | 54042932 | Intron |
| chr6 | 54046578 | 54046892 | Intron |
| chr6 | 54152901 | 54154264 | Distal Intergenic |
| chr6 | 54654072 | 54654856 | Distal Intergenic |
| chr6 | 54682066 | 54682415 | Distal Intergenic |
| chr6 | 54687601 | 54687958 | Distal Intergenic |
| chr6 | 54767329 | 54767886 | Intron |
| chr6 | 54816722 | 54816921 | Distal Intergenic |
| chr6 | 54817001 | 54817478 | Distal Intergenic |
| chr6 | 54826887 | 54827104 | Distal Intergenic |
| chr6 | 56746600 | 56747265 | Intron |
| chr6 | 56747777 | 56748236 | Intron |
| chr6 | 56750164 | 56750700 | Intron |
| chr6 | 56786922 | 56788802 | Intron |
| chr6 | 56806142 | 56807418 | Intron |
| chr6 | 56810046 | 56810565 | Intron |
| chr6 | 56851673 | 56852058 | Intron |
| chr6 | 57653392 | 57653601 | Distal Intergenic |
| chr6 | 57761457 | 57764096 | Distal Intergenic |
| chr6 | 57774270 | 57774898 | Distal Intergenic |
| chr6 | 58790761 | 58791817 | Distal Intergenic |
| chr6 | 58791882 | 58792546 | Distal Intergenic |
| chr6 | 58797495 | 58797912 | Distal Intergenic |
| chr6 | 59252685 | 59252991 | Distal Intergenic |
| chr6 | 59253090 | 59253942 | Distal Intergenic |
| chr6 | 61156914 | 61157559 | Distal Intergenic |
| chr6 | 61175625 | 61175856 | Distal Intergenic |
| chr6 | 61182964 | 61184844 | Distal Intergenic |
| chr6 | 61192514 | 61193216 | Distal Intergenic |
| chr6 | 61196660 | 61197013 | Distal Intergenic |
| chr6 | 61689573 | 61689810 | Distal Intergenic |
| chr6 | 63848036 | 63848581 | Distal Intergenic |
| chr6 | 64066508 | 64066707 | Distal Intergenic |
| chr6 | 64992651 | 64993120 | Intron |
| chr6 | 65540329 | 65540552 | Intron |
| chr6 | 65556996 | 65557240 | Intron |
| chr6 | 66485365 | 66485670 | Distal Intergenic |
| chr6 | 66859670 | 66859881 | Distal Intergenic |

| | | | |
|---|---|---|---|
| chr6 | 66859930 | 66861779 | Distal Intergenic |
| chr6 | 66946693 | 66947591 | Distal Intergenic |
| chr6 | 66963769 | 66965446 | Distal Intergenic |
| chr6 | 66986031 | 66986360 | Distal Intergenic |
| chr6 | 66987600 | 66987854 | Distal Intergenic |
| chr6 | 66990729 | 66992326 | Distal Intergenic |
| chr6 | 67008103 | 67008589 | Distal Intergenic |
| chr6 | 67008783 | 67009004 | Distal Intergenic |
| chr6 | 67020785 | 67022159 | Distal Intergenic |
| chr6 | 67080696 | 67080926 | Distal Intergenic |
| chr6 | 67081077 | 67081443 | Distal Intergenic |
| chr6 | 67083578 | 67085526 | Distal Intergenic |
| chr6 | 67111183 | 67111382 | Distal Intergenic |
| chr6 | 67216904 | 67217598 | Distal Intergenic |
| chr6 | 67217828 | 67218458 | Distal Intergenic |
| chr6 | 67218809 | 67219575 | Distal Intergenic |
| chr6 | 67250899 | 67251280 | Distal Intergenic |
| chr6 | 67251922 | 67252182 | Distal Intergenic |
| chr6 | 67253043 | 67253896 | Distal Intergenic |
| chr6 | 67263570 | 67263974 | Distal Intergenic |
| chr6 | 67384024 | 67386159 | Distal Intergenic |
| chr6 | 67386223 | 67386432 | Distal Intergenic |
| chr6 | 67391191 | 67392203 | Distal Intergenic |
| chr6 | 67404051 | 67404387 | Distal Intergenic |
| chr6 | 67430870 | 67431587 | Distal Intergenic |
| chr6 | 67431636 | 67432016 | Distal Intergenic |
| chr6 | 67480767 | 67485691 | Distal Intergenic |
| chr6 | 70493830 | 70494463 | Intron |
| chr6 | 70494512 | 70494864 | Intron |
| chr6 | 70583058 | 70583877 | Intron |
| chr6 | 70589265 | 70589484 | 5' UTR |
| chr6 | 70719980 | 70720181 | Intron |
| chr6 | 70723736 | 70724359 | Intron |
| chr6 | 70763718 | 70763921 | Intron |
| chr6 | 70774606 | 70775168 | Intron |
| chr6 | 70829775 | 70831023 | Intron |
| chr6 | 71038566 | 71039941 | Distal Intergenic |
| chr6 | 71221767 | 71222305 | Intron |
| chr6 | 71493900 | 71494686 | Intron |
| chr6 | 71540927 | 71541176 | Intron |
| chr6 | 71541252 | 71541837 | Intron |
| chr6 | 71858916 | 71859193 | Distal Intergenic |
| chr6 | 72006323 | 72006730 | Exon |
| chr6 | 72167295 | 72167826 | Distal Intergenic |

| | | | |
|---|---|---|---|
| chr6 | 72167886 | 72168187 | Distal Intergenic |
| chr6 | 72266003 | 72266271 | Distal Intergenic |
| chr6 | 72294799 | 72295072 | Distal Intergenic |
| chr6 | 72297517 | 72297953 | Distal Intergenic |
| chr6 | 72319564 | 72322360 | Distal Intergenic |
| chr6 | 72339610 | 72340057 | Distal Intergenic |
| chr6 | 72340113 | 72340418 | Distal Intergenic |
| chr6 | 72423386 | 72424009 | Distal Intergenic |
| chr6 | 72497773 | 72502819 | Distal Intergenic |
| chr6 | 72566605 | 72566832 | Distal Intergenic |
| chr6 | 72626944 | 72627577 | Intron |
| chr6 | 72627638 | 72628019 | Intron |
| chr6 | 72748400 | 72748624 | Intron |
| chr6 | 72748717 | 72749022 | Intron |
| chr6 | 72806859 | 72807811 | Exon |
| chr6 | 72822335 | 72822876 | Intron |
| chr6 | 72831219 | 72832245 | Intron |
| chr6 | 72838969 | 72839169 | Intron |
| chr6 | 72848398 | 72849600 | Intron |
| chr6 | 72850178 | 72850976 | Intron |
| chr6 | 72876593 | 72877698 | Intron |
| chr6 | 72907731 | 72908221 | Intron |
| chr6 | 73198390 | 73198629 | Distal Intergenic |
| chr6 | 76915037 | 76915575 | Distal Intergenic |
| chr6 | 76916845 | 76917207 | Distal Intergenic |
| chr6 | 81896022 | 81896423 | Distal Intergenic |
| chr6 | 81970731 | 81971208 | Distal Intergenic |
| chr6 | 81982092 | 81982837 | Distal Intergenic |
| chr6 | 81985338 | 81986190 | Distal Intergenic |
| chr6 | 81990886 | 81991089 | Distal Intergenic |
| chr6 | 81991259 | 81991540 | Distal Intergenic |
| chr6 | 81991708 | 81992474 | Distal Intergenic |
| chr6 | 82002250 | 82002914 | Distal Intergenic |
| chr6 | 82047448 | 82048136 | Distal Intergenic |
| chr6 | 82048229 | 82048976 | Distal Intergenic |
| chr6 | 82049083 | 82049471 | Distal Intergenic |
| chr6 | 82812703 | 82813908 | Distal Intergenic |
| chr6 | 83085808 | 83087873 | Distal Intergenic |
| chr6 | 83091902 | 83093781 | Distal Intergenic |
| chr6 | 83094848 | 83095375 | Distal Intergenic |
| chr6 | 83095417 | 83095720 | Distal Intergenic |
| chr6 | 83383251 | 83383451 | Distal Intergenic |
| chr6 | 83385268 | 83385570 | Distal Intergenic |
| chr6 | 83391763 | 83392735 | Distal Intergenic |

| | | | |
|---|---|---|---|
| chr6 | 83392779 | 83393485 | Distal Intergenic |
| chr6 | 83393548 | 83393799 | Distal Intergenic |
| chr6 | 83406903 | 83407596 | Distal Intergenic |
| chr6 | 83439027 | 83439290 | Distal Intergenic |
| chr6 | 83445942 | 83446146 | Distal Intergenic |
| chr6 | 83446186 | 83446611 | Distal Intergenic |
| chr6 | 83446782 | 83447127 | Distal Intergenic |
| chr6 | 83451486 | 83453311 | Distal Intergenic |
| chr6 | 83724573 | 83725013 | Intron |
| chr6 | 84128554 | 84129240 | Intron |
| chr6 | 84129306 | 84129573 | Intron |
| chr6 | 84454425 | 84455272 | Distal Intergenic |
| chr6 | 84493712 | 84493966 | Distal Intergenic |
| chr6 | 84508732 | 84512150 | Distal Intergenic |
| chr6 | 84515038 | 84516844 | Distal Intergenic |
| chr6 | 84521052 | 84521298 | Distal Intergenic |
| chr6 | 84551082 | 84551794 | Distal Intergenic |
| chr6 | 84725846 | 84726560 | Distal Intergenic |
| chr6 | 84842938 | 84843428 | Intron |
| chr6 | 84870330 | 84870667 | Exon |
| chr6 | 84870712 | 84871686 | Exon |
| chr6 | 84963420 | 84964328 | Distal Intergenic |
| chr6 | 85024720 | 85025393 | Distal Intergenic |
| chr6 | 85036644 | 85037207 | Distal Intergenic |
| chr6 | 85037259 | 85038147 | Distal Intergenic |
| chr6 | 85049121 | 85049448 | Distal Intergenic |
| chr6 | 85298340 | 85301155 | Distal Intergenic |
| chr6 | 85316695 | 85317063 | Distal Intergenic |
| chr6 | 85323566 | 85323787 | Distal Intergenic |
| chr6 | 85332340 | 85332748 | Distal Intergenic |
| chr6 | 85401991 | 85402441 | Intron |
| chr6 | 85887383 | 85887612 | Distal Intergenic |
| chr6 | 85910999 | 85911884 | Distal Intergenic |
| chr6 | 86099293 | 86099528 | Intron |
| chr6 | 86099825 | 86100315 | Exon |
| chr6 | 86107511 | 86107920 | Distal Intergenic |
| chr6 | 86130443 | 86130745 | Distal Intergenic |
| chr6 | 86168777 | 86170126 | Intron |
| chr6 | 86268371 | 86269442 | Intron |
| chr6 | 86274008 | 86276076 | 5' UTR |
| chr6 | 86316037 | 86316448 | Downstream |
| chr6 | 86321527 | 86321894 | 3' UTR |
| chr6 | 86344842 | 86345973 | Intron |
| chr6 | 86354326 | 86355633 | Distal Intergenic |

| | | | |
|---|---|---|---|
| chr6 | 86387990 | 86388260 | Promoter |
| chr6 | 86388901 | 86389105 | Promoter |
| chr6 | 86389195 | 86389425 | Promoter |
| chr6 | 86474454 | 86474679 | Distal Intergenic |
| chr6 | 86474902 | 86475386 | Distal Intergenic |
| chr6 | 86475503 | 86475917 | Distal Intergenic |
| chr6 | 86476691 | 86477697 | Distal Intergenic |
| chr6 | 86550243 | 86550472 | Distal Intergenic |
| chr6 | 86555525 | 86555904 | Distal Intergenic |
| chr6 | 86556008 | 86556606 | Distal Intergenic |
| chr6 | 86589845 | 86590662 | Distal Intergenic |
| chr6 | 86799024 | 86799321 | Distal Intergenic |
| chr6 | 86922455 | 86923207 | Distal Intergenic |
| chr6 | 87018738 | 87020091 | Distal Intergenic |
| chr6 | 87136218 | 87137938 | Distal Intergenic |
| chr6 | 87172515 | 87172804 | Distal Intergenic |
| chr6 | 87172930 | 87177728 | Distal Intergenic |
| chr6 | 87186857 | 87188241 | Distal Intergenic |
| chr6 | 87208257 | 87210061 | Distal Intergenic |
| chr6 | 87215118 | 87215543 | Distal Intergenic |
| chr6 | 87221419 | 87222779 | Distal Intergenic |
| chr6 | 87222865 | 87223252 | Distal Intergenic |
| chr6 | 87274949 | 87275950 | Distal Intergenic |
| chr6 | 87336572 | 87336771 | Distal Intergenic |
| chr6 | 87460094 | 87460502 | Distal Intergenic |
| chr6 | 87788907 | 87789517 | Distal Intergenic |
| chr6 | 87789556 | 87789816 | Distal Intergenic |
| chr6 | 87799964 | 87800205 | Intron |
| chr6 | 87800365 | 87800726 | Intron |
| chr6 | 87818637 | 87819011 | Distal Intergenic |
| chr6 | 87994349 | 87994867 | 5' UTR |
| chr6 | 88003529 | 88003734 | Intron |
| chr6 | 88003773 | 88004133 | Intron |
| chr6 | 88415393 | 88415618 | Distal Intergenic |
| chr6 | 88468415 | 88468750 | Distal Intergenic |
| chr6 | 88524153 | 88525028 | Intron |
| chr6 | 88713382 | 88715128 | Distal Intergenic |
| chr6 | 88723509 | 88725374 | Distal Intergenic |
| chr6 | 88874438 | 88874863 | Promoter |
| chr6 | 89363327 | 89364643 | Intron |
| chr6 | 89372835 | 89373087 | Intron |
| chr6 | 89411517 | 89411719 | Intron |
| chr6 | 89502248 | 89502782 | Intron |
| chr6 | 89502824 | 89503069 | Intron |

| | | | |
|---|---|---|---|
| chr6 | 89826098 | 89827548 | Promoter |
| chr6 | 90298620 | 90298872 | Intron |
| chr6 | 90309231 | 90310011 | Intron |
| chr6 | 90310108 | 90311079 | Intron |
| chr6 | 90456993 | 90457199 | Exon |
| chr6 | 91151792 | 91153585 | Distal Intergenic |
| chr6 | 91157645 | 91160266 | Distal Intergenic |
| chr6 | 91160333 | 91161821 | Distal Intergenic |
| chr6 | 91162479 | 91162949 | Distal Intergenic |
| chr6 | 91164396 | 91165271 | Distal Intergenic |
| chr6 | 91182882 | 91183085 | Distal Intergenic |
| chr6 | 91282860 | 91283788 | Intron |
| chr6 | 91322701 | 91322914 | Distal Intergenic |
| chr6 | 91339262 | 91339952 | Distal Intergenic |
| chr6 | 91401905 | 91402566 | Distal Intergenic |
| chr6 | 91404743 | 91404978 | Distal Intergenic |
| chr6 | 91417395 | 91417610 | Distal Intergenic |
| chr6 | 91422884 | 91423207 | Distal Intergenic |
| chr6 | 91423643 | 91424166 | Distal Intergenic |
| chr6 | 91424268 | 91426813 | Distal Intergenic |
| chr6 | 91433230 | 91433471 | Distal Intergenic |
| chr6 | 91439067 | 91439307 | Distal Intergenic |
| chr6 | 91526702 | 91527011 | Distal Intergenic |
| chr6 | 91529110 | 91531562 | Distal Intergenic |
| chr6 | 91596660 | 91597077 | Distal Intergenic |
| chr6 | 91597114 | 91597496 | Distal Intergenic |
| chr6 | 91597533 | 91597970 | Distal Intergenic |
| chr6 | 91602146 | 91602571 | Distal Intergenic |
| chr6 | 91602671 | 91602932 | Distal Intergenic |
| chr6 | 91606489 | 91606688 | Distal Intergenic |
| chr6 | 91638982 | 91639824 | Distal Intergenic |
| chr6 | 91676345 | 91677492 | Distal Intergenic |
| chr6 | 92046825 | 92047277 | Distal Intergenic |
| chr6 | 92189546 | 92191535 | Distal Intergenic |
| chr6 | 92374285 | 92376346 | Intron |
| chr6 | 92377376 | 92377778 | Intron |
| chr6 | 92377828 | 92378067 | Intron |
| chr6 | 92400009 | 92400251 | Exon |
| chr6 | 93207251 | 93207865 | Distal Intergenic |
| chr6 | 93874143 | 93874499 | Distal Intergenic |
| chr6 | 94050376 | 94050793 | Intron |
| chr6 | 94051244 | 94051488 | Intron |
| chr6 | 94105698 | 94106562 | Intron |
| chr6 | 94160681 | 94160965 | Distal Intergenic |

| | | | |
|---|---|---|---|
| chr6 | 94161026 | 94161321 | Distal Intergenic |
| chr6 | 94175150 | 94175752 | Distal Intergenic |
| chr6 | 94175898 | 94176585 | Distal Intergenic |
| chr6 | 94213541 | 94215796 | Distal Intergenic |
| chr6 | 94224862 | 94225543 | Distal Intergenic |
| chr6 | 94225673 | 94225877 | Distal Intergenic |
| chr6 | 94577104 | 94577525 | Distal Intergenic |
| chr6 | 94577843 | 94578083 | Distal Intergenic |
| chr6 | 94578239 | 94578539 | Distal Intergenic |
| chr6 | 94584824 | 94586451 | Distal Intergenic |
| chr6 | 94589169 | 94589483 | Distal Intergenic |
| chr6 | 94589542 | 94590932 | Distal Intergenic |
| chr6 | 94595963 | 94596716 | Distal Intergenic |
| chr6 | 94596769 | 94597046 | Distal Intergenic |
| chr6 | 94597232 | 94597471 | Distal Intergenic |
| chr6 | 94694866 | 94695215 | Distal Intergenic |
| chr6 | 94695425 | 94695624 | Distal Intergenic |
| chr6 | 94695702 | 94696528 | Distal Intergenic |
| chr6 | 96002934 | 96003256 | Distal Intergenic |
| chr6 | 96003817 | 96004254 | Distal Intergenic |
| chr6 | 96021029 | 96021276 | Distal Intergenic |
| chr6 | 97097475 | 97098688 | Distal Intergenic |
| chr6 | 97128165 | 97129593 | Distal Intergenic |
| chr6 | 97129771 | 97130036 | Distal Intergenic |
| chr6 | 97155211 | 97155544 | Distal Intergenic |
| chr6 | 97243045 | 97244919 | 3' UTR |
| chr6 | 97253782 | 97256338 | Intron |
| chr6 | 97260962 | 97263149 | Intron |
| chr6 | 97265200 | 97266733 | Intron |
| chr6 | 97266802 | 97267030 | Intron |
| chr6 | 97305975 | 97306395 | Distal Intergenic |
| chr6 | 97396087 | 97397961 | Intron |
| chr6 | 97456986 | 97457350 | Promoter |
| chr6 | 97460036 | 97462419 | Intron |
| chr6 | 97508804 | 97511233 | Exon |
| chr6 | 97513446 | 97513787 | Intron |
| chr6 | 97513842 | 97514181 | Intron |
| chr6 | 97514325 | 97514551 | Intron |
| chr6 | 97514598 | 97515018 | Intron |
| chr6 | 97515065 | 97515350 | Intron |
| chr6 | 97524590 | 97526143 | Intron |
| chr6 | 97540903 | 97542634 | Intron |
| chr6 | 97545573 | 97545892 | Intron |
| chr6 | 97546036 | 97546792 | Intron |

| | | | |
|---|---|---|---|
| chr6 | 97563084 | 97564147 | Intron |
| chr6 | 97565015 | 97567842 | Intron |
| chr6 | 98899193 | 98900625 | Distal Intergenic |
| chr6 | 98904276 | 98905976 | Distal Intergenic |
| chr6 | 98976958 | 98977493 | Distal Intergenic |
| chr6 | 99034805 | 99035377 | Distal Intergenic |
| chr6 | 99043059 | 99043529 | Distal Intergenic |
| chr6 | 99045635 | 99046532 | Distal Intergenic |
| chr6 | 99149033 | 99149904 | Distal Intergenic |
| chr6 | 99167393 | 99168384 | Distal Intergenic |
| chr6 | 99199770 | 99200116 | Distal Intergenic |
| chr6 | 99293721 | 99294012 | Distal Intergenic |
| chr6 | 99294158 | 99294645 | Distal Intergenic |
| chr6 | 99300166 | 99301154 | Distal Intergenic |
| chr6 | 99320078 | 99320422 | Downstream |
| chr6 | 99325495 | 99325827 | Intron |
| chr6 | 99342720 | 99343232 | Intron |
| chr6 | 99393739 | 99393950 | Intron |
| chr6 | 99396277 | 99398495 | Promoter |
| chr6 | 99415808 | 99417712 | Distal Intergenic |
| chr6 | 99417751 | 99417956 | Distal Intergenic |
| chr6 | 99580274 | 99580676 | Distal Intergenic |
| chr6 | 99612223 | 99612599 | Distal Intergenic |
| chr6 | 99612928 | 99613529 | Distal Intergenic |
| chr6 | 100748679 | 100751247 | Distal Intergenic |
| chr6 | 100784728 | 100784981 | Distal Intergenic |
| chr6 | 100785086 | 100785343 | Distal Intergenic |
| chr6 | 101278753 | 101278976 | Intron |
| chr6 | 101376981 | 101377451 | Distal Intergenic |
| chr6 | 101414653 | 101415734 | Distal Intergenic |
| chr6 | 103294720 | 103296268 | Distal Intergenic |
| chr6 | 103296305 | 103296626 | Distal Intergenic |
| chr6 | 103296782 | 103297249 | Distal Intergenic |
| chr6 | 103309715 | 103309914 | Distal Intergenic |
| chr6 | 103330710 | 103331188 | Distal Intergenic |
| chr6 | 103339301 | 103339500 | Distal Intergenic |
| chr6 | 103350883 | 103351719 | Distal Intergenic |
| chr6 | 103460639 | 103461036 | Distal Intergenic |
| chr6 | 103475443 | 103475759 | Distal Intergenic |
| chr6 | 103475823 | 103476083 | Distal Intergenic |
| chr6 | 105498123 | 105498573 | Intron |
| chr6 | 108018934 | 108019297 | Distal Intergenic |
| chr6 | 108392895 | 108393144 | Intron |
| chr6 | 108397999 | 108398461 | Distal Intergenic |

| | | | |
|---|---|---|---|
| chr6 | 108398707 | 108399026 | Distal Intergenic |
| chr6 | 108401265 | 108402108 | Distal Intergenic |
| chr6 | 108402208 | 108402635 | Distal Intergenic |
| chr6 | 108480901 | 108482573 | Distal Intergenic |
| chr6 | 108484826 | 108485025 | Distal Intergenic |
| chr6 | 108609870 | 108610219 | Distal Intergenic |
| chr6 | 108610705 | 108610904 | Distal Intergenic |
| chr6 | 108615397 | 108615687 | Promoter |
| chr6 | 108654444 | 108656254 | Intron |
| chr6 | 108733091 | 108733368 | Intron |
| chr6 | 108733530 | 108734470 | Intron |
| chr6 | 108744706 | 108746955 | Intron |
| chr6 | 109130604 | 109130848 | Distal Intergenic |
| chr6 | 112981203 | 112981407 | Distal Intergenic |
| chr6 | 112984376 | 112985039 | Distal Intergenic |
| chr6 | 112995656 | 112995998 | Distal Intergenic |
| chr6 | 112996575 | 112996865 | Distal Intergenic |
| chr6 | 112996959 | 112997264 | Distal Intergenic |
| chr6 | 113022701 | 113022925 | Distal Intergenic |
| chr6 | 113023103 | 113023324 | Distal Intergenic |
| chr6 | 113024824 | 113026088 | Distal Intergenic |
| chr6 | 113027509 | 113029478 | Distal Intergenic |
| chr6 | 113029553 | 113029805 | Distal Intergenic |
| chr6 | 113033305 | 113034746 | Distal Intergenic |
| chr6 | 113049081 | 113050220 | Distal Intergenic |
| chr6 | 113116279 | 113117449 | Distal Intergenic |
| chr6 | 113117582 | 113117819 | Distal Intergenic |
| chr6 | 113257534 | 113258380 | Distal Intergenic |
| chr6 | 113327511 | 113328094 | Distal Intergenic |
| chr6 | 113421348 | 113422169 | Distal Intergenic |
| chr6 | 113430164 | 113431867 | Distal Intergenic |
| chr6 | 113436530 | 113437680 | Distal Intergenic |
| chr6 | 113472982 | 113473304 | Distal Intergenic |
| chr6 | 113640480 | 113640996 | Distal Intergenic |
| chr6 | 113645836 | 113646725 | Distal Intergenic |
| chr6 | 114371013 | 114372103 | Intron |
| chr6 | 114373222 | 114374675 | Intron |
| chr6 | 114398552 | 114400317 | Intron |
| chr6 | 114408380 | 114408824 | Intron |
| chr6 | 114705359 | 114706931 | Distal Intergenic |
| chr6 | 114755750 | 114756608 | Distal Intergenic |
| chr6 | 114811424 | 114811665 | Distal Intergenic |
| chr6 | 114816068 | 114816663 | Distal Intergenic |
| chr6 | 114824534 | 114826316 | Distal Intergenic |

| | | | |
|---|---|---|---|
| chr6 | 114832922 | 114834013 | Distal Intergenic |
| chr6 | 114838677 | 114839326 | Distal Intergenic |
| chr6 | 114870249 | 114872236 | Distal Intergenic |
| chr6 | 114880172 | 114880427 | Distal Intergenic |
| chr6 | 114880790 | 114882583 | Distal Intergenic |
| chr6 | 114882803 | 114887652 | Distal Intergenic |
| chr6 | 114889315 | 114889772 | Distal Intergenic |
| chr6 | 114894598 | 114895571 | Distal Intergenic |
| chr6 | 114913844 | 114914243 | Distal Intergenic |
| chr6 | 114914413 | 114914848 | Distal Intergenic |
| chr6 | 114917503 | 114919033 | Distal Intergenic |
| chr6 | 114937409 | 114940141 | Distal Intergenic |
| chr6 | 114955038 | 114956561 | Distal Intergenic |
| chr6 | 114956910 | 114957109 | Distal Intergenic |
| chr6 | 115625467 | 115625764 | Distal Intergenic |
| chr6 | 115626189 | 115626567 | Distal Intergenic |
| chr6 | 115708743 | 115708990 | Distal Intergenic |
| chr6 | 115709066 | 115710158 | Distal Intergenic |
| chr6 | 115758069 | 115759114 | Distal Intergenic |
| chr6 | 115852053 | 115852427 | Distal Intergenic |
| chr6 | 116216111 | 116216813 | Distal Intergenic |
| chr6 | 116356562 | 116358306 | Intron |
| chr6 | 116369689 | 116369966 | Intron |
| chr6 | 117666694 | 117666914 | Intron |
| chr6 | 117791456 | 117791827 | Intron |
| chr6 | 117792192 | 117792406 | Intron |
| chr6 | 117792666 | 117792907 | Intron |
| chr6 | 117792949 | 117793177 | Intron |
| chr6 | 117828968 | 117829199 | Intron |
| chr6 | 117829283 | 117829486 | Intron |
| chr6 | 117829594 | 117829868 | Intron |
| chr6 | 117829978 | 117830534 | Intron |
| chr6 | 117856191 | 117856560 | Intron |
| chr6 | 117857245 | 117858000 | Intron |
| chr6 | 117858203 | 117858896 | Exon |
| chr6 | 117863036 | 117863552 | Intron |
| chr6 | 117865773 | 117867222 | 3' UTR |
| chr6 | 117867331 | 117868396 | Intron |
| chr6 | 118404043 | 118405145 | Intron |
| chr6 | 118405248 | 118405520 | Intron |
| chr6 | 119439953 | 119440183 | Intron |
| chr6 | 119613230 | 119613467 | Intron |
| chr6 | 119732350 | 119733690 | Distal Intergenic |
| chr6 | 119811680 | 119811995 | Promoter |

| | | | |
|---|---|---|---|
| chr6 | 119814248 | 119814974 | Distal Intergenic |
| chr6 | 119874618 | 119875033 | Distal Intergenic |
| chr6 | 119960571 | 119960839 | Distal Intergenic |
| chr6 | 119973790 | 119974002 | Distal Intergenic |
| chr6 | 119986765 | 119987351 | Distal Intergenic |
| chr6 | 119989569 | 119989786 | Distal Intergenic |
| chr6 | 119989845 | 119990281 | Distal Intergenic |
| chr6 | 120048409 | 120048811 | Distal Intergenic |
| chr6 | 120053818 | 120054053 | Distal Intergenic |
| chr6 | 120064709 | 120065275 | Distal Intergenic |
| chr6 | 120066729 | 120067595 | Distal Intergenic |
| chr6 | 120095141 | 120095549 | Distal Intergenic |
| chr6 | 120122043 | 120122959 | Distal Intergenic |
| chr6 | 120123170 | 120123571 | Distal Intergenic |
| chr6 | 120139666 | 120141395 | Distal Intergenic |
| chr6 | 120141503 | 120142691 | Distal Intergenic |
| chr6 | 120142889 | 120143355 | Distal Intergenic |
| chr6 | 120187206 | 120188490 | Distal Intergenic |
| chr6 | 120212235 | 120214961 | Distal Intergenic |
| chr6 | 120307040 | 120307411 | Distal Intergenic |
| chr6 | 120312830 | 120314324 | Distal Intergenic |
| chr6 | 120592021 | 120592420 | Distal Intergenic |
| chr6 | 120747815 | 120750556 | Distal Intergenic |
| chr6 | 120781519 | 120783032 | Distal Intergenic |
| chr6 | 120785983 | 120787536 | Distal Intergenic |
| chr6 | 120815879 | 120816273 | Distal Intergenic |
| chr6 | 120822931 | 120824761 | Distal Intergenic |
| chr6 | 120866938 | 120867288 | Distal Intergenic |
| chr6 | 120877148 | 120877472 | Distal Intergenic |
| chr6 | 120877517 | 120878278 | Distal Intergenic |
| chr6 | 120937186 | 120937963 | Distal Intergenic |
| chr6 | 120945109 | 120948188 | Distal Intergenic |
| chr6 | 120949363 | 120949562 | Distal Intergenic |
| chr6 | 120967156 | 120967509 | Distal Intergenic |
| chr6 | 120967670 | 120968267 | Distal Intergenic |
| chr6 | 120985224 | 120986085 | Distal Intergenic |
| chr6 | 120989359 | 120989755 | Distal Intergenic |
| chr6 | 120993319 | 120995982 | Distal Intergenic |
| chr6 | 121013968 | 121015043 | Distal Intergenic |
| chr6 | 121015209 | 121017952 | Distal Intergenic |
| chr6 | 121018055 | 121018261 | Distal Intergenic |
| chr6 | 121018344 | 121018877 | Distal Intergenic |
| chr6 | 121035092 | 121038825 | Distal Intergenic |
| chr6 | 121143901 | 121144179 | Distal Intergenic |

| | | | |
|---|---|---|---|
| chr6 | 121144225 | 121144998 | Distal Intergenic |
| chr6 | 122259064 | 122260152 | Distal Intergenic |
| chr6 | 122551965 | 122552634 | Distal Intergenic |
| chr6 | 122552706 | 122553196 | Distal Intergenic |
| chr6 | 122760638 | 122761002 | Distal Intergenic |
| chr6 | 122761455 | 122762117 | Downstream |
| chr6 | 122769524 | 122769764 | Intron |
| chr6 | 122776523 | 122776758 | Intron |
| chr6 | 122818966 | 122819166 | Intron |
| chr6 | 122819218 | 122820463 | Intron |
| chr6 | 122954409 | 122955581 | 5' UTR |
| chr6 | 124365225 | 124365540 | Intron |
| chr6 | 124435040 | 124435290 | Intron |
| chr6 | 124633434 | 124634488 | Intron |
| chr6 | 124647093 | 124648147 | Intron |
| chr6 | 124650129 | 124650945 | Intron |
| chr6 | 124662327 | 124662538 | Intron |
| chr6 | 124667094 | 124669494 | Intron |
| chr6 | 124688438 | 124689391 | Intron |
| chr6 | 124689870 | 124690285 | Intron |
| chr6 | 124690361 | 124691853 | Intron |
| chr6 | 124707106 | 124707471 | Intron |
| chr6 | 124755135 | 124756519 | Intron |
| chr6 | 124763229 | 124763514 | Intron |
| chr6 | 124763629 | 124764168 | Intron |
| chr6 | 124905080 | 124905965 | Intron |
| chr6 | 124915494 | 124915701 | Intron |
| chr6 | 124915868 | 124916349 | Intron |
| chr6 | 124989843 | 124990276 | Intron |
| chr6 | 125017739 | 125018125 | Intron |
| chr6 | 125081365 | 125081810 | Intron |
| chr6 | 125110830 | 125111248 | Intron |
| chr6 | 125114505 | 125114992 | Intron |
| chr6 | 125116243 | 125116756 | Intron |
| chr6 | 125116820 | 125117071 | Intron |
| chr6 | 125141921 | 125142890 | Intron |
| chr6 | 125205665 | 125206898 | Distal Intergenic |
| chr6 | 125263658 | 125263911 | Intron |
| chr6 | 125299486 | 125300171 | Intron |
| chr6 | 125300208 | 125300821 | Intron |
| chr6 | 125302220 | 125302466 | Intron |
| chr6 | 125302619 | 125303006 | Intron |
| chr6 | 125304061 | 125304915 | Promoter |
| chr6 | 125327177 | 125328823 | Intron |

| | | | |
|---|---|---|---|
| chr6 | 125363653 | 125363915 | Intron |
| chr6 | 125371602 | 125372543 | Intron |
| chr6 | 125372760 | 125376447 | Intron |
| chr6 | 125383603 | 125384155 | Intron |
| chr6 | 125417836 | 125418862 | Distal Intergenic |
| chr6 | 125428186 | 125429002 | Distal Intergenic |
| chr6 | 125430712 | 125434883 | Distal Intergenic |
| chr6 | 125442432 | 125444209 | Distal Intergenic |
| chr6 | 125453179 | 125453445 | Distal Intergenic |
| chr6 | 125453609 | 125453997 | Distal Intergenic |
| chr6 | 125898085 | 125898408 | Distal Intergenic |
| chr6 | 125948332 | 125948721 | Distal Intergenic |
| chr6 | 125948911 | 125949238 | Distal Intergenic |
| chr6 | 125995309 | 125995910 | Promoter |
| chr6 | 126059420 | 126059623 | Intron |
| chr6 | 126121841 | 126122974 | Intron |
| chr6 | 126188403 | 126189640 | Intron |
| chr6 | 126241498 | 126242149 | Exon |
| chr6 | 126297220 | 126298074 | Intron |
| chr6 | 126315697 | 126317148 | Exon |
| chr6 | 126317193 | 126317401 | Intron |
| chr6 | 126324554 | 126325904 | Intron |
| chr6 | 126325964 | 126326163 | Intron |
| chr6 | 127099036 | 127099497 | Intron |
| chr6 | 127099545 | 127100393 | Intron |
| chr6 | 127275935 | 127276440 | Intron |
| chr6 | 127279195 | 127280364 | Intron |
| chr6 | 127280623 | 127281040 | Intron |
| chr6 | 127325338 | 127326173 | Intron |
| chr6 | 127326238 | 127326470 | Intron |
| chr6 | 127941110 | 127942468 | Distal Intergenic |
| chr6 | 127982277 | 127982508 | Distal Intergenic |
| chr6 | 127993194 | 127993589 | Distal Intergenic |
| chr6 | 127993783 | 127994176 | Distal Intergenic |
| chr6 | 127996320 | 127998663 | Distal Intergenic |
| chr6 | 128011928 | 128013199 | Distal Intergenic |
| chr6 | 128052467 | 128053003 | Intron |
| chr6 | 128054449 | 128055556 | Intron |
| chr6 | 128084956 | 128085179 | Intron |
| chr6 | 128085221 | 128086152 | Intron |
| chr6 | 128092627 | 128093102 | Intron |
| chr6 | 128096714 | 128096994 | Intron |
| chr6 | 128097304 | 128097555 | Intron |
| chr6 | 128232675 | 128232915 | Intron |

| | | | |
|---|---|---|---|
| chr6 | 128246313 | 128246778 | Distal Intergenic |
| chr6 | 128247594 | 128247869 | Distal Intergenic |
| chr6 | 128248289 | 128249463 | Distal Intergenic |
| chr6 | 128289019 | 128290388 | 3' UTR |
| chr6 | 128291810 | 128292362 | Intron |
| chr6 | 128292522 | 128292738 | Intron |
| chr6 | 128311976 | 128312542 | Exon |
| chr6 | 128313566 | 128313765 | Intron |
| chr6 | 128371223 | 128371438 | Intron |
| chr6 | 128371590 | 128371789 | Intron |
| chr6 | 128374997 | 128375196 | Intron |
| chr6 | 128794693 | 128794901 | Intron |
| chr6 | 128912952 | 128913211 | Distal Intergenic |
| chr6 | 128925028 | 128925431 | Distal Intergenic |
| chr6 | 128942796 | 128943026 | Distal Intergenic |
| chr6 | 128943225 | 128943424 | Distal Intergenic |
| chr6 | 128958103 | 128961958 | Distal Intergenic |
| chr6 | 128962483 | 128964524 | Distal Intergenic |
| chr6 | 129048084 | 129048283 | Distal Intergenic |
| chr6 | 129113686 | 129114341 | Distal Intergenic |
| chr6 | 129130496 | 129131450 | Distal Intergenic |
| chr6 | 129131714 | 129131999 | Distal Intergenic |
| chr6 | 129179634 | 129181052 | Distal Intergenic |
| chr6 | 129181256 | 129181490 | Distal Intergenic |
| chr6 | 129181713 | 129183988 | Distal Intergenic |
| chr6 | 129184733 | 129188052 | Distal Intergenic |
| chr6 | 129188120 | 129188371 | Distal Intergenic |
| chr6 | 130013521 | 130013758 | Intron |
| chr6 | 130013809 | 130014043 | Intron |
| chr6 | 131331917 | 131334281 | Intron |
| chr6 | 131337016 | 131337564 | Intron |
| chr6 | 133055379 | 133055706 | Promoter |
| chr6 | 133291401 | 133292412 | Distal Intergenic |
| chr6 | 133503353 | 133504186 | Distal Intergenic |
| chr6 | 133504387 | 133504647 | Distal Intergenic |
| chr6 | 133985061 | 133985577 | Intron |
| chr6 | 133986141 | 133986433 | Intron |
| chr6 | 133986489 | 133986786 | Intron |
| chr6 | 134032540 | 134032981 | Intron |
| chr6 | 134079564 | 134080359 | Intron |
| chr6 | 134517019 | 134517478 | Intron |
| chr6 | 134525381 | 134525606 | Intron |
| chr6 | 134525723 | 134526056 | Intron |
| chr6 | 134701104 | 134702275 | Distal Intergenic |

| | | | |
|---|---|---|---|
| chr6 | 134702389 | 134702607 | Distal Intergenic |
| chr6 | 134870523 | 134870780 | Intron |
| chr6 | 134871113 | 134871678 | Intron |
| chr6 | 134871878 | 134872674 | Intron |
| chr6 | 134872766 | 134875508 | Intron |
| chr6 | 134875759 | 134878592 | Intron |
| chr6 | 134879136 | 134879570 | Intron |
| chr6 | 134940340 | 134941494 | Distal Intergenic |
| chr6 | 134943245 | 134943456 | Distal Intergenic |
| chr6 | 134943804 | 134944125 | Exon |
| chr6 | 134944332 | 134944823 | Distal Intergenic |
| chr6 | 134948465 | 134950103 | Distal Intergenic |
| chr6 | 134950143 | 134951123 | Distal Intergenic |
| chr6 | 134951169 | 134951434 | Distal Intergenic |
| chr6 | 134961778 | 134962187 | Distal Intergenic |
| chr6 | 134974755 | 134975149 | Distal Intergenic |
| chr6 | 135010890 | 135011441 | Distal Intergenic |
| chr6 | 135019847 | 135020706 | Distal Intergenic |
| chr6 | 135200484 | 135200836 | Distal Intergenic |
| chr6 | 135201033 | 135202852 | Distal Intergenic |
| chr6 | 135215842 | 135216082 | Distal Intergenic |
| chr6 | 135216283 | 135216512 | Distal Intergenic |
| chr6 | 135245264 | 135245603 | Intron |
| chr6 | 135245672 | 135246018 | Intron |
| chr6 | 135293752 | 135294794 | Intron |
| chr6 | 135312523 | 135312903 | Intron |
| chr6 | 135312946 | 135314054 | Intron |
| chr6 | 135314138 | 135314469 | Intron |
| chr6 | 135337900 | 135339065 | Intron |
| chr6 | 135454046 | 135455195 | Distal Intergenic |
| chr6 | 136414567 | 136415048 | Intron |
| chr6 | 136415128 | 136415373 | Intron |
| chr6 | 136439176 | 136439443 | Intron |
| chr6 | 136440654 | 136440992 | Intron |
| chr6 | 136489650 | 136490667 | Intron |
| chr6 | 136752164 | 136753869 | Intron |
| chr6 | 136756879 | 136757207 | Intron |
| chr6 | 136757258 | 136758012 | Intron |
| chr6 | 136775631 | 136776775 | Intron |
| chr6 | 136776879 | 136777267 | Intron |
| chr6 | 136878648 | 136879933 | 3' UTR |
| chr6 | 136883234 | 136883474 | Intron |
| chr6 | 136883677 | 136883876 | Exon |
| chr6 | 136885490 | 136886235 | Intron |

| | | | |
|---|---|---|---|
| chr6 | 136886275 | 136887703 | Intron |
| chr6 | 136888260 | 136888577 | Intron |
| chr6 | 136979169 | 136979391 | Intron |
| chr6 | 136979594 | 136979972 | Intron |
| chr6 | 140496273 | 140496583 | Distal Intergenic |
| chr6 | 140496889 | 140497193 | Distal Intergenic |
| chr6 | 140563687 | 140563967 | Distal Intergenic |
| chr6 | 140568775 | 140571950 | Distal Intergenic |
| chr6 | 140572568 | 140572861 | Distal Intergenic |
| chr6 | 140572916 | 140573301 | Distal Intergenic |
| chr6 | 140573540 | 140573869 | Distal Intergenic |
| chr6 | 140574249 | 140574470 | Distal Intergenic |
| chr6 | 142287238 | 142288908 | Intron |
| chr6 | 142289065 | 142289343 | Intron |
| chr6 | 142314424 | 142315503 | Intron |
| chr6 | 142316061 | 142316401 | Intron |
| chr6 | 143116883 | 143117145 | Intron |
| chr6 | 143117282 | 143117552 | Intron |
| chr6 | 143117790 | 143118511 | Intron |
| chr6 | 144850687 | 144851256 | Intron |
| chr6 | 145159921 | 145160255 | Intron |
| chr6 | 145225088 | 145226032 | Distal Intergenic |
| chr6 | 145240321 | 145240550 | Distal Intergenic |
| chr6 | 145921602 | 145921944 | Distal Intergenic |
| chr6 | 145926536 | 145926822 | Distal Intergenic |
| chr6 | 145947258 | 145947493 | 3' UTR |
| chr6 | 146058978 | 146059833 | Intron |
| chr6 | 146069180 | 146069419 | Intron |
| chr6 | 146380595 | 146381398 | Intron |
| chr6 | 146564560 | 146564821 | Intron |
| chr6 | 146564963 | 146565498 | Intron |
| chr6 | 146591213 | 146591496 | Intron |
| chr6 | 146591884 | 146592123 | Intron |
| chr6 | 146741929 | 146742238 | Intron |
| chr6 | 146742294 | 146742774 | Intron |
| chr6 | 146745962 | 146746741 | Intron |
| chr6 | 146838358 | 146838672 | Distal Intergenic |
| chr6 | 146847853 | 146848328 | Distal Intergenic |
| chr6 | 146857041 | 146859814 | Distal Intergenic |
| chr6 | 146883609 | 146883808 | Distal Intergenic |
| chr6 | 146887033 | 146887561 | Distal Intergenic |
| chr6 | 146887680 | 146887973 | Distal Intergenic |
| chr6 | 146904045 | 146904298 | Distal Intergenic |
| chr6 | 146908270 | 146908719 | Distal Intergenic |

| | | | |
|---|---|---|---|
| chr6 | 146933414 | 146933626 | Intron |
| chr6 | 146958706 | 146959175 | Intron |
| chr6 | 147011495 | 147011805 | Intron |
| chr6 | 147021323 | 147022195 | 5' UTR |
| chr6 | 147065997 | 147066377 | Intron |
| chr6 | 147066488 | 147066758 | Intron |
| chr6 | 147147582 | 147149019 | Distal Intergenic |
| chr6 | 147209281 | 147212595 | Intron |
| chr6 | 147212738 | 147213182 | Intron |
| chr6 | 147213355 | 147216324 | Intron |
| chr6 | 147216555 | 147216801 | Intron |
| chr6 | 147217049 | 147217300 | Intron |
| chr6 | 147224640 | 147224909 | Intron |
| chr6 | 147225208 | 147226973 | Intron |
| chr6 | 147241170 | 147242341 | Intron |
| chr6 | 147249116 | 147250305 | Intron |
| chr6 | 147270709 | 147271622 | Intron |
| chr6 | 147272706 | 147273537 | Intron |
| chr6 | 147338210 | 147341269 | Intron |
| chr6 | 147346165 | 147347982 | Intron |
| chr6 | 147361023 | 147363191 | Intron |
| chr6 | 147366919 | 147367282 | Intron |
| chr6 | 147367333 | 147367789 | Intron |
| chr6 | 147374779 | 147375284 | Intron |
| chr6 | 147375404 | 147376139 | Intron |
| chr6 | 147482135 | 147482437 | Intron |
| chr6 | 147482643 | 147483236 | Intron |
| chr6 | 147689923 | 147690785 | Intron |
| chr6 | 147899907 | 147900106 | Distal Intergenic |
| chr6 | 147913302 | 147913649 | Distal Intergenic |
| chr6 | 147913694 | 147913903 | Distal Intergenic |
| chr6 | 148893370 | 148893741 | Distal Intergenic |
| chr6 | 149136471 | 149137177 | Intron |
| chr6 | 149141572 | 149143053 | Intron |
| chr6 | 149208662 | 149210521 | Intron |
| chr6 | 149258034 | 149258719 | Intron |
| chr6 | 149303661 | 149304901 | Intron |
| chr6 | 149316339 | 149316749 | Intron |
| chr7 | 3021789 | 3022179 | Intron |
| chr7 | 3028724 | 3029341 | Intron |
| chr7 | 3067392 | 3068166 | Intron |
| chr7 | 3135027 | 3135938 | Distal Intergenic |
| chr7 | 3149863 | 3151664 | Distal Intergenic |
| chr7 | 3167790 | 3168985 | Distal Intergenic |

| | | | |
|---|---|---|---|
| chr7 | 3169045 | 3169272 | Distal Intergenic |
| chr7 | 3275542 | 3275829 | Distal Intergenic |
| chr7 | 3276025 | 3276238 | Distal Intergenic |
| chr7 | 3289266 | 3293312 | Distal Intergenic |
| chr7 | 3451336 | 3452653 | Intron |
| chr7 | 3498966 | 3499341 | Intron |
| chr7 | 3542187 | 3542634 | Intron |
| chr7 | 3581138 | 3581497 | Intron |
| chr7 | 3616675 | 3617120 | Intron |
| chr7 | 3628360 | 3629191 | Intron |
| chr7 | 3644934 | 3645757 | Intron |
| chr7 | 3645818 | 3646019 | Intron |
| chr7 | 3646234 | 3646561 | Intron |
| chr7 | 3654900 | 3655104 | Intron |
| chr7 | 3655311 | 3655518 | Intron |
| chr7 | 3655596 | 3656046 | Intron |
| chr7 | 3656177 | 3656610 | Intron |
| chr7 | 3658490 | 3658910 | Exon |
| chr7 | 4050257 | 4051232 | Promoter |
| chr7 | 4085661 | 4086417 | Intron |
| chr7 | 4087835 | 4088918 | Intron |
| chr7 | 4101154 | 4101516 | Intron |
| chr7 | 4453412 | 4453765 | Distal Intergenic |
| chr7 | 4740229 | 4740428 | Intron |
| chr7 | 4743978 | 4744889 | Intron |
| chr7 | 4745065 | 4745280 | Intron |
| chr7 | 4764145 | 4765300 | Intron |
| chr7 | 4966464 | 4966908 | Intron |
| chr7 | 4971051 | 4972253 | Intron |
| chr7 | 4989808 | 4990055 | Intron |
| chr7 | 4990186 | 4990649 | Intron |
| chr7 | 5012413 | 5012680 | Promoter |
| chr7 | 5012880 | 5013453 | Promoter |
| chr7 | 5014151 | 5015287 | Promoter |
| chr7 | 5032096 | 5033118 | Intron |
| chr7 | 6106981 | 6107197 | Distal Intergenic |
| chr7 | 13490343 | 13490860 | Distal Intergenic |
| chr7 | 13506626 | 13507594 | Distal Intergenic |
| chr7 | 13507676 | 13508142 | Distal Intergenic |
| chr7 | 13508984 | 13509183 | Distal Intergenic |
| chr7 | 13563520 | 13563954 | Distal Intergenic |
| chr7 | 13595060 | 13595512 | Distal Intergenic |
| chr7 | 13618812 | 13620105 | Distal Intergenic |
| chr7 | 13622519 | 13623523 | Distal Intergenic |

| | | | |
|---|---|---|---|
| chr7 | 13623943 | 13624459 | Distal Intergenic |
| chr7 | 13724185 | 13725215 | Distal Intergenic |
| chr7 | 13863627 | 13864549 | Distal Intergenic |
| chr7 | 13876434 | 13877114 | Distal Intergenic |
| chr7 | 13915581 | 13916598 | Distal Intergenic |
| chr7 | 16528530 | 16531944 | Intron |
| chr7 | 16533248 | 16534937 | Intron |
| chr7 | 16540671 | 16541813 | Intron |
| chr7 | 16541856 | 16542196 | Intron |
| chr7 | 16548236 | 16548616 | Intron |
| chr7 | 16561508 | 16562993 | Intron |
| chr7 | 16569213 | 16569676 | Promoter |
| chr7 | 16613603 | 16614599 | Intron |
| chr7 | 16614667 | 16614866 | Intron |
| chr7 | 16622149 | 16622839 | Distal Intergenic |
| chr7 | 16634101 | 16634674 | Distal Intergenic |
| chr7 | 16637432 | 16637687 | Downstream |
| chr7 | 16637807 | 16638183 | Downstream |
| chr7 | 16642105 | 16642971 | Exon |
| chr7 | 16684151 | 16684384 | Intron |
| chr7 | 16893495 | 16894777 | Distal Intergenic |
| chr7 | 16951905 | 16952104 | Distal Intergenic |
| chr7 | 16952166 | 16952376 | Distal Intergenic |
| chr7 | 16952620 | 16955419 | Distal Intergenic |
| chr7 | 16955584 | 16955784 | Distal Intergenic |
| chr7 | 16984711 | 16984913 | Distal Intergenic |
| chr7 | 16984986 | 16987521 | Distal Intergenic |
| chr7 | 16987564 | 16989155 | Distal Intergenic |
| chr7 | 16989698 | 16992030 | Distal Intergenic |
| chr7 | 17109164 | 17111322 | Distal Intergenic |
| chr7 | 17163242 | 17164128 | Distal Intergenic |
| chr7 | 17167579 | 17167815 | Distal Intergenic |
| chr7 | 17182296 | 17182907 | Distal Intergenic |
| chr7 | 17186703 | 17187785 | Distal Intergenic |
| chr7 | 17199544 | 17199892 | Distal Intergenic |
| chr7 | 17439572 | 17439777 | Distal Intergenic |
| chr7 | 17439877 | 17440188 | Distal Intergenic |
| chr7 | 17590858 | 17591157 | Distal Intergenic |
| chr7 | 17602443 | 17603033 | Distal Intergenic |
| chr7 | 17610298 | 17611965 | Distal Intergenic |
| chr7 | 17612435 | 17612842 | Distal Intergenic |
| chr7 | 17612963 | 17613263 | Distal Intergenic |
| chr7 | 18772246 | 18772929 | Intron |
| chr7 | 18799726 | 18799975 | Intron |

| | | | |
|---|---|---|---|
| chr7 | 18800050 | 18800922 | Intron |
| chr7 | 18821585 | 18823215 | Intron |
| chr7 | 18823642 | 18824508 | Intron |
| chr7 | 19443729 | 19443954 | Distal Intergenic |
| chr7 | 19588448 | 19588897 | Distal Intergenic |
| chr7 | 19741405 | 19741736 | Intron |
| chr7 | 19812639 | 19813335 | Promoter |
| chr7 | 19814368 | 19814609 | Distal Intergenic |
| chr7 | 19855279 | 19858331 | Distal Intergenic |
| chr7 | 19929040 | 19929239 | Distal Intergenic |
| chr7 | 19929579 | 19930752 | Distal Intergenic |
| chr7 | 19934467 | 19934675 | Distal Intergenic |
| chr7 | 19944084 | 19944311 | Distal Intergenic |
| chr7 | 19944840 | 19945055 | Distal Intergenic |
| chr7 | 19945721 | 19946510 | Distal Intergenic |
| chr7 | 19946618 | 19947250 | Distal Intergenic |
| chr7 | 19947303 | 19951694 | Distal Intergenic |
| chr7 | 20043418 | 20045159 | Distal Intergenic |
| chr7 | 20062552 | 20063901 | Distal Intergenic |
| chr7 | 20064095 | 20064414 | Distal Intergenic |
| chr7 | 20093754 | 20093969 | Distal Intergenic |
| chr7 | 20094068 | 20094334 | Distal Intergenic |
| chr7 | 20094419 | 20094817 | Distal Intergenic |
| chr7 | 20146592 | 20147166 | Distal Intergenic |
| chr7 | 20147581 | 20147782 | Distal Intergenic |
| chr7 | 20157885 | 20158686 | Distal Intergenic |
| chr7 | 20158753 | 20159379 | Distal Intergenic |
| chr7 | 20163502 | 20163777 | Distal Intergenic |
| chr7 | 20163832 | 20164536 | Distal Intergenic |
| chr7 | 20164626 | 20164825 | Distal Intergenic |
| chr7 | 20165676 | 20167051 | Distal Intergenic |
| chr7 | 20194769 | 20195367 | Intron |
| chr7 | 20250548 | 20251227 | Intron |
| chr7 | 20289472 | 20289959 | Distal Intergenic |
| chr7 | 20354342 | 20355157 | Distal Intergenic |
| chr7 | 20355209 | 20355610 | Distal Intergenic |
| chr7 | 20382256 | 20382647 | Intron |
| chr7 | 20402082 | 20403883 | 5' UTR |
| chr7 | 20408919 | 20409175 | Intron |
| chr7 | 20409264 | 20409658 | Intron |
| chr7 | 20409805 | 20410223 | Intron |
| chr7 | 20416099 | 20417913 | Exon |
| chr7 | 20462001 | 20462206 | Distal Intergenic |
| chr7 | 24867229 | 24868070 | Intron |

| | | | |
|---|---|---|---|
| chr7 | 24868129 | 24868540 | Intron |
| chr7 | 24897459 | 24898028 | Intron |
| chr7 | 25150436 | 25150852 | Distal Intergenic |
| chr7 | 25171209 | 25172759 | Downstream |
| chr7 | 25179464 | 25180577 | Intron |
| chr7 | 25234259 | 25234516 | Distal Intergenic |
| chr7 | 25244673 | 25245462 | Distal Intergenic |
| chr7 | 25246250 | 25247291 | Distal Intergenic |
| chr7 | 25247505 | 25247963 | Distal Intergenic |
| chr7 | 25255504 | 25256133 | Distal Intergenic |
| chr7 | 25256275 | 25256968 | Distal Intergenic |
| chr7 | 25257030 | 25258138 | Distal Intergenic |
| chr7 | 25265225 | 25266506 | Exon |
| chr7 | 25291800 | 25292241 | Distal Intergenic |
| chr7 | 25315639 | 25315838 | Distal Intergenic |
| chr7 | 25316019 | 25316607 | Distal Intergenic |
| chr7 | 25384306 | 25385226 | Distal Intergenic |
| chr7 | 25386731 | 25387177 | Distal Intergenic |
| chr7 | 25418474 | 25418686 | Distal Intergenic |
| chr7 | 25421285 | 25422329 | Distal Intergenic |
| chr7 | 25422552 | 25423006 | Distal Intergenic |
| chr7 | 25670041 | 25670347 | Intron |
| chr7 | 25670535 | 25671144 | Intron |
| chr7 | 25689308 | 25689878 | Intron |
| chr7 | 25691929 | 25693159 | Intron |
| chr7 | 25988861 | 25992066 | Promoter |
| chr7 | 25993474 | 25995357 | Distal Intergenic |
| chr7 | 26005025 | 26006134 | Distal Intergenic |
| chr7 | 26006480 | 26006796 | Distal Intergenic |
| chr7 | 26033438 | 26033678 | Distal Intergenic |
| chr7 | 26033980 | 26034493 | Distal Intergenic |
| chr7 | 26034637 | 26035369 | Distal Intergenic |
| chr7 | 26053900 | 26055498 | Distal Intergenic |
| chr7 | 26058945 | 26063798 | Distal Intergenic |
| chr7 | 26064070 | 26064435 | Distal Intergenic |
| chr7 | 26064593 | 26066467 | Distal Intergenic |
| chr7 | 26066585 | 26066919 | Distal Intergenic |
| chr7 | 26082715 | 26083045 | Distal Intergenic |
| chr7 | 26178473 | 26179817 | Distal Intergenic |
| chr7 | 26179865 | 26180201 | Distal Intergenic |
| chr7 | 26429425 | 26430524 | Distal Intergenic |
| chr7 | 26472253 | 26474561 | Intron |
| chr7 | 26490569 | 26491516 | Intron |
| chr7 | 26492625 | 26494452 | Intron |

| | | | |
|---|---|---|---|
| chr7 | 26495506 | 26497071 | Intron |
| chr7 | 26497107 | 26497483 | Intron |
| chr7 | 26497686 | 26498023 | Intron |
| chr7 | 26503552 | 26504338 | Intron |
| chr7 | 26538916 | 26539129 | Distal Intergenic |
| chr7 | 26539392 | 26539821 | Distal Intergenic |
| chr7 | 26539865 | 26540253 | Distal Intergenic |
| chr7 | 26565492 | 26567314 | Distal Intergenic |
| chr7 | 26568009 | 26568304 | Distal Intergenic |
| chr7 | 26568357 | 26568568 | Distal Intergenic |
| chr7 | 26569031 | 26569503 | Distal Intergenic |
| chr7 | 26573357 | 26574066 | Exon |
| chr7 | 26616194 | 26617736 | Distal Intergenic |
| chr7 | 26621986 | 26622478 | Distal Intergenic |
| chr7 | 26628497 | 26630288 | Distal Intergenic |
| chr7 | 26687178 | 26687719 | Distal Intergenic |
| chr7 | 26981121 | 26982380 | Intron |
| chr7 | 27776484 | 27777013 | Distal Intergenic |
| chr7 | 27979018 | 27979267 | Intron |
| chr7 | 27980201 | 27980999 | Intron |
| chr7 | 28006961 | 28007297 | Intron |
| chr7 | 28010159 | 28010994 | Intron |
| chr7 | 28012935 | 28013246 | Intron |
| chr7 | 28013336 | 28013616 | Intron |
| chr7 | 28016402 | 28018233 | Intron |
| chr7 | 28076349 | 28076850 | Intron |
| chr7 | 28116043 | 28117545 | Intron |
| chr7 | 28117584 | 28118120 | Intron |
| chr7 | 28140557 | 28140793 | Intron |
| chr7 | 28272026 | 28272920 | Intron |
| chr7 | 28273007 | 28273235 | Intron |
| chr7 | 28273289 | 28273552 | Intron |
| chr7 | 28275491 | 28276650 | Intron |
| chr7 | 28292816 | 28293113 | Distal Intergenic |
| chr7 | 28338302 | 28338857 | Promoter |
| chr7 | 28354106 | 28355553 | Intron |
| chr7 | 28359068 | 28359674 | Intron |
| chr7 | 28376160 | 28376359 | Intron |
| chr7 | 28376820 | 28377475 | Intron |
| chr7 | 28392858 | 28393813 | Intron |
| chr7 | 28516450 | 28516777 | Intron |
| chr7 | 28589586 | 28589811 | Intron |
| chr7 | 28763869 | 28764077 | 3' UTR |
| chr7 | 28946655 | 28947797 | Distal Intergenic |

| | | | |
|---|---|---|---|
| chr7 | 28953107 | 28953430 | Distal Intergenic |
| chr7 | 28954103 | 28954495 | Distal Intergenic |
| chr7 | 29002388 | 29002602 | Distal Intergenic |
| chr7 | 29119479 | 29120350 | Intron |
| chr7 | 29123889 | 29124121 | Intron |
| chr7 | 29135860 | 29136059 | Intron |
| chr7 | 29136256 | 29136693 | Intron |
| chr7 | 29136850 | 29137049 | Intron |
| chr7 | 29156066 | 29156510 | Intron |
| chr7 | 29157086 | 29157649 | Intron |
| chr7 | 29160142 | 29162533 | 5' UTR |
| chr7 | 29162583 | 29164962 | Intron |
| chr7 | 29165292 | 29167195 | Intron |
| chr7 | 29167532 | 29168157 | Intron |
| chr7 | 29203071 | 29205381 | Intron |
| chr7 | 29338087 | 29338286 | Intron |
| chr7 | 29355108 | 29355413 | Intron |
| chr7 | 29355548 | 29355857 | Intron |
| chr7 | 29361587 | 29363307 | Intron |
| chr7 | 29404310 | 29404661 | Intron |
| chr7 | 29586381 | 29588298 | Intron |
| chr7 | 29593061 | 29593291 | Intron |
| chr7 | 29593511 | 29595218 | Intron |
| chr7 | 29634446 | 29635758 | Distal Intergenic |
| chr7 | 29650889 | 29651579 | Distal Intergenic |
| chr7 | 29651772 | 29652242 | Distal Intergenic |
| chr7 | 29652382 | 29652841 | Distal Intergenic |
| chr7 | 29704225 | 29706309 | Intron |
| chr7 | 29706510 | 29706851 | Intron |
| chr7 | 29708799 | 29709520 | Intron |
| chr7 | 29709704 | 29711578 | Intron |
| chr7 | 29713902 | 29717308 | Intron |
| chr7 | 29717442 | 29718058 | Intron |
| chr7 | 29723638 | 29724083 | Promoter |
| chr7 | 29724472 | 29725499 | Promoter |
| chr7 | 29725924 | 29726338 | Promoter |
| chr7 | 29737844 | 29739269 | Intron |
| chr7 | 29742938 | 29743204 | Intron |
| chr7 | 29743272 | 29743842 | Intron |
| chr7 | 29744009 | 29745010 | Intron |
| chr7 | 29745545 | 29745766 | Intron |
| chr7 | 29745916 | 29746534 | Intron |
| chr7 | 29746702 | 29747138 | Intron |
| chr7 | 29747387 | 29748089 | Intron |

| | | | |
|---|---|---|---|
| chr7 | 29754808 | 29757613 | Distal Intergenic |
| chr7 | 29759634 | 29760205 | Intron |
| chr7 | 29767062 | 29767294 | Intron |
| chr7 | 29809941 | 29812340 | Distal Intergenic |
| chr7 | 29964912 | 29965345 | Intron |
| chr7 | 30022797 | 30023301 | Intron |
| chr7 | 30032765 | 30034501 | Distal Intergenic |
| chr7 | 30036168 | 30036617 | Distal Intergenic |
| chr7 | 30036722 | 30037725 | Distal Intergenic |
| chr7 | 30041738 | 30043124 | Distal Intergenic |
| chr7 | 30060802 | 30061207 | Intron |
| chr7 | 30061378 | 30062279 | 3' UTR |
| chr7 | 30126489 | 30128666 | Intron |
| chr7 | 30134350 | 30134578 | Intron |
| chr7 | 30134671 | 30135684 | Intron |
| chr7 | 30135724 | 30138429 | Intron |
| chr7 | 30227195 | 30228512 | Distal Intergenic |
| chr7 | 30229112 | 30230644 | Distal Intergenic |
| chr7 | 30235052 | 30235305 | Distal Intergenic |
| chr7 | 30237942 | 30238343 | Distal Intergenic |
| chr7 | 30239578 | 30239777 | Distal Intergenic |
| chr7 | 30239832 | 30240127 | Distal Intergenic |
| chr7 | 30243061 | 30243648 | Distal Intergenic |
| chr7 | 30247203 | 30247513 | Distal Intergenic |
| chr7 | 30247749 | 30247948 | Distal Intergenic |
| chr7 | 30256088 | 30256483 | Distal Intergenic |
| chr7 | 30256678 | 30257141 | Distal Intergenic |
| chr7 | 30261909 | 30262271 | Distal Intergenic |
| chr7 | 30262319 | 30262722 | Distal Intergenic |
| chr7 | 30265899 | 30267039 | Distal Intergenic |
| chr7 | 30269119 | 30270009 | Distal Intergenic |
| chr7 | 30270121 | 30270520 | Distal Intergenic |
| chr7 | 30290202 | 30291262 | Distal Intergenic |
| chr7 | 30298114 | 30298332 | Distal Intergenic |
| chr7 | 30552970 | 30553200 | Downstream |
| chr7 | 30553346 | 30553664 | Downstream |
| chr7 | 30738523 | 30738808 | Promoter |
| chr7 | 30738906 | 30739177 | Promoter |
| chr7 | 30767219 | 30767717 | Intron |
| chr7 | 30768020 | 30768780 | Intron |
| chr7 | 30879932 | 30880334 | Intron |
| chr7 | 30954369 | 30955093 | Intron |
| chr7 | 30979152 | 30980165 | Distal Intergenic |
| chr7 | 30980299 | 30981183 | Distal Intergenic |

| | | | |
|---|---|---|---|
| chr7 | 30981380 | 30981923 | Distal Intergenic |
| chr7 | 31336757 | 31337337 | Distal Intergenic |
| chr7 | 31337893 | 31338383 | Distal Intergenic |
| chr7 | 31344830 | 31345272 | Distal Intergenic |
| chr7 | 31347450 | 31348029 | Distal Intergenic |
| chr7 | 31348196 | 31348623 | Distal Intergenic |
| chr7 | 31372891 | 31373241 | Distal Intergenic |
| chr7 | 31514822 | 31515062 | Distal Intergenic |
| chr7 | 31533018 | 31533236 | Distal Intergenic |
| chr7 | 31536475 | 31536678 | Distal Intergenic |
| chr7 | 31543997 | 31544517 | Distal Intergenic |
| chr7 | 31544598 | 31545190 | Distal Intergenic |
| chr7 | 31547916 | 31549142 | Distal Intergenic |
| chr7 | 31560842 | 31562104 | Intron |
| chr7 | 31579399 | 31580283 | Intron |
| chr7 | 31580392 | 31580837 | Intron |
| chr7 | 31580905 | 31582375 | Intron |
| chr7 | 31589413 | 31589818 | Intron |
| chr7 | 31601033 | 31602687 | Intron |
| chr7 | 31859086 | 31859461 | Intron |
| chr7 | 31860618 | 31861784 | Intron |
| chr7 | 34915678 | 34917339 | Intron |
| chr7 | 34951767 | 34952015 | Distal Intergenic |
| chr7 | 34952950 | 34953519 | Distal Intergenic |
| chr7 | 34953625 | 34955159 | Distal Intergenic |
| chr7 | 34998015 | 34999096 | Intron |
| chr7 | 35018498 | 35019079 | Intron |
| chr7 | 35051655 | 35052177 | Intron |
| chr7 | 35052463 | 35054673 | Exon |
| chr7 | 35127137 | 35128094 | Intron |
| chr7 | 35163490 | 35164020 | Exon |
| chr7 | 35172902 | 35174586 | Intron |
| chr7 | 35174901 | 35175375 | Intron |
| chr7 | 35414165 | 35415689 | Exon |
| chr7 | 35438829 | 35439028 | Distal Intergenic |
| chr7 | 35439819 | 35440131 | Distal Intergenic |
| chr7 | 35590491 | 35591803 | Distal Intergenic |
| chr7 | 35840818 | 35841201 | Promoter |
| chr7 | 35861307 | 35861773 | Intron |
| chr7 | 35861842 | 35862377 | Intron |
| chr7 | 35869966 | 35870581 | Promoter |
| chr7 | 35912174 | 35912561 | Exon |
| chr7 | 35912635 | 35912916 | Intron |
| chr7 | 36134057 | 36134256 | Distal Intergenic |

| | | | |
|---|---|---|---|
| chr7 | 36180925 | 36181630 | Distal Intergenic |
| chr7 | 38470767 | 38471046 | Intron |
| chr7 | 38480324 | 38480736 | Intron |
| chr7 | 38862367 | 38862577 | Intron |
| chr7 | 38877213 | 38877877 | Intron |
| chr7 | 38969266 | 38969473 | Distal Intergenic |
| chr7 | 38969921 | 38970120 | Distal Intergenic |
| chr7 | 39055969 | 39056175 | Intron |
| chr7 | 46772895 | 46773188 | Distal Intergenic |
| chr7 | 46808101 | 46808365 | Distal Intergenic |
| chr7 | 47121625 | 47123366 | Distal Intergenic |
| chr7 | 47123580 | 47124144 | Distal Intergenic |
| chr7 | 47127903 | 47128748 | Distal Intergenic |
| chr7 | 48341203 | 48341426 | Intron |
| chr7 | 48648959 | 48649177 | Intron |
| chr7 | 50654212 | 50654940 | Downstream |
| chr7 | 50927040 | 50927264 | Distal Intergenic |
| chr7 | 50927374 | 50927747 | Distal Intergenic |
| chr7 | 51501283 | 51501515 | Distal Intergenic |
| chr7 | 51501566 | 51502331 | Distal Intergenic |
| chr7 | 51546363 | 51546860 | Distal Intergenic |
| chr7 | 51660459 | 51660737 | Distal Intergenic |
| chr7 | 51792152 | 51792372 | Distal Intergenic |
| chr7 | 51869967 | 51870180 | Distal Intergenic |
| chr7 | 51917334 | 51918838 | Distal Intergenic |
| chr7 | 52003524 | 52003767 | Distal Intergenic |
| chr7 | 52032235 | 52032811 | Distal Intergenic |
| chr7 | 52071776 | 52071994 | Distal Intergenic |
| chr7 | 52072058 | 52072668 | Distal Intergenic |
| chr7 | 52081949 | 52082226 | Distal Intergenic |
| chr7 | 52104660 | 52105251 | Distal Intergenic |
| chr7 | 52105334 | 52105613 | Distal Intergenic |
| chr7 | 52239526 | 52240164 | Distal Intergenic |
| chr7 | 52242216 | 52243009 | Distal Intergenic |
| chr7 | 52253065 | 52254034 | Distal Intergenic |
| chr7 | 52270876 | 52271132 | Distal Intergenic |
| chr7 | 52271178 | 52271638 | Distal Intergenic |
| chr7 | 52272142 | 52274756 | Distal Intergenic |
| chr7 | 52280707 | 52281460 | Distal Intergenic |
| chr7 | 52306534 | 52306746 | Distal Intergenic |
| chr7 | 52307095 | 52307430 | Distal Intergenic |
| chr7 | 52316884 | 52317772 | Distal Intergenic |
| chr7 | 52379330 | 52379552 | Distal Intergenic |
| chr7 | 52379623 | 52379827 | Distal Intergenic |

| | | | |
|---|---|---|---|
| chr7 | 52380126 | 52380615 | Distal Intergenic |
| chr7 | 52380708 | 52380927 | Distal Intergenic |
| chr7 | 52381620 | 52382666 | Distal Intergenic |
| chr7 | 52382747 | 52384187 | Distal Intergenic |
| chr7 | 52395122 | 52395524 | Distal Intergenic |
| chr7 | 52395724 | 52396607 | Distal Intergenic |
| chr7 | 52408820 | 52409667 | Distal Intergenic |
| chr7 | 52409737 | 52409936 | Distal Intergenic |
| chr7 | 52650336 | 52651030 | Distal Intergenic |
| chr7 | 52668926 | 52669145 | Distal Intergenic |
| chr7 | 52669210 | 52669681 | Distal Intergenic |
| chr7 | 52715154 | 52716787 | Distal Intergenic |
| chr7 | 52717510 | 52717723 | Distal Intergenic |
| chr7 | 52747593 | 52748199 | Distal Intergenic |
| chr7 | 52780779 | 52781281 | Distal Intergenic |
| chr7 | 52781692 | 52782238 | Distal Intergenic |
| chr7 | 52860985 | 52861719 | Distal Intergenic |
| chr7 | 52961424 | 52961888 | Distal Intergenic |
| chr7 | 52962594 | 52962845 | Distal Intergenic |
| chr7 | 52963105 | 52963387 | Distal Intergenic |
| chr7 | 52969526 | 52970480 | Distal Intergenic |
| chr7 | 52970803 | 52971025 | Distal Intergenic |
| chr7 | 52972470 | 52972729 | Distal Intergenic |
| chr7 | 52972781 | 52973945 | Distal Intergenic |
| chr7 | 52974103 | 52974465 | Distal Intergenic |
| chr7 | 53026150 | 53026717 | Distal Intergenic |
| chr7 | 53055599 | 53055963 | Distal Intergenic |
| chr7 | 53057507 | 53058921 | Distal Intergenic |
| chr7 | 53060003 | 53060614 | Distal Intergenic |
| chr7 | 53069451 | 53069674 | Distal Intergenic |
| chr7 | 53128191 | 53128445 | Distal Intergenic |
| chr7 | 53128643 | 53128983 | Distal Intergenic |
| chr7 | 53129184 | 53129573 | Distal Intergenic |
| chr7 | 53176167 | 53176815 | Distal Intergenic |
| chr7 | 54016451 | 54016899 | Distal Intergenic |
| chr7 | 54051612 | 54051904 | Distal Intergenic |
| chr7 | 54069703 | 54070217 | Distal Intergenic |
| chr7 | 54094004 | 54095303 | Distal Intergenic |
| chr7 | 54100261 | 54100627 | Distal Intergenic |
| chr7 | 54100708 | 54101133 | Distal Intergenic |
| chr7 | 54101175 | 54101388 | Distal Intergenic |
| chr7 | 54101569 | 54102414 | Distal Intergenic |
| chr7 | 54173965 | 54175316 | Distal Intergenic |
| chr7 | 54213279 | 54213802 | Distal Intergenic |

| | | | |
|---|---|---|---|
| chr7 | 54263411 | 54263782 | Distal Intergenic |
| chr7 | 54278007 | 54278211 | Distal Intergenic |
| chr7 | 54286317 | 54286839 | Distal Intergenic |
| chr7 | 54286922 | 54287253 | Distal Intergenic |
| chr7 | 54353715 | 54353933 | Distal Intergenic |
| chr7 | 54366033 | 54366383 | Distal Intergenic |
| chr7 | 55704439 | 55705043 | Distal Intergenic |
| chr7 | 55913064 | 55913452 | Intron |
| chr7 | 56017897 | 56018269 | Distal Intergenic |
| chr7 | 56088316 | 56088852 | Exon |
| chr7 | 56115812 | 56116826 | Intron |
| chr7 | 56143422 | 56144523 | Intron |
| chr7 | 56172669 | 56173386 | Promoter |
| chr7 | 56174572 | 56174772 | Promoter |
| chr7 | 56349265 | 56349860 | Distal Intergenic |
| chr7 | 56675752 | 56676323 | Distal Intergenic |
| chr7 | 56775663 | 56777197 | Distal Intergenic |
| chr7 | 56777258 | 56779474 | Distal Intergenic |
| chr7 | 56808282 | 56810491 | Distal Intergenic |
| chr7 | 56810798 | 56811598 | Distal Intergenic |
| chr7 | 56812207 | 56814502 | Distal Intergenic |
| chr7 | 56814568 | 56816509 | Distal Intergenic |
| chr7 | 56817224 | 56817464 | Distal Intergenic |
| chr7 | 56817623 | 56818193 | Distal Intergenic |
| chr7 | 56841285 | 56841495 | Distal Intergenic |
| chr7 | 56841702 | 56842291 | Distal Intergenic |
| chr7 | 56855763 | 56858962 | Distal Intergenic |
| chr7 | 57014725 | 57015103 | Distal Intergenic |
| chr7 | 57015283 | 57015511 | Distal Intergenic |
| chr7 | 57016102 | 57018950 | Distal Intergenic |
| chr7 | 57063957 | 57065116 | Distal Intergenic |
| chr7 | 57137561 | 57137933 | Distal Intergenic |
| chr7 | 57143078 | 57143766 | Distal Intergenic |
| chr7 | 59001217 | 59001557 | Distal Intergenic |
| chr7 | 59169130 | 59169647 | Distal Intergenic |
| chr7 | 59222921 | 59223446 | Distal Intergenic |
| chr7 | 59593027 | 59593655 | Distal Intergenic |
| chr7 | 60483324 | 60484047 | Distal Intergenic |
| chr7 | 61106225 | 61106481 | Distal Intergenic |
| chr7 | 61106578 | 61107610 | Distal Intergenic |
| chr7 | 62103424 | 62103789 | Distal Intergenic |
| chr7 | 62791827 | 62792261 | Distal Intergenic |
| chr7 | 62794910 | 62796126 | Distal Intergenic |
| chr7 | 63097097 | 63097496 | Distal Intergenic |

| | | | |
|---|---|---|---|
| chr7 | 63097666 | 63098779 | Distal Intergenic |
| chr7 | 63114195 | 63114482 | Distal Intergenic |
| chr7 | 63195045 | 63195261 | Distal Intergenic |
| chr7 | 63217930 | 63218252 | Distal Intergenic |
| chr7 | 63305792 | 63306009 | Distal Intergenic |
| chr7 | 65061237 | 65062989 | Distal Intergenic |
| chr7 | 66005230 | 66005708 | Intron |
| chr7 | 66055628 | 66055835 | Intron |
| chr7 | 66055883 | 66057673 | Promoter |
| chr7 | 66077390 | 66078386 | Distal Intergenic |
| chr7 | 66483382 | 66483610 | Intron |
| chr7 | 66483793 | 66483997 | Intron |
| chr7 | 66484160 | 66484422 | Intron |
| chr7 | 66484730 | 66485392 | Intron |
| chr7 | 68659275 | 68660414 | Distal Intergenic |
| chr7 | 69021687 | 69022095 | Distal Intergenic |
| chr7 | 69022215 | 69022634 | Distal Intergenic |
| chr7 | 70627356 | 70628646 | Intron |
| chr7 | 70641976 | 70642672 | Intron |
| chr7 | 71001515 | 71001714 | Intron |
| chr7 | 71084897 | 71085096 | Intron |
| chr7 | 71195494 | 71196695 | Distal Intergenic |
| chr7 | 71196815 | 71197970 | Distal Intergenic |
| chr7 | 71209468 | 71210880 | Distal Intergenic |
| chr7 | 71384504 | 71384703 | Intron |
| chr7 | 71394558 | 71394922 | Intron |
| chr7 | 71399692 | 71400447 | Intron |
| chr7 | 71410406 | 71411394 | Intron |
| chr7 | 71421528 | 71422658 | Intron |
| chr7 | 71505441 | 71505672 | Intron |
| chr7 | 71536904 | 71537180 | Intron |
| chr7 | 72051307 | 72052122 | Intron |
| chr7 | 72612883 | 72613766 | Exon |
| chr7 | 72614844 | 72615423 | Intron |
| chr7 | 73223958 | 73224327 | Distal Intergenic |
| chr7 | 73322656 | 73323755 | Distal Intergenic |
| chr7 | 73324058 | 73324349 | Distal Intergenic |
| chr7 | 73345261 | 73346857 | Distal Intergenic |
| chr7 | 73362953 | 73363313 | Distal Intergenic |
| chr7 | 73369349 | 73370469 | Distal Intergenic |
| chr7 | 73510230 | 73511540 | Promoter |
| chr7 | 73513921 | 73514762 | Intron |
| chr7 | 73515113 | 73515480 | Intron |
| chr7 | 73520275 | 73522443 | Exon |

| | | | |
|---|---|---|---|
| chr7 | 73742753 | 73743224 | Intron |
| chr7 | 73887831 | 73889250 | Intron |
| chr7 | 73891566 | 73891929 | Intron |
| chr7 | 73900317 | 73900632 | Intron |
| chr7 | 73900977 | 73901473 | Intron |
| chr7 | 73901603 | 73902083 | Intron |
| chr7 | 73902200 | 73902515 | Intron |
| chr7 | 73971283 | 73971762 | Intron |
| chr7 | 73971925 | 73973033 | Exon |
| chr7 | 74428412 | 74429166 | Intron |
| chr7 | 74431937 | 74432173 | Exon |
| chr7 | 74432293 | 74433203 | Exon |
| chr7 | 74586959 | 74588055 | Promoter |
| chr7 | 74637135 | 74638665 | Intron |
| chr7 | 74641118 | 74641511 | Intron |
| chr7 | 74649902 | 74650107 | Intron |
| chr7 | 74693309 | 74694218 | Intron |
| chr7 | 74790299 | 74790952 | Intron |
| chr7 | 74791851 | 74792058 | Intron |
| chr7 | 74792301 | 74793192 | Intron |
| chr7 | 74808507 | 74809869 | Intron |
| chr7 | 74981138 | 74982135 | Promoter |
| chr7 | 74982222 | 74982640 | Promoter |
| chr7 | 74988775 | 74989340 | Promoter |
| chr7 | 74998864 | 74999159 | Intron |
| chr7 | 75097433 | 75097734 | Intron |
| chr7 | 75097865 | 75098651 | Intron |
| chr7 | 75099265 | 75099929 | Intron |
| chr7 | 75100298 | 75100526 | Intron |
| chr7 | 75104154 | 75104353 | 5' UTR |
| chr7 | 75104400 | 75105818 | 5' UTR |
| chr7 | 75112476 | 75114913 | Promoter |
| chr7 | 75150971 | 75151836 | Intron |
| chr7 | 75165136 | 75167715 | 3' UTR |
| chr7 | 75278767 | 75279018 | Intron |
| chr7 | 75279103 | 75279488 | Intron |
| chr7 | 75279621 | 75280702 | Intron |
| chr7 | 77160606 | 77162202 | Distal Intergenic |
| chr7 | 77181073 | 77183062 | Intron |
| chr7 | 77288955 | 77291056 | Intron |
| chr7 | 77411274 | 77411525 | Distal Intergenic |
| chr7 | 77411667 | 77411994 | Distal Intergenic |
| chr7 | 77412243 | 77412756 | Distal Intergenic |
| chr7 | 77448729 | 77450253 | Intron |

| | | | |
|---|---|---|---|
| chr7 | 77503747 | 77504235 | Intron |
| chr7 | 77504300 | 77504644 | Intron |
| chr7 | 77504822 | 77505027 | Intron |
| chr7 | 77505296 | 77505782 | Intron |
| chr7 | 77718750 | 77719688 | Intron |
| chr7 | 79262452 | 79262667 | Distal Intergenic |
| chr7 | 79262735 | 79262963 | Distal Intergenic |
| chr7 | 79443173 | 79444948 | Distal Intergenic |
| chr7 | 79470470 | 79471957 | Distal Intergenic |
| chr7 | 79589143 | 79589541 | Distal Intergenic |
| chr7 | 79589609 | 79589845 | Distal Intergenic |
| chr7 | 79589904 | 79590136 | Distal Intergenic |
| chr7 | 79591180 | 79593085 | Distal Intergenic |
| chr7 | 79593234 | 79593433 | Distal Intergenic |
| chr7 | 79637484 | 79639309 | Distal Intergenic |
| chr7 | 79640657 | 79643299 | Distal Intergenic |
| chr7 | 79643438 | 79644529 | Distal Intergenic |
| chr7 | 79644606 | 79645034 | Distal Intergenic |
| chr7 | 79654654 | 79655394 | Distal Intergenic |
| chr7 | 79664499 | 79664733 | Distal Intergenic |
| chr7 | 79735391 | 79735771 | Distal Intergenic |
| chr7 | 79835708 | 79838363 | Intron |
| chr7 | 79840551 | 79840793 | Intron |
| chr7 | 79840846 | 79841354 | Intron |
| chr7 | 80165003 | 80165577 | Intron |
| chr7 | 80173611 | 80174937 | Intron |
| chr7 | 80191954 | 80192610 | Intron |
| chr7 | 80193023 | 80193249 | Intron |
| chr7 | 80378243 | 80378532 | Exon |
| chr7 | 80678122 | 80679579 | Distal Intergenic |
| chr7 | 80680095 | 80680563 | Distal Intergenic |
| chr7 | 80681470 | 80682828 | Distal Intergenic |
| chr7 | 80683001 | 80683716 | Distal Intergenic |
| chr7 | 80683768 | 80684411 | Distal Intergenic |
| chr7 | 80684473 | 80687280 | Distal Intergenic |
| chr7 | 80696496 | 80698500 | Distal Intergenic |
| chr7 | 80698536 | 80698806 | Distal Intergenic |
| chr7 | 80698895 | 80699426 | Distal Intergenic |
| chr7 | 80699578 | 80703171 | Distal Intergenic |
| chr7 | 80703672 | 80704386 | Distal Intergenic |
| chr7 | 80731417 | 80731745 | Distal Intergenic |
| chr7 | 80732191 | 80732392 | Distal Intergenic |
| chr7 | 80732641 | 80735085 | Distal Intergenic |
| chr7 | 80751608 | 80753455 | Distal Intergenic |

| | | | |
|---|---|---|---|
| chr7 | 80760610 | 80763205 | Distal Intergenic |
| chr7 | 80763522 | 80764445 | Distal Intergenic |
| chr7 | 80768861 | 80769221 | Distal Intergenic |
| chr7 | 81317667 | 81317877 | Intron |
| chr7 | 81521496 | 81521718 | Distal Intergenic |
| chr7 | 81609554 | 81610498 | Intron |
| chr7 | 81678402 | 81678601 | Intron |
| chr7 | 82069013 | 82069798 | Intron |
| chr7 | 82538143 | 82538877 | Exon |
| chr7 | 82539619 | 82540342 | Intron |
| chr7 | 82542455 | 82543295 | Intron |
| chr7 | 82543893 | 82544220 | Exon |
| chr7 | 82556069 | 82556887 | Intron |
| chr7 | 82571609 | 82572613 | Intron |
| chr7 | 82786252 | 82789904 | Intron |
| chr7 | 82793194 | 82795562 | Promoter |
| chr7 | 82799954 | 82801212 | Distal Intergenic |
| chr7 | 82811093 | 82812585 | Distal Intergenic |
| chr7 | 82823911 | 82828949 | Distal Intergenic |
| chr7 | 82847829 | 82848231 | Distal Intergenic |
| chr7 | 85927989 | 85928223 | Distal Intergenic |
| chr7 | 85969688 | 85970562 | Distal Intergenic |
| chr7 | 86040787 | 86042080 | Distal Intergenic |
| chr7 | 86058257 | 86058480 | Distal Intergenic |
| chr7 | 86058838 | 86059800 | Distal Intergenic |
| chr7 | 86285335 | 86285709 | Intron |
| chr7 | 86293279 | 86293571 | Intron |
| chr7 | 86294281 | 86294656 | Intron |
| chr7 | 86355022 | 86357217 | Intron |
| chr7 | 86414442 | 86415390 | Intron |
| chr7 | 86415663 | 86416183 | Exon |
| chr7 | 86417627 | 86417826 | Intron |
| chr7 | 86430058 | 86430446 | Intron |
| chr7 | 86433246 | 86433445 | Intron |
| chr7 | 86481248 | 86481508 | Intron |
| chr7 | 87004902 | 87006137 | Exon |
| chr7 | 87025894 | 87026646 | Intron |
| chr7 | 87026806 | 87027005 | Intron |
| chr7 | 87174423 | 87175122 | Intron |
| chr7 | 87204266 | 87204609 | Intron |
| chr7 | 87305378 | 87305739 | Intron |
| chr7 | 87308780 | 87312599 | Intron |
| chr7 | 87320765 | 87321010 | Intron |
| chr7 | 87331136 | 87331751 | Intron |

| | | | |
|---|---|---|---|
| chr7 | 87331901 | 87332424 | Intron |
| chr7 | 87332577 | 87332842 | Intron |
| chr7 | 87332926 | 87337320 | Intron |
| chr7 | 87342399 | 87342863 | Promoter |
| chr7 | 87343504 | 87343995 | Promoter |
| chr7 | 87348993 | 87349384 | Intron |
| chr7 | 87349479 | 87349779 | Intron |
| chr7 | 87350881 | 87354294 | Intron |
| chr7 | 87354355 | 87354577 | Intron |
| chr7 | 87406107 | 87406684 | Intron |
| chr7 | 87414553 | 87415026 | Intron |
| chr7 | 87439265 | 87440036 | Intron |
| chr7 | 87481441 | 87482723 | Intron |
| chr7 | 87484287 | 87484507 | Intron |
| chr7 | 87485245 | 87485455 | Intron |
| chr7 | 87514466 | 87515389 | 5' UTR |
| chr7 | 87515431 | 87515730 | Intron |
| chr7 | 87520050 | 87520345 | Intron |
| chr7 | 87545259 | 87547134 | Distal Intergenic |
| chr7 | 87548289 | 87548591 | Distal Intergenic |
| chr7 | 87584319 | 87584854 | Intron |
| chr7 | 87587009 | 87588059 | Intron |
| chr7 | 87618512 | 87618867 | Intron |
| chr7 | 87860552 | 87861021 | Distal Intergenic |
| chr7 | 87907316 | 87908161 | 3' UTR |
| chr7 | 87918845 | 87919243 | Intron |
| chr7 | 87919374 | 87919774 | Intron |
| chr7 | 87924007 | 87925133 | Intron |
| chr7 | 87931188 | 87932524 | Intron |
| chr7 | 87947792 | 87948231 | Distal Intergenic |
| chr7 | 87968207 | 87969743 | Distal Intergenic |
| chr7 | 87969972 | 87971308 | Distal Intergenic |
| chr7 | 87972667 | 87973813 | Distal Intergenic |
| chr7 | 87980691 | 87981109 | Distal Intergenic |
| chr7 | 87982438 | 87983725 | Distal Intergenic |
| chr7 | 87984279 | 87985023 | Distal Intergenic |
| chr7 | 87985187 | 87985890 | Distal Intergenic |
| chr7 | 88045771 | 88045992 | Distal Intergenic |
| chr7 | 88092257 | 88092697 | Distal Intergenic |
| chr7 | 88137656 | 88138317 | Distal Intergenic |
| chr7 | 88327605 | 88328299 | Distal Intergenic |
| chr7 | 88364303 | 88364530 | Distal Intergenic |
| chr7 | 88364860 | 88365059 | Distal Intergenic |
| chr7 | 88455705 | 88456779 | Intron |

| | | | |
|---|---|---|---|
| chr7 | 88464272 | 88465566 | Intron |
| chr7 | 88933642 | 88934444 | Intron |
| chr7 | 89020176 | 89020552 | Distal Intergenic |
| chr7 | 89063232 | 89063610 | Distal Intergenic |
| chr7 | 89134405 | 89134839 | Distal Intergenic |
| chr7 | 89136215 | 89136693 | Distal Intergenic |
| chr7 | 89136827 | 89137242 | Distal Intergenic |
| chr7 | 89137447 | 89138001 | Distal Intergenic |
| chr7 | 89140385 | 89141006 | Distal Intergenic |
| chr7 | 89141617 | 89141830 | Distal Intergenic |
| chr7 | 89143759 | 89144975 | Distal Intergenic |
| chr7 | 89152237 | 89152485 | Distal Intergenic |
| chr7 | 89152599 | 89153111 | Distal Intergenic |
| chr7 | 89161609 | 89163735 | Distal Intergenic |
| chr7 | 89172996 | 89175509 | Distal Intergenic |
| chr7 | 89175820 | 89176074 | Distal Intergenic |
| chr7 | 89201426 | 89202029 | Distal Intergenic |
| chr7 | 89212724 | 89216228 | Distal Intergenic |
| chr7 | 89216436 | 89217087 | Distal Intergenic |
| chr7 | 89612782 | 89613052 | Distal Intergenic |
| chr7 | 89613123 | 89613375 | Distal Intergenic |
| chr7 | 89628069 | 89628330 | Distal Intergenic |
| chr7 | 89797158 | 89797393 | Promoter |
| chr7 | 89813397 | 89813600 | Intron |
| chr7 | 89813648 | 89813968 | Intron |
| chr7 | 89814130 | 89814333 | Intron |
| chr7 | 89846186 | 89846936 | Intron |
| chr7 | 90043157 | 90043757 | 3' UTR |
| chr7 | 90043824 | 90044145 | 3' UTR |
| chr7 | 90204786 | 90205516 | Intron |
| chr7 | 90205565 | 90205771 | Intron |
| chr7 | 90455012 | 90455249 | Intron |
| chr7 | 91031719 | 91032568 | Distal Intergenic |
| chr7 | 91032901 | 91033290 | Distal Intergenic |
| chr7 | 91111892 | 91112091 | Distal Intergenic |
| chr7 | 91177996 | 91178201 | Distal Intergenic |
| chr7 | 91236637 | 91237284 | Distal Intergenic |
| chr7 | 91266427 | 91266684 | Distal Intergenic |
| chr7 | 91284072 | 91285287 | Distal Intergenic |
| chr7 | 91290420 | 91290648 | Distal Intergenic |
| chr7 | 91332860 | 91334071 | Distal Intergenic |
| chr7 | 91394577 | 91394822 | Distal Intergenic |
| chr7 | 91394907 | 91395901 | Distal Intergenic |
| chr7 | 91613560 | 91615824 | Intron |

| | | | |
|---|---|---|---|
| chr7 | 91669404 | 91670575 | 3' UTR |
| chr7 | 91670937 | 91671305 | 3' UTR |
| chr7 | 91695452 | 91695689 | Intron |
| chr7 | 91827765 | 91827965 | Downstream |
| chr7 | 93635484 | 93638098 | Distal Intergenic |
| chr7 | 95348008 | 95348838 | Distal Intergenic |
| chr7 | 95419830 | 95420475 | Intron |
| chr7 | 95573887 | 95574198 | Intron |
| chr7 | 95574248 | 95574909 | Intron |
| chr7 | 95578429 | 95578717 | Intron |
| chr7 | 95578873 | 95579550 | Intron |
| chr7 | 95580021 | 95580305 | Intron |
| chr7 | 95614394 | 95614844 | Intron |
| chr7 | 95616236 | 95618043 | Exon |
| chr7 | 95623974 | 95624308 | Intron |
| chr7 | 96084033 | 96084255 | Distal Intergenic |
| chr7 | 96326713 | 96327316 | Intron |
| chr7 | 96327356 | 96327621 | Intron |
| chr7 | 96590128 | 96590632 | Distal Intergenic |
| chr7 | 96612386 | 96612663 | Intron |
| chr7 | 96612775 | 96613134 | Intron |
| chr7 | 96665371 | 96665953 | Distal Intergenic |
| chr7 | 96666291 | 96667001 | Distal Intergenic |
| chr7 | 96667322 | 96668270 | Distal Intergenic |
| chr7 | 96671864 | 96672766 | Distal Intergenic |
| chr7 | 96689014 | 96690448 | Distal Intergenic |
| chr7 | 96707605 | 96710320 | Distal Intergenic |
| chr7 | 96712728 | 96712945 | Distal Intergenic |
| chr7 | 96728627 | 96730356 | Distal Intergenic |
| chr7 | 96754626 | 96754865 | Intron |
| chr7 | 96759272 | 96760132 | Intron |
| chr7 | 96824861 | 96825090 | Distal Intergenic |
| chr7 | 96887246 | 96887981 | Distal Intergenic |
| chr7 | 96936283 | 96936488 | Distal Intergenic |
| chr7 | 96957805 | 96958533 | Distal Intergenic |
| chr7 | 97089481 | 97089727 | Distal Intergenic |
| chr7 | 97128091 | 97128676 | Distal Intergenic |
| chr7 | 97129051 | 97129253 | Distal Intergenic |
| chr7 | 97154963 | 97156397 | Distal Intergenic |
| chr7 | 97187902 | 97188190 | Distal Intergenic |
| chr7 | 97190114 | 97190528 | Distal Intergenic |
| chr7 | 97190816 | 97191102 | Distal Intergenic |
| chr7 | 97208049 | 97208637 | Distal Intergenic |
| chr7 | 97244013 | 97244680 | Distal Intergenic |

| | | | |
|---|---|---|---|
| chr7 | 97244798 | 97245068 | Distal Intergenic |
| chr7 | 97245171 | 97245984 | Distal Intergenic |
| chr7 | 97278043 | 97278406 | Distal Intergenic |
| chr7 | 97279194 | 97280798 | Distal Intergenic |
| chr7 | 99345985 | 99347295 | Distal Intergenic |
| chr7 | 99351106 | 99351518 | Distal Intergenic |
| chr7 | 99351569 | 99351772 | Downstream |
| chr7 | 99578856 | 99580016 | Promoter |
| chr7 | 99627858 | 99628122 | Exon |
| chr7 | 99790473 | 99790738 | Intron |
| chr7 | 99815325 | 99815804 | Promoter |
| chr7 | 99816914 | 99818440 | Promoter |
| chr7 | 99847073 | 99847488 | Intron |
| chr7 | 99851615 | 99852202 | Intron |
| chr7 | 99885612 | 99887968 | Distal Intergenic |
| chr7 | 99888015 | 99890001 | Distal Intergenic |
| chr7 | 99923552 | 99925042 | Intron |
| chr7 | 99945182 | 99946338 | Intron |
| chr7 | 99971614 | 99972230 | Promoter |
| chr7 | 99974303 | 99975267 | Intron |
| chr7 | 99985451 | 99986561 | Intron |
| chr7 | 99986662 | 99986867 | Intron |
| chr7 | 100021859 | 100023620 | 5' UTR |
| chr7 | 100023706 | 100023921 | Intron |
| chr7 | 100100090 | 100100372 | Distal Intergenic |
| chr7 | 100101154 | 100101526 | Distal Intergenic |
| chr7 | 103658440 | 103659315 | Distal Intergenic |
| chr7 | 103670333 | 103670578 | Distal Intergenic |
| chr7 | 103670994 | 103671201 | Distal Intergenic |
| chr7 | 103710760 | 103711700 | Distal Intergenic |
| chr7 | 103723771 | 103724026 | Distal Intergenic |
| chr7 | 103856204 | 103856447 | Distal Intergenic |
| chr7 | 104520463 | 104520671 | Intron |
| chr7 | 104659741 | 104660201 | Intron |
| chr7 | 104728729 | 104729603 | Intron |
| chr7 | 104793445 | 104794215 | Intron |
| chr7 | 104849202 | 104849483 | Intron |
| chr7 | 104849652 | 104850239 | Intron |
| chr7 | 104857635 | 104858004 | Intron |
| chr7 | 104919376 | 104919575 | Intron |
| chr7 | 104937054 | 104937259 | Intron |
| chr7 | 104937764 | 104938078 | 5' UTR |
| chr7 | 104938495 | 104938772 | Promoter |
| chr7 | 104940009 | 104941684 | Promoter |

| | | | |
|---|---|---|---|
| chr7 | 104978479 | 104979936 | Intron |
| chr7 | 105264513 | 105264838 | Promoter |
| chr7 | 105265234 | 105265960 | Promoter |
| chr7 | 105319746 | 105319945 | Promoter |
| chr7 | 105326993 | 105327471 | Intron |
| chr7 | 105452385 | 105452871 | Intron |
| chr7 | 105457200 | 105457435 | Intron |
| chr7 | 105457756 | 105457975 | Intron |
| chr7 | 105537241 | 105537537 | Intron |
| chr7 | 105537713 | 105537912 | Intron |
| chr7 | 105539213 | 105539552 | Intron |
| chr7 | 105539634 | 105541055 | Intron |
| chr7 | 105564043 | 105564634 | Intron |
| chr7 | 105640463 | 105640798 | Intron |
| chr7 | 105706920 | 105707944 | Distal Intergenic |
| chr7 | 105708121 | 105708348 | Distal Intergenic |
| chr7 | 105708526 | 105711176 | Distal Intergenic |
| chr7 | 105736370 | 105736656 | Intron |
| chr7 | 105804890 | 105805171 | Distal Intergenic |
| chr7 | 105841239 | 105842090 | Distal Intergenic |
| chr7 | 105842128 | 105842327 | Distal Intergenic |
| chr7 | 105850467 | 105851303 | Distal Intergenic |
| chr7 | 106116151 | 106117927 | Distal Intergenic |
| chr7 | 106146263 | 106147270 | Intron |
| chr7 | 106172095 | 106172936 | Intron |
| chr7 | 106175059 | 106176642 | Intron |
| chr7 | 106233598 | 106233841 | Intron |
| chr7 | 106417731 | 106418733 | Distal Intergenic |
| chr7 | 106424640 | 106426128 | Distal Intergenic |
| chr7 | 106451098 | 106451329 | Distal Intergenic |
| chr7 | 106451955 | 106452183 | Distal Intergenic |
| chr7 | 106452522 | 106453945 | Distal Intergenic |
| chr7 | 106467242 | 106468234 | Distal Intergenic |
| chr7 | 106469422 | 106469703 | Distal Intergenic |
| chr7 | 106470261 | 106471768 | Distal Intergenic |
| chr7 | 106542428 | 106542664 | Intron |
| chr7 | 106556301 | 106556500 | Distal Intergenic |
| chr7 | 106607660 | 106608534 | Distal Intergenic |
| chr7 | 106608675 | 106608882 | Distal Intergenic |
| chr7 | 106614633 | 106614995 | Distal Intergenic |
| chr7 | 106622595 | 106623178 | Distal Intergenic |
| chr7 | 106631774 | 106632032 | Distal Intergenic |
| chr7 | 106974450 | 106975161 | Intron |
| chr7 | 106975369 | 106975636 | Intron |

| | | | |
|---|---|---|---|
| chr7 | 106975803 | 106976812 | Intron |
| chr7 | 107007007 | 107007206 | Intron |
| chr7 | 107095390 | 107095606 | Intron |
| chr7 | 107123109 | 107124208 | Intron |
| chr7 | 107127421 | 107128461 | Intron |
| chr7 | 107578386 | 107578711 | Intron |
| chr7 | 107578859 | 107580062 | Intron |
| chr7 | 107613134 | 107614772 | Exon |
| chr7 | 107739286 | 107740013 | Intron |
| chr7 | 107826484 | 107827150 | Intron |
| chr7 | 107827278 | 107828702 | Intron |
| chr7 | 107988620 | 107988835 | Intron |
| chr7 | 108080792 | 108081059 | Intron |
| chr7 | 108189708 | 108189969 | Distal Intergenic |
| chr7 | 108192460 | 108192930 | Distal Intergenic |
| chr7 | 108193004 | 108193203 | Distal Intergenic |
| chr7 | 108264758 | 108265324 | Distal Intergenic |
| chr7 | 108834132 | 108835248 | Distal Intergenic |
| chr7 | 108985263 | 108985596 | Distal Intergenic |
| chr7 | 108985675 | 108985877 | Distal Intergenic |
| chr7 | 109053672 | 109054628 | Distal Intergenic |
| chr7 | 109081294 | 109082290 | Distal Intergenic |
| chr7 | 109113589 | 109114174 | Distal Intergenic |
| chr7 | 109117438 | 109117645 | Distal Intergenic |
| chr7 | 109117968 | 109118761 | Distal Intergenic |
| chr7 | 109119200 | 109120763 | Distal Intergenic |
| chr7 | 109214027 | 109214293 | Distal Intergenic |
| chr7 | 109371504 | 109371816 | Distal Intergenic |
| chr7 | 109371990 | 109373245 | Distal Intergenic |
| chr7 | 109386989 | 109388167 | Distal Intergenic |
| chr7 | 109398626 | 109399111 | Distal Intergenic |
| chr7 | 109399461 | 109399989 | Distal Intergenic |
| chr7 | 109400820 | 109401189 | Distal Intergenic |
| chr7 | 109537601 | 109538024 | Distal Intergenic |
| chr7 | 109544031 | 109545100 | Distal Intergenic |
| chr7 | 109546878 | 109548958 | Distal Intergenic |
| chr7 | 109590279 | 109591082 | Distal Intergenic |
| chr7 | 109627197 | 109628917 | Distal Intergenic |
| chr7 | 111366323 | 111368181 | 3' UTR |
| chr7 | 111368316 | 111368603 | 3' UTR |
| chr7 | 112884410 | 112884609 | Distal Intergenic |
| chr7 | 112885201 | 112885558 | Distal Intergenic |
| chr7 | 112885861 | 112886084 | Distal Intergenic |
| chr7 | 112886825 | 112887822 | Distal Intergenic |

| | | | |
|---|---|---|---|
| chr7 | 112892348 | 112892555 | Distal Intergenic |
| chr7 | 112892645 | 112893096 | Distal Intergenic |
| chr7 | 113677960 | 113678205 | Distal Intergenic |
| chr7 | 113694732 | 113694944 | Distal Intergenic |
| chr7 | 113700063 | 113701195 | Distal Intergenic |
| chr7 | 114825103 | 114826519 | Distal Intergenic |
| chr7 | 114828715 | 114829335 | Distal Intergenic |
| chr7 | 114829372 | 114829643 | Distal Intergenic |
| chr7 | 114829880 | 114830122 | Distal Intergenic |
| chr7 | 114831147 | 114831700 | Distal Intergenic |
| chr7 | 114869586 | 114870383 | Distal Intergenic |
| chr7 | 116215497 | 116215874 | Distal Intergenic |
| chr7 | 116279072 | 116279771 | Distal Intergenic |
| chr7 | 116280591 | 116281752 | Distal Intergenic |
| chr7 | 116284351 | 116284669 | Distal Intergenic |
| chr7 | 116284805 | 116285064 | Distal Intergenic |
| chr7 | 116285117 | 116286833 | Distal Intergenic |
| chr7 | 116286972 | 116287634 | Distal Intergenic |
| chr7 | 116287897 | 116288122 | Distal Intergenic |
| chr7 | 116288656 | 116290287 | Distal Intergenic |
| chr7 | 116330966 | 116331323 | Intron |
| chr7 | 116575160 | 116575463 | Distal Intergenic |
| chr7 | 116575500 | 116575763 | Distal Intergenic |
| chr7 | 116614503 | 116614799 | Intron |
| chr7 | 116661939 | 116662285 | Intron |
| chr7 | 116662658 | 116663192 | Intron |
| chr7 | 116663369 | 116663935 | Intron |
| chr7 | 116664256 | 116664529 | Intron |
| chr7 | 116664834 | 116665448 | Intron |
| chr7 | 116665684 | 116666078 | Intron |
| chr7 | 116666827 | 116667035 | Intron |
| chr7 | 116690022 | 116690236 | Intron |
| chr7 | 116690834 | 116691033 | Intron |
| chr7 | 116705037 | 116705279 | Intron |
| chr7 | 116705679 | 116705994 | Intron |
| chr7 | 116711655 | 116713162 | Exon |
| chr7 | 116722045 | 116723599 | Intron |
| chr7 | 116752837 | 116753943 | Intron |
| chr7 | 116764919 | 116765233 | Intron |
| chr7 | 116773102 | 116775888 | Exon |
| chr7 | 116779017 | 116779232 | Intron |
| chr7 | 116779290 | 116781875 | Intron |
| chr7 | 117130006 | 117130415 | Intron |
| chr7 | 117152007 | 117152226 | Intron |

| | | | |
|---|---|---|---|
| chr7 | 117152324 | 117152546 | Intron |
| chr7 | 117161934 | 117162988 | Intron |
| chr7 | 117203670 | 117204964 | Exon |
| chr7 | 117205181 | 117206344 | Intron |
| chr7 | 117252430 | 117253266 | Intron |
| chr7 | 117266859 | 117267223 | Intron |
| chr7 | 117267270 | 117267469 | Intron |
| chr7 | 117307897 | 117308291 | 3' UTR |
| chr7 | 117380221 | 117380802 | Intron |
| chr7 | 117380868 | 117382015 | Intron |
| chr7 | 117431258 | 117433900 | Exon |
| chr7 | 117469053 | 117470522 | Intron |
| chr7 | 117478077 | 117479150 | Intron |
| chr7 | 117480717 | 117481215 | Intron |
| chr7 | 117481290 | 117481511 | Intron |
| chr7 | 117574557 | 117575799 | Distal Intergenic |
| chr7 | 117672300 | 117673293 | Distal Intergenic |
| chr7 | 117682366 | 117683398 | Distal Intergenic |
| chr7 | 117683679 | 117683945 | Distal Intergenic |
| chr7 | 117698581 | 117698838 | Distal Intergenic |
| chr7 | 117757393 | 117757762 | Distal Intergenic |
| chr7 | 117812367 | 117812570 | Distal Intergenic |
| chr7 | 117814892 | 117815259 | Distal Intergenic |
| chr7 | 117912830 | 117915134 | Distal Intergenic |
| chr7 | 117916540 | 117918047 | Distal Intergenic |
| chr7 | 117923340 | 117924842 | Distal Intergenic |
| chr7 | 117924882 | 117925385 | Distal Intergenic |
| chr7 | 117928169 | 117929000 | Distal Intergenic |
| chr7 | 117929512 | 117929747 | Distal Intergenic |
| chr7 | 117930314 | 117930872 | Distal Intergenic |
| chr7 | 117930983 | 117931297 | Distal Intergenic |
| chr7 | 117939223 | 117939956 | Distal Intergenic |
| chr7 | 117940423 | 117941883 | Distal Intergenic |
| chr7 | 117950785 | 117951069 | Distal Intergenic |
| chr7 | 117974624 | 117975458 | Distal Intergenic |
| chr7 | 117994964 | 117995328 | Distal Intergenic |
| chr7 | 118084941 | 118085228 | Distal Intergenic |
| chr7 | 118225517 | 118226231 | Distal Intergenic |
| chr7 | 118226859 | 118227391 | Distal Intergenic |
| chr7 | 118265810 | 118266925 | Distal Intergenic |
| chr7 | 118299019 | 118300732 | Distal Intergenic |
| chr7 | 118300798 | 118301848 | Distal Intergenic |
| chr7 | 118388704 | 118389488 | Distal Intergenic |
| chr7 | 118845333 | 118846192 | Distal Intergenic |

| | | | |
|---|---|---|---|
| chr7 | 118859927 | 118861646 | Distal Intergenic |
| chr7 | 118862323 | 118862879 | Distal Intergenic |
| chr7 | 118866974 | 118868421 | Distal Intergenic |
| chr7 | 118868633 | 118868873 | Distal Intergenic |
| chr7 | 118869007 | 118869672 | Distal Intergenic |
| chr7 | 118914899 | 118915532 | Distal Intergenic |
| chr7 | 119120022 | 119120299 | Distal Intergenic |
| chr7 | 119120351 | 119120608 | Distal Intergenic |
| chr7 | 119120719 | 119120968 | Distal Intergenic |
| chr7 | 119167035 | 119167390 | Distal Intergenic |
| chr7 | 119167460 | 119168419 | Distal Intergenic |
| chr7 | 119174050 | 119174269 | Distal Intergenic |
| chr7 | 119284184 | 119284675 | Distal Intergenic |
| chr7 | 119284747 | 119284984 | Distal Intergenic |
| chr7 | 119286903 | 119289003 | Distal Intergenic |
| chr7 | 119300847 | 119301240 | Distal Intergenic |
| chr7 | 119331355 | 119331987 | Distal Intergenic |
| chr7 | 119345455 | 119345840 | Distal Intergenic |
| chr7 | 119345903 | 119347109 | Distal Intergenic |
| chr7 | 119348237 | 119348653 | Distal Intergenic |
| chr7 | 119370179 | 119370881 | Distal Intergenic |
| chr7 | 119371044 | 119371342 | Distal Intergenic |
| chr7 | 119389787 | 119390187 | Distal Intergenic |
| chr7 | 119420286 | 119420910 | Distal Intergenic |
| chr7 | 119441943 | 119444099 | Distal Intergenic |
| chr7 | 119447970 | 119448310 | Distal Intergenic |
| chr7 | 119479628 | 119479841 | Distal Intergenic |
| chr7 | 119519292 | 119519496 | Distal Intergenic |
| chr7 | 119519696 | 119519907 | Distal Intergenic |
| chr7 | 119551994 | 119552298 | Distal Intergenic |
| chr7 | 119560497 | 119560752 | Distal Intergenic |
| chr7 | 119596984 | 119597439 | Distal Intergenic |
| chr7 | 119597610 | 119598012 | Distal Intergenic |
| chr7 | 119598063 | 119598989 | Distal Intergenic |
| chr7 | 119599039 | 119600031 | Distal Intergenic |
| chr7 | 119610763 | 119612267 | Distal Intergenic |
| chr7 | 119642798 | 119643006 | Distal Intergenic |
| chr7 | 119643263 | 119643561 | Distal Intergenic |
| chr7 | 119806323 | 119806927 | Distal Intergenic |
| chr7 | 119840537 | 119840999 | Distal Intergenic |
| chr7 | 120050779 | 120051262 | Intron |
| chr7 | 120147703 | 120147968 | Intron |
| chr7 | 120148899 | 120149871 | Intron |
| chr7 | 120152280 | 120153057 | Intron |

| | | | |
|---|---|---|---|
| chr7 | 120190896 | 120192544 | Intron |
| chr7 | 120207189 | 120207593 | Intron |
| chr7 | 120249849 | 120250481 | Intron |
| chr7 | 120250690 | 120251146 | Intron |
| chr7 | 120277295 | 120277566 | Intron |
| chr7 | 120289956 | 120290276 | Intron |
| chr7 | 120290347 | 120293740 | Intron |
| chr7 | 120294476 | 120295509 | Intron |
| chr7 | 120296614 | 120296986 | Intron |
| chr7 | 120311983 | 120312675 | Intron |
| chr7 | 120329492 | 120330191 | Intron |
| chr7 | 120377328 | 120377975 | Intron |
| chr7 | 120378819 | 120379018 | Intron |
| chr7 | 120379156 | 120380197 | Intron |
| chr7 | 120385847 | 120386188 | Exon |
| chr7 | 120392849 | 120393103 | Distal Intergenic |
| chr7 | 120403425 | 120404155 | Distal Intergenic |
| chr7 | 120644324 | 120644654 | Intron |
| chr7 | 120645384 | 120646418 | Intron |
| chr7 | 120646523 | 120646814 | Intron |
| chr7 | 120686862 | 120687064 | Exon |
| chr7 | 120687158 | 120687733 | Intron |
| chr7 | 121207288 | 121207564 | Distal Intergenic |
| chr7 | 121314578 | 121315886 | Distal Intergenic |
| chr7 | 121319739 | 121320393 | Distal Intergenic |
| chr7 | 121324528 | 121325159 | Distal Intergenic |
| chr7 | 121419188 | 121419387 | Distal Intergenic |
| chr7 | 122347545 | 122348177 | Intron |
| chr7 | 122452559 | 122453158 | Intron |
| chr7 | 122453247 | 122453446 | Intron |
| chr7 | 122453668 | 122455616 | Intron |
| chr7 | 122465867 | 122467302 | Intron |
| chr7 | 122718266 | 122719508 | Distal Intergenic |
| chr7 | 122723477 | 122724250 | Distal Intergenic |
| chr7 | 122724395 | 122724823 | Distal Intergenic |
| chr7 | 122726502 | 122726773 | Distal Intergenic |
| chr7 | 123236848 | 123237249 | Distal Intergenic |
| chr7 | 123243592 | 123243901 | Intron |
| chr7 | 123332974 | 123334123 | Intron |
| chr7 | 123336189 | 123337498 | Exon |
| chr7 | 123340857 | 123341458 | Intron |
| chr7 | 123341623 | 123342090 | Intron |
| chr7 | 123374374 | 123375235 | Intron |
| chr7 | 123477271 | 123477673 | Distal Intergenic |

| | | | |
|---|---|---|---|
| chr7 | 123477757 | 123477956 | Distal Intergenic |
| chr7 | 123587354 | 123587837 | 5' UTR |
| chr7 | 123587913 | 123588477 | Intron |
| chr7 | 124400273 | 124400610 | Intron |
| chr7 | 124400647 | 124400877 | Intron |
| chr7 | 125248602 | 125248946 | Distal Intergenic |
| chr7 | 125259032 | 125259283 | Distal Intergenic |
| chr7 | 125259507 | 125260175 | Distal Intergenic |
| chr7 | 125272776 | 125273088 | Distal Intergenic |
| chr7 | 125273289 | 125273932 | Distal Intergenic |
| chr7 | 125274087 | 125274432 | Distal Intergenic |
| chr7 | 125385996 | 125386795 | Distal Intergenic |
| chr7 | 125386843 | 125387089 | Distal Intergenic |
| chr7 | 125395308 | 125395672 | Distal Intergenic |
| chr7 | 125395909 | 125396141 | Distal Intergenic |
| chr7 | 125420842 | 125421399 | Distal Intergenic |
| chr7 | 125606630 | 125606829 | Distal Intergenic |
| chr7 | 125623559 | 125625481 | Distal Intergenic |
| chr7 | 125625618 | 125626051 | Distal Intergenic |
| chr7 | 125627320 | 125627532 | Distal Intergenic |
| chr7 | 125630032 | 125630284 | Distal Intergenic |
| chr7 | 125630323 | 125630522 | Distal Intergenic |
| chr7 | 125630562 | 125630949 | Distal Intergenic |
| chr7 | 125631040 | 125632585 | Distal Intergenic |
| chr7 | 125634938 | 125635473 | Distal Intergenic |
| chr7 | 125642135 | 125643372 | Distal Intergenic |
| chr7 | 125651256 | 125652221 | Distal Intergenic |
| chr7 | 125712315 | 125712651 | Distal Intergenic |
| chr7 | 126038797 | 126039669 | Distal Intergenic |
| chr7 | 126072925 | 126073225 | Distal Intergenic |
| chr7 | 126073493 | 126073915 | Distal Intergenic |
| chr7 | 126078764 | 126079045 | 3' UTR |
| chr7 | 126079205 | 126079560 | 3' UTR |
| chr7 | 126743030 | 126743396 | Intron |
| chr7 | 127003797 | 127005872 | Distal Intergenic |
| chr7 | 127102846 | 127103766 | Distal Intergenic |
| chr7 | 127797811 | 127798569 | Distal Intergenic |
| chr7 | 127981624 | 127982102 | Intron |
| chr7 | 127982327 | 127982736 | Intron |
| chr7 | 127982785 | 127982984 | Promoter |
| chr7 | 128008182 | 128010842 | Distal Intergenic |
| chr7 | 128012517 | 128013432 | Distal Intergenic |
| chr7 | 128013596 | 128013819 | Distal Intergenic |
| chr7 | 128845879 | 128846101 | Promoter |

| | | | |
|---|---|---|---|
| chr7 | 128853925 | 128855499 | Distal Intergenic |
| chr7 | 129158733 | 129159262 | Distal Intergenic |
| chr7 | 129209565 | 129210549 | Distal Intergenic |
| chr7 | 129210681 | 129210923 | Distal Intergenic |
| chr7 | 129302235 | 129302670 | Intron |
| chr7 | 130113970 | 130114307 | Distal Intergenic |
| chr7 | 130251822 | 130252021 | Intron |
| chr7 | 130252126 | 130252727 | Intron |
| chr7 | 130253459 | 130254511 | Intron |
| chr7 | 130264308 | 130264684 | Intron |
| chr7 | 130267424 | 130267623 | Intron |
| chr7 | 130554772 | 130555084 | Downstream |
| chr7 | 132526304 | 132527337 | Intron |
| chr7 | 132642271 | 132642517 | Intron |
| chr7 | 132642700 | 132643727 | Intron |
| chr7 | 132678021 | 132680294 | Intron |
| chr7 | 132682470 | 132682705 | Intron |
| chr7 | 132682794 | 132683302 | Intron |
| chr7 | 132849671 | 132850852 | Distal Intergenic |
| chr7 | 132859790 | 132860733 | Distal Intergenic |
| chr7 | 133935099 | 133935320 | Intron |
| chr7 | 133935402 | 133936020 | Intron |
| chr7 | 133936118 | 133936518 | Intron |
| chr7 | 134119659 | 134119858 | Distal Intergenic |
| chr7 | 134119920 | 134120233 | Distal Intergenic |
| chr7 | 134321263 | 134321818 | Distal Intergenic |
| chr7 | 134339099 | 134339300 | Intron |
| chr7 | 134339348 | 134339576 | Intron |
| chr7 | 134375718 | 134376530 | Distal Intergenic |
| chr7 | 134613767 | 134615357 | Promoter |
| chr7 | 134622041 | 134622490 | Intron |
| chr7 | 134655537 | 134655799 | Distal Intergenic |
| chr7 | 134656203 | 134656410 | Distal Intergenic |
| chr7 | 134704849 | 134706978 | Intron |
| chr7 | 134707106 | 134707417 | Intron |
| chr7 | 134889066 | 134889622 | 5' UTR |
| chr7 | 135018977 | 135019586 | Distal Intergenic |
| chr7 | 135056170 | 135056721 | Intron |
| chr7 | 135168228 | 135169027 | Intron |
| chr7 | 135169547 | 135169746 | Intron |
| chr7 | 135232617 | 135232844 | Distal Intergenic |
| chr7 | 135232933 | 135233234 | Distal Intergenic |
| chr7 | 135381174 | 135381584 | Intron |
| chr7 | 135461838 | 135464913 | Distal Intergenic |

| | | | |
|---|---|---|---|
| chr7 | 135506364 | 135508802 | Distal Intergenic |
| chr7 | 135540476 | 135541964 | Distal Intergenic |
| chr7 | 135545871 | 135547308 | Distal Intergenic |
| chr7 | 135582912 | 135583535 | Distal Intergenic |
| chr7 | 135603360 | 135603622 | Distal Intergenic |
| chr7 | 135603965 | 135604170 | Distal Intergenic |
| chr7 | 135667077 | 135668244 | Distal Intergenic |
| chr7 | 135668373 | 135668744 | Distal Intergenic |
| chr7 | 135669939 | 135670468 | Distal Intergenic |
| chr7 | 135672306 | 135672664 | Distal Intergenic |
| chr7 | 135672889 | 135673191 | Distal Intergenic |
| chr7 | 135673294 | 135673493 | Distal Intergenic |
| chr7 | 135675425 | 135675671 | Distal Intergenic |
| chr7 | 135676222 | 135676559 | Distal Intergenic |
| chr7 | 135696866 | 135697313 | Distal Intergenic |
| chr7 | 135697358 | 135698242 | Distal Intergenic |
| chr7 | 135765406 | 135767067 | Distal Intergenic |
| chr7 | 135815232 | 135815459 | Distal Intergenic |
| chr7 | 135815646 | 135815989 | Distal Intergenic |
| chr7 | 135816154 | 135816369 | Distal Intergenic |
| chr7 | 136794923 | 136795914 | Intron |
| chr7 | 136796458 | 136796657 | Intron |
| chr7 | 136807355 | 136807600 | Intron |
| chr7 | 136807977 | 136808183 | Intron |
| chr7 | 136809253 | 136810052 | Intron |
| chr7 | 136906505 | 136908212 | Distal Intergenic |
| chr7 | 136908295 | 136908755 | Distal Intergenic |
| chr7 | 136990434 | 136990808 | Intron |
| chr7 | 136990859 | 136991361 | Intron |
| chr7 | 137117759 | 137118151 | Intron |
| chr7 | 137118292 | 137119041 | Intron |
| chr7 | 137175965 | 137176180 | Intron |
| chr7 | 137331893 | 137332305 | Intron |
| chr7 | 137356099 | 137358183 | Intron |
| chr7 | 137358498 | 137360377 | Intron |
| chr7 | 137406633 | 137407673 | Intron |
| chr7 | 137408209 | 137408634 | Intron |
| chr7 | 137432345 | 137432820 | Intron |
| chr7 | 137450955 | 137451406 | Intron |
| chr7 | 137451552 | 137451891 | Intron |
| chr7 | 137452502 | 137452971 | Intron |
| chr7 | 137483276 | 137485451 | Intron |
| chr7 | 137718165 | 137718769 | Intron |
| chr7 | 137767692 | 137768012 | Intron |

| | | | |
|---|---|---|---|
| chr7 | 137788492 | 137789593 | Intron |
| chr7 | 137789659 | 137790214 | Exon |
| chr7 | 137790372 | 137791153 | Intron |
| chr7 | 137804620 | 137804915 | Distal Intergenic |
| chr7 | 137813804 | 137814010 | Distal Intergenic |
| chr7 | 137814113 | 137815443 | Distal Intergenic |
| chr7 | 137816249 | 137817579 | Distal Intergenic |
| chr7 | 137840194 | 137840508 | Distal Intergenic |
| chr7 | 138085639 | 138087223 | Distal Intergenic |
| chr7 | 138089220 | 138090921 | Distal Intergenic |
| chr7 | 138091529 | 138091981 | Distal Intergenic |
| chr7 | 138092026 | 138093234 | Distal Intergenic |
| chr7 | 138093688 | 138095285 | Distal Intergenic |
| chr7 | 138108197 | 138108508 | Distal Intergenic |
| chr7 | 138108585 | 138109305 | Distal Intergenic |
| chr7 | 138120482 | 138120806 | Distal Intergenic |
| chr7 | 138120875 | 138121423 | Distal Intergenic |
| chr7 | 138121544 | 138124357 | Distal Intergenic |
| chr7 | 138125653 | 138125963 | Distal Intergenic |
| chr7 | 138126173 | 138126525 | Distal Intergenic |
| chr7 | 138379025 | 138379795 | Distal Intergenic |
| chr7 | 138505112 | 138505682 | Distal Intergenic |
| chr7 | 138704296 | 138704544 | Distal Intergenic |
| chr7 | 138723526 | 138723795 | Distal Intergenic |
| chr7 | 138724014 | 138724343 | Distal Intergenic |
| chr7 | 138725390 | 138726331 | Downstream |
| chr7 | 139423257 | 139423672 | Intron |
| chr7 | 139425846 | 139426070 | Intron |
| chr7 | 139426145 | 139426556 | Intron |
| chr7 | 139481976 | 139483026 | 5' UTR |
| chr7 | 139508202 | 139508401 | Intron |
| chr7 | 139564634 | 139565050 | Intron |
| chr7 | 139926791 | 139929331 | Distal Intergenic |
| chr7 | 139982139 | 139982894 | Distal Intergenic |
| chr7 | 139982974 | 139983173 | Distal Intergenic |
| chr7 | 139993965 | 139995735 | Distal Intergenic |
| chr7 | 140003506 | 140004106 | Distal Intergenic |
| chr7 | 140004203 | 140004420 | Distal Intergenic |
| chr7 | 140044150 | 140044349 | Intron |
| chr7 | 140123526 | 140123946 | Intron |
| chr7 | 140197428 | 140198821 | Distal Intergenic |
| chr7 | 140198939 | 140199228 | Distal Intergenic |
| chr7 | 140289979 | 140291215 | Intron |
| chr7 | 140304191 | 140305127 | Intron |

| | | | |
|---|---|---|---|
| chr7 | 140313245 | 140313685 | Intron |
| chr7 | 140313732 | 140314343 | Intron |
| chr7 | 140315664 | 140315891 | Intron |
| chr7 | 140710440 | 140710813 | 5' UTR |
| chr7 | 140954355 | 140954582 | Intron |
| chr7 | 140974634 | 140974843 | Intron |
| chr7 | 140985434 | 140985844 | Intron |
| chr7 | 141061578 | 141062022 | Intron |
| chr7 | 141306160 | 141306866 | Intron |
| chr7 | 141422459 | 141422681 | Intron |
| chr7 | 141422727 | 141423927 | 3' UTR |
| chr7 | 141428523 | 141429315 | Intron |
| chr7 | 141452576 | 141453105 | Distal Intergenic |
| chr7 | 141454876 | 141455191 | Distal Intergenic |
| chr7 | 141455387 | 141455586 | Distal Intergenic |
| chr7 | 142436877 | 142437560 | Intron |
| chr7 | 142437596 | 142437796 | Intron |
| chr7 | 142796648 | 142797497 | Distal Intergenic |
| chr7 | 142801612 | 142801954 | Distal Intergenic |
| chr7 | 142843625 | 142844825 | Distal Intergenic |
| chr7 | 142845063 | 142845775 | Distal Intergenic |
| chr7 | 142845907 | 142846271 | Distal Intergenic |
| chr7 | 142850938 | 142851137 | Distal Intergenic |
| chr7 | 142851181 | 142851421 | Distal Intergenic |
| chr7 | 144629377 | 144630855 | Distal Intergenic |
| chr7 | 146098673 | 146099035 | Intron |
| chr7 | 146431489 | 146434237 | Intron |
| chr7 | 146438335 | 146438561 | Intron |
| chr7 | 146438659 | 146438920 | Intron |
| chr7 | 146439136 | 146439539 | Intron |
| chr7 | 146443442 | 146444200 | Intron |
| chr7 | 146466109 | 146466340 | Intron |
| chr7 | 146466455 | 146467113 | Intron |
| chr7 | 146469842 | 146470325 | Intron |
| chr7 | 146470521 | 146470901 | Intron |
| chr7 | 146471220 | 146471575 | Exon |
| chr7 | 146568855 | 146569415 | Intron |
| chr7 | 146600015 | 146601908 | Intron |
| chr7 | 146702097 | 146702324 | Intron |
| chr7 | 146702367 | 146702607 | Intron |
| chr7 | 146717524 | 146717735 | Intron |
| chr7 | 147155945 | 147156752 | Intron |
| chr7 | 147156817 | 147159025 | Intron |
| chr7 | 147179440 | 147179939 | Intron |

| | | | |
|---|---|---|---|
| chr7 | 147180182 | 147180548 | Intron |
| chr7 | 148007681 | 148007883 | Intron |
| chr7 | 148017626 | 148017843 | Intron |
| chr7 | 148020365 | 148020951 | Intron |
| chr7 | 148021152 | 148021594 | Intron |
| chr7 | 148141729 | 148142206 | Distal Intergenic |
| chr7 | 148159115 | 148160405 | Distal Intergenic |
| chr7 | 148167812 | 148168357 | Distal Intergenic |
| chr7 | 148172880 | 148173149 | Distal Intergenic |
| chr7 | 148173246 | 148173995 | Distal Intergenic |
| chr7 | 148195692 | 148195894 | Distal Intergenic |
| chr7 | 148196397 | 148196891 | Distal Intergenic |
| chr7 | 148264322 | 148266562 | Distal Intergenic |
| chr7 | 148353366 | 148353667 | Distal Intergenic |
| chr7 | 148353733 | 148354631 | Distal Intergenic |
| chr7 | 148422760 | 148424430 | Intron |
| chr7 | 148513722 | 148514147 | 3' UTR |
| chr7 | 148517947 | 148518178 | Intron |
| chr7 | 148524895 | 148525250 | Intron |
| chr7 | 148525450 | 148526073 | Exon |
| chr7 | 148526192 | 148526417 | Intron |
| chr7 | 148558245 | 148558527 | Intron |
| chr7 | 148653768 | 148654777 | Distal Intergenic |
| chr7 | 148715754 | 148715953 | Intron |
| chr7 | 148771382 | 148771628 | 5' UTR |
| chr7 | 149245052 | 149245254 | Exon |
| chr7 | 149245963 | 149246373 | Exon |
| chr7 | 149269889 | 149270178 | Intron |
| chr7 | 149270379 | 149272102 | Intron |
| chr7 | 149526281 | 149526488 | Intron |
| chr7 | 149548662 | 149549229 | Intron |
| chr7 | 149549292 | 149549543 | Intron |
| chr7 | 149561892 | 149562109 | 3' UTR |
| chr7 | 149573540 | 149573759 | Intron |
| chr7 | 149579085 | 149579563 | Exon |
| chr7 | 149619298 | 149621934 | Distal Intergenic |
| chr7 | 149621996 | 149623162 | Distal Intergenic |
| chr7 | 150210918 | 150211869 | Promoter |
| chr7 | 150248142 | 150248873 | Distal Intergenic |
| chr7 | 150535872 | 150536356 | Intron |
| chr7 | 150536442 | 150536759 | Intron |
| chr7 | 150542686 | 150543156 | Intron |
| chr7 | 150810567 | 150811325 | Promoter |
| chr7 | 150815663 | 150816021 | Exon |

| | | | |
|---|---|---|---|
| chr7 | 150817401 | 150817854 | Intron |
| chr7 | 150876710 | 150877788 | Intron |
| chr7 | 150923917 | 150924800 | Promoter |
| chr7 | 150925153 | 150925399 | Promoter |
| chr7 | 150971157 | 150971437 | Intron |
| chr7 | 151017499 | 151017775 | Distal Intergenic |
| chr7 | 151017854 | 151018358 | Distal Intergenic |
| chr7 | 151223889 | 151224296 | Distal Intergenic |
| chr7 | 151655311 | 151655599 | Intron |
| chr7 | 151749433 | 151749790 | Intron |
| chr7 | 151749971 | 151750488 | Intron |
| chr7 | 151752659 | 151753019 | Intron |
| chr7 | 151753090 | 151753580 | Intron |
| chr7 | 151753619 | 151753823 | Intron |
| chr7 | 151798484 | 151798846 | Intron |
| chr7 | 151799045 | 151799272 | Intron |
| chr7 | 151869963 | 151870739 | Intron |
| chr7 | 151870970 | 151871603 | Exon |
| chr7 | 151933337 | 151933967 | Intron |
| chr7 | 152126050 | 152126538 | Intron |
| chr7 | 152126817 | 152127551 | Intron |
| chr7 | 152149497 | 152151264 | Distal Intergenic |
| chr7 | 152156312 | 152157382 | Distal Intergenic |
| chr7 | 152157820 | 152158764 | Distal Intergenic |
| chr7 | 152158887 | 152159261 | Distal Intergenic |
| chr7 | 152292155 | 152292544 | Distal Intergenic |
| chr7 | 152297274 | 152297546 | Distal Intergenic |
| chr7 | 152470144 | 152470440 | Intron |
| chr7 | 152470638 | 152471085 | Intron |
| chr7 | 152516307 | 152516875 | Intron |
| chr8 | 3077171 | 3078161 | Intron |
| chr8 | 3393110 | 3393371 | Intron |
| chr8 | 3407490 | 3409835 | Intron |
| chr8 | 3411222 | 3413415 | Intron |
| chr8 | 3417828 | 3418482 | Intron |
| chr8 | 3424385 | 3424784 | Intron |
| chr8 | 3424869 | 3426824 | Intron |
| chr8 | 3428598 | 3429087 | Intron |
| chr8 | 3429126 | 3430559 | Intron |
| chr8 | 3515169 | 3516495 | Intron |
| chr8 | 3631200 | 3631528 | Intron |
| chr8 | 3647213 | 3648233 | Intron |
| chr8 | 3676484 | 3677368 | Intron |
| chr8 | 3991344 | 3991836 | Intron |

| | | | |
|---|---|---|---|
| chr8 | 4013112 | 4015295 | Intron |
| chr8 | 4165349 | 4165904 | Intron |
| chr8 | 4252035 | 4252686 | Intron |
| chr8 | 4259440 | 4259803 | Intron |
| chr8 | 4275334 | 4275963 | Intron |
| chr8 | 4276066 | 4276324 | Intron |
| chr8 | 4323872 | 4324248 | Intron |
| chr8 | 4324493 | 4324866 | Intron |
| chr8 | 4612013 | 4612266 | Intron |
| chr8 | 4612658 | 4613568 | Intron |
| chr8 | 4677953 | 4678413 | Intron |
| chr8 | 4678467 | 4679202 | Intron |
| chr8 | 5283162 | 5284384 | Distal Intergenic |
| chr8 | 7086316 | 7086989 | Distal Intergenic |
| chr8 | 8109968 | 8110423 | Distal Intergenic |
| chr8 | 8186553 | 8188354 | Intron |
| chr8 | 8273426 | 8275417 | Distal Intergenic |
| chr8 | 8275563 | 8275783 | Distal Intergenic |
| chr8 | 8487512 | 8488132 | Distal Intergenic |
| chr8 | 8602941 | 8603429 | Distal Intergenic |
| chr8 | 8650071 | 8650289 | Intron |
| chr8 | 9515117 | 9516988 | Intron |
| chr8 | 9810738 | 9811330 | Distal Intergenic |
| chr8 | 10547517 | 10547745 | Intron |
| chr8 | 10574856 | 10576952 | Distal Intergenic |
| chr8 | 10589073 | 10589669 | Intron |
| chr8 | 10589751 | 10590553 | Intron |
| chr8 | 10629702 | 10630247 | Intron |
| chr8 | 10646751 | 10646950 | Intron |
| chr8 | 10652942 | 10654204 | Intron |
| chr8 | 10664934 | 10665282 | Intron |
| chr8 | 10855791 | 10856062 | Intron |
| chr8 | 10930236 | 10930593 | Intron |
| chr8 | 10930668 | 10930990 | Intron |
| chr8 | 10935108 | 10935848 | Intron |
| chr8 | 10977401 | 10978516 | Intron |
| chr8 | 10982803 | 10983110 | Intron |
| chr8 | 10990687 | 10991580 | Intron |
| chr8 | 10993255 | 10993956 | Intron |
| chr8 | 10994363 | 10994859 | Intron |
| chr8 | 10997253 | 10997527 | Intron |
| chr8 | 11259659 | 11259918 | Intron |
| chr8 | 11263763 | 11264402 | Intron |
| chr8 | 11322558 | 11323496 | Promoter |

| | | | |
|---|---|---|---|
| chr8 | 11346043 | 11346903 | Distal Intergenic |
| chr8 | 11457548 | 11457756 | Distal Intergenic |
| chr8 | 11476576 | 11478061 | Distal Intergenic |
| chr8 | 11497501 | 11497922 | Distal Intergenic |
| chr8 | 11550368 | 11550592 | Intron |
| chr8 | 11632475 | 11634797 | Intron |
| chr8 | 11635918 | 11636279 | Intron |
| chr8 | 12512567 | 12513367 | Intron |
| chr8 | 12516144 | 12517476 | 5' UTR |
| chr8 | 12537464 | 12538250 | Distal Intergenic |
| chr8 | 12544845 | 12545455 | Distal Intergenic |
| chr8 | 12710503 | 12710797 | Distal Intergenic |
| chr8 | 12730503 | 12731691 | Distal Intergenic |
| chr8 | 12756020 | 12756226 | Distal Intergenic |
| chr8 | 12756376 | 12756936 | Distal Intergenic |
| chr8 | 12778858 | 12780282 | Distal Intergenic |
| chr8 | 12996495 | 12997087 | Intron |
| chr8 | 13105757 | 13105976 | Intron |
| chr8 | 13106152 | 13106601 | Intron |
| chr8 | 13139305 | 13140519 | Intron |
| chr8 | 13144788 | 13145002 | Intron |
| chr8 | 13159447 | 13159902 | Intron |
| chr8 | 13160098 | 13160315 | Intron |
| chr8 | 13339171 | 13339730 | Intron |
| chr8 | 13372709 | 13373115 | Promoter |
| chr8 | 13491860 | 13492928 | Distal Intergenic |
| chr8 | 13597464 | 13597965 | Distal Intergenic |
| chr8 | 13624179 | 13625636 | Distal Intergenic |
| chr8 | 13664669 | 13665295 | Distal Intergenic |
| chr8 | 13667886 | 13669143 | Distal Intergenic |
| chr8 | 13694881 | 13696397 | Distal Intergenic |
| chr8 | 13758188 | 13758567 | Distal Intergenic |
| chr8 | 13869449 | 13869650 | Distal Intergenic |
| chr8 | 14321455 | 14321998 | Intron |
| chr8 | 14880571 | 14882068 | Intron |
| chr8 | 14888528 | 14888770 | Intron |
| chr8 | 14913884 | 14914083 | Intron |
| chr8 | 14917111 | 14917868 | Intron |
| chr8 | 17593906 | 17595431 | Intron |
| chr8 | 17599638 | 17601346 | Exon |
| chr8 | 18595502 | 18595887 | Intron |
| chr8 | 18608972 | 18609432 | Intron |
| chr8 | 18619098 | 18619554 | Intron |
| chr8 | 18696034 | 18696677 | Intron |

| | | | |
|---|---|---|---|
| chr8 | 18698698 | 18699951 | Intron |
| chr8 | 18768172 | 18768527 | Intron |
| chr8 | 18846641 | 18846875 | Intron |
| chr8 | 19163239 | 19165075 | Distal Intergenic |
| chr8 | 19193979 | 19194238 | Intron |
| chr8 | 19194279 | 19194953 | Intron |
| chr8 | 19784662 | 19785158 | Distal Intergenic |
| chr8 | 19785358 | 19785769 | Distal Intergenic |
| chr8 | 19892108 | 19892943 | Distal Intergenic |
| chr8 | 20019260 | 20020367 | Intron |
| chr8 | 22044235 | 22044511 | Intron |
| chr8 | 22044564 | 22044800 | Intron |
| chr8 | 22057110 | 22057320 | Intron |
| chr8 | 22057493 | 22058030 | Intron |
| chr8 | 22869327 | 22870583 | Intron |
| chr8 | 23303539 | 23304401 | Intron |
| chr8 | 23325033 | 23325589 | Distal Intergenic |
| chr8 | 23325778 | 23326102 | Distal Intergenic |
| chr8 | 23374689 | 23375896 | Distal Intergenic |
| chr8 | 23587460 | 23588627 | Distal Intergenic |
| chr8 | 23590756 | 23591104 | Distal Intergenic |
| chr8 | 23591339 | 23591927 | Distal Intergenic |
| chr8 | 23601703 | 23602587 | Distal Intergenic |
| chr8 | 23803939 | 23804200 | Distal Intergenic |
| chr8 | 23816456 | 23816733 | Distal Intergenic |
| chr8 | 23864702 | 23866122 | Distal Intergenic |
| chr8 | 23867429 | 23868021 | Distal Intergenic |
| chr8 | 23917569 | 23918125 | Distal Intergenic |
| chr8 | 23918166 | 23918406 | Distal Intergenic |
| chr8 | 23983766 | 23983983 | Distal Intergenic |
| chr8 | 23991229 | 23994151 | Distal Intergenic |
| chr8 | 24118644 | 24121475 | Distal Intergenic |
| chr8 | 24126362 | 24126859 | Distal Intergenic |
| chr8 | 24140811 | 24141723 | Distal Intergenic |
| chr8 | 24254655 | 24255030 | Exon |
| chr8 | 24318568 | 24318848 | Intron |
| chr8 | 24366938 | 24367518 | 3' UTR |
| chr8 | 24499111 | 24499507 | Distal Intergenic |
| chr8 | 24577152 | 24577363 | Distal Intergenic |
| chr8 | 24595850 | 24597189 | Distal Intergenic |
| chr8 | 24626722 | 24627600 | Distal Intergenic |
| chr8 | 24664205 | 24665165 | Distal Intergenic |
| chr8 | 24835373 | 24835663 | Distal Intergenic |
| chr8 | 25469152 | 25469551 | Intron |

| | | | |
|---|---|---|---|
| chr8 | 25469837 | 25470148 | Intron |
| chr8 | 25475033 | 25476155 | Intron |
| chr8 | 25510086 | 25510417 | Intron |
| chr8 | 25515802 | 25516045 | Intron |
| chr8 | 25529472 | 25532120 | Intron |
| chr8 | 25540597 | 25542676 | Intron |
| chr8 | 25548024 | 25549441 | Intron |
| chr8 | 25576701 | 25578030 | Intron |
| chr8 | 25629053 | 25629890 | Intron |
| chr8 | 26085442 | 26086853 | Intron |
| chr8 | 26101314 | 26101586 | Intron |
| chr8 | 26120285 | 26121351 | Intron |
| chr8 | 26125878 | 26126175 | Intron |
| chr8 | 26126376 | 26126896 | Intron |
| chr8 | 26177828 | 26178302 | Intron |
| chr8 | 26178621 | 26179714 | Intron |
| chr8 | 26180304 | 26180592 | Intron |
| chr8 | 26211523 | 26212574 | Exon |
| chr8 | 26532215 | 26532641 | Distal Intergenic |
| chr8 | 26548740 | 26549025 | Distal Intergenic |
| chr8 | 26576277 | 26576605 | Distal Intergenic |
| chr8 | 26579217 | 26579858 | Distal Intergenic |
| chr8 | 26707122 | 26708023 | Intron |
| chr8 | 26708100 | 26708300 | Intron |
| chr8 | 26711773 | 26711992 | Intron |
| chr8 | 26712122 | 26712580 | Intron |
| chr8 | 26712699 | 26713380 | Intron |
| chr8 | 26722493 | 26723626 | Promoter |
| chr8 | 26745715 | 26748037 | Distal Intergenic |
| chr8 | 26762011 | 26764600 | Distal Intergenic |
| chr8 | 26909165 | 26909858 | Distal Intergenic |
| chr8 | 26910143 | 26911190 | Distal Intergenic |
| chr8 | 26911813 | 26912259 | Distal Intergenic |
| chr8 | 26914991 | 26915633 | Distal Intergenic |
| chr8 | 26916334 | 26916534 | Distal Intergenic |
| chr8 | 26916587 | 26917385 | Distal Intergenic |
| chr8 | 26929349 | 26930757 | Distal Intergenic |
| chr8 | 26951109 | 26951316 | Distal Intergenic |
| chr8 | 27229895 | 27230165 | Intron |
| chr8 | 27923268 | 27923723 | Intron |
| chr8 | 27934174 | 27935743 | Intron |
| chr8 | 27939742 | 27940243 | Intron |
| chr8 | 27941681 | 27943292 | Promoter |
| chr8 | 27946153 | 27948146 | Promoter |

| | | | |
|---|---|---|---|
| chr8 | 27996935 | 27998839 | Intron |
| chr8 | 28073347 | 28074230 | Distal Intergenic |
| chr8 | 28084701 | 28085410 | Distal Intergenic |
| chr8 | 28085569 | 28085890 | Distal Intergenic |
| chr8 | 28086232 | 28087450 | Distal Intergenic |
| chr8 | 28087948 | 28089051 | Distal Intergenic |
| chr8 | 28153365 | 28153599 | Distal Intergenic |
| chr8 | 28153639 | 28153903 | Distal Intergenic |
| chr8 | 28259995 | 28260407 | Intron |
| chr8 | 28260459 | 28260856 | Intron |
| chr8 | 28371507 | 28372220 | Intron |
| chr8 | 29720167 | 29720399 | Distal Intergenic |
| chr8 | 32222679 | 32224918 | Intron |
| chr8 | 32259565 | 32260378 | Intron |
| chr8 | 32260476 | 32261002 | Intron |
| chr8 | 32277843 | 32278234 | Intron |
| chr8 | 32278478 | 32279231 | Intron |
| chr8 | 32483652 | 32484190 | Intron |
| chr8 | 32631386 | 32632487 | Distal Intergenic |
| chr8 | 32831129 | 32831339 | Intron |
| chr8 | 32831377 | 32831635 | Intron |
| chr8 | 32866003 | 32866730 | Intron |
| chr8 | 33103765 | 33105203 | Distal Intergenic |
| chr8 | 33117913 | 33118198 | Distal Intergenic |
| chr8 | 33118369 | 33119082 | Distal Intergenic |
| chr8 | 33119327 | 33119535 | Distal Intergenic |
| chr8 | 33147497 | 33147745 | Distal Intergenic |
| chr8 | 33148664 | 33150153 | Distal Intergenic |
| chr8 | 33313200 | 33313924 | Promoter |
| chr8 | 33320273 | 33320656 | Intron |
| chr8 | 33328548 | 33329520 | Intron |
| chr8 | 33329579 | 33331318 | Promoter |
| chr8 | 33331405 | 33332625 | Promoter |
| chr8 | 33374531 | 33377307 | Distal Intergenic |
| chr8 | 33396649 | 33398438 | Distal Intergenic |
| chr8 | 34751830 | 34752823 | Distal Intergenic |
| chr8 | 34764755 | 34765030 | Distal Intergenic |
| chr8 | 34770373 | 34771870 | Distal Intergenic |
| chr8 | 34783519 | 34785240 | Distal Intergenic |
| chr8 | 34798025 | 34799376 | Distal Intergenic |
| chr8 | 34840931 | 34841605 | Distal Intergenic |
| chr8 | 34843334 | 34843673 | Distal Intergenic |
| chr8 | 34912602 | 34913317 | Distal Intergenic |
| chr8 | 35078503 | 35079171 | Distal Intergenic |

| | | | |
|---|---|---|---|
| chr8 | 35170955 | 35171327 | Intron |
| chr8 | 35171542 | 35171741 | Intron |
| chr8 | 35235801 | 35237072 | Intron |
| chr8 | 35350382 | 35351508 | Intron |
| chr8 | 35432740 | 35433194 | Intron |
| chr8 | 35646733 | 35646934 | Intron |
| chr8 | 35653808 | 35654963 | Distal Intergenic |
| chr8 | 35675991 | 35676190 | Distal Intergenic |
| chr8 | 35676257 | 35676508 | Distal Intergenic |
| chr8 | 35676544 | 35676744 | Distal Intergenic |
| chr8 | 35676839 | 35677151 | Distal Intergenic |
| chr8 | 35677192 | 35677977 | Distal Intergenic |
| chr8 | 35685636 | 35685872 | Distal Intergenic |
| chr8 | 35685911 | 35686344 | Distal Intergenic |
| chr8 | 36016152 | 36018149 | Distal Intergenic |
| chr8 | 36027660 | 36027954 | Distal Intergenic |
| chr8 | 36028963 | 36029268 | Distal Intergenic |
| chr8 | 36384593 | 36384792 | Distal Intergenic |
| chr8 | 36626842 | 36628412 | Distal Intergenic |
| chr8 | 36629974 | 36630999 | Distal Intergenic |
| chr8 | 36649505 | 36649746 | Intron |
| chr8 | 36650399 | 36650772 | Intron |
| chr8 | 36676682 | 36677007 | Intron |
| chr8 | 36677320 | 36677792 | Intron |
| chr8 | 36927937 | 36929655 | Distal Intergenic |
| chr8 | 36929698 | 36930110 | Distal Intergenic |
| chr8 | 36937116 | 36937744 | Distal Intergenic |
| chr8 | 36944482 | 36945822 | Distal Intergenic |
| chr8 | 36960947 | 36963142 | Distal Intergenic |
| chr8 | 36963278 | 36965885 | Distal Intergenic |
| chr8 | 36975201 | 36976967 | Distal Intergenic |
| chr8 | 36981224 | 36982408 | Distal Intergenic |
| chr8 | 36982610 | 36984171 | Distal Intergenic |
| chr8 | 37002877 | 37004310 | Distal Intergenic |
| chr8 | 37005895 | 37006193 | Distal Intergenic |
| chr8 | 37033874 | 37034765 | Distal Intergenic |
| chr8 | 37068274 | 37068847 | Distal Intergenic |
| chr8 | 37091230 | 37092294 | Distal Intergenic |
| chr8 | 37157121 | 37157811 | Distal Intergenic |
| chr8 | 37157878 | 37158865 | Distal Intergenic |
| chr8 | 37158913 | 37159112 | Distal Intergenic |
| chr8 | 37159221 | 37159675 | Distal Intergenic |
| chr8 | 37179636 | 37181087 | Distal Intergenic |
| chr8 | 37218451 | 37218746 | Distal Intergenic |

| | | | |
|---|---|---|---|
| chr8 | 37226793 | 37228878 | Distal Intergenic |
| chr8 | 37234663 | 37236498 | Distal Intergenic |
| chr8 | 37237274 | 37239529 | Distal Intergenic |
| chr8 | 37247079 | 37247320 | Distal Intergenic |
| chr8 | 37247429 | 37247959 | Distal Intergenic |
| chr8 | 37249740 | 37250931 | Distal Intergenic |
| chr8 | 37251727 | 37252722 | Distal Intergenic |
| chr8 | 37252841 | 37254172 | Distal Intergenic |
| chr8 | 37287925 | 37289989 | Distal Intergenic |
| chr8 | 37310785 | 37311037 | Distal Intergenic |
| chr8 | 37702724 | 37705709 | Exon |
| chr8 | 37754441 | 37754769 | Intron |
| chr8 | 37790020 | 37791478 | Downstream |
| chr8 | 38025339 | 38025665 | Intron |
| chr8 | 41594567 | 41595036 | Intron |
| chr8 | 41595565 | 41595924 | Intron |
| chr8 | 41595962 | 41597823 | Intron |
| chr8 | 42325064 | 42325755 | Intron |
| chr8 | 42433293 | 42434680 | Distal Intergenic |
| chr8 | 42459563 | 42459852 | Distal Intergenic |
| chr8 | 42460134 | 42460457 | Distal Intergenic |
| chr8 | 42502137 | 42502463 | Distal Intergenic |
| chr8 | 43154515 | 43155140 | Intron |
| chr8 | 45346298 | 45346639 | Distal Intergenic |
| chr8 | 45422255 | 45423724 | Distal Intergenic |
| chr8 | 45540580 | 45540968 | Distal Intergenic |
| chr8 | 45541011 | 45541495 | Distal Intergenic |
| chr8 | 45606918 | 45608081 | Distal Intergenic |
| chr8 | 45755966 | 45756783 | Distal Intergenic |
| chr8 | 45843589 | 45845010 | Distal Intergenic |
| chr8 | 45852589 | 45853108 | Distal Intergenic |
| chr8 | 45884704 | 45885981 | Distal Intergenic |
| chr8 | 45886847 | 45887057 | Distal Intergenic |
| chr8 | 45887507 | 45888314 | Distal Intergenic |
| chr8 | 45890642 | 45891219 | Distal Intergenic |
| chr8 | 45949834 | 45950677 | Distal Intergenic |
| chr8 | 45962242 | 45964170 | Distal Intergenic |
| chr8 | 45967043 | 45969038 | Distal Intergenic |
| chr8 | 45979498 | 45980157 | Distal Intergenic |
| chr8 | 46010178 | 46010495 | Distal Intergenic |
| chr8 | 46018675 | 46018905 | Distal Intergenic |
| chr8 | 46019156 | 46020541 | Distal Intergenic |
| chr8 | 46020620 | 46022053 | Distal Intergenic |
| chr8 | 46022106 | 46022355 | Distal Intergenic |

| | | | |
|---|---|---|---|
| chr8 | 46022511 | 46023195 | Distal Intergenic |
| chr8 | 46023514 | 46024044 | Distal Intergenic |
| chr8 | 46024109 | 46024536 | Distal Intergenic |
| chr8 | 46024912 | 46025442 | Distal Intergenic |
| chr8 | 46044507 | 46044948 | Distal Intergenic |
| chr8 | 46045105 | 46045443 | Distal Intergenic |
| chr8 | 46046034 | 46046755 | Distal Intergenic |
| chr8 | 46046882 | 46047136 | Distal Intergenic |
| chr8 | 46062779 | 46062978 | Distal Intergenic |
| chr8 | 46063386 | 46064772 | Distal Intergenic |
| chr8 | 46086033 | 46086246 | Distal Intergenic |
| chr8 | 46086288 | 46086757 | Distal Intergenic |
| chr8 | 46495728 | 46495946 | Distal Intergenic |
| chr8 | 46614207 | 46614458 | Distal Intergenic |
| chr8 | 46665242 | 46665719 | Distal Intergenic |
| chr8 | 46734364 | 46736935 | Distal Intergenic |
| chr8 | 46804930 | 46805314 | Distal Intergenic |
| chr8 | 47531561 | 47532315 | Distal Intergenic |
| chr8 | 47716031 | 47716231 | Distal Intergenic |
| chr8 | 47799286 | 47799534 | Distal Intergenic |
| chr8 | 47825137 | 47827135 | Distal Intergenic |
| chr8 | 47866089 | 47866491 | Distal Intergenic |
| chr8 | 47866614 | 47866832 | Distal Intergenic |
| chr8 | 47896588 | 47896820 | Distal Intergenic |
| chr8 | 47896879 | 47897078 | Distal Intergenic |
| chr8 | 47951730 | 47952732 | Distal Intergenic |
| chr8 | 47953413 | 47955411 | Distal Intergenic |
| chr8 | 47955469 | 47955677 | Distal Intergenic |
| chr8 | 48303805 | 48304060 | Intron |
| chr8 | 48437861 | 48438270 | Intron |
| chr8 | 48460562 | 48461010 | Intron |
| chr8 | 48464787 | 48465067 | Intron |
| chr8 | 48506302 | 48506820 | Intron |
| chr8 | 48758547 | 48759031 | Intron |
| chr8 | 48760270 | 48760517 | Intron |
| chr8 | 48760778 | 48761370 | Intron |
| chr8 | 49017601 | 49017904 | Distal Intergenic |
| chr8 | 49018002 | 49018402 | Distal Intergenic |
| chr8 | 49059113 | 49060557 | Distal Intergenic |
| chr8 | 49067800 | 49069340 | Distal Intergenic |
| chr8 | 49202025 | 49202677 | Distal Intergenic |
| chr8 | 49633613 | 49633823 | Intron |
| chr8 | 49633915 | 49635742 | Intron |
| chr8 | 49636059 | 49636258 | 3' UTR |

| | | | |
|---|---|---|---|
| chr8 | 49639143 | 49639417 | Intron |
| chr8 | 49639634 | 49639937 | Intron |
| chr8 | 50367977 | 50368195 | Distal Intergenic |
| chr8 | 55301111 | 55301377 | Distal Intergenic |
| chr8 | 55301416 | 55301688 | Distal Intergenic |
| chr8 | 55301825 | 55302027 | Distal Intergenic |
| chr8 | 55510372 | 55510856 | Distal Intergenic |
| chr8 | 55535002 | 55536165 | Intron |
| chr8 | 55536274 | 55536738 | Intron |
| chr8 | 58871791 | 58872193 | Distal Intergenic |
| chr8 | 59029500 | 59029870 | Intron |
| chr8 | 59977445 | 59977824 | Intron |
| chr8 | 59990473 | 59991004 | Intron |
| chr8 | 60068225 | 60068592 | Distal Intergenic |
| chr8 | 60130601 | 60130899 | Distal Intergenic |
| chr8 | 60278942 | 60279175 | Distal Intergenic |
| chr8 | 63308859 | 63309984 | Intron |
| chr8 | 63320227 | 63321861 | Intron |
| chr8 | 63326869 | 63327343 | Intron |
| chr8 | 63338286 | 63338756 | Intron |
| chr8 | 63343738 | 63343948 | Intron |
| chr8 | 63432472 | 63432891 | Intron |
| chr8 | 63432930 | 63434074 | Intron |
| chr8 | 63747330 | 63748108 | Intron |
| chr8 | 63748348 | 63750481 | Intron |
| chr8 | 63817158 | 63817408 | Intron |
| chr8 | 63818556 | 63819290 | Intron |
| chr8 | 63819415 | 63821410 | Intron |
| chr8 | 64057215 | 64059591 | Distal Intergenic |
| chr8 | 64059666 | 64060710 | Distal Intergenic |
| chr8 | 64068677 | 64069244 | Distal Intergenic |
| chr8 | 64069419 | 64070039 | Distal Intergenic |
| chr8 | 64084992 | 64085464 | 5' UTR |
| chr8 | 64085918 | 64086144 | Intron |
| chr8 | 64102175 | 64103397 | Intron |
| chr8 | 64121071 | 64121427 | Intron |
| chr8 | 64143659 | 64144837 | Distal Intergenic |
| chr8 | 64176616 | 64176818 | Distal Intergenic |
| chr8 | 64176966 | 64177408 | Distal Intergenic |
| chr8 | 64261315 | 64261514 | Distal Intergenic |
| chr8 | 64261628 | 64262221 | Distal Intergenic |
| chr8 | 64265792 | 64266197 | Distal Intergenic |
| chr8 | 64267871 | 64270577 | Distal Intergenic |
| chr8 | 64274663 | 64276011 | Distal Intergenic |

| | | | |
|---|---|---|---|
| chr8 | 64281323 | 64283388 | Distal Intergenic |
| chr8 | 67001341 | 67002408 | Intron |
| chr8 | 67021133 | 67021915 | Distal Intergenic |
| chr8 | 67060981 | 67061274 | Intron |
| chr8 | 67113083 | 67113418 | Distal Intergenic |
| chr8 | 67204337 | 67206358 | Distal Intergenic |
| chr8 | 67206488 | 67206698 | Distal Intergenic |
| chr8 | 67206822 | 67207051 | Distal Intergenic |
| chr8 | 67207262 | 67207872 | Distal Intergenic |
| chr8 | 67211222 | 67212262 | Distal Intergenic |
| chr8 | 67469282 | 67469909 | Distal Intergenic |
| chr8 | 67728053 | 67728554 | Intron |
| chr8 | 69205334 | 69205939 | Distal Intergenic |
| chr8 | 69275565 | 69275773 | Intron |
| chr8 | 69281768 | 69282820 | Intron |
| chr8 | 69428887 | 69430895 | Intron |
| chr8 | 70245731 | 70247368 | Distal Intergenic |
| chr8 | 70248727 | 70249735 | Distal Intergenic |
| chr8 | 70249926 | 70250323 | Distal Intergenic |
| chr8 | 70434662 | 70434918 | Intron |
| chr8 | 70454787 | 70455369 | Intron |
| chr8 | 70460399 | 70461583 | Intron |
| chr8 | 70534743 | 70534950 | Intron |
| chr8 | 70535412 | 70535988 | Intron |
| chr8 | 70606727 | 70608120 | Intron |
| chr8 | 70680958 | 70681261 | Intron |
| chr8 | 70681358 | 70681668 | Intron |
| chr8 | 70695569 | 70696005 | Intron |
| chr8 | 70698899 | 70699102 | Intron |
| chr8 | 70701343 | 70701703 | Intron |
| chr8 | 70721465 | 70721903 | Intron |
| chr8 | 70722189 | 70722537 | Intron |
| chr8 | 70767940 | 70768256 | Distal Intergenic |
| chr8 | 70777100 | 70777553 | Distal Intergenic |
| chr8 | 70850595 | 70851124 | Distal Intergenic |
| chr8 | 71551855 | 71552268 | Intron |
| chr8 | 71561381 | 71562780 | Intron |
| chr8 | 71732950 | 71733343 | Distal Intergenic |
| chr8 | 71919063 | 71919298 | Distal Intergenic |
| chr8 | 72371959 | 72374564 | Intron |
| chr8 | 72374624 | 72375380 | Intron |
| chr8 | 72460065 | 72461005 | Distal Intergenic |
| chr8 | 72461213 | 72461516 | Distal Intergenic |
| chr8 | 72463580 | 72463861 | Distal Intergenic |

| | | | |
|---|---|---|---|
| chr8 | 72519383 | 72519775 | Distal Intergenic |
| chr8 | 72520749 | 72521222 | Distal Intergenic |
| chr8 | 72524805 | 72525661 | Distal Intergenic |
| chr8 | 72536008 | 72536910 | Distal Intergenic |
| chr8 | 72745692 | 72746112 | Distal Intergenic |
| chr8 | 72758849 | 72759070 | Intron |
| chr8 | 72759979 | 72760433 | Intron |
| chr8 | 72760626 | 72762626 | Intron |
| chr8 | 72795349 | 72797099 | Intron |
| chr8 | 72797136 | 72797335 | Intron |
| chr8 | 72876483 | 72876682 | Intron |
| chr8 | 72978185 | 72978429 | Intron |
| chr8 | 72978470 | 72978819 | Intron |
| chr8 | 73000147 | 73000641 | Distal Intergenic |
| chr8 | 73001177 | 73001437 | Distal Intergenic |
| chr8 | 73033535 | 73033769 | Distal Intergenic |
| chr8 | 73034388 | 73034803 | Distal Intergenic |
| chr8 | 73046787 | 73047007 | Distal Intergenic |
| chr8 | 73053051 | 73054386 | Distal Intergenic |
| chr8 | 73063949 | 73064202 | Distal Intergenic |
| chr8 | 73085610 | 73086280 | Distal Intergenic |
| chr8 | 73172955 | 73173744 | Distal Intergenic |
| chr8 | 73197678 | 73197881 | Distal Intergenic |
| chr8 | 73201168 | 73201373 | Distal Intergenic |
| chr8 | 73202772 | 73202990 | Distal Intergenic |
| chr8 | 73203478 | 73204123 | Distal Intergenic |
| chr8 | 73206014 | 73206238 | Distal Intergenic |
| chr8 | 73222471 | 73222670 | Distal Intergenic |
| chr8 | 73290275 | 73290558 | Distal Intergenic |
| chr8 | 73290643 | 73290940 | Distal Intergenic |
| chr8 | 73296928 | 73297481 | Distal Intergenic |
| chr8 | 73297553 | 73298279 | Distal Intergenic |
| chr8 | 73300924 | 73301151 | Distal Intergenic |
| chr8 | 73403776 | 73403982 | Distal Intergenic |
| chr8 | 73404153 | 73404770 | Distal Intergenic |
| chr8 | 73420533 | 73421768 | Distal Intergenic |
| chr8 | 73796955 | 73797683 | Intron |
| chr8 | 73800361 | 73800687 | Intron |
| chr8 | 73836562 | 73836888 | Intron |
| chr8 | 73891136 | 73891365 | Distal Intergenic |
| chr8 | 73891539 | 73892124 | Distal Intergenic |
| chr8 | 73894787 | 73895026 | Distal Intergenic |
| chr8 | 74096582 | 74097877 | Distal Intergenic |
| chr8 | 74097967 | 74098524 | Distal Intergenic |

| | | | |
|---|---|---|---|
| chr8 | 74102155 | 74102397 | Distal Intergenic |
| chr8 | 74102903 | 74103367 | Distal Intergenic |
| chr8 | 74104634 | 74105782 | Distal Intergenic |
| chr8 | 74195725 | 74195925 | Distal Intergenic |
| chr8 | 74200032 | 74200237 | Downstream |
| chr8 | 74225842 | 74227363 | Intron |
| chr8 | 74446765 | 74447510 | Intron |
| chr8 | 74659707 | 74660711 | Promoter |
| chr8 | 74668137 | 74668850 | Distal Intergenic |
| chr8 | 74669243 | 74671633 | Distal Intergenic |
| chr8 | 74685259 | 74685639 | Distal Intergenic |
| chr8 | 74931503 | 74932285 | Intron |
| chr8 | 74932342 | 74932545 | Intron |
| chr8 | 74937607 | 74937946 | Intron |
| chr8 | 74999356 | 74999567 | Distal Intergenic |
| chr8 | 74999629 | 74999852 | Distal Intergenic |
| chr8 | 75016820 | 75017232 | Distal Intergenic |
| chr8 | 75034383 | 75035666 | Distal Intergenic |
| chr8 | 75071353 | 75071634 | Distal Intergenic |
| chr8 | 75095106 | 75095735 | Distal Intergenic |
| chr8 | 75178981 | 75179395 | Intron |
| chr8 | 75179456 | 75179984 | Intron |
| chr8 | 75187770 | 75189587 | Intron |
| chr8 | 75647551 | 75647804 | Intron |
| chr8 | 75647977 | 75648872 | Intron |
| chr8 | 75799678 | 75800253 | Distal Intergenic |
| chr8 | 75805619 | 75806189 | Distal Intergenic |
| chr8 | 75808576 | 75808797 | Distal Intergenic |
| chr8 | 75808842 | 75809077 | Distal Intergenic |
| chr8 | 75813371 | 75814350 | Distal Intergenic |
| chr8 | 75814509 | 75814739 | Distal Intergenic |
| chr8 | 77617044 | 77617256 | Promoter |
| chr8 | 77617552 | 77617751 | Exon |
| chr8 | 77632715 | 77633225 | Intron |
| chr8 | 77693355 | 77694013 | Intron |
| chr8 | 78012945 | 78013144 | Distal Intergenic |
| chr8 | 79851107 | 79851323 | Distal Intergenic |
| chr8 | 79945498 | 79945939 | Distal Intergenic |
| chr8 | 79966186 | 79967572 | Distal Intergenic |
| chr8 | 79967873 | 79969013 | Distal Intergenic |
| chr8 | 79982484 | 79983169 | Distal Intergenic |
| chr8 | 80005433 | 80006584 | Distal Intergenic |
| chr8 | 80040475 | 80040975 | Distal Intergenic |
| chr8 | 80301617 | 80301849 | Distal Intergenic |

| | | | |
|---|---|---|---|
| chr8 | 80845746 | 80846071 | Intron |
| chr8 | 80846152 | 80846807 | Intron |
| chr8 | 80846857 | 80847076 | Intron |
| chr8 | 81031957 | 81033049 | Intron |
| chr8 | 81033205 | 81033690 | Intron |
| chr8 | 81033968 | 81035429 | Intron |
| chr8 | 81104784 | 81106047 | Distal Intergenic |
| chr8 | 81108524 | 81109052 | Distal Intergenic |
| chr8 | 81172180 | 81172388 | Distal Intergenic |
| chr8 | 81552414 | 81552648 | 3' UTR |
| chr8 | 81731720 | 81732289 | Intron |
| chr8 | 81732475 | 81733538 | Intron |
| chr8 | 81818245 | 81818540 | Distal Intergenic |
| chr8 | 81873645 | 81875485 | Distal Intergenic |
| chr8 | 82066121 | 82066546 | Distal Intergenic |
| chr8 | 82074376 | 82074611 | Distal Intergenic |
| chr8 | 82085525 | 82085848 | Distal Intergenic |
| chr8 | 82085895 | 82086442 | Distal Intergenic |
| chr8 | 82106821 | 82107083 | Distal Intergenic |
| chr8 | 82235134 | 82235472 | Distal Intergenic |
| chr8 | 83235016 | 83235511 | Distal Intergenic |
| chr8 | 83263915 | 83264134 | Distal Intergenic |
| chr8 | 83375890 | 83376510 | Distal Intergenic |
| chr8 | 83376677 | 83378581 | Distal Intergenic |
| chr8 | 83380261 | 83381240 | Distal Intergenic |
| chr8 | 83381489 | 83382559 | Distal Intergenic |
| chr8 | 83382697 | 83382977 | Distal Intergenic |
| chr8 | 83385472 | 83386322 | Distal Intergenic |
| chr8 | 83390850 | 83392302 | Distal Intergenic |
| chr8 | 83392473 | 83392855 | Distal Intergenic |
| chr8 | 84280532 | 84282111 | Distal Intergenic |
| chr8 | 84373547 | 84373791 | Distal Intergenic |
| chr8 | 84374316 | 84374515 | Distal Intergenic |
| chr8 | 84378189 | 84378738 | Distal Intergenic |
| chr8 | 84389556 | 84390328 | Distal Intergenic |
| chr8 | 84390888 | 84391310 | Distal Intergenic |
| chr8 | 84849612 | 84849914 | Distal Intergenic |
| chr8 | 85297450 | 85297690 | Intron |
| chr8 | 86117544 | 86117743 | Intron |
| chr8 | 86125796 | 86126909 | 3' UTR |
| chr8 | 86131076 | 86131301 | Intron |
| chr8 | 86131406 | 86131791 | Promoter |
| chr8 | 86132435 | 86133475 | Promoter |
| chr8 | 86142866 | 86144284 | Intron |

| | | | |
|---|---|---|---|
| chr8 | 86239147 | 86241093 | 3' UTR |
| chr8 | 86714791 | 86715069 | Distal Intergenic |
| chr8 | 86715108 | 86716645 | Distal Intergenic |
| chr8 | 86721064 | 86722453 | Distal Intergenic |
| chr8 | 86722494 | 86724731 | Distal Intergenic |
| chr8 | 86725074 | 86728752 | 5' UTR |
| chr8 | 86860278 | 86860529 | Distal Intergenic |
| chr8 | 87180323 | 87180747 | Distal Intergenic |
| chr8 | 87180805 | 87182303 | Distal Intergenic |
| chr8 | 87183895 | 87184292 | Distal Intergenic |
| chr8 | 87184340 | 87186014 | Distal Intergenic |
| chr8 | 87356096 | 87356390 | Intron |
| chr8 | 87433677 | 87433927 | Intron |
| chr8 | 87492506 | 87492965 | Exon |
| chr8 | 87493186 | 87493636 | Intron |
| chr8 | 87498354 | 87500181 | Exon |
| chr8 | 87500597 | 87501351 | Exon |
| chr8 | 87501501 | 87501753 | Intron |
| chr8 | 87501799 | 87502069 | Intron |
| chr8 | 87502290 | 87502574 | Intron |
| chr8 | 87503007 | 87506872 | Intron |
| chr8 | 87506971 | 87509748 | Intron |
| chr8 | 87513581 | 87513788 | Intron |
| chr8 | 87540597 | 87541134 | 5' UTR |
| chr8 | 87549178 | 87549425 | Intron |
| chr8 | 87550816 | 87551659 | Intron |
| chr8 | 87556632 | 87556867 | Intron |
| chr8 | 87556919 | 87557453 | Exon |
| chr8 | 87559483 | 87560606 | Promoter |
| chr8 | 87561106 | 87562206 | Intron |
| chr8 | 87562470 | 87562699 | Intron |
| chr8 | 87564839 | 87565054 | Intron |
| chr8 | 87565181 | 87565424 | Intron |
| chr8 | 87584596 | 87584932 | Downstream |
| chr8 | 87595309 | 87595572 | Intron |
| chr8 | 87825848 | 87826116 | Distal Intergenic |
| chr8 | 87861241 | 87862387 | Distal Intergenic |
| chr8 | 87932476 | 87932826 | Intron |
| chr8 | 87953242 | 87953516 | Intron |
| chr8 | 87953567 | 87953791 | Intron |
| chr8 | 87954723 | 87955074 | Intron |
| chr8 | 87955324 | 87955568 | Intron |
| chr8 | 88036828 | 88037959 | Intron |
| chr8 | 88038196 | 88038448 | Intron |

| | | | |
|---|---|---|---|
| chr8 | 88042845 | 88043231 | Intron |
| chr8 | 88061494 | 88062016 | Intron |
| chr8 | 88078084 | 88078802 | Intron |
| chr8 | 88967220 | 88967746 | Distal Intergenic |
| chr8 | 88969581 | 88969840 | Distal Intergenic |
| chr8 | 88978417 | 88979210 | Distal Intergenic |
| chr8 | 89254272 | 89255307 | Intron |
| chr8 | 89269090 | 89269563 | Intron |
| chr8 | 89269630 | 89269904 | Intron |
| chr8 | 89289662 | 89291262 | Intron |
| chr8 | 89291760 | 89292862 | Intron |
| chr8 | 89293012 | 89294966 | Intron |
| chr8 | 89296459 | 89297698 | Intron |
| chr8 | 89408101 | 89408406 | Distal Intergenic |
| chr8 | 90684281 | 90684560 | Distal Intergenic |
| chr8 | 90804790 | 90805614 | Distal Intergenic |
| chr8 | 90806894 | 90807130 | Distal Intergenic |
| chr8 | 90807339 | 90807541 | Distal Intergenic |
| chr8 | 90807647 | 90808990 | Distal Intergenic |
| chr8 | 91160968 | 91162836 | Distal Intergenic |
| chr8 | 91170777 | 91171692 | Distal Intergenic |
| chr8 | 91171903 | 91172641 | Distal Intergenic |
| chr8 | 91172792 | 91175192 | Distal Intergenic |
| chr8 | 91222631 | 91222973 | Distal Intergenic |
| chr8 | 93348737 | 93349930 | Distal Intergenic |
| chr8 | 93352225 | 93352650 | Distal Intergenic |
| chr8 | 93352748 | 93353214 | Distal Intergenic |
| chr8 | 93363786 | 93364487 | Distal Intergenic |
| chr8 | 93364669 | 93364871 | Distal Intergenic |
| chr8 | 93385141 | 93385725 | Distal Intergenic |
| chr8 | 93387953 | 93388749 | Distal Intergenic |
| chr8 | 93399903 | 93401030 | Distal Intergenic |
| chr8 | 93401086 | 93401485 | Distal Intergenic |
| chr8 | 93410987 | 93411278 | Distal Intergenic |
| chr8 | 93697225 | 93697567 | Distal Intergenic |
| chr8 | 93828200 | 93828603 | Distal Intergenic |
| chr8 | 93828760 | 93829072 | Distal Intergenic |
| chr8 | 93919012 | 93920165 | Intron |
| chr8 | 93920337 | 93920569 | Intron |
| chr8 | 93933924 | 93934123 | Intron |
| chr8 | 93934176 | 93934418 | Intron |
| chr8 | 93935440 | 93935969 | Intron |
| chr8 | 94012958 | 94013217 | Intron |
| chr8 | 94013270 | 94014276 | Intron |

| | | | |
|---|---|---|---|
| chr8 | 94016888 | 94017410 | Intron |
| chr8 | 94018825 | 94019129 | Intron |
| chr8 | 94824701 | 94825110 | Intron |
| chr8 | 95012897 | 95013168 | Distal Intergenic |
| chr8 | 95370416 | 95373581 | Distal Intergenic |
| chr8 | 95418673 | 95419539 | Intron |
| chr8 | 95426514 | 95426962 | Intron |
| chr8 | 95554521 | 95555166 | Intron |
| chr8 | 95893155 | 95893763 | 3' UTR |
| chr8 | 95898978 | 95899292 | Intron |
| chr8 | 96464966 | 96465303 | Intron |
| chr8 | 96505187 | 96506454 | Intron |
| chr8 | 96707639 | 96708442 | Intron |
| chr8 | 96708667 | 96709066 | Intron |
| chr8 | 96710182 | 96710554 | Intron |
| chr8 | 97501193 | 97501392 | Distal Intergenic |
| chr8 | 97506584 | 97508366 | Promoter |
| chr8 | 97516496 | 97517335 | Intron |
| chr8 | 97571223 | 97571682 | Intron |
| chr8 | 97615123 | 97615699 | Intron |
| chr8 | 97615862 | 97616087 | Intron |
| chr8 | 97699217 | 97700116 | Intron |
| chr8 | 97872476 | 97872677 | Intron |
| chr8 | 98331656 | 98331915 | Distal Intergenic |
| chr8 | 98372753 | 98373527 | Distal Intergenic |
| chr8 | 106960181 | 106960439 | Distal Intergenic |
| chr8 | 107410211 | 107410542 | Intron |
| chr8 | 107410620 | 107410852 | Intron |
| chr8 | 107426421 | 107427782 | Intron |
| chr8 | 107449589 | 107450649 | Intron |
| chr8 | 107458772 | 107459848 | Promoter |
| chr8 | 107485033 | 107486287 | Intron |
| chr8 | 107683024 | 107683827 | Intron |
| chr8 | 107683868 | 107684119 | Intron |
| chr8 | 107710265 | 107710657 | Intron |
| chr8 | 107751844 | 107752078 | Intron |
| chr8 | 107776884 | 107777308 | Intron |
| chr8 | 107785462 | 107785791 | Distal Intergenic |
| chr8 | 107788177 | 107788460 | Distal Intergenic |
| chr8 | 107821411 | 107821766 | Distal Intergenic |
| chr8 | 107827859 | 107828259 | Distal Intergenic |
| chr8 | 107828407 | 107829125 | Distal Intergenic |
| chr8 | 107829306 | 107829916 | Distal Intergenic |
| chr8 | 108117667 | 108118082 | Distal Intergenic |

| | | | |
|---|---|---|---|
| chr8 | 108160644 | 108161349 | Distal Intergenic |
| chr8 | 108161589 | 108161951 | Distal Intergenic |
| chr8 | 108313493 | 108314071 | Intron |
| chr8 | 108322843 | 108323664 | Intron |
| chr8 | 108374761 | 108375248 | Intron |
| chr8 | 108375305 | 108375736 | Intron |
| chr8 | 108384511 | 108384851 | Intron |
| chr8 | 108385593 | 108385889 | Intron |
| chr8 | 108424390 | 108425107 | Intron |
| chr8 | 108582478 | 108583322 | Distal Intergenic |
| chr8 | 108593224 | 108594341 | Distal Intergenic |
| chr8 | 108658174 | 108658436 | Distal Intergenic |
| chr8 | 108658474 | 108658693 | Distal Intergenic |
| chr8 | 108659274 | 108660030 | Distal Intergenic |
| chr8 | 108660138 | 108660533 | Distal Intergenic |
| chr8 | 108660773 | 108662504 | Distal Intergenic |
| chr8 | 108745824 | 108746177 | Distal Intergenic |
| chr8 | 109042337 | 109045144 | Intron |
| chr8 | 109050493 | 109054597 | Intron |
| chr8 | 109085232 | 109085810 | Intron |
| chr8 | 109087333 | 109088623 | Intron |
| chr8 | 109134560 | 109135070 | Distal Intergenic |
| chr8 | 109135824 | 109136023 | Distal Intergenic |
| chr8 | 109139361 | 109140669 | Distal Intergenic |
| chr8 | 109143019 | 109143279 | Distal Intergenic |
| chr8 | 109143417 | 109143616 | Distal Intergenic |
| chr8 | 109147729 | 109147973 | Distal Intergenic |
| chr8 | 109260828 | 109261062 | Promoter |
| chr8 | 109274979 | 109275485 | Distal Intergenic |
| chr8 | 109292665 | 109293923 | Distal Intergenic |
| chr8 | 109294080 | 109294480 | Distal Intergenic |
| chr8 | 109313033 | 109314365 | Distal Intergenic |
| chr8 | 109333672 | 109334567 | Distal Intergenic |
| chr8 | 109341854 | 109344262 | Distal Intergenic |
| chr8 | 109416249 | 109416927 | Distal Intergenic |
| chr8 | 109477497 | 109478113 | Intron |
| chr8 | 109674167 | 109674446 | Intron |
| chr8 | 109674702 | 109674932 | Intron |
| chr8 | 109730386 | 109730849 | Intron |
| chr8 | 109730890 | 109731240 | Intron |
| chr8 | 109778332 | 109779414 | Intron |
| chr8 | 109779991 | 109780227 | Intron |
| chr8 | 109815263 | 109815867 | Distal Intergenic |
| chr8 | 109927062 | 109927399 | Distal Intergenic |

| | | | |
|---|---|---|---|
| chr8 | 109976954 | 109977267 | Distal Intergenic |
| chr8 | 109981675 | 109981890 | Distal Intergenic |
| chr8 | 110024660 | 110028078 | Distal Intergenic |
| chr8 | 110068988 | 110069325 | Distal Intergenic |
| chr8 | 110112131 | 110112332 | Intron |
| chr8 | 110904129 | 110904328 | Distal Intergenic |
| chr8 | 111246376 | 111247460 | Distal Intergenic |
| chr8 | 111250100 | 111250367 | Distal Intergenic |
| chr8 | 111250473 | 111250881 | Distal Intergenic |
| chr8 | 111288073 | 111290281 | Distal Intergenic |
| chr8 | 112089234 | 112089714 | Distal Intergenic |
| chr8 | 112174537 | 112177021 | Distal Intergenic |
| chr8 | 112199881 | 112200758 | Distal Intergenic |
| chr8 | 112216627 | 112216943 | Distal Intergenic |
| chr8 | 112273541 | 112273959 | Distal Intergenic |
| chr8 | 112274092 | 112274725 | Distal Intergenic |
| chr8 | 112274770 | 112275191 | Distal Intergenic |
| chr8 | 112302835 | 112303809 | Distal Intergenic |
| chr8 | 112334011 | 112334214 | Distal Intergenic |
| chr8 | 112334302 | 112335326 | Distal Intergenic |
| chr8 | 112443608 | 112443916 | Distal Intergenic |
| chr8 | 112444073 | 112445183 | Distal Intergenic |
| chr8 | 112481915 | 112482235 | Distal Intergenic |
| chr8 | 113278557 | 113278960 | Intron |
| chr8 | 113318230 | 113318599 | Exon |
| chr8 | 113348961 | 113349185 | Promoter |
| chr8 | 113369978 | 113370203 | Intron |
| chr8 | 113370432 | 113371107 | Intron |
| chr8 | 113464651 | 113466073 | Intron |
| chr8 | 113557784 | 113558355 | Intron |
| chr8 | 113581143 | 113581342 | Intron |
| chr8 | 113805892 | 113806677 | Intron |
| chr8 | 113861351 | 113861581 | Intron |
| chr8 | 113861902 | 113862429 | Intron |
| chr8 | 113959161 | 113959413 | Intron |
| chr8 | 114061373 | 114061751 | Intron |
| chr8 | 114062229 | 114062428 | Intron |
| chr8 | 114123397 | 114123689 | Intron |
| chr8 | 114123872 | 114124348 | Intron |
| chr8 | 114128815 | 114129036 | Intron |
| chr8 | 114129087 | 114129425 | Intron |
| chr8 | 114250049 | 114251322 | Intron |
| chr8 | 114255437 | 114255943 | Intron |
| chr8 | 114256134 | 114256747 | Intron |

| | | | |
|---|---|---|---|
| chr8 | 114257004 | 114257528 | Intron |
| chr8 | 114269343 | 114271360 | Intron |
| chr8 | 114271548 | 114272365 | Intron |
| chr8 | 114303291 | 114303607 | Intron |
| chr8 | 114303767 | 114306390 | Intron |
| chr8 | 114429618 | 114429848 | Intron |
| chr8 | 114534645 | 114534873 | Distal Intergenic |
| chr8 | 118288822 | 118289241 | Distal Intergenic |
| chr8 | 118294667 | 118295253 | Distal Intergenic |
| chr8 | 118295372 | 118295728 | Distal Intergenic |
| chr8 | 118307167 | 118307415 | Distal Intergenic |
| chr8 | 118378608 | 118379229 | Distal Intergenic |
| chr8 | 118606379 | 118607596 | Distal Intergenic |
| chr8 | 118620358 | 118620590 | Distal Intergenic |
| chr8 | 118620889 | 118621372 | Distal Intergenic |
| chr8 | 118623655 | 118623919 | Distal Intergenic |
| chr8 | 118922217 | 118922438 | Intron |
| chr8 | 119645167 | 119646017 | Intron |
| chr8 | 119646284 | 119646483 | Intron |
| chr8 | 119646560 | 119648866 | Intron |
| chr8 | 119694127 | 119694565 | Intron |
| chr8 | 119781190 | 119781862 | Distal Intergenic |
| chr8 | 119782074 | 119783070 | Distal Intergenic |
| chr8 | 119850502 | 119853152 | Distal Intergenic |
| chr8 | 119857917 | 119858140 | Distal Intergenic |
| chr8 | 119858362 | 119860225 | Distal Intergenic |
| chr8 | 119895666 | 119895934 | Distal Intergenic |
| chr8 | 119896103 | 119896835 | Distal Intergenic |
| chr8 | 119910114 | 119910326 | Distal Intergenic |
| chr8 | 119910453 | 119910820 | Distal Intergenic |
| chr8 | 119910967 | 119911502 | Distal Intergenic |
| chr8 | 120085142 | 120085767 | Intron |
| chr8 | 120243043 | 120243383 | Intron |
| chr8 | 120310606 | 120311391 | Distal Intergenic |
| chr8 | 120810362 | 120811361 | Intron |
| chr8 | 121945305 | 121947213 | Distal Intergenic |
| chr8 | 121952171 | 121952431 | Distal Intergenic |
| chr8 | 121954774 | 121954973 | Distal Intergenic |
| chr8 | 122210669 | 122211133 | Distal Intergenic |
| chr8 | 122211298 | 122212376 | Distal Intergenic |
| chr8 | 122232492 | 122233191 | Distal Intergenic |
| chr8 | 122234255 | 122234481 | Distal Intergenic |
| chr8 | 122335182 | 122335619 | Distal Intergenic |
| chr8 | 122335768 | 122337475 | Distal Intergenic |

| | | | |
|---|---|---|---|
| chr8 | 122338308 | 122338520 | Distal Intergenic |
| chr8 | 122339743 | 122339959 | Distal Intergenic |
| chr8 | 122340116 | 122342300 | Distal Intergenic |
| chr8 | 122352798 | 122354047 | Distal Intergenic |
| chr8 | 122471867 | 122472535 | Distal Intergenic |
| chr8 | 122472737 | 122473065 | Distal Intergenic |
| chr8 | 122473104 | 122473303 | Distal Intergenic |
| chr8 | 122645032 | 122645477 | Intron |
| chr8 | 122645854 | 122646216 | Intron |
| chr8 | 123062308 | 123062547 | Intron |
| chr8 | 123087155 | 123088053 | Intron |
| chr8 | 123806884 | 123807404 | Intron |
| chr8 | 124192057 | 124192418 | Promoter |
| chr8 | 124192563 | 124192922 | Exon |
| chr8 | 124193125 | 124193501 | Promoter |
| chr8 | 124385747 | 124386039 | Intron |
| chr8 | 124386099 | 124386303 | Intron |
| chr8 | 124422159 | 124422409 | Intron |
| chr8 | 124494775 | 124495011 | Distal Intergenic |
| chr8 | 124528051 | 124528390 | Intron |
| chr8 | 124542806 | 124543535 | Promoter |
| chr8 | 124543585 | 124544881 | Promoter |
| chr8 | 124867064 | 124867283 | Intron |
| chr8 | 124867452 | 124867890 | Intron |
| chr8 | 124985356 | 124985706 | Exon |
| chr8 | 125047005 | 125047232 | Intron |
| chr8 | 125047283 | 125048411 | Exon |
| chr8 | 125050134 | 125050743 | Intron |
| chr8 | 125060691 | 125061614 | Intron |
| chr8 | 125072617 | 125072899 | Exon |
| chr8 | 125090935 | 125091222 | Intron |
| chr8 | 125091277 | 125091677 | Intron |
| chr8 | 125092274 | 125092497 | Intron |
| chr8 | 125099193 | 125100747 | Intron |
| chr8 | 125109116 | 125109818 | Exon |
| chr8 | 125429117 | 125430876 | Distal Intergenic |
| chr8 | 125456084 | 125456320 | Distal Intergenic |
| chr8 | 125457845 | 125458082 | Distal Intergenic |
| chr8 | 125522083 | 125522295 | Intron |
| chr8 | 125544813 | 125547486 | Intron |
| chr8 | 125625241 | 125625454 | Intron |
| chr8 | 125627614 | 125627813 | Intron |
| chr8 | 125937168 | 125937768 | Exon |
| chr8 | 125970978 | 125971430 | Distal Intergenic |

| | | | |
|---|---|---|---|
| chr8 | 126407851 | 126408271 | Distal Intergenic |
| chr8 | 126612299 | 126612543 | Distal Intergenic |
| chr8 | 126789224 | 126789470 | Distal Intergenic |
| chr8 | 126789560 | 126789846 | Distal Intergenic |
| chr8 | 126794633 | 126795088 | Distal Intergenic |
| chr8 | 126801489 | 126803163 | Distal Intergenic |
| chr8 | 126854884 | 126855112 | Distal Intergenic |
| chr8 | 126871764 | 126872122 | Distal Intergenic |
| chr8 | 127167920 | 127168125 | Distal Intergenic |
| chr8 | 127168161 | 127168388 | Distal Intergenic |
| chr8 | 127777368 | 127777771 | Distal Intergenic |
| chr8 | 128498326 | 128498563 | Distal Intergenic |
| chr8 | 128519990 | 128520742 | Distal Intergenic |
| chr8 | 129046705 | 129046914 | Intron |
| chr8 | 129046960 | 129047854 | Intron |
| chr8 | 129055141 | 129055552 | Intron |
| chr8 | 129055860 | 129056372 | Intron |
| chr8 | 129063141 | 129063541 | Intron |
| chr8 | 129063757 | 129064481 | Intron |
| chr8 | 129094981 | 129095305 | Intron |
| chr8 | 129113718 | 129114085 | Distal Intergenic |
| chr8 | 129115837 | 129116142 | Distal Intergenic |
| chr8 | 129119041 | 129119262 | Distal Intergenic |
| chr8 | 129119448 | 129120068 | Distal Intergenic |
| chr8 | 129120141 | 129120685 | Distal Intergenic |
| chr8 | 129127271 | 129127522 | Distal Intergenic |
| chr8 | 129137841 | 129139004 | Distal Intergenic |
| chr8 | 129139112 | 129140155 | Distal Intergenic |
| chr8 | 129196622 | 129196950 | Distal Intergenic |
| chr8 | 129208891 | 129209258 | Distal Intergenic |
| chr8 | 129209909 | 129210827 | Distal Intergenic |
| chr8 | 129211070 | 129211884 | Distal Intergenic |
| chr8 | 129363126 | 129363386 | Distal Intergenic |
| chr8 | 129376065 | 129376927 | Distal Intergenic |
| chr8 | 129495102 | 129497034 | Distal Intergenic |
| chr8 | 129552544 | 129553090 | Distal Intergenic |
| chr8 | 129626946 | 129627425 | Distal Intergenic |
| chr8 | 129633250 | 129633669 | Distal Intergenic |
| chr8 | 129633738 | 129634186 | Distal Intergenic |
| chr8 | 129637129 | 129637328 | Distal Intergenic |
| chr8 | 129685890 | 129686616 | Distal Intergenic |
| chr8 | 129750470 | 129751816 | Distal Intergenic |
| chr8 | 129751854 | 129752956 | Distal Intergenic |
| chr8 | 130636952 | 130637405 | Distal Intergenic |

| | | | |
|---|---|---|---|
| chr8 | 130689279 | 130689750 | Distal Intergenic |
| chr8 | 130743280 | 130743675 | Distal Intergenic |
| chr8 | 131143226 | 131143507 | Intron |
| chr8 | 131178566 | 131179014 | Intron |
| chr8 | 131190547 | 131190848 | Intron |
| chr8 | 131191110 | 131191372 | Intron |
| chr8 | 131209041 | 131209374 | Intron |
| chr8 | 131209600 | 131210665 | Intron |
| chr8 | 131210792 | 131211139 | Intron |
| chr8 | 131214341 | 131214540 | Intron |
| chr8 | 131221820 | 131222075 | Intron |
| chr8 | 131226407 | 131226733 | Intron |
| chr8 | 131226772 | 131228060 | Exon |
| chr8 | 131228265 | 131229384 | Intron |
| chr9 | 3403415 | 3404401 | Intron |
| chr9 | 4309054 | 4309433 | Distal Intergenic |
| chr9 | 5309100 | 5309427 | Distal Intergenic |
| chr9 | 6374019 | 6374247 | Distal Intergenic |
| chr9 | 7183829 | 7184489 | Distal Intergenic |
| chr9 | 7387623 | 7388323 | Distal Intergenic |
| chr9 | 7510003 | 7510478 | Distal Intergenic |
| chr9 | 7835902 | 7837020 | Distal Intergenic |
| chr9 | 8003101 | 8003504 | Distal Intergenic |
| chr9 | 8003613 | 8003974 | Distal Intergenic |
| chr9 | 8004177 | 8004549 | Distal Intergenic |
| chr9 | 8004907 | 8005121 | Distal Intergenic |
| chr9 | 8103677 | 8103907 | Distal Intergenic |
| chr9 | 8104067 | 8104393 | Distal Intergenic |
| chr9 | 8104431 | 8104715 | Distal Intergenic |
| chr9 | 8113777 | 8114884 | Distal Intergenic |
| chr9 | 9120380 | 9120579 | Intron |
| chr9 | 9120655 | 9120871 | Intron |
| chr9 | 9171323 | 9171785 | Intron |
| chr9 | 9207408 | 9208074 | Intron |
| chr9 | 9241131 | 9241360 | Intron |
| chr9 | 9241428 | 9242423 | Intron |
| chr9 | 9245764 | 9246406 | Intron |
| chr9 | 11380235 | 11381220 | Distal Intergenic |
| chr9 | 11570268 | 11571375 | Distal Intergenic |
| chr9 | 12016690 | 12017090 | Distal Intergenic |
| chr9 | 12017208 | 12017434 | Distal Intergenic |
| chr9 | 13228486 | 13228721 | Intron |
| chr9 | 13245194 | 13246020 | Intron |
| chr9 | 13258490 | 13260273 | Intron |

| | | | |
|---|---|---|---|
| chr9 | 13298791 | 13299024 | Distal Intergenic |
| chr9 | 13299887 | 13301645 | Distal Intergenic |
| chr9 | 13326965 | 13327867 | Distal Intergenic |
| chr9 | 13329784 | 13330533 | Distal Intergenic |
| chr9 | 13365021 | 13365459 | Distal Intergenic |
| chr9 | 13385264 | 13386109 | Distal Intergenic |
| chr9 | 13464843 | 13465885 | Distal Intergenic |
| chr9 | 13468541 | 13469456 | Distal Intergenic |
| chr9 | 13469532 | 13470086 | Distal Intergenic |
| chr9 | 13473235 | 13473499 | Distal Intergenic |
| chr9 | 13553885 | 13554256 | Distal Intergenic |
| chr9 | 13632195 | 13632439 | Distal Intergenic |
| chr9 | 13632712 | 13633038 | Distal Intergenic |
| chr9 | 13776081 | 13776722 | Distal Intergenic |
| chr9 | 13777576 | 13778259 | Distal Intergenic |
| chr9 | 13778415 | 13778641 | Distal Intergenic |
| chr9 | 13858040 | 13858390 | Distal Intergenic |
| chr9 | 13961802 | 13962018 | Distal Intergenic |
| chr9 | 14019980 | 14020585 | Distal Intergenic |
| chr9 | 14169750 | 14169964 | Intron |
| chr9 | 14170091 | 14171631 | Intron |
| chr9 | 14171697 | 14172995 | Intron |
| chr9 | 14183919 | 14184167 | Intron |
| chr9 | 14184498 | 14184783 | Intron |
| chr9 | 14325466 | 14326944 | Intron |
| chr9 | 14339683 | 14339998 | Intron |
| chr9 | 14362220 | 14362653 | Intron |
| chr9 | 14363695 | 14364117 | Intron |
| chr9 | 14364170 | 14364638 | Intron |
| chr9 | 14372785 | 14373323 | Intron |
| chr9 | 14373536 | 14373779 | Intron |
| chr9 | 14393126 | 14393489 | Intron |
| chr9 | 14408912 | 14409264 | Distal Intergenic |
| chr9 | 14417345 | 14420123 | Distal Intergenic |
| chr9 | 14422410 | 14422713 | Distal Intergenic |
| chr9 | 14465940 | 14468233 | Distal Intergenic |
| chr9 | 14480322 | 14480817 | Distal Intergenic |
| chr9 | 14588684 | 14589022 | Intron |
| chr9 | 14589127 | 14589328 | Intron |
| chr9 | 14834226 | 14834627 | Intron |
| chr9 | 14849040 | 14849352 | Intron |
| chr9 | 14873160 | 14874270 | Intron |
| chr9 | 14923636 | 14924133 | Distal Intergenic |
| chr9 | 14924381 | 14924698 | Distal Intergenic |

| | | | |
|---|---|---|---|
| chr9 | 14926287 | 14927510 | Distal Intergenic |
| chr9 | 14942921 | 14943431 | Distal Intergenic |
| chr9 | 14949895 | 14951047 | Distal Intergenic |
| chr9 | 15042377 | 15042792 | Intron |
| chr9 | 15110796 | 15111015 | Intron |
| chr9 | 15111069 | 15111320 | Intron |
| chr9 | 15111368 | 15112744 | Intron |
| chr9 | 15113726 | 15114074 | Intron |
| chr9 | 15161591 | 15162248 | Distal Intergenic |
| chr9 | 15531954 | 15532193 | Distal Intergenic |
| chr9 | 15539156 | 15539519 | Distal Intergenic |
| chr9 | 15551102 | 15551920 | Promoter |
| chr9 | 17362046 | 17362901 | Intron |
| chr9 | 18096347 | 18097781 | Distal Intergenic |
| chr9 | 19260096 | 19260916 | Intron |
| chr9 | 19261025 | 19261257 | Intron |
| chr9 | 19426532 | 19426959 | Intron |
| chr9 | 20263993 | 20264820 | Distal Intergenic |
| chr9 | 20296791 | 20297243 | Distal Intergenic |
| chr9 | 20297312 | 20297844 | Distal Intergenic |
| chr9 | 20325072 | 20325271 | Distal Intergenic |
| chr9 | 20325664 | 20325873 | Distal Intergenic |
| chr9 | 20411872 | 20412071 | Promoter |
| chr9 | 20447016 | 20447495 | Intron |
| chr9 | 20447675 | 20448228 | Exon |
| chr9 | 20449024 | 20449237 | Intron |
| chr9 | 20449358 | 20449621 | Intron |
| chr9 | 20466169 | 20466399 | Intron |
| chr9 | 20466465 | 20466915 | Intron |
| chr9 | 20532463 | 20533707 | Intron |
| chr9 | 20692659 | 20693089 | Intron |
| chr9 | 20702242 | 20702447 | Intron |
| chr9 | 20702561 | 20702760 | Intron |
| chr9 | 20748893 | 20749625 | Intron |
| chr9 | 20756257 | 20756746 | Intron |
| chr9 | 20877859 | 20878375 | Intron |
| chr9 | 21029542 | 21030033 | Intron |
| chr9 | 21042790 | 21043098 | Distal Intergenic |
| chr9 | 21055396 | 21055925 | Distal Intergenic |
| chr9 | 21062001 | 21062217 | Distal Intergenic |
| chr9 | 21068712 | 21069636 | Distal Intergenic |
| chr9 | 21092066 | 21092366 | Distal Intergenic |
| chr9 | 21136959 | 21138537 | Downstream |
| chr9 | 21141877 | 21142076 | Promoter |

| | | | |
|---|---|---|---|
| chr9 | 21172758 | 21173181 | Distal Intergenic |
| chr9 | 21229307 | 21229779 | Intron |
| chr9 | 21298779 | 21299504 | Distal Intergenic |
| chr9 | 21351824 | 21352108 | Promoter |
| chr9 | 21420056 | 21420352 | Distal Intergenic |
| chr9 | 21420702 | 21421024 | Distal Intergenic |
| chr9 | 21421108 | 21421338 | Distal Intergenic |
| chr9 | 21421503 | 21422207 | Distal Intergenic |
| chr9 | 21474518 | 21474730 | Intron |
| chr9 | 21484396 | 21485398 | Intron |
| chr9 | 21491042 | 21491279 | Intron |
| chr9 | 21491336 | 21491842 | Intron |
| chr9 | 21532831 | 21533920 | Intron |
| chr9 | 21596294 | 21596493 | Distal Intergenic |
| chr9 | 21728115 | 21728429 | Distal Intergenic |
| chr9 | 21728619 | 21729065 | Distal Intergenic |
| chr9 | 22029772 | 22030026 | Intron |
| chr9 | 22030448 | 22032004 | Intron |
| chr9 | 22190357 | 22190578 | Distal Intergenic |
| chr9 | 22190785 | 22191227 | Distal Intergenic |
| chr9 | 22192523 | 22193281 | Distal Intergenic |
| chr9 | 22193352 | 22195534 | Distal Intergenic |
| chr9 | 22210106 | 22210328 | Distal Intergenic |
| chr9 | 23498237 | 23500191 | Distal Intergenic |
| chr9 | 23503381 | 23504133 | Intron |
| chr9 | 24747311 | 24747634 | Distal Intergenic |
| chr9 | 24748095 | 24748477 | Distal Intergenic |
| chr9 | 24748596 | 24749043 | Distal Intergenic |
| chr9 | 24847023 | 24847384 | Distal Intergenic |
| chr9 | 24955880 | 24956736 | Distal Intergenic |
| chr9 | 24957271 | 24957774 | Distal Intergenic |
| chr9 | 25051494 | 25052006 | Distal Intergenic |
| chr9 | 25058957 | 25059932 | Distal Intergenic |
| chr9 | 25060295 | 25061032 | Distal Intergenic |
| chr9 | 25061123 | 25061323 | Distal Intergenic |
| chr9 | 25074371 | 25075085 | Distal Intergenic |
| chr9 | 25399189 | 25400132 | Exon |
| chr9 | 25400283 | 25400667 | Distal Intergenic |
| chr9 | 26806881 | 26807166 | Distal Intergenic |
| chr9 | 26807284 | 26807729 | Distal Intergenic |
| chr9 | 26834561 | 26834803 | Distal Intergenic |
| chr9 | 28716075 | 28716303 | Intron |
| chr9 | 28797652 | 28797925 | Intron |
| chr9 | 28800252 | 28801408 | Intron |

| | | | |
|---|---|---|---|
| chr9 | 28804692 | 28809082 | Intron |
| chr9 | 28811313 | 28813132 | Intron |
| chr9 | 28816154 | 28816448 | Intron |
| chr9 | 28867024 | 28869821 | Intron |
| chr9 | 28875134 | 28875551 | Intron |
| chr9 | 29046668 | 29047243 | Intron |
| chr9 | 29170980 | 29171605 | Intron |
| chr9 | 29301427 | 29302066 | Distal Intergenic |
| chr9 | 30585099 | 30586277 | Distal Intergenic |
| chr9 | 30586521 | 30586804 | Distal Intergenic |
| chr9 | 30587826 | 30591650 | Distal Intergenic |
| chr9 | 30600786 | 30603062 | Distal Intergenic |
| chr9 | 30838227 | 30839054 | Distal Intergenic |
| chr9 | 31010765 | 31012231 | Distal Intergenic |
| chr9 | 31012346 | 31012852 | Distal Intergenic |
| chr9 | 31025652 | 31025851 | Distal Intergenic |
| chr9 | 31050028 | 31051748 | Distal Intergenic |
| chr9 | 31072153 | 31073174 | Distal Intergenic |
| chr9 | 31073245 | 31074157 | Distal Intergenic |
| chr9 | 31087329 | 31088715 | Distal Intergenic |
| chr9 | 31088772 | 31089430 | Distal Intergenic |
| chr9 | 31126400 | 31127402 | Distal Intergenic |
| chr9 | 31127979 | 31130096 | Distal Intergenic |
| chr9 | 31306612 | 31307364 | Distal Intergenic |
| chr9 | 31315070 | 31315286 | Distal Intergenic |
| chr9 | 31315328 | 31316330 | Distal Intergenic |
| chr9 | 31421613 | 31422072 | Distal Intergenic |
| chr9 | 31423255 | 31423539 | Distal Intergenic |
| chr9 | 31458118 | 31458902 | Distal Intergenic |
| chr9 | 31462066 | 31462502 | Distal Intergenic |
| chr9 | 31663703 | 31664576 | Distal Intergenic |
| chr9 | 31805092 | 31806649 | Distal Intergenic |
| chr9 | 31806711 | 31807000 | Distal Intergenic |
| chr9 | 31818489 | 31819421 | Distal Intergenic |
| chr9 | 31863333 | 31864304 | Distal Intergenic |
| chr9 | 31864830 | 31865965 | Distal Intergenic |
| chr9 | 31871092 | 31871299 | Distal Intergenic |
| chr9 | 31923320 | 31924511 | Distal Intergenic |
| chr9 | 31924582 | 31927166 | Distal Intergenic |
| chr9 | 31927203 | 31929180 | Distal Intergenic |
| chr9 | 31966876 | 31967814 | Distal Intergenic |
| chr9 | 31967899 | 31969733 | Distal Intergenic |
| chr9 | 32031224 | 32032264 | Distal Intergenic |
| chr9 | 32032353 | 32032665 | Distal Intergenic |

| | | | |
|---|---|---|---|
| chr9 | 32436742 | 32436979 | Intron |
| chr9 | 32503073 | 32503480 | Promoter |
| chr9 | 32504324 | 32504672 | Intron |
| chr9 | 32512990 | 32513859 | Intron |
| chr9 | 32515797 | 32516224 | Intron |
| chr9 | 32516320 | 32519063 | Intron |
| chr9 | 32521763 | 32522668 | Intron |
| chr9 | 32557279 | 32557967 | Intron |
| chr9 | 32788678 | 32789110 | 3' UTR |
| chr9 | 33666452 | 33666783 | Distal Intergenic |
| chr9 | 34087080 | 34087634 | 3' UTR |
| chr9 | 34087680 | 34088081 | 3' UTR |
| chr9 | 34869089 | 34870000 | Distal Intergenic |
| chr9 | 34924614 | 34926584 | Distal Intergenic |
| chr9 | 34946930 | 34947187 | Distal Intergenic |
| chr9 | 35000878 | 35001386 | Distal Intergenic |
| chr9 | 35006790 | 35007099 | Distal Intergenic |
| chr9 | 35018106 | 35018620 | Distal Intergenic |
| chr9 | 35019111 | 35019481 | Distal Intergenic |
| chr9 | 35055362 | 35056081 | 3' UTR |
| chr9 | 35367093 | 35367320 | Intron |
| chr9 | 36533908 | 36534177 | Distal Intergenic |
| chr9 | 36534269 | 36534599 | Distal Intergenic |
| chr9 | 36535142 | 36535518 | Distal Intergenic |
| chr9 | 36542702 | 36543680 | Distal Intergenic |
| chr9 | 36544274 | 36544554 | Distal Intergenic |
| chr9 | 36603961 | 36604629 | 5' UTR |
| chr9 | 37043217 | 37043546 | Distal Intergenic |
| chr9 | 37155544 | 37155974 | Intron |
| chr9 | 37226183 | 37227019 | Intron |
| chr9 | 37295971 | 37296829 | Intron |
| chr9 | 37463781 | 37465251 | Promoter |
| chr9 | 37541267 | 37542508 | 5' UTR |
| chr9 | 37544060 | 37544377 | Intron |
| chr9 | 37641136 | 37641335 | Distal Intergenic |
| chr9 | 38525849 | 38526089 | Distal Intergenic |
| chr9 | 38526142 | 38526940 | Distal Intergenic |
| chr9 | 40554760 | 40555475 | Intron |
| chr9 | 40555542 | 40556305 | Intron |
| chr9 | 40573186 | 40576550 | Intron |
| chr9 | 40587871 | 40591057 | Intron |
| chr9 | 40602446 | 40604537 | Intron |
| chr9 | 40606416 | 40606638 | Intron |
| chr9 | 40607033 | 40608909 | Intron |

| | | | |
|---|---|---|---|
| chr9 | 40609335 | 40609544 | Intron |
| chr9 | 40609725 | 40609993 | Intron |
| chr9 | 40610132 | 40610373 | 5' UTR |
| chr9 | 40614518 | 40615564 | Intron |
| chr9 | 40615612 | 40615826 | Intron |
| chr9 | 40829481 | 40829715 | Intron |
| chr9 | 41288597 | 41288892 | Distal Intergenic |
| chr9 | 41296137 | 41296538 | Distal Intergenic |
| chr9 | 41316935 | 41317210 | Distal Intergenic |
| chr9 | 41385648 | 41386060 | Distal Intergenic |
| chr9 | 41387761 | 41387985 | Distal Intergenic |
| chr9 | 41388460 | 41388733 | Distal Intergenic |
| chr9 | 41388807 | 41390031 | Distal Intergenic |
| chr9 | 41390136 | 41392806 | Distal Intergenic |
| chr9 | 41392860 | 41394228 | Distal Intergenic |
| chr9 | 41395387 | 41395774 | Distal Intergenic |
| chr9 | 41395886 | 41397206 | Distal Intergenic |
| chr9 | 41543092 | 41544105 | Distal Intergenic |
| chr9 | 41640516 | 41640743 | Distal Intergenic |
| chr9 | 41666515 | 41667089 | Distal Intergenic |
| chr9 | 41816526 | 41817231 | Distal Intergenic |
| chr9 | 41901876 | 41903082 | Distal Intergenic |
| chr9 | 41916070 | 41918737 | Distal Intergenic |
| chr9 | 41988382 | 41988581 | Intron |
| chr9 | 42060142 | 42060726 | Distal Intergenic |
| chr9 | 42060873 | 42061137 | Distal Intergenic |
| chr9 | 42071324 | 42072387 | Distal Intergenic |
| chr9 | 42072722 | 42073122 | Distal Intergenic |
| chr9 | 42073174 | 42073373 | Distal Intergenic |
| chr9 | 42080923 | 42081893 | Distal Intergenic |
| chr9 | 42081961 | 42082182 | Distal Intergenic |
| chr9 | 42269494 | 42269975 | Distal Intergenic |
| chr9 | 42279744 | 42279969 | Distal Intergenic |
| chr9 | 42811097 | 42811646 | Distal Intergenic |
| chr9 | 42811736 | 42812138 | Distal Intergenic |
| chr9 | 42818944 | 42819796 | Distal Intergenic |
| chr9 | 42819869 | 42820068 | Distal Intergenic |
| chr9 | 42822807 | 42824717 | Distal Intergenic |
| chr9 | 42825050 | 42825371 | Distal Intergenic |
| chr9 | 42842422 | 42842679 | Downstream |
| chr9 | 42842830 | 42844143 | Downstream |
| chr9 | 42869624 | 42870612 | Intron |
| chr9 | 42882509 | 42883597 | Exon |
| chr9 | 42906021 | 42907310 | Distal Intergenic |

| | | | |
|---|---|---|---|
| chr9 | 42910089 | 42910916 | Distal Intergenic |
| chr9 | 42913144 | 42913726 | Distal Intergenic |
| chr9 | 43093289 | 43093832 | Exon |
| chr9 | 43108377 | 43109111 | Intron |
| chr9 | 43116678 | 43116984 | Intron |
| chr9 | 43117114 | 43117389 | Intron |
| chr9 | 43117474 | 43117983 | Intron |
| chr9 | 43118299 | 43118541 | Intron |
| chr9 | 43119827 | 43120056 | Intron |
| chr9 | 43120157 | 43121692 | Exon |
| chr9 | 43122146 | 43122544 | Intron |
| chr9 | 43122693 | 43122945 | Intron |
| chr9 | 43123069 | 43123994 | Intron |
| chr9 | 43124087 | 43125138 | Exon |
| chr9 | 43125354 | 43125589 | Exon |
| chr9 | 43126032 | 43127788 | Intron |
| chr9 | 43133509 | 43135234 | Promoter |
| chr9 | 43165406 | 43167782 | Distal Intergenic |
| chr9 | 43349871 | 43350804 | Distal Intergenic |
| chr9 | 43476313 | 43477495 | Distal Intergenic |
| chr9 | 43481355 | 43482604 | Distal Intergenic |
| chr9 | 43482643 | 43483210 | Distal Intergenic |
| chr9 | 43487469 | 43490660 | Distal Intergenic |
| chr9 | 43527559 | 43529549 | Distal Intergenic |
| chr9 | 43552913 | 43554520 | Distal Intergenic |
| chr9 | 43554778 | 43555147 | Distal Intergenic |
| chr9 | 43557410 | 43558335 | Distal Intergenic |
| chr9 | 43558497 | 43559171 | Distal Intergenic |
| chr9 | 43561288 | 43561504 | Distal Intergenic |
| chr9 | 43563869 | 43564213 | Distal Intergenic |
| chr9 | 43564907 | 43565198 | Distal Intergenic |
| chr9 | 43565246 | 43566628 | Distal Intergenic |
| chr9 | 43579611 | 43582984 | Distal Intergenic |
| chr9 | 43583040 | 43583796 | Distal Intergenic |
| chr9 | 43583837 | 43584909 | Distal Intergenic |
| chr9 | 43585008 | 43586823 | Distal Intergenic |
| chr9 | 43590103 | 43590400 | Distal Intergenic |
| chr9 | 43591142 | 43591547 | Distal Intergenic |
| chr9 | 43591596 | 43592366 | Distal Intergenic |
| chr9 | 43592486 | 43593788 | Distal Intergenic |
| chr9 | 43595027 | 43597333 | Distal Intergenic |
| chr9 | 43603763 | 43604631 | Distal Intergenic |
| chr9 | 43604794 | 43605750 | Distal Intergenic |
| chr9 | 43607874 | 43608582 | Distal Intergenic |

| | | | |
|---|---|---|---|
| chr9 | 43609058 | 43610341 | Distal Intergenic |
| chr9 | 43658932 | 43659367 | Distal Intergenic |
| chr9 | 43665666 | 43667124 | Distal Intergenic |
| chr9 | 43681928 | 43682954 | Distal Intergenic |
| chr9 | 43848514 | 43848963 | Intron |
| chr9 | 43920997 | 43921231 | 3' UTR |
| chr9 | 43921313 | 43921585 | 3' UTR |
| chr9 | 43943362 | 43944606 | Distal Intergenic |
| chr9 | 43948585 | 43948927 | Distal Intergenic |
| chr9 | 44006035 | 44006354 | Distal Intergenic |
| chr9 | 44006491 | 44006818 | Distal Intergenic |
| chr9 | 44007064 | 44007302 | Distal Intergenic |
| chr9 | 44007358 | 44008068 | Distal Intergenic |
| chr9 | 44008256 | 44008459 | Distal Intergenic |
| chr9 | 44012819 | 44013187 | Distal Intergenic |
| chr9 | 44015546 | 44016256 | Distal Intergenic |
| chr9 | 44040793 | 44041612 | Distal Intergenic |
| chr9 | 44075907 | 44076214 | Distal Intergenic |
| chr9 | 44076415 | 44077162 | Distal Intergenic |
| chr9 | 44077942 | 44078181 | Distal Intergenic |
| chr9 | 44134311 | 44134854 | Distal Intergenic |
| chr9 | 44141926 | 44142553 | Distal Intergenic |
| chr9 | 44142873 | 44144415 | Distal Intergenic |
| chr9 | 44151631 | 44151833 | Distal Intergenic |
| chr9 | 44214805 | 44215034 | Distal Intergenic |
| chr9 | 44215086 | 44215388 | Distal Intergenic |
| chr9 | 44215435 | 44216495 | Distal Intergenic |
| chr9 | 44286129 | 44287742 | Distal Intergenic |
| chr9 | 44288378 | 44288849 | Distal Intergenic |
| chr9 | 44290545 | 44291027 | Distal Intergenic |
| chr9 | 44291505 | 44292496 | Distal Intergenic |
| chr9 | 44292833 | 44293415 | Distal Intergenic |
| chr9 | 44293563 | 44294611 | Distal Intergenic |
| chr9 | 44294699 | 44297646 | Distal Intergenic |
| chr9 | 44302178 | 44304184 | Distal Intergenic |
| chr9 | 44305142 | 44305997 | Distal Intergenic |
| chr9 | 44306059 | 44306887 | Distal Intergenic |
| chr9 | 44307264 | 44307861 | Distal Intergenic |
| chr9 | 44308264 | 44308576 | Distal Intergenic |
| chr9 | 44309803 | 44310057 | Distal Intergenic |
| chr9 | 44407306 | 44407518 | Distal Intergenic |
| chr9 | 44411730 | 44412272 | Distal Intergenic |
| chr9 | 44412425 | 44412637 | Distal Intergenic |
| chr9 | 44413126 | 44414036 | Distal Intergenic |

| | | | |
|---|---|---|---|
| chr9 | 44524625 | 44525579 | Distal Intergenic |
| chr9 | 44527075 | 44528049 | Distal Intergenic |
| chr9 | 44528232 | 44528496 | Distal Intergenic |
| chr9 | 44728671 | 44729597 | Distal Intergenic |
| chr9 | 44773311 | 44773705 | Distal Intergenic |
| chr9 | 44773826 | 44774241 | Distal Intergenic |
| chr9 | 44835470 | 44836470 | Distal Intergenic |
| chr9 | 44850216 | 44851442 | Exon |
| chr9 | 44854495 | 44854948 | Distal Intergenic |
| chr9 | 44863172 | 44863432 | Distal Intergenic |
| chr9 | 44863615 | 44864129 | Distal Intergenic |
| chr9 | 44867700 | 44869358 | Distal Intergenic |
| chr9 | 45008276 | 45008798 | Distal Intergenic |
| chr9 | 45008856 | 45009067 | Distal Intergenic |
| chr9 | 45012149 | 45012705 | Distal Intergenic |
| chr9 | 45012906 | 45013195 | Distal Intergenic |
| chr9 | 45013252 | 45013536 | Distal Intergenic |
| chr9 | 45017842 | 45018278 | Distal Intergenic |
| chr9 | 45145328 | 45145854 | Distal Intergenic |
| chr9 | 45636740 | 45636970 | Distal Intergenic |
| chr9 | 45653945 | 45654843 | Distal Intergenic |
| chr9 | 45654953 | 45655308 | Distal Intergenic |
| chr9 | 45762252 | 45762451 | Distal Intergenic |
| chr9 | 45762490 | 45763288 | Distal Intergenic |
| chr9 | 45791832 | 45792533 | Distal Intergenic |
| chr9 | 45792834 | 45793328 | Distal Intergenic |
| chr9 | 45854869 | 45855819 | Distal Intergenic |
| chr9 | 45856896 | 45857110 | Distal Intergenic |
| chr9 | 45876792 | 45877009 | Distal Intergenic |
| chr9 | 45877102 | 45877848 | Distal Intergenic |
| chr9 | 45891402 | 45891895 | Distal Intergenic |
| chr9 | 45928313 | 45929572 | Distal Intergenic |
| chr9 | 46090824 | 46091023 | Distal Intergenic |
| chr9 | 46478212 | 46478797 | Distal Intergenic |
| chr9 | 46667637 | 46668744 | Distal Intergenic |
| chr9 | 46769198 | 46769493 | Distal Intergenic |
| chr9 | 46769700 | 46769962 | Distal Intergenic |
| chr9 | 46830015 | 46831206 | Promoter |
| chr9 | 46831311 | 46831622 | Intron |
| chr9 | 46870073 | 46870881 | Distal Intergenic |
| chr9 | 47170423 | 47171330 | Distal Intergenic |
| chr9 | 47360833 | 47361170 | Distal Intergenic |
| chr9 | 47361227 | 47361711 | Distal Intergenic |
| chr9 | 47397111 | 47398951 | Distal Intergenic |

| | | | |
|---|---|---|---|
| chr9 | 47427058 | 47428079 | Distal Intergenic |
| chr9 | 47433840 | 47434132 | Distal Intergenic |
| chr9 | 47510737 | 47511201 | Distal Intergenic |
| chr9 | 47511705 | 47512267 | Distal Intergenic |
| chr9 | 48215836 | 48216543 | Distal Intergenic |
| chr9 | 48302892 | 48303376 | Distal Intergenic |
| chr9 | 49234181 | 49235810 | Distal Intergenic |
| chr9 | 49622106 | 49622428 | Distal Intergenic |
| chr9 | 49692603 | 49693002 | Distal Intergenic |
| chr9 | 49707317 | 49707534 | Distal Intergenic |
| chr9 | 49731512 | 49732419 | Distal Intergenic |
| chr9 | 49732562 | 49732807 | Distal Intergenic |
| chr9 | 49780572 | 49781553 | Distal Intergenic |
| chr9 | 49783964 | 49784948 | Distal Intergenic |
| chr9 | 49788740 | 49789757 | Distal Intergenic |
| chr9 | 49808302 | 49810042 | Distal Intergenic |
| chr9 | 49813806 | 49814806 | Distal Intergenic |
| chr9 | 49900270 | 49900937 | Distal Intergenic |
| chr9 | 50278266 | 50280155 | Distal Intergenic |
| chr9 | 50280937 | 50281936 | Distal Intergenic |
| chr9 | 50284983 | 50285430 | Distal Intergenic |
| chr9 | 50302828 | 50303192 | Distal Intergenic |
| chr9 | 50304322 | 50304912 | Distal Intergenic |
| chr9 | 50328772 | 50329080 | Distal Intergenic |
| chr9 | 50329416 | 50329631 | Distal Intergenic |
| chr9 | 50399172 | 50399414 | Distal Intergenic |
| chr9 | 50412212 | 50412657 | Distal Intergenic |
| chr9 | 50424829 | 50425039 | Distal Intergenic |
| chr9 | 50425184 | 50425591 | Distal Intergenic |
| chr9 | 50489720 | 50490053 | Distal Intergenic |
| chr9 | 50494678 | 50496045 | Distal Intergenic |
| chr9 | 50549398 | 50550355 | Distal Intergenic |
| chr9 | 50554026 | 50554977 | Distal Intergenic |
| chr9 | 50655052 | 50655501 | Distal Intergenic |
| chr9 | 50660385 | 50661919 | Distal Intergenic |
| chr9 | 50662095 | 50662901 | Distal Intergenic |
| chr9 | 50708304 | 50709179 | Distal Intergenic |
| chr9 | 50864863 | 50865144 | Distal Intergenic |
| chr9 | 50866198 | 50866704 | Distal Intergenic |
| chr9 | 50866762 | 50867573 | Distal Intergenic |
| chr9 | 50913584 | 50914547 | Distal Intergenic |
| chr9 | 50977096 | 50978791 | Distal Intergenic |
| chr9 | 50983608 | 50984441 | Distal Intergenic |
| chr9 | 51076920 | 51078756 | Distal Intergenic |

| | | | |
|---|---|---|---|
| chr9 | 51772025 | 51772232 | Distal Intergenic |
| chr9 | 51856250 | 51856712 | Distal Intergenic |
| chr9 | 51857199 | 51857412 | Distal Intergenic |
| chr9 | 51975350 | 51975651 | Distal Intergenic |
| chr9 | 51975828 | 51976367 | Distal Intergenic |
| chr9 | 51980845 | 51981337 | Distal Intergenic |
| chr9 | 51982601 | 51983736 | Distal Intergenic |
| chr9 | 51988545 | 51990413 | Distal Intergenic |
| chr9 | 53071925 | 53072927 | Distal Intergenic |
| chr9 | 53086981 | 53087543 | Distal Intergenic |
| chr9 | 53095093 | 53095995 | Distal Intergenic |
| chr9 | 53098054 | 53098745 | Distal Intergenic |
| chr9 | 53192048 | 53192376 | Distal Intergenic |
| chr9 | 53192530 | 53192992 | Distal Intergenic |
| chr9 | 53216922 | 53217382 | Distal Intergenic |
| chr9 | 54414515 | 54415563 | Distal Intergenic |
| chr9 | 54460981 | 54461418 | Distal Intergenic |
| chr9 | 54490089 | 54491445 | Distal Intergenic |
| chr9 | 54493076 | 54494080 | Distal Intergenic |
| chr9 | 54494857 | 54495476 | Distal Intergenic |
| chr9 | 54496871 | 54497378 | Distal Intergenic |
| chr9 | 54520554 | 54521479 | Distal Intergenic |
| chr9 | 54555293 | 54556024 | Distal Intergenic |
| chr9 | 54570306 | 54570770 | Distal Intergenic |
| chr9 | 54580643 | 54580844 | Distal Intergenic |
| chr9 | 54580920 | 54582319 | Distal Intergenic |
| chr9 | 54642763 | 54643204 | Distal Intergenic |
| chr9 | 54650165 | 54651776 | Distal Intergenic |
| chr9 | 54756647 | 54757411 | Distal Intergenic |
| chr9 | 54757487 | 54757911 | Distal Intergenic |
| chr9 | 54798834 | 54799353 | Distal Intergenic |
| chr9 | 54962531 | 54962786 | Distal Intergenic |
| chr9 | 54962932 | 54963665 | Distal Intergenic |
| chr9 | 54984400 | 54985234 | Distal Intergenic |
| chr9 | 55258270 | 55258471 | Distal Intergenic |
| chr9 | 55258687 | 55259047 | Distal Intergenic |
| chr9 | 55324215 | 55324414 | Distal Intergenic |
| chr9 | 55787976 | 55788971 | Distal Intergenic |
| chr9 | 55789010 | 55789213 | Distal Intergenic |
| chr9 | 55907153 | 55907354 | Distal Intergenic |
| chr9 | 55937013 | 55937961 | Distal Intergenic |
| chr9 | 55938018 | 55939445 | Distal Intergenic |
| chr9 | 55947848 | 55949192 | Distal Intergenic |
| chr9 | 55949312 | 55949577 | Distal Intergenic |

| | | | |
|---|---|---|---|
| chr9 | 55951638 | 55951837 | Distal Intergenic |
| chr9 | 55953897 | 55956306 | Distal Intergenic |
| chr9 | 56007986 | 56008542 | Distal Intergenic |
| chr9 | 56015125 | 56016671 | Distal Intergenic |
| chr9 | 56025069 | 56026268 | Distal Intergenic |
| chr9 | 56027106 | 56028885 | Distal Intergenic |
| chr9 | 56044348 | 56045199 | Distal Intergenic |
| chr9 | 56112659 | 56113006 | Distal Intergenic |
| chr9 | 56113071 | 56113639 | Distal Intergenic |
| chr9 | 56182211 | 56182462 | Distal Intergenic |
| chr9 | 56218436 | 56219133 | Distal Intergenic |
| chr9 | 56240923 | 56241373 | Distal Intergenic |
| chr9 | 56353358 | 56353865 | Distal Intergenic |
| chr9 | 56354019 | 56354720 | Distal Intergenic |
| chr9 | 56356684 | 56358124 | Distal Intergenic |
| chr9 | 56362894 | 56363377 | Distal Intergenic |
| chr9 | 56392934 | 56393242 | Distal Intergenic |
| chr9 | 56400426 | 56400660 | Distal Intergenic |
| chr9 | 56706927 | 56707325 | Distal Intergenic |
| chr9 | 56708617 | 56709075 | Distal Intergenic |
| chr9 | 56748295 | 56749384 | Distal Intergenic |
| chr9 | 56749488 | 56749875 | Distal Intergenic |
| chr9 | 56754675 | 56755675 | Distal Intergenic |
| chr9 | 56755734 | 56756305 | Distal Intergenic |
| chr9 | 56758358 | 56759647 | Distal Intergenic |
| chr9 | 56760021 | 56760220 | Distal Intergenic |
| chr9 | 56760827 | 56762765 | Distal Intergenic |
| chr9 | 56766352 | 56767127 | Distal Intergenic |
| chr9 | 56767234 | 56767522 | Distal Intergenic |
| chr9 | 56771835 | 56772970 | Distal Intergenic |
| chr9 | 56774893 | 56775923 | Distal Intergenic |
| chr9 | 56798297 | 56798640 | Distal Intergenic |
| chr9 | 56799104 | 56799359 | Distal Intergenic |
| chr9 | 56816931 | 56818360 | Distal Intergenic |
| chr9 | 56842888 | 56843676 | Distal Intergenic |
| chr9 | 56843730 | 56844244 | Distal Intergenic |
| chr9 | 56922640 | 56922857 | Distal Intergenic |
| chr9 | 56923258 | 56923556 | Distal Intergenic |
| chr9 | 56978675 | 56979004 | Distal Intergenic |
| chr9 | 57051542 | 57051803 | Distal Intergenic |
| chr9 | 57369621 | 57369950 | Distal Intergenic |
| chr9 | 57385103 | 57385345 | Distal Intergenic |
| chr9 | 57462478 | 57463116 | Distal Intergenic |
| chr9 | 57466533 | 57467328 | Distal Intergenic |

| | | | |
|---|---|---|---|
| chr9 | 57473391 | 57474043 | Distal Intergenic |
| chr9 | 57488417 | 57488910 | Distal Intergenic |
| chr9 | 57612083 | 57612962 | Distal Intergenic |
| chr9 | 57758457 | 57758706 | Distal Intergenic |
| chr9 | 57903653 | 57904560 | Distal Intergenic |
| chr9 | 57979121 | 57979328 | Distal Intergenic |
| chr9 | 58095105 | 58097953 | Distal Intergenic |
| chr9 | 58101045 | 58101339 | Distal Intergenic |
| chr9 | 58103333 | 58104586 | Distal Intergenic |
| chr9 | 58283882 | 58285452 | Distal Intergenic |
| chr9 | 58290421 | 58290765 | Distal Intergenic |
| chr9 | 58295807 | 58296183 | Distal Intergenic |
| chr9 | 58301571 | 58302267 | Distal Intergenic |
| chr9 | 58395224 | 58395458 | Distal Intergenic |
| chr9 | 58396259 | 58398013 | Distal Intergenic |
| chr9 | 58401487 | 58403654 | Distal Intergenic |
| chr9 | 58428861 | 58429264 | Distal Intergenic |
| chr9 | 58429347 | 58429933 | Distal Intergenic |
| chr9 | 58511073 | 58511283 | Distal Intergenic |
| chr9 | 58513032 | 58514018 | Distal Intergenic |
| chr9 | 58554085 | 58554400 | Distal Intergenic |
| chr9 | 59317315 | 59317645 | Distal Intergenic |
| chr9 | 59333128 | 59333944 | Distal Intergenic |
| chr9 | 59334723 | 59334970 | Distal Intergenic |
| chr9 | 59469763 | 59470058 | Distal Intergenic |
| chr9 | 59470096 | 59471181 | Distal Intergenic |
| chr9 | 59488827 | 59489491 | Distal Intergenic |
| chr9 | 59500560 | 59502017 | Distal Intergenic |
| chr9 | 59502112 | 59502603 | Distal Intergenic |
| chr9 | 59503216 | 59504330 | Distal Intergenic |
| chr9 | 59504456 | 59504841 | Distal Intergenic |
| chr9 | 59505062 | 59507695 | Distal Intergenic |
| chr9 | 59507871 | 59508070 | Distal Intergenic |
| chr9 | 59528574 | 59529675 | Distal Intergenic |
| chr9 | 59529755 | 59529971 | Distal Intergenic |
| chr9 | 59534565 | 59536088 | Distal Intergenic |
| chr9 | 59538539 | 59541542 | Distal Intergenic |
| chr9 | 59554252 | 59555817 | Distal Intergenic |
| chr9 | 59580985 | 59581749 | Distal Intergenic |
| chr9 | 59581955 | 59582173 | Distal Intergenic |
| chr9 | 60535579 | 60535809 | Distal Intergenic |
| chr9 | 60685445 | 60685992 | Distal Intergenic |
| chr9 | 61128917 | 61129332 | Distal Intergenic |
| chr9 | 61129511 | 61131584 | Distal Intergenic |

| | | | |
|---|---|---|---|
| chr9 | 61134590 | 61134793 | Distal Intergenic |
| chr9 | 61169505 | 61170282 | Distal Intergenic |
| chr9 | 61170359 | 61170765 | Distal Intergenic |
| chr9 | 61406441 | 61406644 | Distal Intergenic |
| chr9 | 61406845 | 61407189 | Distal Intergenic |
| chr9 | 61517330 | 61517615 | Distal Intergenic |
| chr9 | 61759340 | 61759591 | Distal Intergenic |
| chr9 | 61760781 | 61760999 | Distal Intergenic |
| chr9 | 61794762 | 61795845 | Distal Intergenic |
| chr9 | 61800142 | 61800402 | Distal Intergenic |
| chr9 | 61810735 | 61813837 | Distal Intergenic |
| chr9 | 61813939 | 61814374 | Distal Intergenic |
| chr9 | 61846018 | 61846222 | Distal Intergenic |
| chr9 | 61847242 | 61847472 | Distal Intergenic |
| chr9 | 61847603 | 61848594 | Distal Intergenic |
| chr9 | 62658432 | 62658839 | Distal Intergenic |
| chr9 | 62659045 | 62659541 | Distal Intergenic |
| chr9 | 62664909 | 62666558 | Distal Intergenic |
| chr9 | 62675788 | 62676128 | Distal Intergenic |
| chr9 | 62885174 | 62885402 | Distal Intergenic |
| chr9 | 63427874 | 63428337 | Distal Intergenic |
| chr9 | 63455708 | 63456226 | Distal Intergenic |
| chr9 | 63458805 | 63459013 | Distal Intergenic |
| chr9 | 63494656 | 63495047 | Distal Intergenic |
| chr9 | 63495489 | 63495785 | Distal Intergenic |
| chr9 | 63522442 | 63523355 | Distal Intergenic |
| chr9 | 63525404 | 63525784 | Distal Intergenic |
| chr9 | 63569096 | 63569674 | Distal Intergenic |
| chr9 | 63573372 | 63573774 | Distal Intergenic |
| chr9 | 63593798 | 63594071 | Distal Intergenic |
| chr9 | 63662617 | 63663492 | Distal Intergenic |
| chr9 | 63684628 | 63684828 | Distal Intergenic |
| chr9 | 63840239 | 63841159 | Distal Intergenic |
| chr9 | 63865700 | 63866915 | Distal Intergenic |
| chr9 | 63869599 | 63869798 | Distal Intergenic |
| chr9 | 63897365 | 63898817 | Distal Intergenic |
| chr9 | 64020498 | 64021540 | Distal Intergenic |
| chr9 | 64031922 | 64032455 | Distal Intergenic |
| chr9 | 64050494 | 64050908 | Distal Intergenic |
| chr9 | 64051230 | 64051530 | Distal Intergenic |
| chr9 | 64064371 | 64064771 | Distal Intergenic |
| chr9 | 64065754 | 64066108 | Distal Intergenic |
| chr9 | 64100418 | 64100681 | Distal Intergenic |
| chr9 | 64129453 | 64129836 | Distal Intergenic |

| | | | |
|---|---|---|---|
| chr9 | 64188138 | 64189328 | Distal Intergenic |
| chr9 | 64319051 | 64319257 | Distal Intergenic |
| chr9 | 64548807 | 64552069 | Distal Intergenic |
| chr9 | 64582091 | 64582617 | Distal Intergenic |
| chr9 | 64583346 | 64584563 | Distal Intergenic |
| chr9 | 64692829 | 64694224 | Distal Intergenic |
| chr9 | 64808625 | 64808995 | Distal Intergenic |
| chr9 | 64836055 | 64836310 | Distal Intergenic |
| chr9 | 64841906 | 64842725 | Distal Intergenic |
| chr9 | 64866135 | 64866604 | Distal Intergenic |
| chr9 | 64867661 | 64867869 | Distal Intergenic |
| chr9 | 65049083 | 65049833 | Distal Intergenic |
| chr9 | 65139311 | 65140332 | Distal Intergenic |
| chr9 | 65141108 | 65141829 | Distal Intergenic |
| chr9 | 65193556 | 65194069 | Distal Intergenic |
| chr9 | 65340224 | 65340821 | Distal Intergenic |
| chr9 | 65389147 | 65389920 | Distal Intergenic |
| chr9 | 65397274 | 65399607 | Distal Intergenic |
| chr9 | 65422266 | 65424009 | Distal Intergenic |
| chr9 | 65484953 | 65485325 | Downstream |
| chr9 | 65567747 | 65568250 | Distal Intergenic |
| chr9 | 65624461 | 65625604 | Distal Intergenic |
| chr9 | 65674341 | 65675091 | Distal Intergenic |
| chr9 | 65676015 | 65676262 | Distal Intergenic |
| chr9 | 65676834 | 65677173 | Distal Intergenic |
| chr9 | 65756386 | 65756632 | Distal Intergenic |
| chr9 | 65757038 | 65757679 | Distal Intergenic |
| chr9 | 65757726 | 65758176 | Distal Intergenic |
| chr9 | 65972331 | 65972564 | Distal Intergenic |
| chr9 | 66054586 | 66055033 | Distal Intergenic |
| chr9 | 66065420 | 66066177 | Distal Intergenic |
| chr9 | 66073482 | 66074708 | Distal Intergenic |
| chr9 | 66075390 | 66075682 | Distal Intergenic |
| chr9 | 66075901 | 66077286 | Distal Intergenic |
| chr9 | 66088405 | 66088736 | Distal Intergenic |
| chr9 | 66364664 | 66364876 | Distal Intergenic |
| chr9 | 66439558 | 66440346 | Distal Intergenic |
| chr9 | 66440487 | 66440774 | Distal Intergenic |
| chr9 | 66468236 | 66469449 | Exon |
| chr9 | 66469559 | 66470101 | Distal Intergenic |
| chr9 | 66470456 | 66470732 | Distal Intergenic |
| chr9 | 66568367 | 66568597 | Distal Intergenic |
| chr9 | 66578829 | 66579040 | Distal Intergenic |
| chr9 | 66763748 | 66764354 | Distal Intergenic |

| | | | |
|---|---|---|---|
| chr9 | 66793398 | 66794436 | Distal Intergenic |
| chr9 | 66794488 | 66794875 | Distal Intergenic |
| chr9 | 66823241 | 66823786 | Distal Intergenic |
| chr9 | 66836139 | 66838991 | Distal Intergenic |
| chr9 | 66872487 | 66872694 | Distal Intergenic |
| chr9 | 66872970 | 66873169 | Distal Intergenic |
| chr9 | 66873210 | 66873670 | Distal Intergenic |
| chr9 | 66884438 | 66885457 | Distal Intergenic |
| chr9 | 66885656 | 66886086 | Distal Intergenic |
| chr9 | 66888734 | 66888987 | Distal Intergenic |
| chr9 | 66890925 | 66891124 | Distal Intergenic |
| chr9 | 66893012 | 66893764 | Distal Intergenic |
| chr9 | 66896177 | 66896663 | Distal Intergenic |
| chr9 | 66929405 | 66930024 | Intron |
| chr9 | 66952422 | 66954224 | Distal Intergenic |
| chr9 | 66954334 | 66954558 | Distal Intergenic |
| chr9 | 66955746 | 66956040 | Distal Intergenic |
| chr9 | 67007216 | 67007415 | Distal Intergenic |
| chr9 | 67008030 | 67008300 | Distal Intergenic |
| chr9 | 67013149 | 67013781 | Distal Intergenic |
| chr9 | 67055720 | 67055976 | Distal Intergenic |
| chr9 | 67056337 | 67057465 | Distal Intergenic |
| chr9 | 67071242 | 67072183 | Distal Intergenic |
| chr9 | 67178890 | 67179229 | Distal Intergenic |
| chr9 | 67724795 | 67725263 | Distal Intergenic |
| chr9 | 67725310 | 67725548 | Distal Intergenic |
| chr9 | 67802578 | 67804129 | Distal Intergenic |
| chr9 | 68635226 | 68636179 | Distal Intergenic |
| chr9 | 68636240 | 68636713 | Distal Intergenic |
| chr9 | 68636911 | 68637155 | Distal Intergenic |
| chr9 | 69245832 | 69246031 | Exon |
| chr9 | 69278995 | 69280312 | Distal Intergenic |
| chr9 | 69290060 | 69291382 | Distal Intergenic |
| chr9 | 69301486 | 69308316 | Distal Intergenic |
| chr9 | 69310098 | 69310370 | Distal Intergenic |
| chr9 | 69311170 | 69311370 | Distal Intergenic |
| chr9 | 69311422 | 69311621 | Distal Intergenic |
| chr9 | 69313405 | 69313658 | Distal Intergenic |
| chr9 | 69313696 | 69317339 | Distal Intergenic |
| chr9 | 69319204 | 69319692 | Distal Intergenic |
| chr9 | 69319743 | 69321189 | Distal Intergenic |
| chr9 | 69321226 | 69321516 | Distal Intergenic |
| chr9 | 69895476 | 69896917 | Distal Intergenic |
| chr9 | 70055271 | 70055548 | Distal Intergenic |

| | | | |
|---|---|---|---|
| chr9 | 70055956 | 70056269 | Distal Intergenic |
| chr9 | 70056669 | 70058118 | Distal Intergenic |
| chr9 | 70058271 | 70058658 | Distal Intergenic |
| chr9 | 70064981 | 70065314 | Distal Intergenic |
| chr9 | 70065635 | 70066023 | Distal Intergenic |
| chr9 | 70075265 | 70076131 | Distal Intergenic |
| chr9 | 70077187 | 70078157 | Distal Intergenic |
| chr9 | 70096881 | 70097132 | Distal Intergenic |
| chr9 | 70100581 | 70100780 | Distal Intergenic |
| chr9 | 70100835 | 70101742 | Distal Intergenic |
| chr9 | 70104602 | 70105903 | Distal Intergenic |
| chr9 | 70108400 | 70109753 | Distal Intergenic |
| chr9 | 70138350 | 70138920 | Distal Intergenic |
| chr9 | 70148264 | 70150315 | Distal Intergenic |
| chr9 | 70268937 | 70269384 | Distal Intergenic |
| chr9 | 70357613 | 70358839 | Distal Intergenic |
| chr9 | 70359669 | 70360011 | Distal Intergenic |
| chr9 | 70423517 | 70423771 | Downstream |
| chr9 | 70477678 | 70477877 | Promoter |
| chr9 | 70478900 | 70479397 | Intron |
| chr9 | 70484421 | 70485203 | Exon |
| chr9 | 70528005 | 70528205 | Distal Intergenic |
| chr9 | 70561101 | 70561322 | Distal Intergenic |
| chr9 | 70580520 | 70581537 | Distal Intergenic |
| chr9 | 70599024 | 70600032 | Distal Intergenic |
| chr9 | 70600098 | 70600453 | Distal Intergenic |
| chr9 | 70609148 | 70609416 | Distal Intergenic |
| chr9 | 70682108 | 70683539 | Distal Intergenic |
| chr9 | 70685517 | 70687367 | Distal Intergenic |
| chr9 | 70689771 | 70689977 | Distal Intergenic |
| chr9 | 70690035 | 70690320 | Distal Intergenic |
| chr9 | 70735826 | 70737588 | Distal Intergenic |
| chr9 | 70742074 | 70742482 | Distal Intergenic |
| chr9 | 70742714 | 70743895 | Distal Intergenic |
| chr9 | 70772454 | 70776074 | Distal Intergenic |
| chr9 | 71003364 | 71003583 | Intron |
| chr9 | 71332746 | 71333747 | Intron |
| chr9 | 71418744 | 71419483 | Intron |
| chr9 | 71421810 | 71422035 | Intron |
| chr9 | 71868892 | 71870063 | 3' UTR |
| chr9 | 71923305 | 71925404 | Distal Intergenic |
| chr9 | 71948566 | 71949877 | Intron |
| chr9 | 71958991 | 71959210 | Intron |
| chr9 | 72123088 | 72123630 | Intron |

| | | | |
|---|---|---|---|
| chr9 | 72378622 | 72379867 | Distal Intergenic |
| chr9 | 72380130 | 72380453 | Distal Intergenic |
| chr9 | 72510725 | 72511345 | Intron |
| chr9 | 72549439 | 72550116 | Distal Intergenic |
| chr9 | 72887702 | 72888060 | Intron |
| chr9 | 72915588 | 72916761 | Intron |
| chr9 | 72960793 | 72961200 | Promoter |
| chr9 | 72961442 | 72961731 | Promoter |
| chr9 | 73654168 | 73654544 | Intron |
| chr9 | 73753533 | 73753791 | Intron |
| chr9 | 74866724 | 74868312 | 3' UTR |
| chr9 | 74870747 | 74871197 | Distal Intergenic |
| chr9 | 74871279 | 74871985 | Distal Intergenic |
| chr9 | 74885424 | 74886437 | Distal Intergenic |
| chr9 | 74886557 | 74887822 | Distal Intergenic |
| chr9 | 74932817 | 74933666 | Intron |
| chr9 | 74974751 | 74974996 | Promoter |
| chr9 | 74975132 | 74975453 | Promoter |
| chr9 | 75042105 | 75042304 | Distal Intergenic |
| chr9 | 75043042 | 75044951 | Distal Intergenic |
| chr9 | 75200134 | 75201822 | Intron |
| chr9 | 75214008 | 75214371 | Intron |
| chr9 | 75226433 | 75226639 | Intron |
| chr9 | 75226838 | 75228175 | Intron |
| chr9 | 75232574 | 75234988 | Intron |
| chr9 | 75255410 | 75255688 | Intron |
| chr9 | 75255855 | 75257425 | Intron |
| chr9 | 75257566 | 75257815 | Intron |
| chr9 | 75258123 | 75258353 | Intron |
| chr9 | 75269536 | 75271329 | Intron |
| chr9 | 75271431 | 75271917 | Intron |
| chr9 | 75273757 | 75273956 | Intron |
| chr9 | 75275192 | 75275546 | Intron |
| chr9 | 75275592 | 75275942 | Intron |
| chr9 | 75312675 | 75312905 | Intron |
| chr9 | 75313253 | 75313912 | Intron |
| chr9 | 75313988 | 75314219 | Intron |
| chr9 | 75377417 | 75377723 | Intron |
| chr9 | 75389226 | 75389536 | Intron |
| chr9 | 75389673 | 75389919 | Intron |
| chr9 | 75406794 | 75407104 | Exon |
| chr9 | 75858025 | 75858363 | Distal Intergenic |
| chr9 | 75862391 | 75863598 | Distal Intergenic |
| chr9 | 75871214 | 75872631 | Distal Intergenic |

| | | | |
|---|---|---|---|
| chr9 | 76359377 | 76359623 | Distal Intergenic |
| chr9 | 76362737 | 76363899 | Distal Intergenic |
| chr9 | 76370488 | 76371833 | Distal Intergenic |
| chr9 | 76386738 | 76386971 | Distal Intergenic |
| chr9 | 76387734 | 76390490 | Distal Intergenic |
| chr9 | 76401380 | 76402295 | Distal Intergenic |
| chr9 | 76413357 | 76414916 | Distal Intergenic |
| chr9 | 76415148 | 76415419 | Distal Intergenic |
| chr9 | 76500970 | 76501273 | Distal Intergenic |
| chr9 | 76651718 | 76652173 | Distal Intergenic |
| chr9 | 76652568 | 76653323 | Distal Intergenic |
| chr9 | 77312338 | 77312744 | Distal Intergenic |
| chr9 | 77393635 | 77394044 | Intron |
| chr9 | 77434327 | 77435906 | Exon |
| chr9 | 77549298 | 77549497 | Distal Intergenic |
| chr9 | 77549541 | 77551231 | Distal Intergenic |
| chr9 | 77551316 | 77552539 | Distal Intergenic |
| chr9 | 77558678 | 77559905 | Downstream |
| chr9 | 77592308 | 77592893 | Intron |
| chr9 | 77602818 | 77603135 | Intron |
| chr9 | 77758629 | 77759674 | Intron |
| chr9 | 77833935 | 77834355 | Distal Intergenic |
| chr9 | 78023788 | 78024843 | Distal Intergenic |
| chr9 | 78026576 | 78027058 | Distal Intergenic |
| chr9 | 78031696 | 78032496 | Distal Intergenic |
| chr9 | 78038374 | 78039951 | Distal Intergenic |
| chr9 | 78040011 | 78040257 | Distal Intergenic |
| chr9 | 78040447 | 78040744 | Distal Intergenic |
| chr9 | 78041038 | 78041239 | Distal Intergenic |
| chr9 | 78059650 | 78061041 | Distal Intergenic |
| chr9 | 78077614 | 78078516 | Distal Intergenic |
| chr9 | 78103266 | 78104699 | Distal Intergenic |
| chr9 | 78118714 | 78119590 | Intron |
| chr9 | 78129900 | 78130113 | Intron |
| chr9 | 78324455 | 78325319 | Intron |
| chr9 | 78325443 | 78325678 | Intron |
| chr9 | 78327400 | 78327668 | Intron |
| chr9 | 78328087 | 78328320 | Intron |
| chr9 | 78328522 | 78328782 | Intron |
| chr9 | 78329805 | 78330258 | Intron |
| chr9 | 78350669 | 78352619 | Distal Intergenic |
| chr9 | 78375406 | 78375792 | Distal Intergenic |
| chr9 | 78470042 | 78470269 | Distal Intergenic |
| chr9 | 78470432 | 78471874 | Distal Intergenic |

| | | | |
|---|---|---|---|
| chr9 | 78569141 | 78569468 | Intron |
| chr9 | 78919542 | 78919780 | Intron |
| chr9 | 79013268 | 79014072 | Promoter |
| chr9 | 79014117 | 79014363 | Promoter |
| chr9 | 79063575 | 79063802 | Intron |
| chr9 | 79064167 | 79064407 | Intron |
| chr9 | 79377836 | 79379191 | Promoter |
| chr9 | 79437147 | 79437402 | Intron |
| chr9 | 79496595 | 79497179 | Intron |
| chr9 | 79497221 | 79497524 | Intron |
| chr9 | 79528977 | 79529398 | Distal Intergenic |
| chr9 | 79556448 | 79557425 | Distal Intergenic |
| chr9 | 79561255 | 79563041 | Distal Intergenic |
| chr9 | 79607213 | 79607849 | Distal Intergenic |
| chr9 | 79914701 | 79916267 | Intron |
| chr9 | 80007488 | 80007689 | Intron |
| chr9 | 80581838 | 80582109 | Intron |
| chr9 | 80582363 | 80582621 | Intron |
| chr9 | 80707899 | 80709051 | Distal Intergenic |
| chr9 | 80997348 | 80997610 | Distal Intergenic |
| chr9 | 81019881 | 81020321 | Distal Intergenic |
| chr9 | 82199838 | 82200041 | Intron |
| chr9 | 82200088 | 82200707 | Intron |
| chr9 | 82860265 | 82861642 | Distal Intergenic |
| chr9 | 82862173 | 82862413 | Distal Intergenic |
| chr9 | 82865083 | 82865282 | Distal Intergenic |
| chr9 | 82866818 | 82867661 | Distal Intergenic |
| chr9 | 83126609 | 83126869 | Distal Intergenic |
| chr9 | 83126920 | 83127127 | Distal Intergenic |
| chr9 | 83236280 | 83236822 | Distal Intergenic |
| chr9 | 83241354 | 83241761 | Distal Intergenic |
| chr9 | 83242728 | 83243763 | Distal Intergenic |
| chr9 | 83442147 | 83442904 | Distal Intergenic |
| chr9 | 83456431 | 83460117 | Distal Intergenic |
| chr9 | 83460203 | 83460500 | Distal Intergenic |
| chr9 | 83461510 | 83463294 | Distal Intergenic |
| chr9 | 83463528 | 83463807 | Distal Intergenic |
| chr9 | 83464568 | 83465956 | Distal Intergenic |
| chr9 | 83509716 | 83510680 | Distal Intergenic |
| chr9 | 83649990 | 83650225 | Distal Intergenic |
| chr9 | 83715757 | 83717814 | Distal Intergenic |
| chr9 | 83724552 | 83724840 | Distal Intergenic |
| chr9 | 83776095 | 83776464 | Distal Intergenic |
| chr9 | 83776553 | 83777041 | Distal Intergenic |

| | | | |
|---|---|---|---|
| chr9 | 85391054 | 85391503 | Distal Intergenic |
| chr9 | 85510296 | 85510903 | Distal Intergenic |
| chr9 | 85642593 | 85642794 | Intron |
| chr9 | 85643030 | 85643598 | Intron |
| chr9 | 85714615 | 85714935 | Distal Intergenic |
| chr9 | 85736020 | 85737538 | Distal Intergenic |
| chr9 | 86340895 | 86341113 | Distal Intergenic |
| chr9 | 86361147 | 86361376 | Intron |
| chr9 | 86517989 | 86519369 | Exon |
| chr9 | 86521488 | 86521792 | Intron |
| chr9 | 86521887 | 86522086 | Intron |
| chr9 | 86698257 | 86698823 | Intron |
| chr9 | 86894738 | 86895187 | Exon |
| chr9 | 86897958 | 86898250 | Intron |
| chr9 | 86898356 | 86898870 | Intron |
| chr9 | 87358180 | 87359871 | Intron |
| chr9 | 88220584 | 88221358 | Intron |
| chr9 | 88267881 | 88268345 | Intron |
| chr9 | 88334050 | 88334417 | Intron |
| chr9 | 88334513 | 88334748 | Intron |
| chr9 | 88375893 | 88376172 | Distal Intergenic |
| chr9 | 88392437 | 88393240 | Distal Intergenic |
| chr9 | 88414631 | 88414912 | Distal Intergenic |
| chr9 | 88416723 | 88416934 | Distal Intergenic |
| chr9 | 88417323 | 88417691 | Distal Intergenic |
| chr9 | 88434596 | 88434877 | Intron |
| chr9 | 88614648 | 88614907 | Intron |
| chr9 | 88614990 | 88615202 | Intron |
| chr9 | 89930209 | 89932600 | Distal Intergenic |
| chr9 | 89963414 | 89964469 | Distal Intergenic |
| chr9 | 89964517 | 89965039 | Distal Intergenic |
| chr9 | 90008260 | 90008586 | Distal Intergenic |
| chr9 | 90008768 | 90008967 | Distal Intergenic |
| chr9 | 90108648 | 90109046 | Distal Intergenic |
| chr9 | 90110453 | 90110771 | Distal Intergenic |
| chr9 | 90110829 | 90111073 | Distal Intergenic |
| chr9 | 90111112 | 90111999 | Promoter |
| chr9 | 90130296 | 90130897 | Intron |
| chr9 | 90135182 | 90135780 | Intron |
| chr9 | 90135933 | 90136354 | Intron |
| chr9 | 90143773 | 90145625 | Intron |
| chr9 | 90156755 | 90156963 | Intron |
| chr9 | 90157092 | 90158542 | Intron |
| chr9 | 90246384 | 90246658 | Intron |

| | | | |
|---|---|---|---|
| chr9 | 92436621 | 92436820 | Distal Intergenic |
| chr9 | 92437423 | 92437694 | Distal Intergenic |
| chr9 | 93026525 | 93026903 | Distal Intergenic |
| chr9 | 94329242 | 94329627 | Intron |
| chr9 | 95093156 | 95094254 | Intron |
| chr9 | 95411747 | 95412057 | Exon |
| chr9 | 95412449 | 95412718 | Intron |
| chr9 | 95414502 | 95414939 | Exon |
| chr9 | 95543542 | 95543744 | Distal Intergenic |
| chr9 | 95572745 | 95572952 | Promoter |
| chr9 | 95758386 | 95758690 | Intron |
| chr9 | 95904636 | 95905304 | Distal Intergenic |
| chr9 | 96102983 | 96103182 | Intron |
| chr9 | 96263981 | 96264225 | Intron |
| chr9 | 96281494 | 96282734 | Intron |
| chr9 | 96284057 | 96284638 | Intron |
| chr9 | 96284836 | 96285036 | Intron |
| chr9 | 96313059 | 96313998 | Intron |
| chr9 | 96315682 | 96316027 | Intron |
| chr9 | 96316459 | 96317333 | Intron |
| chr9 | 96317370 | 96317590 | Intron |
| chr9 | 96606564 | 96606832 | Distal Intergenic |
| chr9 | 96614584 | 96614917 | Distal Intergenic |
| chr9 | 96614976 | 96615377 | Distal Intergenic |
| chr9 | 96621031 | 96622031 | Distal Intergenic |
| chr9 | 96651889 | 96652359 | Distal Intergenic |
| chr9 | 96652451 | 96653885 | Distal Intergenic |
| chr9 | 96655718 | 96656487 | Distal Intergenic |
| chr9 | 96656719 | 96656980 | Distal Intergenic |
| chr9 | 96669987 | 96670558 | Distal Intergenic |
| chr9 | 96670600 | 96670877 | Distal Intergenic |
| chr9 | 96672923 | 96673500 | Distal Intergenic |
| chr9 | 96673681 | 96675631 | Distal Intergenic |
| chr9 | 96681343 | 96681542 | Distal Intergenic |
| chr9 | 96681592 | 96682988 | Distal Intergenic |
| chr9 | 96683030 | 96684830 | Distal Intergenic |
| chr9 | 96689465 | 96691365 | Distal Intergenic |
| chr9 | 96691414 | 96691656 | Distal Intergenic |
| chr9 | 96700430 | 96701281 | Distal Intergenic |
| chr9 | 96761029 | 96761681 | Distal Intergenic |
| chr9 | 96761802 | 96762031 | Distal Intergenic |
| chr9 | 96763319 | 96764036 | Distal Intergenic |
| chr9 | 97003863 | 97004568 | Distal Intergenic |
| chr9 | 97176290 | 97176591 | Promoter |

| | | | |
|---|---|---|---|
| chr9 | 97535399 | 97535621 | Exon |
| chr9 | 97535724 | 97535952 | Intron |
| chr9 | 98197804 | 98198174 | Distal Intergenic |
| chr9 | 98214707 | 98215063 | Intron |
| chr9 | 98238273 | 98239317 | Exon |
| chr9 | 98333149 | 98335010 | Distal Intergenic |
| chr9 | 99030994 | 99031363 | Intron |
| chr9 | 99031902 | 99032474 | Intron |
| chr9 | 99032597 | 99033032 | Intron |
| chr9 | 99037598 | 99037896 | Intron |
| chr9 | 99066024 | 99067930 | Distal Intergenic |
| chr9 | 99529972 | 99530171 | Intron |
| chr9 | 99530545 | 99533552 | Intron |
| chr9 | 99620458 | 99621047 | Intron |
| chr9 | 100505575 | 100506891 | Distal Intergenic |
| chr9 | 100542229 | 100542578 | Distal Intergenic |
| chr9 | 100542724 | 100543293 | Distal Intergenic |
| chr9 | 100544995 | 100545433 | Distal Intergenic |
| chr9 | 100979696 | 100981048 | Intron |
| chr9 | 101077623 | 101077843 | Intron |
| chr9 | 101131400 | 101131697 | Intron |
| chr9 | 101131906 | 101132135 | Intron |
| chr9 | 101145527 | 101146980 | Intron |
| chr9 | 101147137 | 101147351 | Intron |
| chr9 | 101167837 | 101168872 | Exon |
| chr9 | 101170666 | 101173280 | Intron |
| chr9 | 101191115 | 101193036 | Intron |
| chr9 | 102535979 | 102536216 | Intron |
| chr9 | 102536393 | 102536630 | Intron |
| chr9 | 102536712 | 102537154 | Intron |
| chr9 | 102540408 | 102540654 | Intron |
| chr9 | 102575429 | 102575840 | Intron |
| chr9 | 102576098 | 102576904 | Intron |
| chr9 | 102639710 | 102639941 | Distal Intergenic |
| chr9 | 102639987 | 102640194 | Distal Intergenic |
| chr9 | 102640388 | 102640732 | Distal Intergenic |
| chr9 | 102663467 | 102666545 | Intron |
| chr9 | 102679329 | 102682071 | Intron |
| chr9 | 102682128 | 102683180 | Intron |
| chr9 | 102683441 | 102683886 | Intron |
| chr9 | 102683928 | 102684358 | Intron |
| chr9 | 102685118 | 102685894 | Intron |
| chr9 | 102707615 | 102708397 | Intron |
| chr9 | 102708912 | 102709210 | Intron |

| | | | |
|---|---|---|---|
| chr9 | 102744735 | 102745376 | 3' UTR |
| chr9 | 102745816 | 102746400 | Intron |
| chr9 | 102874770 | 102875079 | Intron |
| chr9 | 102963397 | 102964424 | Intron |
| chr9 | 102975010 | 102975313 | Intron |
| chr9 | 102984146 | 102984345 | Intron |
| chr9 | 103207555 | 103207832 | Intron |
| chr9 | 103207869 | 103208095 | Intron |
| chr9 | 103274677 | 103275172 | Intron |
| chr9 | 103278603 | 103278876 | Intron |
| chr9 | 103278928 | 103279667 | Exon |
| chr9 | 103987669 | 103987887 | Intron |
| chr9 | 103988001 | 103990081 | Intron |
| chr9 | 104057427 | 104057691 | Intron |
| chr9 | 104154009 | 104154493 | Intron |
| chr9 | 104154621 | 104155006 | Intron |
| chr9 | 104156756 | 104157221 | Intron |
| chr9 | 104239755 | 104240710 | Intron |
| chr9 | 104240794 | 104241048 | Intron |
| chr9 | 104263762 | 104264349 | Distal Intergenic |
| chr9 | 105360912 | 105361486 | Distal Intergenic |
| chr9 | 105362815 | 105363104 | Distal Intergenic |
| chr9 | 105363748 | 105364395 | Distal Intergenic |
| chr9 | 105397738 | 105397943 | Distal Intergenic |
| chr9 | 105595193 | 105595498 | Distal Intergenic |
| chr9 | 105599395 | 105599677 | Distal Intergenic |
| chr9 | 105599770 | 105600210 | Distal Intergenic |
| chr9 | 105604564 | 105604984 | Distal Intergenic |
| chr9 | 105905144 | 105905409 | Exon |
| chr9 | 106074450 | 106074672 | Intron |
| chr9 | 106177252 | 106177536 | Distal Intergenic |
| chr9 | 106261871 | 106262101 | Distal Intergenic |
| chr9 | 106263292 | 106263544 | Distal Intergenic |
| chr9 | 106267871 | 106268261 | Distal Intergenic |
| chr9 | 106268907 | 106269896 | Distal Intergenic |
| chr9 | 106271550 | 106274740 | Distal Intergenic |
| chr9 | 106277192 | 106277414 | Distal Intergenic |
| chr9 | 106279152 | 106281904 | Distal Intergenic |
| chr9 | 106293460 | 106293879 | Distal Intergenic |
| chr9 | 106294032 | 106294868 | Distal Intergenic |
| chr9 | 106295218 | 106295534 | Distal Intergenic |
| chr9 | 106296681 | 106296964 | Distal Intergenic |
| chr9 | 106314244 | 106314597 | Distal Intergenic |
| chr9 | 106314772 | 106315974 | Distal Intergenic |

| | | | |
|---|---|---|---|
| chr9 | 106316062 | 106316311 | Distal Intergenic |
| chr9 | 106378768 | 106378968 | Distal Intergenic |
| chr9 | 106724440 | 106724647 | Distal Intergenic |
| chr9 | 106794586 | 106794949 | Distal Intergenic |
| chr9 | 107181697 | 107181949 | Distal Intergenic |
| chr9 | 107191160 | 107191886 | Distal Intergenic |
| chr9 | 107191958 | 107192184 | Distal Intergenic |
| chr9 | 107200933 | 107201132 | Distal Intergenic |
| chr9 | 107207702 | 107207924 | Distal Intergenic |
| chr9 | 107458117 | 107458745 | Distal Intergenic |
| chr9 | 107458787 | 107459144 | Distal Intergenic |
| chr9 | 107459852 | 107460093 | Distal Intergenic |
| chr9 | 107471021 | 107472549 | Distal Intergenic |
| chr9 | 107477571 | 107479252 | Distal Intergenic |
| chr9 | 107479482 | 107479715 | Distal Intergenic |
| chr9 | 107481661 | 107482127 | Distal Intergenic |
| chr9 | 107482440 | 107482771 | Distal Intergenic |
| chr9 | 107535209 | 107535667 | Promoter |
| chr9 | 107536174 | 107536412 | Promoter |
| chr9 | 107536570 | 107536987 | Promoter |
| chr9 | 107537328 | 107537579 | Promoter |
| chr9 | 107603846 | 107604144 | Intron |
| chr9 | 107605845 | 107606125 | Exon |
| chr9 | 107606209 | 107606571 | Exon |
| chr9 | 107609251 | 107609967 | Intron |
| chr9 | 107610105 | 107610920 | Intron |
| chr9 | 107611174 | 107611611 | Intron |
| chr9 | 107672965 | 107673314 | Intron |
| chr9 | 107673485 | 107673736 | Intron |
| chr9 | 107809674 | 107813548 | Distal Intergenic |
| chr9 | 107982741 | 107983342 | Distal Intergenic |
| chr9 | 108039484 | 108039689 | Intron |
| chr9 | 108166480 | 108167172 | Intron |
| chr9 | 108167235 | 108167487 | Intron |
| chr9 | 108169654 | 108170508 | Intron |
| chr9 | 108174228 | 108175001 | Intron |
| chr9 | 108207903 | 108208347 | Distal Intergenic |
| chr9 | 108240624 | 108241453 | Intron |
| chr9 | 108241652 | 108241964 | Exon |
| chr9 | 108243945 | 108244268 | Intron |
| chr9 | 108244574 | 108244868 | Intron |
| chr9 | 108264864 | 108265305 | Intron |
| chr9 | 108361472 | 108363392 | 5' UTR |
| chr9 | 108393191 | 108393945 | Intron |

| | | | |
|---|---|---|---|
| chr9 | 108394172 | 108394473 | Intron |
| chr9 | 108424028 | 108424241 | Promoter |
| chr9 | 108847078 | 108847736 | Distal Intergenic |
| chr9 | 108850248 | 108852908 | Distal Intergenic |
| chr9 | 108854101 | 108855149 | Distal Intergenic |
| chr9 | 108956776 | 108957018 | Intron |
| chr9 | 108961409 | 108962512 | Intron |
| chr9 | 108996326 | 108997132 | Intron |
| chr9 | 108997218 | 108997620 | Intron |
| chr9 | 108998609 | 108998994 | Intron |
| chr9 | 108999102 | 108999313 | Intron |
| chr9 | 109021497 | 109021778 | Intron |
| chr9 | 109022128 | 109022730 | Intron |
| chr9 | 109798303 | 109798564 | Intron |
| chr9 | 109799273 | 109800365 | Intron |
| chr9 | 110247267 | 110247941 | 3' UTR |
| chr9 | 110251548 | 110251942 | Promoter |
| chr9 | 110251981 | 110253666 | Promoter |
| chr9 | 110309904 | 110310376 | Distal Intergenic |
| chr9 | 110378825 | 110379162 | Distal Intergenic |
| chr9 | 110435418 | 110435723 | Distal Intergenic |
| chr9 | 110435805 | 110436105 | Distal Intergenic |
| chr9 | 110486548 | 110486747 | Distal Intergenic |
| chr9 | 110486976 | 110487353 | Distal Intergenic |
| chr9 | 110487525 | 110488053 | Distal Intergenic |
| chr9 | 110583475 | 110583992 | Distal Intergenic |
| chr9 | 110585821 | 110586270 | Distal Intergenic |
| chr9 | 110586471 | 110587801 | Distal Intergenic |
| chr9 | 110599324 | 110599930 | Distal Intergenic |
| chr9 | 110600077 | 110600321 | Distal Intergenic |
| chr9 | 110612210 | 110612505 | Distal Intergenic |
| chr9 | 110782963 | 110784584 | Distal Intergenic |
| chr9 | 111006324 | 111006625 | Distal Intergenic |
| chr9 | 111006814 | 111007013 | Distal Intergenic |
| chr9 | 111021253 | 111021724 | Distal Intergenic |
| chr9 | 111034273 | 111035102 | Distal Intergenic |
| chr9 | 111051454 | 111051838 | Distal Intergenic |
| chr9 | 111058497 | 111058897 | Distal Intergenic |
| chr9 | 111058935 | 111059236 | Distal Intergenic |
| chr9 | 111066092 | 111066980 | Distal Intergenic |
| chr9 | 111145859 | 111146261 | Distal Intergenic |
| chr9 | 111174404 | 111174685 | Distal Intergenic |
| chr9 | 111174873 | 111175383 | Distal Intergenic |
| chr9 | 111489509 | 111489868 | Distal Intergenic |

| | | | |
|---|---|---|---|
| chr9 | 111634116 | 111636471 | Intron |
| chr9 | 111657527 | 111658182 | Intron |
| chr9 | 112852910 | 112853228 | Promoter |
| chr9 | 113287125 | 113288012 | Intron |
| chr9 | 113578962 | 113580528 | Distal Intergenic |
| chr9 | 113615075 | 113616644 | Distal Intergenic |
| chr9 | 113616944 | 113617178 | Distal Intergenic |
| chr9 | 113638237 | 113641280 | Intron |
| chr9 | 114310028 | 114310235 | Intron |
| chr9 | 115216820 | 115217164 | Intron |
| chr9 | 116060158 | 116060398 | Promoter |
| chr9 | 116061313 | 116063828 | Promoter |
| chr9 | 116083287 | 116084181 | Exon |
| chr9 | 116084435 | 116085078 | Intron |
| chr9 | 116127261 | 116129019 | Intron |
| chr9 | 116130894 | 116131522 | Intron |
| chr9 | 116131998 | 116132957 | 3' UTR |
| chr9 | 116133063 | 116133486 | 3' UTR |
| chr9 | 116193143 | 116193389 | Distal Intergenic |
| chr9 | 116220314 | 116220513 | Intron |
| chr9 | 116220782 | 116221276 | Intron |
| chr9 | 116222352 | 116223890 | 5' UTR |
| chr9 | 116228178 | 116228616 | Intron |
| chr9 | 116229066 | 116229505 | Intron |
| chr9 | 116238349 | 116238857 | Intron |
| chr9 | 116244641 | 116245270 | 5' UTR |
| chr9 | 116251435 | 116251816 | Intron |
| chr9 | 116251962 | 116252496 | Intron |
| chr9 | 116277412 | 116278801 | Intron |
| chr9 | 116278854 | 116279264 | Intron |
| chr9 | 116281052 | 116281547 | Intron |
| chr9 | 116281738 | 116281946 | Intron |
| chr9 | 116292821 | 116293869 | Intron |
| chr9 | 116499746 | 116500007 | Distal Intergenic |
| chr9 | 116500145 | 116500517 | Distal Intergenic |
| chr9 | 116500900 | 116501598 | Distal Intergenic |
| chr9 | 116501678 | 116502158 | Distal Intergenic |
| chr9 | 116525993 | 116526789 | Distal Intergenic |
| chr9 | 116811411 | 116813616 | 3' UTR |
| chr9 | 116818150 | 116818499 | 3' UTR |
| chr9 | 116818539 | 116819608 | 3' UTR |
| chr9 | 116925963 | 116926647 | Intron |
| chr9 | 116930162 | 116930750 | Promoter |
| chr9 | 117060880 | 117061823 | Intron |

| | | | |
|---|---|---|---|
| chr9 | 117106665 | 117107097 | Intron |
| chr9 | 118441668 | 118442262 | Distal Intergenic |
| chr9 | 118918885 | 118919977 | Intron |
| chr9 | 119001687 | 119002275 | Intron |
| chr9 | 119002643 | 119002965 | Intron |
| chr9 | 119003063 | 119004234 | Intron |
| chr9 | 119890198 | 119890690 | Intron |
| chr9 | 119890759 | 119891065 | Intron |
| chr9 | 119923027 | 119923235 | Intron |
| chr9 | 120436125 | 120436717 | Distal Intergenic |
| chr9 | 120479994 | 120480294 | Distal Intergenic |
| chr9 | 120502325 | 120502924 | Distal Intergenic |
| chr9 | 120648459 | 120649007 | Distal Intergenic |
| chr9 | 120827948 | 120828255 | Distal Intergenic |
| chr9 | 120870843 | 120871487 | Distal Intergenic |
| chr9 | 121276076 | 121276449 | Distal Intergenic |
| chr9 | 121276497 | 121276747 | Distal Intergenic |
| chr9 | 121740631 | 121740972 | Distal Intergenic |
| chr9 | 122006644 | 122007159 | Intron |
| chr9 | 122012786 | 122013228 | Intron |
| chr9 | 122013278 | 122014475 | Intron |
| chr9 | 122016641 | 122017477 | Intron |
| chr9 | 122017670 | 122017904 | Intron |
| chr9 | 122019181 | 122019608 | Intron |
| chr9 | 123032625 | 123034204 | Distal Intergenic |
| chr9 | 123045871 | 123046983 | Distal Intergenic |
| chr9 | 123078621 | 123078820 | Distal Intergenic |
| chr9 | 123086617 | 123087138 | Distal Intergenic |
| chr9 | 123092696 | 123092916 | Distal Intergenic |
| chr9 | 123093067 | 123093554 | Distal Intergenic |
| chr9 | 123110592 | 123111125 | Distal Intergenic |
| chr9 | 123111842 | 123112602 | Distal Intergenic |
| chr9 | 123112875 | 123113113 | Distal Intergenic |
| chr9 | 123115830 | 123116846 | Distal Intergenic |
| chr9 | 123122304 | 123123370 | Distal Intergenic |
| chr9 | 123123645 | 123124922 | Distal Intergenic |
| chr9 | 123144771 | 123145075 | Distal Intergenic |
| chr9 | 123145111 | 123145447 | Distal Intergenic |
| chr9 | 123145751 | 123145950 | Distal Intergenic |
| chr9 | 123145992 | 123146304 | Distal Intergenic |
| chr9 | 123146422 | 123146936 | Distal Intergenic |
| chr9 | 123147471 | 123147858 | Distal Intergenic |
| chr9 | 123148752 | 123149019 | Downstream |
| chr9 | 123840102 | 123840468 | Intron |

| | | | |
|---|---|---|---|
| chr9 | 124060930 | 124061129 | Promoter |
| chrX | 5564385 | 5565107 | Distal Intergenic |
| chrX | 5992980 | 5993184 | Intron |
| chrX | 6320789 | 6321075 | Distal Intergenic |
| chrX | 6764800 | 6765104 | Distal Intergenic |
| chrX | 6765309 | 6765908 | Distal Intergenic |
| chrX | 7234717 | 7235127 | Intron |
| chrX | 7235200 | 7236228 | Intron |
| chrX | 7248026 | 7248240 | Intron |
| chrX | 7250346 | 7252291 | Exon |
| chrX | 7299292 | 7299662 | Distal Intergenic |
| chrX | 7316981 | 7318294 | Distal Intergenic |
| chrX | 7340025 | 7341380 | Distal Intergenic |
| chrX | 7366650 | 7367101 | Distal Intergenic |
| chrX | 7418992 | 7419236 | Distal Intergenic |
| chrX | 7461086 | 7461352 | Distal Intergenic |
| chrX | 7722453 | 7722871 | Distal Intergenic |
| chrX | 7723082 | 7723346 | Distal Intergenic |
| chrX | 7778891 | 7779351 | Distal Intergenic |
| chrX | 7799409 | 7800649 | Distal Intergenic |
| chrX | 7800790 | 7801023 | Distal Intergenic |
| chrX | 7806295 | 7806538 | Distal Intergenic |
| chrX | 7806710 | 7807347 | Distal Intergenic |
| chrX | 8779534 | 8779744 | Distal Intergenic |
| chrX | 8798979 | 8799218 | Distal Intergenic |
| chrX | 8799508 | 8800480 | Distal Intergenic |
| chrX | 8800733 | 8800980 | Distal Intergenic |
| chrX | 9229725 | 9231403 | Distal Intergenic |
| chrX | 9236503 | 9240051 | Distal Intergenic |
| chrX | 9386642 | 9386841 | Distal Intergenic |
| chrX | 9682461 | 9682763 | Intron |
| chrX | 9682950 | 9683159 | Exon |
| chrX | 9685085 | 9686853 | 3' UTR |
| chrX | 10294177 | 10294468 | Distal Intergenic |
| chrX | 10294760 | 10295059 | Distal Intergenic |
| chrX | 10295109 | 10295398 | Distal Intergenic |
| chrX | 10295830 | 10296251 | Distal Intergenic |
| chrX | 10567730 | 10568636 | Intron |
| chrX | 10572509 | 10572708 | Intron |
| chrX | 10572774 | 10573004 | Intron |
| chrX | 10578270 | 10578502 | Intron |
| chrX | 10768087 | 10768342 | Intron |
| chrX | 11105440 | 11105676 | Intron |
| chrX | 11106015 | 11106214 | Intron |

| | | | |
|---|---|---|---|
| chrX | 11142682 | 11142975 | Distal Intergenic |
| chrX | 11372013 | 11372289 | Intron |
| chrX | 11644906 | 11645327 | Intron |
| chrX | 11923950 | 11924800 | Distal Intergenic |
| chrX | 11942980 | 11943268 | Distal Intergenic |
| chrX | 12107649 | 12108214 | Distal Intergenic |
| chrX | 12178867 | 12179308 | Intron |
| chrX | 12250468 | 12250751 | Intron |
| chrX | 12250841 | 12251219 | Intron |
| chrX | 12337787 | 12338032 | Intron |
| chrX | 12338070 | 12338860 | Intron |
| chrX | 12338933 | 12339721 | Intron |
| chrX | 12339872 | 12340087 | Intron |
| chrX | 12399404 | 12400524 | Intron |
| chrX | 12447629 | 12447917 | Intron |
| chrX | 12503686 | 12504348 | Intron |
| chrX | 12536534 | 12537113 | Intron |
| chrX | 12555028 | 12555386 | Intron |
| chrX | 12619242 | 12619470 | Intron |
| chrX | 12648886 | 12649264 | Intron |
| chrX | 12649492 | 12649741 | Intron |
| chrX | 12649873 | 12650295 | Intron |
| chrX | 12807316 | 12807515 | Distal Intergenic |
| chrX | 12807560 | 12807911 | Distal Intergenic |
| chrX | 12857172 | 12858702 | Distal Intergenic |
| chrX | 12859201 | 12859499 | Distal Intergenic |
| chrX | 12859697 | 12859949 | Distal Intergenic |
| chrX | 12975540 | 12975749 | Distal Intergenic |
| chrX | 13029835 | 13030037 | Distal Intergenic |
| chrX | 13030238 | 13030946 | Distal Intergenic |
| chrX | 13038979 | 13039463 | Distal Intergenic |
| chrX | 13039737 | 13040383 | Distal Intergenic |
| chrX | 13084090 | 13084538 | Distal Intergenic |
| chrX | 13100856 | 13103070 | Distal Intergenic |
| chrX | 16175033 | 16176441 | Distal Intergenic |
| chrX | 16723067 | 16723266 | Intron |
| chrX | 16724183 | 16724424 | Intron |
| chrX | 16767848 | 16768886 | Intron |
| chrX | 17148463 | 17149034 | Intron |
| chrX | 17149085 | 17149353 | Intron |
| chrX | 17740156 | 17741332 | Intron |
| chrX | 18417706 | 18418070 | Distal Intergenic |
| chrX | 18739716 | 18741489 | Intron |
| chrX | 18742702 | 18745024 | Intron |

| | | | |
|---|---|---|---|
| chrX | 18745407 | 18745935 | Intron |
| chrX | 19264313 | 19265464 | Distal Intergenic |
| chrX | 19314123 | 19314373 | Distal Intergenic |
| chrX | 19314516 | 19314715 | Distal Intergenic |
| chrX | 19314775 | 19314989 | Distal Intergenic |
| chrX | 19481192 | 19481801 | Intron |
| chrX | 19481926 | 19482251 | Intron |
| chrX | 19941732 | 19941942 | Intron |
| chrX | 19995452 | 19996159 | Distal Intergenic |
| chrX | 20003467 | 20003700 | Downstream |
| chrX | 20194428 | 20194643 | Exon |
| chrX | 20194684 | 20195277 | Exon |
| chrX | 20240020 | 20240840 | Intron |
| chrX | 20264089 | 20264466 | Intron |
| chrX | 20264503 | 20264892 | Intron |
| chrX | 20393701 | 20394291 | Distal Intergenic |
| chrX | 20394376 | 20394651 | Distal Intergenic |
| chrX | 20394828 | 20395077 | Distal Intergenic |
| chrX | 20444761 | 20445898 | Distal Intergenic |
| chrX | 20446054 | 20447062 | Distal Intergenic |
| chrX | 20492189 | 20492402 | Distal Intergenic |
| chrX | 20538804 | 20539104 | Distal Intergenic |
| chrX | 21142675 | 21143203 | Distal Intergenic |
| chrX | 21144234 | 21144827 | Distal Intergenic |
| chrX | 21155647 | 21156379 | Distal Intergenic |
| chrX | 21156591 | 21157334 | Distal Intergenic |
| chrX | 33468297 | 33469096 | Distal Intergenic |
| chrX | 33470236 | 33471252 | Distal Intergenic |
| chrX | 33618632 | 33618939 | Distal Intergenic |
| chrX | 33619121 | 33619417 | Distal Intergenic |
| chrX | 33654081 | 33654442 | Distal Intergenic |
| chrX | 33654957 | 33656264 | Distal Intergenic |
| chrX | 33669000 | 33669877 | Distal Intergenic |
| chrX | 33925621 | 33926270 | Distal Intergenic |
| chrX | 34137716 | 34138031 | Distal Intergenic |
| chrX | 34335888 | 34336121 | Distal Intergenic |
| chrX | 34347525 | 34347821 | Distal Intergenic |
| chrX | 34414488 | 34414742 | Distal Intergenic |
| chrX | 34622767 | 34622966 | Distal Intergenic |
| chrX | 34624057 | 34624262 | Distal Intergenic |
| chrX | 34624689 | 34625157 | Distal Intergenic |
| chrX | 34649969 | 34650286 | Intron |
| chrX | 34666995 | 34667201 | Intron |
| chrX | 35395446 | 35395749 | Distal Intergenic |

| | | | |
|---|---|---|---|
| chrX | 35395829 | 35396250 | Distal Intergenic |
| chrX | 35396702 | 35397538 | Distal Intergenic |
| chrX | 35514429 | 35514645 | Distal Intergenic |
| chrX | 35538305 | 35539509 | Distal Intergenic |
| chrX | 35542689 | 35543525 | Distal Intergenic |
| chrX | 35781987 | 35782191 | Distal Intergenic |
| chrX | 35795565 | 35797588 | Distal Intergenic |
| chrX | 35798668 | 35799479 | Distal Intergenic |
| chrX | 35826560 | 35826759 | Distal Intergenic |
| chrX | 35917633 | 35918048 | Distal Intergenic |
| chrX | 35953519 | 35954369 | Intron |
| chrX | 39265115 | 39265392 | Distal Intergenic |
| chrX | 39277419 | 39278300 | Distal Intergenic |
| chrX | 39317486 | 39318159 | Distal Intergenic |
| chrX | 39322855 | 39324334 | Distal Intergenic |
| chrX | 39421411 | 39421707 | Distal Intergenic |
| chrX | 39421974 | 39422929 | Distal Intergenic |
| chrX | 39492545 | 39493762 | Distal Intergenic |
| chrX | 39502665 | 39503108 | Distal Intergenic |
| chrX | 39503907 | 39505290 | Distal Intergenic |
| chrX | 42460138 | 42460351 | Distal Intergenic |
| chrX | 45283123 | 45284337 | Distal Intergenic |
| chrX | 45537995 | 45538232 | Distal Intergenic |
| chrX | 45538424 | 45538817 | Distal Intergenic |
| chrX | 45606588 | 45606896 | Promoter |
| chrX | 45610050 | 45610299 | Distal Intergenic |
| chrX | 45806616 | 45806842 | Distal Intergenic |
| chrX | 45813611 | 45813812 | Distal Intergenic |
| chrX | 45819240 | 45821053 | Distal Intergenic |
| chrX | 47988351 | 47988558 | Intron |
| chrX | 48176957 | 48177723 | Distal Intergenic |
| chrX | 48272116 | 48272552 | Promoter |
| chrX | 48282742 | 48284252 | Distal Intergenic |
| chrX | 48343913 | 48345511 | 3' UTR |
| chrX | 48370469 | 48371049 | 5' UTR |
| chrX | 48372056 | 48372667 | Exon |
| chrX | 48521061 | 48521493 | Distal Intergenic |
| chrX | 48521815 | 48522181 | Distal Intergenic |
| chrX | 48538213 | 48539289 | Distal Intergenic |
| chrX | 48585635 | 48586428 | Distal Intergenic |
| chrX | 48727041 | 48727440 | Distal Intergenic |
| chrX | 48859361 | 48860250 | Promoter |
| chrX | 48860572 | 48860790 | Distal Intergenic |
| chrX | 49056123 | 49056592 | Promoter |

| | | | |
|---|---|---|---|
| chrX | 49517481 | 49517693 | Distal Intergenic |
| chrX | 49573751 | 49574372 | Distal Intergenic |
| chrX | 50266050 | 50266354 | Distal Intergenic |
| chrX | 50340953 | 50341182 | Intron |
| chrX | 50341341 | 50342683 | Exon |
| chrX | 50468397 | 50468617 | Intron |
| chrX | 53260351 | 53261438 | Downstream |
| chrX | 53930920 | 53933003 | Distal Intergenic |
| chrX | 60399410 | 60399896 | Distal Intergenic |
| chrX | 60715706 | 60717329 | Distal Intergenic |
| chrX | 65665530 | 65665790 | Distal Intergenic |
| chrX | 65931896 | 65932095 | Distal Intergenic |
| chrX | 65932173 | 65932373 | Distal Intergenic |
| chrX | 66585241 | 66585555 | Distal Intergenic |
| chrX | 67639084 | 67639484 | Intron |
| chrX | 67639595 | 67639951 | Intron |
| chrX | 69289693 | 69290288 | Distal Intergenic |
| chrX | 69543338 | 69543537 | Intron |
| chrX | 70184712 | 70185829 | Distal Intergenic |
| chrX | 70919751 | 70920524 | Intron |
| chrX | 70920573 | 70920840 | Exon |
| chrX | 70923608 | 70924386 | Intron |
| chrX | 70924428 | 70924776 | Intron |
| chrX | 70927089 | 70927726 | Intron |
| chrX | 70961078 | 70961277 | Distal Intergenic |
| chrX | 70961654 | 70961976 | Distal Intergenic |
| chrX | 70963793 | 70964889 | Distal Intergenic |
| chrX | 71081321 | 71081722 | Distal Intergenic |
| chrX | 71160149 | 71160358 | Intron |
| chrX | 71167396 | 71167707 | Intron |
| chrX | 71211068 | 71213256 | Intron |
| chrX | 71489132 | 71490147 | Downstream |
| chrX | 71490307 | 71490526 | Downstream |
| chrX | 71492242 | 71492441 | Downstream |
| chrX | 71492977 | 71493191 | Exon |
| chrX | 71494073 | 71495226 | 3' UTR |
| chrX | 71495275 | 71495816 | Exon |
| chrX | 71500396 | 71500940 | Distal Intergenic |
| chrX | 71514905 | 71516893 | Distal Intergenic |
| chrX | 71527055 | 71527444 | Promoter |
| chrX | 71528111 | 71528602 | Distal Intergenic |
| chrX | 71614091 | 71614306 | Intron |
| chrX | 71638137 | 71638533 | Intron |
| chrX | 71638646 | 71639290 | Intron |

| | | | |
|---|---|---|---|
| chrX | 71669350 | 71672220 | Intron |
| chrX | 71672467 | 71672681 | Intron |
| chrX | 71674075 | 71675257 | Intron |
| chrX | 72271388 | 72271626 | Distal Intergenic |
| chrX | 72273170 | 72273451 | Distal Intergenic |
| chrX | 72273518 | 72273786 | Distal Intergenic |
| chrX | 72273850 | 72274230 | Distal Intergenic |
| chrX | 72277237 | 72277455 | Distal Intergenic |
| chrX | 72277516 | 72277717 | Distal Intergenic |
| chrX | 72375861 | 72376303 | Distal Intergenic |
| chrX | 72381901 | 72383045 | Distal Intergenic |
| chrX | 72383316 | 72383515 | Distal Intergenic |
| chrX | 72661506 | 72661792 | Distal Intergenic |
| chrX | 72782203 | 72782402 | Promoter |
| chrX | 72783323 | 72783581 | Promoter |
| chrX | 73009450 | 73009764 | Distal Intergenic |
| chrX | 73077950 | 73080483 | Distal Intergenic |
| chrX | 73080526 | 73080810 | Distal Intergenic |
| chrX | 73087565 | 73090159 | Distal Intergenic |
| chrX | 73091013 | 73091271 | Distal Intergenic |
| chrX | 73091475 | 73091709 | Distal Intergenic |
| chrX | 73119324 | 73119629 | Distal Intergenic |
| chrX | 73120027 | 73120431 | Distal Intergenic |
| chrX | 74756994 | 74757425 | Distal Intergenic |
| chrX | 74861772 | 74862379 | Distal Intergenic |
| chrX | 74862808 | 74863267 | Distal Intergenic |
| chrX | 74863358 | 74863804 | Distal Intergenic |
| chrX | 75041051 | 75041256 | Distal Intergenic |
| chrX | 76896368 | 76896612 | Intron |
| chrX | 83016706 | 83017061 | Distal Intergenic |
| chrX | 90779815 | 90780394 | Distal Intergenic |
| chrX | 90780451 | 90780687 | Distal Intergenic |
| chrX | 90877129 | 90877494 | Distal Intergenic |
| chrX | 90879719 | 90880126 | Distal Intergenic |
| chrX | 90919832 | 90920204 | Distal Intergenic |
| chrX | 90920525 | 90921209 | Distal Intergenic |
| chrX | 91305953 | 91306179 | Intron |
| chrX | 91363365 | 91364248 | Intron |
| chrX | 91366433 | 91368283 | Intron |
| chrX | 91384592 | 91384827 | Intron |
| chrX | 91397618 | 91398021 | Intron |
| chrX | 91414946 | 91415562 | Intron |
| chrX | 91423064 | 91423263 | Intron |
| chrX | 92361216 | 92361415 | Distal Intergenic |

| | | | |
|---|---|---|---|
| chrX | 92361616 | 92361840 | Distal Intergenic |
| chrX | 93143775 | 93144120 | Distal Intergenic |
| chrX | 93291414 | 93293844 | Distal Intergenic |
| chrX | 93302143 | 93303132 | Distal Intergenic |
| chrX | 93303292 | 93303988 | Distal Intergenic |
| chrX | 93316119 | 93318425 | Distal Intergenic |
| chrX | 93320862 | 93321082 | Distal Intergenic |
| chrX | 93324932 | 93325483 | Distal Intergenic |
| chrX | 93325527 | 93326290 | Distal Intergenic |
| chrX | 93326359 | 93326656 | Distal Intergenic |
| chrX | 93326696 | 93326997 | Distal Intergenic |
| chrX | 93328030 | 93328633 | Distal Intergenic |
| chrX | 93328693 | 93330861 | Distal Intergenic |
| chrX | 94561095 | 94562068 | Distal Intergenic |
| chrX | 94568019 | 94568293 | Distal Intergenic |
| chrX | 94568341 | 94568599 | Distal Intergenic |
| chrX | 94568828 | 94569071 | Distal Intergenic |
| chrX | 94569119 | 94569569 | Distal Intergenic |
| chrX | 94569618 | 94570311 | Distal Intergenic |
| chrX | 94570907 | 94572542 | Distal Intergenic |
| chrX | 94572611 | 94572869 | Distal Intergenic |
| chrX | 96331187 | 96331820 | Intron |
| chrX | 96331967 | 96332422 | Intron |
| chrX | 96332579 | 96332902 | Intron |
| chrX | 96333018 | 96334391 | Intron |
| chrX | 96334793 | 96335028 | Intron |
| chrX | 96335515 | 96336872 | Intron |
| chrX | 96336920 | 96337699 | Intron |
| chrX | 96338696 | 96339041 | Intron |
| chrX | 96339097 | 96340038 | Intron |
| chrX | 96618040 | 96618480 | Intron |
| chrX | 96618684 | 96619189 | Intron |
| chrX | 96648619 | 96648823 | Intron |
| chrX | 96717720 | 96718447 | Intron |
| chrX | 96799482 | 96805051 | Intron |
| chrX | 96805137 | 96805903 | Intron |
| chrX | 96805961 | 96806258 | Intron |
| chrX | 97821100 | 97821360 | Distal Intergenic |
| chrX | 97882896 | 97883449 | Distal Intergenic |
| chrX | 97911220 | 97911651 | Distal Intergenic |
| chrX | 97911711 | 97912144 | Distal Intergenic |
| chrX | 97912188 | 97912459 | Distal Intergenic |
| chrX | 97933655 | 97933913 | Distal Intergenic |
| chrX | 98624342 | 98624757 | Distal Intergenic |

| | | | |
|---|---|---|---|
| chrX | 98625243 | 98625669 | Distal Intergenic |
| chrX | 98835334 | 98835815 | Intron |
| chrX | 98962473 | 98962672 | Intron |
| chrX | 99383860 | 99384084 | Distal Intergenic |
| chrX | 99384439 | 99384638 | Distal Intergenic |
| chrX | 99579114 | 99579326 | Intron |
| chrX | 99862020 | 99862784 | Distal Intergenic |
| chrX | 100689266 | 100689572 | Distal Intergenic |
| chrX | 100704502 | 100705059 | Distal Intergenic |
| chrX | 100709942 | 100710531 | Distal Intergenic |
| chrX | 100710572 | 100711340 | Distal Intergenic |
| chrX | 100807380 | 100807668 | Intron |
| chrX | 100811546 | 100812036 | Distal Intergenic |
| chrX | 100812411 | 100812610 | Distal Intergenic |
| chrX | 100843620 | 100844572 | Distal Intergenic |
| chrX | 100845839 | 100846136 | Distal Intergenic |
| chrX | 100846736 | 100847150 | Distal Intergenic |
| chrX | 100847519 | 100847756 | Distal Intergenic |
| chrX | 100847820 | 100848786 | Distal Intergenic |
| chrX | 100985397 | 100986736 | Distal Intergenic |
| chrX | 101175354 | 101176580 | Intron |
| chrX | 101678498 | 101678893 | Intron |
| chrX | 101682806 | 101683085 | Intron |
| chrX | 101683127 | 101683612 | Intron |
| chrX | 101683704 | 101683985 | Intron |
| chrX | 103124328 | 103124728 | Distal Intergenic |
| chrX | 103381398 | 103381825 | Intron |
| chrX | 103381969 | 103382274 | Intron |
| chrX | 103382483 | 103382690 | Intron |
| chrX | 103382736 | 103383764 | Intron |
| chrX | 103383808 | 103384064 | Intron |
| chrX | 106028222 | 106028463 | Exon |
| chrX | 109812340 | 109812654 | Distal Intergenic |
| chrX | 115931578 | 115931813 | Distal Intergenic |
| chrX | 126091455 | 126092009 | Distal Intergenic |
| chrX | 126092082 | 126092334 | Distal Intergenic |
| chrX | 131075502 | 131075748 | Distal Intergenic |
| chrX | 131120283 | 131120585 | Distal Intergenic |
| chrX | 132779679 | 132779902 | Intron |
| chrX | 132802400 | 132802861 | Intron |
| chrX | 133275282 | 133275625 | Distal Intergenic |
| chrX | 133555758 | 133556629 | Intron |
| chrX | 136532500 | 136533027 | Distal Intergenic |
| chrX | 137077547 | 137077761 | Distal Intergenic |

| | | | |
|---|---|---|---|
| chrX | 137107362 | 137107962 | Distal Intergenic |
| chrX | 137837741 | 137837959 | Intron |
| chrX | 138771149 | 138771451 | Intron |
| chrX | 138825197 | 138825488 | Intron |
| chrX | 139332077 | 139332349 | Distal Intergenic |
| chrX | 139332388 | 139333512 | Distal Intergenic |
| chrX | 139348070 | 139348457 | Distal Intergenic |
| chrX | 147022897 | 147023208 | Intron |
| chrX | 147511088 | 147511581 | Distal Intergenic |
| chrX | 148236856 | 148237103 | Distal Intergenic |
| chrX | 148238221 | 148238557 | Distal Intergenic |
| chrX | 149138008 | 149138543 | Intron |
| chrX | 149138735 | 149139113 | Intron |
| chrX | 149139284 | 149139575 | Intron |
| chrX | 149151654 | 149152097 | Intron |
| chrX | 149152164 | 149152648 | Intron |
| chrX | 149196782 | 149197401 | Intron |
| chrX | 149197439 | 149197638 | Intron |
| chrX | 151651187 | 151651583 | Distal Intergenic |
| chrX | 151697081 | 151697463 | Distal Intergenic |
| chrX | 156246649 | 156247651 | Distal Intergenic |
| chrX | 156247729 | 156248551 | Distal Intergenic |
| chrX | 156270508 | 156271199 | Distal Intergenic |
| chrX | 156278147 | 156278789 | Distal Intergenic |
| chrX | 156279034 | 156280312 | Distal Intergenic |
| chrX | 156286751 | 156287097 | Distal Intergenic |
| chrX | 156288380 | 156288773 | Distal Intergenic |
| chrX | 160450798 | 160451173 | Distal Intergenic |
| chrX | 160451267 | 160452068 | Distal Intergenic |
| chrX | 160573485 | 160573809 | Distal Intergenic |
| chrX | 160720827 | 160722663 | Distal Intergenic |
| chrX | 160817521 | 160817980 | Distal Intergenic |
| chrX | 160818113 | 160818417 | Distal Intergenic |
| chrX | 160831624 | 160832185 | Distal Intergenic |
| chrX | 160832250 | 160832482 | Distal Intergenic |
| chrX | 163099039 | 163099362 | Distal Intergenic |
| chrX | 163203275 | 163204348 | Distal Intergenic |
| chrX | 163243592 | 163243948 | Distal Intergenic |
| chrX | 163409328 | 163409898 | Distal Intergenic |
| chrX | 163410170 | 163410461 | Distal Intergenic |
| chrX | 163445325 | 163446044 | Distal Intergenic |
| chrX | 163529872 | 163530408 | Distal Intergenic |
| chrX | 163589691 | 163589956 | Distal Intergenic |

**Supplementary Table 4. Summary of Primers**

| | | |
|---|---|---|
| Primers used for real-time quantitative PCR (5' to 3') | | |
| | Forward primer sequence | Reverse primer sequence |
| Mouse *Sox2* | GCGGAGTGGAAACTTTTGTCC | GGGAAGCGTGTACTTATCCTTCT |
| Human *SMAD4* | CTCATGTGATCTATGCCCGTC | AGGTGATACAACTCGTTCGTAGT |
| Human *SOX2* | CACAACTCGGAGATCAGCAA | CTCCGGGAAGCGTGTACTTA |
| *GAPDH* | AATCCCATCACCATCTTCCA | TGGACTCCACGACGTACTCA |
| Primers (5' to 3') used for PCR to verify the knockout of *Sox2/SOX2*-associated loops targeted regions | | |
| Mouse *Sox2*-associated loop targeted enhancer region knockout | TCAGCTTCAACGAAACTCTG | GGCCAATAAATGGCAAGAAG |
| Human *SOX2*-associated loop targeted enhancer region knockout | GTTTTGGTGTCACCCATTGGAG | ATGCAAATTGTTAATGCCCAAAGA |
| Primers (5' to 3') used for 3C-PCR and 3C-qPCR | | |
| Mouse *Sox2*-associated loop | GGTCCTCCCAAACCTAATGT | GGTGGTGTGCCATTGTTTCT |
| Human *SOX2*-associated loop | CCGGGTTCCCAAGAACTAAA | ATGCAAATTGTTAATGCCCA |

| Primers (5' to 3') used for ChIP-PCR and ChIP-qPCR | | |
|---|---|---|
| Mouse *Sox2* promoter region | GGCTTGGGTCTAACTTCTCG | GTGGTGTGCCATTGTTTCTG |
| Mouse *Sox2*-associated loop targeted enhancer region | GGGACGAGTGTGCTTTATGT | AGCAGAGCATCATGGGAATG |

**Supplementary Table 5. Summary of Guide RNA (gRNA) Sequences**

| Summary of guide RNA (gRNA) sequences (5' to 3') | | | |
|---|---|---|---|
| | vector to ligation | gRNA oligo1 | gRNA oligo2 |
| Mouse *Smad4* gRNA | LentiCRIPSRV2 | CACCGGCCAAGTAATCGCGCATCAA | AAACTTGATGCGCGATTACTTGGCC |
| Human *SMAD4* gRNA | | CACCGAACTCTGTACAAAGACCGCG | AAACCGCGGTCTTTGTACAGAGTTC |
| Human *PTEN* gRNA | | CACCGACCGCCAAATTTAATTGCAG | AAACCTGCAATTAAATTTGGCGGTC |
| Mouse *Sox2*-associated loop targeted enhancer region dual-gRNA | Plenti-H1-hU6-hygro-dual-gRNA | ATTTCTAGCTCTAAAACACATTACGGCGAAAAGAATCCGGGAAAGAGTGGTCTC | ATTTCTAGCTCTAAAACCTCGAGCTGCTAAAAGTTCGCGGTGTTTCGTCCTTTCC |
| Mouse *p300* dual-gRNA | | ATTTCTAGCTCTAAAACTCTCGGCGTCCGCCAGCGATCGGGAAAGAGTGGTCTC | ATTTCTAGCTCTAAAACGCCGGTAAAGTGCCTCCAATCGGTGTTTCGTCCTTTCC |

| | | |
|---|---|---|
| Human *SOX2*-associated loop targeted enhancer region dual-gRNA | ATTTCTAGCTCTAAAACTCCACTTACATGCTTCGGTGCGGGAAAGAGTGGTCTC | ATTTCTAGCTCTAAAACCGTTACACCCAATATCTTCCCGGTGTTTCGTCCTTTCC |

**Supplementary Table 6. Summary of siRNA Sequences**

| Summary of guide siRNA sequences (5' to 3') | | |
|---|---|---|
| | sence | anti-sence |
| si-NC (Mouse) | UUCUCCGACAGUGUCACGU(dT)(dT) | ACGUGACACUGUCGGAGAA(dT)(dT) |
| si-*Smad4* (Mouse) | CAGUAUGCGUUUGACUUAA(dT)(dT) | UUAAGUCAAACGCAUACUG(dT)(dT) |
| si-NC (Human) | UUCUCCGAACGUGUCACGU(dT)(dT) | ACGUGACACGUUCGGAGAA(dT)(dT) |
| si-*SMAD4* (Human) | CAGAGUUACUACUUAGACA(dT)(dT) | UGUCUAAGUAGUAACUCUG(dT)(dT) |

**Supplementary Table 7. Summary of Antisense Oligonucleotides (ASO) Sequences**

| Summary of guide siRNA sequences (5' to 3') | |
|---|---|
| | sence |
| NC ASO | GCGUATTATAGCCGAUUAAC |
| *Sox2*-associated enhancer region targeted by ASO | AATGCACATAAAGCACACTC |
